# Unraveling the role of receptor-like protein tyrosine phosphatase PTPRH in cell signaling regulation and biological processes of non-small cell lung cancer

**DOI:** 10.1101/2024.06.13.598886

**Authors:** Mylena M. O. Ortiz, Deeya M. Patel, Jesus A. García-Lerena, Andrew C. Nelson, Matthew Swiatnicki, Eran Andrechek

**Affiliations:** Genetics and Genomic Sciences Program, Michigan State University; Department of Physiology, Michigan State University; Lyman Briggs College, Michigan State University; Cellular and Molecular Biology Program, Michigan State University; Department of Laboratory Medicine and Pathology, University of Minnesota

## Abstract

The balance of protein phosphorylation is often disrupted in cancer, with hyperactivity of kinases and inactivation of phosphatases driving cell proliferation and survival pathways. PTPRH, a protein tyrosine phosphatase, is mutated in ∼5% of non-small cell lung cancers (NSCLC). However, how PTPRH contributes to biological processes and tumorigenesis was unknown. We uncovered PTPRH’s candidate interactors and associated pathways by applying a proximity-dependent biotinylation assay (BioID) and generating a signature transcriptome in two NSCLC cell lines derived from the primary tumor (NCI-H23) or a metastatic site (NCI-H2023), followed by functional validation. Candidate interactors included signaling molecules and structural proteins linked to integrins and focal adhesions, adherens junctions, migration, and the cytoskeleton, in addition to interactions with the receptor tyrosine kinases EGFR, EPHA2, and ROR2, and the phosphatases PTPN3 and PTPRJ. Considering the importance of EGFR in lung cancers and the role of EPHA2 in regulating cell adhesion, we examined how PTPRH regulates their signaling. Overexpression of PTPRH decreased EGFR phosphorylation at tyrosine 1173. It also reduced phospho-EPHA2, with one of the target tyrosine residues identified as the ligand-dependent Y588. At the cellular level, PTPRH and EPHA2 colocalize, with PTPRH gain inducing morphological alterations, such as increased eccentricity, smaller size and changes in the cytoskeleton organization in NCI-H23 cells. These changes are accompanied by increased FAK Y397 phosphorylation, but reduced cell adhesion to the ECM. Additionally, pathway enrichment analysis revealed downregulation of multiple oncogenic, metabolic, and cell adhesion signaling pathways, with increased levels of PTPRH leading to reduced migration in vitro, suppressed tumor growth and lung colonization and tumor differentiation in vivo. Interestingly, some alterations may be independent of PTPRH catalytic activity and tailored to a cell line’s site of origin and genetic background. These results indicate that PTPRH regulates key signaling, structural networks, and tumor behavior with loss facilitating NSCLC progression.

## Introduction

In normal cells, phosphorylation and dephosphorylation are tightly regulated in time and space by kinases and phosphatases, and together they orchestrate fundamental cellular processes, such as proliferation, differentiation, and apoptosis^1,2^. However, this delicate equilibrium is often disrupted in cancer^3^. While mutations in kinases in non-small cell lung cancers (NSCLC) are the focus of numerous studies, such as EGFR, the consequences of mutations in phosphatases are much less well understood. Phosphatases can exhibit characteristics of both tumor suppressors and oncogenes in lung cancer^16^. A well-known example is *PTEN*, an important tumor-suppressor gene mutated in approximately 7% of NSCLC cases. The inactivation of this phosphatase leads to an increase in the PI3K/mTOR/Akt signaling pathway and lower survival rates^5,6^. Conversely, activating mutations in the tyrosine phosphatase *PTPN11/*Shp2 increase phospho-ERK 1/2 levels and activate the MAPK pathway, thus behaving as an oncogene^7^. Recently, our group has shown that point mutations in the receptor-like protein tyrosine phosphatase (RPTP) *PTPRH* are observed in ∼5% of NSCLC patients and over 80% of MMTV-*PyMT* mice, a murine model of breast cancer^8,9^. A mutation that results in inactivation of this phosphatase leads to elevated phosphorylation of EGFR at tyrosine 1173 (alternatively numbered as 1197) and results in activation of the Akt pathway^8–10^. This suggests that PTPRH contributes to negatively regulating the EGFR pathway, however, the precise underlying mechanism is unclear.

PTPRH, also known as SAP-1, is part of the tyrosine phosphatase family, which encompasses over 100 members^11^. Structurally, PTPRH is composed of an extracellular region rich in fibronectin type 3 (FN3) repeats, a hydrophobic transmembrane domain, and a single intracellular phosphatase domain^12,13^. Little is known about PTPRH kinetics and interactors; however, a few associations with tyrosine kinase signaling and cellular processes have been found in different cell types. By dephosphorylating tyrosine residues, PTPRH is suggested to negatively regulate T-cell function^14^, intestinal immunity^15^, integrin-associated pathways^16^, and, more recently, EGFR signaling^8–10,17^. Furthermore, PTPRH has been found to induce apoptotic cell death pathways^18^ and may stimulate c-Src activity^19^.

The role of this phosphatase in cancer, although not fully elucidated, seems to be largely related to the tissue of origin. PTPRH expression varies across and within different cancer types, and conflicting findings have been observed^12,20–22^. For instance, in colorectal cancers, PTPRH plays a causal role in cancer progression, but not initiation, and overexpression was observed in some cases ^23,24^. Conversely, as hepatocarcinoma advances to poorly differentiated stages, a decrease in the phosphatase level was observed^25^. In non-small cell lung cancers, *PTPRH* and *EGFR* mutations were found to be mutually exclusive^8^. Therefore, it is evident that we do not understand the full spectrum of cell signaling regulation by PTPRH and how it stimulates or suppresses tumorigenesis. To bridge this gap, we built a comprehensive map of PTPRH interactors and associated cell signaling networks in lung adenocarcinoma cell lines derived from the primary tumor and a metastatic site. Using multi-omics techniques, we began by investigating protein-protein interactions and the various signaling pathways and biological processes associated with lung cancers in which this phosphatase is involved. Our study revealed novel PTPRH interactors and functions, and we explored how the gain or loss of function of this phosphatase may affect cellular behavior and tumor hallmarks, shedding light on how mutations in this phosphatase may deregulate lung cancer cell signaling pathways.

## Results

### BioID reveals novel PTPRH candidate interactors in lung cancer cells

Little is known about PTPRH interaction partners and how this phosphatase may regulate different biological processes. To address this, we generated a comprehensive map of PTPRH’s protein-protein interactions using the BioID assay. This is a powerful *in-vitro* labeling technique that adds a biotin tag to proteins interacting stably or transiently with the protein of interest in a labeling radius of ∼10 nm^26^ (Fig. 1A). We cloned the BioID2 sequence, followed by an HA tag, into the C-terminus of the PTPRH wild-type (PTPRH-WT) coding sequence, in proximity to its intracellular phosphatase domain. Given that reaction rates for tyrosine phosphatases and the interaction with their substrates can be notoriously fast^27,28^, we also generated a substrate-trapping construct (PTPRH-DACS) containing mutations in critical catalytic residues (**D**986**A/C**1020**S**), allowing us to maximize discovery of candidate interactors (Supplemental Fig. 1A)^24,29,30^. To investigate PTPRH’s interactors in lung cancer cells, one human lung adenocarcinoma cell line derived from the primary site (NCI-H23) in which the *PTPRH* gene was previously knocked out (H23 KO)^8^ and one derived from the lymph node metastatic site (NCI-H2023) were infected with the constructs. Their expression and correct subcellular localization were validated using immunoblot and immunofluorescence (Supplemental Fig. 1B-C, Supplemental Fig. 2A-B). The modified PTPRH was localized mainly in the plasma membrane and cytoplasm, in agreement with the subcellular localization of the endogenous protein. Promiscuous biotinylation was noted in the cells infected with BioID2-only, while some biotinylated proteins were present exclusively in the wild-type or substrate-trapping PTPRH-BioID2 group, suggesting unique interactors (Supplemental Fig. 1D-E). As expected, overexpression of the wild-type construct notably reduced tyrosine phosphorylation of a protein at ∼120 KDa, whereas augmented phosphorylation was observed in the substrate trapping group, validating the presence or loss of catalytic activity, respectively (Supplemental Fig. 1F).

**Figure 1.**
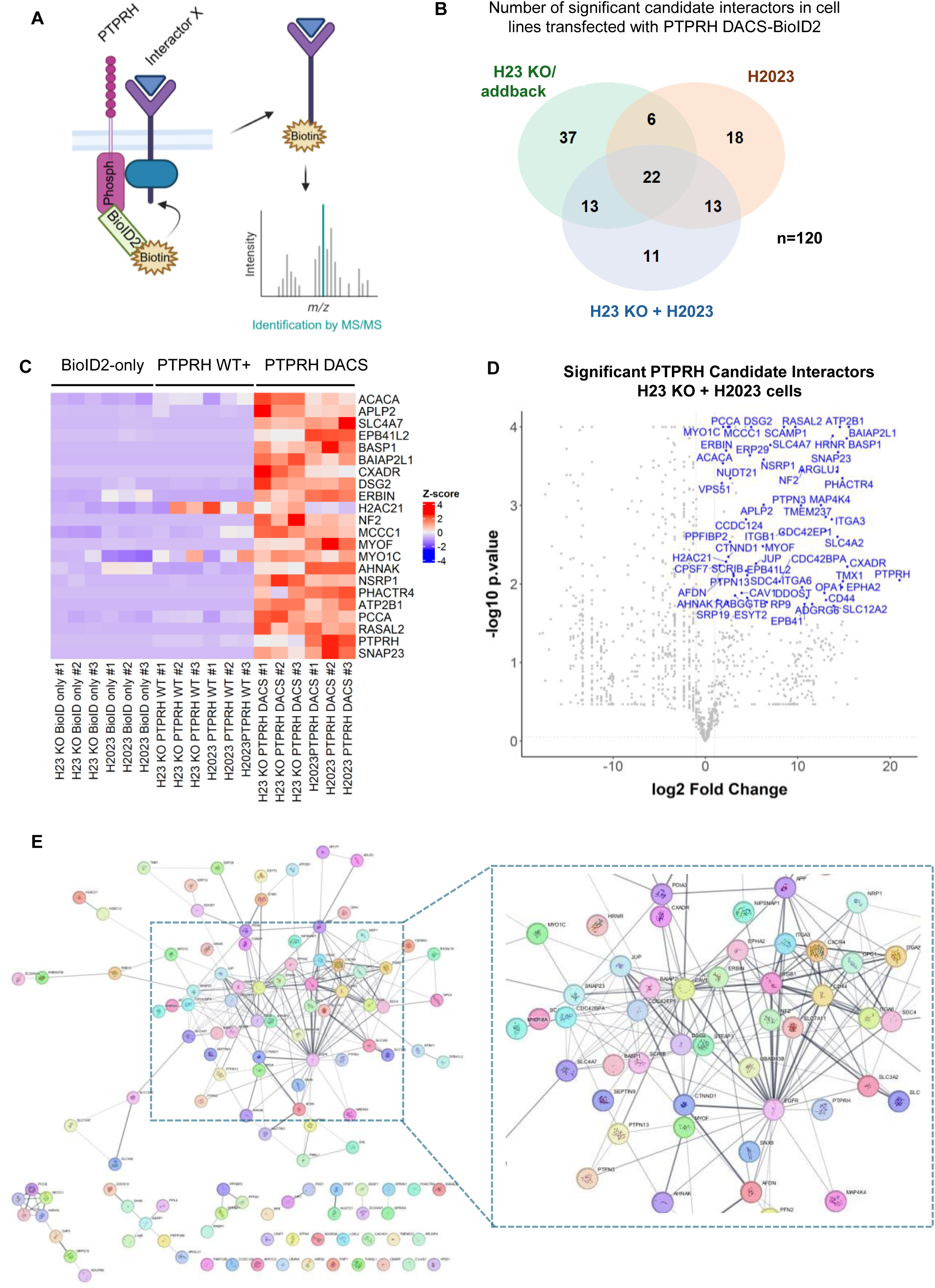
PTPRH interactors revealed by the *in vitro* labeling technique (BioID) in lung adenocarcinoma cell lines. *A)* Schematic representation of the BioID assay. H2023 and H23 KO cell lines transduced with BioID2-HA-only, PTPRH WT-BioID2-HA, PTPRH DACS-BioID2-HA (substrate-trapping mutant), or mock (negative control) were serum-starved for 24h, followed by stimulation with FBS and biotin for 18h. Proteins that interact stably or transiently with PTPRH are biotinylated. These candidate interactors are pulled down and identified by mass spectrometry. *B)* Venn diagram of PTPRH candidate interactors identified in each cell line by the BioID assay. *C)* Heatmap showing the 22 candidate interactors commonly found in all three analyses in PTPRH DACS-BioID2-HA transduced cells (see Figure 1B) and z-scores of normalized total precursor intensity across BioID2-HA-only, PTPRH WT-BioID2-HA, and PTPRH DACS-BioID2-HA groups*. D)* Volcano plots representing the 59 significant PTPRH candidate interactors (labeled in blue) identified in the PTPRH DACS-BioID2-HA vs. BioID-HA-only combinatorial analysis of H2023 and H23 KO cells. *E)* STRING analysis of all PTPRH candidate interactors identified in the cell lines individually or in combination (n=120 proteins).

Rigorous reproduction was observed in the substrate-trapping cell lines (Fig. 1C and 2B), which were chosen for subsequent analysis. In total, 120 unique interacting partners were identified (Fig. 1B and Supplemental Table 1). Individual analysis of H2023 or H23 KO cells revealed a total of 59 and 78 candidate interactors, respectively (Fig. 1D and Supplemental Fig. 4A). When both cell lines were analyzed in combination, 59 candidate interactors achieved significance. Of these candidates, 22 proteins were common to all three analyses (Fig. 1B-C). This agrees with a previous report that employed BioID to screen PTP1B interactors and a limited set of proteins was found to overlap between cell lines^29^, highlighting that protein-protein interactions can be fluid and depend on the cell type and intrinsic cell signaling. Despite NCI-H23 and NCI-H2023 being lung adenocarcinoma cell lines, they differ in mutation signatures. While NCI-H23 is a cell line collected from a primary lung tumor, NCI-H2023 was derived from a metastasis in a lymph node^31,32^. Thus, it is possible that due to intrinsic mechanisms and activated downstream signaling divergence, PTPRH may alter interaction partners to regulate the various pathways as required.

**Figure 2.**
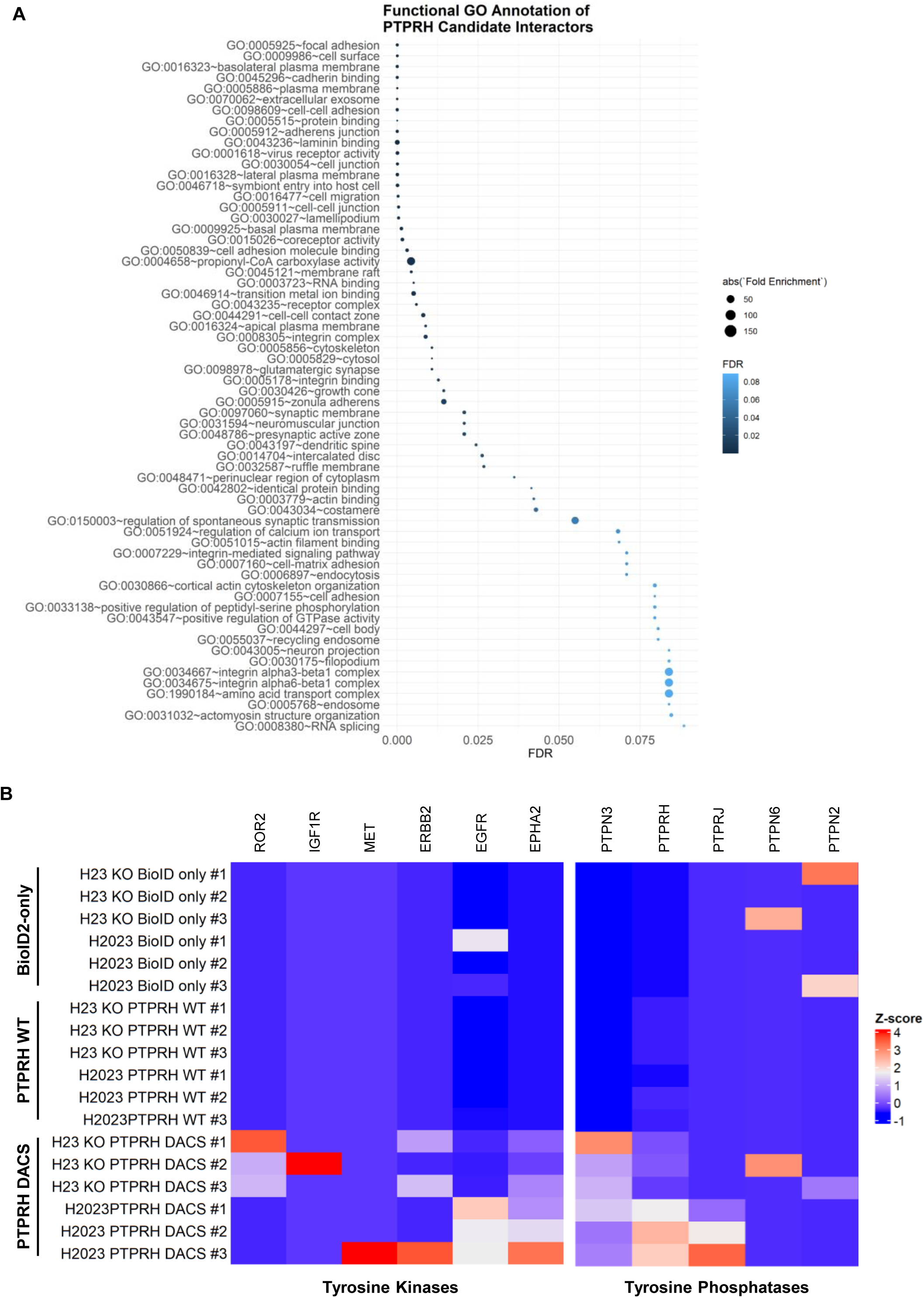
Functional annotation of PTPRH candidate interactors. *A)* Using DAVID, the PTPRH candidate interactors identified in the cell lines individually or in combination (n=120 proteins) were assessed for enrichment of gene ontology (GO) terms. The terms with FDR<0.1 and their fold enrichment absolute values are shown. *B-C)* Heatmaps showing the z-scores of normalized total precursor intensity of tyrosine kinases (B) and tyrosine phosphatases (C) across replicates from BioID2-HA-only, PTPRH WT-BioID2-HA, and PTPRH DACS-BioID2-HA groups.

In the shared targets between cell lines, the assay identified PTPRH itself, the actin motors myosin MYOF and MYO1C, the adhesion molecule CXADR, Neurofibromin 2 (NF2), which is involved in cell-cell adhesion and proliferation, ERBB2 Interacting Protein (ERBIN), and several others. Receptor tyrosine kinase phosphatases like PTPRH are thought to form dimers; thus, it was not surprising that PTPRH was the top-enriched protein in the analyses (2,100,000 fold change, p=0.0089), suggesting that this phosphatase is found as a homodimer in our infected cells^19,33^. Moreover, other candidate interactors are integrin subunits (ITGB1, ITGA2, ITGA3, ITGA6), a member of the MAPK signaling (MAP4K4), the tyrosine phosphatases PTPN3 and PTPN13, the cell adhesion-associated molecules CD44 and p120-catenin (CTNND1), and other prominent pathways.

We next sought to understand whether the candidate proteins interact with each other and form complexes of biological relevance. STRING analysis of the 120 unique interactors revealed one major network and a few smaller ones, with proteins such as EGFR, CAV1, and ITGβ1 being central (Fig. 1E and Supplemental Fig. 3). Importantly, DAVID functional annotation analysis using Gene Ontology (GO) terms revealed that the most enriched pathways are related to focal adhesion, cadherins, binding to integrins and actin, cell-cell adhesion and junction, cytoskeleton organization, the plasma membrane, and even RNA binding and splicing (Fig. 2A and Supplemental Fig. 4B-D).

We also examined the data for evidence of interactions with tyrosine kinases and other protein tyrosine phosphatases. Several groups, including our own, have previously reported that PTPRH regulates the EGFR signaling pathway^8–10^. Consistent with these findings, EGFR emerged as a significant interactor in H23 KO cells, showing a 17,000-fold enrichment (p = 0.0031) (Fig. 2B, Supplemental Fig. 4A, and Supplemental Table 1). Although H2023 cells exhibited a 3.1-fold enrichment for EGFR, the result did not reach statistical significance (p = 0.084). Similarly, ROR2, a tyrosine kinase involved in the non-canonical Wnt/planar cell polarity pathway, was highly enriched in the PTPRH group (32,000-fold in H23 KO and 170-fold in H2023) but not significantly. In both cell lines, we identified significant protein-protein interactions with the tyrosine kinase EPHA2, part of the cell adhesion receptor family, and enrichment of ERBB2, IGF1R, and MET in select replicates. PTPRJ and PTPN3, two phosphatases proposed to regulate EGFR phosphorylation^34,35^, were detected in both cell lines. Taken together, these data have uncovered novel interactions and have reinforced existing interactions for PTPRH.

### EGFR phosphorylation patterns in lung cancer cells *PTPRH* knockout

Since our interactome results revealed a broad range of different pathways and biological processes in which PTPRH might play a role, we explored several pathways in greater depth. A few published observations suggested that PTPRH regulates EGFR signaling, but the full nature of this regulation is still unknown. We hypothesized that PTPRH negatively regulates EGFR. Considering that we identified significant protein-protein interactions of PTPRH with EGFR in NCI-H23 cells, we then used monoclonal H23 *PTPRH* knockout (H23 KO) cells, whose parental cell line is wild-type for *PTPRH* and *EGFR,* to explore this regulatory mechanism. These knockouts were previously generated by inserting a frameshift mutation in the fourth exon of endogenous PTPRH^8^, and the absence of the protein was validated through immunoblot (Supplemental Fig. 8).

Previous data from our lab suggested that PTPRH regulates phosphorylation of EGFR at tyrosine residue 1173 (pEGFR Y1173, also referred to as pEGFR Y1197). To explore this, we first assessed pEGFR Y1173 levels in two independent PTPRH knockout clones following 15 minutes of EGF stimulation. Densitometry analysis of the bands indicated an approximately two-fold increase in pEGFR Y1173 in one clone (H23 KO). However, when normalized to total EGFR levels, neither clone showed a statistically significant difference compared to the parental cells, although a clear upward trend was observed (Supplemental Fig. 9A). Notably, both H23 KO and H23 KO#2 clones exhibited a reduction in total EGFR protein levels. Given that EGFR phosphorylation and activation happen rapidly after ligand binding and dimerization^36,37^, we reasoned that an increase in pEGFR1173 in H23 KO cells could reach a maximum at other time points following EGF stimulation. To test this, we stimulated the parental and the H23 KO cells with EGF for 5, 15, 30, 60, and 120 minutes, and observed the largest difference in pEGFR1173/total EGFR ratios between the two groups at the 15-minute mark (Fig. 3A). Despite not being significant, an increasing trend was again evident. Concomitantly, we also evaluated the activation of AKT, a pathway downstream of EGFR, but no difference in pAKT/total AKT ratios was noted.

**Figure 3.**
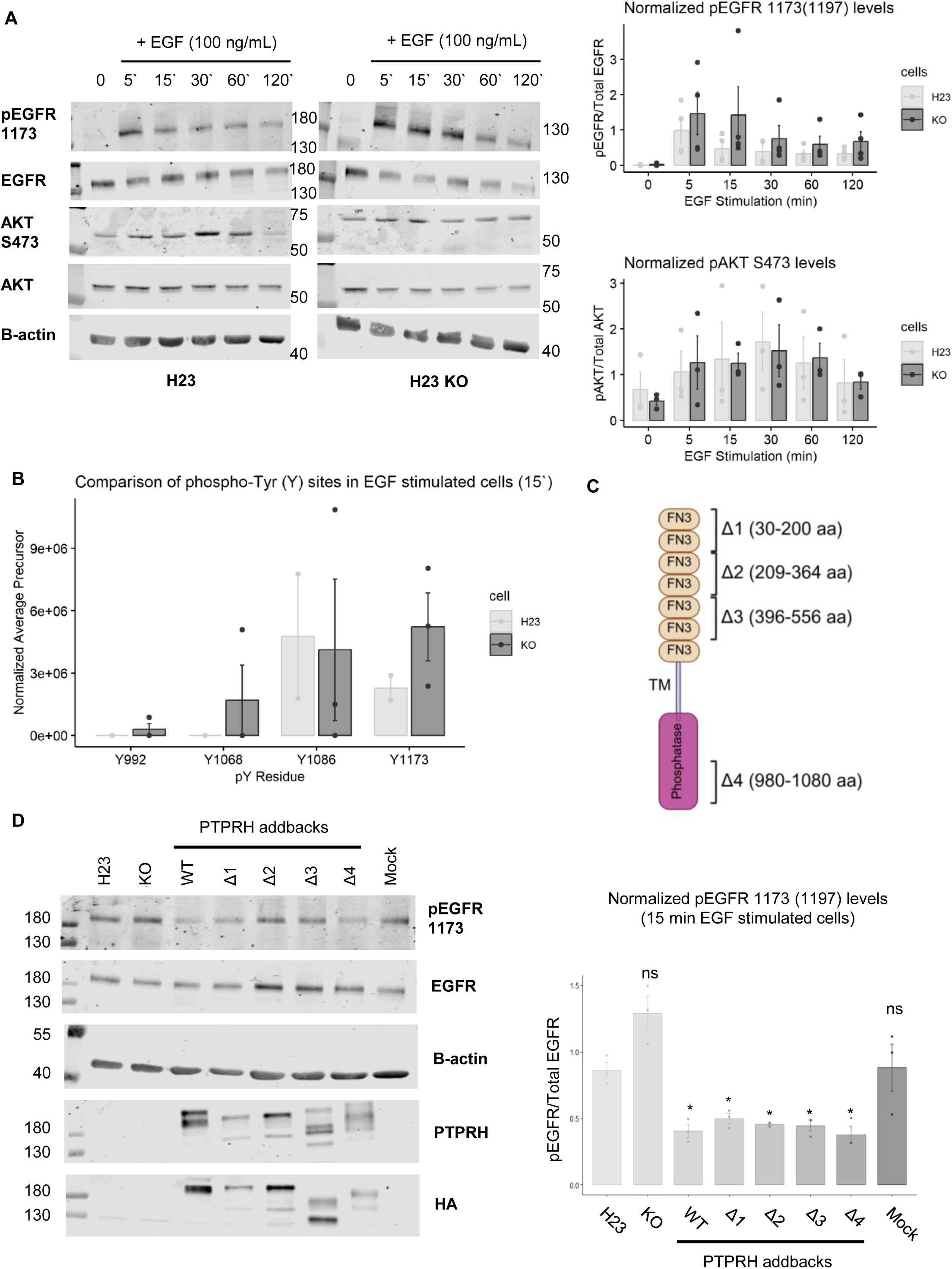
Investigation of pEGFR 1173 (1197) levels in H23 *PTPRH* knockout cell lines and after rescuing PTPRH expression with various mutant constructs. *A)* H23 and H23 KO cells were stimulated with EGF (100 ng/mL), and cell lysates were collected after 5, 15, 30, 60, and 120 minutes. Timepoint 0 represents non-stimulated (untreated) cells. Differences in pEGFR 1173 (1197) (*n*=4/group in each time point) and pAKT S473 (*n*=3/group in each time point) levels were assessed by immunoblot, and representative blots and densitometry are shown. Blots for both cell lines were imaged together to allow for comparisons. *B)* EGFR was immunoprecipitated from H23 (n=2) and H23 KO (n=3) cells stimulated with EGF (100 ng/mL) for 15 minutes, and phosphorylation status was analyzed by MS/MS. Quantitative values of normalized total precursor intensities for the identified phospho-Tyr sites in each cell line are represented in the bar plot (mean +/- s.d.). *C)* Schematic representation of the deletions generated in the PTPRH-WT-HA construct. *D)* Lung adenocarcinoma H23 (parental), H23 KO, H23 KO mock (control for lentivirus transduction), or H23 KO cells in which PTPRH WT or various PTPRH deletions were added back to overexpress the protein were stimulated with EGF (100 ng/mL) for 15 min. The whole cell lysates (*n*=3/group) were used to immunoblot pEGFR 1173 (1197), total EGFR, B-actin, PTPRH, and HA-tag, and the representative blot is shown. Densitometry results for pEGFR 1173 (1197)/total EGFR and pAKT/Total AKT ratio for the replicates are demonstrated as individual dots in the bar plot, in addition to the mean +/- s.d. A t-test was performed to compare each group with the parental cell (H23), and *p < 0.05 was considered a significant result*. Abbreviations:* ns = non-significant; Tyr (Y) = tyrosine

We then began to consider whether PTPRH might dephosphorylate an EGFR tyrosine residue other than 1173. To investigate this, we immunoprecipitated EGFR from the parental NCI-H23 and H23 KO cells stimulated with EGF for 15 minutes, and quantified phospho-tyrosine residues using MS/MS. Our analysis revealed that two tyrosine residues, Y1086 and Y1173, were phosphorylated in most replicates from both groups (Fig. 3B). Consistent with our western blot findings, H23 KO cells exhibited an approximately two-fold increase in pEGFR1173 levels; however, the differences in phosphorylation at the Y1086 and Y1173 sites did not quite reach significance. Notably, phosphorylation of Y992 and Y1068 was detected exclusively in the knockout cells, but only in 1 out of 3 replicates. A slight upregulation of phosphorylation in these two tyrosine residues in the same knockout clone was confirmed by western blot, although the changes were once again not significantly different (Supplemental Fig. 10A-B).

### EGFR phosphorylation at tyrosine 1173 is reduced in lung cancer cells overexpressing PTPRH wild-type or various deletions

Considering that we did not see significant changes in pEGFR when using the PTPRH knockout cells, we reasoned that this may be due to the low PTPRH protein levels in the NCI-H23 parental cells, making it challenging to compare phenotypes when the phosphatase activity is ablated. Indeed, compared to the colorectal cancer cell line HT-29, in which upregulation of PTPRH has been demonstrated^22^, the gene expression levels are dramatically different (Supplemental Fig. 8B). To address this issue, we investigated EGFR phosphorylation patterns in the H23 KO cells in which the PTPRH WT-HA construct was added back and overexpressed. In addition to the wild-type, we also generated several add-back cell lines expressing in-frame deletions in different domains of the same construct. Because the extracellular region of the phosphatase contains multiple fibronectin type III (FN3) domains, we began by deleting pairs of consecutive FN3 domains (constructs Δ1–3). The catalytic site was deleted in Δ4, and a schematic of the mutations is presented in Figure 3C and Supplemental Fig. 11B. Attempts to delete all FN3 domains simultaneously were unsuccessful, likely due to the repetitive nature of sequences within the extracellular region. All deletions were in-frame (Supplemental Fig. 11C), and expression of constructs was confirmed by immunoblot against PTPRH and the HA tag, which showed a smaller molecular weight than the WT counterpart (Fig. 3D, bottom lower panels).

As anticipated, while the H23 KO group tended to increase pEGFR 1173 ratios compared to the parental cells, the difference was not statistically significant. However, rescuing and overexpressing PTPRH WT caused a significant decrease of approximately 2-fold (Fig. 3D). Surprisingly, all PTPRH deletions yielded similar results to the wild-type addback, even when lacking the catalytic site. In addition, while Δ2 and Δ3 addbacks caused a modest upregulation of total EGFR levels and consequent increase in pEGFR1173, when pEGFR1173 levels were normalized by total EGFR expression, both led to a significant downregulation of the phosphorylated ratio. Therefore, it appeared that neither the individual FN3 domains nor the catalytic site of PTPRH was necessary for EGFR Y1173 dephosphorylation. Rather, we hypothesize that the transmembrane domain or multiple FN3 domains in combination are implicated in this regulation. Alternatively, this phosphatase might aid EGFR dephosphorylation indirectly.

We then investigated the effect of PTPRH WT overexpression in the metastatic lung adenocarcinoma cell line H2023. Contrary to the cells from the primary tumor, insertion of the exogenous PTPRH WT-HA construct did not yield significant differences in pEGFR 1173 after 5 or 15 minutes of EGF stimulation (Supplemental Fig. 9B). Taken together, these findings suggest that, although regulation of EGFR levels and phosphorylation can be affected by PTPRH, this regulation might be indirect and dependent on the nature of the cell type.

### PTPRH overexpression reduces cell-substrate adhesion and alters vinculin and pFAK levels

The top result of BioID pointed to PTPRH involvement in focal adhesions; therefore, our next step was to understand the cell signaling mechanism behind it. One report from the early 2000s has demonstrated that this tyrosine phosphatase can induce dephosphorylation of some integrin-associated molecules (focal adhesion kinase (FAK), p130^cas^, and p62^dok^) and alter the number of vinculin and actin stress fibers in focal adhesions^16^. However, this remains an open area of study as no further studies have been reported. Although we did not observe FAK, p130^cas^, or p62^dok^ in the interactome assay, we found significant protein-protein interactions with multiple integrin subunits and myosin.

We used the primary site-derived H23 KO cells or the metastatic H2023 cell line, with two pooled populations overexpressing/added-back PTPRH-WT (referenced as WT+ and WT++) and one the catalytic-deficient (DACS) to first investigate the cells’ ability to adhere to a panel of extracellular matrix (ECM) proteins and assessed activation of key downstream molecules previously linked to the phosphatase. Overall, increased levels of WT tended to reduce adhesion to fibronectin, collagen I, and collagen IV, a phenotype that extends to fibrinogen in the primary-site derived cells (Fig. 4A). However, only collagen I in H2023 cells and fibrinogen in H23 KO presented significant results. Curiously, overexpression of the catalytic-deficient version tended to increase adhesion in the metastatic cell line.

**Figure 4.**
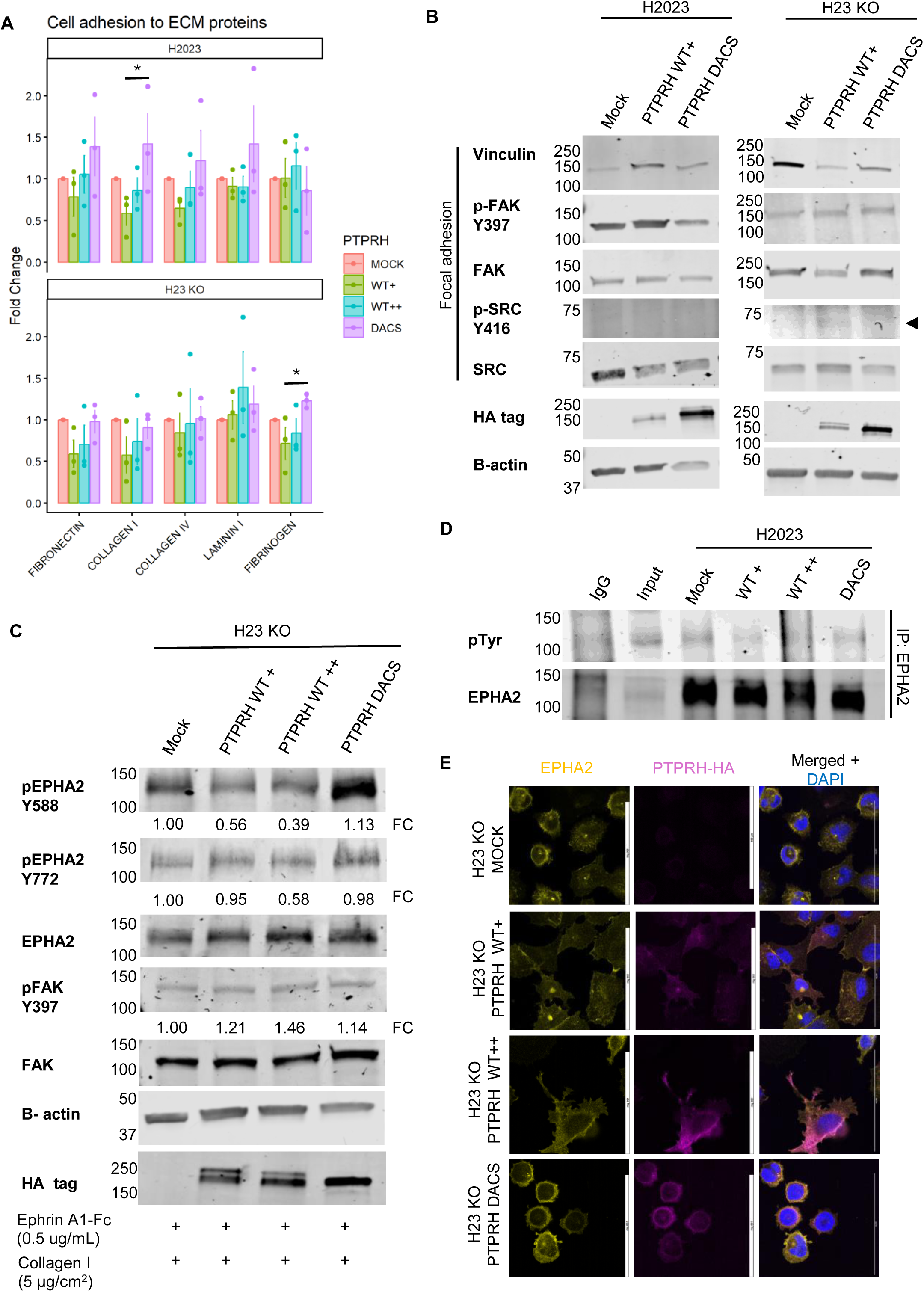
Cell signaling alteration in focal adhesions and EPHA2 phosphorylation induced by PTPRH in lung adenocarcinoma cell lines. *A)* Ability of H23 KO and H2023 cells overexpressing/added-back PTPRH WT-HA or PTPRH DACS-HA to adhere to different extracellular matrix (ECM) substrates (*n=*3/group). Fold-change for each substrate was calculated in comparison to the mock group, and values were plotted on the Y-axis. Statistical analysis was performed using One-Way ANOVA, and a *p<0.05 was considered a significant result. Replicates are demonstrated as individual dots in the bar plot, in addition to the mean +/- s.d. *B)* The same cells were serum starved for 24 hours in uncoated culture dishes, followed by stimulation with complete medium for 18 hours before cell lysis. The immunoblots are representations of protein expression and post-translational modifications of the key molecules involved in focal adhesion. *C)* Ligand-dependent EPHA2 phosphorylation and FAK activation were assessed in H23 KO cells expressing the various addbacks when plated in a thin coat of collagen I (5 µg/cm^2^), serum starved overnight, followed by stimulation with Ephrin A1-Fc (1.5 µg/mL) for 15 minutes. The densitometry of the bands (phospho/total normalized by beta-actin) is represented as a fold change (FC) in comparison to the mock group. *D)* EPHA2 was immunoprecipitated from unstimulated H2023 cells overexpressing PTPRH WT-HA or PTPRH DACS-HA, or mock (lower panel), and total phospho-tyrosine phosphorylation of the tyrosine kinase was assessed by immunoblotting (upper panel). Immunoprecipitation using IgG was used as a negative control, and whole-cell lysate (input) as a positive control. *E)* Representative confocal images of EPHA2 (yellow) and exogenous PTPRH-HA (magenta) distribution and subcellular localization. The individual channels, as well as the merged image + DAPI are shown. *Scale bar* = 100 μm, *magnification* = 100x.

In agreement with the literature, we observed that rescuing PTPRH WT in H23 KO cells caused a significant reduction in vinculin expression (Fig. 4B and Supplemental Fig. 12). Although less prominent, this reduction was also observed in the presence of DACS, suggesting that catalytic activity may not be required. Interestingly, the metastatic H2023 cells overexpressing PTPRH WT or DACS displayed the opposite trend. Furthermore, we also observed mild but significant changes in FAK. WT rescue in H23 KO cells increased the phospho-FAK ratios, a marker for FAK activation. To our surprise, an increase in phospho-FAK was also present in H2023 DACS cells. Despite this, no significant changes were observed in c-SRC activation, another tyrosine kinase downstream of the integrins; hence, the modest levels of activated FAK might not be enough to ultimately trigger c-SRC activity. Given the reduced cell adhesion to the ECM, we hypothesize that FAK is not the main mechanism governing this phenotype, as its activation strengthens focal adhesions. Despite that, these experiments suggest that PTPRH deregulates the balance of focal adhesion molecules to some extent, and disruption of structural proteins like vinculin or even myosin might dictate cell adhesion. Moreover, the catalytic activity and origin of the cell type (primary cancer site vs metastatic) also seem to influence the functional outcome.

### PTPRH targets EPHA2 for dephosphorylation

EPHA2, a tyrosine kinase that arose as a strong PTPRH interactor in both cell lines, plays a regulatory role on focal adhesions, governing cell adhesion to the substrate, cell spreading, morphogenesis, and migration^38–40^. EPHA2 is unique amongst the kinases, as its activation can be triggered by its ligand ephrin-A or be ligand-independent. The ligand-dependent activation, marked by phosphorylation of Y588 and Y772, can either limit integrin signaling and FAK activity or increase focal adhesions depending on the context, revealing a complex regulatory mechanism. Furthermore, EPHA2 is reported to participate in cell-cell junctions^41^. Hence, we sought to understand whether PTPRH regulates cell adhesion by controlling EPHA2 phosphorylation.

Given that one of the ECM substrates that seemed to affect cell adhesion the most is collagen I, we analyzed EPHA2 phosphorylation (pEPHA2) on cells that were plated onto culture dishes coated with collagen I and stimulated with ephrin A1-Fc at a lower (0.5 μg/mL) and higher (1.5 μg/mL) concentration for 15 minutes. In both conditions, rescuing PTPRH WT, but not DACS, in H23 KO cells diminished pEPHA2 ratios at the ligand-dependent tyrosine residue Y588 (Fig. 4C and Supplemental Fig. 13A-B), consistent with its role as a tyrosine phosphatase. Stimulation with a higher concentration of the ligand also slightly decreased pEPHA2 at residue Y772. In H2023 cells, this reduction was observed only in one of the pooled populations overexpressing PTPRH WT, independent of the ligand concentration. Interestingly, and reproducing the behavior observed previously, overexpressing the catalytic-deficient version in these cells promoted the opposite effect and consistently increased pEPHA2 Y588 ratios. To test if the other tyrosine residues could be potential PTPRH targets in this cell line, we pulled down EPHA2 in cells grown until confluency and blotted for total tyrosine phosphorylation. Marked EPHA2 dephosphorylation was observed in the WT+ and WT++ groups compared to mock or DACS (Fig. 4D), confirming that PTPRH targets the tyrosine kinase for dephosphorylation. In agreement with our previous observations, all addback groups slightly elevated pFAK Y397 ratios, including the catalytic-deficient mutant. Therefore, PTPRH-mediated dephosphorylation of EPHA2 might not be solely responsible for FAK activation in our model.

To validate the proximity of the phosphatase to EPHA2, we imaged the subcellular localization of both proteins in the various cell lines using similar conditions as before. Remarkably, the PTPRH-HA and EPHA2 channels presented similar distribution patterns when cells were grown in the presence of collagen I with or without ligand stimulation (Fig. 4E and Supplemental Fig. 14). Proteins were mainly localized in the plasma membrane and cytosol, and several colocalization points were noticed, strongly suggesting that EPHA2 is indeed a substrate for PTPRH.

### Overexpression of PTPRH leads to changes in cell morphology and cytoskeleton organization

Both focal adhesions and EPHA2 phosphorylation cause actin polymerization and rearrangement, leading to changes in the cytoskeleton dynamics, cell morphology, and migration^42,43^. Therefore, we assessed whether changes in the cytoskeleton organization occurred in the cells when PTPRH levels were upregulated. We performed three independent experiments and imaged the F-actin filaments of cells grown in a thin coating of collagen I or stimulated with Ephrin A1-Fc for 5 or 15 minutes. Long filaments linearly organized across the cell body (orange arrows) or at the cell periphery can be seen in several H23 KO cells with rescued PTPRH-WT expression in all conditions (Fig. 5A and Supplemental Fig. 17). In contrast, in control cells (mock) or expressing PTPRH DACS, the F-actin filaments are mostly aggregated around the plasma membrane or densily aggregated within the cytosol, and shorter filaments randomly oriented are spotted throughout the cell body in most cells (white arrows). These differences, however, were less apparent in H2023 cells (Fig. 5B and Supplemental Fig. 17).

**Figure 5.**
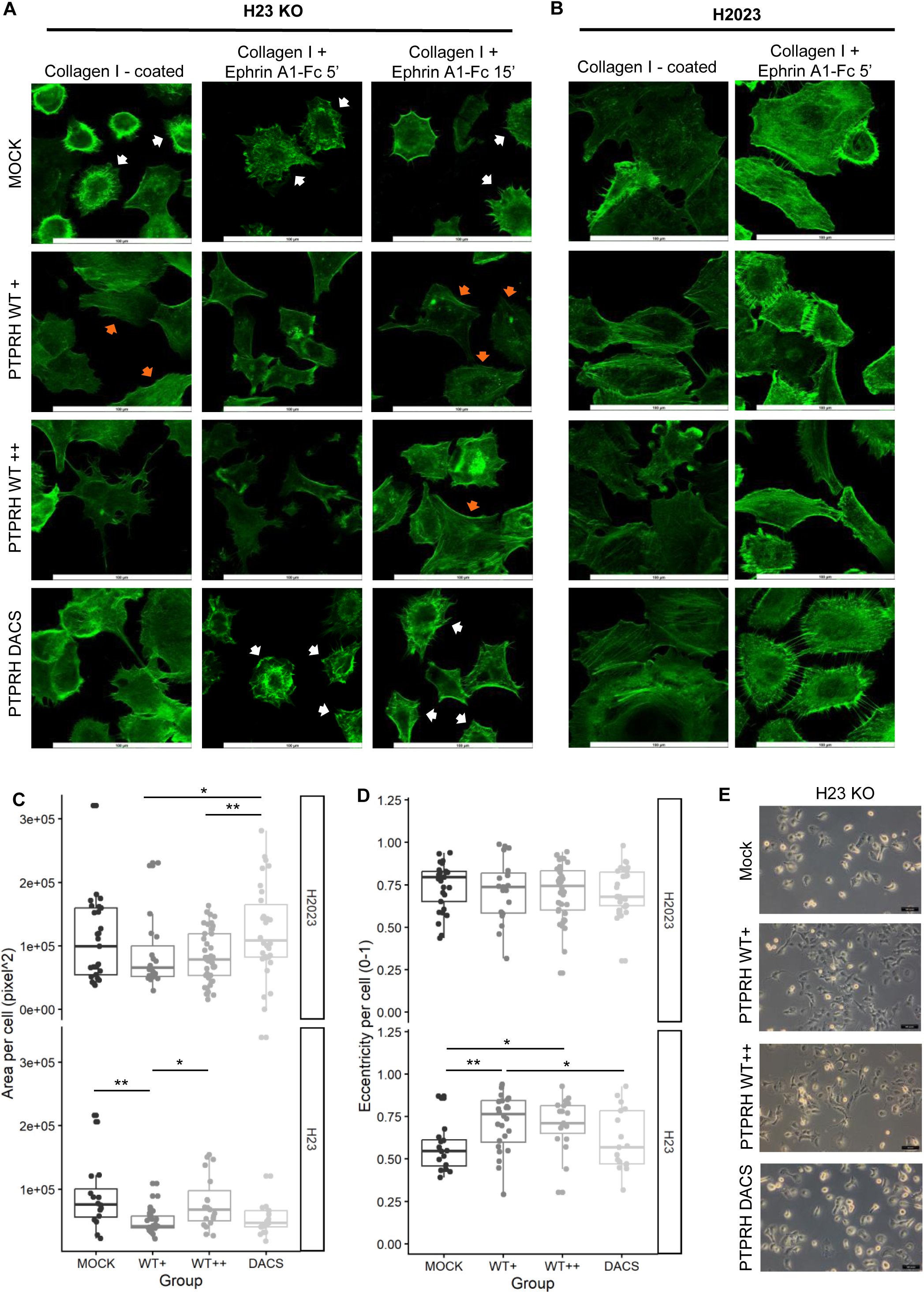
Morphology and cytoskeleton organization of lung adenocarcinoma cell lines overexpressing PTPRH WT or DACS. Images representing cytoskeleton organization and cell morphology of H2023 and H23 KO cells in which PTPRH WT-HA or PTPRH DACS-HA was overexpressed when plated in a thin coat of collagen I (5 µg/cm^2^) with or without stimulation with Ephrin A1-Fc (1.0 µg/mL) for 5 or 15 minutes. Cell shape parameters were extracted based on the confocal images represented in *A* using *Cell Profiler.* The mock cells for each cell line were used as a control*. A-B)* Representative images of F-actin filaments organization. Fields were chosen randomly. Orange arrows indicate cells where long filaments can be observed across the cell body or in the periphery. White arrows demonstrate cells with short, randomly distributed filaments. *Scale bar =*100 μm, *magnification* = 100x. *C)* Area and *D)* eccentricity values in pixels of H23 KO- and H2023-derived cells. Eccentricity is given in the 0-1 range, in which 0 = perfect round shape (circle) and 1= line segment. *E)* Bright field images of H23 KO-derived cells in culture (uncoated dishes). *Scale bar =*500 pixels, *magnification* = 10x. Statistical analysis was performed using One-Way ANOVA. Dots in the plots represent individual cell values, and the mean +/- s.d is shown. *p<0.05, **p<0.01, and ***p<0.001

To accurately quantify morphological changes, we applied Cell Profiler on three image batches of cells grown on uncoated glass slides or coated with collagen I +/- EPHA2 stimulation as indicated. This algorithm outputs several morphological features, but we focused our analysis on the well-described ones, such as cell area, perimeter, eccentricity, solidity, compactness, and mean radius. While some feature measurements varied between the batches (Supplemental Fig. 15), we identified a few common alterations, which were generally independent of the condition. In the H23 KO cells, rescuing PTPRH WT expression resulted in a loss of eccentricity, with cell spreading becoming visually noticeable (Fig. 5C). Given that a characteristic of ligand-dependent EPHA2 phosphorylation is cell rounding^44–46^, this observation fits with the decreased phospho-EPHA2 levels. The eccentricity is also evidenced by bright field images of live cells cultured in a monolayer, and long and thin cell protrusions were primarily observed in the PTPRH WT+ or WT++ groups (Fig. 5E and Supplemental Fig. 16). Furthermore, these cells have decreased cell size (Fig. 5D), and a reduced perimeter and compactness were observed in most batches (Supplemental Fig.15). Some of these morphological features also applied to the DACS group, although the changes were less prevalent and often non-significant. For the H2023 metastatic cell line, the only consistent morphological change was in cell area. Interestingly, overexpression of PTPRH-DACS leads to wider cells and augmented perimeter, while the WT counterpart causes the cell area to fluctuate in size, with smaller cells and perimeter being observed in two of the batches (Fig. 5D and Supplemental Fig. 15). Hence, we demonstrated that PTPRH can induce changes in the organization of the cytoskeleton, which is accompanied by alterations in the cell shape.

### PTPRH downregulates several adhesion, oncogenic, and metabolic pathways

To further shed light on pathway alterations induced by PTPRH modulation in cancer cells, we performed RNA-sequencing on the same series of lung adenocarcinoma cell lines overexpressing PTPRH or *PTPRH* knockout. Principal component analysis (PCA) clustered cells primarily based on PTPRH status (PC1- 82% variance)(Fig. 6A). PC2 reflects 9% of the variance and further separate the clusters based on the transcriptome of the original cell line. Surprisingly, the H23 KO cells cluster closely to the H23 PTPRH overexpression cells (Supplemental Fig. 5A). This may reflect a drift of phenotype caused by the ablation of PTPRH after a few passages.

**Figure 6.**
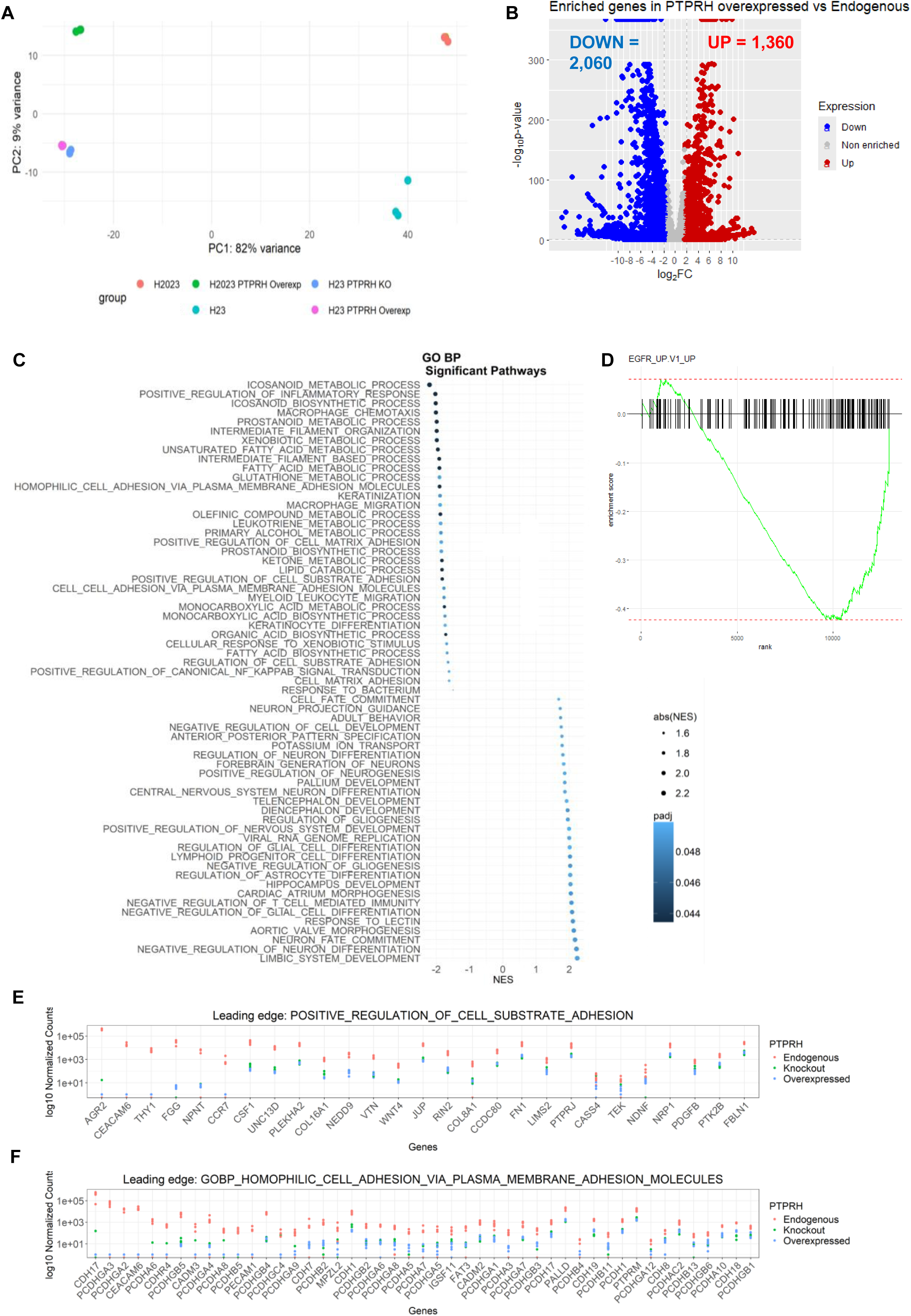
Transcriptome analysis of lung adenocarcinoma cell lines overexpressing PTPRH wild-type. H2023 and H23 cell lines expressing endogenous or overexpressed PTPRH WT levels and H23 *PTPRH* knockout cells were serum-starved for 24h, followed by stimulation with FBS for 16h prior to RNA extraction (n=3 replicates/group). *A)* PCA plot of counts normalized by variance stabilizing transformation showing PC1 and PC2. *B)* Volcano plot of differentially expressed genes in the PTPRH overexpression cell lines compared to cell lines expressing endogenous PTPRH (parental cells) with padj < 0.05. Genes were considered upregulated (red) when fold change > 2 and downregulated (blue) when log2 fold change < -2. *C)* GSEA plots showing significantly enriched pathways in the PTPRH overexpression cell lines using the Gene Ontology (GO)(C5 MSigDB) gene set. *D)* Enrichment GSEA plot for EGFR_UP.V1_UP pathway in PTPRH WT overexpressing cell lines. *E-F*) Normalized counts of leading-edge genes in the indicated pathways when PTPRH levels are modulated, ordered by log2 fold-change.

Differential expression analysis demonstrated 1,360 upregulated genes and 2,060 downregulated genes using a 2-fold change filter in PTPRH overexpressing cells compared to parental cells (Fig. 6B, Supplemental Fig. 5C, and Supplemental Table 2). These represent PTPRH signature genes, and the top 30 genes up- and downregulated are shown in Supplemental Fig.7A. To understand how the differentially expressed genes (DEG) modulated by PTPRH are involved in biological pathways and cancer, we performed pathway enrichment analyses using the Molecular Signature (MSigDB) and GO databases. Reinforcing our previous results that PTPRH regulates cell adhesion, Gene Set Enrichment Analysis (GSEA) using the GO BP database revealed that the most downregulated processes were those related to cell-cell and cell-substrate adhesion, as well as migration, intermediate filament organization, and multiple metabolic pathways (Fig. 6C). In contrast, pathways related to development, differentiation, and cell fate were upregulated. To uncover which adhesion molecules responded more strongly to changes in PTPRH level, we then plotted the normalized counts of the leading-edge genes linked to positive regulation of cell-substrate adhesion and homophilic cell adhesion via the plasma membrane (Fig. 6E-F). 27 genes linked to cell-substrate adhesion, such as collagens (COL16A1, COL8A1), fibronectin I (FN1), CEACAM6, and 47 genes linked to cell-cell adhesion, such as a wide variety of protocadherins and cadherins, presented a significant decrease in gene expression in cells overexpressing the phosphatase.

**Figure 7.**
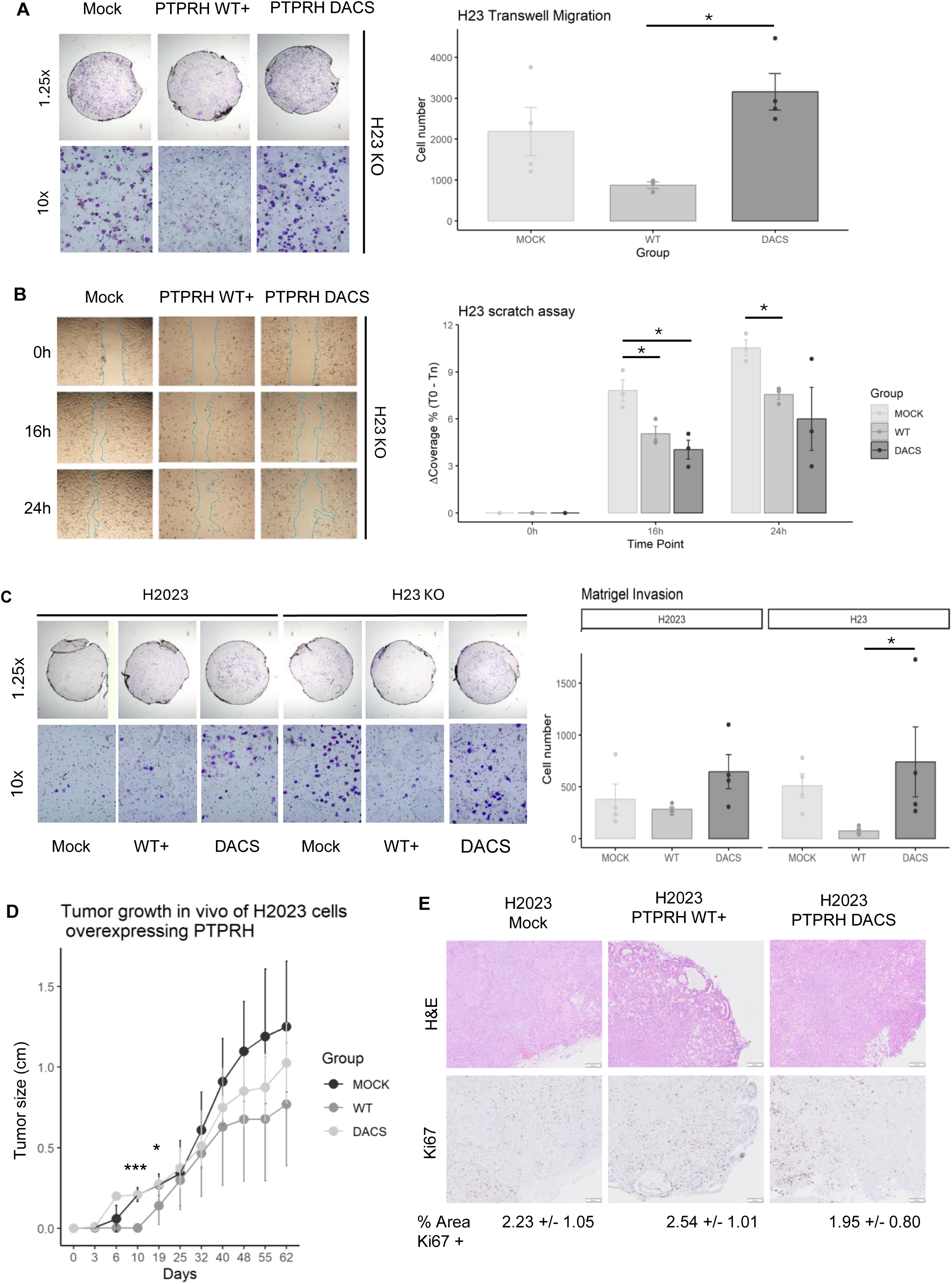
Functional validation of PTPRH role in migration, invasion, and tumor suppression. The ability of cells to migrate or invade *in vitro* upon PTPRH WT-HA or DACS-HA overexpression in H23 KO and H2023 cells was assessed through different functional assays. *A)* Transwell migration after 24 hours. Membranes from mock, WT, and DACS groups were stained with crystal violet for visualization of the migrating cells, imaged using 1.25x and 10x magnification. The total number of migrating cells in each replicate per group is demonstrated in the bar plot (n=4 replicates/group), and images are representative. *B)* Scratch was imaged at time point 0, and migration of the cells towards the scratch area was monitored after 16 and 24 hours (n=3 replicates/group). The covered area in the scratch over time (T_n_) for each replicate was calculated using the time point 0h (T_0_) as a reference and given as a percentage in the bar plot*. C)* Cells were allowed to invade through a Matrigel-coated membrane for 24 hours before crystal violet staining of the membranes and quantification of invading cells (bar plots). Membranes were imaged using 1.25x and 10x magnification (n=4 replicates/group). *D)* Tumor growth in the flank of NOD-SCID mice injected with H2023-derived cells was monitored over time, and the tumor size in *cm* is demonstrated. The experiment started with n=5 mice/group. On day 40, one mouse from the mock group and three from the PTPRH-DACS group were found dead. *E)* Representative histology (H&E staining) and Ki67 immunohistochemistry of flank tumors overexpressing PTPRH WT-HA or DACS-HA using 20x magnification (scale bar = 200 μm). The percentage of total area that presented Ki67 positive staining and quantification of migration and invasion assays are given as the mean +/- standard deviation. Statistical analysis was performed using One-Way ANOVA. *p<0.05, **p<0.01, and ***p<0.001

Importantly, multiple oncogenic pathways were also significantly downregulated in these cells, including NFE2L2, KRAS, P53, ERBB2, and others (Supplemental Fig. 7B). We also observed significant downregulation of genes involved in the EGFR (NES = - 1.55) and AKT (NES = -1.53) pathways, in agreement with our hypothesis and previous findings that PTPRH negatively regulates EGFR and downstream signaling pathways (Fig. 6D). Further, downregulation of apical junctions, estrogen response, and metabolism (adipogenesis, xenobiotic, and fatty acid metabolism), etc., and upregulation of proliferation pathways and the WNT-Beta-catenin signaling were also noted in the hallmark MSigDB geneset (Supplemental Fig. 7C).

We next asked whether PTPRH overexpression would modulate the expression of other protein tyrosine phosphatases (PTPs) and tyrosine kinases (TKs). We interrogated the RNA-seq data based on the genes that were present in a DEG analysis and found that 34 PTPs and 45 TKs were differentially expressed in the PTPRH-WT overexpressing cell lines (Supplemental Fig. 6A-B). Overall, 24 PTPs were downregulated, suggesting redundancy in function among the PTPs. Interestingly, most TKs presented increased gene expression levels (n = 28), while 17 were found to be downregulated.

Additionally, given that PTPRH may be involved in RNA-splicing, we reanalyzed our transcriptomic data using a splicing-aware pipeline^47^. 55 differentially expressed transcripts varied in isoform abundance across the various PTPRH modulation strategies (Supplemental Table 3), reinforcing that the phosphatase may play non-canonical roles beyond protein dephosphorylation. An example of a differentially spliced gene is MPHOSPH10 (Supplemental Fig. 7D), a protein that is specifically phosphorylated during the M-phase of mitosis, leading to cell structural modifications^48^.

### PTPRH alters the migratory behavior of lung cancer cells

Considering the deregulated cell signaling pathways that were observed with gain or loss of function of PTPRH in the lung cancer cells, we asked whether the combination of these alterations would be sufficient to suppress or accelerate various cancer hallmarks. The conflicting findings around PTPRH suggest that this phosphatase can display both phenotypes depending on the cellular context and tissue. Given that changes in cell adhesion and cell shape can affect cellular motility, we began by measuring the migratory behavior of the cells. Rescuing PTPRH WT expression in H23 KO cells consistently suppressed migration after 16 or 24 hours in wound healing and transwell assays (Fig. 7A-B and Supplemental Fig. 18-19). Interestingly, rescuing PTPRH DACS expression also reproduced similar wound healing behavior, but failed to decrease transwell migration. In the metastatic cell line H2023, overexpression of the wild-type protein inhibited migration in 2 out of 4 experiments compared to mock cells, while the remaining trials showed no statistically significant effect. However, at all times, the PTPRH WT overexpression group lagged behind the cells overexpressing the catalytically inactive construct (PTPRH DACS).

Additionally, we evaluated the invasion through a Matrigel-coated membrane *in vitro*. Significant differences were observed between PTPRH WT and DACS rescue in H23 KO cells, with a reduction in the number of invading cells in the wild-type group (Fig.7C and Supplemental Fig. 20). However, no differences were observed in comparison to the mock control or among the H2023-derived cell lines.

### PTPRH suppresses tumor growth, dedifferentiation, and prolongs tumor latency *in vivo*

Finally, we interrogated whether PTPRH could alter tumor growth *in vivo.* For this purpose, we generated xenografts of the various cell lines using the immunodeficient NOD-SCID mouse model, and the tumor growth in the flank was monitored over time until the endpoint. Strikingly, xenografts derived from H23 KO-derived cells exhibited minimal growth and did not form palpable tumors until day 48 (data not shown). In marked contrast, palpable tumors were observed for the H2023 mock and PTPRH DACS groups on day 6 post-injection (Fig. 7D). In agreement with the lack of catalytic activity, these two groups showed comparable tumor growth curves until day 25. Notably, tumor growth in H2023 xenografts overexpressing PTPRH WT started later, on day 19, suggesting that the increased levels of the phosphatase prolong tumor latency. Tumors from this group were significantly smaller up to the end of the experiment compared to mock, and significant differences were found on days 10 and 19, suggesting that PTPRH behaves as a tumor suppressor *in vivo*. Due to the delayed growth of the H23 KO-derived tumors, all mice were sacrificed at the endpoint defined by the H2023 xenograft timeline. Necropsy revealed that, among the H23 KO groups, 2/5 mice injected with mock cells and 3/5 injected with PTPRH-DACS cells developed small tumors. In contrast, only 1/5 mice injected with H23 KO cells overexpressing PTPRH WT exhibited tumor formation (Supplemental Fig. 21A-C).

Histologically, tumors were non-mucinous with a predominant solid, high-grade growth pattern across all conditions and pushing invasive borders (Fig.7E and Supplemental Fig.21D). However, variable proportions of admixed acinar growth were noticed in the H2023 series: tumors derived from mock and PTPRH-DACS were 5-10% acinar and 90-95% solid, while PTPRH WT presented higher gland formation, with the acinar proportion ranging from 20-80% (Supplemental Table 4). Interestingly, these tumors did not present significant changes in the proliferation marker Ki67, suggesting that overexpression of the phosphatase might suppress the tumor growth by altering cell growth or tumor progression instead. Likewise, despite the mitotic rate being mildly elevated in all groups, no qualitative differences were observed after careful assessment of the tumors’ histological features. Together, these results show that PTPRH can improve tumor differentiation in the metastatic cell line and potentially lead to a better prognosis; nonetheless, conclusions should be drawn with care, as these are histologic qualitative impressions.

To confirm the tumor-suppressive feature and assess the ability of the lung adenocarcinoma cells overexpressing PTPRH to colonize the lungs, luciferase-labeled H2023 cell lines were tracked over the course of 34 days following tail vein injection in the NOD-SCID mice (Fig. 8A and Supplemental Fig. 22). Images from day 0 (30 minutes post-injection) showed a strong bioluminescence signal in the lungs of animals in all groups, confirming that the cells had reached the bloodstream and were in circulation. As expected, the cells that failed to colonize the organ were cleared out by day 8. Strikingly, the results were similar to the xenografts. On days 8 and 22, colonization was detected in the lungs of the mice that received the mock or PTPRH catalytic-dead cells. The bioluminescence total flux between these days also increased similarly for the two groups, suggesting that the cells were proliferating over time. Meanwhile, the total flux from the PTPRH WT overexpression group was lower and similar to the negative controls. Unfortunately, halfway through the experiment, one mouse from each group (except mock) was lost due to unknown causes. Despite that, conclusions for the later time points could still be drawn from the remaining animals. At the endpoint (day 34), only one mouse from the PTPRH WT overexpression group presented a detectable bioluminescence and smaller total flux than mock or DACS, suggesting that the phosphatase inhibits the ability of cells to colonize the lungs (Fig. 8B).

**Figure 8.**
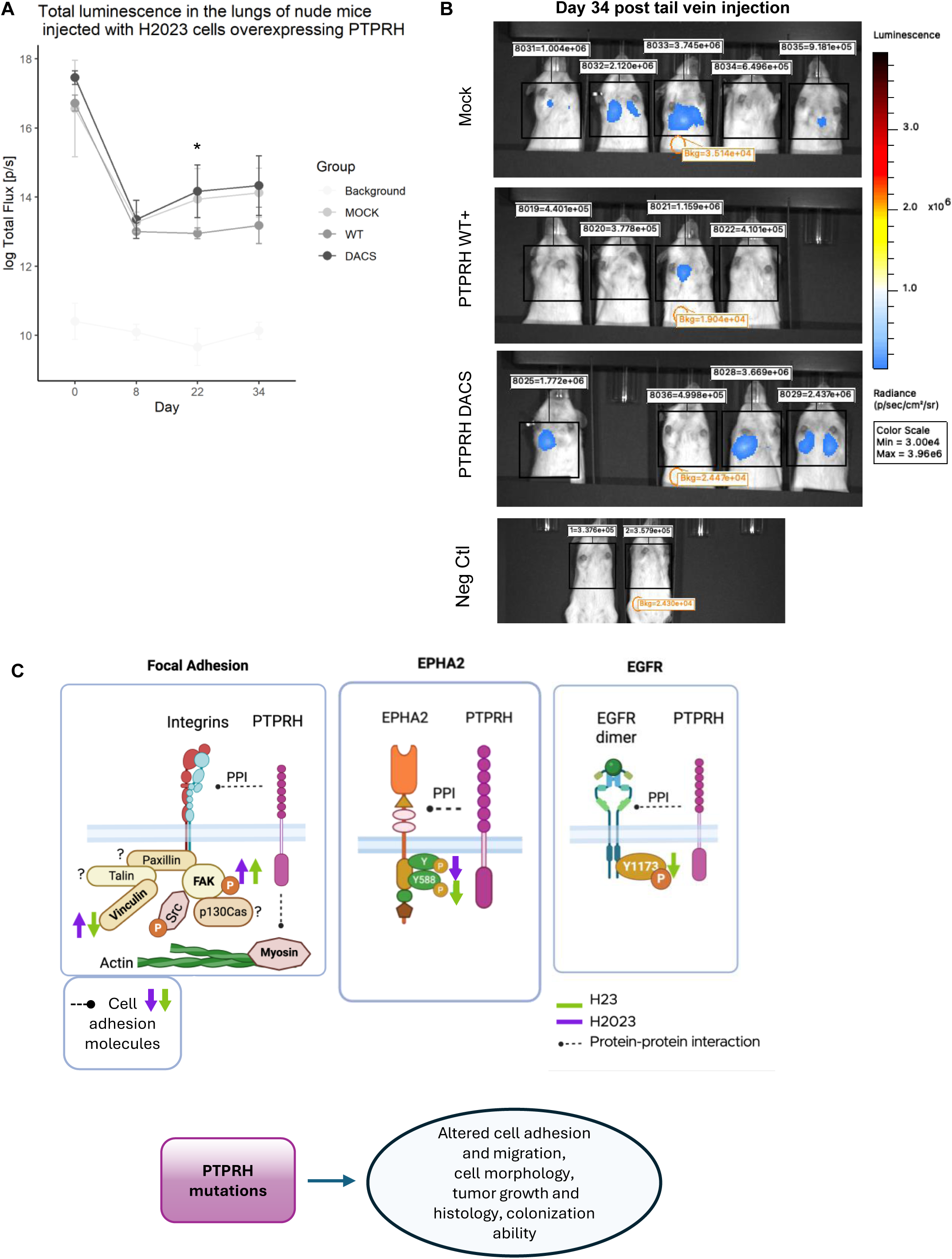
Lung colonization and cell signaling alterations induced by PTPRH overexpression in the primary and metastatic-derived lung adenocarcinoma cell lines. *A)* NOD-SCID mice were injected in the tail vein with various H2023-derived cells labeled with mCherry and luciferase. The total bioluminescent signal in the chest was measured for each mouse in each group at four time points, and their group average and standard deviation are represented in the plot as the log of total flux (p/s= photons per second) over time. Statistical analysis was performed using One-Way ANOVA. *p<0.05, **p<0.01, and ***p<0.001. *B)* Images showing the bioluminescence signal in the lungs of each NOD-SCID mouse 34 days post-injection. Their individual total flux (p/s) was calculated within the regions of interest (black boxes), and the values are demonstrated above each animal following their IDs. The total flux of a background area (orange circles) and the lungs of two negative control mice (not injected with cells) are also shown in the picture. *C)* Summary picture representing some of the key cell signaling alterations induced by overexpression of PTPRH wild-type in NCI-H23 and NCI-H2023 cells. Protein-protein interactions with integrins, myosin, EGFR, EPHA2, and cell adhesion molecules highlight PTPRH’s participation in adhesion processes and the EGFR pathway. In the primary tumor-derived cells (H23 cells), PTPRH overexpression is associated with reduced vinculin expression, a slight increase in pFAK, reduced EPHA2 phosphorylation at the ligand-dependent tyrosine site Y588, and decreased phosphorylation of EGFR at tyrosine 1173 (pEGFR Y1173). In the metastatic cell line (H2023), the phosphatase might behave differently in some aspects, with both vinculin and pFAK increasing upon PTPRH overexpression. EPHA2 tyrosine phosphorylation is also reduced, although the precise sites are yet to be uncovered. Interestingly, alterations in pEGFR and pFAK seem to be independent of PTPRH’s catalytic activity. Furthermore, multiple cell-cell and cell-substrate adhesion molecules are downregulated upon PTPRH overexpression. The distinct signaling alterations in each cell line result in changes in cell adhesion, migration, cellular morphology, cytoskeletal organization, tumor growth and differentiation, and ability to colonize distant sites.

## Discussion

Little is known about the role of the receptor-like protein tyrosine phosphatase PTPRH in the regulation of various cell signaling pathways and how alterations may contribute to tumorigenesis. To bridge this gap, we built a comprehensive protein-protein interaction network based on stable and transient interactions and evaluated how the gain or loss of the phosphatase activity would disrupt the various biological and cellular processes in lung cancer cells from the primary tumor (NCI-H23) and a metastatic site (NCI-H2023). Here, we report novel candidate interactors, roles, and mechanisms of PTPRH in pathways linked to cell-substrate adhesion, cell-cell junctions, differentiation, metabolism, tumor suppression, and others, and confirm its participation in regulating EGFR and EPHA2 phosphorylation levels (Fig. 8C).

We identified 120 unique candidate interactors that significantly interact with PTPRH in our BioID assay. While 28 of these were commonly found in the two non-small cell lung adenocarcinoma cell lines, in agreement with previous studies, we found that some protein-protein interactions were specific to one or the other cell type, suggesting that the phosphatase can behave differently according to cellular context (e.g., primary vs metastatic tumor). Despite this variance, the PTPRH interacting proteins are heavily involved in cell adhesion, cell-cell junctions, the cytoskeleton, and the plasma membrane processes in both cell lines. Furthermore, although some proteins did not reach statistical significance, they were exclusively enriched in the PTPRH-transfected group; therefore, we do not exclude the possibility that these might also be interactors or substrates. Interestingly, we observed multiple protein-protein interactions with signaling molecules in the integrin-associated pathway (i.e., focal adhesion) and cell-cell junctions. In addition, novel pathways linked to RNA (binding, processing, and splicing) were enriched at both the protein and RNA levels and validated through quantification of transcript isoforms. However, how PTPRH may participate in RNA splicing remains to be elucidated.

Very few studies on PTPRH have been published so far. To our knowledge, only one, dated from the early 2000s, has suggested PTPRH to be a regulator of focal adhesions^16^. Since then, this has remained an open area of study, as the mechanisms behind it are largely unknown. Using human pancreatic carcinoma cells, Noguchi *et al*.have shown that overexpression of the phosphatase disrupted cellular responses mediated by integrins, such as cell growth and motility, by dephosphorylation of p130^cas^, FAK, and p62^dok^^16^. While p130^cas^ was proposed to be a potential substrate of the phosphatase, dephosphorylation of FAK and p62^dok^ seemed to be indirect, as none of these proteins were co-immunoprecipitated with PTPRH. Indeed, in our experiments, neither one of them was identified as a candidate interactor. Instead, members from the ITGA and ITGB superfamily and myosin were present, suggesting that in lung cancer cells, PTPRH mediates regulation of focal adhesions by interaction with the integrins, structural proteins, or even EPHA2. Interestingly, our results demonstrated that vinculin and active FAK can be affected when the phosphatase expression is increased under basal conditions or when stimulated with collagen I and the EPHA2 ligand. The nature of this modulation, however, was modest, varied in the primary tumor compared to metastasis, and may be independent of the phosphatase activity. In agreement with our findings in NCI-H23 cells, Noguchi *et al* also reported disruption of actin fiber organization and reduced vinculin in mouse embryonic fibroblasts when PTPRH wild-type is overexpressed.

Herein, we report for the first time that PTPRH interacts and dephosphorylates EPHA2 ligand-dependent tyrosine sites, as these two proteins occupy similar locations in the cell. Several lines of evidence support that, while activation of the ligand-dependent pathway (canonical) is tumor-suppressive, activation of the ligand-independent pathway (non-canonical), characterized by Ser897 phosphorylation by AKT, is oncogenic and facilitates tumor formation, progression, and chemoresistence^49,50^. In the ligand-dependent pathway, EPHA2 is activated by binding to its ligand ephrin A1, which is anchored to the cell surface of neighboring cells^51^. Following this, EPHA2 dimerizes and is phosphorylated at tyrosine residues Y588 and Y772, located in the activation loop. In one proposed model, its activation inhibits integrin signaling and limits FAK activity, which, in reciprocity, phosphorylates EPHA2 Y772 in a ligand-independent manner, creating a negative feedback loop^52^. In another, ligand-dependent activation increases focal adhesion number, upregulates pFAK, and induces cellular contractions dependent on myosin, directing cellular motility^40,53^. Hence, the outcome of EPHA2 pathway activation is dependent on the cellular context. In our study, it seems that this relationship follows a negative feedback, with decreased pEPHA2 leading to increased pFAK. Nonetheless, the fact that the catalytic-deficient/substrate trapping group also has the potential to activate FAK reveals that this dynamic is fluid and more complex than we anticipated.

We demonstrated that the cytoskeleton reorganization and morphological changes in PTPRH-overexpressing cells, which include reduced area but elongated and asymmetrical shape in the primary site-derived cells, could result from interference with cell adhesion pathways, reduced levels of EPHA2 phosphorylation, or another mechanism yet to be identified. Increased phosphorylation of ligand-dependent EPHA2 tyrosine sites is generally associated with Rho-dependent contraction of actin/myosin, leading to cell rounding and repulsion^44–46^. Changes in cell morphology induced by PTPRH have been previously seen in Chinese ovarian cancer cells^16^ and hepatocarcinoma cell lines^25^. In the latter, PTPRH overexpression caused cells to appear flattened and less polarized. Notably, those studies relied solely on visual assessment without employing quantitative methods, unlike the unbiased, quantitative approach used in our analysis. Additionally, both studies found that PTPRH overexpression suppresses cell migration, in agreement with our findings. Cellular migration is a multifactorial process that requires coordination among adhesion molecules, focal adhesion assembly and disassembly, and cytoskeleton rearrangement to provide traction^54^. Hence, we believe that the limited cell motility can be attributed to decreased cell-substrate and cell-cell adhesion strength, as demonstrated by our *in vitro* assays and transcriptomics data, and cell shape and cytoskeleton alterations. Diminished pEPHA2 and increased FAK activity could potentially augment cell adhesion. Despite this controversy, we propose a model in which the downregulation of multiple genes in adhesion pathways, such as in the cadherin family and ECM proteins, in association with disrupted morphology, ultimately prevails and determines the functional outcome.

Multiple lines of evidence suggest that PTPRH participates in the regulation of the EGFR pathway^8,10,17,55^. In the HER2-positive breast cancer cell line SKBR-3, PTPRH was shown to be only partly involved in EGFR dephosphorylation, raising the hypothesis that more than one phosphatase is in charge of controlling the inactivation of the EGFR pathway^55^. Supporting this, we noted a significant protein-protein interaction of PTPRH with EGFR in the NCI-H23 cells. Our findings suggest that PTPRH affects EGFR levels and phosphorylation status at tyrosine 1173 (1197) position in that same cell line, in agreement with our previous publication^8^, with the maximum phosphorylation peak happening 15 minutes after EGF stimulation. However, no significant differences were observed when phospho-EGFR was assessed in the metastatic NCI-H2023 cells. Surprisingly, it appeared that none of the fibronectin domains in groups of two or the phosphatase domain was required in this process. Therefore, we hypothesize that other regions of the phosphatase might mediate the dephosphorylation, and that PTPRH participates in this reaction indirectly or with the aid of other phosphatases. PTPRO, for example, negatively regulates EGFR through direct interaction with SRC rather than with the tyrosine kinase^56^. Several tyrosine phosphatases have been described as EGFR regulators, including *PTPN2*^57^, *PTPN6* (SHP-1)^58,59^, *PTPN11* (SHP-2)^60^, *PTPN12*^61^, *PTPRK*^62,63^, *PTPRJ*^64,65^, and PTPN3^35^. Interestingly, PTPRJ dephosphorylates EGFR at the same tyrosine residue as PTPRH in a pH-dependent manner^64^. As PTPRJ is strongly downregulated in our PTPRH overexpressed cell lines, and our BioID assay suggests that PTPRH interacts with PTPRJ as well as with PTPN3, we believe that these phosphatases have redundant roles or even work in conjunction.

We recognize that the literature on PTPRH is contradictory. Many factors could explain this, ranging from variations in research models, such as mouse *versus* human models, to cell lines derived from different tissues or cultivated under different conditions and at different passages^66,67^. The differences in mutation types (point mutations vs frameshift and indels)^68^ and compensation mechanisms by other protein tyrosine phosphatases can also contribute to the contradictions in the literature^69^. Herein, we observed that modulation of PTPRH expression alone caused only a few-fold difference in many experiments. We reason that this mild change might not be enough to produce a strong phenotype under all circumstances. Supporting this, studies in mice and *Drosophila* revealed that ablation of phosphatase genes leads to mild or no change in phenotype, likely due to redundancy in tyrosine phosphatase functions^23,69,70^. Indeed, in our study, we observed the up- and downregulation of several other PTPs when PTPRH is overexpressed.

In addition to EGFR and ROR2, several other tyrosine kinases presented significant gene expression changes upon PTPRH overexpression, including many Ephrin receptors, while some others were also found to interact with PTPRH, such as ERBB2, IGF1R, and MET in select replicates. More studies are needed to determine whether PTPRH can alter the phosphorylation patterns of the kinases, downstream signaling, and induce a phenotype.

Overall, we found that PTPRH behaves as a tumor suppressor and reduces tumor growth and colonization to the lungs, prolongs tumor latency, and stimulates tumor differentiation *in vivo*. Indeed, it has been shown that when PTPRH is ablated in the NCI-H23 cells, it accelerates tumor growth and proliferation *in vitro*^8^. In our study, the tumors did not present changes in the proliferation marker Ki67; therefore, we believe that the suppression of tumorigenesis is linked to changes in cell growth instead. In addition to all the alterations discussed here, the downregulation of several oncogenic signatures and hallmark pathways in NSCLC cells overexpressing PTPRH establishes a link for how this phosphatase may suppress lung cancer progression and is worthy of deeper investigation. Taken together, inactivation of the phosphatase may well be an advantageous event for lung cancer progression. This is a marked contrast in comparison to colorectal cancers, in which this phosphatase is suggested to participate in the tumorigenesis^20,23^. Given the variety of mechanisms in which PTPRH is involved and their complexity, further study of protein tyrosine phosphatases and their regulation in cancer is required.

## Methods

### Cell culture

Cells were acquired from ATCC, and H23 PTPRH knockout cells were generated as described previously^8^. NCI-H23-derived cells were cultured in RPMI medium (Corning #50-020-PC), 10% fetal bovine serum, 1% Penicillin-Streptomycin, 23.8 mM sodium bicarbonate, 25 mM HEPES, and 1 mM sodium pyruvate. NCI-H2023-derived lineages were cultured in DMEM-F12/HEPES medium (Gibco #11330032), 5% fetal bovine serum (FBS), 1% Penicillin-Streptomycin, 30nM sodium selenite, 10 nM hydrocortisone, 10 nM beta-estradiol, 0.01 mg/mL transferrin, 0.005 mg/mL insulin, and 2 mM L-glutamine. 293T cells were cultured in DMEM high glucose medium (Sigma-Aldrich #56436C), 10% FBS, 1% Penicillin-Streptomycin. Cells were incubated at 37°C and 5% CO_2._

### Cell stimulation and protein lysate preparation

For analysis of EGFR phosphorylation, recombinant human EGF (Gibco #PHG0311) was supplemented to the medium at 100 ng/mL. Cells were incubated at 37°C and 5% CO_2_ for 2, 5, 10, 15, 30, 60, or 120 minutes. To evaluate EPHA2 phosphorylation, cells were seeded in thin-coated collagen I (Gibco #A10483-01) culture dishes at 5 μg/cm^2^, serum starved overnight, followed by stimulation with complete medium + Ephrin A1-Fc (R&D systems #6417-A1-050) at the indicated concentration (0.5-1.5 μg/mL) for 15 minutes at 37°C and 5% CO_2_. When stimulation was not intended, cells were cultured in complete medium until the desired confluency.

Cells were washed twice with ice-cold HEPES-Buffered Saline (HBS), scraped from the dish, and resuspended in ice-cold HBS. The cell suspension was centrifuged at 1,000 x g for 5 minutes at 4°C, followed by lysis in ice with ice-cold Pierce RIPA buffer (immunoblot and LC/MS/MS experiments) (Thermo Scientific #89901) or IP lysis/wash buffer (immunoprecipitation experiments) (Pierce Classic Magnetic IP/Co-IP kit #88804) + Halt Protease and Phosphatase inhibitor (Thermo Scientific #78441) for 20 minutes. Cell extracts were centrifuged at 16,000 x g for 20 minutes at 4°C, and the supernatant collected. Protein quantification was performed using Pierce BCA protein assay (Thermo Scientific #23225) according to the manufacturer’s instructions.

### Immunoblotting and Densitometry

Immunoblotting was performed as described elsewhere^71^. The following dilutions were used for the primary antibodies: PTPRH 1:1000 (Abcam #ab231767), pTyr 1000 (1:2000, Cell Signaling #8954), pEGFR 1173 1:500 (Cell Signaling #4407), EGFR 1:1000 (Cell Signaling #4267), B-actin 1:1000 (Cell Signaling #4967), HA Tag 1:1000 (Cell Signaling #3724), anti-Biotin 1:1000 (Cell Signaling #5597), pAKT S473 1:2000 (Cell Signaling #4060), AKT 1:1000 (Cell Signaling #4685), Vinculin 1:1000 (Cell Signaling #13901), pFAK Y397 1:1000 (Cell Signaling #8556), FAK 1:1000 (Cell Signaling #13009), pSRC Y416 1:1000 (Cell Signaling #2101), SRC 1:1000 (Cell Signaling #2108), EPHA2 (1:1000, Cell Signaling #6997), pEPHA2 Y772 (1:1000, Cell Signaling #8244), or pEPHA2 Y588 (1:1000, Cell Signaling #12677). Bands were detected using fluorescent IRDye® 800CW goat anti-rabbit (LICOR #926-32211) secondary antibody diluted 1:10000 in TBS-Tween 0.1% and 5% BSA. Blots were imaged using the LICOR Odyssey M. Densitometry was performed using ImageJ 1.54p software as described elsewhere^72^. The grey mean values for the target proteins were corrected by the endogenous control (e.g., Beta-actin) for data normalization.

### Immunoprecipitation

Equal amounts of protein lysates were incubated with the primary antibody against EPHA2 (1:100, Cell Signaling #6997) or EGFR (1:50, Cell Signaling #4267) using end-to-end rotation at 4°C overnight. The antigen-antibody complex was then immunoprecipitated using 50μl of Dynabeads Protein A (Invitrogen #10006D) for 30 minutes using end-to-end rotation at room temperature, washed four times with the washing buffer provided in the kit, and eluted (denaturing elution) following the manufacturer’s instructions. Immunoprecipitated proteins were submitted to immunoblot or LC/MS/MS for downstream analysis.

### Plasmids

pCMV-VSV-G was a gift from Bob Weinberg (Addgene plasmid # 8454), pCMV delta R8.2 was a gift from Didier Trono (Addgene plasmid # 12263), and MCS-BioID2-HA pBabe-puro was a gift from Kyle Roux (Addgene plasmid #120308). The lentiviral PTPRH WT-HA plasmid was prebuilt by Addgene. Briefly, the lentiCas9-Blast plasmid (Addgene plasmid # 52962), a gift from Feng Zhang, was modified to replace the Cas9 sequence with the human *PTPRH* coding sequence (CCDS 33110.1) and a C-terminal HA tag. A map of the generated plasmid is provided in the Supplemental Figure 11A.

### Site-Directed Mutagenesis

To generate PTPRH constructs with deletions in various sites, primers were designed using the NEBaseChanger software (https://nebasechanger.neb.com/) using the human *PTPRH* coding sequence (CCDS 33110.1) as a template. The sequence of the primers is described in the table below. Mutagenesis was carried out in the PTPRH WT-HA plasmid using the Q5 Site-Direct Mutagenesis Kit (New England Biolabs #E0554) according to the manufacturer’s instructions. Constructs containing *in-frame* mutations in the intended site were used for lentivirus generation and cell transduction.

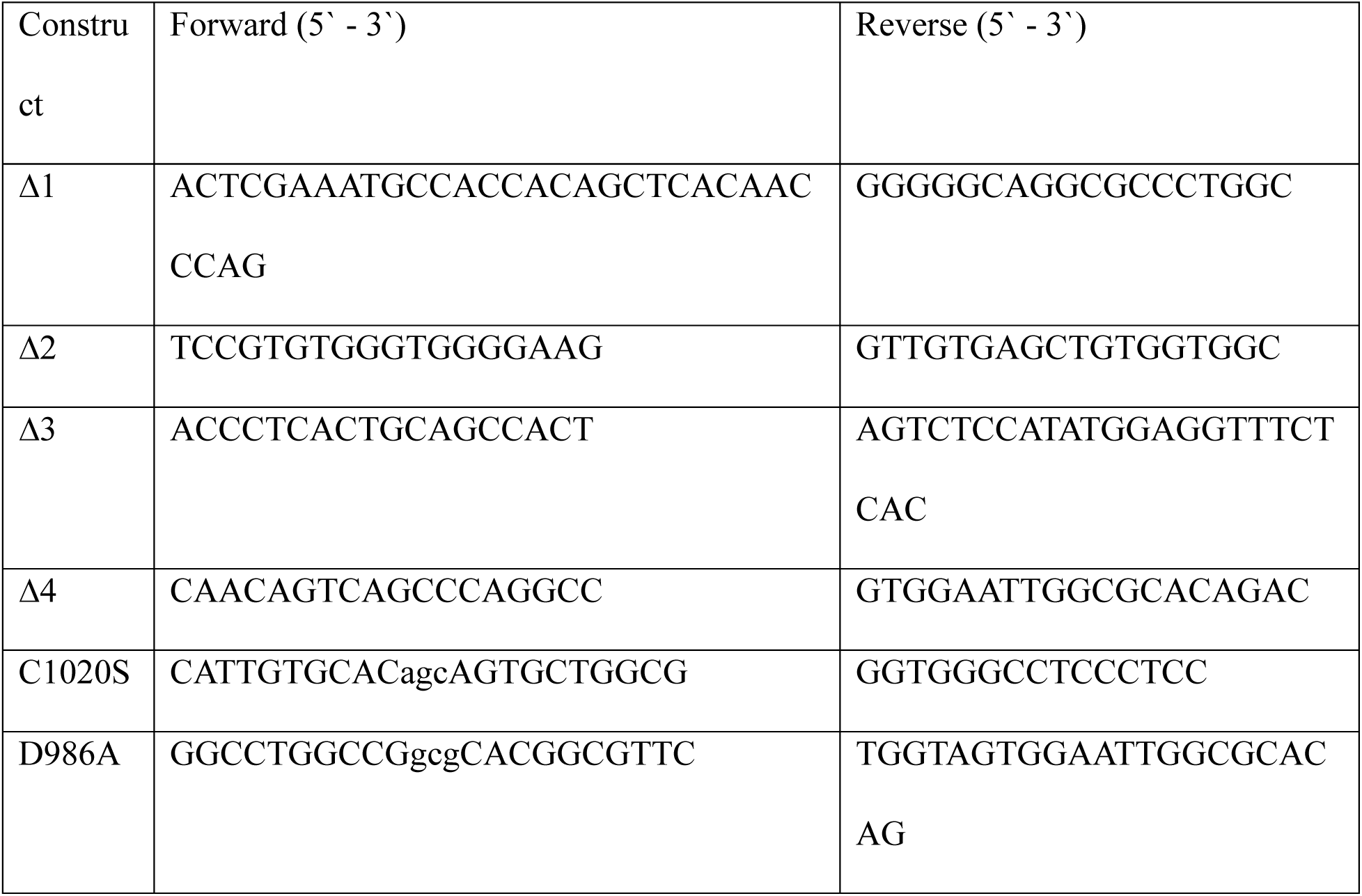

### Lentivirus Generation and Cell Transduction

293T cells were seeded on 10 cm dishes on the day before the transfection. The next day, pCMV-VSV-G, pCMV delta R8.2, and transfer plasmid were mixed with PEI Prime (Sigma-Aldrich #919012) in Opti-MEM medium (Gibco #11058021) in a 1 DNA: 3 PEI ratio and added to the cells. Mock viruses were generated using pCMV-VSV-G and pCMV delta R8.2 only. After 6 hours of incubation, the medium was discarded, and cells were replenished with a complete DMEM high-glucose medium. The supernatant containing the lentiviruses was harvested at 48 and 72 hours after transfection and filtered through a 0.45 µm Millex PES filter (Millipore Sigma #SLHPR33RS). The lentiviruses were concentrated 10x using Lenti-X Concentrator (Takara Bio #631232) following the manufacturer’s instructions and stored at -80°C.

For the transduction step, target cells were plated in 6-well plates using 10^5^ cells/well on day 1. The next day, when cells were ∼40% confluent, a mixture of complete media, Viral Entry Transduction Enhancer (abm #G515), and an equal volume of lentiviruses was added. On day 5, we started the selection of infected cells with blasticidin (Dot Scientific Inc. #3513-03-9) at 4.0 or 8.0 μg/mL, as dictated by the killing curve. The selection was stopped when all parental cells were dead.

### BioID2 experiment

BioID2-HA was cloned into the C-terminal of *PTPRH*-WT, *PTPRH*-DACS, or inserted into an empty lentiviral backbone using the Golden Gate method. Assembly reaction was performed at a 1:2 vector: insert ratio using the NEBuilder HiFi DNA Assembly Cloning Kit (New England Biolabs #E5520) according to the manufacturer’s protocol.

NCI-H2023 and H23 KO cells were transduced with BioID2-HA-only, *PTPRH* WT-HA-BioID2, *PTPRH* DACS-HA-BioID2, or mock lentiviruses. Cells expressing the BioID2 constructs were selected by treatment with blasticidin. When ∼ 60% confluency was reached, cells were serum-starved for 24 hours. After the incubation period, cells were replenished with a complete medium containing FBS and 50 μM biotin (Thermo Scientific #29129) and lysed after 18 hours. Biotinylated proteins were immunoiprecipitated using MagReSyn Streptavidin MS beads (ReSyn Bio #MR-STV002) equilibrated in Pierce RIPA buffer + Halt Protease and Phosphatase inhibitor (Thermo Scientific #78441) for 24 hours at 4°C using end-to-end rotation. The washing of the antigen-bead complex was performed following the manufacturer’s instructions. Next, the immunoprecipitated fraction was run in LC/MS/MS for protein identification. The analysis of PTPRH interactors adhered to the recommendations outlined by Roux *et al* ^73,74^. Briefly, the identified proteins underwent initial filtration to exclude those commonly observed in biotinylation experiments, referencing the CRAPome database. Those with a CRAPome score > 300 were considered contaminants and removed from the analysis. Candidate interactors were defined as having a positive fold change in normalized total precursor intensity in the PTPRH -BioID2 group when compared to BioID2-only and p < 0.1 after Benjamini-Hochberg multi-hypothesis correction. STRING analysis of these candidate interactors was done using Cytoscape 3.10.2.

### Mass Spectrometry

LC/MS/MS was performed at the MSU Mass Spectrometry Core. On-bead antibody-bound proteins were first washed 3 times with 50 mM ammonium bicarbonate and digested with trypsin (5 ng/μL, incubation at 37°C for 6 hours). The solution was acidified using 1% trifluoroacetic acid and centrifuged at 14000 x g. After supernatant collection, the peptides were concentrated by solid phase extraction using StageTips^75^ and dried by vacuum centrifugation. An injection of 10 μL of dried peptides resuspended in 2% acetonitrile, and 0.1% trifluoroacetic acid was conducted using the Thermo Scientific EASY-nLC 1200 system and 0.1mm x 20 mm Acclaim™ PepMap™ C18 RSLC trapping columns (Thermo Scientific), followed by a 5-minute wash with buffer A (99.9% Water/0.1% Formic Acid). Peptides were eluted for 35 minutes onto a 0.075 mm x 500 mm Acclaim™ PepMap™ RSLC resolving column (Thermo Scientific) using a linear gradient ranging from 5-28% of buffer B (80% Acetonitrile/0.1% Formic Acid/19.9% Water) for 24 minutes. Next, column wash was performed using 90% buffer B for 10 minutes (constant flow rate = 300nl/min). Column temperature was kept constant at 50°C with the aid of an integrated column oven-PRSO-V2 (Sonation GmbH). Using a FlexSpray ion source, the peptides were then sprayed into the Q-Exactive mass spectrometer (Thermo Scientific), and survey scans of 45000 resolution (based on m/z 200) were collected using the Orbi trap. The predominant 15 ions from each survey underwent an automatic higher-energy collision-induced dissociation (HCD), and the resulting fragment spectra were collected using a resolution of 7500. These spectra were then transformed into a peak list by the software Mascot Distiller v2.8.3. Using the Mascot^2^ searching algorithm v 2.8.0.1, the peaks list was compared to a human protein database containing all sequences from UniProt and supplemented with frequent laboratory contaminants (downloaded from www.thegpm.org, cRAP project). Parameters were set to allow up to 2 missed tryptic sites, fixed modification of carbamidomethyl cysteine, variable modification of oxidation of Methionine and/or phosphorylation of Tyrosine, MS/MS tolerance of 0.02 Da, and peptide tolerance +/-10 ppm. Following, the output list was analyzed using Scaffold v5.3.0. Proteins identified with 99% protein threshold, 95% peptide threshold, and 2 minimum peptides were considered true. Quantitative analysis was performed based on normalized total precursor intensity, and missing values were replaced by imputation (1,000 or 10,000).

### Immunofluorescence and Confocal Imaging

Cells were seeded in 6-well plates containing glass coverslips or 8-well glass culture slides (Falcon #354108). Thin-coating with collagen I (5 µg/cm^2^, Gibco #A10483-01) and Ephrin A1-Fc (R&D systems #6417-A1-050) stimulation was used when indicated. Cells were allowed to grow until the desired confluency and fixed in 4% paraformaldehyde (PFA, pH 7.4) for 10 minutes at RT and rinsed with ice-cold PBS three times. Next, cells were permeabilized using PBS-Tween 20 0.4% for 15 min, washed three times with PBS for 5 min/wash, and blocked with 1% BSA/22.52 mg/mL glycine in PBS 0.1% Tween 20 for 30 minutes. The slides were then incubated with primary antibodies against HA tag (1:100, Cell Signaling #2367) or EPHA2 (1:200, Cell Signaling #6997) diluted in 1% BSA/ PBS-Tween 20 0.1% for one to two hours in a humidified chamber at RT. Following three washes with PBS for 5 minutes/each, Alexa Fluor 680 donkey anti-mouse (1:400, Invitrogen #A-10038), Alexa Fluor 594 goat anti-rabbit (1:400, Invitrogen #A-11012), and Alexa Fluor 488 Phalloidin (1:400, Invitrogen #A-12379) diluted in 1% BSA/ PBS-Tween 20 0.1% was added to the coverslips and incubated for 1 hour at RT. Slides were mounted to glass slides with Prolong Gold Antifade Mountant with DAPI (Invitrogen #P36935).

Images were acquired in the inverted Leica Stellaris confocal microscope using the HC PL APO CS2 63x/1.40 or HC PL APO CS2 100x/1.40 oil objectives, an 8-bit or 16-bit detector, and the laser lines 499, 590, and 681 nm with constant intensity in the 2-10 range, depending on the channel. All image parameters (e.g., laser intensity and gain) were kept the same, and a minimum of 5 random fields per replicate were imaged. Images were processed with ImageJ 1.54p.

### Cell morphology analysis

Confocal images were obtained as described above, and individual cells from multiple random fields of each replicate were treated as objects for the analysis. To quantitatively measure the morphological features on the grey scale images, the Cell Profiler 4.2.8 software was used. First, the nuclei of the cells were segmented using the DAPI channel through the ‘IdentifyPrimaryObjects’ pipeline. Next, the cell contour was segmented based on the F-actin staining using the Propagation method and Minimum Cross-Entropy thresholding method from ‘IdentifySecondaryObjects’. If the automated segmentation was not accurate, the option ‘IdentifyObjectsManually’ was used as an alternative. Following, the cytosplasm was segmented through the ‘IdentifyTertiaryObjects’ and the area and shape features of the segmented objects were outputted in the ‘MeasureObjectSizeShapè pipeline using default parameters. Quantitative values for each feature and object were given in pixels and used as input in R Studio for statistical analysis and plotting.

### H&E Staining

Tissue samples fixed in 10% formalin and embedded in paraffin were sectioned at 4-5 microns. Sections were dried in a 56°C slide incubator to ensure adherence to the slides for 2 – 24 hours and stained on the Leica Autostainer XL using the Hematoxylin and Eosin staining method. For this, slides were covered with Xylene for 5 minutes twice, followed by two incubations with absolute ethanol for 5 minutes each, two incubations with 95% ethanol for 2 minutes each, and rinsed with tap water. Next, slides were stained with CATHE Hematoxilyn 1:10 (Biocare) for 1.5 minutes, followed by a 10 – 15 second differentiation in 1% aqueous glacial acetic acid and tap water for 2 minutes. The slides were then covered in 95% ethanol for 2 minutes, stained with 1% Alcoholic Eosin-Phloxine B for 2 minutes, followed by a change of 95% ethanol for 2 minutes, 100% ethanol four times for 2 minutes each, and four changes of Xylene for 2 minutes each before coverslipping with mounting media. Slides were scanned using the Olympus SLIDEVIEW VS200 at 20x magnification and bright field.

### qPCR

NCI-H23, A427, and HT-29 cells were seeded as triplicates in 6-well plates (2 x 10^5^ cells/well) in complete medium. On the following morning, EGF was added to each well (500 ng/mL) and cells were incubated for 18 hours. Total RNA was extracted using the RNeasy Mini Kit (Qiagen #74104) according to the manufacturer’s instructions. cDNA generation and qPCR were performed in one-step using 100 ng of template, 100 nM of primers (PTPRH, F: 5’ TGTGCGTCCACATACCCAGA 3’, R: 5’ CAGCCTCCCCACATGAAGAT 3’; GAPDH, F: 5’ TCATGACCACAGTGGATGCC 3’, R: 5’ GGAGTTGCTGTTGAAGTCGC 3’), and Power SYBR™ Green RNA-to-CT™ 1-Step Kit (Applied Biosystems #4391178) following the manufacturer’s protocol. ΔΔ Ct analysis was performed as described by Livak & Schmittgen.^76^

### RNA-sequencing

NCI-H2023 and H23 KO-derived cells were seeded in triplicate in 6-well plates (10^5^ cells/well), serum starved for 24 hours, and replenished with complete medium for 16 hours. Total RNA was collected using RNeasy Mini Kit (Qiagen #74104) according to the manufacturer’s protocol. Library preparation and paired-end RNA sequencing (Illumina PE150) were conducted by Novogene using NovaSeq X Plus Series (Illumina) using a minimum coverage of 30M reads per sample. The quality of the raw reads was assessed through *FastQC* 0.11.7 and trimmed using *Trimmomatic* 0.39. Trimmed reads were aligned against the human genome (grch38) using *Hisat2* 2.1.0 and a count matrix of raw reads to be used as input for DESeq2 was generated by *StringTie* 2.1.3 as described in the *StringTie* documentation. Differential expression analysis was performed using DESeq2 1.42.1, and differentially expressed genes (DEG) with padj < 0.05 were considered significant. The *fgsea* package 1.32.4 was used for Gene Set Enrichment Analysis on DEG and MSigDB gene sets, including hallmark, C5 gene ontology, and C6 oncogenic signature. Pathways with padj < 0.05 were considered enriched and reported in our analysis.

For differential expression of transcripts and isoform abundance analysis, we followed the pipeline described by Pertea *et al*^47^. Briefly, reads mapped by *Hisat* were fed into *StringTie* for transcript assembly and abundance estimation using isoform discovery parameters. Following, isoform-level differential expression analysis and plotting were performed using *Ballgown* 2.38.0. Low-abundance genes were removed from the differential analysis, and only those with q<0.1 were considered significant.

### Transwell migration and invasion assay

Transwell migration and invasion assays were performed using 24-well companion plates containing Corning® Control inserts (#354578) or BioCoat™ Matrigel® Invasion Chamber inserts (#354480), respectively, with pore sizes equal to 8 μm. Before the assay, the invasion chambers were rehydrated with complete serum-free media for two hours. 25.000 cells in 500 μL of serum-free complete medium were seeded into each insert in quadruplicate, and 750 μL of FBS-containing medium was added as a chemoattractant in the well. Cells were incubated at 37°C, 5% CO2 atmosphere, for 24 hours. After the incubation period, non-migrating/non-invading cells on the upper surface of the membrane were removed with a swab. Migrating or invading cells on the lower surface of the membrane were fixed using 10% formalin for 30 minutes and stained with crystal violet for 15 minutes, washed, and air-dried. Cell counting of each replicate was conducted using the bright field of the Olympus BX41 microscope. Images of the entire lower surface of the membranes were acquired in the same microscope using 1.25x magnification, while representative fields were imaged at 10x. Images were processed with ImageJ 1.54p.

### Wound healing assay

3-4 x 10^5^ cells/well were seeded in complete medium in triplicate in a 6-well plate to reach 90-95% confluency on the day of the assay. Following, the cells were treated with Mitomycin C at 5 μg/ml and incubated for 2 hours at 37°C, 5% CO2 atmosphere to halt proliferation. Cells were washed twice with PBS, and a scratch was manually made in the middle of the well from top to bottom using a p200 tip. The scratch was imaged over time at 0h, 16h, and 24h (Experiment #1), or 0h, 25h, 48h (Experiment #2) time points using a 5x objective and bright field of the Evos Cell Imaging System or Zeiss Invertoskop 40C microscope. To ensure consistency between images, a black mark at the bottom of the well near the scratch was added and served as a guide to position the scratch in the microscope. The covered area within the scratch over time was calculated using the Wound Healing Size Tool plugin for ImageJ 1.54p as previously described by the authors^77^.

### Cell-ECM adhesion assay

Cell suspensions at 0.5×10^6^ cells/mL density in serum-free medium were added to the CytoSelect^TM^ 48-well cell adhesion assay plates (ECM array, colorimetric) (Cell Biolabs Inc #CBA-070) in triplicate (*n=*3/group) and incubated for 1 hour at 37°C, 5% CO2 atmosphere. BSA-coated wells were used as a negative control. Protocol was performed according to the manufacturer’s recommendations. OD was measured at 560nm wavelength using the plate reader SpectraMax iD3® (Molecular Devices).

### In vivo xenografts and bioluminescence imaging

Immunodeficient NOD-SCID female mice between 8-10 weeks old were purchased from Jackson Labs and Charles River Laboratories and acclimated for two to three weeks before experiments. Mice were injected subcutaneously with 8×10^5^ cells (H23 KO-derived cell lines) or 4 ×10^5^ cells (H2023-derived cell lines) diluted in sterile PBS into the right flank (n=5 mice/group). Tumor sizes were measured weekly using digital calipers along two perpendicular axes: length (the longest dimension) and width (the shorter dimension). When one or more mice reached the endpoint, defined as a tumor size equal to 2 cm in any measurement or when the tumors ulcerated, all animals were sacrificed. Animals were euthanized using CO_2_ inhalation followed by cervical dislocation as a secondary method, and necropsy was immediately performed. To avoid overestimations, tumor volumes were measured *ex vivo* by using the formula *Volume = 0.5 ×length × (width)*^2^. Following, tumors were formalin-fixed and embedded in paraffin for H&E and IHC histological analysis.

Before the tail vein injections, H2023-derived cells were stably labeled with the mCherry and Luciferase genes using lentiviruses (Addgene #200101). The transduced cells were cell-sorted for mCherry-positive cells using the Sony MA900. On the day of the experiment, 1x 10^5^ sorted cells diluted in sterile PBS were injected into the tail vein of NOD-SCID female mice (n=5 mice/group). For the bioluminescence imaging, the animals received sterile D-luciferin (GoldBio, #115144-35-9) intraperitoneally at 5 μl/g of body weight, and images were taken 15 minutes later, while the mice were under anesthesia by isoflurane inhalation. Image acquisition was performed in the IVIS Spectrum (Revvity®) imaging system using open filter and autoexposure settings. Mice were imaged at four time points after the tail vein injections: at 30 minutes (day 0) and after 8, 22, and 34 days (endpoint). The lungs were collected at the endpoint, inflated, formalin-fixed, and embedded in paraffin for H&E staining. Images were analyzed using the Living Image software (Revvity®). Regions of interest of the same size were drawn around the chest area, and the total flux [photons/second] within those areas was calculated for each animal at each time point using the calibrated units.

Mice were maintained in individually ventilated cages under pathogen-free conditions, and weight loss and behavior were monitored weekly. All animal studies were approved by the Institutional Animal Care and Use Committee (IACUC) under protocol number PROTO202400120.

### Data Visualization and Statistical Analysis

Data visualization and statistical tests were performed in R version 4.3.3. * p < 0.05 was considered a significant result.

### Data Availability

RNA-sequencing results are publicly available on the GEO repository under accession number GSE270167.

### Supporting Information

This article contains supporting information.

## Acknowledgments

We would like to thank Douglas Whitten from the MSU Proteomics Core, Dr. Sandra O’Reilly from the MSU IVIS Core, Dr. Melinda Frame from the MSU Center for Advanced Microscopy, and Amy Porter from the MSU Histology Lab for their guidance and expertise with the MS/MS data analysis, *in vivo* experimental design, and imaging. We also would like to acknowledge the members of the Andrechek lab – Carson Broeker, John Vusich, Anthony Schulte, Morgan Atkins, and Sarah Marei– as well as Dr. Amy Ralston, Dr. Sachi Horibata, Dr. Ana Vazquez, and Dr. Dohun Pyeon for providing insightful opinions about this research project.

## Authors’ contributions

MO, DP, and JGL performed the experiments. MS contributed to the experimental design, ACN provided the histological descriptions, and MO and ERA contributed to the experimental design and drafting of the manuscript.

This work was supported with NIH R01CA160514 to E.R.A

