## Supplemental Figures for "Unraveling the role of receptor-like protein tyrosine phosphatase PTPRH in cell signaling regulation and biological processes of non-small cell lung cancer"

- Supplemental Figure 1
- Supplemental Figure 2
- Supplemental Figure 3
- Supplemental Figure 4
- Supplemental Figure 5
- Supplemental Figure 6
- Supplemental Figure 7
- Supplemental Figure 8
- Supplemental Figure 9
- Supplemental Figure 10
- Supplemental Figure 11
- Supplemental Figure 12
- Supplemental Figure 13
- Supplemental Figure 14
- Supplemental Figure 15
- Supplemental Figure 16
- Supplemental Figure 17
- Supplemental Figure 18
- Supplemental Figure 19
- Supplemental Figure 20
- Supplemental Figure 21
- Supplemental Figure 22

## S-1

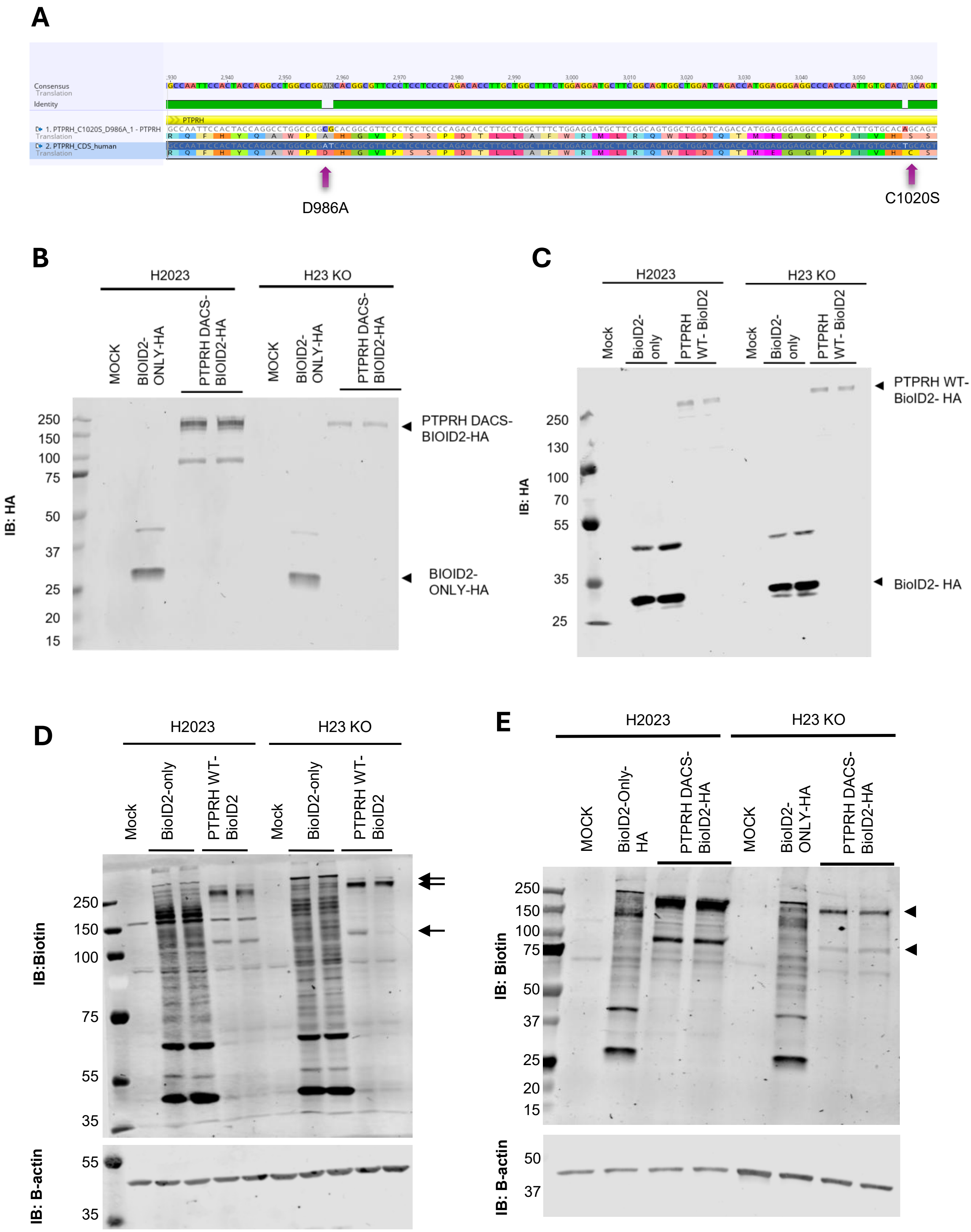
A
Consensus
Translation
Identity
PTPRH
1. PTPRH_C1020S_D986A_1 - PTPRH
Translation
2. PTPRH_CDS_human
Translation
D986A
C1020S
B
H2023
H23 KO
MOCK
BIOID2-ONLY-HA
PTPRH DACS-BIOID2-HA
MOCK
BIOID2-ONLY-HA
PTPRH DACS-BIOID2-HA
IB: HA
250
150
100
75
50
37
25
20
15
PTPRH DACS-BIOID2-HA
BIOID2-ONLY-HA
C
H2023
H23 KO
Mock
BioID2-only
PTPRH WT- BioID2
Mock
BioID2-only
PTPRH WT- BioID2
IB: HA
250
130
100
70
55
35
25
PTPRH WT-BioID2- HA
BioID2- HA
D
H2023
H23 KO
Mock
BioID2-only
PTPRH WT-BioID2
Mock
BioID2-only
PTPRH WT-BioID2
IB:Biotin
250
150
100
75
55
35
IB: B-actin
55
35
E
H2023
H23 KO
MOCK
BioID2-Only-HA
PTPRH DACS-BioID2-HA
MOCK
BioID2-ONLY-HA
PTPRH DACS-BioID2-HA
IB: Biotin
250
150
100
75
50
37
25
20
15
IB: B-actin
50
37

#### S-1 *(continued)*

**F**

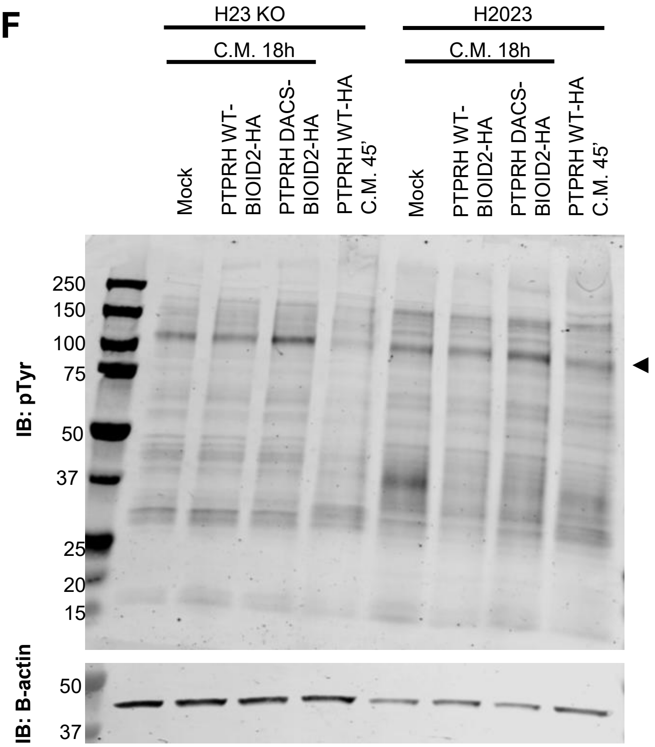


**Supplemental Figure 1. Validation of BioID2 transduced cells.** H23 and H2023 cells were transduced with mock, BioID2-only-HA, PTPRH WT-BioID2-HA, or PTPRH DACS-BioID2-HA and immunoblots against the HA tag, biotin, and phospho-tyrosine were performed to confirm expression of the constructs and functional validation. *A)* Sequence alignment of the substrate trapping (*PTPRH*-DACS) construct again the human *PTPRH* wild-type coding sequence*.* The consensus sequence and nucleotide position are shown at the top. *B)* Immunoblot showing expression of PTPRH DACS-BioID2-HA and BioID2-only-HA constructs. *C)* Immunoblot showing expression of PTPRH WT-BioID2-HA and BioID2-only-HA constructs. *D - F)* Biotinylation and tyrosine phosphorylation pattern in cells expressing BioD2-only-HA, PTPRH WT-BioID2-HA, and PTPRH DACS-BioID2-HA. The arrows indicate bands matching unique or presenting elevated levels in the PTPRH group. *Abbreviations:* IB = immunoblot, C.M. = complete media

## S-2 A

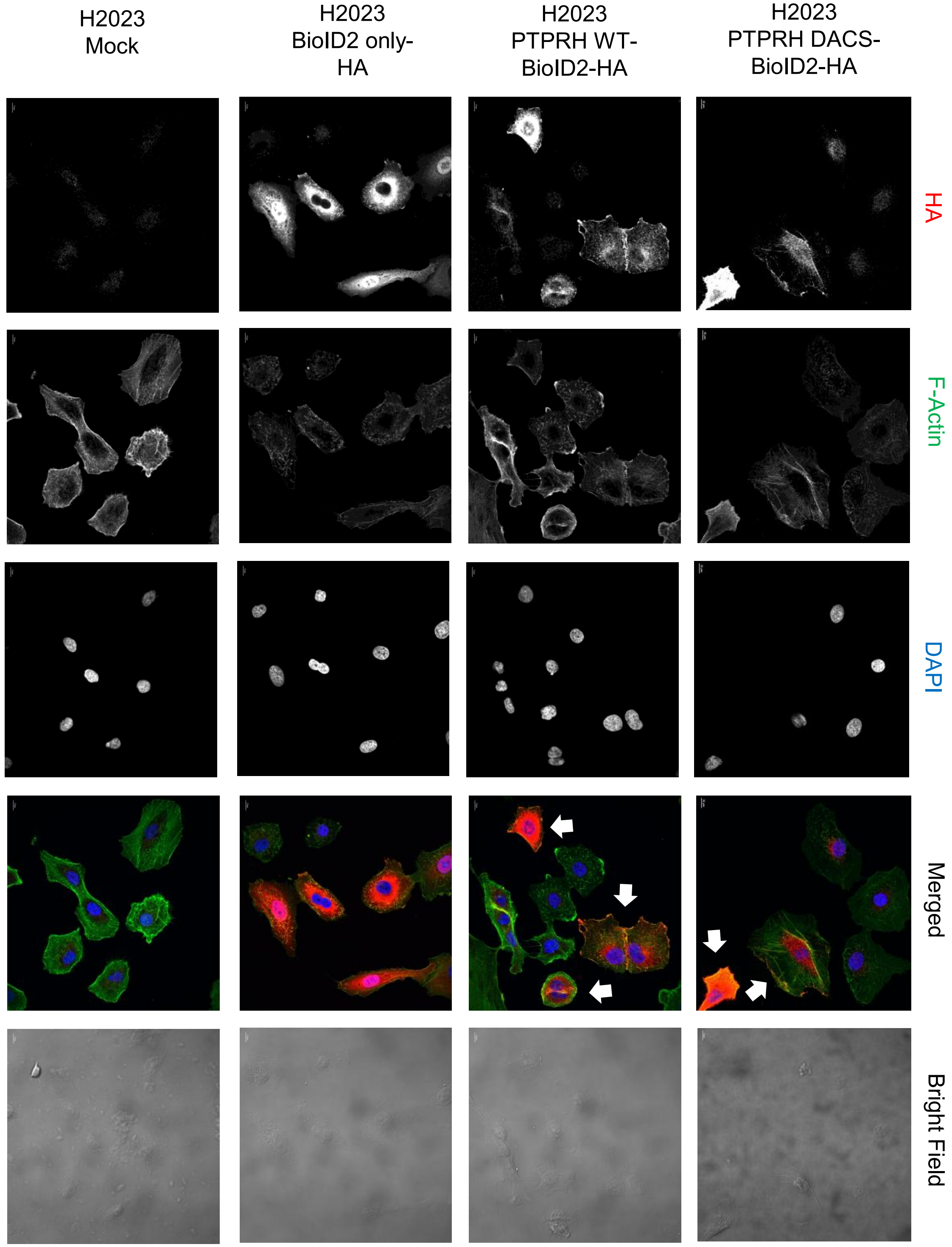


**Supplemental Figure 2. Subcellular localization of BioID2 constructs in lung adenocarcinoma cells.** H2023 (a) and H23 *PTPRH* knockout (b) cell lines transduced with BioID2-HA-only, PTPRH WT-BioID2-HA, PTPRH DACS-BioID2-HA or mock (control) were stained with antibody against the HA tag. Subcellular localization of protein is shown in red. DAPI staining is shown in blue, and F-actin in green. Representative cells showing expression of PTPRH WT-BioID2-HA are indicated by arrows. Images are representative fields using 60x magnification, scale bar =10 µm.

## S-2 B

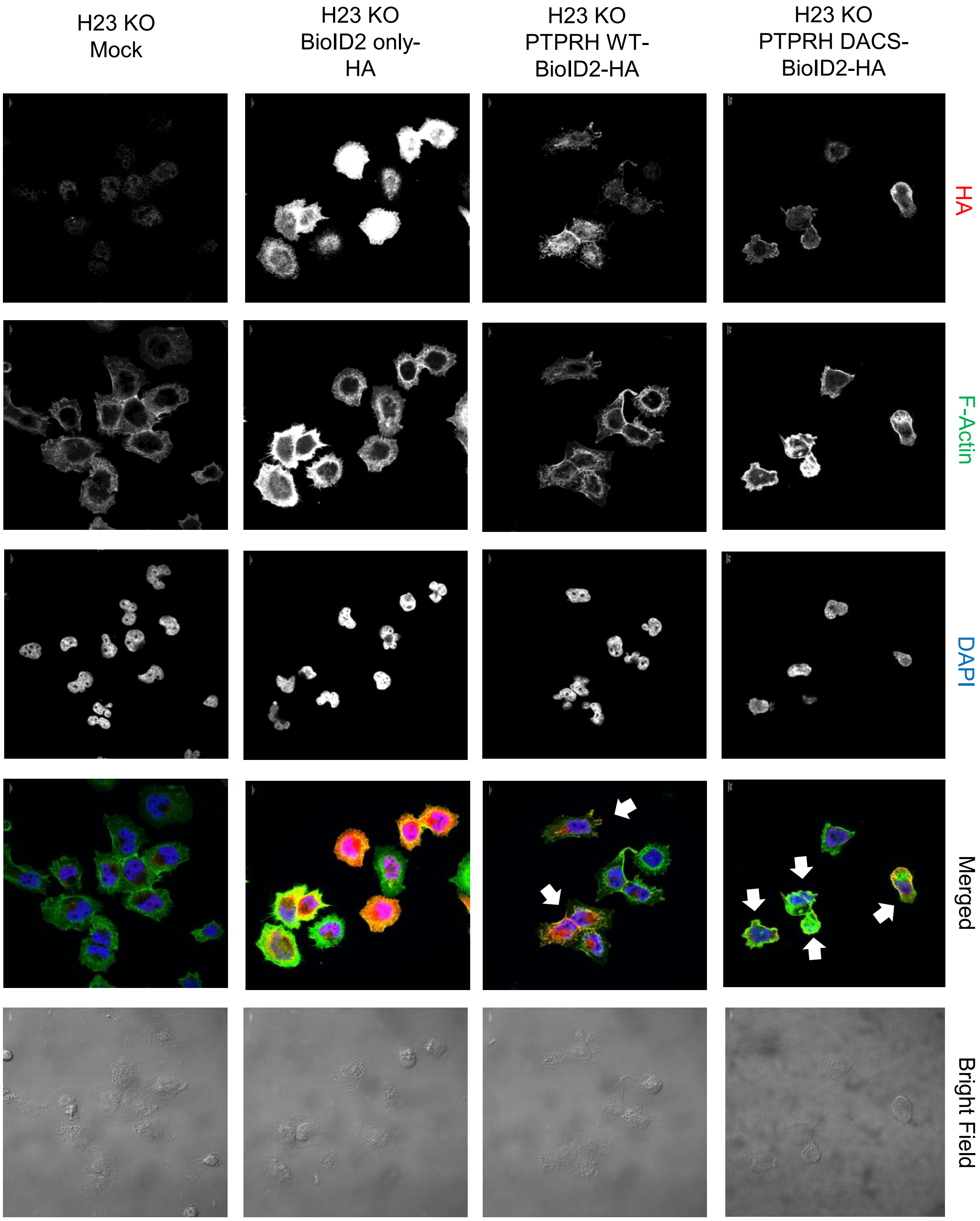

H23 KO
Mock
H23 KO
BioID2 only-
HA
H23 KO
PTPRH WT-
BioID2-HA
H23 KO
PTPRH DACS-
BioID2-HA
HA
F-Actin
DAPI
Merged
Bright Field

## S-3

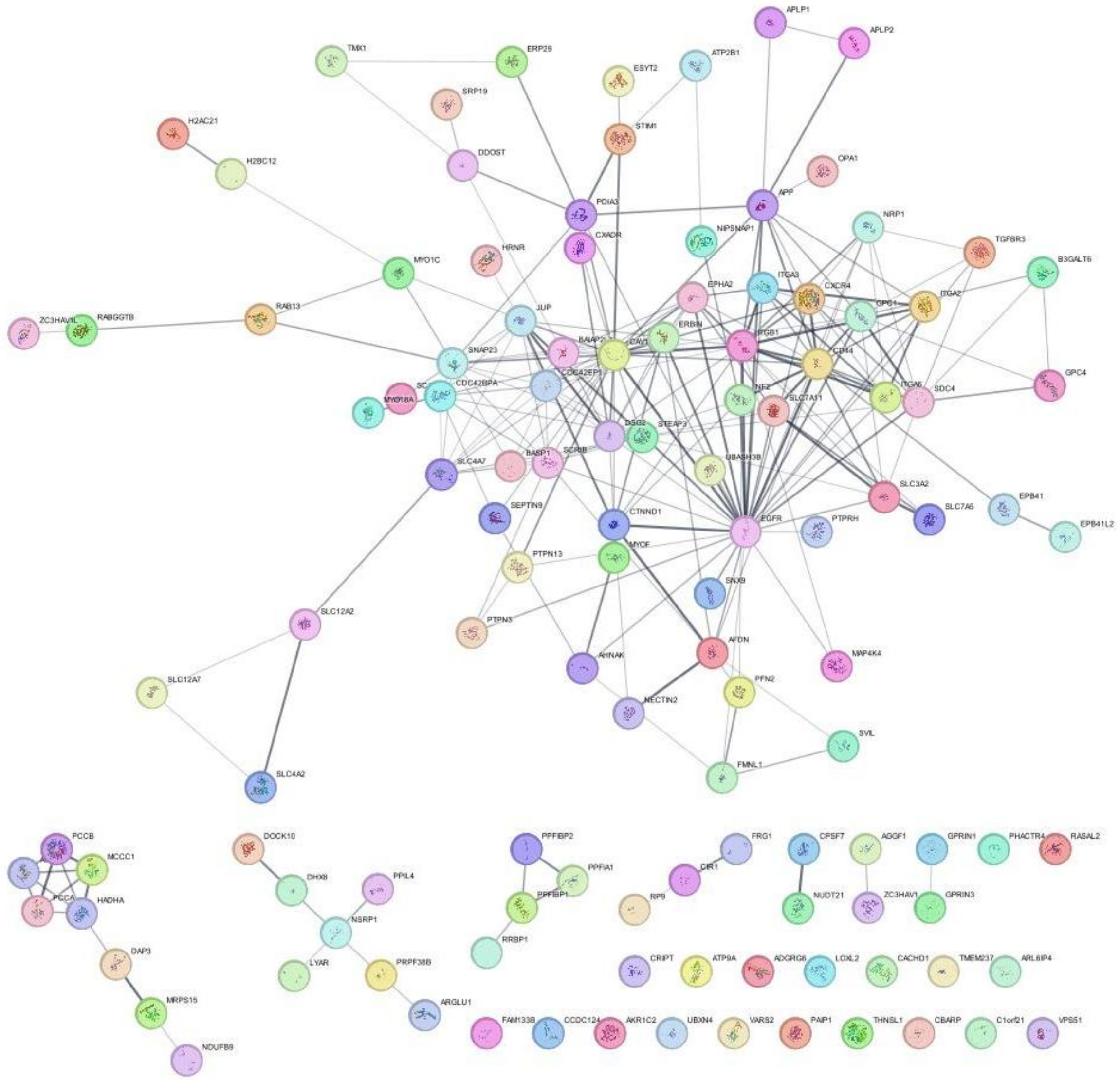


**Supplemental Figure 3. STRING analysis of candidate PTPRH interactors identified in H2023 and H23 PTPRH knockout cell lines transduced with PTPRH DACS-BioID2-HA.**

## S-4

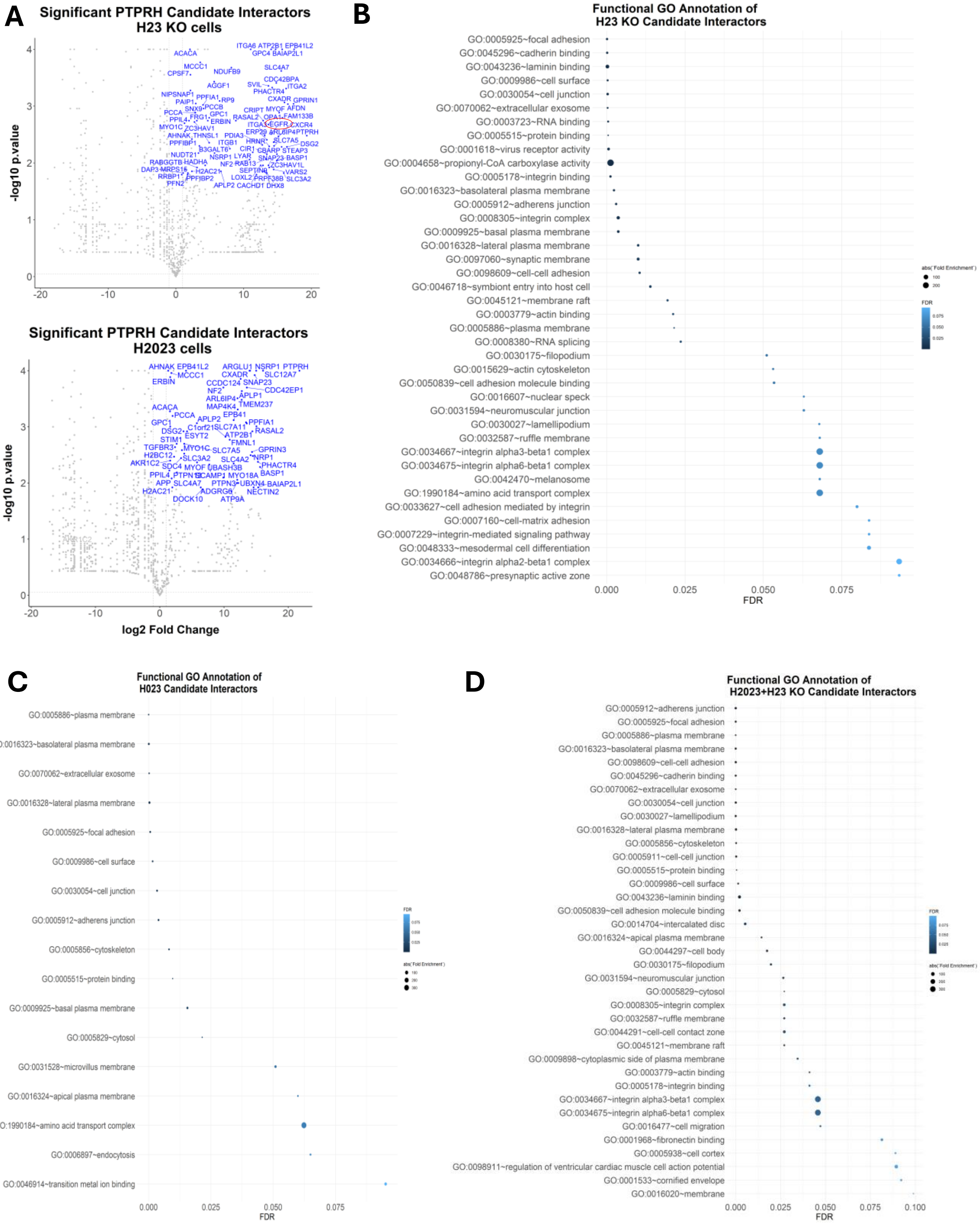


**Supplemental Figure 4. Functional annotation of PTPRH candidate interactors.** *A)* Significant candidate interactors identified in H23 KO and H2023 cells transfected with PTPRH DACS-BioID2 constructs. *B-D)* Using DAVID, the PTPRH candidate interactors identified in the H23 KO (n= 78 proteins) and H2023 (n=59 proteins) cells or the H23 KO + H2023 combinatorial analysis (n=59 proteins) were assessed for enrichment of gene ontology (GO) terms. The terms with FDR<0.1 and their fold enrichment absolute values are shown

## S-5

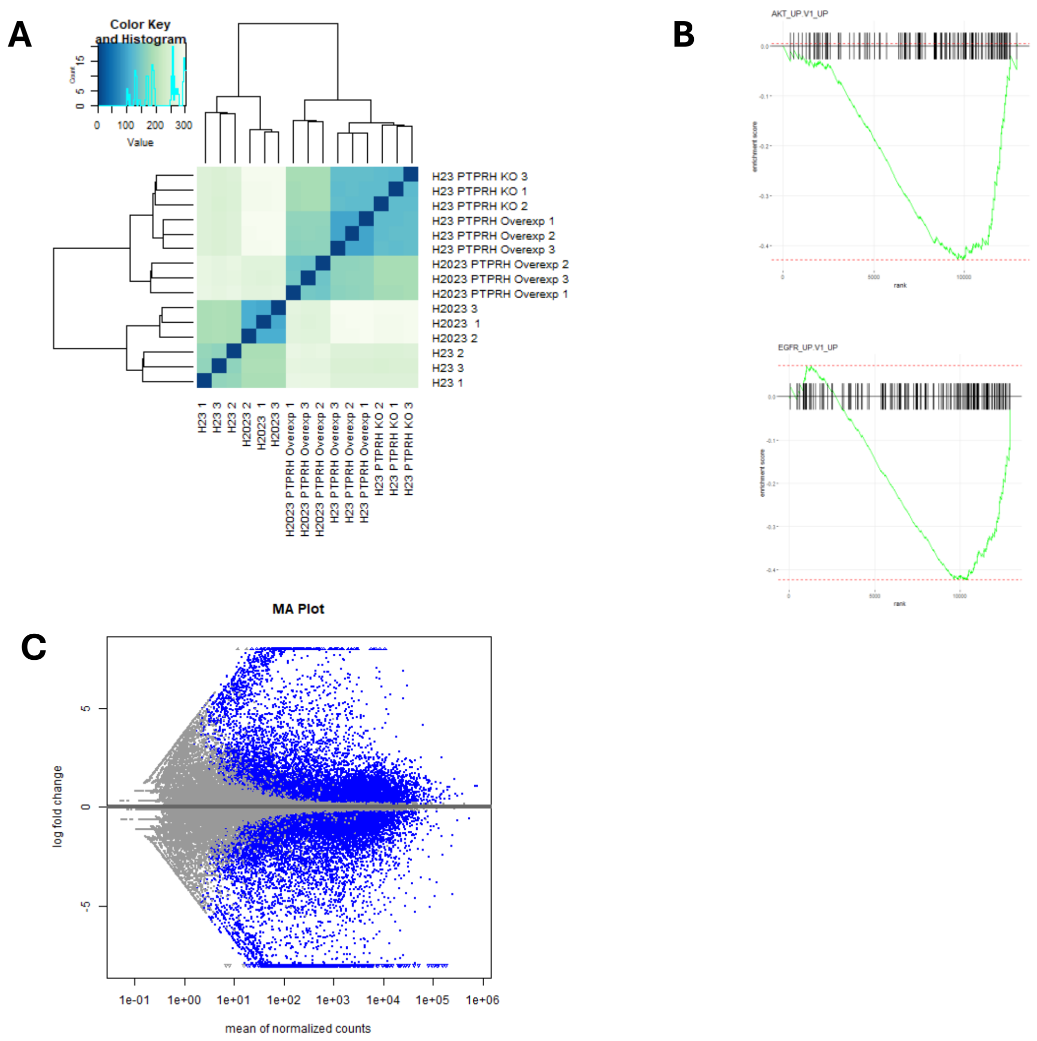


**Supplemental Figure 5. Transcriptome analysis of lung adenocarcinoma cell lines overexpressing PTPRH WT.** *A)* Hierarchical clustering analysis of technical replicates based on their normalized gene counts (rlog transformation). *B)* Normalized enrichment score (NES) plot of AKT_UP.V1_UP and EGFR_UP.V1_UP gene sets in PTPRH overexpression cell lines. *C)* MA plot of log2 fold change and mean of normalized counts.

<u>**S-6**</u>

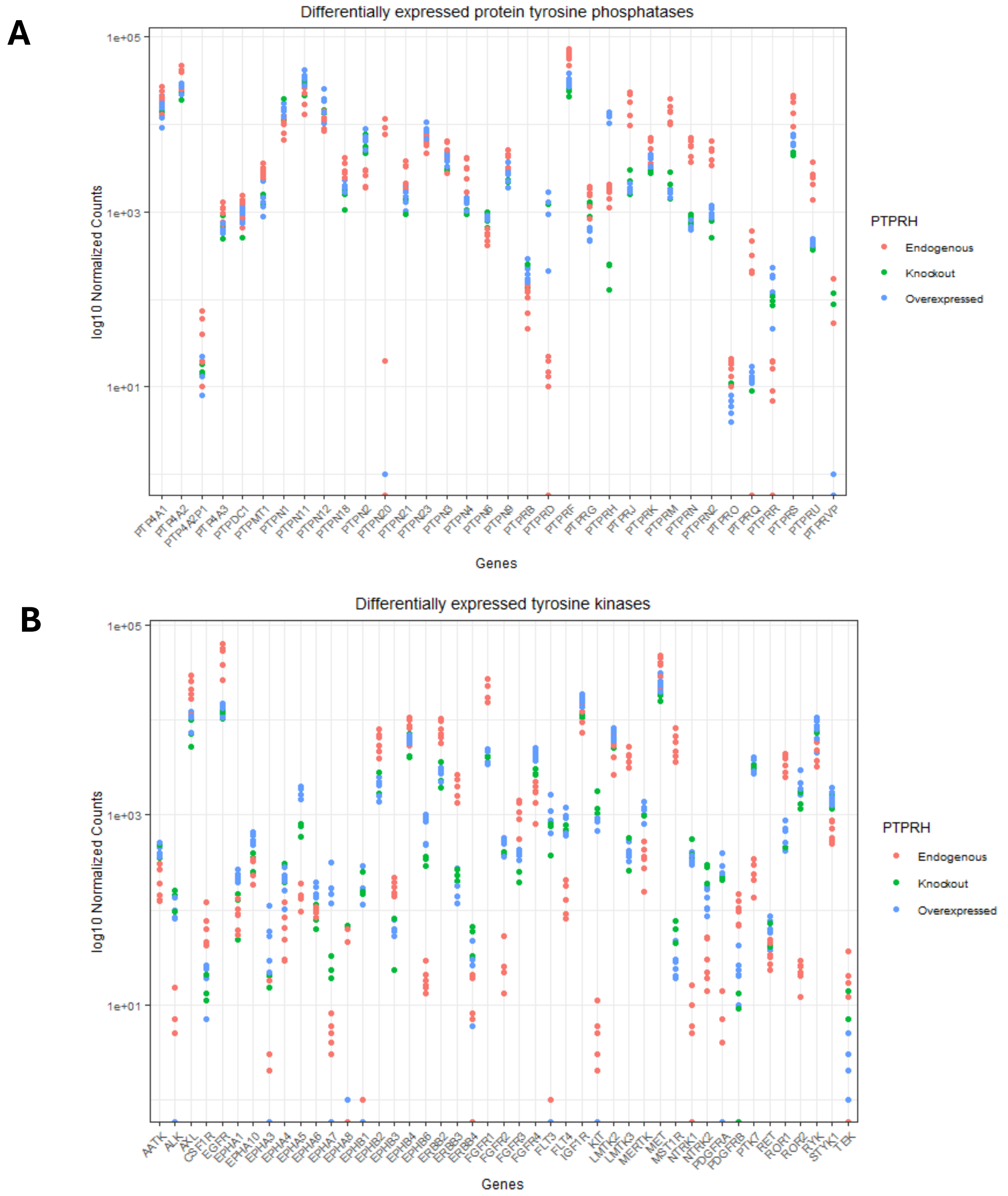


**Supplemental Figure 6. DEG analysis of tyrosine phosphatases and tyrosine kinases in lung adenocarcinoma cell lines overexpressing PTPRH WT.** Expression of differentially expressed protein tyrosine phosphatases *(A)* and tyrosine kinases *(B)* are represented as log10 normalized counts (padj <0.05).

# S-7 A

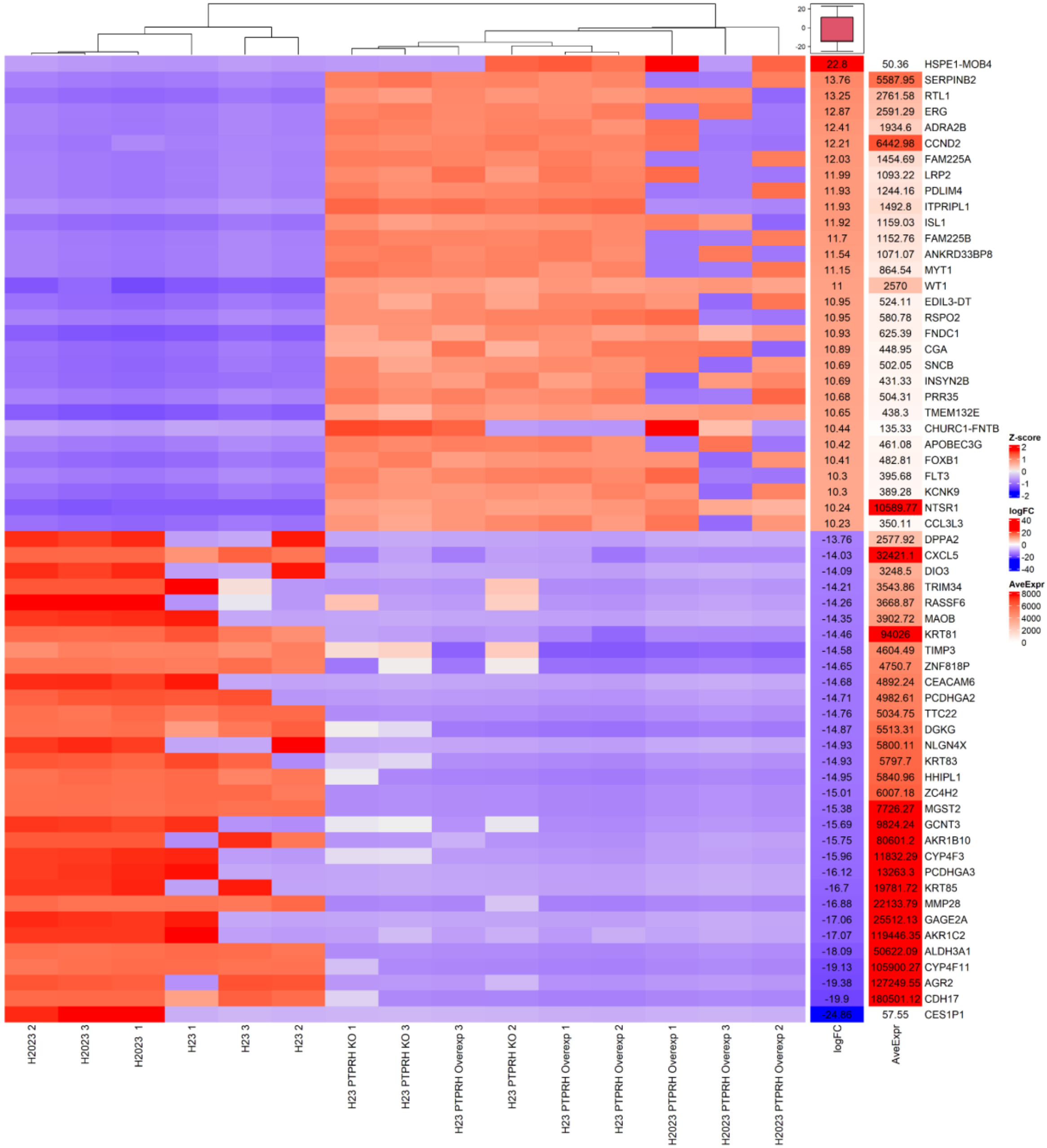
20
0
-20
22.8 50.36 HSPE1-MOB4
13.76 5587.95 SERPINB2
13.25 2761.58 RTL1
12.87 2591.29 ERG
12.41 1934.6 ADRA2B
12.21 6442.98 CCND2
12.03 1454.69 FAM225A
11.99 1093.22 LRP2
11.93 1244.16 PDLIM4
11.93 1492.8 ITPRIPL1
11.92 1159.03 ISL1
11.7 1152.76 FAM225B
11.54 1071.07 ANKRD33BP8
11.15 864.54 MYT1
11 2570 WT1
10.95 524.11 EDIL3-DT
10.95 580.78 RSPO2
10.93 625.39 FNDC1
10.89 448.95 CGA
10.69 502.05 SNCB
10.69 431.33 INSYN2B
10.68 504.31 PRR35
10.65 438.3 TMEM132E
10.44 135.33 CHURC1-FNTB
10.42 461.08 APOBEC3G
10.41 482.81 FOXB1
10.3 395.68 FLT3
10.3 389.28 KCNK9
10.24 10589.77 NTSR1
10.23 350.11 CCL3L3
-13.76 2577.92 DPPA2
-14.03 32421.1 CXCL5
-14.09 3248.5 DIO3
-14.21 3543.86 TRIM34
-14.26 3668.87 RASSF6
-14.35 3902.72 MAOB
-14.46 94026 KRT81
-14.58 4604.49 TIMP3
-14.65 4750.7 ZNF818P
-14.68 4892.24 CEACAM6
-14.71 4982.61 PCDHGA2
-14.76 5034.75 TTC22
-14.87 5513.31 DGKG
-14.93 5800.11 NLGN4X
-14.93 5797.7 KRT83
-14.95 5840.96 HHIPL1
-15.01 6007.18 ZC4H2
-15.38 7726.27 MGST2
-15.69 9824.24 GCNT3
-15.75 80601.2 AKR1B10
-15.96 11832.29 CYP4F3
-16.12 13263.3 PCDHGA3
-16.7 19781.72 KRT85
-16.88 22133.79 MMP28
-17.06 25512.13 GAGE2A
-17.07 119446.35 AKR1C2
-18.09 50622.09 ALDH3A1
-19.13 105900.27 CYP4F11
-19.38 127249.55 AGR2
-19.9 180501.12 CDH17
-24.88 57.55 CES1P1
H2023 2
H2023 3
H2023 1
H23 1
H23 3
H23 2
H23 PTPRH KO 1
H23 PTPRH KO 3
H23 PTPRH Overexp 3
H23 PTPRH KO 2
H23 PTPRH Overexp 1
H23 PTPRH Overexp 2
H2023 PTPRH Overexp 1
H2023 PTPRH Overexp 3
H2023 PTPRH Overexp 2
logFC
AveExpr
Z-score
2
1
0
-1
-2
logFC
40
20
0
-20
-40
AveExpr
8000
6000
4000
2000
0

#### S-7 *(continued)*

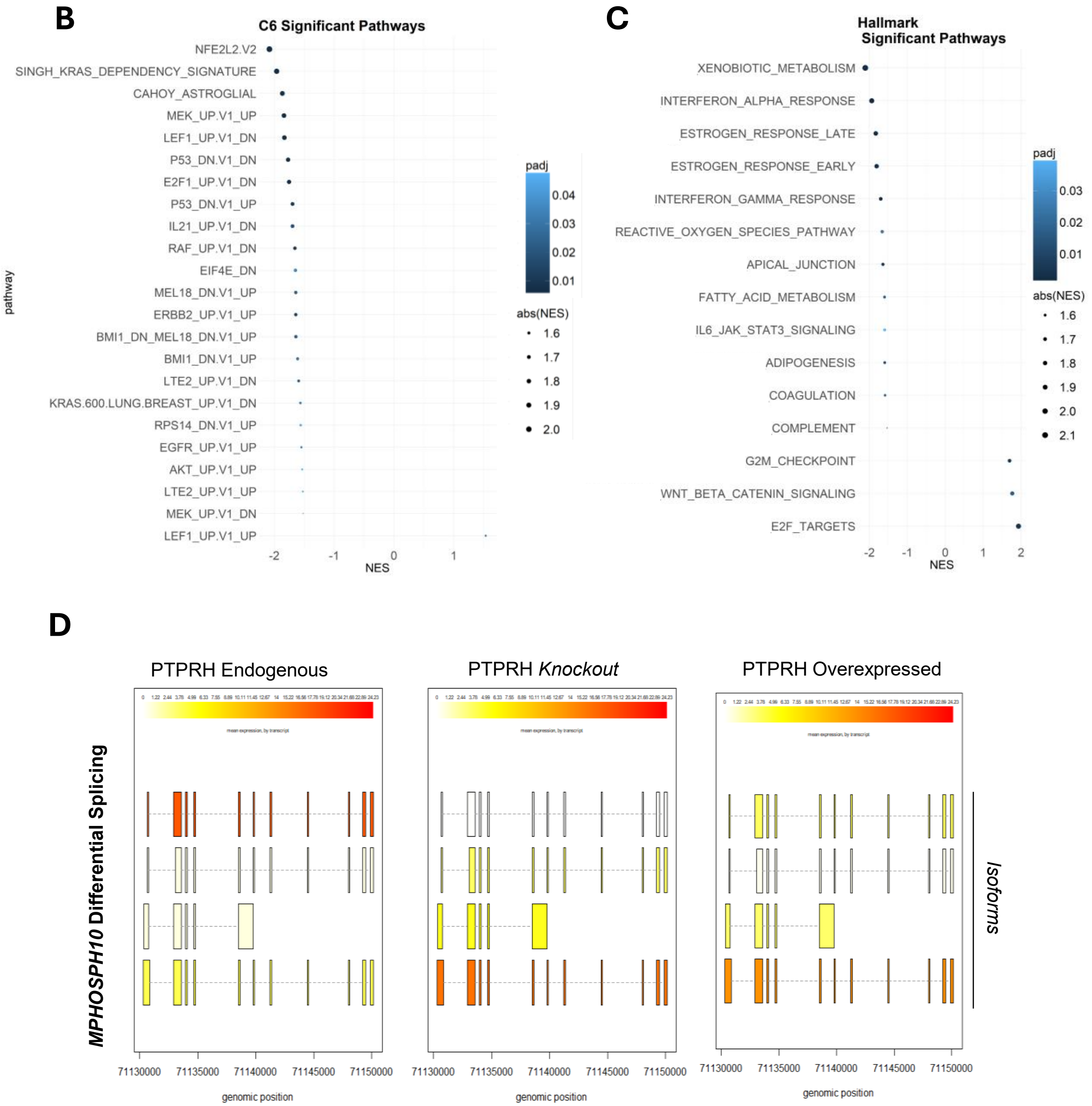


**Supplemental Figure 7. Transcriptional modulation by PTPRH overexpression.** *A)* Heatmap of top 30 most upregulated and downregulated differentially expressed genes (padj <0.05) in PTPRH overexpression cell lines. Log2 fold change (logFC), average expression (AveExpr), and Z-score for each gene are shown. Technical replicates of samples were clustered using hierarchical clustering analysis. *B-C)* Gene Set Enrichment Analysis using C6 oncogenic signature gene set (B) and the Hallmark (C) gene sets on differentially expressed genes. *D*) An example of how the *MPHOSPH10* transcripts vary in isoform abundance when PTPRH wild-type is endogenously expressed, knocked out, or overexpressed due to differential splicing.

# S-8

**A**

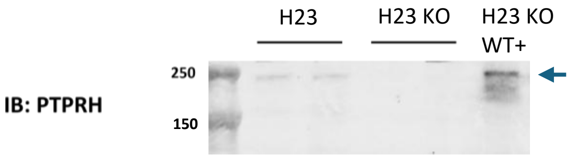


**B**

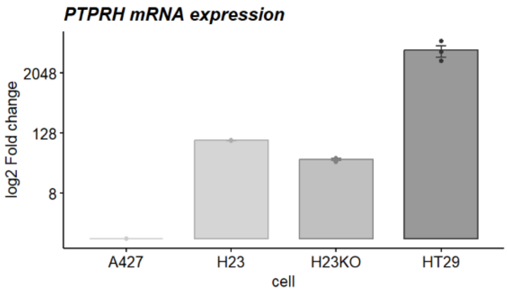


**Supplemental Figure 8. Endogenous PTPRH expression in lung adenocarcinoma cell lines.** *A)* Immunoblot against PTPRH in H23, H23 *PTPRH* KO, and H23 *PTPRH* KO cells in which PTPRH wild-type construct was added back. *B)* Log2 fold change of PTPRH mRNA expression in A427 (used as a reference sample), HT-29 cells (a colorectal carcinoma cell line overexpressing endogenous PTPRH, used as positive control), H23, and H23 *PTPRH* KO cells (n=3/cell line). As expected, mRNA levels were still detected even in the H23 KO cells, since our designed primers target a region of the phosphatase before the frameshift mutation that inserted to knockout PTPRH. Replicates are represented as individual dots in the bar plot, and the mean+/- s.d. is shown. *Abbreviations:* IB = immunoblot, WT = wild-type, H23 KO = H23 *PTPRH* KO

**S-9**

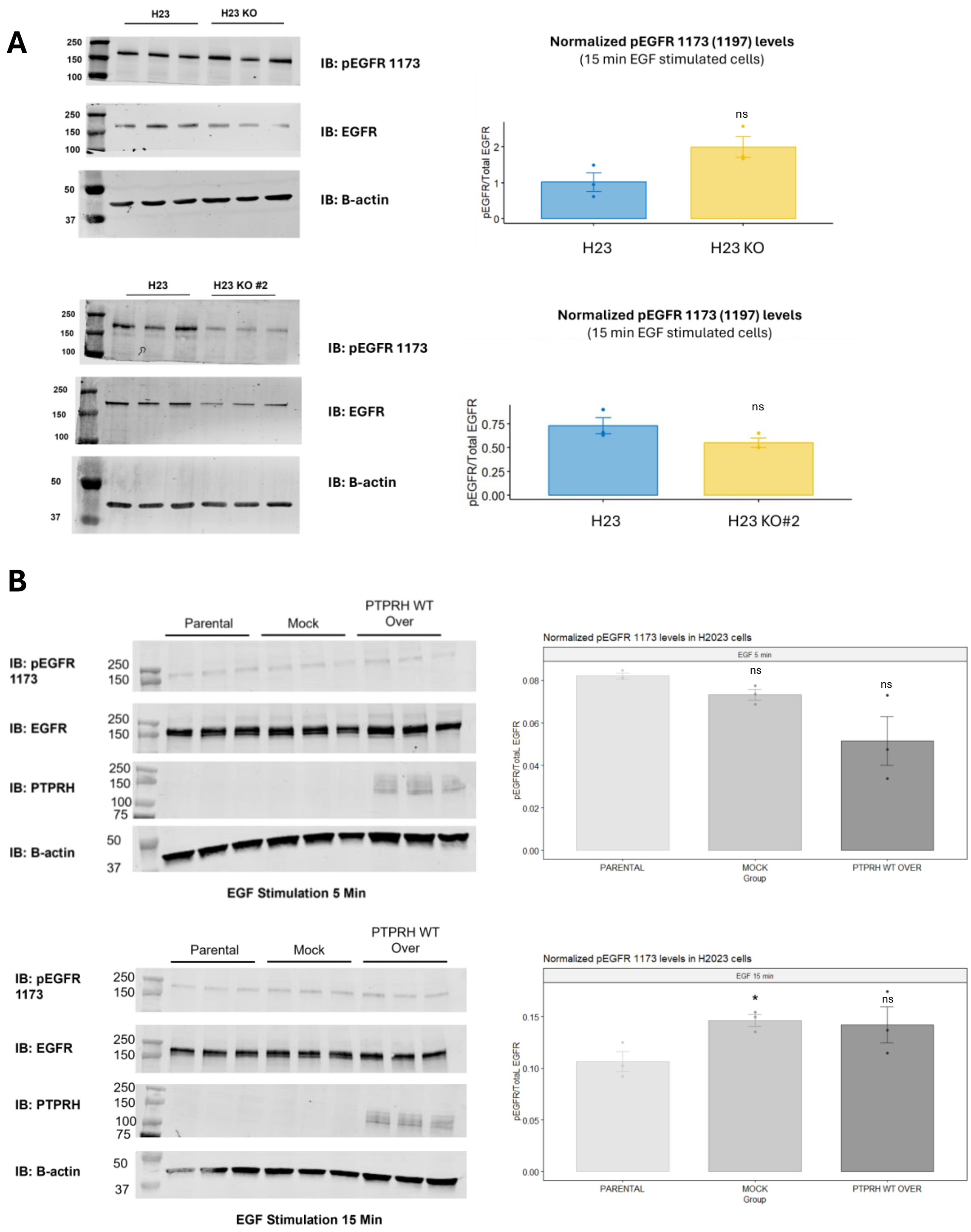


**Supplemental Figure. 9. pEGFR 1173 (1197) expression in lung adenocarcinoma cell lines**. Cells were screened for differences in pEGFR 1173 (1197) and total EGFR levels EGF (100 ng/mL) stimulation by immunoblot *A)* Comparison between H23 cells (parental) and two *H23 PTPRH* KO clones (KO and KO #2) (n=3/cell line) after 15 min of EGF stimulation. T-test was performed to compare the *PTPRH* KO group to the parental cell (H23) and p<0.05 was considered as a threshold for significancy. *B)* Comparison between H2023 (parental), H2023 Mock, and H2023 PTPRH overexpression cells (n=3/cell line) after 5 or 15 min of EGF stimulation. T-test was performed to compare the different groups to the parental cell line and p<0.05 was considered as a threshold for significancy. Results of densitometry for pEGFR 1173 (1197)/total EGFR ratio for the replicates are demonstrated as individual dots in the bar plots (mean +/- s.d.). *Abbreviations:* IB = immunoblot, ns = non-significant

## S-10

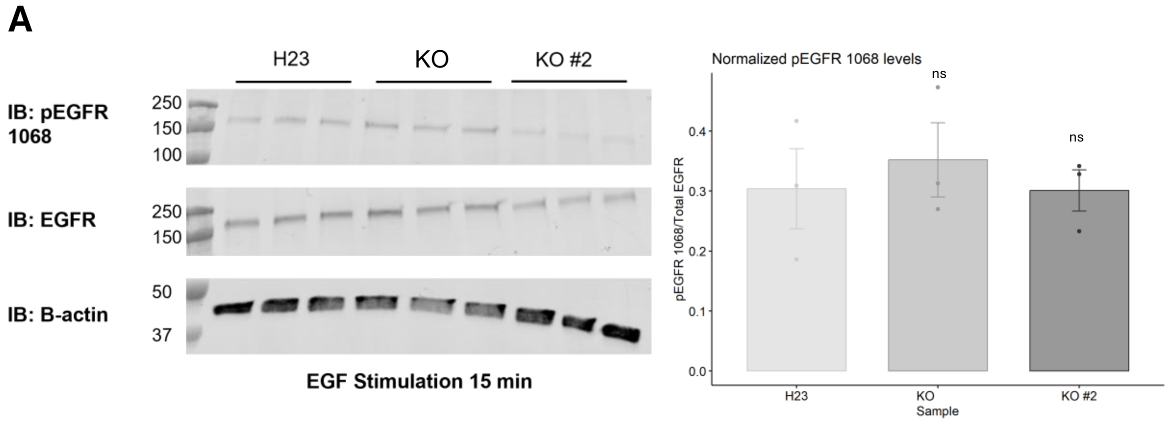


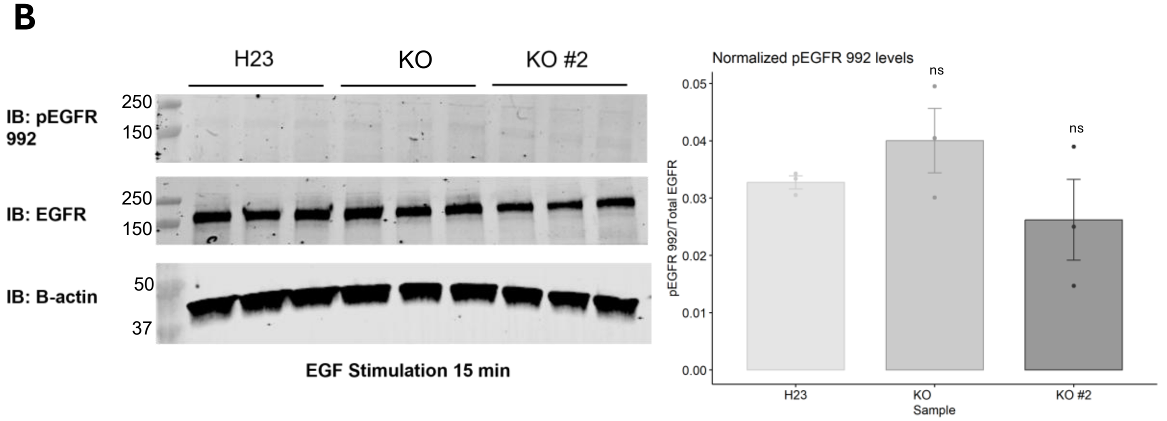


**Supplemental Figure. 10. pEGFR 1068 and pEGFR 992 expression in H23 *PTPRH* knockout cell lines**. Cells were screened for differences in pEGFR 1068 (*A*) and pEGFR 992 (*B*) after 15 minutes of EGF (100 ng/mL) stimulation by immunoblot. T-test was performed to compare the *PTPRH* KO clones (KO and KO #2) with the parental cell line (H23) and p<0.05 was considered a significant result. Results of densitometry for pEGFR/Total EGFR ratios of the replicates (n=3/group) are demonstrated as individual dots in the bar plots (mean +/- s.d.). *Abbreviations:* IB = immunoblot, ns = non-significant

# S-11

A

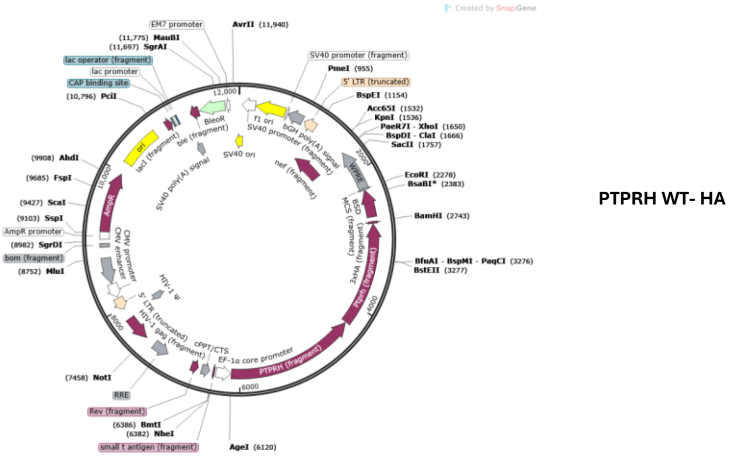


B

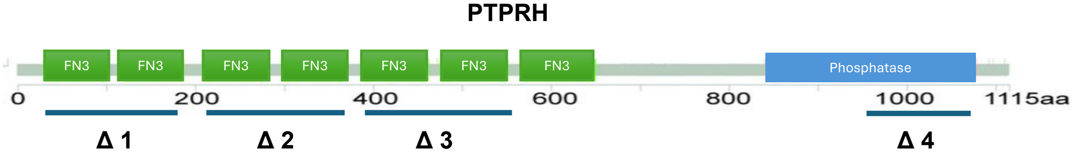


| Construct name | Description | Deletion Position |
|---|---|---|
| PTPRH-HA Δ1 | Deletion of fibronectin3 domains 1 & 2 | 30 – 200 aa (90-600 nt) |
| PTPRH-HA Δ2 | Deletion of fibronectin3 domains 3 & 4 | 209 – 364 aa (627-1092 nt) |
| PTPRH-HA Δ3 | Deletion of fibronectin3 domains 5 & 6 | 396 – 556 aa (1188-1668 nt) |
| PTPRH-HA Δ4 | Deletion of the end portion of the phosphatase domain | 980 – 1080 aa (2940-3240 nt) |

C

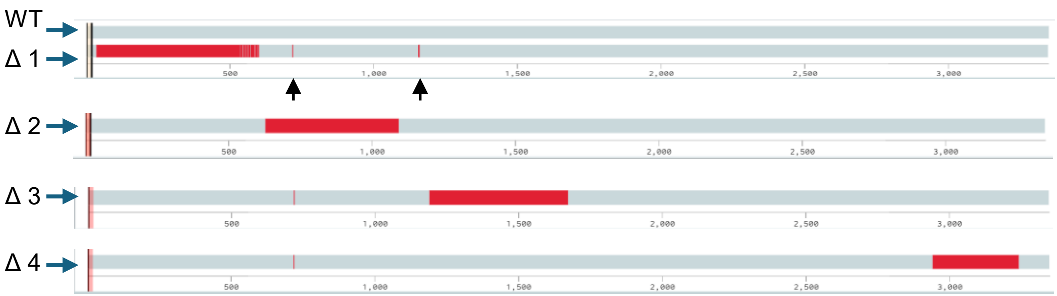


**Supplemental Figure 11. Sequencing of PTPRH constructs.** *A)* Annotated map of the PTPRH WT- HA plasmid containing the PTPRH human coding sequence. *B)* Description of the deletions inserted in the the fibronectin3 or the phosphatase domains of the *PTPRH* sequence contained in this plasmid. *C)* The generated constructs harboring deletions were aligned against the human PTPRH WT coding sequence. The red portions in the alignment represent the mismatched (deleted) regions. Numbers represent the nucleotide position. The arrows indicate positions where a single-nucleotide variation was found. A synonymous mutation is present across all constructs (c.717 G>C), besides a non-synonymous mutation in the Δ 1 construct (c.1157 C>A).

# S-12

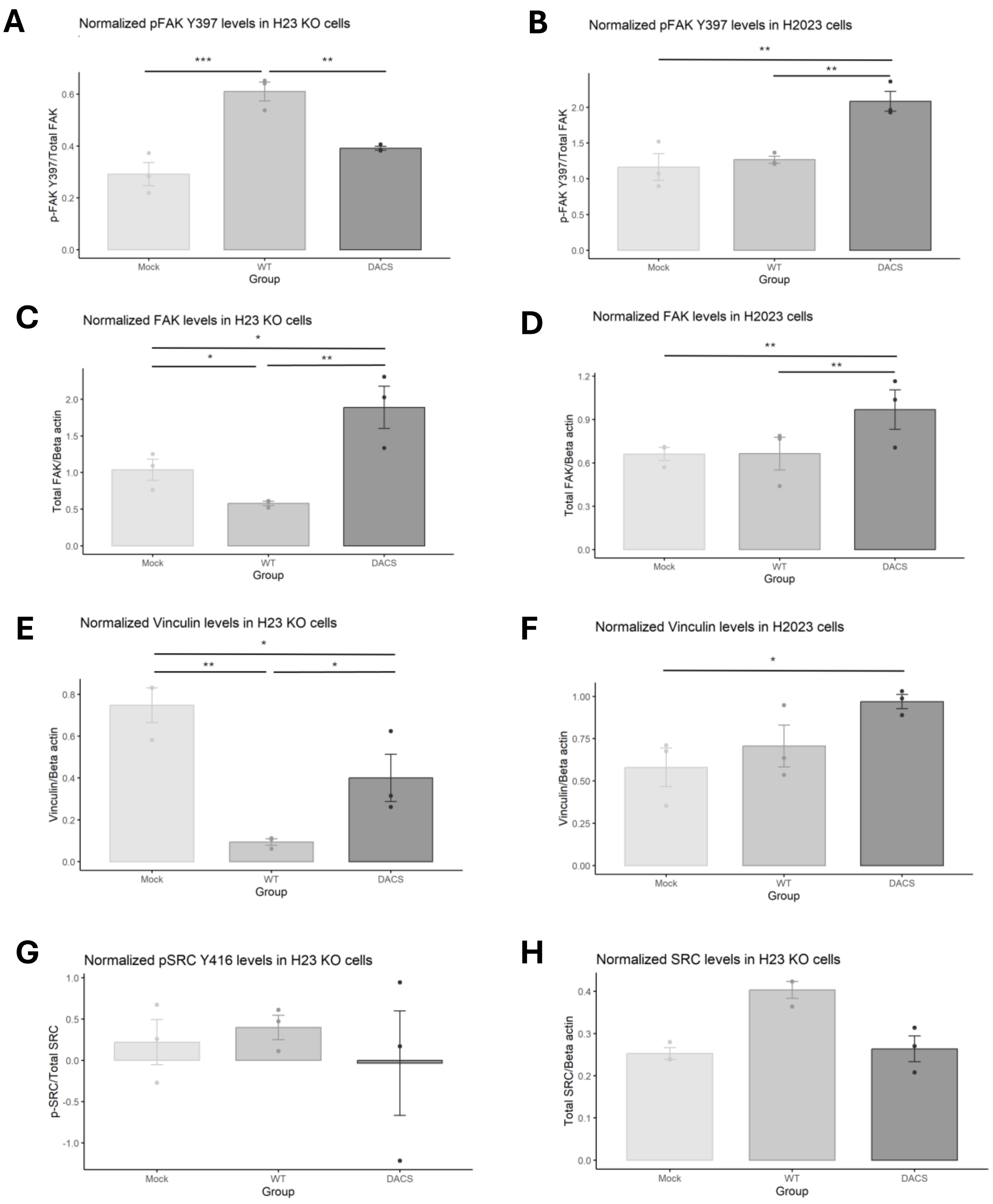


**Supplemental Figure 12. Densitometry of immunoblots for focal adhesion signaling molecules.** *A-H)* Immunoblots represented in *A* were quantified through the densitometry method. Protein expression was normalized by the beta-actin levels and the phosphorylated fraction calculated based on the normalized total levels of the protein. Statistical analysis was performed using One-Way ANOVA. Dots in the plots represent individual values for the replicates (n=3/group), and the mean +/- s.d is shown. *p<0.05, **p<0.01, and ***p<0.001

**S-13**

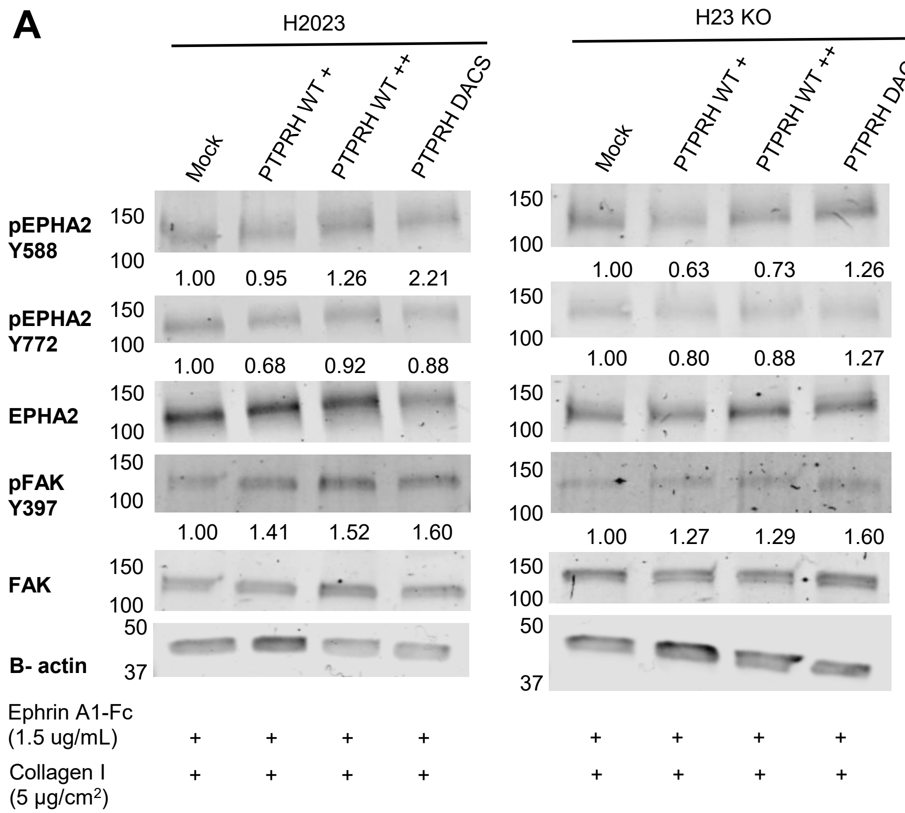


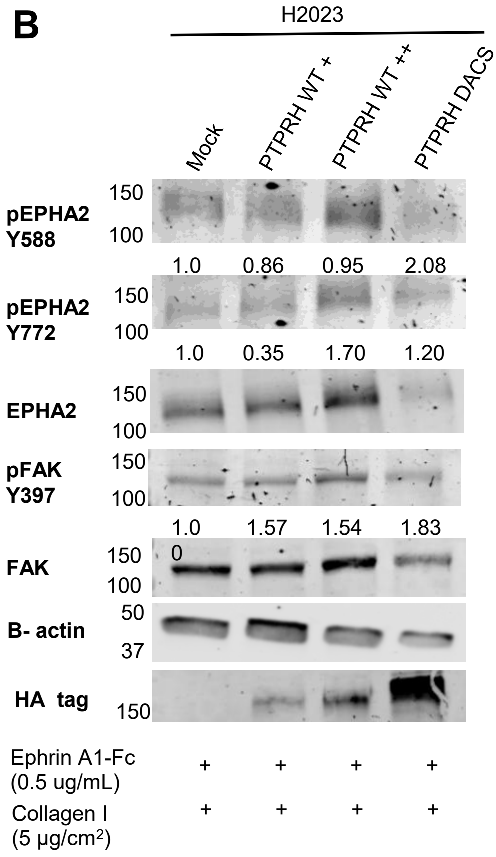


**Supplemental Figure 13. Phosphorylation of EPHA2 and FAK in lung adenocarcinoma cells overexpressing PTPRH WT or DACS.** *A-B)* Cells were grown in dishes thin-coated with collagen I, serum starved overnight, and stimulated with 1.5 ug/mL (*A*) or 0.5 ug/mL (*B*) of Ephrin A1-Fc for 15 minutes. Phosphorylation of EPHA2 at Y588 and Y772 (ligand-dependent pathway) and FAK at Y397 (activation marker) sites were assessed in the cell lysates by immunoblot. The densitometry of the bands are represented as fold change in comparison to the mock group.

## S-14

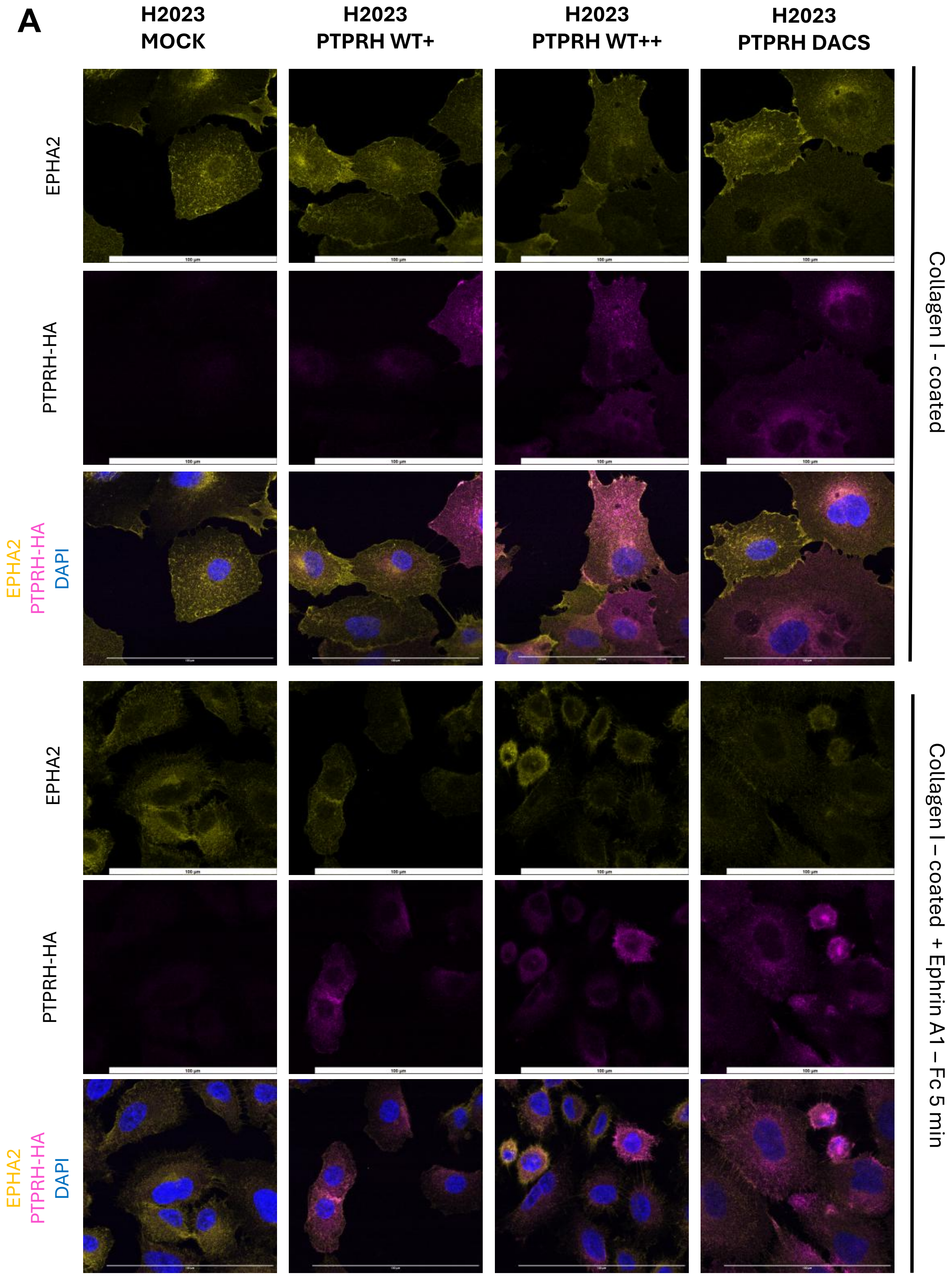

A
H2023 MOCK
H2023 PTPRH WT+
H2023 PTPRH WT++
H2023 PTPRH DACS
EPHA2
PTPRH-HA
EPHA2
PTPRH-HA
DAPI
EPHA2
PTPRH-HA
EPHA2
PTPRH-HA
DAPI
Collagen I - coated
Collagen I – coated + Ephrin A1 – Fc 5 min

#### S-14 *(continued)*

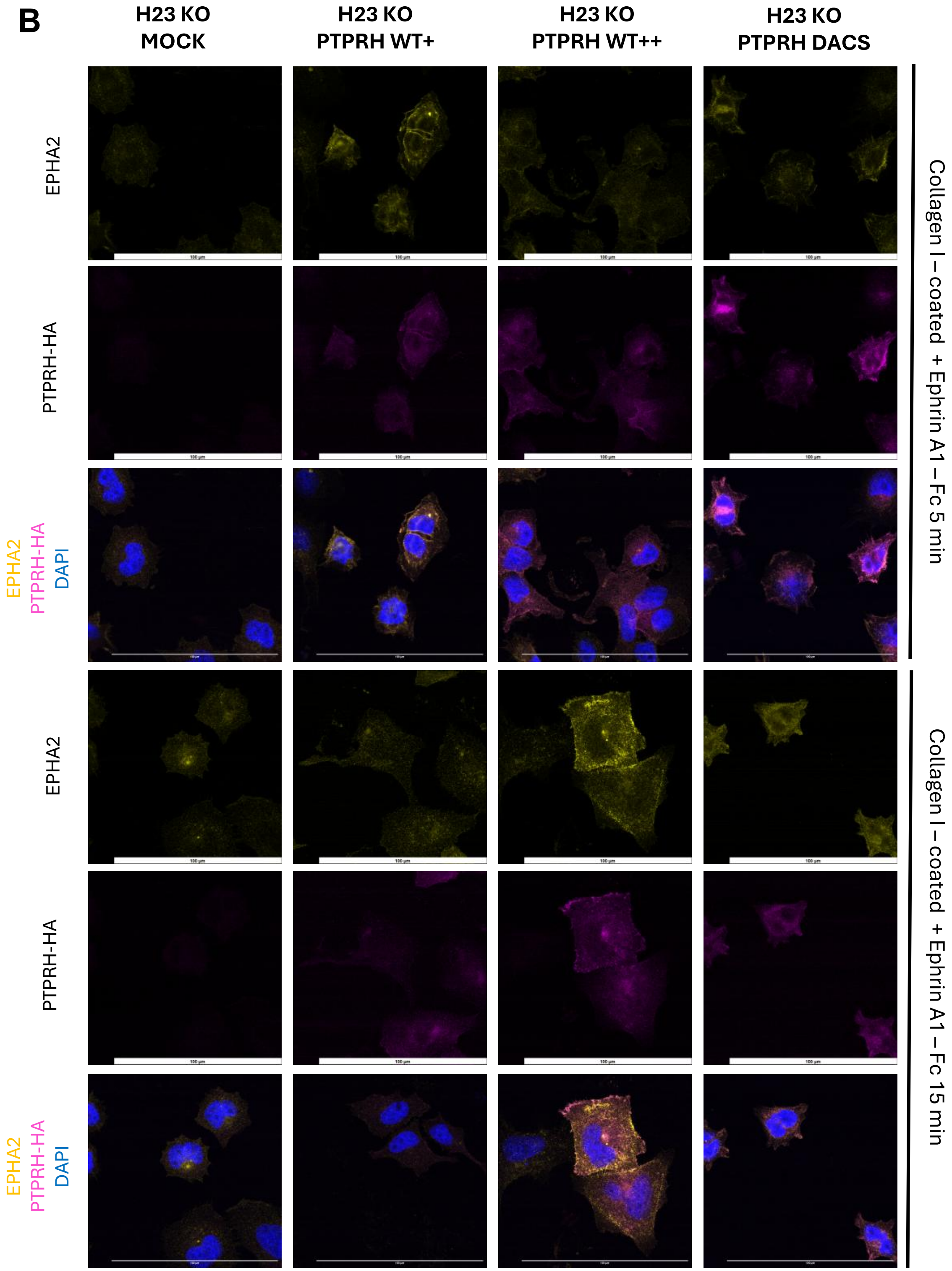


**Supplemental Figure 14. Spatial distribution of exogenous PTPRH-HA (WT or substrate-trapping/DACS) and EPHA2 in lung adenocarcinoma cells lines.** *A-B)* Representative images of EPHA2 (yellow) and exogenous PTPRH constructs tagged with HA (magenta) when cells are plated in over a thin-coat of collagen I with and without Ephrin A1-Fc stimulation (1.0 ug/mL) for the indicated time. *Scale bar* =100 µm, *magnification* = 100x.

**S-15**

**A**

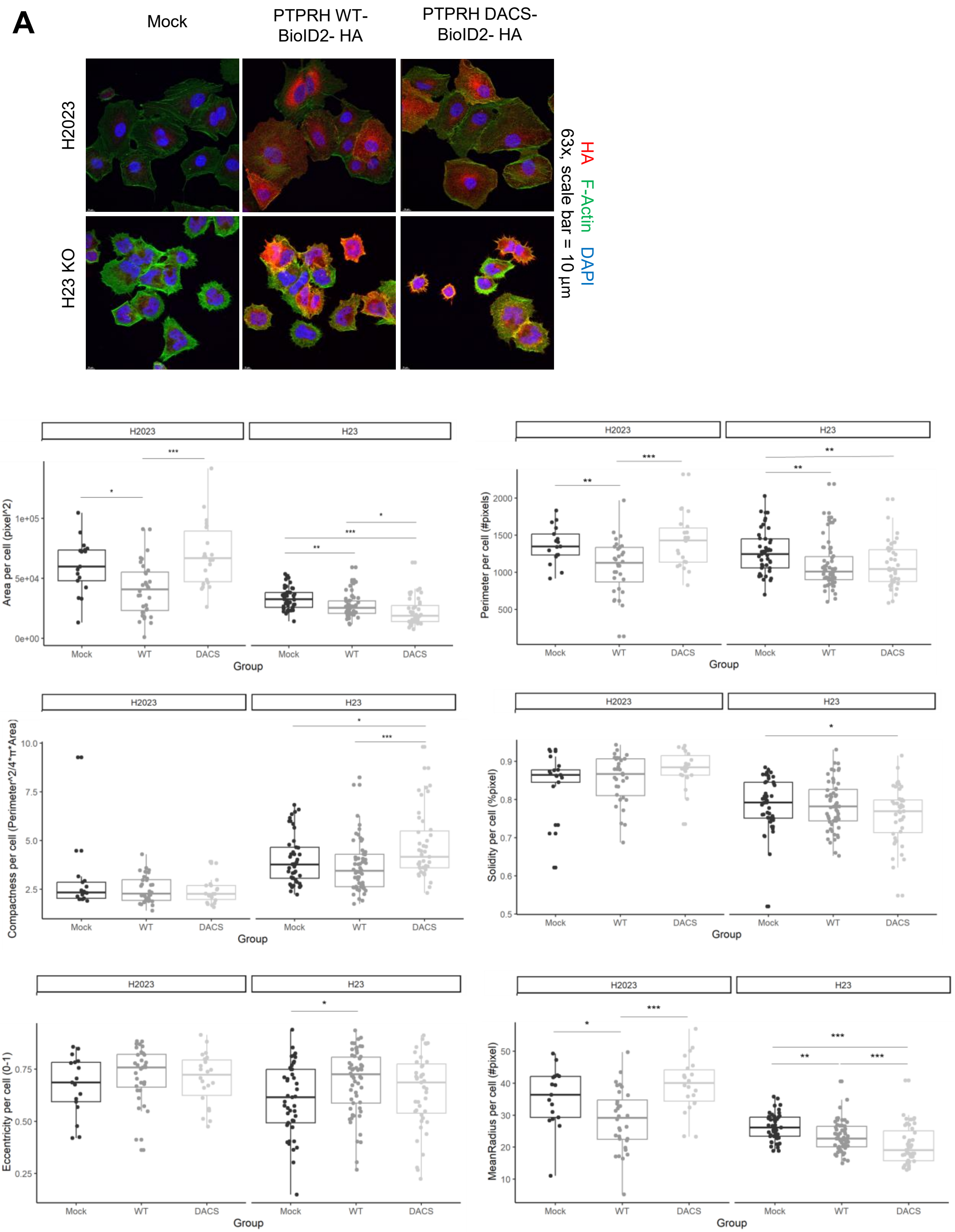
Mock
PTPRH WT-BioID2- HA
PTPRH DACS-BioID2- HA
H2023
H23 KO
HA F-Actin DAPI
63x, scale bar = 10 µm
H2023
H23
Area per cell (pixel^2)
Perimeter per cell (#pixels)
Compactness per cell (Perimeter^2/4*π*Area)
Solidity per cell (%pixel)
Eccentricity per cell (0-1)
MeanRadius per cell (#pixel)
Mock
WT
DACS
Group

#### ***S-15 (continued)***

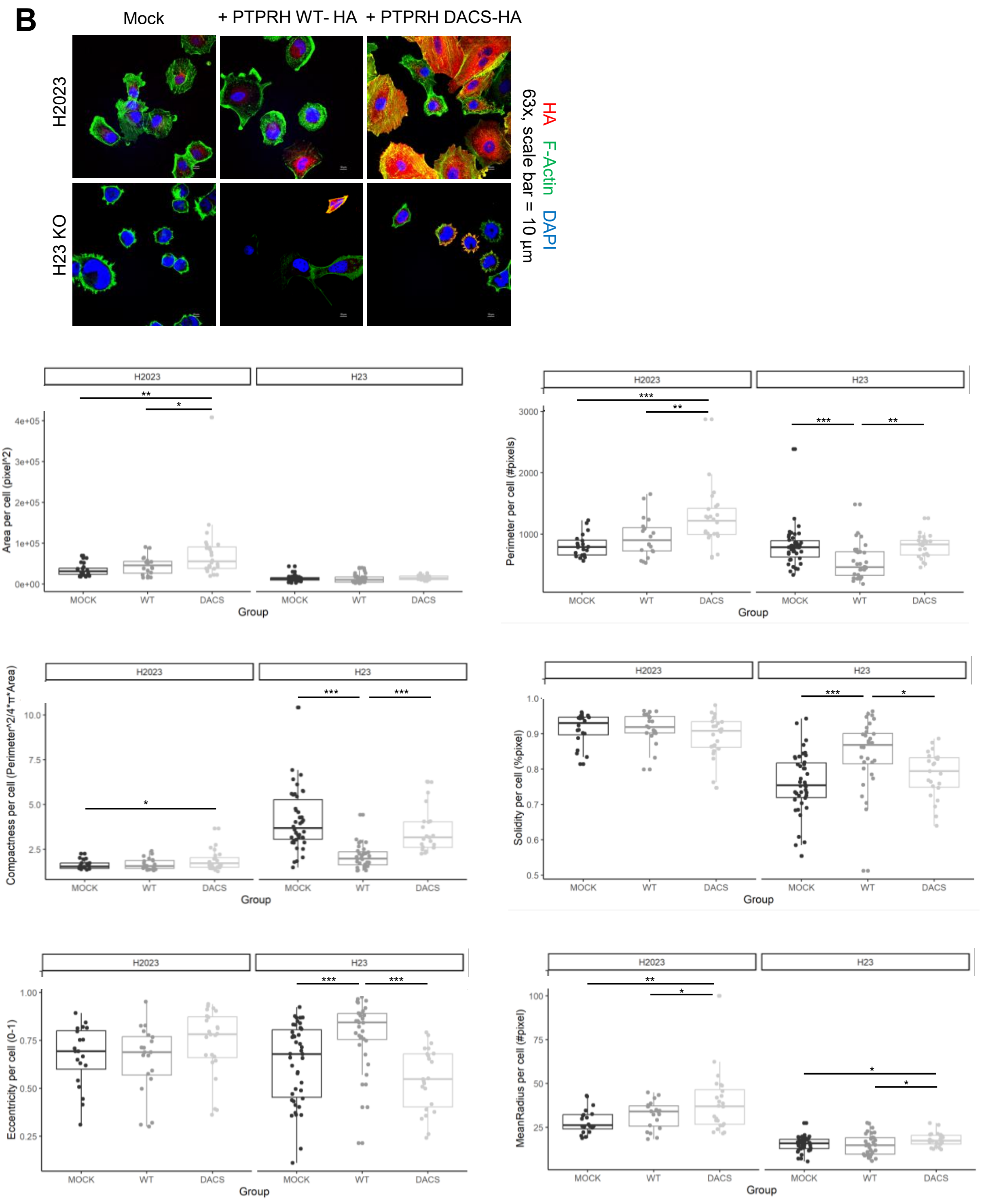

### S-15 *(continued)*

C

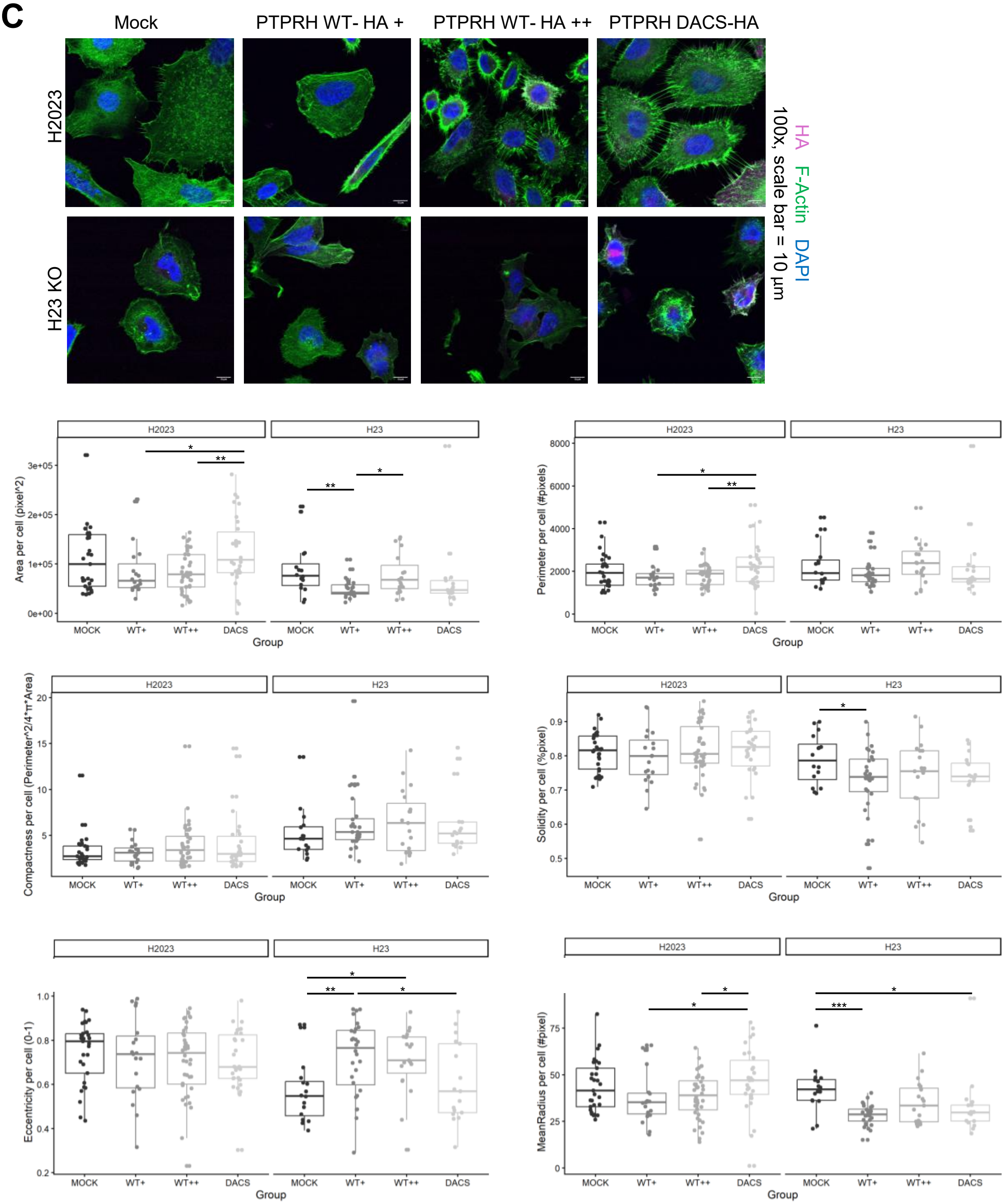

**Supplemental Figure 15. Size and shape measurements of lung adenocarcinoma cell lines overexpressing PTPRH WT-HA or PTPRH-DACS-HA.** Cell morphology parameters of H2023 and H23 KO cells in which exogenous PTPRH WT-HA or PTPRH DACS-HA with (*A*) or without the BioID2 tag (*B*-*C*) was overexpressed were extracted based on the confocal images using *Cell Profiler.* Cells were grown in glass slides (*A*-*B*) or glass slides thin-coated with collagen I (5 µg/cm$^2$) and stimulated with Ephrin A1-Fc (1.0 µg/mL) for 5 or 15 minutes (*C*). Representative images are demonstrated in the upper panel. The mock cells for each cell line were used as a control. Plots representing the area, perimeter, compactness, solidity, eccentricity, and mean radius per cell for each cell line are shown. Quantification values are represented in pixels and eccentricity is given in the 0-1 range, in which 0 = perfect round shape (circle) and 1= line segment. Statistical analysis was performed using One-Way ANOVA. Dots in the plots represent individual cell values, and the mean +/- s.d is shown. *p<0.05, **p<0.01, and ***p<0.001.

# S-16

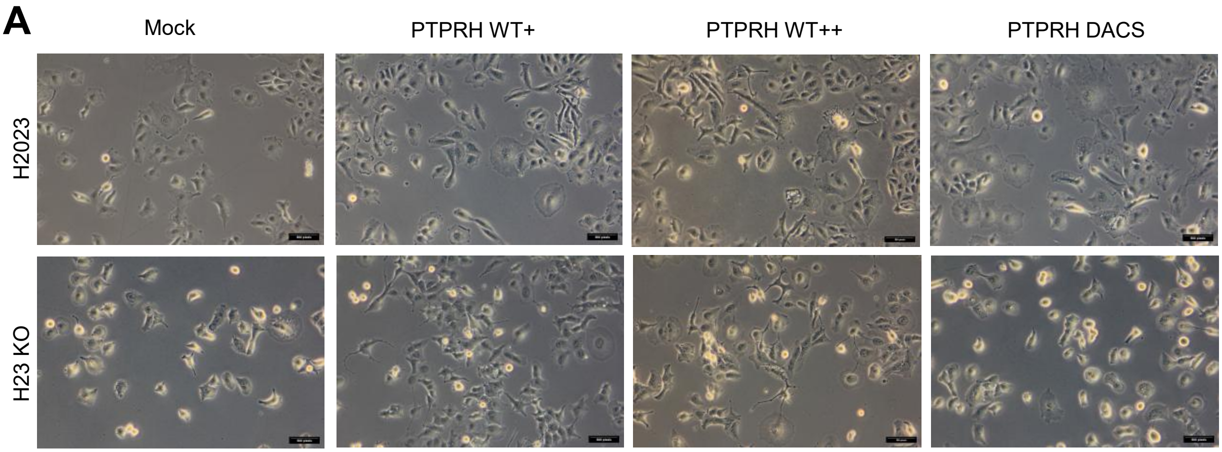


**Supplemental Figure 16. Bright field images of lung adenocarcinoma cell lines overexpressing PTPRH WT or DACS**. Images of live cells in monolayer culture. *Scale bar*= 500 pixels, *magnification* = 10x.

**S-17**

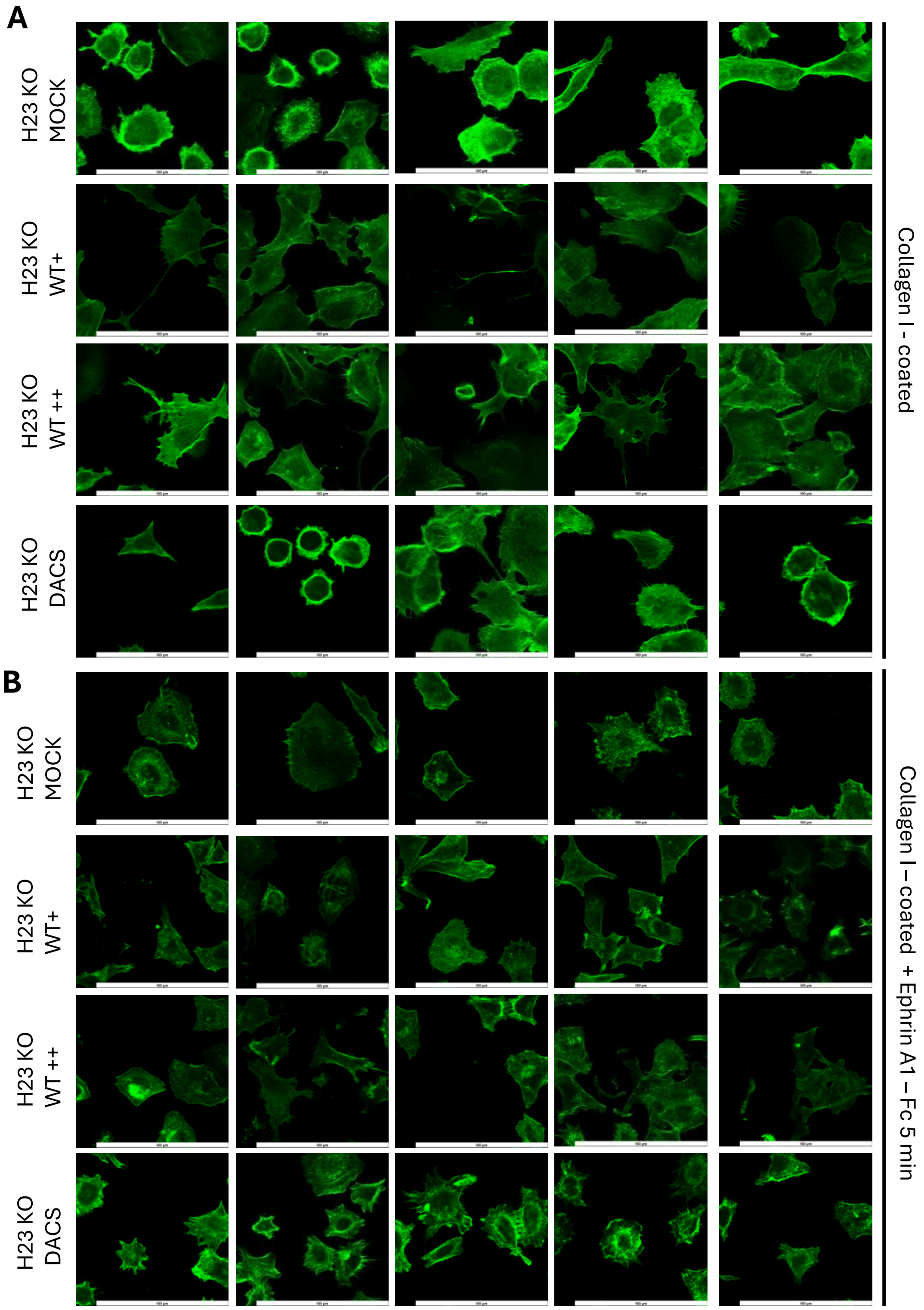
A
H23 KO MOCK
H23 KO WT+
H23 KO WT ++
H23 KO DACS
Collagen I - coated
B
H23 KO MOCK
H23 KO WT+
H23 KO WT ++
H23 KO DACS
Collagen I – coated + Ephrin A1 – Fc 5 min

#### S-17 *(continued)*

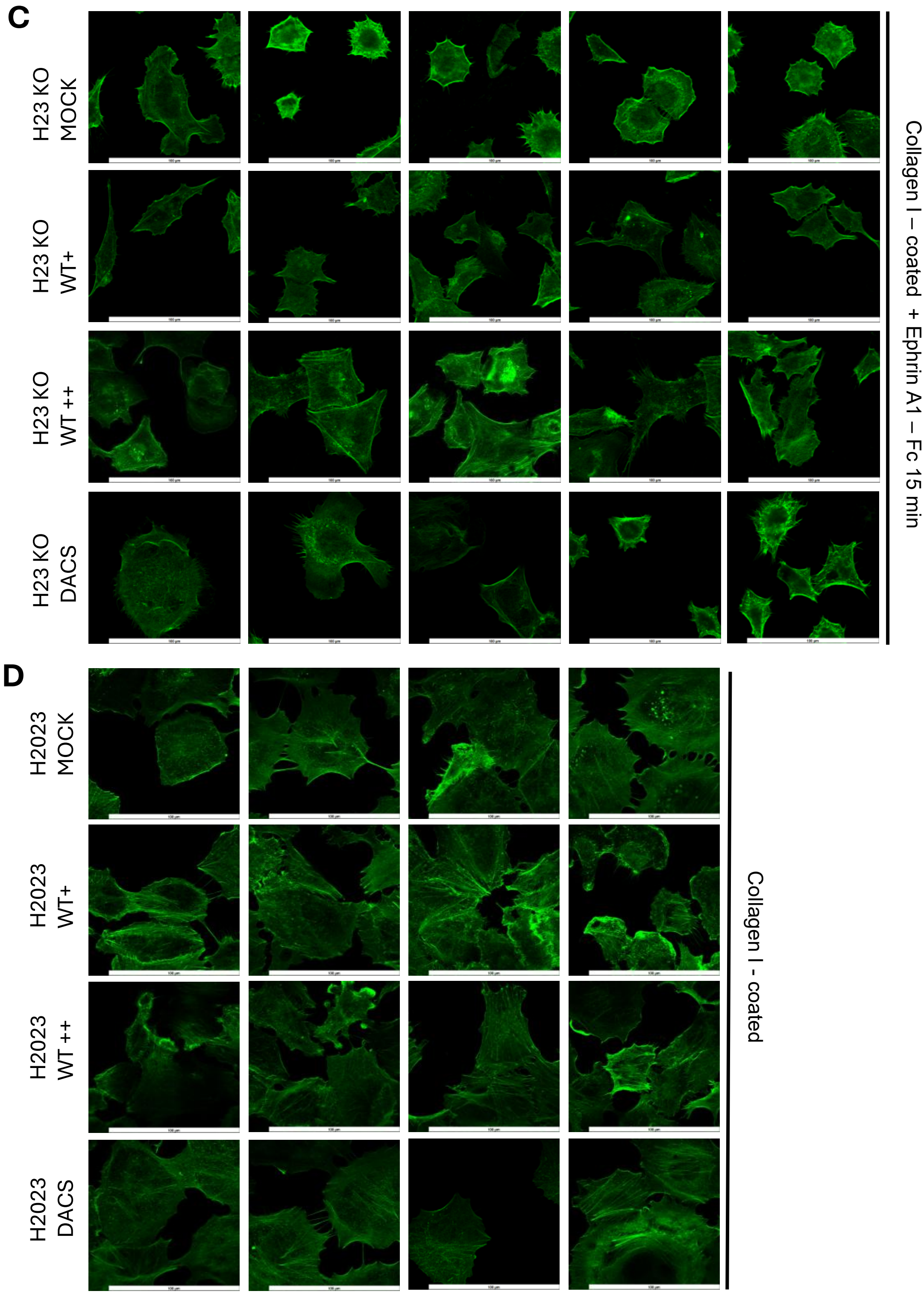

#### S-17 *(continued)*

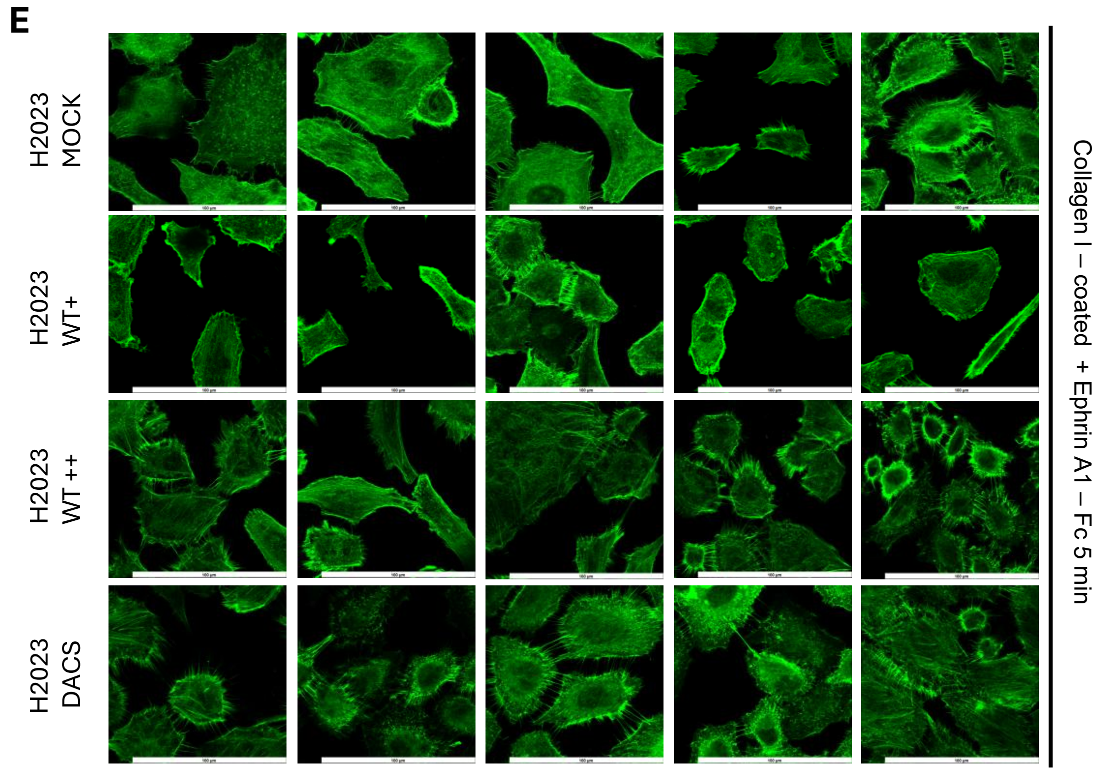


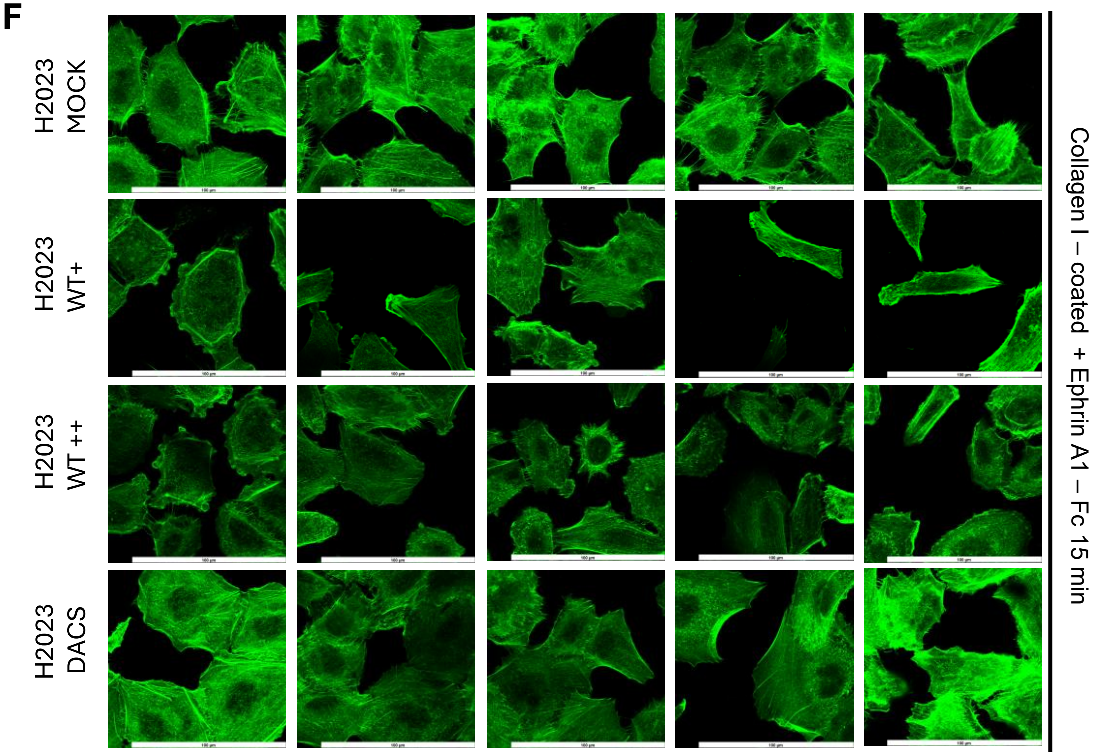


**Supplemental Figure 17. Cytoskeleton organization of H23 *PTPRH* KO and H2023 cells in which PTPRH WT-HA or PTPRH-DACS-HA expression was rescued/overexpressed.** *A-F)* Images of F-actin filaments organization when cells are plated over a thin-coat of collagen I (5 µg/cm$^2$) with and without Ephrin A1-Fc stimulation (1.0 ug/mL) for the indicated time. *Scale bar* =100 µm, *magnification* = 100x

## S-18

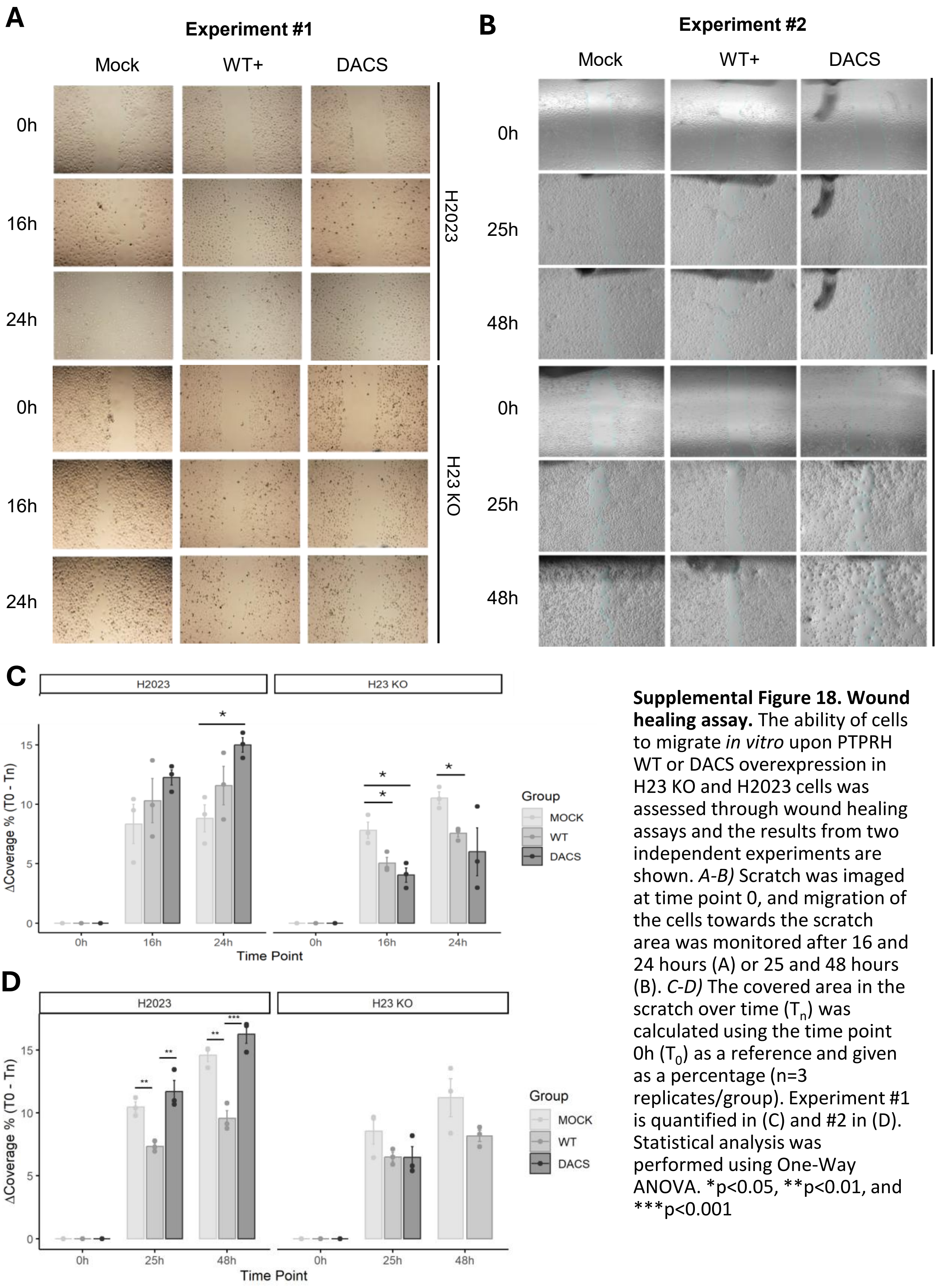


**Supplemental Figure 18. Wound healing assay.** The ability of cells to migrate *in vitro* upon PTPRH WT or DACS overexpression in H23 KO and H2023 cells was assessed through wound healing assays and the results from two independent experiments are shown. *A-B)* Scratch was imaged at time point 0, and migration of the cells towards the scratch area was monitored after 16 and 24 hours (A) or 25 and 48 hours (B). *C-D)* The covered area in the scratch over time ($T_n$) was calculated using the time point 0h ($T_0$) as a reference and given as a percentage (n=3 replicates/group). Experiment #1 is quantified in (C) and #2 in (D). Statistical analysis was performed using One-Way ANOVA. *p<0.05, **p<0.01, and ***p<0.001

**S-19**

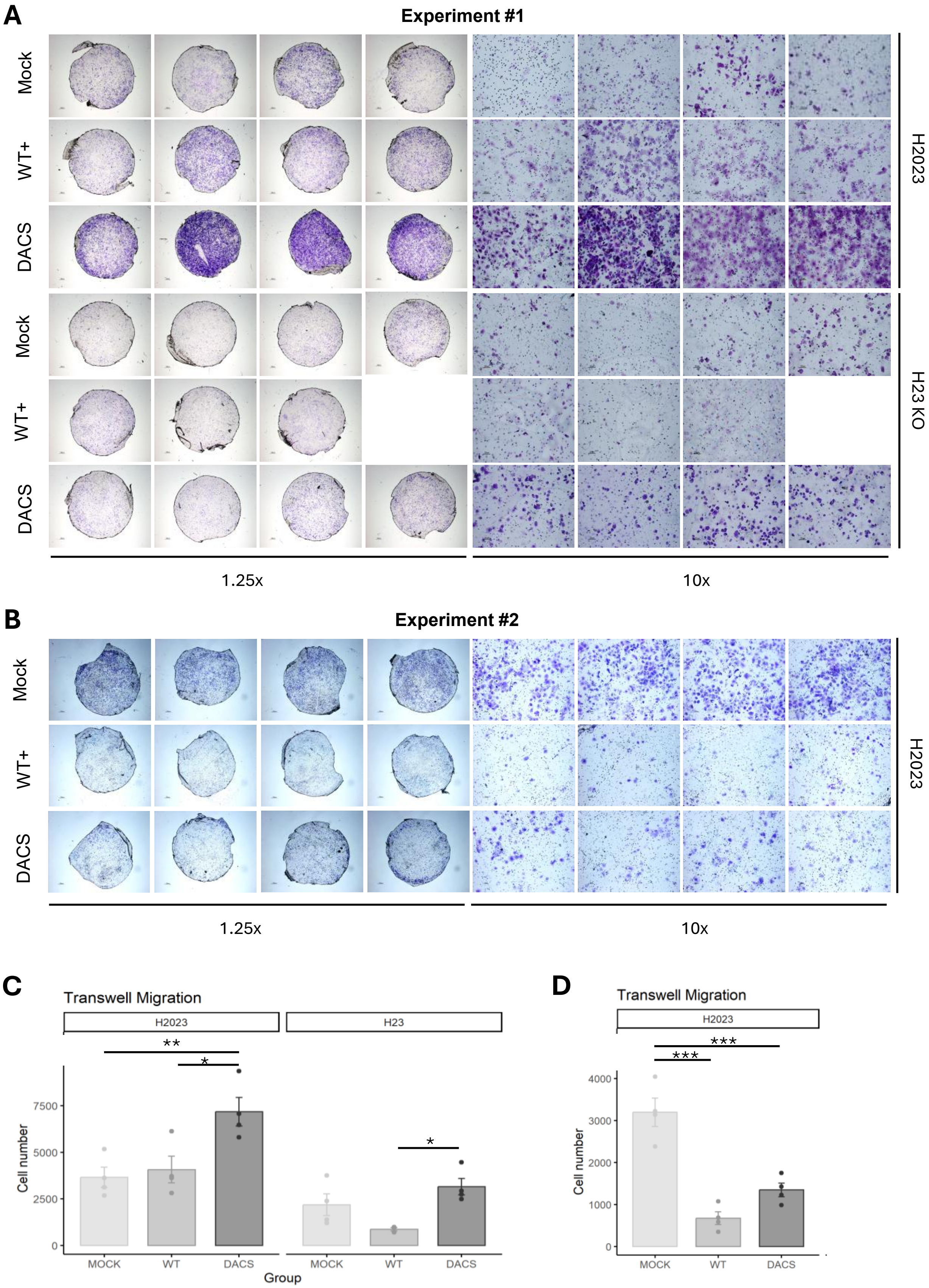

A
Experiment #1
Mock
WT+
DACS
Mock
WT+
DACS
H2023
H23 KO
1.25x
10x
B
Experiment #2
Mock
WT+
DACS
H2023
1.25x
10x
C
Transwell Migration
H2023
H23
**
*
*
Cell number
7500
5000
2500
0
MOCK
WT
DACS
MOCK
WT
DACS
Group
D
Transwell Migration
H2023
***
***
Cell number
4000
3000
2000
1000
0
MOCK
WT
DACS

**Supplemental Figure 19. Transwell migration.** The ability of cells to migrate *in vitro* upon PTPRH WT or DACS overexpression in H23 KO and H2023 cells was assessed through transwell assays. Membranes contained 8 µm pore sizes . *A-B)* After 24 hours, membranes from mock, WT, and DACS groups were stained with crystal violet for visualization of the migrating cells. Two independent experiments were performed to evaluate the migration behavior of H2023-derived cells. All replicates were imaged using 1.25x and 10x magnification. *C-D)* The total number of migrating cells in each replicate per group is demonstrated in the bar plot (n=4 replicates/group) and the mean +/- s.d is shown. Experiment #1 is quantified in (C) and #2 in (D). Statistical analysis was performed using One-Way ANOVA. *$p<0.05$, **$p<0.01$, and ***$p<0.001$

## S-20

**A**

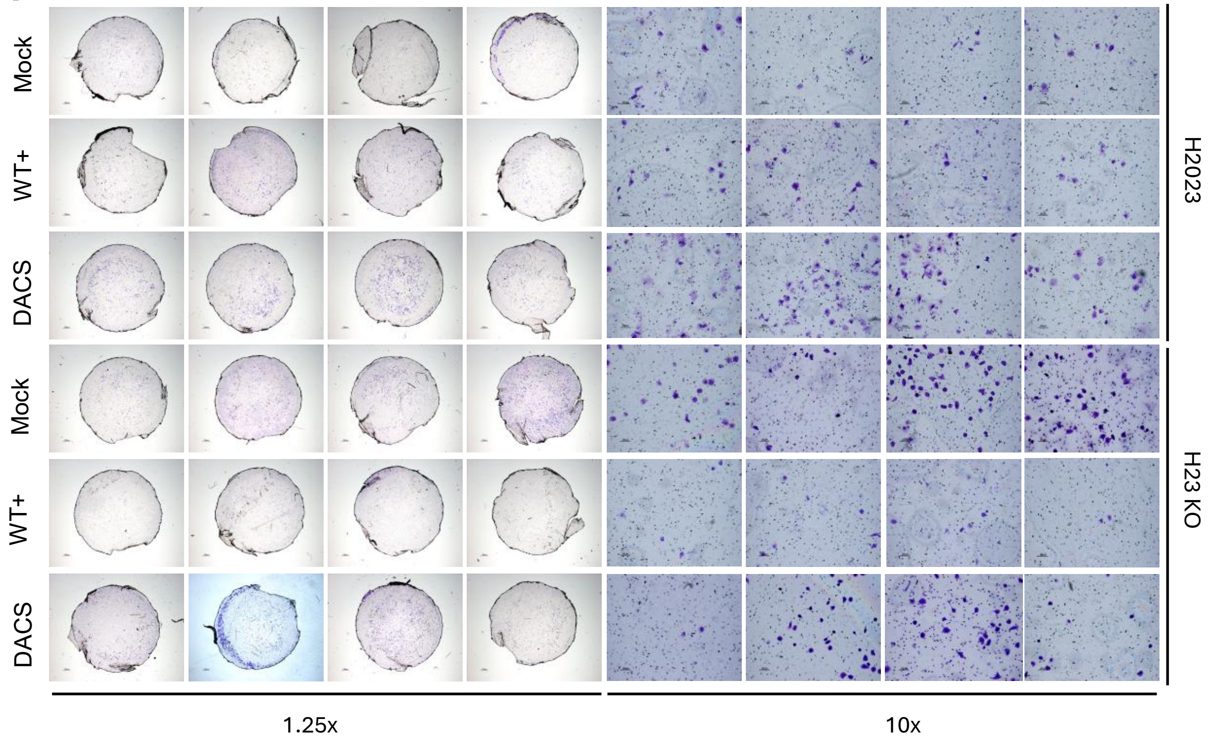


**B**

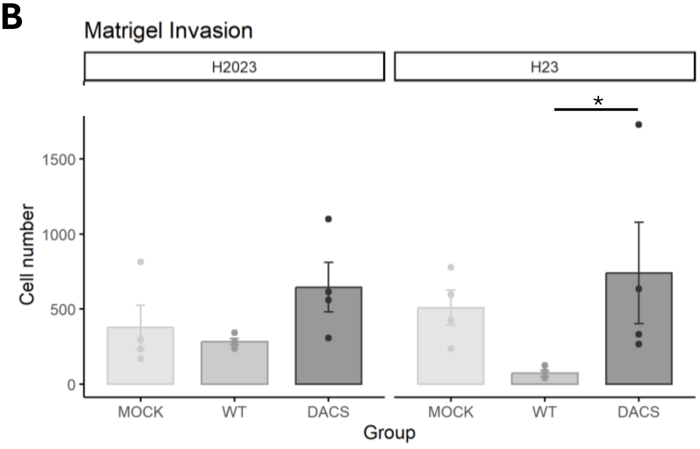


**Supplemental Figure 20. Evaluation of in *vitro* invasion .** The ability of invade *in vitro* upon PTPRH WT or DACS overexpression in H23 KO and H2023 cells was assessed through transwell assays. Membranes contained 8 µm pore sizes and were coated with Matrigel. *A)* After 24 hours, membranes from mock, WT, and DACS groups were stained with crystal violet for visualization of the migrating cells. All replicates were imaged using 1.25x and 10x magnification. *B)* The total number of invading cells in each replicate per group is demonstrated in the bar plot (n=4 replicates/group) and the mean +/- s.d is shown. Statistical analysis was performed using One-Way ANOVA. *p<0.05, **p<0.01, and ***p<0.001

## S-21

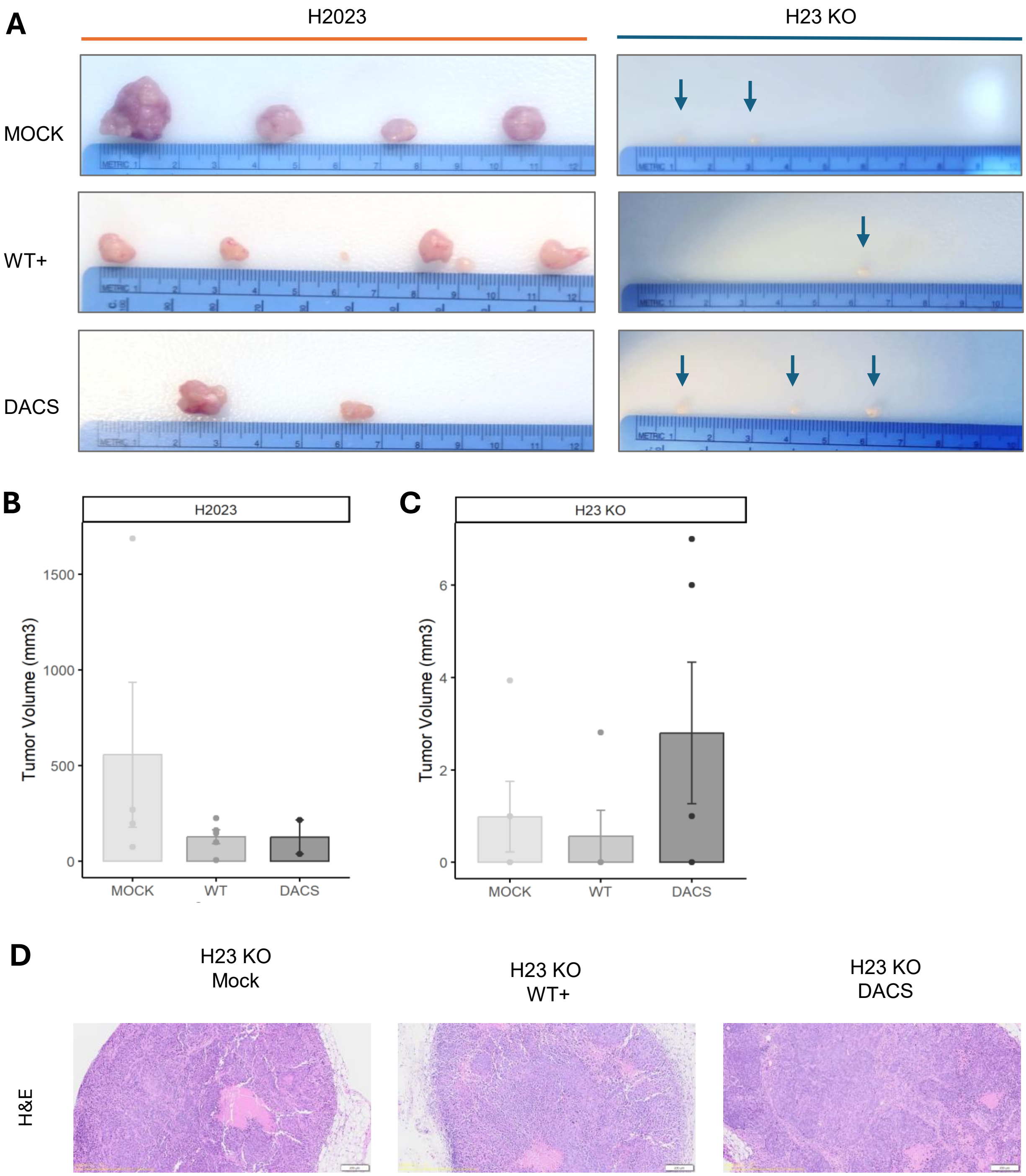


**Supplemental Figure 21. Tumor growth of lung adenocarcinoma cells overexpressing PTPRH WT or DACS *in vivo*.** *A)* Images showing the tumors collected from each NOD-SCID mouse injected with the various cell lines at the endpoint (day 62). The experiment started with n=5 mice/group. On day 62, H2023 mock group contained 4 mice, H2023 PTPRH-DACS group contained 2 mice, and H2023 PTPRH-WT, five mice. *B-C*) The tumor volume of the tumors in (*A*) were calculated using the length and width and demonstrated in the bar plot. The mean +/- s.d is shown. Statistical analysis was performed using One-Way ANOVA. *p<0.05, **p<0.01, and ***p<0.001. *D)* Representative histology (H&E staining) of flank tumors overexpressing PTPRH WT or DACS using 4x magnification (scale bar = 200 µm).

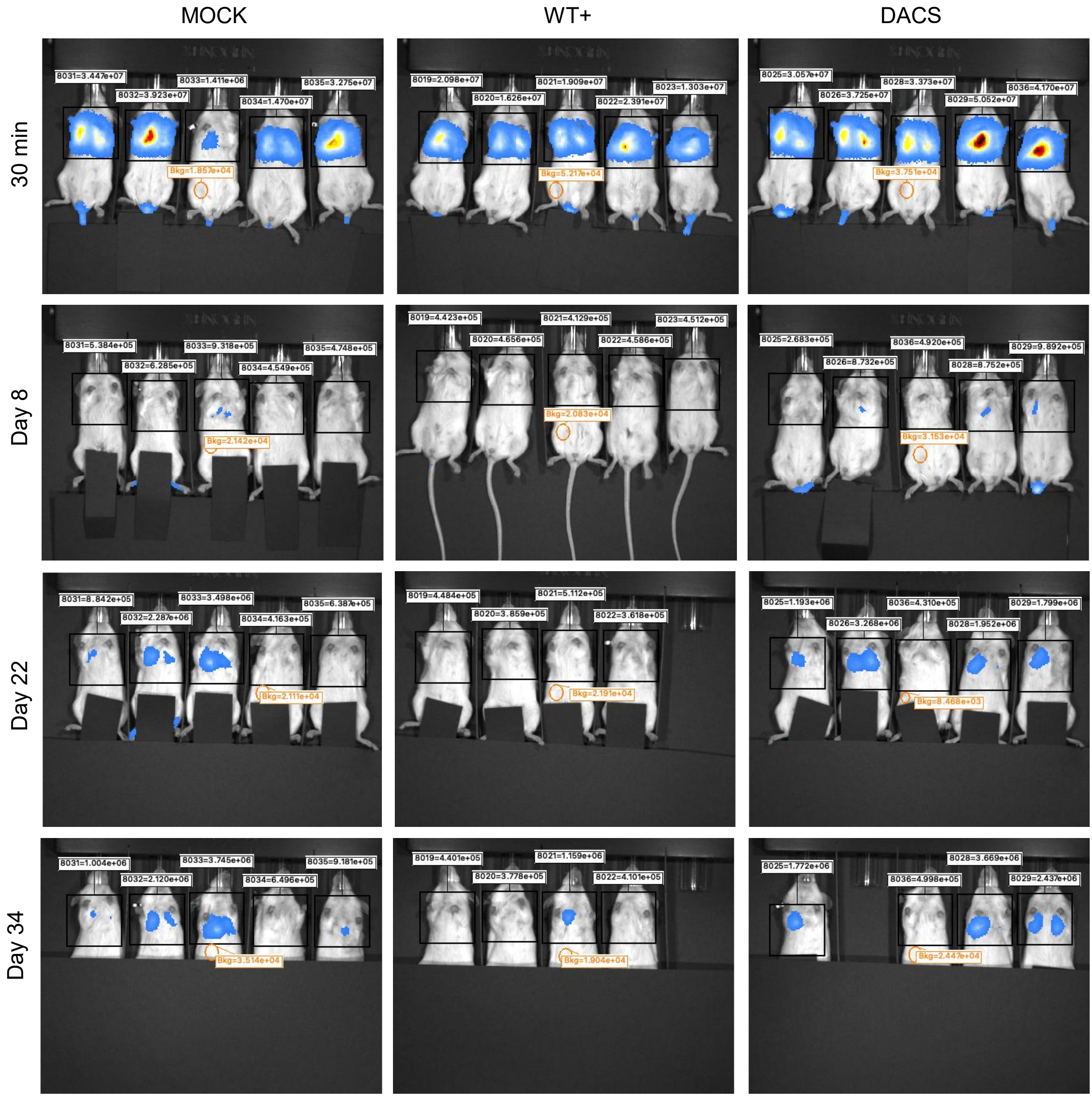


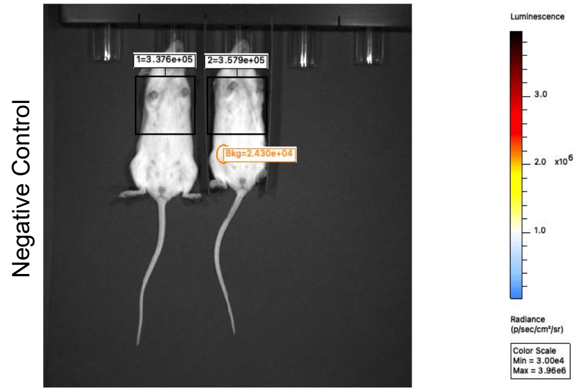


**Supplemental Figure 22. Colonization of H2023 cells overexpressing PTPRH WT or DACS in the lungs over time.** Images showing the luminescence signal in the lungs of each NOD-SCID mouse after tail vein injection of H2023 cells labeled with mCherry and luciferase. The individual luminescence is expressed as total flux (radiance units, photons per second (p/s)) and was calculated within the regions of interest (black boxes). The total flux of a background area (orange circles) and the lungs of two negative control mice (not injected) are also shown.
