## Supplemental Table 1 for "Unraveling the role of receptor-like protein tyrosine phosphatase PTPRH in cell signaling regulation and biological processes of non-small cell lung cancer"

Andrechek - Ortiz Combined Data Feb - June 2024

| # | Identified Proteins (1233) | Acc Num | Alt ID | CRAPome Score | MW | Protein Grouping Ambiguity | T-test p-value | FC by Category | Quantitative Profile | Quantitative Value (Normalized Total Precursor Intensity) BioID H2023 BioID Only | BioID H2023 BioID #1 | BioID H2023 BioID #3 | PTPRH H2023 PTPRH WT | PTPRH H2023 PTPRH 5+6+10 | PTPRH H2023 PTPRH 4+7+8 |
|---|---|---|---|---|---|---|---|---|---|---|---|---|---|---|---|
| 1 | Receptor-type tyrosine-protein phosphatase H OS=Ho | Q9HD43 | PTPRH | 0 | 122 kDa | | 0.12 | 110000 [] | | 1,000.00 | 1,000.00 | 1,000.00 | 1865800 | 1.95E+08 | 1.38E+08 |
| 2 | Thrombospondin-1 OS=Homo sapiens OX=9606 GN= | P07996 | THBS1 | 14 | 129 kDa | | 0.22 | 100000 [] | | 1,000.00 | 1,000.00 | 1,000.00 | 2.38E+08 | 2.48E+07 | 3.84E+07 |
| 3 | Aly/REF export factor OS=Homo sapiens OX=9606 G | E9PB61 (+1) | ALYREF | 390 | 28 kDa | | 0.37 | 14000 [] | | 1,000.00 | 1,000.00 | 1,000.00 | 1,000.00 | 4.20E+07 | 1,000.00 |
| 4 | Glyceraldehyde-3-phosphate dehydrogenase OS=Hom | P04406 | GAPDH | 458 | 36 kDa | | 0.027 | 13000 [] | | 1,000.00 | 1,000.00 | 1,000.00 | 1.76E+07 | 5574600 | 1.72E+07 |
| 5 | Peptidyl-prolyl cis-trans isomerase A OS=Homo sapier | P62937 (+1) | PPIA | 439 | 18 kDa | | 0.14 | 13000 [] | | 1,000.00 | 1,000.00 | 1,000.00 | 1,000.00 | 2.34E+07 | 1.46E+07 |
| 6 | Hemoglobin subunit alpha 2 (Fragment) OS=Homo sa | A0A2R8Y7C0 (+2) | HBA2 | 90 | 14 kDa | | 0.12 | 10000 [] | | 1,000.00 | 1,000.00 | 1,000.00 | 1,000.00 | 1.61E+07 | 1.45E+07 |
| 7 | Liprin-beta-1 OS=Homo sapiens OX=9606 GN=PPFII | Q86W92 | PPFIBP1 | 23 | 114 kDa | TRUE | 0.15 | 10000 [] | | 1,000.00 | 1,000.00 | 1,000.00 | 1,000.00 | 1.93E+07 | 1.07E+07 |
| 8 | Protein disulfide-isomerase A4 OS=Homo sapiens OX | P13667 | PDIA4 | 168 | 73 kDa | | 0.37 | 9800 [] | | 1,000.00 | 1,000.00 | 1,000.00 | 2.93E+07 | 1,000.00 | 1,000.00 |
| 9 | Transcriptional activator protein Pur-alpha OS=Homo | Q00577 | PURA | 55 | 35 kDa | TRUE | 0.12 | 8100 [] | | 1,000.00 | 1,000.00 | 1,000.00 | 1,000.00 | 1.37E+07 | 1.04E+07 |
| 10 | Cluster of phosphopyruvate hydratase OS=Homo sapi | A0A2R8Y6G6 [6] | ENO1 | 478 | 47 kDa | TRUE | 0.12 | 7400 [] | | 1,000.00 | 1,000.00 | 1,000.00 | 1,000.00 | 1.07E+07 | 1.13E+07 |
| 11 | Desmoplakin OS=Homo sapiens OX=9606 GN=DSP 1 | P15924 | DSP | 328 | 332 kDa | | 0.24 | 6200 [] | | 1,000.00 | 1,000.00 | 1,000.00 | 3567900 | 1,000.00 | 1.52E+07 |
| 12 | 116 kDa U5 small nuclear ribonucleoprotein compone | Q15029 | EFTUD2 | 381 | 109 kDa | TRUE | 0.37 | 6200 [] | | 1,000.00 | 1,000.00 | 1,000.00 | 1.85E+07 | 1,000.00 | 1,000.00 |
| 13 | Cluster of Serine/threonine-protein phosphatase 6 regu | O15084 [3] | ANKRD28 | 53 | 113 kDa | TRUE | 0.18 | 6000 [] | | 1,000.00 | 1,000.00 | 1,000.00 | 1,000.00 | 5076600 | 1.28E+07 |
| 14 | U3 small nucleolar RNA-associated protein 18 homolo | Q9Y5J1 | UTP18 | 64 | 62 kDa | | 0.16 | 5900 [] | | 1,000.00 | 1,000.00 | 1,000.00 | 5869300 | 1.18E+07 | 1,000.00 |
| 15 | Thioredoxin-dependent peroxide reductase, mitochond | P30048 | PRDX3 | 315 | 28 kDa | | 0.21 | 5100 [] | | 1,000.00 | 1,000.00 | 1,000.00 | 1,000.00 | 3663000 | 1.16E+07 |
| 16 | DNA polymerase delta interacting protein 3 OS=Hom | F6VRR5 (+3) | POLDIP3 | 247 | 48 kDa | | 0.37 | 4400 [] | | 1,000.00 | 1,000.00 | 1,000.00 | 1.31E+07 | 1,000.00 | 1,000.00 |
| 17 | Spermatogenesis-associated serine-rich protein 2 OS=I | Q86XZ4 | SPATS2 | 24 | 60 kDa | | 0.37 | 4200 [] | | 1,000.00 | 1,000.00 | 1,000.00 | 1.25E+07 | 1,000.00 | 1,000.00 |
| 18 | Helicase with zinc finger domain 2 OS=Homo sapiens | Q9BYK8 | HELZ2 | 4 | 295 kDa | | 0.3 | 4100 [] | | 1,000.00 | 1,000.00 | 1,000.00 | 1.10E+07 | 1339000 | 1,000.00 |
| 19 | Regulator of microtubule dynamics 3 (Fragment) OS=I | H0YLG5 (+2) | RMDN3 | 2 | 31 kDa | | 0.37 | 4000 [] | | 1,000.00 | 1,000.00 | 1,000.00 | 1.21E+07 | 1,000.00 | 1,000.00 |
| 20 | Cluster of NOP56 ribonucleoprotein (Fragment) OS=I | H0YDU4 [4] | NOP56 | 208 | 28 kDa | TRUE | 0.32 | 3800 [] | | 1,000.00 | 1,000.00 | 1,000.00 | 1,000.00 | 865590 | 1.07E+07 |
| 21 | Heat shock protein family B (small) member 1 OS=Ho | A0A6Q8PFK8 (+2) | HSPB1 | 128 | 22 kDa | | 0.23 | 3700 [] | | 1,000.00 | 1,000.00 | 1,000.00 | 1,000.00 | 2449400 | 8788500 |
| 22 | Cluster of Proline-, glutamic acid- and leucine-rich pro | C9JFV4 [4] | PELP1 | 87 | 125 kDa | TRUE | 0.23 | 3300 [] | | 1,000.00 | 1,000.00 | 1,000.00 | 7656200 | 2148700 | 1,000.00 |
| 23 | WD repeat domain 6 OS=Homo sapiens OX=9606 GN | A0A087X295 (+2) | WDR6 | 12 | 125 kDa | | 0.37 | 2700 [] | | 1,000.00 | 1,000.00 | 1,000.00 | 8082400 | 1,000.00 | 1,000.00 |
| 24 | RNA helicase OS=Homo sapiens OX=9606 GN=DDX | B7Z6D5 (+1) | DDX27 | 107 | 87 kDa | | 0.37 | 2600 [] | | 1,000.00 | 1,000.00 | 1,000.00 | 1,000.00 | 1,000.00 | 7709900 |
| 25 | Heterogeneous nuclear ribonucleoprotein A0 OS=Hon | Q13151 | HNRNPA0 | 405 | 31 kDa | | 0.15 | 2500 [] | | 1,000.00 | 1,000.00 | 1,000.00 | 1,000.00 | 4826500 | 2580800 |
| 26 | Cluster of Tumor protein p53 binding protein 1 OS=H | A6NNK5 [4] | TP53BP1 | 146 | 209 kDa | TRUE | 0.37 | 2200 [] | | 1,000.00 | 1,000.00 | 1,000.00 | 6542000 | 1,000.00 | 1,000.00 |
| 27 | Fatty acyl-CoA reductase 1 OS=Homo sapiens OX=96 | Q8WVX9 | FAR1 | 26 | 59 kDa | | 0.37 | 1700 [] | | 1,000.00 | 1,000.00 | 1,000.00 | 4968500 | 1,000.00 | 1,000.00 |
| 28 | Ribosome biogenesis protein BMS1 homolog OS=Hon | Q14692 | BMS1 | 141 | 146 kDa | | 0.37 | 1600 [] | | 1,000.00 | 1,000.00 | 1,000.00 | 1,000.00 | 1,000.00 | 4780500 |
| 29 | Cluster of E3 ubiquitin-protein ligase OS=Homo sapie | A0A6Q8PGP5 [2] | NEDD4L | 3 | 109 kDa | TRUE | 0.37 | 1500 [] | | 1,000.00 | 1,000.00 | 1,000.00 | 4447700 | 1,000.00 | 1,000.00 |

| | | | | | | | | | | | | | | | |
|---|---|---|---|---|---|---|---|---|---|---|---|---|---|---|---|
| 31 | Myosin VA OS=Homo sapiens OX=9606 GN=MYO5 | G3V394 (+1) | MYO5A | 52 | 214 kDa | | 0.37 | 1400 | [] | 1,000.00 | 1,000.00 | 1,000.00 | 4048500 | 1,000.00 | 1,000.00 |
| 32 | E3 ubiquitin-protein ligase RNF31 OS=Homo sapiens | Q96EP0 | RNF31 | 0 | 120 kDa | | 0.37 | 1200 | [] | 1,000.00 | 1,000.00 | 1,000.00 | 3665100 | 1,000.00 | 1,000.00 |
| 33 | Threonine synthase-like 1 OS=Homo sapiens OX=960 | Q8IYQ7 | THNSL1 | 2 | 83 kDa | | 0.37 | 1200 | [] | 1,000.00 | 1,000.00 | 1,000.00 | 1,000.00 | 3629900 | 1,000.00 |
| 35 | Nuclear factor NF-kappa-B p100 subunit OS=Homo sa | Q00653 | NFKB2 | 11 | 97 kDa | | 0.37 | 1100 | [] | 1,000.00 | 1,000.00 | 1,000.00 | 3182400 | 1,000.00 | 1,000.00 |
| 36 | Protein arginine N-methyltransferase 5 OS=Homo sapi | O14744 | PRMT5 | 285 | 73 kDa | | 0.32 | 930 | [] | 1,000.00 | 1,000.00 | 1,000.00 | 230580 | 1,000.00 | 2568200 |
| 37 | Disks large homolog 5 OS=Homo sapiens OX=9606 C | Q8TDM6 | DLG5 | 69 | 214 kDa | | 0.37 | 680 | [] | 1,000.00 | 1,000.00 | 1,000.00 | 2049200 | 1,000.00 | 1,000.00 |
| 38 | DNA (cytosine-5)-methyltransferase OS=Homo sapien | A0A7I2V490 (+5) | DNMT1 | 185 | 187 kDa | | 0.37 | 560 | [] | 1,000.00 | 1,000.00 | 1,000.00 | 1665900 | 1,000.00 | 1,000.00 |
| 39 | RNA-binding protein 5 OS=Homo sapiens OX=9606 C | P52756 | RBM5 | 99 | 92 kDa | | 0.37 | 460 | [] | 1,000.00 | 1,000.00 | 1,000.00 | 1392700 | 1,000.00 | 1,000.00 |
| 40 | Probable ATP-dependent RNA helicase DDX56 OS=H | Q9NY93 | DDX56 | 115 | 62 kDa | | 0.37 | 340 | [] | 1,000.00 | 1,000.00 | 1,000.00 | 1004700 | 1,000.00 | 1,000.00 |
| 41 | Zinc finger CCCH-type containing 11A OS=Homo sap | E9PQ61 (+1) | ZC3H11A | 161 | 73 kDa | TRUE | 0.37 | 260 | [] | 1,000.00 | 1,000.00 | 1,000.00 | 785300 | 1,000.00 | 1,000.00 |
| 42 | Cluster of Profilin-1 OS=Homo sapiens OX=9606 GN | P07737 [2] | PFN1 | 308 | 15 kDa | TRUE | 0.37 | 260 | [] | 1,000.00 | 1,000.00 | 1,000.00 | 1,000.00 | 779520 | 1,000.00 |
| 43 | Cluster of Kinesin-like protein KIF18A OS=Homo sap | Q8NI77 [3] | KIF18A | 24 | 102 kDa | TRUE | 0.37 | 160 | [] | 1,000.00 | 1,000.00 | 1,000.00 | 1,000.00 | 474820 | 1,000.00 |
| 45 | 39S ribosomal protein L50, mitochondrial OS=Homo s | Q8N5N7 | MRPL50 | 35 | 18 kDa | | 0.068 | 35 | [] | 1129500 | 1,000.00 | 1,000.00 | 1.83E+07 | 1.89E+07 | 2937000 |
| 46 | Sideroflexin-1 OS=Homo sapiens OX=9606 GN=SFX | Q9H9B4 | SFXN1 | 140 | 36 kDa | | 0.37 | 25 | [] | 1,000.00 | 1,000.00 | 1,000.00 | 71783 | 1,000.00 | 1,000.00 |
| 50 | Exonuclease 3'-5' domain-containing protein 2 OS=Ho | Q9NVH0 | EXD2 | 3 | 70 kDa | | 0.087 | 11 | [] | 1154000 | 1,000.00 | 1,000.00 | 4471400 | 1235800 | 6968900 |
| 49 | dCTP pyrophosphatase 1 OS=Homo sapiens OX=960 | Q9H773 | DCTPP1 | 107 | 19 kDa | | 0.16 | 11 | [] | 6326800 | 510020 | 1007900 | 1,000.00 | 3.66E+07 | 5.23E+07 |
| 51 | 40S ribosomal protein S19 OS=Homo sapiens OX=96 | P39019 | RPS19 | 334 | 16 kDa | | 0.42 | 11 | [] | 1,000.00 | 446110 | 1,000.00 | 1,000.00 | 1,000.00 | 4800300 |
| 52 | RNA-binding protein 25 OS=Homo sapiens OX=9606 | P49756 | RBM25 | 237 | 100 kDa | | 0.42 | 10 | [] | 1,000.00 | 1,000.00 | 869090 | 1,000.00 | 1,000.00 | 8954800 |
| 53 | Cluster of KN motif and ankyrin repeat domain-contai | Q63ZY3 [2] | KANK2 | 87 | 91 kDa | TRUE | 0.42 | 8.7 | [] | 1623600 | 361800 | 1,000.00 | 1.71E+07 | 182780 | 1,000.00 |
| 54 | Cluster of Serine/threonine-protein phosphatase 2A 55 | P63151 [2] | PPP2R2A | 304 | 52 kDa | TRUE | 0.18 | 8.6 | [] | 2130300 | 1,000.00 | 1,000.00 | 1,000.00 | 7182700 | 1.11E+07 |
| 55 | Calponin-3 OS=Homo sapiens OX=9606 GN=CNN3 I | Q15417 | CNN3 | 213 | 36 kDa | | 0.36 | 8.5 | [] | 3.01E+07 | 1.24E+07 | 1.93E+07 | 4.74E+08 | 2.40E+07 | 2.88E+07 |
| 56 | La-related protein 4B OS=Homo sapiens OX=9606 G | Q92615 | LARP4B | 86 | 81 kDa | | 0.43 | 8.1 | [] | 1,000.00 | 400890 | 1,000.00 | 1,000.00 | 3246400 | 1,000.00 |
| 57 | Myosin IC OS=Homo sapiens OX=9606 GN=MYO1C | F5H6E2 (+1) | MYO1C | 152 | 119 kDa | | 0.069 | 7.9 | [] | 4.74E+07 | 5.36E+07 | 7452000 | 1.17E+08 | 2.78E+08 | 4.65E+08 |
| 58 | 3'-5' RNA helicase YTHDC2 OS=Homo sapiens OX= | Q9H6S0 | YTHDC2 | 133 | 160 kDa | | 0.0052 | 7.7 | [] | 3170000 | 7033000 | 1,000.00 | 3.06E+07 | 2.88E+07 | 1.91E+07 |
| 59 | Cluster of Myosin IB OS=Homo sapiens OX=9606 GN | E9PDF6 [3] | MYO1B | 159 | 128 kDa | TRUE | 0.015 | 7.7 | [] | 7.73E+07 | 1.69E+07 | 2.29E+07 | 1.90E+08 | 3.09E+08 | 4.02E+08 |
| 60 | Heat shock protein HSP 90-beta OS=Homo sapiens O | P08238 | HSP90AB1 | 573 | 83 kDa | TRUE | 0.058 | 7.7 | [] | 1,000.00 | 8825300 | 1,000.00 | 1.18E+07 | 2.06E+07 | 3.52E+07 |
| 61 | Glial fibrillary acidic protein (Fragment) OS=Homo sa | K7EPT8 | GFAP | 367 | 8 kDa | | 0.18 | 7.6 | [] | 1,000.00 | 2966500 | 1,000.00 | 1,000.00 | 1.33E+07 | 9272300 |
| 63 | Cluster of Phosphatidylinositol-4-phosphate 5-kinase t | A6PW57 [4] | PIP5K1A | 20 | 61 kDa | TRUE | 0.27 | 7.4 | [] | 1,000.00 | 1,000.00 | 2020300 | 1,000.00 | 1.13E+07 | 3624800 |
| 62 | 26S proteasome regulatory subunit 6B OS=Homo sapi | P43686 | PSMC4 | 210 | 47 kDa | TRUE | 0.44 | 7.4 | [] | 138690 | 1,000.00 | 1,000.00 | 1,000.00 | 1042300 | 1,000.00 |
| 65 | Cluster of Aldo-keto reductase family 1 member C2 O | B4DK69 [3] | AKR1C2 | 10 | 34 kDa | TRUE | 0.17 | 7.1 | [] | 6121700 | 4721100 | 2144100 | 1,000.00 | 3.94E+07 | 5.25E+07 |
| 64 | Protocadherin Fat 1 OS=Homo sapiens OX=9606 GN | Q14517 (+1) | FAT1 | 3 | 506 kDa | | 0.44 | 7.1 | [] | 1721900 | 1,000.00 | 1,000.00 | 1.22E+07 | 1,000.00 | 1,000.00 |
| 66 | Dihydrolipoamide branched chain transacylase E2 OS= | A0A7P0T9W1 (+2) | DBT | 143 | 33 kDa | | 0.15 | 6.9 | [] | 2308900 | 1,000.00 | 1,000.00 | 7702200 | 7729100 | 461410 |
| 67 | HBS1-like protein OS=Homo sapiens OX=9606 GN=I | Q9Y450 | HBS1L | 94 | 75 kDa | | 0.1 | 6.6 | [] | 1163900 | 1,000.00 | 3977400 | 2570200 | 1.48E+07 | 1.68E+07 |

| | | | | | | | | | | | | | | | |
|---|---|---|---|---|---|---|---|---|---|---|---|---|---|---|---|
| 68 | Heat shock protein HSP 90-alpha OS=Homo sapiens ( | P07900 | HSP90AA1 | 565 | 85 kDa | TRUE | 0.03 | 6.5 | [] | 1,000.00 | 8825300 | 1,000.00 | 1.18E+07 | 2.06E+07 | 2.52E+07 |
| 70 | Transcription elongation factor SPT6 OS=Homo sapie | Q7KZ85 | SUPT6H | 85 | 199 kDa | | 0.017 | 6.1 | [] | 1,000.00 | 4924100 | 1,000.00 | 7447800 | 1.21E+07 | 1.04E+07 |
| 69 | Nucleolar protein 16 OS=Homo sapiens OX=9606 GN | A0A0C4DGU5 (+3) | NOP16 | 57 | 26 kDa | | 0.046 | 6.1 | [] | 7928700 | 1,000.00 | 1,000.00 | 2.30E+07 | 9279900 | 1.65E+07 |
| 71 | LINE-1 retrotransposable element ORF1 protein OS=I | Q9UN81 | L1RE1 | 5 | 40 kDa | | 0.079 | 6.1 | [] | 4703100 | 6730000 | 1,000.00 | 3.88E+07 | 1.77E+07 | 1.28E+07 |
| 73 | N-acylneuraminate cytidylyltransferase OS=Homo sapi | Q8NFW8 | CMAS | 57 | 48 kDa | | 0.0053 | 5.8 | [] | 1.38E+07 | 1.26E+07 | 1,000.00 | 5.73E+07 | 5.70E+07 | 3.84E+07 |
| 72 | 60S ribosomal protein L34 OS=Homo sapiens OX=96 | P49207 | RPL34 | 247 | 13 kDa | | 0.28 | 5.8 | [] | 4147800 | 8.11E+07 | 1.82E+07 | 4.50E+08 | 1.37E+07 | 1.38E+08 |
| 74 | Protein transport protein SEC23 OS=Homo sapiens O | A0A2R8YFH5 (+1) | SEC23B | 95 | 84 kDa | TRUE | 0.45 | 5.7 | [] | 8879000 | 3151800 | 2192700 | 1,000.00 | 7.91E+07 | 1479400 |
| 76 | Transcription factor A, mitochondrial OS=Homo sapie | Q00059 | TFAM | 107 | 29 kDa | | 0.031 | 4.8 | [] | 1,000.00 | 3460200 | 7130200 | 2.33E+07 | 1.10E+07 | 1.65E+07 |
| 77 | A-kinase anchoring protein 8 OS=Homo sapiens OX=9 | A0A7P0T893 (+1) | AKAP8 | 111 | 75 kDa | | 0.34 | 4.8 | [] | 748000 | 1,000.00 | 1913300 | 1.02E+07 | 2275200 | 181580 |
| 79 | ATP binding cassette subfamily D member 3 OS=Hom | A0A3B3ITW3 (+1) | ABCD3 | 148 | 75 kDa | | 0.18 | 4.6 | [] | 1,000.00 | 1636200 | 5503000 | 2131400 | 1.96E+07 | 1.10E+07 |
| 80 | FYVE and coiled-coil domain-containing protein 1 OS | Q9BQS8 | FYCO1 | 5 | 167 kDa | TRUE | 0.32 | 4.5 | [] | 1968400 | 1,000.00 | 310840 | 7784500 | 2517400 | 1,000.00 |
| 81 | Cluster of 60S ribosomal protein L36a-like OS=Homo | Q969Q0 [3] | RPL36AL | 203 | 12 kDa | TRUE | 0.36 | 4.5 | [] | 1,000.00 | 1,000.00 | 7583400 | 1,000.00 | 6632400 | 2.75E+07 |
| 82 | Semaphorin-3C OS=Homo sapiens OX=9606 GN=SE | Q99985 | SEMA3C | 0 | 85 kDa | | 0.14 | 4.4 | [] | 2708100 | 1856700 | 1,000.00 | 1.16E+07 | 2219800 | 6376200 |
| 83 | Junction plakoglobin OS=Homo sapiens OX=9606 GN | P14923 | JUP | 244 | 82 kDa | | 0.41 | 4.2 | [] | 1757700 | 1437300 | 1899300 | 1.88E+07 | 2420500 | 1,000.00 |
| 85 | 28S rRNA (cytosine-C(5))-methyltransferase OS=Hon | Q96P11 | NSUN5 | 12 | 47 kDa | | 0.27 | 4 | [] | 951260 | 1,000.00 | 1,000.00 | 1,000.00 | 2233100 | 1533100 |
| 84 | Very-long-chain enoyl-CoA reductase OS=Homo sapi | Q9NZ01 | TECR | 192 | 36 kDa | | 0.34 | 4 | [] | 1686400 | 1,000.00 | 1,000.00 | 1,000.00 | 1727600 | 4994100 |
| 86 | Keratin, type I cytoskeletal 9 OS=Homo sapiens OX=9 | P35527 | KRT9 | 577 | 62 kDa | TRUE | 0.0049 | 3.9 | [] | 4.30E+07 | 7.50E+07 | 1.60E+08 | 4.21E+08 | 3.69E+08 | 3.06E+08 |
| 87 | Keratin, type II cytoskeletal 4 OS=Homo sapiens OX= | P19013 | KRT4 | 391 | 56 kDa | TRUE | 0.31 | 3.9 | [] | 6939800 | 3.96E+07 | 1,000.00 | 1,000.00 | 5.40E+07 | 1.27E+08 |
| 88 | Cluster of Keratin 10 OS=Homo sapiens OX=9606 GN | A0A1B0GVI3 [12] | KRT10 | 616 | 63 kDa | TRUE | 0.00024 | 3.8 | BioID low, ptprh high | 2.33E+08 | 1.53E+08 | 1.79E+08 | 7.40E+08 | 6.50E+08 | 7.67E+08 |
| 92 | Cluster of Tropomyosin 3 OS=Homo sapiens OX=960 | A0A087WWU8 [5] | TPM3 | 354 | 26 kDa | TRUE | 0.17 | 3.8 | [] | 8736300 | 3878200 | 1,000.00 | 2.62E+07 | 3668900 | 1.76E+07 |
| 89 | 60S ribosomal protein L7-like 1 OS=Homo sapiens O | Q6DKI1 (+1) | RPL7L1 | 54 | 30 kDa | | 0.24 | 3.8 | [] | 831180 | 657520 | 1,000.00 | 1,000.00 | 3177700 | 2460300 |
| 91 | Elongator complex protein 3 OS=Homo sapiens OX=9 | Q9H9T3 | ELP3 | 23 | 62 kDa | | 0.3 | 3.8 | [] | 1,000.00 | 2407700 | 1,000.00 | 1,000.00 | 3246800 | 5817100 |
| 93 | Activating signal cointegrator 1 complex subunit 2 OS | Q9H1I8 | ASCC2 | 19 | 86 kDa | | 0.49 | 3.8 | [] | 4915800 | 1,000.00 | 1,000.00 | 1.77E+07 | 772450 | 1,000.00 |
| 90 | Pyruvate kinase PKM OS=Homo sapiens OX=9606 G | P14618 | PKM | 536 | 58 kDa | | 0.52 | 3.8 | [] | 849960 | 1,000.00 | 1,000.00 | 3217400 | 1,000.00 | 1,000.00 |
| 96 | THUMP domain-containing protein 1 OS=Homo sapie | Q9NXG2 | THUMPD1 | 14 | 39 kDa | | 0.081 | 3.7 | [] | 9258600 | 1,000.00 | 1,000.00 | 7915500 | 1.22E+07 | 1.41E+07 |
| 94 | MSL complex subunit 1 OS=Homo sapiens OX=9606 | J3KSZ8 (+2) | MSL1 | 20 | 66 kDa | | 0.52 | 3.7 | [] | 1,000.00 | 1,000.00 | 2542200 | 9423500 | 1,000.00 | 1,000.00 |
| 95 | Dolichol-phosphate mannosyltransferase subunit 1 OS | H0Y368 (+2) | DPM1 | 107 | 33 kDa | | 0.52 | 3.7 | [] | 1,000.00 | 2433800 | 1,000.00 | 1,000.00 | 9016500 | 1,000.00 |
| 97 | Adenosine deaminase RNA specific OS=Homo sapiens | A0A7P0Z4F9 (+1) | ADAR | 235 | 102 kDa | | 0.25 | 3.6 | [] | 6235300 | 1704500 | 1,000.00 | 1,000.00 | 1.58E+07 | 1.29E+07 |
| 99 | Zinc finger protein 24 OS=Homo sapiens OX=9606 G | P17028 | ZNF24 | 22 | 42 kDa | | 0.37 | 3.6 | [] | 1,000.00 | 1,000.00 | 1467200 | 3851900 | 1375500 | 1,000.00 |
| 101 | Ubiquitin-40S ribosomal protein S27a OS=Homo sapi | P62979 (+1) | RPS27A | 478 | 18 kDa | | 0.082 | 3.5 | [] | 1,000.00 | 8264200 | 6328400 | 2.58E+07 | 1.14E+07 | 1.33E+07 |
| 100 | Signal recognition particle 14 kDa protein OS=Homo s | P37108 | SRP14 | 338 | 15 kDa | | 0.39 | 3.5 | [] | 2.03E+07 | 1.11E+07 | 1,000.00 | 1,000.00 | 8.90E+07 | 2.19E+07 |
| 102 | Putative ATP-dependent RNA helicase DHX57 OS=H | Q6P158 | DHX57 | 26 | 156 kDa | TRUE | 0.29 | 3.4 | [] | 1.34E+07 | 3.09E+07 | 1.58E+07 | 1.97E+07 | 1.44E+08 | 4.02E+07 |
| 103 | enoyl-CoA hydratase OS=Homo sapiens OX=9606 GN | H0YFD6 (+1) | HADHA | 154 | 86 kDa | | 0.54 | 3.4 | [] | 880790 | 1,000.00 | 1,000.00 | 1,000.00 | 2963900 | 1,000.00 |

| | | | | | | | | | | | | | | | |
|---|---|---|---|---|---|---|---|---|---|---|---|---|---|---|---|
| 104 | 39S ribosomal protein L38, mitochondrial OS=Homo s | Q96DV4 | MRPL38 | 34 | 45 kDa | | 0.051 | 3.3 | [] | 2763400 | 1.16E+07 | 5.63E+07 | 6.71E+07 | 6.68E+07 | 9.61E+07 |
| 105 | 40S ribosomal protein S18 OS=Homo sapiens OX=960 | P62269 | RPS18 | 399 | 18 kDa | | 0.065 | 3.2 | [] | 1.70E+07 | 1.84E+07 | 5.37E+07 | 4.97E+07 | 1.14E+08 | 1.25E+08 |
| 106 | Cluster of Heterogeneous nuclear ribonucleoprotein C | B4DY08 [10] | HNRNPC | 444 | 32 kDa | TRUE | 0.087 | 3.2 | [] | 2.62E+07 | 1,000.00 | 2.08E+07 | 3.35E+07 | 4.19E+07 | 7.73E+07 |
| 107 | Pre-mRNA processing factor 40 homolog A OS=Hom | A0A7N4I394 (+1) | PRPF40A | 195 | 112 kDa | | 0.28 | 3.2 | [] | 7694400 | 1.38E+07 | 7097800 | 1,000.00 | 3.50E+07 | 5.53E+07 |
| 109 | 40S ribosomal protein S25 OS=Homo sapiens OX=960 | P62851 | RPS25 | 356 | 14 kDa | | 0.033 | 3.1 | [] | 1.94E+08 | 2.62E+08 | 6.87E+07 | 6.86E+08 | 3.51E+08 | 5.84E+08 |
| 110 | ATP-dependent RNA helicase DDX55 OS=Homo sapi | Q8NHQ9 | DDX55 | 81 | 69 kDa | | 0.31 | 3 | [] | 7293500 | 1.04E+07 | 4847000 | 4.76E+07 | 1.16E+07 | 7856600 |
| 111 | ADP-ribosylation factor GTPase-activating protein 1 C | Q8N6T3 | ARFGAP1 | 112 | 45 kDa | | 0.56 | 3 | [] | 2535000 | 1,000.00 | 1,000.00 | 7520500 | 1,000.00 | 1,000.00 |
| 116 | Heterogeneous nuclear ribonucleoprotein U-like protei | Q9BUJ2 | HNRNPUL1 | 254 | 96 kDa | | 0.11 | 2.9 | [] | 1.13E+08 | 1.79E+08 | 2.00E+08 | 7.56E+08 | 3.27E+08 | 3.20E+08 |
| 112 | Myb-binding protein 1A OS=Homo sapiens OX=9606 | Q9BQG0 | MYBBP1A | 301 | 149 kDa | | 0.26 | 2.9 | [] | 709690 | 1.01E+07 | 1,000.00 | 3980700 | 9291700 | 1.85E+07 |
| 113 | Heterochromatin protein 1-binding protein 3 OS=Hom | Q5SSJ5 | HP1BP3 | 214 | 61 kDa | | 0.31 | 2.9 | [] | 4066600 | 1.77E+07 | 3208900 | 935510 | 2.65E+07 | 4.56E+07 |
| 115 | Cluster of U3 small nucleolar RNA-associated protein | Q9BVJ6 [2] | UTP14A | 117 | 88 kDa | TRUE | 0.36 | 2.9 | [] | 1995700 | 733940 | 7014400 | 2.07E+07 | 2809900 | 4650300 |
| 114 | Nucleolar protein 14 OS=Homo sapiens OX=9606 GN | P78316 | NOP14 | 46 | 98 kDa | | 0.38 | 2.9 | [] | 1,000.00 | 929560 | 930500 | 1,000.00 | 1431200 | 3984800 |
| 118 | Keratin, type II cytoskeletal 1 OS=Homo sapiens OX= | P04264 | KRT1 | 671 | 66 kDa | TRUE | 0.011 | 2.7 | [] | 1.96E+08 | 2.90E+08 | 4.58E+08 | 8.97E+08 | 1.01E+09 | 6.79E+08 |
| 121 | Cluster of Polyadenylate-binding protein OS=Homo sa | A0A7I2V4L7 [4] | PABPC4 | 376 | 67 kDa | TRUE | 0.034 | 2.7 | [] | 1.70E+08 | 8.31E+07 | 1.45E+08 | 4.84E+08 | 2.64E+08 | 3.21E+08 |
| 120 | NADH dehydrogenase [ubiquinone] 1 alpha subcompl | A0A087WXC5 (+17 | NDUFA10 | 67 | 42 kDa | | 0.21 | 2.7 | [] | 3874300 | 2134200 | 1,000.00 | 1648800 | 8447200 | 6261100 |
| 123 | Elongation factor 2 OS=Homo sapiens OX=9606 GN= | P13639 | EEF2 | 488 | 95 kDa | TRUE | 0.32 | 2.7 | [] | 3190700 | 1,000.00 | 1526600 | 5639600 | 1,000.00 | 6902900 |
| 122 | Sphingosine-1-phosphate lyase 1 OS=Homo sapiens O | O95470 | SGPL1 | 15 | 64 kDa | | 0.38 | 2.7 | [] | 1,000.00 | 1.26E+07 | 3813900 | 1,000.00 | 2.92E+07 | 1.45E+07 |
| 127 | Cluster of Nucleoporin 155 OS=Homo sapiens OX=96 | E9PF10 [3] | NUP155 | 175 | 148 kDa | TRUE | 0.0024 | 2.6 | [] | 2.20E+07 | 4.39E+07 | 2.64E+07 | 7.98E+07 | 7.66E+07 | 8.67E+07 |
| 126 | 60S ribosomal protein L36 OS=Homo sapiens OX=960 | Q9Y3U8 | RPL36 | 221 | 12 kDa | | 0.018 | 2.6 | [] | 8.09E+07 | 1.46E+08 | 1.72E+08 | 2.63E+08 | 4.33E+08 | 3.54E+08 |
| 130 | 60S ribosomal protein L27a OS=Homo sapiens OX=9 | P46776 | RPL27A | 430 | 17 kDa | | 0.1 | 2.6 | [] | 5.21E+08 | 1.64E+08 | 6.77E+08 | 1.77E+09 | 9.64E+08 | 7.89E+08 |
| 124 | Probable ATP-dependent RNA helicase DHX40 OS=H | Q8IX18 | DHX40 | 54 | 89 kDa | | 0.34 | 2.6 | [] | 1,000.00 | 6741400 | 2415900 | 1,000.00 | 9775800 | 1.44E+07 |
| 125 | Protein transport protein Sec24B OS=Homo sapiens O | O95487 | SEC24B | 139 | 137 kDa | | 0.41 | 2.6 | [] | 9554000 | 1,000.00 | 1076100 | 1.94E+07 | 8659200 | 1,000.00 |
| 128 | Probable 28S rRNA (cytosine(4447)-C(5))-methyltrans | P46087 | NOP2 | 276 | 89 kDa | | 0.45 | 2.6 | [] | 670940 | 8564700 | 6244200 | 3.24E+07 | 1,000.00 | 8047000 |
| 129 | Transcription factor Gibbin OS=Homo sapiens OX=96 | Q5TGY3 | AHDC1 | 7 | 168 kDa | | 0.6 | 2.6 | [] | 1,000.00 | 1,000.00 | 2001300 | 5179400 | 1,000.00 | 1,000.00 |
| 132 | Protein transport protein Sec24C OS=Homo sapiens O | P53992 | SEC24C | 122 | 118 kDa | TRUE | 0.094 | 2.5 | [] | 5677100 | 1.52E+07 | 2.47E+07 | 5.38E+07 | 3.94E+07 | 2.23E+07 |
| 131 | Twinkle mtDNA helicase OS=Homo sapiens OX=960 | Q96RR1 | TWNK | 11 | 77 kDa | | 0.6 | 2.5 | [] | 1,000.00 | 1,000.00 | 2158800 | 5474600 | 1,000.00 | 1,000.00 |
| 134 | Eukaryotic translation initiation factor 2 alpha kinase 2 | A0A7P0TBA9 (+1) | EIF2AK2 | 76 | 61 kDa | | 0.035 | 2.4 | [] | 2.78E+07 | 1.35E+07 | 8649100 | 3.07E+07 | 4.76E+07 | 4.35E+07 |
| 137 | 40S ribosomal protein SA OS=Homo sapiens OX=960 | A0A0C4DG17 (+2) | RPSA | 335 | 33 kDa | | 0.13 | 2.4 | [] | 1.02E+07 | 2.02E+07 | 4.94E+07 | 9.22E+07 | 5.28E+07 | 4.39E+07 |
| 133 | Cluster of 60S ribosomal protein L13a (Fragment) OS= | M0QYS1 [2] | RPL13A | 294 | 24 kDa | TRUE | 0.22 | 2.4 | [] | 3.33E+08 | 8.71E+08 | 1.17E+09 | 3.41E+09 | 1.02E+09 | 1.36E+09 |
| 136 | Caveolae-associated protein 1 OS=Homo sapiens OX= | Q6NZI2 | CAVIN1 | 27 | 43 kDa | | 0.35 | 2.4 | [] | 1.31E+07 | 2.15E+07 | 1.66E+07 | 8.46E+07 | 1.79E+07 | 1.90E+07 |
| 138 | Zinc finger protein 281 OS=Homo sapiens OX=9606 ( | Q9Y2X9 | ZNF281 | 107 | 97 kDa | | 0.35 | 2.4 | [] | 7.38E+07 | 3.31E+07 | 1.36E+08 | 3.88E+08 | 1.15E+08 | 7.01E+07 |
| 139 | Polyadenylate-binding protein-interacting protein 1 OS | Q9H074 | PAIP1 | 26 | 54 kDa | | 0.44 | 2.4 | [] | 2627800 | 1,000.00 | 1,000.00 | 3551700 | 1,000.00 | 2642300 |
| 135 | KH-type splicing regulatory protein OS=Homo sapiens | M0R0C6 (+2) | KHSRP | 403 | 77 kDa | TRUE | 0.52 | 2.4 | [] | 3787100 | 3.26E+07 | 2.24E+07 | 1.25E+08 | 1.03E+07 | 8053200 |

| | | | | | | | | | | | | | | | |
|---|---|---|---|---|---|---|---|---|---|---|---|---|---|---|---|
| 141 | 39S ribosomal protein L19, mitochondrial OS=Homo s | P49406 | MRPL19 | 59 | 34 kDa | | 0.14 | 2.3 | [] | 4753200 | 2513000 | 1,000.00 | 4579300 | 4416700 | 7549800 |
| 140 | Cluster of Peroxiredoxin-1 (Fragment) OS=Homo sapi | A0A0A0MSI0 [4] | PRDX1 | 549 | 19 kDa | TRUE | 0.23 | 2.3 | [] | 4.93E+07 | 1.59E+07 | 1202100 | 8.06E+07 | 2.97E+07 | 4.49E+07 |
| 143 | Cluster of KH domain-containing, RNA-binding, signa | Q07666 [2] | KHDRBS1 | 348 | 48 kDa | TRUE | 0.33 | 2.3 | [] | 7.83E+08 | 6.31E+08 | 1.67E+09 | 4.56E+09 | 1.05E+09 | 1.34E+09 |
| 142 | Polyglutamine-binding protein 1 OS=Homo sapiens OX | O60828 | PQBP1 | 67 | 30 kDa | | 0.43 | 2.3 | [] | 5068400 | 2627500 | 6510400 | 2.41E+07 | 2504800 | 5604500 |
| 152 | Ribosomal protein L32 OS=Homo sapiens OX=9606 C | F8W727 (+1) | RPL32 | 184 | 18 kDa | | 0.06 | 2.2 | [] | 1.04E+08 | 1.91E+08 | 9.29E+07 | 2.11E+08 | 3.71E+08 | 2.52E+08 |
| 145 | 28S ribosomal protein S9, mitochondrial OS=Homo sa | P82933 | MRPS9 | 72 | 46 kDa | | 0.1 | 2.2 | [] | 2.86E+07 | 1.27E+08 | 1.08E+08 | 1.14E+08 | 2.48E+08 | 2.27E+08 |
| 144 | Src substrate cortactin OS=Homo sapiens OX=9606 G | Q14247 | CTTN | 318 | 62 kDa | | 0.15 | 2.2 | [] | 6.04E+07 | 6.80E+07 | 8.44E+07 | 2.52E+08 | 9.11E+07 | 1.33E+08 |
| 150 | Constitutive coactivator of PPAR-gamma-like protein | Q9NZB2 | FAM120A | 159 | 122 kDa | | 0.15 | 2.2 | [] | 1.18E+08 | 1.01E+08 | 9.10E+07 | 3.53E+08 | 1.95E+08 | 1.22E+08 |
| 149 | Tricarboxylate transport protein, mitochondrial OS=Ho | P53007 | SLC25A1 | 175 | 34 kDa | | 0.27 | 2.2 | [] | 1.13E+07 | 1.16E+07 | 1.72E+07 | 5126000 | 3.79E+07 | 4.45E+07 |
| 146 | Cluster of Pre-mRNA 3'-end-processing factor FIP1 O | Q6UN15 [2] | FIP1L1 | 219 | 67 kDa | TRUE | 0.31 | 2.2 | [] | 1,000.00 | 1,000.00 | 1.59E+07 | 1.43E+07 | 9572700 | 1.12E+07 |
| 151 | 39S ribosomal protein L41, mitochondrial OS=Homo s | Q8IXM3 | MRPL41 | 67 | 15 kDa | | 0.33 | 2.2 | [] | 3.64E+07 | 1103700 | 1.31E+07 | 8330700 | 5.28E+07 | 4.79E+07 |
| 147 | Cluster of Metadherin OS=Homo sapiens OX=9606 G | E5RJU9 [3] | MTDH | 179 | 58 kDa | TRUE | 0.55 | 2.2 | [] | 2.36E+07 | 1.62E+07 | 4.25E+07 | 1.61E+08 | 1.09E+07 | 9710200 |
| 148 | Eukaryotic translation initiation factor 3 subunit G OS= | O75821 | EIF3G | 289 | 36 kDa | | 0.65 | 2.2 | [] | 2.17E+07 | 1,000.00 | 1,000.00 | 4.75E+07 | 1,000.00 | 1,000.00 |
| 156 | 39S ribosomal protein L24, mitochondrial OS=Homo s | Q96A35 | MRPL24 | 42 | 25 kDa | | 0.026 | 2.1 | [] | 1.92E+07 | 6.59E+07 | 4.97E+07 | 9.57E+07 | 9.92E+07 | 8.71E+07 |
| 153 | Insulin-like growth factor 2 mRNA-binding protein 1 C | Q9NZI8 | IGF2BP1 | 343 | 63 kDa | TRUE | 0.036 | 2.1 | [] | 5.37E+07 | 1.33E+08 | 1.13E+08 | 2.11E+08 | 1.68E+08 | 2.64E+08 |
| 162 | Double-stranded RNA-binding protein Staufen homolo | O95793 | STAU1 | 228 | 63 kDa | TRUE | 0.21 | 2.1 | [] | 2684800 | 2.63E+07 | 7729700 | 3.42E+07 | 1.91E+07 | 2.21E+07 |
| 155 | Fatty acid synthase OS=Homo sapiens OX=9606 GN= | A0A0U1RQF0 (+1) | FASN | 436 | 273 kDa | | 0.33 | 2.1 | [] | 3621000 | 5697800 | 3262200 | 1.70E+07 | 3344700 | 6319200 |
| 160 | Cluster of F-actin-capping protein subunit alpha-1 OS= | P52907 [5] | CAPZA1 | 318 | 33 kDa | TRUE | 0.38 | 2.1 | [] | 5830100 | 6328900 | 5824800 | 1,000.00 | 1.56E+07 | 2.16E+07 |
| 154 | Angiogenic factor with G patch and FHA domains 1 O | Q8N302 | AGGF1 | 2 | 81 kDa | | 0.4 | 2.1 | [] | 1,000.00 | 3412300 | 6938400 | 1,000.00 | 1.06E+07 | 1.15E+07 |
| 158 | Transcriptional activator protein Pur-beta OS=Homo s | Q96QR8 | PURB | 57 | 33 kDa | TRUE | 0.49 | 2.1 | [] | 1,000.00 | 1,000.00 | 2289400 | 2505000 | 2271700 | 1,000.00 |
| 159 | Translocation associated membrane protein 1 OS=Hon | G3XAN4 (+1) | TRAM1 | 4 | 33 kDa | | 0.53 | 2.1 | [] | 1.49E+07 | 1,000.00 | 6407100 | 3.49E+07 | 9435900 | 1,000.00 |
| 157 | Cluster of SWI/SNF-related matrix-associated actin-de | O60264 [4] | SMARCA5 | 210 | 122 kDa | TRUE | 0.64 | 2.1 | [] | 1,000.00 | 5048300 | 2316900 | 1.54E+07 | 1,000.00 | 1,000.00 |
| 169 | Cluster of Myosin light chain 6 OS=Homo sapiens OX | B7Z6Z4 [8] | MYL6 | 411 | 27 kDa | TRUE | 0.028 | 2 | [] | 1.27E+08 | 1.94E+08 | 2.58E+08 | 4.49E+08 | 3.77E+08 | 3.11E+08 |
| 170 | Cluster of Myosin-14 OS=Homo sapiens OX=9606 GN | Q7Z406 [2] | MYH14 | 318 | 228 kDa | TRUE | 0.042 | 2 | [] | 6.01E+08 | 8.65E+08 | 5.41E+08 | 9.32E+08 | 1.43E+09 | 1.56E+09 |
| 163 | 40S ribosomal protein S27 OS=Homo sapiens OX=96 | P42677 | RPS27 | 411 | 9 kDa | TRUE | 0.14 | 2 | [] | 2.27E+07 | 1.18E+08 | 1.77E+08 | 1.81E+08 | 1.71E+08 | 2.96E+08 |
| 167 | 40S ribosomal protein S15a OS=Homo sapiens OX=9 | P62244 | RPS15A | 401 | 15 kDa | | 0.18 | 2 | [] | 1.12E+07 | 6.93E+07 | 1.12E+08 | 1.54E+08 | 1.50E+08 | 7.71E+07 |
| 165 | 60S ribosomal protein L6 OS=Homo sapiens OX=960 | Q02878 | RPL6 | 432 | 33 kDa | | 0.2 | 2 | [] | 1.24E+09 | 8.01E+08 | 1.17E+09 | 3.59E+09 | 1.59E+09 | 1.33E+09 |
| 164 | Mitochondrial ribosomal protein L39 (Fragment) OS=I | C9JG87 (+1) | MRPL39 | 77 | 34 kDa | | 0.53 | 2 | [] | 1,000.00 | 7169600 | 1,000.00 | 1785500 | 2526500 | 1.02E+07 |
| 168 | Carnitine O-palmitoyltransferase 1, liver isoform OS=I | P50416 | CPT1A | 9 | 88 kDa | | 0.64 | 2 | [] | 1,000.00 | 4617200 | 1.44E+07 | 1,000.00 | 3.54E+07 | 2020200 |
| 166 | Melanoma-associated antigen 4 OS=Homo sapiens OX | P43358 | MAGEA4 | 10 | 35 kDa | | 0.65 | 2 | [] | 1226700 | 966650 | 1,000.00 | 1,000.00 | 4368200 | 1,000.00 |
| 186 | 40S ribosomal protein S16 OS=Homo sapiens OX=96 | P62249 | RPS16 | 411 | 16 kDa | | 0.004 | 1.9 | [] | 6.60E+08 | 7.94E+08 | 4.93E+08 | 1.15E+09 | 1.21E+09 | 1.28E+09 |
| 172 | 60S ribosomal protein L21 OS=Homo sapiens OX=96 | P46778 | RPL21 | 329 | 19 kDa | | 0.0078 | 1.9 | [] | 5.97E+08 | 6.85E+08 | 5.16E+08 | 1.21E+09 | 9.66E+08 | 1.31E+09 |
| 171 | Cluster of Elongation factor 1-delta OS=Homo sapiens | A0A087X1X7 [3] | EEF1D | 353 | 69 kDa | TRUE | 0.026 | 1.9 | [] | 4.05E+08 | 4.85E+08 | 7.98E+08 | 9.04E+08 | 1.15E+09 | 1.23E+09 |

| | | | | | | | | | | | | | | | |
|---|---|---|---|---|---|---|---|---|---|---|---|---|---|---|---|
| 188 | Methylcrotonoyl-CoA carboxylase subunit alpha, mitoc | Q96RQ3 | MCCC1 | 280 | 80 kDa | | 0.043 | 1.9 | [] | 1.85E+08 | 2.74E+08 | 2.29E+08 | 3.05E+08 | 5.11E+08 | 4.67E+08 |
| 184 | G protein nucleolar 3 like OS=Homo sapiens OX=960 | A0A6I8PL80 (+2) | GNL3L | 46 | 61 kDa | | 0.1 | 1.9 | [] | 6896700 | 1.13E+07 | 1.30E+07 | 2.70E+07 | 1.78E+07 | 1.36E+07 |
| 176 | 60S ribosomal protein L10a OS=Homo sapiens OX=9 | P62906 | RPL10A | 280 | 25 kDa | | 0.12 | 1.9 | [] | 3.50E+08 | 2.24E+08 | 1.08E+08 | 5.99E+08 | 3.41E+08 | 3.68E+08 |
| 182 | Elongation factor 1-gamma OS=Homo sapiens OX=96 | P26641 | EEF1G | 461 | 50 kDa | | 0.15 | 1.9 | [] | 9.44E+08 | 1.01E+09 | 1.04E+09 | 9.03E+08 | 2.19E+09 | 2.55E+09 |
| 180 | 60S ribosomal protein L23 OS=Homo sapiens OX=96 | P62829 | RPL23 | 507 | 15 kDa | | 0.16 | 1.9 | [] | 6.71E+08 | 7.27E+08 | 6.18E+08 | 1.95E+09 | 8.01E+08 | 1.08E+09 |
| 178 | 60S ribosomal protein L35a OS=Homo sapiens OX=9 | P18077 | RPL35A | 220 | 13 kDa | | 0.18 | 1.9 | [] | 7.01E+07 | 1.52E+08 | 1.35E+08 | 3.28E+08 | 1.17E+08 | 2.36E+08 |
| 175 | Replication initiator 1 (Fragment) OS=Homo sapiens C | C9JYJ8 (+2) | REPIN1 | 2 | 44 kDa | | 0.23 | 1.9 | [] | 5794000 | 8174600 | 1.55E+07 | 8690200 | 1.96E+07 | 2.82E+07 |
| 190 | 60S ribosomal protein L12 OS=Homo sapiens OX=96 | P30050 | RPL12 | 397 | 18 kDa | | 0.23 | 1.9 | [] | 1.08E+08 | 1.22E+08 | 1.63E+08 | 3.94E+08 | 1.42E+08 | 1.91E+08 |
| 189 | 40S ribosomal protein S21 OS=Homo sapiens OX=96 | P63220 (+1) | RPS21 | 252 | 9 kDa | | 0.34 | 1.9 | [] | 4198400 | 1,000.00 | 6970200 | 1.07E+07 | 3071700 | 7016400 |
| 174 | Signal recognition particle 9 OS=Homo sapiens OX=9 | E9PE20 (+3) | SRP9 | 208 | 8 kDa | | 0.43 | 1.9 | [] | 2690900 | 1.20E+07 | 4101500 | 1,000.00 | 1.77E+07 | 1.86E+07 |
| 181 | Phosphatidylserine synthase 1 OS=Homo sapiens OX= | P48651 | PTDSS1 | 14 | 56 kDa | | 0.53 | 1.9 | [] | 1.78E+07 | 2.35E+07 | 1,000.00 | 1,000.00 | 5.71E+07 | 2.12E+07 |
| 183 | Growth arrest and DNA damage-inducible proteins-int | Q8TAE8 | GADD45GIP1 | 57 | 25 kDa | | 0.59 | 1.9 | [] | 2.65E+08 | 5.60E+07 | 3.02E+07 | 5.33E+08 | 5.00E+07 | 7.58E+07 |
| 185 | Coatomer subunit alpha OS=Homo sapiens OX=9606 | A0A3B3IT15 (+1) | COPA | 202 | 136 kDa | | 0.6 | 1.9 | [] | 9.47E+07 | 1.44E+08 | 9.25E+07 | 1.81E+07 | 5.38E+08 | 6.25E+07 |
| 179 | PDZ and LIM domain protein 5 OS=Homo sapiens O | Q96HC4 | PDLIM5 | 116 | 64 kDa | | 0.67 | 1.9 | [] | 1.05E+07 | 679720 | 1,000.00 | 2.02E+07 | 1,000.00 | 1173800 |
| 173 | Protein Red OS=Homo sapiens OX=9606 GN=IK PE= | Q13123 | IK | 134 | 66 kDa | | 0.69 | 1.9 | [] | 1,000.00 | 460740 | 1,000.00 | 1,000.00 | 1,000.00 | 896520 |
| 187 | Basement membrane-specific heparan sulfate proteogly | P98160 | HSPG2 | 8 | 469 kDa | | 0.7 | 1.9 | [] | 1,000.00 | 1,000.00 | 3593500 | 6699900 | 1,000.00 | 1,000.00 |
| 191 | 60S ribosomal protein L27 OS=Homo sapiens OX=96 | P61353 | RPL27 | 324 | 16 kDa | | 0.037 | 1.8 | [] | 3.05E+08 | 2.72E+08 | 2.04E+08 | 5.97E+08 | 3.90E+08 | 4.36E+08 |
| 192 | Arginine/proline rich coiled-coil 1, isoform CRA_b OS | D3DPS3 (+1) | MAP7D1 | 60 | 93 kDa | | 0.044 | 1.8 | [] | 4.61E+07 | 2.18E+07 | 5.02E+07 | 6.35E+07 | 8.49E+07 | 6.66E+07 |
| 193 | 60S ribosomal protein L35 OS=Homo sapiens OX=96 | P42766 | RPL35 | 302 | 15 kDa | | 0.11 | 1.8 | [] | 1.55E+08 | 2.83E+08 | 2.33E+08 | 2.58E+08 | 5.34E+08 | 4.28E+08 |
| 197 | 40S ribosomal protein S27 OS=Homo sapiens OX=96 | C9JLI6 (+2) | RPS27L | 372 | 11 kDa | TRUE | 0.17 | 1.8 | [] | 2.27E+07 | 9.39E+07 | 1.44E+08 | 1.81E+08 | 1.13E+08 | 1.76E+08 |
| 194 | Elongation factor 1-beta OS=Homo sapiens OX=9606 | P24534 | EEF1B2 | 379 | 25 kDa | TRUE | 0.42 | 1.8 | [] | 7.77E+07 | 1.44E+08 | 1.35E+08 | 8732900 | 3.50E+08 | 2.91E+08 |
| 199 | Calcium load-activated calcium channel OS=Homo sap | J3QQY2 (+1) | TMCO1 | 30 | 11 kDa | | 0.42 | 1.8 | [] | 5938800 | 376590 | 8029400 | 1.43E+07 | 9269000 | 2102600 |
| 196 | Propionyl-CoA carboxylase subunit beta OS=Homo sa | C9JQS9 (+4) | PCCB | 263 | 61 kDa | | 0.45 | 1.8 | [] | 644530 | 1.75E+07 | 3.96E+07 | 1.13E+07 | 3.19E+07 | 6.11E+07 |
| 195 | G patch domain-containing protein 4 OS=Homo sapie | Q5T3I0 | GPATCH4 | 114 | 50 kDa | | 0.54 | 1.8 | [] | 1,000.00 | 4145900 | 9874600 | 1.82E+07 | 5908000 | 1394300 |
| 201 | Squalene synthase OS=Homo sapiens OX=9606 GN=I | P37268 | FDFT1 | 35 | 48 kDa | | 0.58 | 1.8 | [] | 1,000.00 | 6988000 | 694150 | 1,000.00 | 7291100 | 6176900 |
| 200 | Target of EGR1 protein 1 OS=Homo sapiens OX=960 | Q96GM8 | TOE1 | 31 | 57 kDa | | 0.68 | 1.8 | [] | 5001100 | 2877300 | 7524400 | 2.67E+07 | 1,000.00 | 590370 |
| 198 | Cluster of Stem-loop binding protein OS=Homo sapie | F8W8D3 [4] | SLBP | 15 | 32 kDa | TRUE | 0.72 | 1.8 | [] | 1,000.00 | 346730 | 1,000.00 | 1,000.00 | 1,000.00 | 626990 |
| 210 | Cluster of Myosin-9 OS=Homo sapiens OX=9606 GN | P35579 [3] | MYH9 | 448 | 227 kDa | TRUE | 0.12 | 1.7 | [] | 8.48E+09 | 8.46E+09 | 6.75E+09 | 8.08E+09 | 1.68E+10 | 1.48E+10 |
| 206 | 39S ribosomal protein L9, mitochondrial OS=Homo sa | Q9BYD2 | MRPL9 | 44 | 30 kDa | | 0.16 | 1.7 | [] | 3.00E+07 | 4.01E+07 | 4.42E+07 | 7.66E+07 | 8.39E+07 | 3.50E+07 |
| 204 | Cluster of Acetyl-CoA carboxylase 1 OS=Homo sapie | Q13085 [2] | ACACA | 296 | 266 kDa | TRUE | 0.22 | 1.7 | [] | 9.42E+08 | 8.33E+08 | 6.03E+08 | 6.44E+08 | 1.50E+09 | 1.92E+09 |
| 207 | Cluster of 40S ribosomal protein S2 OS=Homo sapien | P15880 [2] | RPS2 | 447 | 31 kDa | TRUE | 0.25 | 1.7 | [] | 3.09E+08 | 3.40E+08 | 5.16E+08 | 1.02E+09 | 4.63E+08 | 4.71E+08 |
| 209 | 60S ribosomal protein L38 OS=Homo sapiens OX=96 | P63173 | RPL38 | 300 | 8 kDa | | 0.3 | 1.7 | [] | 3.57E+08 | 5.38E+07 | 1.04E+08 | 2.62E+08 | 3.33E+08 | 2.68E+08 |
| 205 | 28S ribosomal protein S21, mitochondrial OS=Homo s | P82921 | MRPS21 | 52 | 11 kDa | | 0.31 | 1.7 | [] | 3.32E+07 | 1.33E+08 | 2.26E+07 | 7.77E+07 | 1.27E+08 | 1.18E+08 |

| | | | | | | | | | | | | | | | |
|---|---|---|---|---|---|---|---|---|---|---|---|---|---|---|---|
| 208 | 60S ribosomal protein L7a OS=Homo sapiens OX=96( | P62424 | RPL7A | 398 | 30 kDa | | 0.33 | 1.7 | [] | 1.51E+09 | 1.21E+09 | 1.41E+09 | 3.96E+09 | 1.33E+09 | 1.64E+09 |
| 203 | Ribosomal protein S11 OS=Homo sapiens OX=9606 ( | M0QZC5 (+1) | RPS11 | 340 | 14 kDa | | 0.5 | 1.7 | [] | 3.64E+08 | 1.00E+08 | 3.46E+08 | 9.76E+08 | 2.14E+08 | 2.11E+08 |
| 202 | Casein kinase 1 alpha 1 OS=Homo sapiens OX=9606 ( | A0A590UJ43 (+3) | CSNK1A1 | 70 | 49 kDa | | 0.62 | 1.7 | [] | 1,000.00 | 3190600 | 1,000.00 | 1,000.00 | 3452200 | 2111500 |
| 211 | Small subunit processome component 20 homolog OS= | O75691 | UTP20 | 43 | 318 kDa | | 0.75 | 1.7 | [] | 1,000.00 | 1,000.00 | 2806600 | 4653300 | 1,000.00 | 1,000.00 |
| 232 | Cluster of Polyadenylate-binding protein OS=Homo sa | A0A7I2V4N4 [4] | PABPC1 | 408 | 64 kDa | TRUE | 0.036 | 1.6 | [] | 4.08E+08 | 2.41E+08 | 3.74E+08 | 5.85E+08 | 5.43E+08 | 4.68E+08 |
| 213 | E3 ubiquitin-protein ligase TRIM56 OS=Homo sapiens | Q9BRZ2 | TRIM56 | 26 | 81 kDa | | 0.069 | 1.6 | [] | 6264400 | 2817700 | 7466500 | 9701500 | 8861200 | 8598500 |
| 231 | 39S ribosomal protein L2, mitochondrial OS=Homo sa | Q5T653 | MRPL2 | 43 | 33 kDa | | 0.084 | 1.6 | [] | 8.07E+07 | 3.65E+07 | 6.01E+07 | 8.49E+07 | 1.06E+08 | 8.59E+07 |
| 216 | Cluster of Heterogeneous nuclear ribonucleoprotein H | G8JLB6 [3] | HNRNPH1 | 558 | 51 kDa | TRUE | 0.088 | 1.6 | [] | 9.40E+08 | 6.56E+08 | 5.22E+08 | 1.43E+09 | 1.01E+09 | 9.76E+08 |
| 234 | Cluster of 60S ribosomal protein L13 OS=Homo sapie | P26373 [2] | RPL13 | 474 | 24 kDa | TRUE | 0.1 | 1.6 | [] | 1.19E+09 | 1.16E+09 | 1.49E+09 | 2.57E+09 | 1.47E+09 | 1.93E+09 |
| 223 | 40S ribosomal protein S8 OS=Homo sapiens OX=960( | P62241 (+1) | RPS8 | 498 | 24 kDa | | 0.11 | 1.6 | [] | 1.81E+09 | 1.09E+09 | 1.75E+09 | 3.17E+09 | 1.89E+09 | 2.34E+09 |
| 233 | 60S ribosomal protein L24 OS=Homo sapiens OX=96( | P83731 | RPL24 | 447 | 18 kDa | | 0.16 | 1.6 | [] | 6.67E+08 | 8.56E+08 | 1.23E+09 | 1.92E+09 | 1.21E+09 | 1.16E+09 |
| 212 | Ras GTPase-activating-like protein IQGAP1 OS=Hom | P46940 | IQGAP1 | 197 | 189 kDa | TRUE | 0.25 | 1.6 | [] | 1.44E+07 | 7544600 | 6109000 | 8124400 | 1.83E+07 | 1.98E+07 |
| 219 | 60S ribosomal protein L11 OS=Homo sapiens OX=96( | P62913 (+1) | RPL11 | 495 | 20 kDa | | 0.29 | 1.6 | [] | 3.76E+08 | 8.86E+07 | 6.63E+08 | 7.12E+08 | 4.31E+08 | 6.64E+08 |
| 217 | Cluster of La-related protein 1 OS=Homo sapiens OX= | Q6PKG0 [2] | LARP1 | 251 | 124 kDa | TRUE | 0.39 | 1.6 | [] | 3.52E+07 | 3.44E+07 | 9.10E+07 | 1.39E+08 | 4.48E+07 | 7.44E+07 |
| 226 | Family with sequence similarity 98 member A OS=Hor | E9PH82 (+1) | FAM98A | 112 | 34 kDa | | 0.41 | 1.6 | [] | 7502600 | 1.69E+07 | 1.30E+07 | 6031600 | 3.15E+07 | 2.17E+07 |
| 224 | Cluster of 39S ribosomal protein S18a, mitochondrial ( | Q9NVS2 [2] | MRPS18A | 41 | 22 kDa | TRUE | 0.42 | 1.6 | [] | 2968200 | 1.04E+07 | 1.75E+07 | 2.67E+07 | 9910700 | 1.24E+07 |
| 229 | Epiplakin 1 OS=Homo sapiens OX=9606 GN=EPPK1 | A0A075B730 (+1) | EPPK1 | 301 | 553 kDa | TRUE | 0.42 | 1.6 | [] | 1,000.00 | 8.87E+07 | 5.64E+07 | 4.76E+07 | 7.55E+07 | 1.04E+08 |
| 218 | Zinc finger CCCH-type containing 4 (Fragment) OS=F | M0QY97 (+1) | ZC3H4 | 134 | 96 kDa | | 0.49 | 1.6 | [] | 1.58E+08 | 7.07E+07 | 2.41E+08 | 1.65E+08 | 1.08E+08 | 4.80E+08 |
| 214 | 28S ribosomal protein S11, mitochondrial OS=Homo s | P82912 | MRPS11 | 60 | 21 kDa | | 0.5 | 1.6 | [] | 6.44E+07 | 1.48E+08 | 7.48E+07 | 3.11E+08 | 1.05E+08 | 5.40E+07 |
| 227 | Endophilin-A2 OS=Homo sapiens OX=9606 GN=SH3 | Q99961 | SH3GL1 | 109 | 41 kDa | | 0.51 | 1.6 | [] | 5819100 | 4654800 | 5161100 | 1,000.00 | 1.28E+07 | 1.19E+07 |
| 228 | Cluster of Protein transport protein Sec16A OS=Homo | O15027 [2] | SEC16A | 179 | 252 kDa | TRUE | 0.58 | 1.6 | [] | 6.27E+07 | 4304100 | 2.56E+07 | 9.78E+07 | 2.28E+07 | 2.54E+07 |
| 230 | Zinc finger protein RFP OS=Homo sapiens OX=9606 ( | P14373 | TRIM27 | 9 | 58 kDa | | 0.59 | 1.6 | [] | 1,000.00 | 3217700 | 3119800 | 1,000.00 | 5913500 | 3989900 |
| 222 | 40S ribosomal protein S12 OS=Homo sapiens OX=96( | P25398 | RPS12 | 382 | 15 kDa | | 0.63 | 1.6 | [] | 7666200 | 1,000.00 | 1,000.00 | 1225700 | 5073200 | 5913600 |
| 235 | BUB3 mitotic checkpoint protein (Fragment) OS=Hon | J3QT28 (+1) | BUB3 | 298 | 32 kDa | | 0.63 | 1.6 | [] | 5283700 | 1.23E+07 | 1,000.00 | 1.88E+07 | 1217800 | 7267700 |
| 221 | RNA binding motif protein X-linked (Fragment) OS=F | H0Y6E7 (+2) | RBMX | 377 | 32 kDa | | 0.7 | 1.6 | [] | 1,000.00 | 1,000.00 | 6047400 | 1,000.00 | 2538300 | 7132600 |
| 215 | Cluster of Sodium/potassium-transporting ATPase sub | B1AKY9 [3] | ATP1A2 | 275 | 111 kDa | TRUE | 0.73 | 1.6 | [] | 1,000.00 | 1,000.00 | 1.18E+07 | 1.76E+07 | 1,000.00 | 1721100 |
| 220 | PHD finger protein 6 OS=Homo sapiens OX=9606 GN | A0A0D9SGE8 (+1) | PHF6 | 156 | 41 kDa | | 0.74 | 1.6 | [] | 1,000.00 | 1762100 | 2.12E+07 | 3.37E+07 | 1,000.00 | 2976400 |
| 225 | Heterogeneous nuclear ribonucleoprotein L-like OS=H | Q8WVV9 | HNRNPLL | 80 | 60 kDa | TRUE | 0.75 | 1.6 | [] | 5744100 | 1673300 | 1,000.00 | 1,000.00 | 1.18E+07 | 1,000.00 |
| 239 | Ribosomal protein L18 (Fragment) OS=Homo sapiens | J3QQ67 (+1) | RPL18 | 399 | 22 kDa | | 0.031 | 1.5 | [] | 8.23E+08 | 1.11E+09 | 1.15E+09 | 1.41E+09 | 1.44E+09 | 1.79E+09 |
| 249 | Cluster of 60S ribosomal protein L4 OS=Homo sapien | P36578 [2] | RPL4 | 444 | 48 kDa | TRUE | 0.04 | 1.5 | [] | 1.81E+09 | 1.18E+09 | 1.26E+09 | 2.21E+09 | 1.91E+09 | 2.06E+09 |
| 236 | 40S ribosomal protein S3a OS=Homo sapiens OX=96( | P61247 | RPS3A | 434 | 30 kDa | | 0.071 | 1.5 | [] | 2.62E+08 | 4.79E+08 | 5.80E+08 | 6.45E+08 | 7.34E+08 | 6.59E+08 |
| 237 | 39S ribosomal protein L15, mitochondrial OS=Homo s | Q9P015 | MRPL15 | 56 | 33 kDa | | 0.17 | 1.5 | [] | 1.97E+08 | 1.62E+08 | 2.07E+08 | 2.34E+08 | 2.27E+08 | 4.01E+08 |
| 244 | Cluster of 40S ribosomal protein S26 OS=Homo sapie | P62854 [2] | RPS26 | 375 | 13 kDa | TRUE | 0.22 | 1.5 | [] | 3.38E+08 | 3.34E+08 | 2.45E+08 | 6.56E+08 | 3.68E+08 | 3.48E+08 |

| | | | | | | | | | | | | | | | |
|---|---|---|---|---|---|---|---|---|---|---|---|---|---|---|---|
| 248 | Cluster of RNA helicase OS=Homo sapiens OX=9606 | A0A0D9SF53 [14] | DDX3X | 498 | 81 kDa | TRUE | 0.3 | 1.5 | [] | 9.44E+08 | 9.27E+08 | 4.66E+08 | 6.41E+08 | 1.48E+09 | 1.29E+09 |
| 250 | 60S ribosomal protein L17 OS=Homo sapiens OX=96( | P18621 | RPL17 | 424 | 21 kDa | | 0.39 | 1.5 | [] | 5.99E+08 | 3.48E+08 | 4.94E+08 | 2.76E+08 | 8.67E+08 | 9.49E+08 |
| 240 | Cytoskeleton-associated protein 4 OS=Homo sapiens ( | Q07065 | CKAP4 | 168 | 66 kDa | | 0.4 | 1.5 | [] | 1.92E+07 | 2.71E+07 | 1.00E+08 | 7.84E+07 | 6.34E+07 | 7.84E+07 |
| 247 | Cleavage and polyadenylation specificity factor subunit | F8WJN3 (+1) | CPSF6 | 299 | 52 kDa | | 0.47 | 1.5 | [] | 2.69E+07 | 1.02E+08 | 1.04E+08 | 9.22E+07 | 6.25E+07 | 1.87E+08 |
| 238 | Dolichyl-diphosphooligosaccharide--protein glycosyltra | P04843 | RPN1 | 246 | 69 kDa | | 0.49 | 1.5 | [] | 1.13E+07 | 1.09E+07 | 890590 | 1.90E+07 | 5381000 | 1.07E+07 |
| 246 | Catenin delta 1 OS=Homo sapiens OX=9606 GN=CT! | C9JZR2 (+2) | CTNND1 | 99 | 105 kDa | | 0.53 | 1.5 | [] | 4990300 | 4708900 | 3348900 | 1.22E+07 | 3714900 | 3279300 |
| 245 | Activating signal cointegrator 1 OS=Homo sapiens OX | Q15650 | TRIP4 | 7 | 66 kDa | | 0.6 | 1.5 | [] | 7403300 | 3219100 | 8230400 | 1.92E+07 | 7445200 | 1528600 |
| 241 | Clathrin heavy chain OS=Homo sapiens OX=9606 GN | A0A087WVQ6 (+1) | CLTC | 406 | 192 kDa | | 0.61 | 1.5 | [] | 9540200 | 1631600 | 1,000.00 | 5240900 | 2953900 | 8618600 |
| 242 | Protein SPT2 homolog OS=Homo sapiens OX=9606 ( | Q68D10 | SPTY2D1 | 6 | 76 kDa | | 0.65 | 1.5 | [] | 1,000.00 | 1523600 | 7534200 | 3896200 | 8289900 | 1426300 |
| 253 | 60S ribosomal protein L7 OS=Homo sapiens OX=960 | P18124 | RPL7 | 335 | 29 kDa | | 0.015 | 1.4 | [] | 1.47E+09 | 1.33E+09 | 1.60E+09 | 2.37E+09 | 1.91E+09 | 2.03E+09 |
| 262 | 60S ribosomal protein L18a OS=Homo sapiens OX=9( | Q02543 | RPL18A | 300 | 21 kDa | | 0.048 | 1.4 | [] | 1.49E+08 | 1.89E+08 | 1.28E+08 | 1.99E+08 | 2.10E+08 | 2.38E+08 |
| 251 | Cluster of E3 ubiquitin-protein ligase OS=Homo sapie | A0A6Q8PHK0 [2] | TRIP12 | 116 | 228 kDa | TRUE | 0.19 | 1.4 | [] | 1.75E+07 | 2.04E+07 | 2.66E+07 | 4.19E+07 | 2.38E+07 | 2.77E+07 |
| 252 | 40S ribosomal protein S9 OS=Homo sapiens OX=960 | P46781 | RPS9 | 388 | 23 kDa | | 0.21 | 1.4 | [] | 8.15E+08 | 1.71E+09 | 8.24E+08 | 1.35E+09 | 1.68E+09 | 1.78E+09 |
| 259 | 39S ribosomal protein L21, mitochondrial OS=Homo s | Q7Z2W9 | MRPL21 | 45 | 23 kDa | | 0.21 | 1.4 | [] | 6.87E+07 | 6.66E+07 | 1.14E+08 | 1.06E+08 | 1.49E+08 | 9.49E+07 |
| 261 | Cluster of RNA-binding protein 39 OS=Homo sapiens | Q14498 [2] | RBM39 | 366 | 59 kDa | TRUE | 0.22 | 1.4 | [] | 2.93E+07 | 4.02E+07 | 7.15E+07 | 6.73E+07 | 6.33E+07 | 6.52E+07 |
| 265 | Protein THEM6 OS=Homo sapiens OX=9606 GN=TH | Q8WUY1 | THEM6 | 4 | 24 kDa | | 0.4 | 1.4 | [] | 1505600 | 383460 | 1960700 | 1978600 | 1895900 | 1396400 |
| 269 | Cluster of ATPase family AAA domain-containing pro | Q9NVI7 [3] | ATAD3A | 229 | 71 kDa | TRUE | 0.4 | 1.4 | [] | 1.55E+08 | 1.93E+08 | 3.69E+08 | 2.10E+08 | 3.43E+08 | 4.17E+08 |
| 272 | Sorting nexin 9 OS=Homo sapiens OX=9606 GN=SN | A0A7P0T8C7 (+7) | SNX9 | 73 | 71 kDa | | 0.45 | 1.4 | [] | 2.49E+07 | 1.55E+07 | 2.02E+07 | 4.26E+07 | 1.53E+07 | 2.38E+07 |
| 255 | 28S ribosomal protein S25, mitochondrial OS=Homo s | P82663 | MRPS25 | 21 | 20 kDa | | 0.46 | 1.4 | [] | 4.21E+07 | 9.63E+07 | 6.93E+07 | 4.43E+07 | 9.53E+07 | 1.58E+08 |
| 271 | 60S ribosomal protein L9 OS=Homo sapiens OX=960 | P32969 | RPL9 | 390 | 22 kDa | | 0.47 | 1.4 | [] | 3.81E+08 | 2.38E+08 | 4.86E+07 | 3.44E+08 | 2.78E+08 | 2.81E+08 |
| 264 | Mitochondrial ribosomal protein L27 (Fragment) OS=I | H7C5U8 (+1) | MRPL27 | 41 | 15 kDa | | 0.48 | 1.4 | [] | 7026000 | 2.53E+07 | 4.47E+07 | 4.73E+07 | 2.96E+07 | 2.92E+07 |
| 268 | La-related protein 4 OS=Homo sapiens OX=9606 GN | Q71RC2 | LARP4 | 131 | 81 kDa | | 0.53 | 1.4 | [] | 2.91E+07 | 1.99E+07 | 1075100 | 2.46E+07 | 1.83E+07 | 2.48E+07 |
| 256 | 40S ribosomal protein S13 OS=Homo sapiens OX=96( | P62277 | RPS13 | 311 | 17 kDa | | 0.54 | 1.4 | [] | 3.83E+07 | 1.03E+08 | 8.09E+07 | 1.74E+07 | 1.44E+08 | 1.57E+08 |
| 260 | Cluster of Importin subunit alpha-5 OS=Homo sapiens | P52294 [2] | KPNA1 | 181 | 60 kDa | TRUE | 0.63 | 1.4 | [] | 2.02E+07 | 1.24E+07 | 1.60E+07 | 6008800 | 1.56E+07 | 4.64E+07 |
| 257 | 28S ribosomal protein S24, mitochondrial OS=Homo s | Q96EL2 | MRPS24 | 12 | 19 kDa | | 0.66 | 1.4 | [] | 4313300 | 4055200 | 5473600 | 1,000.00 | 1.41E+07 | 5660900 |
| 254 | DnaJ homolog subfamily C member 16 OS=Homo sap | Q5TDG9 (+2) | DNAJC16 | 1 | 69 kDa | | 0.7 | 1.4 | [] | 1,000.00 | 1,000.00 | 1.50E+07 | 3474800 | 9077200 | 8925100 |
| 258 | Replication factor C subunit 5 OS=Homo sapiens OX= | P40937 | RFC5 | 92 | 38 kDa | | 0.71 | 1.4 | [] | 1.41E+07 | 3.31E+07 | 1.51E+07 | 1782100 | 1.61E+07 | 7.11E+07 |
| 263 | Transforming growth factor beta receptor type 3 OS=I | Q03167 | TGFBR3 | 1 | 94 kDa | | 0.72 | 1.4 | [] | 3402200 | 4214800 | 1,000.00 | 1,000.00 | 3230700 | 7277800 |
| 267 | >sp\|Q9UJS0\|CMC2_HUMAN Calcium-binding mitocl | sp\|Q9UJS0\|CMC2_I | SLC25A13 | 160 | ? | TRUE | 0.73 | 1.4 | [] | 7.43E+07 | 4.28E+07 | 1.92E+07 | 2.53E+07 | 1.46E+08 | 1.43E+07 |
| 270 | rRNA methyltransferase 3, mitochondrial OS=Homo s | Q9HC36 | MRM3 | 39 | 47 kDa | | 0.77 | 1.4 | [] | 1,000.00 | 1,000.00 | 5402800 | 1776500 | 4314600 | 1214200 |
| 266 | Cluster of Matrin-3 OS=Homo sapiens OX=9606 GN= | A0A0R4J2E8 [4] | MATR3 | 441 | 95 kDa | TRUE | 0.8 | 1.4 | [] | 9441600 | 1,000.00 | 1,000.00 | 537850 | 9668900 | 2710100 |
| 276 | 60S ribosomal protein L8 OS=Homo sapiens OX=960 | P62917 | RPL8 | 390 | 28 kDa | | 0.17 | 1.3 | [] | 2.49E+08 | 1.68E+08 | 2.09E+08 | 2.47E+08 | 3.50E+08 | 2.41E+08 |
| 291 | Heterogeneous nuclear ribonucleoprotein U-like protei | Q1KMD3 | HNRNPUL2 | 135 | 85 kDa | | 0.2 | 1.3 | [] | 2.20E+07 | 2.35E+07 | 1.72E+07 | 2.11E+07 | 2.66E+07 | 3.30E+07 |

| | | | | | | | | | | | | | | | |
|---|---|---|---|---|---|---|---|---|---|---|---|---|---|---|---|
| 280 | Mitochondrial ribosomal protein L4 (Fragment) OS=H | K7ES61 (+1) | MRPL4 | 72 | 34 kDa | | 0.24 | 1.3 | [] | 1.01E+08 | 9.35E+07 | 1.32E+08 | 1.73E+08 | 1.62E+08 | 9.86E+07 |
| 283 | Insulin-like growth factor 2 mRNA-binding protein 3 C | O00425 | IGF2BP3 | 288 | 64 kDa | TRUE | 0.24 | 1.3 | [] | 2.45E+08 | 1.83E+08 | 1.38E+08 | 2.88E+08 | 1.89E+08 | 2.66E+08 |
| 285 | Heterogeneous nuclear ribonucleoprotein F OS=Homo | P52597 | HNRNPF | 528 | 46 kDa | TRUE | 0.27 | 1.3 | [] | 4.52E+08 | 3.43E+08 | 2.21E+08 | 3.58E+08 | 5.22E+08 | 4.49E+08 |
| 289 | Cluster of Fibrillarin (Fragment) OS=Homo sapiens OX | M0QXL5 [4] | FBL | 387 | 27 kDa | TRUE | 0.3 | 1.3 | [] | 2.49E+07 | 2.81E+07 | 2.84E+07 | 2.28E+07 | 3.68E+07 | 4.63E+07 |
| 281 | Cluster of Actin beta (Fragment) OS=Homo sapiens O | A0A2R8Y793 [6] | ACTB | 667 | 34 kDa | TRUE | 0.34 | 1.3 | [] | 2.35E+09 | 3.53E+09 | 2.87E+09 | 2.25E+09 | 4.84E+09 | 4.49E+09 |
| 273 | Cell cycle associated protein 1 (Fragment) OS=Homo s | E9PLA9 (+1) | CAPRIN1 | 282 | 20 kDa | | 0.35 | 1.3 | [] | 1.12E+08 | 9.02E+07 | 2.27E+08 | 2.23E+08 | 1.95E+08 | 1.59E+08 |
| 277 | 28S ribosomal protein S18b, mitochondrial OS=Homo | Q9Y676 | MRPS18B | 67 | 29 kDa | | 0.36 | 1.3 | [] | 8.04E+07 | 7.78E+07 | 1.74E+08 | 1.80E+08 | 1.29E+08 | 1.33E+08 |
| 282 | Cluster of 60S ribosomal protein L26 OS=Homo sapie | P61254 [2] | RPL26 | 312 | 17 kDa | TRUE | 0.37 | 1.3 | [] | 4.79E+07 | 8.89E+07 | 1.11E+08 | 7.24E+07 | 1.21E+08 | 1.32E+08 |
| 293 | General transcription factor IIIC subunit 5 OS=Homo s | Q5T7U1 (+1) | GTF3C5 | 136 | 52 kDa | | 0.46 | 1.3 | [] | 1.11E+07 | 1.09E+07 | 1.97E+07 | 1.97E+07 | 1.05E+07 | 2.30E+07 |
| 274 | Cluster of 60S ribosomal protein L3 OS=Homo sapien | P39023 [2] | RPL3 | 402 | 46 kDa | TRUE | 0.48 | 1.3 | [] | 2.66E+09 | 1.28E+09 | 5.98E+08 | 1.53E+09 | 2.44E+09 | 2.13E+09 |
| 288 | Transcription elongation factor SPT5 OS=Homo sapie | O00267 | SUPT5H | 158 | 121 kDa | | 0.48 | 1.3 | [] | 1.18E+08 | 6.65E+07 | 3.17E+07 | 6.99E+07 | 1.01E+08 | 1.11E+08 |
| 295 | GTP-binding protein 4 OS=Homo sapiens OX=9606 G | Q9BZE4 | GTPBP4 | 186 | 74 kDa | | 0.49 | 1.3 | [] | 2.31E+07 | 5.73E+07 | 3.76E+07 | 6.25E+07 | 3.15E+07 | 5.47E+07 |
| 278 | Cluster of Tubulin beta chain OS=Homo sapiens OX=9 | P07437 [10] | TUBB | 685 | 50 kDa | TRUE | 0.56 | 1.3 | [] | 2.04E+09 | 9.36E+08 | 8.01E+08 | 2.70E+09 | 9.09E+08 | 1.43E+09 |
| 275 | Heterogeneous nuclear ribonucleoprotein R OS=Homo | O43390 | HNRNPR | 422 | 71 kDa | TRUE | 0.59 | 1.3 | [] | 3.20E+07 | 1.44E+08 | 1.85E+08 | 1.06E+08 | 1.10E+08 | 2.66E+08 |
| 279 | 39S ribosomal protein L23, mitochondrial OS=Homo s | Q16540 | MRPL23 | 38 | 18 kDa | | 0.62 | 1.3 | [] | 1.98E+07 | 4.61E+07 | 1.31E+07 | 1.17E+07 | 5.67E+07 | 3.67E+07 |
| 296 | Testis-specific Y-encoded-like protein 5 OS=Homo sa | Q86VY4 | TSPYL5 | 2 | 45 kDa | | 0.65 | 1.3 | [] | 2.72E+07 | 8335700 | 9706700 | 2.82E+07 | 1.01E+07 | 1.86E+07 |
| 292 | 60S ribosomal protein L30 (Fragment) OS=Homo sapi | E5RI99 (+1) | RPL30 | 255 | 13 kDa | | 0.67 | 1.3 | [] | 2.10E+07 | 3.38E+07 | 4410200 | 9852900 | 3.87E+07 | 2.74E+07 |
| 290 | Small nuclear ribonucleoprotein Sm D3 OS=Homo sap | P62318 | SNRPD3 | 377 | 14 kDa | | 0.71 | 1.3 | [] | 4863700 | 8890600 | 1,000.00 | 1598100 | 8816300 | 7455400 |
| 287 | YTH domain-containing family protein 2 OS=Homo sa | Q9Y5A9 | YTHDF2 | 250 | 62 kDa | TRUE | 0.73 | 1.3 | [] | 8274300 | 5961700 | 1.70E+07 | 1,000.00 | 2.74E+07 | 1.33E+07 |
| 286 | Cluster of Y-box-binding protein 1 OS=Homo sapiens | P67809 [2] | YBX1 | 448 | 36 kDa | TRUE | 0.77 | 1.3 | [] | 1.49E+07 | 1,000.00 | 5.14E+07 | 8410000 | 1.88E+07 | 5.93E+07 |
| 284 | Protein RCC2 OS=Homo sapiens OX=9606 GN=RCC | Q9P258 | RCC2 | 336 | 56 kDa | | 0.82 | 1.3 | [] | 8087400 | 2.90E+07 | 4.45E+07 | 2656900 | 1.02E+08 | 2183300 |
| 294 | NADH dehydrogenase [ubiquinone] 1 alpha subcomple | Q16795 | NDUFA9 | 35 | 43 kDa | | 0.87 | 1.3 | [] | 5989700 | 1,000.00 | 1,000.00 | 1,000.00 | 7614100 | 1,000.00 |
| 302 | Cluster of 40S ribosomal protein S6 OS=Homo sapiens | P62753 [2] | RPS6 | 466 | 29 kDa | TRUE | 0.034 | 1.2 | [] | 1.36E+09 | 1.46E+09 | 1.33E+09 | 1.51E+09 | 1.80E+09 | 1.77E+09 |
| 312 | Cluster of Keratin, type II cytoskeletal 8 OS=Homo sa | P05787 [17] | KRT8 | 545 | 54 kDa | TRUE | 0.11 | 1.2 | [] | 2.09E+09 | 1.87E+09 | 1.60E+09 | 2.07E+09 | 2.18E+09 | 2.21E+09 |
| 306 | RNA helicase OS=Homo sapiens OX=9606 GN=DHX | H7BXY3 (+1) | DHX30 | 188 | 131 kDa | | 0.19 | 1.2 | [] | 1.20E+08 | 1.47E+08 | 1.72E+08 | 1.93E+08 | 1.65E+08 | 1.65E+08 |
| 316 | Cluster of Interleukin enhancer-binding factor 3 OS=H | Q12906 [3] | ILF3 | 379 | 95 kDa | TRUE | 0.52 | 1.2 | [] | 1.87E+08 | 3.12E+08 | 3.20E+08 | 3.70E+08 | 2.31E+08 | 3.46E+08 |
| 297 | Ribosomal protein S14 (Fragment) OS=Homo sapiens | A0A2R8Y811 (+1) | RPS14 | 514 | 16 kDa | | 0.57 | 1.2 | [] | 3.45E+08 | 4.59E+08 | 9.35E+08 | 9.59E+08 | 5.70E+08 | 6.13E+08 |
| 300 | General transcription factor 3C polypeptide 1 OS=Hom | Q12789 | GTF3C1 | 134 | 239 kDa | | 0.59 | 1.2 | [] | 4899600 | 5960800 | 9580500 | 4124000 | 1.18E+07 | 9199500 |
| 310 | Activating signal cointegrator 1 complex subunit 3 OS= | Q8N3C0 | ASCC3 | 93 | 251 kDa | | 0.59 | 1.2 | [] | 7.88E+07 | 3.53E+07 | 9.51E+07 | 1.02E+08 | 7.29E+07 | 6.99E+07 |
| 318 | Cluster of Heterogeneous nuclear ribonucleoprotein K | P61978 [2] | HNRNPK | 581 | 51 kDa | TRUE | 0.59 | 1.2 | [] | 1.31E+09 | 1.36E+09 | 1.78E+09 | 2.42E+09 | 1.33E+09 | 1.38E+09 |
| 304 | 39S ribosomal protein L22, mitochondrial OS=Homo s | E7ESL0 (+2) | MRPL22 | 72 | 24 kDa | | 0.63 | 1.2 | [] | 3.90E+07 | 8.90E+07 | 1.63E+08 | 9.07E+07 | 1.14E+08 | 1.49E+08 |
| 314 | Cluster of ADP/ATP translocase 2 OS=Homo sapiens | P05141 [3] | SLC25A5 | 525 | 33 kDa | TRUE | 0.68 | 1.2 | [] | 3.98E+08 | 4.14E+08 | 3.05E+08 | 1.82E+08 | 5.02E+08 | 6.10E+08 |
| 301 | ATP binding cassette subfamily F member 1 OS=Homo | Q5STZ8 (+1) | ABCF1 | 288 | 96 kDa | | 0.7 | 1.2 | [] | 6490300 | 3735900 | 1.37E+07 | 3287300 | 1.34E+07 | 1.26E+07 |

| | | | | | | | | | | | | | | | |
|---|---|---|---|---|---|---|---|---|---|---|---|---|---|---|---|
| 303 | Cluster of Zinc finger CCCH-type antiviral protein 1 O | Q7Z2W4 [2] | ZC3HAV1 | 210 | 101 kDa | TRUE | 0.71 | 1.2 | [] | 2.56E+08 | 2.34E+08 | 5.12E+08 | 7.49E+08 | 2.42E+08 | 2.37E+08 |
| 319 | 40S ribosomal protein S20 OS=Homo sapiens OX=96( | P60866 | RPS20 | 383 | 13 kDa | | 0.73 | 1.2 | [] | 2.29E+08 | 9.35E+07 | 1.20E+08 | 2.60E+08 | 1.12E+08 | 1.38E+08 |
| 315 | 28S ribosomal protein S31, mitochondrial OS=Homo s | Q92665 | MRPS31 | 92 | 45 kDa | | 0.74 | 1.2 | [] | 8.62E+07 | 8.97E+07 | 5.94E+07 | 2.45E+07 | 1.37E+08 | 1.11E+08 |
| 307 | 40S ribosomal protein S24 OS=Homo sapiens OX=96( | A0A2R8Y849 (+3) | RPS24 | 353 | 15 kDa | | 0.75 | 1.2 | [] | 8.29E+08 | 4.86E+08 | 1.10E+08 | 2.45E+08 | 8.30E+08 | 6.21E+08 |
| 313 | Cluster of Polypyrimidine tract-binding protein 1 OS=I | P26599 [2] | PTBP1 | 438 | 60 kDa | TRUE | 0.76 | 1.2 | [] | 6.54E+08 | 4.01E+08 | 1.31E+08 | 2.79E+08 | 6.89E+08 | 4.10E+08 |
| 299 | Mitochondrial ribosomal protein L43 OS=Homo sapie | B1AL05 (+2) | MRPL43 | 47 | 21 kDa | | 0.77 | 1.2 | [] | 1.76E+07 | 1,000.00 | 6.68E+07 | 3.06E+07 | 3.41E+07 | 3.88E+07 |
| 309 | Signal recognition particle subunit SRP72 OS=Homo s | O76094 | SRP72 | 168 | 75 kDa | | 0.79 | 1.2 | [] | 3.60E+07 | 5.34E+07 | 3.62E+07 | 1,000.00 | 8.78E+07 | 6.10E+07 |
| 298 | Probable ATP-dependent RNA helicase DHX37 OS=H | Q8IY37 | DHX37 | 65 | 130 kDa | | 0.82 | 1.2 | [] | 2578100 | 4969700 | 2.91E+07 | 2.26E+07 | 2.23E+07 | 1,000.00 |
| 311 | Cluster of Ataxin-2-like protein OS=Homo sapiens OX | Q8WWM7 [2] | ATXN2L | 305 | 113 kDa | TRUE | 0.82 | 1.2 | [] | 2.93E+07 | 2.69E+07 | 8.47E+07 | 1.05E+08 | 2.34E+07 | 3.61E+07 |
| 308 | LIM and SH3 domain protein 1 OS=Homo sapiens OX | Q14847 | LASP1 | 60 | 30 kDa | | 0.86 | 1.2 | [] | 1,000.00 | 1498400 | 2492600 | 3700300 | 1047500 | 1,000.00 |
| 305 | Amyloid beta precursor like protein 2 OS=Homo sapie | Q06481 | APLP2 | 2 | 87 kDa | | 0.91 | 1.2 | [] | 1863500 | 1,000.00 | 1,000.00 | 1,000.00 | 2227400 | 1,000.00 |
| 317 | Protein-glutamine gamma-glutamyltransferase 2 OS=H | P21980 | TGM2 | 10 | 77 kDa | | 0.92 | 1.2 | [] | 2441700 | 1,000.00 | 1,000.00 | 1,000.00 | 2511600 | 308350 |
| 321 | Ribosomal protein L19 OS=Homo sapiens OX=9606 ( | A0A7I2YQG2 (+3) | RPL19 | 458 | 26 kDa | | 0.68 | 1.1 | [] | 1.03E+09 | 1.41E+09 | 1.27E+09 | 6.61E+08 | 1.63E+09 | 1.95E+09 |
| 325 | Cluster of 40S ribosomal protein S4, X isoform OS=H | P62701 [3] | RPS4X | 451 | 30 kDa | TRUE | 0.71 | 1.1 | [] | 7.54E+08 | 5.53E+08 | 4.37E+08 | 4.04E+08 | 9.78E+08 | 5.96E+08 |
| 339 | Serine/threonine-protein kinase VRK2 OS=Homo sapi | Q86Y07 | VRK2 | 36 | 58 kDa | | 0.74 | 1.1 | [] | 1.37E+07 | 3.24E+07 | 1.79E+07 | 1.98E+07 | 2.48E+07 | 2.57E+07 |
| 324 | FAU ubiquitin like and ribosomal protein S30 fusion O | E9PR30 (+1) | FAU | 320 | 11 kDa | | 0.76 | 1.1 | [] | 2.23E+07 | 2.99E+07 | 4.07E+07 | 5.10E+07 | 4.32E+07 | 1.14E+07 |
| 332 | 39S ribosomal protein S30, mitochondrial OS=Homo s | Q9NP92 | MRPS30 | 21 | 50 kDa | | 0.77 | 1.1 | [] | 3.54E+07 | 1.51E+07 | 2.26E+07 | 3.94E+07 | 2.51E+07 | 1.68E+07 |
| 320 | Transducin beta like 2 OS=Homo sapiens OX=9606 G | E9PF19 (+1) | TBL2 | 47 | 46 kDa | | 0.78 | 1.1 | [] | 1.34E+07 | 4.33E+07 | 8.00E+07 | 7.54E+07 | 4.43E+07 | 3.72E+07 |
| 335 | 60S ribosomal protein L23a OS=Homo sapiens OX=9( | P62750 | RPL23A | 418 | 18 kDa | | 0.79 | 1.1 | [] | 2.66E+08 | 1.62E+08 | 4.91E+08 | 3.33E+08 | 2.37E+08 | 4.47E+08 |
| 323 | 60S ribosomal protein L31 OS=Homo sapiens OX=96( | B7Z4C8 (+4) | RPL31 | 335 | 15 kDa | | 0.8 | 1.1 | [] | 3.29E+07 | 1.96E+07 | 2867300 | 1.32E+07 | 2.09E+07 | 2.92E+07 |
| 326 | Eukaryotic translation initiation factor 5B OS=Homo s | A0A087WUT6 (+1) | EIF5B | 200 | 139 kDa | | 0.8 | 1.1 | [] | 1.09E+08 | 4.27E+07 | 2.15E+07 | 8.02E+07 | 6.75E+07 | 4.78E+07 |
| 331 | 60S ribosomal protein L10 OS=Homo sapiens OX=96( | P27635 | RPL10 | 369 | 25 kDa | | 0.8 | 1.1 | [] | 7.13E+08 | 5.64E+08 | 3.16E+08 | 3.18E+08 | 5.13E+08 | 9.43E+08 |
| 329 | Ribosomal protein L22 (Fragment) OS=Homo sapiens | K7ELC4 (+3) | RPL22 | 429 | 9 kDa | | 0.83 | 1.1 | [] | 1.41E+07 | 7.41E+07 | 1.36E+08 | 9.91E+07 | 6.26E+07 | 8.79E+07 |
| 327 | ELAV-like protein 1 OS=Homo sapiens OX=9606 GN | Q15717 | ELAVL1 | 322 | 36 kDa | | 0.84 | 1.1 | [] | 2064600 | 1.41E+07 | 1.64E+07 | 4422900 | 1.44E+07 | 1.77E+07 |
| 328 | tRNA methyltransferase 10 homolog C OS=Homo sap | Q7L0Y3 | TRMT10C | 76 | 47 kDa | | 0.84 | 1.1 | [] | 2.45E+07 | 2.07E+07 | 1.83E+07 | 8335600 | 4.66E+07 | 1.61E+07 |
| 352 | 28S ribosomal protein S15, mitochondrial OS=Homo s | P82914 | MRPS15 | 39 | 30 kDa | | 0.84 | 1.1 | [] | 9.02E+07 | 6.79E+07 | 1.15E+08 | 1.51E+08 | 7.30E+07 | 6.86E+07 |
| 330 | Acyl-CoA:lysophosphatidylglycerol acyltransferase 1 C | Q92604 | LPGAT1 | 3 | 43 kDa | | 0.87 | 1.1 | [] | 1223500 | 2919000 | 2403200 | 1,000.00 | 4480300 | 2813700 |
| 333 | Peptidyl-prolyl cis-trans isomerase-like 4 OS=Homo sa | Q8WUA2 | PPIL4 | 213 | 57 kDa | | 0.87 | 1.1 | [] | 2.16E+07 | 7.23E+07 | 5.05E+07 | 1,000.00 | 8.51E+07 | 7.55E+07 |
| 340 | FMR1 autosomal homolog 1 OS=Homo sapiens OX=9 | B4DXZ6 (+1) | FXR1 | 140 | 68 kDa | TRUE | 0.87 | 1.1 | [] | 3.00E+07 | 8.07E+07 | 1.49E+08 | 3.18E+07 | 9.60E+07 | 1.57E+08 |
| 349 | Cluster of Ribosome-binding protein 1 OS=Homo sapi | Q9P2E9 [2] | RRBP1 | 172 | 152 kDa | TRUE | 0.87 | 1.1 | [] | 1.66E+08 | 3.64E+08 | 7.28E+08 | 6.53E+08 | 3.19E+08 | 3.84E+08 |
| 351 | Sequestosome-1 OS=Homo sapiens OX=9606 GN=SC | Q13501 | SQSTM1 | 138 | 48 kDa | | 0.87 | 1.1 | [] | 2.33E+08 | 1.62E+08 | 1.85E+08 | 4.78E+07 | 3.21E+08 | 2.55E+08 |
| 342 | Drebrin-like protein OS=Homo sapiens OX=9606 GN= | Q9UJU6 | DBNL | 146 | 48 kDa | | 0.88 | 1.1 | [] | 1,000.00 | 3.31E+07 | 5.64E+07 | 3.64E+07 | 3.42E+07 | 2.70E+07 |
| 353 | Pentatricopeptide repeat domain-containing protein 3, | Q96EY7 | PTCD3 | 83 | 79 kDa | | 0.88 | 1.1 | [] | 1.01E+08 | 1.13E+08 | 1.93E+08 | 5.06E+07 | 1.61E+08 | 2.24E+08 |

| | | | | | | | | | | | | | | | |
|---|---|---|---|---|---|---|---|---|---|---|---|---|---|---|---|
| 347 | 60S ribosomal protein L14 OS=Homo sapiens OX=96 | P50914 | RPL14 | 333 | 23 kDa | | 0.89 | 1.1 | [] | 4.19E+08 | 2.43E+08 | 1.49E+08 | 4.79E+07 | 4.92E+08 | 3.37E+08 |
| 354 | 28S ribosomal protein S35, mitochondrial OS=Homo s | P82673 | MRPS35 | 74 | 37 kDa | | 0.89 | 1.1 | [] | 7.66E+07 | 7.02E+07 | 4.37E+07 | 2.75E+07 | 1.17E+08 | 5.89E+07 |
| 338 | Leucine rich pentatricopeptide repeat containing OS=H | A0A804HIG4 (+7) | LRPPRC | 249 | 155 kDa | | 0.9 | 1.1 | [] | 1.03E+07 | 9379800 | 1.44E+07 | 634420 | 2.90E+07 | 7741300 |
| 341 | U4/U6 small nuclear ribonucleoprotein Prp31 OS=Hor | Q8WWY3 | PRPF31 | 251 | 55 kDa | | 0.9 | 1.1 | [] | 8017500 | 7503100 | 5490300 | 5970000 | 1.70E+07 | 1,000.00 |
| 346 | Tripartite motif containing 25 OS=Homo sapiens OX= | A0A3B3IUA7 (+1) | TRIM25 | 136 | 51 kDa | | 0.9 | 1.1 | [] | 1.67E+07 | 1.76E+07 | 5170300 | 2.10E+07 | 2.18E+07 | 1,000.00 |
| 334 | Tripartite motif-containing protein 26 OS=Homo sapie | Q12899 | TRIM26 | 13 | 62 kDa | | 0.91 | 1.1 | [] | 1.27E+07 | 1,000.00 | 5335700 | 1,000.00 | 1.32E+07 | 6701500 |
| 343 | ATP-binding cassette sub-family F member 2 OS=Hon | Q9UG63 | ABCF2 | 137 | 71 kDa | | 0.91 | 1.1 | [] | 3.65E+07 | 8393400 | 6308300 | 1832600 | 2.70E+07 | 2.69E+07 |
| 345 | 24-dehydrocholesterol reductase OS=Homo sapiens O | A0A0A0MTI1 (+1) | DHCR24 | 9 | 61 kDa | | 0.92 | 1.1 | [] | 1,000.00 | 4238500 | 3819600 | 1,000.00 | 3545600 | 5208500 |
| 322 | Dolichyl-diphosphooligosaccharide--protein glycosyltra | P04844 | RPN2 | 179 | 69 kDa | | 0.93 | 1.1 | [] | 3033300 | 1,000.00 | 1,000.00 | 3468100 | 1,000.00 | 1,000.00 |
| 336 | Chromobox protein homolog 3 OS=Homo sapiens OX | Q13185 | CBX3 | 309 | 21 kDa | | 0.93 | 1.1 | [] | 1,000.00 | 1.23E+07 | 1,000.00 | 1,000.00 | 6599800 | 7031600 |
| 348 | Poly(A) RNA polymerase, mitochondrial OS=Homo sa | Q9NVV4 | MTPAP | 59 | 66 kDa | | 0.93 | 1.1 | [] | 1,000.00 | 3726200 | 8103500 | 1,000.00 | 6685300 | 6080400 |
| 337 | Agrin OS=Homo sapiens OX=9606 GN=AGRN PE=1 | A0A494C0G5 | AGRN | 5 | 203 kDa | | 0.94 | 1.1 | [] | 1779600 | 1011900 | 1,000.00 | 1,000.00 | 3068400 | 1,000.00 |
| 350 | Cluster of Y-box-binding protein 2 OS=Homo sapiens | Q9Y2T7 [2] | YBX2 | 343 | 39 kDa | TRUE | 0.94 | 1.1 | [] | 1,000.00 | 1,000.00 | 4.54E+07 | 1.50E+07 | 1.16E+07 | 2.22E+07 |
| 357 | Cluster of DNA-directed RNA polymerase, mitochond | O00411 [2] | POLRMT | 49 | 139 kDa | TRUE | 0.94 | 1.1 | [] | 5.21E+07 | 1.15E+07 | 3.71E+07 | 1.48E+07 | 7.67E+07 | 1.43E+07 |
| 344 | Cluster of 40S ribosomal protein S7 OS=Homo sapien: | P62081 [2] | RPS7 | 368 | 22 kDa | TRUE | 0.95 | 1.1 | [] | 3.83E+07 | 1905700 | 1,000.00 | 1,000.00 | 3.60E+07 | 7786800 |
| 356 | DnaJ homolog subfamily C member 10 OS=Homo sapi | A0A7P0TAQ9 (+3) | DNAJC10 | 131 | 88 kDa | | 0.95 | 1.1 | [] | 1636500 | 1,000.00 | 2046300 | 2413000 | 1466200 | 1,000.00 |
| 355 | Nucleolar protein 58 OS=Homo sapiens OX=9606 GN | Q9Y2X3 | NOP58 | 246 | 60 kDa | | 0.96 | 1.1 | [] | 1,000.00 | 1,000.00 | 7609800 | 2499200 | 1,000.00 | 5535200 |
| 473 | Cluster of Elongation factor 1-alpha 1 OS=Homo sapi | P68104 [3] | EEF1A1 | 653 | 50 kDa | TRUE | 0.86 | 1 | [] | 2.73E+08 | 2.10E+08 | 1.62E+08 | 1.59E+08 | 2.48E+08 | 2.15E+08 |
| 472 | 39S ribosomal protein L14, mitochondrial OS=Homo s | Q6P1L8 | MRPL14 | 74 | 16 kDa | | 0.9 | 1 | [] | 1.74E+07 | 1.19E+07 | 1.15E+07 | 1.91E+07 | 9680100 | 1.07E+07 |
| 475 | 60S acidic ribosomal protein P0 OS=Homo sapiens O | P05388 | RPLP0 | 436 | 34 kDa | | 0.9 | 1 | [] | 2.17E+07 | 4.40E+07 | 2.72E+07 | 2.35E+07 | 2.33E+07 | 4.24E+07 |
| 362 | La-related protein 7 OS=Homo sapiens OX=9606 GN | Q4G0J3 | LARP7 | 92 | 67 kDa | | 0.92 | 1 | [] | 5.11E+07 | 4.31E+07 | 6.87E+07 | 8.74E+07 | 3.56E+07 | 4.52E+07 |
| 367 | Interleukin enhancer binding factor 2 OS=Homo sapier | B4DY09 (+1) | ILF2 | 313 | 39 kDa | | 0.93 | 1 | [] | 1.91E+08 | 1.30E+08 | 2.14E+08 | 2.20E+08 | 1.88E+08 | 1.37E+08 |
| 358 | 39S ribosomal protein L47, mitochondrial OS=Homo s | Q9HD33 | MRPL47 | 40 | 29 kDa | | 0.94 | 1 | [] | 9.79E+07 | 2.32E+08 | 3.19E+07 | 4.15E+07 | 2.09E+08 | 1.29E+08 |
| 366 | Cluster of Propionyl-CoA carboxylase alpha chain, mit | P05165 [3] | PCCA | 284 | 80 kDa | TRUE | 0.95 | 1 | [] | 1.81E+08 | 2.62E+08 | 2.07E+08 | 6.49E+07 | 3.01E+08 | 3.01E+08 |
| 370 | Aspartyl/asparaginyl beta-hydroxylase OS=Homo sapi | Q12797 | ASPH | 28 | 86 kDa | | 0.97 | 1 | [] | 5.15E+08 | 7.23E+08 | 3.91E+08 | 4.64E+08 | 6.91E+08 | 4.87E+08 |
| 471 | Peptidyl-tRNA hydrolase 1 homolog (Fragment) OS=H | A0A096LP22 (+2) | PTRH1 | 2 | 19 kDa | | 0.97 | 1 | [] | 9029200 | 1.20E+07 | 1,000.00 | 1,000.00 | 1.11E+07 | 9369700 |
| 474 | Cluster of Ankyrin repeat and KH domain-containing p | Q8IWZ3 [2] | ANKHD1 | 130 | 269 kDa | TRUE | 0.97 | 1 | [] | 1.30E+07 | 1,000.00 | 6270000 | 1.57E+07 | 1,000.00 | 2842100 |
| 359 | Aminoacyl tRNA synthase complex-interacting multifu | Q12904 | AIMP1 | 163 | 34 kDa | | 0.98 | 1 | [] | 3263100 | 1,000.00 | 1,000.00 | 1,000.00 | 3403900 | 1,000.00 |
| 360 | Protein disulfide-isomerase OS=Homo sapiens OX=96 | A0A7P0TA35 | P4HB | 247 | 61 kDa | | 0.98 | 1 | [] | 1,000.00 | 1282700 | 1,000.00 | 1,000.00 | 1,000.00 | 1336800 |
| 361 | 39S ribosomal protein L13, mitochondrial OS=Homo s | Q9BYD1 | MRPL13 | 49 | 21 kDa | | 0.98 | 1 | [] | 1,000.00 | 6128200 | 1,000.00 | 1,000.00 | 4963400 | 1380100 |
| 363 | Splicing factor 3B subunit 3 OS=Homo sapiens OX=96 | Q15393 | SF3B3 | 322 | 136 kDa | | 0.98 | 1 | [] | 1,000.00 | 5008900 | 9.75E+07 | 1953800 | 5.24E+07 | 5.15E+07 |
| 364 | Pre-mRNA-splicing factor ATP-dependent RNA helica | Q92620 | DHX38 | 110 | 141 kDa | | 0.98 | 1 | [] | 1166000 | 1,000.00 | 1,000.00 | 1,000.00 | 1201100 | 1,000.00 |
| 365 | Acyl-CoA 6-desaturase OS=Homo sapiens OX=9606 ( | O95864 | FADS2 | 4 | 52 kDa | | 0.98 | 1 | [] | 1,000.00 | 1.71E+07 | 1,000.00 | 1,000.00 | 1.20E+07 | 5582800 |

| | | | | | | | | | | | | | | | |
|---|---|---|---|---|---|---|---|---|---|---|---|---|---|---|---|
| 369 | 28S ribosomal protein S5, mitochondrial OS=Homo sa | P82675 | MRPS5 | 66 | 48 kDa | | 0.98 | 1 | [] | 2.37E+08 | 1.71E+08 | 1.51E+08 | 6.75E+07 | 2.79E+08 | 2.19E+08 |
| 371 | Cluster of Actinin alpha 1 OS=Homo sapiens OX=960 | A0A7I2V4Y4 [11] | ACTN1 | 314 | 104 kDa | TRUE | 0.98 | 1 | [] | 8.78E+07 | 5.93E+07 | 1.55E+08 | 1.18E+08 | 1.04E+08 | 8.18E+07 |
| 372 | Mitochondrial ribosomal protein S22 OS=Homo sapie | A0A7P0MKV3 (+2) | MRPS22 | 90 | 41 kDa | | 0.98 | 1 | [] | 1.45E+08 | 2.72E+08 | 2.68E+08 | 1.33E+08 | 2.81E+08 | 2.77E+08 |
| 467 | 28S ribosomal protein S29, mitochondrial OS=Homo s | P51398 | DAP3 | 96 | 46 kDa | | 0.98 | 1 | [] | 9.72E+07 | 4.39E+07 | 9.75E+07 | 8.43E+07 | 5.09E+07 | 1.02E+08 |
| 469 | Golgi integral membrane protein 4 OS=Homo sapiens | O00461 | GOLIM4 | 10 | 82 kDa | | 0.98 | 1 | [] | 2.45E+07 | 3.16E+07 | 6.77E+07 | 5.14E+07 | 4.42E+07 | 2.69E+07 |
| 468 | ATP-dependent RNA helicase OS=Homo sapiens OX= | G3V529 (+1) | DDX24 | 138 | 91 kDa | | 0.99 | 1 | [] | 1,000.00 | 1,000.00 | 4836200 | 1,000.00 | 2727300 | 2068100 |
| 373 | Poly(rC) binding protein 2 OS=Homo sapiens OX=96 | F8VZX2 (+2) | PCBP2 | 475 | 34 kDa | TRUE | 1 | 1 | [] | 1.32E+07 | 1.32E+07 | 3.06E+07 | 5.02E+07 | 1,000.00 | 6986900 |
| 375 | E3 ubiquitin-protein ligase NEDD4 OS=Homo sapiens | P46934 | NEDD4 | 3 | 149 kDa | TRUE | 1 | 1 | [] | 1,000.00 | 1,000.00 | 1,000.00 | 1,000.00 | 1,000.00 | 1,000.00 |
| 376 | Ribosomal L1 domain containing 1 (Fragment) OS=Ho | J3QSV6 (+1) | RSL1D1 | 277 | 48 kDa | | 1 | 1 | [] | 1,000.00 | 1,000.00 | 1,000.00 | 1,000.00 | 1,000.00 | 1,000.00 |
| 377 | Testis-expressed protein 10 OS=Homo sapiens OX=96 | Q9NXF1 | TEX10 | 115 | 106 kDa | | 1 | 1 | [] | 1,000.00 | 1,000.00 | 1,000.00 | 1,000.00 | 1,000.00 | 1,000.00 |
| 378 | Spectrin repeat containing nuclear envelope protein 2 | G3V5X4 (+1) | SYNE2 | 26 | 788 kDa | | 1 | 1 | [] | 1,000.00 | 1,000.00 | 1,000.00 | 1,000.00 | 1,000.00 | 1,000.00 |
| 379 | Paraneoplastic antigen Ma2 OS=Homo sapiens OX=9 | Q9UL42 | PNMA2 | 24 | 42 kDa | | 1 | 1 | [] | 1,000.00 | 1,000.00 | 1,000.00 | 1,000.00 | 1,000.00 | 1,000.00 |
| 380 | Ribosome biogenesis protein BRX1 homolog OS=Hon | Q8TDN6 | BRIX1 | 178 | 41 kDa | | 1 | 1 | [] | 1,000.00 | 1,000.00 | 1,000.00 | 1,000.00 | 1,000.00 | 1,000.00 |
| 381 | Serine/threonine-protein kinase RIO1 OS=Homo sapie | Q9BRS2 | RIOK1 | 153 | 66 kDa | | 1 | 1 | [] | 1,000.00 | 1,000.00 | 1,000.00 | 1,000.00 | 1,000.00 | 1,000.00 |
| 382 | Transgelin-2 OS=Homo sapiens OX=9606 GN=TAGL | P37802 | TAGLN2 | 410 | 22 kDa | | 1 | 1 | [] | 1,000.00 | 1,000.00 | 1,000.00 | 1,000.00 | 1,000.00 | 1,000.00 |
| 383 | tRNA (guanine(6)-N2)-methyltransferase THUMP3 O | Q9BV44 | THUMPD3 | 12 | 57 kDa | | 1 | 1 | [] | 1,000.00 | 1,000.00 | 1,000.00 | 1,000.00 | 1,000.00 | 1,000.00 |
| 384 | C-1-tetrahydrofolate synthase, cytoplasmic (Fragment) | A0A494C1T2 (+4) | MTHFD1 | 311 | 70 kDa | | 1 | 1 | [] | 1,000.00 | 1,000.00 | 1,000.00 | 1,000.00 | 1,000.00 | 1,000.00 |
| 385 | EBNA1 binding protein 2 OS=Homo sapiens OX=960 | H7C2Q8 (+1) | EBNA1BP2 | 137 | 41 kDa | | 1 | 1 | [] | 1,000.00 | 1,000.00 | 1,000.00 | 1,000.00 | 1,000.00 | 1,000.00 |
| 386 | Zinc finger and BTB domain-containing protein 1 OS= | Q9Y2K1 | ZBTB1 | 0 | 82 kDa | | 1 | 1 | [] | 1,000.00 | 1,000.00 | 1,000.00 | 1,000.00 | 1,000.00 | 1,000.00 |
| 387 | DNA mismatch repair protein Msh6 OS=Homo sapien | P52701 | MSH6 | 222 | 153 kDa | | 1 | 1 | [] | 1,000.00 | 1,000.00 | 1,000.00 | 1,000.00 | 1,000.00 | 1,000.00 |
| 388 | Proto-oncogene c-Rel OS=Homo sapiens OX=9606 G | Q04864 | REL | 1 | 69 kDa | | 1 | 1 | [] | 1,000.00 | 1,000.00 | 1,000.00 | 1,000.00 | 1,000.00 | 1,000.00 |
| 389 | Serine/threonine-protein kinase RIO2 OS=Homo sapie | Q9BVS4 | RIOK2 | 14 | 63 kDa | | 1 | 1 | [] | 1,000.00 | 1,000.00 | 1,000.00 | 1,000.00 | 1,000.00 | 1,000.00 |
| 390 | A-kinase anchor protein 12 OS=Homo sapiens OX=96 | Q02952 | AKAP12 | 182 | 191 kDa | | 1 | 1 | [] | 1,000.00 | 1,000.00 | 1,000.00 | 1,000.00 | 1,000.00 | 1,000.00 |
| 391 | Testis-specific Y-encoded-like protein 2 OS=Homo sa | Q9H2G4 | TSPYL2 | 1 | 79 kDa | | 1 | 1 | [] | 1,000.00 | 1,000.00 | 1,000.00 | 1,000.00 | 1,000.00 | 1,000.00 |
| 392 | Cluster of Metastasis associated 1 family member 3 OS | E7EQY4 [5] | MTA3 | 166 | 59 kDa | TRUE | 1 | 1 | [] | 1,000.00 | 1,000.00 | 1,000.00 | 1,000.00 | 1,000.00 | 1,000.00 |
| 393 | Dishevelled segment polarity protein 2 OS=Homo sapi | I3L2N2 (+1) | DVL2 | 130 | 78 kDa | | 1 | 1 | [] | 1,000.00 | 1,000.00 | 1,000.00 | 1,000.00 | 1,000.00 | 1,000.00 |
| 394 | Protein angel homolog 1 OS=Homo sapiens OX=9606 | Q9UNK9 | ANGEL1 | 0 | 75 kDa | | 1 | 1 | [] | 1,000.00 | 1,000.00 | 1,000.00 | 1,000.00 | 1,000.00 | 1,000.00 |
| 395 | Protein ELFN1 OS=Homo sapiens OX=9606 GN=EL | P0C7U0 | ELFN1 | 0 | 90 kDa | | 1 | 1 | [] | 1,000.00 | 1,000.00 | 1,000.00 | 1,000.00 | 1,000.00 | 1,000.00 |
| 396 | Cluster of Poly(A)-specific ribonuclease PARN OS=H | A0A494C0W0 [3] | PARN | 30 | 76 kDa | TRUE | 1 | 1 | [] | 1,000.00 | 1,000.00 | 1,000.00 | 1,000.00 | 1,000.00 | 1,000.00 |
| 397 | Nuclear speckle splicing regulatory protein 1 OS=Hon | Q9H0G5 | NSRP1 | 131 | 66 kDa | | 1 | 1 | [] | 1,000.00 | 1,000.00 | 1,000.00 | 1,000.00 | 1,000.00 | 1,000.00 |
| 398 | Aurora kinase A OS=Homo sapiens OX=9606 GN=A | O14965 | AURKA | 24 | 46 kDa | | 1 | 1 | [] | 1,000.00 | 1,000.00 | 1,000.00 | 1,000.00 | 1,000.00 | 1,000.00 |
| 399 | Probable RNA-binding protein 19 OS=Homo sapiens | Q9Y4C8 | RBM19 | 23 | 107 kDa | | 1 | 1 | [] | 1,000.00 | 1,000.00 | 1,000.00 | 1,000.00 | 1,000.00 | 1,000.00 |
| 400 | BRCA1 associated protein OS=Homo sapiens OX=96 | J3KNN7 (+1) | BRAP | 93 | 64 kDa | | 1 | 1 | [] | 1,000.00 | 1,000.00 | 1,000.00 | 1,000.00 | 1,000.00 | 1,000.00 |

| | | | | | | | | | | | | | | | |
|---|---|---|---|---|---|---|---|---|---|---|---|---|---|---|---|
| 401 | Dyskerin pseudouridine synthase 1 OS=Homo sapiens | C9IYT0 (+2) | DKC1 | 186 | 43 kDa | | 1 | 1 | [] | 1,000.00 | 1,000.00 | 1,000.00 | 1,000.00 | 1,000.00 | 1,000.00 |
| 402 | Putative pre-mRNA-splicing factor ATP-dependent RN | Q7L7V1 | DHX32 | 7 | 84 kDa | | 1 | 1 | [] | 1,000.00 | 1,000.00 | 1,000.00 | 1,000.00 | 1,000.00 | 1,000.00 |
| 403 | Probable ATP-dependent RNA helicase DDX41 OS=H | Q9UJV9 | DDX41 | 171 | 70 kDa | | 1 | 1 | [] | 1,000.00 | 1,000.00 | 1,000.00 | 1,000.00 | 1,000.00 | 1,000.00 |
| 404 | Sentrin-specific protease 3 OS=Homo sapiens OX=96( | Q9H4L4 | SENP3 | 136 | 65 kDa | | 1 | 1 | [] | 1,000.00 | 1,000.00 | 1,000.00 | 1,000.00 | 1,000.00 | 1,000.00 |
| 405 | Cluster of Eukaryotic translation initiation factor 2 sub | H0YJS4 [2] | EIF2S1 | 240 | 29 kDa | TRUE | 1 | 1 | [] | 1,000.00 | 1,000.00 | 1,000.00 | 1,000.00 | 1,000.00 | 1,000.00 |
| 406 | NADH dehydrogenase [ubiquinone] 1 beta subcomplex | A0A7I2V620 (+2) | NDUFB9 | 6 | 20 kDa | | 1 | 1 | [] | 1,000.00 | 1,000.00 | 1,000.00 | 1,000.00 | 1,000.00 | 1,000.00 |
| 407 | Apoptosis inducing factor mitochondria associated 1 O | A0A6Q8PFA7 (+10 | AIFM1 | 226 | 50 kDa | | 1 | 1 | [] | 1,000.00 | 1,000.00 | 1,000.00 | 1,000.00 | 1,000.00 | 1,000.00 |
| 408 | RNA-binding protein NOB1 OS=Homo sapiens OX=9 | Q9ULX3 | NOB1 | 32 | 47 kDa | | 1 | 1 | [] | 1,000.00 | 1,000.00 | 1,000.00 | 1,000.00 | 1,000.00 | 1,000.00 |
| 409 | EMAP like 4 OS=Homo sapiens OX=9606 GN=EML4 | B5MBZ0 (+1) | EML4 | 69 | 110 kDa | | 1 | 1 | [] | 1,000.00 | 1,000.00 | 1,000.00 | 1,000.00 | 1,000.00 | 1,000.00 |
| 410 | Cluster of Unconventional myosin-Ie OS=Homo sapier | Q12965 [2] | MYO1E | 30 | 127 kDa | TRUE | 1 | 1 | [] | 1,000.00 | 1,000.00 | 1,000.00 | 1,000.00 | 1,000.00 | 1,000.00 |
| 411 | SCY1 like pseudokinase 1 OS=Homo sapiens OX=96( | E9PK59 (+3) | SCYL1 | 12 | 86 kDa | | 1 | 1 | [] | 1,000.00 | 1,000.00 | 1,000.00 | 1,000.00 | 1,000.00 | 1,000.00 |
| 412 | Cluster of PTPRF interacting protein alpha 1 (Fragmen | A0A2R8Y7R9 [8] | PPFIA1 | 14 | 139 kDa | TRUE | 1 | 1 | [] | 1,000.00 | 1,000.00 | 1,000.00 | 1,000.00 | 1,000.00 | 1,000.00 |
| 413 | Methylosome subunit pICln OS=Homo sapiens OX=9( | E9PJF4 (+3) | CLNS1A | 205 | 20 kDa | | 1 | 1 | [] | 1,000.00 | 1,000.00 | 1,000.00 | 1,000.00 | 1,000.00 | 1,000.00 |
| 414 | Centrosomal protein 170 (Fragment) OS=Homo sapier | H0Y2V6 | CEP170 | 145 | 172 kDa | | 1 | 1 | [] | 1,000.00 | 1,000.00 | 1,000.00 | 1,000.00 | 1,000.00 | 1,000.00 |
| 415 | NEDD4 E3 ubiquitin protein ligase OS=Homo sapiens | A0A3B3IUC1 | NEDD4 | 3 | 9 kDa | | 1 | 1 | [] | 1,000.00 | 1,000.00 | 1,000.00 | 1,000.00 | 1,000.00 | 1,000.00 |
| 416 | GTPase activating protein and VPS9 domains 1 OS=H | F8W9S7 (+1) | GAPVD1 | 125 | 157 kDa | | 1 | 1 | [] | 1,000.00 | 1,000.00 | 1,000.00 | 1,000.00 | 1,000.00 | 1,000.00 |
| 417 | Ribosome biogenesis regulatory protein homolog OS=] | Q15050 | RRS1 | 93 | 41 kDa | | 1 | 1 | [] | 1,000.00 | 1,000.00 | 1,000.00 | 1,000.00 | 1,000.00 | 1,000.00 |
| 418 | Protein LLP homolog OS=Homo sapiens OX=9606 G] | Q9BRT6 | LLPH | 43 | 15 kDa | | 1 | 1 | [] | 1,000.00 | 1,000.00 | 1,000.00 | 1,000.00 | 1,000.00 | 1,000.00 |
| 419 | Zinc finger FYVE domain-containing protein 16 OS=H | Q7Z3T8 | ZFYVE16 | 72 | 169 kDa | | 1 | 1 | [] | 1,000.00 | 1,000.00 | 1,000.00 | 1,000.00 | 1,000.00 | 1,000.00 |
| 420 | Suppressor of glucose, autophagy associated 1 (Fragm | H0YDM2 (+1) | SOGA1 | 7 | 142 kDa | | 1 | 1 | [] | 1,000.00 | 1,000.00 | 1,000.00 | 1,000.00 | 1,000.00 | 1,000.00 |
| 421 | Serine/threonine-protein phosphatase 6 catalytic subun | O00743 | PPP6C | 37 | 35 kDa | | 1 | 1 | [] | 1,000.00 | 1,000.00 | 1,000.00 | 1,000.00 | 1,000.00 | 1,000.00 |
| 422 | tRNA (guanine(10)-N2)-methyltransferase homolog O | Q7Z4G4 | TRMT11 | 24 | 53 kDa | | 1 | 1 | [] | 1,000.00 | 1,000.00 | 1,000.00 | 1,000.00 | 1,000.00 | 1,000.00 |
| 423 | DNA-binding protein SMUBP-2 OS=Homo sapiens O | P38935 | IGHMBP2 | 3 | 109 kDa | TRUE | 1 | 1 | [] | 1,000.00 | 1,000.00 | 1,000.00 | 1,000.00 | 1,000.00 | 1,000.00 |
| 424 | Hyaluronan mediated motility receptor OS=Homo sapi | O75330 | HMMR | 12 | 84 kDa | | 1 | 1 | [] | 1,000.00 | 1,000.00 | 1,000.00 | 1,000.00 | 1,000.00 | 1,000.00 |
| 425 | Nucleolar protein interacting with the FHA domain of ] | C9J6C5 (+1) | NIFK | 84 | 20 kDa | | 1 | 1 | [] | 1,000.00 | 1,000.00 | 1,000.00 | 1,000.00 | 1,000.00 | 1,000.00 |
| 426 | Cluster of Ubiquitin carboxyl-terminal hydrolase 15 OS | Q9Y4E8 [2] | USP15 | 182 | 112 kDa | TRUE | 1 | 1 | [] | 1,000.00 | 1,000.00 | 1,000.00 | 1,000.00 | 1,000.00 | 1,000.00 |
| 427 | Ubiquitin-conjugating enzyme E2 R1 OS=Homo sapier | P49427 | CDC34 | 0 | 27 kDa | | 1 | 1 | [] | 1,000.00 | 1,000.00 | 1,000.00 | 1,000.00 | 1,000.00 | 1,000.00 |
| 428 | RING-type E3 ubiquitin transferase OS=Homo sapiens | A0A087WTW0 (+2) | UHRF1 | 89 | 97 kDa | | 1 | 1 | [] | 1,000.00 | 1,000.00 | 1,000.00 | 1,000.00 | 1,000.00 | 1,000.00 |
| 429 | Chromosome 20 open reading frame 129 OS=Homo sa | A0A087WXK8 (+1) | FAM83D | 1 | 68 kDa | | 1 | 1 | [] | 1,000.00 | 1,000.00 | 1,000.00 | 1,000.00 | 1,000.00 | 1,000.00 |
| 430 | Assembly factor for spindle microtubules OS=Homo sa | A0A7P0Z491 (+2) | ASPM | 23 | 418 kDa | | 1 | 1 | [] | 1,000.00 | 1,000.00 | 1,000.00 | 1,000.00 | 1,000.00 | 1,000.00 |
| 431 | Pre-mRNA-processing factor 39 OS=Homo sapiens O | Q86UA1 | PRPF39 | 38 | 78 kDa | | 1 | 1 | [] | 1,000.00 | 1,000.00 | 1,000.00 | 1,000.00 | 1,000.00 | 1,000.00 |
| 432 | Protein PAT1 homolog 1 OS=Homo sapiens OX=960( | Q86TB9 | PATL1 | 25 | 87 kDa | | 1 | 1 | [] | 1,000.00 | 1,000.00 | 1,000.00 | 1,000.00 | 1,000.00 | 1,000.00 |
| 433 | Amyloid beta (A4) protein-binding, family B, member 2 | G5E9Y1 (+2) | APBB2 | 0 | 81 kDa | | 1 | 1 | [] | 1,000.00 | 1,000.00 | 1,000.00 | 1,000.00 | 1,000.00 | 1,000.00 |

| | | | | | | | | | | | | | | |
|---|---|---|---|---|---|---|---|---|---|---|---|---|---|---|
| 434 | Disks large-associated protein 5 OS=Homo sapiens OX | Q15398 | DLGAP5 | 79 | 95 kDa | | 1 | 1 | [] | 1,000.00 | 1,000.00 | 1,000.00 | 1,000.00 | 1,000.00 | 1,000.00 |
| 435 | Zinc finger and BTB domain-containing protein 43 OS | O43298 (+1) | ZBTB43 | 0 | 53 kDa | | 1 | 1 | [] | 1,000.00 | 1,000.00 | 1,000.00 | 1,000.00 | 1,000.00 | 1,000.00 |
| 436 | Probable ATP-dependent RNA helicase DDX31 OS=H | Q9H8H2 (+1) | DDX31 | 40 | 94 kDa | | 1 | 1 | [] | 1,000.00 | 1,000.00 | 1,000.00 | 1,000.00 | 1,000.00 | 1,000.00 |
| 437 | Cluster of Formin like 2 OS=Homo sapiens OX=9606 | C9IZY8 [4] | FMNL2 | 8 | 53 kDa | TRUE | 1 | 1 | [] | 1,000.00 | 1,000.00 | 1,000.00 | 1,000.00 | 1,000.00 | 1,000.00 |
| 438 | WD repeat and coiled-coil-containing protein OS=Hon | Q9H6R7 | WDCP | 6 | 79 kDa | | 1 | 1 | [] | 1,000.00 | 1,000.00 | 1,000.00 | 1,000.00 | 1,000.00 | 1,000.00 |
| 439 | Thymidine kinase (Fragment) OS=Homo sapiens OX= | K7ERJ1 (+3) | TK1 | 14 | 21 kDa | | 1 | 1 | [] | 1,000.00 | 1,000.00 | 1,000.00 | 1,000.00 | 1,000.00 | 1,000.00 |
| 440 | Cluster of Protein bicaudal D homolog 2 OS=Homo sa | Q8TD16 [3] | BICD2 | 20 | 94 kDa | TRUE | 1 | 1 | [] | 1,000.00 | 1,000.00 | 1,000.00 | 1,000.00 | 1,000.00 | 1,000.00 |
| 441 | DNA polymerase epsilon catalytic subunit OS=Homo s | F5H1D6 (+1) | POLE | 24 | 258 kDa | | 1 | 1 | [] | 1,000.00 | 1,000.00 | 1,000.00 | 1,000.00 | 1,000.00 | 1,000.00 |
| 442 | Pleckstrin homology domain-containing family A meml | Q9HAU0 | PLEKHA5 | 65 | 127 kDa | | 1 | 1 | [] | 1,000.00 | 1,000.00 | 1,000.00 | 1,000.00 | 1,000.00 | 1,000.00 |
| 443 | Cluster of Apolipoprotein B mRNA editing enzyme cat | Q6ICH2 [3] | APOBEC3D | 23 | 24 kDa | TRUE | 1 | 1 | [] | 1,000.00 | 1,000.00 | 1,000.00 | 1,000.00 | 1,000.00 | 1,000.00 |
| 444 | Calponin (Fragment) OS=Homo sapiens OX=9606 GN | A0A087X1X5 | CNN2 | 164 | 21 kDa | | 1 | 1 | [] | 1,000.00 | 1,000.00 | 1,000.00 | 1,000.00 | 1,000.00 | 1,000.00 |
| 445 | Thrombospondin 3 OS=Homo sapiens OX=9606 GN= | F5H4Z8 (+1) | THBS3 | 0 | 103 kDa | | 1 | 1 | [] | 1,000.00 | 1,000.00 | 1,000.00 | 1,000.00 | 1,000.00 | 1,000.00 |
| 446 | Cluster of ATP-dependent 6-phosphofructokinase, plat | Q01813 [2] | PFKP | 201 | 86 kDa | TRUE | 1 | 1 | [] | 1,000.00 | 1,000.00 | 1,000.00 | 1,000.00 | 1,000.00 | 1,000.00 |
| 447 | Activating signal cointegrator 1 complex subunit 1 OS | A0A5F9ZH79 (+4) | ASCC1 | 27 | 37 kDa | | 1 | 1 | [] | 1,000.00 | 1,000.00 | 1,000.00 | 1,000.00 | 1,000.00 | 1,000.00 |
| 448 | Cluster of non-specific serine/threonine protein kinase | A0A0A6YYC0 [4] | RPS6KA4 | 2 | 78 kDa | TRUE | 1 | 1 | [] | 1,000.00 | 1,000.00 | 1,000.00 | 1,000.00 | 1,000.00 | 1,000.00 |
| 449 | Ribosomal oxygenase 1 OS=Homo sapiens OX=9606 | Q9H6W3 | RIOX1 | 8 | 71 kDa | | 1 | 1 | [] | 1,000.00 | 1,000.00 | 1,000.00 | 1,000.00 | 1,000.00 | 1,000.00 |
| 450 | LSM family member 14B (Fragment) OS=Homo sapie | Q5TBP9 (+1) | LSM14B | 29 | 27 kDa | | 1 | 1 | [] | 1,000.00 | 1,000.00 | 1,000.00 | 1,000.00 | 1,000.00 | 1,000.00 |
| 451 | SURP and G-patch domain containing 2 OS=Homo sa | M0R2Z9 (+2) | SUGP2 | 136 | 122 kDa | | 1 | 1 | [] | 1,000.00 | 1,000.00 | 1,000.00 | 1,000.00 | 1,000.00 | 1,000.00 |
| 452 | Probable ubiquitin carboxyl-terminal hydrolase FAF-X | Q93008 | USP9X | 177 | 290 kDa | | 1 | 1 | [] | 1,000.00 | 1,000.00 | 1,000.00 | 1,000.00 | 1,000.00 | 1,000.00 |
| 453 | Arginine and glutamate-rich protein 1 OS=Homo sapie | Q9NWB6 | ARGLU1 | 150 | 33 kDa | | 1 | 1 | [] | 1,000.00 | 1,000.00 | 1,000.00 | 1,000.00 | 1,000.00 | 1,000.00 |
| 454 | Microtubule cross-linking factor 1 OS=Homo sapiens | Q9Y4B5 | MTCL1 | 1 | 210 kDa | | 1 | 1 | [] | 1,000.00 | 1,000.00 | 1,000.00 | 1,000.00 | 1,000.00 | 1,000.00 |
| 455 | Condensin complex subunit 2 OS=Homo sapiens OX= | E9PHA2 (+1) | NCAPH | 212 | 82 kDa | | 1 | 1 | [] | 1,000.00 | 1,000.00 | 1,000.00 | 1,000.00 | 1,000.00 | 1,000.00 |
| 456 | Lethal(2) giant larvae protein homolog 1 OS=Homo sa | Q15334 | LLGL1 | 19 | 115 kDa | | 1 | 1 | [] | 1,000.00 | 1,000.00 | 1,000.00 | 1,000.00 | 1,000.00 | 1,000.00 |
| 458 | Cingulin OS=Homo sapiens OX=9606 GN=CGN PE= | Q9P2M7 | CGN | 15 | 137 kDa | | 1 | 1 | [] | 1,000.00 | 1,000.00 | 1,000.00 | 1,000.00 | 1,000.00 | 1,000.00 |
| 459 | Zinc finger protein 629 OS=Homo sapiens OX=9606 | Q9UEG4 | ZNF629 | 23 | 97 kDa | TRUE | 1 | 1 | [] | 1,000.00 | 1,000.00 | 1,000.00 | 1,000.00 | 1,000.00 | 1,000.00 |
| 460 | Period circadian regulator 1 OS=Homo sapiens OX=9 | J3KTM2 (+1) | PER1 | 0 | 88 kDa | | 1 | 1 | [] | 1,000.00 | 1,000.00 | 1,000.00 | 1,000.00 | 1,000.00 | 1,000.00 |
| 461 | RNA polymerase II-associated protein 3 OS=Homo sa | Q9H6T3 | RPAP3 | 114 | 76 kDa | | 1 | 1 | [] | 1,000.00 | 1,000.00 | 1,000.00 | 1,000.00 | 1,000.00 | 1,000.00 |
| 462 | UDP-glucose 6-dehydrogenase OS=Homo sapiens OX | O60701 | UGDH | 19 | 55 kDa | | 1 | 1 | [] | 1,000.00 | 1,000.00 | 1,000.00 | 1,000.00 | 1,000.00 | 1,000.00 |
| 463 | Raf-1 proto-oncogene, serine/threonine kinase (Fragm | H7C155 (+1) | RAF1 | 43 | 47 kDa | | 1 | 1 | [] | 1,000.00 | 1,000.00 | 1,000.00 | 1,000.00 | 1,000.00 | 1,000.00 |
| 464 | Sperm-associated antigen 5 OS=Homo sapiens OX=96 | Q96R06 | SPAG5 | 89 | 134 kDa | TRUE | 1 | 1 | [] | 1,000.00 | 1,000.00 | 1,000.00 | 1,000.00 | 1,000.00 | 1,000.00 |
| 466 | Segment polarity protein dishevelled homolog DVL-1 | O14640 | DVL1 | 85 | 75 kDa | | 1 | 1 | [] | 1,000.00 | 1,000.00 | 1,000.00 | 1,000.00 | 1,000.00 | 1,000.00 |
| 516 | Ribonucleases P/MRP protein subunit POP1 OS=Hom | Q99575 | POP1 | 73 | 115 kDa | | 0.43 | 0.9 | [] | 1.93E+08 | 2.75E+08 | 2.98E+08 | 2.50E+08 | 1.61E+08 | 2.43E+08 |
| 508 | Cluster of UDP-glucose:glycoprotein glucosyltransfera | Q9NYU2 [2] | UGGT1 | 69 | 177 kDa | TRUE | 0.56 | 0.9 | [] | 1.39E+07 | 1.85E+07 | 9686100 | 9931900 | 1.37E+07 | 1.31E+07 |

| | | | | | | | | | | | | | | | |
|---|---|---|---|---|---|---|---|---|---|---|---|---|---|---|---|
| 497 | 60S ribosomal protein L28 OS=Homo sapiens OX=96 | P46779 | RPL28 | 316 | 16 kDa | | 0.59 | 0.9 | [] | 2.12E+08 | 2.20E+08 | 3.54E+08 | 2.32E+08 | 2.06E+08 | 2.61E+08 |
| 485 | Cluster of 40S ribosomal protein S3 OS=Homo sapien | P23396 [2] | RPS3 | 491 | 27 kDa | TRUE | 0.6 | 0.9 | [] | 5.78E+08 | 5.03E+08 | 6.50E+08 | 4.46E+08 | 5.86E+08 | 5.90E+08 |
| 512 | Keratin, type I cytoskeletal 18 OS=Homo sapiens OX= | P05783 | KRT18 | 471 | 48 kDa | TRUE | 0.6 | 0.9 | [] | 1.44E+09 | 1.36E+09 | 1.13E+09 | 1.73E+09 | 7.56E+08 | 8.91E+08 |
| 517 | A-kinase anchoring protein 8 like OS=Homo sapiens C | A0A7P0TAE3 (+1) | AKAP8L | 85 | 68 kDa | | 0.7 | 0.9 | [] | 1.47E+07 | 2.43E+07 | 1.53E+07 | 4297100 | 1.95E+07 | 2.25E+07 |
| 498 | Microtubule associated protein 1A OS=Homo sapiens | E9PGC8 (+1) | MAP1A | 113 | 331 kDa | TRUE | 0.71 | 0.9 | [] | 1.28E+07 | 1.74E+07 | 2.25E+07 | 2.23E+07 | 8797500 | 1.59E+07 |
| 510 | Delta(14)-sterol reductase LBR OS=Homo sapiens O | A0A494C1L1 (+1) | LBR | 241 | 47 kDa | | 0.71 | 0.9 | [] | 2.39E+07 | 9126400 | 1.16E+07 | 1.69E+07 | 1.13E+07 | 1.04E+07 |
| 476 | HCG1984214, isoform CRA_a OS=Homo sapiens OX | I3L0E3 (+1) | hCG_1984214 | 64 | 26 kDa | | 0.74 | 0.9 | [] | 3.68E+07 | 5.15E+07 | 4.01E+07 | 3.66E+07 | 3.61E+07 | 4.92E+07 |
| 502 | RNA-splicing ligase RtcB homolog OS=Homo sapiens | Q9Y3I0 | RTCB | 290 | 55 kDa | | 0.76 | 0.9 | [] | 2.53E+07 | 2.83E+07 | 1.66E+07 | 8058800 | 3.51E+07 | 1.87E+07 |
| 509 | RNA cytosine C(5)-methyltransferase NSUN2 OS=Ho | Q08J23 | NSUN2 | 254 | 86 kDa | | 0.77 | 0.9 | [] | 3.48E+07 | 3.10E+07 | 1.14E+07 | 6295000 | 3.62E+07 | 2.43E+07 |
| 501 | FMR1-interacting protein NUFIP2 OS=Homo sapiens | Q7Z417 | NUFIP2 | 223 | 76 kDa | | 0.78 | 0.9 | [] | 6.87E+07 | 5.98E+07 | 1.70E+08 | 1.11E+08 | 5.22E+07 | 9.98E+07 |
| 489 | Glypican-1 OS=Homo sapiens OX=9606 GN=GPC1 P | P35052 | GPC1 | 9 | 62 kDa | | 0.79 | 0.9 | [] | 1.93E+08 | 1.88E+08 | 1.73E+08 | 8.22E+07 | 2.19E+08 | 2.14E+08 |
| 514 | Large subunit GTPase 1 homolog OS=Homo sapiens ( | Q9H089 | LSG1 | 85 | 75 kDa | | 0.79 | 0.9 | [] | 1.64E+07 | 7.72E+07 | 2.61E+07 | 2.02E+07 | 3.77E+07 | 4.47E+07 |
| 518 | 40S ribosomal protein S28 OS=Homo sapiens OX=96 | P62857 | RPS28 | 404 | 8 kDa | | 0.79 | 0.9 | [] | 3155100 | 4294400 | 1191700 | 1,000.00 | 3751100 | 3597100 |
| 495 | 40S ribosomal protein S23 OS=Homo sapiens OX=96 | P62266 | RPS23 | 413 | 16 kDa | | 0.8 | 0.9 | [] | 8.39E+08 | 4.88E+08 | 2.77E+08 | 3.31E+08 | 6.16E+08 | 5.07E+08 |
| 511 | Cluster of RNA-binding protein 47 OS=Homo sapiens | A0AV96 [2] | RBM47 | 2 | 64 kDa | TRUE | 0.8 | 0.9 | [] | 1.80E+08 | 6.60E+07 | 3.82E+07 | 5.57E+07 | 1.16E+08 | 7.42E+07 |
| 477 | Pyruvate carboxylase, mitochondrial OS=Homo sapien | P11498 | PC | 306 | 130 kDa | | 0.81 | 0.9 | [] | 4.34E+09 | 4.61E+09 | 3.05E+09 | 2.55E+09 | 4.57E+09 | 4.27E+09 |
| 504 | Phosphate carrier protein, mitochondrial OS=Homo sa | Q00325 | SLC25A3 | 362 | 40 kDa | | 0.81 | 0.9 | [] | 4.84E+07 | 8.06E+07 | 2.22E+08 | 1.32E+08 | 9.88E+07 | 7.73E+07 |
| 507 | Calumenin OS=Homo sapiens OX=9606 GN=CALU P | O43852 | CALU | 147 | 37 kDa | | 0.81 | 0.9 | [] | 3.83E+07 | 5.49E+07 | 6.12E+07 | 1,000.00 | 8.48E+07 | 5.01E+07 |
| 482 | X-ray repair cross-complementing protein 6 OS=Hom | P12956 | XRCC6 | 388 | 70 kDa | | 0.82 | 0.9 | [] | 1.10E+08 | 7.59E+07 | 4.47E+07 | 6.90E+07 | 7.65E+07 | 7.13E+07 |
| 491 | methylcrotonoyl-CoA carboxylase OS=Homo sapiens | A0A804HKE8 (+1) | MCCC2 | 251 | 58 kDa | | 0.82 | 0.9 | [] | 8477600 | 1.39E+07 | 1.59E+07 | 4862100 | 1.45E+07 | 1.59E+07 |
| 500 | Mitochondrial ribosomal protein S34 OS=Homo sapie | C9JJ19 (+1) | MRPS34 | 58 | 26 kDa | | 0.82 | 0.9 | [] | 9.00E+07 | 6.87E+07 | 4.49E+07 | 5836400 | 1.06E+08 | 6.84E+07 |
| 496 | 60S ribosomal protein L5 OS=Homo sapiens OX=960 | P46777 | RPL5 | 395 | 34 kDa | | 0.83 | 0.9 | [] | 4.80E+07 | 2.41E+08 | 1.36E+08 | 5.98E+07 | 1.51E+08 | 1.69E+08 |
| 506 | 28S ribosomal protein S16, mitochondrial OS=Homo s | Q9Y3D3 | MRPS16 | 26 | 15 kDa | | 0.83 | 0.9 | [] | 1.47E+07 | 1.78E+07 | 3.41E+07 | 1,000.00 | 2.08E+07 | 3.74E+07 |
| 513 | Cluster of DNA topoisomerase 1 OS=Homo sapiens O | P11387 [2] | TOP1 | 285 | 91 kDa | TRUE | 0.85 | 0.9 | [] | 3524100 | 8.14E+07 | 1.68E+08 | 4257800 | 8.56E+07 | 1.27E+08 |
| 488 | Ubiquitin carboxyl-terminal hydrolase 10 OS=Homo sa | Q14694 | USP10 | 114 | 87 kDa | | 0.86 | 0.9 | [] | 7.21E+07 | 2.85E+07 | 6.03E+07 | 2.62E+07 | 4.57E+07 | 7.80E+07 |
| 483 | 60S ribosomal protein L15 OS=Homo sapiens OX=96 | P61313 | RPL15 | 365 | 24 kDa | | 0.87 | 0.9 | [] | 1.59E+08 | 2.94E+08 | 1.71E+08 | 8.19E+07 | 2.70E+08 | 2.34E+08 |
| 484 | Synaptotagmin binding cytoplasmic RNA interacting p | A0A7I2YQN2 | SYNCRIP | 459 | 63 kDa | TRUE | 0.87 | 0.9 | [] | 2.03E+08 | 2.56E+08 | 4.51E+08 | 1.67E+08 | 2.77E+08 | 4.11E+08 |
| 499 | Probable ATP-dependent RNA helicase DDX47 OS=H | Q9H0S4 | DDX47 | 96 | 51 kDa | | 0.87 | 0.9 | [] | 4289000 | 1.25E+07 | 5111500 | 1,000.00 | 1.29E+07 | 6611000 |
| 492 | Protein HEXIM1 OS=Homo sapiens OX=9606 GN=H | O94992 | HEXIM1 | 77 | 41 kDa | | 0.89 | 0.9 | [] | 1.87E+07 | 3.88E+07 | 8.27E+07 | 9053900 | 6.88E+07 | 5.04E+07 |
| 515 | SRSF protein kinase 1 (Fragment) OS=Homo sapiens | H3BLV9 (+1) | SRPK1 | 157 | 76 kDa | TRUE | 0.89 | 0.9 | [] | 96928 | 1.13E+07 | 1.03E+07 | 1.78E+07 | 794590 | 1,000.00 |
| 503 | Cluster of PATJ crumbs cell polarity complex compon | A0A2R8Y549 [4] | PATJ | 8 | 205 kDa | TRUE | 0.9 | 0.9 | [] | 2.37E+07 | 980450 | 6965800 | 2.15E+07 | 5905600 | 422710 |
| 505 | Cluster of Profilin OS=Homo sapiens OX=9606 GN=P | C9J712 [4] | PFN2 | 186 | 10 kDa | TRUE | 0.91 | 0.9 | [] | 1091800 | 1,000.00 | 3.33E+07 | 1,000.00 | 1.47E+07 | 1.54E+07 |
| 486 | Cluster of Spermatogenesis associated serine rich 2 lik | F8W6C2 [3] | SPATS2L | 34 | 30 kDa | TRUE | 0.92 | 0.9 | [] | 4422700 | 6677800 | 3631500 | 528670 | 1.01E+07 | 3182800 |

| | | | | | | | | | | | | | | | |
|---|---|---|---|---|---|---|---|---|---|---|---|---|---|---|---|
| 494 | Poly(rC)-binding protein 1 OS=Homo sapiens OX=96( | Q15365 | PCBP1 | 472 | 37 kDa | TRUE | 0.92 | 0.9 | [] | 1.65E+07 | 1.32E+07 | 3.29E+07 | 5.02E+07 | 1,000.00 | 6986900 |
| 479 | Heterogeneous nuclear ribonucleoprotein D OS=Homo | D6RAF8 (+3) | HNRNPD | 489 | 23 kDa | TRUE | 0.94 | 0.9 | [] | 4765300 | 2.55E+07 | 5002600 | 1.59E+07 | 1.64E+07 | 1047000 |
| 478 | 28S ribosomal protein S2, mitochondrial OS=Homo sa | Q9Y399 | MRPS2 | 65 | 33 kDa | | 0.95 | 0.9 | [] | 1,000.00 | 6588000 | 3910800 | 1,000.00 | 7333100 | 2607100 |
| 487 | Cluster of Probable ATP-dependent RNA helicase DD | Q9Y2R4 [2] | DDX52 | 101 | 67 kDa | TRUE | 0.96 | 0.9 | [] | 6354200 | 1603300 | 1,000.00 | 1,000.00 | 7435600 | 1,000.00 |
| 490 | Heterogeneous nuclear ribonucleoprotein A2/B1 OS=I | A0A024RA28 (+7) | HNRNPA2B1 | 536 | 31 kDa | | 0.96 | 0.9 | [] | 1,000.00 | 1,000.00 | 1.21E+07 | 1.11E+07 | 1,000.00 | 1,000.00 |
| 480 | Cluster of NFKB activating protein (Fragment) OS=H | A0A494C050 [3] | NKAP | 125 | 47 kDa | TRUE | 0.97 | 0.9 | [] | 1,000.00 | 1,000.00 | 1473700 | 1,000.00 | 1,000.00 | 1391600 |
| 549 | 28S ribosomal protein S14, mitochondrial OS=Homo s | O60783 | MRPS14 | 52 | 15 kDa | | 0.0006 | 0.8 | [] | 9.07E+07 | 9.62E+07 | 9.04E+07 | 6.70E+07 | 7.16E+07 | 7.03E+07 |
| 530 | Heterogeneous nuclear ribonucleoprotein U OS=Homo | Q00839 | HNRNPU | 578 | 91 kDa | | 0.18 | 0.8 | [] | 1.71E+09 | 1.80E+09 | 2.14E+09 | 1.85E+09 | 1.27E+09 | 1.51E+09 |
| 535 | Plectin OS=Homo sapiens OX=9606 GN=PLEC PE=1 | Q15149 | PLEC | 233 | 532 kDa | TRUE | 0.3 | 0.8 | [] | 4.88E+09 | 6.05E+09 | 5.45E+09 | 2.66E+09 | 5.51E+09 | 4.91E+09 |
| 546 | Cluster of Sulfide:quinone oxidoreductase, mitochondr | Q9Y6N5 [2] | SQOR | 2 | 50 kDa | TRUE | 0.43 | 0.8 | [] | 1.03E+08 | 5.10E+07 | 5.14E+07 | 6.66E+07 | 4.69E+07 | 4.22E+07 |
| 524 | Spliceosome associated factor 3, U4/U6 recycling prot | A0A499FI31 (+1) | SART3 | 198 | 112 kDa | | 0.46 | 0.8 | [] | 9.93E+07 | 1.54E+08 | 1.84E+08 | 1.51E+08 | 8.72E+07 | 1.24E+08 |
| 542 | Hexokinase HKDC1 OS=Homo sapiens OX=9606 GN | Q2TB90 | HKDC1 | 36 | 103 kDa | TRUE | 0.51 | 0.8 | [] | 1.90E+07 | 6979800 | 2.21E+07 | 1.22E+07 | 1.63E+07 | 8537400 |
| 529 | Cluster of Heterogeneous nuclear ribonucleoprotein M | A0A087X0X3 [3] | HNRNPM | 520 | 78 kDa | TRUE | 0.55 | 0.8 | [] | 7.18E+07 | 1.34E+08 | 1.56E+08 | 1.41E+08 | 8.03E+07 | 7.56E+07 |
| 538 | Cluster of Probable ATP-dependent RNA helicase DD | J3KTA4 [3] | DDX5 | 528 | 69 kDa | TRUE | 0.57 | 0.8 | [] | 3.42E+08 | 9.58E+08 | 4.87E+08 | 3.29E+08 | 5.71E+08 | 5.18E+08 |
| 539 | 39S ribosomal protein L44, mitochondrial OS=Homo s | Q9H9J2 | MRPL44 | 59 | 38 kDa | | 0.59 | 0.8 | [] | 4.31E+07 | 5.99E+07 | 6.01E+07 | 8943100 | 7.34E+07 | 4.66E+07 |
| 548 | Beta-2-syntrophin OS=Homo sapiens OX=9606 GN=S | Q13425 | SNTB2 | 26 | 58 kDa | TRUE | 0.59 | 0.8 | [] | 2.18E+07 | 1.01E+07 | 3.72E+07 | 2.54E+07 | 6100600 | 2.06E+07 |
| 537 | Nitric oxide-associated protein 1 OS=Homo sapiens O | Q8NC60 | NOA1 | 3 | 78 kDa | | 0.6 | 0.8 | [] | 2.20E+07 | 1.67E+07 | 1.02E+07 | 1.32E+07 | 2.12E+07 | 4524300 |
| 536 | Ankycorbin OS=Homo sapiens OX=9606 GN=RAI14 | Q9P0K7 | RAI14 | 121 | 110 kDa | | 0.64 | 0.8 | [] | 7.55E+07 | 6.05E+07 | 1.41E+08 | 1.91E+07 | 1.10E+08 | 9.08E+07 |
| 550 | Serine and arginine rich splicing factor 3 OS=Homo sa | A0A087X2D0 (+1) | SRSF3 | 392 | 10 kDa | | 0.65 | 0.8 | [] | 2.47E+07 | 1.54E+07 | 7487400 | 1,000.00 | 2.23E+07 | 1.34E+07 |
| 525 | Dimethyladenosine transferase 1, mitochondrial OS=H | Q8WVM0 | TFB1M | 8 | 40 kDa | | 0.66 | 0.8 | [] | 3.21E+07 | 2.16E+07 | 7030700 | 1.97E+07 | 1.49E+07 | 1.56E+07 |
| 531 | 39S ribosomal protein L16, mitochondrial OS=Homo s | Q9NX20 | MRPL16 | 57 | 28 kDa | | 0.66 | 0.8 | [] | 2.55E+07 | 3.31E+07 | 2.82E+07 | 1561300 | 2.90E+07 | 3.99E+07 |
| 547 | Replication factor C subunit 1 OS=Homo sapiens OX= | P35251 | RFC1 | 129 | 128 kDa | | 0.67 | 0.8 | [] | 2.35E+07 | 4.21E+07 | 1.09E+08 | 7.82E+07 | 3.00E+07 | 2.36E+07 |
| 533 | 39S ribosomal protein L20, mitochondrial OS=Homo s | Q9BYC9 | MRPL20 | 23 | 17 kDa | | 0.69 | 0.8 | [] | 1.21E+07 | 2.13E+07 | 2.41E+07 | 1.99E+07 | 1,000.00 | 2.63E+07 |
| 545 | Small nuclear ribonucleoprotein D1 polypeptide OS=H | J3QLI9 (+1) | SNRPD1 | 356 | 8 kDa | | 0.74 | 0.8 | [] | 5.15E+07 | 1.20E+07 | 3398900 | 1.02E+07 | 2.22E+07 | 1.82E+07 |
| 519 | Heterogeneous nuclear ribonucleoprotein D like OS=H | A0A087WUK2 (+1) | HNRNPDL | 369 | 40 kDa | TRUE | 0.75 | 0.8 | [] | 4765300 | 1.27E+07 | 1.57E+07 | 3410100 | 8201200 | 1.65E+07 |
| 528 | Poly [ADP-ribose] polymerase 1 OS=Homo sapiens O | P09874 | PARP1 | 438 | 113 kDa | | 0.76 | 0.8 | [] | 1.43E+07 | 7.05E+07 | 1.45E+08 | 5.08E+07 | 9.98E+07 | 3.75E+07 |
| 520 | Speckle targeted PIP5K1A-regulated poly(A) polymer | F5H0R1 (+1) | TUT1 | 0 | 98 kDa | | 0.77 | 0.8 | [] | 6832300 | 1.49E+07 | 6538500 | 1,000.00 | 1.31E+07 | 1.06E+07 |
| 527 | mRNA cap guanine-N7 methyltransferase OS=Homo s | O43148 | RNMT | 81 | 55 kDa | | 0.82 | 0.8 | [] | 4214700 | 3517600 | 1.37E+07 | 1,000.00 | 1.35E+07 | 4154500 |
| 534 | General transcription factor 3C polypeptide 2 OS=Hor | Q8WUA4 | GTF3C2 | 23 | 101 kDa | | 0.82 | 0.8 | [] | 7874400 | 5235400 | 7960000 | 1.69E+07 | 1,000.00 | 1,000.00 |
| 541 | Myosin X OS=Homo sapiens OX=9606 GN=MYO10 | A0A0A0MQX1 (+1 | MYO10 | 3 | 239 kDa | | 0.82 | 0.8 | [] | 2.56E+07 | 1,000.00 | 2736900 | 3998200 | 1.19E+07 | 6157500 |
| 543 | 39S ribosomal protein L32, mitochondrial OS=Homo s | Q9BYC8 | MRPL32 | 31 | 21 kDa | | 0.84 | 0.8 | [] | 2.26E+07 | 1,000.00 | 1,000.00 | 1,000.00 | 8589200 | 8669700 |
| 540 | Putative RNA-binding protein Luc7-like 2 OS=Homo s | Q9Y383 | LUC7L2 | 332 | 47 kDa | | 0.85 | 0.8 | [] | 1,000.00 | 5317600 | 1.93E+07 | 1,000.00 | 1.94E+07 | 1,000.00 |
| 544 | Coatomer subunit beta OS=Homo sapiens OX=9606 C | P53618 | COPB1 | 278 | 107 kDa | | 0.85 | 0.8 | [] | 1,000.00 | 7978900 | 1,000.00 | 5519700 | 573250 | 1,000.00 |

| | | | | | | | | | | | | | | | |
|---|---|---|---|---|---|---|---|---|---|---|---|---|---|---|---|
| 532 | MAP7 domain-containing protein 3 OS=Homo sapiens | Q8IWC1 | MAP7D3 | 129 | 98 kDa | | 0.86 | 0.8 | [] | 1,000.00 | 1763100 | 5663700 | 1,000.00 | 1,000.00 | 6005000 |
| 523 | E3 ubiquitin-protein ligase RNF14 OS=Homo sapiens | Q9UBS8 | RNF14 | 0 | 54 kDa | | 0.87 | 0.8 | [] | 793510 | 1575400 | 1,000.00 | 1,000.00 | 1960400 | 1,000.00 |
| 522 | Ubiquitin carboxyl-terminal hydrolase 5 OS=Homo sap | P45974 | USP5 | 148 | 96 kDa | | 0.88 | 0.8 | [] | 1,000.00 | 1,000.00 | 6927000 | 2479000 | 1,000.00 | 3269000 |
| 521 | Zinc finger and BTB domain-containing protein 21 OS | Q9ULJ3 | ZBTB21 | 95 | 119 kDa | | 0.89 | 0.8 | [] | 2850700 | 1,000.00 | 1.83E+07 | 1.77E+07 | 1,000.00 | 1,000.00 |
| 526 | ATP-dependent DNA/RNA helicase DHX36 OS=Hom | Q9H2U1 | DHX36 | 78 | 115 kDa | TRUE | 0.9 | 0.8 | [] | 1,000.00 | 8046500 | 1,000.00 | 1,000.00 | 6619400 | 1,000.00 |
| 559 | Mitochondrial ribosomal protein S7 OS=Homo sapiens | J3QLS3 (+1) | MRPS7 | 76 | 32 kDa | | 0.27 | 0.7 | [] | 1.38E+08 | 7.31E+07 | 8.63E+07 | 5.83E+07 | 8.39E+07 | 7.44E+07 |
| 567 | Mitochondrial import inner membrane translocase subu | O14925 | TIMM23 | 51 | 22 kDa | | 0.29 | 0.7 | [] | 9138300 | 1.35E+07 | 1.74E+07 | 8283000 | 1.35E+07 | 6418600 |
| 574 | Cluster of Double-stranded RNA-binding protein Stau | Q9NUL3 [2] | STAU2 | 121 | 63 kDa | TRUE | 0.31 | 0.7 | [] | 8.71E+07 | 1.21E+08 | 2.11E+08 | 1.12E+08 | 1.06E+08 | 6.28E+07 |
| 560 | RNA helicase OS=Homo sapiens OX=9606 GN=DDX | A0A1X7SBZ2 (+2) | DDX17 | 523 | 80 kDa | TRUE | 0.4 | 0.7 | [] | 2.62E+08 | 7.16E+08 | 4.22E+08 | 2.77E+08 | 3.83E+08 | 3.58E+08 |
| 551 | Cluster of ATP-dependent RNA helicase DHX15 OS= | O43143 [5] | DHX15 | 432 | 91 kDa | TRUE | 0.44 | 0.7 | [] | 2.72E+08 | 9.53E+07 | 1.86E+08 | 1.07E+08 | 1.39E+08 | 1.68E+08 |
| 572 | THAP domain-containing protein 11 OS=Homo sapier | Q96EK4 | THAP11 | 2 | 34 kDa | | 0.46 | 0.7 | [] | 1039200 | 1446100 | 1106200 | 1,000.00 | 966490 | 1502200 |
| 566 | Cluster of Transitional endoplasmic reticulum ATPase | A0A7P0TAW3 [3] | VCP | 314 | 86 kDa | TRUE | 0.47 | 0.7 | [] | 1.25E+09 | 9.06E+08 | 5.68E+08 | 1.76E+08 | 1.10E+09 | 6.48E+08 |
| 568 | Serine/threonine-protein phosphatase PGAM5, mitoch | Q96HS1 | PGAM5 | 223 | 32 kDa | | 0.47 | 0.7 | [] | 1.17E+07 | 2.68E+07 | 4.65E+07 | 2.17E+07 | 1.22E+07 | 2.55E+07 |
| 571 | Nuclear factor 1 OS=Homo sapiens OX=9606 GN=NI | A0A286YEX4 (+1) | NFIC | 59 | 58 kDa | TRUE | 0.55 | 0.7 | [] | 2.51E+07 | 7988400 | 7612400 | 5975900 | 1.50E+07 | 7177300 |
| 569 | Signal recognition particle subunit SRP68 OS=Homo s | Q9UHB9 | SRP68 | 192 | 71 kDa | | 0.58 | 0.7 | [] | 1.48E+08 | 5.15E+07 | 5.67E+07 | 1.09E+07 | 1.11E+08 | 5.62E+07 |
| 573 | X-ray repair cross-complementing protein 5 OS=Homo | P13010 | XRCC5 | 428 | 83 kDa | | 0.6 | 0.7 | [] | 1.28E+08 | 4.82E+07 | 2.53E+07 | 1.82E+07 | 8.76E+07 | 3.13E+07 |
| 579 | Rac GTPase-activating protein 1 OS=Homo sapiens O | Q9H0H5 | RACGAP1 | 72 | 71 kDa | | 0.6 | 0.7 | [] | 1,000.00 | 2.20E+07 | 5.11E+07 | 1.62E+07 | 1.23E+07 | 1.94E+07 |
| 563 | Cluster of Protein NipSnap homolog 1 OS=Homo sapi | Q9BPW8 [2] | NIPSNAP1 | 40 | 33 kDa | TRUE | 0.61 | 0.7 | [] | 6129700 | 1.33E+07 | 4778100 | 1.20E+07 | 3194100 | 2212000 |
| 554 | RNA cytidine acetyltransferase OS=Homo sapiens OX | Q9H0A0 | NAT10 | 196 | 116 kDa | | 0.66 | 0.7 | [] | 3370400 | 2371000 | 1552000 | 1,000.00 | 1275200 | 4131500 |
| 552 | Mitochondrial ribosomal protein L3 OS=Homo sapiens | E7ETU7 (+2) | MRPL3 | 39 | 42 kDa | | 0.67 | 0.7 | [] | 8.05E+07 | 8060800 | 5.02E+07 | 4276000 | 4.85E+07 | 5.09E+07 |
| 562 | BBX high mobility group box domain containing OS=I | C9JA69 (+1) | BBX | 69 | 99 kDa | | 0.67 | 0.7 | [] | 8397300 | 2742000 | 8601900 | 1.13E+07 | 3035200 | 1,000.00 |
| 565 | Syndecan-1 OS=Homo sapiens OX=9606 GN=SDC1 I | P18827 | SDC1 | 2 | 32 kDa | | 0.67 | 0.7 | [] | 1.33E+07 | 5.84E+07 | 4.96E+07 | 1,000.00 | 6.92E+07 | 1.77E+07 |
| 561 | pre-rRNA 2'-O-ribose RNA methyltransferase FTSJ3 ( | Q8IY81 | FTSJ3 | 184 | 97 kDa | | 0.69 | 0.7 | [] | 1,000.00 | 5501700 | 7425600 | 1,000.00 | 4554500 | 4827700 |
| 575 | Far upstream element-binding protein 3 OS=Homo sap | Q96I24 | FUBP3 | 212 | 62 kDa | TRUE | 0.7 | 0.7 | [] | 3201500 | 1722300 | 1.86E+07 | 3902800 | 1.17E+07 | 1,000.00 |
| 576 | WW domain-binding protein 4 OS=Homo sapiens OX= | O75554 | WBP4 | 17 | 43 kDa | | 0.7 | 0.7 | [] | 1.49E+07 | 3.79E+07 | 2161900 | 3.46E+07 | 1,000.00 | 1546400 |
| 570 | Cluster of Myoferlin OS=Homo sapiens OX=9606 GN | Q9NZM1 [2] | MYOF | 29 | 235 kDa | TRUE | 0.71 | 0.7 | [] | 3027200 | 2866500 | 1,000.00 | 3660300 | 421910 | 1,000.00 |
| 578 | Translation machinery-associated protein 16 (Fragmen | H0Y9X1 (+1) | TMA16 | 85 | 28 kDa | | 0.72 | 0.7 | [] | 1.52E+07 | 5019900 | 1,000.00 | 1,000.00 | 1.21E+07 | 1164200 |
| 558 | 39S ribosomal protein L1, mitochondrial OS=Homo sa | Q9BYD6 | MRPL1 | 43 | 37 kDa | | 0.76 | 0.7 | [] | 1,000.00 | 1760000 | 7370400 | 2562300 | 1,000.00 | 4100300 |
| 553 | DNA-3-methyladenine glycosylase II (Fragment) OS=I | A2IDA3 (+1) | MPG | 37 | 27 kDa | | 0.78 | 0.7 | [] | 4522600 | 3414500 | 187100 | 302060 | 5745500 | 1,000.00 |
| 556 | Leucine-rich repeat flightless-interacting protein 1 OS= | Q32MZ4 | LRRFIP1 | 158 | 89 kDa | | 0.78 | 0.7 | [] | 1,000.00 | 5.96E+08 | 3.31E+08 | 1,000.00 | 6.78E+08 | 1,000.00 |
| 577 | Glutamate-rich WD repeat-containing protein 1 OS=H | Q9BQ67 | GRWD1 | 160 | 49 kDa | | 0.79 | 0.7 | [] | 1,000.00 | 3719000 | 1,000.00 | 1,000.00 | 2446700 | 1,000.00 |
| 557 | CDK5 regulatory subunit-associated protein 3 OS=Ho | Q96JB5 | CDK5RAP3 | 24 | 57 kDa | | 0.82 | 0.7 | [] | 2230000 | 5.43E+07 | 1503800 | 1,000.00 | 3.85E+07 | 3909800 |
| 555 | U4/U6.U5 tri-snRNP-associated protein 1 OS=Homo s | O43290 | SART1 | 233 | 90 kDa | | 0.84 | 0.7 | [] | 1,000.00 | 1,000.00 | 2.56E+07 | 1.89E+07 | 1,000.00 | 1,000.00 |

| | | | | | | | | | | | | | | | |
|---|---|---|---|---|---|---|---|---|---|---|---|---|---|---|---|
| 595 | Cluster of Heat shock cognate 71 kDa protein OS=Ho | P11142 [8] | HSPA8 | 703 | 71 kDa | TRUE | 0.1 | 0.6 | [] | 2.55E+08 | 4.40E+08 | 4.37E+08 | 2.30E+08 | 1.29E+08 | 2.85E+08 |
| 588 | Cytochrome b-c1 complex subunit 2, mitochondrial O | P22695 | UQCRC2 | 190 | 48 kDa | | 0.24 | 0.6 | [] | 2.28E+07 | 9994000 | 2.98E+07 | 1.49E+07 | 1.04E+07 | 1.26E+07 |
| 602 | Ribosomal protein L37a OS=Homo sapiens OX=9606 | C9J4Z3 (+1) | RPL37A | 200 | 8 kDa | | 0.25 | 0.6 | [] | 8.96E+07 | 2.93E+07 | 4.87E+07 | 3.38E+07 | 2.11E+07 | 3.81E+07 |
| 599 | Leucine-rich repeat-containing protein 59 OS=Homo s | Q96AG4 | LRRC59 | 175 | 35 kDa | | 0.27 | 0.6 | [] | 8558200 | 1.04E+07 | 1.14E+07 | 390220 | 1.20E+07 | 4611100 |
| 603 | Plasminogen activator inhibitor 1 RNA-binding protein | Q8NC51 | SERBP1 | 468 | 45 kDa | | 0.31 | 0.6 | [] | 5.53E+07 | 3.76E+07 | 5.94E+07 | 6.32E+07 | 1.90E+07 | 2022300 |
| 596 | Fibroblast growth factor OS=Homo sapiens OX=9606 | A0A0A0MQV6 (+1 | FGF2 | 14 | 31 kDa | | 0.33 | 0.6 | [] | 1.10E+07 | 1.39E+07 | 2.34E+07 | 1.79E+07 | 9488600 | 1,000.00 |
| 604 | Cluster of Kinesin-like protein OS=Homo sapiens OX= | F5H3M2 [6] | KIFC3 | 0 | 81 kDa | TRUE | 0.38 | 0.6 | [] | 2797500 | 806860 | 2055700 | 2171800 | 952380 | 1,000.00 |
| 598 | Voltage dependent anion channel 2 (Fragment) OS=H | A0A0A0MR02 (+1) | VDAC2 | 256 | 30 kDa | | 0.4 | 0.6 | [] | 7828200 | 5.83E+07 | 7.30E+07 | 3.06E+07 | 8682700 | 3.89E+07 |
| 600 | Cluster of Replication factor C subunit 2 OS=Homo sa | P35250 [2] | RFC2 | 128 | 39 kDa | TRUE | 0.51 | 0.6 | [] | 1,000.00 | 2.49E+07 | 5.63E+07 | 1.00E+07 | 1.42E+07 | 2.13E+07 |
| 582 | General transcription factor IIIC subunit 3 OS=Homo | A0A494C1S7 (+1) | GTF3C3 | 113 | 71 kDa | | 0.58 | 0.6 | [] | 7103100 | 1.70E+07 | 8175200 | 1.81E+07 | 739730 | 1582800 |
| 586 | Proline rich coiled-coil 2C OS=Homo sapiens OX=96 | E7EPN9 (+1) | PRRC2C | 223 | 309 kDa | TRUE | 0.58 | 0.6 | [] | 4.69E+07 | 5007700 | 5.10E+07 | 5.38E+07 | 2182700 | 7320100 |
| 601 | DNA-directed RNA polymerase subunit OS=Homo sa | A0A6Q8PGB0 (+1) | POLR2A | 175 | 217 kDa | | 0.58 | 0.6 | [] | 6603900 | 1409300 | 1.24E+07 | 1.14E+07 | 1,000.00 | 1,000.00 |
| 585 | Phenylalanyl-tRNA synthetase OS=Homo sapiens OX= | A0A3B3ITR6 (+1) | FARS2 | 2 | 50 kDa | | 0.59 | 0.6 | [] | 5053000 | 2923500 | 6816500 | 9114400 | 1,000.00 | 1,000.00 |
| 587 | Elongator complex protein 1 OS=Homo sapiens OX=9 | A0A6Q8PHC9 (+1) | ELP1 | 63 | 152 kDa | | 0.6 | 0.6 | [] | 2755900 | 2483300 | 938500 | 1,000.00 | 1,000.00 | 3792900 |
| 590 | Cluster of Tubulin alpha-1B chain OS=Homo sapiens | P68363 [3] | TUBA1B | 694 | 50 kDa | TRUE | 0.6 | 0.6 | [] | 1.52E+09 | 2.70E+08 | 1.01E+08 | 5.31E+08 | 3.29E+08 | 2.68E+08 |
| 594 | 60 kDa heat shock protein, mitochondrial OS=Homo s | A0A7I2V2X6 (+2) | HSPD1 | 520 | 58 kDa | | 0.6 | 0.6 | [] | 6135200 | 3445100 | 1,000.00 | 5054700 | 1,000.00 | 449780 |
| 597 | >sp\|O75746\|CMC1_HUMAN Calcium-binding mitoch | sp\|O75746\|CMC1_ | SLC25A12 | 66 | ? | TRUE | 0.6 | 0.6 | [] | 1,000.00 | 2817000 | 4180700 | 3954000 | 1,000.00 | 1,000.00 |
| 592 | TRIO and F-actin-binding protein OS=Homo sapiens | Q9H2D6 | TRIOBP | 9 | 261 kDa | TRUE | 0.63 | 0.6 | [] | 1,000.00 | 3108400 | 4597700 | 4559700 | 1,000.00 | 1,000.00 |
| 593 | Aurora kinase A-interacting protein OS=Homo sapiens | Q9NWT8 | AURKAIP1 | 7 | 22 kDa | | 0.63 | 0.6 | [] | 1,000.00 | 8912700 | 4.05E+07 | 1,000.00 | 9954800 | 1.86E+07 |
| 581 | Protein PRRC2A OS=Homo sapiens OX=9606 GN=P | P48634 | PRRC2A | 212 | 229 kDa | TRUE | 0.64 | 0.6 | [] | 1.43E+07 | 2353500 | 2.25E+07 | 2.29E+07 | 2022100 | 1,000.00 |
| 605 | Multiple myeloma tumor-associated protein 2 OS=Ho | Q9BU76 | MMTAG2 | 230 | 29 kDa | | 0.64 | 0.6 | [] | 1,000.00 | 2289400 | 1.73E+07 | 1,000.00 | 4259500 | 6523100 |
| 589 | DNA ligase 3 OS=Homo sapiens OX=9606 GN=LIG3 | P49916 | LIG3 | 101 | 113 kDa | | 0.67 | 0.6 | [] | 1022100 | 6563500 | 1,000.00 | 1,000.00 | 3002300 | 1546300 |
| 591 | Serine/threonine-protein phosphatase 2A catalytic sub | P67775 | PPP2CA | 193 | 36 kDa | | 0.74 | 0.6 | [] | 8762500 | 1,000.00 | 1,000.00 | 1,000.00 | 5200900 | 1,000.00 |
| 583 | Structural maintenance of chromosomes protein OS=H | G8JLG1 (+1) | SMC1A | 248 | 141 kDa | | 0.77 | 0.6 | [] | 1,000.00 | 1,000.00 | 2199100 | 1383000 | 1,000.00 | 1,000.00 |
| 580 | Protein LTV1 homolog OS=Homo sapiens OX=9606 | Q96GA3 | LTV1 | 137 | 55 kDa | | 0.78 | 0.6 | [] | 1,000.00 | 2103700 | 1,000.00 | 1,000.00 | 1352800 | 1,000.00 |
| 607 | Microtubule-associated protein 1B OS=Homo sapiens | P46821 | MAP1B | 260 | 271 kDa | TRUE | 0.019 | 0.5 | [] | 1.01E+09 | 8.72E+08 | 6.99E+08 | 5.72E+08 | 4.21E+08 | 4.12E+08 |
| 624 | PIN2/TERF1-interacting telomerase inhibitor 1 OS=H | Q96BK5 | PINX1 | 36 | 37 kDa | | 0.04 | 0.5 | [] | 1.05E+08 | 7.29E+07 | 1.20E+08 | 3.35E+07 | 5.24E+07 | 6.40E+07 |
| 612 | Serine-threonine kinase receptor-associated protein O | Q9Y3F4 | STRAP | 319 | 38 kDa | | 0.056 | 0.5 | [] | 8713600 | 1.49E+07 | 1.09E+07 | 4757900 | 5585300 | 7857600 |
| 627 | Eukaryotic initiation factor 4A-I OS=Homo sapiens O | P60842 | EIF4A1 | 479 | 46 kDa | TRUE | 0.075 | 0.5 | [] | 2.33E+07 | 2.36E+07 | 1.17E+07 | 7365100 | 1.11E+07 | 1.07E+07 |
| 621 | ATP-dependent DNA helicase Q1 OS=Homo sapiens | P46063 | RECQL | 69 | 73 kDa | | 0.18 | 0.5 | [] | 1.23E+07 | 2.07E+07 | 1.82E+07 | 6928000 | 1.74E+07 | 1597000 |
| 625 | Eukaryotic translation initiation factor 3 subunit A OS | Q14152 | EIF3A | 287 | 167 kDa | | 0.26 | 0.5 | [] | 1.24E+08 | 7.10E+07 | 2.65E+07 | 3.81E+07 | 4.13E+07 | 3.19E+07 |
| 622 | CAD protein OS=Homo sapiens OX=9606 GN=CAD | F8VPD4 (+1) | CAD | 387 | 236 kDa | TRUE | 0.33 | 0.5 | [] | 1.28E+08 | 9.68E+07 | 1.68E+07 | 1.39E+07 | 5.07E+07 | 5.74E+07 |
| 631 | Developmentally-regulated GTP-binding protein 1 OS | Q9Y295 | DRG1 | 172 | 41 kDa | | 0.38 | 0.5 | [] | 4894400 | 4076300 | 1.84E+07 | 3346800 | 6548300 | 3486800 |

| | | | | | | | | | | | | | | | |
|---|---|---|---|---|---|---|---|---|---|---|---|---|---|---|---|
| 609 | GTPase Era, mitochondrial OS=Homo sapiens OX=96 | O75616 | ERAL1 | 40 | 48 kDa | | 0.4 | 0.5 | [] | 2.27E+07 | 2.94E+07 | 7.96E+07 | 1,000.00 | 3.48E+07 | 3.58E+07 |
| 626 | Eukaryotic translation initiation factor 3 subunit B OS= | P55884 | EIF3B | 282 | 92 kDa | | 0.41 | 0.5 | [] | 7.35E+07 | 2.43E+07 | 1.22E+07 | 1.12E+07 | 3.09E+07 | 1.31E+07 |
| 613 | 40S ribosomal protein S15 OS=Homo sapiens OX=96( | A0A0B4J2B4 (+4) | RPS15 | 239 | 14 kDa | | 0.42 | 0.5 | [] | 4.65E+07 | 2.78E+07 | 1.28E+07 | 1,000.00 | 3.84E+07 | 7361600 |
| 608 | Protein disulfide-isomerase A6 OS=Homo sapiens OX= | Q15084 | PDIA6 | 357 | 48 kDa | | 0.43 | 0.5 | [] | 3.27E+08 | 8.22E+07 | 9.07E+07 | 1.55E+08 | 4.85E+07 | 6.62E+07 |
| 634 | Uncharacterized protein C11orf98 OS=Homo sapiens ( | E9PRG8 | C11orf98 | 88 | 14 kDa | | 0.43 | 0.5 | [] | 1,000.00 | 1.19E+07 | 1.75E+07 | 1,000.00 | 4738000 | 9457700 |
| 640 | 28S ribosomal protein S12, mitochondrial OS=Homo s | O15235 | MRPS12 | 19 | 15 kDa | | 0.43 | 0.5 | [] | 1,000.00 | 9259200 | 5035500 | 1,000.00 | 4233100 | 2351300 |
| 614 | Myosin light chain 1/3, skeletal muscle isoform OS=Hc | P05976 (+1) | MYL1 | 103 | 21 kDa | | 0.47 | 0.5 | [] | 3.57E+07 | 4.75E+07 | 1,000.00 | 1.39E+07 | 1,000.00 | 2.94E+07 |
| 639 | Probable ATP-dependent RNA helicase DDX28 OS=H | Q9NUL7 | DDX28 | 17 | 60 kDa | | 0.49 | 0.5 | [] | 2.53E+07 | 2.30E+07 | 1.39E+08 | 1.04E+07 | 7.04E+07 | 6469000 |
| 616 | Mitochondrial ribosomal protein S27 OS=Homo sapier | G5EA06 (+1) | MRPS27 | 86 | 41 kDa | | 0.5 | 0.5 | [] | 8225600 | 1.33E+08 | 3.74E+07 | 9260400 | 3.73E+07 | 4.53E+07 |
| 629 | E3 ubiquitin-protein ligase OS=Homo sapiens OX=96( | A0A590UJQ1 (+2) | ITCH | 7 | 103 kDa | TRUE | 0.5 | 0.5 | [] | 1,000.00 | 6151200 | 1.62E+07 | 1,000.00 | 5371800 | 5631100 |
| 633 | GIT ArfGAP 1 OS=Homo sapiens OX=9606 GN=GIT | A0A0C4DGN6 (+2) | GIT1 | 29 | 83 kDa | TRUE | 0.5 | 0.5 | [] | 1,000.00 | 1.12E+07 | 9161000 | 9832500 | 1,000.00 | 1,000.00 |
| 615 | Acylamino-acid-releasing enzyme OS=Homo sapiens C | C9JIF9 (+1) | APEH | 35 | 82 kDa | | 0.51 | 0.5 | [] | 1390000 | 560370 | 347000 | 1,000.00 | 1184700 | 1,000.00 |
| 619 | Derlin OS=Homo sapiens OX=9606 GN=DERL1 PE= | B4E1G1 (+2) | DERL1 | 6 | 17 kDa | | 0.51 | 0.5 | [] | 7573600 | 1.62E+07 | 1,000.00 | 1,000.00 | 8555300 | 3634800 |
| 611 | Regulator of nonsense transcripts 1 OS=Homo sapiens | Q92900 | UPF1 | 117 | 124 kDa | | 0.52 | 0.5 | [] | 5.81E+07 | 6277000 | 1.23E+07 | 2.09E+07 | 1.41E+07 | 5636100 |
| 632 | ATP-dependent RNA helicase OS=Homo sapiens OX= | A0A7I2V430 (+2) | DDX1 | 331 | 78 kDa | | 0.54 | 0.5 | [] | 8.35E+07 | 1.09E+07 | 1.72E+07 | 1,000.00 | 5.14E+07 | 2684000 |
| 635 | Transcription elongation factor, mitochondrial OS=Ho | Q96QE5 | TEFM | 0 | 42 kDa | | 0.55 | 0.5 | [] | 3923100 | 841160 | 1,000.00 | 1,000.00 | 1134300 | 1160800 |
| 641 | Probable ATP-dependent RNA helicase DDX23 OS=H | Q9BUQ8 | DDX23 | 144 | 96 kDa | | 0.55 | 0.5 | [] | 1,000.00 | 587460 | 2174700 | 1254900 | 1,000.00 | 1,000.00 |
| 620 | Myosin phosphatase Rho interacting protein OS=Hom | A0A494BZV2 | MPRIP | 86 | 274 kDa | TRUE | 0.57 | 0.5 | [] | 2459000 | 4.74E+07 | 2.48E+08 | 3730700 | 6.31E+07 | 8.41E+07 |
| 628 | Emerin OS=Homo sapiens OX=9606 GN=EMD PE=1 | P50402 | EMD | 270 | 29 kDa | | 0.59 | 0.5 | [] | 1,000.00 | 8765900 | 1824300 | 4708600 | 528100 | 1,000.00 |
| 617 | RNA-binding protein 6 OS=Homo sapiens OX=9606 ( | P78332 | RBM6 | 80 | 129 kDa | | 0.6 | 0.5 | [] | 2.18E+07 | 3339600 | 1,000.00 | 1,000.00 | 7439400 | 5489100 |
| 630 | RNA-binding protein 14 OS=Homo sapiens OX=9606 | Q96PK6 | RBM14 | 293 | 69 kDa | | 0.64 | 0.5 | [] | 1,000.00 | 1,000.00 | 4.59E+07 | 1.22E+07 | 1024200 | 9218300 |
| 636 | Beta-1,3-galactosyltransferase 6 OS=Homo sapiens O) | Q96L58 | B3GALT6 | 1 | 37 kDa | | 0.64 | 0.5 | [] | 1,000.00 | 1,000.00 | 1.44E+07 | 1,000.00 | 2146800 | 4798000 |
| 610 | [3-methyl-2-oxobutanoate dehydrogenase [lipoamide]] | O14874 | BCKDK | 49 | 46 kDa | | 0.66 | 0.5 | [] | 1,000.00 | 1512600 | 1.07E+07 | 6556400 | 1,000.00 | 1,000.00 |
| 637 | Pre-mRNA-processing factor 6 OS=Homo sapiens OX | O94906 | PRPF6 | 180 | 107 kDa | | 0.66 | 0.5 | [] | 1,000.00 | 1,000.00 | 6445700 | 1,000.00 | 1,000.00 | 3069000 |
| 638 | Chromosome 8 open reading frame 33 OS=Homo sapi | A0A3B3IRR6 (+3) | C8orf33 | 59 | 24 kDa | | 0.66 | 0.5 | [] | 1,000.00 | 1,000.00 | 5393200 | 1,000.00 | 2524300 | 1,000.00 |
| 623 | Protein arginine methyltransferase 1 OS=Homo sapien: | E9PKG1 (+1) | PRMT1 | 262 | 38 kDa | | 0.68 | 0.5 | [] | 2293800 | 1,000.00 | 1,000.00 | 1,000.00 | 1156500 | 1,000.00 |
| 642 | Elongation factor Tu, mitochondrial OS=Homo sapien: | P49411 | TUFM | 408 | 50 kDa | | 0.0064 | 0.4 | [] | 2.85E+08 | 3.53E+08 | 2.90E+08 | 1.43E+08 | 1.77E+08 | 9.21E+07 |
| 670 | Cluster of FUS RNA binding protein OS=Homo sapier | H3BPE7 [3] | FUS | 428 | 53 kDa | TRUE | 0.043 | 0.4 | [] | 2.47E+07 | 3.07E+07 | 3.65E+07 | 1,000.00 | 1.88E+07 | 1.47E+07 |
| 664 | Endoplasmic reticulum chaperone BiP OS=Homo sapi | P11021 | HSPA5 | 661 | 72 kDa | TRUE | 0.058 | 0.4 | [] | 3.07E+08 | 2.53E+08 | 5.12E+08 | 1.90E+08 | 1.21E+08 | 1.01E+08 |
| 643 | Stress-70 protein, mitochondrial OS=Homo sapiens O) | A0A7I2V2G2 (+2) | HSPA9 | 552 | 67 kDa | | 0.068 | 0.4 | [] | 6.15E+07 | 1.36E+08 | 1.01E+08 | 3.47E+07 | 3.92E+07 | 5.66E+07 |
| 669 | Cms1 ribosomal small subunit homolog (Fragment) OS | C9J384 (+1) | CMSS1 | 8 | 26 kDa | | 0.092 | 0.4 | [] | 1.37E+07 | 6302900 | 1.66E+07 | 7722000 | 3640400 | 2242600 |
| 655 | G-rich RNA sequence binding factor 1 (Fragment) OS= | H0Y8R1 (+3) | GRSF1 | 114 | 47 kDa | | 0.095 | 0.4 | [] | 6.33E+07 | 3.49E+07 | 2.92E+07 | 1.14E+07 | 2.60E+07 | 1.53E+07 |
| 671 | Surfeit locus protein 4 OS=Homo sapiens OX=9606 G | O15260 (+1) | SURF4 | 158 | 30 kDa | | 0.11 | 0.4 | [] | 2.11E+07 | 8831000 | 1.50E+07 | 1,000.00 | 1.02E+07 | 6149900 |

| | | | | | | | | | | | | | | | |
|---|---|---|---|---|---|---|---|---|---|---|---|---|---|---|---|
| 656 | Splicing factor 3B subunit 1 OS=Homo sapiens OX=9( | O75533 | SF3B1 | 361 | 146 kDa | | 0.12 | 0.4 | [] | 2.55E+07 | 1.30E+07 | 2.73E+07 | 1,000.00 | 1.69E+07 | 1.02E+07 |
| 666 | Nucleolar RNA helicase 2 OS=Homo sapiens OX=960 | Q9NR30 | DDX21 | 438 | 87 kDa | TRUE | 0.13 | 0.4 | [] | 6.04E+07 | 5.11E+07 | 1.36E+08 | 2.69E+07 | 3.20E+07 | 3.50E+07 |
| 654 | Cluster of Utrophin OS=Homo sapiens OX=9606 GN= | P46939 [2] | UTRN | 83 | 394 kDa | TRUE | 0.14 | 0.4 | [] | 1.30E+08 | 5.73E+07 | 8.76E+07 | 7.76E+07 | 2.92E+07 | 7425600 |
| 674 | Coatomer subunit gamma-1 OS=Homo sapiens OX=9( | Q9Y678 | COPG1 | 228 | 98 kDa | TRUE | 0.15 | 0.4 | [] | 4.43E+07 | 1.47E+07 | 2.08E+07 | 1.12E+07 | 1.47E+07 | 2698400 |
| 659 | Hydroxysteroid 17-beta dehydrogenase 10 OS=Homo | A0A804HHW5 (+1) | HSD17B10 | 290 | 22 kDa | | 0.16 | 0.4 | [] | 1.47E+07 | 1.79E+07 | 1.41E+07 | 1.68E+07 | 2171200 | 1,000.00 |
| 662 | Importin subunit alpha OS=Homo sapiens OX=9606 G | A0A7I2V351 (+4) | KPNA2 | 367 | 55 kDa | | 0.16 | 0.4 | [] | 1.27E+07 | 4217600 | 1.79E+07 | 6023100 | 4279100 | 3448500 |
| 673 | ATP-dependent RNA helicase A OS=Homo sapiens O | Q08211 | DHX9 | 470 | 141 kDa | | 0.16 | 0.4 | [] | 2.99E+07 | 7.69E+07 | 3.38E+07 | 1,000.00 | 2.61E+07 | 2.44E+07 |
| 672 | Heterogeneous nuclear ribonucleoprotein A3 OS=Hon | A0A7I2V2R3 (+3) | HNRNPA3 | 447 | 35 kDa | | 0.18 | 0.4 | [] | 1.09E+07 | 3152100 | 1.42E+07 | 1,000.00 | 4162700 | 6074600 |
| 675 | Solute carrier family 3 member 2 OS=Homo sapiens O | A0A7P0TAN7 (+5) | SLC3A2 | 189 | 67 kDa | | 0.18 | 0.4 | [] | 8.82E+07 | 6.14E+07 | 2.74E+07 | 1,000.00 | 5.03E+07 | 1.28E+07 |
| 667 | Heterogeneous nuclear ribonucleoprotein H3 OS=Hon | P31942 | HNRNPH3 | 239 | 37 kDa | | 0.2 | 0.4 | [] | 1.29E+07 | 8944100 | 8181800 | 1,000.00 | 1,000.00 | 1.13E+07 |
| 646 | Microtubule associated protein 7 OS=Homo sapiens O | A0A087WZ40 (+1) | MAP7 | 109 | 88 kDa | | 0.25 | 0.4 | [] | 1.47E+07 | 3.51E+07 | 5.46E+07 | 379780 | 3.14E+07 | 1.31E+07 |
| 658 | 60S ribosomal protein L29 OS=Homo sapiens OX=96( | A0A3B3ITT5 (+1) | RPL29 | 399 | 19 kDa | | 0.25 | 0.4 | [] | 1.18E+08 | 2.64E+08 | 5.06E+08 | 2.42E+08 | 3.73E+07 | 8.35E+07 |
| 645 | Cluster of Ras-related protein Rab-10 OS=Homo sapie | P61026 [6] | RAB10 | 176 | 23 kDa | TRUE | 0.27 | 0.4 | [] | 3499600 | 4892300 | 4022400 | 1,000.00 | 5360000 | 1,000.00 |
| 677 | Cleavage and polyadenylation specificity factor subunit | O43809 | NUDT21 | 295 | 26 kDa | | 0.27 | 0.4 | [] | 9251600 | 2.97E+07 | 7567400 | 727520 | 1.22E+07 | 3439100 |
| 660 | Serine/arginine repetitive matrix protein 2 OS=Homo s | Q9UQ35 | SRRM2 | 324 | 300 kDa | | 0.28 | 0.4 | [] | 2.89E+07 | 3.95E+07 | 1.25E+08 | 2.58E+07 | 1.84E+07 | 3.41E+07 |
| 644 | Cyclin-T1 OS=Homo sapiens OX=9606 GN=CCNT1 | O60563 | CCNT1 | 133 | 81 kDa | | 0.29 | 0.4 | [] | 1.47E+07 | 1.15E+07 | 2.10E+07 | 2.05E+07 | 1,000.00 | 1,000.00 |
| 648 | rRNA adenine N(6)-methyltransferase OS=Homo sapi( | A0A494C0G7 (+1) | DIMT1 | 51 | 36 kDa | | 0.32 | 0.4 | [] | 1.30E+07 | 7386000 | 1562800 | 932000 | 6537300 | 1879800 |
| 650 | Helicase MOV-10 OS=Homo sapiens OX=9606 GN=! | Q9HCE1 | MOV10 | 87 | 114 kDa | | 0.32 | 0.4 | [] | 8483000 | 1.49E+07 | 3.18E+07 | 2.04E+07 | 3042600 | 1,000.00 |
| 663 | 39S ribosomal protein L37, mitochondrial OS=Homo s | Q9BZE1 (+1) | MRPL37 | 61 | 48 kDa | | 0.34 | 0.4 | [] | 1231300 | 1.60E+07 | 9415500 | 1,000.00 | 1466500 | 8847200 |
| 665 | Death-inducer obliterator 1 OS=Homo sapiens OX=96 | Q9BTC0 | DIDO1 | 140 | 244 kDa | | 0.39 | 0.4 | [] | 6612100 | 474350 | 1.10E+07 | 6933400 | 1,000.00 | 1,000.00 |
| 668 | DNA replication licensing factor MCM6 OS=Homo sa | Q14566 | MCM6 | 282 | 93 kDa | | 0.39 | 0.4 | [] | 2764300 | 2922800 | 1.85E+07 | 3944100 | 3717400 | 1375500 |
| 649 | Probable helicase senataxin OS=Homo sapiens OX=96 | Q7Z333 | SETX | 43 | 303 kDa | | 0.4 | 0.4 | [] | 1,000.00 | 1.66E+07 | 3.34E+07 | 1.09E+07 | 8914800 | 1477000 |
| 661 | Very-long-chain (3R)-3-hydroxyacyl-CoA dehydratase | Q9P035 | HACD3 | 176 | 43 kDa | | 0.43 | 0.4 | [] | 2.95E+07 | 3888300 | 7494300 | 1,000.00 | 1776900 | 1.47E+07 |
| 676 | Thyroid hormone receptor associated protein 3 OS=H( | A0A3B3ITZ9 (+1) | THRAP3 | 382 | 104 kDa | | 0.44 | 0.4 | [] | 3159000 | 1,000.00 | 1.01E+07 | 1,000.00 | 4702300 | 1,000.00 |
| 657 | Cluster of Serine/threonine-protein phosphatase 2A 65 | P30153 [2] | PPP2R1A | 316 | 65 kDa | TRUE | 0.45 | 0.4 | [] | 1420300 | 1.74E+07 | 3490700 | 1,000.00 | 4121600 | 5029500 |
| 653 | Ring finger protein 213 OS=Homo sapiens OX=9606 ( | A0A0A0MTC1 (+2) | RNF213 | 23 | 596 kDa | TRUE | 0.47 | 0.4 | [] | 2.14E+08 | 1.73E+07 | 4.29E+07 | 9.26E+07 | 1.85E+07 | 2726700 |
| 651 | Patatin like phospholipase domain containing 6 OS=H( | A0A384DVU0 (+1) | PNPLA6 | 9 | 150 kDa | | 0.53 | 0.4 | [] | 1473200 | 1.01E+08 | 1.28E+07 | 1,000.00 | 1.45E+07 | 3.38E+07 |
| 652 | Nuclear RNA export factor 2 OS=Homo sapiens OX=! | Q9GZY0 | NXF2 | 0 | 72 kDa | | 0.55 | 0.4 | [] | 953920 | 1,000.00 | 5935300 | 1,000.00 | 1,000.00 | 2880800 |
| 703 | Neuroblast differentiation-associated protein AHNAK | Q09666 | AHNAK | 212 | 629 kDa | | 0.00021 | 0.3 | BioID high, ptprh low | 2.10E+10 | 2.11E+10 | 2.03E+10 | 7.99E+09 | 4.24E+09 | 5.04E+09 |
| 704 | Annexin OS=Homo sapiens OX=9606 GN=ANXA2 P | H0YMW4 (+1) | ANXA2 | 398 | 42 kDa | | 0.0065 | 0.3 | [] | 6.67E+08 | 7.09E+08 | 4.65E+08 | 1.09E+08 | 2.47E+08 | 1.41E+08 |
| 711 | Guanine nucleotide-binding protein-like 3 OS=Homo s | Q9BVP2 | GNL3 | 256 | 62 kDa | | 0.016 | 0.3 | [] | 2.62E+07 | 2.04E+07 | 1.61E+07 | 2011000 | 1.03E+07 | 3665900 |
| 694 | Poly [ADP-ribose] polymerase tankyrase-1 OS=Homo | O95271 | TNKS | 1 | 142 kDa | | 0.024 | 0.3 | [] | 5.78E+07 | 4.31E+07 | 3.79E+07 | 1.22E+07 | 2.67E+07 | 2574900 |
| 714 | 182 kDa tankyrase-1-binding protein OS=Homo sapier | Q9C0C2 | TNKS1BP1 | 85 | 182 kDa | | 0.024 | 0.3 | [] | 1.34E+08 | 2.23E+08 | 1.68E+08 | 9.68E+07 | 1.85E+07 | 1.69E+07 |

| | | | | | | | | | | | | | | | |
|---|---|---|---|---|---|---|---|---|---|---|---|---|---|---|---|
| 699 | Cluster of Dystonin OS=Homo sapiens OX=9606 GN= | A0A7P0T890 [6] | DST | 84 | 888 kDa | TRUE | 0.048 | 0.3 | [] | 7.83E+07 | 9.36E+07 | 1.45E+08 | 7410100 | 1.74E+07 | 6.51E+07 |
| 691 | Pre-rRNA-processing protein TSR1 homolog OS=Hor | Q2NL82 | TSR1 | 148 | 92 kDa | | 0.069 | 0.3 | [] | 4.57E+07 | 3.17E+07 | 5.27E+07 | 1307000 | 4382800 | 3.41E+07 |
| 681 | Uncharacterized protein KIAA1522 OS=Homo sapien: | Q9P206 | KIAA1522 | 5 | 107 kDa | | 0.079 | 0.3 | [] | 3.61E+07 | 4.07E+07 | 1.44E+07 | 6213900 | 1.61E+07 | 8282100 |
| 705 | Kinesin-like protein (Fragment) OS=Homo sapiens OX | A0A6Q8PGH7 | KIF2A | 105 | 78 kDa | TRUE | 0.082 | 0.3 | [] | 4711800 | 2777400 | 4953100 | 1,000.00 | 3343300 | 1,000.00 |
| 697 | Mitochondrial 2-oxoglutarate/malate carrier protein OS | Q02978 | SLC25A11 | 290 | 34 kDa | | 0.083 | 0.3 | [] | 7828800 | 7982800 | 1.55E+07 | 1,000.00 | 2177200 | 6813700 |
| 701 | Cleavage and polyadenylation specific factor 7 (Fragm | F5H669 (+1) | CPSF7 | 154 | 41 kDa | | 0.084 | 0.3 | [] | 1.28E+07 | 1.76E+07 | 3.21E+07 | 1,000.00 | 7458200 | 1.01E+07 |
| 695 | Cluster of Protein flightless-1 homolog OS=Homo sap | Q13045 [2] | FLII | 81 | 145 kDa | TRUE | 0.1 | 0.3 | [] | 2.21E+07 | 7865200 | 2.98E+07 | 7168200 | 8110900 | 2532300 |
| 707 | Mitochondrial ribosomal protein L45 OS=Homo sapier | A0A087X2D5 (+1) | MRPL45 | 41 | 35 kDa | | 0.1 | 0.3 | [] | 2.14E+07 | 2.93E+07 | 5.62E+07 | 2268600 | 2.28E+07 | 3263300 |
| 696 | Cluster of Sarcoplasmic/endoplasmic reticulum calciun | P16615 [2] | ATP2A2 | 302 | 115 kDa | TRUE | 0.11 | 0.3 | [] | 9.76E+07 | 3.13E+07 | 6.76E+07 | 4.45E+07 | 8177900 | 4940400 |
| 692 | RRP12-like protein OS=Homo sapiens OX=9606 GN= | Q5JTH9 | RRP12 | 176 | 144 kDa | | 0.12 | 0.3 | [] | 5.45E+07 | 3.58E+07 | 1.78E+07 | 1,000.00 | 9652800 | 2.33E+07 |
| 712 | Striatin OS=Homo sapiens OX=9606 GN=STRN PE= | O43815 | STRN | 38 | 86 kDa | TRUE | 0.13 | 0.3 | [] | 2.56E+07 | 8.80E+07 | 7.46E+07 | 4.77E+07 | 1,000.00 | 1,000.00 |
| 683 | Cluster of Receptor of activated protein C kinase 1 OS | P63244 [2] | RACK1 | 352 | 35 kDa | TRUE | 0.14 | 0.3 | [] | 2.42E+07 | 2.09E+07 | 5049200 | 3720800 | 9046100 | 3809100 |
| 709 | Eukaryotic translation initiation factor 3 subunit C-like | B5ME19 (+1) | EIF3CL | 274 | 105 kDa | | 0.14 | 0.3 | [] | 1.67E+07 | 1.13E+07 | 3.58E+07 | 1.42E+07 | 724830 | 1759400 |
| 698 | Ataxin 2 OS=Homo sapiens OX=9606 GN=ATXN2 P | A0A2R8Y5A6 (+10 | ATXN2 | 116 | 126 kDa | | 0.15 | 0.3 | [] | 8913900 | 2.99E+07 | 4.78E+07 | 1.20E+07 | 5191300 | 7648700 |
| 706 | Cluster of Fragile X messenger ribonucleoprotein 1 OS | A8MQB8 [3] | FMR1 | 131 | 66 kDa | TRUE | 0.18 | 0.3 | [] | 6946900 | 1.86E+07 | 3.09E+07 | 1,000.00 | 1.51E+07 | 1,000.00 |
| 684 | Nucleophosmin OS=Homo sapiens OX=9606 GN=NP | A0A7I2V5S2 (+1) | NPM1 | 554 | 34 kDa | | 0.21 | 0.3 | [] | 2.54E+08 | 1.31E+08 | 5.40E+07 | 1,000.00 | 1.04E+08 | 4.13E+07 |
| 690 | Eukaryotic peptide chain release factor subunit 1 OS=I | P62495 | ETF1 | 167 | 49 kDa | | 0.22 | 0.3 | [] | 2.65E+07 | 1.18E+07 | 6096100 | 1,000.00 | 1.14E+07 | 2152600 |
| 700 | Protein argonaute-2 OS=Homo sapiens OX=9606 GN | Q9UKV8 | AGO2 | 92 | 97 kDa | | 0.24 | 0.3 | [] | 1,000.00 | 8429500 | 8723900 | 1,000.00 | 2077300 | 2772800 |
| 686 | LIM and calponin homology domains 1 OS=Homo sap | D6RD46 (+1) | LIMCH1 | 89 | 110 kDa | | 0.25 | 0.3 | [] | 1229100 | 3.91E+07 | 5.51E+07 | 5545400 | 1.24E+07 | 1.29E+07 |
| 678 | ATP-dependent RNA helicase DDX18 OS=Homo sapi | Q9NVP1 | DDX18 | 203 | 75 kDa | | 0.26 | 0.3 | [] | 7813200 | 2.84E+07 | 3.60E+07 | 1,000.00 | 1,000.00 | 2.53E+07 |
| 680 | Proteasome 26S subunit, ATPase 2 OS=Homo sapiens | C9JX88 (+1) | PSMC2 | 313 | 48 kDa | | 0.26 | 0.3 | [] | 2.18E+07 | 8114000 | 3.83E+07 | 1,000.00 | 2.30E+07 | 1,000.00 |
| 702 | Glucose-6-phosphate 1-dehydrogenase OS=Homo sap | P11413 | G6PD | 89 | 59 kDa | | 0.28 | 0.3 | [] | 6508300 | 1,000.00 | 9462000 | 4040200 | 154530 | 254340 |
| 689 | Cluster of ATP-dependent RNA helicase DDX54 OS= | Q8TDD1 [2] | DDX54 | 148 | 99 kDa | TRUE | 0.32 | 0.3 | [] | 4939400 | 1,000.00 | 3183800 | 1,000.00 | 2503400 | 1,000.00 |
| 693 | Importin subunit beta-1 OS=Homo sapiens OX=9606 ( | Q14974 | KPNB1 | 418 | 97 kDa | | 0.32 | 0.3 | [] | 686770 | 2.21E+07 | 8737800 | 1,000.00 | 5409600 | 4193600 |
| 685 | 7SK snRNA methylphosphate capping enzyme OS=Hc | Q7L2J0 | MEPCE | 88 | 74 kDa | | 0.33 | 0.3 | [] | 3.52E+07 | 9796600 | 3714500 | 2238000 | 3959900 | 9767100 |
| 679 | 39S ribosomal protein L28, mitochondrial OS=Homo s | Q13084 | MRPL28 | 42 | 30 kDa | | 0.36 | 0.3 | [] | 4864600 | 7576800 | 1,000.00 | 1,000.00 | 4260100 | 1,000.00 |
| 710 | Cluster of Nucleolar GTP-binding protein 2 OS=Homc | Q13823 [2] | GNL2 | 138 | 84 kDa | TRUE | 0.38 | 0.3 | [] | 1,000.00 | 2.20E+07 | 4886700 | 1,000.00 | 2454200 | 4475600 |
| 713 | RNA-binding protein FXR2 OS=Homo sapiens OX=9( | P51116 | FXR2 | 139 | 74 kDa | TRUE | 0.39 | 0.3 | [] | 1,000.00 | 2.23E+07 | 9.82E+07 | 1,000.00 | 1,000.00 | 3.04E+07 |
| 687 | Cluster of Tyrosine 3-monooxygenase/tryptophan 5-mc | E9PG15 [7] | YWHAQ | 410 | 17 kDa | TRUE | 0.5 | 0.3 | [] | 1,000.00 | 2099000 | 1.85E+07 | 1,000.00 | 6628800 | 1,000.00 |
| 708 | ATPase WRNIP1 OS=Homo sapiens OX=9606 GN=V | Q96S55 | WRNIP1 | 111 | 72 kDa | | 0.52 | 0.3 | [] | 5033200 | 1,000.00 | 1,000.00 | 1327500 | 1,000.00 | 1,000.00 |
| 682 | Importin-9 OS=Homo sapiens OX=9606 GN=IPO9 PI | Q96P70 | IPO9 | 120 | 116 kDa | | 0.55 | 0.3 | [] | 2946100 | 1,000.00 | 1,000.00 | 1,000.00 | 391900 | 584680 |
| 730 | Carbamoyl-phosphate synthase [ammonia], mitochondi | P31327 | CPS1 | 82 | 165 kDa | TRUE | 0.0062 | 0.2 | [] | 2.30E+08 | 1.76E+08 | 1.69E+08 | 7066400 | 8.01E+07 | 2.72E+07 |
| 733 | Cold shock domain-containing protein E1 OS=Homo s | O75534 | CSDE1 | 191 | 89 kDa | | 0.007 | 0.2 | [] | 3.71E+07 | 3.78E+07 | 5.47E+07 | 1,000.00 | 1.08E+07 | 1.25E+07 |

| | | | | | | | | | | | | | | | |
|---|---|---|---|---|---|---|---|---|---|---|---|---|---|---|---|
| 724 | General transcription factor IIi OS=Homo sapiens OX= | A0A494C1K3 (+1) | GTF2I | 312 | 131 kDa | | 0.02 | 0.2 | [] | 8.79E+07 | 1.88E+08 | 1.51E+08 | 3.19E+07 | 3.29E+07 | 3.24E+07 |
| 731 | Band 4.1-like protein 5 OS=Homo sapiens OX=9606 ( | Q9HCM4 | EPB41L5 | 9 | 82 kDa | | 0.048 | 0.2 | [] | 4.91E+07 | 3.04E+07 | 8.26E+07 | 8829700 | 1.29E+07 | 1.03E+07 |
| 720 | Eukaryotic translation initiation factor 4 gamma 1 OS= | E7EUU4 (+1) | EIF4G1 | 293 | 172 kDa | TRUE | 0.068 | 0.2 | [] | 5.09E+07 | 1.67E+07 | 4.34E+07 | 1,000.00 | 9441300 | 1.64E+07 |
| 725 | Ras GTPase-activating protein-binding protein 1 OS=I | Q13283 | G3BP1 | 353 | 52 kDa | TRUE | 0.07 | 0.2 | [] | 1.97E+08 | 8.82E+07 | 8.00E+07 | 1.45E+07 | 4.34E+07 | 2.30E+07 |
| 726 | Obg-like ATPase 1 OS=Homo sapiens OX=9606 GN= | J3KQ32 (+1) | OLA1 | 120 | 47 kDa | | 0.077 | 0.2 | [] | 8291500 | 3285900 | 5562300 | 1,000.00 | 3667500 | 1,000.00 |
| 738 | Protein NipSnap homolog 2 OS=Homo sapiens OX=9( | O75323 | NIPSNAP2 | 24 | 34 kDa | TRUE | 0.11 | 0.2 | [] | 8979800 | 5.25E+07 | 5.38E+07 | 1,000.00 | 1,000.00 | 1.80E+07 |
| 739 | Anterior gradient protein 2 homolog OS=Homo sapien | O95994 | AGR2 | 0 | 20 kDa | | 0.11 | 0.2 | [] | 2.70E+07 | 2.88E+07 | 3637000 | 1,000.00 | 4366000 | 4547400 |
| 729 | Splicing factor, proline- and glutamine-rich OS=Homo | P23246 | SFPQ | 484 | 76 kDa | TRUE | 0.13 | 0.2 | [] | 5366700 | 944400 | 4362300 | 1,000.00 | 2125800 | 1,000.00 |
| 716 | Nucleolin OS=Homo sapiens OX=9606 GN=NCL PE= | A0A7I2V428 (+3) | NCL | 527 | 75 kDa | | 0.14 | 0.2 | [] | 3.28E+08 | 1.53E+08 | 7.18E+07 | 5.88E+07 | 3.46E+07 | 4.10E+07 |
| 737 | Zinc finger protein 768 OS=Homo sapiens OX=9606 ( | H3BS42 (+1) | ZNF768 | 22 | 57 kDa | TRUE | 0.18 | 0.2 | [] | 1,000.00 | 4905300 | 5295400 | 1,000.00 | 1595100 | 1,000.00 |
| 735 | Cluster of valine--tRNA ligase OS=Homo sapiens OX= | A0A140T8Y0 [4] | VARS2 | 5 | 118 kDa | TRUE | 0.2 | 0.2 | [] | 1,000.00 | 8750800 | 1.11E+07 | 3292000 | 1,000.00 | 1,000.00 |
| 734 | Probable ATP-dependent RNA helicase DDX6 OS=Hc | P26196 | DDX6 | 303 | 54 kDa | | 0.21 | 0.2 | [] | 2400800 | 3566300 | 1,000.00 | 1,000.00 | 1005200 | 1,000.00 |
| 722 | Ubiquitin carboxyl-terminal hydrolase 7 OS=Homo sap | A0A669KBL1 (+2) | USP7 | 201 | 132 kDa | | 0.24 | 0.2 | [] | 2689400 | 3366400 | 1,000.00 | 1,000.00 | 1383400 | 1,000.00 |
| 717 | Desmoglein-2 OS=Homo sapiens OX=9606 GN=DSG | Q14126 | DSG2 | 113 | 122 kDa | | 0.3 | 0.2 | [] | 8860800 | 4232200 | 1,000.00 | 1,000.00 | 3167700 | 1,000.00 |
| 718 | Cell division cycle 5-like protein OS=Homo sapiens O? | Q99459 | CDC5L | 209 | 92 kDa | | 0.43 | 0.2 | [] | 551750 | 2729400 | 2.83E+07 | 6752400 | 1,000.00 | 834450 |
| 736 | U2 snRNP associated SURP domain containing OS=H | E7ET15 (+1) | U2SURP | 262 | 118 kDa | | 0.46 | 0.2 | [] | 1.97E+07 | 1,000.00 | 1,000.00 | 1,000.00 | 1,000.00 | 3214500 |
| 727 | BCL2 associated transcription factor 1 OS=Homo sapi | A0A1W2PQ43 (+5) | BCLAF1 | 390 | 81 kDa | | 0.48 | 0.2 | [] | 1,000.00 | 47314 | 1.06E+07 | 1,000.00 | 1,000.00 | 2257900 |
| 728 | RNA helicase OS=Homo sapiens OX=9606 GN=DDX | A0A7P0T9T8 (+1) | DDX20 | 108 | 83 kDa | | 0.48 | 0.2 | [] | 1.41E+07 | 1,000.00 | 1,000.00 | 2880400 | 1,000.00 | 1,000.00 |
| 721 | Dynein cytoplasmic 1 heavy chain 1 OS=Homo sapiens | A0A2R8Y5T0 (+6) | DYNC1H1 | 345 | 528 kDa | | 0.49 | 0.2 | [] | 6841700 | 1,000.00 | 1,000.00 | 1,000.00 | 839610 | 725880 |
| 719 | Coatomer subunit gamma-2 OS=Homo sapiens OX=9( | Q9UBF2 | COPG2 | 224 | 98 kDa | TRUE | 0.5 | 0.2 | [] | 8814000 | 1,000.00 | 1,000.00 | 2085900 | 1,000.00 | 1,000.00 |
| 758 | Enhancer of mRNA-decapping protein 4 OS=Homo sa | Q6P2E9 | EDC4 | 163 | 152 kDa | | 0.0082 | 0.1 | [] | 6.81E+07 | 1.13E+08 | 6.66E+07 | 9231600 | 8729700 | 8042100 |
| 748 | Golgin subfamily B member 1 OS=Homo sapiens OX= | Q14789 | GOLGB1 | 30 | 376 kDa | | 0.011 | 0.1 | [] | 1.61E+07 | 2.61E+07 | 2.00E+07 | 8113100 | 1,000.00 | 1,000.00 |
| 753 | Cluster of Heterogeneous nuclear ribonucleoprotein L | A0A3B3ITJ4 [2] | HNRNPL | 460 | 59 kDa | TRUE | 0.031 | 0.1 | [] | 1.78E+07 | 1.05E+07 | 7297600 | 2346100 | 1,000.00 | 1766500 |
| 747 | T-complex protein 1 subunit zeta OS=Homo sapiens C | P40227 | CCT6A | 413 | 58 kDa | | 0.033 | 0.1 | [] | 5144300 | 5846700 | 1.13E+07 | 1,000.00 | 1169300 | 1746500 |
| 754 | EWS RNA binding protein 1 OS=Homo sapiens OX=9 | B0QYK0 (+1) | EWSR1 | 301 | 65 kDa | | 0.035 | 0.1 | [] | 1.09E+07 | 2.81E+07 | 1.60E+07 | 1,000.00 | 3532300 | 2686800 |
| 749 | DNA-dependent protein kinase catalytic subunit OS=H | P78527 | PRKDC | 406 | 469 kDa | | 0.041 | 0.1 | [] | 2.48E+07 | 6.84E+07 | 4.30E+07 | 1.33E+07 | 362710 | 3883800 |
| 742 | Cluster of T-complex protein 1 subunit alpha OS=Hon | P17987 [2] | TCP1 | 432 | 60 kDa | TRUE | 0.06 | 0.1 | [] | 1.13E+07 | 9750300 | 2.39E+07 | 6305000 | 1,000.00 | 1,000.00 |
| 751 | Smoothelin OS=Homo sapiens OX=9606 GN=SMTN | A0A087WVP4 (+2) | SMTN | 11 | 105 kDa | | 0.089 | 0.1 | [] | 3351100 | 1.62E+07 | 1.80E+07 | 4766800 | 1,000.00 | 1,000.00 |
| 756 | Kinesin-like protein KIFC1 OS=Homo sapiens OX=96 | Q9BW19 | KIFC1 | 78 | 74 kDa | | 0.093 | 0.1 | [] | 1.06E+07 | 1647800 | 9589500 | 2419700 | 1,000.00 | 1,000.00 |
| 746 | 5'-3' exoribonuclease 2 OS=Homo sapiens OX=9606 ( | Q9H0D6 | XRN2 | 276 | 109 kDa | | 0.13 | 0.1 | [] | 1.03E+07 | 740040 | 1.18E+07 | 1657700 | 1352800 | 1,000.00 |
| 744 | Golgin subfamily A member 2 OS=Homo sapiens OX= | A0A6Q8KRG2 (+1) | GOLGA2 | 71 | 112 kDa | | 0.15 | 0.1 | [] | 1.06E+09 | 6.14E+08 | 8.74E+07 | 1.58E+08 | 4.62E+07 | 4.14E+07 |
| 755 | Erlin-1 OS=Homo sapiens OX=9606 GN=ERLIN1 PE | O75477 | ERLIN1 | 81 | 39 kDa | | 0.16 | 0.1 | [] | 1,000.00 | 1.07E+07 | 1.38E+07 | 1,000.00 | 2721800 | 1,000.00 |
| 741 | Protein S100-A10 OS=Homo sapiens OX=9606 GN=S | P60903 | S100A10 | 17 | 11 kDa | | 0.22 | 0.1 | [] | 8.64E+07 | 2.30E+08 | 1.17E+07 | 1,000.00 | 2.43E+07 | 2.24E+07 |

| | | | | | | | | | | | | | | | |
|---|---|---|---|---|---|---|---|---|---|---|---|---|---|---|---|
| 752 | Protein phosphatase 1G OS=Homo sapiens OX=9606 | O15355 | PPM1G | 155 | 59 kDa | | 0.22 | 0.1 | [] | 1,000.00 | 2162300 | 4908600 | 1,000.00 | 1,000.00 | 834830 |
| 760 | Citron Rho-interacting kinase OS=Homo sapiens OX= | O14578 | CIT | 10 | 231 kDa | | 0.25 | 0.1 | [] | 1,000.00 | 2.03E+07 | 6633100 | 1,000.00 | 1,000.00 | 2760500 |
| 740 | Kinectin OS=Homo sapiens OX=9606 GN=KTN1 PE= | Q86UP2 | KTN1 | 132 | 156 kDa | | 0.26 | 0.1 | [] | 820200 | 9247300 | 2.85E+07 | 4740000 | 1,000.00 | 833030 |
| 762 | TAF6-like RNA polymerase II p300/CBP-associated fa | Q9Y6J9 | TAF6L | 32 | 68 kDa | | 0.27 | 0.1 | [] | 1,000.00 | 4061600 | 1.48E+07 | 1822700 | 1,000.00 | 1,000.00 |
| 750 | GTP-binding protein 10 OS=Homo sapiens OX=9606 | A4D1E9 | GTPBP10 | 8 | 43 kDa | | 0.33 | 0.1 | [] | 2128400 | 1,000.00 | 382640 | 321220 | 1,000.00 | 1,000.00 |
| 743 | Family with sequence similarity 83 member H OS=Hor | A0A494C1T9 (+2) | FAM83H | 6 | 149 kDa | | 0.39 | 0.1 | [] | 5061800 | 1,000.00 | 6.63E+07 | 583990 | 4766300 | 4607500 |
| 759 | serine--tRNA ligase OS=Homo sapiens OX=9606 GN= | M0QWZ7 (+1) | SARS2 | 75 | 58 kDa | | 0.4 | 0.1 | [] | 4101700 | 1.35E+08 | 1,000.00 | 1,000.00 | 1.08E+07 | 3612700 |
| 761 | Sideroflexin-3 OS=Homo sapiens OX=9606 GN=SFX | Q9BWM7 (+1) | SFXN3 | 31 | 36 kDa | | 0.42 | 0.1 | [] | 1,000.00 | 1,000.00 | 5.04E+07 | 4681100 | 201880 | 1,000.00 |
| 757 | Zinc finger CCCH domain-containing protein 15 OS=H | Q8WU90 | ZC3H15 | 149 | 49 kDa | | 0.43 | 0.1 | [] | 3.65E+07 | 1,000.00 | 1,000.00 | 3226700 | 792320 | 1,000.00 |
| 770 | Cluster of Lamina-associated polypeptide 2, isoform al | P42166 [2] | TMPO | 386 | 75 kDa | TRUE | 0.0026 | 0.09 | [] | 1.30E+08 | 1.30E+08 | 1.91E+08 | 8855500 | 1.63E+07 | 1.47E+07 |
| 768 | Splicing factor 3B subunit 2 OS=Homo sapiens OX=9 | Q13435 | SF3B2 | 317 | 100 kDa | | 0.0056 | 0.09 | [] | 3.04E+07 | 2.32E+07 | 4.11E+07 | 2205400 | 4772700 | 1417200 |
| 773 | Cluster of Protein numb homolog OS=Homo sapiens C | P49757 [2] | NUMB | 42 | 71 kDa | TRUE | 0.11 | 0.09 | [] | 4.35E+07 | 3546800 | 3.63E+07 | 7261100 | 1,000.00 | 1,000.00 |
| 772 | A-kinase anchor protein 13 OS=Homo sapiens OX=96 | Q12802 | AKAP13 | 32 | 308 kDa | | 0.14 | 0.09 | [] | 2.21E+07 | 4400600 | 7485500 | 2965800 | 1,000.00 | 1,000.00 |
| 764 | Upstream binding transcription factor OS=Homo sapie | E9PKP7 | UBTF | 68 | 87 kDa | | 0.15 | 0.09 | [] | 2778300 | 1.65E+07 | 3.27E+07 | 1,000.00 | 1,000.00 | 4847100 |
| 767 | GTP-binding nuclear protein Ran OS=Homo sapiens C | B5MDF5 (+2) | RAN | 440 | 26 kDa | | 0.16 | 0.09 | [] | 1,000.00 | 9078700 | 1.34E+07 | 1,000.00 | 1,000.00 | 2020500 |
| 765 | Staphylococcal nuclease domain-containing protein 1 C | Q7KZF4 | SND1 | 265 | 102 kDa | | 0.22 | 0.09 | [] | 1,000.00 | 8108300 | 3274100 | 1,000.00 | 1055100 | 1,000.00 |
| 771 | Large neutral amino acids transporter small subunit 1 C | Q01650 | SLC7A5 | 73 | 55 kDa | TRUE | 0.24 | 0.09 | [] | 1,000.00 | 1.62E+07 | 4.79E+07 | 1,000.00 | 5630700 | 1,000.00 |
| 763 | Zinc finger CCHC domain-containing protein 8 OS=Ho | Q6NZY4 | ZCCHC8 | 138 | 79 kDa | | 0.37 | 0.09 | [] | 1,000.00 | 1.77E+07 | 1468500 | 1,000.00 | 1805800 | 1,000.00 |
| 769 | Zinc finger CCCH domain-containing protein 7A OS=H | Q8IWR0 | ZC3H7A | 24 | 111 kDa | | 0.38 | 0.09 | [] | 3.20E+07 | 315440 | 1744600 | 1,000.00 | 3022500 | 1,000.00 |
| 766 | GRB10-interacting GYF protein 2 OS=Homo sapiens C | Q6Y7W6 | GIGYF2 | 156 | 150 kDa | | 0.41 | 0.09 | [] | 1,000.00 | 2.86E+08 | 3646300 | 2.69E+07 | 1,000.00 | 1,000.00 |
| 777 | Proliferation marker protein Ki-67 OS=Homo sapiens C | P46013 | MKI67 | 246 | 359 kDa | | 0.11 | 0.08 | [] | 3746600 | 1.92E+07 | 3.16E+07 | 1593300 | 1751100 | 1004900 |
| 774 | Tight junction protein ZO-3 OS=Homo sapiens OX=96 | O95049 | TJP3 | 3 | 101 kDa | | 0.14 | 0.08 | [] | 1346000 | 7385100 | 1.49E+07 | 1998000 | 1,000.00 | 1,000.00 |
| 775 | DNA replication licensing factor MCM3 OS=Homo sa | P25205 | MCM3 | 322 | 91 kDa | | 0.17 | 0.08 | [] | 1875800 | 1,000.00 | 3202600 | 1,000.00 | 300190 | 127600 |
| 776 | Rho guanine nucleotide exchange factor 2 OS=Homo s | Q92974 | ARHGEF2 | 78 | 112 kDa | | 0.31 | 0.08 | [] | 1,000.00 | 2.83E+07 | 5088900 | 1,000.00 | 2705800 | 1,000.00 |
| 784 | Spectrin beta chain, non-erythrocytic 1 OS=Homo sapi | Q01082 | SPTBN1 | 250 | 275 kDa | TRUE | 0.00035 | 0.07 | BioID high, ptprh low | 8.56E+07 | 9.02E+07 | 7.67E+07 | 1.72E+07 | 1,000.00 | 1,000.00 |
| 782 | Gem-associated protein 5 OS=Homo sapiens OX=960 | Q8TEQ6 | GEMIN5 | 145 | 169 kDa | | 0.0072 | 0.07 | [] | 5.05E+07 | 2.83E+07 | 5.29E+07 | 6483300 | 2847100 | 1,000.00 |
| 783 | Cluster of Eukaryotic translation initiation factor 4 gan | A0A0U1RQK7 [3] | EIF4G3 | 181 | 195 kDa | TRUE | 0.013 | 0.07 | [] | 3.92E+07 | 2.15E+07 | 2.28E+07 | 1,000.00 | 5833000 | 1,000.00 |
| 781 | Pericentriolar material 1 OS=Homo sapiens OX=9606 | A0A5H1ZRS1 (+1) | PCM1 | 127 | 228 kDa | | 0.078 | 0.07 | [] | 2027300 | 5738100 | 1.01E+07 | 1,000.00 | 585780 | 689810 |
| 778 | Kinesin-like protein OS=Homo sapiens OX=9606 GN= | A0A7I2V5Y5 (+2) | KIF23 | 136 | 112 kDa | | 0.14 | 0.07 | [] | 1,000.00 | 9.59E+07 | 9.60E+07 | 1,000.00 | 1.41E+07 | 1,000.00 |
| 780 | Exosome component 10 OS=Homo sapiens OX=9606 | Q01780 | EXOSC10 | 210 | 101 kDa | | 0.27 | 0.07 | [] | 2.05E+07 | 1061600 | 3518200 | 1787400 | 1,000.00 | 1,000.00 |
| 785 | Calcium homeostasis endoplasmic reticulum protein OS | J3QK89 (+1) | CHERP | 217 | 105 kDa | | 0.4 | 0.07 | [] | 1,000.00 | 3.33E+07 | 1,000.00 | 1,000.00 | 1,000.00 | 2242500 |
| 779 | Chromosome transmission fidelity factor 18 OS=Homo | A0A0D9SF58 (+2) | CHTF18 | 10 | 129 kDa | | 0.41 | 0.07 | [] | 5565000 | 1,000.00 | 1,000.00 | 1,000.00 | 404380 | 1,000.00 |
| 786 | F-actin-capping protein subunit beta OS=Homo sapien | B1AK88 | CAPZB | 366 | 34 kDa | | 0.071 | 0.06 | [] | 7489300 | 4.14E+07 | 3.07E+07 | 5158600 | 1,000.00 | 1,000.00 |

| | | | | | | | | | | | | | | | |
|---|---|---|---|---|---|---|---|---|---|---|---|---|---|---|---|
| 787 | 26S proteasome non-ATPase regulatory subunit 2 OS= | Q13200 | PSMD2 | 271 | 100 kDa | | 0.17 | 0.06 | [] | 1.08E+07 | 1504300 | 3166600 | 1,000.00 | 500710 | 498340 |
| 791 | Transcription intermediary factor 1-beta OS=Homo sa | Q13263 | TRIM28 | 466 | 89 kDa | | 0.0049 | 0.05 | [] | 1.70E+08 | 1.01E+08 | 1.15E+08 | 1.79E+07 | 1554400 | 1,000.00 |
| 788 | Cluster of General transcription and DNA repair factor | A0A2R8Y6W8 [3] | ERCC3 | 2 | 92 kDa | TRUE | 0.021 | 0.05 | [] | 3405300 | 5977800 | 2592300 | 371710 | 259710 | 1,000.00 |
| 792 | cAMP-dependent protein kinase type II-alpha regulato | P13861 | PRKAR2A | 140 | 46 kDa | | 0.11 | 0.05 | [] | 6.36E+07 | 6.97E+07 | 3532300 | 2914800 | 3486800 | 1,000.00 |
| 790 | NF-X1-type zinc finger protein NFXL1 OS=Homo sap | Q6ZNB6 | NFXL1 | 9 | 101 kDa | | 0.27 | 0.05 | [] | 7599400 | 1,000.00 | 3.24E+07 | 1,000.00 | 2044000 | 1,000.00 |
| 789 | E3 ubiquitin-protein ligase RNF216 OS=Homo sapiens | Q9NWF9 | RNF216 | 0 | 99 kDa | | 0.4 | 0.05 | [] | 1.32E+08 | 1,000.00 | 1,000.00 | 6854500 | 1,000.00 | 1,000.00 |
| 800 | Tight junction protein 1 OS=Homo sapiens OX=9606 | A0A087X0K9 (+5) | TJP1 | 134 | 197 kDa | | 0.019 | 0.04 | [] | 3.46E+07 | 3.27E+07 | 1.36E+07 | 1,000.00 | 3113500 | 1,000.00 |
| 796 | Cluster of Ubiquitin associated protein 2 OS=Homo sa | A0A8C8MQ70 [3] | UBAP2 | 134 | 112 kDa | TRUE | 0.036 | 0.04 | [] | 1.94E+07 | 4.74E+07 | 6.62E+07 | 5488400 | 1,000.00 | 1,000.00 |
| 793 | Protein AHNAK2 OS=Homo sapiens OX=9606 GN=/ | Q8IVF2 | AHNAK2 | 69 | 617 kDa | | 0.051 | 0.04 | [] | 9.76E+08 | 1.18E+09 | 2.77E+09 | 1.00E+08 | 5.51E+07 | 6.42E+07 |
| 797 | Cluster of Microtubule-associated protein OS=Homo s | A0A669KB77 [3] | MAP2 | 2 | 217 kDa | TRUE | 0.088 | 0.04 | [] | 1.08E+08 | 3.39E+07 | 3.30E+07 | 1,000.00 | 2761600 | 4334800 |
| 799 | 5'-3' exoribonuclease 1 OS=Homo sapiens OX=9606 ( | Q8IZH2 | XRN1 | 105 | 194 kDa | | 0.13 | 0.04 | [] | 7.68E+07 | 6664500 | 3.74E+07 | 2512600 | 2340000 | 1,000.00 |
| 794 | S1 RNA-binding domain-containing protein 1 OS=Hor | Q8N5C6 | SRBD1 | 44 | 112 kDa | | 0.16 | 0.04 | [] | 2.09E+07 | 103150 | 1.12E+07 | 1,000.00 | 1,000.00 | 1402900 |
| 795 | Cluster of Inositol 1,4,5-trisphosphate receptor type 3 | Q14573 [7] | ITPR3 | 21 | 304 kDa | TRUE | 0.26 | 0.04 | [] | 1499300 | 1.01E+07 | 5.16E+07 | 2728000 | 1,000.00 | 1,000.00 |
| 798 | Protein ECT2 OS=Homo sapiens OX=9606 GN=ECT | Q9H8V3 | ECT2 | 4 | 104 kDa | | 0.37 | 0.04 | [] | 1,000.00 | 1,000.00 | 72639 | 1,000.00 | 1,000.00 | 1,000.00 |
| 801 | Microtubule-associated protein OS=Homo sapiens OX | E7EVA0 | MAP4 | 312 | 245 kDa | | 0.01 | 0.03 | [] | 6.63E+08 | 3.06E+08 | 6.08E+08 | 3.62E+07 | 9870500 | 7301200 |
| 803 | Afadin, adherens junction formation factor OS=Homo | A8MQ02 | AFDN | 116 | 202 kDa | | 0.036 | 0.03 | [] | 2.13E+07 | 7.07E+07 | 7.65E+07 | 4343200 | 1,000.00 | 1,000.00 |
| 802 | NIMA related kinase 9 OS=Homo sapiens OX=9606 ( | A0A7I2V5R1 (+1) | NEK9 | 89 | 108 kDa | | 0.37 | 0.03 | [] | 1,000.00 | 95165 | 1,000.00 | 1,000.00 | 1,000.00 | 1,000.00 |
| 804 | Kinesin-like protein KIF14 OS=Homo sapiens OX=96 | Q15058 | KIF14 | 65 | 186 kDa | | 0.00033 | 0.02 | BioID high, ptprh low | 4104400 | 3834200 | 3095900 | 1,000.00 | 264860 | 1,000.00 |
| 807 | Erbin OS=Homo sapiens OX=9606 GN=ERBIN PE=1 | Q96RT1 | ERBIN | 107 | 158 kDa | | 0.0014 | 0.02 | [] | 9.61E+07 | 1.32E+08 | 1.48E+08 | 6177800 | 1488700 | 1,000.00 |
| 810 | Far upstream element binding protein 1 OS=Homo sap | E9PEB5 (+1) | FUBP1 | 240 | 69 kDa | TRUE | 0.084 | 0.02 | [] | 3110800 | 2.48E+07 | 1.63E+07 | 1,000.00 | 783140 | 1,000.00 |
| 805 | Cluster of E3 SUMO-protein ligase RanBP2 OS=Hom | P49792 [3] | RANBP2 | 268 | 358 kDa | TRUE | 0.086 | 0.02 | [] | 2.58E+07 | 6.42E+07 | 1.37E+08 | 5348000 | 1,000.00 | 1,000.00 |
| 808 | Stromal interaction molecule 1 OS=Homo sapiens OX | E9PNJ4 (+3) | STIM1 | 7 | 58 kDa | | 0.17 | 0.02 | [] | 1.89E+07 | 1311400 | 6431300 | 1,000.00 | 1,000.00 | 538660 |
| 811 | Replication factor C subunit 4 OS=Homo sapiens OX= | P35249 | RFC4 | 162 | 40 kDa | | 0.29 | 0.02 | [] | 1,000.00 | 9493900 | 6.23E+07 | 1,000.00 | 1167400 | 1,000.00 |
| 809 | Cluster of Cytoplasmic FMR1-interacting protein 1 OS | Q7L576 [7] | CYFIP1 | 217 | 145 kDa | TRUE | 0.36 | 0.02 | [] | 4.63E+07 | 1,000.00 | 1481000 | 1,000.00 | 961200 | 1,000.00 |
| 806 | Cluster of Filamin A OS=Homo sapiens OX=9606 GN | Q60FE5 [5] | FLNA | 508 | 278 kDa | TRUE | < 0.00010 | 0.02 | BioID high, ptprh low | 5.37E+09 | 5.94E+09 | 5.81E+09 | 2.54E+08 | 5.88E+07 | 7.36E+07 |
| 813 | E3 ubiquitin-protein ligase UBR5 OS=Homo sapiens ( | O95071 | UBR5 | 44 | 309 kDa | | 0.2 | 0.01 | [] | 3.05E+07 | 4751600 | 4891600 | 466560 | 1,000.00 | 1,000.00 |
| 815 | DNA replication licensing factor MCM7 OS=Homo sa | P33993 | MCM7 | 328 | 81 kDa | | 0.31 | 0.01 | [] | 8999400 | 1.30E+08 | 4809900 | 1,000.00 | 1411600 | 1,000.00 |
| 814 | Epidermal growth factor receptor OS=Homo sapiens ( | P00533 (+2) | EGFR | 32 | 134 kDa | | 0.32 | 0.01 | [] | 1,000.00 | 2.31E+08 | 2.13E+07 | 1,000.00 | 1,000.00 | 2639800 |
| 812 | DNA topoisomerase II binding protein 1 OS=Homo sa | A0A2R8YD63 (+1) | TOPBP1 | 17 | 170 kDa | | 0.37 | 0.01 | [] | 1,000.00 | 1,000.00 | 232470 | 1,000.00 | 1,000.00 | 1,000.00 |
| 817 | Spectrin alpha, non-erythrocytic 1 OS=Homo sapiens ( | A0A0D9SF54 (+2) | SPTAN1 | 324 | 283 kDa | TRUE | 0.0055 | 0.008 | [] | 9.67E+07 | 1.47E+08 | 8.23E+07 | 2552600 | 1,000.00 | 1,000.00 |
| 816 | Nibrin OS=Homo sapiens OX=9606 GN=NBN PE=1 S | O60934 | NBN | 112 | 85 kDa | | 0.37 | 0.008 | [] | 1,000.00 | 1,000.00 | 362850 | 1,000.00 | 1,000.00 | 1,000.00 |
| 819 | RNA helicase OS=Homo sapiens OX=9606 GN=EIF4 | I3L3H2 (+1) | EIF4A3 | 418 | 45 kDa | TRUE | 0.078 | 0.007 | [] | 1.98E+07 | 2.20E+07 | 2409600 | 1,000.00 | 1,000.00 | 329130 |
| 820 | A-kinase anchoring protein 9 OS=Homo sapiens OX=9 | A0A0A0MRF6 (+8) | AKAP9 | 16 | 456 kDa | | 0.0011 | 0.006 | [] | 9.89E+07 | 9.61E+07 | 6.69E+07 | 1,000.00 | 1,000.00 | 1440800 |

| | | | | | | | | | | | | | | | |
|---|---|---|---|---|---|---|---|---|---|---|---|---|---|---|---|
| 821 | Small nuclear ribonucleoprotein-associated proteins B | P14678 | SNRPB | 308 | 25 kDa | | 0.37 | 0.005 | [] | 544520 | 1,000.00 | 1,000.00 | 1,000.00 | 1,000.00 | 1,000.00 |
| 822 | Transcription initiation factor TFIID subunit 4 (Fragm | A0A0G2JRY5 (+1) | TAF4 | 109 | 50 kDa | | 0.37 | 0.005 | [] | 1,000.00 | 1,000.00 | 557420 | 1,000.00 | 1,000.00 | 1,000.00 |
| 823 | Heterogeneous nuclear ribonucleoprotein A/B OS=Ho | D6R9P3 (+2) | HNRNPAB | 442 | 30 kDa | | 0.37 | 0.005 | [] | 644360 | 1,000.00 | 1,000.00 | 1,000.00 | 1,000.00 | 1,000.00 |
| 824 | Cluster of GTP-binding protein 1 OS=Homo sapiens C | O00178 [2] | GTPBP1 | 47 | 72 kDa | TRUE | 0.37 | 0.004 | [] | 708650 | 1,000.00 | 1,000.00 | 1,000.00 | 1,000.00 | 1,000.00 |
| 825 | ATP-dependent RNA helicase DHX29 OS=Homo sapi | Q7Z478 | DHX29 | 109 | 155 kDa | | 0.0044 | 0.003 | [] | 1.27E+08 | 9.05E+07 | 7.01E+07 | 1,000.00 | 951240 | 1,000.00 |
| 826 | Transcription factor CP2 (Fragment) OS=Homo sapier | F8VWL0 (+1) | TFCP2 | 87 | 44 kDa | | 0.37 | 0.003 | [] | 950020 | 1,000.00 | 1,000.00 | 1,000.00 | 1,000.00 | 1,000.00 |
| 827 | Cell growth-regulating nucleolar protein OS=Homo sa | Q9NX58 | LYAR | 94 | 44 kDa | | 0.37 | 0.003 | [] | 1,000.00 | 1008300 | 1,000.00 | 1,000.00 | 1,000.00 | 1,000.00 |
| 828 | General transcription factor 3C polypeptide 4 OS=Hon | Q9UKN8 | GTF3C4 | 150 | 92 kDa | | 0.37 | 0.003 | [] | 1101700 | 1,000.00 | 1,000.00 | 1,000.00 | 1,000.00 | 1,000.00 |
| 829 | NF-kappa-B-repressing factor OS=Homo sapiens OX= | O15226 | NKRF | 168 | 78 kDa | | 0.37 | 0.003 | [] | 1,000.00 | 1,000.00 | 1135500 | 1,000.00 | 1,000.00 | 1,000.00 |
| 836 | Cellular tumor antigen p53 OS=Homo sapiens OX=96 | A0A0U1RQC9 (+1) | TP53 | 131 | 46 kDa | | 0.35 | 0.002 | [] | 1,000.00 | 54638 | 1620700 | 1,000.00 | 1,000.00 | 1,000.00 |
| 830 | Dimethyladenosine transferase 2, mitochondrial OS=H | Q9H5Q4 | TFB2M | 9 | 45 kDa | | 0.37 | 0.002 | [] | 1265500 | 1,000.00 | 1,000.00 | 1,000.00 | 1,000.00 | 1,000.00 |
| 831 | Cell division cycle protein 20 homolog OS=Homo sapi | Q12834 | CDC20 | 9 | 55 kDa | | 0.37 | 0.002 | [] | 1372200 | 1,000.00 | 1,000.00 | 1,000.00 | 1,000.00 | 1,000.00 |
| 832 | Talin-1 OS=Homo sapiens OX=9606 GN=TLN1 PE= | Q9Y490 | TLN1 | 217 | 270 kDa | TRUE | 0.37 | 0.002 | [] | 1,000.00 | 1,000.00 | 1496000 | 1,000.00 | 1,000.00 | 1,000.00 |
| 833 | DNA polymerase delta subunit 3 OS=Homo sapiens O | Q15054 | POLD3 | 86 | 51 kDa | | 0.37 | 0.002 | [] | 1,000.00 | 1499200 | 1,000.00 | 1,000.00 | 1,000.00 | 1,000.00 |
| 834 | Ankyrin repeat domain-containing protein 50 OS=Hon | Q9ULJ7 | ANKRD50 | 2 | 156 kDa | | 0.37 | 0.002 | [] | 1635700 | 1,000.00 | 1,000.00 | 1,000.00 | 1,000.00 | 1,000.00 |
| 835 | Transcription elongation regulator 1 OS=Homo sapien | A0A7P0T8N8 (+2) | TCERG1 | 183 | 126 kDa | | 0.37 | 0.002 | [] | 1,000.00 | 1668800 | 1,000.00 | 1,000.00 | 1,000.00 | 1,000.00 |
| 837 | non-specific serine/threonine protein kinase OS=Homo | F5GWT4 (+1) | WNK1 | 109 | 225 kDa | | 0.37 | 0.002 | [] | 1686600 | 1,000.00 | 1,000.00 | 1,000.00 | 1,000.00 | 1,000.00 |
| 838 | PCF11 cleavage and polyadenylation factor subunit (Fr | E9PQ01 (+1) | PCF11 | 80 | 112 kDa | | 0.37 | 0.002 | [] | 1,000.00 | 1,000.00 | 1751900 | 1,000.00 | 1,000.00 | 1,000.00 |
| 844 | DnaJ heat shock protein family (Hsp40) member B1 O | A0A7I2V3K7 (+1) | DNAJB1 | 143 | 30 kDa | | 0.013 | 0.001 | [] | 1260200 | 1156200 | 519190 | 1,000.00 | 1,000.00 | 1,000.00 |
| 840 | Voltage-dependent anion-selective channel protein 3 O | Q9Y277 | VDAC3 | 226 | 31 kDa | | 0.12 | 0.001 | [] | 1200900 | 996160 | 1,000.00 | 1,000.00 | 1,000.00 | 1,000.00 |
| 845 | Cluster of CTP synthase OS=Homo sapiens OX=9606 | A0A3B3IRQ8 [3] | CTPS1 | 312 | 50 kDa | TRUE | 0.12 | 0.001 | [] | 1549000 | 1,000.00 | 1409800 | 1,000.00 | 1,000.00 | 1,000.00 |
| 846 | Zinc finger and BTB domain-containing protein 10 OS | Q96DT7 | ZBTB10 | 24 | 95 kDa | | 0.19 | 0.001 | [] | 874510 | 2260600 | 1,000.00 | 1,000.00 | 1,000.00 | 1,000.00 |
| 839 | Dehydrogenase/reductase 7B (Fragment) OS=Homo sa | J3KRS1 | DHRS7B | 26 | 14 kDa | | 0.37 | 0.001 | [] | 1,000.00 | 1,000.00 | 2004900 | 1,000.00 | 1,000.00 | 1,000.00 |
| 841 | Prune homolog 2 with BCH domain OS=Homo sapien | A0A088AWP5 (+3) | PRUNE2 | 1 | 338 kDa | | 0.37 | 0.001 | [] | 1,000.00 | 1,000.00 | 2673100 | 1,000.00 | 1,000.00 | 1,000.00 |
| 842 | Holliday junction recognition protein (Fragment) OS=I | C9JWC4 (+1) | HJURP | 2 | 64 kDa | | 0.37 | 0.001 | [] | 1,000.00 | 1,000.00 | 2685800 | 1,000.00 | 1,000.00 | 1,000.00 |
| 843 | Exosome complex component RRP40 OS=Homo sapi | Q9NQT5 | EXOSC3 | 137 | 30 kDa | | 0.37 | 0.001 | [] | 2705300 | 1,000.00 | 1,000.00 | 1,000.00 | 1,000.00 | 1,000.00 |
| 850 | Prohibitin OS=Homo sapiens OX=9606 GN=PHB2 PE | J3KPX7 (+1) | PHB2 | 243 | 29 kDa | | 0.17 | 0.0009 | [] | 1,000.00 | 2451200 | 1070800 | 1,000.00 | 1,000.00 | 1,000.00 |
| 849 | FSHD region gene 1 (Fragment) OS=Homo sapiens O | E9PRR7 (+1) | FRG1 | 138 | 13 kDa | | 0.37 | 0.0009 | [] | 1,000.00 | 1,000.00 | 3433100 | 1,000.00 | 1,000.00 | 1,000.00 |
| 851 | Nuclear pore complex protein Nup153 OS=Homo sapi | P49790 | NUP153 | 200 | 154 kDa | | 0.37 | 0.0009 | [] | 3526800 | 1,000.00 | 1,000.00 | 1,000.00 | 1,000.00 | 1,000.00 |
| 854 | Peptidyl-prolyl cis-trans isomerase FKBP8 OS=Homo | Q14318 | FKBP8 | 82 | 45 kDa | | 0.25 | 0.0008 | [] | 1,000.00 | 714210 | 3193000 | 1,000.00 | 1,000.00 | 1,000.00 |
| 852 | Cluster of Dedicator of cytokinesis 10 OS=Homo sapi | A0A2R8YD85 [3] | DOCK10 | 4 | 251 kDa | TRUE | 0.37 | 0.0008 | [] | 1,000.00 | 3592400 | 1,000.00 | 1,000.00 | 1,000.00 | 1,000.00 |
| 853 | RuvB-like 1 OS=Homo sapiens OX=9606 GN=RUVB | Q9Y265 | RUVBL1 | 378 | 50 kDa | | 0.37 | 0.0008 | [] | 3653400 | 1,000.00 | 1,000.00 | 1,000.00 | 1,000.00 | 1,000.00 |
| 863 | Cluster of CCR4-NOT transcription complex subunit 1 | A5YKK6 [2] | CNOT1 | 130 | 267 kDa | TRUE | 0.046 | 0.0007 | [] | 9.85E+07 | 2.05E+08 | 3.68E+08 | 1,000.00 | 440030 | 1,000.00 |

| | | | | | | | | | | | | | | | |
|---|---|---|---|---|---|---|---|---|---|---|---|---|---|---|---|
| 859 | Extended synaptotagmin 2 OS=Homo sapiens OX=96( | A0A499FIX8 (+3) | ESYT2 | 77 | 94 kDa | | 0.12 | 0.0007 | [] | 1916000 | 2399900 | 1,000.00 | 1,000.00 | 1,000.00 | 1,000.00 |
| 855 | Kinase D interacting substrate 220 OS=Homo sapiens | A0A1W2PPB7 (+3) | KIDINS220 | 58 | 147 kDa | | 0.17 | 0.0007 | [] | 1243700 | 2802100 | 1,000.00 | 1,000.00 | 1,000.00 | 1,000.00 |
| 857 | TAR DNA-binding protein 43 (Fragment) OS=Homo s | A0A087WX29 (+1) | TARDBP | 324 | 27 kDa | | 0.27 | 0.0007 | [] | 653370 | 3434300 | 1,000.00 | 1,000.00 | 1,000.00 | 1,000.00 |
| 864 | Cluster of VPS53 subunit of GARP complex OS=Hom | A0A7P0T874 [8] | VPS53 | 23 | 76 kDa | TRUE | 0.33 | 0.0007 | [] | 4227400 | 1,000.00 | 325860 | 1,000.00 | 1,000.00 | 1,000.00 |
| 856 | Eukaryotic translation initiation factor 4B OS=Homo s | E7EX17 (+1) | EIF4B | 344 | 70 kDa | | 0.37 | 0.0007 | [] | 1,000.00 | 1,000.00 | 4083000 | 1,000.00 | 1,000.00 | 1,000.00 |
| 858 | Cluster of Zinc finger CCCH-type with G patch domai | Q8N5A5 [2] | ZGPAT | 7 | 57 kDa | TRUE | 0.37 | 0.0007 | [] | 1,000.00 | 1,000.00 | 4131000 | 1,000.00 | 1,000.00 | 1,000.00 |
| 860 | Catenin beta 1 OS=Homo sapiens OX=9606 GN=CTN | A0A2R8Y543 (+5) | CTNNB1 | 78 | 78 kDa | | 0.37 | 0.0007 | [] | 1,000.00 | 1,000.00 | 4360800 | 1,000.00 | 1,000.00 | 1,000.00 |
| 861 | Signal transducer and activator of transcription OS=H( | A0A669KB53 (+6) | STAT1 | 109 | 91 kDa | | 0.37 | 0.0007 | [] | 4463300 | 1,000.00 | 1,000.00 | 1,000.00 | 1,000.00 | 1,000.00 |
| 862 | U5 small nuclear ribonucleoprotein 200 kDa helicase C | O75643 | SNRNP200 | 333 | 245 kDa | | 0.37 | 0.0007 | [] | 1,000.00 | 1,000.00 | 4517400 | 1,000.00 | 1,000.00 | 1,000.00 |
| 865 | Serine/threonine-protein kinase PLK1 OS=Homo sapi | P53350 | PLK1 | 54 | 68 kDa | | 0.37 | 0.0007 | [] | 1,000.00 | 4569600 | 1,000.00 | 1,000.00 | 1,000.00 | 1,000.00 |
| 873 | Treacle protein OS=Homo sapiens OX=9606 GN=TC( | Q13428 | TCOF1 | 332 | 152 kDa | | 0.12 | 0.0006 | [] | 1,000.00 | 2717600 | 2652600 | 1,000.00 | 1,000.00 | 1,000.00 |
| 871 | Paraspeckle component 1 OS=Homo sapiens OX=960( | Q8WXF1 | PSPC1 | 217 | 59 kDa | | 0.13 | 0.0006 | [] | 2997000 | 2075000 | 1,000.00 | 1,000.00 | 1,000.00 | 1,000.00 |
| 867 | Double-strand break repair protein OS=Homo sapiens | F8W7U8 (+1) | MRE11 | 191 | 81 kDa | | 0.31 | 0.0006 | [] | 1,000.00 | 4348200 | 444700 | 1,000.00 | 1,000.00 | 1,000.00 |
| 866 | Transcription initiation factor TFIID subunit 9 OS=Ho | Q16594 | TAF9 | 94 | 29 kDa | | 0.37 | 0.0006 | [] | 1,000.00 | 1,000.00 | 4686500 | 1,000.00 | 1,000.00 | 1,000.00 |
| 868 | Cluster of Protein transport protein SEC23 OS=Homo | F5H365 [3] | SEC23A | 125 | 83 kDa | TRUE | 0.37 | 0.0006 | [] | 4953100 | 1,000.00 | 1,000.00 | 1,000.00 | 1,000.00 | 1,000.00 |
| 869 | AP-3 complex subunit delta-1 OS=Homo sapiens OX= | O14617 (+1) | AP3D1 | 198 | 130 kDa | | 0.37 | 0.0006 | [] | 5010400 | 1,000.00 | 1,000.00 | 1,000.00 | 1,000.00 | 1,000.00 |
| 870 | DnaJ homolog subfamily A member 1 OS=Homo sapie | P31689 | DNAJA1 | 243 | 45 kDa | | 0.37 | 0.0006 | [] | 5027900 | 1,000.00 | 1,000.00 | 1,000.00 | 1,000.00 | 1,000.00 |
| 872 | Signal transducer and activator of transcription OS=H( | A0A7I2V2G1 (+11) | STAT3 | 136 | 88 kDa | | 0.37 | 0.0006 | [] | 1,000.00 | 5302900 | 1,000.00 | 1,000.00 | 1,000.00 | 1,000.00 |
| 874 | Glutathione peroxidase 1 OS=Homo sapiens OX=960( | P07203 | GPX1 | 68 | 22 kDa | | 0.37 | 0.0006 | [] | 1,000.00 | 1,000.00 | 5416600 | 1,000.00 | 1,000.00 | 1,000.00 |
| 878 | Tyrosine-protein phosphatase non-receptor type 13 OS | Q12923 | PTPN13 | 61 | 277 kDa | | 0.12 | 0.0005 | [] | 2766000 | 1,000.00 | 3306200 | 1,000.00 | 1,000.00 | 1,000.00 |
| 877 | DNA mismatch repair protein Msh2 OS=Homo sapien: | P43246 | MSH2 | 202 | 105 kDa | | 0.2 | 0.0005 | [] | 1453700 | 4530200 | 42811 | 1,000.00 | 1,000.00 | 1,000.00 |
| 875 | IQ motif containing GTPase activating protein 3 OS=F | F2Z2E2 (+1) | IQGAP3 | 22 | 179 kDa | | 0.37 | 0.0005 | [] | 5648800 | 1,000.00 | 1,000.00 | 1,000.00 | 1,000.00 | 1,000.00 |
| 876 | DDB1- and CUL4-associated factor 1 OS=Homo sapi( | Q9Y4B6 | DCAF1 | 7 | 169 kDa | | 0.37 | 0.0005 | [] | 5661400 | 1,000.00 | 1,000.00 | 1,000.00 | 1,000.00 | 1,000.00 |
| 879 | Trio Rho guanine nucleotide exchange factor OS=Hon | E7EWP2 (+1) | TRIO | 62 | 263 kDa | TRUE | 0.37 | 0.0005 | [] | 6527300 | 1,000.00 | 1,000.00 | 1,000.00 | 1,000.00 | 1,000.00 |
| 881 | Mediator of RNA polymerase II transcription subunit 1 | Q93074 (+1) | MED12 | 12 | 243 kDa | | 0.047 | 0.0004 | [] | 843510 | 3584600 | 2311900 | 1,000.00 | 1,000.00 | 1,000.00 |
| 891 | Cluster of Chromodomain-helicase-DNA-binding prot( | Q9HCK8 [4] | CHD8 | 118 | 291 kDa | TRUE | 0.091 | 0.0004 | [] | 3882900 | 298100 | 4382800 | 1,000.00 | 1,000.00 | 1,000.00 |
| 885 | ATP-dependent RNA helicase DDX42 OS=Homo sapi | Q86XP3 | DDX42 | 199 | 103 kDa | | 0.12 | 0.0004 | [] | 1,000.00 | 3572800 | 3872300 | 1,000.00 | 1,000.00 | 1,000.00 |
| 882 | Leukocyte receptor cluster member 8 OS=Homo sapie | A0A087WUE4 (+3) | LENG8 | 104 | 96 kDa | | 0.14 | 0.0004 | [] | 2663400 | 4256900 | 1,000.00 | 1,000.00 | 1,000.00 | 1,000.00 |
| 886 | Laminin subunit alpha-5 OS=Homo sapiens OX=9606 | O15230 | LAMA5 | 11 | 400 kDa | | 0.17 | 0.0004 | [] | 330850 | 1756700 | 5432100 | 1,000.00 | 1,000.00 | 1,000.00 |
| 883 | Cluster of Homeobox protein cut-like OS=Homo sapie | A0A2R8Y852 [4] | CUX1 | 98 | 159 kDa | TRUE | 0.28 | 0.0004 | [] | 5921800 | 1,000.00 | 1008200 | 1,000.00 | 1,000.00 | 1,000.00 |
| 880 | MAX dimerization protein MGA (Fragment) OS=Hon | A0A8C8P5L8 | MGA | 124 | 283 kDa | | 0.29 | 0.0004 | [] | 801430 | 1,000.00 | 5911300 | 1,000.00 | 1,000.00 | 1,000.00 |
| 884 | Cell division cycle and apoptosis regulator protein 1 O: | Q8IX12 | CCAR1 | 103 | 133 kDa | | 0.33 | 0.0004 | [] | 494510 | 1,000.00 | 6661000 | 1,000.00 | 1,000.00 | 1,000.00 |
| 887 | SWI/SNF related, matrix associated, actin dependent r | F8VXC8 (+1) | SMARCC2 | 164 | 136 kDa | | 0.37 | 0.0004 | [] | 1,000.00 | 7714500 | 1,000.00 | 1,000.00 | 1,000.00 | 1,000.00 |

| | | | | | | | | | | | | | | | |
|---|---|---|---|---|---|---|---|---|---|---|---|---|---|---|---|
| 889 | Amino acid transporter OS=Homo sapiens OX=9606 ( | M0QXM4 (+1) | SLC1A5 | 236 | 39 kDa | | 0.37 | 0.0004 | [] | 8005300 | 1,000.00 | 1,000.00 | 1,000.00 | 1,000.00 | 1,000.00 |
| 890 | Serine/threonine-protein phosphatase 6 regulatory anky | Q8NB46 | ANKRD52 | 10 | 115 kDa | | 0.37 | 0.0004 | [] | 8526700 | 1,000.00 | 1,000.00 | 1,000.00 | 1,000.00 | 1,000.00 |
| 905 | UDP-N-acetylglucosamine--peptide N-acetylglucosam | O15294 | OGT | 162 | 117 kDa | | 0.11 | 0.0003 | [] | 1680800 | 2247300 | 7578800 | 1,000.00 | 1,000.00 | 1,000.00 |
| 906 | FK506-binding protein 15 OS=Homo sapiens OX=960 | Q5T1M5 | FKBP15 | 46 | 134 kDa | | 0.11 | 0.0003 | [] | 7679800 | 2233900 | 1782800 | 1,000.00 | 1,000.00 | 1,000.00 |
| 895 | 26S proteasome regulatory subunit 4 OS=Homo sapier | P62191 | PSMC1 | 195 | 49 kDa | TRUE | 0.12 | 0.0003 | [] | 5850200 | 1465200 | 1389700 | 1,000.00 | 1,000.00 | 1,000.00 |
| 900 | Metastasis-associated protein MTA2 OS=Homo sapier | O94776 | MTA2 | 206 | 75 kDa | TRUE | 0.13 | 0.0003 | [] | 4102700 | 1,000.00 | 6072300 | 1,000.00 | 1,000.00 | 1,000.00 |
| 892 | Nucleolar and spindle-associated protein 1 OS=Homo | Q9BXS6 | NUSAP1 | 52 | 49 kDa | | 0.15 | 0.0003 | [] | 1439300 | 1089000 | 6093500 | 1,000.00 | 1,000.00 | 1,000.00 |
| 904 | Proteasome 26S subunit, ATPase 3 (Fragment) OS=H | E9PKD5 (+3) | PSMC3 | 225 | 35 kDa | | 0.15 | 0.0003 | [] | 7384300 | 3960900 | 1,000.00 | 1,000.00 | 1,000.00 | 1,000.00 |
| 894 | Sickle tail protein homolog OS=Homo sapiens OX=96 | Q5T5P2 | KIAA1217 | 46 | 214 kDa | | 0.16 | 0.0003 | [] | 773920 | 1712900 | 6212600 | 1,000.00 | 1,000.00 | 1,000.00 |
| 899 | Round spermatid basic protein 1 OS=Homo sapiens O | A0A087WWP8 (+1) | RSBN1 | 50 | 85 kDa | TRUE | 0.19 | 0.0003 | [] | 1,000.00 | 2790000 | 7156700 | 1,000.00 | 1,000.00 | 1,000.00 |
| 903 | Vacuolar protein sorting-associated protein 51 homolo | Q9UID3 | VPS51 | 44 | 86 kDa | | 0.2 | 0.0003 | [] | 8363800 | 1,000.00 | 2945900 | 1,000.00 | 1,000.00 | 1,000.00 |
| 902 | Cluster of MutL homolog 1 (Fragment) OS=Homo sap | A0A669KAW3 [2] | MLH1 | 105 | 77 kDa | TRUE | 0.22 | 0.0003 | [] | 2517600 | 8631700 | 1,000.00 | 1,000.00 | 1,000.00 | 1,000.00 |
| 893 | Cytoskeleton-associated protein 5 OS=Homo sapiens ( | Q14008 | CKAP5 | 196 | 226 kDa | | 0.34 | 0.0003 | [] | 1,000.00 | 460940 | 8227700 | 1,000.00 | 1,000.00 | 1,000.00 |
| 896 | Acidic leucine-rich nuclear phosphoprotein 32 family n | Q92688 | ANP32B | 114 | 29 kDa | | 0.37 | 0.0003 | [] | 8862400 | 1,000.00 | 1,000.00 | 1,000.00 | 1,000.00 | 1,000.00 |
| 897 | GRAM domain containing 1A OS=Homo sapiens OX= | M0QZ12 (+1) | GRAMD1A | 26 | 90 kDa | | 0.37 | 0.0003 | [] | 1,000.00 | 1,000.00 | 9021200 | 1,000.00 | 1,000.00 | 1,000.00 |
| 898 | Eukaryotic elongation factor 2 kinase OS=Homo sapie | O00418 | EEF2K | 30 | 82 kDa | | 0.37 | 0.0003 | [] | 9024500 | 1,000.00 | 1,000.00 | 1,000.00 | 1,000.00 | 1,000.00 |
| 929 | WASH complex subunit 5 OS=Homo sapiens OX=96( | E7EQI7 (+1) | WASHC5 | 12 | 117 kDa | | 0.0041 | 0.0002 | [] | 7583400 | 7184300 | 4191100 | 1,000.00 | 1,000.00 | 1,000.00 |
| 928 | CD2-associated protein OS=Homo sapiens OX=9606 ( | Q9Y5K6 | CD2AP | 147 | 71 kDa | | 0.0061 | 0.0002 | [] | 4062200 | 6676300 | 8116400 | 1,000.00 | 1,000.00 | 1,000.00 |
| 912 | Vacuolar protein sorting-associated protein 52 homolo | Q8N1B4 | VPS52 | 12 | 82 kDa | | 0.0063 | 0.0002 | [] | 2784100 | 4774600 | 5599000 | 1,000.00 | 1,000.00 | 1,000.00 |
| 916 | Splicing factor 1 OS=Homo sapiens OX=9606 GN=SF | A0A7P0T9U7 (+1) | SF1 | 256 | 80 kDa | | 0.0082 | 0.0002 | [] | 6600400 | 4260300 | 3335600 | 1,000.00 | 1,000.00 | 1,000.00 |
| 924 | Cluster of Dedicator of cytokinesis protein 7 OS=Hom | Q96N67 [2] | DOCK7 | 103 | 243 kDa | TRUE | 0.033 | 0.0002 | [] | 8674400 | 5289500 | 2684600 | 1,000.00 | 1,000.00 | 1,000.00 |
| 919 | Replication protein A 70 kDa DNA-binding subunit OS | P27694 | RPA1 | 242 | 68 kDa | | 0.14 | 0.0002 | [] | 1.08E+07 | 2653000 | 2067500 | 1,000.00 | 1,000.00 | 1,000.00 |
| 922 | Cluster of T-complex protein 1 subunit theta OS=Hom | P50990 [2] | CCT8 | 451 | 60 kDa | TRUE | 0.14 | 0.0002 | [] | 976620 | 4191500 | 1.10E+07 | 1,000.00 | 1,000.00 | 1,000.00 |
| 925 | Anillin OS=Homo sapiens OX=9606 GN=ANLN PE= | Q9NQW6 | ANLN | 72 | 124 kDa | | 0.15 | 0.0002 | [] | 3564000 | 1903900 | 1.27E+07 | 1,000.00 | 1,000.00 | 1,000.00 |
| 921 | Protein TANC1 OS=Homo sapiens OX=9606 GN=TA | Q9C0D5 | TANC1 | 38 | 202 kDa | | 0.18 | 0.0002 | [] | 1,000.00 | 4865400 | 1.13E+07 | 1,000.00 | 1,000.00 | 1,000.00 |
| 917 | Aftiphilin OS=Homo sapiens OX=9606 GN=AFTPH F | Q6ULP2 | AFTPH | 34 | 102 kDa | | 0.21 | 0.0002 | [] | 1.10E+07 | 1,000.00 | 3454500 | 1,000.00 | 1,000.00 | 1,000.00 |
| 907 | DNA-directed RNA polymerase subunit beta OS=Hon | C9J2Y9 (+1) | POLR2B | 130 | 133 kDa | | 0.24 | 0.0002 | [] | 2462900 | 1,000.00 | 1.03E+07 | 1,000.00 | 1,000.00 | 1,000.00 |
| 911 | Cell division cycle 73 OS=Homo sapiens OX=9606 GN | A0A3B3IRP5 (+1) | CDC73 | 162 | 54 kDa | | 0.24 | 0.0002 | [] | 1.05E+07 | 1,000.00 | 2552700 | 1,000.00 | 1,000.00 | 1,000.00 |
| 913 | Integrator complex subunit 1 OS=Homo sapiens OX=9 | Q8N201 | INTS1 | 28 | 244 kDa | | 0.24 | 0.0002 | [] | 1356400 | 1.08E+07 | 1138600 | 1,000.00 | 1,000.00 | 1,000.00 |
| 927 | Structural maintenance of chromosomes protein 2 OS= | O95347 | SMC2 | 194 | 136 kDa | | 0.27 | 0.0002 | [] | 1,000.00 | 2956400 | 1.56E+07 | 1,000.00 | 1,000.00 | 1,000.00 |
| 909 | Cluster of AP-2 complex subunit alpha-1 OS=Homo sa | O95782 [3] | AP2A1 | 80 | 108 kDa | TRUE | 0.29 | 0.0002 | [] | 1.14E+07 | 467160 | 1077800 | 1,000.00 | 1,000.00 | 1,000.00 |
| 908 | Splicing factor 45 OS=Homo sapiens OX=9606 GN=F | Q96I25 | RBM17 | 206 | 45 kDa | | 0.31 | 0.0002 | [] | 1166700 | 1,000.00 | 1.17E+07 | 1,000.00 | 1,000.00 | 1,000.00 |
| 926 | Cluster of Acyl-coenzyme A thioesterase 9, mitochond | Q9Y305 [2] | ACOT9 | 63 | 50 kDa | TRUE | 0.31 | 0.0002 | [] | 1,000.00 | 1.66E+07 | 1657200 | 1,000.00 | 1,000.00 | 1,000.00 |

| | | | | | | | | | | | | | | | |
|---|---|---|---|---|---|---|---|---|---|---|---|---|---|---|---|
| 914 | Elongin-A OS=Homo sapiens OX=9606 GN=ELOA P | Q14241 (+1) | ELOA | 122 | 90 kDa | | 0.32 | 0.0002 | [] | 1.23E+07 | 1115800 | 1,000.00 | 1,000.00 | 1,000.00 | 1,000.00 |
| 923 | Golgin subfamily A member 5 OS=Homo sapiens OX= | Q8TBA6 | GOLGA5 | 67 | 83 kDa | | 0.35 | 0.0002 | [] | 1,000.00 | 564890 | 1.58E+07 | 1,000.00 | 1,000.00 | 1,000.00 |
| 910 | TPD52 like 2 OS=Homo sapiens OX=9606 GN=TPD5 | A0A087WYR3 | TPD52L2 | 68 | 24 kDa | | 0.37 | 0.0002 | [] | 1,000.00 | 1,000.00 | 1.30E+07 | 1,000.00 | 1,000.00 | 1,000.00 |
| 915 | RNA helicase OS=Homo sapiens OX=9606 GN=DDX | A0A0C4DG89 | DDX46 | 302 | 117 kDa | | 0.37 | 0.0002 | [] | 1,000.00 | 1,000.00 | 1.37E+07 | 1,000.00 | 1,000.00 | 1,000.00 |
| 918 | Polyhomeotic-like protein 3 OS=Homo sapiens OX=9 | Q8NDX5 | PHC3 | 46 | 106 kDa | | 0.37 | 0.0002 | [] | 1,000.00 | 1,000.00 | 1.45E+07 | 1,000.00 | 1,000.00 | 1,000.00 |
| 920 | Proline-rich protein PRCC OS=Homo sapiens OX=96 | Q92733 | PRCC | 81 | 52 kDa | | 0.37 | 0.0002 | [] | 1.58E+07 | 1,000.00 | 1,000.00 | 1,000.00 | 1,000.00 | 1,000.00 |
| 930 | Bifunctional 3'-5' exonuclease/ATP-dependent helicase | Q14191 | WRN | 52 | 162 kDa | | 0.0063 | 0.0001 | [] | 4475000 | 8902300 | 6755500 | 1,000.00 | 1,000.00 | 1,000.00 |
| 946 | G-patch domain and KOW motifs-containing protein C | Q92917 | GPKOW | 107 | 52 kDa | | 0.0068 | 0.0001 | [] | 9707700 | 5725100 | 1.17E+07 | 1,000.00 | 1,000.00 | 1,000.00 |
| 945 | Ran GTPase-activating protein 1 OS=Homo sapiens O | P46060 | RANGAP1 | 283 | 64 kDa | | 0.015 | 0.0001 | [] | 6615200 | 1.33E+07 | 6930500 | 1,000.00 | 1,000.00 | 1,000.00 |
| 939 | WD repeat and HMG-box DNA-binding protein 1 OS= | O75717 | WDHD1 | 107 | 126 kDa | | 0.022 | 0.0001 | [] | 8099600 | 4259000 | 1.21E+07 | 1,000.00 | 1,000.00 | 1,000.00 |
| 938 | CCR4-NOT transcription complex subunit 9 OS=Hom | Q92600 | CNOT9 | 31 | 34 kDa | | 0.052 | 0.0001 | [] | 4624300 | 1.38E+07 | 5509000 | 1,000.00 | 1,000.00 | 1,000.00 |
| 944 | Cluster of Forkhead box protein K1 OS=Homo sapiens | P85037 [2] | FOXK1 | 38 | 75 kDa | TRUE | 0.093 | 0.0001 | [] | 1.11E+07 | 1.40E+07 | 971310 | 1,000.00 | 1,000.00 | 1,000.00 |
| 936 | T-complex protein 1 subunit beta OS=Homo sapiens C | P78371 | CCT2 | 428 | 57 kDa | | 0.12 | 0.0001 | [] | 1.15E+07 | 1.20E+07 | 1,000.00 | 1,000.00 | 1,000.00 | 1,000.00 |
| 942 | G3BP stress granule assembly factor 2 OS=Homo sapi | A0A7I2YQD9 (+1) | G3BP2 | 214 | 48 kDa | TRUE | 0.12 | 0.0001 | [] | 1.37E+07 | 1,000.00 | 1.13E+07 | 1,000.00 | 1,000.00 | 1,000.00 |
| 931 | ATP-citrate synthase OS=Homo sapiens OX=9606 GN | P53396 | ACLY | 296 | 121 kDa | | 0.13 | 0.0001 | [] | 8314600 | 1.26E+07 | 1,000.00 | 1,000.00 | 1,000.00 | 1,000.00 |
| 949 | Eukaryotic translation initiation factor 4 gamma 2 OS= | P78344 | EIF4G2 | 119 | 102 kDa | | 0.13 | 0.0001 | [] | 1,000.00 | 1.23E+07 | 1.84E+07 | 1,000.00 | 1,000.00 | 1,000.00 |
| 948 | Wings apart-like protein homolog OS=Homo sapiens C | Q7Z5K2 | WAPL | 123 | 133 kDa | | 0.19 | 0.0001 | [] | 1,000.00 | 7860100 | 2.10E+07 | 1,000.00 | 1,000.00 | 1,000.00 |
| 932 | Chloride channel CLIC-like protein 1 OS=Homo sapie | A0A6Q8PEZ7 (+7) | CLCC1 | 103 | 64 kDa | | 0.25 | 0.0001 | [] | 1032600 | 2631500 | 1.80E+07 | 1,000.00 | 1,000.00 | 1,000.00 |
| 933 | WASH complex subunit 2C OS=Homo sapiens OX=9 | A0A096LPC5 (+3) | WASHC2C | 94 | 147 kDa | | 0.25 | 0.0001 | [] | 1.85E+07 | 1191700 | 2484000 | 1,000.00 | 1,000.00 | 1,000.00 |
| 940 | Cluster of DNA helicase OS=Homo sapiens OX=9606 | A0A0C4DGG9 [19] | CHD4 | 256 | 220 kDa | TRUE | 0.26 | 0.0001 | [] | 2808100 | 2.07E+07 | 1266400 | 1,000.00 | 1,000.00 | 1,000.00 |
| 947 | Syndecan-4 OS=Homo sapiens OX=9606 GN=SDC4 I | P31431 | SDC4 | 6 | 22 kDa | | 0.26 | 0.0001 | [] | 2606200 | 1725500 | 2.37E+07 | 1,000.00 | 1,000.00 | 1,000.00 |
| 934 | Nuclear receptor corepressor 2 OS=Homo sapiens OX | C9J0Q5 (+3) | NCOR2 | 56 | 273 kDa | | 0.27 | 0.0001 | [] | 1,000.00 | 3420900 | 1.95E+07 | 1,000.00 | 1,000.00 | 1,000.00 |
| 943 | T-complex protein 1 subunit epsilon OS=Homo sapien | P48643 | CCT5 | 457 | 60 kDa | | 0.3 | 0.0001 | [] | 2.29E+07 | 1,000.00 | 2907700 | 1,000.00 | 1,000.00 | 1,000.00 |
| 935 | Chromosome-associated kinesin KIF4A OS=Homo sa | O95239 | KIF4A | 136 | 140 kDa | | 0.31 | 0.0001 | [] | 2353800 | 1,000.00 | 2.08E+07 | 1,000.00 | 1,000.00 | 1,000.00 |
| 950 | GATA zinc finger domain containing 2B OS=Homo sa | A0A0U1RRM1 (+1) | GATAD2B | 178 | 63 kDa | | 0.34 | 0.0001 | [] | 1,000.00 | 1501500 | 2.94E+07 | 1,000.00 | 1,000.00 | 1,000.00 |
| 937 | A-kinase anchor protein 1, mitochondrial OS=Homo sa | Q92667 | AKAP1 | 120 | 97 kDa | | < 0.00010 | 0.0001 | BioID high, ptprh low | 8018800 | 7199300 | 8458000 | 1,000.00 | 1,000.00 | 1,000.00 |
| 941 | MICOS complex subunit MIC60 OS=Homo sapiens O | B9A067 (+3) | IMMT | 93 | 79 kDa | | < 0.00010 | 0.0001 | BioID high, ptprh low | 7256700 | 8942300 | 8632800 | 1,000.00 | 1,000.00 | 1,000.00 |
| 952 | ELKS/RAB6-interacting/CAST family member 1 OS=I | G8JLD3 (+2) | ERC1 | 115 | 125 kDa | | 0.00017 | 0.00009 | BioID high, ptprh low | 9843700 | 1.09E+07 | 1.26E+07 | 1,000.00 | 1,000.00 | 1,000.00 |
| 953 | Telomere-associated protein RIF1 OS=Homo sapiens C | Q5UIP0 | RIF1 | 170 | 274 kDa | | 0.1 | 0.00009 | [] | 2256600 | 1.15E+07 | 2.13E+07 | 1,000.00 | 1,000.00 | 1,000.00 |
| 951 | Cluster of Protein scribble homolog OS=Homo sapiens | A0A0G2JNZ2 [5] | SCRIB | 109 | 175 kDa | TRUE | 0.18 | 0.00009 | [] | 2.30E+07 | 1,000.00 | 9577600 | 1,000.00 | 1,000.00 | 1,000.00 |
| 956 | Crystallin beta-gamma domain containing 1 OS=Homo | A0A0J9YWL0 (+1) | CRYBG1 | 11 | 232 kDa | TRUE | 0.00029 | 0.00008 | BioID high, ptprh low | 1.05E+07 | 1.37E+07 | 1.38E+07 | 1,000.00 | 1,000.00 | 1,000.00 |
| 957 | Cluster of CCR4-NOT transcription complex subunit 3 | B7Z6J7 [4] | CNOT3 | 23 | 77 kDa | TRUE | 0.008 | 0.00008 | [] | 1.23E+07 | 9148400 | 1.84E+07 | 1,000.00 | 1,000.00 | 1,000.00 |
| 954 | Protein lin-54 homolog OS=Homo sapiens OX=9606 C | Q6MZP7 | LIN54 | 37 | 79 kDa | | 0.17 | 0.00008 | [] | 5653000 | 4032000 | 2.71E+07 | 1,000.00 | 1,000.00 | 1,000.00 |

| | | | | | | | | | | | | | | | |
|---|---|---|---|---|---|---|---|---|---|---|---|---|---|---|---|
| 955 | Dyslexia-associated protein KIAA0319-like protein (Fr | E7EN73 (+1) | KIAA0319L | 0 | 113 kDa | | < 0.00010 | 0.00008 | BioID high, ptprh low | 1.23E+07 | 1.23E+07 | 1.28E+07 | 1,000.00 | 1,000.00 | 1,000.00 |
| 959 | Cluster of DNA replication licensing factor MCM4 OS | A0A3B3IT92 [4] | MCM4 | 293 | 101 kDa | TRUE | 0.0018 | 0.00007 | [] | 1.46E+07 | 1.20E+07 | 1.90E+07 | 1,000.00 | 1,000.00 | 1,000.00 |
| 958 | Splicing factor U2AF 65 kDa subunit OS=Homo sapie | P26368 | U2AF2 | 231 | 54 kDa | | 0.2 | 0.00007 | [] | 3.22E+07 | 1.13E+07 | 1,000.00 | 1,000.00 | 1,000.00 | 1,000.00 |
| 960 | Mitochondrial import inner membrane translocase subu | Q3ZCQ8 | TIMM50 | 354 | 40 kDa | | 0.27 | 0.00006 | [] | 3319100 | 4.09E+07 | 3427700 | 1,000.00 | 1,000.00 | 1,000.00 |
| 961 | Cysteine-rich protein 2-binding protein OS=Homo sapi | A0A075B6H4 (+2) | KAT14 | 7 | 89 kDa | | 0.37 | 0.00006 | [] | 1,000.00 | 1,000.00 | 5.40E+07 | 1,000.00 | 1,000.00 | 1,000.00 |
| 963 | CCR4-NOT transcription complex subunit 2 OS=Hom | F8VUB4 (+2) | CNOT2 | 31 | 57 kDa | | 0.016 | 0.00005 | [] | 1.78E+07 | 1.50E+07 | 3.26E+07 | 1,000.00 | 1,000.00 | 1,000.00 |
| 962 | RuvB-like 2 OS=Homo sapiens OX=9606 GN=RUVB | Q9Y230 | RUVBL2 | 310 | 51 kDa | | 0.087 | 0.00005 | [] | 3121700 | 2.58E+07 | 3.49E+07 | 1,000.00 | 1,000.00 | 1,000.00 |
| 966 | Cluster of calcium/calmodulin-dependent protein kinas | A0A2Q3DQE3 [8] | CAMK2G | 34 | 62 kDa | TRUE | 0.0081 | 0.00004 | [] | 1.59E+07 | 2.76E+07 | 3.39E+07 | 1,000.00 | 1,000.00 | 1,000.00 |
| 967 | Nuclear receptor corepressor 1 OS=Homo sapiens OX | O75376 | NCOR1 | 101 | 270 kDa | | 0.028 | 0.00004 | [] | 2.71E+07 | 1.40E+07 | 4.27E+07 | 1,000.00 | 1,000.00 | 1,000.00 |
| 965 | Extended synaptotagmin-1 OS=Homo sapiens OX=96( | Q9BSJ8 | ESYT1 | 173 | 123 kDa | | 0.059 | 0.00004 | [] | 4.08E+07 | 2.50E+07 | 8136600 | 1,000.00 | 1,000.00 | 1,000.00 |
| 972 | Glucocorticoid receptor OS=Homo sapiens OX=9606 | P04150 | NR3C1 | 89 | 86 kDa | | 0.0049 | 0.00003 | [] | 3.82E+07 | 2.27E+07 | 4.33E+07 | 1,000.00 | 1,000.00 | 1,000.00 |
| 974 | Structural maintenance of chromosomes flexible hinge | A6NHR9 | SMCHD1 | 155 | 226 kDa | | 0.0057 | 0.00003 | [] | 4.24E+07 | 4.85E+07 | 2.47E+07 | 1,000.00 | 1,000.00 | 1,000.00 |
| 971 | Thyroid receptor-interacting protein 11 OS=Homo sap | Q15643 | TRIP11 | 52 | 228 kDa | | 0.007 | 0.00003 | [] | 2.65E+07 | 2.90E+07 | 4.79E+07 | 1,000.00 | 1,000.00 | 1,000.00 |
| 975 | Reticulon-4 OS=Homo sapiens OX=9606 GN=RTN4 | Q9NQC3 | RTN4 | 169 | 130 kDa | | 0.0098 | 0.00003 | [] | 2.28E+07 | 4.56E+07 | 5.06E+07 | 1,000.00 | 1,000.00 | 1,000.00 |
| 968 | Caldesmon 1 OS=Homo sapiens OX=9606 GN=CALI | E7EX44 (+1) | CALD1 | 158 | 64 kDa | | 0.045 | 0.00003 | [] | 4.64E+07 | 1.12E+07 | 3.04E+07 | 1,000.00 | 1,000.00 | 1,000.00 |
| 969 | Cluster of Nucleoporin 214 OS=Homo sapiens OX=96 | A0A494C1F2 [3] | NUP214 | 211 | 212 kDa | TRUE | 0.091 | 0.00003 | [] | 1.87E+07 | 1.43E+07 | 5.69E+07 | 1,000.00 | 1,000.00 | 1,000.00 |
| 973 | Survival of motor neuron 2, centromeric OS=Homo sa | A0A1W2PRV5 (+2) | SMN2 | 123 | 30 kDa | | 0.17 | 0.00003 | [] | 1,000.00 | 3.40E+07 | 7.32E+07 | 1,000.00 | 1,000.00 | 1,000.00 |
| 976 | Erythrocyte membrane protein band 4.1 like 2 OS=Ho | E9PHY5 (+1) | EPB41L2 | 191 | 104 kDa | TRUE | 0.028 | 0.00002 | [] | 2.99E+07 | 6.68E+07 | 2.89E+07 | 1,000.00 | 1,000.00 | 1,000.00 |
| 977 | Cluster of Cyclin-dependent kinase 13 OS=Homo sapi | Q14004 [9] | CDK13 | 193 | 165 kDa | TRUE | 0.053 | 0.00002 | [] | 2.80E+07 | 2.84E+07 | 7.72E+07 | 1,000.00 | 1,000.00 | 1,000.00 |
| 979 | RNA-binding protein 27 OS=Homo sapiens OX=9606 | Q9P2N5 | RBM27 | 165 | 119 kDa | | 0.071 | 0.00002 | [] | 3.51E+07 | 4.28E+07 | 1.19E+08 | 1,000.00 | 1,000.00 | 1,000.00 |
| 978 | ATP-dependent RNA helicase DDX50 OS=Homo sapi | Q9BQ39 | DDX50 | 344 | 83 kDa | TRUE | 0.23 | 0.00002 | [] | 9215000 | 2.35E+07 | 1.31E+08 | 1,000.00 | 1,000.00 | 1,000.00 |
| 982 | Non-POU domain containing octamer binding OS=Ho | A0A7I2V535 | NONO | 505 | 58 kDa | TRUE | 0.00034 | 0.00001 | BioID high, ptprh low | 8.91E+07 | 1.20E+08 | 9.85E+07 | 1,000.00 | 1,000.00 | 1,000.00 |
| 980 | Ubiquitin-associated protein 2-like OS=Homo sapiens | Q14157 | UBAP2L | 322 | 115 kDa | | 0.00099 | 0.00001 | [] | 9.30E+07 | 6.31E+07 | 7.23E+07 | 1,000.00 | 1,000.00 | 1,000.00 |
| 981 | CDK5 regulatory subunit-associated protein 2 OS=Ho | Q96SN8 | CDK5RAP2 | 11 | 215 kDa | | 0.15 | 0.00001 | [] | 1.03E+07 | 7.70E+07 | 1.92E+08 | 1,000.00 | 1,000.00 | 1,000.00 |
| 983 | Palladin OS=Homo sapiens OX=9606 GN=PALLD PE | Q8WX93 | PALLD | 85 | 151 kDa | | < 0.00010 | 0.000008 | BioID high, ptprh low | 1.26E+08 | 1.17E+08 | 1.39E+08 | 1,000.00 | 1,000.00 | 1,000.00 |
| 984 | Cluster of Host cell factor 1 OS=Homo sapiens OX=9( | P51610 [2] | HCFC1 | 278 | 209 kDa | TRUE | 0.05 | 0.000007 | [] | 1.03E+08 | 7.31E+07 | 2.32E+08 | 1,000.00 | 1,000.00 | 1,000.00 |
| 985 | Cluster of High density lipoprotein binding protein OS | A0A024R4E5 [3] | HDLBP | 218 | 141 kDa | TRUE | 0.014 | 0.000005 | [] | 2.02E+08 | 3.11E+08 | 1.32E+08 | 1,000.00 | 1,000.00 | 1,000.00 |
