## Supplemental Table 2 for "Unraveling the role of receptor-like protein tyrosine phosphatase PTPRH in cell signaling regulation and biological processes of non-small cell lung cancer"

| | baseMean | log2FC | lfcSE | stat | pvalue | padj |
|---|---|---|---|---|---|---|
| ENSG00000225972\|MTND1P23 | 8609.5982 | 6.4668 | ##### | 44.2580 | 0.0000E+00 | 0.0000E+00 |
| ENSG00000142657\|PGD | 44294.3026 | -4.6117 | ##### | ####### | 0.0000E+00 | 0.0000E+00 |
| ENSG00000162522\|KIAA1522 | 11239.1564 | -5.1980 | ##### | ####### | 0.0000E+00 | 0.0000E+00 |
| ENSG00000184371\|CSF1 | 12047.3332 | -7.8059 | ##### | ####### | 0.0000E+00 | 0.0000E+00 |
| ENSG00000085552\|IGSF9 | 2489.2789 | 5.8328 | ##### | 41.0772 | 0.0000E+00 | 0.0000E+00 |
| ENSG00000163362\|INAVA | 3515.6864 | -5.7961 | ##### | ####### | 0.0000E+00 | 0.0000E+00 |
| ENSG00000143850\|PLEKHA6 | 1542.0420 | -5.7663 | ##### | ####### | 0.0000E+00 | 0.0000E+00 |
| ENSG00000196878\|LAMB3 | 4265.6679 | -7.1051 | ##### | ####### | 0.0000E+00 | 0.0000E+00 |
| ENSG00000265190\|ANXA8 | 1854.4007 | -7.4535 | ##### | ####### | 0.0000E+00 | 0.0000E+00 |
| ENSG00000204175\|GPRIN2 | 10373.9536 | -4.9803 | ##### | ####### | 0.0000E+00 | 0.0000E+00 |
| ENSG00000122862\|SRGN | 28077.7050 | -9.6256 | ##### | ####### | 0.0000E+00 | 0.0000E+00 |
| ENSG00000042286\|AIFM2 | 3231.7848 | -5.5598 | ##### | ####### | 0.0000E+00 | 0.0000E+00 |
| ENSG00000107738\|VSIR | 5415.8075 | -7.3724 | ##### | ####### | 0.0000E+00 | 0.0000E+00 |
| ENSG00000107742\|SPOCK2 | 24916.5343 | 6.8804 | ##### | 49.8014 | 0.0000E+00 | 0.0000E+00 |
| ENSG00000156113\|KCNMA1 | 8455.8578 | 6.8303 | ##### | 42.7778 | 0.0000E+00 | 0.0000E+00 |
| ENSG00000023839\|ABCC2 | 5277.3092 | -5.7157 | ##### | ####### | 0.0000E+00 | 0.0000E+00 |
| ENSG00000134824\|FADS2 | 14525.2437 | -5.4361 | ##### | ####### | 0.0000E+00 | 0.0000E+00 |
| ENSG00000167995\|BEST1 | 41195.9240 | ####### | ##### | ####### | 0.0000E+00 | 0.0000E+00 |
| ENSG00000006534\|ALDH3B1 | 4433.5420 | -5.2919 | ##### | ####### | 0.0000E+00 | 0.0000E+00 |
| ENSG00000227097\|RPS28P7 | 7249.7655 | 6.5290 | ##### | 43.4366 | 0.0000E+00 | 0.0000E+00 |
| ENSG00000249628\|LINC00942 | 3218.3928 | -7.8251 | ##### | ####### | 0.0000E+00 | 0.0000E+00 |
| ENSG00000182326\|C1S | 25080.4897 | -7.4959 | ##### | ####### | 0.0000E+00 | 0.0000E+00 |
| ENSG00000008394\|MGST1 | 21487.9525 | -9.0956 | ##### | ####### | 0.0000E+00 | 0.0000E+00 |
| ENSG00000134548\|SPX | 2129.0860 | -7.4420 | ##### | ####### | 0.0000E+00 | 0.0000E+00 |
| ENSG00000123095\|BHLHE41 | 4141.0728 | 4.5878 | ##### | 41.3370 | 0.0000E+00 | 0.0000E+00 |
| ENSG00000135480\|KRT7 | ########## | ####### | ##### | ####### | 0.0000E+00 | 0.0000E+00 |
| ENSG00000149948\|HMGA2 | 11769.6917 | 4.9430 | ##### | 44.9230 | 0.0000E+00 | 0.0000E+00 |
| ENSG00000133636\|NTS | 4348.4504 | -6.7200 | ##### | ####### | 0.0000E+00 | 0.0000E+00 |
| ENSG00000136010\|ALDH1L2 | 2118.4881 | -5.6585 | ##### | ####### | 0.0000E+00 | 0.0000E+00 |
| ENSG00000111275\|ALDH2 | 4141.3335 | -6.6297 | ##### | ####### | 0.0000E+00 | 0.0000E+00 |
| ENSG00000127863\|TNFRSF19 | 1743.4820 | 6.4925 | ##### | 39.0082 | 0.0000E+00 | 0.0000E+00 |
| ENSG00000120694\|HSPH1 | 24793.0344 | 5.5007 | ##### | 44.2833 | 0.0000E+00 | 0.0000E+00 |
| ENSG00000087303\|NID2 | 4338.9118 | -5.8821 | ##### | ####### | 0.0000E+00 | 0.0000E+00 |
| ENSG00000185215\|TNFAIP2 | 14006.3363 | -4.8154 | ##### | ####### | 0.0000E+00 | 0.0000E+00 |
| ENSG00000074410\|CA12 | 9400.5631 | -7.6669 | ##### | ####### | 0.0000E+00 | 0.0000E+00 |
| ENSG00000102854\|MSLN | 5005.8932 | -9.9742 | ##### | ####### | 0.0000E+00 | 0.0000E+00 |
| ENSG00000103485\|QPRT | 6357.8553 | -6.6686 | ##### | ####### | 0.0000E+00 | 0.0000E+00 |
| ENSG00000205336\|ADGRG1 | 6756.9202 | -5.1609 | ##### | ####### | 0.0000E+00 | 0.0000E+00 |
| ENSG00000103257\|SLC7A5 | ########## | -3.8007 | ##### | ####### | 0.0000E+00 | 0.0000E+00 |
| ENSG00000171345\|KRT19 | 10897.2246 | -9.3571 | ##### | ####### | 0.0000E+00 | 0.0000E+00 |
| ENSG00000141753\|IGFBP4 | 42882.1345 | -5.3162 | ##### | ####### | 0.0000E+00 | 0.0000E+00 |
| ENSG00000173801\|JUP | 8155.8645 | -4.7232 | ##### | ####### | 0.0000E+00 | 0.0000E+00 |
| ENSG00000092929\|UNC13D | 4176.5552 | -7.0141 | ##### | ####### | 0.0000E+00 | 0.0000E+00 |
| ENSG00000173918\|C1QTNF1 | 4612.6658 | 5.1945 | ##### | 37.5271 | 0.0000E+00 | 0.0000E+00 |
| ENSG00000101680\|LAMA1 | 11147.3749 | 5.5140 | ##### | 44.8109 | 0.0000E+00 | 0.0000E+00 |
| ENSG00000053747\|LAMA3 | 6011.3237 | -5.8684 | ##### | ####### | 0.0000E+00 | 0.0000E+00 |
| ENSG00000099812\|MISP | 4993.7366 | -6.7967 | ##### | ####### | 0.0000E+00 | 0.0000E+00 |
| ENSG00000125730\|C3 | 17332.1989 | -7.6923 | ##### | ####### | 0.0000E+00 | 0.0000E+00 |
| ENSG00000167815\|PRDX2 | 3876.0100 | -5.4473 | ##### | ####### | 0.0000E+00 | 0.0000E+00 |
| ENSG00000024422\|EHD2 | 16528.5708 | -4.6193 | ##### | ####### | 0.0000E+00 | 0.0000E+00 |
| ENSG00000115155\|OTOF | 3029.9074 | ####### | ##### | ####### | 0.0000E+00 | 0.0000E+00 |
| ENSG00000183023\|SLC8A1 | 2397.7535 | 5.4733 | ##### | 40.5473 | 0.0000E+00 | 0.0000E+00 |
| ENSG00000144218\|AFF3 | 1890.7787 | 5.9847 | ##### | 41.0034 | 0.0000E+00 | 0.0000E+00 |
| ENSG00000163083\|INHBB | 1789.7909 | -6.2102 | ##### | ####### | 0.0000E+00 | 0.0000E+00 |
| ENSG00000115919\|KYNU | 5489.4196 | -8.6909 | ##### | ####### | 0.0000E+00 | 0.0000E+00 |
| ENSG00000115844\|DLX2 | 2928.4667 | 8.7745 | ##### | 39.0190 | 0.0000E+00 | 0.0000E+00 |
| ENSG00000021826\|CPS1 | 27196.5361 | -5.0044 | ##### | ####### | 0.0000E+00 | 0.0000E+00 |
| ENSG00000081052\|COL4A4 | 3997.3800 | -7.4017 | ##### | ####### | 0.0000E+00 | 0.0000E+00 |
| ENSG00000115648\|MLPH | 4849.7490 | -7.8504 | ##### | ####### | 0.0000E+00 | 0.0000E+00 |
| ENSG00000160209\|PDXK | 17572.5775 | -3.6583 | ##### | ####### | 0.0000E+00 | 0.0000E+00 |
| ENSG00000074416\|MGLL | 7308.4270 | -6.4646 | ##### | ####### | 0.0000E+00 | 0.0000E+00 |
| ENSG00000047457\|CP | 5403.1045 | -8.1751 | ##### | ####### | 0.0000E+00 | 0.0000E+00 |

| | | | | | | |
|---|---|---|---|---|---|---|
| ENSG00000091490\|SEL1L3 | 2793.3230 | -4.7754 | ##### | ####### | 0.0000E+00 | 0.0000E+00 |
| ENSG00000064042\|LIMCH1 | 6768.5370 | -5.4049 | ##### | ####### | 0.0000E+00 | 0.0000E+00 |
| ENSG00000118785\|SPP1 | 22165.1813 | -7.6276 | ##### | ####### | 0.0000E+00 | 0.0000E+00 |
| ENSG00000052802\|MSMO1 | 6018.1197 | -3.9528 | ##### | ####### | 0.0000E+00 | 0.0000E+00 |
| ENSG00000164220\|F2RL2 | 2753.9046 | -5.6718 | ##### | ####### | 0.0000E+00 | 0.0000E+00 |
| ENSG00000120708\|TGFBI | 21696.8802 | -6.2612 | ##### | ####### | 0.0000E+00 | 0.0000E+00 |
| ENSG00000152377\|SPOCK1 | 9323.1284 | 6.0927 | ##### | 45.2798 | 0.0000E+00 | 0.0000E+00 |
| ENSG00000120738\|EGR1 | 13721.3562 | 5.7024 | ##### | 41.6384 | 0.0000E+00 | 0.0000E+00 |
| ENSG00000111846\|GCNT2 | 1320.5651 | -6.3942 | ##### | ####### | 0.0000E+00 | 0.0000E+00 |
| ENSG00000185760\|KCNQ5 | 2879.1779 | 8.8990 | ##### | 38.3739 | 0.0000E+00 | 0.0000E+00 |
| ENSG00000272398\|CD24 | 8634.9245 | -6.3081 | ##### | ####### | 0.0000E+00 | 0.0000E+00 |
| ENSG00000112769\|LAMA4 | 3358.4717 | -5.7490 | ##### | ####### | 0.0000E+00 | 0.0000E+00 |
| ENSG00000146374\|RSPO3 | 8987.7381 | -5.3060 | ##### | ####### | 0.0000E+00 | 0.0000E+00 |
| ENSG00000146674\|IGFBP3 | 43145.1754 | -7.8959 | ##### | ####### | 0.0000E+00 | 0.0000E+00 |
| ENSG00000197343\|ZNF655 | 3028.9150 | 8.1590 | ##### | 38.8526 | 0.0000E+00 | 0.0000E+00 |
| ENSG00000106366\|SERPINE1 | 17351.5841 | -5.8095 | ##### | ####### | 0.0000E+00 | 0.0000E+00 |
| ENSG00000091129\|NRCAM | 1846.8451 | -7.1421 | ##### | ####### | 0.0000E+00 | 0.0000E+00 |
| ENSG00000128591\|FLNC | 13156.9008 | -6.9190 | ##### | ####### | 0.0000E+00 | 0.0000E+00 |
| ENSG00000164690\|SHH | 7353.5489 | 7.0218 | ##### | 41.8039 | 0.0000E+00 | 0.0000E+00 |
| ENSG00000275342\|PRAG1 | 5789.0121 | -4.9716 | ##### | ####### | 0.0000E+00 | 0.0000E+00 |
| ENSG00000077782\|FGFR1 | 11119.2462 | -2.4459 | ##### | ####### | 0.0000E+00 | 0.0000E+00 |
| ENSG00000169499\|PLEKHA2 | 8066.0579 | -5.4114 | ##### | ####### | 0.0000E+00 | 0.0000E+00 |
| ENSG00000198363\|ASPH | 85309.5790 | -5.0646 | ##### | ####### | 0.0000E+00 | 0.0000E+00 |
| ENSG00000180543\|TSPYL5 | 5610.1821 | -8.5330 | ##### | ####### | 0.0000E+00 | 0.0000E+00 |
| ENSG00000147852\|VLDLR | 2621.5336 | -6.6465 | ##### | ####### | 0.0000E+00 | 0.0000E+00 |
| ENSG00000196730\|DAPK1 | 3798.4521 | -5.1298 | ##### | ####### | 0.0000E+00 | 0.0000E+00 |
| ENSG00000047644\|WWC3 | 9568.1024 | ####### | ##### | ####### | 0.0000E+00 | 0.0000E+00 |
| ENSG00000204382\|XAGE1B | 8410.6105 | 9.9395 | ##### | 39.2042 | 0.0000E+00 | 0.0000E+00 |
| ENSG00000204379\|XAGE1A | 7167.4124 | 9.8655 | ##### | 44.3078 | 0.0000E+00 | 0.0000E+00 |
| ENSG00000022267\|FHL1 | 30153.6911 | 6.1074 | ##### | 47.4377 | 0.0000E+00 | 0.0000E+00 |
| ENSG00000276256 | 843.9883 | -5.1149 | ##### | ####### | 0.0000E+00 | 0.0000E+00 |
| ENSG00000140937\|CDH11 | 3660.1334 | 5.8484 | ##### | 37.4621 | 3.8217E-307 | ######### |
| ENSG00000132470\|ITGB4 | 3891.7821 | -4.8700 | ##### | ####### | 1.6169E-304 | ######### |
| ENSG00000189067\|LITAF | 12657.1376 | -4.6325 | ##### | ####### | 1.1797E-303 | ######### |
| ENSG00000253953\|PCDHGB4 | 2681.9859 | -7.9153 | ##### | ####### | 1.8823E-303 | ######### |
| ENSG00000198113\|TOR4A | 2452.4637 | -4.9090 | ##### | ####### | 2.3951E-303 | ######### |
| ENSG00000244509\|APOBEC3C | 2100.0662 | 6.3176 | ##### | 37.1569 | 3.3973E-302 | ######### |
| ENSG00000138772\|ANXA3 | 3649.7380 | -5.4589 | ##### | ####### | 5.1837E-302 | ######### |
| ENSG00000021355\|SERPINB1 | 3305.6607 | -5.4451 | ##### | ####### | 1.5603E-300 | ######### |
| ENSG00000065833\|ME1 | 10727.5862 | -4.5581 | ##### | ####### | 1.3273E-299 | ######### |
| ENSG00000153029\|MR1 | 1377.0851 | -5.5376 | ##### | ####### | 2.2402E-299 | ######### |
| ENSG00000075043\|KCNQ2 | 1940.4547 | 6.7193 | ##### | 36.9776 | 2.6202E-299 | ######### |
| ENSG00000115457\|IGFBP2 | 12636.3860 | 4.6701 | ##### | 36.6900 | 1.0549E-294 | ######### |
| ENSG00000169418\|NPR1 | 5614.9243 | -4.8204 | ##### | ####### | 2.1428E-293 | ######### |
| ENSG00000249395\|CASC9 | 1202.9707 | -5.2374 | ##### | ####### | 4.7116E-293 | ######### |
| ENSG00000089820\|ARHGAP4 | 3305.2369 | 5.3951 | ##### | 36.4112 | 2.8332E-290 | ######### |
| ENSG00000143631\|FLG | 2561.4034 | 5.3386 | ##### | 36.3314 | 5.1727E-289 | ######### |
| ENSG00000219438\|TAFA5 | 1170.5799 | -8.3614 | ##### | ####### | 2.6201E-288 | ######### |
| ENSG00000164733\|CTSB | ########## | -4.5200 | ##### | ####### | 3.9320E-288 | ######### |
| ENSG00000087086\|FTL | ########## | -4.4118 | ##### | ####### | 1.7501E-286 | ######### |
| ENSG00000128833\|MYO5C | 945.2093 | -6.5712 | ##### | ####### | 1.2323E-285 | ######### |
| ENSG00000291110\|TRIM16L | 4726.4426 | -4.8304 | ##### | ####### | 1.4520E-282 | ######### |
| ENSG00000177283\|FZD8 | 1267.3031 | 4.5663 | ##### | 35.8926 | 3.9844E-282 | ######### |
| ENSG00000173221\|GLRX | 3812.4961 | -4.8622 | ##### | ####### | 5.4946E-282 | ######### |
| ENSG00000106853\|PTGR1 | 29061.9411 | -4.3818 | ##### | ####### | 4.5814E-281 | ######### |
| ENSG00000066735\|KIF26A | 2091.3517 | 5.0131 | ##### | 35.8048 | 9.2922E-281 | ######### |
| ENSG00000188643\|S100A16 | 2549.5612 | -6.2914 | ##### | ####### | 4.7856E-279 | ######### |
| ENSG00000257671\|KRT7-AS | 1605.7107 | -7.4232 | ##### | ####### | 2.7216E-277 | ######### |
| ENSG00000076864\|RAP1GAP | 3024.0972 | -4.5403 | ##### | ####### | 2.8035E-276 | ######### |
| ENSG00000276547\|PCDHGB5 | 1895.3344 | -9.3780 | ##### | ####### | 9.2483E-276 | ######### |
| ENSG00000074181\|NOTCH3 | 9639.4650 | -4.3469 | ##### | ####### | 2.3220E-275 | ######### |
| ENSG00000165124\|SVEP1 | 3394.7832 | -4.4309 | ##### | ####### | 3.7589E-275 | ######### |
| ENSG00000185477\|GPRIN3 | 1540.7981 | -9.1770 | ##### | ####### | 4.2644E-273 | ######### |

| | | | | | | |
|---|---|---|---|---|---|---|
| ENSG00000139318\|DUSP6 | 4558.4532 | 3.8256 | ##### | 35.1524 | 1.0689E-270 | ######### |
| ENSG00000039068\|CDH1 | 3494.8594 | -5.7478 | ##### | ####### | 8.1427E-270 | ######### |
| ENSG00000106804\|C5 | 4144.2667 | -4.7668 | ##### | ####### | 1.0722E-269 | ######### |
| ENSG00000138449\|SLC40A1 | 2495.5719 | -5.8254 | ##### | ####### | 9.4831E-269 | ######### |
| ENSG00000145335\|SNCA | 2325.3698 | 4.8387 | ##### | 35.0217 | 1.0531E-268 | ######### |
| ENSG00000125966\|MMP24 | 4826.3412 | -5.0053 | ##### | ####### | 4.6049E-267 | ######### |
| ENSG00000159399\|HK2 | 3528.1260 | 4.5266 | ##### | 34.7488 | 1.4457E-264 | ######### |
| ENSG00000224078\|SNHG14 | 2001.6156 | 6.5391 | ##### | 34.6268 | 9.9829E-263 | ######### |
| ENSG00000167191\|GPRC5B | 5245.2619 | -4.3377 | ##### | ####### | 7.0351E-260 | ######### |
| ENSG00000065320\|NTN1 | 1617.8717 | 7.4410 | ##### | 34.3999 | 2.5344E-259 | ######### |
| ENSG00000169071\|ROR2 | 1074.3108 | 6.3893 | ##### | 34.3593 | 1.0216E-258 | ######### |
| ENSG00000118046\|STK11 | 2398.8296 | 5.0658 | ##### | 34.2607 | 3.0201E-257 | ######### |
| ENSG00000154545\|MAGED4 | 1113.4490 | 6.0047 | ##### | 34.2048 | 2.0486E-256 | ######### |
| ENSG00000159403\|C1R | 1790.8692 | -5.1520 | ##### | ####### | 2.6589E-256 | ######### |
| ENSG00000183760\|ACP7 | 1442.1299 | 6.9966 | ##### | 34.1120 | 4.8992E-255 | ######### |
| ENSG00000188064\|WNT7B | 5344.8774 | -4.5718 | ##### | ####### | 7.5367E-255 | ######### |
| ENSG00000105339\|DENND3 | 2688.7836 | -5.2769 | ##### | ####### | 1.6365E-254 | ######### |
| ENSG00000205213\|LGR4 | 4031.5469 | -4.2497 | ##### | ####### | 2.0542E-254 | ######### |
| ENSG00000142798\|HSPG2 | 20813.7929 | -4.0920 | ##### | ####### | 2.3164E-254 | ######### |
| ENSG00000164251\|F2RL1 | 11001.1458 | 4.2224 | ##### | 34.0396 | 5.7763E-254 | ######### |
| ENSG00000112297\|CRYBG1 | 2678.9300 | -4.8051 | ##### | ####### | 4.6148E-253 | ######### |
| ENSG00000110237\|ARHGEF17 | 3524.4753 | 4.6148 | ##### | 33.9561 | 9.9067E-253 | ######### |
| ENSG00000168679\|SLC16A4 | 2105.3511 | -5.7566 | ##### | ####### | 2.2476E-252 | ######### |
| ENSG00000074527\|NTN4 | 3096.9920 | -4.4870 | ##### | ####### | 2.3942E-252 | ######### |
| ENSG00000211448\|DIO2 | 7071.1159 | -4.1885 | ##### | ####### | 2.0873E-251 | ######### |
| ENSG00000113389\|NPR3 | 1903.7480 | -7.7137 | ##### | ####### | 3.0335E-251 | ######### |
| ENSG00000264230\|ANXA8L1 | 1985.0136 | -8.1104 | ##### | ####### | 9.9593E-250 | ######### |
| ENSG00000125848\|FLRT3 | 1622.6447 | -5.4028 | ##### | ####### | 1.6496E-247 | ######### |
| ENSG00000173391\|OLR1 | 2365.1950 | 4.7072 | ##### | 33.5928 | 2.1385E-247 | ######### |
| ENSG00000183092\|BEGAIN | 1751.2692 | 4.9255 | ##### | 33.4803 | 9.3350E-246 | ######### |
| ENSG00000172572\|PDE3A | 3729.9697 | -4.1628 | ##### | ####### | 1.3879E-244 | ######### |
| ENSG00000144821\|MYH15 | 2968.4329 | 6.1957 | ##### | 33.3715 | 3.5504E-244 | ######### |
| ENSG00000197415\|VEPH1 | 783.6270 | -5.3906 | ##### | ####### | 8.6768E-244 | ######### |
| ENSG00000170579\|DLGAP1 | 906.7586 | 5.0965 | ##### | 33.3105 | 2.7213E-243 | ######### |
| ENSG00000166086\|JAM3 | 2227.1560 | 4.6889 | ##### | 33.2926 | 4.9362E-243 | ######### |
| ENSG00000144063\|MALL | 4083.4461 | -8.1172 | ##### | ####### | 2.2821E-241 | ######### |
| ENSG00000105852\|PON3 | 1233.2194 | -5.9672 | ##### | ####### | 4.2883E-241 | ######### |
| ENSG00000067167\|TRAM1 | 21214.7430 | -3.4611 | ##### | ####### | 1.5050E-240 | ######### |
| ENSG00000250986\|LINC02600 | 1311.8335 | 5.2337 | ##### | 33.1020 | 2.7837E-240 | ######### |
| ENSG00000132718\|SYT11 | 1861.2475 | 4.4975 | ##### | 33.0707 | 7.8475E-240 | ######### |
| ENSG00000132967\|HMGB1P5 | 2983.7349 | 4.5451 | ##### | 33.0693 | 8.2082E-240 | ######### |
| ENSG00000218336\|TENM3 | 2232.4608 | 4.8257 | ##### | 33.0614 | 1.0685E-239 | ######### |
| ENSG00000197937\|ZNF347 | 1129.7695 | -4.9839 | ##### | ####### | 7.2165E-238 | ######### |
| ENSG00000018236\|CNTN1 | 2744.5736 | -7.5981 | ##### | ####### | 1.1310E-237 | ######### |
| ENSG00000137819\|PAQR5 | 4020.3025 | -4.5242 | ##### | ####### | 4.1213E-236 | ######### |
| ENSG00000164308\|ERAP2 | 1858.2773 | -5.5810 | ##### | ####### | 1.2047E-235 | ######### |
| ENSG00000156299\|TIAM1 | 1276.9504 | 6.1555 | ##### | 32.7570 | 2.4172E-235 | ######### |
| ENSG00000158315\|RHBDL2 | 983.3092 | -8.5956 | ##### | ####### | 5.1115E-235 | ######### |
| ENSG00000115594\|IL1R1 | 5755.7690 | -3.8431 | ##### | ####### | 1.1989E-234 | ######### |
| ENSG00000164684\|ZNF704 | 1827.6296 | -5.2377 | ##### | ####### | 5.9098E-234 | ######### |
| ENSG00000095383\|TBC1D2 | 2397.9295 | -4.7482 | ##### | ####### | 6.7834E-233 | ######### |
| ENSG00000106780\|MEGF9 | 2552.6158 | -4.8239 | ##### | ####### | 5.8211E-232 | ######### |
| ENSG00000164976\|MYORG | 1492.5374 | -4.6592 | ##### | ####### | 6.0351E-232 | ######### |
| ENSG00000183305\|MAGEA2B | 1752.2457 | ####### | ##### | ####### | 8.8628E-232 | ######### |
| ENSG00000169554\|ZEB2 | 977.5338 | 5.1574 | ##### | 32.3921 | 3.5517E-230 | ######### |
| ENSG00000049323\|LTBP1 | 5367.4622 | -4.0409 | ##### | ####### | 8.4738E-230 | ######### |
| ENSG00000130720\|FIBCD1 | 913.6745 | -4.7939 | ##### | ####### | 1.3918E-229 | ######### |
| ENSG00000187243\|MAGED4B | 1080.3443 | 5.8054 | ##### | 32.3491 | 1.4295E-229 | ######### |
| ENSG00000272405 | 3077.2148 | -4.1744 | ##### | ####### | 2.9630E-229 | ######### |
| ENSG00000079459\|FDFT1 | 37734.6495 | -3.8430 | ##### | ####### | 3.9222E-229 | ######### |
| ENSG00000135678\|CPM | 9974.2433 | -3.8083 | ##### | ####### | 5.0735E-229 | ######### |
| ENSG00000198959\|TGM2 | 85864.9746 | -3.6780 | ##### | ####### | 7.7630E-229 | ######### |
| ENSG00000169031\|COL4A3 | 1454.1259 | -5.2402 | ##### | ####### | 1.0284E-228 | ######### |

| | | | | | | |
|---|---|---|---|---|---|---|
| ENSG00000206538\|VGLL3 | 4177.2083 | -4.0474 | ##### | ####### | 2.6740E-228 | ######### |
| ENSG00000180537\|RNF182 | 2012.0933 | 6.5140 | ##### | 32.2551 | 2.9847E-228 | ######### |
| ENSG00000114200\|BCHE | 1800.7344 | -6.4956 | ##### | ####### | 6.6374E-228 | ######### |
| ENSG00000116729\|WLS | 3568.4959 | 4.4809 | ##### | 32.1707 | 4.5295E-227 | ######### |
| ENSG00000138347\|MYPN | 3535.3489 | 4.0381 | ##### | 32.1119 | 3.0040E-226 | ######### |
| ENSG00000163053\|SLC16A14 | 1588.8383 | -7.8179 | ##### | ####### | 7.5643E-226 | ######### |
| ENSG00000154188\|ANGPT1 | 2245.2268 | -4.4303 | ##### | ####### | 7.6652E-226 | ######### |
| ENSG00000272602\|ZNF595 | 1331.0078 | -9.3848 | ##### | ####### | 1.1114E-225 | ######### |
| ENSG00000001617\|SEMA3F | 1783.5469 | -5.8355 | ##### | ####### | 4.6113E-225 | ######### |
| ENSG00000115267\|IFIH1 | 1525.5508 | -4.1507 | ##### | ####### | 2.7894E-224 | ######### |
| ENSG00000105419\|MEIS3 | 2510.7824 | -4.9887 | ##### | ####### | 2.9348E-224 | ######### |
| ENSG00000167766\|ZNF83 | 1393.2585 | -4.7153 | ##### | ####### | 4.6147E-224 | ######### |
| ENSG00000196562\|SULF2 | 1565.1309 | 5.2151 | ##### | 31.8373 | 1.9760E-222 | ######### |
| ENSG00000205413\|SAMD9 | 1411.2728 | -7.0778 | ##### | ####### | 7.7047E-222 | ######### |
| ENSG00000090013\|BLVRB | 8549.4747 | -4.0562 | ##### | ####### | 9.0565E-222 | ######### |
| ENSG00000237973\|MTCO1P12 | 63336.3016 | 4.1238 | ##### | 31.6948 | 1.8342E-220 | ######### |
| ENSG00000253368\|TRNP1 | 9021.7681 | -3.9746 | ##### | ####### | 8.8384E-220 | ######### |
| ENSG00000076716\|GPC4 | 817.7145 | 5.0747 | ##### | 31.6286 | 1.4935E-219 | ######### |
| ENSG00000130208\|APOC1 | 1304.9400 | -4.4054 | ##### | ####### | 2.6376E-219 | ######### |
| ENSG00000128242\|GAL3ST1 | 1155.3644 | -7.6898 | ##### | ####### | 1.5569E-218 | ######### |
| ENSG00000159450\|TCHH | 874.6703 | 5.7147 | ##### | 31.5423 | 2.2869E-218 | ######### |
| ENSG00000221963\|APOL6 | 1264.2533 | -4.7705 | ##### | ####### | 3.3065E-218 | ######### |
| ENSG00000164120\|HPGD | 842.2039 | -6.5210 | ##### | ####### | 1.5500E-217 | ######### |
| ENSG00000253958\|CLDN23 | 1698.7630 | -4.3190 | ##### | ####### | 1.9217E-217 | ######### |
| ENSG00000198732\|SMOC1 | 1302.4522 | -4.4880 | ##### | ####### | 1.8207E-216 | ######### |
| ENSG00000178445\|GLDC | 2365.2744 | -3.0944 | ##### | ####### | 2.7474E-216 | ######### |
| ENSG00000167244\|IGF2 | 3221.2065 | 6.5135 | ##### | 31.3754 | 4.3782E-216 | ######### |
| ENSG00000148175\|STOM | 11138.5793 | -2.1567 | ##### | ####### | 5.6300E-216 | ######### |
| ENSG00000181790\|ADGRB1 | 2115.5677 | 7.8202 | ##### | 31.3428 | 1.2190E-215 | ######### |
| ENSG00000111816\|FRK | 1876.0264 | -5.5308 | ##### | ####### | 1.4664E-215 | ######### |
| ENSG00000268606\|MAGEA2 | 1815.9359 | ####### | ##### | ####### | 2.7587E-215 | ######### |
| ENSG00000122870\|BICC1 | 5380.1452 | -3.4574 | ##### | ####### | 4.5827E-215 | ######### |
| ENSG00000132613\|MTSS2 | 4667.4181 | -4.0277 | ##### | ####### | 4.4678E-214 | ######### |
| ENSG00000091592\|NLRP1 | 909.5221 | -5.3194 | ##### | ####### | 2.0456E-213 | ######### |
| ENSG00000182809\|CRIP2 | 3404.4014 | -4.0626 | ##### | ####### | 2.9527E-213 | ######### |
| ENSG00000167779\|IGFBP6 | 1571.0738 | -4.2829 | ##### | ####### | 3.4223E-213 | ######### |
| ENSG00000173349\|SFT2D3 | 889.4155 | -4.2253 | ##### | ####### | 9.9331E-213 | ######### |
| ENSG00000077092\|RARB | 539.4864 | -5.9914 | ##### | ####### | 1.0177E-212 | ######### |
| ENSG00000173698\|ADGRG2 | 850.7513 | -5.8076 | ##### | ####### | 1.6736E-212 | ######### |
| ENSG00000148426\|PROSER2 | 2235.2283 | -3.9067 | ##### | ####### | 1.9423E-212 | ######### |
| ENSG00000104325\|DECR1 | 3248.5183 | -4.1853 | ##### | ####### | 3.7725E-212 | ######### |
| ENSG00000113504\|SLC12A7 | 4823.7691 | -3.7691 | ##### | ####### | 2.9150E-211 | ######### |
| ENSG00000275832\|ARHGAP23 | 2204.0805 | -4.3852 | ##### | ####### | 7.0795E-211 | ######### |
| ENSG00000145632\|PLK2 | 21423.4874 | -2.8893 | ##### | ####### | 1.0729E-210 | ######### |
| ENSG00000135046\|ANXA1 | 29755.4467 | -2.2407 | ##### | ####### | 1.3000E-210 | ######### |
| ENSG00000020633\|RUNX3 | 1688.9304 | 4.8328 | ##### | 30.9227 | 5.9129E-210 | ######### |
| ENSG00000100979\|PLTP | 9810.0432 | 4.0045 | ##### | 30.9111 | 8.4853E-210 | ######### |
| ENSG00000188112\|C6orf132 | 960.8012 | -6.7987 | ##### | ####### | 4.0911E-209 | ######### |
| ENSG00000104490\|NCALD | 751.8921 | -6.5487 | ##### | ####### | 6.3382E-209 | ######### |
| ENSG00000129116\|PALLD | 8707.2922 | -3.6993 | ##### | ####### | 7.2441E-209 | ######### |
| ENSG00000149380\|P4HA3 | 1232.7760 | -4.7854 | ##### | ####### | 7.9637E-209 | ######### |
| ENSG00000153234\|NR4A2 | 3728.1364 | 3.5240 | ##### | 30.8355 | 8.7679E-209 | ######### |
| ENSG00000171772\|SYCE1 | 570.2386 | -6.9837 | ##### | ####### | 1.5381E-208 | ######### |
| ENSG00000198053\|SIRPA | 2328.1700 | -3.8195 | ##### | ####### | 1.5637E-208 | ######### |
| ENSG00000168874\|ATOH8 | 1138.2786 | -4.3868 | ##### | ####### | 3.6437E-208 | ######### |
| ENSG00000121005\|CRISPLD1 | 2129.6519 | -4.1518 | ##### | ####### | 4.5054E-208 | ######### |
| ENSG00000081923\|ATP8B1 | 2238.5596 | -4.4670 | ##### | ####### | 7.3164E-208 | ######### |
| ENSG00000029534\|ANK1 | 858.4054 | 5.0169 | ##### | 30.7499 | 1.2255E-207 | ######### |
| ENSG00000176153\|GPX2 | 3826.3183 | ####### | ##### | ####### | 4.7561E-207 | ######### |
| ENSG00000157613\|CREB3L1 | 559.5427 | -3.9342 | ##### | ####### | 9.6458E-206 | ######### |
| ENSG00000118257\|NRP2 | 1649.3229 | -4.4741 | ##### | ####### | 1.4005E-205 | ######### |
| ENSG00000171557\|FGG | 12203.2382 | ####### | ##### | ####### | 2.6288E-205 | ######### |
| ENSG00000186907\|RTN4RL2 | 1246.8785 | -4.1014 | ##### | ####### | 5.8230E-205 | ######### |

| | | | | | | |
|---|---|---|---|---|---|---|
| ENSG00000101188\|NTSR1 | 10589.7744 | 10.2419 | ##### | 30.4916 | 3.3671E-204 | ######### |
| ENSG00000171246\|NPTX1 | 18755.4429 | 3.5373 | ##### | 30.4812 | 4.6217E-204 | ######### |
| ENSG00000178860\|MSC | 1624.5935 | -4.0488 | ##### | ####### | 1.1442E-203 | ######### |
| ENSG00000204176\|SYT15 | 1112.4374 | -4.6841 | ##### | ####### | 3.5351E-202 | ######### |
| ENSG00000171951\|SCG2 | 1189.7237 | 4.4600 | ##### | 30.2844 | 1.8425E-201 | ######### |
| ENSG00000198121\|LPAR1 | 2776.7451 | -4.0690 | ##### | ####### | 1.0901E-200 | ######### |
| ENSG00000162496\|DHRS3 | 2694.6377 | -6.9074 | ##### | ####### | 5.3823E-200 | ######### |
| ENSG00000196154\|S100A4 | 3617.8716 | -4.0351 | ##### | ####### | 1.0773E-199 | ######### |
| ENSG00000131668\|BARX1 | 1284.7837 | 5.1302 | ##### | 30.1318 | 1.8556E-199 | ######### |
| ENSG00000146592\|CREB5 | 1339.7255 | 6.1720 | ##### | 30.0945 | 5.7219E-199 | ######### |
| ENSG00000136068\|FLNB | 56765.7170 | -3.5451 | ##### | ####### | 1.4332E-198 | ######### |
| ENSG00000185070\|FLRT2 | 2182.6435 | 6.6948 | ##### | 30.0493 | 2.2279E-198 | ######### |
| ENSG00000092820\|EZR | 23176.1959 | -1.7022 | ##### | ####### | 2.5243E-198 | ######### |
| ENSG00000204442\|NALF1 | 737.5983 | -5.2455 | ##### | ####### | 7.3241E-198 | ######### |
| ENSG00000198838\|RYR3 | 935.9908 | 4.7100 | ##### | 29.9729 | 2.2111E-197 | ######### |
| ENSG00000105711\|SCN1B | 899.9988 | -4.6573 | ##### | ####### | 8.9255E-196 | ######### |
| ENSG00000124920\|MYRF | 2627.0715 | -4.1670 | ##### | ####### | 3.3325E-195 | ######### |
| ENSG00000147905\|ZCCHC7 | 3454.4638 | 2.0943 | ##### | 29.7966 | 4.3297E-195 | ######### |
| ENSG00000115414\|FN1 | 10402.1414 | -3.6474 | ##### | ####### | 8.8252E-194 | ######### |
| ENSG00000168077\|SCARA3 | 2463.9836 | -3.7701 | ##### | ####### | 1.4472E-193 | ######### |
| ENSG00000205426\|KRT81 | 94026.0002 | ####### | ##### | ####### | 2.2499E-193 | ######### |
| ENSG00000178538\|CA8 | 1129.1445 | 4.9969 | ##### | 29.6555 | 2.8798E-193 | ######### |
| ENSG00000111907\|TPD52L1 | 1424.0597 | -4.3690 | ##### | ####### | 4.9880E-193 | ######### |
| ENSG00000156959\|LHFPL4 | 875.7565 | 6.1597 | ##### | 29.6224 | 7.6915E-193 | ######### |
| ENSG00000110711\|AIP | 7662.2896 | 2.5516 | ##### | 29.6003 | 1.4790E-192 | ######### |
| ENSG00000058063\|ATP11B | 2611.4609 | -3.8254 | ##### | ####### | 2.4642E-192 | ######### |
| ENSG00000158050\|DUSP2 | 2515.0193 | 4.5021 | ##### | 29.5708 | 3.5450E-192 | ######### |
| ENSG00000119280\|C1orf198 | 1353.5229 | -4.4515 | ##### | ####### | 4.5007E-192 | ######### |
| ENSG00000134775\|FHOD3 | 1806.9109 | 4.0859 | ##### | 29.5574 | 5.2812E-192 | ######### |
| ENSG00000091622\|PITPNM3 | 644.9655 | -4.8881 | ##### | ####### | 5.4678E-192 | ######### |
| ENSG00000141527\|CARD14 | 905.1494 | -5.2951 | ##### | ####### | 4.1043E-191 | ######### |
| ENSG00000148143\|ZNF462 | 2095.0408 | 3.9050 | ##### | 29.3795 | 1.0027E-189 | ######### |
| ENSG00000159167\|STC1 | 6641.0914 | 3.9549 | ##### | 29.3589 | 1.8402E-189 | ######### |
| ENSG00000100968\|NFATC4 | 1066.0212 | 5.4710 | ##### | 29.3333 | 3.9003E-189 | ######### |
| ENSG00000144355\|DLX1 | 2231.8314 | 8.0219 | ##### | 29.3039 | 9.2613E-189 | ######### |
| ENSG00000063660\|GPC1 | 22745.0822 | -3.6141 | ##### | ####### | 2.0340E-188 | ######### |
| ENSG00000164078\|MST1R | 2362.5904 | -7.8523 | ##### | ####### | 1.2797E-187 | ######### |
| ENSG00000132170\|PPARG | 729.6711 | -4.2944 | ##### | ####### | 1.4945E-187 | ######### |
| ENSG00000118849\|RARRES1 | 651.8736 | -4.8337 | ##### | ####### | 2.9281E-187 | ######### |
| ENSG00000261115\|TMEM178B | 1412.4638 | -4.4201 | ##### | ####### | 3.0435E-187 | ######### |
| ENSG00000178038\|ALS2CL | 1686.4664 | -3.9801 | ##### | ####### | 1.6468E-186 | ######### |
| ENSG00000170962\|PDGFD | 1045.8347 | -6.4894 | ##### | ####### | 1.9340E-186 | ######### |
| ENSG00000160216\|AGPAT3 | 4246.7098 | -3.5185 | ##### | ####### | 3.0537E-186 | ######### |
| ENSG00000144746\|ARL6IP5 | 5582.1368 | -3.5451 | ##### | ####### | 1.6834E-185 | ######### |
| ENSG00000113657\|DPYSL3 | 2188.6877 | 4.7571 | ##### | 29.0465 | 1.7015E-185 | ######### |
| ENSG00000164181\|ELOVL7 | 1328.9715 | -4.0332 | ##### | ####### | 3.9395E-184 | ######### |
| ENSG00000229953\|BCAN-AS2 | 952.3383 | -4.3231 | ##### | ####### | 5.8423E-184 | ######### |
| ENSG00000111319\|SCNN1A | 791.9619 | -5.8136 | ##### | ####### | 1.2055E-183 | ######### |
| ENSG00000109113\|RAB34 | 6282.6017 | 9.4654 | ##### | 28.8870 | 1.7374E-183 | ######### |
| ENSG00000172927\|MYEOV | 1271.6527 | -4.3693 | ##### | ####### | 1.1321E-182 | ######### |
| ENSG00000088992\|TESC | 2640.7359 | -3.6729 | ##### | ####### | 1.7121E-182 | ######### |
| ENSG00000109321\|AREG | 4371.7995 | -3.5593 | ##### | ####### | 2.1021E-182 | ######### |
| ENSG00000162545\|CAMK2N1 | 1641.3037 | -4.5078 | ##### | ####### | 3.1035E-182 | ######### |
| ENSG00000204257\|HLA-DMA | 509.4730 | -4.6518 | ##### | ####### | 3.3553E-182 | ######### |
| ENSG00000167642\|SPINT2 | 26658.9556 | -3.4591 | ##### | ####### | 3.7898E-182 | ######### |
| ENSG00000070808\|CAMK2A | 682.6071 | -5.4297 | ##### | ####### | 4.7112E-182 | ######### |
| ENSG00000151012\|SLC7A11 | 11874.9016 | -3.5016 | ##### | ####### | 2.2147E-181 | ######### |
| ENSG00000229644\|NAMPTP1 | 3124.5963 | -3.4365 | ##### | ####### | 3.3204E-181 | ######### |
| ENSG00000213626\|LBH | 1550.9341 | -4.5508 | ##### | ####### | 4.4159E-181 | ######### |
| ENSG00000163430\|FSTL1 | 9351.4772 | -3.3641 | ##### | ####### | 7.8651E-181 | ######### |
| ENSG00000129422\|MTUS1 | 1295.0282 | -3.8505 | ##### | ####### | 4.8999E-180 | ######### |
| ENSG00000009694\|TENM1 | 568.4856 | -5.8621 | ##### | ####### | 7.1633E-180 | ######### |
| ENSG00000160117\|ANKLE1 | 1163.3217 | 4.6693 | ##### | 28.5785 | 1.2421E-179 | ######### |

| | | | | | | |
|---|---|---|---|---|---|---|
| ENSG00000168453\|HR | 809.1287 | -4.1814 | ##### | ####### | 1.3980E-179 | ######### |
| ENSG00000170345\|FOS | 2304.4334 | 4.0499 | ##### | 28.4875 | 1.6757E-178 | ######### |
| ENSG00000007264\|MATK | 1772.9405 | 4.6376 | ##### | 28.4840 | 1.8486E-178 | ######### |
| ENSG00000135362\|PRR5L | 843.1803 | 4.4146 | ##### | 28.4332 | 7.8704E-178 | ######### |
| ENSG00000109072\|VTN | 619.1534 | -4.9365 | ##### | ####### | 8.0886E-177 | ######### |
| ENSG00000226887\|ERVMER34-1 | 1166.1315 | 4.0645 | ##### | 28.3277 | 1.5763E-176 | ######### |
| ENSG00000116133\|DHCR24 | 33335.9800 | -3.2614 | ##### | ####### | 1.1085E-175 | ######### |
| ENSG00000157851\|DPYSL5 | 694.3647 | 5.1496 | ##### | 28.2531 | 1.3034E-175 | ######### |
| ENSG00000065534\|MYLK | 5563.7014 | -3.4969 | ##### | ####### | 1.9662E-174 | ######### |
| ENSG00000124249\|KCNK15 | 3404.8524 | 3.5805 | ##### | 28.1231 | 5.1121E-174 | ######### |
| ENSG00000148677\|ANKRD1 | 1563.2751 | -5.0303 | ##### | ####### | 2.9565E-172 | ######### |
| ENSG00000187800\|PEAR1 | 558.6027 | -5.0899 | ##### | ####### | 5.2101E-172 | ######### |
| ENSG00000115112\|TFCP2L1 | 1139.2324 | -3.8764 | ##### | ####### | 1.0743E-171 | ######### |
| ENSG00000188641\|DPYD | 2307.5244 | -3.5789 | ##### | ####### | 1.5367E-171 | ######### |
| ENSG00000185950\|IRS2 | 1908.5786 | -3.8316 | ##### | ####### | 5.2311E-171 | ######### |
| ENSG00000189221\|MAOA | 1603.5975 | -3.7911 | ##### | ####### | 9.2560E-171 | ######### |
| ENSG00000143819\|EPHX1 | 11697.6458 | -3.3696 | ##### | ####### | 3.0090E-170 | ######### |
| ENSG00000107317\|PTGDS | 1667.3800 | -4.1812 | ##### | ####### | 2.5538E-169 | ######### |
| ENSG00000042493\|CAPG | 1027.3381 | -4.7070 | ##### | ####### | 3.4221E-169 | ######### |
| ENSG00000163132\|MSX1 | 1246.2220 | 4.1423 | ##### | 27.6695 | 1.6244E-168 | ######### |
| ENSG00000132669\|RIN2 | 1323.0632 | -4.6817 | ##### | ####### | 1.6916E-168 | ######### |
| ENSG00000162104\|ADCY9 | 2052.6306 | -3.7091 | ##### | ####### | 1.7167E-168 | ######### |
| ENSG00000091986\|CCDC80 | 2422.6823 | -3.7628 | ##### | ####### | 3.1933E-168 | ######### |
| ENSG00000105255\|FSD1 | 4927.9034 | 3.8486 | ##### | 27.6437 | 3.3190E-168 | ######### |
| ENSG00000165685\|TMEM52B | 1341.9005 | 7.6701 | ##### | 27.6065 | 9.2913E-168 | ######### |
| ENSG00000151892\|GFRA1 | 867.4355 | -4.4595 | ##### | ####### | 2.9601E-167 | ######### |
| ENSG00000106733\|NMRK1 | 933.8414 | -4.1076 | ##### | ####### | 6.1772E-167 | ######### |
| ENSG00000121413\|ZSCAN18 | 2848.5574 | -2.7625 | ##### | ####### | 2.2532E-166 | ######### |
| ENSG00000134222\|PSRC1 | 4621.8840 | 2.3129 | ##### | 27.4882 | 2.4265E-166 | ######### |
| ENSG00000130021\|PUDP | 1483.4074 | -3.8431 | ##### | ####### | 3.1203E-166 | ######### |
| ENSG00000173267\|SNCG | 925.1591 | -9.4279 | ##### | ####### | 4.4481E-166 | ######### |
| ENSG00000165272\|AQP3 | 873.3655 | -5.2940 | ##### | ####### | 8.8283E-166 | ######### |
| ENSG00000151790\|TDO2 | 447.0534 | -7.1161 | ##### | ####### | 3.3294E-165 | ######### |
| ENSG00000164649\|CDCA7L | 3951.6144 | 3.4857 | ##### | 27.3761 | 5.2800E-165 | ######### |
| ENSG00000139629\|GALNT6 | 2669.2625 | 3.7378 | ##### | 27.3131 | 2.9664E-164 | ######### |
| ENSG00000176485\|PLAAT3 | 1611.6351 | -2.5396 | ##### | ####### | 1.7757E-163 | ######### |
| ENSG00000121966\|CXCR4 | 1039.6756 | 7.6571 | ##### | 27.2444 | 1.9379E-163 | ######### |
| ENSG00000118640\|VAMP8 | 1905.3259 | -3.5850 | ##### | ####### | 3.4377E-163 | ######### |
| ENSG00000180818\|HOXC10 | 4088.5727 | 3.7467 | ##### | 27.2164 | 4.1502E-163 | ######### |
| ENSG00000105289\|TJP3 | 939.8705 | -4.1724 | ##### | ####### | 6.5765E-163 | ######### |
| ENSG00000106034\|CPED1 | 684.0649 | -7.4521 | ##### | ####### | 1.2465E-162 | ######### |
| ENSG00000105472\|CLEC11A | 3982.7815 | 3.7139 | ##### | 27.1677 | 1.5636E-162 | ######### |
| ENSG00000085733\|CTTN | 43741.1985 | 1.6864 | ##### | 27.1511 | 2.4561E-162 | ######### |
| ENSG00000198538\|ZNF28 | 1879.0376 | -3.7915 | ##### | ####### | 4.6466E-162 | ######### |
| ENSG00000166741\|NNMT | 6567.2109 | -3.2915 | ##### | ####### | 1.7286E-161 | ######### |
| ENSG00000079215\|SLC1A3 | 790.4823 | 7.3165 | ##### | 27.0363 | 5.5366E-161 | ######### |
| ENSG00000143614\|GATAD2B | 6666.5818 | 2.2050 | ##### | 27.0113 | 1.0891E-160 | ######### |
| ENSG00000196549\|MME | 6779.1086 | 3.2746 | ##### | 27.0003 | 1.4672E-160 | ######### |
| ENSG00000183153\|GJD3 | 994.2809 | -3.7926 | ##### | ####### | 4.9151E-160 | ######### |
| ENSG00000157110\|RBPMS | 1627.8179 | -3.8726 | ##### | ####### | 9.6849E-160 | ######### |
| ENSG00000154065\|ANKRD29 | 1437.9292 | -5.1334 | ##### | ####### | 1.0506E-159 | ######### |
| ENSG00000126561\|STAT5A | 1182.2796 | -5.4837 | ##### | ####### | 1.5750E-159 | ######### |
| ENSG00000089127\|OAS1 | 872.6063 | -4.9347 | ##### | ####### | 1.7676E-159 | ######### |
| ENSG00000125637\|PSD4 | 1722.7402 | -3.7554 | ##### | ####### | 2.5014E-159 | ######### |
| ENSG00000132792\|CTNNBL1 | 7930.6604 | 1.7643 | ##### | 26.8729 | 4.5539E-159 | ######### |
| ENSG00000139970\|RTN1 | 988.5293 | 7.9781 | ##### | 26.8686 | 5.1141E-159 | ######### |
| ENSG00000140961\|OSGIN1 | 3530.2926 | -3.3841 | ##### | ####### | 1.2895E-158 | ######### |
| ENSG00000074047\|GLI2 | 1584.8550 | -3.9124 | ##### | ####### | 3.6434E-158 | ######### |
| ENSG00000131037\|EPS8L1 | 668.9534 | -6.8911 | ##### | ####### | 5.9682E-158 | ######### |
| ENSG00000179111\|HES7 | 1170.0486 | 3.8619 | ##### | 26.7709 | 7.0489E-158 | ######### |
| ENSG00000167106\|EEIG1 | 8661.2376 | -3.1637 | ##### | ####### | 8.4530E-158 | ######### |
| ENSG00000111859\|NEDD9 | 1053.9709 | -5.1264 | ##### | ####### | 9.3884E-158 | ######### |
| ENSG00000132561\|MATN2 | 1331.5702 | -3.7671 | ##### | ####### | 2.1813E-157 | ######### |

| | | | | | | |
|---|---|---|---|---|---|---|
| ENSG00000166147\|FBN1 | 3566.0933 | -3.4019 | ##### | ####### | 8.0919E-157 | ######### |
| ENSG00000134531\|EMP1 | 3422.1681 | -3.4988 | ##### | ####### | 1.0156E-156 | ######### |
| ENSG00000125798\|FOXA2 | 593.7090 | -4.9204 | ##### | ####### | 2.4051E-156 | ######### |
| ENSG00000277758\|SYT15B | 1465.0603 | -3.5884 | ##### | ####### | 4.1699E-156 | ######### |
| ENSG00000279569 | 2176.4840 | -3.7468 | ##### | ####### | 5.9005E-156 | ######### |
| ENSG00000227036\|LINC00511 | 862.2726 | -3.5472 | ##### | ####### | 1.2508E-155 | ######### |
| ENSG00000198576\|ARC | 703.4060 | 4.5852 | ##### | 26.5672 | 1.6257E-155 | ######### |
| ENSG00000168280\|KIF5C | 646.2794 | 5.8818 | ##### | 26.5602 | 1.9602E-155 | ######### |
| ENSG00000183111\|ARHGEF37 | 1082.4746 | -4.4390 | ##### | ####### | 1.3432E-154 | ######### |
| ENSG00000149177\|PTPRJ | 7969.5756 | -3.2732 | ##### | ####### | 3.0768E-154 | ######### |
| ENSG00000152952\|PLOD2 | 12531.4574 | -3.2288 | ##### | ####### | 4.7997E-154 | ######### |
| ENSG00000112308\|C6orf62 | 12382.5584 | 1.5596 | ##### | 26.4288 | 6.4002E-154 | ######### |
| ENSG00000235531\|MSC-AS1 | 1821.1237 | -3.5122 | ##### | ####### | 1.1559E-153 | ######### |
| ENSG00000124212\|PTGIS | 663.8582 | -4.2864 | ##### | ####### | 1.3729E-153 | ######### |
| ENSG00000138271\|GPR87 | 774.5128 | 5.6995 | ##### | 26.3808 | 2.2736E-153 | ######### |
| ENSG00000169297\|NR0B1 | 1581.5645 | -3.5968 | ##### | ####### | 2.8306E-153 | ######### |
| ENSG00000129925\|PGAP6 | 4405.1739 | -2.5932 | ##### | ####### | 9.6613E-153 | ######### |
| ENSG00000136205\|TNS3 | 17301.6949 | -3.1014 | ##### | ####### | 1.5382E-152 | ######### |
| ENSG00000001036\|FUCA2 | 6210.5370 | -3.3242 | ##### | ####### | 4.6980E-152 | ######### |
| ENSG00000179862\|CITED4 | 2779.8595 | -3.2236 | ##### | ####### | 1.2116E-151 | ######### |
| ENSG00000138413\|IDH1 | 9619.2398 | -2.6711 | ##### | ####### | 1.8922E-151 | ######### |
| ENSG00000109814\|UGDH | 14115.8708 | -3.1732 | ##### | ####### | 2.5615E-151 | ######### |
| ENSG00000144278\|GALNT13 | 4579.3810 | -3.2152 | ##### | ####### | 7.8722E-151 | ######### |
| ENSG00000102452\|NALCN | 746.4817 | -4.0450 | ##### | ####### | 2.0727E-150 | ######### |
| ENSG00000171631\|P2RY6 | 870.9642 | -4.4821 | ##### | ####### | 4.2973E-150 | ######### |
| ENSG00000001084\|GCLC | 13514.4834 | -2.9506 | ##### | ####### | 4.3561E-150 | ######### |
| ENSG00000071575\|TRIB2 | 1190.2418 | 6.2873 | ##### | 26.0906 | 4.6631E-150 | ######### |
| ENSG00000060982\|BCAT1 | 2100.6863 | 3.9802 | ##### | 26.0092 | 3.8996E-149 | ######### |
| ENSG00000149485\|FADS1 | 14933.1407 | -2.5520 | ##### | ####### | 3.9272E-149 | ######### |
| ENSG00000142303\|ADAMTS10 | 1083.9597 | -4.8026 | ##### | ####### | 7.0966E-149 | ######### |
| ENSG00000173482\|PTPRM | 6632.1303 | -3.1769 | ##### | ####### | 8.7922E-149 | ######### |
| ENSG00000166394\|CYB5R2 | 946.9911 | 4.1903 | ##### | 25.9378 | 2.4959E-148 | ######### |
| ENSG00000167644\|C19orf33 | 4948.8119 | -3.4488 | ##### | ####### | 3.3306E-148 | ######### |
| ENSG00000108176\|DNAJC12 | 435.5341 | -5.3784 | ##### | ####### | 4.9935E-148 | ######### |
| ENSG00000101846\|STS | 883.4547 | -3.8292 | ##### | ####### | 9.0871E-148 | ######### |
| ENSG00000154529\|CNTNAP3B | 505.8094 | -4.7741 | ##### | ####### | 9.9537E-148 | ######### |
| ENSG00000164379\|FOXQ1 | 378.1442 | -5.1846 | ##### | ####### | 1.0693E-147 | ######### |
| ENSG00000154553\|PDLIM3 | 469.8067 | 5.1142 | ##### | 25.8723 | 1.3641E-147 | ######### |
| ENSG00000106070\|GRB10 | 3593.9267 | -2.7458 | ##### | ####### | 1.6408E-147 | ######### |
| ENSG00000184937\|WT1 | 2570.0037 | 11.0022 | ##### | 25.8531 | 2.2446E-147 | ######### |
| ENSG00000183780\|SLC35F3 | 1121.9381 | -4.3289 | ##### | ####### | 2.6503E-147 | ######### |
| ENSG00000183098\|GPC6 | 1707.7602 | -3.9039 | ##### | ####### | 1.0551E-146 | ######### |
| ENSG00000112576\|CCND3 | 18584.0262 | -3.1393 | ##### | ####### | 1.5316E-146 | ######### |
| ENSG00000065361\|ERBB3 | 897.1939 | -3.6735 | ##### | ####### | 2.5052E-146 | ######### |
| ENSG00000183691\|NOG | 754.7703 | 5.6150 | ##### | 25.7577 | 2.6414E-146 | ######### |
| ENSG00000170500\|LONRF2 | 5238.1515 | 3.4630 | ##### | 25.7139 | 8.1766E-146 | ######### |
| ENSG00000144712\|CAND2 | 643.1116 | 3.9072 | ##### | 25.7007 | 1.1476E-145 | ######### |
| ENSG00000130558\|OLFM1 | 2468.3160 | 3.6145 | ##### | 25.6569 | 3.5402E-145 | ######### |
| ENSG00000145476\|CYP4V2 | 783.5675 | -5.0253 | ##### | ####### | 4.2298E-145 | ######### |
| ENSG00000129354\|AP1M2 | 1635.5612 | -3.5644 | ##### | ####### | 8.0685E-145 | ######### |
| ENSG00000137842\|TMEM62 | 890.6987 | -3.7816 | ##### | ####### | 2.0279E-144 | ######### |
| ENSG00000101282\|RSPO4 | 636.7136 | 6.1436 | ##### | 25.5855 | 2.2124E-144 | ######### |
| ENSG00000128298\|BAIAP2L2 | 628.5501 | -4.1472 | ##### | ####### | 4.3268E-144 | ######### |
| ENSG00000276850 | 675.7454 | -5.2099 | ##### | ####### | 4.9054E-144 | ######### |
| ENSG00000182704\|TSKU | 7355.4500 | -3.2204 | ##### | ####### | 7.6282E-144 | ######### |
| ENSG00000142235\|LMTK3 | 1808.8543 | -3.4195 | ##### | ####### | 9.6429E-143 | ######### |
| ENSG00000285294\|LINC00842 | 929.9272 | -3.4806 | ##### | ####### | 1.3844E-142 | ######### |
| ENSG00000105223\|PLD3 | 17727.7921 | -3.1235 | ##### | ####### | 1.5935E-142 | ######### |
| ENSG00000099810\|MTAP | 2421.2095 | 3.4025 | ##### | 25.4164 | 1.6634E-142 | ######### |
| ENSG00000078900\|TP73 | 564.2015 | 6.4827 | ##### | 25.4072 | 2.0987E-142 | ######### |
| ENSG00000119866\|BCL11A | 1122.2764 | 8.5807 | ##### | 25.3890 | 3.3385E-142 | ######### |
| ENSG00000169213\|RAB3B | 2425.8480 | -3.6890 | ##### | ####### | 1.2369E-141 | ######### |
| ENSG00000100625\|SIX4 | 1384.9519 | 2.4459 | ##### | 25.3337 | 1.3602E-141 | ######### |

| | | | | | | |
|---|---|---|---|---|---|---|
| ENSG00000135750\|KCNK1 | 942.0531 | -7.0680 | ##### | ####### | 5.0207E-141 | ######### |
| ENSG00000082397\|EPB41L3 | 1214.6948 | 3.5642 | ##### | 25.2788 | 5.4698E-141 | ######### |
| ENSG00000138496\|PARP9 | 1160.6708 | -3.6834 | ##### | ####### | 5.5353E-141 | ######### |
| ENSG00000116604\|MEF2D | 14416.6463 | 1.8573 | ##### | 25.2383 | 1.5240E-140 | ######### |
| ENSG00000155980\|KIF5A | 685.2634 | 3.6219 | ##### | 25.2343 | 1.6864E-140 | ######### |
| ENSG00000267121\|FMNL1-DT | 731.1827 | -4.2958 | ##### | ####### | 1.9321E-140 | ######### |
| ENSG00000170382\|LRRN2 | 1151.1260 | -3.7057 | ##### | ####### | 2.1525E-140 | ######### |
| ENSG00000099337\|KCNK6 | 721.4118 | -3.9356 | ##### | ####### | 4.2485E-140 | ######### |
| ENSG00000162591\|MEGF6 | 2489.2521 | -3.4848 | ##### | ####### | 5.9146E-140 | ######### |
| ENSG00000138639\|ARHGAP24 | 629.6603 | -3.6439 | ##### | ####### | 1.1427E-139 | ######### |
| ENSG00000010704\|HFE | 710.6441 | 7.8297 | ##### | 25.1367 | 1.9767E-139 | ######### |
| ENSG00000163898\|LIPH | 1339.3029 | -6.2646 | ##### | ####### | 2.8911E-139 | ######### |
| ENSG00000226950\|DANCR | 5638.0663 | 3.1381 | ##### | 25.1207 | 2.9577E-139 | ######### |
| ENSG00000169432\|SCN9A | 617.1278 | 4.1071 | ##### | 25.0759 | 9.1097E-139 | ######### |
| ENSG00000157570\|TSPAN18 | 912.6205 | 3.8292 | ##### | 25.0622 | 1.2869E-138 | ######### |
| ENSG00000130508\|PXDN | 5558.7689 | 3.1861 | ##### | 25.0600 | 1.3593E-138 | ######### |
| ENSG00000158445\|KCNB1 | 1314.4181 | 4.0411 | ##### | 25.0435 | 2.0545E-138 | ######### |
| ENSG00000183722\|LHFPL6 | 1112.9588 | 2.8354 | ##### | 25.0298 | 2.8947E-138 | ######### |
| ENSG00000189223\|PAX8-AS1 | 1952.6945 | 4.3659 | ##### | 25.0190 | 3.7946E-138 | ######### |
| ENSG00000169908\|TM4SF1 | 31761.4284 | -3.0298 | ##### | ####### | 9.8944E-138 | ######### |
| ENSG00000128710\|HOXD10 | 440.8690 | 6.8833 | ##### | 24.9544 | 1.9122E-137 | ######### |
| ENSG00000005108\|THSD7A | 1213.9183 | -4.5156 | ##### | ####### | 2.3838E-137 | ######### |
| ENSG00000122691\|TWIST1 | 1748.3717 | 3.3193 | ##### | 24.9234 | 4.1446E-137 | ######### |
| ENSG00000134247\|PTGFRN | 2576.2453 | -3.1368 | ##### | ####### | 1.3836E-136 | ######### |
| ENSG00000112655\|PTK7 | 2009.0749 | 3.6639 | ##### | 24.8620 | 1.9174E-136 | ######### |
| ENSG00000168528\|SERINC2 | 2294.4497 | -3.8509 | ##### | ####### | 2.3686E-136 | ######### |
| ENSG00000187714\|SLC18A3 | 464.4468 | 4.7435 | ##### | 24.8278 | 4.4891E-136 | ######### |
| ENSG00000135052\|GOLM1 | 16457.3749 | -2.9896 | ##### | ####### | 7.1177E-136 | ######### |
| ENSG00000163364\|LINC01116 | 1081.3898 | 3.7374 | ##### | 24.7816 | 1.4151E-135 | ######### |
| ENSG00000198910\|L1CAM | 5987.6935 | 2.9395 | ##### | 24.7811 | 1.4334E-135 | ######### |
| ENSG00000225968\|ELFN1 | 763.3304 | 3.9365 | ##### | 24.7763 | 1.6166E-135 | ######### |
| ENSG00000213793\|ZNF888 | 517.1709 | -4.9473 | ##### | ####### | 2.5916E-135 | ######### |
| ENSG00000134962\|KLB | 353.1915 | -6.3161 | ##### | ####### | 3.8825E-135 | ######### |
| ENSG00000119888\|EPCAM | 578.0529 | -3.8068 | ##### | ####### | 5.7275E-135 | ######### |
| ENSG00000105357\|MYH14 | 4936.4749 | -3.0237 | ##### | ####### | 6.4913E-135 | ######### |
| ENSG00000119508\|NR4A3 | 1994.4551 | 3.0290 | ##### | 24.7033 | 9.8638E-135 | ######### |
| ENSG00000084636\|COL16A1 | 414.3483 | -5.2857 | ##### | ####### | 1.6331E-134 | ######### |
| ENSG00000130402\|ACTN4 | 64109.7894 | -2.1615 | ##### | ####### | 2.2286E-134 | ######### |
| ENSG00000119669\|IRF2BPL | 11182.5036 | 2.3278 | ##### | 24.6591 | 2.9406E-134 | ######### |
| ENSG00000140853\|NLRC5 | 506.6785 | -4.9084 | ##### | ####### | 3.6188E-134 | ######### |
| ENSG00000205809\|KLRC2 | 615.0641 | 4.9194 | ##### | 24.6466 | 4.0001E-134 | ######### |
| ENSG00000163661\|PTX3 | 3133.0154 | 3.6750 | ##### | 24.6381 | 4.9310E-134 | ######### |
| ENSG00000184347\|SLIT3 | 3714.1414 | 3.0180 | ##### | 24.5738 | 2.4069E-133 | ######### |
| ENSG00000156453\|PCDH1 | 709.7376 | -3.4038 | ##### | ####### | 4.4134E-133 | ######### |
| ENSG00000100031\|GGT1 | 1892.1478 | -7.0781 | ##### | ####### | 5.5996E-133 | ######### |
| ENSG00000133401\|PDZD2 | 833.6754 | -4.7770 | ##### | ####### | 6.7852E-133 | ######### |
| ENSG00000026559\|KCNG1 | 3231.8144 | 3.1129 | ##### | 24.5217 | 8.6663E-133 | ######### |
| ENSG00000245680\|ZNF585B | 861.2361 | -2.9917 | ##### | ####### | 1.0664E-132 | ######### |
| ENSG00000070404\|FSTL3 | 13497.3182 | -2.4630 | ##### | ####### | 1.2466E-132 | ######### |
| ENSG00000142694\|EVA1B | 784.6722 | 5.1978 | ##### | 24.4983 | 1.5388E-132 | ######### |
| ENSG00000172159\|FRMD3 | 1506.2455 | -5.9797 | ##### | ####### | 1.6161E-132 | ######### |
| ENSG00000162878\|PKDCC | 926.9172 | -3.5097 | ##### | ####### | 1.7947E-132 | ######### |
| ENSG00000100504\|PYGL | 20874.7107 | 1.2643 | ##### | 24.4838 | 2.1999E-132 | ######### |
| ENSG00000056998\|GYG2 | 717.3260 | -9.4323 | ##### | ####### | 2.4274E-132 | ######### |
| ENSG00000154237\|LRRK1 | 1664.1456 | -3.2457 | ##### | ####### | 3.5432E-132 | ######### |
| ENSG00000099219\|ERMP1 | 2898.9165 | -3.0255 | ##### | ####### | 4.0413E-132 | ######### |
| ENSG00000130513\|GDF15 | 1110.0627 | -3.5892 | ##### | ####### | 1.1039E-131 | ######### |
| ENSG00000080503\|SMARCA2 | 1229.1945 | -3.1725 | ##### | ####### | 4.3433E-131 | ######### |
| ENSG00000181885\|CLDN7 | 378.7687 | -7.5606 | ##### | ####### | 8.2762E-131 | ######### |
| ENSG00000121281\|ADCY7 | 1958.5075 | -3.4479 | ##### | ####### | 1.5999E-130 | ######### |
| ENSG00000161681\|SHANK1 | 433.8687 | 4.5899 | ##### | 24.3070 | 1.6543E-130 | ######### |
| ENSG00000184564\|SLITRK6 | 612.5916 | 5.8004 | ##### | 24.2993 | 1.9930E-130 | ######### |
| ENSG00000177106\|EPS8L2 | 1962.8111 | -3.2257 | ##### | ####### | 2.5072E-130 | ######### |

| | | | | | | |
|---|---|---|---|---|---|---|
| ENSG00000105499\|PLA2G4C | 709.2763 | -3.8797 | ##### | ####### | 3.1768E-130 | ######### |
| ENSG00000137968\|SLC44A5 | 497.3000 | -4.1492 | ##### | ####### | 4.3642E-130 | ######### |
| ENSG00000106976\|DNM1 | 3977.4142 | -2.9900 | ##### | ####### | 7.9383E-130 | ######### |
| ENSG00000106537\|TSPAN13 | 6308.2040 | -2.9813 | ##### | ####### | 1.0991E-129 | ######### |
| ENSG00000166501\|PRKCB | 1611.9128 | 3.5435 | ##### | 24.2141 | 1.5815E-129 | ######### |
| ENSG00000184867\|ARMCX2 | 2436.5995 | 3.2091 | ##### | 24.2088 | 1.7985E-129 | ######### |
| ENSG00000167799\|NUDT8 | 1114.7605 | 3.4065 | ##### | 24.2039 | 2.0249E-129 | ######### |
| ENSG00000164176\|EDIL3 | 1046.7624 | 6.0340 | ##### | 24.2019 | 2.1257E-129 | ######### |
| ENSG00000147257\|GPC3 | 487.4741 | 4.8282 | ##### | 24.1973 | 2.3779E-129 | ######### |
| ENSG00000162458\|FBLIM1 | 6501.6274 | -2.8843 | ##### | ####### | 7.9923E-129 | ######### |
| ENSG00000069011\|PITX1 | 1001.1534 | -3.7281 | ##### | ####### | 1.2546E-128 | ######### |
| ENSG00000125740\|FOSB | 641.8720 | 4.8149 | ##### | 24.1132 | 1.8177E-128 | ######### |
| ENSG00000185201\|IFITM2 | 681.4581 | -6.5742 | ##### | ####### | 1.9909E-128 | ######### |
| ENSG00000196793\|ZNF239 | 381.7579 | -3.8719 | ##### | ####### | 2.8246E-128 | ######### |
| ENSG00000187244\|BCAM | 2541.7797 | -3.1899 | ##### | ####### | 3.1784E-128 | ######### |
| ENSG00000170537\|TMC7 | 973.3733 | -4.0221 | ##### | ####### | 6.1804E-128 | ######### |
| ENSG00000117245\|KIF17 | 503.7453 | -4.8480 | ##### | ####### | 7.4812E-128 | ######### |
| ENSG00000104687\|GSR | 20446.2392 | -2.9831 | ##### | ####### | 7.8167E-128 | ######### |
| ENSG00000076344\|RGS11 | 636.0228 | -3.5638 | ##### | ####### | 8.0253E-128 | ######### |
| ENSG00000111052\|LIN7A | 1362.6134 | -5.0802 | ##### | ####### | 9.4889E-128 | ######### |
| ENSG00000215612\|HMX1 | 718.6696 | 7.5409 | ##### | 24.0237 | 1.5717E-127 | ######### |
| ENSG00000198879\|SFMBT2 | 684.3344 | 3.6852 | ##### | 24.0080 | 2.2919E-127 | ######### |
| ENSG00000161249\|DMKN | 2679.0214 | 3.1135 | ##### | 23.9850 | 3.9860E-127 | ######### |
| ENSG00000272068\|BCAN-AS1 | 430.2771 | -4.5433 | ##### | ####### | 8.3578E-127 | ######### |
| ENSG00000163694\|RBM47 | 3519.8104 | -3.0990 | ##### | ####### | 1.0962E-126 | ######### |
| ENSG00000155850\|SLC26A2 | 17813.5751 | -2.8547 | ##### | ####### | 1.8593E-126 | ######### |
| ENSG00000109452\|INPP4B | 1019.4479 | -3.6240 | ##### | ####### | 4.4179E-126 | ######### |
| ENSG00000164574\|GALNT10 | 18268.3472 | -2.8198 | ##### | ####### | 6.4346E-126 | ######### |
| ENSG00000155465\|SLC7A7 | 427.2578 | -5.2575 | ##### | ####### | 6.8345E-126 | ######### |
| ENSG00000196352\|CD55 | 8615.4709 | -2.4903 | ##### | ####### | 1.0735E-125 | ######### |
| ENSG00000261786 | 668.3163 | -4.1322 | ##### | ####### | 2.2324E-125 | ######### |
| ENSG00000099250\|NRP1 | 8737.3618 | -2.9547 | ##### | ####### | 4.5405E-125 | ######### |
| ENSG00000160223\|ICOSLG | 709.8774 | 3.6868 | ##### | 23.7752 | 6.0274E-125 | ######### |
| ENSG00000166963\|MAP1A | 2559.8780 | -3.3299 | ##### | ####### | 6.7711E-125 | ######### |
| ENSG00000111261\|MANSC1 | 1306.8899 | -3.5740 | ##### | ####### | 8.2546E-125 | ######### |
| ENSG00000123609\|NMI | 1910.1086 | -3.2343 | ##### | ####### | 9.6428E-125 | ######### |
| ENSG00000127561\|SYNGR3 | 1045.3143 | -3.5994 | ##### | ####### | 1.4249E-124 | ######### |
| ENSG00000071991\|CDH19 | 768.2885 | -3.4481 | ##### | ####### | 1.6572E-124 | ######### |
| ENSG00000198848\|CES1 | 22079.3842 | ####### | ##### | ####### | 1.7032E-124 | ######### |
| ENSG00000262454\|MIR193BHG | 639.2001 | -4.6503 | ##### | ####### | 4.0236E-124 | ######### |
| ENSG00000065621\|GSTO2 | 312.4254 | -4.9314 | ##### | ####### | 4.0990E-124 | ######### |
| ENSG00000121578\|B4GALT4 | 3836.2758 | -2.7850 | ##### | ####### | 4.9223E-124 | ######### |
| ENSG00000250510\|GPR162 | 486.7271 | 4.1122 | ##### | 23.6850 | 5.1458E-124 | ######### |
| ENSG00000135100\|HNF1A | 454.9273 | -4.0584 | ##### | ####### | 6.6577E-124 | ######### |
| ENSG00000089199\|CHGB | 1382.6303 | 4.7128 | ##### | 23.6634 | 8.5822E-124 | ######### |
| ENSG00000105270\|CLIP3 | 882.5580 | -3.6164 | ##### | ####### | 1.8565E-123 | ######### |
| ENSG00000285517 | 1650.6047 | 3.3737 | ##### | 23.6192 | 2.4449E-123 | ######### |
| ENSG00000151150\|ANK3 | 797.2483 | -3.6707 | ##### | ####### | 3.0115E-123 | ######### |
| ENSG00000184898\|RBM43 | 726.7555 | -4.2155 | ##### | ####### | 4.3113E-123 | ######### |
| ENSG00000101255\|TRIB3 | 2398.9332 | -2.9204 | ##### | ####### | 5.3470E-123 | ######### |
| ENSG00000023171\|GRAMD1B | 20917.7067 | 2.7869 | ##### | 23.5718 | 7.5067E-123 | ######### |
| ENSG00000130702\|LAMA5 | 20484.2822 | -2.8991 | ##### | ####### | 1.5357E-122 | ######### |
| ENSG00000171517\|LPAR3 | 740.8235 | 4.6268 | ##### | 23.5232 | 2.3599E-122 | ######### |
| ENSG00000161940\|BCL6B | 569.9575 | 4.1649 | ##### | 23.4428 | 1.5662E-121 | ######### |
| ENSG00000158089\|GALNT14 | 1796.0077 | 3.7486 | ##### | 23.4247 | 2.3935E-121 | ######### |
| ENSG00000115828\|QPCT | 1188.7320 | 4.4773 | ##### | 23.4097 | 3.4013E-121 | ######### |
| ENSG00000132429\|POPDC3 | 1262.8422 | -3.0778 | ##### | ####### | 4.1743E-121 | ######### |
| ENSG00000116678\|LEPR | 1161.2152 | -3.2631 | ##### | ####### | 4.3963E-121 | ######### |
| ENSG00000104626\|ERI1 | 11778.0483 | -2.8583 | ##### | ####### | 7.4587E-121 | ######### |
| ENSG00000163840\|DTX3L | 3036.5595 | -3.0234 | ##### | ####### | 1.0232E-120 | ######### |
| ENSG00000124225\|PMEPA1 | 20173.9280 | 2.3894 | ##### | 23.3458 | 1.5221E-120 | ######### |
| ENSG00000103534\|TMC5 | 656.1071 | -3.7203 | ##### | ####### | 3.1951E-120 | ######### |
| ENSG00000180530\|NRIP1 | 4087.3350 | -2.9593 | ##### | ####### | 6.8534E-120 | ######### |

| | | | | | | |
|---|---|---|---|---|---|---|
| ENSG00000237975\|FLG-AS1 | 846.2673 | 5.7188 | ##### | 23.2804 | 6.9991E-120 | ######### |
| ENSG00000259976 | 438.4965 | -3.9048 | ##### | ####### | 9.5300E-120 | ######### |
| ENSG00000064012\|CASP8 | 1901.5605 | -3.1250 | ##### | ####### | 1.2659E-119 | ######### |
| ENSG00000168003\|SLC3A2 | 31440.3570 | -2.8101 | ##### | ####### | 1.3197E-119 | ######### |
| ENSG00000105989\|WNT2 | 1133.6628 | 3.6362 | ##### | 23.2421 | 1.7102E-119 | ######### |
| ENSG00000073150\|PANX2 | 1198.0500 | -3.4744 | ##### | ####### | 2.9324E-119 | ######### |
| ENSG00000142227\|EMP3 | 8783.4313 | -2.8643 | ##### | ####### | 3.6385E-119 | ######### |
| ENSG00000067840\|PDZD4 | 668.3280 | 4.1177 | ##### | 23.1895 | 5.8067E-119 | ######### |
| ENSG00000096060\|FKBP5 | 6468.1223 | -2.8546 | ##### | ####### | 1.0386E-118 | ######### |
| ENSG00000277117 | 459.9926 | 4.5204 | ##### | 23.1569 | 1.2398E-118 | ######### |
| ENSG00000145284\|SCD5 | 863.3196 | 4.8659 | ##### | 23.1395 | 1.8557E-118 | ######### |
| ENSG00000140526\|ABHD2 | 15798.7037 | -2.7890 | ##### | ####### | 3.7120E-118 | ######### |
| ENSG00000008277\|ADAM22 | 706.4048 | 3.7482 | ##### | 23.1012 | 4.5020E-118 | ######### |
| ENSG00000019186\|CYP24A1 | 21261.2924 | -2.6172 | ##### | ####### | 4.5527E-118 | ######### |
| ENSG00000078399\|HOXA9 | 324.5733 | -4.8487 | ##### | ####### | 5.7315E-118 | ######### |
| ENSG00000134602\|STK26 | 4953.1744 | 3.0856 | ##### | 23.0901 | 5.8210E-118 | ######### |
| ENSG00000106460\|TMEM106B | 11392.8987 | -2.8568 | ##### | ####### | 1.2674E-117 | ######### |
| ENSG00000121716\|PILRB | 3222.4249 | 3.2165 | ##### | 23.0473 | 1.5638E-117 | ######### |
| ENSG00000113790\|EHHADH | 948.4104 | -3.4133 | ##### | ####### | 2.8049E-117 | ######### |
| ENSG00000054356\|PTPRN | 2615.9092 | -3.0325 | ##### | ####### | 7.9238E-117 | ######### |
| ENSG00000002834\|LASP1 | 37242.5670 | -1.7027 | ##### | ####### | 8.3463E-117 | ######### |
| ENSG00000167306\|MYO5B | 1340.8655 | -4.8271 | ##### | ####### | 8.8269E-117 | ######### |
| ENSG00000160211\|G6PD | 36947.2079 | -2.6540 | ##### | ####### | 2.1291E-116 | ######### |
| ENSG00000139112\|GABARAPL1 | 12493.5823 | 1.7730 | ##### | 22.9336 | 2.1460E-116 | ######### |
| ENSG00000164761\|TNFRSF11B | 2020.0257 | 2.8647 | ##### | 22.8699 | 9.2562E-116 | ######### |
| ENSG00000133687\|TMTC1 | 700.9772 | 4.4472 | ##### | 22.8532 | 1.3582E-115 | ######### |
| ENSG00000141429\|GALNT1 | 9244.0726 | -2.8279 | ##### | ####### | 1.5688E-115 | ######### |
| ENSG00000131711\|MAP1B | 9672.5844 | -2.7313 | ##### | ####### | 1.7940E-115 | ######### |
| ENSG00000123094\|RASSF8 | 13219.1428 | 2.6454 | ##### | 22.8142 | 3.3166E-115 | ######### |
| ENSG00000147155\|EBP | 1319.6229 | 4.6974 | ##### | 22.8060 | 3.9934E-115 | ######### |
| ENSG00000078018\|MAP2 | 3655.5272 | -2.9611 | ##### | ####### | 1.0817E-114 | ######### |
| ENSG00000054277\|OPN3 | 9904.3334 | -3.1625 | ##### | ####### | 1.5594E-114 | ######### |
| ENSG00000173193\|PARP14 | 2495.5211 | -3.2075 | ##### | ####### | 2.0080E-114 | ######### |
| ENSG00000100095\|SEZ6L | 828.5770 | 4.2513 | ##### | 22.7319 | 2.1679E-114 | ######### |
| ENSG00000140044\|JDP2 | 686.9114 | -3.7067 | ##### | ####### | 4.6081E-114 | ######### |
| ENSG00000167772\|ANGPTL4 | 5620.8451 | -2.7559 | ##### | ####### | 9.4219E-114 | ######### |
| ENSG00000106605\|BLVRA | 2971.7257 | -2.8768 | ##### | ####### | 4.4935E-113 | ######### |
| ENSG00000074964\|ARHGEF10L | 1396.5344 | -3.0613 | ##### | ####### | 7.1074E-113 | ######### |
| ENSG00000278384 | 1129.0214 | 3.4577 | ##### | 22.5762 | 7.4282E-113 | ######### |
| ENSG00000167972\|ABCA3 | 6157.8262 | -2.7369 | ##### | ####### | 1.4327E-112 | ######### |
| ENSG00000138646\|HERC5 | 1379.3672 | 2.7533 | ##### | 22.5193 | 2.6830E-112 | ######### |
| ENSG00000101049\|SGK2 | 439.7025 | -5.0755 | ##### | ####### | 2.6894E-112 | ######### |
| ENSG00000165105\|RASEF | 345.7833 | -6.4695 | ##### | ####### | 3.5480E-112 | ######### |
| ENSG00000138166\|DUSP5 | 4953.3188 | -2.9748 | ##### | ####### | 3.6082E-112 | ######### |
| ENSG00000102554\|KLF5 | 2870.8358 | -2.8538 | ##### | ####### | 5.5797E-112 | ######### |
| ENSG00000183770\|FOXL2 | 665.6699 | 6.8413 | ##### | 22.4792 | 6.6290E-112 | ######### |
| ENSG00000111981\|ULBP1 | 511.8310 | 4.2968 | ##### | 22.4547 | 1.1513E-111 | ######### |
| ENSG00000174307\|PHLDA3 | 2041.6625 | -3.0509 | ##### | ####### | 1.6615E-111 | ######### |
| ENSG00000130477\|UNC13A | 2009.2462 | -2.8790 | ##### | ####### | 1.8890E-111 | ######### |
| ENSG00000146166\|LGSN | 1917.6621 | -2.9686 | ##### | ####### | 2.0604E-111 | ######### |
| ENSG00000182534\|MXRA7 | 4495.0957 | 3.0191 | ##### | 22.4208 | 2.4693E-111 | ######### |
| ENSG00000104361\|NIPAL2 | 1879.6603 | -2.9084 | ##### | ####### | 2.5756E-111 | ######### |
| ENSG00000090975\|PITPNM2 | 2094.5647 | -2.8881 | ##### | ####### | 2.7348E-111 | ######### |
| ENSG00000158104\|HPD | 3024.9112 | -9.5462 | ##### | ####### | 2.9605E-111 | ######### |
| ENSG00000040275\|SPDL1 | 7796.0304 | 2.0539 | ##### | 22.4035 | 3.6360E-111 | ######### |
| ENSG00000179083\|FAM133A | 535.8944 | -6.5576 | ##### | ####### | 6.3747E-111 | ######### |
| ENSG00000188910\|GJB3 | 857.7910 | -3.2673 | ##### | ####### | 8.3899E-111 | ######### |
| ENSG00000187446\|CHP1 | 10764.6324 | -2.0656 | ##### | ####### | 1.3912E-110 | ######### |
| ENSG00000278709\|NKILA | 2030.3665 | 3.2301 | ##### | 22.3077 | 3.1101E-110 | ######### |
| ENSG00000077942\|FBLN1 | 12190.5511 | -2.7578 | ##### | ####### | 3.6434E-110 | ######### |
| ENSG00000151491\|EPS8 | 2426.7383 | -2.9133 | ##### | ####### | 5.2732E-110 | ######### |
| ENSG00000080031\|PTPRH | 5066.7649 | 2.7467 | ##### | 22.2786 | 5.9537E-110 | ######### |
| ENSG00000120833\|SOCS2 | 3558.6561 | 3.0096 | ##### | 22.2725 | 6.8234E-110 | ######### |

| | | | | | | |
|---|---|---|---|---|---|---|
| ENSG00000188763\|FZD9 | 282.9978 | -6.8729 | ##### | ####### | 9.5180E-110 | ######### |
| ENSG00000249199 | 365.6564 | -3.9100 | ##### | ####### | 1.2657E-109 | ######### |
| ENSG00000180628\|PCGF5 | 3663.1997 | -2.8958 | ##### | ####### | 1.7785E-109 | ######### |
| ENSG00000180340\|FZD2 | 2160.7833 | -2.1651 | ##### | ####### | 1.9175E-109 | ######### |
| ENSG00000134569\|LRP4 | 1092.9319 | 3.7357 | ##### | 22.1970 | 3.6686E-109 | ######### |
| ENSG00000293339 | 7910.4715 | 2.2383 | ##### | 22.1828 | 5.0358E-109 | ######### |
| ENSG00000189190\|ZNF600 | 541.7247 | -3.8797 | ##### | ####### | 5.6099E-109 | ######### |
| ENSG00000138829\|FBN2 | 1502.7029 | -3.2107 | ##### | ####### | 6.7194E-109 | ######### |
| ENSG00000173227\|SYT12 | 537.3753 | -3.6650 | ##### | ####### | 9.5608E-109 | ######### |
| ENSG00000167363\|FN3K | 501.7662 | -3.8092 | ##### | ####### | 1.8257E-108 | ######### |
| ENSG00000221926\|TRIM16 | 4282.1375 | -3.1622 | ##### | ####### | 2.1640E-108 | ######### |
| ENSG00000005513\|SOX8 | 313.4196 | 4.6816 | ##### | 22.1161 | 2.2105E-108 | ######### |
| ENSG00000050438\|SLC4A8 | 1211.5553 | 2.9327 | ##### | 22.0963 | 3.4285E-108 | ######### |
| ENSG00000108602\|ALDH3A1 | 50622.0902 | ####### | ##### | ####### | 3.5458E-108 | ######### |
| ENSG00000214548\|MEG3 | 323.7264 | 4.8385 | ##### | 22.0829 | 4.6194E-108 | ######### |
| ENSG00000279414\|CDRT15P9 | 342.3518 | 6.5069 | ##### | 22.0759 | 5.3887E-108 | ######### |
| ENSG00000204604\|ZNF468 | 1589.7816 | -2.8928 | ##### | ####### | 5.5542E-108 | ######### |
| ENSG00000114771\|AADAC | 208.5509 | -5.3775 | ##### | ####### | 5.7248E-108 | ######### |
| ENSG00000143512\|HHIPL2 | 1113.0275 | -3.1614 | ##### | ####### | 1.1264E-107 | ######### |
| ENSG00000111321\|LTBR | 15097.6070 | ####### | ##### | ####### | 1.4698E-107 | ######### |
| ENSG00000149328\|GLB1L2 | 1344.5953 | 3.2075 | ##### | 22.0145 | 2.0931E-107 | ######### |
| ENSG00000170961\|HAS2 | 1948.9976 | 2.8979 | ##### | 21.9854 | 3.9759E-107 | ######### |
| ENSG00000152078\|TLCD4 | 1699.3382 | -3.0179 | ##### | ####### | 5.4745E-107 | ######### |
| ENSG00000086061\|DNAJA1 | 34307.8771 | 1.5406 | ##### | 21.9690 | 5.6963E-107 | ######### |
| ENSG00000101447\|FAM83D | 8643.4400 | 2.2015 | ##### | 21.9424 | 1.0246E-106 | ######### |
| ENSG00000025708\|TYMP | 7376.3160 | -2.6508 | ##### | ####### | 1.0762E-106 | ######### |
| ENSG00000157601\|MX1 | 303.0103 | -4.4584 | ##### | ####### | 1.1530E-106 | ######### |
| ENSG00000235437\|LINC01278 | 723.6585 | -3.1820 | ##### | ####### | 1.7200E-106 | ######### |
| ENSG00000072133\|RPS6KA6 | 1874.0080 | 2.7035 | ##### | 21.9119 | 1.9994E-106 | ######### |
| ENSG00000189369\|GSPT2 | 853.3675 | 3.8905 | ##### | 21.8638 | 5.7404E-106 | ######### |
| ENSG00000289911 | 543.1004 | -3.2990 | ##### | ####### | 6.0818E-106 | ######### |
| ENSG00000135905\|DOCK10 | 2740.2569 | -2.7634 | ##### | ####### | 9.5015E-106 | ######### |
| ENSG00000163082\|SGPP2 | 412.8755 | -4.4149 | ##### | ####### | 1.0523E-105 | ######### |
| ENSG00000152669\|CCNO | 916.2500 | -3.1900 | ##### | ####### | 1.1772E-105 | ######### |
| ENSG00000115009\|CCL20 | 1739.6850 | 5.3078 | ##### | 21.8304 | 1.1933E-105 | ######### |
| ENSG00000087258\|GNAO1 | 1134.3519 | 3.2233 | ##### | 21.7901 | 2.8778E-105 | ######### |
| ENSG00000182585\|EPGN | 1215.8894 | 2.9609 | ##### | 21.7379 | 8.9986E-105 | ######### |
| ENSG00000005102\|MEOX1 | 1420.7800 | 9.4878 | ##### | 21.7376 | 9.0460E-105 | ######### |
| ENSG00000121152\|NCAPH | 9644.1004 | 2.6569 | ##### | 21.7130 | 1.5479E-104 | ######### |
| ENSG00000099282\|TSPAN15 | 2919.0256 | -2.0628 | ##### | ####### | 1.6725E-104 | ######### |
| ENSG00000138944\|SHISAL1 | 861.1097 | -3.5228 | ##### | ####### | 1.8025E-104 | ######### |
| ENSG00000162722\|TRIM58 | 2183.5641 | 3.0343 | ##### | 21.7012 | 1.9988E-104 | ######### |
| ENSG00000142552\|RCN3 | 462.6724 | -4.0113 | ##### | ####### | 3.3538E-104 | ######### |
| ENSG00000124102\|PI3 | 607.0423 | -5.8506 | ##### | ####### | 3.4172E-104 | ######### |
| ENSG00000104427\|ZC2HC1A | 1489.4160 | -3.0826 | ##### | ####### | 4.2515E-104 | ######### |
| ENSG00000157483\|MYO1E | 8654.2481 | -2.5735 | ##### | ####### | 5.9900E-104 | ######### |
| ENSG00000169740\|ZNF32 | 809.7845 | -3.1097 | ##### | ####### | 6.2948E-104 | ######### |
| ENSG00000178695\|KCTD12 | 730.9325 | 3.0360 | ##### | 21.6315 | 9.0766E-104 | ######### |
| ENSG00000175591\|P2RY2 | 873.9134 | -3.3929 | ##### | ####### | 9.3570E-104 | ######### |
| ENSG00000178919\|FOXE1 | 5843.3702 | 2.9076 | ##### | 21.6286 | 9.6598E-104 | ######### |
| ENSG00000196535\|MYO18A | 4607.3476 | -2.7497 | ##### | ####### | 1.4682E-103 | ######### |
| ENSG00000007237\|GAS7 | 846.1267 | 4.0305 | ##### | 21.5906 | 2.2003E-103 | ######### |
| ENSG00000197565\|COL4A6 | 1925.3953 | 3.4224 | ##### | 21.5835 | 2.5679E-103 | ######### |
| ENSG00000079739\|PGM1 | 9025.8275 | -2.6510 | ##### | ####### | 3.1070E-103 | ######### |
| ENSG00000180263\|FGD6 | 6039.1934 | -2.6281 | ##### | ####### | 4.3190E-103 | ######### |
| ENSG00000128713\|HOXD11 | 313.3659 | 5.6279 | ##### | 21.5354 | 7.2603E-103 | ######### |
| ENSG00000187098\|MITF | 1121.4660 | -3.3712 | ##### | ####### | 8.8888E-103 | ######### |
| ENSG00000169242\|EFNA1 | 7560.8031 | 2.7469 | ##### | 21.5021 | 1.4881E-102 | ######### |
| ENSG00000173281\|PPP1R3B | 3583.1387 | -2.7767 | ##### | ####### | 2.1711E-102 | ######### |
| ENSG00000047230\|CTPS2 | 805.1551 | -3.2385 | ##### | ####### | 2.3817E-102 | ######### |
| ENSG00000003147\|ICA1 | 722.5324 | -3.2322 | ##### | ####### | 2.7912E-102 | ######### |
| ENSG00000253537\|PCDHGA7 | 320.4642 | -4.5604 | ##### | ####### | 5.4556E-102 | ######### |
| ENSG00000073060\|SCARB1 | 15640.5627 | -2.0174 | ##### | ####### | 7.7453E-102 | ######### |

| | | | | | | |
|---|---|---|---|---|---|---|
| ENSG00000175793\|SFN | 17456.0764 | -2.5888 | ##### | ####### | 9.2790E-102 | ######### |
| ENSG00000176293\|ZNF135 | 988.4614 | -3.1648 | ##### | ####### | 1.3200E-101 | ######### |
| ENSG00000122729\|ACO1 | 3501.7073 | -2.6338 | ##### | ####### | 1.6666E-101 | ######### |
| ENSG00000148120\|AOPEP | 1554.2392 | -2.8607 | ##### | ####### | 1.9280E-101 | ######### |
| ENSG00000132164\|SLC6A11 | 405.6467 | 6.0878 | ##### | 21.3767 | 2.2032E-101 | ######### |
| ENSG00000114270\|COL7A1 | 2133.8441 | -2.8819 | ##### | ####### | 3.3957E-101 | 1.4529E-99 |
| ENSG00000096433\|ITPR3 | 18228.3256 | -2.5443 | ##### | ####### | 8.3954E-101 | 3.5870E-99 |
| ENSG00000117525\|F3 | 606.2356 | -3.3076 | ##### | ####### | 1.1411E-100 | 4.8684E-99 |
| ENSG00000111206\|FOXM1 | 27039.8568 | 1.7749 | ##### | 21.2853 | 1.5536E-100 | 6.6190E-99 |
| ENSG00000196116\|TDRD7 | 1379.2793 | -2.7514 | ##### | ####### | 4.4815E-100 | 1.9065E-98 |
| ENSG00000268686 | 314.8270 | 4.3699 | ##### | 21.2329 | 4.7409E-100 | 2.0140E-98 |
| ENSG00000107249\|GLIS3 | 1461.7112 | -2.7053 | ##### | ####### | 5.1894E-100 | 2.2014E-98 |
| ENSG00000271303\|SRXN1 | 33147.6356 | -2.6079 | ##### | ####### | 9.8181E-100 | 4.1591E-98 |
| ENSG00000173762\|CD7 | 429.9115 | 6.7199 | ##### | 21.1864 | 1.2749E-99 | 5.3932E-98 |
| ENSG00000205133\|TRIQK | 1646.0888 | -2.9004 | ##### | ####### | 1.8193E-99 | 7.6853E-98 |
| ENSG00000123684\|LPGAT1 | 10885.4287 | -2.4801 | ##### | ####### | 2.3385E-99 | 9.8644E-98 |
| ENSG00000131981\|LGALS3 | 6262.8912 | -2.6914 | ##### | ####### | 3.6845E-99 | 1.5520E-97 |
| ENSG00000035862\|TIMP2 | 16518.0886 | -2.5876 | ##### | ####### | 5.8243E-99 | 2.4499E-97 |
| ENSG00000151468\|CCDC3 | 466.3315 | 4.4141 | ##### | 21.1005 | 7.8635E-99 | 3.3030E-97 |
| ENSG00000106123\|EPHB6 | 367.3666 | 5.2398 | ##### | 21.0950 | 8.8328E-99 | 3.7049E-97 |
| ENSG00000126351\|THRA | 1753.7139 | -2.3022 | ##### | ####### | 1.0871E-98 | 4.5534E-97 |
| ENSG00000164093\|PITX2 | 396.7037 | 5.0104 | ##### | 21.0833 | 1.1317E-98 | 4.7336E-97 |
| ENSG00000173210\|ABLIM3 | 787.9977 | -3.0141 | ##### | ####### | 1.2536E-98 | 5.2361E-97 |
| ENSG00000119698\|PPP4R4 | 523.6069 | 5.6103 | ##### | 21.0757 | 1.3302E-98 | 5.5484E-97 |
| ENSG00000170421\|KRT8 | ########## | -2.4062 | ##### | ####### | 1.3431E-98 | 5.5942E-97 |
| ENSG00000280287 | 416.8315 | 3.6226 | ##### | 21.0697 | 1.5096E-98 | 6.2791E-97 |
| ENSG00000169116\|PARM1 | 613.2204 | -2.8155 | ##### | ####### | 1.5418E-98 | 6.4041E-97 |
| ENSG00000088826\|SMOX | 3748.9173 | -2.6192 | ##### | ####### | 1.7838E-98 | 7.3993E-97 |
| ENSG00000221829\|FANCG | 5190.5996 | 2.1957 | ##### | 21.0584 | 1.9169E-98 | 7.9402E-97 |
| ENSG00000178980\|SELENOW | 8992.8793 | -2.5311 | ##### | ####### | 2.2429E-98 | 9.2777E-97 |
| ENSG00000196542\|SPTSSB | 199.3120 | -5.3532 | ##### | ####### | 2.3438E-98 | 9.6817E-97 |
| ENSG00000122863\|CHST3 | 6372.7642 | -2.5072 | ##### | ####### | 4.2294E-98 | 1.7446E-96 |
| ENSG00000135083\|CCNJL | 807.8860 | -3.1665 | ##### | ####### | 5.0171E-98 | 2.0667E-96 |
| ENSG00000108175\|ZMIZ1 | 2842.3271 | -2.6235 | ##### | ####### | 6.5209E-98 | 2.6825E-96 |
| ENSG00000105738\|SIPA1L3 | 4102.7126 | -2.5723 | ##### | ####### | 7.2384E-98 | 2.9735E-96 |
| ENSG00000117791\|MTARC2 | 313.4935 | -6.2461 | ##### | ####### | 9.4964E-98 | 3.8957E-96 |
| ENSG00000112972\|HMGCS1 | 14936.5512 | -2.2078 | ##### | ####### | 1.1473E-97 | 4.7003E-96 |
| ENSG00000147676\|MAL2 | 1271.0554 | -2.7545 | ##### | ####### | 1.3368E-97 | 5.4688E-96 |
| ENSG00000126001\|CEP250 | 8492.6737 | 2.4822 | ##### | 20.9592 | 1.5474E-97 | 6.3220E-96 |
| ENSG00000173120\|KDM2A | 29418.6504 | 2.4548 | ##### | 20.9488 | 1.9252E-97 | 7.8545E-96 |
| ENSG00000184922\|FMNL1 | 8651.7804 | -2.5335 | ##### | ####### | 2.0788E-97 | 8.4699E-96 |
| ENSG00000064932\|SBNO2 | 9527.0313 | 2.3720 | ##### | 20.9329 | 2.6871E-97 | 1.0933E-95 |
| ENSG00000101265\|RASSF2 | 699.8935 | -3.0869 | ##### | ####### | 2.9250E-97 | 1.1885E-95 |
| ENSG00000118495\|PLAGL1 | 443.3052 | -3.3661 | ##### | ####### | 3.6298E-97 | 1.4729E-95 |
| ENSG00000242086\|MUC20-OT1 | 3825.1701 | 2.7377 | ##### | 20.9169 | 3.7584E-97 | 1.5230E-95 |
| ENSG00000263528\|IKBKE | 565.3663 | -8.1152 | ##### | ####### | 3.9386E-97 | 1.5939E-95 |
| ENSG00000131398\|KCNC3 | 1915.7785 | 2.6764 | ##### | 20.9102 | 4.3271E-97 | 1.7487E-95 |
| ENSG00000155307\|SAMSN1 | 244.4025 | -6.2987 | ##### | ####### | 4.6537E-97 | 1.8781E-95 |
| ENSG00000225614\|ZNF469 | 1060.8162 | -3.1705 | ##### | ####### | 6.5729E-97 | 2.6491E-95 |
| ENSG00000198467\|TPM2 | 57103.9028 | 2.5062 | ##### | 20.8818 | 7.8350E-97 | 3.1536E-95 |
| ENSG00000182986\|ZNF320 | 1074.0139 | -3.0117 | ##### | ####### | 9.1054E-97 | 3.6599E-95 |
| ENSG00000150961\|SEC24D | 3964.3775 | -2.5490 | ##### | ####### | 1.0014E-96 | 4.0199E-95 |
| ENSG00000265933\|LINC00668 | 855.0485 | 3.3580 | ##### | 20.8695 | 1.0141E-96 | 4.0651E-95 |
| ENSG00000147224\|PRPS1 | 3097.6728 | -2.7113 | ##### | ####### | 1.0774E-96 | 4.3133E-95 |
| ENSG00000158710\|TAGLN2 | ########## | 2.5182 | ##### | 20.8548 | 1.3799E-96 | 5.5167E-95 |
| ENSG00000000971\|CFH | 3306.5207 | -9.8351 | ##### | ####### | 1.6108E-96 | 6.4315E-95 |
| ENSG00000164294\|GPX8 | 5158.3166 | -2.5064 | ##### | ####### | 1.9534E-96 | 7.7889E-95 |
| ENSG00000115232\|ITGA4 | 1496.7444 | -5.4455 | ##### | ####### | 5.0451E-96 | 2.0090E-94 |
| ENSG00000290682\|STAG3L5P | 1249.4434 | 3.1167 | ##### | 20.7919 | 5.1248E-96 | 2.0380E-94 |
| ENSG00000123143\|PKN1 | 8530.0823 | -2.0781 | ##### | ####### | 6.7362E-96 | 2.6752E-94 |
| ENSG00000166689\|PLEKHA7 | 270.6287 | -4.5129 | ##### | ####### | 8.7517E-96 | 3.4710E-94 |
| ENSG00000160460\|SPTBN4 | 635.2696 | -3.1739 | ##### | ####### | 1.2836E-95 | 5.0844E-94 |
| ENSG00000118402\|ELOVL4 | 751.3469 | 3.3643 | ##### | 20.7473 | 1.2972E-95 | 5.1313E-94 |

| | | | | | | |
|---|---|---|---|---|---|---|
| ENSG00000090376\|IRAK3 | 856.8616 | -3.4594 | ##### | ####### | 1.3527E-95 | 5.3436E-94 |
| ENSG00000168917\|SLC35G2 | 1434.9523 | -2.7065 | ##### | ####### | 1.5361E-95 | 6.0603E-94 |
| ENSG00000100439\|ABHD4 | 2969.6813 | -2.6388 | ##### | ####### | 1.5951E-95 | 6.2847E-94 |
| ENSG00000060656\|PTPRU | 1285.6647 | -2.7029 | ##### | ####### | 1.7414E-95 | 6.8520E-94 |
| ENSG00000253293\|HOXA10 | 728.1641 | -5.3515 | ##### | ####### | 2.8133E-95 | 1.1055E-93 |
| ENSG00000067445\|TRO | 757.8349 | 3.3575 | ##### | 20.7084 | 2.9120E-95 | 1.1428E-93 |
| ENSG00000185483\|ROR1 | 1769.7012 | -2.5960 | ##### | ####### | 3.1077E-95 | 1.2180E-93 |
| ENSG00000092621\|PHGDH | 1010.4050 | -2.8257 | ##### | ####### | 3.5072E-95 | 1.3728E-93 |
| ENSG00000168758\|SEMA4C | 6524.9559 | 2.4250 | ##### | 20.6931 | 4.0009E-95 | 1.5639E-93 |
| ENSG00000266208\|GJD3-AS1 | 706.0542 | -4.0311 | ##### | ####### | 5.0121E-95 | 1.9567E-93 |
| ENSG00000143409\|MINDY1 | 542.9721 | -4.4296 | ##### | ####### | 5.4497E-95 | 2.1247E-93 |
| ENSG00000108846\|ABCC3 | 17521.9720 | -2.5963 | ##### | ####### | 7.1213E-95 | 2.7729E-93 |
| ENSG00000121743\|GJA3 | 775.7570 | 6.0816 | ##### | 20.6560 | 8.6233E-95 | 3.3533E-93 |
| ENSG00000243649\|CFB | 436.1356 | -4.7047 | ##### | ####### | 1.0296E-94 | 3.9987E-93 |
| ENSG00000135828\|RNASEL | 461.2753 | -3.2152 | ##### | ####### | 1.3527E-94 | 5.2465E-93 |
| ENSG00000228630\|HOTAIR | 573.4034 | 3.3485 | ##### | 20.6336 | 1.3705E-94 | 5.3087E-93 |
| ENSG00000154134\|ROBO3 | 1049.0657 | 3.4318 | ##### | 20.6280 | 1.5397E-94 | 5.9564E-93 |
| ENSG00000162909\|CAPN2 | 33530.7027 | -1.9695 | ##### | ####### | 2.1604E-94 | 8.3468E-93 |
| ENSG00000120254\|MTHFD1L | 11154.4995 | 2.4364 | ##### | 20.6066 | 2.3972E-94 | 9.2496E-93 |
| ENSG00000122642\|FKBP9 | 13791.2187 | -1.5898 | ##### | ####### | 3.2380E-94 | 1.2478E-92 |
| ENSG00000174799\|CEP135 | 3138.2579 | 2.5634 | ##### | 20.5860 | 3.6616E-94 | 1.4092E-92 |
| ENSG00000145685\|LHFPL2 | 4066.9381 | -1.9127 | ##### | ####### | 3.7945E-94 | 1.4585E-92 |
| ENSG00000250899 | 3326.5378 | 2.5997 | ##### | 20.5802 | 4.1276E-94 | 1.5844E-92 |
| ENSG00000136158\|SPRY2 | 1784.2702 | 2.7172 | ##### | 20.5697 | 5.1268E-94 | 1.9655E-92 |
| ENSG00000163013\|FBXO41 | 2489.2283 | 2.3646 | ##### | 20.5696 | 5.1361E-94 | 1.9665E-92 |
| ENSG00000111012\|CYP27B1 | 1678.6278 | 2.9334 | ##### | 20.5671 | 5.4080E-94 | 2.0680E-92 |
| ENSG00000203727\|SAMD5 | 1170.6532 | -2.8677 | ##### | ####### | 6.9318E-94 | 2.6473E-92 |
| ENSG00000224189\|HAGLR | 388.9858 | 3.7223 | ##### | 20.5537 | 7.1353E-94 | 2.7215E-92 |
| ENSG00000179403\|VWA1 | 2303.8551 | -2.7193 | ##### | ####### | 7.6855E-94 | 2.9277E-92 |
| ENSG00000161653\|NAGS | 837.9075 | 3.3032 | ##### | 20.5386 | 9.7288E-94 | 3.7013E-92 |
| ENSG00000163873\|GRIK3 | 808.2848 | 8.5832 | ##### | 20.5305 | 1.1507E-93 | 4.3723E-92 |
| ENSG00000164742\|ADCY1 | 930.8541 | 3.3355 | ##### | 20.5193 | 1.4490E-93 | 5.4988E-92 |
| ENSG00000213625\|LEPROT | 8347.9273 | -2.0647 | ##### | ####### | 1.4915E-93 | 5.6527E-92 |
| ENSG00000065809\|FAM107B | 5419.3268 | -2.4126 | ##### | ####### | 1.8566E-93 | 7.0276E-92 |
| ENSG00000105963\|ADAP1 | 584.8037 | -3.6761 | ##### | ####### | 1.9168E-93 | 7.2460E-92 |
| ENSG00000108515\|ENO3 | 4327.8018 | 2.5857 | ##### | 20.4924 | 2.5151E-93 | 9.4962E-92 |
| ENSG00000160213\|CSTB | 7018.6634 | -2.6131 | ##### | ####### | 2.9917E-93 | 1.1281E-91 |
| ENSG00000101104\|PABPC1L | 1584.6143 | 2.6780 | ##### | 20.4813 | 3.1616E-93 | 1.1907E-91 |
| ENSG00000178913\|TAF7 | 14213.7871 | -1.4125 | ##### | ####### | 3.1893E-93 | 1.1996E-91 |
| ENSG00000136770\|DNAJC1 | 4326.4789 | -2.4956 | ##### | ####### | 3.1976E-93 | 1.2012E-91 |
| ENSG00000078687\|TNRC6C | 936.9748 | 2.6754 | ##### | 20.4796 | 3.2702E-93 | 1.2269E-91 |
| ENSG00000118946\|PCDH17 | 428.7171 | -3.7391 | ##### | ####### | 3.9659E-93 | 1.4861E-91 |
| ENSG00000111788\|DDX12B | 1804.6928 | 2.5340 | ##### | 20.4596 | 4.9379E-93 | 1.8480E-91 |
| ENSG00000161011\|SQSTM1 | 83157.0374 | -2.4969 | ##### | ####### | 5.1482E-93 | 1.9243E-91 |
| ENSG00000231806\|PCAT7 | 306.8845 | -3.3686 | ##### | ####### | 6.7062E-93 | 2.5035E-91 |
| ENSG00000125731\|SH2D3A | 992.4816 | -2.7617 | ##### | ####### | 6.9136E-93 | 2.5777E-91 |
| ENSG00000147592\|LACTB2 | 3135.3724 | -2.6000 | ##### | ####### | 8.1072E-93 | 3.0190E-91 |
| ENSG00000085831\|TTC39A | 507.5373 | -5.8185 | ##### | ####### | 1.0684E-92 | 3.9737E-91 |
| ENSG00000170837\|GPR27 | 265.0448 | -3.8755 | ##### | ####### | 1.2908E-92 | 4.7948E-91 |
| ENSG00000124788\|ATXN1 | 1861.2028 | 2.5283 | ##### | 20.4072 | 1.4423E-92 | 5.3509E-91 |
| ENSG00000143621\|ILF2 | 48997.5051 | 1.9175 | ##### | 20.4058 | 1.4857E-92 | 5.5050E-91 |
| ENSG00000182871\|COL18A1 | 6373.3960 | -2.4362 | ##### | ####### | 1.8013E-92 | 6.6663E-91 |
| ENSG00000187185 | 455.7963 | -3.3532 | ##### | ####### | 1.8175E-92 | 6.7179E-91 |
| ENSG00000076554\|TPD52 | 12918.1651 | -2.5108 | ##### | ####### | 1.8853E-92 | 6.9599E-91 |
| ENSG00000267062 | 192.1047 | -5.2391 | ##### | ####### | 2.1169E-92 | 7.8050E-91 |
| ENSG00000134285\|FKBP11 | 1949.5928 | 2.8245 | ##### | 20.3437 | 5.2832E-92 | 1.9455E-90 |
| ENSG00000136167\|LCP1 | 317.9323 | -3.9832 | ##### | ####### | 5.3599E-92 | 1.9713E-90 |
| ENSG00000145242\|EPHA5 | 852.3876 | 3.3972 | ##### | 20.3291 | 7.1119E-92 | 2.6125E-90 |
| ENSG00000186806\|VSIG10L | 1780.2582 | -2.9511 | ##### | ####### | 9.3502E-92 | 3.4305E-90 |
| ENSG00000139410\|SDSL | 1278.9730 | -2.7755 | ##### | ####### | 1.0098E-91 | 3.7004E-90 |
| ENSG00000167216\|KATNAL2 | 803.0433 | -3.0322 | ##### | ####### | 2.0126E-91 | 7.3661E-90 |
| ENSG00000198513\|ATL1 | 866.1409 | -2.6579 | ##### | ####### | 2.0408E-91 | 7.4601E-90 |
| ENSG00000172932\|ANKRD13D | 4273.1705 | 2.5150 | ##### | 20.2730 | 2.2249E-91 | 8.1231E-90 |

| | | | | | | |
|---|---|---|---|---|---|---|
| ENSG00000120899\|PTK2B | 1152.0336 | -2.7705 | ##### | ####### | 2.3047E-91 | 8.4043E-90 |
| ENSG00000101955\|SRPX | 389.0870 | 4.2574 | ##### | 20.2578 | 3.0303E-91 | 1.1036E-89 |
| ENSG00000173320\|STOX2 | 705.4763 | -3.0199 | ##### | ####### | 3.0907E-91 | 1.1243E-89 |
| ENSG00000155961\|RAB39B | 318.2408 | 4.2109 | ##### | 20.2411 | 4.2563E-91 | 1.5464E-89 |
| ENSG00000164116\|GUCY1A1 | 465.6486 | 3.2492 | ##### | 20.2342 | 4.8941E-91 | 1.7760E-89 |
| ENSG00000109819\|PPARGC1A | 590.4338 | -4.9533 | ##### | ####### | 5.1927E-91 | 1.8820E-89 |
| ENSG00000198715\|GLMP | 1412.5889 | -2.8131 | ##### | ####### | 6.2189E-91 | 2.2508E-89 |
| ENSG00000054793\|ATP9A | 3022.9598 | -2.4213 | ##### | ####### | 6.2251E-91 | 2.2508E-89 |
| ENSG00000101144\|BMP7 | 1942.0632 | 9.3086 | ##### | 20.2157 | 7.1298E-91 | 2.5747E-89 |
| ENSG00000119138\|KLF9 | 602.5127 | -2.8926 | ##### | ####### | 1.3008E-90 | 4.6918E-89 |
| ENSG00000129595\|EPB41L4A | 926.8405 | -3.2484 | ##### | ####### | 1.4523E-90 | 5.2320E-89 |
| ENSG00000160190\|SLC37A1 | 799.7590 | -3.1767 | ##### | ####### | 2.2059E-90 | 7.9373E-89 |
| ENSG00000106031\|HOXA13 | 756.7681 | -3.3239 | ##### | ####### | 2.9239E-90 | 1.0508E-88 |
| ENSG00000173898\|SPTBN2 | 2246.2375 | -2.5404 | ##### | ####### | 3.6046E-90 | 1.2939E-88 |
| ENSG00000104643\|MTMR9 | 7380.5558 | -2.3853 | ##### | ####### | 3.6832E-90 | 1.3205E-88 |
| ENSG00000291109\|CES1P1 | 2168.8633 | -9.5423 | ##### | ####### | 4.1422E-90 | 1.4833E-88 |
| ENSG00000233621\|ZC3H12A-DT | 579.2482 | -3.4964 | ##### | ####### | 4.4429E-90 | 1.5891E-88 |
| ENSG00000160801\|PTH1R | 254.7141 | -4.4125 | ##### | ####### | 4.7086E-90 | 1.6821E-88 |
| ENSG00000291133 | 983.6687 | 3.2648 | ##### | 20.1085 | 6.2235E-90 | 2.2206E-88 |
| ENSG00000213190\|MLLT11 | 4818.8026 | -2.5252 | ##### | ####### | 9.3951E-90 | 3.3483E-88 |
| ENSG00000163931\|TKT | 50789.1972 | -2.1220 | ##### | ####### | 1.2144E-89 | 4.3228E-88 |
| ENSG00000152495\|CAMK4 | 697.9733 | 4.3454 | ##### | 20.0621 | 1.5830E-89 | 5.6281E-88 |
| ENSG00000110888\|CAPRIN2 | 4527.3131 | 2.4985 | ##### | 20.0484 | 2.0844E-89 | 7.4019E-88 |
| ENSG00000144893\|MED12L | 336.0738 | 4.1208 | ##### | 20.0398 | 2.4770E-89 | 8.7856E-88 |
| ENSG00000132031\|MATN3 | 1345.4181 | -2.7113 | ##### | ####### | 3.4512E-89 | 1.2227E-87 |
| ENSG00000204876 | 780.0664 | 6.8513 | ##### | 20.0127 | 4.2680E-89 | 1.5102E-87 |
| ENSG00000293322 | 240.0568 | 5.0227 | ##### | 19.9989 | 5.6315E-89 | 1.9904E-87 |
| ENSG00000123358\|NR4A1 | 9468.7738 | 2.3959 | ##### | 19.9836 | 7.6515E-89 | 2.7011E-87 |
| ENSG00000144959\|NCEH1 | 8790.0663 | -2.6464 | ##### | ####### | 7.7759E-89 | 2.7418E-87 |
| ENSG00000154760\|SLFN13 | 1900.9899 | -2.6627 | ##### | ####### | 8.2549E-89 | 2.9072E-87 |
| ENSG00000280441 | 742.9800 | 3.2003 | ##### | 19.9797 | 8.2660E-89 | 2.9077E-87 |
| ENSG00000106789\|CORO2A | 672.9778 | -4.3859 | ##### | ####### | 9.1359E-89 | 3.2099E-87 |
| ENSG00000072832\|CRMP1 | 1591.6960 | 2.5767 | ##### | 19.9720 | 9.6477E-89 | 3.3858E-87 |
| ENSG00000182752\|PAPPA | 2433.2560 | -2.5129 | ##### | ####### | 1.3534E-88 | 4.7440E-87 |
| ENSG00000087510\|TFAP2C | 5646.4783 | 2.5101 | ##### | 19.9534 | 1.3994E-88 | 4.8995E-87 |
| ENSG00000040341\|STAU2 | 2757.1641 | -2.5005 | ##### | ####### | 1.4125E-88 | 4.9397E-87 |
| ENSG00000135476\|ESPL1 | 8240.3557 | 2.4284 | ##### | 19.9490 | 1.5300E-88 | 5.3444E-87 |
| ENSG00000103449\|SALL1 | 714.4498 | 3.5107 | ##### | 19.9435 | 1.7069E-88 | 5.9551E-87 |
| ENSG00000274070\|CASTOR2 | 1824.1142 | -2.6227 | ##### | ####### | 1.8137E-88 | 6.3204E-87 |
| ENSG00000163638\|ADAMTS9 | 991.3218 | -4.9680 | ##### | ####### | 2.6965E-88 | 9.3859E-87 |
| ENSG00000170017\|ALCAM | 8985.7871 | -2.4118 | ##### | ####### | 4.7594E-88 | 1.6547E-86 |
| ENSG00000106772\|PRUNE2 | 245.3328 | -4.4339 | ##### | ####### | 5.7961E-88 | 2.0128E-86 |
| ENSG00000115604\|IL18R1 | 448.8332 | -3.7890 | ##### | ####### | 7.5900E-88 | 2.6327E-86 |
| ENSG00000049239\|H6PD | 6151.5897 | -2.4529 | ##### | ####### | 8.9468E-88 | 3.0997E-86 |
| ENSG00000120093\|HOXB3 | 636.4164 | 3.4797 | ##### | 19.8556 | 9.8493E-88 | 3.4085E-86 |
| ENSG00000164056\|SPRY1 | 873.8769 | 3.1389 | ##### | 19.8432 | 1.2609E-87 | 4.3583E-86 |
| ENSG00000139737\|SLAIN1 | 316.6192 | 4.4230 | ##### | 19.8336 | 1.5266E-87 | 5.2707E-86 |
| ENSG00000163535\|SGO2 | 6277.0435 | 2.4820 | ##### | 19.8318 | 1.5832E-87 | 5.4599E-86 |
| ENSG00000142494\|SLC47A1 | 1619.6955 | -3.3111 | ##### | ####### | 1.6740E-87 | 5.7663E-86 |
| ENSG00000175161\|CADM2 | 365.9268 | -4.7919 | ##### | ####### | 1.7079E-87 | 5.8763E-86 |
| ENSG00000171552\|BCL2L1 | 8661.2951 | -2.5490 | ##### | ####### | 1.7320E-87 | 5.9525E-86 |
| ENSG00000138835\|RGS3 | 931.2552 | -2.7555 | ##### | ####### | 2.4311E-87 | 8.3453E-86 |
| ENSG00000178685\|PARP10 | 609.6517 | -2.9566 | ##### | ####### | 2.6858E-87 | 9.2092E-86 |
| ENSG00000137331\|IER3 | 27634.0606 | 2.3923 | ##### | 19.8034 | 2.7823E-87 | 9.5289E-86 |
| ENSG00000138685\|FGF2 | 8813.3206 | -2.4108 | ##### | ####### | 3.1941E-87 | 1.0927E-85 |
| ENSG00000205116\|TMEM88B | 376.0271 | 3.5898 | ##### | 19.7948 | 3.2988E-87 | 1.1272E-85 |
| ENSG00000197375\|SLC22A5 | 1335.6103 | -2.5931 | ##### | ####### | 4.1477E-87 | 1.4157E-85 |
| ENSG00000196632\|WNK3 | 490.6424 | 4.9909 | ##### | 19.7704 | 5.3505E-87 | 1.8241E-85 |
| ENSG00000167930\|FAM234A | 3878.3512 | -2.4675 | ##### | ####### | 6.8149E-87 | 2.3207E-85 |
| ENSG00000166473\|PKD1L2 | 199.0532 | -3.8821 | ##### | ####### | 8.7389E-87 | 2.9725E-85 |
| ENSG00000070371\|CLTCL1 | 1323.5982 | -2.6657 | ##### | ####### | 9.2974E-87 | 3.1589E-85 |
| ENSG00000187942\|LDLRAD2 | 695.6763 | -3.3805 | ##### | ####### | 9.7008E-87 | 3.2922E-85 |
| ENSG00000167861\|HID1 | 446.7669 | -3.0920 | ##### | ####### | 1.0200E-86 | 3.4578E-85 |

| | | | | | | |
|---|---|---|---|---|---|---|
| ENSG00000143294\|PRCC | 14393.7520 | 1.8145 | ##### | 19.7325 | 1.1343E-86 | 3.8407E-85 |
| ENSG00000155099\|PIP4P2 | 1633.6953 | -2.5793 | ##### | ####### | 1.2928E-86 | 4.3727E-85 |
| ENSG00000126603\|GLIS2 | 5572.3564 | -1.3786 | ##### | ####### | 1.5270E-86 | 5.1588E-85 |
| ENSG00000112183\|RBM24 | 2940.1084 | 2.5603 | ##### | 19.7167 | 1.5488E-86 | 5.2267E-85 |
| ENSG00000139278\|GLIPR1 | 4398.3340 | -3.8384 | ##### | ####### | 1.8472E-86 | 6.2265E-85 |
| ENSG00000105889\|STEAP1B | 364.3004 | 4.1138 | ##### | 19.7028 | 2.0416E-86 | 6.8740E-85 |
| ENSG00000154328\|NEIL2 | 6111.7334 | -1.8948 | ##### | ####### | 3.4860E-86 | 1.1724E-84 |
| ENSG00000187678\|SPRY4 | 7277.8185 | 2.6695 | ##### | 19.6740 | 3.6044E-86 | 1.2109E-84 |
| ENSG00000156515\|HK1 | 13634.8380 | -2.4425 | ##### | ####### | 4.6113E-86 | 1.5474E-84 |
| ENSG00000126016\|AMOT | 534.9001 | 3.0630 | ##### | 19.6331 | 8.0642E-86 | 2.7030E-84 |
| ENSG00000060138\|YBX3 | 43264.1235 | 2.3391 | ##### | 19.6306 | 8.4654E-86 | 2.8343E-84 |
| ENSG00000116833\|NR5A2 | 2400.8402 | -2.4363 | ##### | ####### | 9.0155E-86 | 3.0151E-84 |
| ENSG00000172531\|PPP1CA | 35329.9110 | 2.3385 | ##### | 19.6179 | 1.0884E-85 | 3.6358E-84 |
| ENSG00000095397\|WHRN | 2623.2854 | 2.5760 | ##### | 19.6169 | 1.1090E-85 | 3.7005E-84 |
| ENSG00000108771\|DHX58 | 210.6498 | -4.2948 | ##### | ####### | 1.5793E-85 | 5.2641E-84 |
| ENSG00000276043\|UHRF1 | 24034.0152 | 2.2786 | ##### | 19.5956 | 1.6862E-85 | 5.6141E-84 |
| ENSG00000105835\|NAMPT | 26824.3654 | -2.3548 | ##### | ####### | 1.8046E-85 | 6.0016E-84 |
| ENSG00000235501\|CNN3-DT | 556.5811 | -3.9789 | ##### | ####### | 2.0271E-85 | 6.7341E-84 |
| ENSG00000175063\|UBE2C | 11845.5935 | 2.5047 | ##### | 19.5859 | 2.0378E-85 | 6.7621E-84 |
| ENSG00000186197\|EDARADD | 290.9595 | 4.9550 | ##### | 19.5848 | 2.0824E-85 | 6.9025E-84 |
| ENSG00000113083\|LOX | 347.7510 | 3.7263 | ##### | 19.5744 | 2.5567E-85 | 8.4652E-84 |
| ENSG00000266714\|MYO15B | 873.3392 | -2.7107 | ##### | ####### | 2.8725E-85 | 9.5002E-84 |
| ENSG00000130600\|H19 | 905.5385 | -2.7760 | ##### | ####### | 3.7163E-85 | 1.2277E-83 |
| ENSG00000176890\|TYMS | 35797.9830 | 2.2789 | ##### | 19.5524 | 3.9342E-85 | 1.2983E-83 |
| ENSG00000158856\|DMTN | 957.9819 | -2.7263 | ##### | ####### | 4.9131E-85 | 1.6195E-83 |
| ENSG00000157617\|C2CD2 | 3341.3796 | -2.5701 | ##### | ####### | 5.2885E-85 | 1.7414E-83 |
| ENSG00000115361\|ACADL | 172.2458 | -6.5878 | ##### | ####### | 5.6315E-85 | 1.8523E-83 |
| ENSG00000103222\|ABCC1 | 11239.6555 | -2.3193 | ##### | ####### | 6.7171E-85 | 2.2069E-83 |
| ENSG00000166402\|TUB | 1233.8705 | 2.7261 | ##### | 19.5177 | 7.7603E-85 | 2.5468E-83 |
| ENSG00000116667\|C1orf21 | 428.2802 | -3.2024 | ##### | ####### | 1.0482E-84 | 3.4364E-83 |
| ENSG00000267056 | 11336.2598 | ####### | ##### | ####### | 1.7595E-84 | 5.7617E-83 |
| ENSG00000067113\|PLPP1 | 2145.4613 | -2.4611 | ##### | ####### | 2.0189E-84 | 6.6038E-83 |
| ENSG00000161714\|PLCD3 | 6293.7958 | -2.4014 | ##### | ####### | 2.5222E-84 | 8.2412E-83 |
| ENSG00000197857\|ZNF44 | 295.4944 | -3.6582 | ##### | ####### | 2.5947E-84 | 8.4689E-83 |
| ENSG00000041353\|RAB27B | 6818.5276 | -2.3198 | ##### | ####### | 4.0034E-84 | 1.3053E-82 |
| ENSG00000273729 | 666.9745 | 2.9493 | ##### | 19.4309 | 4.2273E-84 | 1.3768E-82 |
| ENSG00000010292\|NCAPD2 | 38010.6559 | 2.2428 | ##### | 19.3899 | 9.3952E-84 | 3.0565E-82 |
| ENSG00000174233\|ADCY6 | 6100.2942 | -2.3286 | ##### | ####### | 1.1869E-83 | 3.8573E-82 |
| ENSG00000164823\|OSGIN2 | 12230.5842 | -2.4102 | ##### | ####### | 1.3028E-83 | 4.2291E-82 |
| ENSG00000184985\|SORCS2 | 355.6799 | -4.1289 | ##### | ####### | 1.7360E-83 | 5.6293E-82 |
| ENSG00000175265\|GOLGA8A | 3185.5130 | 2.4963 | ##### | 19.3530 | 1.9245E-83 | 6.2338E-82 |
| ENSG00000206341\|HLA-H | 5150.1295 | 2.4728 | ##### | 19.3474 | 2.1430E-83 | 6.9341E-82 |
| ENSG00000006756\|ARSD | 1790.9796 | -2.6331 | ##### | ####### | 2.2733E-83 | 7.3477E-82 |
| ENSG00000147889\|CDKN2A | 2862.2599 | 2.4679 | ##### | 19.3386 | 2.5430E-83 | 8.2104E-82 |
| ENSG00000186523\|FAM86B1 | 2361.5012 | -2.4583 | ##### | ####### | 2.7298E-83 | 8.8041E-82 |
| ENSG00000197580\|BCO2 | 403.8862 | -3.6365 | ##### | ####### | 2.7603E-83 | 8.8927E-82 |
| ENSG00000198157\|HMGN5 | 356.1097 | -3.5543 | ##### | ####### | 3.4061E-83 | 1.0961E-81 |
| ENSG00000163131\|CTSS | 799.1739 | -2.7929 | ##### | ####### | 3.9178E-83 | 1.2581E-81 |
| ENSG00000069482\|GAL | 1978.7179 | 2.6389 | ##### | 19.3163 | 3.9178E-83 | 1.2581E-81 |
| ENSG00000187498\|COL4A1 | 1720.8643 | -2.5352 | ##### | ####### | 5.4415E-83 | 1.7456E-81 |
| ENSG00000169964\|TMEM42 | 352.7873 | -4.4255 | ##### | ####### | 6.0330E-83 | 1.9332E-81 |
| ENSG00000112320\|SOBP | 250.3038 | 3.8057 | ##### | 19.2938 | 6.0527E-83 | 1.9375E-81 |
| ENSG00000112343\|TRIM38 | 426.0611 | -5.0449 | ##### | ####### | 6.1794E-83 | 1.9759E-81 |
| ENSG00000136944\|LMX1B | 337.2459 | 3.8199 | ##### | 19.2913 | 6.3558E-83 | 2.0301E-81 |
| ENSG00000105379\|ETFB | 9463.2654 | -2.2490 | ##### | ####### | 6.5010E-83 | 2.0743E-81 |
| ENSG00000225138\|SLC9A3-AS1 | 570.9911 | -2.6374 | ##### | ####### | 6.7343E-83 | 2.1464E-81 |
| ENSG00000155093\|PTPRN2 | 2429.9458 | -2.4147 | ##### | ####### | 1.0106E-82 | 3.2178E-81 |
| ENSG00000179046\|TRIML2 | 303.7298 | 5.7233 | ##### | 19.2664 | 1.0280E-82 | 3.2695E-81 |
| ENSG00000103021\|CCDC113 | 1128.1499 | -2.7975 | ##### | ####### | 1.2341E-82 | 3.9209E-81 |
| ENSG00000160282\|FTCD | 454.1705 | 3.2404 | ##### | 19.2556 | 1.2680E-82 | 4.0244E-81 |
| ENSG00000128641\|MYO1B | 14891.4796 | -2.2872 | ##### | ####### | 1.4724E-82 | 4.6682E-81 |
| ENSG00000143674\|MAP3K21 | 1704.1988 | -2.5178 | ##### | ####### | 1.5681E-82 | 4.9663E-81 |
| ENSG00000104549\|SQLE | 8966.7475 | -1.2605 | ##### | ####### | 2.3047E-82 | 7.2913E-81 |

| | | | | | | |
|---|---|---|---|---|---|---|
| ENSG00000131626\|PPFIA1 | 16171.1807 | 1.9505 | ##### | 19.2143 | 2.8078E-82 | 8.8734E-81 |
| ENSG00000106236\|NPTX2 | 3590.4884 | 2.5288 | ##### | 19.2076 | 3.1964E-82 | 1.0091E-80 |
| ENSG00000243646\|IL10RB | 2367.2649 | -2.8336 | ##### | ####### | 3.2359E-82 | 1.0205E-80 |
| ENSG00000290387\|SORD2P | 1701.4717 | 2.5054 | ##### | 19.1964 | 3.9656E-82 | 1.2493E-80 |
| ENSG00000103528\|SYT17 | 932.4301 | -2.7017 | ##### | ####### | 4.6255E-82 | 1.4557E-80 |
| ENSG00000105655\|ISYNA1 | 9468.0862 | 2.3952 | ##### | 19.1838 | 5.0593E-82 | 1.5905E-80 |
| ENSG00000116191\|RALGPS2 | 2996.1000 | -2.4222 | ##### | ####### | 6.2996E-82 | 1.9783E-80 |
| ENSG00000047315\|POLR2B | 28461.1751 | 1.8761 | ##### | 19.1688 | 6.7507E-82 | 2.1177E-80 |
| ENSG00000107438\|PDLIM1 | 5202.3185 | -2.4433 | ##### | ####### | 7.7171E-82 | 2.4184E-80 |
| ENSG00000108821\|COL1A1 | 3387.2451 | 2.3921 | ##### | 19.1500 | 9.6708E-82 | 3.0275E-80 |
| ENSG00000221890\|NPTXR | 2828.3067 | -2.4442 | ##### | ####### | 1.3049E-81 | 4.0808E-80 |
| ENSG00000070882\|OSBPL3 | 2255.4354 | -2.3100 | ##### | ####### | 1.9277E-81 | 6.0222E-80 |
| ENSG00000113719\|ERGIC1 | 14551.3254 | -1.3069 | ##### | ####### | 2.6253E-81 | 8.1929E-80 |
| ENSG00000006025\|OSBPL7 | 329.5556 | -6.1591 | ##### | ####### | 4.1761E-81 | 1.3019E-79 |
| ENSG00000138311\|ZNF365 | 394.7835 | 3.2446 | ##### | 19.0679 | 4.6658E-81 | 1.4530E-79 |
| ENSG00000110675\|ELMOD1 | 226.3014 | 5.3573 | ##### | 19.0588 | 5.5548E-81 | 1.7281E-79 |
| ENSG00000137767\|SQOR | 4348.8439 | -2.5709 | ##### | ####### | 5.9391E-81 | 1.8449E-79 |
| ENSG00000112208\|BAG2 | 4848.2979 | -2.5288 | ##### | ####### | 5.9427E-81 | 1.8449E-79 |
| ENSG00000286037 | 2771.8055 | -3.2225 | ##### | ####### | 6.0412E-81 | 1.8735E-79 |
| ENSG00000250056\|LINC01018 | 237.0394 | 4.0529 | ##### | 19.0377 | 8.3041E-81 | 2.5726E-79 |
| ENSG00000213853\|EMP2 | 7374.6757 | -2.3735 | ##### | ####### | 8.6046E-81 | 2.6630E-79 |
| ENSG00000114378\|HYAL1 | 169.1062 | -5.0514 | ##### | ####### | 1.0249E-80 | 3.1687E-79 |
| ENSG00000104324\|CPQ | 461.7457 | -3.3194 | ##### | ####### | 1.0304E-80 | 3.1822E-79 |
| ENSG00000171848\|RRM2 | 28713.4923 | 1.5501 | ##### | 19.0164 | 1.2470E-80 | 3.8472E-79 |
| ENSG00000168487\|BMP1 | 2444.8600 | -2.3820 | ##### | ####### | 1.2592E-80 | 3.8809E-79 |
| ENSG00000129654\|FOXJ1 | 1172.5416 | 2.9222 | ##### | 19.0128 | 1.3374E-80 | 4.1177E-79 |
| ENSG00000116396\|KCNC4 | 4452.6980 | 2.4272 | ##### | 19.0068 | 1.4984E-80 | 4.6086E-79 |
| ENSG00000166670\|MMP10 | 336.2724 | 3.8453 | ##### | 18.9907 | 2.0377E-80 | 6.2609E-79 |
| ENSG00000146670\|CDCA5 | 6965.1499 | 1.3455 | ##### | 18.9879 | 2.1476E-80 | 6.5918E-79 |
| ENSG00000163235\|TGFA | 682.0810 | -2.7563 | ##### | ####### | 2.7429E-80 | 8.4103E-79 |
| ENSG00000169752\|NRG4 | 263.4107 | -5.5657 | ##### | ####### | 2.9894E-80 | 9.1567E-79 |
| ENSG00000196263\|ZNF471 | 385.6938 | -4.1796 | ##### | ####### | 3.9593E-80 | 1.2115E-78 |
| ENSG00000134470\|IL15RA | 401.8179 | -4.8775 | ##### | ####### | 4.1760E-80 | 1.2765E-78 |
| ENSG00000163485\|ADORA1 | 473.9754 | 4.2150 | ##### | 18.9350 | 5.8677E-80 | 1.7918E-78 |
| ENSG00000272070 | 164.7676 | -5.6445 | ##### | ####### | 6.2867E-80 | 1.9178E-78 |
| ENSG00000153395\|LPCAT1 | 6588.8181 | -2.2473 | ##### | ####### | 6.7099E-80 | 2.0448E-78 |
| ENSG00000051620\|HEBP2 | 2086.7501 | -2.3286 | ##### | ####### | 6.8892E-80 | 2.0973E-78 |
| ENSG00000087077\|TRIP6 | 394.4632 | -3.2509 | ##### | ####### | 7.4343E-80 | 2.2609E-78 |
| ENSG00000110075\|PPP6R3 | 18827.5847 | 2.3036 | ##### | 18.9189 | 7.9626E-80 | 2.4191E-78 |
| ENSG00000078401\|EDN1 | 753.8480 | -2.9041 | ##### | ####### | 1.1610E-79 | 3.5236E-78 |
| ENSG00000085871\|MGST2 | 7726.2660 | ####### | ##### | ####### | 1.1854E-79 | 3.5942E-78 |
| ENSG00000007384\|RHBDF1 | 1589.9860 | -2.4616 | ##### | ####### | 1.5171E-79 | 4.5952E-78 |
| ENSG00000168785\|TSPAN5 | 1093.7106 | 2.7696 | ##### | 18.8826 | 1.5872E-79 | 4.8027E-78 |
| ENSG00000166816\|LDHD | 316.3060 | -3.1910 | ##### | ####### | 1.8222E-79 | 5.5080E-78 |
| ENSG00000008513\|ST3GAL1 | 3086.2555 | -2.3415 | ##### | ####### | 2.0709E-79 | 6.2536E-78 |
| ENSG00000173273\|TNKS | 9280.9488 | -2.2606 | ##### | ####### | 2.1200E-79 | 6.3953E-78 |
| ENSG00000196526\|AFAP1 | 5432.9177 | -2.2104 | ##### | ####### | 2.1377E-79 | 6.4422E-78 |
| ENSG00000073008\|PVR | 14543.9594 | -2.2052 | ##### | ####### | 2.2988E-79 | 6.9208E-78 |
| ENSG00000158769\|F11R | 7359.3270 | -2.3104 | ##### | ####### | 2.6178E-79 | 7.8732E-78 |
| ENSG00000133030\|MPRIP | 21900.1195 | -2.2007 | ##### | ####### | 2.9708E-79 | 8.9259E-78 |
| ENSG00000004478\|FKBP4 | 74406.0015 | 1.3801 | ##### | 18.8401 | 3.5419E-79 | 1.0631E-77 |
| ENSG00000245275\|SAP30L-AS1 | 371.9413 | -3.2858 | ##### | ####### | 7.5764E-79 | 2.2718E-77 |
| ENSG00000149212\|SESN3 | 753.6143 | -2.7621 | ##### | ####### | 8.3412E-79 | 2.4986E-77 |
| ENSG00000135956\|TMEM127 | 8735.6286 | 1.5055 | ##### | 18.7660 | 1.4338E-78 | 4.2882E-77 |
| ENSG00000169439\|SDC2 | 1473.0042 | -3.0188 | ##### | ####### | 1.4344E-78 | 4.2882E-77 |
| ENSG00000006704\|GTF2IRD1 | 2212.7610 | -2.5623 | ##### | ####### | 1.4729E-78 | 4.3987E-77 |
| ENSG00000162552\|WNT4 | 131.8358 | -4.7513 | ##### | ####### | 2.3932E-78 | 7.1400E-77 |
| ENSG00000112983\|BRD8 | 6671.9123 | 1.9641 | ##### | 18.7216 | 3.3007E-78 | 9.8380E-77 |
| ENSG00000117226\|GBP3 | 248.1601 | -3.8627 | ##### | ####### | 4.7803E-78 | 1.4234E-76 |
| ENSG00000161298\|ZNF382 | 227.7114 | -5.2044 | ##### | ####### | 5.0362E-78 | 1.4981E-76 |
| ENSG00000108947\|EFNB3 | 184.9358 | 4.9658 | ##### | 18.6915 | 5.8059E-78 | 1.7253E-76 |
| ENSG00000256235\|SMIM3 | 682.1084 | 3.2230 | ##### | 18.6758 | 7.7876E-78 | 2.3119E-76 |
| ENSG00000133466\|C1QTNF6 | 2222.9329 | -2.3642 | ##### | ####### | 1.1442E-77 | 3.3934E-76 |

| | | | | | | |
|---|---|---|---|---|---|---|
| ENSG00000177181\|RIMKLA | 794.6963 | -2.5727 | ##### | ####### | 1.1880E-77 | 3.5197E-76 |
| ENSG00000186871\|ERCC6L | 4065.7293 | 2.2019 | ##### | 18.6453 | 1.3792E-77 | 4.0824E-76 |
| ENSG00000137628\|DDX60 | 571.8546 | -2.8729 | ##### | ####### | 1.6578E-77 | 4.9020E-76 |
| ENSG00000171621\|SPSB1 | 2335.4313 | -2.4917 | ##### | ####### | 1.8902E-77 | 5.5836E-76 |
| ENSG00000160285\|LSS | 4894.3288 | -2.3232 | ##### | ####### | 2.0166E-77 | 5.9510E-76 |
| ENSG00000085719\|CPNE3 | 6104.9587 | -2.2966 | ##### | ####### | 3.2687E-77 | 9.6367E-76 |
| ENSG00000152784\|PRDM8 | 678.0106 | 2.6255 | ##### | 18.5895 | 3.9075E-77 | 1.1509E-75 |
| ENSG00000127585\|FBXL16 | 1388.9772 | -2.6298 | ##### | ####### | 3.9516E-77 | 1.1627E-75 |
| ENSG00000187123\|LYPD6 | 381.9609 | 4.4113 | ##### | 18.5856 | 4.2063E-77 | 1.2364E-75 |
| ENSG00000177459\|ERICH5 | 306.2549 | -8.2509 | ##### | ####### | 7.2766E-77 | 2.1368E-75 |
| ENSG00000241945\|PWP2 | 5064.4567 | 2.4192 | ##### | 18.5531 | 7.6950E-77 | 2.2575E-75 |
| ENSG00000291003 | 1609.7500 | 2.6975 | ##### | 18.5420 | 9.4691E-77 | 2.7752E-75 |
| ENSG00000105559\|PLEKHA4 | 527.6650 | -3.3713 | ##### | ####### | 1.0304E-76 | 3.0169E-75 |
| ENSG00000171522\|PTGER4 | 1803.1245 | 2.4820 | ##### | 18.5367 | 1.0441E-76 | 3.0541E-75 |
| ENSG00000113569\|NUP155 | 20872.5791 | 2.1147 | ##### | 18.5193 | 1.4442E-76 | 4.2202E-75 |
| ENSG00000130294\|KIF1A | 15533.2329 | 2.2102 | ##### | 18.5123 | 1.6419E-76 | 4.7932E-75 |
| ENSG00000112242\|E2F3 | 6824.1711 | 2.2079 | ##### | 18.5096 | 1.7282E-76 | 5.0402E-75 |
| ENSG00000214193\|SH3D21 | 2067.4515 | 2.6841 | ##### | 18.5051 | 1.8799E-76 | 5.4773E-75 |
| ENSG00000178726\|THBD | 717.6061 | -2.9647 | ##### | ####### | 2.5057E-76 | 7.2938E-75 |
| ENSG00000103710\|RASL12 | 206.6089 | -4.4617 | ##### | ####### | 2.5923E-76 | 7.5383E-75 |
| ENSG00000115041\|KCNIP3 | 250.4400 | 3.6125 | ##### | 18.4871 | 2.6233E-76 | 7.6211E-75 |
| ENSG00000169583\|CLIC3 | 380.6112 | -4.1726 | ##### | ####### | 4.1076E-76 | 1.1922E-74 |
| ENSG00000079150\|FKBP7 | 393.6008 | -3.7271 | ##### | ####### | 4.3963E-76 | 1.2747E-74 |
| ENSG00000095539\|SEMA4G | 907.8235 | -2.6724 | ##### | ####### | 5.3274E-76 | 1.5432E-74 |
| ENSG00000022567\|SLC45A4 | 4863.5578 | -2.3091 | ##### | ####### | 5.5914E-76 | 1.6181E-74 |
| ENSG00000157388\|CACNA1D | 283.3687 | -3.2964 | ##### | ####### | 5.9177E-76 | 1.7109E-74 |
| ENSG00000086062\|B4GALT1 | 13366.8619 | -2.1798 | ##### | ####### | 5.9263E-76 | 1.7117E-74 |
| ENSG00000213949\|ITGA1 | 1761.4542 | 2.2968 | ##### | 18.4327 | 7.1744E-76 | 2.0702E-74 |
| ENSG00000137135\|ARHGEF39 | 5076.0418 | 2.4889 | ##### | 18.4194 | 9.1779E-76 | 2.6457E-74 |
| ENSG00000204262\|COL5A2 | 35101.7598 | -2.1516 | ##### | ####### | 1.6608E-75 | 4.7829E-74 |
| ENSG00000100462\|PRMT5 | 24367.3640 | 1.2330 | ##### | 18.3855 | 1.7172E-75 | 4.9407E-74 |
| ENSG00000140950\|MEAK7 | 4349.9940 | -2.4735 | ##### | ####### | 1.9472E-75 | 5.5971E-74 |
| ENSG00000189337\|KAZN | 1106.9017 | 2.9362 | ##### | 18.3776 | 1.9856E-75 | 5.7018E-74 |
| ENSG00000104231\|ZFAND1 | 2255.7767 | -2.4347 | ##### | ####### | 2.0701E-75 | 5.9388E-74 |
| ENSG00000172340\|SUCLG2 | 3677.7963 | -2.3396 | ##### | ####### | 2.4644E-75 | 7.0632E-74 |
| ENSG00000118898\|PPL | 1121.5597 | -2.5843 | ##### | ####### | 2.5861E-75 | 7.4049E-74 |
| ENSG00000117036\|ETV3 | 7452.2753 | 2.2725 | ##### | 18.3632 | 2.5896E-75 | 7.4078E-74 |
| ENSG00000260807\|CEROX1 | 192.2809 | 6.1574 | ##### | 18.3609 | 2.7025E-75 | 7.7234E-74 |
| ENSG00000169894\|MUC3A | 1978.9197 | -2.4751 | ##### | ####### | 3.0341E-75 | 8.6628E-74 |
| ENSG00000256771\|ZNF253 | 235.3351 | -5.1006 | ##### | ####### | 3.0827E-75 | 8.7932E-74 |
| ENSG00000085999\|RAD54L | 3567.3420 | 1.5705 | ##### | 18.3487 | 3.3798E-75 | 9.6316E-74 |
| ENSG00000196092\|PAX5 | 228.2043 | 6.1974 | ##### | 18.3404 | 3.9417E-75 | 1.1222E-73 |
| ENSG00000105246\|EBI3 | 546.4982 | 3.1748 | ##### | 18.3379 | 4.1220E-75 | 1.1724E-73 |
| ENSG00000154146\|NRGN | 823.6495 | 2.8688 | ##### | 18.3324 | 4.5670E-75 | 1.2977E-73 |
| ENSG00000206262\|FOXL2NB | 302.4006 | 7.6051 | ##### | 18.3230 | 5.4212E-75 | 1.5390E-73 |
| ENSG00000155254\|MARVELD1 | 9482.2901 | -1.5328 | ##### | ####### | 5.5409E-75 | 1.5715E-73 |
| ENSG00000164010\|ERMAP | 1569.1413 | -2.4259 | ##### | ####### | 6.0528E-75 | 1.7150E-73 |
| ENSG00000146530\|VWDE | 887.1747 | -2.5534 | ##### | ####### | 7.7445E-75 | 2.1923E-73 |
| ENSG00000266302 | 661.3998 | ####### | ##### | ####### | 8.0994E-75 | 2.2906E-73 |
| ENSG00000174808\|BTC | 494.3703 | -3.2008 | ##### | ####### | 9.3351E-75 | 2.6376E-73 |
| ENSG00000132970\|WASF3 | 1503.0583 | 2.5886 | ##### | 18.2907 | 9.8222E-75 | 2.7726E-73 |
| ENSG00000126970\|ZC4H2 | 6007.1788 | ####### | ##### | ####### | 2.1210E-74 | 5.9813E-73 |
| ENSG00000153832\|FBXO36 | 331.6529 | -2.9952 | ##### | ####### | 2.2617E-74 | 6.3723E-73 |
| ENSG00000112715\|VEGFA | 4773.8442 | -1.7723 | ##### | ####### | 2.9147E-74 | 8.2043E-73 |
| ENSG00000101193\|GID8 | 5271.4909 | 2.1076 | ##### | 18.2309 | 2.9371E-74 | 8.2596E-73 |
| ENSG00000106689\|LHX2 | 1308.6833 | 2.2381 | ##### | 18.2301 | 2.9754E-74 | 8.3595E-73 |
| ENSG00000267432\|DNAH17-AS1 | 702.8342 | 2.6307 | ##### | 18.2267 | 3.1704E-74 | 8.8989E-73 |
| ENSG00000166897\|ELFN2 | 1189.1490 | -2.5687 | ##### | ####### | 3.7346E-74 | 1.0473E-72 |
| ENSG00000218048 | 332.2337 | -3.0980 | ##### | ####### | 3.9825E-74 | 1.1157E-72 |
| ENSG00000115109\|EPB41L5 | 3118.8137 | -2.2264 | ##### | ####### | 4.9941E-74 | 1.3978E-72 |
| ENSG00000205808\|PLPP6 | 875.2894 | -2.7275 | ##### | ####### | 5.3484E-74 | 1.4956E-72 |
| ENSG00000160714\|UBE2Q1 | 14602.1020 | 1.5715 | ##### | 18.1757 | 8.0436E-74 | 2.2472E-72 |
| ENSG00000176623\|RMDN1 | 2559.2856 | -2.2279 | ##### | ####### | 8.3539E-74 | 2.3317E-72 |

| | | | | | | |
|---|---|---|---|---|---|---|
| ENSG00000070182\|SPTB | 488.6163 | 3.2073 | ##### | 18.1685 | 9.1643E-74 | 2.5555E-72 |
| ENSG00000170522\|ELOVL6 | 2667.5647 | -2.2458 | ##### | ####### | 1.0577E-73 | 2.9467E-72 |
| ENSG00000228709\|LINC02575 | 382.1452 | 5.3495 | ##### | 18.1529 | 1.2190E-73 | 3.3930E-72 |
| ENSG00000132676\|DAP3 | 20103.3037 | 1.4327 | ##### | 18.1521 | 1.2360E-73 | 3.4371E-72 |
| ENSG00000135709\|KIAA0513 | 1047.4449 | -2.4889 | ##### | ####### | 1.4575E-73 | 4.0492E-72 |
| ENSG00000112238\|PRDM13 | 448.9214 | 2.9048 | ##### | 18.1294 | 1.8682E-73 | 5.1854E-72 |
| ENSG00000130821\|SLC6A8 | 6853.0121 | 2.1462 | ##### | 18.1206 | 2.1917E-73 | 6.0775E-72 |
| ENSG00000124217\|MOCS3 | 2003.2280 | 1.2559 | ##### | 18.1117 | 2.5766E-73 | 7.1383E-72 |
| ENSG00000143384\|MCL1 | 36208.1257 | 1.4701 | ##### | 18.1116 | 2.5824E-73 | 7.1476E-72 |
| ENSG00000205277\|MUC12 | 168.3879 | -5.1254 | ##### | ####### | 2.8248E-73 | 7.8114E-72 |
| ENSG00000280179 | 215.6496 | -3.8239 | ##### | ####### | 2.8369E-73 | 7.8377E-72 |
| ENSG00000121068\|TBX2 | 445.8799 | -3.8590 | ##### | ####### | 2.9751E-73 | 8.2117E-72 |
| ENSG00000165617\|DACT1 | 246.6387 | 4.4153 | ##### | 18.0971 | 3.3575E-73 | 9.2587E-72 |
| ENSG00000164050\|PLXNB1 | 1666.2974 | -2.4755 | ##### | ####### | 4.2734E-73 | 1.1773E-71 |
| ENSG00000173432\|SAA1 | 276.1364 | -5.0294 | ##### | ####### | 4.5609E-73 | 1.2554E-71 |
| ENSG00000237172\|B3GNT9 | 902.2974 | -2.6096 | ##### | ####### | 4.9580E-73 | 1.3634E-71 |
| ENSG00000196569\|LAMA2 | 518.6182 | 3.3344 | ##### | 18.0669 | 5.8042E-73 | 1.5947E-71 |
| ENSG00000172201\|ID4 | 1048.2509 | 2.5423 | ##### | 18.0590 | 6.7082E-73 | 1.8414E-71 |
| ENSG00000276386\|CNTNAP3P2 | 241.9517 | -5.2205 | ##### | ####### | 6.8758E-73 | 1.8856E-71 |
| ENSG00000059804\|SLC2A3 | 9216.2040 | 2.2237 | ##### | 18.0533 | 7.4351E-73 | 2.0371E-71 |
| ENSG00000164039\|BDH2 | 523.4020 | -2.7236 | ##### | ####### | 7.5605E-73 | 2.0696E-71 |
| ENSG00000146555\|SDK1 | 396.7488 | 3.5500 | ##### | 18.0518 | 7.6370E-73 | 2.0886E-71 |
| ENSG00000132563\|REEP2 | 199.0280 | 4.3713 | ##### | 18.0469 | 8.3520E-73 | 2.2821E-71 |
| ENSG00000293515 | 309.5599 | -5.2285 | ##### | ####### | 1.4370E-72 | 3.9228E-71 |
| ENSG00000253731\|PCDHGA6 | 196.4150 | -5.3378 | ##### | ####### | 1.6460E-72 | 4.4893E-71 |
| ENSG00000243749\|TMEM35B | 1230.7953 | -2.5033 | ##### | ####### | 1.6705E-72 | 4.5519E-71 |
| ENSG00000162139\|NEU3 | 2411.0277 | 1.9260 | ##### | 18.0046 | 1.7929E-72 | 4.8810E-71 |
| ENSG00000104432\|IL7 | 140.0010 | -6.5750 | ##### | ####### | 2.1850E-72 | 5.9430E-71 |
| ENSG00000141458\|NPC1 | 14892.3860 | -2.1480 | ##### | ####### | 4.8780E-72 | 1.3250E-70 |
| ENSG00000136783\|NIPSNAP3A | 934.3480 | -2.5089 | ##### | ####### | 4.8803E-72 | 1.3250E-70 |
| ENSG00000034152\|MAP2K3 | 9291.9079 | 1.8925 | ##### | 17.9458 | 5.1774E-72 | 1.4044E-70 |
| ENSG00000049130\|KITLG | 3388.2638 | -2.4349 | ##### | ####### | 5.3072E-72 | 1.4383E-70 |
| ENSG00000233058\|ATP13A3-DT | 216.3928 | -4.4266 | ##### | ####### | 5.4236E-72 | 1.4685E-70 |
| ENSG00000108960\|MMD | 4138.1845 | -2.1563 | ##### | ####### | 5.5594E-72 | 1.5039E-70 |
| ENSG00000166920\|C15orf48 | 15727.0852 | 2.1456 | ##### | 17.9381 | 5.9495E-72 | 1.6080E-70 |
| ENSG00000170545\|SMAGP | 2222.9270 | -2.4359 | ##### | ####### | 6.4602E-72 | 1.7444E-70 |
| ENSG00000198952\|SMG5 | 25920.9587 | 2.1480 | ##### | 17.9222 | 7.9112E-72 | 2.1343E-70 |
| ENSG00000213020\|ZNF611 | 692.2723 | -2.6828 | ##### | ####### | 8.2335E-72 | 2.2192E-70 |
| ENSG00000006555\|TTC22 | 5034.7473 | ####### | ##### | ####### | 1.0258E-71 | 2.7625E-70 |
| ENSG00000125266\|EFNB2 | 816.9459 | -2.8578 | ##### | ####### | 1.0734E-71 | 2.8879E-70 |
| ENSG00000154548\|SRSF12 | 242.5312 | -4.3568 | ##### | ####### | 1.0889E-71 | 2.9270E-70 |
| ENSG00000196517\|SLC6A9 | 1560.7479 | -2.3790 | ##### | ####### | 1.4286E-71 | 3.8367E-70 |
| ENSG00000173156\|RHOD | 1036.0697 | -2.5255 | ##### | ####### | 1.4372E-71 | 3.8564E-70 |
| ENSG00000133943\|DGLUCY | 616.2381 | -3.0602 | ##### | ####### | 1.4439E-71 | 3.8709E-70 |
| ENSG00000115353\|TACR1 | 320.8419 | -3.7143 | ##### | ####### | 1.5951E-71 | 4.2726E-70 |
| ENSG00000153814\|JAZF1 | 1208.2627 | -2.3931 | ##### | ####### | 1.7995E-71 | 4.8155E-70 |
| ENSG00000107819\|SFXN3 | 2854.2653 | -2.3060 | ##### | ####### | 2.2735E-71 | 6.0785E-70 |
| ENSG00000084090\|STARD7 | 24094.8723 | 1.5581 | ##### | 17.8593 | 2.4463E-71 | 6.5348E-70 |
| ENSG00000272636\|DOC2B | 437.8088 | -3.8904 | ##### | ####### | 2.8665E-71 | 7.6504E-70 |
| ENSG00000166831\|RBPMS2 | 752.5724 | -2.6673 | ##### | ####### | 3.0328E-71 | 8.0871E-70 |
| ENSG00000166311\|SMPD1 | 929.9570 | -2.4977 | ##### | ####### | 3.9183E-71 | 1.0439E-69 |
| ENSG00000104450\|SPAG1 | 1536.9730 | -2.4299 | ##### | ####### | 4.1330E-71 | 1.1001E-69 |
| ENSG00000169122\|FAM110B | 574.7385 | 4.2488 | ##### | 17.8286 | 4.2396E-71 | 1.1275E-69 |
| ENSG00000163900\|TMEM41A | 2007.4773 | -2.2673 | ##### | ####### | 4.5069E-71 | 1.1975E-69 |
| ENSG00000066248\|NGEF | 1398.7964 | -2.3886 | ##### | ####### | 5.8052E-71 | 1.5411E-69 |
| ENSG00000134516\|DOCK2 | 281.2521 | 3.4463 | ##### | 17.8058 | 6.3749E-71 | 1.6908E-69 |
| ENSG00000188343\|CIBAR1 | 736.7454 | -2.3345 | ##### | ####### | 7.3786E-71 | 1.9553E-69 |
| ENSG00000109089\|CDR2L | 6140.3173 | -2.1984 | ##### | ####### | 8.0419E-71 | 2.1292E-69 |
| ENSG00000158246\|TENT5B | 582.2168 | -3.0336 | ##### | ####### | 9.4443E-71 | 2.4983E-69 |
| ENSG00000080493\|SLC4A4 | 817.7039 | -2.9466 | ##### | ####### | 9.6170E-71 | 2.5417E-69 |
| ENSG00000102265\|TIMP1 | 10630.3437 | 2.2318 | ##### | 17.7791 | 1.0258E-70 | 2.7087E-69 |
| ENSG00000069188\|SDK2 | 379.1750 | 4.3194 | ##### | 17.7782 | 1.0420E-70 | 2.7491E-69 |
| ENSG00000241322\|FBXW10B | 246.3649 | -3.8410 | ##### | ####### | 1.1474E-70 | 3.0244E-69 |

| | | | | | | |
|---|---|---|---|---|---|---|
| ENSG00000185361\|TNFAIP8L1 | 3679.7295 | 2.4538 | ##### | 17.7605 | 1.4305E-70 | 3.7674E-69 |
| ENSG00000162627\|SNX7 | 2788.4704 | -2.3304 | ##### | ####### | 1.4525E-70 | 3.8219E-69 |
| ENSG00000141576\|RNF157 | 1441.9045 | -2.5962 | ##### | ####### | 1.6948E-70 | 4.4555E-69 |
| ENSG00000107104\|KANK1 | 1955.4150 | -2.3760 | ##### | ####### | 1.8407E-70 | 4.8348E-69 |
| ENSG00000120705\|ETF1 | 24024.8390 | 1.3672 | ##### | 17.7430 | 1.9518E-70 | 5.1221E-69 |
| ENSG00000055813\|CCDC85A | 333.3846 | -4.2988 | ##### | ####### | 2.2010E-70 | 5.7710E-69 |
| ENSG00000182492\|BGN | 440.5591 | 5.8637 | ##### | 17.7288 | 2.5151E-70 | 6.5890E-69 |
| ENSG00000111639\|MRPL51 | 16993.5273 | 1.4571 | ##### | 17.7275 | 2.5735E-70 | 6.7360E-69 |
| ENSG00000106049\|HIBADH | 6059.2941 | -2.2116 | ##### | ####### | 3.3463E-70 | 8.7512E-69 |
| ENSG00000174125\|TLR1 | 321.4929 | -3.5620 | ##### | ####### | 4.1065E-70 | 1.0730E-68 |
| ENSG00000273760 | 245.6831 | -3.3180 | ##### | ####### | 4.4724E-70 | 1.1676E-68 |
| ENSG00000138162\|TACC2 | 5337.1564 | -2.0904 | ##### | ####### | 5.4556E-70 | 1.4230E-68 |
| ENSG00000176658\|MYO1D | 2040.0599 | -2.2121 | ##### | ####### | 6.4791E-70 | 1.6885E-68 |
| ENSG00000143153\|ATP1B1 | 7918.0012 | -2.2016 | ##### | ####### | 8.7147E-70 | 2.2691E-68 |
| ENSG00000116406\|EDEM3 | 4333.8109 | -2.1441 | ##### | ####### | 1.2836E-69 | 3.3393E-68 |
| ENSG00000067057\|PFKP | 37096.4641 | -1.2557 | ##### | ####### | 1.6036E-69 | 4.1654E-68 |
| ENSG00000109099\|PMP22 | 4717.3339 | -2.2488 | ##### | ####### | 1.6039E-69 | 4.1654E-68 |
| ENSG00000073849\|ST6GAL1 | 1063.0852 | -2.5114 | ##### | ####### | 2.0068E-69 | 5.2070E-68 |
| ENSG00000136052\|SLC41A2 | 2137.6530 | -2.2574 | ##### | ####### | 2.0435E-69 | 5.2978E-68 |
| ENSG00000131669\|NINJ1 | 1570.1335 | -2.3597 | ##### | ####### | 2.1846E-69 | 5.6586E-68 |
| ENSG00000100234\|TIMP3 | 4604.4892 | ####### | ##### | ####### | 2.2601E-69 | 5.8490E-68 |
| ENSG00000148411\|NACC2 | 6033.0509 | -2.1247 | ##### | ####### | 2.5435E-69 | 6.5768E-68 |
| ENSG00000198346\|ZNF813 | 694.1163 | -2.6710 | ##### | ####### | 3.5532E-69 | 9.1796E-68 |
| ENSG00000185404\|SP140L | 467.2591 | -6.1081 | ##### | ####### | 4.3917E-69 | 1.1336E-67 |
| ENSG00000164430\|CGAS | 6242.8306 | 2.0812 | ##### | 17.5483 | 6.1254E-69 | 1.5797E-67 |
| ENSG00000112319\|EYA4 | 1911.5489 | -2.2515 | ##### | ####### | 6.4118E-69 | 1.6522E-67 |
| ENSG00000280734\|LINC01232 | 189.3649 | -5.1192 | ##### | ####### | 6.8073E-69 | 1.7526E-67 |
| ENSG00000112149\|CD83 | 799.0484 | 3.0100 | ##### | 17.5415 | 6.9037E-69 | 1.7759E-67 |
| ENSG00000274021 | 553.7023 | 3.2136 | ##### | 17.5337 | 7.9257E-69 | 2.0370E-67 |
| ENSG00000166033\|HTRA1 | 2510.8249 | 2.2245 | ##### | 17.5241 | 9.3849E-69 | 2.4100E-67 |
| ENSG00000181031\|RPH3AL | 717.0821 | -2.7924 | ##### | ####### | 1.2042E-68 | 3.0896E-67 |
| ENSG00000103811\|CTSH | 2141.8148 | -2.5306 | ##### | ####### | 1.2272E-68 | 3.1460E-67 |
| ENSG00000198400\|NTRK1 | 231.0508 | 5.7587 | ##### | 17.5082 | 1.2408E-68 | 3.1780E-67 |
| ENSG00000073350\|LLGL2 | 2234.3239 | -2.3268 | ##### | ####### | 1.3064E-68 | 3.3432E-67 |
| ENSG00000164659\|ELAPOR2 | 582.9471 | -3.3397 | ##### | ####### | 1.5402E-68 | 3.9381E-67 |
| ENSG00000082781\|ITGB5 | 20464.9060 | -2.0875 | ##### | ####### | 1.6242E-68 | 4.1494E-67 |
| ENSG00000143624\|INTS3 | 18589.0836 | 2.0910 | ##### | 17.4752 | 2.2131E-68 | 5.6491E-67 |
| ENSG00000145012\|LPP | 3876.9597 | -2.1216 | ##### | ####### | 2.3347E-68 | 5.9544E-67 |
| ENSG00000196511\|TPK1 | 196.4395 | -4.4634 | ##### | ####### | 2.3664E-68 | 6.0302E-67 |
| ENSG00000164949\|GEM | 3019.7454 | -2.1449 | ##### | ####### | 2.4699E-68 | 6.2883E-67 |
| ENSG00000165821\|SALL2 | 575.7128 | 3.3803 | ##### | 17.4563 | 3.0833E-68 | 7.8434E-67 |
| ENSG00000173599\|PC | 2639.7052 | -2.2082 | ##### | ####### | 3.5114E-68 | 8.9249E-67 |
| ENSG00000067066\|SP100 | 1205.6307 | -7.6287 | ##### | ####### | 3.6695E-68 | 9.3188E-67 |
| ENSG00000290700 | 185.6134 | 3.7360 | ##### | 17.4421 | 3.9556E-68 | 1.0037E-66 |
| ENSG00000133107\|TRPC4 | 539.8387 | 2.8471 | ##### | 17.4402 | 4.0883E-68 | 1.0365E-66 |
| ENSG00000075275\|CELSR1 | 6398.4756 | -2.1672 | ##### | ####### | 4.7264E-68 | 1.1972E-66 |
| ENSG00000170445\|HARS1 | 10031.6055 | 1.3672 | ##### | 17.4210 | 5.7145E-68 | 1.4463E-66 |
| ENSG00000162976\|SLC66A3 | 1370.6509 | -2.3522 | ##### | ####### | 7.1010E-68 | 1.7957E-66 |
| ENSG00000188385\|JAKMIP3 | 383.9172 | -3.2175 | ##### | ####### | 7.2351E-68 | 1.8281E-66 |
| ENSG00000213563\|C8orf82 | 1820.2936 | -2.1593 | ##### | ####### | 8.7864E-68 | 2.2181E-66 |
| ENSG00000137393\|RNF144B | 149.1849 | -4.6864 | ##### | ####### | 8.9253E-68 | 2.2513E-66 |
| ENSG00000126787\|DLGAP5 | 8929.0391 | 2.1165 | ##### | 17.3861 | 1.0518E-67 | 2.6509E-66 |
| ENSG00000052126\|PLEKHA5 | 1946.0149 | 2.1129 | ##### | 17.3673 | 1.4593E-67 | 3.6746E-66 |
| ENSG00000138678\|GPAT3 | 3300.1236 | -2.1424 | ##### | ####### | 1.6301E-67 | 4.1015E-66 |
| ENSG00000230615 | 162.6867 | -4.2851 | ##### | ####### | 1.6522E-67 | 4.1535E-66 |
| ENSG00000257913\|DDN-AS1 | 512.8343 | 2.9504 | ##### | 17.3601 | 1.6544E-67 | 4.1555E-66 |
| ENSG00000169991\|IFFO2 | 2418.5318 | -2.1830 | ##### | ####### | 1.6986E-67 | 4.2629E-66 |
| ENSG00000213047\|DENND1B | 724.5336 | -2.8744 | ##### | ####### | 2.0294E-67 | 5.0888E-66 |
| ENSG00000131459\|GFPT2 | 753.5188 | -3.1518 | ##### | ####### | 2.2213E-67 | 5.5653E-66 |
| ENSG00000154359\|LONRF1 | 2290.4222 | -2.1512 | ##### | ####### | 2.4013E-67 | 6.0113E-66 |
| ENSG00000183117\|CSMD1 | 303.0360 | -3.8152 | ##### | ####### | 2.6880E-67 | 6.7233E-66 |
| ENSG00000125170\|DOK4 | 1143.6953 | -2.3626 | ##### | ####### | 2.7555E-67 | 6.8865E-66 |
| ENSG00000172922\|RNASEH2C | 3882.5637 | 1.7874 | ##### | 17.3281 | 2.8882E-67 | 7.2121E-66 |

| | | | | | | |
|---|---|---|---|---|---|---|
| ENSG00000258376\|PAPLN-AS1 | 406.0526 | 5.8920 | ##### | 17.3189 | 3.3892E-67 | 8.4561E-66 |
| ENSG00000117500\|TMED5 | 10222.9435 | -1.2098 | ##### | ####### | 3.6557E-67 | 9.1133E-66 |
| ENSG00000185112\|FAM43A | 605.7388 | 3.2009 | ##### | 17.3112 | 3.8749E-67 | 9.6517E-66 |
| ENSG00000144028\|SNRNP200 | 57280.9782 | 2.0284 | ##### | 17.3060 | 4.2362E-67 | 1.0543E-65 |
| ENSG00000180376\|CCDC66 | 1032.5832 | 2.2005 | ##### | 17.3025 | 4.5048E-67 | 1.1202E-65 |
| ENSG00000143258\|USP21 | 5186.9267 | 1.8749 | ##### | 17.2987 | 4.8104E-67 | 1.1952E-65 |
| ENSG00000176463\|SLCO3A1 | 973.5981 | 2.5991 | ##### | 17.2965 | 4.9943E-67 | 1.2398E-65 |
| ENSG00000008086\|CDKL5 | 712.0389 | -3.1691 | ##### | ####### | 5.0630E-67 | 1.2559E-65 |
| ENSG00000111331\|OAS3 | 3769.8203 | -2.1275 | ##### | ####### | 5.3142E-67 | 1.3171E-65 |
| ENSG00000157557\|ETS2 | 2927.5780 | -2.1824 | ##### | ####### | 5.7900E-67 | 1.4338E-65 |
| ENSG00000115129\|TP53I3 | 3722.4013 | -2.2049 | ##### | ####### | 7.6316E-67 | 1.8883E-65 |
| ENSG00000109861\|CTSC | 7277.2255 | -2.1536 | ##### | ####### | 7.6753E-67 | 1.8975E-65 |
| ENSG00000069974\|RAB27A | 996.5047 | -2.5599 | ##### | ####### | 8.9243E-67 | 2.2045E-65 |
| ENSG00000166592\|RRAD | 285.4401 | -3.0361 | ##### | ####### | 9.7815E-67 | 2.4142E-65 |
| ENSG00000100714\|MTHFD1 | 31148.6557 | 1.2408 | ##### | 17.2435 | 1.2516E-66 | 3.0865E-65 |
| ENSG00000072422\|RHOBTB1 | 1497.1861 | -2.3854 | ##### | ####### | 1.4614E-66 | 3.6009E-65 |
| ENSG00000162616\|DNAJB4 | 4290.2068 | -2.0932 | ##### | ####### | 1.4815E-66 | 3.6475E-65 |
| ENSG00000153132\|CLGN | 615.0509 | -2.5816 | ##### | ####### | 1.7100E-66 | 4.2066E-65 |
| ENSG00000069424\|KCNAB2 | 1943.0413 | -2.2514 | ##### | ####### | 3.0052E-66 | 7.3870E-65 |
| ENSG00000164867\|NOS3 | 1575.2303 | 7.4885 | ##### | 17.1889 | 3.2177E-66 | 7.9028E-65 |
| ENSG00000025434\|NR1H3 | 452.9612 | -2.5642 | ##### | ####### | 3.4292E-66 | 8.4153E-65 |
| ENSG00000137821\|LRRC49 | 1035.2465 | 2.3554 | ##### | 17.1756 | 4.0465E-66 | 9.9219E-65 |
| ENSG00000186767\|SPIN4 | 1358.0224 | -2.2701 | ##### | ####### | 4.4247E-66 | 1.0840E-64 |
| ENSG00000265666\|RARA-AS1 | 257.8239 | -3.1485 | ##### | ####### | 4.9700E-66 | 1.2166E-64 |
| ENSG00000124588\|NQO2 | 3585.4910 | -2.1904 | ##### | ####### | 5.2762E-66 | 1.2905E-64 |
| ENSG00000111145\|ELK3 | 15458.2382 | 2.1702 | ##### | 17.1535 | 5.9180E-66 | 1.4463E-64 |
| ENSG00000118515\|SGK1 | 9237.7317 | 2.1271 | ##### | 17.1521 | 6.0605E-66 | 1.4799E-64 |
| ENSG00000152229\|PSTPIP2 | 538.3971 | -3.9853 | ##### | ####### | 6.1326E-66 | 1.4963E-64 |
| ENSG00000196843\|ARID5A | 1506.0652 | 2.2951 | ##### | 17.0972 | 1.5569E-65 | 3.7958E-64 |
| ENSG00000246705\|H2AJ | 3908.7731 | -2.1217 | ##### | ####### | 1.5670E-65 | 3.8171E-64 |
| ENSG00000251151\|HOXC-AS3 | 219.2938 | 4.4625 | ##### | 17.0941 | 1.6417E-65 | 3.9958E-64 |
| ENSG00000167508\|MVD | 4348.2004 | -2.1811 | ##### | ####### | 1.6561E-65 | 4.0277E-64 |
| ENSG00000139132\|FGD4 | 726.8525 | -2.6102 | ##### | ####### | 1.7126E-65 | 4.1618E-64 |
| ENSG00000112118\|MCM3 | 19112.9279 | 1.6742 | ##### | 17.0885 | 1.8068E-65 | 4.3870E-64 |
| ENSG00000196975\|ANXA4 | 8687.4982 | -2.0376 | ##### | ####### | 1.8717E-65 | 4.5409E-64 |
| ENSG00000109046\|WSB1 | 6625.7726 | 1.3998 | ##### | 17.0753 | 2.2651E-65 | 5.4909E-64 |
| ENSG00000146648\|EGFR | 27201.7453 | -2.0666 | ##### | ####### | 2.4203E-65 | 5.8624E-64 |
| ENSG00000133985\|TTC9 | 609.8383 | -2.7279 | ##### | ####### | 3.2592E-65 | 7.8880E-64 |
| ENSG00000167792\|NDUFV1 | 18168.1496 | 2.0088 | ##### | 17.0456 | 3.7712E-65 | 9.1197E-64 |
| ENSG00000100364\|KIAA0930 | 8513.3375 | -1.9769 | ##### | ####### | 4.4470E-65 | 1.0745E-63 |
| ENSG00000189410\|SH2D5 | 1308.9376 | -2.2990 | ##### | ####### | 4.7257E-65 | 1.1409E-63 |
| ENSG00000204396\|VWA7 | 343.8737 | -3.4323 | ##### | ####### | 5.9151E-65 | 1.4269E-63 |
| ENSG00000068724\|TTC7A | 4959.9861 | -2.0181 | ##### | ####### | 6.5955E-65 | 1.5898E-63 |
| ENSG00000160767\|ENTREP3 | 10588.2480 | 2.0490 | ##### | 17.0109 | 6.8151E-65 | 1.6414E-63 |
| ENSG00000089351\|GRAMD1A | 10051.9126 | -1.7010 | ##### | ####### | 7.1555E-65 | 1.7220E-63 |
| ENSG00000170899\|GSTA4 | 761.4102 | -3.2503 | ##### | ####### | 7.7139E-65 | 1.8549E-63 |
| ENSG00000111110\|PPM1H | 1800.6956 | -2.1568 | ##### | ####### | 8.8501E-65 | 2.1264E-63 |
| ENSG00000262001\|DLGAP1-AS2 | 1669.5132 | 2.3393 | ##### | 16.9928 | 9.2794E-65 | 2.2277E-63 |
| ENSG00000132026\|RTBDN | 236.1119 | 4.1905 | ##### | 16.9770 | 1.2158E-64 | 2.9165E-63 |
| ENSG00000110092\|CCND1 | 46947.5024 | 1.4589 | ##### | 16.9759 | 1.2379E-64 | 2.9670E-63 |
| ENSG00000109586\|GALNT7 | 4814.5758 | -2.0233 | ##### | ####### | 1.4092E-64 | 3.3749E-63 |
| ENSG00000153064\|BANK1 | 246.8071 | -3.4512 | ##### | ####### | 1.4381E-64 | 3.4413E-63 |
| ENSG00000164855\|TMEM184A | 3557.1055 | -1.9720 | ##### | ####### | 1.5267E-64 | 3.6505E-63 |
| ENSG00000161513\|FDXR | 2250.1204 | 2.2813 | ##### | 16.9634 | 1.5312E-64 | 3.6584E-63 |
| ENSG00000041982\|TNC | 7225.5208 | 2.1057 | ##### | 16.9562 | 1.7330E-64 | 4.1373E-63 |
| ENSG00000107719\|PALD1 | 1408.2941 | 2.1623 | ##### | 16.9555 | 1.7526E-64 | 4.1807E-63 |
| ENSG00000131089\|ARHGEF9 | 973.2763 | -2.3173 | ##### | ####### | 1.8176E-64 | 4.3323E-63 |
| ENSG00000205861\|PCOTH | 218.2317 | 3.3003 | ##### | 16.9483 | 1.9812E-64 | 4.7184E-63 |
| ENSG00000160741\|CRTC2 | 7868.9412 | 2.1369 | ##### | 16.9306 | 2.6780E-64 | 6.3727E-63 |
| ENSG00000056972\|TRAF3IP2 | 710.9198 | -2.5508 | ##### | ####### | 2.7201E-64 | 6.4678E-63 |
| ENSG00000104894\|CD37 | 168.4092 | -3.4250 | ##### | ####### | 3.0440E-64 | 7.2322E-63 |
| ENSG00000101057\|MYBL2 | 16058.9602 | 1.6822 | ##### | 16.9125 | 3.6380E-64 | 8.6367E-63 |
| ENSG00000071073\|MGAT4A | 1183.0492 | 2.5395 | ##### | 16.9077 | 3.9503E-64 | 9.3707E-63 |

| | | | | | | |
|---|---|---|---|---|---|---|
| ENSG00000164619\|BMPER | 1070.8812 | -2.3585 | ##### | ####### | 3.9958E-64 | 9.4711E-63 |
| ENSG00000157214\|STEAP2 | 2637.0006 | -2.2067 | ##### | ####### | 4.3106E-64 | 1.0209E-62 |
| ENSG00000281376\|ABALON | 1508.0314 | -2.3529 | ##### | ####### | 4.4012E-64 | 1.0415E-62 |
| ENSG00000116574\|RHOU | 989.6877 | -2.3613 | ##### | ####### | 4.5075E-64 | 1.0659E-62 |
| ENSG00000100401\|RANGAP1 | 16878.1166 | 1.3537 | ##### | 16.8998 | 4.5147E-64 | 1.0667E-62 |
| ENSG00000153404\|PLEKHG4B | 327.0666 | 3.2273 | ##### | 16.8877 | 5.5429E-64 | 1.3086E-62 |
| ENSG00000087245\|MMP2 | 1181.3389 | -6.7536 | ##### | ####### | 5.6412E-64 | 1.3308E-62 |
| ENSG00000164136\|IL15 | 319.2922 | -4.1050 | ##### | ####### | 5.8750E-64 | 1.3848E-62 |
| ENSG00000132692\|BCAN | 325.8668 | -3.0527 | ##### | ####### | 6.5083E-64 | 1.5329E-62 |
| ENSG00000164124\|TMEM144 | 862.5609 | -2.2140 | ##### | ####### | 6.9026E-64 | 1.6245E-62 |
| ENSG00000132915\|PDE6A | 302.5088 | -3.8131 | ##### | ####### | 8.4680E-64 | 1.9913E-62 |
| ENSG00000205542\|TMSB4X | 46641.0030 | -1.3528 | ##### | ####### | 9.2183E-64 | 2.1660E-62 |
| ENSG00000003436\|TFPI | 13166.4396 | -2.0672 | ##### | ####### | 9.3438E-64 | 2.1938E-62 |
| ENSG00000133019\|CHRM3 | 527.9069 | -2.4161 | ##### | ####### | 1.0717E-63 | 2.5143E-62 |
| ENSG00000164237\|CMBL | 9173.5224 | -2.0348 | ##### | ####### | 1.1628E-63 | 2.7257E-62 |
| ENSG00000116661\|FBXO2 | 4224.6743 | -2.0078 | ##### | ####### | 1.3802E-63 | 3.2330E-62 |
| ENSG00000128564\|VGF | 2079.2332 | 2.1656 | ##### | 16.8287 | 1.5045E-63 | 3.5214E-62 |
| ENSG00000100908\|EMC9 | 2354.2252 | 2.1714 | ##### | 16.8138 | 1.9339E-63 | 4.5228E-62 |
| ENSG00000113231\|PDE8B | 420.7873 | -3.5156 | ##### | ####### | 1.9828E-63 | 4.6336E-62 |
| ENSG00000187231\|SESTD1 | 1113.0990 | -2.3077 | ##### | ####### | 2.1940E-63 | 5.1230E-62 |
| ENSG00000120129\|DUSP1 | 4363.5070 | -2.1256 | ##### | ####### | 2.4490E-63 | 5.7141E-62 |
| ENSG00000168615\|ADAM9 | 28761.1781 | -1.9795 | ##### | ####### | 2.6713E-63 | 6.2279E-62 |
| ENSG00000072163\|LIMS2 | 418.5562 | -3.3282 | ##### | ####### | 2.9281E-63 | 6.8213E-62 |
| ENSG00000112984\|KIF20A | 7721.5555 | 2.1053 | ##### | 16.7881 | 2.9819E-63 | 6.9411E-62 |
| ENSG00000119408\|NEK6 | 3844.1845 | -2.1151 | ##### | ####### | 3.3775E-63 | 7.8559E-62 |
| ENSG00000267601 | 342.4361 | -2.8159 | ##### | ####### | 3.4006E-63 | 7.9034E-62 |
| ENSG00000134369\|NAV1 | 1986.1430 | -2.1804 | ##### | ####### | 4.1929E-63 | 9.7374E-62 |
| ENSG00000180891\|CUEDC1 | 4147.4237 | -2.1030 | ##### | ####### | 4.7444E-63 | 1.1009E-61 |
| ENSG00000020181\|ADGRA2 | 787.9292 | 3.3851 | ##### | 16.7560 | 5.1235E-63 | 1.1880E-61 |
| ENSG00000110002\|VWA5A | 1896.6484 | 2.2326 | ##### | 16.7532 | 5.3644E-63 | 1.2429E-61 |
| ENSG00000105971\|CAV2 | 13706.8827 | -1.6513 | ##### | ####### | 5.4669E-63 | 1.2656E-61 |
| ENSG00000188916\|INSYN2A | 219.8984 | -3.8219 | ##### | ####### | 5.8173E-63 | 1.3457E-61 |
| ENSG00000124795\|DEK | 16685.9024 | 1.9850 | ##### | 16.7288 | 8.0890E-63 | 1.8698E-61 |
| ENSG00000080854\|IGSF9B | 511.6098 | 4.1264 | ##### | 16.7249 | 8.6331E-63 | 1.9940E-61 |
| ENSG00000233251 | 196.1386 | -5.1378 | ##### | ####### | 1.0273E-62 | 2.3709E-61 |
| ENSG00000116141\|MARK1 | 475.1702 | 3.3203 | ##### | 16.7095 | 1.1181E-62 | 2.5785E-61 |
| ENSG00000134463\|ECHDC3 | 674.1801 | -2.6666 | ##### | ####### | 1.1650E-62 | 2.6845E-61 |
| ENSG00000119686\|FLVCR2 | 303.3249 | -3.6852 | ##### | ####### | 1.2603E-62 | 2.9021E-61 |
| ENSG00000135114\|OASL | 171.2039 | -5.1814 | ##### | ####### | 1.2842E-62 | 2.9548E-61 |
| ENSG00000137054\|POLR1E | 6327.4890 | 2.0692 | ##### | 16.6956 | 1.4100E-62 | 3.2417E-61 |
| ENSG00000213337\|ANKRD39 | 1973.5719 | 2.2024 | ##### | 16.6909 | 1.5256E-62 | 3.5048E-61 |
| ENSG00000230650 | 273.9386 | -3.3873 | ##### | ####### | 1.6217E-62 | 3.7226E-61 |
| ENSG00000167674\|HDGFL2 | 9730.2567 | 1.9484 | ##### | 16.6836 | 1.7252E-62 | 3.9573E-61 |
| ENSG00000005243\|COPZ2 | 415.9848 | -2.7378 | ##### | ####### | 1.8773E-62 | 4.3028E-61 |
| ENSG00000197461\|PDGFA | 693.7153 | -2.5009 | ##### | ####### | 2.3411E-62 | 5.3618E-61 |
| ENSG00000162849\|KIF26B | 174.3988 | 3.5590 | ##### | 16.6543 | 2.8144E-62 | 6.4408E-61 |
| ENSG00000232358 | 202.5875 | 3.4227 | ##### | 16.6509 | 2.9789E-62 | 6.8120E-61 |
| ENSG00000272620\|AFAP1-AS1 | 2318.9716 | -2.3069 | ##### | ####### | 2.9985E-62 | 6.8517E-61 |
| ENSG00000123124\|WWP1 | 4156.7149 | -1.9587 | ##### | ####### | 4.0425E-62 | 9.2301E-61 |
| ENSG00000268883\|PNMA6B | 243.7229 | 2.8019 | ##### | 16.6248 | 4.6061E-62 | 1.0509E-60 |
| ENSG00000135596\|MICAL1 | 2096.4710 | -2.1214 | ##### | ####### | 5.1645E-62 | 1.1774E-60 |
| ENSG00000197308\|GATA3-AS1 | 307.8052 | 2.8092 | ##### | 16.6168 | 5.2627E-62 | 1.1989E-60 |
| ENSG00000130826\|DKC1 | 21272.7236 | 1.3075 | ##### | 16.6020 | 6.7421E-62 | 1.5347E-60 |
| ENSG00000137691\|CFAP300 | 458.4614 | -2.5711 | ##### | ####### | 7.2186E-62 | 1.6419E-60 |
| ENSG00000116132\|PRRX1 | 464.9540 | 2.7582 | ##### | 16.5943 | 7.6630E-62 | 1.7417E-60 |
| ENSG00000166145\|SPINT1 | 490.7003 | -3.1108 | ##### | ####### | 7.8745E-62 | 1.7884E-60 |
| ENSG00000269994\|LINC02893 | 283.9419 | -5.3992 | ##### | ####### | 7.9678E-62 | 1.8082E-60 |
| ENSG00000164930\|FZD6 | 2959.5002 | -2.1222 | ##### | ####### | 8.3884E-62 | 1.9022E-60 |
| ENSG00000073331\|ALPK1 | 545.4568 | -2.5170 | ##### | ####### | 9.4465E-62 | 2.1405E-60 |
| ENSG00000166508\|MCM7 | 38668.8740 | 1.4722 | ##### | 16.5720 | 1.1109E-61 | 2.5154E-60 |
| ENSG00000261578 | 336.7383 | -2.6717 | ##### | ####### | 1.1186E-61 | 2.5307E-60 |
| ENSG00000291131 | 2072.4987 | ####### | ##### | ####### | 1.5829E-61 | 3.5787E-60 |
| ENSG00000117407\|ARTN | 569.3753 | 2.4385 | ##### | 16.5502 | 1.5950E-61 | 3.6033E-60 |

| | | | | | | |
|---|---|---|---|---|---|---|
| ENSG00000101417\|PXMP4 | 546.4144 | -2.7756 | ##### | ####### | 1.6171E-61 | 3.6505E-60 |
| ENSG00000181381\|DDX60L | 1042.9216 | -2.2387 | ##### | ####### | 1.6317E-61 | 3.6805E-60 |
| ENSG00000189420\|ZFP92 | 571.5763 | 2.5127 | ##### | 16.5411 | 1.8550E-61 | 4.1812E-60 |
| ENSG00000280435 | 208.4659 | -4.2201 | ##### | ####### | 1.9499E-61 | 4.3918E-60 |
| ENSG00000040731\|CDH10 | 279.9183 | 3.4971 | ##### | 16.5368 | 1.9933E-61 | 4.4861E-60 |
| ENSG00000169756\|LIMS1 | 8731.9214 | -1.9964 | ##### | ####### | 2.1568E-61 | 4.8503E-60 |
| ENSG00000085276\|MECOM | 1042.2932 | -2.5784 | ##### | ####### | 2.2051E-61 | 4.9554E-60 |
| ENSG00000145526\|CDH18 | 414.6302 | -2.7092 | ##### | ####### | 2.6374E-61 | 5.9222E-60 |
| ENSG00000205181\|LINC00654 | 121.0955 | -4.7482 | ##### | ####### | 2.7731E-61 | 6.2223E-60 |
| ENSG00000168672\|LRATD2 | 1217.2173 | 2.9193 | ##### | 16.4999 | 3.6746E-61 | 8.2388E-60 |
| ENSG00000007968\|E2F2 | 3433.7225 | 1.4141 | ##### | 16.4944 | 4.0233E-61 | 9.0139E-60 |
| ENSG00000147475\|ERLIN2 | 3357.9010 | -1.7203 | ##### | ####### | 4.6288E-61 | 1.0363E-59 |
| ENSG00000114698\|PLSCR4 | 428.0426 | -5.5181 | ##### | ####### | 4.6547E-61 | 1.0413E-59 |
| ENSG00000101191\|DIDO1 | 9471.9100 | 1.9931 | ##### | 16.4806 | 5.0558E-61 | 1.1302E-59 |
| ENSG00000171843\|MLLT3 | 748.8257 | -2.4196 | ##### | ####### | 5.1808E-61 | 1.1572E-59 |
| ENSG00000008256\|CYTH3 | 4404.5344 | -1.8023 | ##### | ####### | 5.7231E-61 | 1.2774E-59 |
| ENSG00000125864\|BFSP1 | 512.5219 | -2.7016 | ##### | ####### | 6.0647E-61 | 1.3527E-59 |
| ENSG00000290692 | 178.0977 | -3.6963 | ##### | ####### | 6.3865E-61 | 1.4234E-59 |
| ENSG00000163735\|CXCL5 | 32421.0955 | ####### | ##### | ####### | 6.8303E-61 | 1.5212E-59 |
| ENSG00000171757\|LRRC34 | 197.1841 | 4.1723 | ##### | 16.4578 | 7.3698E-61 | 1.6401E-59 |
| ENSG00000198246\|SLC29A3 | 930.4601 | -2.3755 | ##### | ####### | 7.5989E-61 | 1.6898E-59 |
| ENSG00000179604\|CDC42EP4 | 4216.3638 | -2.0579 | ##### | ####### | 8.0601E-61 | 1.7910E-59 |
| ENSG00000100106\|TRIOBP | 5048.2643 | -1.9491 | ##### | ####### | 8.2135E-61 | 1.8238E-59 |
| ENSG00000286273 | 190.7998 | 4.3926 | ##### | 16.4474 | 8.7592E-61 | 1.9435E-59 |
| ENSG00000238279 | 192.5998 | 2.8608 | ##### | 16.4430 | 9.4214E-61 | 2.0889E-59 |
| ENSG00000051596\|THOC3 | 7214.2596 | 2.0340 | ##### | 16.4301 | 1.1638E-60 | 2.5784E-59 |
| ENSG00000197299\|BLM | 2151.0289 | 1.6607 | ##### | 16.4249 | 1.2694E-60 | 2.8103E-59 |
| ENSG00000259583\|ALDH1A3-AS1 | 502.7690 | -2.5777 | ##### | ####### | 1.4289E-60 | 3.1592E-59 |
| ENSG00000197405\|C5AR1 | 376.9258 | -5.0181 | ##### | ####### | 1.4291E-60 | 3.1592E-59 |
| ENSG00000124789\|NUP153 | 15638.4091 | 1.9724 | ##### | 16.4122 | 1.5647E-60 | 3.4563E-59 |
| ENSG00000108679\|LGALS3BP | 395.0644 | -2.7575 | ##### | ####### | 1.7522E-60 | 3.8678E-59 |
| ENSG00000185189\|NRBP2 | 3413.8738 | -1.9033 | ##### | ####### | 1.8016E-60 | 3.9737E-59 |
| ENSG00000171298\|GAA | 7742.6591 | -1.9297 | ##### | ####### | 1.8770E-60 | 4.1369E-59 |
| ENSG00000144029\|MRPS5 | 9645.3216 | 1.6874 | ##### | 16.3981 | 1.9720E-60 | 4.3431E-59 |
| ENSG00000198055\|GRK6 | 5568.4432 | 2.0318 | ##### | 16.3948 | 2.0844E-60 | 4.5873E-59 |
| ENSG00000069702\|TGFBR3 | 2328.0053 | -2.1799 | ##### | ####### | 2.1830E-60 | 4.8007E-59 |
| ENSG00000070814\|TCOF1 | 19201.5043 | 1.9096 | ##### | 16.3876 | 2.3436E-60 | 5.1502E-59 |
| ENSG00000143549\|TPM3 | 87741.3335 | 1.2910 | ##### | 16.3843 | 2.4779E-60 | 5.4414E-59 |
| ENSG00000062282\|DGAT2 | 3296.4537 | 2.0373 | ##### | 16.3819 | 2.5769E-60 | 5.6546E-59 |
| ENSG00000256802\|LCIIAR | 586.0805 | -2.2862 | ##### | ####### | 2.8072E-60 | 6.1554E-59 |
| ENSG00000293490\|SLC66A1L | 318.3044 | -3.2499 | ##### | ####### | 2.8399E-60 | 6.2225E-59 |
| ENSG00000228672\|PROB1 | 920.9978 | 2.2878 | ##### | 16.3734 | 2.9614E-60 | 6.4840E-59 |
| ENSG00000115170\|ACVR1 | 3491.6409 | -2.1008 | ##### | ####### | 2.9917E-60 | 6.5456E-59 |
| ENSG00000254285\|KRT8P3 | 216.3123 | -3.1509 | ##### | ####### | 3.2468E-60 | 7.0986E-59 |
| ENSG00000103260\|METRN | 5469.4796 | -1.9594 | ##### | ####### | 3.7076E-60 | 8.1001E-59 |
| ENSG00000122435\|TRMT13 | 1638.3611 | -2.1262 | ##### | ####### | 4.0094E-60 | 8.7529E-59 |
| ENSG00000249550\|LINC01234 | 237.3027 | -4.0745 | ##### | ####### | 4.3739E-60 | 9.5417E-59 |
| ENSG00000121900\|TMEM54 | 3822.2033 | -2.0704 | ##### | ####### | 4.7321E-60 | 1.0316E-58 |
| ENSG00000131746\|TNS4 | 291.2178 | -6.1733 | ##### | ####### | 5.1509E-60 | 1.1220E-58 |
| ENSG00000111665\|CDCA3 | 7138.4491 | 2.0599 | ##### | 16.3395 | 5.1650E-60 | 1.1243E-58 |
| ENSG00000145555\|MYO10 | 15971.8958 | -1.9636 | ##### | ####### | 5.4355E-60 | 1.1823E-58 |
| ENSG00000173041\|ZNF680 | 405.7310 | -2.5946 | ##### | ####### | 5.7534E-60 | 1.2505E-58 |
| ENSG00000118655\|DCLRE1B | 3594.8613 | 1.1880 | ##### | 16.3298 | 6.0573E-60 | 1.3157E-58 |
| ENSG00000259948 | 261.0711 | -2.6183 | ##### | ####### | 6.7942E-60 | 1.4746E-58 |
| ENSG00000072210\|ALDH3A2 | 10817.9589 | -2.0430 | ##### | ####### | 7.2047E-60 | 1.5626E-58 |
| ENSG00000158555\|GDPD5 | 685.9996 | -2.4829 | ##### | ####### | 7.8572E-60 | 1.7029E-58 |
| ENSG00000104763\|ASAH1 | 6467.0572 | -2.0340 | ##### | ####### | 8.4266E-60 | 1.8249E-58 |
| ENSG00000166886\|NAB2 | 5514.1391 | 1.9819 | ##### | 16.3035 | 9.3229E-60 | 2.0176E-58 |
| ENSG00000124313\|IQSEC2 | 1828.6367 | -2.0424 | ##### | ####### | 1.0422E-59 | 2.2538E-58 |
| ENSG00000114854\|TNNC1 | 256.1759 | -5.0622 | ##### | ####### | 1.2343E-59 | 2.6674E-58 |
| ENSG00000135549\|PKIB | 366.3917 | -6.8699 | ##### | ####### | 1.4541E-59 | 3.1401E-58 |
| ENSG00000229931\|ATXN1-AS1 | 231.0275 | 4.0896 | ##### | 16.2723 | 1.5516E-59 | 3.3482E-58 |
| ENSG00000135269\|TES | 8058.5755 | -2.0085 | ##### | ####### | 1.7473E-59 | 3.7677E-58 |

| | | | | | | |
|---|---|---|---|---|---|---|
| ENSG00000272341 | 190.2459 | 4.3546 | ##### | 16.2550 | 2.0602E-59 | 4.4392E-58 |
| ENSG00000102580\|DNAJC3 | 3904.7384 | -2.0186 | ##### | ####### | 2.1268E-59 | 4.5794E-58 |
| ENSG00000225362\|CT62 | 148.1444 | 4.8536 | ##### | 16.2484 | 2.2923E-59 | 4.9322E-58 |
| ENSG00000167700\|MFSD3 | 1278.5865 | -2.1577 | ##### | ####### | 2.4747E-59 | 5.3210E-58 |
| ENSG00000103196\|CRISPLD2 | 329.9479 | -2.8247 | ##### | ####### | 2.6701E-59 | 5.7369E-58 |
| ENSG00000164695\|CHMP4C | 895.1332 | -2.3006 | ##### | ####### | 3.0176E-59 | 6.4787E-58 |
| ENSG00000131171\|SH3BGRL | 3582.9247 | -6.6946 | ##### | ####### | 3.2040E-59 | 6.8741E-58 |
| ENSG00000137261\|KIAA0319 | 633.9425 | -2.4636 | ##### | ####### | 3.5947E-59 | 7.7067E-58 |
| ENSG00000186567\|CEACAM19 | 327.0804 | -3.8697 | ##### | ####### | 3.9854E-59 | 8.5384E-58 |
| ENSG00000147604\|RPL7 | 41757.4367 | -1.4050 | ##### | ####### | 4.9753E-59 | 1.0651E-57 |
| ENSG00000138759\|FRAS1 | 380.3468 | -2.7464 | ##### | ####### | 5.0695E-59 | 1.0845E-57 |
| ENSG00000163536\|SERPINI1 | 172.9263 | -3.2744 | ##### | ####### | 5.3103E-59 | 1.1348E-57 |
| ENSG00000188732\|FAM221A | 123.6952 | -6.8827 | ##### | ####### | 5.3122E-59 | 1.1348E-57 |
| ENSG00000069020\|MAST4 | 810.5414 | -2.7190 | ##### | ####### | 5.5579E-59 | 1.1865E-57 |
| ENSG00000066468\|FGFR2 | 289.2933 | 3.8531 | ##### | 16.1937 | 5.5851E-59 | 1.1914E-57 |
| ENSG00000165359\|INTS6L | 1272.9426 | 2.1863 | ##### | 16.1937 | 5.5909E-59 | 1.1918E-57 |
| ENSG00000100311\|PDGFB | 352.8676 | -2.8897 | ##### | ####### | 5.9108E-59 | 1.2591E-57 |
| ENSG00000006611\|USH1C | 1844.7112 | ####### | ##### | ####### | 6.0936E-59 | 1.2971E-57 |
| ENSG00000139174\|PRICKLE1 | 500.3738 | -2.8438 | ##### | ####### | 6.3530E-59 | 1.3514E-57 |
| ENSG00000143013\|LMO4 | 4422.8321 | -1.9735 | ##### | ####### | 6.9722E-59 | 1.4820E-57 |
| ENSG00000196756\|SNHG17 | 5003.5356 | 2.0856 | ##### | 16.1796 | 7.0213E-59 | 1.4914E-57 |
| ENSG00000009307\|CSDE1 | 82980.0177 | 1.2593 | ##### | 16.1752 | 7.5401E-59 | 1.6005E-57 |
| ENSG00000273604\|EPOP | 3858.6960 | 1.5461 | ##### | 16.1688 | 8.3719E-59 | 1.7758E-57 |
| ENSG00000124228\|DDX27 | 17300.6103 | 1.7833 | ##### | 16.1686 | 8.3996E-59 | 1.7804E-57 |
| ENSG00000162129\|CLPB | 4628.0943 | 1.5739 | ##### | 16.1514 | 1.1099E-58 | 2.3509E-57 |
| ENSG00000162039\|MEIOB | 2684.3613 | -9.6882 | ##### | ####### | 1.1115E-58 | 2.3527E-57 |
| ENSG00000152527\|PLEKHH2 | 372.0012 | -2.7572 | ##### | ####### | 1.2268E-58 | 2.5947E-57 |
| ENSG00000271503\|CCL5 | 704.7776 | 2.8149 | ##### | 16.1447 | 1.2370E-58 | 2.6146E-57 |
| ENSG00000168763\|CNNM3 | 5030.6076 | 1.6762 | ##### | 16.1365 | 1.4142E-58 | 2.9869E-57 |
| ENSG00000257594\|GALNT4 | 1266.0341 | -2.3396 | ##### | ####### | 1.4811E-58 | 3.1261E-57 |
| ENSG00000162772\|ATF3 | 1770.9774 | 2.1143 | ##### | 16.1306 | 1.5545E-58 | 3.2785E-57 |
| ENSG00000108840\|HDAC5 | 1932.0248 | 2.1517 | ##### | 16.1151 | 1.9971E-58 | 4.2092E-57 |
| ENSG00000173530\|TNFRSF10D | 7432.2153 | 2.0343 | ##### | 16.1096 | 2.1849E-58 | 4.6017E-57 |
| ENSG00000143198\|MGST3 | 6188.0313 | -1.9723 | ##### | ####### | 2.4598E-58 | 5.1770E-57 |
| ENSG00000242265\|PEG10 | 17775.7055 | -1.9278 | ##### | ####### | 2.8417E-58 | 5.9765E-57 |
| ENSG00000227124\|ZNF717 | 257.2582 | -2.8749 | ##### | ####### | 3.2347E-58 | 6.7983E-57 |
| ENSG00000143382\|ADAMTSL4 | 1611.3520 | -2.0283 | ##### | ####### | 4.0990E-58 | 8.6087E-57 |
| ENSG00000117859\|OSBPL9 | 7507.0380 | -1.2932 | ##### | ####### | 4.1360E-58 | 8.6804E-57 |
| ENSG00000035681\|NSMAF | 3070.6491 | -2.0078 | ##### | ####### | 4.2592E-58 | 8.9326E-57 |
| ENSG00000064393\|HIPK2 | 4621.1135 | -1.9772 | ##### | ####### | 4.4943E-58 | 9.4191E-57 |
| ENSG00000251493\|FOXD1 | 2856.5626 | 2.0512 | ##### | 16.0635 | 4.6011E-58 | 9.6362E-57 |
| ENSG00000177989\|CIMAP1B | 459.1310 | -2.6263 | ##### | ####### | 4.8110E-58 | 1.0069E-56 |
| ENSG00000135924\|DNAJB2 | 9599.5804 | 1.9884 | ##### | 16.0590 | 4.9445E-58 | 1.0341E-56 |
| ENSG00000155849\|ELMO1 | 290.7241 | -2.9941 | ##### | ####### | 7.1605E-58 | 1.4965E-56 |
| ENSG00000196872\|CRACDL | 820.4839 | 2.4292 | ##### | 16.0341 | 7.3895E-58 | 1.5433E-56 |
| ENSG00000167996\|FTH1 | ########## | -1.9668 | ##### | ####### | 9.4290E-58 | 1.9678E-56 |
| ENSG00000273297\|LINC03060 | 308.2497 | -2.7796 | ##### | ####### | 9.7019E-58 | 2.0234E-56 |
| ENSG00000071242\|RPS6KA2 | 507.1595 | 2.5526 | ##### | 16.0131 | 1.0355E-57 | 2.1581E-56 |
| ENSG00000104312\|RIPK2 | 2822.3051 | -2.0271 | ##### | ####### | 1.1632E-57 | 2.4225E-56 |
| ENSG00000133083\|DCLK1 | 7327.3250 | 1.9023 | ##### | 16.0044 | 1.1913E-57 | 2.4794E-56 |
| ENSG00000163468\|CCT3 | 83467.0978 | 1.3700 | ##### | 16.0028 | 1.2218E-57 | 2.5410E-56 |
| ENSG00000102893\|PHKB | 4782.4112 | -1.9503 | ##### | ####### | 1.5215E-57 | 3.1621E-56 |
| ENSG00000108984\|MAP2K6 | 413.6560 | -2.8074 | ##### | ####### | 1.6741E-57 | 3.4768E-56 |
| ENSG00000171227\|TMEM37 | 333.3103 | -4.7272 | ##### | ####### | 1.8065E-57 | 3.7492E-56 |
| ENSG00000159899\|NPR2 | 168.3496 | -3.9969 | ##### | ####### | 1.9118E-57 | 3.9650E-56 |
| ENSG00000164509\|IL31RA | 1082.4526 | 2.7116 | ##### | 15.9702 | 2.0619E-57 | 4.2732E-56 |
| ENSG00000213025\|COX20P1 | 224.3158 | 4.1136 | ##### | 15.9676 | 2.1490E-57 | 4.4508E-56 |
| ENSG00000186088\|GSAP | 690.8328 | -2.3479 | ##### | ####### | 2.5617E-57 | 5.3019E-56 |
| ENSG00000100867\|DHRS2 | 471.4989 | 2.8338 | ##### | 15.9493 | 2.8793E-57 | 5.9550E-56 |
| ENSG00000107099\|DOCK8 | 254.3524 | -3.4273 | ##### | ####### | 3.0066E-57 | 6.2139E-56 |
| ENSG00000146918\|NCAPG2 | 16024.7629 | 1.9042 | ##### | 15.9407 | 3.3082E-57 | 6.8326E-56 |
| ENSG00000242498\|ARPIN | 1571.1195 | -2.0446 | ##### | ####### | 3.3129E-57 | 6.8375E-56 |
| ENSG00000157657\|ZNF618 | 2451.6888 | -2.0901 | ##### | ####### | 3.5845E-57 | 7.3930E-56 |

| | | | | | | |
|---|---|---|---|---|---|---|
| ENSG00000171813\|PWWP2B | 1547.6925 | -2.1303 | ##### | ####### | 3.7226E-57 | 7.6725E-56 |
| ENSG00000204264\|PSMB8 | 2408.7173 | -1.9308 | ##### | ####### | 4.2877E-57 | 8.8313E-56 |
| ENSG00000188660\|LINC00319 | 215.9970 | -3.0806 | ##### | ####### | 4.8463E-57 | 9.9749E-56 |
| ENSG00000291100 | 3027.8187 | -1.3638 | ##### | ####### | 5.4254E-57 | 1.1159E-55 |
| ENSG00000147642\|SYBU | 1571.1153 | -2.1618 | ##### | ####### | 6.1040E-57 | 1.2546E-55 |
| ENSG00000243232\|PCDHAC2 | 506.5854 | -3.0337 | ##### | ####### | 6.4700E-57 | 1.3289E-55 |
| ENSG00000221990\|EXOC3-AS1 | 243.6270 | -2.8470 | ##### | ####### | 8.4784E-57 | 1.7403E-55 |
| ENSG00000152076\|CCDC74B | 2010.2487 | 2.1332 | ##### | 15.8785 | 8.9237E-57 | 1.8304E-55 |
| ENSG00000120149\|MSX2 | 286.0568 | 2.8731 | ##### | 15.8782 | 8.9727E-57 | 1.8392E-55 |
| ENSG00000101187\|SLCO4A1 | 6326.0660 | 1.8670 | ##### | 15.8777 | 9.0493E-57 | 1.8536E-55 |
| ENSG00000181577\|LINC03040 | 165.6811 | -7.9658 | ##### | ####### | 1.0596E-56 | 2.1689E-55 |
| ENSG00000086289\|EPDR1 | 8580.7422 | -1.9774 | ##### | ####### | 1.0666E-56 | 2.1818E-55 |
| ENSG00000160588\|MPZL3 | 186.0642 | -3.1599 | ##### | ####### | 1.2852E-56 | 2.6271E-55 |
| ENSG00000164924\|YWHAZ | 79542.8742 | -1.3841 | ##### | ####### | 1.2935E-56 | 2.6424E-55 |
| ENSG00000152137\|HSPB8 | 612.9786 | -2.5150 | ##### | ####### | 1.4009E-56 | 2.8596E-55 |
| ENSG00000163444\|TMEM183A | 5387.7385 | -0.9697 | ##### | ####### | 1.7856E-56 | 3.6425E-55 |
| ENSG00000116711\|PLA2G4A | 843.4764 | -2.2526 | ##### | ####### | 1.8447E-56 | 3.7605E-55 |
| ENSG00000135451\|TROAP | 5043.5152 | 1.9939 | ##### | 15.8287 | 1.9717E-56 | 4.0166E-55 |
| ENSG00000164741\|DLC1 | 1651.9017 | 2.0870 | ##### | 15.8266 | 2.0388E-56 | 4.1506E-55 |
| ENSG00000196754\|S100A2 | 717.9737 | 2.3893 | ##### | 15.8263 | 2.0497E-56 | 4.1699E-55 |
| ENSG00000156804\|FBXO32 | 271.1735 | -3.1038 | ##### | ####### | 2.0589E-56 | 4.1858E-55 |
| ENSG00000150471\|ADGRL3 | 170.6920 | -3.0462 | ##### | ####### | 2.0821E-56 | 4.2302E-55 |
| ENSG00000221968\|FADS3 | 2602.2816 | -2.0033 | ##### | ####### | 2.1989E-56 | 4.4643E-55 |
| ENSG00000113448\|PDE4D | 4972.2020 | -2.0079 | ##### | ####### | 2.2352E-56 | 4.5349E-55 |
| ENSG00000102897\|LYRM1 | 1569.8001 | -2.1261 | ##### | ####### | 2.3466E-56 | 4.7577E-55 |
| ENSG00000142634\|EFHD2 | 16289.6102 | -1.8121 | ##### | ####### | 2.3519E-56 | 4.7653E-55 |
| ENSG00000128512\|DOCK4 | 3451.6459 | -2.0424 | ##### | ####### | 2.4787E-56 | 5.0188E-55 |
| ENSG00000113368\|LMNB1 | 9048.5968 | 1.4571 | ##### | 15.8122 | 2.5628E-56 | 5.1855E-55 |
| ENSG00000050730\|TNIP3 | 228.1448 | -3.7242 | ##### | ####### | 2.8151E-56 | 5.6922E-55 |
| ENSG00000196739\|COL27A1 | 2191.7530 | -2.0229 | ##### | ####### | 2.8200E-56 | 5.6982E-55 |
| ENSG00000228232\|GAPDHP1 | 893.1777 | 2.3377 | ##### | 15.8014 | 3.0405E-56 | 6.1396E-55 |
| ENSG00000106351\|AGFG2 | 2049.7165 | -2.2397 | ##### | ####### | 3.4160E-56 | 6.8931E-55 |
| ENSG00000130829\|DUSP9 | 1078.9009 | 2.3188 | ##### | 15.7892 | 3.6913E-56 | 7.4438E-55 |
| ENSG00000025293\|PHF20 | 14689.1811 | 1.7243 | ##### | 15.7877 | 3.7794E-56 | 7.6162E-55 |
| ENSG00000166106\|ADAMTS15 | 762.2335 | -2.4958 | ##### | ####### | 4.0352E-56 | 8.1262E-55 |
| ENSG00000013375\|PGM3 | 7377.4826 | -1.9204 | ##### | ####### | 4.1570E-56 | 8.3603E-55 |
| ENSG00000186480\|INSIG1 | 2162.0822 | -1.9684 | ##### | ####### | 4.1571E-56 | 8.3603E-55 |
| ENSG00000125629\|INSIG2 | 1463.3588 | -2.0863 | ##### | ####### | 4.3504E-56 | 8.7432E-55 |
| ENSG00000276223 | 172.8696 | 3.1434 | ##### | 15.7605 | 5.8198E-56 | 1.1689E-54 |
| ENSG00000179674\|ARL14 | 1306.4767 | 2.2617 | ##### | 15.7576 | 6.0942E-56 | 1.2231E-54 |
| ENSG00000128311\|TST | 1767.5749 | -2.1387 | ##### | ####### | 6.3921E-56 | 1.2821E-54 |
| ENSG00000273706\|LHX1 | 206.7320 | -3.2990 | ##### | ####### | 6.5556E-56 | 1.3140E-54 |
| ENSG00000145349\|CAMK2D | 2098.6148 | -1.9421 | ##### | ####### | 8.9740E-56 | 1.7975E-54 |
| ENSG00000119929\|CUTC | 1720.7185 | -1.9060 | ##### | ####### | 8.9922E-56 | 1.7999E-54 |
| ENSG00000197183\|NOL4L | 1138.4265 | -2.2177 | ##### | ####### | 1.0227E-55 | 2.0458E-54 |
| ENSG00000141524\|TMC6 | 1595.3147 | -2.0253 | ##### | ####### | 1.0801E-55 | 2.1591E-54 |
| ENSG00000107968\|MAP3K8 | 1453.5783 | 2.2272 | ##### | 15.7202 | 1.0993E-55 | 2.1961E-54 |
| ENSG00000065717\|TLE2 | 4676.7822 | -1.9202 | ##### | ####### | 1.1021E-55 | 2.2002E-54 |
| ENSG00000028137\|TNFRSF1B | 282.1385 | -2.6944 | ##### | ####### | 1.2766E-55 | 2.5468E-54 |
| ENSG00000162337\|LRP5 | 6100.9702 | -1.8736 | ##### | ####### | 1.5901E-55 | 3.1702E-54 |
| ENSG00000099994\|SUSD2 | 336.5932 | -3.2744 | ##### | ####### | 1.7563E-55 | 3.4991E-54 |
| ENSG00000075461\|CACNG4 | 1669.8377 | 2.2486 | ##### | 15.6622 | 2.7411E-55 | 5.4576E-54 |
| ENSG00000254535\|PABPC4L | 1361.3858 | ####### | ##### | ####### | 3.0349E-55 | 6.0384E-54 |
| ENSG00000181104\|F2R | 2998.1257 | 1.9576 | ##### | 15.6490 | 3.3758E-55 | 6.7123E-54 |
| ENSG00000121691\|CAT | 2675.7774 | -1.9663 | ##### | ####### | 3.6192E-55 | 7.1915E-54 |
| ENSG00000140451\|PIF1 | 979.1284 | 2.2764 | ##### | 15.6409 | 3.8347E-55 | 7.6146E-54 |
| ENSG00000129636\|ITFG1 | 4086.5100 | -1.8613 | ##### | ####### | 3.8707E-55 | 7.6810E-54 |
| ENSG00000082126\|MPP4 | 243.6160 | 3.7343 | ##### | 15.6383 | 3.9951E-55 | 7.9226E-54 |
| ENSG00000116580\|GON4L | 9372.9329 | 1.8442 | ##### | 15.6352 | 4.1914E-55 | 8.3064E-54 |
| ENSG00000171533\|MAP6 | 201.2562 | 4.1659 | ##### | 15.6350 | 4.2060E-55 | 8.3297E-54 |
| ENSG00000082497\|SERTAD4 | 1135.3926 | -2.1310 | ##### | ####### | 4.5270E-55 | 8.9596E-54 |
| ENSG00000138356\|AOX1 | 967.0614 | -2.0873 | ##### | ####### | 4.9993E-55 | 9.8877E-54 |
| ENSG00000109572\|CLCN3 | 5812.7711 | -1.8718 | ##### | ####### | 5.6603E-55 | 1.1188E-53 |

| | | | | | | |
|---|---|---|---|---|---|---|
| ENSG00000168404\|MLKL | 2224.7673 | -1.9785 | ##### | ####### | 5.8943E-55 | 1.1642E-53 |
| ENSG00000102034\|ELF4 | 2066.2624 | -1.9907 | ##### | ####### | 6.4128E-55 | 1.2658E-53 |
| ENSG00000181019\|NQO1 | 61619.8579 | -1.6575 | ##### | ####### | 6.7108E-55 | 1.3238E-53 |
| ENSG00000143590\|EFNA3 | 942.4877 | 2.1178 | ##### | 15.6033 | 6.9174E-55 | 1.3636E-53 |
| ENSG00000178342\|KCNG2 | 183.0521 | -3.3271 | ##### | ####### | 6.9405E-55 | 1.3673E-53 |
| ENSG00000175906\|ARL4D | 1412.7985 | -2.1362 | ##### | ####### | 7.1350E-55 | 1.4047E-53 |
| ENSG00000167670\|CHAF1A | 9719.3988 | 1.7875 | ##### | 15.5955 | 7.8060E-55 | 1.5357E-53 |
| ENSG00000159164\|SV2A | 1433.1443 | 1.9771 | ##### | 15.5902 | 8.4855E-55 | 1.6683E-53 |
| ENSG00000125378\|BMP4 | 171.8617 | 4.9119 | ##### | 15.5851 | 9.1979E-55 | 1.8072E-53 |
| ENSG00000117543\|DPH5 | 1978.7473 | -2.0440 | ##### | ####### | 9.2431E-55 | 1.8149E-53 |
| ENSG00000101109\|STK4 | 11332.8064 | 1.8866 | ##### | 15.5746 | 1.0831E-54 | 2.1253E-53 |
| ENSG00000081377\|CDC14B | 2129.6936 | -1.9234 | ##### | ####### | 1.0879E-54 | 2.1332E-53 |
| ENSG00000214357\|NEURL1B | 1067.9341 | 2.2119 | ##### | 15.5631 | 1.2963E-54 | 2.5403E-53 |
| ENSG00000138771\|SHROOM3 | 1613.2729 | -1.9383 | ##### | ####### | 1.4007E-54 | 2.7430E-53 |
| ENSG00000234678\|ELF3-AS1 | 367.2211 | -2.7886 | ##### | ####### | 1.4409E-54 | 2.8199E-53 |
| ENSG00000143369\|ECM1 | 352.5816 | -2.9233 | ##### | ####### | 1.9718E-54 | 3.8564E-53 |
| ENSG00000239887\|C1orf226 | 394.0316 | -2.8390 | ##### | ####### | 2.1584E-54 | 4.2186E-53 |
| ENSG00000197905\|TEAD4 | 7378.9523 | 1.8900 | ##### | 15.5250 | 2.3509E-54 | 4.5918E-53 |
| ENSG00000171858\|RPS21 | 38192.9338 | 2.0169 | ##### | 15.5215 | 2.4801E-54 | 4.8411E-53 |
| ENSG00000129226\|CD68 | 1312.0355 | -2.1263 | ##### | ####### | 2.7256E-54 | 5.3168E-53 |
| ENSG00000176697\|BDNF | 1088.2984 | -1.6935 | ##### | ####### | 2.9509E-54 | 5.7525E-53 |
| ENSG00000166377\|ATP9B | 2081.4048 | -1.9430 | ##### | ####### | 3.9275E-54 | 7.6513E-53 |
| ENSG00000134072\|CAMK1 | 388.1556 | -3.2185 | ##### | ####### | 4.4941E-54 | 8.7494E-53 |
| ENSG00000087087\|SRRT | 13893.5799 | 1.0053 | ##### | 15.4802 | 4.7223E-54 | 9.1877E-53 |
| ENSG00000197043\|ANXA6 | 4622.5609 | -1.8763 | ##### | ####### | 4.8003E-54 | 9.3333E-53 |
| ENSG00000109674\|NEIL3 | 2182.0571 | -1.9876 | ##### | ####### | 6.3976E-54 | 1.2431E-52 |
| ENSG00000162434\|JAK1 | 27394.3196 | -1.8377 | ##### | ####### | 6.6746E-54 | 1.2961E-52 |
| ENSG00000260083\|MIR762HG | 294.4784 | -2.8168 | ##### | ####### | 6.8402E-54 | 1.3274E-52 |
| ENSG00000171914\|TLN2 | 1580.6972 | -2.1365 | ##### | ####### | 7.7617E-54 | 1.5052E-52 |
| ENSG00000136108\|CKAP2 | 8252.0333 | 1.5639 | ##### | 15.4461 | 8.0138E-54 | 1.5531E-52 |
| ENSG00000128016\|ZFP36 | 1981.7725 | -1.9988 | ##### | ####### | 8.3375E-54 | 1.6148E-52 |
| ENSG00000128274\|A4GALT | 964.7692 | -2.1719 | ##### | ####### | 9.1389E-54 | 1.7688E-52 |
| ENSG00000204869\|IGFL4 | 224.7712 | 5.3589 | ##### | 15.4304 | 1.0221E-53 | 1.9771E-52 |
| ENSG00000159228\|CBR1 | 11425.8164 | -1.7762 | ##### | ####### | 1.0927E-53 | 2.1122E-52 |
| ENSG00000167658\|EEF2 | ########## | 0.9878 | ##### | 15.4043 | 1.5323E-53 | 2.9600E-52 |
| ENSG00000172260\|NEGR1 | 353.9332 | -2.6069 | ##### | ####### | 1.7588E-53 | 3.3953E-52 |
| ENSG00000112029\|FBXO5 | 4408.4554 | 1.9438 | ##### | 15.3949 | 1.7705E-53 | 3.4157E-52 |
| ENSG00000144118\|RALB | 7811.7121 | -1.8406 | ##### | ####### | 1.7778E-53 | 3.4276E-52 |
| ENSG00000090238\|YPEL3 | 153.6875 | -4.2484 | ##### | ####### | 1.9574E-53 | 3.7715E-52 |
| ENSG00000103152\|MPG | 2936.7424 | -1.9514 | ##### | ####### | 1.9898E-53 | 3.8314E-52 |
| ENSG00000167695\|TLCD3A | 1743.6533 | -1.9172 | ##### | ####### | 2.1195E-53 | 4.0786E-52 |
| ENSG00000138193\|PLCE1 | 1810.4915 | -1.9949 | ##### | ####### | 2.2019E-53 | 4.2344E-52 |
| ENSG00000085662\|AKR1B1 | 36445.7217 | -1.7988 | ##### | ####### | 2.4382E-53 | 4.6859E-52 |
| ENSG00000127507\|ADGRE2 | 674.2888 | 3.1907 | ##### | 15.3679 | 2.6877E-53 | 5.1619E-52 |
| ENSG00000121316\|PLBD1 | 1496.9733 | -1.8860 | ##### | ####### | 3.1479E-53 | 6.0418E-52 |
| ENSG00000137267\|TUBB2A | 3198.5506 | -1.8881 | ##### | ####### | 3.5108E-53 | 6.7342E-52 |
| ENSG00000149639\|SOGA1 | 10748.4643 | 2.0128 | ##### | 15.3429 | 3.9493E-53 | 7.5704E-52 |
| ENSG00000277701 | 408.4065 | 2.3585 | ##### | 15.3417 | 4.0248E-53 | 7.7101E-52 |
| ENSG00000175832\|ETV4 | 7030.9167 | 1.9779 | ##### | 15.3414 | 4.0433E-53 | 7.7406E-52 |
| ENSG00000160803\|UBQLN4 | 16098.5571 | 1.8903 | ##### | 15.3384 | 4.2360E-53 | 8.1044E-52 |
| ENSG00000122705\|CLTA | 21579.2735 | 1.8080 | ##### | 15.3278 | 4.9896E-53 | 9.5400E-52 |
| ENSG00000164296\|TIGD6 | 908.8495 | -2.1430 | ##### | ####### | 5.3372E-53 | 1.0198E-51 |
| ENSG00000001461\|NIPAL3 | 1308.6477 | -2.1517 | ##### | ####### | 5.5102E-53 | 1.0522E-51 |
| ENSG00000129680\|MAP7D3 | 4644.9713 | 1.9865 | ##### | 15.3173 | 5.8597E-53 | 1.1182E-51 |
| ENSG00000101190\|TCFL5 | 2335.2099 | 1.9252 | ##### | 15.3153 | 6.0388E-53 | 1.1516E-51 |
| ENSG00000132821\|VSTM2L | 912.0831 | -1.5885 | ##### | ####### | 6.0728E-53 | 1.1574E-51 |
| ENSG00000253910\|PCDHGB2 | 238.4192 | -5.3507 | ##### | ####### | 7.1624E-53 | 1.3642E-51 |
| ENSG00000107798\|LIPA | 5637.2896 | -1.8673 | ##### | ####### | 7.4627E-53 | 1.4205E-51 |
| ENSG00000117335\|CD46 | 12704.7649 | -1.5713 | ##### | ####### | 7.7233E-53 | 1.4691E-51 |
| ENSG00000163623\|NKX6-1 | 354.4555 | 3.3888 | ##### | 15.2880 | 9.1988E-53 | 1.7487E-51 |
| ENSG00000053918\|KCNQ1 | 179.2228 | 3.8633 | ##### | 15.2860 | 9.4815E-53 | 1.8013E-51 |
| ENSG00000256043\|CTSO | 262.8677 | -4.1405 | ##### | ####### | 1.0082E-52 | 1.9141E-51 |
| ENSG00000170088\|TMEM192 | 1461.3756 | -2.0682 | ##### | ####### | 1.0543E-52 | 2.0005E-51 |

| | | | | | | |
|---|---|---|---|---|---|---|
| ENSG00000151917\|BEND6 | 480.8585 | -2.1219 | ##### | ####### | 1.2335E-52 | 2.3389E-51 |
| ENSG00000165424\|ZCCHC24 | 955.9118 | -2.0387 | ##### | ####### | 1.5397E-52 | 2.9176E-51 |
| ENSG00000102996\|MMP15 | 2909.3303 | -1.8850 | ##### | ####### | 1.8387E-52 | 3.4822E-51 |
| ENSG00000204019\|CT83 | 362.0691 | -2.6547 | ##### | ####### | 1.8470E-52 | 3.4955E-51 |
| ENSG00000203930\|LINC00632 | 378.1447 | 3.8436 | ##### | 15.2381 | 1.9743E-52 | 3.7341E-51 |
| ENSG00000124532\|MRS2 | 3172.2751 | 1.3755 | ##### | 15.2332 | 2.1293E-52 | 4.0248E-51 |
| ENSG00000214944\|ARHGEF28 | 2430.2870 | -1.9838 | ##### | ####### | 2.3738E-52 | 4.4841E-51 |
| ENSG00000159263\|SIM2 | 459.0251 | -2.8967 | ##### | ####### | 2.4245E-52 | 4.5769E-51 |
| ENSG00000235109\|ZSCAN31 | 1188.4330 | ####### | ##### | ####### | 2.7542E-52 | 5.1961E-51 |
| ENSG00000170542\|SERPINB9 | 1249.8811 | -1.9985 | ##### | ####### | 3.0182E-52 | 5.6906E-51 |
| ENSG00000147324\|MFHAS1 | 7834.9791 | -1.8714 | ##### | ####### | 3.3597E-52 | 6.3304E-51 |
| ENSG00000175183\|CSRP2 | 1054.6083 | 2.2447 | ##### | 15.1918 | 4.0049E-52 | 7.5413E-51 |
| ENSG00000125775\|SDCBP2 | 125.5401 | -3.5564 | ##### | ####### | 5.3839E-52 | 1.0132E-50 |
| ENSG00000132740\|IGHMBP2 | 3535.0705 | 1.8311 | ##### | 15.1706 | 5.5405E-52 | 1.0420E-50 |
| ENSG00000293110 | 193.5919 | -3.1171 | ##### | ####### | 5.6337E-52 | 1.0589E-50 |
| ENSG00000149503\|INCENP | 6118.9866 | 1.8106 | ##### | 15.1592 | 6.5830E-52 | 1.2365E-50 |
| ENSG00000197555\|SIPA1L1 | 4045.5985 | 1.8631 | ##### | 15.1579 | 6.7143E-52 | 1.2604E-50 |
| ENSG00000166250\|CLMP | 7089.0086 | -1.8635 | ##### | ####### | 7.2715E-52 | 1.3641E-50 |
| ENSG00000159348\|CYB5R1 | 2040.3151 | -1.9116 | ##### | ####### | 7.6253E-52 | 1.4296E-50 |
| ENSG00000105928\|GSDME | 1130.3404 | -1.3561 | ##### | ####### | 8.4389E-52 | 1.5811E-50 |
| ENSG00000122359\|ANXA11 | 10251.4245 | -1.8846 | ##### | ####### | 8.6750E-52 | 1.6243E-50 |
| ENSG00000108622\|ICAM2 | 152.0838 | -4.8504 | ##### | ####### | 9.0283E-52 | 1.6894E-50 |
| ENSG00000274349\|ZNF658 | 270.3284 | -2.9386 | ##### | ####### | 9.3665E-52 | 1.7516E-50 |
| ENSG00000165280\|VCP | 71679.2641 | 1.4611 | ##### | 15.1349 | 9.5299E-52 | 1.7810E-50 |
| ENSG00000183856\|IQGAP3 | 7917.2487 | 1.8248 | ##### | 15.1325 | 9.8941E-52 | 1.8479E-50 |
| ENSG00000071054\|MAP4K4 | 14651.6584 | -1.8235 | ##### | ####### | 1.0474E-51 | 1.9551E-50 |
| ENSG00000165046\|LETM2 | 197.6712 | -2.9638 | ##### | ####### | 1.1686E-51 | 2.1798E-50 |
| ENSG00000120437\|ACAT2 | 1917.5476 | 5.2016 | ##### | 15.1162 | 1.2657E-51 | 2.3596E-50 |
| ENSG00000197818\|SLC9A8 | 1903.0210 | -1.3580 | ##### | ####### | 1.2720E-51 | 2.3699E-50 |
| ENSG00000204584 | 130.7761 | -3.7401 | ##### | ####### | 1.3228E-51 | 2.4629E-50 |
| ENSG00000172613\|RAD9A | 2874.2795 | 1.8241 | ##### | 15.1126 | 1.3384E-51 | 2.4905E-50 |
| ENSG00000154511\|DIPK1A | 1490.1399 | -2.1954 | ##### | ####### | 1.3800E-51 | 2.5662E-50 |
| ENSG00000165895\|ARHGAP42 | 1517.6183 | -2.1549 | ##### | ####### | 1.4114E-51 | 2.6231E-50 |
| ENSG00000088836\|SLC4A11 | 592.6597 | -2.2788 | ##### | ####### | 1.4247E-51 | 2.6461E-50 |
| ENSG00000144381\|HSPD1 | 78682.4501 | 1.2090 | ##### | 15.1072 | 1.4508E-51 | 2.6929E-50 |
| ENSG00000123388\|HOXC11 | 449.0537 | 2.4801 | ##### | 15.1035 | 1.5366E-51 | 2.8504E-50 |
| ENSG00000112964\|GHR | 306.2223 | 3.1670 | ##### | 15.0962 | 1.7160E-51 | 3.1811E-50 |
| ENSG00000180257\|ZNF816 | 387.4619 | -2.3863 | ##### | ####### | 1.9666E-51 | 3.6434E-50 |
| ENSG00000152778\|IFIT5 | 1023.4060 | -2.0437 | ##### | ####### | 1.9742E-51 | 3.6553E-50 |
| ENSG00000175768\|TOMM5 | 12525.3366 | 1.8332 | ##### | 15.0822 | 2.1215E-51 | 3.9256E-50 |
| ENSG00000250303\|LINC02762 | 217.2212 | -3.6385 | ##### | ####### | 2.2659E-51 | 4.1901E-50 |
| ENSG00000115425\|PECR | 1217.9715 | -2.1061 | ##### | ####### | 2.2810E-51 | 4.2154E-50 |
| ENSG00000261373\|VPS9D1-AS1 | 2511.1526 | 1.9097 | ##### | 15.0764 | 2.3144E-51 | 4.2745E-50 |
| ENSG00000115183\|TANC1 | 4048.2784 | -1.8396 | ##### | ####### | 2.4289E-51 | 4.4832E-50 |
| ENSG00000137124\|ALDH1B1 | 8198.4454 | 1.8178 | ##### | 15.0709 | 2.5176E-51 | 4.6441E-50 |
| ENSG00000117394\|SLC2A1 | 7219.3034 | -1.6268 | ##### | ####### | 2.5349E-51 | 4.6731E-50 |
| ENSG00000109738\|GLRB | 402.6126 | -2.6416 | ##### | ####### | 2.5748E-51 | 4.7438E-50 |
| ENSG00000099377\|HSD3B7 | 4119.3048 | -2.0636 | ##### | ####### | 2.7105E-51 | 4.9907E-50 |
| ENSG00000116852\|KIF21B | 412.3473 | -3.2118 | ##### | ####### | 2.8770E-51 | 5.2941E-50 |
| ENSG00000102384\|CENPI | 3549.2291 | 1.7875 | ##### | 15.0614 | 2.9074E-51 | 5.3467E-50 |
| ENSG00000156162\|DPY19L4 | 6242.7576 | -1.8337 | ##### | ####### | 2.9152E-51 | 5.3577E-50 |
| ENSG00000133135\|RNF128 | 687.6777 | 2.1438 | ##### | 15.0571 | 3.1020E-51 | 5.6976E-50 |
| ENSG00000007372\|PAX6 | 219.3740 | 5.8238 | ##### | 15.0546 | 3.2223E-51 | 5.9148E-50 |
| ENSG00000198453\|ZNF568 | 738.1552 | -2.2480 | ##### | ####### | 3.3947E-51 | 6.2275E-50 |
| ENSG00000124882\|EREG | 5071.7280 | 1.9756 | ##### | 15.0476 | 3.5802E-51 | 6.5638E-50 |
| ENSG00000119431\|HDHD3 | 480.7334 | 6.0289 | ##### | 15.0467 | 3.6295E-51 | 6.6500E-50 |
| ENSG00000240694\|PNMA2 | 4700.2503 | 1.8165 | ##### | 15.0449 | 3.7308E-51 | 6.8315E-50 |
| ENSG00000182890\|GLUD2 | 159.8734 | -3.1932 | ##### | ####### | 3.7594E-51 | 6.8795E-50 |
| ENSG00000099625\|CBARP | 1544.3231 | 1.9466 | ##### | 15.0118 | 6.1453E-51 | 1.1239E-49 |
| ENSG00000127990\|SGCE | 3858.4011 | -1.8772 | ##### | ####### | 6.6822E-51 | 1.2213E-49 |
| ENSG00000105245\|NUMBL | 4431.9173 | -1.8704 | ##### | ####### | 7.1393E-51 | 1.3041E-49 |
| ENSG00000156504\|PABIR2 | 5774.8178 | 1.8030 | ##### | 14.9953 | 7.8825E-51 | 1.4389E-49 |
| ENSG00000188906\|LRRK2 | 381.2318 | -2.7960 | ##### | ####### | 9.2546E-51 | 1.6884E-49 |

| | | | | | | |
|---|---|---|---|---|---|---|
| ENSG00000073803\|MAP3K13 | 1477.8792 | -2.0269 | ##### | ####### | 9.6284E-51 | 1.7555E-49 |
| ENSG00000176533\|GNG7 | 190.7469 | 3.6664 | ##### | 14.9757 | 1.0584E-50 | 1.9285E-49 |
| ENSG00000131069\|ACSS2 | 2506.9931 | -1.9164 | ##### | ####### | 1.1649E-50 | 2.1213E-49 |
| ENSG00000125726\|CD70 | 1057.9023 | 2.0818 | ##### | 14.9678 | 1.1921E-50 | 2.1696E-49 |
| ENSG00000124466\|LYPD3 | 222.9584 | -3.1914 | ##### | ####### | 1.2542E-50 | 2.2811E-49 |
| ENSG00000084731\|KIF3C | 3785.1689 | -1.9305 | ##### | ####### | 1.2928E-50 | 2.3499E-49 |
| ENSG00000274290\|H2BC6 | 375.9583 | 2.4615 | ##### | 14.9570 | 1.4031E-50 | 2.5489E-49 |
| ENSG00000165449\|SLC16A9 | 397.4587 | 2.5370 | ##### | 14.9523 | 1.5054E-50 | 2.7331E-49 |
| ENSG00000111224\|PARP11 | 474.4036 | 2.4526 | ##### | 14.9307 | 2.0794E-50 | 3.7729E-49 |
| ENSG00000141655\|TNFRSF11A | 300.7148 | 2.8036 | ##### | 14.9260 | 2.2319E-50 | 4.0471E-49 |
| ENSG00000127325\|BEST3 | 170.8833 | 3.2799 | ##### | 14.9172 | 2.5474E-50 | 4.6164E-49 |
| ENSG00000173621\|LRFN4 | 3999.9616 | -1.8194 | ##### | ####### | 2.5633E-50 | 4.6424E-49 |
| ENSG00000256940\|PPP1R14B-AS1 | 415.2897 | -2.4336 | ##### | ####### | 2.7592E-50 | 4.9942E-49 |
| ENSG00000183186\|C2CD4C | 654.6884 | 2.1594 | ##### | 14.9074 | 2.9517E-50 | 5.3393E-49 |
| ENSG00000177788\|RAB4A-AS1 | 1031.7877 | -1.9889 | ##### | ####### | 3.1238E-50 | 5.6472E-49 |
| ENSG00000095203\|EPB41L4B | 2616.9006 | -1.8716 | ##### | ####### | 3.7426E-50 | 6.7619E-49 |
| ENSG00000108582\|CPD | 8653.8877 | -1.7852 | ##### | ####### | 4.3206E-50 | 7.8014E-49 |
| ENSG00000136999\|CCN3 | 1268.8972 | 2.1122 | ##### | 14.8794 | 4.4827E-50 | 8.0892E-49 |
| ENSG00000115255\|REEP6 | 958.8322 | 2.2413 | ##### | 14.8675 | 5.3608E-50 | 9.6679E-49 |
| ENSG00000147119\|CHST7 | 467.8854 | -2.2918 | ##### | ####### | 6.2242E-50 | 1.1218E-48 |
| ENSG00000156017\|CARNMT1 | 2603.2866 | -1.1728 | ##### | ####### | 7.0830E-50 | 1.2758E-48 |
| ENSG00000153885\|KCTD15 | 5773.6513 | -1.4482 | ##### | ####### | 7.3497E-50 | 1.3231E-48 |
| ENSG00000104722\|NEFM | 198.6220 | -3.2654 | ##### | ####### | 7.6031E-50 | 1.3679E-48 |
| ENSG00000141682\|PMAIP1 | 1129.3580 | 2.2260 | ##### | 14.8407 | 7.9929E-50 | 1.4371E-48 |
| ENSG00000033327\|GAB2 | 668.5497 | -2.3496 | ##### | ####### | 8.2591E-50 | 1.4841E-48 |
| ENSG00000162415\|ZSWIM5 | 541.6206 | -2.3510 | ##### | ####### | 8.3137E-50 | 1.4930E-48 |
| ENSG00000057704\|TMCC3 | 553.3551 | -2.5531 | ##### | ####### | 8.4596E-50 | 1.5183E-48 |
| ENSG00000133104\|SPART | 3296.3767 | -1.8747 | ##### | ####### | 8.5774E-50 | 1.5385E-48 |
| ENSG00000080986\|NDC80 | 4360.1615 | 1.7675 | ##### | 14.8338 | 8.8626E-50 | 1.5887E-48 |
| ENSG00000188549\|CCDC9B | 1528.6453 | -1.9550 | ##### | ####### | 9.0589E-50 | 1.6229E-48 |
| ENSG00000243701\|DUBR | 816.2419 | 2.0563 | ##### | 14.8321 | 9.0913E-50 | 1.6278E-48 |
| ENSG00000143398\|PIP5K1A | 15371.1423 | 1.4046 | ##### | 14.8253 | 1.0060E-49 | 1.8002E-48 |
| ENSG00000079308\|TNS1 | 292.5696 | -2.8915 | ##### | ####### | 1.0337E-49 | 1.8486E-48 |
| ENSG00000214402\|LCNL1 | 322.9603 | -2.6171 | ##### | ####### | 1.0356E-49 | 1.8509E-48 |
| ENSG00000183578\|TNFAIP8L3 | 184.8639 | -3.2229 | ##### | ####### | 1.0697E-49 | 1.9106E-48 |
| ENSG00000154102\|C16orf74 | 477.0966 | -2.2593 | ##### | ####### | 1.0708E-49 | 1.9115E-48 |
| ENSG00000167555\|ZNF528 | 266.8233 | -2.9551 | ##### | ####### | 1.1591E-49 | 2.0679E-48 |
| ENSG00000073910\|FRY | 407.7602 | 2.7377 | ##### | 14.8125 | 1.2155E-49 | 2.1661E-48 |
| ENSG00000156467\|UQCRB | 9298.0453 | -1.7595 | ##### | ####### | 1.2156E-49 | 2.1661E-48 |
| ENSG00000114450\|GNB4 | 6123.0269 | -1.7613 | ##### | ####### | 1.4518E-49 | 2.5855E-48 |
| ENSG00000115221\|ITGB6 | 161.3883 | -4.2690 | ##### | ####### | 1.4941E-49 | 2.6591E-48 |
| ENSG00000166387\|PPFIBP2 | 741.9421 | -2.1077 | ##### | ####### | 1.5918E-49 | 2.8314E-48 |
| ENSG00000145912\|NHP2 | 17528.8239 | 1.7657 | ##### | 14.7845 | 1.8431E-49 | 3.2765E-48 |
| ENSG00000114480\|GBE1 | 2555.2191 | -1.8982 | ##### | ####### | 2.1870E-49 | 3.8855E-48 |
| ENSG00000178904\|DPY19L3 | 1905.5040 | -1.8904 | ##### | ####### | 2.5810E-49 | 4.5827E-48 |
| ENSG00000226711\|FAM66C | 262.4567 | 2.9358 | ##### | 14.7602 | 2.6442E-49 | 4.6921E-48 |
| ENSG00000103202\|NME4 | 7534.1467 | -1.7846 | ##### | ####### | 2.6935E-49 | 4.7768E-48 |
| ENSG00000171903\|CYP4F11 | ########## | ####### | ##### | ####### | 2.8614E-49 | 5.0715E-48 |
| ENSG00000134897\|BIVM | 1229.5351 | -2.0188 | ##### | ####### | 2.9131E-49 | 5.1602E-48 |
| ENSG00000291106\|SNHG33 | 614.5595 | -2.4228 | ##### | ####### | 3.3805E-49 | 5.9845E-48 |
| ENSG00000160953\|PWWP3A | 8791.5852 | 1.4785 | ##### | 14.7413 | 3.4998E-49 | 6.1920E-48 |
| ENSG00000163659\|TIPARP | 9543.2653 | 1.8197 | ##### | 14.7284 | 4.2376E-49 | 7.4930E-48 |
| ENSG00000196372\|ASB13 | 1797.6749 | -1.8160 | ##### | ####### | 4.3854E-49 | 7.7496E-48 |
| ENSG00000288075 | 139.1457 | 4.4790 | ##### | 14.7251 | 4.4449E-49 | 7.8503E-48 |
| ENSG00000177156\|TALDO1 | 13072.8959 | -1.7332 | ##### | ####### | 4.9350E-49 | 8.7107E-48 |
| ENSG00000176444\|CLK2 | 4162.8408 | 1.6042 | ##### | 14.7168 | 5.0313E-49 | 8.8754E-48 |
| ENSG00000103855\|CD276 | 3834.3227 | -1.8441 | ##### | ####### | 6.2235E-49 | 1.0972E-47 |
| ENSG00000104823\|ECH1 | 13125.4040 | -1.8056 | ##### | ####### | 6.7926E-49 | 1.1968E-47 |
| ENSG00000125841\|NRSN2 | 3534.6107 | -1.4471 | ##### | ####### | 7.1715E-49 | 1.2628E-47 |
| ENSG00000130958\|SLC35D2 | 1243.4046 | -2.0705 | ##### | ####### | 7.3253E-49 | 1.2892E-47 |
| ENSG00000135899\|SP110 | 629.8263 | -2.0840 | ##### | ####### | 8.5362E-49 | 1.5014E-47 |
| ENSG00000233237\|LINC00472 | 1084.9336 | 2.1480 | ##### | 14.6749 | 9.3303E-49 | 1.6401E-47 |
| ENSG00000136367\|ZFHX2 | 476.8635 | 2.2968 | ##### | 14.6722 | 9.7134E-49 | 1.7064E-47 |

| | | | | | | |
|---|---|---|---|---|---|---|
| ENSG00000088387\|DOCK9 | 2739.3370 | -1.8684 | ##### | ####### | 1.0778E-48 | 1.8924E-47 |
| ENSG00000143569\|UBAP2L | 22335.3391 | 1.0671 | ##### | 14.6642 | 1.0935E-48 | 1.9188E-47 |
| ENSG00000129451\|KLK10 | 305.3116 | -2.5746 | ##### | ####### | 1.1204E-48 | 1.9648E-47 |
| ENSG00000139200\|PIANP | 232.2620 | 4.1246 | ##### | 14.6604 | 1.1556E-48 | 2.0253E-47 |
| ENSG00000131018\|SYNE1 | 1915.3866 | -2.0588 | ##### | ####### | 1.1917E-48 | 2.0873E-47 |
| ENSG00000183421\|RIPK4 | 4272.8789 | -1.8704 | ##### | ####### | 1.2337E-48 | 2.1597E-47 |
| ENSG00000213214\|ARHGEF35 | 700.2979 | -2.1619 | ##### | ####### | 1.2597E-48 | 2.2038E-47 |
| ENSG00000226608\|FTLP3 | 248.3559 | -3.4808 | ##### | ####### | 1.2684E-48 | 2.2178E-47 |
| ENSG00000090776\|EFNB1 | 1437.3866 | 1.6686 | ##### | 14.6507 | 1.3343E-48 | 2.3317E-47 |
| ENSG00000113580\|NR3C1 | 14138.4201 | -1.7090 | ##### | ####### | 1.3931E-48 | 2.4330E-47 |
| ENSG00000228801\|PCMTD1-DT | 507.7737 | -2.2502 | ##### | ####### | 1.4317E-48 | 2.4990E-47 |
| ENSG00000020577\|SAMD4A | 2976.0182 | 1.9086 | ##### | 14.6454 | 1.4412E-48 | 2.5140E-47 |
| ENSG00000137474\|MYO7A | 323.0761 | 2.5661 | ##### | 14.6402 | 1.5567E-48 | 2.7140E-47 |
| ENSG00000108813\|DLX4 | 590.5305 | 2.4507 | ##### | 14.6401 | 1.5587E-48 | 2.7158E-47 |
| ENSG00000128602\|SMO | 1256.5621 | 1.9675 | ##### | 14.6399 | 1.5631E-48 | 2.7221E-47 |
| ENSG00000172725\|CORO1B | 9025.4625 | 1.7440 | ##### | 14.6374 | 1.6212E-48 | 2.8199E-47 |
| ENSG00000114315\|HES1 | 2628.6478 | 1.8581 | ##### | 14.6374 | 1.6205E-48 | 2.8199E-47 |
| ENSG00000015413\|DPEP1 | 111.4378 | -7.0019 | ##### | ####### | 1.6534E-48 | 2.8741E-47 |
| ENSG00000163872\|YEATS2 | 4075.3108 | -1.7918 | ##### | ####### | 1.6840E-48 | 2.9257E-47 |
| ENSG00000054967\|RELT | 2530.1440 | 1.8409 | ##### | 14.6215 | 2.0495E-48 | 3.5586E-47 |
| ENSG00000151690\|MFSD6 | 2477.2107 | -1.8339 | ##### | ####### | 2.2902E-48 | 3.9743E-47 |
| ENSG00000293477\|CRYZL2P | 480.5228 | -2.3831 | ##### | ####### | 2.3548E-48 | 4.0816E-47 |
| ENSG00000291079 | 480.4827 | -2.3548 | ##### | ####### | 2.3535E-48 | 4.0816E-47 |
| ENSG00000154380\|ENAH | 19701.0766 | -1.7155 | ##### | ####### | 2.3714E-48 | 4.1080E-47 |
| ENSG00000168724\|DNAJC21 | 9112.6281 | 1.1305 | ##### | 14.6102 | 2.4196E-48 | 4.1890E-47 |
| ENSG00000189060\|H1-0 | 18185.9569 | 1.7711 | ##### | 14.6013 | 2.7544E-48 | 4.7660E-47 |
| ENSG00000286388 | 636.8765 | -2.2204 | ##### | ####### | 2.9951E-48 | 5.1795E-47 |
| ENSG00000158864\|NDUFS2 | 20251.0819 | 1.5493 | ##### | 14.5949 | 3.0282E-48 | 5.2336E-47 |
| ENSG00000291139\|DPY19L2P2 | 955.9794 | 2.1249 | ##### | 14.5826 | 3.6235E-48 | 6.2588E-47 |
| ENSG00000116260\|QSOX1 | 10940.5111 | -1.3085 | ##### | ####### | 3.7575E-48 | 6.4866E-47 |
| ENSG00000111077\|TNS2 | 1933.5714 | -1.9414 | ##### | ####### | 3.9485E-48 | 6.8124E-47 |
| ENSG00000116016\|EPAS1 | 23024.4022 | -1.6224 | ##### | ####### | 4.0603E-48 | 7.0012E-47 |
| ENSG00000105997\|HOXA3 | 662.1678 | 2.1379 | ##### | 14.5687 | 4.4399E-48 | 7.6514E-47 |
| ENSG00000286931 | 129.9286 | 3.2981 | ##### | 14.5668 | 4.5676E-48 | 7.8669E-47 |
| ENSG00000129103\|SUMF2 | 11814.7117 | -1.6060 | ##### | ####### | 4.5832E-48 | 7.8892E-47 |
| ENSG00000121753\|ADGRB2 | 2890.7029 | -1.7288 | ##### | ####### | 5.0431E-48 | 8.6758E-47 |
| ENSG00000172716\|SLFN11 | 3931.1492 | -1.7848 | ##### | ####### | 5.0697E-48 | 8.7165E-47 |
| ENSG00000235590\|GNAS-AS1 | 328.3152 | 4.1162 | ##### | 14.5595 | 5.0836E-48 | 8.7355E-47 |
| ENSG00000111641\|NOP2 | 21368.1414 | 1.7095 | ##### | 14.5592 | 5.1088E-48 | 8.7736E-47 |
| ENSG00000101811\|CSTF2 | 4852.6725 | 1.0462 | ##### | 14.5583 | 5.1737E-48 | 8.8800E-47 |
| ENSG00000116544\|DLGAP3 | 300.0233 | 2.8921 | ##### | 14.5552 | 5.4144E-48 | 9.2879E-47 |
| ENSG00000269825 | 292.7966 | -2.6361 | ##### | ####### | 6.0010E-48 | 1.0288E-46 |
| ENSG00000138768\|USO1 | 7704.1689 | -1.7447 | ##### | ####### | 6.0527E-48 | 1.0371E-46 |
| ENSG00000102181\|CD99L2 | 3521.4322 | -1.7699 | ##### | ####### | 6.6270E-48 | 1.1348E-46 |
| ENSG00000197497\|ZNF665 | 951.8936 | ####### | ##### | ####### | 7.3785E-48 | 1.2628E-46 |
| ENSG00000150455\|TIRAP | 780.7754 | -2.0376 | ##### | ####### | 8.0727E-48 | 1.3808E-46 |
| ENSG00000092096\|SLC22A17 | 1012.2672 | 2.0194 | ##### | 14.5267 | 8.2042E-48 | 1.4025E-46 |
| ENSG00000107338\|SHB | 8318.3369 | 1.8638 | ##### | 14.5254 | 8.3675E-48 | 1.4296E-46 |
| ENSG00000115738\|ID2 | 6448.2193 | 1.8558 | ##### | 14.5230 | 8.6676E-48 | 1.4800E-46 |
| ENSG00000253250\|C8orf88 | 678.5287 | -8.6218 | ##### | ####### | 9.1128E-48 | 1.5552E-46 |
| ENSG00000153786\|ZDHHC7 | 9015.9180 | -1.4944 | ##### | ####### | 1.0530E-47 | 1.7959E-46 |
| ENSG00000119946\|CNNM1 | 1282.1385 | -1.9041 | ##### | ####### | 1.1500E-47 | 1.9603E-46 |
| ENSG00000165757\|JCAD | 2105.8253 | -1.8453 | ##### | ####### | 1.2932E-47 | 2.2032E-46 |
| ENSG00000213694\|S1PR3 | 3846.5082 | -1.7745 | ##### | ####### | 1.3060E-47 | 2.2237E-46 |
| ENSG00000169851\|PCDH7 | 7422.3691 | 1.7446 | ##### | 14.4946 | 1.3106E-47 | 2.2303E-46 |
| ENSG00000214078\|CPNE1 | 18526.3180 | 1.7618 | ##### | 14.4901 | 1.3992E-47 | 2.3796E-46 |
| ENSG00000060491\|OGFR | 8193.5480 | 1.7387 | ##### | 14.4887 | 1.4283E-47 | 2.4278E-46 |
| ENSG00000118523\|CCN2 | 8795.7599 | 1.1782 | ##### | 14.4876 | 1.4514E-47 | 2.4657E-46 |
| ENSG00000165097\|KDM1B | 3458.4941 | 1.6990 | ##### | 14.4850 | 1.5079E-47 | 2.5601E-46 |
| ENSG00000214491\|SEC14L6 | 147.4260 | -3.4806 | ##### | ####### | 1.7276E-47 | 2.9315E-46 |
| ENSG00000140403\|DNAJA4 | 270.4363 | -3.7005 | ##### | ####### | 1.9236E-47 | 3.2622E-46 |
| ENSG00000115226\|FNDC4 | 517.1454 | -2.2332 | ##### | ####### | 2.0754E-47 | 3.5177E-46 |
| ENSG00000171067\|C11orf24 | 6962.8573 | 1.7760 | ##### | 14.4614 | 2.1249E-47 | 3.5995E-46 |

| | | | | | | |
|---|---|---|---|---|---|---|
| ENSG00000127946\|HIP1 | 5559.9465 | -1.7747 | ##### | ####### | 2.7565E-47 | 4.6669E-46 |
| ENSG00000078269\|SYNJ2 | 6692.2210 | 1.7255 | ##### | 14.4410 | 2.8567E-47 | 4.8338E-46 |
| ENSG00000084733\|RAB10 | 21478.8671 | -0.8601 | ##### | ####### | 3.3726E-47 | 5.7034E-46 |
| ENSG00000272913\|LINC03052 | 740.1851 | 2.3480 | ##### | 14.4266 | 3.5179E-47 | 5.9458E-46 |
| ENSG00000197345\|MRPL21 | 6011.5820 | 1.7725 | ##### | 14.4245 | 3.6273E-47 | 6.1272E-46 |
| ENSG00000087586\|AURKA | 13861.1601 | 1.5890 | ##### | 14.4235 | 3.6793E-47 | 6.2116E-46 |
| ENSG00000165283\|STOML2 | 20283.1194 | 1.7396 | ##### | 14.4214 | 3.7938E-47 | 6.4013E-46 |
| ENSG00000107371\|EXOSC3 | 4671.4563 | 1.7920 | ##### | 14.4189 | 3.9338E-47 | 6.6337E-46 |
| ENSG00000099957\|P2RX6 | 197.9547 | -3.5201 | ##### | ####### | 4.2293E-47 | 7.1281E-46 |
| ENSG00000168447\|SCNN1B | 213.3693 | -3.3330 | ##### | ####### | 4.5096E-47 | 7.5961E-46 |
| ENSG00000162777\|DENND2D | 627.2616 | -2.2432 | ##### | ####### | 5.0916E-47 | 8.5716E-46 |
| ENSG00000137962\|ARHGAP29 | 7608.4455 | -1.8150 | ##### | ####### | 5.3530E-47 | 9.0067E-46 |
| ENSG00000055147\|FAM114A2 | 3304.9418 | -1.8571 | ##### | ####### | 5.3772E-47 | 9.0423E-46 |
| ENSG00000129353\|SLC44A2 | 3332.3908 | -1.7677 | ##### | ####### | 5.7255E-47 | 9.6224E-46 |
| ENSG00000111962\|UST | 556.4035 | 2.2063 | ##### | 14.3866 | 6.2764E-47 | 1.0542E-45 |
| ENSG00000204673\|AKT1S1 | 9859.6025 | -1.3695 | ##### | ####### | 6.7208E-47 | 1.1283E-45 |
| ENSG00000106484\|MEST | 4943.1339 | 1.4967 | ##### | 14.3796 | 6.9477E-47 | 1.1657E-45 |
| ENSG00000233184\|DPH5-DT | 758.4735 | -2.2066 | ##### | ####### | 7.3737E-47 | 1.2365E-45 |
| ENSG00000117151\|CTBS | 2188.5933 | -1.7749 | ##### | ####### | 8.0644E-47 | 1.3515E-45 |
| ENSG00000289810 | 248.2039 | 3.0158 | ##### | 14.3672 | 8.3181E-47 | 1.3933E-45 |
| ENSG00000130699\|TAF4 | 3434.5108 | 1.7450 | ##### | 14.3636 | 8.7562E-47 | 1.4658E-45 |
| ENSG00000135940\|COX5B | 10421.8606 | 1.7477 | ##### | 14.3607 | 9.1359E-47 | 1.5285E-45 |
| ENSG00000214826\|DDX12P | 1286.2745 | 1.9063 | ##### | 14.3601 | 9.2105E-47 | 1.5402E-45 |
| ENSG00000196275\|GTF2IRD2 | 270.5337 | -2.7091 | ##### | ####### | 9.4685E-47 | 1.5824E-45 |
| ENSG00000277149\|TYW1B | 358.0143 | 2.9150 | ##### | 14.3545 | 9.9859E-47 | 1.6679E-45 |
| ENSG00000130669\|PAK4 | 9436.2348 | -1.0920 | ##### | ####### | 9.9929E-47 | 1.6682E-45 |
| ENSG00000128165\|ADM2 | 310.3555 | -2.3556 | ##### | ####### | 1.0391E-46 | 1.7336E-45 |
| ENSG00000155816\|FMN2 | 309.5455 | 2.6345 | ##### | 14.3492 | 1.0776E-46 | 1.7969E-45 |
| ENSG00000080573\|COL5A3 | 124.3188 | -3.7685 | ##### | ####### | 1.0831E-46 | 1.8051E-45 |
| ENSG00000092758\|COL9A3 | 283.9289 | 2.5850 | ##### | 14.3469 | 1.1146E-46 | 1.8565E-45 |
| ENSG00000162650\|ATXN7L2 | 1431.3330 | 1.8701 | ##### | 14.3436 | 1.1692E-46 | 1.9464E-45 |
| ENSG00000109107\|ALDOC | 275.2218 | -3.0553 | ##### | ####### | 1.2134E-46 | 2.0189E-45 |
| ENSG00000254093\|PINX1 | 3410.3306 | -1.8415 | ##### | ####### | 1.2193E-46 | 2.0275E-45 |
| ENSG00000118276\|B4GALT6 | 4448.0055 | 1.8338 | ##### | 14.3320 | 1.3818E-46 | 2.2965E-45 |
| ENSG00000185885\|IFITM1 | 161.7438 | -2.9672 | ##### | ####### | 1.4480E-46 | 2.4051E-45 |
| ENSG00000163975\|MELTF | 4095.7749 | -1.3879 | ##### | ####### | 1.6244E-46 | 2.6966E-45 |
| ENSG00000124191\|TOX2 | 708.1884 | 2.3643 | ##### | 14.3139 | 1.7911E-46 | 2.9717E-45 |
| ENSG00000168040\|FADD | 8887.8294 | 1.7140 | ##### | 14.3130 | 1.8141E-46 | 3.0083E-45 |
| ENSG00000184792\|OSBP2 | 681.9090 | -2.1539 | ##### | ####### | 1.9377E-46 | 3.2113E-45 |
| ENSG00000107185\|RGP1 | 12151.1221 | 1.7178 | ##### | 14.2950 | 2.3510E-46 | 3.8943E-45 |
| ENSG00000074219\|TEAD2 | 3813.7393 | -1.7787 | ##### | ####### | 2.6204E-46 | 4.3380E-45 |
| ENSG00000129245\|FXR2 | 10018.2026 | 0.9113 | ##### | 14.2824 | 2.8175E-46 | 4.6617E-45 |
| ENSG00000259884\|NR4A1AS | 267.7382 | 2.7905 | ##### | 14.2770 | 3.0447E-46 | 5.0350E-45 |
| ENSG00000074800\|ENO1 | ########## | -1.4112 | ##### | ####### | 3.4700E-46 | 5.7351E-45 |
| ENSG00000170379\|TCAF2 | 148.2326 | 3.1064 | ##### | 14.2553 | 4.1538E-46 | 6.8614E-45 |
| ENSG00000149289\|ZC3H12C | 1081.4751 | -2.0077 | ##### | ####### | 4.1865E-46 | 6.9115E-45 |
| ENSG00000198729\|PPP1R14C | 555.0654 | 2.0754 | ##### | 14.2535 | 4.2616E-46 | 7.0316E-45 |
| ENSG00000089177\|KIF16B | 1808.5212 | -1.8284 | ##### | ####### | 4.2982E-46 | 7.0880E-45 |
| ENSG00000013364\|MVP | 12918.4024 | -1.6102 | ##### | ####### | 4.5802E-46 | 7.5490E-45 |
| ENSG00000183287\|CCBE1 | 462.7112 | 2.2395 | ##### | 14.2477 | 4.6308E-46 | 7.6282E-45 |
| ENSG00000246695\|RASSF8-AS1 | 1622.6792 | 1.9018 | ##### | 14.2413 | 5.0755E-46 | 8.3560E-45 |
| ENSG00000184661\|CDCA2 | 3295.3568 | 1.7330 | ##### | 14.2271 | 6.2209E-46 | 1.0236E-44 |
| ENSG00000130935\|NOL11 | 11487.6886 | 0.9767 | ##### | 14.2230 | 6.5932E-46 | 1.0843E-44 |
| ENSG00000075651\|PLD1 | 1524.6345 | -1.9277 | ##### | ####### | 6.7291E-46 | 1.1060E-44 |
| ENSG00000137492\|THAP12 | 6383.7538 | 1.3344 | ##### | 14.2046 | 8.5742E-46 | 1.4085E-44 |
| ENSG00000068903\|SIRT2 | 1378.6913 | -1.8520 | ##### | ####### | 8.8054E-46 | 1.4457E-44 |
| ENSG00000103723\|AP3B2 | 490.7303 | 2.4387 | ##### | 14.1983 | 9.3873E-46 | 1.5404E-44 |
| ENSG00000167657\|DAPK3 | 10143.1642 | 1.7526 | ##### | 14.1973 | 9.5222E-46 | 1.5617E-44 |
| ENSG00000088002\|SULT2B1 | 337.3167 | -3.4802 | ##### | ####### | 9.5697E-46 | 1.5686E-44 |
| ENSG00000132481\|TRIM47 | 6851.5841 | -1.7525 | ##### | ####### | 9.9103E-46 | 1.6235E-44 |
| ENSG00000112992\|NNT | 1963.8977 | -8.5793 | ##### | ####### | 1.0015E-45 | 1.6398E-44 |
| ENSG00000110066\|KMT5B | 3858.7577 | 1.5864 | ##### | 14.1719 | 1.3682E-45 | 2.2389E-44 |
| ENSG00000175634\|RPS6KB2 | 7600.5825 | 1.7507 | ##### | 14.1612 | 1.5922E-45 | 2.6041E-44 |

| | | | | | | |
|---|---|---|---|---|---|---|
| ENSG00000119042\|SATB2 | 2004.7309 | 1.9046 | ##### | 14.1564 | 1.7062E-45 | 2.7890E-44 |
| ENSG00000253729\|PRKDC | 37519.4402 | 1.7249 | ##### | 14.1548 | 1.7449E-45 | 2.8507E-44 |
| ENSG00000081189\|MEF2C | 487.5991 | 2.1800 | ##### | 14.1512 | 1.8348E-45 | 2.9959E-44 |
| ENSG00000253676\|TAGLN2P1 | 234.2482 | 2.5699 | ##### | 14.1504 | 1.8559E-45 | 3.0287E-44 |
| ENSG00000170214\|ADRA1B | 763.6905 | 2.0537 | ##### | 14.1490 | 1.8953E-45 | 3.0915E-44 |
| ENSG00000231607\|DLEU2 | 1219.1470 | 2.1112 | ##### | 14.1482 | 1.9172E-45 | 3.1255E-44 |
| ENSG00000070214\|SLC44A1 | 8783.6484 | -1.6948 | ##### | ####### | 1.9939E-45 | 3.2487E-44 |
| ENSG00000105755\|ETHE1 | 2597.5546 | -1.7958 | ##### | ####### | 2.0284E-45 | 3.3031E-44 |
| ENSG00000105854\|PON2 | 13769.2817 | -1.7521 | ##### | ####### | 2.1700E-45 | 3.5317E-44 |
| ENSG00000221923\|ZNF880 | 770.0689 | -1.9036 | ##### | ####### | 2.2655E-45 | 3.6851E-44 |
| ENSG00000160447\|PKN3 | 1308.4310 | -1.8698 | ##### | ####### | 2.5798E-45 | 4.1941E-44 |
| ENSG00000180998\|GPR137C | 316.0757 | 2.7676 | ##### | 14.1250 | 2.6659E-45 | 4.3317E-44 |
| ENSG00000205002\|AARD | 182.1112 | 3.1063 | ##### | 14.1239 | 2.7071E-45 | 4.3962E-44 |
| ENSG00000143469\|SYT14 | 638.9543 | 3.3037 | ##### | 14.1194 | 2.8853E-45 | 4.6831E-44 |
| ENSG00000136813\|ECPAS | 9470.7400 | -1.6468 | ##### | ####### | 3.1687E-45 | 5.1403E-44 |
| ENSG00000110057\|UNC93B1 | 6631.9175 | -1.5711 | ##### | ####### | 3.4198E-45 | 5.5447E-44 |
| ENSG00000070610\|GBA2 | 5591.6265 | 1.7408 | ##### | 14.1071 | 3.4342E-45 | 5.5650E-44 |
| ENSG00000126458\|RRAS | 6672.7543 | -1.7325 | ##### | ####### | 3.5513E-45 | 5.7516E-44 |
| ENSG00000250120\|PCDHA10 | 203.2941 | -2.7677 | ##### | ####### | 3.7095E-45 | 6.0045E-44 |
| ENSG00000183258\|DDX41 | 6820.2880 | 1.1721 | ##### | 14.1016 | 3.7146E-45 | 6.0096E-44 |
| ENSG00000241878\|PISD | 2854.0384 | -1.8930 | ##### | ####### | 3.7214E-45 | 6.0173E-44 |
| ENSG00000133639\|BTG1 | 5701.7075 | -1.7406 | ##### | ####### | 3.9202E-45 | 6.3353E-44 |
| ENSG00000010310\|GIPR | 254.5770 | -2.7229 | ##### | ####### | 3.9630E-45 | 6.4010E-44 |
| ENSG00000170365\|SMAD1 | 1216.3156 | -1.9562 | ##### | ####### | 3.9756E-45 | 6.4179E-44 |
| ENSG00000078237\|TIGAR | 7623.7392 | 1.8086 | ##### | 14.0966 | 3.9845E-45 | 6.4288E-44 |
| ENSG00000101596\|SMCHD1 | 15441.5539 | 1.6946 | ##### | 14.0936 | 4.1611E-45 | 6.7101E-44 |
| ENSG00000229043\|ZFAND2A-DT | 791.4975 | -1.9740 | ##### | ####### | 4.6502E-45 | 7.4947E-44 |
| ENSG00000173614\|NMNAT1 | 1554.8629 | -1.9754 | ##### | ####### | 4.7152E-45 | 7.5955E-44 |
| ENSG00000143036\|SLC44A3 | 687.7522 | -2.0574 | ##### | ####### | 5.0429E-45 | 8.1190E-44 |
| ENSG00000260877 | 130.0528 | -2.7084 | ##### | ####### | 5.4391E-45 | 8.7520E-44 |
| ENSG00000184602\|SNN | 951.8894 | -2.1025 | ##### | ####### | 5.5818E-45 | 8.9769E-44 |
| ENSG00000161791\|FMNL3 | 3492.1575 | 1.8295 | ##### | 14.0726 | 5.5978E-45 | 8.9977E-44 |
| ENSG00000174938\|SEZ6L2 | 4430.7704 | -1.1533 | ##### | ####### | 5.6611E-45 | 9.0946E-44 |
| ENSG00000109576\|AADAT | 413.3357 | -2.4532 | ##### | ####### | 5.7250E-45 | 9.1923E-44 |
| ENSG00000134574\|DDB2 | 2834.5061 | 1.6979 | ##### | 14.0709 | 5.7327E-45 | 9.1997E-44 |
| ENSG00000189171\|S100A13 | 1058.8396 | -2.0078 | ##### | ####### | 5.8448E-45 | 9.3746E-44 |
| ENSG00000257524 | 1051.2908 | -3.4148 | ##### | ####### | 6.1295E-45 | 9.8259E-44 |
| ENSG00000254827\|SLC22A18AS | 177.1931 | -4.7019 | ##### | ####### | 6.1479E-45 | 9.8502E-44 |
| ENSG00000182118\|FAM89A | 726.9088 | -2.0554 | ##### | ####### | 6.1818E-45 | 9.8991E-44 |
| ENSG00000170190\|SLC16A5 | 1601.1899 | -1.8905 | ##### | ####### | 7.0161E-45 | 1.1229E-43 |
| ENSG00000230606\|APPAT | 682.6594 | 2.4436 | ##### | 14.0527 | 7.4170E-45 | 1.1864E-43 |
| ENSG00000160753\|RUSC1 | 6258.5249 | 1.7574 | ##### | 14.0445 | 8.3238E-45 | 1.3308E-43 |
| ENSG00000183873\|SCN5A | 88.0654 | 5.7931 | ##### | 14.0419 | 8.6376E-45 | 1.3802E-43 |
| ENSG00000164023\|SGMS2 | 3348.1983 | -1.7333 | ##### | ####### | 8.7922E-45 | 1.4042E-43 |
| ENSG00000141736\|ERBB2 | 4730.7330 | -1.6776 | ##### | ####### | 8.8170E-45 | 1.4074E-43 |
| ENSG00000076356\|PLXNA2 | 1218.1306 | -1.8498 | ##### | ####### | 9.4235E-45 | 1.5034E-43 |
| ENSG00000203697\|CAPN8 | 84.1757 | -6.2599 | ##### | ####### | 9.6813E-45 | 1.5437E-43 |
| ENSG00000132510\|KDM6B | 3508.3500 | 1.7095 | ##### | 14.0284 | 1.0444E-44 | 1.6644E-43 |
| ENSG00000164880\|INTS1 | 27601.5641 | -1.6476 | ##### | ####### | 1.0504E-44 | 1.6730E-43 |
| ENSG00000115290\|GRB14 | 1554.1193 | -1.8415 | ##### | ####### | 1.0951E-44 | 1.7433E-43 |
| ENSG00000169169\|CPT1C | 1190.9431 | 1.8543 | ##### | 14.0233 | 1.1228E-44 | 1.7865E-43 |
| ENSG00000090339\|ICAM1 | 13034.8798 | -1.6675 | ##### | ####### | 1.1607E-44 | 1.8457E-43 |
| ENSG00000274847\|MAFIP | 1576.8393 | 1.9207 | ##### | 14.0204 | 1.1694E-44 | 1.8587E-43 |
| ENSG00000139329\|LUM | 268.8533 | 2.8112 | ##### | 14.0040 | 1.4729E-44 | 2.3399E-43 |
| ENSG00000274333 | 470.5400 | -1.5053 | ##### | ####### | 1.5769E-44 | 2.5036E-43 |
| ENSG00000034677\|RNF19A | 2957.6973 | -1.7678 | ##### | ####### | 1.6056E-44 | 2.5479E-43 |
| ENSG00000181982\|CCDC149 | 531.3511 | -2.0487 | ##### | ####### | 1.7442E-44 | 2.7664E-43 |
| ENSG00000128342\|LIF | 7045.9293 | 1.7769 | ##### | 13.9912 | 1.7646E-44 | 2.7973E-43 |
| ENSG00000163755\|HPS3 | 5917.6218 | -1.6653 | ##### | ####### | 1.9550E-44 | 3.0974E-43 |
| ENSG00000158023\|CFAP251 | 277.6723 | 2.7134 | ##### | 13.9833 | 1.9717E-44 | 3.1221E-43 |
| ENSG00000112541\|PDE10A | 4384.3782 | -1.7155 | ##### | ####### | 1.9880E-44 | 3.1463E-43 |
| ENSG00000085415\|SEH1L | 11202.1383 | 1.7245 | ##### | 13.9820 | 2.0069E-44 | 3.1745E-43 |
| ENSG00000087842\|PIR | 2690.1099 | -1.8172 | ##### | ####### | 2.0574E-44 | 3.2528E-43 |

| | | | | | | |
|---|---|---|---|---|---|---|
| ENSG00000291050\|TEKT4P2 | 2329.2326 | 1.8358 | ##### | 13.9784 | 2.1106E-44 | 3.3351E-43 |
| ENSG00000110987\|BCL7A | 1183.6603 | -1.9093 | ##### | ####### | 2.1464E-44 | 3.3899E-43 |
| ENSG00000137210\|TMEM14B | 8546.2123 | 1.7175 | ##### | 13.9769 | 2.1563E-44 | 3.4037E-43 |
| ENSG00000031691\|CENPQ | 1082.2403 | 1.7794 | ##### | 13.9755 | 2.2001E-44 | 3.4710E-43 |
| ENSG00000013588\|GPRC5A | 38469.9073 | -1.7239 | ##### | ####### | 2.2042E-44 | 3.4756E-43 |
| ENSG00000143079\|CTTNBP2NL | 4123.8208 | 1.2934 | ##### | 13.9734 | 2.2649E-44 | 3.5695E-43 |
| ENSG00000131507\|NDFIP1 | 12117.0817 | -1.6719 | ##### | ####### | 2.3064E-44 | 3.6329E-43 |
| ENSG00000184451\|CCR10 | 411.0394 | 2.4158 | ##### | 13.9684 | 2.4294E-44 | 3.8247E-43 |
| ENSG00000291141\|EMBP1 | 202.2559 | -2.6181 | ##### | ####### | 2.4395E-44 | 3.8386E-43 |
| ENSG00000138696\|BMPR1B | 790.3147 | -1.9963 | ##### | ####### | 2.6662E-44 | 4.1930E-43 |
| ENSG00000096654\|ZNF184 | 753.1783 | 1.7974 | ##### | 13.9581 | 2.8082E-44 | 4.4140E-43 |
| ENSG00000156869\|FRRS1 | 524.8622 | -2.3920 | ##### | ####### | 2.8139E-44 | 4.4207E-43 |
| ENSG00000232907\|DLGAP4-AS1 | 142.9698 | 2.9408 | ##### | 13.9579 | 2.8165E-44 | 4.4224E-43 |
| ENSG00000112378\|PERP | 15861.4245 | -1.7007 | ##### | ####### | 2.8952E-44 | 4.5437E-43 |
| ENSG00000101445\|PPP1R16B | 152.3439 | 3.6574 | ##### | 13.9559 | 2.8974E-44 | 4.5448E-43 |
| ENSG00000256268\|LINC02454 | 116.5167 | 5.0542 | ##### | 13.9538 | 2.9829E-44 | 4.6763E-43 |
| ENSG00000063127\|SLC6A16 | 97.6060 | -3.6947 | ##### | ####### | 3.1255E-44 | 4.8974E-43 |
| ENSG00000185386\|MAPK11 | 943.0702 | -1.9901 | ##### | ####### | 3.2142E-44 | 5.0336E-43 |
| ENSG00000198799\|LRIG2 | 2360.1548 | 1.6544 | ##### | 13.9451 | 3.3690E-44 | 5.2733E-43 |
| ENSG00000177954\|RPS27 | 35019.1806 | 2.0018 | ##### | 13.9305 | 4.1333E-44 | 6.4662E-43 |
| ENSG00000290318 | 610.3253 | -3.0271 | ##### | ####### | 4.1606E-44 | 6.5056E-43 |
| ENSG00000144224\|UBXN4 | 10472.0232 | -1.6685 | ##### | ####### | 4.1776E-44 | 6.5287E-43 |
| ENSG00000163251\|FZD5 | 1324.2850 | -1.8924 | ##### | ####### | 5.1535E-44 | 8.0497E-43 |
| ENSG00000164109\|MAD2L1 | 7900.9875 | 1.6118 | ##### | 13.9124 | 5.3241E-44 | 8.3117E-43 |
| ENSG00000086300\|SNX10 | 2370.9191 | -1.7728 | ##### | ####### | 5.4665E-44 | 8.5296E-43 |
| ENSG00000112137\|PHACTR1 | 238.0995 | 3.4665 | ##### | 13.9092 | 5.5732E-44 | 8.6916E-43 |
| ENSG00000168795\|ZBTB5 | 3677.3974 | 1.7604 | ##### | 13.8995 | 6.3800E-44 | 9.9446E-43 |
| ENSG00000151553\|FHIP2A | 2143.4484 | -1.8161 | ##### | ####### | 6.6360E-44 | 1.0338E-42 |
| ENSG00000144021\|CIAO1 | 13066.5100 | 1.4464 | ##### | 13.8934 | 6.9443E-44 | 1.0813E-42 |
| ENSG00000140396\|NCOA2 | 2654.9423 | -1.7991 | ##### | ####### | 7.4149E-44 | 1.1540E-42 |
| ENSG00000219626\|FAM228B | 224.5480 | -2.7721 | ##### | ####### | 7.7669E-44 | 1.2081E-42 |
| ENSG00000158106\|RHPN1 | 2818.1416 | 1.7002 | ##### | 13.8842 | 7.9027E-44 | 1.2286E-42 |
| ENSG00000086544\|ITPKC | 1505.7044 | -1.8236 | ##### | ####### | 8.0046E-44 | 1.2438E-42 |
| ENSG00000163376\|KBTBD8 | 329.6925 | 2.8376 | ##### | 13.8826 | 8.0769E-44 | 1.2544E-42 |
| ENSG00000065357\|DGKA | 1056.5854 | -1.7607 | ##### | ####### | 8.5180E-44 | 1.3222E-42 |
| ENSG00000102030\|NAA10 | 9903.0036 | 1.6018 | ##### | 13.8749 | 8.9965E-44 | 1.3957E-42 |
| ENSG00000138074\|SLC5A6 | 10108.8790 | 1.2483 | ##### | 13.8701 | 9.6079E-44 | 1.4898E-42 |
| ENSG00000188505\|NCCRP1 | 140.7772 | -4.9920 | ##### | ####### | 1.0006E-43 | 1.5507E-42 |
| ENSG00000152284\|TCF7L1 | 766.5646 | 1.8336 | ##### | 13.8549 | 1.1876E-43 | 1.8396E-42 |
| ENSG00000023909\|GCLM | 14234.4369 | -1.6555 | ##### | ####### | 1.1934E-43 | 1.8477E-42 |
| ENSG00000103876\|FAH | 1993.7732 | -1.7454 | ##### | ####### | 1.2382E-43 | 1.9159E-42 |
| ENSG00000258441\|LINC00641 | 1301.1820 | 1.9282 | ##### | 13.8415 | 1.4315E-43 | 2.2140E-42 |
| ENSG00000272752\|STAG3L5P-PVRIG2P-PILRB | 3041.8914 | 1.7956 | ##### | 13.8405 | 1.4519E-43 | 2.2443E-42 |
| ENSG00000173207\|CKS1B | 17829.4891 | 1.7098 | ##### | 13.8398 | 1.4671E-43 | 2.2667E-42 |
| ENSG00000132680\|KHDC4 | 7529.2575 | 1.7351 | ##### | 13.8300 | 1.6796E-43 | 2.5936E-42 |
| ENSG00000136802\|LRRC8A | 6501.3530 | -1.6975 | ##### | ####### | 1.8823E-43 | 2.9051E-42 |
| ENSG00000109929\|SC5D | 3566.4947 | -1.7013 | ##### | ####### | 1.9349E-43 | 2.9849E-42 |
| ENSG00000196604\|POTEF | 85.8479 | 4.8196 | ##### | 13.8191 | 1.9562E-43 | 3.0161E-42 |
| ENSG00000146950\|SHROOM2 | 1299.9939 | ####### | ##### | ####### | 2.0086E-43 | 3.0953E-42 |
| ENSG00000106868\|SUSD1 | 986.8539 | -2.0784 | ##### | ####### | 2.0943E-43 | 3.2257E-42 |
| ENSG00000287887 | 117.7231 | 3.5963 | ##### | 13.8120 | 2.1578E-43 | 3.3218E-42 |
| ENSG00000203705\|TATDN3 | 680.0141 | -1.8818 | ##### | ####### | 2.1988E-43 | 3.3832E-42 |
| ENSG00000198223\|CSF2RA | 128.9047 | -4.0413 | ##### | ####### | 2.3502E-43 | 3.6142E-42 |
| ENSG00000180336\|MEIOC | 4153.0428 | 1.7122 | ##### | 13.8058 | 2.3517E-43 | 3.6148E-42 |
| ENSG00000253320\|MAILR | 165.3218 | -2.8049 | ##### | ####### | 2.5595E-43 | 3.9321E-42 |
| ENSG00000185567\|AHNAK2 | 10699.6907 | -1.6961 | ##### | ####### | 2.9126E-43 | 4.4723E-42 |
| ENSG00000117115\|PADI2 | 285.5088 | 2.4338 | ##### | 13.7891 | 2.9631E-43 | 4.5474E-42 |
| ENSG00000236107\|SCN1A-AS1 | 88.9207 | -4.4560 | ##### | ####### | 2.9748E-43 | 4.5631E-42 |
| ENSG00000134986\|NREP | 1366.2600 | 2.4298 | ##### | 13.7866 | 3.0693E-43 | 4.7055E-42 |
| ENSG00000103154\|NECAB2 | 147.7078 | -3.5961 | ##### | ####### | 3.2717E-43 | 5.0133E-42 |
| ENSG00000221955\|SLC12A8 | 284.3642 | -4.3274 | ##### | ####### | 3.2772E-43 | 5.0192E-42 |
| ENSG00000167550\|RHEBL1 | 330.0369 | 2.6297 | ##### | 13.7812 | 3.3071E-43 | 5.0623E-42 |
| ENSG00000280414 | 141.5986 | 4.6404 | ##### | 13.7756 | 3.5743E-43 | 5.4685E-42 |

| | | | | | | |
|---|---|---|---|---|---|---|
| ENSG00000181143\|MUC16 | 317.0835 | -2.7856 | ##### | ####### | 3.6592E-43 | 5.5957E-42 |
| ENSG00000151689\|INPP1 | 1472.4666 | -1.8743 | ##### | ####### | 4.2373E-43 | 6.4764E-42 |
| ENSG00000278318\|ZNF229 | 1775.5318 | -1.7427 | ##### | ####### | 4.2581E-43 | 6.5048E-42 |
| ENSG00000154864\|PIEZO2 | 2505.9684 | -1.9073 | ##### | ####### | 4.3132E-43 | 6.5856E-42 |
| ENSG00000129514\|FOXA1 | 918.8485 | -1.8182 | ##### | ####### | 5.1180E-43 | 7.8103E-42 |
| ENSG00000149596\|JPH2 | 303.3920 | 2.6273 | ##### | 13.7455 | 5.4221E-43 | 8.2703E-42 |
| ENSG00000101745\|ANKRD12 | 3580.0164 | 1.7631 | ##### | 13.7372 | 6.0788E-43 | 9.2671E-42 |
| ENSG00000185697\|MYBL1 | 9123.6648 | 1.4145 | ##### | 13.7344 | 6.3198E-43 | 9.6297E-42 |
| ENSG00000116337\|AMPD2 | 18032.5446 | 1.2249 | ##### | 13.7313 | 6.5978E-43 | 1.0048E-41 |
| ENSG00000171766\|GATM | 751.4026 | 2.1764 | ##### | 13.7303 | 6.6829E-43 | 1.0173E-41 |
| ENSG00000132849\|PATJ | 2588.0860 | -1.7454 | ##### | ####### | 6.8306E-43 | 1.0392E-41 |
| ENSG00000129195\|PIMREG | 3259.6618 | 1.7910 | ##### | 13.7249 | 7.1980E-43 | 1.0946E-41 |
| ENSG00000130529\|TRPM4 | 2471.8951 | -1.7563 | ##### | ####### | 7.2230E-43 | 1.0978E-41 |
| ENSG00000225921\|NOL7 | 6982.6163 | 1.6157 | ##### | 13.7242 | 7.2755E-43 | 1.1052E-41 |
| ENSG00000259877\|ZNF778-DT | 235.5032 | -2.6904 | ##### | ####### | 8.0309E-43 | 1.2193E-41 |
| ENSG00000024526\|DEPDC1 | 7605.8823 | 0.8750 | ##### | 13.7155 | 8.1956E-43 | 1.2437E-41 |
| ENSG00000077935\|SMC1B | 137.4484 | -4.0131 | ##### | ####### | 8.4974E-43 | 1.2889E-41 |
| ENSG00000048649\|RSF1 | 6853.2641 | 1.6461 | ##### | 13.7098 | 8.8670E-43 | 1.3442E-41 |
| ENSG00000123136\|DDX39A | 8941.6751 | 1.7420 | ##### | 13.7084 | 9.0383E-43 | 1.3695E-41 |
| ENSG00000105880\|DLX5 | 169.4545 | 5.5336 | ##### | 13.6963 | 1.0678E-42 | 1.6172E-41 |
| ENSG00000181830\|SLC35C1 | 2099.5161 | -1.7277 | ##### | ####### | 1.1119E-42 | 1.6831E-41 |
| ENSG00000156026\|MCU | 3338.6967 | -1.6572 | ##### | ####### | 1.2288E-42 | 1.8590E-41 |
| ENSG00000198542\|ITGBL1 | 1458.5583 | -1.8583 | ##### | ####### | 1.2696E-42 | 1.9198E-41 |
| ENSG00000272831 | 401.2347 | -2.9107 | ##### | ####### | 1.3929E-42 | 2.1052E-41 |
| ENSG00000135540\|NHSL1 | 898.0085 | 1.9020 | ##### | 13.6745 | 1.4425E-42 | 2.1791E-41 |
| ENSG00000197971\|MBP | 2755.2259 | -1.7214 | ##### | ####### | 1.4483E-42 | 2.1867E-41 |
| ENSG00000177685\|CRACR2B | 787.3427 | 2.0548 | ##### | 13.6729 | 1.4746E-42 | 2.2253E-41 |
| ENSG00000157193\|LRP8 | 6372.2956 | -1.6651 | ##### | ####### | 1.5181E-42 | 2.2897E-41 |
| ENSG00000108819\|PPP1R9B | 16109.7553 | 1.5596 | ##### | 13.6648 | 1.6469E-42 | 2.4829E-41 |
| ENSG00000171617\|ENC1 | 10162.7880 | 1.7156 | ##### | 13.6621 | 1.7095E-42 | 2.5758E-41 |
| ENSG00000267327\|LINC03092 | 292.7623 | 2.5666 | ##### | 13.6617 | 1.7184E-42 | 2.5880E-41 |
| ENSG00000291126 | 290.8350 | 2.3005 | ##### | 13.6609 | 1.7387E-42 | 2.6172E-41 |
| ENSG00000143772\|ITPKB | 1815.4858 | 1.7620 | ##### | 13.6601 | 1.7568E-42 | 2.6431E-41 |
| ENSG00000120519\|SLC10A7 | 350.5637 | -2.2778 | ##### | ####### | 1.7589E-42 | 2.6450E-41 |
| ENSG00000104140\|RHOV | 3424.4785 | 1.6162 | ##### | 13.6594 | 1.7735E-42 | 2.6656E-41 |
| ENSG00000164751\|PEX2 | 1888.7277 | -1.7695 | ##### | ####### | 1.9229E-42 | 2.8887E-41 |
| ENSG00000285382 | 153.8058 | -3.4371 | ##### | ####### | 1.9369E-42 | 2.9083E-41 |
| ENSG00000136014\|USP44 | 461.7568 | -2.4205 | ##### | ####### | 1.9925E-42 | 2.9902E-41 |
| ENSG00000157445\|CACNA2D3 | 138.8374 | -3.0732 | ##### | ####### | 1.9981E-42 | 2.9972E-41 |
| ENSG00000205517\|RGL3 | 653.3116 | -2.0916 | ##### | ####### | 2.0093E-42 | 3.0124E-41 |
| ENSG00000272121 | 98.3760 | -3.6084 | ##### | ####### | 2.5016E-42 | 3.7486E-41 |
| ENSG00000109265\|CRACD | 622.7158 | 2.3582 | ##### | 13.6312 | 2.6134E-42 | 3.9141E-41 |
| ENSG00000118680\|MYL12B | 52922.7498 | 1.6783 | ##### | 13.6310 | 2.6202E-42 | 3.9225E-41 |
| ENSG00000005700\|IBTK | 8090.8770 | -1.6101 | ##### | ####### | 2.7846E-42 | 4.1665E-41 |
| ENSG00000221869\|CEBPD | 1360.3340 | -1.8668 | ##### | ####### | 2.8252E-42 | 4.2251E-41 |
| ENSG00000257702\|LBX2-AS1 | 447.5924 | -2.5044 | ##### | ####### | 2.8553E-42 | 4.2680E-41 |
| ENSG00000100523\|DDHD1 | 2630.0253 | 1.2450 | ##### | 13.6237 | 2.8948E-42 | 4.3248E-41 |
| ENSG00000126882\|FAM78A | 315.8669 | -2.0123 | ##### | ####### | 3.1443E-42 | 4.6952E-41 |
| ENSG00000112144\|CILK1 | 905.6006 | -1.8419 | ##### | ####### | 3.2406E-42 | 4.8366E-41 |
| ENSG00000164099\|PRSS12 | 283.7121 | 3.0709 | ##### | 13.6098 | 3.5043E-42 | 5.2276E-41 |
| ENSG00000163702\|IL17RC | 1330.4162 | -1.8657 | ##### | ####### | 3.5925E-42 | 5.3565E-41 |
| ENSG00000178460\|MCMDC2 | 267.8681 | -2.8306 | ##### | ####### | 3.6296E-42 | 5.4090E-41 |
| ENSG00000253738\|OTUD6B-AS1 | 1564.8666 | -1.8673 | ##### | ####### | 3.8538E-42 | 5.7404E-41 |
| ENSG00000291237\|SOD2 | 15224.3674 | -1.5856 | ##### | ####### | 3.9806E-42 | 5.9262E-41 |
| ENSG00000108511\|HOXB6 | 452.8433 | -2.3620 | ##### | ####### | 4.2473E-42 | 6.3201E-41 |
| ENSG00000144339\|TMEFF2 | 311.3976 | 2.9318 | ##### | 13.5870 | 4.7823E-42 | 7.1127E-41 |
| ENSG00000104805\|NUCB1 | 17911.2633 | -1.6548 | ##### | ####### | 5.0922E-42 | 7.5699E-41 |
| ENSG00000168542\|COL3A1 | 865.3123 | -2.0822 | ##### | ####### | 5.4111E-42 | 8.0399E-41 |
| ENSG00000112419\|PHACTR2 | 1365.5221 | -1.9415 | ##### | ####### | 6.3167E-42 | 9.3808E-41 |
| ENSG00000291035\|RRP7BP | 1192.2340 | 1.8132 | ##### | 13.5639 | 6.5537E-42 | 9.7279E-41 |
| ENSG00000181222\|POLR2A | 29342.9146 | 1.6028 | ##### | 13.5634 | 6.6001E-42 | 9.7919E-41 |
| ENSG00000254829 | 812.8062 | 11.2548 | ##### | 13.5628 | 6.6511E-42 | 9.8627E-41 |
| ENSG00000177406\|NINJ2-AS1 | 222.4667 | -2.5192 | ##### | ####### | 6.7496E-42 | 1.0004E-40 |

| | | | | | | |
|---|---|---|---|---|---|---|
| ENSG00000139131\|YARS2 | 5588.4998 | 1.5275 | ##### | 13.5557 | 7.3280E-42 | 1.0856E-40 |
| ENSG00000139146\|SINHCAF | 9059.7622 | 1.5843 | ##### | 13.5484 | 8.0984E-42 | 1.1991E-40 |
| ENSG00000178999\|AURKB | 4567.0015 | 1.7732 | ##### | 13.5483 | 8.1069E-42 | 1.1998E-40 |
| ENSG00000056487\|PHF21B | 289.2038 | 2.3778 | ##### | 13.5457 | 8.4047E-42 | 1.2432E-40 |
| ENSG00000256894 | 156.1156 | 3.3920 | ##### | 13.5402 | 9.0561E-42 | 1.3389E-40 |
| ENSG00000168209\|DDIT4 | 8950.7834 | -1.6365 | ##### | ####### | 9.8521E-42 | 1.4559E-40 |
| ENSG00000126218\|F10 | 131.1933 | 5.0346 | ##### | 13.5326 | 1.0043E-41 | 1.4833E-40 |
| ENSG00000204389\|HSPA1A | 30336.5558 | 1.5871 | ##### | 13.5324 | 1.0069E-41 | 1.4864E-40 |
| ENSG00000161202\|DVL3 | 4876.5826 | -1.3943 | ##### | ####### | 1.0505E-41 | 1.5500E-40 |
| ENSG00000135069\|PSAT1 | 8210.3032 | -1.6097 | ##### | ####### | 1.0586E-41 | 1.5612E-40 |
| ENSG00000169783\|LINGO1 | 298.9474 | 2.7323 | ##### | 13.5126 | 1.3182E-41 | 1.9431E-40 |
| ENSG00000115295\|CLIP4 | 3202.1967 | -1.7171 | ##### | ####### | 1.3218E-41 | 1.9475E-40 |
| ENSG00000289194 | 564.3801 | 2.2336 | ##### | 13.5123 | 1.3228E-41 | 1.9479E-40 |
| ENSG00000129534\|MIS18BP1 | 4026.5784 | 1.6949 | ##### | 13.5110 | 1.3466E-41 | 1.9821E-40 |
| ENSG00000156587\|UBE2L6 | 1162.4304 | -1.8670 | ##### | ####### | 1.3478E-41 | 1.9829E-40 |
| ENSG00000129566\|TEP1 | 3563.3473 | 1.6935 | ##### | 13.5093 | 1.3776E-41 | 2.0257E-40 |
| ENSG00000128683\|GAD1 | 452.9105 | 2.4081 | ##### | 13.5049 | 1.4634E-41 | 2.1508E-40 |
| ENSG00000106688\|SLC1A1 | 348.8399 | -2.2252 | ##### | ####### | 1.5647E-41 | 2.2986E-40 |
| ENSG00000116171\|SCP2 | 5728.2226 | -1.6517 | ##### | ####### | 1.7394E-41 | 2.5539E-40 |
| ENSG00000114126\|TFDP2 | 3156.1543 | -1.4695 | ##### | ####### | 1.8296E-41 | 2.6850E-40 |
| ENSG00000074657\|ZNF532 | 5520.1212 | -1.6157 | ##### | ####### | 1.8619E-41 | 2.7311E-40 |
| ENSG00000198468\|FLVCR1-DT | 428.0011 | -2.1410 | ##### | ####### | 1.8637E-41 | 2.7324E-40 |
| ENSG00000293257 | 193.9838 | -2.4939 | ##### | ####### | 1.8850E-41 | 2.7623E-40 |
| ENSG00000196368\|NUDT11 | 978.2946 | 1.8518 | ##### | 13.4847 | 1.9245E-41 | 2.8188E-40 |
| ENSG00000086619\|ERO1B | 648.1627 | 1.9535 | ##### | 13.4810 | 2.0243E-41 | 2.9636E-40 |
| ENSG00000064205\|CCN5 | 982.3581 | -6.3302 | ##### | ####### | 2.0447E-41 | 2.9919E-40 |
| ENSG00000277013\|SUNO1 | 101.0061 | 5.1638 | ##### | 13.4790 | 2.0794E-41 | 3.0413E-40 |
| ENSG00000117620\|SLC35A3 | 2421.0252 | -1.6818 | ##### | ####### | 2.3580E-41 | 3.4470E-40 |
| ENSG00000104447\|TRPS1 | 371.1196 | 2.3793 | ##### | 13.4677 | 2.4243E-41 | 3.5421E-40 |
| ENSG00000118058\|KMT2A | 6808.9173 | 1.7177 | ##### | 13.4666 | 2.4583E-41 | 3.5901E-40 |
| ENSG00000185973\|TMLHE | 808.4438 | -1.9048 | ##### | ####### | 2.6224E-41 | 3.8279E-40 |
| ENSG00000100219\|XBP1 | 5174.3737 | -1.7073 | ##### | ####### | 3.0039E-41 | 4.3826E-40 |
| ENSG00000142168\|SOD1 | 9199.3856 | -1.6477 | ##### | ####### | 3.1034E-41 | 4.5256E-40 |
| ENSG00000197451\|HNRNPAB | 35441.1746 | 1.6412 | ##### | 13.4458 | 3.2568E-41 | 4.7470E-40 |
| ENSG00000271447\|MMP28 | 22133.7933 | ####### | ##### | ####### | 3.6038E-41 | 5.2501E-40 |
| ENSG00000137710\|RDX | 14431.1559 | -1.6129 | ##### | ####### | 4.1034E-41 | 5.9750E-40 |
| ENSG00000163449\|TMEM169 | 231.4522 | -2.6679 | ##### | ####### | 4.3778E-41 | 6.3714E-40 |
| ENSG00000176619\|LMNB2 | 41539.7802 | 1.5295 | ##### | 13.4236 | 4.4008E-41 | 6.4018E-40 |
| ENSG00000272711\|HK2-DT | 364.1923 | 3.1262 | ##### | 13.4231 | 4.4281E-41 | 6.4385E-40 |
| ENSG00000076382\|SPAG5 | 12766.2530 | 1.6056 | ##### | 13.4178 | 4.7559E-41 | 6.9116E-40 |
| ENSG00000278974 | 842.8351 | 1.8347 | ##### | 13.4175 | 4.7762E-41 | 6.9377E-40 |
| ENSG00000254721 | 2835.6118 | 1.6545 | ##### | 13.4073 | 5.4782E-41 | 7.9537E-40 |
| ENSG00000204388\|HSPA1B | 24059.6340 | 1.4419 | ##### | 13.4058 | 5.5884E-41 | 8.1098E-40 |
| ENSG00000140545\|MFGE8 | 6897.8497 | -1.6319 | ##### | ####### | 5.6937E-41 | 8.2585E-40 |
| ENSG00000184428\|TOP1MT | 4783.7334 | 1.6179 | ##### | 13.3951 | 6.4583E-41 | 9.3630E-40 |
| ENSG00000179361\|ARID3B | 1488.9072 | 1.8753 | ##### | 13.3925 | 6.6884E-41 | 9.6919E-40 |
| ENSG00000290782\|CFL1P1 | 119.8875 | 3.6816 | ##### | 13.3870 | 7.2056E-41 | 1.0436E-39 |
| ENSG00000184939\|ZFP90 | 1617.3287 | -1.7723 | ##### | ####### | 7.6342E-41 | 1.1052E-39 |
| ENSG00000000419\|DPM1 | 5665.1312 | 1.5542 | ##### | 13.3817 | 7.7355E-41 | 1.1193E-39 |
| ENSG00000116962\|NID1 | 392.7735 | -2.4420 | ##### | ####### | 8.5588E-41 | 1.2378E-39 |
| ENSG00000164414\|SLC35A1 | 1119.6381 | -1.7597 | ##### | ####### | 8.9296E-41 | 1.2908E-39 |
| ENSG00000167600\|CYP2S1 | 1436.5564 | -1.8102 | ##### | ####### | 9.5370E-41 | 1.3780E-39 |
| ENSG00000143341\|HMCN1 | 456.5475 | 5.3828 | ##### | 13.3597 | 1.0392E-40 | 1.5007E-39 |
| ENSG00000108654\|DDX5 | 43011.8021 | 1.0002 | ##### | 13.3593 | 1.0458E-40 | 1.5096E-39 |
| ENSG00000198900\|TOP1 | 23787.0443 | 1.5100 | ##### | 13.3550 | 1.1076E-40 | 1.5980E-39 |
| ENSG00000126777\|KTN1 | 51485.4647 | 1.5802 | ##### | 13.3547 | 1.1121E-40 | 1.6037E-39 |
| ENSG00000283064\|H2BC6-AS1 | 354.9133 | 2.6350 | ##### | 13.3537 | 1.1278E-40 | 1.6255E-39 |
| ENSG00000143256\|PFDN2 | 11558.5370 | 1.5562 | ##### | 13.3529 | 1.1389E-40 | 1.6408E-39 |
| ENSG00000114541\|FRMD4B | 200.7981 | -2.9164 | ##### | ####### | 1.4160E-40 | 2.0390E-39 |
| ENSG00000181472\|ZBTB2 | 4190.8621 | 1.7091 | ##### | 13.3314 | 1.5209E-40 | 2.1889E-39 |
| ENSG00000144655\|CSRNP1 | 2135.6525 | 1.5644 | ##### | 13.3257 | 1.6401E-40 | 2.3595E-39 |
| ENSG00000150782\|IL18 | 4843.6059 | -1.6014 | ##### | ####### | 1.7345E-40 | 2.4940E-39 |
| ENSG00000054282\|SDCCAG8 | 677.1992 | -1.9022 | ##### | ####### | 1.7450E-40 | 2.5079E-39 |

| | | | | | | |
|---|---|---|---|---|---|---|
| ENSG00000137812\|KNL1 | 3975.1117 | 1.6622 | ##### | 13.3189 | 1.7974E-40 | 2.5820E-39 |
| ENSG00000104904\|OAZ1 | 70696.2148 | 1.5579 | ##### | 13.3177 | 1.8255E-40 | 2.6211E-39 |
| ENSG00000111186\|WNT5B | 239.1018 | -2.5782 | ##### | ####### | 1.8763E-40 | 2.6928E-39 |
| ENSG00000237523\|LINC00857 | 584.3118 | -2.1273 | ##### | ####### | 1.9633E-40 | 2.8162E-39 |
| ENSG00000157470\|FAM81A | 1001.2073 | -1.8212 | ##### | ####### | 2.2466E-40 | 3.2211E-39 |
| ENSG00000105552\|BCAT2 | 3209.6648 | -1.6224 | ##### | ####### | 2.3462E-40 | 3.3622E-39 |
| ENSG00000181418\|DDN | 421.2051 | 2.2980 | ##### | 13.2962 | 2.4352E-40 | 3.4882E-39 |
| ENSG00000264717\|NPY4R2 | 251.9265 | -6.3770 | ##### | ####### | 2.6090E-40 | 3.7353E-39 |
| ENSG00000149600\|COMMD7 | 2512.9556 | -1.7156 | ##### | ####### | 2.6135E-40 | 3.7399E-39 |
| ENSG00000215039\|CD27-AS1 | 414.1221 | 2.0596 | ##### | 13.2897 | 2.6564E-40 | 3.7995E-39 |
| ENSG00000071246\|VASH1 | 1992.0616 | 1.6409 | ##### | 13.2862 | 2.7836E-40 | 3.9796E-39 |
| ENSG00000225546\|LINC02476 | 366.4452 | ####### | ##### | ####### | 2.7977E-40 | 3.9978E-39 |
| ENSG00000182612\|TSPAN10 | 585.9805 | -2.0339 | ##### | ####### | 2.8055E-40 | 4.0070E-39 |
| ENSG00000251442\|LINC01094 | 488.7321 | ####### | ##### | ####### | 2.8837E-40 | 4.1168E-39 |
| ENSG00000204044\|SLC12A5-AS1 | 307.4472 | 3.2435 | ##### | 13.2790 | 3.0640E-40 | 4.3720E-39 |
| ENSG00000197892\|KIF13B | 1423.3388 | -1.8970 | ##### | ####### | 3.2402E-40 | 4.6213E-39 |
| ENSG00000137491\|SLCO2B1 | 503.4168 | -7.0490 | ##### | ####### | 3.3070E-40 | 4.7142E-39 |
| ENSG00000188158\|NHS | 1768.0341 | 1.7651 | ##### | 13.2681 | 3.5423E-40 | 5.0472E-39 |
| ENSG00000170954\|ZNF415 | 91.4132 | -5.7812 | ##### | ####### | 3.7727E-40 | 5.3730E-39 |
| ENSG00000164647\|STEAP1 | 1784.2052 | -1.7299 | ##### | ####### | 3.9046E-40 | 5.5581E-39 |
| ENSG00000064199\|SPA17 | 614.9089 | -1.9036 | ##### | ####### | 4.5873E-40 | 6.5269E-39 |
| ENSG00000179913\|B3GNT3 | 472.3622 | -2.5007 | ##### | ####### | 4.6869E-40 | 6.6655E-39 |
| ENSG00000136002\|ARHGEF4 | 698.0520 | 2.1881 | ##### | 13.2400 | 5.1515E-40 | 7.3227E-39 |
| ENSG00000139946\|PELI2 | 215.9546 | -2.4543 | ##### | ####### | 5.8535E-40 | 8.3166E-39 |
| ENSG00000119125\|GDA | 3241.3520 | 1.6485 | ##### | 13.2291 | 5.9579E-40 | 8.4609E-39 |
| ENSG00000145391\|SETD7 | 7422.9998 | -1.5984 | ##### | ####### | 5.9845E-40 | 8.4946E-39 |
| ENSG00000213066\|CEP43 | 3275.1784 | 1.6443 | ##### | 13.2271 | 6.1194E-40 | 8.6820E-39 |
| ENSG00000217236\|SP9 | 362.3487 | 6.0839 | ##### | 13.2256 | 6.2419E-40 | 8.8516E-39 |
| ENSG00000144868\|TMEM108 | 118.6625 | 4.1801 | ##### | 13.2243 | 6.3514E-40 | 9.0026E-39 |
| ENSG00000197429\|IPP | 1379.0539 | -1.7484 | ##### | ####### | 6.4911E-40 | 9.1962E-39 |
| ENSG00000288784\|CTAGE4 | 144.6491 | -3.0109 | ##### | ####### | 6.4985E-40 | 9.2024E-39 |
| ENSG00000165591\|FAAH2 | 574.9989 | ####### | ##### | ####### | 6.5884E-40 | 9.3252E-39 |
| ENSG00000197653\|DNAH10 | 345.7587 | 2.5048 | ##### | 13.2123 | 7.4523E-40 | 1.0543E-38 |
| ENSG00000104320\|NBN | 10755.8728 | -1.3598 | ##### | ####### | 7.5985E-40 | 1.0745E-38 |
| ENSG00000117448\|AKR1A1 | 4207.6061 | -1.5525 | ##### | ####### | 7.9301E-40 | 1.1208E-38 |
| ENSG00000231187\|SYT15-AS1 | 559.7603 | -2.1384 | ##### | ####### | 8.1679E-40 | 1.1539E-38 |
| ENSG00000151611\|MMAA | 388.1258 | -2.0836 | ##### | ####### | 8.7450E-40 | 1.2348E-38 |
| ENSG00000138642\|HERC6 | 418.1641 | -2.1071 | ##### | ####### | 9.6600E-40 | 1.3634E-38 |
| ENSG00000277200\|CCDC92B | 187.9382 | 3.2825 | ##### | 13.1830 | 1.0990E-39 | 1.5504E-38 |
| ENSG00000121410\|A1BG | 443.8612 | -2.0952 | ##### | ####### | 1.1040E-39 | 1.5566E-38 |
| ENSG00000112312\|GMNN | 3125.6552 | 1.6618 | ##### | 13.1826 | 1.1045E-39 | 1.5566E-38 |
| ENSG00000183091\|NEB | 225.0671 | -3.2848 | ##### | ####### | 1.1315E-39 | 1.5939E-38 |
| ENSG00000184271\|POU6F1 | 314.2384 | -2.3748 | ##### | ####### | 1.1401E-39 | 1.6053E-38 |
| ENSG00000164535\|DAGLB | 3926.8250 | -1.6222 | ##### | ####### | 1.1624E-39 | 1.6359E-38 |
| ENSG00000196782\|MAML3 | 722.9146 | -2.0918 | ##### | ####### | 1.1781E-39 | 1.6572E-38 |
| ENSG00000123104\|ITPR2 | 5973.3584 | 1.6143 | ##### | 13.1775 | 1.1817E-39 | 1.6615E-38 |
| ENSG00000176396\|EID2 | 1919.4490 | -1.6358 | ##### | ####### | 1.1860E-39 | 1.6668E-38 |
| ENSG00000165929\|TC2N | 380.7218 | -2.1375 | ##### | ####### | 1.2204E-39 | 1.7144E-38 |
| ENSG00000278041 | 194.0660 | 3.6373 | ##### | 13.1744 | 1.2321E-39 | 1.7300E-38 |
| ENSG00000196497\|IPO4 | 17166.7467 | 1.7133 | ##### | 13.1723 | 1.2662E-39 | 1.7771E-38 |
| ENSG00000213401\|MAGEA12 | 890.2531 | -9.5492 | ##### | ####### | 1.2845E-39 | 1.8019E-38 |
| ENSG00000163590\|PPM1L | 1509.3675 | 1.7216 | ##### | 13.1711 | 1.2872E-39 | 1.8049E-38 |
| ENSG00000291076\|AADACP1 | 185.7104 | -3.0165 | ##### | ####### | 1.3061E-39 | 1.8305E-38 |
| ENSG00000147526\|TACC1 | 7252.6060 | -1.6196 | ##### | ####### | 1.3520E-39 | 1.8938E-38 |
| ENSG00000143401\|ANP32E | 20922.5561 | 1.5640 | ##### | 13.1653 | 1.3904E-39 | 1.9467E-38 |
| ENSG00000101040\|ZMYND8 | 3950.8288 | 1.6152 | ##### | 13.1622 | 1.4472E-39 | 2.0254E-38 |
| ENSG00000253873\|PCDHGA11 | 777.5949 | 3.3345 | ##### | 13.1582 | 1.5278E-39 | 2.1371E-38 |
| ENSG00000166349\|RAG1 | 155.6928 | -5.0811 | ##### | ####### | 1.5306E-39 | 2.1401E-38 |
| ENSG00000090889\|KIF4A | 10576.2524 | 1.5829 | ##### | 13.1477 | 1.7532E-39 | 2.4501E-38 |
| ENSG00000186684\|CYP27C1 | 1237.9918 | -1.7297 | ##### | ####### | 1.7676E-39 | 2.4690E-38 |
| ENSG00000079112\|CDH17 | ########## | ####### | ##### | ####### | 1.9913E-39 | 2.7803E-38 |
| ENSG00000168958\|MFF | 9123.5174 | -1.6611 | ##### | ####### | 2.1259E-39 | 2.9668E-38 |
| ENSG00000147853\|AK3 | 2137.3127 | -1.6688 | ##### | ####### | 2.2463E-39 | 3.1334E-38 |

| | | | | | | |
|---|---|---|---|---|---|---|
| ENSG00000099864\|PALM | 2571.5947 | -1.6408 | ##### | ####### | 2.3047E-39 | 3.2134E-38 |
| ENSG00000107957\|SH3PXD2A | 265.7856 | -2.3659 | ##### | ####### | 2.5087E-39 | 3.4962E-38 |
| ENSG00000037280\|FLT4 | 519.5538 | 2.6238 | ##### | 13.1196 | 2.5422E-39 | 3.5412E-38 |
| ENSG00000031003\|FAM13B | 3660.3211 | 1.6732 | ##### | 13.1160 | 2.6654E-39 | 3.7111E-38 |
| ENSG00000167889\|MGAT5B | 856.9508 | 1.8754 | ##### | 13.1098 | 2.8939E-39 | 4.0273E-38 |
| ENSG00000143578\|CREB3L4 | 1040.3800 | 1.8872 | ##### | 13.1044 | 3.1055E-39 | 4.3198E-38 |
| ENSG00000117266\|CDK18 | 210.6677 | 2.4269 | ##### | 13.1009 | 3.2526E-39 | 4.5222E-38 |
| ENSG00000163701\|IL17RE | 129.7169 | -3.7181 | ##### | ####### | 3.3002E-39 | 4.5863E-38 |
| ENSG00000092969\|TGFB2 | 3568.9636 | -1.6089 | ##### | ####### | 3.4169E-39 | 4.7463E-38 |
| ENSG00000204371\|EHMT2 | 11683.3828 | 1.3694 | ##### | 13.0902 | 3.7439E-39 | 5.1982E-38 |
| ENSG00000126821\|SGPP1 | 9101.6581 | -1.6019 | ##### | ####### | 3.8234E-39 | 5.3061E-38 |
| ENSG00000187775\|DNAH17 | 549.0754 | 2.0105 | ##### | 13.0882 | 3.8485E-39 | 5.3384E-38 |
| ENSG00000001460\|STPG1 | 667.0401 | -1.8979 | ##### | ####### | 4.4191E-39 | 6.1270E-38 |
| ENSG00000265817\|FSBP | 440.6835 | -2.0076 | ##### | ####### | 4.5813E-39 | 6.3490E-38 |
| ENSG00000151876\|FBXO4 | 536.1578 | -2.0168 | ##### | ####### | 4.7460E-39 | 6.5742E-38 |
| ENSG00000259363 | 151.3255 | 2.8335 | ##### | 13.0711 | 4.8168E-39 | 6.6692E-38 |
| ENSG00000099194\|SCD | 43321.9529 | -1.5781 | ##### | ####### | 4.8888E-39 | 6.7658E-38 |
| ENSG00000137970\|RPL7P9 | 3374.2392 | -1.5671 | ##### | ####### | 5.2500E-39 | 7.2622E-38 |
| ENSG00000189195\|BTBD8 | 225.4927 | -2.5249 | ##### | ####### | 5.7527E-39 | 7.9539E-38 |
| ENSG00000165304\|MELK | 7837.9123 | 1.6408 | ##### | 13.0572 | 5.7824E-39 | 7.9913E-38 |
| ENSG00000132535\|DLG4 | 2603.7138 | 1.6608 | ##### | 13.0567 | 5.8184E-39 | 8.0373E-38 |
| ENSG00000148444\|COMMD3 | 2579.5285 | -1.6723 | ##### | ####### | 5.9106E-39 | 8.1608E-38 |
| ENSG00000131650\|KREMEN2 | 1047.6854 | 1.9136 | ##### | 13.0542 | 6.0172E-39 | 8.3042E-38 |
| ENSG00000135094\|SDS | 91.8544 | -4.0005 | ##### | ####### | 6.2772E-39 | 8.6591E-38 |
| ENSG00000196576\|PLXNB2 | 15711.4934 | -1.5335 | ##### | ####### | 6.3903E-39 | 8.8109E-38 |
| ENSG00000237289\|CKMT1B | 800.1359 | -1.8602 | ##### | ####### | 6.3956E-39 | 8.8143E-38 |
| ENSG00000271122\|HERPUD2-AS1 | 1556.4870 | -1.7702 | ##### | ####### | 6.7793E-39 | 9.3388E-38 |
| ENSG00000168014\|C2CD3 | 4397.8096 | 1.5644 | ##### | 13.0409 | 7.1580E-39 | 9.8559E-38 |
| ENSG00000254726\|MEX3A | 1945.3920 | 1.6910 | ##### | 13.0387 | 7.3673E-39 | 1.0139E-37 |
| ENSG00000158691\|ZSCAN12 | 476.4580 | 2.0874 | ##### | 13.0384 | 7.3955E-39 | 1.0173E-37 |
| ENSG00000064102\|INTS13 | 8124.3758 | 0.9308 | ##### | 13.0379 | 7.4476E-39 | 1.0240E-37 |
| ENSG00000116983\|HPCAL4 | 155.6071 | 3.2571 | ##### | 13.0371 | 7.5309E-39 | 1.0350E-37 |
| ENSG00000118418\|HMGN3 | 5173.2339 | 1.5838 | ##### | 13.0312 | 8.1341E-39 | 1.1174E-37 |
| ENSG00000114861\|FOXP1 | 545.8502 | -2.0573 | ##### | ####### | 8.1706E-39 | 1.1219E-37 |
| ENSG00000262655\|SPON1 | 225.5257 | -4.7832 | ##### | ####### | 8.2319E-39 | 1.1298E-37 |
| ENSG00000133103\|COG6 | 1529.5464 | -1.6961 | ##### | ####### | 8.2877E-39 | 1.1369E-37 |
| ENSG00000198464\|ZNF480 | 1533.6842 | -1.7184 | ##### | ####### | 8.3631E-39 | 1.1468E-37 |
| ENSG00000166704\|ZNF606 | 703.9483 | -1.8589 | ##### | ####### | 8.5977E-39 | 1.1784E-37 |
| ENSG00000113108\|APBB3 | 787.5897 | 2.0400 | ##### | 13.0224 | 9.1279E-39 | 1.2505E-37 |
| ENSG00000118855\|MFSD1 | 5070.1587 | -1.5859 | ##### | ####### | 9.2320E-39 | 1.2642E-37 |
| ENSG00000117477\|CCDC181 | 180.6579 | 3.1080 | ##### | 13.0120 | 1.0456E-38 | 1.4311E-37 |
| ENSG00000257621\|PSMA3-AS1 | 1371.6375 | 1.4826 | ##### | 13.0109 | 1.0611E-38 | 1.4510E-37 |
| ENSG00000129347\|KRI1 | 5829.5085 | 1.6149 | ##### | 13.0109 | 1.0611E-38 | 1.4510E-37 |
| ENSG00000005884\|ITGA3 | 39018.9497 | -1.3790 | ##### | ####### | 1.1238E-38 | 1.5360E-37 |
| ENSG00000197077\|KIAA1671 | 4375.0021 | -1.7104 | ##### | ####### | 1.2639E-38 | 1.7267E-37 |
| ENSG00000204131\|NHSL2 | 260.7272 | 2.9839 | ##### | 12.9971 | 1.2711E-38 | 1.7358E-37 |
| ENSG00000155542\|SETD9 | 1149.7731 | -1.2313 | ##### | ####### | 1.3013E-38 | 1.7762E-37 |
| ENSG00000181291\|TMEM132E | 438.3024 | 10.6472 | ##### | 12.9949 | 1.3073E-38 | 1.7836E-37 |
| ENSG00000145506\|NKD2 | 642.2458 | -1.7811 | ##### | ####### | 1.3534E-38 | 1.8456E-37 |
| ENSG00000164694\|FNDC1 | 625.3881 | 10.9259 | ##### | 12.9845 | 1.4980E-38 | 2.0418E-37 |
| ENSG00000178401\|DNAJC22 | 284.2202 | -2.2179 | ##### | ####### | 1.5301E-38 | 2.0847E-37 |
| ENSG00000137075\|RNF38 | 2887.8545 | 1.6855 | ##### | 12.9798 | 1.5935E-38 | 2.1701E-37 |
| ENSG00000002586\|CD99 | 3730.3849 | -1.6712 | ##### | ####### | 1.6600E-38 | 2.2596E-37 |
| ENSG00000172828\|CES3 | 119.9031 | -2.8994 | ##### | ####### | 1.6653E-38 | 2.2659E-37 |
| ENSG00000140479\|PCSK6 | 1448.9022 | -1.7750 | ##### | ####### | 1.6756E-38 | 2.2788E-37 |
| ENSG00000289755 | 763.0991 | -2.1412 | ##### | ####### | 1.7610E-38 | 2.3938E-37 |
| ENSG00000203867\|RBM20 | 166.5922 | -3.1972 | ##### | ####### | 1.8412E-38 | 2.5017E-37 |
| ENSG00000104870\|FCGRT | 843.0320 | -1.8903 | ##### | ####### | 1.9283E-38 | 2.6189E-37 |
| ENSG00000169231\|THBS3 | 7040.6054 | 1.5378 | ##### | 12.9619 | 2.0128E-38 | 2.7323E-37 |
| ENSG00000164236\|ANKRD33B | 5303.8462 | -1.5859 | ##### | ####### | 2.0632E-38 | 2.7995E-37 |
| ENSG00000106524\|ANKMY2 | 2057.1578 | -1.6202 | ##### | ####### | 2.0676E-38 | 2.8043E-37 |
| ENSG00000155265\|GOLGA7B | 563.3893 | 2.1271 | ##### | 12.9549 | 2.2044E-38 | 2.9871E-37 |
| ENSG00000168476\|REEP4 | 3923.5755 | 1.5944 | ##### | 12.9549 | 2.2041E-38 | 2.9871E-37 |

| | | | | | | |
|---|---|---|---|---|---|---|
| ENSG00000147650\|LRP12 | 3974.2433 | -1.6385 | ##### | ####### | 2.2420E-38 | 3.0367E-37 |
| ENSG00000082512\|TRAF5 | 620.0693 | -1.8323 | ##### | ####### | 2.3125E-38 | 3.1307E-37 |
| ENSG00000135519\|KCNH3 | 413.2880 | -1.9567 | ##### | ####### | 2.6321E-38 | 3.5617E-37 |
| ENSG00000076770\|MBNL3 | 540.0778 | 1.9944 | ##### | 12.9397 | 2.6853E-38 | 3.6322E-37 |
| ENSG00000214160\|ALG3 | 4797.8157 | -1.6525 | ##### | ####### | 2.7424E-38 | 3.7076E-37 |
| ENSG00000254635\|WAC-AS1 | 1604.0123 | -1.5879 | ##### | ####### | 3.1246E-38 | 4.2224E-37 |
| ENSG00000172164\|SNTB1 | 329.8622 | -2.0199 | ##### | ####### | 3.1306E-38 | 4.2286E-37 |
| ENSG00000008952\|SEC62 | 7815.7982 | -1.5707 | ##### | ####### | 3.1321E-38 | 4.2288E-37 |
| ENSG00000189068\|VSTM1 | 143.9629 | 3.2476 | ##### | 12.9246 | 3.2685E-38 | 4.4109E-37 |
| ENSG00000149218\|ENDOD1 | 2014.5720 | -1.3601 | ##### | ####### | 3.4420E-38 | 4.6430E-37 |
| ENSG00000136830\|NIBAN2 | 21263.1359 | -1.4582 | ##### | ####### | 3.6014E-38 | 4.8558E-37 |
| ENSG00000156298\|TSPAN7 | 483.0048 | -3.9517 | ##### | ####### | 3.6576E-38 | 4.9293E-37 |
| ENSG00000198719\|DLL1 | 147.6635 | 3.7951 | ##### | 12.9119 | 3.8590E-38 | 5.1984E-37 |
| ENSG00000171649\|ZIK1 | 56.8724 | -5.2968 | ##### | ####### | 4.2340E-38 | 5.7010E-37 |
| ENSG00000109445\|ZNF330 | 1835.9849 | -1.6265 | ##### | ####### | 4.6055E-38 | 6.1984E-37 |
| ENSG00000115756\|HPCAL1 | 7950.8474 | -1.1514 | ##### | ####### | 4.8867E-38 | 6.5739E-37 |
| ENSG00000147100\|SLC16A2 | 4332.0498 | 1.6269 | ##### | 12.8917 | 5.0097E-38 | 6.7364E-37 |
| ENSG00000101972\|STAG2 | 11146.7881 | 1.5136 | ##### | 12.8872 | 5.3147E-38 | 7.1433E-37 |
| ENSG00000140564\|FURIN | 4422.6658 | -1.5732 | ##### | ####### | 5.4441E-38 | 7.3140E-37 |
| ENSG00000168297\|PXK | 2093.9416 | 1.7929 | ##### | 12.8849 | 5.4736E-38 | 7.3503E-37 |
| ENSG00000105767\|CADM4 | 633.7592 | -1.8284 | ##### | ####### | 5.8187E-38 | 7.8101E-37 |
| ENSG00000130700\|GATA5 | 150.7050 | 4.3756 | ##### | 12.8776 | 6.0213E-38 | 8.0785E-37 |
| ENSG00000273373\|SLC16A4-AS1 | 261.7295 | 2.4817 | ##### | 12.8652 | 7.0624E-38 | 9.4709E-37 |
| ENSG00000164190\|NIPBL | 11624.1778 | 1.5343 | ##### | 12.8605 | 7.5112E-38 | 1.0068E-36 |
| ENSG00000180425\|C11orf71 | 434.9696 | -2.2350 | ##### | ####### | 7.5412E-38 | 1.0104E-36 |
| ENSG00000101000\|PROCR | 10040.5104 | -1.5738 | ##### | ####### | 8.8001E-38 | 1.1785E-36 |
| ENSG00000166750\|SLFN5 | 2415.4674 | -1.6742 | ##### | ####### | 1.0669E-37 | 1.4282E-36 |
| ENSG00000104728\|ARHGEF10 | 5719.1240 | 1.4949 | ##### | 12.8244 | 1.1975E-37 | 1.6023E-36 |
| ENSG00000112977\|DAP | 7104.1587 | -1.5047 | ##### | ####### | 1.2243E-37 | 1.6374E-36 |
| ENSG00000116663\|FBXO6 | 1331.6412 | -1.7932 | ##### | ####### | 1.2443E-37 | 1.6635E-36 |
| ENSG00000147416\|ATP6V1B2 | 4573.2869 | 1.1858 | ##### | 12.8210 | 1.2500E-37 | 1.6703E-36 |
| ENSG00000008283\|CYB561 | 4293.4516 | -1.5225 | ##### | ####### | 1.3421E-37 | 1.7926E-36 |
| ENSG00000206560\|ANKRD28 | 7504.3635 | 1.6028 | ##### | 12.8063 | 1.5117E-37 | 2.0182E-36 |
| ENSG00000162745\|OLFML2B | 520.8366 | -1.9031 | ##### | ####### | 1.5274E-37 | 2.0383E-36 |
| ENSG00000169914\|OTUD3 | 1652.8766 | 1.7340 | ##### | 12.8037 | 1.5630E-37 | 2.0848E-36 |
| ENSG00000013573\|DDX11 | 7526.7107 | 1.5065 | ##### | 12.7993 | 1.6539E-37 | 2.2050E-36 |
| ENSG00000130332\|LSM7 | 6064.2923 | 1.5614 | ##### | 12.7991 | 1.6596E-37 | 2.2117E-36 |
| ENSG00000141965\|FEM1A | 7412.9926 | 1.3039 | ##### | 12.7989 | 1.6623E-37 | 2.2144E-36 |
| ENSG00000145147\|SLIT2 | 431.2623 | -2.5446 | ##### | ####### | 1.6785E-37 | 2.2348E-36 |
| ENSG00000158636\|EMSY | 2841.0317 | 1.6078 | ##### | 12.7876 | 1.9229E-37 | 2.5591E-36 |
| ENSG00000136111\|TBC1D4 | 1970.9939 | 1.6764 | ##### | 12.7826 | 2.0516E-37 | 2.7292E-36 |
| ENSG00000186283\|TOR3A | 3458.9002 | -1.3836 | ##### | ####### | 2.0827E-37 | 2.7694E-36 |
| ENSG00000105464\|GRIN2D | 4389.4959 | -1.5546 | ##### | ####### | 2.1560E-37 | 2.8656E-36 |
| ENSG00000236699\|ARHGEF38 | 82.4488 | -3.2389 | ##### | ####### | 2.2252E-37 | 2.9562E-36 |
| ENSG00000196814\|MVB12B | 729.8499 | -1.8558 | ##### | ####### | 2.2371E-37 | 2.9707E-36 |
| ENSG00000196611\|MMP1 | 629.7980 | 2.9024 | ##### | 12.7750 | 2.2618E-37 | 3.0022E-36 |
| ENSG00000123685\|BATF3 | 606.3935 | 2.0458 | ##### | 12.7743 | 2.2809E-37 | 3.0262E-36 |
| ENSG00000134444\|RELCH | 3291.2933 | 1.5562 | ##### | 12.7741 | 2.2876E-37 | 3.0337E-36 |
| ENSG00000105810\|CDK6 | 3903.1446 | -1.6392 | ##### | ####### | 2.3650E-37 | 3.1349E-36 |
| ENSG00000127616\|SMARCA4 | 1943.9549 | 1.6049 | ##### | 12.7703 | 2.4031E-37 | 3.1840E-36 |
| ENSG00000145901\|TNIP1 | 9655.3951 | -1.5279 | ##### | ####### | 2.6464E-37 | 3.5049E-36 |
| ENSG00000104774\|MAN2B1 | 2226.3762 | -1.6329 | ##### | ####### | 2.6840E-37 | 3.5531E-36 |
| ENSG00000134198\|TSPAN2 | 182.7890 | 5.8142 | ##### | 12.7579 | 2.8164E-37 | 3.7267E-36 |
| ENSG00000119471\|HSDL2 | 3340.6252 | -1.5979 | ##### | ####### | 2.8474E-37 | 3.7661E-36 |
| ENSG00000289725 | 341.5089 | ####### | ##### | ####### | 2.9218E-37 | 3.8627E-36 |
| ENSG00000104953\|TLE6 | 781.4140 | -1.7675 | ##### | ####### | 2.9387E-37 | 3.8834E-36 |
| ENSG00000196976\|LAGE3 | 5807.7644 | 1.6198 | ##### | 12.7533 | 2.9873E-37 | 3.9458E-36 |
| ENSG00000128059\|PPAT | 8925.4882 | 1.5544 | ##### | 12.7512 | 3.0686E-37 | 4.0514E-36 |
| ENSG00000149084\|HSD17B12 | 6855.6802 | -1.5925 | ##### | ####### | 3.3197E-37 | 4.3811E-36 |
| ENSG00000158201\|ABHD3 | 2712.0542 | -1.6610 | ##### | ####### | 3.5487E-37 | 4.6812E-36 |
| ENSG00000111058\|ACSS3 | 96.0581 | -3.9985 | ##### | ####### | 3.7938E-37 | 5.0022E-36 |
| ENSG00000134057\|CCNB1 | 16401.1965 | 1.3264 | ##### | 12.7339 | 3.8319E-37 | 5.0503E-36 |
| ENSG00000123178\|SPRYD7 | 2838.3622 | -1.5802 | ##### | ####### | 3.8359E-37 | 5.0533E-36 |

| | | | | | | |
|---|---|---|---|---|---|---|
| ENSG00000064419\|TNPO3 | 11232.9147 | 0.7840 | ##### | 12.7323 | 3.9125E-37 | 5.1520E-36 |
| ENSG00000185561\|TLCD2 | 335.9817 | -2.0592 | ##### | ####### | 3.9855E-37 | 5.2458E-36 |
| ENSG00000234685\|NUS1P2 | 101.4967 | 5.3367 | ##### | 12.7302 | 4.0175E-37 | 5.2855E-36 |
| ENSG00000186625\|KATNA1 | 2744.5624 | 1.6328 | ##### | 12.7301 | 4.0250E-37 | 5.2931E-36 |
| ENSG00000198431\|TXNRD1 | ########## | -1.4955 | ##### | ####### | 4.1312E-37 | 5.4304E-36 |
| ENSG00000182759\|MAFA | 163.6861 | 2.6737 | ##### | 12.7248 | 4.3070E-37 | 5.6590E-36 |
| ENSG00000234160 | 319.6599 | 2.3681 | ##### | 12.7199 | 4.5849E-37 | 6.0215E-36 |
| ENSG00000143162\|CREG1 | 3644.9734 | -1.6181 | ##### | ####### | 5.2090E-37 | 6.8382E-36 |
| ENSG00000175130\|MARCKSL1 | 6178.5039 | -1.5580 | ##### | ####### | 5.3551E-37 | 7.0268E-36 |
| ENSG00000261183\|SPINT1-AS1 | 279.8374 | ####### | ##### | ####### | 5.4025E-37 | 7.0860E-36 |
| ENSG00000186185\|KIF18B | 6753.2425 | 1.4955 | ##### | 12.7038 | 5.6358E-37 | 7.3886E-36 |
| ENSG00000105426\|PTPRS | 10171.9053 | -1.5069 | ##### | ####### | 5.7353E-37 | 7.5158E-36 |
| ENSG00000182165\|TP53TG1 | 585.5664 | -1.8360 | ##### | ####### | 5.8601E-37 | 7.6759E-36 |
| ENSG00000198182\|ZNF607 | 852.9531 | -1.6146 | ##### | ####### | 5.9758E-37 | 7.8241E-36 |
| ENSG00000121236\|TRIM6 | 1898.2348 | -1.6007 | ##### | ####### | 6.0828E-37 | 7.9606E-36 |
| ENSG00000287839 | 219.2016 | -3.5330 | ##### | ####### | 6.3698E-37 | 8.3327E-36 |
| ENSG00000261655 | 275.7402 | ####### | ##### | ####### | 6.6275E-37 | 8.6660E-36 |
| ENSG00000203485\|INF2 | 9112.3810 | -0.9498 | ##### | ####### | 6.7572E-37 | 8.8317E-36 |
| ENSG00000148408\|CACNA1B | 334.8775 | 3.7841 | ##### | 12.6831 | 7.3328E-37 | 9.5798E-36 |
| ENSG00000103174\|NAGPA | 1560.6585 | -1.1362 | ##### | ####### | 7.6169E-37 | 9.9467E-36 |
| ENSG00000105325\|FZR1 | 6856.3491 | 1.5297 | ##### | 12.6768 | 7.9556E-37 | 1.0384E-35 |
| ENSG00000168062\|BATF2 | 215.0653 | -3.2521 | ##### | ####### | 7.9702E-37 | 1.0399E-35 |
| ENSG00000173237\|C11orf86 | 91.5106 | -3.6966 | ##### | ####### | 7.9780E-37 | 1.0405E-35 |
| ENSG00000175581\|MRPL48 | 1402.3526 | 1.7364 | ##### | 12.6600 | 9.8486E-37 | 1.2838E-35 |
| ENSG00000172985\|SH3RF3 | 258.9766 | 2.5718 | ##### | 12.6570 | 1.0228E-36 | 1.3327E-35 |
| ENSG00000187140\|FOXD3 | 357.8766 | -2.0745 | ##### | ####### | 1.0686E-36 | 1.3917E-35 |
| ENSG00000168792\|ABHD15 | 2106.5477 | -1.6033 | ##### | ####### | 1.1069E-36 | 1.4410E-35 |
| ENSG00000232445\|EMSLR | 1150.6249 | 1.9273 | ##### | 12.6505 | 1.1123E-36 | 1.4474E-35 |
| ENSG00000144908\|ALDH1L1 | 278.4512 | 2.3839 | ##### | 12.6481 | 1.1461E-36 | 1.4908E-35 |
| ENSG00000186469\|GNG2 | 169.2146 | -5.3085 | ##### | ####### | 1.1722E-36 | 1.5241E-35 |
| ENSG00000173905\|GOLIM4 | 5048.9785 | -1.5347 | ##### | ####### | 1.1787E-36 | 1.5318E-35 |
| ENSG00000134755\|DSC2 | 1439.2079 | -1.6789 | ##### | ####### | 1.2508E-36 | 1.6248E-35 |
| ENSG00000163683\|SMIM14 | 2736.1410 | -1.5364 | ##### | ####### | 1.2532E-36 | 1.6273E-35 |
| ENSG00000168566\|SNRNP48 | 1884.3563 | 1.6619 | ##### | 12.6329 | 1.3901E-36 | 1.8043E-35 |
| ENSG00000070756\|PABPC1 | ########## | -0.9010 | ##### | ####### | 1.3983E-36 | 1.8141E-35 |
| ENSG00000166913\|YWHAB | 43178.0817 | 1.1866 | ##### | 12.6291 | 1.4603E-36 | 1.8937E-35 |
| ENSG00000122912\|SLC25A16 | 1347.7318 | -1.7724 | ##### | ####### | 1.4887E-36 | 1.9297E-35 |
| ENSG00000117305\|HMGCL | 1082.3649 | -1.7508 | ##### | ####### | 1.5581E-36 | 2.0187E-35 |
| ENSG00000099817\|POLR2E | 18595.5117 | 1.0305 | ##### | 12.6218 | 1.6005E-36 | 2.0729E-35 |
| ENSG00000130830\|MPP1 | 856.2806 | -1.6589 | ##### | ####### | 1.7075E-36 | 2.2104E-35 |
| ENSG00000102981\|PARD6A | 429.4951 | 2.4812 | ##### | 12.6130 | 1.7911E-36 | 2.3176E-35 |
| ENSG00000099204\|ABLIM1 | 8837.2511 | -1.5073 | ##### | ####### | 1.8315E-36 | 2.3689E-35 |
| ENSG00000113810\|SMC4 | 26837.4120 | 1.4846 | ##### | 12.6024 | 2.0488E-36 | 2.6488E-35 |
| ENSG00000137285\|TUBB2B | 1806.1805 | -1.6082 | ##### | ####### | 2.1088E-36 | 2.7252E-35 |
| ENSG00000129484\|PARP2 | 3394.4009 | 1.2555 | ##### | 12.5988 | 2.1437E-36 | 2.7691E-35 |
| ENSG00000165102\|HGSNAT | 1911.0156 | -1.5638 | ##### | ####### | 2.3175E-36 | 2.9923E-35 |
| ENSG00000262198 | 118.5638 | 7.3647 | ##### | 12.5917 | 2.3450E-36 | 3.0266E-35 |
| ENSG00000204128\|C2orf72 | 492.8351 | -1.9601 | ##### | ####### | 2.4209E-36 | 3.1231E-35 |
| ENSG00000140873\|ADAMTS18 | 206.0193 | 2.3441 | ##### | 12.5847 | 2.5638E-36 | 3.3061E-35 |
| ENSG00000275778 | 428.2348 | 2.0196 | ##### | 12.5818 | 2.6586E-36 | 3.4268E-35 |
| ENSG00000064666\|CNN2 | 23099.6997 | 1.5030 | ##### | 12.5704 | 3.0703E-36 | 3.9558E-35 |
| ENSG00000196628\|TCF4 | 825.8709 | 1.7353 | ##### | 12.5696 | 3.1012E-36 | 3.9938E-35 |
| ENSG00000112996\|MRPS30 | 11187.2685 | 1.4377 | ##### | 12.5684 | 3.1489E-36 | 4.0535E-35 |
| ENSG00000213614\|HEXA | 4406.1827 | -1.5405 | ##### | ####### | 3.1832E-36 | 4.0960E-35 |
| ENSG00000157927\|RADIL | 596.1421 | 1.7778 | ##### | 12.5665 | 3.2279E-36 | 4.1516E-35 |
| ENSG00000052795\|FNIP2 | 727.4622 | -1.8209 | ##### | ####### | 3.3453E-36 | 4.3008E-35 |
| ENSG00000166676\|TVP23A | 84.4837 | -3.7647 | ##### | ####### | 3.8363E-36 | 4.9299E-35 |
| ENSG00000115318\|LOXL3 | 660.7953 | 2.4657 | ##### | 12.5471 | 4.1256E-36 | 5.2994E-35 |
| ENSG00000230798\|FOXD3-AS1 | 215.6068 | -2.6025 | ##### | ####### | 4.4872E-36 | 5.7614E-35 |
| ENSG00000129932\|DOHH | 3745.2455 | 1.5376 | ##### | 12.5336 | 4.8890E-36 | 6.2745E-35 |
| ENSG00000183864\|TOB2 | 6054.1914 | 1.0660 | ##### | 12.5297 | 5.1335E-36 | 6.5856E-35 |
| ENSG00000084092\|NOA1 | 4651.6863 | 1.5378 | ##### | 12.5291 | 5.1765E-36 | 6.6379E-35 |
| ENSG00000124181\|PLCG1 | 7043.6473 | 1.5269 | ##### | 12.5286 | 5.2081E-36 | 6.6755E-35 |

| | | | | | | |
|---|---|---|---|---|---|---|
| ENSG00000174292\|TNK1 | 216.8769 | -2.2415 | ##### | ####### | 5.4567E-36 | 6.9912E-35 |
| ENSG00000106714\|CNTNAP3 | 775.5021 | -1.8262 | ##### | ####### | 5.6688E-36 | 7.2599E-35 |
| ENSG00000133216\|EPHB2 | 3612.0335 | -1.7195 | ##### | ####### | 5.7297E-36 | 7.3346E-35 |
| ENSG00000198554\|WDHD1 | 6101.4143 | 1.5189 | ##### | 12.5198 | 5.8205E-36 | 7.4477E-35 |
| ENSG00000165650\|PDZD8 | 8275.4819 | -1.4718 | ##### | ####### | 5.8950E-36 | 7.5398E-35 |
| ENSG00000137752\|CASP1 | 87.4096 | -4.4660 | ##### | ####### | 5.9029E-36 | 7.5467E-35 |
| ENSG00000160959\|LRRC14 | 3051.0861 | -1.3273 | ##### | ####### | 6.0924E-36 | 7.7856E-35 |
| ENSG00000088756\|ARHGAP28 | 1828.1961 | -1.6578 | ##### | ####### | 6.3405E-36 | 8.0992E-35 |
| ENSG00000280832\|GSEC | 309.2147 | -2.0878 | ##### | ####### | 6.6083E-36 | 8.4377E-35 |
| ENSG00000130433\|CACNG6 | 1147.9133 | -1.7927 | ##### | ####### | 6.8162E-36 | 8.6994E-35 |
| ENSG00000171421\|MRPL36 | 1540.2952 | -1.6891 | ##### | ####### | 7.0815E-36 | 9.0341E-35 |
| ENSG00000225830\|ERCC6 | 2766.1589 | 1.5428 | ##### | 12.5036 | 7.1336E-36 | 9.0967E-35 |
| ENSG00000211584\|SLC48A1 | 2786.7689 | -1.6066 | ##### | ####### | 7.1858E-36 | 9.1593E-35 |
| ENSG00000153574\|RPIA | 4751.5076 | 1.5707 | ##### | 12.4979 | 7.6634E-36 | 9.7639E-35 |
| ENSG00000153246\|PLA2R1 | 504.6858 | -2.0649 | ##### | ####### | 7.7326E-36 | 9.8479E-35 |
| ENSG00000156052\|GNAQ | 4116.1336 | -1.5429 | ##### | ####### | 8.1986E-36 | 1.0437E-34 |
| ENSG00000293137\|MSL3P1 | 957.7136 | -1.6352 | ##### | ####### | 9.1219E-36 | 1.1607E-34 |
| ENSG00000123836\|PFKFB2 | 1659.9821 | -1.6065 | ##### | ####### | 9.5523E-36 | 1.2150E-34 |
| ENSG00000078070\|MCCC1 | 1115.8587 | -1.7143 | ##### | ####### | 1.0279E-35 | 1.3069E-34 |
| ENSG00000100591\|AHSA1 | 16292.2102 | 1.4992 | ##### | 12.4692 | 1.0993E-35 | 1.3971E-34 |
| ENSG00000137040\|RANBP6 | 1611.7277 | -1.5616 | ##### | ####### | 1.1142E-35 | 1.4154E-34 |
| ENSG00000174428\|GTF2IRD2B | 807.8073 | -1.6615 | ##### | ####### | 1.1519E-35 | 1.4626E-34 |
| ENSG00000160679\|CHTOP | 14114.8952 | 1.3718 | ##### | 12.4634 | 1.1821E-35 | 1.5004E-34 |
| ENSG00000111203\|ITFG2 | 2958.4817 | 1.8947 | ##### | 12.4589 | 1.2503E-35 | 1.5863E-34 |
| ENSG00000161405\|IKZF3 | 234.2277 | 3.0514 | ##### | 12.4503 | 1.3929E-35 | 1.7664E-34 |
| ENSG00000181444\|ZNF467 | 2193.7399 | -1.5187 | ##### | ####### | 1.4143E-35 | 1.7927E-34 |
| ENSG00000055118\|KCNH2 | 166.6530 | 2.8596 | ##### | 12.4452 | 1.4849E-35 | 1.8815E-34 |
| ENSG00000178150\|ZNF114 | 303.0846 | -2.1041 | ##### | ####### | 1.4894E-35 | 1.8864E-34 |
| ENSG00000143158\|MPC2 | 6081.6635 | -1.5335 | ##### | ####### | 1.4959E-35 | 1.8938E-34 |
| ENSG00000026950\|BTN3A1 | 598.3118 | -1.8007 | ##### | ####### | 1.5710E-35 | 1.9880E-34 |
| ENSG00000160785\|SLC25A44 | 7140.0690 | 1.5207 | ##### | 12.4390 | 1.6052E-35 | 2.0304E-34 |
| ENSG00000152492\|CCDC50 | 4157.1532 | -1.5602 | ##### | ####### | 1.7421E-35 | 2.2027E-34 |
| ENSG00000111801\|BTN3A3 | 165.8879 | -2.6141 | ##### | ####### | 1.7727E-35 | 2.2405E-34 |
| ENSG00000157368\|IL34 | 108.1999 | -3.9017 | ##### | ####### | 1.8891E-35 | 2.3865E-34 |
| ENSG00000104884\|ERCC2 | 5975.8273 | -1.2773 | ##### | ####### | 1.9068E-35 | 2.4078E-34 |
| ENSG00000291224 | 594.2446 | 2.8006 | ##### | 12.4230 | 1.9607E-35 | 2.4750E-34 |
| ENSG00000261215 | 173.0676 | 2.7399 | ##### | 12.4210 | 2.0104E-35 | 2.5365E-34 |
| ENSG00000115268\|RPS15 | 47019.6299 | 1.4804 | ##### | 12.4203 | 2.0268E-35 | 2.5562E-34 |
| ENSG00000043093\|DCUN1D1 | 2711.3359 | -1.5893 | ##### | ####### | 2.1353E-35 | 2.6919E-34 |
| ENSG00000158286\|RNF207 | 1838.8493 | -1.7350 | ##### | ####### | 2.1440E-35 | 2.7017E-34 |
| ENSG00000154721\|JAM2 | 190.5335 | 5.1134 | ##### | 12.4148 | 2.1714E-35 | 2.7350E-34 |
| ENSG00000158850\|B4GALT3 | 7931.7757 | 1.5144 | ##### | 12.4107 | 2.2868E-35 | 2.8792E-34 |
| ENSG00000249859\|PVT1 | 5583.3147 | 1.5688 | ##### | 12.4032 | 2.5097E-35 | 3.1585E-34 |
| ENSG00000164778\|EN2 | 183.2472 | 2.8997 | ##### | 12.4007 | 2.5912E-35 | 3.2598E-34 |
| ENSG00000173020\|GRK2 | 10615.6015 | 1.4434 | ##### | 12.3979 | 2.6816E-35 | 3.3720E-34 |
| ENSG00000149716\|LTO1 | 3255.0537 | 1.5707 | ##### | 12.3948 | 2.7878E-35 | 3.5041E-34 |
| ENSG00000233927\|RPS28 | 8192.5215 | -1.6701 | ##### | ####### | 2.8273E-35 | 3.5522E-34 |
| ENSG00000120725\|SIL1 | 2562.0837 | -1.6085 | ##### | ####### | 2.8412E-35 | 3.5682E-34 |
| ENSG00000269001\|ZNF818P | 4750.7009 | ####### | ##### | ####### | 2.9532E-35 | 3.7073E-34 |
| ENSG00000115963\|RND3 | 6322.9168 | -1.3405 | ##### | ####### | 2.9767E-35 | 3.7352E-34 |
| ENSG00000049656\|CLPTM1L | 15608.1518 | -1.5048 | ##### | ####### | 3.6368E-35 | 4.5617E-34 |
| ENSG00000165376\|CLDN2 | 49.8872 | -5.1657 | ##### | ####### | 3.6701E-35 | 4.6015E-34 |
| ENSG00000143333\|RGS16 | 307.2943 | 2.6070 | ##### | 12.3707 | 3.7654E-35 | 4.7190E-34 |
| ENSG00000172270\|BSG | 67341.4437 | 1.4420 | ##### | 12.3696 | 3.8166E-35 | 4.7812E-34 |
| ENSG00000117528\|ABCD3 | 6582.6221 | -1.5128 | ##### | ####### | 3.9648E-35 | 4.9648E-34 |
| ENSG00000279488 | 177.8869 | 2.5184 | ##### | 12.3647 | 4.0580E-35 | 5.0793E-34 |
| ENSG00000132819\|RBM38 | 4036.7455 | 1.5398 | ##### | 12.3642 | 4.0807E-35 | 5.1056E-34 |
| ENSG00000058085\|LAMC2 | 1528.4364 | -2.1166 | ##### | ####### | 4.2802E-35 | 5.3529E-34 |
| ENSG00000232320\|MXRA7P1 | 218.9616 | 2.9030 | ##### | 12.3576 | 4.4298E-35 | 5.5377E-34 |
| ENSG00000147586\|MRPS28 | 1605.7148 | -1.5434 | ##### | ####### | 4.7078E-35 | 5.8828E-34 |
| ENSG00000112763\|BTN2A1 | 2625.8776 | 1.5453 | ##### | 12.3524 | 4.7297E-35 | 5.9077E-34 |
| ENSG00000038002\|AGA | 676.1436 | -1.7795 | ##### | ####### | 4.7847E-35 | 5.9738E-34 |
| ENSG00000267980 | 105.5633 | 3.4058 | ##### | 12.3500 | 4.8730E-35 | 6.0816E-34 |

| | | | | | | |
|---|---|---|---|---|---|---|
| ENSG00000058866\|DGKG | 5513.3070 | ####### | ##### | ####### | 5.3230E-35 | 6.6404E-34 |
| ENSG00000152208\|GRID2 | 152.1723 | -2.5842 | ##### | ####### | 5.3338E-35 | 6.6511E-34 |
| ENSG00000189319\|FAM53B | 2325.2475 | -1.3450 | ##### | ####### | 5.4586E-35 | 6.8039E-34 |
| ENSG00000188452\|CERKL | 305.9460 | -2.2646 | ##### | ####### | 5.9058E-35 | 7.3582E-34 |
| ENSG00000104356\|POP1 | 3356.9389 | -1.6000 | ##### | ####### | 5.9364E-35 | 7.3932E-34 |
| ENSG00000133703\|KRAS | 13198.9447 | 1.4606 | ##### | 12.3296 | 6.2788E-35 | 7.8164E-34 |
| ENSG00000242612\|DECR2 | 1645.1202 | -1.6418 | ##### | ####### | 6.2918E-35 | 7.8294E-34 |
| ENSG00000135862\|LAMC1 | 22002.0615 | -1.4492 | ##### | ####### | 6.4803E-35 | 8.0606E-34 |
| ENSG00000155974\|GRIP1 | 1419.0966 | 1.6714 | ##### | 12.3245 | 6.6876E-35 | 8.3150E-34 |
| ENSG00000101189\|MRGBP | 5877.3251 | 1.5420 | ##### | 12.3142 | 7.5966E-35 | 9.4412E-34 |
| ENSG00000167964\|RAB26 | 216.4594 | -2.7139 | ##### | ####### | 8.0232E-35 | 9.9631E-34 |
| ENSG00000134030\|CTIF | 4459.3693 | -1.5076 | ##### | ####### | 8.0201E-35 | 9.9631E-34 |
| ENSG00000079819\|EPB41L2 | 5042.1033 | -1.4560 | ##### | ####### | 8.3718E-35 | 1.0392E-33 |
| ENSG00000137203\|TFAP2A | 5585.5886 | 1.5208 | ##### | 12.3028 | 8.7519E-35 | 1.0859E-33 |
| ENSG00000143641\|GALNT2 | 27546.4665 | -1.3307 | ##### | ####### | 8.9687E-35 | 1.1123E-33 |
| ENSG00000227640\|SOX21-AS1 | 139.4443 | -3.3243 | ##### | ####### | 9.3401E-35 | 1.1579E-33 |
| ENSG00000175920\|DOK7 | 89.0267 | -4.1878 | ##### | ####### | 9.8105E-35 | 1.2157E-33 |
| ENSG00000219507\|FTH1P8 | 121.0529 | -2.5702 | ##### | ####### | 1.0452E-34 | 1.2947E-33 |
| ENSG00000105519\|CAPS | 292.7712 | -2.3879 | ##### | ####### | 1.0613E-34 | 1.3141E-33 |
| ENSG00000162695\|SLC30A7 | 2817.0108 | -1.5472 | ##### | ####### | 1.1077E-34 | 1.3710E-33 |
| ENSG00000127666\|TICAM1 | 2707.3490 | 1.5412 | ##### | 12.2832 | 1.1147E-34 | 1.3791E-33 |
| ENSG00000051108\|HERPUD1 | 4094.4979 | -1.5015 | ##### | ####### | 1.1701E-34 | 1.4470E-33 |
| ENSG00000116539\|ASH1L | 9103.2863 | 1.5308 | ##### | 12.2779 | 1.1910E-34 | 1.4723E-33 |
| ENSG00000007402\|CACNA2D2 | 208.1543 | 2.3121 | ##### | 12.2726 | 1.2708E-34 | 1.5702E-33 |
| ENSG00000118596\|SLC16A7 | 3871.0092 | -1.5652 | ##### | ####### | 1.3050E-34 | 1.6119E-33 |
| ENSG00000125995\|ROMO1 | 7664.3415 | 1.5239 | ##### | 12.2703 | 1.3083E-34 | 1.6152E-33 |
| ENSG00000111790\|FGFR1OP2 | 4786.8526 | 1.4838 | ##### | 12.2689 | 1.3312E-34 | 1.6429E-33 |
| ENSG00000258366\|RTEL1 | 3209.0301 | 1.5559 | ##### | 12.2600 | 1.4842E-34 | 1.8307E-33 |
| ENSG00000177383\|MAGEF1 | 2354.7205 | -1.5416 | ##### | ####### | 1.4847E-34 | 1.8307E-33 |
| ENSG00000107485\|GATA3 | 5082.0598 | 1.5582 | ##### | 12.2585 | 1.5123E-34 | 1.8640E-33 |
| ENSG00000137100\|DCTN3 | 3439.2278 | 1.5196 | ##### | 12.2552 | 1.5748E-34 | 1.9403E-33 |
| ENSG00000168071\|CCDC88B | 435.2913 | -1.8412 | ##### | ####### | 1.7873E-34 | 2.2011E-33 |
| ENSG00000100266\|PACSIN2 | 7179.8482 | -1.4437 | ##### | ####### | 1.8160E-34 | 2.2356E-33 |
| ENSG00000237943\|PRKCQ-AS1 | 244.9991 | 2.1983 | ##### | 12.2426 | 1.8405E-34 | 2.2648E-33 |
| ENSG00000241081\|RPL22P2 | 74.8510 | 4.5730 | ##### | 12.2416 | 1.8637E-34 | 2.2924E-33 |
| ENSG00000141759\|TXNL4A | 11258.3415 | 0.8611 | ##### | 12.2395 | 1.9131E-34 | 2.3522E-33 |
| ENSG00000162738\|VANGL2 | 132.5379 | -3.1465 | ##### | ####### | 1.9932E-34 | 2.4497E-33 |
| ENSG00000105287\|PRKD2 | 2804.5197 | -1.2870 | ##### | ####### | 2.1070E-34 | 2.5885E-33 |
| ENSG00000101782\|RIOK3 | 4776.3172 | -1.3543 | ##### | ####### | 2.2280E-34 | 2.7359E-33 |
| ENSG00000271335\|CCNY-AS1 | 130.4517 | 2.5778 | ##### | 12.2268 | 2.2358E-34 | 2.7444E-33 |
| ENSG00000063322\|MED29 | 4729.3316 | -1.4331 | ##### | ####### | 2.2574E-34 | 2.7698E-33 |
| ENSG00000265763\|ZNF488 | 594.4043 | -1.7778 | ##### | ####### | 2.3387E-34 | 2.8684E-33 |
| ENSG00000109184\|DCUN1D4 | 6301.6650 | 1.3763 | ##### | 12.2229 | 2.3446E-34 | 2.8744E-33 |
| ENSG00000127824\|TUBA4A | 16064.6109 | -1.5060 | ##### | ####### | 2.3927E-34 | 2.9322E-33 |
| ENSG00000231721\|LINC-PINT | 1096.6197 | 1.6186 | ##### | 12.2204 | 2.4180E-34 | 2.9620E-33 |
| ENSG00000247626\|MARS2 | 1966.5272 | 1.6052 | ##### | 12.2141 | 2.6143E-34 | 3.2011E-33 |
| ENSG00000135749\|PCNX2 | 1564.4005 | -1.5456 | ##### | ####### | 3.0526E-34 | 3.7363E-33 |
| ENSG00000133740\|E2F5 | 2319.5965 | -1.5863 | ##### | ####### | 3.0699E-34 | 3.7559E-33 |
| ENSG00000057252\|SOAT1 | 5795.1680 | -1.4925 | ##### | ####### | 3.2227E-34 | 3.9413E-33 |
| ENSG00000118407\|FILIP1 | 410.8913 | -4.4241 | ##### | ####### | 3.2393E-34 | 3.9600E-33 |
| ENSG00000237649\|KIFC1 | 6026.1224 | 1.3319 | ##### | 12.1914 | 3.4538E-34 | 4.2205E-33 |
| ENSG00000184640\|SEPTIN9 | 19203.8833 | -1.4099 | ##### | ####### | 3.4559E-34 | 4.2213E-33 |
| ENSG00000014123\|UFL1 | 2477.6661 | -1.3710 | ##### | ####### | 3.5995E-34 | 4.3948E-33 |
| ENSG00000174871\|CNIH2 | 265.5839 | 2.4962 | ##### | 12.1834 | 3.8108E-34 | 4.6510E-33 |
| ENSG00000124177\|CHD6 | 4170.9279 | 1.5194 | ##### | 12.1778 | 4.0833E-34 | 4.9815E-33 |
| ENSG00000250133\|HOXC-AS2 | 79.7497 | 4.5286 | ##### | 12.1705 | 4.4659E-34 | 5.4460E-33 |
| ENSG00000100142\|POLR2F | 4215.9029 | 1.1052 | ##### | 12.1684 | 4.5806E-34 | 5.5837E-33 |
| ENSG00000232677\|LINC00665 | 590.8639 | -1.7854 | ##### | ####### | 4.6857E-34 | 5.7094E-33 |
| ENSG00000197165\|SULT1A2 | 63.3469 | -4.5470 | ##### | ####### | 4.7864E-34 | 5.8297E-33 |
| ENSG00000177707\|NECTIN3 | 2931.3863 | -1.5769 | ##### | ####### | 4.8406E-34 | 5.8934E-33 |
| ENSG00000177706\|FAM20C | 2986.8542 | -1.4928 | ##### | ####### | 5.0152E-34 | 6.1035E-33 |
| ENSG00000272273\|IER3-AS1 | 1654.9553 | 2.2725 | ##### | 12.1527 | 5.5487E-34 | 6.7499E-33 |
| ENSG00000164934\|DCAF13 | 10142.6795 | -1.1902 | ##### | ####### | 5.8828E-34 | 7.1535E-33 |

| | | | | | | |
|---|---|---|---|---|---|---|
| ENSG00000197121\|PGAP1 | 2687.0084 | 1.6129 | ##### | 12.1413 | 6.3841E-34 | 7.7599E-33 |
| ENSG00000124787\|RPP40 | 1303.2437 | 1.1922 | ##### | 12.1388 | 6.5820E-34 | 7.9972E-33 |
| ENSG00000111087\|GLI1 | 944.0528 | -1.6378 | ##### | ####### | 6.8718E-34 | 8.3459E-33 |
| ENSG00000130023\|ERMARD | 685.7444 | -1.6692 | ##### | ####### | 7.1578E-34 | 8.6898E-33 |
| ENSG00000183196\|CHST6 | 391.2383 | 2.2984 | ##### | 12.1268 | 7.6161E-34 | 9.2387E-33 |
| ENSG00000128050\|PAICS | 35287.4354 | 1.3172 | ##### | 12.1268 | 7.6144E-34 | 9.2387E-33 |
| ENSG00000143375\|CGN | 2580.3671 | 1.5399 | ##### | 12.1244 | 7.8422E-34 | 9.5091E-33 |
| ENSG00000101276\|SLC52A3 | 306.8838 | -1.9927 | ##### | ####### | 8.9101E-34 | 1.0799E-32 |
| ENSG00000224807\|DUX4L9 | 271.2373 | 9.9859 | ##### | 12.1133 | 8.9824E-34 | 1.0883E-32 |
| ENSG00000277283\|RAB35-AS1 | 287.8282 | -2.2752 | ##### | ####### | 9.1471E-34 | 1.1078E-32 |
| ENSG00000146090\|RASGEF1C | 66.9479 | 4.4866 | ##### | 12.1116 | 9.1697E-34 | 1.1101E-32 |
| ENSG00000112559\|MDFI | 391.0609 | 2.1183 | ##### | 12.1095 | 9.4059E-34 | 1.1382E-32 |
| ENSG00000184160\|ADRA2C | 1983.7961 | 1.5287 | ##### | 12.1081 | 9.5735E-34 | 1.1580E-32 |
| ENSG00000174749\|FAM241A | 938.8549 | -1.7282 | ##### | ####### | 9.6075E-34 | 1.1616E-32 |
| ENSG00000290399 | 175.0129 | -2.4633 | ##### | ####### | 9.8282E-34 | 1.1879E-32 |
| ENSG00000256040\|PAPPA-AS1 | 164.9834 | -2.4770 | ##### | ####### | 1.0179E-33 | 1.2298E-32 |
| ENSG00000141985\|SH3GL1 | 13408.5780 | 1.0552 | ##### | 12.0940 | 1.1366E-33 | 1.3727E-32 |
| ENSG00000102048\|ASB9 | 574.4384 | -1.5721 | ##### | ####### | 1.2160E-33 | 1.4679E-32 |
| ENSG00000032219\|ARID4A | 1785.3193 | 1.5233 | ##### | 12.0877 | 1.2266E-33 | 1.4800E-32 |
| ENSG00000122971\|ACADS | 601.8228 | -1.7705 | ##### | ####### | 1.2842E-33 | 1.5490E-32 |
| ENSG00000168843\|FSTL5 | 334.1852 | -2.0882 | ##### | ####### | 1.3456E-33 | 1.6224E-32 |
| ENSG00000288612 | 240.4387 | -3.7457 | ##### | ####### | 1.3543E-33 | 1.6322E-32 |
| ENSG00000196214\|ZNF766 | 1271.0609 | -1.6239 | ##### | ####### | 1.3707E-33 | 1.6513E-32 |
| ENSG00000125319\|HROB | 1426.1912 | 1.6231 | ##### | 12.0782 | 1.3765E-33 | 1.6576E-32 |
| ENSG00000148450\|MSRB2 | 958.7650 | -1.6192 | ##### | ####### | 1.5589E-33 | 1.8765E-32 |
| ENSG00000163162\|RNF149 | 3520.8031 | -1.4081 | ##### | ####### | 1.6252E-33 | 1.9556E-32 |
| ENSG00000147649\|MTDH | 15898.5860 | -1.4636 | ##### | ####### | 1.6732E-33 | 2.0124E-32 |
| ENSG00000006377\|DLX6 | 202.7600 | 2.5463 | ##### | 12.0560 | 1.8030E-33 | 2.1677E-32 |
| ENSG00000170266\|GLB1 | 3206.9424 | -1.4938 | ##### | ####### | 1.8250E-33 | 2.1933E-32 |
| ENSG00000215784\|FAM72D | 1389.9914 | 1.5689 | ##### | 12.0516 | 1.9021E-33 | 2.2850E-32 |
| ENSG00000163507\|CIP2A | 7237.5387 | 1.3918 | ##### | 12.0495 | 1.9519E-33 | 2.3439E-32 |
| ENSG00000105220\|GPI | 37848.2718 | -1.4215 | ##### | ####### | 1.9691E-33 | 2.3637E-32 |
| ENSG00000172663\|TMEM134 | 2111.3237 | 1.5829 | ##### | 12.0464 | 2.0262E-33 | 2.4312E-32 |
| ENSG00000184508\|HDDC3 | 802.4120 | -2.1240 | ##### | ####### | 2.0941E-33 | 2.5117E-32 |
| ENSG00000135547\|HEY2 | 209.1406 | 2.4392 | ##### | 12.0404 | 2.1788E-33 | 2.6121E-32 |
| ENSG00000142327\|RNPEPL1 | 5308.8729 | -1.4374 | ##### | ####### | 2.2711E-33 | 2.7218E-32 |
| ENSG00000290994\|FAM86C2P | 1045.4486 | 1.5226 | ##### | 12.0354 | 2.3155E-33 | 2.7738E-32 |
| ENSG00000159461\|AMFR | 7510.6554 | -1.4739 | ##### | ####### | 2.3929E-33 | 2.8655E-32 |
| ENSG00000183779\|ZNF703 | 1045.9924 | -1.6905 | ##### | ####### | 2.4112E-33 | 2.8862E-32 |
| ENSG00000143412\|ANXA9 | 120.6408 | -2.5977 | ##### | ####### | 2.5283E-33 | 3.0251E-32 |
| ENSG00000159208\|CIART | 186.8555 | 2.4713 | ##### | 12.0213 | 2.7462E-33 | 3.2846E-32 |
| ENSG00000162813\|BPNT1 | 2240.0021 | -1.5356 | ##### | ####### | 2.7992E-33 | 3.3466E-32 |
| ENSG00000163577\|EIF5A2 | 1274.4550 | -1.6182 | ##### | ####### | 2.8808E-33 | 3.4427E-32 |
| ENSG00000101751\|POLI | 1356.1759 | -1.6407 | ##### | ####### | 3.2083E-33 | 3.8327E-32 |
| ENSG00000167323\|STIM1 | 4248.6952 | -1.4810 | ##### | ####### | 3.2460E-33 | 3.8762E-32 |
| ENSG00000165389\|SPTSSA | 4412.8014 | -1.4737 | ##### | ####### | 3.4295E-33 | 4.0936E-32 |
| ENSG00000166707\|ZCCHC18 | 277.4584 | 9.8783 | ##### | 12.0013 | 3.4994E-33 | 4.1754E-32 |
| ENSG00000119917\|IFIT3 | 222.1170 | -2.8126 | ##### | ####### | 3.6253E-33 | 4.3239E-32 |
| ENSG00000101442\|ACTR5 | 2482.2926 | 1.4996 | ##### | 11.9956 | 3.7480E-33 | 4.4685E-32 |
| ENSG00000186951\|PPARA | 1266.5810 | -1.6062 | ##### | ####### | 3.8356E-33 | 4.5711E-32 |
| ENSG00000133131\|MORC4 | 3251.7779 | 1.4626 | ##### | 11.9936 | 3.8391E-33 | 4.5734E-32 |
| ENSG00000100985\|MMP9 | 944.9647 | 1.8554 | ##### | 11.9933 | 3.8502E-33 | 4.5848E-32 |
| ENSG00000171867\|PRNP | 16047.0640 | 1.4297 | ##### | 11.9880 | 4.1086E-33 | 4.8905E-32 |
| ENSG00000116857\|TMEM9 | 4551.3325 | -1.4670 | ##### | ####### | 4.3033E-33 | 5.1202E-32 |
| ENSG00000151065\|DCP1B | 1448.7992 | 1.6210 | ##### | 11.9829 | 4.3669E-33 | 5.1938E-32 |
| ENSG00000167671\|UBXN6 | 9222.7573 | 1.4531 | ##### | 11.9826 | 4.3857E-33 | 5.2142E-32 |
| ENSG00000099849\|RASSF7 | 1770.5614 | -1.5594 | ##### | ####### | 4.6296E-33 | 5.5020E-32 |
| ENSG00000150593\|PDCD4 | 2972.5688 | -1.5739 | ##### | ####### | 4.7414E-33 | 5.6325E-32 |
| ENSG00000130635\|COL5A1 | 1524.9083 | -1.9785 | ##### | ####### | 4.9259E-33 | 5.8494E-32 |
| ENSG00000178381\|ZFAND2A | 2496.1779 | -1.5572 | ##### | ####### | 5.0523E-33 | 5.9971E-32 |
| ENSG00000100092\|SH3BP1 | 1272.9365 | 2.1372 | ##### | 11.9680 | 5.2288E-33 | 6.2041E-32 |
| ENSG00000175938\|ORAI3 | 705.0472 | -1.8375 | ##### | ####### | 5.3104E-33 | 6.2985E-32 |
| ENSG00000291087\|CRYBB2P1 | 750.2736 | 1.7492 | ##### | 11.9633 | 5.5348E-33 | 6.5620E-32 |

| | | | | | | |
|---|---|---|---|---|---|---|
| ENSG00000133731\|IMPA1 | 3226.1005 | -1.4810 | ##### | ####### | 5.8149E-33 | 6.8914E-32 |
| ENSG00000081019\|RSBN1 | 3225.3117 | 1.4578 | ##### | 11.9572 | 5.9554E-33 | 7.0551E-32 |
| ENSG00000155313\|USP25 | 3302.6291 | -1.4997 | ##### | ####### | 5.9922E-33 | 7.0959E-32 |
| ENSG00000064703\|DDX20 | 4781.4912 | 1.5107 | ##### | 11.9557 | 6.0640E-33 | 7.1780E-32 |
| ENSG00000161921\|CXCL16 | 1340.0598 | -1.5735 | ##### | ####### | 6.0792E-33 | 7.1932E-32 |
| ENSG00000100302\|RASD2 | 800.2978 | 1.7494 | ##### | 11.9529 | 6.2714E-33 | 7.4176E-32 |
| ENSG00000111364\|DDX55 | 4028.1952 | 0.8420 | ##### | 11.9405 | 7.2771E-33 | 8.6038E-32 |
| ENSG00000160888\|IER2 | 23741.2036 | 1.4229 | ##### | 11.9383 | 7.4699E-33 | 8.8281E-32 |
| ENSG00000140511\|HAPLN3 | 777.8124 | -1.7260 | ##### | ####### | 7.7873E-33 | 9.1997E-32 |
| ENSG00000181652\|ATG9B | 297.6070 | 3.5416 | ##### | 11.9310 | 8.1605E-33 | 9.6368E-32 |
| ENSG00000162924\|REL | 4712.3585 | 1.5744 | ##### | 11.9274 | 8.5179E-33 | 1.0055E-31 |
| ENSG00000135111\|TBX3 | 1207.2905 | 1.8989 | ##### | 11.9262 | 8.6456E-33 | 1.0202E-31 |
| ENSG00000172137\|CALB2 | 365.5348 | -4.6405 | ##### | ####### | 8.6782E-33 | 1.0236E-31 |
| ENSG00000163382\|NAXE | 25370.9000 | 1.6087 | ##### | 11.9211 | 9.1905E-33 | 1.0836E-31 |
| ENSG00000277991 | 428.2412 | -1.7230 | ##### | ####### | 9.2496E-33 | 1.0901E-31 |
| ENSG00000259781\|HMGB1P6 | 5477.7862 | 1.5255 | ##### | 11.9169 | 9.6668E-33 | 1.1388E-31 |
| ENSG00000110628\|SLC22A18 | 1097.7078 | -1.5544 | ##### | ####### | 9.6981E-33 | 1.1421E-31 |
| ENSG00000117519\|CNN3 | 7387.0718 | -1.4240 | ##### | ####### | 9.9478E-33 | 1.1710E-31 |
| ENSG00000278599\|TBC1D3E | 265.0913 | 2.2844 | ##### | 11.9143 | 9.9678E-33 | 1.1729E-31 |
| ENSG00000116514\|RNF19B | 3551.2864 | -1.3542 | ##### | ####### | 1.0559E-32 | 1.2420E-31 |
| ENSG00000142156\|COL6A1 | 3809.3782 | -1.4669 | ##### | ####### | 1.0785E-32 | 1.2681E-31 |
| ENSG00000165156\|ZHX1 | 2679.5952 | -1.5478 | ##### | ####### | 1.1603E-32 | 1.3638E-31 |
| ENSG00000161082\|CELF5 | 382.6078 | 2.4163 | ##### | 11.8897 | 1.3389E-32 | 1.5731E-31 |
| ENSG00000224536\|CSRP1-AS1 | 148.9516 | -2.6787 | ##### | ####### | 1.3817E-32 | 1.6226E-31 |
| ENSG00000138778\|CENPE | 5527.6522 | 1.5159 | ##### | 11.8841 | 1.4315E-32 | 1.6805E-31 |
| ENSG00000176225\|RTTN | 2919.6006 | 1.4211 | ##### | 11.8831 | 1.4488E-32 | 1.7001E-31 |
| ENSG00000141447\|OSBPL1A | 2519.7261 | -1.5674 | ##### | ####### | 1.4756E-32 | 1.7309E-31 |
| ENSG00000163412\|EIF4E3 | 134.4316 | -3.1623 | ##### | ####### | 1.5019E-32 | 1.7610E-31 |
| ENSG00000153879\|CEBPG | 4688.8169 | -1.4471 | ##### | ####### | 1.5266E-32 | 1.7893E-31 |
| ENSG00000254402\|LRRC24 | 497.9820 | -1.8527 | ##### | ####### | 1.5440E-32 | 1.8090E-31 |
| ENSG00000147872\|PLIN2 | 1548.0795 | -1.5109 | ##### | ####### | 1.6183E-32 | 1.8953E-31 |
| ENSG00000070087\|PFN2 | 13037.1339 | -1.4299 | ##### | ####### | 1.6951E-32 | 1.9845E-31 |
| ENSG00000159231\|CBR3 | 564.2879 | -1.8199 | ##### | ####### | 1.7206E-32 | 2.0135E-31 |
| ENSG00000137496\|IL18BP | 1941.2540 | 1.6161 | ##### | 11.8606 | 1.8966E-32 | 2.2186E-31 |
| ENSG00000266916\|ZNF793-AS1 | 98.7946 | -4.0192 | ##### | ####### | 1.8973E-32 | 2.2186E-31 |
| ENSG00000061918\|GUCY1B1 | 442.2533 | 2.2872 | ##### | 11.8602 | 1.9044E-32 | 2.2260E-31 |
| ENSG00000140577\|CRTC3 | 2483.2697 | -1.5516 | ##### | ####### | 1.9394E-32 | 2.2660E-31 |
| ENSG00000139428\|MMAB | 1393.0124 | -1.5134 | ##### | ####### | 1.9691E-32 | 2.2998E-31 |
| ENSG00000115042\|FAHD2A | 1806.8490 | 1.5570 | ##### | 11.8553 | 2.0205E-32 | 2.3590E-31 |
| ENSG00000120709\|FAM53C | 6659.2559 | 1.9281 | ##### | 11.8546 | 2.0373E-32 | 2.3777E-31 |
| ENSG00000138336\|TET1 | 171.1858 | -2.6248 | ##### | ####### | 2.0446E-32 | 2.3852E-31 |
| ENSG00000255046 | 170.0857 | -5.5542 | ##### | ####### | 2.1259E-32 | 2.4792E-31 |
| ENSG00000171992\|SYNPO | 6118.4847 | -1.4283 | ##### | ####### | 2.1788E-32 | 2.5398E-31 |
| ENSG00000256671\|LIMS4 | 200.0418 | -2.5327 | ##### | ####### | 2.3550E-32 | 2.7441E-31 |
| ENSG00000055208\|TAB2 | 10714.8046 | 1.4361 | ##### | 11.8423 | 2.3579E-32 | 2.7465E-31 |
| ENSG00000278996 | 159.7606 | 2.3191 | ##### | 11.8372 | 2.5060E-32 | 2.9178E-31 |
| ENSG00000162241\|SLC25A45 | 604.2692 | 1.6947 | ##### | 11.8361 | 2.5406E-32 | 2.9570E-31 |
| ENSG00000175606\|TMEM70 | 2380.3164 | -1.4825 | ##### | ####### | 2.6122E-32 | 3.0391E-31 |
| ENSG00000245910\|SNHG6 | 6054.7972 | -1.4621 | ##### | ####### | 2.6377E-32 | 3.0676E-31 |
| ENSG00000167797\|CDK2AP2 | 6425.4427 | 1.4516 | ##### | 11.8311 | 2.6952E-32 | 3.1332E-31 |
| ENSG00000125962\|ARMCX5 | 1900.0579 | 1.5580 | ##### | 11.8281 | 2.7924E-32 | 3.2450E-31 |
| ENSG00000108639\|SYNGR2 | 20658.7696 | -1.4076 | ##### | ####### | 3.1126E-32 | 3.6156E-31 |
| ENSG00000156508\|EEF1A1 | ########## | 1.0798 | ##### | 11.8189 | 3.1182E-32 | 3.6208E-31 |
| ENSG00000124207\|CSE1L | 28366.0028 | 1.2284 | ##### | 11.8070 | 3.5910E-32 | 4.1681E-31 |
| ENSG00000012171\|SEMA3B | 227.5056 | -2.0134 | ##### | ####### | 3.6593E-32 | 4.2457E-31 |
| ENSG00000100949\|RABGGTA | 2212.3034 | 1.5370 | ##### | 11.8038 | 3.7293E-32 | 4.3253E-31 |
| ENSG00000221947\|XKR9 | 87.6725 | -6.7582 | ##### | ####### | 3.7799E-32 | 4.3822E-31 |
| ENSG00000124214\|STAU1 | 16126.6094 | 1.2689 | ##### | 11.8019 | 3.8140E-32 | 4.4201E-31 |
| ENSG00000260280\|SLX1B-SULT1A4 | 1406.3124 | 1.6756 | ##### | 11.8001 | 3.8996E-32 | 4.5176E-31 |
| ENSG00000146233\|CYP39A1 | 174.8139 | -4.3805 | ##### | ####### | 4.1522E-32 | 4.8083E-31 |
| ENSG00000148680\|HTR7 | 204.2166 | 3.2374 | ##### | 11.7936 | 4.2129E-32 | 4.8766E-31 |
| ENSG00000181481\|RNF135 | 2054.7569 | -1.5103 | ##### | ####### | 4.2420E-32 | 4.9084E-31 |
| ENSG00000230082\|PRRT3-AS1 | 264.1278 | -2.0950 | ##### | ####### | 4.5704E-32 | 5.2864E-31 |

| | | | | | | |
|---|---|---|---|---|---|---|
| ENSG00000146067\|FAM193B | 2087.3410 | 1.5314 | ##### | 11.7861 | 4.6015E-32 | 5.3203E-31 |
| ENSG00000105656\|ELL | 2293.4332 | 1.1889 | ##### | 11.7859 | 4.6156E-32 | 5.3345E-31 |
| ENSG00000119392\|GLE1 | 4936.2089 | -0.7319 | ##### | ####### | 4.6555E-32 | 5.3786E-31 |
| ENSG00000005187\|ACSM3 | 400.1156 | -4.1219 | ##### | ####### | 4.6606E-32 | 5.3823E-31 |
| ENSG00000128656\|CHN1 | 1925.2833 | 1.4884 | ##### | 11.7806 | 4.9126E-32 | 5.6712E-31 |
| ENSG00000227218 | 89.6272 | -3.5982 | ##### | ####### | 5.1988E-32 | 5.9993E-31 |
| ENSG00000172009\|THOP1 | 11421.2259 | 1.3608 | ##### | 11.7739 | 5.3196E-32 | 6.1363E-31 |
| ENSG00000111913\|RIPOR2 | 286.0390 | 9.8125 | ##### | 11.7710 | 5.5061E-32 | 6.3490E-31 |
| ENSG00000104915\|STX10 | 5084.2184 | 1.4441 | ##### | 11.7703 | 5.5543E-32 | 6.4021E-31 |
| ENSG00000079337\|RAPGEF3 | 407.4939 | -1.8524 | ##### | ####### | 5.6972E-32 | 6.5644E-31 |
| ENSG00000146247\|PHIP | 7327.6704 | 1.4112 | ##### | 11.7673 | 5.7532E-32 | 6.6263E-31 |
| ENSG00000204616\|TRIM31 | 194.1108 | -5.1041 | ##### | ####### | 5.8321E-32 | 6.7145E-31 |
| ENSG00000205593\|DENND6B | 341.9567 | -2.0963 | ##### | ####### | 6.0074E-32 | 6.9138E-31 |
| ENSG00000245248\|USP2-AS1 | 392.5298 | 1.9183 | ##### | 11.7625 | 6.0940E-32 | 7.0106E-31 |
| ENSG00000174136\|RGMB | 1857.0192 | 1.5183 | ##### | 11.7527 | 6.8391E-32 | 7.8648E-31 |
| ENSG00000155034\|FBXL18 | 8908.2055 | -1.4359 | ##### | ####### | 6.9691E-32 | 8.0112E-31 |
| ENSG00000088179\|PTPN4 | 1880.9439 | -1.4738 | ##### | ####### | 7.4828E-32 | 8.5985E-31 |
| ENSG00000110697\|PITPNM1 | 11113.4925 | 1.3346 | ##### | 11.7432 | 7.6556E-32 | 8.7936E-31 |
| ENSG00000174628\|IQCK | 770.9005 | -1.5630 | ##### | ####### | 8.0010E-32 | 9.1868E-31 |
| ENSG00000115380\|EFEMP1 | 9609.3221 | -1.0887 | ##### | ####### | 8.0864E-32 | 9.2813E-31 |
| ENSG00000114853\|ZBTB47 | 686.1193 | -1.7368 | ##### | ####### | 8.1553E-32 | 9.3568E-31 |
| ENSG00000251136\|RIPK2-DT | 358.9345 | -1.9008 | ##### | ####### | 8.4712E-32 | 9.7156E-31 |
| ENSG00000170037\|CNTROB | 4213.9334 | 1.4288 | ##### | 11.7344 | 8.4966E-32 | 9.7409E-31 |
| ENSG00000161036\|LRWD1 | 5776.1193 | 1.3965 | ##### | 11.7310 | 8.8455E-32 | 1.0137E-30 |
| ENSG00000290720\|ZNF542P | 1998.9611 | -1.5214 | ##### | ####### | 9.1499E-32 | 1.0482E-30 |
| ENSG00000037749\|MFAP3 | 5669.9112 | -1.4250 | ##### | ####### | 9.9161E-32 | 1.1355E-30 |
| ENSG00000290872 | 426.6727 | 2.0735 | ##### | 11.7211 | 9.9377E-32 | 1.1376E-30 |
| ENSG00000137714\|FDX1 | 2017.0268 | -1.4788 | ##### | ####### | 9.9417E-32 | 1.1376E-30 |
| ENSG00000132694\|ARHGEF11 | 9726.8005 | 1.3961 | ##### | 11.7183 | 1.0269E-31 | 1.1745E-30 |
| ENSG00000245648\|KLRK1-AS1 | 368.8574 | 4.9289 | ##### | 11.7135 | 1.0868E-31 | 1.2426E-30 |
| ENSG00000139832\|RAB20 | 153.1146 | -2.6770 | ##### | ####### | 1.0917E-31 | 1.2477E-30 |
| ENSG00000106571\|GLI3 | 266.2858 | 3.1598 | ##### | 11.7106 | 1.1248E-31 | 1.2851E-30 |
| ENSG00000158796\|DEDD | 6022.2992 | 1.2793 | ##### | 11.7100 | 1.1328E-31 | 1.2938E-30 |
| ENSG00000167693\|NXN | 4316.0012 | 1.4843 | ##### | 11.7052 | 1.1981E-31 | 1.3678E-30 |
| ENSG00000152253\|SPC25 | 1485.2260 | 1.8248 | ##### | 11.7043 | 1.2112E-31 | 1.3822E-30 |
| ENSG00000232187\|FTH1P7 | 107.2138 | -2.7573 | ##### | ####### | 1.2744E-31 | 1.4538E-30 |
| ENSG00000267809\|NDUFV2P1 | 107.2271 | 3.0283 | ##### | 11.6986 | 1.2953E-31 | 1.4770E-30 |
| ENSG00000135776\|ABCB10 | 2338.5275 | -1.4322 | ##### | ####### | 1.3178E-31 | 1.5021E-30 |
| ENSG00000146830\|GIGYF1 | 5643.5340 | 1.3676 | ##### | 11.6956 | 1.3423E-31 | 1.5295E-30 |
| ENSG00000186432\|KPNA4 | 18301.3687 | 1.1560 | ##### | 11.6950 | 1.3519E-31 | 1.5398E-30 |
| ENSG00000126217\|MCF2L | 110.5419 | 3.5008 | ##### | 11.6944 | 1.3611E-31 | 1.5498E-30 |
| ENSG00000285872 | 123.3574 | 2.6113 | ##### | 11.6942 | 1.3654E-31 | 1.5540E-30 |
| ENSG00000182552\|RWDD4 | 2028.1672 | -1.2909 | ##### | ####### | 1.5130E-31 | 1.7214E-30 |
| ENSG00000143367\|TUFT1 | 5013.8213 | -1.4450 | ##### | ####### | 1.5762E-31 | 1.7926E-30 |
| ENSG00000230487\|PSMG3-AS1 | 1074.2798 | -1.5776 | ##### | ####### | 1.7286E-31 | 1.9652E-30 |
| ENSG00000106692\|FKTN | 2715.1376 | -1.4207 | ##### | ####### | 1.7839E-31 | 2.0272E-30 |
| ENSG00000175701\|MTLN | 2170.1103 | -1.5070 | ##### | ####### | 1.8123E-31 | 2.0587E-30 |
| ENSG00000154447\|SH3RF1 | 1372.2690 | -1.6999 | ##### | ####### | 1.8697E-31 | 2.1232E-30 |
| ENSG00000004487\|KDM1A | 10704.1016 | 0.8624 | ##### | 11.6646 | 1.9336E-31 | 2.1949E-30 |
| ENSG00000188227\|ZNF793 | 334.1337 | -2.6391 | ##### | ####### | 1.9616E-31 | 2.2258E-30 |
| ENSG00000293473\|CENPBD1P | 1249.1641 | -1.5569 | ##### | ####### | 1.9698E-31 | 2.2343E-30 |
| ENSG00000274419\|TBC1D3D | 346.6000 | 2.0611 | ##### | 11.6609 | 2.0191E-31 | 2.2893E-30 |
| ENSG00000241935\|HOGA1 | 76.7715 | -4.6446 | ##### | ####### | 2.0247E-31 | 2.2948E-30 |
| ENSG00000185187\|SIGIRR | 1050.0089 | -1.6374 | ##### | ####### | 2.0884E-31 | 2.3661E-30 |
| ENSG00000027697\|IFNGR1 | 4534.7612 | -1.4427 | ##### | ####### | 2.1055E-31 | 2.3845E-30 |
| ENSG00000167178\|ISLR2 | 194.9499 | -2.7226 | ##### | ####### | 2.1418E-31 | 2.4248E-30 |
| ENSG00000293402 | 191.8710 | 3.1285 | ##### | 11.6501 | 2.2907E-31 | 2.5923E-30 |
| ENSG00000135976\|ANKRD36 | 502.5487 | 1.7899 | ##### | 11.6371 | 2.6681E-31 | 3.0183E-30 |
| ENSG00000140534\|TICRR | 2394.6603 | 1.4743 | ##### | 11.6366 | 2.6852E-31 | 3.0364E-30 |
| ENSG00000090989\|EXOC1 | 5730.1259 | 1.3736 | ##### | 11.6359 | 2.7073E-31 | 3.0603E-30 |
| ENSG00000260691\|ANKRD20A1 | 57.9890 | -6.8661 | ##### | ####### | 3.0429E-31 | 3.4384E-30 |
| ENSG00000285935 | 309.6041 | 2.3842 | ##### | 11.6254 | 3.0598E-31 | 3.4562E-30 |
| ENSG00000169692\|AGPAT2 | 4490.8279 | -1.4253 | ##### | ####### | 3.0803E-31 | 3.4780E-30 |

| | | | | | | |
|---|---|---|---|---|---|---|
| ENSG00000176945\|MUC20 | 116.5266 | -3.3502 | ##### | ####### | 3.0834E-31 | 3.4802E-30 |
| ENSG00000115758\|ODC1 | 41021.7711 | 1.4221 | ##### | 11.6222 | 3.1776E-31 | 3.5852E-30 |
| ENSG00000130638\|ATXN10 | 11915.0942 | -1.3916 | ##### | ####### | 3.2903E-31 | 3.7095E-30 |
| ENSG00000135297\|MTO1 | 3617.5680 | 1.4938 | ##### | 11.6193 | 3.2899E-31 | 3.7095E-30 |
| ENSG00000293413 | 415.9023 | 2.3890 | ##### | 11.6130 | 3.5382E-31 | 3.9875E-30 |
| ENSG00000145819\|ARHGAP26 | 2127.7945 | -1.4376 | ##### | ####### | 3.6306E-31 | 4.0901E-30 |
| ENSG00000235609 | 265.1735 | -2.2688 | ##### | ####### | 3.6919E-31 | 4.1576E-30 |
| ENSG00000274226\|TBC1D3H | 198.8124 | 2.1315 | ##### | 11.6084 | 3.7333E-31 | 4.2026E-30 |
| ENSG00000204174\|NPY4R | 191.3167 | -5.2512 | ##### | ####### | 3.7489E-31 | 4.2186E-30 |
| ENSG00000171793\|CTPS1 | 22164.1786 | -1.4625 | ##### | ####### | 3.8258E-31 | 4.3035E-30 |
| ENSG00000153707\|PTPRD | 703.4404 | 6.2813 | ##### | 11.6027 | 3.9914E-31 | 4.4881E-30 |
| ENSG00000069849\|ATP1B3 | 21590.0032 | 1.3760 | ##### | 11.5980 | 4.2162E-31 | 4.7390E-30 |
| ENSG00000130816\|DNMT1 | 25129.0201 | 1.3536 | ##### | 11.5964 | 4.2961E-31 | 4.8271E-30 |
| ENSG00000088881\|EBF4 | 232.2752 | 2.2441 | ##### | 11.5959 | 4.3249E-31 | 4.8576E-30 |
| ENSG00000126870\|DYNC2I1 | 2843.4046 | 1.3763 | ##### | 11.5950 | 4.3664E-31 | 4.9024E-30 |
| ENSG00000168939\|SPRY3 | 181.4648 | -2.2533 | ##### | ####### | 4.6475E-31 | 5.2160E-30 |
| ENSG00000225630\|MTND2P28 | 35614.0158 | 1.4889 | ##### | 11.5843 | 4.9473E-31 | 5.5504E-30 |
| ENSG00000011638\|LDAF1 | 849.3348 | -1.6898 | ##### | ####### | 4.9975E-31 | 5.6046E-30 |
| ENSG00000259498\|TPM1-AS | 731.7159 | 2.0117 | ##### | 11.5832 | 5.0168E-31 | 5.6241E-30 |
| ENSG00000183763\|TRAIP | 1198.9565 | 1.6111 | ##### | 11.5829 | 5.0311E-31 | 5.6381E-30 |
| ENSG00000288752 | 166.4217 | -2.9901 | ##### | ####### | 5.0520E-31 | 5.6594E-30 |
| ENSG00000159840\|ZYX | 54825.1077 | 1.4487 | ##### | 11.5813 | 5.1284E-31 | 5.7427E-30 |
| ENSG00000104765\|BNIP3L | 5927.0355 | 1.4210 | ##### | 11.5810 | 5.1439E-31 | 5.7579E-30 |
| ENSG00000163513\|TGFBR2 | 6602.3641 | -1.3779 | ##### | ####### | 5.1804E-31 | 5.7967E-30 |
| ENSG00000245526\|LINC00461 | 321.4416 | 10.1607 | ##### | 11.5776 | 5.3500E-31 | 5.9841E-30 |
| ENSG00000145882\|PCYOX1L | 1106.3655 | -1.6860 | ##### | ####### | 5.4022E-31 | 6.0403E-30 |
| ENSG00000170099\|SERPINA6 | 213.2991 | ####### | ##### | ####### | 5.5057E-31 | 6.1538E-30 |
| ENSG00000242173\|ARHGDIG | 88.8385 | -4.1254 | ##### | ####### | 5.5366E-31 | 6.1860E-30 |
| ENSG00000153815\|CMIP | 12809.9663 | -0.8984 | ##### | ####### | 5.6292E-31 | 6.2870E-30 |
| ENSG00000137449\|CPEB2 | 1562.1995 | -1.4717 | ##### | ####### | 6.1357E-31 | 6.8502E-30 |
| ENSG00000290678 | 79.2528 | -3.7988 | ##### | ####### | 6.3215E-31 | 7.0550E-30 |
| ENSG00000149243\|KLHL35 | 593.0134 | 1.6701 | ##### | 11.5610 | 6.4956E-31 | 7.2466E-30 |
| ENSG00000151458\|ANKRD50 | 3571.1896 | 1.4648 | ##### | 11.5588 | 6.6635E-31 | 7.4311E-30 |
| ENSG00000125285\|SOX21 | 110.2213 | -3.2806 | ##### | ####### | 7.1804E-31 | 8.0046E-30 |
| ENSG00000171408\|PDE7B | 670.8872 | -1.7593 | ##### | ####### | 7.1952E-31 | 8.0181E-30 |
| ENSG00000198805\|PNP | 12783.0358 | 1.4361 | ##### | 11.5462 | 7.7173E-31 | 8.5967E-30 |
| ENSG00000235721 | 95.9119 | -3.2722 | ##### | ####### | 7.8115E-31 | 8.6983E-30 |
| ENSG00000130066\|SAT1 | 11346.0728 | 1.4314 | ##### | 11.5438 | 7.9360E-31 | 8.8337E-30 |
| ENSG00000150347\|ARID5B | 1701.1411 | 1.5166 | ##### | 11.5431 | 7.9992E-31 | 8.9008E-30 |
| ENSG00000230212\|CBR1-AS1 | 1183.5001 | -1.5894 | ##### | ####### | 8.0859E-31 | 8.9939E-30 |
| ENSG00000121671\|CRY2 | 744.2590 | 1.4584 | ##### | 11.5405 | 8.2440E-31 | 9.1663E-30 |
| ENSG00000293468\|ATE1OSP | 61.8872 | -3.8518 | ##### | ####### | 8.3715E-31 | 9.3046E-30 |
| ENSG00000150054\|MPP7 | 698.7007 | 1.8480 | ##### | 11.5297 | 9.3450E-31 | 1.0383E-29 |
| ENSG00000107829\|FBXW4 | 1599.2585 | -1.4294 | ##### | ####### | 9.8396E-31 | 1.0928E-29 |
| ENSG00000103381\|CPPED1 | 1281.4939 | -1.4336 | ##### | ####### | 1.0084E-30 | 1.1195E-29 |
| ENSG00000156697\|UTP14A | 6992.2233 | 1.2841 | ##### | 11.5197 | 1.0495E-30 | 1.1647E-29 |
| ENSG00000154582\|ELOC | 5518.1453 | -1.4152 | ##### | ####### | 1.1087E-30 | 1.2300E-29 |
| ENSG00000013810\|TACC3 | 10879.7882 | 1.3906 | ##### | 11.5149 | 1.1093E-30 | 1.2301E-29 |
| ENSG00000147669\|POLR2K | 3423.5696 | -1.0425 | ##### | ####### | 1.1205E-30 | 1.2421E-29 |
| ENSG00000154080\|CHST9 | 57.3511 | -4.8610 | ##### | ####### | 1.2142E-30 | 1.3456E-29 |
| ENSG00000166396\|SERPINB7 | 282.0022 | 2.2439 | ##### | 11.5067 | 1.2207E-30 | 1.3523E-29 |
| ENSG00000162688\|AGL | 2419.8802 | -1.5046 | ##### | ####### | 1.2729E-30 | 1.4090E-29 |
| ENSG00000166257\|SCN3B | 227.3068 | 2.3804 | ##### | 11.5031 | 1.2727E-30 | 1.4090E-29 |
| ENSG00000259187 | 143.3756 | 2.2036 | ##### | 11.5012 | 1.3016E-30 | 1.4402E-29 |
| ENSG00000100916\|BRMS1L | 1607.2193 | 1.5108 | ##### | 11.4965 | 1.3734E-30 | 1.5192E-29 |
| ENSG00000163660\|CCNL1 | 4010.1553 | 1.4102 | ##### | 11.4951 | 1.3957E-30 | 1.5432E-29 |
| ENSG00000079841\|RIMS1 | 68.1219 | 5.1971 | ##### | 11.4903 | 1.4752E-30 | 1.6306E-29 |
| ENSG00000132356\|PRKAA1 | 15547.3877 | 1.3661 | ##### | 11.4888 | 1.5024E-30 | 1.6600E-29 |
| ENSG00000128567\|PODXL | 2653.8064 | 1.4008 | ##### | 11.4879 | 1.5174E-30 | 1.6760E-29 |
| ENSG00000168916\|ZNF608 | 350.8196 | 2.0147 | ##### | 11.4875 | 1.5253E-30 | 1.6840E-29 |
| ENSG00000168993\|CPLX1 | 507.6081 | 2.0497 | ##### | 11.4856 | 1.5584E-30 | 1.7197E-29 |
| ENSG00000106025\|TSPAN12 | 2030.0515 | 1.5574 | ##### | 11.4856 | 1.5587E-30 | 1.7197E-29 |
| ENSG00000233705\|SLC26A4-AS1 | 76.5515 | 5.8212 | ##### | 11.4854 | 1.5621E-30 | 1.7228E-29 |

| | | | | | | |
|---|---|---|---|---|---|---|
| ENSG00000077943\|ITGA8 | 85.2169 | 4.7903 | ##### | 11.4842 | 1.5835E-30 | 1.7457E-29 |
| ENSG00000213123\|DYNLT2B | 464.4120 | -1.7665 | ##### | ####### | 1.5951E-30 | 1.7579E-29 |
| ENSG00000158747\|NBL1 | 4674.2821 | -1.3940 | ##### | ####### | 1.6757E-30 | 1.8460E-29 |
| ENSG00000242732\|RTL5 | 1060.6901 | 1.6104 | ##### | 11.4781 | 1.6991E-30 | 1.8711E-29 |
| ENSG00000280254 | 105.3045 | -2.5519 | ##### | ####### | 1.7081E-30 | 1.8803E-29 |
| ENSG00000186787\|SPIN2B | 370.7315 | -1.7465 | ##### | ####### | 1.7486E-30 | 1.9242E-29 |
| ENSG00000064607\|SUGP2 | 8373.8318 | 1.3572 | ##### | 11.4742 | 1.7777E-30 | 1.9555E-29 |
| ENSG00000196081\|ZNF724 | 141.2397 | -2.5249 | ##### | ####### | 1.7986E-30 | 1.9770E-29 |
| ENSG00000168306\|ACOX2 | 265.3129 | -2.0746 | ##### | ####### | 1.7983E-30 | 1.9770E-29 |
| ENSG00000154059\|IMPACT | 2833.0037 | -1.4439 | ##### | ####### | 1.8420E-30 | 2.0239E-29 |
| ENSG00000101849\|TBL1X | 3161.6654 | -1.3864 | ##### | ####### | 1.8917E-30 | 2.0778E-29 |
| ENSG00000129460\|NGDN | 2945.4451 | 1.2279 | ##### | 11.4680 | 1.9106E-30 | 2.0978E-29 |
| ENSG00000138385\|SSB | 17555.8108 | 1.0370 | ##### | 11.4676 | 1.9180E-30 | 2.1051E-29 |
| ENSG00000173926\|MARCHF3 | 326.5530 | 2.1763 | ##### | 11.4617 | 2.0555E-30 | 2.2553E-29 |
| ENSG00000150630\|VEGFC | 1077.3959 | -1.6096 | ##### | ####### | 2.0911E-30 | 2.2934E-29 |
| ENSG00000148057\|IDNK | 140.0094 | -3.1027 | ##### | ####### | 2.1241E-30 | 2.3288E-29 |
| ENSG00000115084\|SLC35F5 | 8141.4840 | -1.4213 | ##### | ####### | 2.2065E-30 | 2.4183E-29 |
| ENSG00000242539 | 439.5952 | -1.7953 | ##### | ####### | 2.2821E-30 | 2.5001E-29 |
| ENSG00000138650\|PCDH10 | 174.4595 | 2.5914 | ##### | 11.4449 | 2.4951E-30 | 2.7325E-29 |
| ENSG00000177556\|ATOX1 | 2389.8955 | 1.4856 | ##### | 11.4436 | 2.5318E-30 | 2.7717E-29 |
| ENSG00000164885\|CDK5 | 2758.4073 | 1.5031 | ##### | 11.4384 | 2.6886E-30 | 2.9423E-29 |
| ENSG00000171792\|RHNO1 | 4442.0828 | 1.4501 | ##### | 11.4283 | 3.0204E-30 | 3.3042E-29 |
| ENSG00000150403\|TMCO3 | 6022.4243 | -1.4170 | ##### | ####### | 3.0338E-30 | 3.3176E-29 |
| ENSG00000196961\|AP2A1 | 11131.1566 | -1.3096 | ##### | ####### | 3.1607E-30 | 3.4551E-29 |
| ENSG00000143319\|ISG20L2 | 11627.2068 | 1.3318 | ##### | 11.4233 | 3.1996E-30 | 3.4964E-29 |
| ENSG00000153012\|LGI2 | 2746.4167 | 1.5020 | ##### | 11.4212 | 3.2773E-30 | 3.5800E-29 |
| ENSG00000131620\|ANO1 | 89.7550 | -3.7722 | ##### | ####### | 3.4937E-30 | 3.8150E-29 |
| ENSG00000213906\|LTB4R2 | 511.0114 | 1.9827 | ##### | 11.4154 | 3.5015E-30 | 3.8221E-29 |
| ENSG00000214184\|GCC2-AS1 | 61.3120 | -3.4478 | ##### | ####### | 3.5635E-30 | 3.8883E-29 |
| ENSG00000145246\|ATP10D | 1735.2419 | -1.6215 | ##### | ####### | 3.6543E-30 | 3.9860E-29 |
| ENSG00000124570\|SERPINB6 | 5646.1517 | -1.3049 | ##### | ####### | 3.6610E-30 | 3.9918E-29 |
| ENSG00000170439\|TMT1B | 1011.2791 | -1.5046 | ##### | ####### | 3.6738E-30 | 4.0043E-29 |
| ENSG00000176401\|EID2B | 482.6507 | -1.8865 | ##### | ####### | 3.7298E-30 | 4.0639E-29 |
| ENSG00000181467\|RAP2B | 3428.7150 | 1.3583 | ##### | 11.4096 | 3.7458E-30 | 4.0798E-29 |
| ENSG00000185753\|CXorf38 | 1137.1870 | -1.5047 | ##### | ####### | 3.7786E-30 | 4.1140E-29 |
| ENSG00000091136\|LAMB1 | 21790.8675 | -1.3238 | ##### | ####### | 3.8629E-30 | 4.2043E-29 |
| ENSG00000145833\|DDX46 | 13038.1287 | 1.3845 | ##### | 11.4029 | 4.0434E-30 | 4.3992E-29 |
| ENSG00000125741\|OPA3 | 5070.1929 | -1.3828 | ##### | ####### | 4.1011E-30 | 4.4603E-29 |
| ENSG00000130005\|GAMT | 1553.5405 | 1.5174 | ##### | 11.4001 | 4.1744E-30 | 4.5383E-29 |
| ENSG00000255571\|MIR9-3HG | 185.2638 | 2.6705 | ##### | 11.3956 | 4.3958E-30 | 4.7773E-29 |
| ENSG00000137142\|IGFBPL1 | 1188.5520 | 1.5494 | ##### | 11.3953 | 4.4111E-30 | 4.7922E-29 |
| ENSG00000167074\|TEF | 387.5537 | 2.1178 | ##### | 11.3934 | 4.5088E-30 | 4.8966E-29 |
| ENSG00000086589\|RBM22 | 4651.0332 | 0.8631 | ##### | 11.3928 | 4.5408E-30 | 4.9296E-29 |
| ENSG00000175395\|ZNF25 | 510.1564 | -1.6671 | ##### | ####### | 4.5852E-30 | 4.9759E-29 |
| ENSG00000125868\|DSTN | 29530.0655 | -1.4067 | ##### | ####### | 4.7609E-30 | 5.1647E-29 |
| ENSG00000174705\|SH3PXD2B | 3856.6753 | 1.4552 | ##### | 11.3870 | 4.8544E-30 | 5.2642E-29 |
| ENSG00000123810\|B9D2 | 460.7084 | -1.5886 | ##### | ####### | 5.0513E-30 | 5.4758E-29 |
| ENSG00000179981\|TSHZ1 | 761.7634 | -1.6336 | ##### | ####### | 5.5201E-30 | 5.9817E-29 |
| ENSG00000133275\|CSNK1G2 | 8847.5186 | 0.9911 | ##### | 11.3739 | 5.6378E-30 | 6.1071E-29 |
| ENSG00000225206\|MIR137HG | 595.2045 | -2.2103 | ##### | ####### | 5.7159E-30 | 6.1895E-29 |
| ENSG00000149809\|TM7SF2 | 877.2089 | -1.6984 | ##### | ####### | 5.8205E-30 | 6.3005E-29 |
| ENSG00000010810\|FYN | 802.8352 | 1.6891 | ##### | 11.3696 | 5.9263E-30 | 6.4126E-29 |
| ENSG00000099889\|ARVCF | 336.4654 | 1.8437 | ##### | 11.3680 | 6.0362E-30 | 6.5292E-29 |
| ENSG00000079335\|CDC14A | 1245.8573 | 1.5543 | ##### | 11.3650 | 6.2437E-30 | 6.7512E-29 |
| ENSG00000198753\|PLXNB3 | 368.7870 | 1.9460 | ##### | 11.3643 | 6.2980E-30 | 6.8075E-29 |
| ENSG00000175305\|CCNE2 | 1004.5081 | -1.8510 | ##### | ####### | 6.6531E-30 | 7.1887E-29 |
| ENSG00000246763\|RGMB-AS1 | 353.7507 | 10.0954 | ##### | 11.3583 | 6.7440E-30 | 7.2842E-29 |
| ENSG00000139289\|PHLDA1 | 21908.5479 | 0.8032 | ##### | 11.3573 | 6.8245E-30 | 7.3685E-29 |
| ENSG00000146083\|RNF44 | 6652.7174 | 0.8310 | ##### | 11.3525 | 7.2112E-30 | 7.7833E-29 |
| ENSG00000103226\|NOMO3 | 7487.3044 | -1.3896 | ##### | ####### | 7.5948E-30 | 8.1943E-29 |
| ENSG00000198964\|SGMS1 | 2437.8566 | -1.1814 | ##### | ####### | 7.6433E-30 | 8.2437E-29 |
| ENSG00000158987\|RAPGEF6 | 3046.9311 | 1.4428 | ##### | 11.3450 | 7.8513E-30 | 8.4650E-29 |
| ENSG00000163904\|SENP2 | 2793.7140 | -1.4319 | ##### | ####### | 8.1407E-30 | 8.7738E-29 |

| | | | | | | |
|---|---|---|---|---|---|---|
| ENSG00000198589\|LRBA | 3799.0722 | -1.3957 | ##### | ####### | 8.5469E-30 | 9.2083E-29 |
| ENSG00000073737\|DHRS9 | 101.1651 | -3.2344 | ##### | ####### | 8.5946E-30 | 9.2563E-29 |
| ENSG00000232759\|STEAP1B-AS1 | 130.9982 | 3.8221 | ##### | 11.3362 | 8.6843E-30 | 9.3495E-29 |
| ENSG00000139618\|BRCA2 | 2067.9309 | 1.4455 | ##### | 11.3360 | 8.7026E-30 | 9.3658E-29 |
| ENSG00000007171\|NOS2 | 221.7502 | 4.2485 | ##### | 11.3326 | 9.0449E-30 | 9.7307E-29 |
| ENSG00000115541\|HSPE1 | 17447.6993 | 1.3802 | ##### | 11.3240 | 9.9796E-30 | 1.0733E-28 |
| ENSG00000154734\|ADAMTS1 | 1123.5556 | -1.4801 | ##### | ####### | 1.0219E-29 | 1.0986E-28 |
| ENSG00000006016\|CRLF1 | 562.1511 | 1.8946 | ##### | 11.3196 | 1.0497E-29 | 1.1280E-28 |
| ENSG00000216490\|IFI30 | 2805.3149 | -1.4350 | ##### | ####### | 1.0594E-29 | 1.1381E-28 |
| ENSG00000119922\|IFIT2 | 185.0106 | -3.0496 | ##### | ####### | 1.1038E-29 | 1.1854E-28 |
| ENSG00000160710\|ADAR | 44840.5736 | 1.3150 | ##### | 11.3151 | 1.1047E-29 | 1.1859E-28 |
| ENSG00000008382\|MPND | 1377.3410 | 1.5205 | ##### | 11.3131 | 1.1300E-29 | 1.2126E-28 |
| ENSG00000197965\|MPZL1 | 13923.6453 | -1.2054 | ##### | ####### | 1.1467E-29 | 1.2301E-28 |
| ENSG00000152270\|PDE3B | 1258.4918 | 1.5317 | ##### | 11.3117 | 1.1479E-29 | 1.2309E-28 |
| ENSG00000213699\|SLC35F6 | 8032.5379 | -1.3832 | ##### | ####### | 1.2226E-29 | 1.3106E-28 |
| ENSG00000112759\|SLC29A1 | 6935.8233 | 1.1322 | ##### | 11.3006 | 1.3029E-29 | 1.3961E-28 |
| ENSG00000287352 | 180.2221 | -9.9490 | ##### | ####### | 1.3956E-29 | 1.4949E-28 |
| ENSG00000150990\|DHX37 | 15435.6630 | 1.3136 | ##### | 11.2941 | 1.4026E-29 | 1.5019E-28 |
| ENSG00000164107\|HAND2 | 141.4793 | -9.6007 | ##### | ####### | 1.4224E-29 | 1.5225E-28 |
| ENSG00000270055 | 272.1490 | -2.1409 | ##### | ####### | 1.4250E-29 | 1.5248E-28 |
| ENSG00000145687\|SSBP2 | 433.1118 | -1.8447 | ##### | ####### | 1.4299E-29 | 1.5295E-28 |
| ENSG00000262772\|LINC01977 | 101.3748 | 3.0294 | ##### | 11.2924 | 1.4315E-29 | 1.5307E-28 |
| ENSG00000290818 | 224.7861 | -2.0946 | ##### | ####### | 1.4421E-29 | 1.5414E-28 |
| ENSG00000141738\|GRB7 | 482.5691 | -1.7019 | ##### | ####### | 1.4753E-29 | 1.5764E-28 |
| ENSG00000136731\|UGGT1 | 14971.3480 | -1.3574 | ##### | ####### | 1.5678E-29 | 1.6746E-28 |
| ENSG00000163482\|STK36 | 2907.4002 | 1.0982 | ##### | 11.2819 | 1.6114E-29 | 1.7206E-28 |
| ENSG00000043355\|ZIC2 | 784.4972 | 1.5415 | ##### | 11.2814 | 1.6207E-29 | 1.7299E-28 |
| ENSG00000105576\|TNPO2 | 10001.7100 | 1.3849 | ##### | 11.2791 | 1.6650E-29 | 1.7765E-28 |
| ENSG00000183479\|TREX2 | 1181.1807 | 2.6798 | ##### | 11.2758 | 1.7284E-29 | 1.8435E-28 |
| ENSG00000177380\|PPFIA3 | 1913.9630 | -1.4342 | ##### | ####### | 1.7616E-29 | 1.8782E-28 |
| ENSG00000112079\|STK38 | 5821.9452 | -1.0501 | ##### | ####### | 1.8799E-29 | 2.0037E-28 |
| ENSG00000164048\|ZNF589 | 610.3819 | 1.7628 | ##### | 11.2666 | 1.9192E-29 | 2.0448E-28 |
| ENSG00000204956\|PCDHGA1 | 286.4115 | -4.7842 | ##### | ####### | 1.9573E-29 | 2.0847E-28 |
| ENSG00000050327\|ARHGEF5 | 1708.4256 | -1.4016 | ##### | ####### | 2.0500E-29 | 2.1826E-28 |
| ENSG00000143106\|PSMA5 | 19487.8022 | 0.8181 | ##### | 11.2606 | 2.0527E-29 | 2.1847E-28 |
| ENSG00000204525\|HLA-C | 16275.6221 | -1.3415 | ##### | ####### | 2.0781E-29 | 2.2110E-28 |
| ENSG00000169762\|TAPT1 | 859.0895 | -1.5066 | ##### | ####### | 2.1100E-29 | 2.2441E-28 |
| ENSG00000164611\|PTTG1 | 5419.5514 | 1.4264 | ##### | 11.2571 | 2.1370E-29 | 2.2720E-28 |
| ENSG00000169918\|OTUD7A | 200.0557 | -4.3669 | ##### | ####### | 2.2975E-29 | 2.4418E-28 |
| ENSG00000013561\|RNF14 | 6873.8310 | -1.3809 | ##### | ####### | 2.3299E-29 | 2.4754E-28 |
| ENSG00000185272\|RBM11 | 94.2205 | -4.7682 | ##### | ####### | 2.3765E-29 | 2.5240E-28 |
| ENSG00000084207\|GSTP1 | 47233.2267 | 1.3827 | ##### | 11.2445 | 2.4658E-29 | 2.6179E-28 |
| ENSG00000289405 | 310.1466 | 2.1316 | ##### | 11.2432 | 2.5013E-29 | 2.6546E-28 |
| ENSG00000272173 | 1166.1275 | 1.7152 | ##### | 11.2345 | 2.7615E-29 | 2.9297E-28 |
| ENSG00000136868\|SLC31A1 | 5340.5113 | -1.3100 | ##### | ####### | 2.7677E-29 | 2.9352E-28 |
| ENSG00000277067 | 728.6557 | -1.4341 | ##### | ####### | 2.8299E-29 | 3.0001E-28 |
| ENSG00000165912\|PACSIN3 | 2974.9343 | 1.3484 | ##### | 11.2314 | 2.8595E-29 | 3.0305E-28 |
| ENSG00000187186 | 256.6184 | 2.1591 | ##### | 11.2229 | 3.1468E-29 | 3.3338E-28 |
| ENSG00000178826\|TMEM139 | 162.2465 | -2.9053 | ##### | ####### | 3.2081E-29 | 3.3975E-28 |
| ENSG00000117143\|UAP1 | 11433.1101 | 1.0589 | ##### | 11.2201 | 3.2500E-29 | 3.4407E-28 |
| ENSG00000067221\|STOML1 | 464.0823 | -1.8359 | ##### | ####### | 3.2652E-29 | 3.4555E-28 |
| ENSG00000187189\|TSPYL4 | 1288.7566 | -1.5509 | ##### | ####### | 3.4572E-29 | 3.6574E-28 |
| ENSG00000144724\|PTPRG | 1054.1361 | -1.4992 | ##### | ####### | 3.6514E-29 | 3.8615E-28 |
| ENSG00000026103\|FAS | 863.7931 | -1.4672 | ##### | ####### | 3.6906E-29 | 3.9016E-28 |
| ENSG00000172845\|SP3 | 10933.3888 | 1.3763 | ##### | 11.2044 | 3.8804E-29 | 4.1008E-28 |
| ENSG00000165175\|MID1IP1 | 4328.8934 | -1.4205 | ##### | ####### | 3.8823E-29 | 4.1013E-28 |
| ENSG00000128655\|PDE11A | 82.1567 | -3.1095 | ##### | ####### | 4.0278E-29 | 4.2536E-28 |
| ENSG00000198763\|MT-ND2 | ########## | 1.3975 | ##### | 11.2006 | 4.0508E-29 | 4.2763E-28 |
| ENSG00000162426\|SLC45A1 | 1042.2519 | -1.4819 | ##### | ####### | 4.3338E-29 | 4.5735E-28 |
| ENSG00000103089\|FA2H | 1449.6808 | ####### | ##### | ####### | 4.3545E-29 | 4.5937E-28 |
| ENSG00000060558\|GNA15 | 137.2812 | -2.4558 | ##### | ####### | 4.4211E-29 | 4.6623E-28 |
| ENSG00000159733\|ZFYVE28 | 814.5784 | -1.6664 | ##### | ####### | 4.5775E-29 | 4.8255E-28 |
| ENSG00000114209\|PDCD10 | 2483.7757 | -1.4783 | ##### | ####### | 4.6094E-29 | 4.8575E-28 |

| | | | | | | |
|---|---|---|---|---|---|---|
| ENSG00000156675\|RAB11FIP1 | 3777.7955 | -1.3498 | ##### | ####### | 5.0228E-29 | 5.2912E-28 |
| ENSG00000101457\|DNTTIP1 | 4359.1292 | 1.2690 | ##### | 11.1814 | 5.0264E-29 | 5.2931E-28 |
| ENSG00000102910\|LONP2 | 6261.1020 | -1.3469 | ##### | ####### | 5.0841E-29 | 5.3520E-28 |
| ENSG00000197355\|UAP1L1 | 727.5656 | -1.5958 | ##### | ####### | 5.3887E-29 | 5.6707E-28 |
| ENSG00000095002\|MSH2 | 10943.8391 | 1.1543 | ##### | 11.1749 | 5.4138E-29 | 5.6951E-28 |
| ENSG00000105613\|MAST1 | 601.8746 | 1.6281 | ##### | 11.1726 | 5.5553E-29 | 5.8419E-28 |
| ENSG00000259207\|ITGB3 | 583.6181 | 1.7691 | ##### | 11.1685 | 5.8166E-29 | 6.1124E-28 |
| ENSG00000171604\|CXXC5 | 510.2001 | -1.7263 | ##### | ####### | 5.8149E-29 | 6.1124E-28 |
| ENSG00000179528\|LBX2 | 175.3667 | -2.1299 | ##### | ####### | 5.8392E-29 | 6.1340E-28 |
| ENSG00000198026\|ZNF335 | 4156.7997 | 1.3673 | ##### | 11.1668 | 5.9298E-29 | 6.2269E-28 |
| ENSG00000105248\|YJU2 | 2600.6362 | 1.3791 | ##### | 11.1664 | 5.9542E-29 | 6.2504E-28 |
| ENSG00000232653\|GOLGA8N | 117.6408 | -2.3895 | ##### | ####### | 6.0930E-29 | 6.3938E-28 |
| ENSG00000102007\|PLP2 | 11390.5798 | -1.3344 | ##### | ####### | 6.3779E-29 | 6.6905E-28 |
| ENSG00000153214\|TMEM87B | 4016.0104 | -1.4181 | ##### | ####### | 6.4352E-29 | 6.7482E-28 |
| ENSG00000112701\|SENP6 | 9298.8635 | 1.2747 | ##### | 11.1577 | 6.5687E-29 | 6.8857E-28 |
| ENSG00000005022\|SLC25A5 | 41355.5243 | 1.2221 | ##### | 11.1549 | 6.7759E-29 | 7.1005E-28 |
| ENSG00000175984\|DENND2C | 379.2283 | 1.8827 | ##### | 11.1517 | 7.0223E-29 | 7.3561E-28 |
| ENSG00000120075\|HOXB5 | 123.4132 | -2.5240 | ##### | ####### | 7.2883E-29 | 7.6320E-28 |
| ENSG00000079134\|THOC1 | 4265.2766 | 1.4049 | ##### | 11.1481 | 7.3170E-29 | 7.6594E-28 |
| ENSG00000161381\|PLXDC1 | 87.4465 | 5.5316 | ##### | 11.1474 | 7.3695E-29 | 7.7116E-28 |
| ENSG00000161010\|MRNIP | 9142.5293 | -1.6979 | ##### | ####### | 7.6029E-29 | 7.9531E-28 |
| ENSG00000152936\|LMNTD1 | 97.8877 | 4.0357 | ##### | 11.1445 | 7.6183E-29 | 7.9665E-28 |
| ENSG00000132541\|RIDA | 1552.0923 | -1.5026 | ##### | ####### | 7.6339E-29 | 7.9799E-28 |
| ENSG00000174099\|MSRB3 | 7352.9081 | -0.7880 | ##### | ####### | 7.6521E-29 | 7.9961E-28 |
| ENSG00000120318\|ARAP3 | 1633.5058 | -1.4749 | ##### | ####### | 7.8211E-29 | 8.1699E-28 |
| ENSG00000290796 | 285.1954 | 2.0522 | ##### | 11.1415 | 7.8797E-29 | 8.2282E-28 |
| ENSG00000139173\|TMEM117 | 362.6865 | -1.8444 | ##### | ####### | 8.1262E-29 | 8.4827E-28 |
| ENSG00000272669 | 99.2678 | 2.6809 | ##### | 11.1383 | 8.1664E-29 | 8.5216E-28 |
| ENSG00000169230\|PRELID1 | 35132.8047 | 1.3295 | ##### | 11.1279 | 9.1775E-29 | 9.5734E-28 |
| ENSG00000028203\|VEZT | 6792.4829 | -1.2133 | ##### | ####### | 9.5662E-29 | 9.9753E-28 |
| ENSG00000170468\|RIOX1 | 2866.8993 | 1.4023 | ##### | 11.1178 | 1.0274E-28 | 1.0710E-27 |
| ENSG00000174282\|ZBTB4 | 3087.5216 | -1.3448 | ##### | ####### | 1.0406E-28 | 1.0844E-27 |
| ENSG00000070759\|TESK2 | 178.1441 | 2.2245 | ##### | 11.1141 | 1.0717E-28 | 1.1159E-27 |
| ENSG00000065609\|SNAP91 | 271.0322 | -2.0312 | ##### | ####### | 1.0713E-28 | 1.1159E-27 |
| ENSG00000145736\|GTF2H2 | 2776.5021 | 1.4012 | ##### | 11.1105 | 1.1153E-28 | 1.1610E-27 |
| ENSG00000099256\|PRTFDC1 | 2552.3982 | 1.3695 | ##### | 11.1065 | 1.1659E-28 | 1.2132E-27 |
| ENSG00000066629\|EML1 | 2366.5197 | -1.4174 | ##### | ####### | 1.1769E-28 | 1.2243E-27 |
| ENSG00000287064 | 87.5036 | 3.9764 | ##### | 11.1052 | 1.1840E-28 | 1.2312E-27 |
| ENSG00000010017\|RANBP9 | 3783.1588 | 1.3843 | ##### | 11.1043 | 1.1960E-28 | 1.2432E-27 |
| ENSG00000153933\|DGKE | 514.6675 | -1.8728 | ##### | ####### | 1.2543E-28 | 1.3034E-27 |
| ENSG00000286596 | 1122.3703 | 1.8822 | ##### | 11.0974 | 1.2916E-28 | 1.3417E-27 |
| ENSG00000151470\|C4orf33 | 255.9698 | -2.3980 | ##### | ####### | 1.3002E-28 | 1.3502E-27 |
| ENSG00000068366\|ACSL4 | 7450.0053 | 1.3730 | ##### | 11.0967 | 1.3020E-28 | 1.3516E-27 |
| ENSG00000181240\|SLC25A41 | 139.1578 | -9.4574 | ##### | ####### | 1.3044E-28 | 1.3536E-27 |
| ENSG00000147535\|PLPP5 | 1713.6836 | -1.4278 | ##### | ####### | 1.3194E-28 | 1.3686E-27 |
| ENSG00000137831\|UACA | 2846.2338 | -1.3895 | ##### | ####### | 1.3502E-28 | 1.4002E-27 |
| ENSG00000110315\|RNF141 | 2024.2345 | -1.3714 | ##### | ####### | 1.3694E-28 | 1.4195E-27 |
| ENSG00000215375\|MYL5 | 561.6521 | -2.2276 | ##### | ####### | 1.3732E-28 | 1.4230E-27 |
| ENSG00000008735\|MAPK8IP2 | 1756.7246 | -1.4477 | ##### | ####### | 1.4084E-28 | 1.4590E-27 |
| ENSG00000115825\|PRKD3 | 5460.2990 | 1.3997 | ##### | 11.0861 | 1.4656E-28 | 1.5177E-27 |
| ENSG00000213928\|IRF9 | 559.9836 | -1.8092 | ##### | ####### | 1.4764E-28 | 1.5283E-27 |
| ENSG00000126804\|ZBTB1 | 6416.8890 | 1.3835 | ##### | 11.0841 | 1.4984E-28 | 1.5505E-27 |
| ENSG00000142961\|MOB3C | 609.9947 | -1.5725 | ##### | ####### | 1.5740E-28 | 1.6283E-27 |
| ENSG00000185650\|ZFP36L1 | 11204.9673 | -1.3280 | ##### | ####### | 1.6449E-28 | 1.7010E-27 |
| ENSG00000139211\|AMIGO2 | 2353.5343 | -1.2313 | ##### | ####### | 1.7134E-28 | 1.7713E-27 |
| ENSG00000135002\|RFK | 5221.3376 | -1.1105 | ##### | ####### | 1.7435E-28 | 1.8017E-27 |
| ENSG00000053702\|NRIP2 | 68.9412 | 3.2301 | ##### | 11.0683 | 1.7875E-28 | 1.8465E-27 |
| ENSG00000186889\|TMEM17 | 322.9244 | -1.7533 | ##### | ####### | 1.8147E-28 | 1.8733E-27 |
| ENSG00000168079\|SCARA5 | 196.0548 | -2.5274 | ##### | ####### | 1.8144E-28 | 1.8733E-27 |
| ENSG00000132549\|VPS13B | 2697.9222 | -1.4227 | ##### | ####### | 1.8859E-28 | 1.9462E-27 |
| ENSG00000279720\|SOWAHCP2 | 121.3483 | 3.3958 | ##### | 11.0631 | 1.8943E-28 | 1.9542E-27 |
| ENSG00000123374\|CDK2 | 10767.5369 | 0.8511 | ##### | 11.0594 | 1.9740E-28 | 2.0356E-27 |
| ENSG00000150907\|FOXO1 | 437.4909 | -1.7912 | ##### | ####### | 2.0153E-28 | 2.0775E-27 |

| | | | | | | |
|---|---|---|---|---|---|---|
| ENSG00000100321\|SYNGR1 | 634.0700 | -1.6015 | ##### | ####### | 2.0204E-28 | 2.0821E-27 |
| ENSG00000105993\|DNAJB6 | 14123.9253 | 0.9545 | ##### | 11.0556 | 2.0604E-28 | 2.1226E-27 |
| ENSG00000160072\|ATAD3B | 11438.9692 | 0.9538 | ##### | 11.0509 | 2.1706E-28 | 2.2354E-27 |
| ENSG00000099804\|CDC34 | 8848.0208 | 1.3530 | ##### | 11.0502 | 2.1871E-28 | 2.2516E-27 |
| ENSG00000270673\|YTHDF3-DT | 96.4431 | -2.5933 | ##### | ####### | 2.2001E-28 | 2.2642E-27 |
| ENSG00000269604 | 5058.3073 | 1.2006 | ##### | 11.0484 | 2.2308E-28 | 2.2950E-27 |
| ENSG00000291082\|HERC2P9 | 565.8575 | 1.6545 | ##### | 11.0467 | 2.2746E-28 | 2.3392E-27 |
| ENSG00000065268\|WDR18 | 5236.6842 | 1.4015 | ##### | 11.0398 | 2.4558E-28 | 2.5247E-27 |
| ENSG00000177302\|TOP3A | 5295.6509 | 1.0526 | ##### | 11.0328 | 2.6535E-28 | 2.7270E-27 |
| ENSG00000254470\|AP5B1 | 5440.9644 | 1.3160 | ##### | 11.0325 | 2.6648E-28 | 2.7377E-27 |
| ENSG00000099954\|CECR2 | 171.7182 | 2.7317 | ##### | 11.0313 | 2.7003E-28 | 2.7732E-27 |
| ENSG00000198736\|MSRB1 | 2559.6693 | -1.4328 | ##### | ####### | 2.7742E-28 | 2.8482E-27 |
| ENSG00000100599\|RIN3 | 803.3474 | -1.5299 | ##### | ####### | 2.8302E-28 | 2.9046E-27 |
| ENSG00000175691\|ZNF77 | 391.8486 | 1.9012 | ##### | 11.0258 | 2.8695E-28 | 2.9435E-27 |
| ENSG00000229390\|MICD | 90.6353 | 4.6221 | ##### | 11.0258 | 2.8701E-28 | 2.9435E-27 |
| ENSG00000185513\|L3MBTL1 | 667.5495 | 1.6139 | ##### | 11.0256 | 2.8763E-28 | 2.9489E-27 |
| ENSG00000170606\|HSPA4 | 28879.8582 | 1.2973 | ##### | 11.0256 | 2.8773E-28 | 2.9489E-27 |
| ENSG00000286619\|PM20D1-AS1 | 149.6585 | -4.7155 | ##### | ####### | 2.9087E-28 | 2.9800E-27 |
| ENSG00000170989\|S1PR1 | 220.2843 | -2.2837 | ##### | ####### | 2.9663E-28 | 3.0381E-27 |
| ENSG00000138083\|SIX3 | 51.2661 | -4.9909 | ##### | ####### | 3.0543E-28 | 3.1270E-27 |
| ENSG00000141040\|ZNF287 | 453.5567 | -1.7646 | ##### | ####### | 3.0935E-28 | 3.1662E-27 |
| ENSG00000196220\|SRGAP3 | 146.8202 | -3.1013 | ##### | ####### | 3.1220E-28 | 3.1942E-27 |
| ENSG00000197114\|ZGPAT | 2830.9738 | 1.4561 | ##### | 11.0177 | 3.1381E-28 | 3.2095E-27 |
| ENSG00000071564\|TCF3 | 15291.6950 | 1.2257 | ##### | 11.0164 | 3.1868E-28 | 3.2583E-27 |
| ENSG00000188738\|FSIP2 | 99.1542 | -3.3213 | ##### | ####### | 3.2437E-28 | 3.3153E-27 |
| ENSG00000277363\|SRCIN1 | 834.0351 | 1.5749 | ##### | 11.0124 | 3.3312E-28 | 3.4036E-27 |
| ENSG00000055130\|CUL1 | 12183.4189 | 1.0246 | ##### | 11.0102 | 3.4110E-28 | 3.4839E-27 |
| ENSG00000143575\|HAX1 | 10776.8446 | 1.1381 | ##### | 11.0059 | 3.5808E-28 | 3.6561E-27 |
| ENSG00000253304\|TMEM200B | 1497.8687 | -1.5604 | ##### | ####### | 3.7473E-28 | 3.8247E-27 |
| ENSG00000125458\|NT5C | 3507.2478 | -1.0971 | ##### | ####### | 3.8090E-28 | 3.8865E-27 |
| ENSG00000185269\|NOTUM | 227.5701 | 1.9767 | ##### | 10.9994 | 3.8478E-28 | 3.9246E-27 |
| ENSG00000224167\|LINC01357 | 321.2561 | 2.2306 | ##### | 10.9962 | 3.9872E-28 | 4.0655E-27 |
| ENSG00000256751\|PLBD1-AS1 | 252.2203 | -2.3922 | ##### | ####### | 4.0639E-28 | 4.1423E-27 |
| ENSG00000173950\|XXYLT1 | 1736.4426 | -1.1444 | ##### | ####### | 4.1489E-28 | 4.2275E-27 |
| ENSG00000132361\|CLUH | 22012.1215 | 1.2830 | ##### | 10.9912 | 4.2115E-28 | 4.2898E-27 |
| ENSG00000134215\|VAV3 | 1455.8591 | -1.4425 | ##### | ####### | 4.4412E-28 | 4.5222E-27 |
| ENSG00000185306\|C12orf56 | 414.3927 | -1.8725 | ##### | ####### | 4.6819E-28 | 4.7657E-27 |
| ENSG00000047579\|DTNBP1 | 1046.4091 | 1.5601 | ##### | 10.9805 | 4.7429E-28 | 4.8261E-27 |
| ENSG00000185339\|TCN2 | 1940.5620 | ####### | ##### | ####### | 4.8968E-28 | 4.9810E-27 |
| ENSG00000006007\|GDE1 | 7058.9300 | -1.3321 | ##### | ####### | 4.9530E-28 | 5.0365E-27 |
| ENSG00000184497\|TMEM255B | 118.0490 | 2.8730 | ##### | 10.9729 | 5.1599E-28 | 5.2451E-27 |
| ENSG00000283828 | 112.7676 | 3.7228 | ##### | 10.9712 | 5.2577E-28 | 5.3427E-27 |
| ENSG00000206053\|JPT2 | 15445.0161 | -0.9141 | ##### | ####### | 5.3031E-28 | 5.3870E-27 |
| ENSG00000010361\|FUZ | 1417.9305 | 1.4628 | ##### | 10.9698 | 5.3391E-28 | 5.4217E-27 |
| ENSG00000006638\|TBXA2R | 594.9932 | -1.6317 | ##### | ####### | 5.3557E-28 | 5.4367E-27 |
| ENSG00000105298\|CACTIN | 6687.8339 | 1.2750 | ##### | 10.9693 | 5.3709E-28 | 5.4503E-27 |
| ENSG00000076928\|ARHGEF1 | 7572.5789 | -1.2803 | ##### | ####### | 5.5066E-28 | 5.5861E-27 |
| ENSG00000162734\|PEA15 | 23115.0031 | 1.2617 | ##### | 10.9669 | 5.5158E-28 | 5.5935E-27 |
| ENSG00000152939\|MARVELD2 | 437.7239 | -1.9414 | ##### | ####### | 5.5690E-28 | 5.6456E-27 |
| ENSG00000148704\|VAX1 | 288.1655 | 1.8495 | ##### | 10.9633 | 5.7338E-28 | 5.8107E-27 |
| ENSG00000160703\|NLRX1 | 1404.3781 | -1.2244 | ##### | ####### | 5.8157E-28 | 5.8917E-27 |
| ENSG00000168214\|RBPJ | 6116.0463 | 1.1390 | ##### | 10.9618 | 5.8321E-28 | 5.9062E-27 |
| ENSG00000123213\|NLN | 5683.2169 | -1.1027 | ##### | ####### | 5.9413E-28 | 6.0148E-27 |
| ENSG00000244687\|UBE2V1 | 16732.2306 | 1.3635 | ##### | 10.9595 | 5.9858E-28 | 6.0578E-27 |
| ENSG00000204685\|STARD7-AS1 | 945.6108 | 1.5091 | ##### | 10.9530 | 6.4296E-28 | 6.5048E-27 |
| ENSG00000198894\|CIPC | 2565.7068 | 1.4216 | ##### | 10.9519 | 6.5073E-28 | 6.5811E-27 |
| ENSG00000231064\|THBS3-AS1 | 538.1927 | 1.8777 | ##### | 10.9498 | 6.6582E-28 | 6.7315E-27 |
| ENSG00000146112\|PPP1R18 | 6732.2678 | 1.4012 | ##### | 10.9491 | 6.7084E-28 | 6.7800E-27 |
| ENSG00000079313\|REXO1 | 7735.5851 | 1.3423 | ##### | 10.9478 | 6.8099E-28 | 6.8802E-27 |
| ENSG00000214485\|RPL7P1 | 4198.4518 | -1.3916 | ##### | ####### | 7.1007E-28 | 7.1716E-27 |
| ENSG00000075218\|GTSE1 | 4610.8224 | 1.3701 | ##### | 10.9429 | 7.1848E-28 | 7.2541E-27 |
| ENSG00000084463\|WBP11 | 20253.2298 | 1.0410 | ##### | 10.9416 | 7.2930E-28 | 7.3608E-27 |
| ENSG00000105568\|PPP2R1A | 25324.7588 | -1.3261 | ##### | ####### | 7.7340E-28 | 7.8032E-27 |

| | | | | | | |
|---|---|---|---|---|---|---|
| ENSG00000177084\|POLE | 10279.6030 | 1.3048 | ##### | 10.9350 | 7.8368E-28 | 7.9043E-27 |
| ENSG00000163864\|NMNAT3 | 94.5980 | -3.1333 | ##### | ####### | 8.0327E-28 | 8.0991E-27 |
| ENSG00000092140\|G2E3 | 3843.7059 | 1.1527 | ##### | 10.9320 | 8.1018E-28 | 8.1661E-27 |
| ENSG00000080839\|RBL1 | 3757.0647 | 1.3115 | ##### | 10.9316 | 8.1430E-28 | 8.2048E-27 |
| ENSG00000186235\|LINC02610 | 112.4858 | 2.4691 | ##### | 10.9301 | 8.2779E-28 | 8.3379E-27 |
| ENSG00000112742\|TTK | 3497.8358 | 1.3522 | ##### | 10.9296 | 8.3210E-28 | 8.3785E-27 |
| ENSG00000144366\|GULP1 | 5999.0414 | -1.3053 | ##### | ####### | 8.3305E-28 | 8.3853E-27 |
| ENSG00000055483\|USP36 | 10642.1354 | 1.2837 | ##### | 10.9262 | 8.6370E-28 | 8.6908E-27 |
| ENSG00000187994\|RINL | 928.1129 | -7.6933 | ##### | ####### | 8.6521E-28 | 8.7031E-27 |
| ENSG00000163374\|YY1AP1 | 8987.7437 | 1.2357 | ##### | 10.9232 | 8.9247E-28 | 8.9743E-27 |
| ENSG00000131263\|RLIM | 8456.8038 | 1.2824 | ##### | 10.9227 | 8.9761E-28 | 9.0229E-27 |
| ENSG00000251095 | 167.9721 | 5.5817 | ##### | 10.9155 | 9.7202E-28 | 9.7676E-27 |
| ENSG00000203706\|SERTAD4-AS1 | 236.5221 | -1.8190 | ##### | ####### | 1.0214E-27 | 1.0261E-26 |
| ENSG00000122378\|PRXL2A | 2712.0221 | -1.3174 | ##### | ####### | 1.0235E-27 | 1.0278E-26 |
| ENSG00000266074\|BAHCC1 | 2780.2991 | -1.3838 | ##### | ####### | 1.0666E-27 | 1.0707E-26 |
| ENSG00000196912\|ANKRD36B | 354.0259 | 1.7012 | ##### | 10.9059 | 1.0800E-27 | 1.0838E-26 |
| ENSG00000113460\|BRIX1 | 12295.2784 | 1.3308 | ##### | 10.9054 | 1.0864E-27 | 1.0898E-26 |
| ENSG00000198911\|SREBF2 | 9850.1113 | -1.3006 | ##### | ####### | 1.1073E-27 | 1.1105E-26 |
| ENSG00000123562\|MORF4L2 | 29956.3516 | 0.9038 | ##### | 10.9009 | 1.1415E-27 | 1.1444E-26 |
| ENSG00000204390\|HSPA1L | 334.5121 | 2.0251 | ##### | 10.8992 | 1.1629E-27 | 1.1654E-26 |
| ENSG00000235831\|BHLHE40-AS1 | 132.2060 | 2.6311 | ##### | 10.8965 | 1.1971E-27 | 1.1993E-26 |
| ENSG00000140057\|AK7 | 231.9494 | -2.8123 | ##### | ####### | 1.2698E-27 | 1.2717E-26 |
| ENSG00000138411\|HECW2 | 588.8310 | -1.6490 | ##### | ####### | 1.2831E-27 | 1.2846E-26 |
| ENSG00000163739\|CXCL1 | 3100.2639 | 1.4896 | ##### | 10.8886 | 1.3064E-27 | 1.3075E-26 |
| ENSG00000104381\|GDAP1 | 801.6722 | -1.5466 | ##### | ####### | 1.3207E-27 | 1.3213E-26 |
| ENSG00000187742\|SECISBP2 | 2425.1187 | -0.8430 | ##### | ####### | 1.3547E-27 | 1.3549E-26 |
| ENSG00000164830\|OXR1 | 1850.6440 | -1.4189 | ##### | ####### | 1.3831E-27 | 1.3829E-26 |
| ENSG00000179387\|ELMOD2 | 2006.3097 | -1.3717 | ##### | ####### | 1.4638E-27 | 1.4631E-26 |
| ENSG00000100614\|PPM1A | 4579.2560 | 1.1830 | ##### | 10.8775 | 1.4752E-27 | 1.4740E-26 |
| ENSG00000059378\|PARP12 | 2625.5584 | -1.2272 | ##### | ####### | 1.4869E-27 | 1.4851E-26 |
| ENSG00000129968\|ABHD17A | 7596.8179 | 1.1284 | ##### | 10.8765 | 1.4911E-27 | 1.4889E-26 |
| ENSG00000096872\|IFT74 | 811.4578 | -1.5233 | ##### | ####### | 1.5011E-27 | 1.4984E-26 |
| ENSG00000169855\|ROBO1 | 7045.3046 | -1.3077 | ##### | ####### | 1.5026E-27 | 1.4993E-26 |
| ENSG00000111642\|CHD4 | 37584.9777 | 1.2624 | ##### | 10.8751 | 1.5141E-27 | 1.5103E-26 |
| ENSG00000091009\|RBM27 | 5334.1429 | 1.4051 | ##### | 10.8741 | 1.5321E-27 | 1.5278E-26 |
| ENSG00000182253\|SYNM | 5420.2552 | 1.3634 | ##### | 10.8732 | 1.5474E-27 | 1.5425E-26 |
| ENSG00000072682\|P4HA2 | 3347.5242 | -1.3490 | ##### | ####### | 1.5680E-27 | 1.5625E-26 |
| ENSG00000179163\|FUCA1 | 1114.4181 | -1.5375 | ##### | ####### | 1.6172E-27 | 1.6110E-26 |
| ENSG00000160410\|SHKBP1 | 7150.5871 | -0.8175 | ##### | ####### | 1.6406E-27 | 1.6338E-26 |
| ENSG00000138792\|ENPEP | 193.4151 | -2.6321 | ##### | ####### | 1.6701E-27 | 1.6626E-26 |
| ENSG00000281649\|EBLN3P | 6732.0895 | 1.3069 | ##### | 10.8657 | 1.6791E-27 | 1.6709E-26 |
| ENSG00000101335\|MYL9 | 9209.2708 | 1.3632 | ##### | 10.8650 | 1.6931E-27 | 1.6843E-26 |
| ENSG00000164941\|INTS8 | 7044.0850 | -1.2964 | ##### | ####### | 1.7126E-27 | 1.7032E-26 |
| ENSG00000108854\|SMURF2 | 8056.6780 | 1.3075 | ##### | 10.8610 | 1.7684E-27 | 1.7581E-26 |
| ENSG00000181026\|AEN | 5396.7839 | 1.3597 | ##### | 10.8589 | 1.8090E-27 | 1.7979E-26 |
| ENSG00000164512\|ANKRD55 | 70.2006 | 3.6466 | ##### | 10.8502 | 1.9890E-27 | 1.9761E-26 |
| ENSG00000105825\|TFPI2 | ########## | 1.2581 | ##### | 10.8501 | 1.9911E-27 | 1.9775E-26 |
| ENSG00000197321\|SVIL | 12567.2983 | 1.2944 | ##### | 10.8475 | 2.0492E-27 | 2.0345E-26 |
| ENSG00000005249\|PRKAR2B | 725.9143 | 1.5471 | ##### | 10.8472 | 2.0553E-27 | 2.0400E-26 |
| ENSG00000088280\|ASAP3 | 1157.6221 | -1.4922 | ##### | ####### | 2.0730E-27 | 2.0568E-26 |
| ENSG00000116704\|SLC35D1 | 3686.5116 | -1.3425 | ##### | ####### | 2.1068E-27 | 2.0897E-26 |
| ENSG00000128011\|LRFN1 | 1273.1679 | -1.4618 | ##### | ####### | 2.2129E-27 | 2.1941E-26 |
| ENSG00000213366\|GSTM2 | 102.4373 | -2.9257 | ##### | ####### | 2.2728E-27 | 2.2528E-26 |
| ENSG00000114796\|KLHL24 | 570.5989 | -1.5599 | ##### | ####### | 2.2807E-27 | 2.2599E-26 |
| ENSG00000141499\|WRAP53 | 2658.8244 | 1.3842 | ##### | 10.8376 | 2.2835E-27 | 2.2619E-26 |
| ENSG00000126746\|ZNF384 | 10232.8961 | 1.1492 | ##### | 10.8364 | 2.3149E-27 | 2.2922E-26 |
| ENSG00000133863\|TEX15 | 2488.8232 | 1.3767 | ##### | 10.8355 | 2.3374E-27 | 2.3138E-26 |
| ENSG00000172731\|LRRC20 | 1376.0210 | -1.3862 | ##### | ####### | 2.3408E-27 | 2.3164E-26 |
| ENSG00000158966\|CACHD1 | 3151.1296 | 1.2953 | ##### | 10.8342 | 2.3715E-27 | 2.3460E-26 |
| ENSG00000111602\|TIMELESS | 14293.3097 | 1.2413 | ##### | 10.8341 | 2.3732E-27 | 2.3469E-26 |
| ENSG00000175895\|PLEKHF2 | 712.3219 | -1.6122 | ##### | ####### | 2.3925E-27 | 2.3652E-26 |
| ENSG00000276476\|LINC00540 | 823.3080 | ####### | ##### | ####### | 2.4019E-27 | 2.3736E-26 |
| ENSG00000143816\|WNT9A | 1521.1778 | 1.3510 | ##### | 10.8321 | 2.4257E-27 | 2.3964E-26 |

| | | | | | | |
|---|---|---|---|---|---|---|
| ENSG00000241627\|UBQLN4P1 | 307.8028 | 1.8546 | ##### | 10.8311 | 2.4521E-27 | 2.4217E-26 |
| ENSG00000132938\|MTUS2 | 75.6290 | 4.7663 | ##### | 10.8310 | 2.4533E-27 | 2.4220E-26 |
| ENSG00000159063\|ALG8 | 5574.5844 | 1.3386 | ##### | 10.8293 | 2.4998E-27 | 2.4672E-26 |
| ENSG00000122696\|SLC25A51 | 1748.5343 | 1.3237 | ##### | 10.8292 | 2.5047E-27 | 2.4711E-26 |
| ENSG00000144362\|PHOSPHO2 | 264.9633 | 1.9382 | ##### | 10.8241 | 2.6461E-27 | 2.6098E-26 |
| ENSG00000261924 | 142.5977 | 2.9504 | ##### | 10.8237 | 2.6591E-27 | 2.6218E-26 |
| ENSG00000168268\|NT5DC2 | 9159.7524 | 1.1999 | ##### | 10.8182 | 2.8213E-27 | 2.7808E-26 |
| ENSG00000113838\|TBCCD1 | 796.0826 | -1.5169 | ##### | ####### | 2.8709E-27 | 2.8287E-26 |
| ENSG00000176871\|WSB2 | 9675.7816 | -1.2787 | ##### | ####### | 2.9170E-27 | 2.8733E-26 |
| ENSG00000172197\|MBOAT1 | 754.1063 | 1.5745 | ##### | 10.8125 | 3.0023E-27 | 2.9562E-26 |
| ENSG00000261614\|YBX3P1 | 118.5731 | 2.7315 | ##### | 10.8124 | 3.0083E-27 | 2.9612E-26 |
| ENSG00000153531\|ADPRHL1 | 577.7643 | -1.5911 | ##### | ####### | 3.0324E-27 | 2.9839E-26 |
| ENSG00000167371\|PRRT2 | 215.3630 | 2.3173 | ##### | 10.8108 | 3.0589E-27 | 3.0090E-26 |
| ENSG00000143067\|ZNF697 | 1411.3313 | 1.4342 | ##### | 10.8093 | 3.1105E-27 | 3.0588E-26 |
| ENSG00000112031\|MTRF1L | 5902.4129 | 1.2782 | ##### | 10.8088 | 3.1286E-27 | 3.0756E-26 |
| ENSG00000085274\|MYNN | 1410.5551 | -1.4386 | ##### | ####### | 3.1327E-27 | 3.0786E-26 |
| ENSG00000163069\|SGCB | 6947.7785 | 1.0373 | ##### | 10.8030 | 3.3315E-27 | 3.2728E-26 |
| ENSG00000172292\|CERS6 | 8997.0537 | 1.4243 | ##### | 10.8005 | 3.4232E-27 | 3.3618E-26 |
| ENSG00000100503\|NIN | 5934.7117 | 1.3251 | ##### | 10.7988 | 3.4864E-27 | 3.4228E-26 |
| ENSG00000293176\|RPSAP52 | 238.7186 | 9.3005 | ##### | 10.7966 | 3.5719E-27 | 3.5056E-26 |
| ENSG00000145386\|CCNA2 | 9299.4475 | 1.3256 | ##### | 10.7963 | 3.5820E-27 | 3.5144E-26 |
| ENSG00000143862\|ARL8A | 1878.9021 | -1.4123 | ##### | ####### | 3.7568E-27 | 3.6846E-26 |
| ENSG00000204959\|ARHGEF34P | 928.7840 | -1.5166 | ##### | ####### | 3.8417E-27 | 3.7667E-26 |
| ENSG00000240409\|MTATP8P1 | 101.1267 | 2.6381 | ##### | 10.7892 | 3.8713E-27 | 3.7945E-26 |
| ENSG00000293188 | 357.6592 | -1.7287 | ##### | ####### | 3.8817E-27 | 3.8034E-26 |
| ENSG00000077684\|JADE1 | 4147.5839 | -1.2926 | ##### | ####### | 3.8860E-27 | 3.8064E-26 |
| ENSG00000101210\|EEF1A2 | 25465.0038 | 1.2951 | ##### | 10.7884 | 3.9069E-27 | 3.8255E-26 |
| ENSG00000128309\|MPST | 3649.8112 | -1.3133 | ##### | ####### | 3.9090E-27 | 3.8264E-26 |
| ENSG00000165792\|METTL17 | 5047.4021 | 1.2372 | ##### | 10.7815 | 4.2096E-27 | 4.1193E-26 |
| ENSG00000087299\|L2HGDH | 2153.8969 | 1.3289 | ##### | 10.7798 | 4.2870E-27 | 4.1936E-26 |
| ENSG00000171798\|KNDC1 | 273.0943 | 1.9886 | ##### | 10.7757 | 4.4839E-27 | 4.3848E-26 |
| ENSG00000135116\|HRK | 90.7610 | -4.5973 | ##### | ####### | 4.5446E-27 | 4.4427E-26 |
| ENSG00000105323\|HNRNPUL1 | 31272.8980 | -0.6220 | ##### | ####### | 4.6338E-27 | 4.5284E-26 |
| ENSG00000143842\|SOX13 | 2861.9873 | 1.1249 | ##### | 10.7726 | 4.6364E-27 | 4.5295E-26 |
| ENSG00000267787 | 289.1622 | 1.5883 | ##### | 10.7680 | 4.8763E-27 | 4.7623E-26 |
| ENSG00000163710\|PCOLCE2 | 3924.8972 | 1.3891 | ##### | 10.7654 | 5.0146E-27 | 4.8958E-26 |
| ENSG00000123080\|CDKN2C | 1130.3102 | -1.5252 | ##### | ####### | 5.2357E-27 | 5.1099E-26 |
| ENSG00000101076\|HNF4A | 568.2112 | ####### | ##### | ####### | 5.2934E-27 | 5.1646E-26 |
| ENSG00000162599\|NFIA | 1699.6298 | -1.3875 | ##### | ####### | 5.3370E-27 | 5.2054E-26 |
| ENSG00000126368\|NR1D1 | 1254.8348 | 1.4610 | ##### | 10.7546 | 5.6397E-27 | 5.4989E-26 |
| ENSG00000288736 | 44.8175 | -5.9261 | ##### | ####### | 5.6634E-27 | 5.5201E-26 |
| ENSG00000102763\|VWA8 | 1782.9890 | -1.3916 | ##### | ####### | 5.7030E-27 | 5.5569E-26 |
| ENSG00000278730 | 1636.6291 | -1.3953 | ##### | ####### | 5.7073E-27 | 5.5593E-26 |
| ENSG00000180573\|H2AC6 | 3581.1678 | 1.1218 | ##### | 10.7510 | 5.8602E-27 | 5.7064E-26 |
| ENSG00000182218\|HHIPL1 | 5840.9605 | ####### | ##### | ####### | 5.8920E-27 | 5.7355E-26 |
| ENSG00000167969\|ECI1 | 4370.2596 | -1.2852 | ##### | ####### | 6.2909E-27 | 6.1218E-26 |
| ENSG00000177576\|C18orf32 | 2106.2515 | -1.5166 | ##### | ####### | 6.4158E-27 | 6.2413E-26 |
| ENSG00000153363\|LINC00467 | 408.1595 | -1.6805 | ##### | ####### | 6.4225E-27 | 6.2458E-26 |
| ENSG00000111581\|NUP107 | 10016.6530 | 1.0980 | ##### | 10.7410 | 6.5354E-27 | 6.3535E-26 |
| ENSG00000004468\|CD38 | 439.0377 | -1.8791 | ##### | ####### | 6.6260E-27 | 6.4395E-26 |
| ENSG00000204611\|ZNF616 | 640.8123 | -1.5148 | ##### | ####### | 6.8088E-27 | 6.6150E-26 |
| ENSG00000134186\|PRPF38B | 7204.9738 | 1.3233 | ##### | 10.7343 | 7.0270E-27 | 6.8247E-26 |
| ENSG00000111252\|SH2B3 | 2152.8314 | 1.3731 | ##### | 10.7342 | 7.0362E-27 | 6.8315E-26 |
| ENSG00000068078\|FGFR3 | 600.6512 | -1.5245 | ##### | ####### | 7.0766E-27 | 6.8685E-26 |
| ENSG00000130706\|ADRM1 | 33116.0227 | 1.3055 | ##### | 10.7330 | 7.1266E-27 | 6.9148E-26 |
| ENSG00000101670\|LIPG | 388.1840 | 2.1163 | ##### | 10.7311 | 7.2709E-27 | 7.0525E-26 |
| ENSG00000106829\|TLE4 | 1030.2521 | -1.9773 | ##### | ####### | 7.3915E-27 | 7.1672E-26 |
| ENSG00000186868\|MAPT | 655.5892 | 1.5488 | ##### | 10.7214 | 8.0768E-27 | 7.8290E-26 |
| ENSG00000049192\|ADAMTS6 | 237.4437 | 2.5389 | ##### | 10.7203 | 8.1749E-27 | 7.9216E-26 |
| ENSG00000206503\|HLA-A | 23593.1388 | 1.2633 | ##### | 10.7151 | 8.6418E-27 | 8.3713E-26 |
| ENSG00000279196 | 1475.1561 | -1.7817 | ##### | ####### | 8.8580E-27 | 8.5780E-26 |
| ENSG00000182791\|CCDC87 | 71.0756 | 3.1883 | ##### | 10.7123 | 8.9101E-27 | 8.6257E-26 |
| ENSG00000159267\|HLCS | 2629.4141 | -1.3132 | ##### | ####### | 8.9431E-27 | 8.6547E-26 |

| | | | | | | |
|---|---|---|---|---|---|---|
| ENSG00000152926\|ZNF117 | 224.2768 | -2.8911 | ##### | ####### | 9.1651E-27 | 8.8667E-26 |
| ENSG00000154277\|UCHL1 | 28173.0683 | -1.1110 | ##### | ####### | 9.2881E-27 | 8.9828E-26 |
| ENSG00000230733 | 1213.9421 | -1.5252 | ##### | ####### | 9.3389E-27 | 9.0290E-26 |
| ENSG00000100297\|MCM5 | 14342.3004 | 0.8739 | ##### | 10.7074 | 9.3946E-27 | 9.0800E-26 |
| ENSG00000253210\|DENND3-AS1 | 82.8748 | -2.8985 | ##### | ####### | 9.6477E-27 | 9.3215E-26 |
| ENSG00000197724\|PHF2 | 2020.2827 | -1.3586 | ##### | ####### | 1.0214E-26 | 9.8657E-26 |
| ENSG00000145002\|FAM86B2 | 882.4457 | -1.3759 | ##### | ####### | 1.0701E-26 | 1.0332E-25 |
| ENSG00000058453\|CROCC | 1422.3751 | 1.3422 | ##### | 10.6948 | 1.0768E-26 | 1.0394E-25 |
| ENSG00000272688 | 756.5762 | 1.6272 | ##### | 10.6925 | 1.1038E-26 | 1.0652E-25 |
| ENSG00000100784\|RPS6KA5 | 474.1895 | 1.8793 | ##### | 10.6910 | 1.1215E-26 | 1.0818E-25 |
| ENSG00000085741\|WNT11 | 128.2759 | -2.4494 | ##### | ####### | 1.1324E-26 | 1.0920E-25 |
| ENSG00000137936\|BCAR3 | 5821.8403 | -1.3636 | ##### | ####### | 1.2120E-26 | 1.1684E-25 |
| ENSG00000143321\|HDGF | 66860.8072 | 0.9753 | ##### | 10.6820 | 1.2364E-26 | 1.1916E-25 |
| ENSG00000136810\|TXN | 33929.7350 | -1.2979 | ##### | ####### | 1.2437E-26 | 1.1981E-25 |
| ENSG00000105438\|KDELR1 | 17722.0049 | -1.3222 | ##### | ####### | 1.3012E-26 | 1.2532E-25 |
| ENSG00000257653 | 105.4564 | -2.3866 | ##### | ####### | 1.3267E-26 | 1.2773E-25 |
| ENSG00000067715\|SYT1 | 2055.3184 | -1.3660 | ##### | ####### | 1.3627E-26 | 1.3115E-25 |
| ENSG00000156471\|PTDSS1 | 9992.1926 | -1.1231 | ##### | ####### | 1.3893E-26 | 1.3367E-25 |
| ENSG00000186522\|SEPTIN10 | 7825.9474 | -1.3102 | ##### | ####### | 1.4266E-26 | 1.3721E-25 |
| ENSG00000140521\|POLG | 5851.1140 | 1.2275 | ##### | 10.6670 | 1.4521E-26 | 1.3962E-25 |
| ENSG00000197915\|HRNR | 237.7006 | 3.6871 | ##### | 10.6645 | 1.4922E-26 | 1.4343E-25 |
| ENSG00000071282\|LMCD1 | 634.0910 | -1.5548 | ##### | ####### | 1.5362E-26 | 1.4761E-25 |
| ENSG00000169683\|LRRC45 | 1269.3982 | 1.3501 | ##### | 10.6603 | 1.5602E-26 | 1.4987E-25 |
| ENSG00000144815\|NXPE3 | 4665.6019 | -1.3420 | ##### | ####### | 1.5947E-26 | 1.5314E-25 |
| ENSG00000168676\|KCTD19 | 147.4906 | 3.7427 | ##### | 10.6481 | 1.7792E-26 | 1.7080E-25 |
| ENSG00000175354\|PTPN2 | 4669.3484 | 1.3160 | ##### | 10.6468 | 1.8054E-26 | 1.7326E-25 |
| ENSG00000223891\|OSER1-DT | 479.3164 | 1.5932 | ##### | 10.6451 | 1.8378E-26 | 1.7632E-25 |
| ENSG00000165355\|FBXO33 | 1445.0786 | 1.3847 | ##### | 10.6448 | 1.8445E-26 | 1.7690E-25 |
| ENSG00000139800\|ZIC5 | 464.2706 | 1.6927 | ##### | 10.6415 | 1.9096E-26 | 1.8308E-25 |
| ENSG00000081803\|CADPS2 | 605.2789 | -1.5586 | ##### | ####### | 1.9737E-26 | 1.8917E-25 |
| ENSG00000183114\|FAM43B | 494.6966 | 1.6623 | ##### | 10.6384 | 1.9756E-26 | 1.8922E-25 |
| ENSG00000238062\|SPATA3-AS1 | 101.3869 | 3.3699 | ##### | 10.6384 | 1.9754E-26 | 1.8922E-25 |
| ENSG00000110328\|GALNT18 | 815.1259 | -1.5390 | ##### | ####### | 2.0183E-26 | 1.9325E-25 |
| ENSG00000128708\|HAT1 | 8245.0163 | 1.2565 | ##### | 10.6354 | 2.0387E-26 | 1.9515E-25 |
| ENSG00000259803\|SLC22A31 | 132.1759 | 2.8917 | ##### | 10.6345 | 2.0593E-26 | 1.9705E-25 |
| ENSG00000236901\|MIR600HG | 411.0763 | 1.8333 | ##### | 10.6322 | 2.1106E-26 | 2.0190E-25 |
| ENSG00000033050\|ABCF2 | 17808.3912 | 0.7193 | ##### | 10.6315 | 2.1271E-26 | 2.0341E-25 |
| ENSG00000104969\|SGTA | 11376.2955 | 1.1007 | ##### | 10.6313 | 2.1322E-26 | 2.0383E-25 |
| ENSG00000143543\|JTB | 12815.5457 | 1.2826 | ##### | 10.6274 | 2.2211E-26 | 2.1227E-25 |
| ENSG00000100196\|KDELR3 | 1976.2029 | -1.3968 | ##### | ####### | 2.4755E-26 | 2.3651E-25 |
| ENSG00000116741\|RGS2 | 1617.0276 | -1.4576 | ##### | ####### | 2.6061E-26 | 2.4890E-25 |
| ENSG00000151348\|EXT2 | 4755.7343 | -1.3203 | ##### | ####### | 2.6307E-26 | 2.5110E-25 |
| ENSG00000157350\|ST3GAL2 | 5784.7926 | -1.3058 | ##### | ####### | 2.6309E-26 | 2.5110E-25 |
| ENSG00000137266\|SLC22A23 | 2232.9959 | -1.3048 | ##### | ####### | 2.7307E-26 | 2.6055E-25 |
| ENSG00000114999\|TTL | 6723.4453 | -1.2906 | ##### | ####### | 2.7707E-26 | 2.6429E-25 |
| ENSG00000083454\|P2RX5 | 1730.9729 | 1.4737 | ##### | 10.6064 | 2.7831E-26 | 2.6538E-25 |
| ENSG00000115548\|KDM3A | 5750.2766 | 1.3505 | ##### | 10.6018 | 2.9224E-26 | 2.7858E-25 |
| ENSG00000267607\|ICAM4-AS1 | 1245.0647 | -1.4576 | ##### | ####### | 3.0224E-26 | 2.8801E-25 |
| ENSG00000122965\|RBM19 | 6676.8639 | 1.2552 | ##### | 10.5934 | 3.1988E-26 | 3.0473E-25 |
| ENSG00000183248\|PRR36 | 222.0971 | 2.2673 | ##### | 10.5866 | 3.4392E-26 | 3.2752E-25 |
| ENSG00000131409\|LRRC4B | 532.4202 | 1.7602 | ##### | 10.5856 | 3.4766E-26 | 3.3098E-25 |
| ENSG00000156384\|SFR1 | 817.5431 | -1.4872 | ##### | ####### | 3.5247E-26 | 3.3545E-25 |
| ENSG00000113456\|RAD1 | 5709.3286 | 0.9978 | ##### | 10.5821 | 3.6077E-26 | 3.4324E-25 |
| ENSG00000229852\|KHDC1-AS1 | 614.9333 | 1.4138 | ##### | 10.5811 | 3.6463E-26 | 3.4680E-25 |
| ENSG00000149557\|FEZ1 | 453.5846 | 2.0781 | ##### | 10.5794 | 3.7148E-26 | 3.5320E-25 |
| ENSG00000107960\|STN1 | 1380.1179 | -1.3878 | ##### | ####### | 3.7807E-26 | 3.5936E-25 |
| ENSG00000015475\|BID | 1972.7320 | -1.3608 | ##### | ####### | 3.7994E-26 | 3.6103E-25 |
| ENSG00000131351\|HAUS8 | 1631.8882 | 1.3634 | ##### | 10.5771 | 3.8058E-26 | 3.6151E-25 |
| ENSG00000183346\|CABCOCO1 | 180.5052 | -4.1650 | ##### | ####### | 3.9459E-26 | 3.7470E-25 |
| ENSG00000115468\|EFHD1 | 325.7498 | 1.8392 | ##### | 10.5732 | 3.9679E-26 | 3.7667E-25 |
| ENSG00000236212 | 89.9167 | -3.5446 | ##### | ####### | 4.0429E-26 | 3.8367E-25 |
| ENSG00000197948\|FCHSD1 | 2367.7671 | -1.3384 | ##### | ####### | 4.0774E-26 | 3.8682E-25 |
| ENSG00000211455\|STK38L | 3942.0726 | 1.2952 | ##### | 10.5690 | 4.1490E-26 | 3.9349E-25 |

| | | | | | | |
|---|---|---|---|---|---|---|
| ENSG00000106479\|ZNF862 | 877.9325 | -1.5249 | ##### | ####### | 4.1521E-26 | 3.9366E-25 |
| ENSG00000167767\|KRT80 | 2668.7304 | -1.3500 | ##### | ####### | 4.2007E-26 | 3.9814E-25 |
| ENSG00000143643\|TTC13 | 1744.6336 | -1.3274 | ##### | ####### | 4.3445E-26 | 4.1164E-25 |
| ENSG00000115091\|ACTR3 | 25590.0402 | -0.9017 | ##### | ####### | 4.3850E-26 | 4.1535E-25 |
| ENSG00000158560\|DYNC1I1 | 105.2478 | -2.7805 | ##### | ####### | 4.3864E-26 | 4.1535E-25 |
| ENSG00000157227\|MMP14 | 504.1105 | -1.4979 | ##### | ####### | 4.4131E-26 | 4.1775E-25 |
| ENSG00000143786\|CNIH3 | 242.1573 | 1.8830 | ##### | 10.5626 | 4.4416E-26 | 4.2030E-25 |
| ENSG00000178202\|POGLUT3 | 3666.3576 | -1.2896 | ##### | ####### | 4.5764E-26 | 4.3293E-25 |
| ENSG00000137076\|TLN1 | 47450.2646 | 1.2361 | ##### | 10.5585 | 4.6420E-26 | 4.3899E-25 |
| ENSG00000175697\|GPR156 | 161.1255 | -2.1763 | ##### | ####### | 4.6816E-26 | 4.4260E-25 |
| ENSG00000103647\|CORO2B | 144.5338 | 3.1487 | ##### | 10.5516 | 4.9927E-26 | 4.7185E-25 |
| ENSG00000174010\|KLHL15 | 2236.0345 | 1.3134 | ##### | 10.5511 | 5.0206E-26 | 4.7434E-25 |
| ENSG00000213903\|LTB4R | 801.7223 | 1.5838 | ##### | 10.5488 | 5.1427E-26 | 4.8572E-25 |
| ENSG00000164292\|RHOBTB3 | 4406.3588 | -1.3061 | ##### | ####### | 5.2287E-26 | 4.9369E-25 |
| ENSG00000111752\|PHC1 | 10545.9484 | 1.4106 | ##### | 10.5441 | 5.4095E-26 | 5.1060E-25 |
| ENSG00000163071\|SPATA18 | 104.9947 | 3.6564 | ##### | 10.5418 | 5.5408E-26 | 5.2283E-25 |
| ENSG00000279753 | 1591.5843 | 1.8632 | ##### | 10.5412 | 5.5803E-26 | 5.2639E-25 |
| ENSG00000111731\|C2CD5 | 4055.4217 | 1.2485 | ##### | 10.5356 | 5.9189E-26 | 5.5815E-25 |
| ENSG00000065154\|OAT | 8621.0221 | 1.3203 | ##### | 10.5337 | 6.0424E-26 | 5.6944E-25 |
| ENSG00000244486\|SCARF2 | 388.2736 | 1.8133 | ##### | 10.5337 | 6.0407E-26 | 5.6944E-25 |
| ENSG00000247556\|OIP5-AS1 | 7555.4373 | -1.2955 | ##### | ####### | 6.0949E-26 | 5.7421E-25 |
| ENSG00000043591\|ADRB1 | 185.8659 | 2.1552 | ##### | 10.5315 | 6.1861E-26 | 5.8262E-25 |
| ENSG00000115145\|STAM2 | 3588.1535 | -1.2911 | ##### | ####### | 6.2016E-26 | 5.8389E-25 |
| ENSG00000114416\|FXR1 | 16127.7781 | 1.0323 | ##### | 10.5307 | 6.2342E-26 | 5.8678E-25 |
| ENSG00000158793\|NIT1 | 4035.2044 | 1.2964 | ##### | 10.5304 | 6.2551E-26 | 5.8856E-25 |
| ENSG00000181045\|SLC26A11 | 1455.5939 | -1.3695 | ##### | ####### | 6.3782E-26 | 5.9996E-25 |
| ENSG00000132589\|FLOT2 | 6781.3010 | -1.2806 | ##### | ####### | 6.5044E-26 | 6.1163E-25 |
| ENSG00000198919\|DZIP3 | 3769.2387 | 1.2689 | ##### | 10.5200 | 6.9839E-26 | 6.5651E-25 |
| ENSG00000210154\|MT-TD | 77.6472 | -2.4960 | ##### | ####### | 7.0311E-26 | 6.6074E-25 |
| ENSG00000131446\|MGAT1 | 11073.6270 | 1.1049 | ##### | 10.5144 | 7.4174E-26 | 6.9682E-25 |
| ENSG00000164040\|PGRMC2 | 4815.7866 | -1.2221 | ##### | ####### | 7.6397E-26 | 7.1748E-25 |
| ENSG00000203279 | 97.4717 | -2.5091 | ##### | ####### | 7.9178E-26 | 7.4337E-25 |
| ENSG00000236081\|ELFN1-AS1 | 54.4220 | -7.2124 | ##### | ####### | 8.0452E-26 | 7.5509E-25 |
| ENSG00000161996\|WDR90 | 4171.2477 | 1.3170 | ##### | 10.5054 | 8.1591E-26 | 7.6554E-25 |
| ENSG00000124201\|ZNFX1 | 8394.9061 | 1.2367 | ##### | 10.4999 | 8.6479E-26 | 8.1115E-25 |
| ENSG00000179899\|PHC1P1 | 800.3642 | 1.5117 | ##### | 10.4913 | 9.4708E-26 | 8.8806E-25 |
| ENSG00000062038\|CDH3 | 709.2240 | -1.5071 | ##### | ####### | 9.6200E-26 | 9.0176E-25 |
| ENSG00000023734\|STRAP | 32015.1634 | 0.8366 | ##### | 10.4867 | 9.9409E-26 | 9.3155E-25 |
| ENSG00000164038\|SLC9B2 | 1159.2337 | -1.4016 | ##### | ####### | 1.0010E-25 | 9.3769E-25 |
| ENSG00000171823\|FBXL14 | 2806.8795 | 1.3999 | ##### | 10.4852 | 1.0107E-25 | 9.4649E-25 |
| ENSG00000225978\|HAR1A | 141.5237 | 2.8251 | ##### | 10.4832 | 1.0316E-25 | 9.6578E-25 |
| ENSG00000287603 | 93.4523 | -9.0016 | ##### | ####### | 1.0345E-25 | 9.6824E-25 |
| ENSG00000152766\|ANKRD22 | 127.7505 | -4.0929 | ##### | ####### | 1.0736E-25 | 1.0045E-24 |
| ENSG00000059758\|CDK17 | 4493.2057 | 1.3132 | ##### | 10.4790 | 1.0783E-25 | 1.0086E-24 |
| ENSG00000235890\|TSPEAR-AS1 | 169.5874 | 3.1027 | ##### | 10.4787 | 1.0817E-25 | 1.0114E-24 |
| ENSG00000089847\|ANKRD24 | 264.0416 | -1.8631 | ##### | ####### | 1.0864E-25 | 1.0155E-24 |
| ENSG00000267141 | 55.8445 | 4.0457 | ##### | 10.4716 | 1.1663E-25 | 1.0898E-24 |
| ENSG00000151503\|NCAPD3 | 10500.4092 | 1.2901 | ##### | 10.4679 | 1.2130E-25 | 1.1331E-24 |
| ENSG00000116584\|ARHGEF2 | 10230.2641 | 1.2208 | ##### | 10.4675 | 1.2188E-25 | 1.1382E-24 |
| ENSG00000162368\|CMPK1 | 13517.4575 | -1.1384 | ##### | ####### | 1.2254E-25 | 1.1440E-24 |
| ENSG00000232859\|LYRM9 | 135.5974 | -2.4863 | ##### | ####### | 1.2356E-25 | 1.1532E-24 |
| ENSG00000135968\|GCC2 | 4313.2745 | -1.2728 | ##### | ####### | 1.2469E-25 | 1.1634E-24 |
| ENSG00000173262\|SLC2A14 | 168.1737 | 1.9742 | ##### | 10.4620 | 1.2904E-25 | 1.2035E-24 |
| ENSG00000184216\|IRAK1 | 35008.4819 | 1.0961 | ##### | 10.4612 | 1.3019E-25 | 1.2139E-24 |
| ENSG00000196305\|IARS1 | 21912.9009 | -0.8308 | ##### | ####### | 1.3329E-25 | 1.2424E-24 |
| ENSG00000166548\|TK2 | 1497.4237 | -1.4005 | ##### | ####### | 1.3487E-25 | 1.2568E-24 |
| ENSG00000206127\|GOLGA8O | 78.3758 | -2.6360 | ##### | ####### | 1.3977E-25 | 1.3020E-24 |
| ENSG00000174718\|RESF1 | 3347.1545 | 1.2686 | ##### | 10.4511 | 1.4488E-25 | 1.3492E-24 |
| ENSG00000152582\|SPEF2 | 427.4037 | 1.5079 | ##### | 10.4491 | 1.4788E-25 | 1.3767E-24 |
| ENSG00000186416\|NKRF | 3899.0842 | 1.2975 | ##### | 10.4490 | 1.4805E-25 | 1.3779E-24 |
| ENSG00000198718\|TOGARAM1 | 1658.9937 | 1.3995 | ##### | 10.4488 | 1.4845E-25 | 1.3811E-24 |
| ENSG00000227262\|HCG4B | 112.9278 | 3.3712 | ##### | 10.4440 | 1.5615E-25 | 1.4523E-24 |
| ENSG00000105229\|PIAS4 | 3534.2976 | 1.2605 | ##### | 10.4437 | 1.5653E-25 | 1.4554E-24 |

| | | | | | | |
|---|---|---|---|---|---|---|
| ENSG00000251432\|LINC02615 | 145.6356 | -2.4152 | ##### | ####### | 1.6855E-25 | 1.5667E-24 |
| ENSG00000107874\|CUEDC2 | 3055.7213 | -1.2932 | ##### | ####### | 1.6900E-25 | 1.5703E-24 |
| ENSG00000005486\|RHBDD2 | 4544.1130 | -1.2747 | ##### | ####### | 1.6928E-25 | 1.5725E-24 |
| ENSG00000124784\|RIOK1 | 2233.8577 | 1.3464 | ##### | 10.4359 | 1.6994E-25 | 1.5782E-24 |
| ENSG00000206337\|HCP5 | 111.4865 | -2.7245 | ##### | ####### | 1.7137E-25 | 1.5910E-24 |
| ENSG00000177602\|HASPIN | 1791.8763 | 1.3977 | ##### | 10.4345 | 1.7257E-25 | 1.6015E-24 |
| ENSG00000142039\|CCDC97 | 2635.2363 | -1.3401 | ##### | ####### | 1.7262E-25 | 1.6015E-24 |
| ENSG00000066777\|ARFGEF1 | 5661.9099 | -1.2733 | ##### | ####### | 1.8015E-25 | 1.6709E-24 |
| ENSG00000198890\|PRMT6 | 2777.3307 | -1.1035 | ##### | ####### | 1.8024E-25 | 1.6712E-24 |
| ENSG00000188130\|MAPK12 | 2501.0709 | -1.3212 | ##### | ####### | 1.8142E-25 | 1.6816E-24 |
| ENSG00000004399\|PLXND1 | 4462.4156 | -1.2160 | ##### | ####### | 1.8327E-25 | 1.6982E-24 |
| ENSG00000084764\|MAPRE3 | 1527.2058 | -1.3554 | ##### | ####### | 1.9400E-25 | 1.7971E-24 |
| ENSG00000004700\|RECQL | 21386.9921 | 1.2137 | ##### | 10.4216 | 1.9750E-25 | 1.8289E-24 |
| ENSG00000186448\|ZNF197 | 345.6683 | -1.6937 | ##### | ####### | 1.9937E-25 | 1.8457E-24 |
| ENSG00000267519\|MIR23AHG | 3823.7659 | 1.2761 | ##### | 10.4203 | 2.0023E-25 | 1.8530E-24 |
| ENSG00000147601\|TERF1 | 2620.4401 | -1.2861 | ##### | ####### | 2.0968E-25 | 1.9399E-24 |
| ENSG00000135074\|ADAM19 | 370.6435 | 1.5558 | ##### | 10.4147 | 2.1241E-25 | 1.9646E-24 |
| ENSG00000104375\|STK3 | 1931.2492 | -1.3288 | ##### | ####### | 2.2123E-25 | 2.0455E-24 |
| ENSG00000137807\|KIF23 | 7842.8512 | 1.3067 | ##### | 10.4093 | 2.2477E-25 | 2.0776E-24 |
| ENSG00000153208\|MERTK | 784.2431 | 1.5638 | ##### | 10.4090 | 2.2557E-25 | 2.0844E-24 |
| ENSG00000107897\|ACBD5 | 4200.2477 | -1.2518 | ##### | ####### | 2.2664E-25 | 2.0936E-24 |
| ENSG00000233695\|GAS6-AS1 | 368.4238 | 1.7121 | ##### | 10.4079 | 2.2809E-25 | 2.1063E-24 |
| ENSG00000138399\|FASTKD1 | 3338.7001 | 1.3581 | ##### | 10.4073 | 2.2974E-25 | 2.1209E-24 |
| ENSG00000123595\|RAB9A | 1470.0559 | -1.3966 | ##### | ####### | 2.3401E-25 | 2.1597E-24 |
| ENSG00000241489 | 577.2898 | 3.9018 | ##### | 10.4052 | 2.3467E-25 | 2.1651E-24 |
| ENSG00000144583\|MARCHF4 | 9306.1947 | 1.5835 | ##### | 10.4045 | 2.3639E-25 | 2.1803E-24 |
| ENSG00000181634\|TNFSF15 | 67.9497 | -4.7629 | ##### | ####### | 2.3724E-25 | 2.1875E-24 |
| ENSG00000170417\|TMEM182 | 755.2018 | -1.4919 | ##### | ####### | 2.3753E-25 | 2.1895E-24 |
| ENSG00000070423\|RNF126 | 7033.7269 | 1.3304 | ##### | 10.4038 | 2.3815E-25 | 2.1945E-24 |
| ENSG00000184916\|JAG2 | 5930.1188 | -1.2470 | ##### | ####### | 2.4273E-25 | 2.2360E-24 |
| ENSG00000103540\|CCP110 | 4224.7734 | 1.2641 | ##### | 10.4014 | 2.4426E-25 | 2.2482E-24 |
| ENSG00000152475\|ZNF837 | 119.8252 | -2.2862 | ##### | ####### | 2.4428E-25 | 2.2482E-24 |
| ENSG00000147689\|FAM83A | 7303.1931 | ####### | ##### | ####### | 2.4416E-25 | 2.2482E-24 |
| ENSG00000104976\|SNAPC2 | 3066.3541 | 1.2894 | ##### | 10.3979 | 2.5352E-25 | 2.3325E-24 |
| ENSG00000198948\|MFAP3L | 892.9495 | -1.5106 | ##### | ####### | 2.6272E-25 | 2.4164E-24 |
| ENSG00000178233\|TMEM151B | 66.8789 | 3.2301 | ##### | 10.3927 | 2.6764E-25 | 2.4609E-24 |
| ENSG00000233493\|TMEM238 | 210.9462 | -2.1617 | ##### | ####### | 2.6988E-25 | 2.4808E-24 |
| ENSG00000198807\|PAX9 | 429.5122 | 1.6207 | ##### | 10.3911 | 2.7230E-25 | 2.5023E-24 |
| ENSG00000170921\|TANC2 | 5406.7665 | 1.2777 | ##### | 10.3908 | 2.7300E-25 | 2.5079E-24 |
| ENSG00000149658\|YTHDF1 | 8363.7066 | 0.8905 | ##### | 10.3894 | 2.7697E-25 | 2.5436E-24 |
| ENSG00000129911\|KLF16 | 10125.9231 | 1.2555 | ##### | 10.3860 | 2.8703E-25 | 2.6352E-24 |
| ENSG00000160161\|CILP2 | 198.8671 | -2.0252 | ##### | ####### | 2.8807E-25 | 2.6439E-24 |
| ENSG00000180447\|GAS1 | 223.6530 | -2.3680 | ##### | ####### | 3.0764E-25 | 2.8226E-24 |
| ENSG00000163479\|SSR2 | 28722.9768 | 1.0787 | ##### | 10.3783 | 3.1129E-25 | 2.8543E-24 |
| ENSG00000160336\|ZNF761 | 2013.0372 | -1.2667 | ##### | ####### | 3.1122E-25 | 2.8543E-24 |
| ENSG00000175215\|CTDSP2 | 12701.5334 | 0.7270 | ##### | 10.3769 | 3.1598E-25 | 2.8965E-24 |
| ENSG00000172086\|KRCC1 | 1761.7427 | -1.3345 | ##### | ####### | 3.3054E-25 | 3.0290E-24 |
| ENSG00000163795\|ZNF513 | 841.6401 | -1.4603 | ##### | ####### | 3.3774E-25 | 3.0941E-24 |
| ENSG00000133678\|TMEM254 | 1272.3271 | -1.5732 | ##### | ####### | 3.4650E-25 | 3.1733E-24 |
| ENSG00000198929\|NOS1AP | 142.3729 | -2.4042 | ##### | ####### | 3.5184E-25 | 3.2213E-24 |
| ENSG00000170260\|ZNF212 | 1654.6397 | 1.2712 | ##### | 10.3653 | 3.5652E-25 | 3.2631E-24 |
| ENSG00000101407\|TTI1 | 6119.1217 | 1.2553 | ##### | 10.3639 | 3.6185E-25 | 3.3109E-24 |
| ENSG00000097007\|ABL1 | 10180.5885 | -1.2491 | ##### | ####### | 3.7285E-25 | 3.4105E-24 |
| ENSG00000274276 | 2504.8865 | -1.3046 | ##### | ####### | 3.7982E-25 | 3.4731E-24 |
| ENSG00000100003\|SEC14L2 | 1375.1571 | -1.3144 | ##### | ####### | 3.8822E-25 | 3.5489E-24 |
| ENSG00000131437\|KIF3A | 2560.7761 | 1.3582 | ##### | 10.3565 | 3.9104E-25 | 3.5736E-24 |
| ENSG00000006747\|SCIN | 193.6124 | -2.4551 | ##### | ####### | 4.0851E-25 | 3.7321E-24 |
| ENSG00000101347\|SAMHD1 | 3062.5446 | 1.3621 | ##### | 10.3511 | 4.1366E-25 | 3.7780E-24 |
| ENSG00000196407\|THEM5 | 95.9105 | -9.0391 | ##### | ####### | 4.1577E-25 | 3.7961E-24 |
| ENSG00000224389\|C4B | 191.5855 | -2.1352 | ##### | ####### | 4.1741E-25 | 3.8099E-24 |
| ENSG00000137106\|GRHPR | 10075.4001 | 1.1714 | ##### | 10.3405 | 4.6206E-25 | 4.2161E-24 |
| ENSG00000126524\|SBDS | 8010.2904 | -1.2625 | ##### | ####### | 4.6713E-25 | 4.2611E-24 |
| ENSG00000107937\|GTPBP4 | 13792.0026 | 0.8708 | ##### | 10.3375 | 4.7688E-25 | 4.3487E-24 |

| | | | | | | |
|---|---|---|---|---|---|---|
| ENSG00000274383 | 84.1050 | 2.7596 | ##### | 10.3366 | 4.8108E-25 | 4.3857E-24 |
| ENSG00000138398\|PPIG | 7089.5921 | 1.2236 | ##### | 10.3328 | 5.0096E-25 | 4.5655E-24 |
| ENSG00000185519\|FAM131C | 630.6712 | -1.5468 | ##### | ####### | 5.2056E-25 | 4.7427E-24 |
| ENSG00000157873\|TNFRSF14 | 111.7891 | 2.4953 | ##### | 10.3278 | 5.2742E-25 | 4.8037E-24 |
| ENSG00000115271\|GCA | 1069.2271 | -1.3992 | ##### | ####### | 5.2866E-25 | 4.8135E-24 |
| ENSG00000198796\|ALPK2 | 442.1742 | -2.5123 | ##### | ####### | 5.2892E-25 | 4.8145E-24 |
| ENSG00000173821\|RNF213 | 18863.7576 | -1.3082 | ##### | ####### | 5.5423E-25 | 5.0432E-24 |
| ENSG00000212864\|RNF208 | 233.8681 | -2.2007 | ##### | ####### | 5.6850E-25 | 5.1715E-24 |
| ENSG00000073417\|PDE8A | 4790.0553 | 1.2319 | ##### | 10.3192 | 5.7684E-25 | 5.2458E-24 |
| ENSG00000118922\|KLF12 | 260.3850 | 3.5892 | ##### | 10.3164 | 5.9415E-25 | 5.4016E-24 |
| ENSG00000188177\|ZC3H6 | 517.6611 | -1.5752 | ##### | ####### | 6.1370E-25 | 5.5777E-24 |
| ENSG00000064787\|BCAS1 | 195.2317 | -3.4200 | ##### | ####### | 6.1914E-25 | 5.6254E-24 |
| ENSG00000170577\|SIX2 | 712.8588 | ####### | ##### | ####### | 6.2593E-25 | 5.6853E-24 |
| ENSG00000225793\|TMEM30A-DT | 161.4481 | -2.4831 | ##### | ####### | 6.2917E-25 | 5.7130E-24 |
| ENSG00000198915\|RASGEF1A | 1628.2728 | -1.3507 | ##### | ####### | 6.4815E-25 | 5.8836E-24 |
| ENSG00000065183\|WDR3 | 16746.8447 | 1.2218 | ##### | 10.3056 | 6.6466E-25 | 6.0316E-24 |
| ENSG00000076924\|XAB2 | 5369.3577 | 0.9618 | ##### | 10.3056 | 6.6488E-25 | 6.0318E-24 |
| ENSG00000171681\|ATF7IP | 4301.6759 | 1.2935 | ##### | 10.3027 | 6.8545E-25 | 6.2165E-24 |
| ENSG00000237686\|SCIRT | 107.0984 | -3.0349 | ##### | ####### | 6.9235E-25 | 6.2772E-24 |
| ENSG00000156875\|MFSD14A | 3000.7021 | -1.3454 | ##### | ####### | 6.9894E-25 | 6.3350E-24 |
| ENSG00000131773\|KHDRBS3 | 455.3732 | 1.8252 | ##### | 10.2988 | 7.1382E-25 | 6.4679E-24 |
| ENSG00000070388\|FGF22 | 85.0645 | 3.2052 | ##### | 10.2968 | 7.2831E-25 | 6.5970E-24 |
| ENSG00000233654\|NEMP2-DT | 110.2862 | -2.8043 | ##### | ####### | 7.2851E-25 | 6.5970E-24 |
| ENSG00000123575\|FAM199X | 11469.1230 | 1.2622 | ##### | 10.2937 | 7.5232E-25 | 6.8106E-24 |
| ENSG00000138190\|EXOC6 | 1458.5367 | -1.3148 | ##### | ####### | 7.5781E-25 | 6.8582E-24 |
| ENSG00000062194\|GPBP1 | 8335.7885 | -1.2236 | ##### | ####### | 8.1963E-25 | 7.4154E-24 |
| ENSG00000156983\|BRPF1 | 3123.0987 | 1.2662 | ##### | 10.2839 | 8.3272E-25 | 7.5316E-24 |
| ENSG00000137801\|THBS1 | 18507.8150 | 1.2289 | ##### | 10.2837 | 8.3439E-25 | 7.5444E-24 |
| ENSG00000100097\|LGALS1 | 17576.1030 | -1.1931 | ##### | ####### | 8.3787E-25 | 7.5735E-24 |
| ENSG00000120949\|TNFRSF8 | 97.8403 | 3.1848 | ##### | 10.2813 | 8.5562E-25 | 7.7317E-24 |
| ENSG00000175806\|MSRA | 1551.3140 | -1.3949 | ##### | ####### | 8.6066E-25 | 7.7748E-24 |
| ENSG00000152193\|OBI1 | 1746.3748 | 1.3372 | ##### | 10.2794 | 8.7280E-25 | 7.8821E-24 |
| ENSG00000223572\|CKMT1A | 722.2340 | -1.5196 | ##### | ####### | 8.7633E-25 | 7.9116E-24 |
| ENSG00000139998\|RAB15 | 4588.8390 | -1.2284 | ##### | ####### | 8.8736E-25 | 8.0087E-24 |
| ENSG00000288754 | 58.7631 | -3.9028 | ##### | ####### | 8.9152E-25 | 8.0439E-24 |
| ENSG00000138095\|LRPPRC | 46602.1370 | 1.1987 | ##### | 10.2755 | 9.0862E-25 | 8.1957E-24 |
| ENSG00000105523\|FAM83E | 70.5114 | -3.1654 | ##### | ####### | 9.2586E-25 | 8.3487E-24 |
| ENSG00000163050\|COQ8A | 1915.5814 | 1.3403 | ##### | 10.2716 | 9.4624E-25 | 8.5298E-24 |
| ENSG00000273796\|BNAT1 | 43.4361 | -3.9818 | ##### | ####### | 9.7836E-25 | 8.8168E-24 |
| ENSG00000083168\|KAT6A | 4058.8942 | 1.2648 | ##### | 10.2658 | 1.0044E-24 | 9.0483E-24 |
| ENSG00000165480\|SKA3 | 2430.7028 | 1.3144 | ##### | 10.2652 | 1.0107E-24 | 9.1025E-24 |
| ENSG00000270170\|NCBP2AS2 | 1568.0672 | -1.3195 | ##### | ####### | 1.0235E-24 | 9.2156E-24 |
| ENSG00000196123\|MATCAP1 | 873.6062 | -1.4850 | ##### | ####### | 1.0409E-24 | 9.3693E-24 |
| ENSG00000124193\|SRSF6 | 14210.6367 | 1.0575 | ##### | 10.2619 | 1.0464E-24 | 9.4153E-24 |
| ENSG00000276077 | 468.6807 | -1.4313 | ##### | ####### | 1.0781E-24 | 9.6984E-24 |
| ENSG00000131023\|LATS1 | 5479.0538 | 1.2619 | ##### | 10.2589 | 1.0797E-24 | 9.7097E-24 |
| ENSG00000206129\|LINC03069 | 56.3360 | 3.6512 | ##### | 10.2560 | 1.1117E-24 | 9.9944E-24 |
| ENSG00000143257\|NR1I3 | 182.7991 | 2.1614 | ##### | 10.2559 | 1.1131E-24 | 1.0004E-23 |
| ENSG00000158882\|TOMM40L | 3311.9573 | 1.2650 | ##### | 10.2540 | 1.1353E-24 | 1.0200E-23 |
| ENSG00000182870\|GALNT9 | 256.8950 | -1.8106 | ##### | ####### | 1.1365E-24 | 1.0208E-23 |
| ENSG00000081760\|AACS | 4096.8255 | -1.2324 | ##### | ####### | 1.1508E-24 | 1.0333E-23 |
| ENSG00000290791 | 722.1426 | 1.6192 | ##### | 10.2521 | 1.1580E-24 | 1.0394E-23 |
| ENSG00000157021\|CIBAR1P1 | 111.0257 | -2.4504 | ##### | ####### | 1.2120E-24 | 1.0877E-23 |
| ENSG00000153071\|DAB2 | 3081.0660 | 1.3317 | ##### | 10.2398 | 1.3144E-24 | 1.1792E-23 |
| ENSG00000212719\|LINC02693 | 2685.0348 | 1.2333 | ##### | 10.2377 | 1.3436E-24 | 1.2050E-23 |
| ENSG00000196943\|NOP9 | 5755.2315 | 1.2525 | ##### | 10.2321 | 1.4235E-24 | 1.2763E-23 |
| ENSG00000108523\|RNF167 | 6867.0923 | 1.2471 | ##### | 10.2301 | 1.4532E-24 | 1.3022E-23 |
| ENSG00000280018 | 40.2696 | -5.4488 | ##### | ####### | 1.4532E-24 | 1.3022E-23 |
| ENSG00000149657\|LSM14B | 4400.2325 | 1.2670 | ##### | 10.2255 | 1.5240E-24 | 1.3651E-23 |
| ENSG00000143570\|SLC39A1 | 23899.5062 | 1.1179 | ##### | 10.2239 | 1.5495E-24 | 1.3876E-23 |
| ENSG00000196335\|STK31 | 134.0050 | -2.6274 | ##### | ####### | 1.5630E-24 | 1.3992E-23 |
| ENSG00000250320\|EDIL3-DT | 524.1098 | 10.9542 | ##### | 10.2228 | 1.5673E-24 | 1.4027E-23 |
| ENSG00000161203\|AP2M1 | 15462.7730 | -1.2549 | ##### | ####### | 1.5796E-24 | 1.4132E-23 |

| | | | | | | |
|---|---|---|---|---|---|---|
| ENSG00000245662\|LINC02211 | 188.7549 | -1.7861 | ##### | ####### | 1.6213E-24 | 1.4501E-23 |
| ENSG00000173334\|TRIB1 | 4326.2461 | 1.1997 | ##### | 10.2195 | 1.6219E-24 | 1.4502E-23 |
| ENSG00000273270\|LINC03072 | 523.5186 | -1.4090 | ##### | ####### | 1.6332E-24 | 1.4599E-23 |
| ENSG00000164300\|SERINC5 | 1578.6076 | -1.3633 | ##### | ####### | 1.6597E-24 | 1.4832E-23 |
| ENSG00000184305\|CCSER1 | 60.1780 | 5.0459 | ##### | 10.2160 | 1.6810E-24 | 1.5017E-23 |
| ENSG00000291152\|SBDSP1 | 3262.7648 | -1.2794 | ##### | ####### | 1.7229E-24 | 1.5387E-23 |
| ENSG00000188070\|ZFTA | 1235.2464 | 1.4273 | ##### | 10.2115 | 1.7609E-24 | 1.5722E-23 |
| ENSG00000013441\|CLK1 | 1858.0154 | 1.1568 | ##### | 10.2108 | 1.7741E-24 | 1.5835E-23 |
| ENSG00000109944\|JHY | 693.6935 | -1.4639 | ##### | ####### | 1.7751E-24 | 1.5839E-23 |
| ENSG00000159128\|IFNGR2 | 8366.1345 | 1.2546 | ##### | 10.2101 | 1.7875E-24 | 1.5945E-23 |
| ENSG00000177025\|C19orf18 | 212.7163 | ####### | ##### | ####### | 1.8278E-24 | 1.6299E-23 |
| ENSG00000119986\|AVPI1 | 3150.9757 | -1.3297 | ##### | ####### | 1.8630E-24 | 1.6609E-23 |
| ENSG00000213523\|SRA1 | 3292.8819 | 1.0130 | ##### | 10.2027 | 1.9282E-24 | 1.7185E-23 |
| ENSG00000143630\|HCN3 | 1775.4762 | 1.4287 | ##### | 10.2013 | 1.9563E-24 | 1.7429E-23 |
| ENSG00000135636\|DYSF | 654.0882 | -6.2571 | ##### | ####### | 2.0190E-24 | 1.7983E-23 |
| ENSG00000088247\|KHSRP | 26251.7556 | 0.9894 | ##### | 10.1974 | 2.0358E-24 | 1.8128E-23 |
| ENSG00000148358\|GPR107 | 13115.8699 | -1.1896 | ##### | ####### | 2.0573E-24 | 1.8314E-23 |
| ENSG00000238258 | 50.3457 | -3.4308 | ##### | ####### | 2.0799E-24 | 1.8509E-23 |
| ENSG00000085788\|DDHD2 | 3085.0667 | -1.2379 | ##### | ####### | 2.0954E-24 | 1.8641E-23 |
| ENSG00000169519\|METTL15 | 1194.7704 | -1.3503 | ##### | ####### | 2.1169E-24 | 1.8827E-23 |
| ENSG00000154118\|JPH3 | 835.0732 | 1.4853 | ##### | 10.1929 | 2.1338E-24 | 1.8972E-23 |
| ENSG00000204209\|DAXX | 9848.8588 | 1.1378 | ##### | 10.1894 | 2.2102E-24 | 1.9645E-23 |
| ENSG00000106541\|AGR2 | ########## | ####### | ##### | ####### | 2.3104E-24 | 2.0529E-23 |
| ENSG00000184154\|LRRC51 | 543.2523 | 1.6848 | ##### | 10.1849 | 2.3158E-24 | 2.0572E-23 |
| ENSG00000183150\|GPR19 | 177.6085 | 1.8954 | ##### | 10.1848 | 2.3181E-24 | 2.0586E-23 |
| ENSG00000103254\|ANTKMT | 1808.2142 | -1.2778 | ##### | ####### | 2.3788E-24 | 2.1119E-23 |
| ENSG00000164114\|MAP9 | 2197.2279 | -1.3351 | ##### | ####### | 2.4512E-24 | 2.1755E-23 |
| ENSG00000178974\|FBXO34 | 2977.4294 | 1.2636 | ##### | 10.1793 | 2.4532E-24 | 2.1766E-23 |
| ENSG00000112695\|COX7A2 | 17241.0056 | 1.2675 | ##### | 10.1770 | 2.5116E-24 | 2.2277E-23 |
| ENSG00000165556\|CDX2 | 351.1085 | 1.7560 | ##### | 10.1764 | 2.5270E-24 | 2.2408E-23 |
| ENSG00000131019\|ULBP3 | 712.2565 | 1.3347 | ##### | 10.1750 | 2.5651E-24 | 2.2739E-23 |
| ENSG00000198837\|DENND4B | 9398.8330 | 1.1986 | ##### | 10.1739 | 2.5922E-24 | 2.2972E-23 |
| ENSG00000136122\|BORA | 812.1086 | 1.4925 | ##### | 10.1686 | 2.7370E-24 | 2.4248E-23 |
| ENSG00000231312\|MAP4K3-DT | 239.9591 | -2.0289 | ##### | ####### | 2.7398E-24 | 2.4266E-23 |
| ENSG00000072110\|ACTN1 | 53369.5042 | 0.7522 | ##### | 10.1675 | 2.7701E-24 | 2.4527E-23 |
| ENSG00000154319\|FAM167A | 403.7328 | -1.6991 | ##### | ####### | 2.8534E-24 | 2.5257E-23 |
| ENSG00000235194\|PPP1R3E | 460.3644 | 1.5417 | ##### | 10.1602 | 2.9834E-24 | 2.6400E-23 |
| ENSG00000142945\|KIF2C | 9561.7486 | 1.2504 | ##### | 10.1578 | 3.0579E-24 | 2.7051E-23 |
| ENSG00000203326\|ZNF525 | 692.7710 | -1.4524 | ##### | ####### | 3.4563E-24 | 3.0567E-23 |
| ENSG00000107862\|GBF1 | 5908.2308 | -1.2152 | ##### | ####### | 3.4832E-24 | 3.0795E-23 |
| ENSG00000010818\|HIVEP2 | 3231.8786 | 1.2919 | ##### | 10.1423 | 3.5852E-24 | 3.1687E-23 |
| ENSG00000099968\|BCL2L13 | 4330.9313 | -1.1067 | ##### | ####### | 3.6260E-24 | 3.2039E-23 |
| ENSG00000141542\|RAB40B | 455.6419 | -1.8967 | ##### | ####### | 3.6367E-24 | 3.2123E-23 |
| ENSG00000101019\|UQCC1 | 6077.8578 | 1.1213 | ##### | 10.1394 | 3.6920E-24 | 3.2603E-23 |
| ENSG00000132716\|DCAF8 | 14079.2856 | 1.1752 | ##### | 10.1394 | 3.6931E-24 | 3.2603E-23 |
| ENSG00000144642\|RBMS3 | 407.5091 | -1.6625 | ##### | ####### | 3.7116E-24 | 3.2756E-23 |
| ENSG00000148225\|WDR31 | 443.3617 | -1.5396 | ##### | ####### | 3.7544E-24 | 3.3124E-23 |
| ENSG00000213347\|MXD3 | 3312.0300 | 1.6074 | ##### | 10.1351 | 3.8598E-24 | 3.4044E-23 |
| ENSG00000158470\|B4GALT5 | 13775.1927 | -0.9573 | ##### | ####### | 3.8960E-24 | 3.4353E-23 |
| ENSG00000101654\|RNMT | 7069.1606 | 1.2025 | ##### | 10.1326 | 3.9603E-24 | 3.4910E-23 |
| ENSG00000119630\|PGF | 210.4854 | 2.5151 | ##### | 10.1309 | 4.0299E-24 | 3.5513E-23 |
| ENSG00000175221\|MED16 | 4496.5602 | 0.9248 | ##### | 10.1289 | 4.1124E-24 | 3.6229E-23 |
| ENSG00000139263\|LRIG3 | 1325.6015 | -1.3144 | ##### | ####### | 4.2127E-24 | 3.7102E-23 |
| ENSG00000162341\|TPCN2 | 2876.4766 | 1.3143 | ##### | 10.1264 | 4.2177E-24 | 3.7135E-23 |
| ENSG00000165959\|CLMN | 838.8604 | -1.3968 | ##### | ####### | 4.3064E-24 | 3.7905E-23 |
| ENSG00000170458\|CD14 | 173.9872 | 2.7339 | ##### | 10.1241 | 4.3200E-24 | 3.8013E-23 |
| ENSG00000145990\|GFOD1 | 410.9253 | -1.7484 | ##### | ####### | 4.3280E-24 | 3.8073E-23 |
| ENSG00000172992\|DCAKD | 1441.6201 | -1.3417 | ##### | ####### | 4.4073E-24 | 3.8758E-23 |
| ENSG00000238266\|LINC00707 | 563.9358 | 1.4872 | ##### | 10.1215 | 4.4366E-24 | 3.9005E-23 |
| ENSG00000123096\|SSPN | 1231.4769 | 1.2985 | ##### | 10.1205 | 4.4823E-24 | 3.9395E-23 |
| ENSG00000165475\|CRYL1 | 336.3889 | -1.7330 | ##### | ####### | 4.5229E-24 | 3.9740E-23 |
| ENSG00000235961\|PNMA6A | 436.6609 | 1.6207 | ##### | 10.1188 | 4.5589E-24 | 4.0045E-23 |
| ENSG00000178700\|DHFR2 | 928.2126 | -1.4871 | ##### | ####### | 4.5839E-24 | 4.0252E-23 |

| | | | | | | |
|---|---|---|---|---|---|---|
| ENSG00000285693 | 122.1191 | 2.7377 | ##### | 10.1182 | 4.5854E-24 | 4.0254E-23 |
| ENSG00000106415\|GLCCI1 | 1198.3011 | 1.2839 | ##### | 10.1169 | 4.6471E-24 | 4.0783E-23 |
| ENSG00000204634\|TBC1D8 | 5850.7091 | -1.2116 | ##### | ####### | 4.8029E-24 | 4.2138E-23 |
| ENSG00000198133\|TMEM229B | 104.3294 | -2.2866 | ##### | ####### | 4.8178E-24 | 4.2256E-23 |
| ENSG00000133398\|MED10 | 5475.2442 | 1.3057 | ##### | 10.1081 | 5.0854E-24 | 4.4590E-23 |
| ENSG00000169131\|ZNF354A | 968.0922 | 1.4582 | ##### | 10.1059 | 5.2039E-24 | 4.5616E-23 |
| ENSG00000065308\|TRAM2 | 6408.9510 | -1.1814 | ##### | ####### | 5.2624E-24 | 4.6115E-23 |
| ENSG00000091262\|ABCC6 | 120.8746 | -3.0996 | ##### | ####### | 5.2845E-24 | 4.6295E-23 |
| ENSG00000170315\|UBB | 83557.6491 | 0.8902 | ##### | 10.1023 | 5.3956E-24 | 4.7255E-23 |
| ENSG00000116985\|BMP8B | 1061.6319 | -1.4152 | ##### | ####### | 5.4364E-24 | 4.7598E-23 |
| ENSG00000107262\|BAG1 | 4103.7762 | -1.2546 | ##### | ####### | 5.6131E-24 | 4.9131E-23 |
| ENSG00000185305\|ARL15 | 862.6325 | -1.3242 | ##### | ####### | 5.7298E-24 | 5.0137E-23 |
| ENSG00000135346\|CGA | 448.9529 | 10.8866 | ##### | 10.0949 | 5.8164E-24 | 5.0881E-23 |
| ENSG00000116641\|DOCK7 | 4675.9690 | -1.2210 | ##### | ####### | 5.8247E-24 | 5.0938E-23 |
| ENSG00000164087\|POC1A | 2173.8114 | 1.3225 | ##### | 10.0904 | 6.0920E-24 | 5.3260E-23 |
| ENSG00000080845\|DLGAP4 | 8689.9768 | 1.1457 | ##### | 10.0888 | 6.1915E-24 | 5.4114E-23 |
| ENSG00000284906 | 1529.4186 | 1.2651 | ##### | 10.0882 | 6.2307E-24 | 5.4441E-23 |
| ENSG00000156042\|CFAP70 | 122.1162 | -3.0852 | ##### | ####### | 6.4326E-24 | 5.6189E-23 |
| ENSG00000275111\|ZNF2 | 789.8082 | 1.6142 | ##### | 10.0803 | 6.7546E-24 | 5.8985E-23 |
| ENSG00000090530\|P3H2 | 6022.8781 | -1.1540 | ##### | ####### | 6.8125E-24 | 5.9472E-23 |
| ENSG00000196724\|ZNF418 | 151.7024 | -4.4835 | ##### | ####### | 6.8339E-24 | 5.9642E-23 |
| ENSG00000165644\|COMTD1 | 744.1619 | -1.3959 | ##### | ####### | 7.0108E-24 | 6.1168E-23 |
| ENSG00000269226\|TMSB15C | 123.5874 | 2.1992 | ##### | 10.0752 | 7.1121E-24 | 6.2034E-23 |
| ENSG00000136271\|DDX56 | 10180.9805 | 1.1134 | ##### | 10.0743 | 7.1735E-24 | 6.2551E-23 |
| ENSG00000160678\|S100A1 | 175.1342 | 3.0473 | ##### | 10.0733 | 7.2520E-24 | 6.3217E-23 |
| ENSG00000100055\|CYTH4 | 96.6413 | 3.4664 | ##### | 10.0716 | 7.3797E-24 | 6.4312E-23 |
| ENSG00000109854\|HTATIP2 | 6270.9064 | -1.2046 | ##### | ####### | 7.4445E-24 | 6.4857E-23 |
| ENSG00000103490\|PYCARD | 901.4295 | -1.4068 | ##### | ####### | 7.6244E-24 | 6.6405E-23 |
| ENSG00000106462\|EZH2 | 5936.6064 | 1.2186 | ##### | 10.0635 | 8.0083E-24 | 6.9728E-23 |
| ENSG00000171954\|CYP4F22 | 98.8954 | 3.5605 | ##### | 10.0634 | 8.0182E-24 | 6.9794E-23 |
| ENSG00000133142\|TCEAL4 | 5147.7674 | 1.2489 | ##### | 10.0631 | 8.0427E-24 | 6.9987E-23 |
| ENSG00000099260\|PALMD | 716.8335 | -1.4387 | ##### | ####### | 8.1650E-24 | 7.1030E-23 |
| ENSG00000179456\|ZBTB18 | 1177.2207 | 1.3275 | ##### | 10.0604 | 8.2647E-24 | 7.1877E-23 |
| ENSG00000111247\|RAD51AP1 | 2739.7227 | 1.2965 | ##### | 10.0599 | 8.3055E-24 | 7.2211E-23 |
| ENSG00000119979\|DENND10 | 2307.1907 | -1.2205 | ##### | ####### | 8.4384E-24 | 7.3345E-23 |
| ENSG00000124160\|NCOA5 | 5265.8175 | 1.1282 | ##### | 10.0525 | 8.9557E-24 | 7.7819E-23 |
| ENSG00000160867\|FGFR4 | 2841.1162 | 1.2652 | ##### | 10.0500 | 9.1899E-24 | 7.9807E-23 |
| ENSG00000137145\|DENND4C | 1563.2155 | -1.2786 | ##### | ####### | 9.1877E-24 | 7.9807E-23 |
| ENSG00000181588\|MEX3D | 3404.6557 | 1.2219 | ##### | 10.0446 | 9.7057E-24 | 8.4262E-23 |
| ENSG00000136874\|STX17 | 1525.4499 | -1.4522 | ##### | ####### | 9.8266E-24 | 8.5286E-23 |
| ENSG00000072803\|FBXW11 | 7307.1171 | 1.1939 | ##### | 10.0424 | 9.9191E-24 | 8.6065E-23 |
| ENSG00000116455\|WDR77 | 14762.3700 | 1.2123 | ##### | 10.0396 | 1.0207E-23 | 8.8537E-23 |
| ENSG00000106993\|CDC37L1 | 870.8937 | -1.5049 | ##### | ####### | 1.0234E-23 | 8.8749E-23 |
| ENSG00000242588 | 4228.3432 | 1.2319 | ##### | 10.0371 | 1.0471E-23 | 9.0773E-23 |
| ENSG00000103148\|NPRL3 | 3544.8830 | -1.2322 | ##### | ####### | 1.0837E-23 | 9.3917E-23 |
| ENSG00000185909\|KLHDC8B | 567.4376 | -1.4490 | ##### | ####### | 1.0941E-23 | 9.4794E-23 |
| ENSG00000145220\|LYAR | 4786.1053 | 1.2804 | ##### | 10.0325 | 1.0970E-23 | 9.5019E-23 |
| ENSG00000116353\|MECR | 2029.4612 | 1.2397 | ##### | 10.0315 | 1.1083E-23 | 9.5943E-23 |
| ENSG00000101115\|SALL4 | 315.0687 | 1.6720 | ##### | 10.0315 | 1.1083E-23 | 9.5943E-23 |
| ENSG00000187094\|CCK | 85.7880 | -8.8750 | ##### | ####### | 1.1588E-23 | 1.0028E-22 |
| ENSG00000179388\|EGR3 | 58.1664 | 4.6785 | ##### | 10.0267 | 1.1628E-23 | 1.0060E-22 |
| ENSG00000150995\|ITPR1 | 2082.2741 | 1.2664 | ##### | 10.0248 | 1.1861E-23 | 1.0259E-22 |
| ENSG00000268006\|PTOV1-AS1 | 439.0473 | -1.5582 | ##### | ####### | 1.2114E-23 | 1.0474E-22 |
| ENSG00000074317\|SNCB | 502.0485 | 10.6920 | ##### | 10.0195 | 1.2513E-23 | 1.0816E-22 |
| ENSG00000130818\|ZNF426 | 462.1391 | 1.4734 | ##### | 10.0192 | 1.2550E-23 | 1.0845E-22 |
| ENSG00000165458\|INPPL1 | 19143.2967 | 1.1740 | ##### | 10.0189 | 1.2593E-23 | 1.0879E-22 |
| ENSG00000091073\|DTX2 | 1475.3908 | -1.2898 | ##### | ####### | 1.2911E-23 | 1.1150E-22 |
| ENSG00000185813\|PCYT2 | 2074.3894 | -1.2618 | ##### | ####### | 1.3078E-23 | 1.1291E-22 |
| ENSG00000064547\|LPAR2 | 549.4079 | -1.4688 | ##### | ####### | 1.3269E-23 | 1.1453E-22 |
| ENSG00000286938 | 139.4869 | -9.5184 | ##### | ####### | 1.3458E-23 | 1.1613E-22 |
| ENSG00000090520\|DNAJB11 | 4665.0153 | -1.2304 | ##### | ####### | 1.3544E-23 | 1.1680E-22 |
| ENSG00000102078\|SLC25A14 | 831.4460 | 1.4262 | ##### | 10.0117 | 1.3541E-23 | 1.1680E-22 |
| ENSG00000279495 | 142.0543 | 2.2173 | ##### | 10.0095 | 1.3843E-23 | 1.1934E-22 |

| | | | | | | |
|---|---|---|---|---|---|---|
| ENSG00000247400\|DNAJC3-DT | 155.7910 | -3.1094 | ##### | ####### | 1.3982E-23 | 1.2051E-22 |
| ENSG00000281207\|SLFNL1-AS1 | 335.0083 | -1.6485 | ##### | ####### | 1.5106E-23 | 1.3017E-22 |
| ENSG00000157212\|PAXIP1 | 5043.2436 | 1.2336 | ##### | 10.0008 | 1.5112E-23 | 1.3017E-22 |
| ENSG00000158458\|NRG2 | 984.7459 | 1.4730 | ##### | 9.9999 | 1.5257E-23 | 1.3139E-22 |
| ENSG00000101361\|NOP56 | 19012.4361 | 0.8489 | ##### | 9.9988 | 1.5423E-23 | 1.3278E-22 |
| ENSG00000109805\|NCAPG | 7965.5333 | 1.2414 | ##### | 9.9987 | 1.5446E-23 | 1.3294E-22 |
| ENSG00000180008\|SOCS4 | 6366.1919 | 1.2168 | ##### | 9.9983 | 1.5508E-23 | 1.3343E-22 |
| ENSG00000102081\|FMR1 | 4116.9547 | 1.2525 | ##### | 9.9982 | 1.5520E-23 | 1.3350E-22 |
| ENSG00000196867\|ZFP28 | 890.9254 | -1.3819 | ##### | -9.9980 | 1.5552E-23 | 1.3374E-22 |
| ENSG00000166188\|ZNF319 | 2050.7506 | -1.2572 | ##### | -9.9963 | 1.5820E-23 | 1.3600E-22 |
| ENSG00000063245\|EPN1 | 10779.7162 | -0.9411 | ##### | -9.9943 | 1.6147E-23 | 1.3877E-22 |
| ENSG00000196839\|ADA | 5804.6628 | 1.1930 | ##### | 9.9923 | 1.6465E-23 | 1.4146E-22 |
| ENSG00000164543\|STK17A | 3490.6025 | -1.2188 | ##### | -9.9913 | 1.6637E-23 | 1.4290E-22 |
| ENSG00000168056\|LTBP3 | 5108.3960 | -1.1965 | ##### | -9.9912 | 1.6662E-23 | 1.4307E-22 |
| ENSG00000146250\|PRSS35 | 96.0514 | -2.8877 | ##### | -9.9886 | 1.7097E-23 | 1.4677E-22 |
| ENSG00000197620\|EOLA1 | 1422.9653 | 1.3475 | ##### | 9.9863 | 1.7493E-23 | 1.5012E-22 |
| ENSG00000128335\|APOL2 | 1913.7921 | -1.2472 | ##### | -9.9849 | 1.7755E-23 | 1.5233E-22 |
| ENSG00000237846 | 161.9032 | 2.1150 | ##### | 9.9844 | 1.7832E-23 | 1.5295E-22 |
| ENSG00000136848\|DAB2IP | 2429.2020 | -1.2619 | ##### | -9.9835 | 1.8008E-23 | 1.5441E-22 |
| ENSG00000120314\|WDR55 | 6924.7899 | 1.0854 | ##### | 9.9824 | 1.8205E-23 | 1.5605E-22 |
| ENSG00000007944\|MYLIP | 559.6544 | 1.6536 | ##### | 9.9817 | 1.8322E-23 | 1.5701E-22 |
| ENSG00000076685\|NT5C2 | 7167.6101 | -1.2637 | ##### | -9.9789 | 1.8860E-23 | 1.6157E-22 |
| ENSG00000135454\|B4GALNT1 | 14208.1827 | -1.1770 | ##### | -9.9774 | 1.9143E-23 | 1.6395E-22 |
| ENSG00000232555 | 418.0029 | ####### | ##### | -9.9734 | 1.9928E-23 | 1.7062E-22 |
| ENSG00000182010\|RTKN2 | 567.4401 | 1.5489 | ##### | 9.9731 | 1.9984E-23 | 1.7106E-22 |
| ENSG00000119723\|COQ6 | 3124.8979 | 1.2328 | ##### | 9.9717 | 2.0263E-23 | 1.7339E-22 |
| ENSG00000179715\|PCED1B | 161.0723 | -2.3769 | ##### | -9.9709 | 2.0443E-23 | 1.7489E-22 |
| ENSG00000148773\|MKI67 | 15554.8996 | 1.2269 | ##### | 9.9704 | 2.0533E-23 | 1.7561E-22 |
| ENSG00000172508\|CARNS1 | 106.7897 | 2.2912 | ##### | 9.9698 | 2.0656E-23 | 1.7661E-22 |
| ENSG00000260916\|CCPG1 | 4316.7693 | -1.2718 | ##### | -9.9694 | 2.0751E-23 | 1.7737E-22 |
| ENSG00000153066\|TXNDC11 | 3539.5802 | -1.2208 | ##### | -9.9693 | 2.0759E-23 | 1.7739E-22 |
| ENSG00000260805 | 82.3326 | -2.9137 | ##### | -9.9677 | 2.1103E-23 | 1.8028E-22 |
| ENSG00000169504\|CLIC4 | 14080.7882 | -1.1569 | ##### | -9.9627 | 2.2197E-23 | 1.8957E-22 |
| ENSG00000210194\|MT-TE | 2030.1057 | 1.3446 | ##### | 9.9597 | 2.2868E-23 | 1.9524E-22 |
| ENSG00000287151\|C2orf27A | 206.3000 | 1.8873 | ##### | 9.9589 | 2.3060E-23 | 1.9682E-22 |
| ENSG00000092201\|SUPT16H | 22834.1964 | 1.1836 | ##### | 9.9584 | 2.3172E-23 | 1.9772E-22 |
| ENSG00000181827\|RFX7 | 3008.3688 | 1.1791 | ##### | 9.9540 | 2.4225E-23 | 2.0665E-22 |
| ENSG00000164032\|H2AZ1 | 31860.9378 | 1.1487 | ##### | 9.9493 | 2.5391E-23 | 2.1653E-22 |
| ENSG00000119707\|RBM25 | 11579.0748 | 1.1655 | ##### | 9.9474 | 2.5880E-23 | 2.2064E-22 |
| ENSG00000164105\|SAP30 | 488.2079 | -1.3971 | ##### | -9.9461 | 2.6226E-23 | 2.2353E-22 |
| ENSG00000196428\|TSC22D2 | 3197.0294 | 1.2853 | ##### | 9.9453 | 2.6430E-23 | 2.2516E-22 |
| ENSG00000237125\|HAND2-AS1 | 110.9697 | -9.2461 | ##### | -9.9453 | 2.6433E-23 | 2.2516E-22 |
| ENSG00000254349\|MIR2052HG | 78.7243 | -2.8557 | ##### | -9.9443 | 2.6717E-23 | 2.2752E-22 |
| ENSG00000293060 | 87.4858 | -2.9100 | ##### | -9.9396 | 2.8003E-23 | 2.3840E-22 |
| ENSG00000228300\|FAM174C | 4573.0365 | 1.2225 | ##### | 9.9370 | 2.8728E-23 | 2.4451E-22 |
| ENSG00000254004\|ZNF260 | 2876.2931 | -1.2275 | ##### | -9.9369 | 2.8750E-23 | 2.4462E-22 |
| ENSG00000113575\|PPP2CA | 20383.2817 | 1.0693 | ##### | 9.9341 | 2.9593E-23 | 2.5172E-22 |
| ENSG00000168505\|GBX2 | 1049.7329 | 1.5035 | ##### | 9.9338 | 2.9672E-23 | 2.5232E-22 |
| ENSG00000178115\|GOLGA8Q | 104.7380 | -2.4367 | ##### | -9.9331 | 2.9886E-23 | 2.5407E-22 |
| ENSG00000174137\|FAM53A | 462.9079 | 1.4277 | ##### | 9.9293 | 3.1049E-23 | 2.6389E-22 |
| ENSG00000099622\|CIRBP | 4831.0068 | 1.1823 | ##### | 9.9290 | 3.1137E-23 | 2.6455E-22 |
| ENSG00000089693\|MLF2 | 39922.1077 | 1.0855 | ##### | 9.9273 | 3.1667E-23 | 2.6898E-22 |
| ENSG00000179477\|ALOX12B | 91.8918 | 2.6788 | ##### | 9.9264 | 3.1961E-23 | 2.7140E-22 |
| ENSG00000111875\|ASF1A | 1331.8164 | -1.4304 | ##### | -9.9255 | 3.2245E-23 | 2.7373E-22 |
| ENSG00000109466\|KLHL2 | 1617.0846 | -1.2422 | ##### | -9.9239 | 3.2754E-23 | 2.7798E-22 |
| ENSG00000115163\|CENPA | 2719.0627 | 1.2836 | ##### | 9.9230 | 3.3066E-23 | 2.8054E-22 |
| ENSG00000237004\|ZNRF2P1 | 89.0121 | -2.3146 | ##### | -9.9219 | 3.3444E-23 | 2.8367E-22 |
| ENSG00000119048\|UBE2B | 3294.0741 | 1.2491 | ##### | 9.9216 | 3.3521E-23 | 2.8424E-22 |
| ENSG00000104613\|INTS10 | 6374.9720 | 1.1601 | ##### | 9.9183 | 3.4645E-23 | 2.9369E-22 |
| ENSG00000138794\|CASP6 | 845.0769 | -1.3584 | ##### | -9.9113 | 3.7180E-23 | 3.1509E-22 |
| ENSG00000206432\|TMEM200C | 251.3577 | -1.7839 | ##### | -9.9088 | 3.8137E-23 | 3.2311E-22 |
| ENSG00000075568\|TMEM131 | 8584.0925 | 1.1842 | ##### | 9.9076 | 3.8577E-23 | 3.2675E-22 |
| ENSG00000070718\|AP3M2 | 2223.5727 | 1.3325 | ##### | 9.9062 | 3.9132E-23 | 3.3135E-22 |

| | | | | | | |
|---|---|---|---|---|---|---|
| ENSG00000198598\|MMP17 | 1187.4586 | 1.3761 | ##### | 9.8986 | 4.2200E-23 | 3.5723E-22 |
| ENSG00000234975\|FTH1P2 | 96.0822 | -2.4618 | ##### | -9.8975 | 4.2700E-23 | 3.6136E-22 |
| ENSG00000146151\|HMGCLL1 | 130.2541 | -2.5543 | ##### | -9.8967 | 4.3023E-23 | 3.6399E-22 |
| ENSG00000197070\|ARRDC1 | 1870.5284 | -1.2054 | ##### | -9.8941 | 4.4171E-23 | 3.7359E-22 |
| ENSG00000166002\|SMCO4 | 747.2179 | -1.2987 | ##### | -9.8926 | 4.4813E-23 | 3.7892E-22 |
| ENSG00000104413\|ESRP1 | 53.5476 | -4.0355 | ##### | -9.8926 | 4.4827E-23 | 3.7893E-22 |
| ENSG00000100906\|NFKBIA | 10819.5277 | 1.2400 | ##### | 9.8910 | 4.5540E-23 | 3.8485E-22 |
| ENSG00000197106\|SLC6A17 | 4511.2048 | 1.2136 | ##### | 9.8909 | 4.5571E-23 | 3.8500E-22 |
| ENSG00000163297\|ANTXR2 | 2413.5895 | 1.2911 | ##### | 9.8890 | 4.6459E-23 | 3.9239E-22 |
| ENSG00000276317 | 157.6887 | 9.0525 | ##### | 9.8864 | 4.7674E-23 | 4.0254E-22 |
| ENSG00000092094\|OSGEP | 1426.4798 | 1.3314 | ##### | 9.8856 | 4.8070E-23 | 4.0577E-22 |
| ENSG00000148158\|SNX30 | 1022.7637 | -1.3061 | ##### | -9.8841 | 4.8809E-23 | 4.1189E-22 |
| ENSG00000239911\|PRKAG2-AS1 | 168.5672 | 2.6743 | ##### | 9.8833 | 4.9191E-23 | 4.1500E-22 |
| ENSG00000028116\|VRK2 | 5267.7566 | -1.2386 | ##### | -9.8816 | 5.0033E-23 | 4.2198E-22 |
| ENSG00000164073\|MFSD8 | 1079.9112 | -1.3451 | ##### | -9.8803 | 5.0693E-23 | 4.2743E-22 |
| ENSG00000116771\|AGMAT | 908.0713 | -1.3243 | ##### | -9.8791 | 5.1294E-23 | 4.3238E-22 |
| ENSG00000149115\|TNKS1BP1 | 7322.1043 | -1.1814 | ##### | -9.8777 | 5.2029E-23 | 4.3844E-22 |
| ENSG00000145494\|NDUFS6 | 3660.5441 | -1.2788 | ##### | -9.8767 | 5.2515E-23 | 4.4242E-22 |
| ENSG00000151746\|BICD1 | 3665.6735 | 1.2002 | ##### | 9.8704 | 5.5965E-23 | 4.7135E-22 |
| ENSG00000171956\|FOXB1 | 482.8122 | 10.4101 | ##### | 9.8700 | 5.6163E-23 | 4.7288E-22 |
| ENSG00000127837\|AAMP | 13092.9264 | 1.1084 | ##### | 9.8694 | 5.6521E-23 | 4.7576E-22 |
| ENSG00000079616\|KIF22 | 10615.4878 | 1.1937 | ##### | 9.8691 | 5.6680E-23 | 4.7696E-22 |
| ENSG00000100941\|PNN | 9896.9626 | 1.1301 | ##### | 9.8674 | 5.7660E-23 | 4.8507E-22 |
| ENSG00000167548\|KMT2D | 10169.0044 | 1.3002 | ##### | 9.8658 | 5.8576E-23 | 4.9264E-22 |
| ENSG00000163462\|TRIM46 | 1703.2513 | 1.3015 | ##### | 9.8652 | 5.8921E-23 | 4.9540E-22 |
| ENSG00000123815\|COQ8B | 2163.3680 | -1.2611 | ##### | -9.8634 | 6.0002E-23 | 5.0436E-22 |
| ENSG00000227741 | 129.5110 | 2.0218 | ##### | 9.8629 | 6.0274E-23 | 5.0650E-22 |
| ENSG00000095752\|IL11 | 366.2187 | -2.2384 | ##### | -9.8604 | 6.1790E-23 | 5.1909E-22 |
| ENSG00000100116\|GCAT | 466.8214 | -1.3912 | ##### | -9.8601 | 6.1974E-23 | 5.2049E-22 |
| ENSG00000125812\|GZF1 | 1972.4392 | 1.1959 | ##### | 9.8565 | 6.4244E-23 | 5.3940E-22 |
| ENSG00000276707 | 70.9084 | -8.3591 | ##### | -9.8562 | 6.4419E-23 | 5.4072E-22 |
| ENSG00000222036\|POTEM | 33.3922 | -3.8368 | ##### | -9.8548 | 6.5343E-23 | 5.4825E-22 |
| ENSG00000187187\|ZNF546 | 185.3157 | -1.8491 | ##### | -9.8548 | 6.5353E-23 | 5.4825E-22 |
| ENSG00000271581 | 33.8981 | -4.8447 | ##### | -9.8542 | 6.5745E-23 | 5.5139E-22 |
| ENSG00000161671\|EMC10 | 16330.4754 | -0.9006 | ##### | -9.8535 | 6.6203E-23 | 5.5507E-22 |
| ENSG00000100162\|CENPM | 1412.8273 | 1.3010 | ##### | 9.8526 | 6.6806E-23 | 5.5997E-22 |
| ENSG00000182054\|IDH2 | 2284.5483 | -1.2227 | ##### | -9.8523 | 6.6993E-23 | 5.6137E-22 |
| ENSG00000143390\|RFX5 | 9136.1687 | 1.2166 | ##### | 9.8469 | 7.0691E-23 | 5.9220E-22 |
| ENSG00000267160 | 743.2985 | 1.5763 | ##### | 9.8460 | 7.1294E-23 | 5.9708E-22 |
| ENSG00000008441\|NFIX | 699.6997 | -1.3368 | ##### | -9.8432 | 7.3334E-23 | 6.1400E-22 |
| ENSG00000257225 | 123.3913 | -2.7944 | ##### | -9.8431 | 7.3449E-23 | 6.1479E-22 |
| ENSG00000132017\|DCAF15 | 3713.1915 | 1.2384 | ##### | 9.8420 | 7.4215E-23 | 6.2102E-22 |
| ENSG00000186897\|C1QL4 | 330.2025 | 1.8080 | ##### | 9.8420 | 7.4243E-23 | 6.2108E-22 |
| ENSG00000138764\|CCNG2 | 771.0917 | 1.4233 | ##### | 9.8367 | 7.8249E-23 | 6.5442E-22 |
| ENSG00000106069\|CHN2 | 100.7162 | 3.1369 | ##### | 9.8364 | 7.8444E-23 | 6.5586E-22 |
| ENSG00000106683\|LIMK1 | 3190.7110 | -1.2397 | ##### | -9.8362 | 7.8651E-23 | 6.5741E-22 |
| ENSG00000218891\|ZNF579 | 3798.1369 | -1.2571 | ##### | -9.8361 | 7.8673E-23 | 6.5741E-22 |
| ENSG00000145416\|MARCHF1 | 105.5671 | -2.8129 | ##### | -9.8320 | 8.2007E-23 | 6.8507E-22 |
| ENSG00000066855\|MTFR1 | 5892.2245 | -1.1483 | ##### | -9.8317 | 8.2230E-23 | 6.8675E-22 |
| ENSG00000168538\|TRAPPC11 | 1915.3444 | -1.2236 | ##### | -9.8236 | 8.9116E-23 | 7.4405E-22 |
| ENSG00000183496\|MEX3B | 990.3296 | 1.3379 | ##### | 9.8203 | 9.2030E-23 | 7.6816E-22 |
| ENSG00000143355\|LHX9 | 146.2862 | -3.4266 | ##### | -9.8181 | 9.4128E-23 | 7.8545E-22 |
| ENSG00000126814\|TRMT5 | 2842.7740 | 0.9632 | ##### | 9.8175 | 9.4695E-23 | 7.8996E-22 |
| ENSG00000162704\|ARPC5 | 14056.9078 | -0.9771 | ##### | -9.8161 | 9.5947E-23 | 8.0018E-22 |
| ENSG00000269439\|PGLS-DT | 153.0851 | -2.4083 | ##### | -9.8156 | 9.6463E-23 | 8.0426E-22 |
| ENSG00000012174\|MBTPS2 | 3402.1481 | -1.2215 | ##### | -9.8145 | 9.7555E-23 | 8.1314E-22 |
| ENSG00000263020 | 1825.7260 | 1.8046 | ##### | 9.8138 | 9.8152E-23 | 8.1789E-22 |
| ENSG00000267454\|ZNF582-DT | 68.8887 | -2.9011 | ##### | -9.8133 | 9.8736E-23 | 8.2252E-22 |
| ENSG00000117480\|FAAH | 212.6730 | -4.4519 | ##### | -9.8086 | 1.0342E-22 | 8.6127E-22 |
| ENSG00000156011\|PSD3 | 1291.8270 | -1.2972 | ##### | -9.8082 | 1.0378E-22 | 8.6410E-22 |
| ENSG00000287287 | 55.8731 | -3.6043 | ##### | -9.8065 | 1.0561E-22 | 8.7909E-22 |
| ENSG00000268061\|NAPA-AS1 | 318.9987 | -2.1640 | ##### | -9.8028 | 1.0945E-22 | 9.1080E-22 |
| ENSG00000274750\|H3C6 | 445.4586 | 1.5802 | ##### | 9.8024 | 1.0995E-22 | 9.1471E-22 |

| | | | | | | |
|---|---|---|---|---|---|---|
| ENSG00000223496\|EXOSC6 | 3299.6140 | -1.2455 | ##### | -9.8016 | 1.1081E-22 | 9.2160E-22 |
| ENSG00000277443\|MARCKS | 11549.2042 | -1.1748 | ##### | -9.8003 | 1.1228E-22 | 9.3349E-22 |
| ENSG00000161277\|THAP8 | 277.3807 | -1.5695 | ##### | -9.7994 | 1.1323E-22 | 9.4113E-22 |
| ENSG00000236283 | 43.5635 | -8.8612 | ##### | -9.7990 | 1.1371E-22 | 9.4486E-22 |
| ENSG00000234961 | 734.2348 | 1.5459 | ##### | 9.7976 | 1.1524E-22 | 9.5735E-22 |
| ENSG00000289688 | 394.9086 | 10.5822 | ##### | 9.7916 | 1.2232E-22 | 1.0158E-21 |
| ENSG00000104897\|SF3A2 | 7887.8555 | 1.2089 | ##### | 9.7902 | 1.2410E-22 | 1.0303E-21 |
| ENSG00000097033\|SH3GLB1 | 7428.2878 | -1.2514 | ##### | -9.7901 | 1.2416E-22 | 1.0306E-21 |
| ENSG00000204767\|INSYN2B | 431.3337 | 10.6906 | ##### | 9.7869 | 1.2815E-22 | 1.0634E-21 |
| ENSG00000131389\|SLC6A6 | 10268.7505 | -1.1887 | ##### | -9.7862 | 1.2911E-22 | 1.0711E-21 |
| ENSG00000076067\|RBMS2 | 5255.6938 | -1.2203 | ##### | -9.7860 | 1.2926E-22 | 1.0720E-21 |
| ENSG00000176731\|RBIS | 1964.1581 | -1.3024 | ##### | -9.7851 | 1.3041E-22 | 1.0813E-21 |
| ENSG00000135312\|HTR1B | 428.4502 | 1.5216 | ##### | 9.7835 | 1.3252E-22 | 1.0985E-21 |
| ENSG00000231245\|C1DP1 | 166.7456 | 2.5299 | ##### | 9.7808 | 1.3610E-22 | 1.1278E-21 |
| ENSG00000159433\|STARD9 | 635.6797 | 1.3883 | ##### | 9.7801 | 1.3712E-22 | 1.1359E-21 |
| ENSG00000258908 | 228.7998 | 1.8715 | ##### | 9.7795 | 1.3792E-22 | 1.1422E-21 |
| ENSG00000133195\|SLC39A11 | 2168.9057 | -1.1868 | ##### | -9.7788 | 1.3885E-22 | 1.1497E-21 |
| ENSG00000254685\|FPGT | 1565.6240 | -1.2840 | ##### | -9.7773 | 1.4089E-22 | 1.1662E-21 |
| ENSG00000227627 | 864.8309 | 1.6362 | ##### | 9.7765 | 1.4198E-22 | 1.1749E-21 |
| ENSG00000227053\|MUC12-AS1 | 114.9980 | -7.5781 | ##### | -9.7692 | 1.5272E-22 | 1.2634E-21 |
| ENSG00000256663 | 1470.4886 | 1.3344 | ##### | 9.7687 | 1.5344E-22 | 1.2690E-21 |
| ENSG00000289234\|ZNF496-DT | 357.9708 | -1.5660 | ##### | -9.7634 | 1.6159E-22 | 1.3361E-21 |
| ENSG00000072041\|SLC6A15 | 2720.5343 | 1.2059 | ##### | 9.7627 | 1.6275E-22 | 1.3453E-21 |
| ENSG00000248383\|PCDHAC1 | 1214.8260 | 1.2824 | ##### | 9.7624 | 1.6323E-22 | 1.3489E-21 |
| ENSG00000104154\|SLC30A4 | 613.3215 | -1.3649 | ##### | -9.7596 | 1.6781E-22 | 1.3864E-21 |
| ENSG00000054965\|FAM168A | 7043.2156 | 1.1599 | ##### | 9.7567 | 1.7260E-22 | 1.4256E-21 |
| ENSG00000138119\|MYOF | 46875.2184 | -1.1427 | ##### | -9.7559 | 1.7402E-22 | 1.4369E-21 |
| ENSG00000198353\|HOXC4 | 100.9459 | 2.4509 | ##### | 9.7553 | 1.7504E-22 | 1.4449E-21 |
| ENSG00000189077\|TMEM120A | 1541.5666 | -1.3237 | ##### | -9.7551 | 1.7537E-22 | 1.4472E-21 |
| ENSG00000106013\|ANKRD7 | 106.3766 | 2.3217 | ##### | 9.7543 | 1.7680E-22 | 1.4586E-21 |
| ENSG00000161204\|ABCF3 | 3439.9494 | -1.1826 | ##### | -9.7538 | 1.7768E-22 | 1.4655E-21 |
| ENSG00000250337\|PURPL | 382.6528 | -1.5753 | ##### | -9.7516 | 1.8167E-22 | 1.4979E-21 |
| ENSG00000134668\|SPOCD1 | 127.4872 | -1.9838 | ##### | -9.7511 | 1.8250E-22 | 1.5044E-21 |
| ENSG00000104388\|RAB2A | 11911.6653 | -1.1853 | ##### | -9.7490 | 1.8635E-22 | 1.5357E-21 |
| ENSG00000113916\|BCL6 | 652.5857 | -1.4042 | ##### | -9.7482 | 1.8778E-22 | 1.5471E-21 |
| ENSG00000139154\|AEBP2 | 5994.1531 | 0.7073 | ##### | 9.7481 | 1.8803E-22 | 1.5487E-21 |
| ENSG00000144554\|FANCD2 | 4265.0212 | 1.1878 | ##### | 9.7462 | 1.9159E-22 | 1.5775E-21 |
| ENSG00000185624\|P4HB | 59730.3849 | -0.8869 | ##### | -9.7440 | 1.9570E-22 | 1.6110E-21 |
| ENSG00000164074\|ABHD18 | 646.0910 | -1.5045 | ##### | -9.7422 | 1.9911E-22 | 1.6386E-21 |
| ENSG00000167105\|TMEM92 | 455.9638 | -6.0546 | ##### | -9.7391 | 2.0531E-22 | 1.6891E-21 |
| ENSG00000182240\|BACE2 | 3318.8085 | -1.1772 | ##### | -9.7369 | 2.0981E-22 | 1.7257E-21 |
| ENSG00000247317\|LY6E-DT | 79.7918 | -2.6619 | ##### | -9.7356 | 2.1254E-22 | 1.7477E-21 |
| ENSG00000171160\|MORN4 | 802.1371 | -1.3663 | ##### | -9.7345 | 2.1497E-22 | 1.7671E-21 |
| ENSG00000204882\|GPR20 | 50.8663 | -5.7142 | ##### | -9.7330 | 2.1801E-22 | 1.7916E-21 |
| ENSG00000196466\|ZNF799 | 231.2041 | -1.6760 | ##### | -9.7308 | 2.2291E-22 | 1.8314E-21 |
| ENSG00000103888\|CEMIP | 143.5001 | -1.8523 | ##### | -9.7307 | 2.2302E-22 | 1.8318E-21 |
| ENSG00000123427\|EEF1AKMT3 | 795.1316 | -1.3409 | ##### | -9.7304 | 2.2381E-22 | 1.8378E-21 |
| ENSG00000261602 | 659.4509 | -1.7419 | ##### | -9.7303 | 2.2390E-22 | 1.8380E-21 |
| ENSG00000214176\|PLEKHM1P1 | 82.2751 | 4.5135 | ##### | 9.7276 | 2.2995E-22 | 1.8872E-21 |
| ENSG00000247498\|GPRC5D-AS1 | 895.2469 | 1.4408 | ##### | 9.7266 | 2.3216E-22 | 1.9048E-21 |
| ENSG00000172403\|SYNPO2 | 364.4166 | 2.3408 | ##### | 9.7248 | 2.3640E-22 | 1.9391E-21 |
| ENSG00000169427\|KCNK9 | 389.2758 | 10.3033 | ##### | 9.7229 | 2.4073E-22 | 1.9740E-21 |
| ENSG00000176182\|MYPOP | 1038.0823 | -1.3270 | ##### | -9.7224 | 2.4200E-22 | 1.9839E-21 |
| ENSG00000278704 | 244.3345 | 1.7207 | ##### | 9.7207 | 2.4614E-22 | 2.0173E-21 |
| ENSG00000138604\|GLCE | 2526.6191 | -1.2475 | ##### | -9.7199 | 2.4806E-22 | 2.0325E-21 |
| ENSG00000175356\|SCUBE2 | 87.6770 | -2.5099 | ##### | -9.7194 | 2.4927E-22 | 2.0418E-21 |
| ENSG00000182718\|ANXA2 | 89858.6146 | -1.1661 | ##### | -9.7149 | 2.6046E-22 | 2.1329E-21 |
| ENSG00000171443\|ZNF524 | 865.7105 | -1.3327 | ##### | -9.7143 | 2.6202E-22 | 2.1451E-21 |
| ENSG00000102241\|HTATSF1 | 10501.7490 | 1.1011 | ##### | 9.7123 | 2.6728E-22 | 2.1875E-21 |
| ENSG00000120159\|CAAP1 | 1257.8326 | -1.2877 | ##### | -9.7114 | 2.6957E-22 | 2.2057E-21 |
| ENSG00000100650\|SRSF5 | 10699.9770 | 1.0007 | ##### | 9.7108 | 2.7127E-22 | 2.2190E-21 |
| ENSG00000165501\|LRR1 | 2007.6432 | 1.2871 | ##### | 9.7107 | 2.7149E-22 | 2.2202E-21 |
| ENSG00000198039\|ZNF273 | 202.1019 | -1.6802 | ##### | -9.7104 | 2.7218E-22 | 2.2251E-21 |

| | | | | | | |
|---|---|---|---|---|---|---|
| ENSG00000105939\|ZC3HAV1 | 4698.8948 | -1.1539 | ##### | -9.7104 | 2.7224E-22 | 2.2251E-21 |
| ENSG00000140455\|USP3 | 2681.9311 | -1.2179 | ##### | -9.7091 | 2.7567E-22 | 2.2525E-21 |
| ENSG00000160949\|TONSL | 5712.2169 | 1.0514 | ##### | 9.7070 | 2.8160E-22 | 2.3003E-21 |
| ENSG00000279605 | 5360.1763 | 1.2690 | ##### | 9.7067 | 2.8220E-22 | 2.3046E-21 |
| ENSG00000166510\|CCDC68 | 1486.3829 | -1.3006 | ##### | -9.7057 | 2.8509E-22 | 2.3275E-21 |
| ENSG00000092421\|SEMA6A | 270.7884 | 1.9094 | ##### | 9.7054 | 2.8583E-22 | 2.3330E-21 |
| ENSG00000236467\|KCNMA1-AS1 | 117.2691 | 8.2746 | ##### | 9.6974 | 3.0942E-22 | 2.5248E-21 |
| ENSG00000197261\|C6orf141 | 233.0064 | 1.7680 | ##### | 9.6954 | 3.1555E-22 | 2.5741E-21 |
| ENSG00000122966\|CIT | 5441.7664 | 1.2085 | ##### | 9.6948 | 3.1714E-22 | 2.5864E-21 |
| ENSG00000180921\|FAM83H | 8704.3457 | -1.1240 | ##### | -9.6940 | 3.1981E-22 | 2.6075E-21 |
| ENSG00000260428\|SCX | 115.6583 | -2.0415 | ##### | -9.6911 | 3.2884E-22 | 2.6804E-21 |
| ENSG00000196371\|FUT4 | 1066.9039 | -1.4146 | ##### | -9.6891 | 3.3559E-22 | 2.7346E-21 |
| ENSG00000248932\|COPB2-DT | 215.5671 | -1.9869 | ##### | -9.6888 | 3.3651E-22 | 2.7413E-21 |
| ENSG00000186350\|RXRA | 4359.6182 | -1.1893 | ##### | -9.6884 | 3.3781E-22 | 2.7512E-21 |
| ENSG00000137497\|NUMA1 | 22862.1730 | 1.1373 | ##### | 9.6868 | 3.4306E-22 | 2.7932E-21 |
| ENSG00000176222\|ZNF404 | 163.8120 | -1.9737 | ##### | -9.6858 | 3.4660E-22 | 2.8212E-21 |
| ENSG00000135821\|GLUL | 6551.1120 | 0.9302 | ##### | 9.6840 | 3.5276E-22 | 2.8706E-21 |
| ENSG00000167123\|CERCAM | 2521.8226 | -1.1880 | ##### | -9.6836 | 3.5402E-22 | 2.8801E-21 |
| ENSG00000100629\|CEP128 | 1051.3300 | 1.3571 | ##### | 9.6822 | 3.5874E-22 | 2.9177E-21 |
| ENSG00000163762\|TM4SF18 | 110.9992 | -3.4149 | ##### | -9.6807 | 3.6430E-22 | 2.9621E-21 |
| ENSG00000107554\|DNMBP | 7273.5320 | 1.1830 | ##### | 9.6759 | 3.8158E-22 | 3.1018E-21 |
| ENSG00000076108\|BAZ2A | 19230.4926 | 1.1468 | ##### | 9.6739 | 3.8944E-22 | 3.1648E-21 |
| ENSG00000174370\|KCNJ5-AS1 | 339.8529 | 1.8187 | ##### | 9.6737 | 3.8988E-22 | 3.1666E-21 |
| ENSG00000001630\|CYP51A1 | 11082.0515 | -1.1910 | ##### | -9.6737 | 3.8987E-22 | 3.1666E-21 |
| ENSG00000268471\|MIR4453HG | 209.1775 | -1.7701 | ##### | -9.6727 | 3.9383E-22 | 3.1978E-21 |
| ENSG00000221983\|UBA52 | 31077.1882 | 1.1405 | ##### | 9.6685 | 4.1046E-22 | 3.3320E-21 |
| ENSG00000263753\|LINC00667 | 1842.8134 | -1.2015 | ##### | -9.6651 | 4.2431E-22 | 3.4435E-21 |
| ENSG00000104967\|NOVA2 | 168.4226 | 2.4213 | ##### | 9.6646 | 4.2641E-22 | 3.4596E-21 |
| ENSG00000162714\|ZNF496 | 2058.6038 | -1.2077 | ##### | -9.6641 | 4.2839E-22 | 3.4747E-21 |
| ENSG00000100888\|CHD8 | 8990.2145 | 1.1503 | ##### | 9.6622 | 4.3626E-22 | 3.5376E-21 |
| ENSG00000136156\|ITM2B | 10578.0096 | -1.1423 | ##### | -9.6602 | 4.4487E-22 | 3.6064E-21 |
| ENSG00000112739\|PRPF4B | 6320.6703 | 1.1853 | ##### | 9.6586 | 4.5220E-22 | 3.6649E-21 |
| ENSG00000165655\|ZNF503 | 2080.5838 | 1.2205 | ##### | 9.6584 | 4.5273E-22 | 3.6682E-21 |
| ENSG00000110717\|NDUFS8 | 9582.6846 | 1.1950 | ##### | 9.6577 | 4.5592E-22 | 3.6930E-21 |
| ENSG00000164169\|PRMT9 | 535.5937 | -1.4228 | ##### | -9.6575 | 4.5673E-22 | 3.6986E-21 |
| ENSG00000293185\|TPM3P9 | 688.6753 | -1.4144 | ##### | -9.6551 | 4.6772E-22 | 3.7865E-21 |
| ENSG00000138279\|ANXA7 | 11642.4694 | -0.7744 | ##### | -9.6526 | 4.7945E-22 | 3.8804E-21 |
| ENSG00000274933\|TBC1D3I | 205.4278 | 1.7982 | ##### | 9.6524 | 4.8014E-22 | 3.8850E-21 |
| ENSG00000132906\|CASP9 | 828.8529 | -1.3237 | ##### | -9.6512 | 4.8585E-22 | 3.9301E-21 |
| ENSG00000116017\|ARID3A | 2076.3806 | 1.2459 | ##### | 9.6501 | 4.9104E-22 | 3.9710E-21 |
| ENSG00000243335\|KCTD7 | 1132.2643 | 1.3613 | ##### | 9.6491 | 4.9605E-22 | 4.0104E-21 |
| ENSG00000164609\|SLU7 | 3618.5936 | 1.2080 | ##### | 9.6452 | 5.1510E-22 | 4.1633E-21 |
| ENSG00000179348\|GATA2 | 3664.6430 | 1.3159 | ##### | 9.6440 | 5.2109E-22 | 4.2106E-21 |
| ENSG00000100813\|ACIN1 | 17934.9904 | 1.1211 | ##### | 9.6408 | 5.3773E-22 | 4.3439E-21 |
| ENSG00000197563\|PIGN | 2769.7853 | -1.1771 | ##### | -9.6403 | 5.4029E-22 | 4.3634E-21 |
| ENSG00000185838\|GNB1L | 784.8194 | 1.3649 | ##### | 9.6394 | 5.4512E-22 | 4.4012E-21 |
| ENSG00000117724\|CENPF | 11575.8204 | 1.1587 | ##### | 9.6385 | 5.4999E-22 | 4.4393E-21 |
| ENSG00000164163\|ABCE1 | 14048.4320 | -1.1530 | ##### | -9.6361 | 5.6288E-22 | 4.5421E-21 |
| ENSG00000198786\|MT-ND5 | ########## | 1.2180 | ##### | 9.6354 | 5.6653E-22 | 4.5704E-21 |
| ENSG00000176597\|B3GNT5 | 2594.4284 | -1.1898 | ##### | -9.6354 | 5.6695E-22 | 4.5725E-21 |
| ENSG00000120963\|ZNF706 | 3485.7025 | -1.2317 | ##### | -9.6323 | 5.8392E-22 | 4.7081E-21 |
| ENSG00000137575\|SDCBP | 14171.2871 | -1.1805 | ##### | -9.6283 | 6.0712E-22 | 4.8939E-21 |
| ENSG00000244187\|TMEM141 | 1369.4407 | -1.2522 | ##### | -9.6249 | 6.2744E-22 | 5.0563E-21 |
| ENSG00000166450\|PRTG | 438.8302 | -1.5675 | ##### | -9.6213 | 6.5033E-22 | 5.2393E-21 |
| ENSG00000141401\|IMPA2 | 8150.3719 | -1.1959 | ##### | -9.6202 | 6.5732E-22 | 5.2942E-21 |
| ENSG00000160055\|TMEM234 | 693.7286 | -1.3990 | ##### | -9.6194 | 6.6233E-22 | 5.3331E-21 |
| ENSG00000110536\|PTPMT1 | 1968.5475 | -1.2699 | ##### | -9.6189 | 6.6540E-22 | 5.3564E-21 |
| ENSG00000275496 | 311.1766 | -1.4699 | ##### | -9.6180 | 6.7095E-22 | 5.3996E-21 |
| ENSG00000074201\|CLNS1A | 7945.0136 | 1.1820 | ##### | 9.6163 | 6.8247E-22 | 5.4908E-21 |
| ENSG00000125845\|BMP2 | 476.8325 | 1.6389 | ##### | 9.6153 | 6.8902E-22 | 5.5420E-21 |
| ENSG00000144354\|CDCA7 | 6776.0730 | 1.1613 | ##### | 9.6147 | 6.9308E-22 | 5.5732E-21 |
| ENSG00000130544\|ZNF557 | 687.8422 | 1.3913 | ##### | 9.6076 | 7.4234E-22 | 5.9677E-21 |
| ENSG00000180758\|GPR157 | 1437.7772 | -1.2350 | ##### | -9.6061 | 7.5334E-22 | 6.0545E-21 |

| | | | | | | |
|---|---|---|---|---|---|---|
| ENSG00000185127\|C6orf120 | 4061.2723 | -1.2125 | ##### | -9.6039 | 7.6944E-22 | 6.1822E-21 |
| ENSG00000136710\|CCDC115 | 1654.5467 | -1.1721 | ##### | -9.6029 | 7.7691E-22 | 6.2405E-21 |
| ENSG00000118242\|MREG | 2085.1313 | 1.0472 | ##### | 9.5996 | 8.0236E-22 | 6.4433E-21 |
| ENSG00000140682\|TGFB1I1 | 882.7554 | -1.5154 | ##### | -9.5989 | 8.0766E-22 | 6.4841E-21 |
| ENSG00000198369\|SPRED2 | 4893.6071 | 1.1406 | ##### | 9.5980 | 8.1529E-22 | 6.5435E-21 |
| ENSG00000242574\|HLA-DMB | 125.9810 | -2.1982 | ##### | -9.5976 | 8.1843E-22 | 6.5670E-21 |
| ENSG00000143553\|SNAPIN | 3883.7905 | 1.0096 | ##### | 9.5949 | 8.3995E-22 | 6.7379E-21 |
| ENSG00000114631\|PODXL2 | 2516.7523 | -1.2006 | ##### | -9.5946 | 8.4233E-22 | 6.7552E-21 |
| ENSG00000271646\|IRF2-DT | 91.7849 | -2.7534 | ##### | -9.5939 | 8.4788E-22 | 6.7978E-21 |
| ENSG00000132823\|OSER1 | 3386.0134 | 1.2334 | ##### | 9.5890 | 8.8965E-22 | 7.1308E-21 |
| ENSG00000142173\|COL6A2 | 3675.8024 | -1.1428 | ##### | -9.5887 | 8.9221E-22 | 7.1494E-21 |
| ENSG00000228288\|PCAT6 | 74.1218 | -2.5491 | ##### | -9.5880 | 8.9846E-22 | 7.1976E-21 |
| ENSG00000185101\|ANO9 | 92.7612 | -3.1462 | ##### | -9.5862 | 9.1350E-22 | 7.3161E-21 |
| ENSG00000254656\|RTL1 | 2761.5777 | 13.2524 | ##### | 9.5857 | 9.1847E-22 | 7.3539E-21 |
| ENSG00000149679\|CABLES2 | 1341.9191 | 1.1637 | ##### | 9.5851 | 9.2361E-22 | 7.3931E-21 |
| ENSG00000198535\|C2CD4A | 478.3740 | ####### | ##### | -9.5806 | 9.6458E-22 | 7.7190E-21 |
| ENSG00000229891\|LINC01315 | 103.9190 | -4.2079 | ##### | -9.5797 | 9.7322E-22 | 7.7860E-21 |
| ENSG00000107566\|ERLIN1 | 7691.1455 | -1.1573 | ##### | -9.5777 | 9.9251E-22 | 7.9382E-21 |
| ENSG00000119699\|TGFB3 | 220.8471 | -1.9293 | ##### | -9.5756 | 1.0128E-21 | 8.0981E-21 |
| ENSG00000039123\|MTREX | 15180.3877 | -1.1054 | ##### | -9.5737 | 1.0310E-21 | 8.2418E-21 |
| ENSG00000046604\|DSG2 | 20950.8146 | -1.1197 | ##### | -9.5705 | 1.0638E-21 | 8.5016E-21 |
| ENSG00000115073\|ACTR1B | 3495.9182 | 0.9112 | ##### | 9.5675 | 1.0947E-21 | 8.7463E-21 |
| ENSG00000144744\|UBA3 | 3745.3453 | -1.1717 | ##### | -9.5638 | 1.1352E-21 | 9.0674E-21 |
| ENSG00000163684\|RPP14 | 3271.7757 | 1.1901 | ##### | 9.5582 | 1.1979E-21 | 9.5652E-21 |
| ENSG00000139977\|NAA30 | 4396.1582 | 1.1359 | ##### | 9.5576 | 1.2047E-21 | 9.6173E-21 |
| ENSG00000153046\|CDYL | 3281.0824 | 1.1649 | ##### | 9.5575 | 1.2069E-21 | 9.6324E-21 |
| ENSG00000099284\|MACROH2A2 | 1412.7253 | -1.2745 | ##### | -9.5561 | 1.2228E-21 | 9.7569E-21 |
| ENSG00000172493\|AFF1 | 3116.9346 | -1.2164 | ##### | -9.5559 | 1.2247E-21 | 9.7692E-21 |
| ENSG00000184227\|ACOT1 | 955.2577 | 1.2999 | ##### | 9.5556 | 1.2281E-21 | 9.7940E-21 |
| ENSG00000172262\|ZNF131 | 6397.4532 | 1.0770 | ##### | 9.5553 | 1.2317E-21 | 9.8197E-21 |
| ENSG00000276085\|CCL3L3 | 350.1104 | 10.2335 | ##### | 9.5531 | 1.2583E-21 | 1.0029E-20 |
| ENSG00000169258\|GPRIN1 | 2195.0440 | 1.2310 | ##### | 9.5525 | 1.2657E-21 | 1.0085E-20 |
| ENSG00000280071 | 1725.2365 | -1.2967 | ##### | -9.5506 | 1.2895E-21 | 1.0272E-20 |
| ENSG00000146834\|MEPCE | 7138.0323 | 1.1034 | ##### | 9.5493 | 1.3057E-21 | 1.0399E-20 |
| ENSG00000005448\|WDR54 | 2671.7863 | 1.2049 | ##### | 9.5492 | 1.3068E-21 | 1.0404E-20 |
| ENSG00000102931\|ARL2BP | 9896.1233 | -1.5759 | ##### | -9.5480 | 1.3225E-21 | 1.0527E-20 |
| ENSG00000290453 | 89.7023 | -2.6832 | ##### | -9.5478 | 1.3252E-21 | 1.0544E-20 |
| ENSG00000242372\|EIF6 | 22966.0396 | 1.1607 | ##### | 9.5478 | 1.3253E-21 | 1.0544E-20 |
| ENSG00000181191\|PJA1 | 2478.1331 | 0.9525 | ##### | 9.5452 | 1.3586E-21 | 1.0806E-20 |
| ENSG00000244198\|ARHGEF35-AS1 | 504.6656 | -1.7309 | ##### | -9.5442 | 1.3712E-21 | 1.0902E-20 |
| ENSG00000253645 | 74.6160 | -4.1202 | ##### | -9.5436 | 1.3800E-21 | 1.0969E-20 |
| ENSG00000224721\|FLVCR2-AS1 | 30.7187 | -4.1717 | ##### | -9.5422 | 1.3985E-21 | 1.1114E-20 |
| ENSG00000224407 | 865.6524 | -1.3615 | ##### | -9.5390 | 1.4418E-21 | 1.1455E-20 |
| ENSG00000142675\|CNKSR1 | 201.4574 | 2.1956 | ##### | 9.5388 | 1.4452E-21 | 1.1479E-20 |
| ENSG00000197712\|FAM114A1 | 4591.5104 | -1.1637 | ##### | -9.5380 | 1.4556E-21 | 1.1558E-20 |
| ENSG00000197323\|TRIM33 | 7550.9781 | 1.1778 | ##### | 9.5370 | 1.4704E-21 | 1.1673E-20 |
| ENSG00000115616\|SLC9A2 | 643.6210 | -1.4113 | ##### | -9.5367 | 1.4746E-21 | 1.1703E-20 |
| ENSG00000162775\|RBM15 | 2056.5195 | 1.2332 | ##### | 9.5358 | 1.4873E-21 | 1.1801E-20 |
| ENSG00000131747\|TOP2A | 34567.6534 | 1.1405 | ##### | 9.5334 | 1.5219E-21 | 1.2073E-20 |
| ENSG00000106080\|FKBP14 | 5141.9778 | -1.1503 | ##### | -9.5324 | 1.5372E-21 | 1.2190E-20 |
| ENSG00000158805\|ZNF276 | 4021.5580 | 1.1530 | ##### | 9.5292 | 1.5856E-21 | 1.2571E-20 |
| ENSG00000164323\|CFAP97 | 3806.4149 | -1.0829 | ##### | -9.5282 | 1.5997E-21 | 1.2679E-20 |
| ENSG00000182827\|ACBD3 | 6045.2763 | -0.9359 | ##### | -9.5277 | 1.6087E-21 | 1.2747E-20 |
| ENSG00000183569\|SERHL2 | 249.1839 | 1.5331 | ##### | 9.5259 | 1.6363E-21 | 1.2962E-20 |
| ENSG00000143995\|MEIS1 | 220.4851 | 1.7522 | ##### | 9.5258 | 1.6382E-21 | 1.2974E-20 |
| ENSG00000014919\|COX15 | 5586.3789 | -0.9445 | ##### | -9.5231 | 1.6805E-21 | 1.3305E-20 |
| ENSG00000198752\|CDC42BPB | 18000.7123 | -1.0935 | ##### | -9.5206 | 1.7218E-21 | 1.3629E-20 |
| ENSG00000185090\|MANEAL | 3827.3625 | -1.1702 | ##### | -9.5188 | 1.7519E-21 | 1.3864E-20 |
| ENSG00000285796 | 192.5432 | 2.0848 | ##### | 9.5180 | 1.7656E-21 | 1.3968E-20 |
| ENSG00000196792\|STRN3 | 3721.7263 | 1.1097 | ##### | 9.5163 | 1.7952E-21 | 1.4199E-20 |
| ENSG00000278834 | 640.3495 | -1.7097 | ##### | -9.5115 | 1.8801E-21 | 1.4866E-20 |
| ENSG00000198382\|UVRAG | 2811.7311 | 1.2204 | ##### | 9.5088 | 1.9292E-21 | 1.5251E-20 |
| ENSG00000172465\|TCEAL1 | 1832.2427 | 1.1075 | ##### | 9.5078 | 1.9474E-21 | 1.5390E-20 |

| | | | | | | |
|---|---|---|---|---|---|---|
| ENSG00000155561\|NUP205 | 20531.0499 | 1.0943 | ##### | 9.5076 | 1.9514E-21 | 1.5417E-20 |
| ENSG00000176845\|METRNL | 371.1867 | -1.5202 | ##### | -9.5056 | 1.9887E-21 | 1.5708E-20 |
| ENSG00000162614\|NEXN | 649.0324 | 1.3631 | ##### | 9.5049 | 2.0030E-21 | 1.5817E-20 |
| ENSG00000138134\|STAMBPL1 | 1599.5550 | -1.2900 | ##### | -9.5038 | 2.0230E-21 | 1.5971E-20 |
| ENSG00000286190 | 139.8087 | 2.7461 | ##### | 9.5038 | 2.0244E-21 | 1.5976E-20 |
| ENSG00000268812\|LIF-AS2 | 126.9740 | 1.9254 | ##### | 9.5037 | 2.0248E-21 | 1.5976E-20 |
| ENSG00000143622\|RIT1 | 2976.4259 | -1.1855 | ##### | -9.5016 | 2.0662E-21 | 1.6299E-20 |
| ENSG00000163072\|NOSTRIN | 217.9380 | -2.0856 | ##### | -9.4983 | 2.1327E-21 | 1.6819E-20 |
| ENSG00000170175\|CHRNB1 | 879.7798 | 1.3050 | ##### | 9.4960 | 2.1821E-21 | 1.7204E-20 |
| ENSG00000082898\|XPO1 | 37613.5971 | 1.1306 | ##### | 9.4956 | 2.1898E-21 | 1.7260E-20 |
| ENSG00000104415\|CCN4 | 83.2865 | -8.8310 | ##### | -9.4954 | 2.1944E-21 | 1.7292E-20 |
| ENSG00000284948 | 165.5801 | -2.0846 | ##### | -9.4947 | 2.2091E-21 | 1.7403E-20 |
| ENSG00000134996\|OSTF1 | 1088.2379 | -1.2321 | ##### | -9.4946 | 2.2098E-21 | 1.7404E-20 |
| ENSG00000233276\|GPX1 | 3225.0594 | ####### | ##### | -9.4928 | 2.2492E-21 | 1.7709E-20 |
| ENSG00000184402\|SS18L1 | 2635.9394 | 1.1646 | ##### | 9.4927 | 2.2521E-21 | 1.7727E-20 |
| ENSG00000059769\|DNAJC25 | 1879.1900 | -1.1873 | ##### | -9.4905 | 2.3000E-21 | 1.8100E-20 |
| ENSG00000177679\|SRRM3 | 696.3817 | -1.3166 | ##### | -9.4902 | 2.3067E-21 | 1.8144E-20 |
| ENSG00000283764\|MIR6850 | 34.7337 | 4.5196 | ##### | 9.4902 | 2.3065E-21 | 1.8144E-20 |
| ENSG00000280064 | 543.4912 | -1.3580 | ##### | -9.4859 | 2.4035E-21 | 1.8899E-20 |
| ENSG00000215252\|GOLGA8B | 3042.1843 | 1.2262 | ##### | 9.4858 | 2.4054E-21 | 1.8909E-20 |
| ENSG00000255717\|SNHG1 | 3948.4291 | 1.2170 | ##### | 9.4842 | 2.4427E-21 | 1.9198E-20 |
| ENSG00000114867\|EIF4G1 | 48627.3856 | -1.0793 | ##### | -9.4827 | 2.4777E-21 | 1.9468E-20 |
| ENSG00000168310\|IRF2 | 1207.8610 | -1.2386 | ##### | -9.4810 | 2.5182E-21 | 1.9781E-20 |
| ENSG00000164171\|ITGA2 | 12600.7421 | 1.1939 | ##### | 9.4799 | 2.5447E-21 | 1.9984E-20 |
| ENSG00000149091\|DGKZ | 5207.3919 | 1.0488 | ##### | 9.4756 | 2.6525E-21 | 2.0825E-20 |
| ENSG00000181359\|HSP90AA6P | 31.3671 | 4.0099 | ##### | 9.4740 | 2.6939E-21 | 2.1144E-20 |
| ENSG00000151640\|DPYSL4 | 346.2596 | -1.5430 | ##### | -9.4721 | 2.7427E-21 | 2.1521E-20 |
| ENSG00000177426\|TGIF1 | 11821.3666 | 1.1265 | ##### | 9.4702 | 2.7942E-21 | 2.1920E-20 |
| ENSG00000229950\|TFAP2A-AS1 | 218.2170 | 1.9194 | ##### | 9.4676 | 2.8623E-21 | 2.2448E-20 |
| ENSG00000182600\|SNORC | 439.4966 | -1.5097 | ##### | -9.4658 | 2.9145E-21 | 2.2852E-20 |
| ENSG00000183773\|AIFM3 | 134.4306 | -2.2531 | ##### | -9.4654 | 2.9251E-21 | 2.2928E-20 |
| ENSG00000154153\|RETREG1 | 309.8854 | 1.6820 | ##### | 9.4651 | 2.9327E-21 | 2.2982E-20 |
| ENSG00000011478\|QPCTL | 1200.3621 | -1.3198 | ##### | -9.4634 | 2.9823E-21 | 2.3365E-20 |
| ENSG00000076513\|ANKRD13A | 4213.9448 | 1.1292 | ##### | 9.4564 | 3.1883E-21 | 2.4972E-20 |
| ENSG00000181163\|NPM1 | 92795.9664 | 1.1506 | ##### | 9.4557 | 3.2078E-21 | 2.5118E-20 |
| ENSG00000108848\|LUC7L3 | 9221.8321 | 1.1380 | ##### | 9.4551 | 3.2284E-21 | 2.5273E-20 |
| ENSG00000112941\|TENT4A | 4545.3230 | 1.1524 | ##### | 9.4534 | 3.2809E-21 | 2.5677E-20 |
| ENSG00000273154 | 1471.9816 | 1.2971 | ##### | 9.4533 | 3.2844E-21 | 2.5698E-20 |
| ENSG00000261064\|LINC02256 | 38.6536 | -5.4882 | ##### | -9.4524 | 3.3123E-21 | 2.5910E-20 |
| ENSG00000115350\|POLE4 | 3648.0202 | 0.8585 | ##### | 9.4521 | 3.3198E-21 | 2.5961E-20 |
| ENSG00000153250\|RBMS1 | 7008.2218 | -1.1042 | ##### | -9.4513 | 3.3472E-21 | 2.6168E-20 |
| ENSG00000003056\|M6PR | 22920.0199 | 1.1214 | ##### | 9.4491 | 3.4176E-21 | 2.6712E-20 |
| ENSG00000213085\|CFAP45 | 199.3820 | 1.9518 | ##### | 9.4467 | 3.4979E-21 | 2.7333E-20 |
| ENSG00000167645\|YIF1B | 6018.3633 | -1.1957 | ##### | -9.4466 | 3.5015E-21 | 2.7353E-20 |
| ENSG00000145730\|PAM | 6494.4920 | 1.1001 | ##### | 9.4458 | 3.5272E-21 | 2.7547E-20 |
| ENSG00000197956\|S100A6 | 51201.2244 | 1.1264 | ##### | 9.4420 | 3.6558E-21 | 2.8544E-20 |
| ENSG00000173531\|MST1 | 337.7472 | 1.6417 | ##### | 9.4399 | 3.7299E-21 | 2.9115E-20 |
| ENSG00000135778\|NTPCR | 2835.2284 | -1.1756 | ##### | -9.4398 | 3.7366E-21 | 2.9160E-20 |
| ENSG00000165548\|TMEM63C | 517.2401 | 1.5988 | ##### | 9.4386 | 3.7793E-21 | 2.9485E-20 |
| ENSG00000248527\|MTATP6P1 | 49119.3021 | 1.1922 | ##### | 9.4377 | 3.8121E-21 | 2.9733E-20 |
| ENSG00000118363\|SPCS2 | 9433.6869 | 1.1497 | ##### | 9.4351 | 3.9065E-21 | 3.0462E-20 |
| ENSG00000183161\|FANCF | 773.1120 | -1.3213 | ##### | -9.4347 | 3.9196E-21 | 3.0556E-20 |
| ENSG00000163508\|EOMES | 139.0491 | 2.0540 | ##### | 9.4307 | 4.0743E-21 | 3.1753E-20 |
| ENSG00000081026\|MAGI3 | 1499.7772 | -1.2044 | ##### | -9.4296 | 4.1147E-21 | 3.2053E-20 |
| ENSG00000122042\|UBL3 | 2571.9118 | -1.1887 | ##### | -9.4296 | 4.1148E-21 | 3.2053E-20 |
| ENSG00000135144\|DTX1 | 109.4924 | 2.6071 | ##### | 9.4290 | 4.1390E-21 | 3.2232E-20 |
| ENSG00000163491\|NEK10 | 224.1700 | 1.6956 | ##### | 9.4261 | 4.2546E-21 | 3.3124E-20 |
| ENSG00000145216\|FIP1L1 | 6417.1170 | 0.8534 | ##### | 9.4259 | 4.2646E-21 | 3.3184E-20 |
| ENSG00000008083\|JARID2 | 1894.2590 | 1.2321 | ##### | 9.4259 | 4.2640E-21 | 3.3184E-20 |
| ENSG00000130758\|MAP3K10 | 2028.2871 | -1.1828 | ##### | -9.4236 | 4.3591E-21 | 3.3911E-20 |
| ENSG00000168491\|CCDC110 | 334.1713 | 10.2119 | ##### | 9.4219 | 4.4280E-21 | 3.4438E-20 |
| ENSG00000138760\|SCARB2 | 10880.8373 | -0.9813 | ##### | -9.4211 | 4.4656E-21 | 3.4722E-20 |
| ENSG00000138036\|DYNC2LI1 | 887.7602 | -1.3495 | ##### | -9.4191 | 4.5487E-21 | 3.5358E-20 |

| | | | | | | |
|---|---|---|---|---|---|---|
| ENSG00000180104\|EXOC3 | 3504.8514 | -1.2317 | ##### | -9.4173 | 4.6272E-21 | 3.5959E-20 |
| ENSG00000233762\|RPS15P4 | 897.6714 | 1.3914 | ##### | 9.4159 | 4.6909E-21 | 3.6445E-20 |
| ENSG00000164366\|CCDC127 | 1593.0634 | -1.2048 | ##### | -9.4148 | 4.7418E-21 | 3.6830E-20 |
| ENSG00000224531\|SMIM13 | 3509.4420 | 1.1733 | ##### | 9.4138 | 4.7845E-21 | 3.7152E-20 |
| ENSG00000123607\|TTC21B | 1783.4163 | 1.1736 | ##### | 9.4135 | 4.8004E-21 | 3.7266E-20 |
| ENSG00000177675\|CD163L1 | 352.3216 | 1.7590 | ##### | 9.4099 | 4.9638E-21 | 3.8525E-20 |
| ENSG00000174579\|MSL2 | 2906.0770 | 1.1935 | ##### | 9.4098 | 4.9695E-21 | 3.8559E-20 |
| ENSG00000102226\|USP11 | 6488.5762 | 1.1640 | ##### | 9.4089 | 5.0113E-21 | 3.8874E-20 |
| ENSG00000156970\|BUB1B | 6043.0805 | 1.1772 | ##### | 9.4078 | 5.0670E-21 | 3.9295E-20 |
| ENSG00000143429\|LSP1P4 | 163.1486 | 2.1653 | ##### | 9.4069 | 5.1099E-21 | 3.9618E-20 |
| ENSG00000115524\|SF3B1 | 28544.2070 | 1.0947 | ##### | 9.4055 | 5.1805E-21 | 4.0154E-20 |
| ENSG00000036672\|USP2 | 449.4798 | 1.4316 | ##### | 9.4028 | 5.3141E-21 | 4.1180E-20 |
| ENSG00000122481\|RWDD3 | 643.1526 | -1.2669 | ##### | -9.4024 | 5.3343E-21 | 4.1325E-20 |
| ENSG00000131016\|AKAP12 | 53142.4412 | 1.1034 | ##### | 9.4017 | 5.3712E-21 | 4.1600E-20 |
| ENSG00000145907\|G3BP1 | 30336.6089 | -0.8921 | ##### | -9.3994 | 5.4881E-21 | 4.2495E-20 |
| ENSG00000255301 | 100.9328 | -3.9570 | ##### | -9.3967 | 5.6300E-21 | 4.3582E-20 |
| ENSG00000196639\|HRH1 | 2321.3395 | 1.2098 | ##### | 9.3966 | 5.6366E-21 | 4.3622E-20 |
| ENSG00000254143 | 332.2188 | 10.0498 | ##### | 9.3964 | 5.6440E-21 | 4.3668E-20 |
| ENSG00000120925\|RNF170 | 957.7361 | -1.3474 | ##### | -9.3943 | 5.7615E-21 | 4.4565E-20 |
| ENSG00000277969 | 200.5105 | -1.6498 | ##### | -9.3932 | 5.8184E-21 | 4.4994E-20 |
| ENSG00000104341\|LAPTM4B | 17600.5745 | -1.1287 | ##### | -9.3920 | 5.8854E-21 | 4.5500E-20 |
| ENSG00000140548\|ZNF710 | 1244.1267 | -1.2469 | ##### | -9.3888 | 6.0702E-21 | 4.6917E-20 |
| ENSG00000160746\|ANO10 | 1705.3081 | -1.1772 | ##### | -9.3872 | 6.1593E-21 | 4.7593E-20 |
| ENSG00000146666\|LINC00525 | 455.5960 | ####### | ##### | -9.3868 | 6.1856E-21 | 4.7784E-20 |
| ENSG00000157456\|CCNB2 | 3514.7825 | 1.1920 | ##### | 9.3853 | 6.2727E-21 | 4.8444E-20 |
| ENSG00000138741\|TRPC3 | 60.6057 | 3.7439 | ##### | 9.3841 | 6.3457E-21 | 4.8995E-20 |
| ENSG00000259953\|LINC02977 | 223.8001 | -1.8805 | ##### | -9.3824 | 6.4484E-21 | 4.9775E-20 |
| ENSG00000141068\|KSR1 | 1379.6925 | -1.2249 | ##### | -9.3753 | 6.8994E-21 | 5.3243E-20 |
| ENSG00000122008\|POLK | 2234.0766 | -1.2205 | ##### | -9.3735 | 7.0191E-21 | 5.4153E-20 |
| ENSG00000104812\|GYS1 | 5156.4922 | -0.6911 | ##### | -9.3700 | 7.2506E-21 | 5.5925E-20 |
| ENSG00000064961\|HMG20B | 9146.2359 | 0.8519 | ##### | 9.3690 | 7.3252E-21 | 5.6485E-20 |
| ENSG00000157353\|FCSK | 1415.9127 | -1.1783 | ##### | -9.3689 | 7.3283E-21 | 5.6494E-20 |
| ENSG00000003249\|DBNDD1 | 2324.1122 | -1.2450 | ##### | -9.3653 | 7.5844E-21 | 5.8454E-20 |
| ENSG00000116299\|ELAPOR1 | 115.8578 | -2.1272 | ##### | -9.3623 | 7.8021E-21 | 6.0116E-20 |
| ENSG00000180694\|TMEM64 | 4679.3773 | -1.1870 | ##### | -9.3609 | 7.9088E-21 | 6.0922E-20 |
| ENSG00000197782\|ZNF780A | 1170.2174 | -1.2067 | ##### | -9.3604 | 7.9433E-21 | 6.1172E-20 |
| ENSG00000143878\|RHOB | 4626.1232 | -1.1855 | ##### | -9.3575 | 8.1678E-21 | 6.2885E-20 |
| ENSG00000131051\|RBM39 | 15748.0347 | 1.0216 | ##### | 9.3563 | 8.2559E-21 | 6.3547E-20 |
| ENSG00000108784\|NAGLU | 2428.3747 | -1.2046 | ##### | -9.3542 | 8.4273E-21 | 6.4849E-20 |
| ENSG00000250731\|TPM3P6 | 86.8906 | -2.6183 | ##### | -9.3532 | 8.5064E-21 | 6.5442E-20 |
| ENSG00000168612\|ZSWIM1 | 1210.9576 | 1.2731 | ##### | 9.3514 | 8.6492E-21 | 6.6523E-20 |
| ENSG00000155970\|MICU3 | 451.1110 | 1.4679 | ##### | 9.3506 | 8.7158E-21 | 6.7018E-20 |
| ENSG00000235052 | 63.3218 | 2.5941 | ##### | 9.3487 | 8.8704E-21 | 6.8189E-20 |
| ENSG00000136371\|MTHFS | 1305.2406 | -1.2785 | ##### | -9.3470 | 9.0150E-21 | 6.9283E-20 |
| ENSG00000184897\|H1-10 | 18729.0093 | 1.1129 | ##### | 9.3468 | 9.0346E-21 | 6.9416E-20 |
| ENSG00000158773\|USF1 | 4671.2017 | 1.1820 | ##### | 9.3439 | 9.2848E-21 | 7.1319E-20 |
| ENSG00000124172\|ATP5F1E | 9177.2290 | 1.1376 | ##### | 9.3403 | 9.6035E-21 | 7.3748E-20 |
| ENSG00000143365\|RORC | 295.8663 | ####### | ##### | -9.3369 | 9.9189E-21 | 7.6151E-20 |
| ENSG00000251583\|MFFP2 | 50.8155 | -3.4812 | ##### | -9.3332 | 1.0272E-20 | 7.8840E-20 |
| ENSG00000156709\|AIFM1 | 7988.4647 | 1.0488 | ##### | 9.3330 | 1.0288E-20 | 7.8945E-20 |
| ENSG00000138448\|ITGAV | 47713.1962 | -1.1222 | ##### | -9.3313 | 1.0463E-20 | 8.0263E-20 |
| ENSG00000109756\|RAPGEF2 | 3138.0484 | -1.1908 | ##### | -9.3270 | 1.0896E-20 | 8.3568E-20 |
| ENSG00000181274\|FRAT2 | 601.4114 | -1.3474 | ##### | -9.3262 | 1.0975E-20 | 8.4154E-20 |
| ENSG00000143486\|EIF2D | 3701.6703 | -1.1868 | ##### | -9.3256 | 1.1039E-20 | 8.4618E-20 |
| ENSG00000287763 | 301.6358 | 9.9933 | ##### | 9.3217 | 1.1448E-20 | 8.7732E-20 |
| ENSG00000230071\|RPL4P6 | 73.2556 | -2.6848 | ##### | -9.3216 | 1.1461E-20 | 8.7789E-20 |
| ENSG00000106852\|LHX6 | 205.6360 | 1.7543 | ##### | 9.3216 | 1.1461E-20 | 8.7789E-20 |
| ENSG00000143324\|XPR1 | 7462.9660 | -1.1211 | ##### | -9.3205 | 1.1582E-20 | 8.8693E-20 |
| ENSG00000107331\|ABCA2 | 5821.0863 | -1.1197 | ##### | -9.3191 | 1.1730E-20 | 8.9799E-20 |
| ENSG00000260630\|SNAI3-AS1 | 353.2836 | -1.4447 | ##### | -9.3178 | 1.1883E-20 | 9.0949E-20 |
| ENSG00000167994\|RAB3IL1 | 631.6545 | -1.3450 | ##### | -9.3119 | 1.2559E-20 | 9.6102E-20 |
| ENSG00000119328\|ABITRAM | 1617.4797 | -1.2385 | ##### | -9.3114 | 1.2621E-20 | 9.6548E-20 |
| ENSG00000177674\|AGTRAP | 5634.7401 | -1.1573 | ##### | -9.3091 | 1.2892E-20 | 9.8599E-20 |

| | | | | | | |
|---|---|---|---|---|---|---|
| ENSG00000100767\|PAPLN | 994.9080 | -1.2411 | ##### | -9.3065 | 1.3211E-20 | 1.0101E-19 |
| ENSG00000040933\|INPP4A | 3232.3920 | 1.1552 | ##### | 9.3033 | 1.3611E-20 | 1.0404E-19 |
| ENSG00000121270\|ABCC11 | 40.1432 | -3.2192 | ##### | -9.3025 | 1.3718E-20 | 1.0483E-19 |
| ENSG00000163347\|CLDN1 | 19790.3365 | 0.9645 | ##### | 9.3007 | 1.3957E-20 | 1.0663E-19 |
| ENSG00000107020\|PLGRKT | 807.7955 | -1.3169 | ##### | -9.2959 | 1.4596E-20 | 1.1149E-19 |
| ENSG00000213462\|ERV3-1 | 325.8996 | -2.8936 | ##### | -9.2941 | 1.4844E-20 | 1.1335E-19 |
| ENSG00000099624\|ATP5F1D | 6622.2928 | 1.1579 | ##### | 9.2931 | 1.4989E-20 | 1.1443E-19 |
| ENSG00000171889\|MIR31HG | 424.3242 | 2.2235 | ##### | 9.2926 | 1.5060E-20 | 1.1494E-19 |
| ENSG00000213923\|CSNK1E | 9922.1645 | 0.9173 | ##### | 9.2924 | 1.5079E-20 | 1.1506E-19 |
| ENSG00000114805\|PLCH1 | 799.3244 | -1.3120 | ##### | -9.2920 | 1.5136E-20 | 1.1546E-19 |
| ENSG00000158604\|TMED4 | 8113.7481 | -0.6964 | ##### | -9.2915 | 1.5216E-20 | 1.1604E-19 |
| ENSG00000089692\|LAG3 | 243.2135 | 1.6311 | ##### | 9.2894 | 1.5523E-20 | 1.1835E-19 |
| ENSG00000164144\|ARFIP1 | 1799.4494 | -1.1673 | ##### | -9.2885 | 1.5651E-20 | 1.1930E-19 |
| ENSG00000166881\|NEMP1 | 6182.2406 | 1.0815 | ##### | 9.2883 | 1.5679E-20 | 1.1948E-19 |
| ENSG00000172819\|RARG | 3918.4164 | -1.1479 | ##### | -9.2873 | 1.5826E-20 | 1.2057E-19 |
| ENSG00000186019\|ZNF225-AS1 | 500.9721 | -1.7013 | ##### | -9.2864 | 1.5962E-20 | 1.2158E-19 |
| ENSG00000238243\|OR2W3 | 75.2798 | 3.3485 | ##### | 9.2852 | 1.6141E-20 | 1.2291E-19 |
| ENSG00000162825\|NBPF20 | 1729.3614 | 1.1735 | ##### | 9.2839 | 1.6335E-20 | 1.2435E-19 |
| ENSG00000100744\|GSKIP | 1451.5919 | -1.1395 | ##### | -9.2835 | 1.6405E-20 | 1.2486E-19 |
| ENSG00000167900\|TK1 | 9575.3812 | 1.1488 | ##### | 9.2825 | 1.6558E-20 | 1.2599E-19 |
| ENSG00000198960\|ARMCX6 | 1705.7964 | 1.2643 | ##### | 9.2801 | 1.6939E-20 | 1.2885E-19 |
| ENSG00000105281\|SLC1A5 | 26996.6910 | -0.9884 | ##### | -9.2759 | 1.7615E-20 | 1.3396E-19 |
| ENSG00000205309\|NT5M | 303.3123 | -1.5705 | ##### | -9.2752 | 1.7723E-20 | 1.3475E-19 |
| ENSG00000171763\|AFG2B | 1954.6823 | 1.1458 | ##### | 9.2743 | 1.7874E-20 | 1.3586E-19 |
| ENSG00000166228\|PCBD1 | 4800.7729 | -1.1428 | ##### | -9.2742 | 1.7899E-20 | 1.3602E-19 |
| ENSG00000196741\|LINC01560 | 138.7763 | -2.0736 | ##### | -9.2700 | 1.8610E-20 | 1.4139E-19 |
| ENSG00000274808\|TBC1D3B | 74.2467 | 2.5475 | ##### | 9.2687 | 1.8842E-20 | 1.4311E-19 |
| ENSG00000166446\|CDYL2 | 104.3932 | -3.7797 | ##### | -9.2666 | 1.9217E-20 | 1.4593E-19 |
| ENSG00000291054 | 185.9896 | 2.0513 | ##### | 9.2627 | 1.9928E-20 | 1.5129E-19 |
| ENSG00000088325\|TPX2 | 19115.3125 | 1.0020 | ##### | 9.2613 | 2.0190E-20 | 1.5323E-19 |
| ENSG00000167468\|GPX4 | 25465.5412 | 1.0966 | ##### | 9.2598 | 2.0485E-20 | 1.5543E-19 |
| ENSG00000146409\|SLC18B1 | 952.2716 | -1.2074 | ##### | -9.2540 | 2.1615E-20 | 1.6397E-19 |
| ENSG00000130270\|ATP8B3 | 1178.2428 | 1.2486 | ##### | 9.2524 | 2.1961E-20 | 1.6655E-19 |
| ENSG00000170571\|EMB | 857.9458 | 1.2572 | ##### | 9.2454 | 2.3442E-20 | 1.7774E-19 |
| ENSG00000203999\|LINC01270 | 672.0073 | 1.4136 | ##### | 9.2384 | 2.5028E-20 | 1.8971E-19 |
| ENSG00000145703\|IQGAP2 | 172.6853 | 2.4930 | ##### | 9.2373 | 2.5284E-20 | 1.9161E-19 |
| ENSG00000163811\|WDR43 | 12963.5499 | 0.9554 | ##### | 9.2360 | 2.5597E-20 | 1.9393E-19 |
| ENSG00000140299\|BNIP2 | 4665.9725 | 1.1607 | ##### | 9.2325 | 2.6429E-20 | 2.0018E-19 |
| ENSG00000254221\|PCDHGB1 | 183.8365 | -2.6637 | ##### | -9.2304 | 2.6954E-20 | 2.0410E-19 |
| ENSG00000156510\|HKDC1 | 2416.0954 | -8.9258 | ##### | -9.2278 | 2.7629E-20 | 2.0916E-19 |
| ENSG00000291158\|LINC00869 | 768.8767 | 1.2241 | ##### | 9.2265 | 2.7951E-20 | 2.1154E-19 |
| ENSG00000204520\|MICA | 2557.5870 | -1.0611 | ##### | -9.2260 | 2.8091E-20 | 2.1255E-19 |
| ENSG00000104783\|KCNN4 | 1794.6249 | -1.0431 | ##### | -9.2223 | 2.9082E-20 | 2.1999E-19 |
| ENSG00000119574\|ZBTB45 | 1248.7124 | -1.2635 | ##### | -9.2211 | 2.9404E-20 | 2.2237E-19 |
| ENSG00000068912\|ERLEC1 | 5231.8694 | -0.7474 | ##### | -9.2197 | 2.9799E-20 | 2.2530E-19 |
| ENSG00000205138\|SDHAF1 | 1110.2924 | -1.1929 | ##### | -9.2180 | 3.0275E-20 | 2.2885E-19 |
| ENSG00000176834\|VSIG10 | 3356.4876 | -1.1240 | ##### | -9.2174 | 3.0448E-20 | 2.3010E-19 |
| ENSG00000152102\|FAM168B | 20523.2736 | -1.0828 | ##### | -9.2155 | 3.0989E-20 | 2.3413E-19 |
| ENSG00000197635\|DPP4 | 89.8531 | -2.8641 | ##### | -9.2153 | 3.1038E-20 | 2.3443E-19 |
| ENSG00000260007 | 736.9595 | 1.5779 | ##### | 9.2146 | 3.1246E-20 | 2.3595E-19 |
| ENSG00000090861\|AARS1 | 16844.7455 | -1.0768 | ##### | -9.2129 | 3.1757E-20 | 2.3975E-19 |
| ENSG00000165152\|PGAP4 | 1840.9846 | -1.1534 | ##### | -9.2115 | 3.2171E-20 | 2.4280E-19 |
| ENSG00000198756\|COLGALT2 | 166.9947 | -1.8073 | ##### | -9.2081 | 3.3189E-20 | 2.5042E-19 |
| ENSG00000100568\|VTI1B | 11176.3663 | 0.7105 | ##### | 9.2081 | 3.3207E-20 | 2.5050E-19 |
| ENSG00000101161\|PRPF6 | 12119.1940 | 1.0806 | ##### | 9.2070 | 3.3551E-20 | 2.5303E-19 |
| ENSG00000011132\|APBA3 | 2201.8978 | 1.1916 | ##### | 9.2068 | 3.3593E-20 | 2.5328E-19 |
| ENSG00000054938\|CHRDL2 | 1079.7439 | ####### | ##### | -9.2036 | 3.4622E-20 | 2.6097E-19 |
| ENSG00000179085\|DPM3 | 2312.1422 | -1.1251 | ##### | -9.2034 | 3.4685E-20 | 2.6138E-19 |
| ENSG00000138821\|SLC39A8 | 5325.8540 | 0.7603 | ##### | 9.2031 | 3.4777E-20 | 2.6201E-19 |
| ENSG00000117632\|STMN1 | 14183.5306 | 1.1326 | ##### | 9.2023 | 3.5049E-20 | 2.6399E-19 |
| ENSG00000138606\|SHF | 206.6906 | 1.5344 | ##### | 9.2010 | 3.5453E-20 | 2.6697E-19 |
| ENSG00000168010\|ATG16L2 | 991.0121 | 1.2382 | ##### | 9.2002 | 3.5725E-20 | 2.6895E-19 |
| ENSG00000174780\|SRP72 | 20679.9381 | 0.9666 | ##### | 9.1996 | 3.5944E-20 | 2.7053E-19 |

| | | | | | | |
|---|---|---|---|---|---|---|
| ENSG00000170442\|KRT86 | 2081.9494 | -8.6239 | ##### | -9.1969 | 3.6840E-20 | 2.7721E-19 |
| ENSG00000235174\|RPL39P3 | 1580.3015 | 1.2851 | ##### | 9.1942 | 3.7776E-20 | 2.8418E-19 |
| ENSG00000092853\|CLSPN | 6333.7878 | 1.0871 | ##### | 9.1932 | 3.8119E-20 | 2.8669E-19 |
| ENSG00000262712 | 838.1609 | -1.3042 | ##### | -9.1890 | 3.9662E-20 | 2.9822E-19 |
| ENSG00000160991\|ORAI2 | 4052.7080 | -1.1215 | ##### | -9.1886 | 3.9794E-20 | 2.9913E-19 |
| ENSG00000251350\|LINC02475 | 42.0557 | -3.3507 | ##### | -9.1853 | 4.1041E-20 | 3.0843E-19 |
| ENSG00000100575\|TIMM9 | 2538.8453 | 1.1548 | ##### | 9.1812 | 4.2639E-20 | 3.2035E-19 |
| ENSG00000188215\|DCUN1D3 | 1472.3921 | -1.1909 | ##### | -9.1798 | 4.3204E-20 | 3.2452E-19 |
| ENSG00000109971\|HSPA8 | ########## | 0.7366 | ##### | 9.1796 | 4.3271E-20 | 3.2494E-19 |
| ENSG00000110080\|ST3GAL4 | 2840.7900 | -1.1246 | ##### | -9.1758 | 4.4838E-20 | 3.3663E-19 |
| ENSG00000158435\|CNOT11 | 6053.7203 | -1.1148 | ##### | -9.1746 | 4.5314E-20 | 3.4011E-19 |
| ENSG00000187109\|NAP1L1 | 44834.0051 | 0.7226 | ##### | 9.1723 | 4.6308E-20 | 3.4749E-19 |
| ENSG00000153823\|PID1 | 176.1525 | -2.1070 | ##### | -9.1719 | 4.6456E-20 | 3.4851E-19 |
| ENSG00000130202\|NECTIN2 | 7764.5750 | -1.0160 | ##### | -9.1711 | 4.6831E-20 | 3.5123E-19 |
| ENSG00000169375\|SIN3A | 7945.3799 | 1.0698 | ##### | 9.1704 | 4.7114E-20 | 3.5327E-19 |
| ENSG00000198074\|AKR1B10 | 80601.2030 | ####### | ##### | -9.1697 | 4.7414E-20 | 3.5542E-19 |
| ENSG00000145826\|LECT2 | 76.5666 | 8.0023 | ##### | 9.1692 | 4.7649E-20 | 3.5710E-19 |
| ENSG00000116991\|SIPA1L2 | 1729.3152 | -1.1929 | ##### | -9.1682 | 4.8078E-20 | 3.6022E-19 |
| ENSG00000134716\|CYP2J2 | 168.9372 | 2.3772 | ##### | 9.1643 | 4.9872E-20 | 3.7357E-19 |
| ENSG00000146540\|C7orf50 | 5125.9684 | -1.0877 | ##### | -9.1640 | 5.0005E-20 | 3.7447E-19 |
| ENSG00000131013\|PPIL4 | 4119.8574 | 1.1279 | ##### | 9.1633 | 5.0329E-20 | 3.7681E-19 |
| ENSG00000182606\|TRAK1 | 2435.8882 | -1.1461 | ##### | -9.1595 | 5.2127E-20 | 3.9017E-19 |
| ENSG00000117713\|ARID1A | 9969.7813 | 1.0992 | ##### | 9.1590 | 5.2375E-20 | 3.9193E-19 |
| ENSG00000167601\|AXL | 13320.6323 | -1.1621 | ##### | -9.1588 | 5.2492E-20 | 3.9270E-19 |
| ENSG00000119900\|OGFRL1 | 11304.9888 | 1.1191 | ##### | 9.1583 | 5.2721E-20 | 3.9432E-19 |
| ENSG00000001497\|LAS1L | 8185.3526 | 1.1212 | ##### | 9.1538 | 5.4958E-20 | 4.1094E-19 |
| ENSG00000279738 | 403.8411 | 3.3556 | ##### | 9.1514 | 5.6190E-20 | 4.2005E-19 |
| ENSG00000178694\|NSUN3 | 1709.5954 | -1.1021 | ##### | -9.1465 | 5.8789E-20 | 4.3937E-19 |
| ENSG00000110721\|CHKA | 3321.0255 | 1.1396 | ##### | 9.1462 | 5.8996E-20 | 4.4081E-19 |
| ENSG00000164465\|DCBLD1 | 1453.5236 | -1.1659 | ##### | -9.1440 | 6.0208E-20 | 4.4976E-19 |
| ENSG00000115966\|ATF2 | 3688.1274 | 1.1360 | ##### | 9.1425 | 6.1012E-20 | 4.5565E-19 |
| ENSG00000182087\|TMEM259 | 18369.0484 | 1.0251 | ##### | 9.1376 | 6.3858E-20 | 4.7678E-19 |
| ENSG00000075399\|VPS9D1 | 1323.7086 | -1.2295 | ##### | -9.1374 | 6.3968E-20 | 4.7748E-19 |
| ENSG00000185825\|BCAP31 | 23558.5146 | 0.8403 | ##### | 9.1354 | 6.5158E-20 | 4.8624E-19 |
| ENSG00000163349\|HIPK1 | 10476.3437 | 1.0886 | ##### | 9.1319 | 6.7331E-20 | 5.0233E-19 |
| ENSG00000130707\|ASS1 | 1492.5943 | -1.2088 | ##### | -9.1310 | 6.7891E-20 | 5.0639E-19 |
| ENSG00000244038\|DDOST | 24862.5523 | 0.5066 | ##### | 9.1301 | 6.8446E-20 | 5.1040E-19 |
| ENSG00000170523\|KRT83 | 5797.6974 | ####### | ##### | -9.1233 | 7.2858E-20 | 5.4316E-19 |
| ENSG00000188761\|BCL2L15 | 114.8756 | -4.0864 | ##### | -9.1207 | 7.4643E-20 | 5.5633E-19 |
| ENSG00000198498\|TMA16 | 1880.6951 | -1.1624 | ##### | -9.1199 | 7.5195E-20 | 5.6030E-19 |
| ENSG00000112365\|ZBTB24 | 2340.2932 | 1.1515 | ##### | 9.1166 | 7.7525E-20 | 5.7752E-19 |
| ENSG00000176720\|BOK | 3870.8426 | 1.1325 | ##### | 9.1115 | 8.1228E-20 | 6.0496E-19 |
| ENSG00000101166\|PRELID3B | 13454.7654 | 0.8986 | ##### | 9.1046 | 8.6576E-20 | 6.4462E-19 |
| ENSG00000166483\|WEE1 | 2722.9036 | 1.1474 | ##### | 9.1021 | 8.8563E-20 | 6.5925E-19 |
| ENSG00000125676\|THOC2 | 9848.0951 | 1.0858 | ##### | 9.1018 | 8.8868E-20 | 6.6136E-19 |
| ENSG00000146094\|DOK3 | 741.0061 | 1.3111 | ##### | 9.1000 | 9.0324E-20 | 6.7203E-19 |
| ENSG00000102144\|PGK1 | 58291.5726 | 1.0284 | ##### | 9.0996 | 9.0668E-20 | 6.7442E-19 |
| ENSG00000106003\|LFNG | 501.7098 | -1.4747 | ##### | -9.0987 | 9.1400E-20 | 6.7969E-19 |
| ENSG00000283378\|CNTNAP3C | 208.8590 | -1.7441 | ##### | -9.0982 | 9.1834E-20 | 6.8276E-19 |
| ENSG00000153944\|MSI2 | 5242.6889 | -1.1055 | ##### | -9.0972 | 9.2705E-20 | 6.8906E-19 |
| ENSG00000105429\|MEGF8 | 4266.3764 | -1.1522 | ##### | -9.0962 | 9.3515E-20 | 6.9490E-19 |
| ENSG00000291113 | 2317.1050 | 2.9096 | ##### | 9.0924 | 9.6879E-20 | 7.1973E-19 |
| ENSG00000249915\|PDCD6 | 4524.1147 | -1.1359 | ##### | -9.0874 | 1.0144E-19 | 7.5342E-19 |
| ENSG00000144645\|OSBPL10 | 1768.0815 | -1.1990 | ##### | -9.0863 | 1.0247E-19 | 7.6085E-19 |
| ENSG00000173110\|HSPA6 | 433.9460 | 1.4536 | ##### | 9.0862 | 1.0258E-19 | 7.6151E-19 |
| ENSG00000276900 | 215.0534 | -1.5631 | ##### | -9.0841 | 1.0456E-19 | 7.7579E-19 |
| ENSG00000160796\|NBEAL2 | 6020.3272 | -1.0769 | ##### | -9.0841 | 1.0456E-19 | 7.7579E-19 |
| ENSG00000019582\|CD74 | 81.8129 | -3.2282 | ##### | -9.0831 | 1.0552E-19 | 7.8278E-19 |
| ENSG00000161533\|ACOX1 | 3102.5839 | -1.1011 | ##### | -9.0817 | 1.0692E-19 | 7.9295E-19 |
| ENSG00000102043\|MTMR8 | 240.8745 | ####### | ##### | -9.0813 | 1.0733E-19 | 7.9579E-19 |
| ENSG00000151718\|WWC2 | 5492.5573 | -1.1172 | ##### | -9.0808 | 1.0779E-19 | 7.9902E-19 |
| ENSG00000115942\|ORC2 | 3777.0536 | 0.9052 | ##### | 9.0795 | 1.0909E-19 | 8.0842E-19 |
| ENSG00000161960\|EIF4A1 | 73029.1460 | 0.8571 | ##### | 9.0775 | 1.1105E-19 | 8.2276E-19 |

| | | | | | | |
|---|---|---|---|---|---|---|
| ENSG00000124140\|SLC12A5 | 375.6475 | 1.4636 | ##### | 9.0766 | 1.1206E-19 | 8.3001E-19 |
| ENSG00000174500\|GCSAM | 73.3745 | 4.4177 | ##### | 9.0747 | 1.1401E-19 | 8.4427E-19 |
| ENSG00000149262\|INTS4 | 2938.9805 | 1.1340 | ##### | 9.0745 | 1.1424E-19 | 8.4572E-19 |
| ENSG00000293241\|GTF2IP4 | 6336.1176 | -1.0941 | ##### | -9.0711 | 1.1784E-19 | 8.7220E-19 |
| ENSG00000258634 | 1758.2650 | 1.1399 | ##### | 9.0684 | 1.2083E-19 | 8.9412E-19 |
| ENSG00000106799\|TGFBR1 | 5469.6971 | -1.0830 | ##### | -9.0675 | 1.2181E-19 | 9.0114E-19 |
| ENSG00000141642\|ELAC1 | 398.2760 | -1.4626 | ##### | -9.0661 | 1.2331E-19 | 9.1198E-19 |
| ENSG00000242814 | 28.8038 | -4.2640 | ##### | -9.0648 | 1.2479E-19 | 9.2271E-19 |
| ENSG00000100065\|CARD10 | 2447.8273 | -1.1658 | ##### | -9.0634 | 1.2648E-19 | 9.3497E-19 |
| ENSG00000033170\|FUT8 | 3598.2181 | 1.1003 | ##### | 9.0629 | 1.2701E-19 | 9.3870E-19 |
| ENSG00000290792 | 1383.4531 | 1.2268 | ##### | 9.0628 | 1.2712E-19 | 9.3927E-19 |
| ENSG00000231991\|ANXA2P2 | 4556.1888 | -1.1417 | ##### | -9.0625 | 1.2746E-19 | 9.4152E-19 |
| ENSG00000198963\|RORB | 85.7751 | 3.6047 | ##### | 9.0591 | 1.3151E-19 | 9.7119E-19 |
| ENSG00000187653\|TMSB4XP8 | 329.5296 | -1.3096 | ##### | -9.0585 | 1.3228E-19 | 9.7668E-19 |
| ENSG00000140332\|TLE3 | 2953.6871 | 1.0797 | ##### | 9.0582 | 1.3258E-19 | 9.7859E-19 |
| ENSG00000168016\|TRANK1 | 374.0522 | -1.4778 | ##### | -9.0541 | 1.3774E-19 | 1.0165E-18 |
| ENSG00000141854\|MISP3 | 264.7755 | -1.4958 | ##### | -9.0534 | 1.3853E-19 | 1.0221E-18 |
| ENSG00000171503\|ETFDH | 1590.0103 | -1.1145 | ##### | -9.0470 | 1.4696E-19 | 1.0840E-18 |
| ENSG00000247993\|FOXD1-AS1 | 107.0567 | 2.3503 | ##### | 9.0466 | 1.4749E-19 | 1.0876E-18 |
| ENSG00000273786 | 77.1778 | 8.9793 | ##### | 9.0466 | 1.4755E-19 | 1.0878E-18 |
| ENSG00000102287\|GABRE | 1043.3919 | -1.2308 | ##### | -9.0413 | 1.5483E-19 | 1.1412E-18 |
| ENSG00000101236\|RNF24 | 2025.5227 | -1.1778 | ##### | -9.0398 | 1.5692E-19 | 1.1563E-18 |
| ENSG00000291205 | 120.8160 | 1.9707 | ##### | 9.0386 | 1.5871E-19 | 1.1692E-18 |
| ENSG00000140416\|TPM1 | 28308.4346 | -0.6998 | ##### | -9.0380 | 1.5962E-19 | 1.1756E-18 |
| ENSG00000291049\|FAM86B3P | 237.8906 | -1.8812 | ##### | -9.0365 | 1.6170E-19 | 1.1906E-18 |
| ENSG00000227081\|RPS27P3 | 3464.3653 | 1.1757 | ##### | 9.0341 | 1.6529E-19 | 1.2167E-18 |
| ENSG00000102349\|KLF8 | 386.3738 | -1.4655 | ##### | -9.0338 | 1.6585E-19 | 1.2206E-18 |
| ENSG00000198624\|CCDC69 | 117.6025 | 2.2397 | ##### | 9.0330 | 1.6709E-19 | 1.2294E-18 |
| ENSG00000166681\|BEX3 | 10008.7646 | 1.0760 | ##### | 9.0310 | 1.7016E-19 | 1.2516E-18 |
| ENSG00000213430\|HSPD1P1 | 608.7298 | 1.2488 | ##### | 9.0288 | 1.7358E-19 | 1.2765E-18 |
| ENSG00000162882\|HAAO | 235.3702 | 9.5515 | ##### | 9.0287 | 1.7370E-19 | 1.2771E-18 |
| ENSG00000108830\|RND2 | 154.9267 | 2.2716 | ##### | 9.0276 | 1.7551E-19 | 1.2900E-18 |
| ENSG00000117475\|BLZF1 | 2175.5566 | -1.1215 | ##### | -9.0262 | 1.7768E-19 | 1.3057E-18 |
| ENSG00000166503\|HDGFL3 | 4143.1577 | -0.8953 | ##### | -9.0254 | 1.7903E-19 | 1.3153E-18 |
| ENSG00000118579\|MED28 | 3790.4303 | 1.1352 | ##### | 9.0250 | 1.7974E-19 | 1.3202E-18 |
| ENSG00000293141 | 164.9717 | -1.6833 | ##### | -9.0249 | 1.7993E-19 | 1.3213E-18 |
| ENSG00000240445\|FOXO3B | 924.2229 | 1.2358 | ##### | 9.0232 | 1.8275E-19 | 1.3417E-18 |
| ENSG00000107372\|ZFAND5 | 17507.7791 | -1.0525 | ##### | -9.0217 | 1.8524E-19 | 1.3596E-18 |
| ENSG00000055163\|CYFIP2 | 365.6489 | 1.6650 | ##### | 9.0214 | 1.8574E-19 | 1.3629E-18 |
| ENSG00000227051\|C14orf132 | 263.5366 | 1.7149 | ##### | 9.0184 | 1.9092E-19 | 1.4006E-18 |
| ENSG00000197961\|ZNF121 | 2482.9944 | 1.1758 | ##### | 9.0157 | 1.9556E-19 | 1.4343E-18 |
| ENSG00000274414 | 91.0547 | -2.6127 | ##### | -9.0157 | 1.9561E-19 | 1.4343E-18 |
| ENSG00000105866\|SP4 | 1417.5413 | 1.2111 | ##### | 9.0152 | 1.9653E-19 | 1.4407E-18 |
| ENSG00000172409\|CLP1 | 1542.7569 | 0.7428 | ##### | 9.0115 | 2.0332E-19 | 1.4901E-18 |
| ENSG00000186106\|ANKRD46 | 818.7789 | -1.2982 | ##### | -9.0100 | 2.0609E-19 | 1.5100E-18 |
| ENSG00000167100\|SAMD14 | 884.0803 | 1.2760 | ##### | 9.0099 | 2.0628E-19 | 1.5110E-18 |
| ENSG00000224577\|LINC01117 | 68.5837 | 2.7192 | ##### | 9.0085 | 2.0893E-19 | 1.5301E-18 |
| ENSG00000117122\|MFAP2 | 315.0250 | 1.6451 | ##### | 9.0073 | 2.1122E-19 | 1.5465E-18 |
| ENSG00000100938\|GMPR2 | 5023.7266 | 1.0981 | ##### | 9.0065 | 2.1278E-19 | 1.5575E-18 |
| ENSG00000136327\|NKX2-8 | 225.5746 | 1.6959 | ##### | 9.0062 | 2.1335E-19 | 1.5613E-18 |
| ENSG00000121039\|RDH10 | 3457.3985 | -1.1571 | ##### | -9.0057 | 2.1426E-19 | 1.5676E-18 |
| ENSG00000270964\|MAP2K5-DT | 90.3917 | -4.0588 | ##### | -9.0045 | 2.1656E-19 | 1.5840E-18 |
| ENSG00000132199\|ENOSF1 | 11805.5778 | 1.2177 | ##### | 9.0031 | 2.1935E-19 | 1.6040E-18 |
| ENSG00000267801 | 900.0228 | -1.3452 | ##### | -8.9998 | 2.2615E-19 | 1.6534E-18 |
| ENSG00000236915\|CLCA4-AS1 | 74.3446 | -3.7780 | ##### | -8.9967 | 2.3265E-19 | 1.7005E-18 |
| ENSG00000286067 | 341.9666 | 1.3792 | ##### | 8.9962 | 2.3375E-19 | 1.7081E-18 |
| ENSG00000168067\|MAP4K2 | 1329.9121 | 1.1526 | ##### | 8.9931 | 2.4028E-19 | 1.7554E-18 |
| ENSG00000154856\|APCDD1 | 56.4702 | 3.1318 | ##### | 8.9931 | 2.4035E-19 | 1.7554E-18 |
| ENSG00000134954\|ETS1 | 12251.7846 | 1.1389 | ##### | 8.9911 | 2.4481E-19 | 1.7876E-18 |
| ENSG00000180769\|WDFY3-AS2 | 114.4461 | -2.0765 | ##### | -8.9873 | 2.5328E-19 | 1.8490E-18 |
| ENSG00000263327\|TAPT1-AS1 | 92.1632 | -2.5672 | ##### | -8.9866 | 2.5505E-19 | 1.8615E-18 |
| ENSG00000271361\|HTATSF1P2 | 478.2692 | -6.8380 | ##### | -8.9862 | 2.5583E-19 | 1.8667E-18 |
| ENSG00000251127 | 217.5579 | ####### | ##### | -8.9841 | 2.6088E-19 | 1.9031E-18 |

| | | | | | | |
|---|---|---|---|---|---|---|
| ENSG00000106333\|PCOLCE | 866.6690 | 1.2651 | ##### | 8.9839 | 2.6141E-19 | 1.9064E-18 |
| ENSG00000189164\|ZNF527 | 449.4776 | -1.3235 | ##### | -8.9834 | 2.6261E-19 | 1.9148E-18 |
| ENSG00000108773\|KAT2A | 9682.4804 | 0.8389 | ##### | 8.9830 | 2.6354E-19 | 1.9211E-18 |
| ENSG00000163348\|PYGO2 | 7489.7765 | 1.0514 | ##### | 8.9801 | 2.7041E-19 | 1.9706E-18 |
| ENSG00000264772 | 1121.9395 | 1.7430 | ##### | 8.9794 | 2.7233E-19 | 1.9842E-18 |
| ENSG00000292998\|MIR3171HG | 223.1355 | ####### | ##### | -8.9792 | 2.7271E-19 | 1.9864E-18 |
| ENSG00000116954\|RRAGC | 1990.8137 | -1.2013 | ##### | -8.9792 | 2.7282E-19 | 1.9867E-18 |
| ENSG00000145911\|N4BP3 | 738.3132 | 1.2962 | ##### | 8.9791 | 2.7288E-19 | 1.9867E-18 |
| ENSG00000169291\|SHE | 66.6111 | 3.8355 | ##### | 8.9790 | 2.7333E-19 | 1.9895E-18 |
| ENSG00000151366\|NDUFC2 | 7698.9358 | 1.0751 | ##### | 8.9751 | 2.8312E-19 | 2.0603E-18 |
| ENSG00000129559\|NEDD8 | 8958.8717 | 1.0591 | ##### | 8.9746 | 2.8430E-19 | 2.0683E-18 |
| ENSG00000147255\|IGSF1 | 147.0443 | -3.5486 | ##### | -8.9745 | 2.8453E-19 | 2.0695E-18 |
| ENSG00000275719 | 2141.2508 | -1.2086 | ##### | -8.9745 | 2.8474E-19 | 2.0706E-18 |
| ENSG00000111057\|KRT18 | ########## | -1.0656 | ##### | -8.9695 | 2.9784E-19 | 2.1653E-18 |
| ENSG00000102871\|TRADD | 1684.9219 | -1.2138 | ##### | -8.9686 | 3.0034E-19 | 2.1830E-18 |
| ENSG00000116199\|FAM20B | 6267.2475 | -1.0889 | ##### | -8.9674 | 3.0360E-19 | 2.2061E-18 |
| ENSG00000089159\|PXN | 46925.7351 | 1.1152 | ##### | 8.9673 | 3.0388E-19 | 2.2076E-18 |
| ENSG00000127928\|GNGT1 | 68.8672 | -2.4311 | ##### | -8.9672 | 3.0409E-19 | 2.2086E-18 |
| ENSG00000142188\|TMEM50B | 1355.4492 | -1.0933 | ##### | -8.9649 | 3.1066E-19 | 2.2558E-18 |
| ENSG00000223745\|CCDC18-AS1 | 420.5559 | -1.4172 | ##### | -8.9634 | 3.1484E-19 | 2.2855E-18 |
| ENSG00000196369\|SRGAP2B | 2959.9568 | 1.1037 | ##### | 8.9606 | 3.2302E-19 | 2.3444E-18 |
| ENSG00000089916\|GPATCH2L | 4139.0265 | 1.0876 | ##### | 8.9603 | 3.2369E-19 | 2.3486E-18 |
| ENSG00000112175\|BMP5 | 1636.4062 | 1.2170 | ##### | 8.9601 | 3.2445E-19 | 2.3536E-18 |
| ENSG00000143442\|POGZ | 5970.1692 | 1.1190 | ##### | 8.9589 | 3.2790E-19 | 2.3780E-18 |
| ENSG00000087266\|SH3BP2 | 718.2972 | 1.2710 | ##### | 8.9583 | 3.2978E-19 | 2.3911E-18 |
| ENSG00000117616\|RSRP1 | 1498.2306 | 1.2523 | ##### | 8.9580 | 3.3062E-19 | 2.3966E-18 |
| ENSG00000213015\|ZNF580 | 1256.5074 | -1.2180 | ##### | -8.9574 | 3.3253E-19 | 2.4098E-18 |
| ENSG00000164904\|ALDH7A1 | 3150.1148 | 1.1029 | ##### | 8.9548 | 3.4020E-19 | 2.4649E-18 |
| ENSG00000136490\|LIMD2 | 388.0700 | -1.4810 | ##### | -8.9521 | 3.4874E-19 | 2.5261E-18 |
| ENSG00000251359\|WWC2-AS2 | 54.1394 | -2.9493 | ##### | -8.9513 | 3.5121E-19 | 2.5434E-18 |
| ENSG00000291044 | 102.8035 | -1.8521 | ##### | -8.9510 | 3.5231E-19 | 2.5507E-18 |
| ENSG00000163040\|CCDC74A | 1733.4675 | 1.2168 | ##### | 8.9500 | 3.5539E-19 | 2.5724E-18 |
| ENSG00000279170\|TSTD3 | 363.1929 | -1.3641 | ##### | -8.9479 | 3.6226E-19 | 2.6215E-18 |
| ENSG00000259943\|ZMPSTE24-DT | 308.7096 | -1.7191 | ##### | -8.9472 | 3.6461E-19 | 2.6379E-18 |
| ENSG00000141668\|CBLN2 | 110.1406 | 2.2897 | ##### | 8.9453 | 3.7098E-19 | 2.6833E-18 |
| ENSG00000135404\|CD63 | 19202.1372 | -1.0782 | ##### | -8.9408 | 3.8645E-19 | 2.7945E-18 |
| ENSG00000144306\|SCRN3 | 1449.1984 | -1.1939 | ##### | -8.9406 | 3.8694E-19 | 2.7974E-18 |
| ENSG00000152749\|GPR180 | 2683.6093 | -0.8225 | ##### | -8.9371 | 3.9959E-19 | 2.8881E-18 |
| ENSG00000143494\|VASH2 | 174.5358 | -1.7678 | ##### | -8.9315 | 4.2029E-19 | 3.0370E-18 |
| ENSG00000088538\|DOCK3 | 1009.5592 | 1.2142 | ##### | 8.9287 | 4.3101E-19 | 3.1137E-18 |
| ENSG00000169946\|ZFPM2 | 153.3787 | -2.3016 | ##### | -8.9283 | 4.3252E-19 | 3.1239E-18 |
| ENSG00000125746\|EML2 | 3142.0288 | -1.0819 | ##### | -8.9276 | 4.3539E-19 | 3.1439E-18 |
| ENSG00000140807\|NKD1 | 140.8811 | 2.6521 | ##### | 8.9276 | 4.3552E-19 | 3.1440E-18 |
| ENSG00000237036\|ZEB1-AS1 | 595.5006 | -1.2689 | ##### | -8.9270 | 4.3786E-19 | 3.1601E-18 |
| ENSG00000137693\|YAP1 | 20046.3679 | 1.0496 | ##### | 8.9267 | 4.3875E-19 | 3.1658E-18 |
| ENSG00000260528\|FAM157C | 74.9135 | 2.1857 | ##### | 8.9256 | 4.4327E-19 | 3.1977E-18 |
| ENSG00000163918\|RFC4 | 2412.5164 | -1.0633 | ##### | -8.9246 | 4.4712E-19 | 3.2247E-18 |
| ENSG00000226800\|CACTIN-AS1 | 696.2394 | 1.7419 | ##### | 8.9238 | 4.5038E-19 | 3.2474E-18 |
| ENSG00000115053\|NCL | ########## | 0.9326 | ##### | 8.9225 | 4.5575E-19 | 3.2853E-18 |
| ENSG00000139350\|NEDD1 | 4497.8483 | 1.1168 | ##### | 8.9223 | 4.5676E-19 | 3.2918E-18 |
| ENSG00000170469\|SPATA24 | 218.5775 | 1.6429 | ##### | 8.9212 | 4.6146E-19 | 3.3249E-18 |
| ENSG00000254389\|RHPN1-AS1 | 231.2203 | 1.5686 | ##### | 8.9203 | 4.6489E-19 | 3.3488E-18 |
| ENSG00000056277\|ZNF280C | 1179.8184 | 1.2390 | ##### | 8.9185 | 4.7248E-19 | 3.4026E-18 |
| ENSG00000173726\|TOMM20 | 13469.9712 | -1.0888 | ##### | -8.9169 | 4.7955E-19 | 3.4520E-18 |
| ENSG00000008405\|CRY1 | 4121.2832 | 1.1261 | ##### | 8.9169 | 4.7957E-19 | 3.4520E-18 |
| ENSG00000277156\|TOMM40P1 | 357.9685 | 1.5399 | ##### | 8.9159 | 4.8402E-19 | 3.4832E-18 |
| ENSG00000153560\|UBP1 | 8631.1015 | 1.0566 | ##### | 8.9155 | 4.8549E-19 | 3.4929E-18 |
| ENSG00000176994\|SMCR8 | 5133.2744 | 1.1216 | ##### | 8.9142 | 4.9136E-19 | 3.5343E-18 |
| ENSG00000148842\|CNNM2 | 5038.0307 | -1.1331 | ##### | -8.9138 | 4.9332E-19 | 3.5476E-18 |
| ENSG00000101052\|IFT52 | 3129.9116 | 1.1450 | ##### | 8.9119 | 5.0169E-19 | 3.6068E-18 |
| ENSG00000179240\|GVQW3 | 407.9613 | 1.4462 | ##### | 8.9103 | 5.0914E-19 | 3.6595E-18 |
| ENSG00000293115\|CENPBD2P | 3218.4766 | -1.1033 | ##### | -8.9061 | 5.2848E-19 | 3.7976E-18 |
| ENSG00000103018\|CYB5B | 9599.6158 | -1.0137 | ##### | -8.9053 | 5.3256E-19 | 3.8261E-18 |

| | | | | | | |
|---|---|---|---|---|---|---|
| ENSG00000120549\|KIAA1217 | 861.8315 | -1.2647 | ##### | -8.9049 | 5.3454E-19 | 3.8394E-18 |
| ENSG00000166128\|RAB8B | 1556.5815 | -1.1637 | ##### | -8.9046 | 5.3561E-19 | 3.8461E-18 |
| ENSG00000283761 | 224.8786 | -3.4361 | ##### | -8.9030 | 5.4361E-19 | 3.9026E-18 |
| ENSG00000235934 | 64.6988 | 3.5407 | ##### | 8.9012 | 5.5267E-19 | 3.9667E-18 |
| ENSG00000290376\|HERC2P3 | 715.8153 | -2.6105 | ##### | -8.9008 | 5.5428E-19 | 3.9773E-18 |
| ENSG00000290908 | 484.3241 | 1.3503 | ##### | 8.9001 | 5.5799E-19 | 4.0029E-18 |
| ENSG00000206567 | 521.9376 | 1.3379 | ##### | 8.8999 | 5.5911E-19 | 4.0100E-18 |
| ENSG00000111640\|GAPDH | ########## | 1.0754 | ##### | 8.8994 | 5.6128E-19 | 4.0246E-18 |
| ENSG00000232821 | 229.5204 | 9.4976 | ##### | 8.8977 | 5.7008E-19 | 4.0868E-18 |
| ENSG00000233016\|SNHG7 | 3864.8261 | -0.9932 | ##### | -8.8973 | 5.7242E-19 | 4.1025E-18 |
| ENSG00000109118\|PHF12 | 5720.1601 | 0.9501 | ##### | 8.8967 | 5.7519E-19 | 4.1214E-18 |
| ENSG00000273088 | 207.5936 | 9.4452 | ##### | 8.8964 | 5.7669E-19 | 4.1312E-18 |
| ENSG00000165417\|GTF2A1 | 6139.5133 | 1.1272 | ##### | 8.8960 | 5.7909E-19 | 4.1474E-18 |
| ENSG00000231770\|TMEM44-AS1 | 286.4774 | -1.5387 | ##### | -8.8932 | 5.9373E-19 | 4.2512E-18 |
| ENSG00000136628\|EPRS1 | 19722.7037 | -1.0317 | ##### | -8.8928 | 5.9568E-19 | 4.2641E-18 |
| ENSG00000198918\|RPL39 | 18181.3130 | 1.0633 | ##### | 8.8922 | 5.9936E-19 | 4.2894E-18 |
| ENSG00000131187\|F12 | 2294.8938 | 1.1618 | ##### | 8.8918 | 6.0133E-19 | 4.3025E-18 |
| ENSG00000265916 | 61.7386 | 7.5970 | ##### | 8.8914 | 6.0322E-19 | 4.3150E-18 |
| ENSG00000187239\|FNBP1 | 3931.1574 | -1.0700 | ##### | -8.8905 | 6.0860E-19 | 4.3524E-18 |
| ENSG00000198646\|NCOA6 | 5889.2460 | 1.0616 | ##### | 8.8865 | 6.3056E-19 | 4.5084E-18 |
| ENSG00000107745\|MICU1 | 3491.5807 | -0.9436 | ##### | -8.8847 | 6.4091E-19 | 4.5813E-18 |
| ENSG00000232931\|LINC00342 | 251.8255 | 1.4731 | ##### | 8.8837 | 6.4653E-19 | 4.6204E-18 |
| ENSG00000171130\|ATP6V0E2 | 4574.7145 | 1.0994 | ##### | 8.8821 | 6.5633E-19 | 4.6893E-18 |
| ENSG00000265972\|TXNIP | 258.6676 | -1.6019 | ##### | -8.8816 | 6.5895E-19 | 4.7069E-18 |
| ENSG00000268942\|CKS1BP3 | 204.6786 | 1.6441 | ##### | 8.8816 | 6.5932E-19 | 4.7084E-18 |
| ENSG00000137992\|DBT | 1735.3615 | -1.1364 | ##### | -8.8814 | 6.6018E-19 | 4.7135E-18 |
| ENSG00000275993 | 16236.0428 | -1.0622 | ##### | -8.8786 | 6.7685E-19 | 4.8313E-18 |
| ENSG00000117395\|EBNA1BP2 | 27412.8557 | 1.0534 | ##### | 8.8781 | 6.8014E-19 | 4.8536E-18 |
| ENSG00000130244\|FAM98C | 917.6298 | -1.2505 | ##### | -8.8756 | 6.9567E-19 | 4.9633E-18 |
| ENSG00000168765\|GSTM4 | 1467.7053 | -1.1685 | ##### | -8.8748 | 7.0044E-19 | 4.9961E-18 |
| ENSG00000183032\|SLC25A21 | 254.6466 | 1.3910 | ##### | 8.8738 | 7.0675E-19 | 5.0399E-18 |
| ENSG00000113119\|TMCO6 | 951.2183 | 1.2993 | ##### | 8.8721 | 7.1757E-19 | 5.1158E-18 |
| ENSG00000093009\|CDC45 | 3369.5353 | 1.1198 | ##### | 8.8713 | 7.2291E-19 | 5.1527E-18 |
| ENSG00000198945\|L3MBTL3 | 733.5159 | 1.3136 | ##### | 8.8704 | 7.2856E-19 | 5.1917E-18 |
| ENSG00000159792\|PSKH1 | 3115.7804 | -1.0840 | ##### | -8.8688 | 7.3907E-19 | 5.2653E-18 |
| ENSG00000146802\|TMEM168 | 4352.0081 | -1.1116 | ##### | -8.8682 | 7.4371E-19 | 5.2971E-18 |
| ENSG00000164045\|CDC25A | 3200.0937 | 1.0773 | ##### | 8.8662 | 7.5664E-19 | 5.3879E-18 |
| ENSG00000197619\|ZNF615 | 595.7046 | -1.3553 | ##### | -8.8656 | 7.6063E-19 | 5.4151E-18 |
| ENSG00000107223\|EDF1 | 13772.3477 | -1.0827 | ##### | -8.8640 | 7.7195E-19 | 5.4944E-18 |
| ENSG00000130147\|SH3BP4 | 6783.8218 | 1.0917 | ##### | 8.8637 | 7.7413E-19 | 5.5085E-18 |
| ENSG00000166532\|RIMKLB | 3964.4951 | 1.1274 | ##### | 8.8636 | 7.7485E-19 | 5.5124E-18 |
| ENSG00000237054\|PRMT5-AS1 | 844.5749 | 1.2808 | ##### | 8.8626 | 7.8172E-19 | 5.5599E-18 |
| ENSG00000196205\|EEF1A1P5 | 23939.9893 | 1.0435 | ##### | 8.8601 | 7.9955E-19 | 5.6854E-18 |
| ENSG00000012983\|MAP4K5 | 6229.9171 | 1.0675 | ##### | 8.8598 | 8.0154E-19 | 5.6982E-18 |
| ENSG00000164877\|MICALL2 | 3132.6627 | -1.0929 | ##### | -8.8582 | 8.1297E-19 | 5.7770E-18 |
| ENSG00000081248\|CACNA1S | 199.7484 | ####### | ##### | -8.8582 | 8.1302E-19 | 5.7770E-18 |
| ENSG00000290126 | 96.6516 | 2.1159 | ##### | 8.8563 | 8.2680E-19 | 5.8735E-18 |
| ENSG00000215302\|WHAMMP4 | 155.0728 | -1.6223 | ##### | -8.8551 | 8.3600E-19 | 5.9375E-18 |
| ENSG00000286161 | 54.2195 | 3.4134 | ##### | 8.8543 | 8.4201E-19 | 5.9787E-18 |
| ENSG00000177738\|ANXA2R-OT1 | 920.1266 | 1.2857 | ##### | 8.8512 | 8.6550E-19 | 6.1440E-18 |
| ENSG00000206418\|RAB12 | 5151.2566 | 1.0817 | ##### | 8.8504 | 8.7194E-19 | 6.1883E-18 |
| ENSG00000051341\|POLQ | 2780.2357 | 1.1163 | ##### | 8.8478 | 8.9281E-19 | 6.3350E-18 |
| ENSG00000042445\|RETSAT | 4899.8700 | -1.0585 | ##### | -8.8463 | 9.0468E-19 | 6.4177E-18 |
| ENSG00000187624\|LIAT1 | 708.6577 | 1.2990 | ##### | 8.8462 | 9.0584E-19 | 6.4243E-18 |
| ENSG00000117360\|PRPF3 | 5780.7918 | 1.0954 | ##### | 8.8438 | 9.2547E-19 | 6.5620E-18 |
| ENSG00000090565\|RAB11FIP3 | 2843.8739 | -1.1073 | ##### | -8.8432 | 9.3045E-19 | 6.5958E-18 |
| ENSG00000213023\|SYT3 | 345.3517 | 1.4174 | ##### | 8.8370 | 9.8359E-19 | 6.9708E-18 |
| ENSG00000179611\|DGKZP1 | 551.8339 | 1.2323 | ##### | 8.8331 | 1.0186E-18 | 7.2170E-18 |
| ENSG00000249626\|PPP4R2P4 | 52.9107 | -3.0327 | ##### | -8.8327 | 1.0220E-18 | 7.2392E-18 |
| ENSG00000104332\|SFRP1 | 1077.5801 | 1.1860 | ##### | 8.8318 | 1.0303E-18 | 7.2968E-18 |
| ENSG00000124766\|SOX4 | 4753.0698 | 1.0364 | ##### | 8.8317 | 1.0310E-18 | 7.2999E-18 |
| ENSG00000157181\|ODR4 | 2625.1199 | -1.1166 | ##### | -8.8304 | 1.0432E-18 | 7.3843E-18 |
| ENSG00000134287\|ARF3 | 30003.0508 | 0.6319 | ##### | 8.8291 | 1.0552E-18 | 7.4675E-18 |

| | | | | | | |
|---|---|---|---|---|---|---|
| ENSG00000149150\|SLC43A1 | 90.9816 | -3.2519 | ##### | -8.8280 | 1.0652E-18 | 7.5368E-18 |
| ENSG00000152804\|HHEX | 673.9300 | -1.2819 | ##### | -8.8260 | 1.0852E-18 | 7.6765E-18 |
| ENSG00000120756\|PLS1 | 3420.2823 | -1.1034 | ##### | -8.8259 | 1.0858E-18 | 7.6787E-18 |
| ENSG00000182362\|YBEY | 765.3105 | -1.2320 | ##### | -8.8256 | 1.0887E-18 | 7.6975E-18 |
| ENSG00000118507\|AKAP7 | 291.0993 | -1.5887 | ##### | -8.8248 | 1.0963E-18 | 7.7496E-18 |
| ENSG00000115257\|PCSK4 | 293.8962 | 1.7922 | ##### | 8.8230 | 1.1144E-18 | 7.8754E-18 |
| ENSG00000126522\|ASL | 2763.3953 | -1.0908 | ##### | -8.8225 | 1.1193E-18 | 7.9084E-18 |
| ENSG00000164933\|SLC25A32 | 5052.2147 | -1.0886 | ##### | -8.8216 | 1.1283E-18 | 7.9697E-18 |
| ENSG00000216819\|TUBB2BP1 | 153.8972 | -1.6900 | ##### | -8.8204 | 1.1402E-18 | 8.0518E-18 |
| ENSG00000134851\|TMEM165 | 10321.7646 | 1.0663 | ##### | 8.8197 | 1.1476E-18 | 8.1025E-18 |
| ENSG00000293271\|SLC44A3-AS1 | 175.9216 | -1.9783 | ##### | -8.8188 | 1.1565E-18 | 8.1634E-18 |
| ENSG00000143457\|GOLPH3L | 1806.0574 | 1.1626 | ##### | 8.8186 | 1.1589E-18 | 8.1783E-18 |
| ENSG00000134183\|GNAT2 | 53.5380 | 2.7061 | ##### | 8.8174 | 1.1716E-18 | 8.2661E-18 |
| ENSG00000067064\|IDI1 | 5427.3134 | -1.1145 | ##### | -8.8167 | 1.1793E-18 | 8.3182E-18 |
| ENSG00000227117\|HORMAD2-AS1 | 33.0326 | -4.0810 | ##### | -8.8123 | 1.2259E-18 | 8.6448E-18 |
| ENSG00000167646\|DNAAF3 | 1064.2124 | 1.0671 | ##### | 8.8095 | 1.2576E-18 | 8.8662E-18 |
| ENSG00000115241\|PPM1G | 22855.5759 | 0.9724 | ##### | 8.8085 | 1.2680E-18 | 8.9375E-18 |
| ENSG00000101901\|ALG13 | 2432.6627 | 1.1583 | ##### | 8.8076 | 1.2780E-18 | 9.0060E-18 |
| ENSG00000225914\|TSBP1-AS1 | 107.8460 | 2.7684 | ##### | 8.8062 | 1.2950E-18 | 9.1233E-18 |
| ENSG00000228022\|HCG20 | 114.6934 | 2.9170 | ##### | 8.8054 | 1.3040E-18 | 9.1846E-18 |
| ENSG00000261526 | 334.8974 | 1.4039 | ##### | 8.8045 | 1.3144E-18 | 9.2557E-18 |
| ENSG00000137601\|NEK1 | 1029.7415 | -1.1472 | ##### | -8.8028 | 1.3339E-18 | 9.3908E-18 |
| ENSG00000196465\|MYL6B | 3708.1498 | 1.0771 | ##### | 8.8026 | 1.3373E-18 | 9.4128E-18 |
| ENSG00000138777\|PPA2 | 2741.9479 | -1.1300 | ##### | -8.8011 | 1.3549E-18 | 9.5343E-18 |
| ENSG00000035664\|DAPK2 | 154.3732 | -1.6776 | ##### | -8.8000 | 1.3676E-18 | 9.6216E-18 |
| ENSG00000275765 | 170.5748 | -1.9910 | ##### | -8.7997 | 1.3719E-18 | 9.6495E-18 |
| ENSG00000119661\|DNAL1 | 2214.9493 | 1.1375 | ##### | 8.7967 | 1.4085E-18 | 9.9043E-18 |
| ENSG00000101158\|NELFCD | 16407.8763 | 1.1742 | ##### | 8.7951 | 1.4287E-18 | 1.0044E-17 |
| ENSG00000225077\|ICMT-DT | 164.6116 | 2.4302 | ##### | 8.7951 | 1.4291E-18 | 1.0045E-17 |
| ENSG00000198208\|RPS6KL1 | 60.8351 | -3.4049 | ##### | -8.7932 | 1.4539E-18 | 1.0216E-17 |
| ENSG00000130479\|MAP1S | 5230.7593 | 1.0994 | ##### | 8.7931 | 1.4550E-18 | 1.0222E-17 |
| ENSG00000099331\|MYO9B | 10172.2277 | 1.0526 | ##### | 8.7912 | 1.4797E-18 | 1.0393E-17 |
| ENSG00000120451\|SNX19 | 3566.3330 | -1.0919 | ##### | -8.7888 | 1.5114E-18 | 1.0613E-17 |
| ENSG00000214717\|ZBED1 | 3789.6414 | -1.0916 | ##### | -8.7863 | 1.5461E-18 | 1.0854E-17 |
| ENSG00000120253\|NUP43 | 6451.6532 | 1.0724 | ##### | 8.7857 | 1.5538E-18 | 1.0906E-17 |
| ENSG00000042753\|AP2S1 | 7946.8771 | -1.0589 | ##### | -8.7855 | 1.5564E-18 | 1.0922E-17 |
| ENSG00000016082\|ISL1 | 1159.0334 | 11.9215 | ##### | 8.7852 | 1.5606E-18 | 1.0948E-17 |
| ENSG00000167880\|EVPL | 1594.6450 | -1.1405 | ##### | -8.7838 | 1.5805E-18 | 1.1085E-17 |
| ENSG00000139044\|B4GALNT3 | 260.2575 | 1.7528 | ##### | 8.7823 | 1.6013E-18 | 1.1228E-17 |
| ENSG00000204899\|MZT1 | 2991.8102 | 1.1327 | ##### | 8.7818 | 1.6088E-18 | 1.1278E-17 |
| ENSG00000124813\|RUNX2 | 371.0367 | 1.3876 | ##### | 8.7812 | 1.6175E-18 | 1.1337E-17 |
| ENSG00000119559\|C19orf25 | 2774.4966 | 1.0872 | ##### | 8.7810 | 1.6200E-18 | 1.1351E-17 |
| ENSG00000081853\|PCDHGA2 | 4982.6089 | ####### | ##### | -8.7760 | 1.6940E-18 | 1.1868E-17 |
| ENSG00000168066\|SF1 | 16818.8659 | 0.9635 | ##### | 8.7741 | 1.7222E-18 | 1.2062E-17 |
| ENSG00000136842\|TMOD1 | 488.2879 | -3.2061 | ##### | -8.7737 | 1.7283E-18 | 1.2102E-17 |
| ENSG00000143373\|ZNF687 | 4574.6131 | 1.0927 | ##### | 8.7732 | 1.7368E-18 | 1.2159E-17 |
| ENSG00000184634\|MED12 | 5061.3829 | 1.0783 | ##### | 8.7707 | 1.7749E-18 | 1.2423E-17 |
| ENSG00000107404\|DVL1 | 13097.9427 | -1.0398 | ##### | -8.7696 | 1.7932E-18 | 1.2548E-17 |
| ENSG00000229847\|EMX2OS | 179.0359 | 9.3672 | ##### | 8.7694 | 1.7958E-18 | 1.2563E-17 |
| ENSG00000092098\|RNF31 | 3042.5395 | 1.3089 | ##### | 8.7681 | 1.8177E-18 | 1.2713E-17 |
| ENSG00000232611 | 113.3601 | 2.2208 | ##### | 8.7675 | 1.8262E-18 | 1.2770E-17 |
| ENSG00000162407\|PLPP3 | 2984.5346 | -1.0558 | ##### | -8.7671 | 1.8326E-18 | 1.2811E-17 |
| ENSG00000196950\|SLC39A10 | 5119.8357 | -1.1023 | ##### | -8.7656 | 1.8585E-18 | 1.2989E-17 |
| ENSG00000167904\|TMEM68 | 1651.7974 | -1.1800 | ##### | -8.7654 | 1.8614E-18 | 1.3007E-17 |
| ENSG00000172244\|C5orf34 | 1661.0134 | 1.1468 | ##### | 8.7630 | 1.9005E-18 | 1.3277E-17 |
| ENSG00000147133\|TAF1 | 4291.7486 | 1.0621 | ##### | 8.7612 | 1.9311E-18 | 1.3487E-17 |
| ENSG00000198914\|POU3F3 | 154.5487 | 2.3658 | ##### | 8.7610 | 1.9359E-18 | 1.3518E-17 |
| ENSG00000126953\|TIMM8A | 1840.2871 | 1.1385 | ##### | 8.7607 | 1.9403E-18 | 1.3546E-17 |
| ENSG00000104936\|DMPK | 4088.7167 | -1.0814 | ##### | -8.7606 | 1.9420E-18 | 1.3554E-17 |
| ENSG00000122574\|WIPF3 | 309.8326 | -1.4664 | ##### | -8.7595 | 1.9618E-18 | 1.3689E-17 |
| ENSG00000186314\|PRELID2 | 348.7981 | -1.4948 | ##### | -8.7582 | 1.9832E-18 | 1.3835E-17 |
| ENSG00000104826\|LHB | 89.3807 | 2.1954 | ##### | 8.7569 | 2.0061E-18 | 1.3992E-17 |
| ENSG00000125968\|ID1 | 15789.5117 | -1.0474 | ##### | -8.7537 | 2.0651E-18 | 1.4400E-17 |

| | | | | | | |
|---|---|---|---|---|---|---|
| ENSG00000105556\|MIER2 | 3040.7891 | 1.1010 | ##### | 8.7531 | 2.0750E-18 | 1.4466E-17 |
| ENSG00000125912\|NCLN | 22523.8574 | 0.5412 | ##### | 8.7510 | 2.1153E-18 | 1.4740E-17 |
| ENSG00000085185\|BCORL1 | 3024.8813 | 1.1062 | ##### | 8.7510 | 2.1153E-18 | 1.4740E-17 |
| ENSG00000127311\|HELB | 700.7251 | 1.4547 | ##### | 8.7501 | 2.1324E-18 | 1.4855E-17 |
| ENSG00000054392\|HHAT | 480.7780 | -1.2299 | ##### | -8.7491 | 2.1500E-18 | 1.4974E-17 |
| ENSG00000110013\|SIAE | 2435.8125 | -1.1129 | ##### | -8.7490 | 2.1534E-18 | 1.4994E-17 |
| ENSG00000233006\|MIR3936HG | 105.9069 | -2.6010 | ##### | -8.7479 | 2.1730E-18 | 1.5127E-17 |
| ENSG00000132612\|VPS4A | 10699.1661 | -0.9591 | ##### | -8.7473 | 2.1850E-18 | 1.5208E-17 |
| ENSG00000266094\|RASSF5 | 336.1213 | -1.6142 | ##### | -8.7456 | 2.2187E-18 | 1.5439E-17 |
| ENSG00000257497\|GLIPR1-AS1 | 149.2318 | 1.8407 | ##### | 8.7443 | 2.2449E-18 | 1.5617E-17 |
| ENSG00000156711\|MAPK13 | 117.1092 | -2.4495 | ##### | -8.7428 | 2.2740E-18 | 1.5816E-17 |
| ENSG00000289357 | 133.1344 | 1.6789 | ##### | 8.7421 | 2.2890E-18 | 1.5916E-17 |
| ENSG00000180318\|ALX1 | 319.5737 | -1.4109 | ##### | -8.7413 | 2.3049E-18 | 1.6023E-17 |
| ENSG00000140718\|FTO | 3144.8527 | -1.0886 | ##### | -8.7400 | 2.3310E-18 | 1.6201E-17 |
| ENSG00000181894\|ZNF329 | 1155.0264 | -1.1897 | ##### | -8.7365 | 2.4045E-18 | 1.6708E-17 |
| ENSG00000139697\|SBNO1 | 12130.2156 | 1.0440 | ##### | 8.7355 | 2.4269E-18 | 1.6859E-17 |
| ENSG00000255650\|FAM222A-AS1 | 45.8510 | 2.8080 | ##### | 8.7346 | 2.4461E-18 | 1.6989E-17 |
| ENSG00000152127\|MGAT5 | 7189.7839 | -1.0526 | ##### | -8.7339 | 2.4593E-18 | 1.7077E-17 |
| ENSG00000179409\|GEMIN4 | 6660.7164 | 0.7626 | ##### | 8.7338 | 2.4622E-18 | 1.7093E-17 |
| ENSG00000166938\|DIS3L | 2140.6633 | -1.0741 | ##### | -8.7317 | 2.5079E-18 | 1.7406E-17 |
| ENSG00000160606\|TLCD1 | 2253.9105 | 1.1368 | ##### | 8.7310 | 2.5252E-18 | 1.7522E-17 |
| ENSG00000112276\|BVES | 804.3790 | -1.2252 | ##### | -8.7285 | 2.5797E-18 | 1.7896E-17 |
| ENSG00000140743\|CDR2 | 2553.7363 | -1.0653 | ##### | -8.7275 | 2.6026E-18 | 1.8051E-17 |
| ENSG00000263513\|FAM72C | 1245.3819 | 1.1526 | ##### | 8.7256 | 2.6476E-18 | 1.8359E-17 |
| ENSG00000082153\|BZW1 | 40674.6956 | 0.7372 | ##### | 8.7256 | 2.6485E-18 | 1.8360E-17 |
| ENSG00000132688\|NES | 500.3876 | -1.3148 | ##### | -8.7245 | 2.6726E-18 | 1.8523E-17 |
| ENSG00000169604\|ANTXR1 | 16069.0909 | -0.7301 | ##### | -8.7208 | 2.7620E-18 | 1.9138E-17 |
| ENSG00000134146\|DPH6 | 380.6689 | -1.4664 | ##### | -8.7202 | 2.7771E-18 | 1.9238E-17 |
| ENSG00000028310\|BRD9 | 2811.9138 | -1.0616 | ##### | -8.7201 | 2.7806E-18 | 1.9258E-17 |
| ENSG00000088305\|DNMT3B | 1153.4320 | -1.2081 | ##### | -8.7173 | 2.8492E-18 | 1.9729E-17 |
| ENSG00000169446\|MMGT1 | 5212.5253 | 1.1050 | ##### | 8.7157 | 2.8908E-18 | 2.0012E-17 |
| ENSG00000166833\|NAV2 | 1380.7436 | -1.3115 | ##### | -8.7155 | 2.8943E-18 | 2.0032E-17 |
| ENSG00000165698\|SPACA9 | 519.8306 | -1.2935 | ##### | -8.7134 | 2.9479E-18 | 2.0398E-17 |
| ENSG00000265784 | 436.2881 | -2.7968 | ##### | -8.7129 | 2.9609E-18 | 2.0479E-17 |
| ENSG00000155026\|RSPH10B | 155.0840 | 1.7197 | ##### | 8.7129 | 2.9610E-18 | 2.0479E-17 |
| ENSG00000133056\|PIK3C2B | 3107.7343 | -1.1324 | ##### | -8.7128 | 2.9650E-18 | 2.0501E-17 |
| ENSG00000151694\|ADAM17 | 8641.9688 | -1.0494 | ##### | -8.7128 | 2.9655E-18 | 2.0501E-17 |
| ENSG00000138623\|SEMA7A | 324.2783 | -1.6162 | ##### | -8.7118 | 2.9910E-18 | 2.0673E-17 |
| ENSG00000197568\|ANKRD13C-DT | 367.8355 | -1.4221 | ##### | -8.7107 | 3.0198E-18 | 2.0867E-17 |
| ENSG00000180198\|RCC1 | 8924.5316 | 0.9451 | ##### | 8.7093 | 3.0584E-18 | 2.1129E-17 |
| ENSG00000144847\|IGSF11 | 75.5731 | -4.8778 | ##### | -8.7092 | 3.0601E-18 | 2.1136E-17 |
| ENSG00000242715\|CCDC169 | 201.3308 | -1.8284 | ##### | -8.7046 | 3.1863E-18 | 2.2002E-17 |
| ENSG00000126247\|CAPNS1 | 26868.9903 | -1.0515 | ##### | -8.7036 | 3.2159E-18 | 2.2202E-17 |
| ENSG00000152223\|EPG5 | 2725.1403 | -1.1175 | ##### | -8.7017 | 3.2683E-18 | 2.2558E-17 |
| ENSG00000162063\|CCNF | 4402.1391 | 1.1012 | ##### | 8.7009 | 3.2940E-18 | 2.2730E-17 |
| ENSG00000121289\|CEP89 | 2685.8969 | -1.1171 | ##### | -8.6985 | 3.3634E-18 | 2.3204E-17 |
| ENSG00000123407\|HOXC12 | 619.2902 | 1.4282 | ##### | 8.6963 | 3.4293E-18 | 2.3652E-17 |
| ENSG00000196187\|TMEM63A | 1907.1830 | -1.1000 | ##### | -8.6937 | 3.5095E-18 | 2.4201E-17 |
| ENSG00000188690\|UROS | 3380.0056 | -1.0442 | ##### | -8.6926 | 3.5437E-18 | 2.4430E-17 |
| ENSG00000230882 | 78.4153 | -2.2157 | ##### | -8.6921 | 3.5575E-18 | 2.4520E-17 |
| ENSG00000095209\|TMEM38B | 2321.5048 | -1.1362 | ##### | -8.6920 | 3.5625E-18 | 2.4548E-17 |
| ENSG00000049759\|NEDD4L | 8075.8749 | -1.0123 | ##### | -8.6902 | 3.6196E-18 | 2.4936E-17 |
| ENSG00000290418\|TPTEP1 | 610.1920 | ####### | ##### | -8.6888 | 3.6634E-18 | 2.5233E-17 |
| ENSG00000126773\|PCNX4 | 6856.2408 | 0.8968 | ##### | 8.6881 | 3.6843E-18 | 2.5370E-17 |
| ENSG00000086200\|IPO11 | 4362.2975 | 1.0581 | ##### | 8.6851 | 3.7844E-18 | 2.6054E-17 |
| ENSG00000176532\|PRR15 | 130.5165 | -3.9615 | ##### | -8.6803 | 3.9482E-18 | 2.7175E-17 |
| ENSG00000186399\|GOLGA8R | 62.0510 | -2.2356 | ##### | -8.6795 | 3.9737E-18 | 2.7344E-17 |
| ENSG00000288953 | 124.8786 | -2.1739 | ##### | -8.6787 | 4.0027E-18 | 2.7537E-17 |
| ENSG00000167703\|SLC43A2 | 902.4206 | -1.2415 | ##### | -8.6760 | 4.1004E-18 | 2.8203E-17 |
| ENSG00000289295 | 102.5282 | 2.0193 | ##### | 8.6727 | 4.2197E-18 | 2.9017E-17 |
| ENSG00000111671\|SPSB2 | 2740.2191 | -1.0877 | ##### | -8.6698 | 4.3295E-18 | 2.9766E-17 |
| ENSG00000173402\|DAG1 | 18806.3729 | -1.0267 | ##### | -8.6686 | 4.3758E-18 | 3.0077E-17 |
| ENSG00000125952\|MAX | 6629.4553 | 1.0552 | ##### | 8.6649 | 4.5185E-18 | 3.1051E-17 |

| | | | | | | |
|---|---|---|---|---|---|---|
| ENSG00000164919\|COX6C | 8971.0925 | -1.0405 | ##### | -8.6628 | 4.6029E-18 | 3.1623E-17 |
| ENSG00000138430\|OLA1 | 9662.3516 | 1.1310 | ##### | 8.6620 | 4.6347E-18 | 3.1834E-17 |
| ENSG00000065054\|NHERF2 | 1604.2406 | -1.1053 | ##### | -8.6589 | 4.7644E-18 | 3.2717E-17 |
| ENSG00000152192\|POU4F1 | 206.8063 | 4.8839 | ##### | 8.6585 | 4.7812E-18 | 3.2826E-17 |
| ENSG00000137563\|GGH | 7905.5336 | -1.0780 | ##### | -8.6581 | 4.7965E-18 | 3.2923E-17 |
| ENSG00000170456\|DENND5B | 10750.7624 | 1.0433 | ##### | 8.6581 | 4.7979E-18 | 3.2925E-17 |
| ENSG00000064195\|DLX3 | 172.5506 | 1.9924 | ##### | 8.6569 | 4.8458E-18 | 3.3246E-17 |
| ENSG00000005961\|ITGA2B | 275.5195 | 1.7227 | ##### | 8.6546 | 4.9460E-18 | 3.3926E-17 |
| ENSG00000116990\|MYCL | 358.0904 | -1.4772 | ##### | -8.6540 | 4.9723E-18 | 3.4098E-17 |
| ENSG00000136114\|THSD1 | 1032.3141 | -1.2095 | ##### | -8.6527 | 5.0307E-18 | 3.4491E-17 |
| ENSG00000104889\|RNASEH2A | 3640.2255 | 1.0868 | ##### | 8.6505 | 5.1287E-18 | 3.5154E-17 |
| ENSG00000226889\|ATF6-DT | 494.3274 | 1.2402 | ##### | 8.6491 | 5.1901E-18 | 3.5568E-17 |
| ENSG00000205730\|ITPRIPL2 | 9119.9071 | -1.0341 | ##### | -8.6397 | 5.6355E-18 | 3.8611E-17 |
| ENSG00000147251\|DOCK11 | 536.8643 | 1.3063 | ##### | 8.6386 | 5.6905E-18 | 3.8978E-17 |
| ENSG00000133134\|BEX2 | 326.1664 | 1.5447 | ##### | 8.6378 | 5.7323E-18 | 3.9256E-17 |
| ENSG00000115875\|SRSF7 | 12481.3878 | 1.0549 | ##### | 8.6377 | 5.7351E-18 | 3.9258E-17 |
| ENSG00000172058\|SERF1A | 543.2969 | -1.4481 | ##### | -8.6377 | 5.7353E-18 | 3.9258E-17 |
| ENSG00000162999\|DUSP19 | 664.1125 | -1.1941 | ##### | -8.6353 | 5.8594E-18 | 4.0099E-17 |
| ENSG00000185630\|PBX1 | 1556.0158 | -1.1099 | ##### | -8.6316 | 6.0516E-18 | 4.1405E-17 |
| ENSG00000176490\|DIRAS1 | 1492.4785 | 1.1011 | ##### | 8.6307 | 6.0975E-18 | 4.1709E-17 |
| ENSG00000187609\|EXD3 | 369.8951 | -1.3688 | ##### | -8.6303 | 6.1171E-18 | 4.1833E-17 |
| ENSG00000135919\|SERPINE2 | 9930.0627 | -0.9995 | ##### | -8.6290 | 6.1888E-18 | 4.2314E-17 |
| ENSG00000103044\|HAS3 | 4306.8318 | -1.1902 | ##### | -8.6289 | 6.1966E-18 | 4.2358E-17 |
| ENSG00000242960\|FTH1P23 | 34.0302 | -2.8549 | ##### | -8.6288 | 6.2003E-18 | 4.2364E-17 |
| ENSG00000125351\|UPF3B | 2902.6794 | 0.9264 | ##### | 8.6288 | 6.2000E-18 | 4.2364E-17 |
| ENSG00000134871\|COL4A2 | 3918.4619 | -1.0585 | ##### | -8.6277 | 6.2616E-18 | 4.2773E-17 |
| ENSG00000164168\|TMEM184C | 2315.8037 | -1.0770 | ##### | -8.6259 | 6.3597E-18 | 4.3433E-17 |
| ENSG00000141456\|PELP1 | 10933.1334 | 0.7889 | ##### | 8.6242 | 6.4522E-18 | 4.4055E-17 |
| ENSG00000102172\|SMS | 11106.8180 | 0.6690 | ##### | 8.6216 | 6.6036E-18 | 4.5078E-17 |
| ENSG00000119537\|KDSR | 4914.9359 | 0.7547 | ##### | 8.6204 | 6.6737E-18 | 4.5546E-17 |
| ENSG00000142046\|TMEM91 | 120.4875 | -2.2901 | ##### | -8.6202 | 6.6850E-18 | 4.5613E-17 |
| ENSG00000223751 | 254.7156 | 9.0256 | ##### | 8.6198 | 6.7044E-18 | 4.5735E-17 |
| ENSG00000226476\|LINC01748 | 166.1406 | -2.6636 | ##### | -8.6157 | 6.9516E-18 | 4.7410E-17 |
| ENSG00000115884\|SDC1 | 15934.9436 | -0.9414 | ##### | -8.6154 | 6.9700E-18 | 4.7525E-17 |
| ENSG00000116525\|TRIM62 | 693.0750 | 1.2955 | ##### | 8.6091 | 7.3617E-18 | 5.0184E-17 |
| ENSG00000147162\|OGT | 10902.3906 | 1.0504 | ##### | 8.6051 | 7.6256E-18 | 5.1971E-17 |
| ENSG00000104213\|PDGFRL | 474.8778 | 1.7041 | ##### | 8.6042 | 7.6872E-18 | 5.2379E-17 |
| ENSG00000143603\|KCNN3 | 1027.7649 | 8.6179 | ##### | 8.6012 | 7.8857E-18 | 5.3720E-17 |
| ENSG00000262539 | 36.3277 | 3.2459 | ##### | 8.6009 | 7.9066E-18 | 5.3850E-17 |
| ENSG00000189152\|GRAPL | 46.1406 | -7.7664 | ##### | -8.6009 | 7.9092E-18 | 5.3855E-17 |
| ENSG00000166166\|TRMT61A | 3539.0252 | 1.0935 | ##### | 8.5993 | 8.0175E-18 | 5.4580E-17 |
| ENSG00000234545\|FAM133B | 1584.5387 | 0.9946 | ##### | 8.5990 | 8.0422E-18 | 5.4736E-17 |
| ENSG00000118705\|RPN2 | 44492.3723 | 0.7402 | ##### | 8.5984 | 8.0853E-18 | 5.5017E-17 |
| ENSG00000153406\|NMRAL1 | 2477.7422 | -1.1012 | ##### | -8.5908 | 8.6374E-18 | 5.8760E-17 |
| ENSG00000175352\|NRIP3 | 428.8827 | 1.4962 | ##### | 8.5903 | 8.6757E-18 | 5.9007E-17 |
| ENSG00000162804\|SNED1 | 296.5741 | -1.4961 | ##### | -8.5867 | 8.9512E-18 | 6.0867E-17 |
| ENSG00000121904\|CSMD2 | 314.5956 | 9.0083 | ##### | 8.5859 | 9.0163E-18 | 6.1296E-17 |
| ENSG00000159335\|PTMS | 47186.8055 | 1.0345 | ##### | 8.5842 | 9.1509E-18 | 6.2197E-17 |
| ENSG00000145945\|FAM50B | 1254.1790 | 0.9721 | ##### | 8.5831 | 9.2363E-18 | 6.2763E-17 |
| ENSG00000164938\|TP53INP1 | 373.3052 | -1.9680 | ##### | -8.5830 | 9.2461E-18 | 6.2815E-17 |
| ENSG00000182901\|RGS7 | 57.8863 | 2.9574 | ##### | 8.5811 | 9.3930E-18 | 6.3798E-17 |
| ENSG00000179918\|SEPHS2 | 5162.4685 | -1.0087 | ##### | -8.5775 | 9.6973E-18 | 6.5851E-17 |
| ENSG00000115946\|PNO1 | 9214.7314 | 0.7085 | ##### | 8.5760 | 9.8206E-18 | 6.6672E-17 |
| ENSG00000154654\|NCAM2 | 543.6226 | -1.4105 | ##### | -8.5755 | 9.8625E-18 | 6.6942E-17 |
| ENSG00000064300\|NGFR | 161.1101 | 2.4744 | ##### | 8.5751 | 9.8993E-18 | 6.7177E-17 |
| ENSG00000143653\|SCCPDH | 3407.5144 | -1.0320 | ##### | -8.5632 | 1.0978E-17 | 7.4482E-17 |
| ENSG00000143924\|EML4 | 15740.4439 | 1.0154 | ##### | 8.5629 | 1.1005E-17 | 7.4647E-17 |
| ENSG00000089335\|ZNF302 | 1406.3653 | -1.1730 | ##### | -8.5601 | 1.1273E-17 | 7.6447E-17 |
| ENSG00000176170\|SPHK1 | 3479.2604 | 1.1493 | ##### | 8.5596 | 1.1321E-17 | 7.6757E-17 |
| ENSG00000105122\|RASAL3 | 60.4722 | 7.7761 | ##### | 8.5593 | 1.1354E-17 | 7.6960E-17 |
| ENSG00000152904\|GGPS1 | 1845.3116 | -1.1040 | ##### | -8.5589 | 1.1398E-17 | 7.7240E-17 |
| ENSG00000239670 | 36.3249 | 7.6135 | ##### | 8.5588 | 1.1401E-17 | 7.7244E-17 |
| ENSG00000164687\|FABP5 | 6487.0690 | -1.0795 | ##### | -8.5585 | 1.1438E-17 | 7.7479E-17 |

| | | | | | | |
|---|---|---|---|---|---|---|
| ENSG00000061676\|NCKAP1 | 30203.1658 | -0.7886 | ##### | -8.5580 | 1.1484E-17 | 7.7768E-17 |
| ENSG00000113638\|TTC33 | 2684.5391 | 0.9379 | ##### | 8.5579 | 1.1498E-17 | 7.7850E-17 |
| ENSG00000287200 | 92.9143 | -2.0503 | ##### | -8.5568 | 1.1600E-17 | 7.8520E-17 |
| ENSG00000157349\|DDX19B | 2952.9254 | -1.0688 | ##### | -8.5565 | 1.1632E-17 | 7.8719E-17 |
| ENSG00000253661\|ZFHX4-AS1 | 160.6637 | -9.7714 | ##### | -8.5564 | 1.1649E-17 | 7.8817E-17 |
| ENSG00000158941\|CCAR2 | 16902.9729 | 1.0024 | ##### | 8.5538 | 1.1910E-17 | 8.0564E-17 |
| ENSG00000254206\|NPIPB11 | 412.8691 | 1.2836 | ##### | 8.5489 | 1.2422E-17 | 8.4007E-17 |
| ENSG00000235703\|EOLA2-DT | 97.9074 | 2.0653 | ##### | 8.5447 | 1.2884E-17 | 8.7117E-17 |
| ENSG00000144455\|SUMF1 | 1354.4050 | -1.0861 | ##### | -8.5441 | 1.2952E-17 | 8.7553E-17 |
| ENSG00000233799 | 35.3077 | 7.7171 | ##### | 8.5425 | 1.3130E-17 | 8.8737E-17 |
| ENSG00000065970\|FOXJ2 | 4564.9468 | 0.7857 | ##### | 8.5421 | 1.3176E-17 | 8.9026E-17 |
| ENSG00000198695\|MT-ND6 | 54214.5146 | 0.9241 | ##### | 8.5420 | 1.3197E-17 | 8.9147E-17 |
| ENSG00000082068\|WDR70 | 3932.2614 | 1.0464 | ##### | 8.5410 | 1.3309E-17 | 8.9883E-17 |
| ENSG00000145592\|RPL37 | 44480.5448 | 1.0946 | ##### | 8.5395 | 1.3482E-17 | 9.1032E-17 |
| ENSG00000124635\|H2BC11 | 553.3764 | 1.3812 | ##### | 8.5391 | 1.3532E-17 | 9.1347E-17 |
| ENSG00000278921\|EPB41L4A-DT | 48.7097 | -3.5623 | ##### | -8.5387 | 1.3572E-17 | 9.1600E-17 |
| ENSG00000173264\|GPR137 | 3156.9839 | 1.0852 | ##### | 8.5382 | 1.3636E-17 | 9.2007E-17 |
| ENSG00000181218\|H2AC25 | 1168.7287 | -1.1626 | ##### | -8.5366 | 1.3821E-17 | 9.3239E-17 |
| ENSG00000196267\|ZNF836 | 165.7429 | -1.5558 | ##### | -8.5362 | 1.3876E-17 | 9.3587E-17 |
| ENSG00000011485\|PPP5C | 8315.1549 | -0.9557 | ##### | -8.5352 | 1.3997E-17 | 9.4382E-17 |
| ENSG00000228549 | 119.2119 | 2.0017 | ##### | 8.5344 | 1.4094E-17 | 9.5012E-17 |
| ENSG00000214063\|TSPAN4 | 4515.0745 | -1.0125 | ##### | -8.5325 | 1.4321E-17 | 9.6522E-17 |
| ENSG00000237522\|NONOP2 | 87.6014 | 2.2015 | ##### | 8.5317 | 1.4419E-17 | 9.7162E-17 |
| ENSG00000226742\|HSBP1L1 | 828.3361 | -1.1413 | ##### | -8.5314 | 1.4458E-17 | 9.7405E-17 |
| ENSG00000125772\|GPCPD1 | 852.8334 | -1.1766 | ##### | -8.5303 | 1.4598E-17 | 9.8325E-17 |
| ENSG00000288748 | 42.6739 | 7.2512 | ##### | 8.5299 | 1.4649E-17 | 9.8644E-17 |
| ENSG00000108219\|TSPAN14 | 4245.8120 | -1.0511 | ##### | -8.5294 | 1.4711E-17 | 9.9042E-17 |
| ENSG00000234964\|FABP5P7 | 630.5297 | -1.2377 | ##### | -8.5288 | 1.4783E-17 | 9.9502E-17 |
| ENSG00000163781\|TOPBP1 | 9685.3980 | 1.0324 | ##### | 8.5271 | 1.5007E-17 | 1.0099E-16 |
| ENSG00000129675\|ARHGEF6 | 158.8556 | -2.4801 | ##### | -8.5261 | 1.5133E-17 | 1.0181E-16 |
| ENSG00000087085\|ACHE | 432.2184 | -1.3196 | ##### | -8.5250 | 1.5283E-17 | 1.0280E-16 |
| ENSG00000183833\|CFAP91 | 128.0601 | -1.9131 | ##### | -8.5222 | 1.5660E-17 | 1.0531E-16 |
| ENSG00000072071\|ADGRL1 | 6787.8976 | 1.0160 | ##### | 8.5163 | 1.6475E-17 | 1.1077E-16 |
| ENSG00000112379\|ARFGEF3 | 581.9590 | 1.3109 | ##### | 8.5155 | 1.6581E-17 | 1.1146E-16 |
| ENSG00000286638 | 69.2378 | 2.9883 | ##### | 8.5155 | 1.6588E-17 | 1.1147E-16 |
| ENSG00000136826\|KLF4 | 3002.5514 | -1.0885 | ##### | -8.5095 | 1.7473E-17 | 1.1740E-16 |
| ENSG00000165028\|NIPSNAP3B | 91.7298 | -2.1823 | ##### | -8.5089 | 1.7556E-17 | 1.1793E-16 |
| ENSG00000050165\|DKK3 | 1072.8664 | -1.1674 | ##### | -8.5062 | 1.7966E-17 | 1.2066E-16 |
| ENSG00000183066\|WBP2NL | 21.6556 | -4.3352 | ##### | -8.5058 | 1.8040E-17 | 1.2112E-16 |
| ENSG00000148935\|GAS2 | 62.5655 | -2.6113 | ##### | -8.5008 | 1.8832E-17 | 1.2641E-16 |
| ENSG00000189266\|PNRC2 | 5313.3705 | 0.9903 | ##### | 8.4995 | 1.9043E-17 | 1.2780E-16 |
| ENSG00000161692\|DBF4B | 3062.1080 | 1.0820 | ##### | 8.4989 | 1.9146E-17 | 1.2847E-16 |
| ENSG00000149294\|NCAM1 | 1207.5795 | -7.3670 | ##### | -8.4980 | 1.9293E-17 | 1.2942E-16 |
| ENSG00000214999 | 150.7972 | 2.2700 | ##### | 8.4979 | 1.9312E-17 | 1.2952E-16 |
| ENSG00000247095\|MIR210HG | 62.6558 | -2.3349 | ##### | -8.4971 | 1.9432E-17 | 1.3030E-16 |
| ENSG00000258818\|RNASE4 | 583.6190 | -1.3134 | ##### | -8.4967 | 1.9507E-17 | 1.3077E-16 |
| ENSG00000119596\|YLPM1 | 5942.4172 | 1.0214 | ##### | 8.4947 | 1.9843E-17 | 1.3299E-16 |
| ENSG00000123240\|OPTN | 4373.6148 | -1.0336 | ##### | -8.4938 | 2.0006E-17 | 1.3405E-16 |
| ENSG00000106344\|RBM28 | 7934.3085 | 0.9613 | ##### | 8.4935 | 2.0050E-17 | 1.3432E-16 |
| ENSG00000156976\|EIF4A2 | 11077.0392 | -1.0019 | ##### | -8.4923 | 2.0266E-17 | 1.3573E-16 |
| ENSG00000186891\|TNFRSF18 | 62.0549 | 2.3741 | ##### | 8.4918 | 2.0338E-17 | 1.3618E-16 |
| ENSG00000129187\|DCTD | 5589.2970 | -1.0831 | ##### | -8.4907 | 2.0544E-17 | 1.3754E-16 |
| ENSG00000146733\|PSPH | 3126.3095 | -0.8866 | ##### | -8.4896 | 2.0741E-17 | 1.3883E-16 |
| ENSG00000087053\|MTMR2 | 5296.9089 | -1.0622 | ##### | -8.4886 | 2.0908E-17 | 1.3991E-16 |
| ENSG00000156172\|CFAP418 | 900.1373 | -1.1340 | ##### | -8.4860 | 2.1382E-17 | 1.4305E-16 |
| ENSG00000025772\|TOMM34 | 5822.6058 | 1.0272 | ##### | 8.4846 | 2.1650E-17 | 1.4481E-16 |
| ENSG00000138686\|BBS7 | 1886.8381 | 1.0850 | ##### | 8.4834 | 2.1864E-17 | 1.4621E-16 |
| ENSG00000213390\|ARHGAP19 | 2491.1106 | 1.0755 | ##### | 8.4830 | 2.1946E-17 | 1.4672E-16 |
| ENSG00000100731\|PCNX1 | 7271.2848 | 1.0044 | ##### | 8.4803 | 2.2454E-17 | 1.5009E-16 |
| ENSG00000143870\|PDIA6 | 52516.7399 | 0.5354 | ##### | 8.4772 | 2.3070E-17 | 1.5417E-16 |
| ENSG00000173171\|MTX1 | 6039.9294 | 1.0183 | ##### | 8.4771 | 2.3085E-17 | 1.5423E-16 |
| ENSG00000267751\|BSG-AS1 | 359.5935 | 1.3932 | ##### | 8.4757 | 2.3372E-17 | 1.5612E-16 |
| ENSG00000109686\|SH3D19 | 3916.5723 | -1.0107 | ##### | -8.4734 | 2.3835E-17 | 1.5918E-16 |

| | | | | | | |
|---|---|---|---|---|---|---|
| ENSG00000147231\|RADX | 484.3014 | 1.2855 | ##### | 8.4725 | 2.4018E-17 | 1.6036E-16 |
| ENSG00000163378\|EOGT | 1466.2210 | -1.1366 | ##### | -8.4724 | 2.4039E-17 | 1.6046E-16 |
| ENSG00000279406 | 112.1157 | -1.9299 | ##### | -8.4707 | 2.4402E-17 | 1.6285E-16 |
| ENSG00000233608\|TWIST2 | 793.9803 | 0.8327 | ##### | 8.4691 | 2.4722E-17 | 1.6495E-16 |
| ENSG00000197208\|SLC22A4 | 392.4487 | -1.3267 | ##### | -8.4675 | 2.5068E-17 | 1.6722E-16 |
| ENSG00000141480\|ARRB2 | 3310.2148 | 1.0696 | ##### | 8.4674 | 2.5091E-17 | 1.6734E-16 |
| ENSG00000162600\|OMA1 | 1026.4694 | -1.1772 | ##### | -8.4673 | 2.5108E-17 | 1.6742E-16 |
| ENSG00000259985 | 376.2470 | 1.4231 | ##### | 8.4672 | 2.5146E-17 | 1.6763E-16 |
| ENSG00000259031 | 38.7156 | 3.6330 | ##### | 8.4626 | 2.6152E-17 | 1.7430E-16 |
| ENSG00000096384\|HSP90AB1 | ########## | 0.8497 | ##### | 8.4621 | 2.6260E-17 | 1.7498E-16 |
| ENSG00000102858\|MGRN1 | 4719.7722 | -1.0569 | ##### | -8.4619 | 2.6300E-17 | 1.7520E-16 |
| ENSG00000108669\|CYTH1 | 4226.9458 | 1.0115 | ##### | 8.4614 | 2.6423E-17 | 1.7599E-16 |
| ENSG00000214367\|HAUS3 | 1574.0385 | 1.1093 | ##### | 8.4613 | 2.6444E-17 | 1.7609E-16 |
| ENSG00000155367\|PPM1J | 267.5612 | 1.7116 | ##### | 8.4599 | 2.6761E-17 | 1.7816E-16 |
| ENSG00000105443\|CYTH2 | 4853.8534 | -1.0156 | ##### | -8.4583 | 2.7139E-17 | 1.8063E-16 |
| ENSG00000234616\|JRK | 3341.7758 | 1.0522 | ##### | 8.4533 | 2.8321E-17 | 1.8846E-16 |
| ENSG00000165309\|ARMC3 | 84.0795 | -2.3034 | ##### | -8.4501 | 2.9115E-17 | 1.9370E-16 |
| ENSG00000138622\|HCN4 | 155.7673 | -2.2709 | ##### | -8.4497 | 2.9205E-17 | 1.9426E-16 |
| ENSG00000121931\|LRIF1 | 3312.4858 | -0.9326 | ##### | -8.4486 | 2.9493E-17 | 1.9613E-16 |
| ENSG00000155330\|C16orf87 | 949.8554 | -1.1572 | ##### | -8.4484 | 2.9534E-17 | 1.9636E-16 |
| ENSG00000026297\|RNASET2 | 821.3683 | -1.3159 | ##### | -8.4479 | 2.9668E-17 | 1.9720E-16 |
| ENSG00000161217\|PCYT1A | 6790.2982 | -1.0172 | ##### | -8.4438 | 3.0727E-17 | 2.0419E-16 |
| ENSG00000164620\|RELL2 | 817.8298 | -1.2479 | ##### | -8.4434 | 3.0827E-17 | 2.0482E-16 |
| ENSG00000212123\|PRR22 | 377.0287 | 1.3401 | ##### | 8.4405 | 3.1599E-17 | 2.0990E-16 |
| ENSG00000178951\|ZBTB7A | 7761.0443 | 1.0030 | ##### | 8.4399 | 3.1770E-17 | 2.1099E-16 |
| ENSG00000279619 | 138.3145 | 9.0774 | ##### | 8.4396 | 3.1834E-17 | 2.1137E-16 |
| ENSG00000165704\|HPRT1 | 4773.0341 | 1.0486 | ##### | 8.4389 | 3.2043E-17 | 2.1270E-16 |
| ENSG00000161328\|LRRC56 | 199.5465 | 1.5562 | ##### | 8.4382 | 3.2219E-17 | 2.1383E-16 |
| ENSG00000168894\|RNF181 | 3639.7568 | 0.5904 | ##### | 8.4354 | 3.3019E-17 | 2.1909E-16 |
| ENSG00000131386\|GALNT15 | 153.1997 | 9.0326 | ##### | 8.4333 | 3.3600E-17 | 2.2289E-16 |
| ENSG00000103512\|NOMO1 | 16013.1850 | -0.9550 | ##### | -8.4297 | 3.4667E-17 | 2.2992E-16 |
| ENSG00000164096\|C4orf3 | 3069.4353 | -1.0941 | ##### | -8.4296 | 3.4692E-17 | 2.3003E-16 |
| ENSG00000150281\|CTF1 | 133.2775 | -2.9135 | ##### | -8.4291 | 3.4839E-17 | 2.3096E-16 |
| ENSG00000279232 | 694.0557 | 1.2581 | ##### | 8.4271 | 3.5441E-17 | 2.3489E-16 |
| ENSG00000213412\|HNRNPA1P33 | 62.9430 | -2.9293 | ##### | -8.4262 | 3.5697E-17 | 2.3654E-16 |
| ENSG00000113716\|HMGXB3 | 11621.6838 | -0.9997 | ##### | -8.4245 | 3.6239E-17 | 2.4008E-16 |
| ENSG00000272163 | 105.3251 | -1.7512 | ##### | -8.4243 | 3.6294E-17 | 2.4039E-16 |
| ENSG00000260920 | 294.5068 | -1.4638 | ##### | -8.4217 | 3.7116E-17 | 2.4578E-16 |
| ENSG00000083520\|DIS3 | 4843.0447 | 0.9886 | ##### | 8.4207 | 3.7426E-17 | 2.4778E-16 |
| ENSG00000188266\|HYKK | 213.0655 | -1.6675 | ##### | -8.4191 | 3.7932E-17 | 2.5107E-16 |
| ENSG00000100526\|CDKN3 | 2713.2613 | 1.0538 | ##### | 8.4176 | 3.8443E-17 | 2.5440E-16 |
| ENSG00000185808\|PIGP | 902.5652 | -1.2383 | ##### | -8.4175 | 3.8476E-17 | 2.5456E-16 |
| ENSG00000163719\|MTMR14 | 2695.3587 | 1.0098 | ##### | 8.4163 | 3.8871E-17 | 2.5712E-16 |
| ENSG00000186298\|PPP1CC | 18065.8755 | 0.6015 | ##### | 8.4138 | 3.9692E-17 | 2.6245E-16 |
| ENSG00000223509\|WHAMMP1 | 60.6023 | -4.1132 | ##### | -8.4138 | 3.9694E-17 | 2.6245E-16 |
| ENSG00000174173\|TRMT10C | 3720.7858 | 1.0819 | ##### | 8.4124 | 4.0177E-17 | 2.6558E-16 |
| ENSG00000104899\|AMH | 663.8937 | -1.2536 | ##### | -8.4117 | 4.0414E-17 | 2.6709E-16 |
| ENSG00000151240\|DIP2C | 807.7931 | -1.1380 | ##### | -8.4114 | 4.0499E-17 | 2.6759E-16 |
| ENSG00000168743\|NPNT | 2106.3290 | ####### | ##### | -8.4102 | 4.0947E-17 | 2.7049E-16 |
| ENSG00000233757\|ZNF892 | 339.5811 | 1.7551 | ##### | 8.4098 | 4.1085E-17 | 2.7134E-16 |
| ENSG00000154001\|PPP2R5E | 9950.6427 | 1.0384 | ##### | 8.4094 | 4.1203E-17 | 2.7206E-16 |
| ENSG00000157077\|ZFYVE9 | 1798.4457 | -1.0891 | ##### | -8.4083 | 4.1596E-17 | 2.7459E-16 |
| ENSG00000151632\|AKR1C2 | ########## | ####### | ##### | -8.4079 | 4.1738E-17 | 2.7547E-16 |
| ENSG00000259345 | 54.7593 | 3.0575 | ##### | 8.4070 | 4.2058E-17 | 2.7752E-16 |
| ENSG00000160207\|HSF2BP | 204.2120 | 1.6045 | ##### | 8.4065 | 4.2244E-17 | 2.7869E-16 |
| ENSG00000177943\|MAMDC4 | 1317.2711 | -1.1155 | ##### | -8.4058 | 4.2513E-17 | 2.8040E-16 |
| ENSG00000088682\|COQ9 | 3728.1416 | -0.9752 | ##### | -8.4037 | 4.3265E-17 | 2.8530E-16 |
| ENSG00000170085\|SIMC1 | 904.4267 | 1.1394 | ##### | 8.4023 | 4.3774E-17 | 2.8859E-16 |
| ENSG00000103550\|KNOP1 | 6930.3516 | -1.0028 | ##### | -8.4014 | 4.4114E-17 | 2.9075E-16 |
| ENSG00000108439\|PNPO | 4149.3930 | -1.0100 | ##### | -8.4014 | 4.4121E-17 | 2.9075E-16 |
| ENSG00000101940\|WDR13 | 2821.2182 | -1.0537 | ##### | -8.4005 | 4.4458E-17 | 2.9290E-16 |
| ENSG00000228049\|POLR2J2 | 251.6888 | 1.5403 | ##### | 8.3979 | 4.5435E-17 | 2.9927E-16 |
| ENSG00000285155 | 133.2898 | -9.5025 | ##### | -8.3972 | 4.5732E-17 | 3.0116E-16 |

| | | | | | | |
|---|---|---|---|---|---|---|
| ENSG00000164663\|USP49 | 628.2063 | -1.2588 | ##### | -8.3969 | 4.5837E-17 | 3.0179E-16 |
| ENSG00000114529\|C3orf52 | 980.8761 | 1.2057 | ##### | 8.3967 | 4.5918E-17 | 3.0226E-16 |
| ENSG00000116586\|LAMTOR2 | 3913.8349 | 1.0502 | ##### | 8.3963 | 4.6070E-17 | 3.0319E-16 |
| ENSG00000272143\|FGF14-AS2 | 78.6178 | -1.9814 | ##### | -8.3961 | 4.6154E-17 | 3.0368E-16 |
| ENSG00000118965\|WDR35 | 5454.6644 | 1.0163 | ##### | 8.3958 | 4.6262E-17 | 3.0432E-16 |
| ENSG00000107537\|PHYH | 1862.9048 | -1.0570 | ##### | -8.3958 | 4.6272E-17 | 3.0432E-16 |
| ENSG00000170390\|DCLK2 | 825.5216 | -1.0197 | ##### | -8.3956 | 4.6348E-17 | 3.0475E-16 |
| ENSG00000166439\|RNF169 | 3661.6542 | 0.9046 | ##### | 8.3955 | 4.6411E-17 | 3.0510E-16 |
| ENSG00000116761\|CTH | 798.6834 | -1.1401 | ##### | -8.3952 | 4.6496E-17 | 3.0559E-16 |
| ENSG00000147536\|GINS4 | 4992.0344 | 1.1272 | ##### | 8.3942 | 4.6916E-17 | 3.0829E-16 |
| ENSG00000175467\|SART1 | 7751.9166 | 0.9290 | ##### | 8.3910 | 4.8189E-17 | 3.1658E-16 |
| ENSG00000115421\|PAPOLG | 2084.6655 | 1.1028 | ##### | 8.3898 | 4.8677E-17 | 3.1971E-16 |
| ENSG00000101413\|RPRD1B | 5117.1192 | 0.9777 | ##### | 8.3881 | 4.9390E-17 | 3.2433E-16 |
| ENSG00000187790\|FANCM | 1536.6836 | 1.1054 | ##### | 8.3870 | 4.9883E-17 | 3.2749E-16 |
| ENSG00000064225\|ST3GAL6 | 249.6145 | 1.5049 | ##### | 8.3853 | 5.0585E-17 | 3.3203E-16 |
| ENSG00000013016\|EHD3 | 2256.9928 | 1.0294 | ##### | 8.3828 | 5.1682E-17 | 3.3915E-16 |
| ENSG00000112773\|TENT5A | 1824.4881 | 1.0945 | ##### | 8.3825 | 5.1814E-17 | 3.3994E-16 |
| ENSG00000285437\|POLR2J3 | 330.3169 | 1.9991 | ##### | 8.3802 | 5.2840E-17 | 3.4660E-16 |
| ENSG00000143891\|GALM | 1025.4124 | -1.1240 | ##### | -8.3799 | 5.2986E-17 | 3.4748E-16 |
| ENSG00000119705\|SLIRP | 3919.5005 | 1.0414 | ##### | 8.3790 | 5.3372E-17 | 3.4994E-16 |
| ENSG00000032444\|PNPLA6 | 11656.6875 | 0.7517 | ##### | 8.3783 | 5.3706E-17 | 3.5205E-16 |
| ENSG00000188897 | 55.0896 | -2.5392 | ##### | -8.3781 | 5.3805E-17 | 3.5262E-16 |
| ENSG00000175193\|PARL | 2490.4110 | -1.1589 | ##### | -8.3744 | 5.5483E-17 | 3.6354E-16 |
| ENSG00000108106\|UBE2S | 30517.2928 | 1.0017 | ##### | 8.3712 | 5.7047E-17 | 3.7370E-16 |
| ENSG00000138658\|ZGRF1 | 1012.9744 | 1.3108 | ##### | 8.3711 | 5.7068E-17 | 3.7376E-16 |
| ENSG00000174827\|PDZK1 | 105.2872 | -1.8111 | ##### | -8.3699 | 5.7657E-17 | 3.7753E-16 |
| ENSG00000102401\|ARMCX3 | 4065.0440 | 0.9428 | ##### | 8.3683 | 5.8437E-17 | 3.8256E-16 |
| ENSG00000184990\|SIVA1 | 4163.0650 | 1.0328 | ##### | 8.3683 | 5.8458E-17 | 3.8261E-16 |
| ENSG00000104960\|PTOV1 | 13826.8052 | -1.0229 | ##### | -8.3683 | 5.8471E-17 | 3.8261E-16 |
| ENSG00000237310\|LINC03011 | 83.5007 | -2.2606 | ##### | -8.3666 | 5.9299E-17 | 3.8795E-16 |
| ENSG00000147316\|MCPH1 | 1633.0045 | 1.0803 | ##### | 8.3665 | 5.9331E-17 | 3.8807E-16 |
| ENSG00000166788\|SAAL1 | 1531.7110 | 1.1260 | ##### | 8.3649 | 6.0178E-17 | 3.9352E-16 |
| ENSG00000140297\|GCNT3 | 9824.2390 | ####### | ##### | -8.3642 | 6.0538E-17 | 3.9579E-16 |
| ENSG00000166974\|MAPRE2 | 2363.0136 | -1.0379 | ##### | -8.3622 | 6.1577E-17 | 4.0250E-16 |
| ENSG00000205693\|MANSC4 | 46.6224 | 4.2646 | ##### | 8.3618 | 6.1791E-17 | 4.0381E-16 |
| ENSG00000123191\|ATP7B | 1149.0503 | -1.1328 | ##### | -8.3611 | 6.2160E-17 | 4.0604E-16 |
| ENSG00000170142\|UBE2E1 | 4934.5257 | -1.0275 | ##### | -8.3611 | 6.2148E-17 | 4.0604E-16 |
| ENSG00000108370\|RGS9 | 416.1507 | -1.4610 | ##### | -8.3604 | 6.2513E-17 | 4.0825E-16 |
| ENSG00000204962\|PCDHA8 | 70.8532 | -8.5614 | ##### | -8.3586 | 6.3467E-17 | 4.1440E-16 |
| ENSG00000018610\|STEEP1 | 3245.7575 | 1.0319 | ##### | 8.3581 | 6.3711E-17 | 4.1590E-16 |
| ENSG00000114767\|RRP9 | 6724.9688 | 1.0070 | ##### | 8.3559 | 6.4915E-17 | 4.2366E-16 |
| ENSG00000160352\|ZNF714 | 517.6120 | 4.3880 | ##### | 8.3541 | 6.5955E-17 | 4.3036E-16 |
| ENSG00000095794\|CREM | 2426.9573 | 1.0742 | ##### | 8.3515 | 6.7418E-17 | 4.3981E-16 |
| ENSG00000146909\|NOM1 | 4635.2127 | 1.0357 | ##### | 8.3509 | 6.7723E-17 | 4.4170E-16 |
| ENSG00000132824\|SERINC3 | 16795.6622 | 0.7498 | ##### | 8.3499 | 6.8348E-17 | 4.4568E-16 |
| ENSG00000232818\|RPS2P32 | 111.8820 | -3.2449 | ##### | -8.3492 | 6.8708E-17 | 4.4793E-16 |
| ENSG00000116120\|FARSB | 6546.8704 | -1.0427 | ##### | -8.3469 | 7.0049E-17 | 4.5657E-16 |
| ENSG00000181804\|SLC9A9 | 84.6625 | -4.1170 | ##### | -8.3454 | 7.0993E-17 | 4.6262E-16 |
| ENSG00000131759\|RARA | 2047.1786 | -1.0499 | ##### | -8.3452 | 7.1065E-17 | 4.6299E-16 |
| ENSG00000204681\|GABBR1 | 1026.6037 | 1.2034 | ##### | 8.3429 | 7.2478E-17 | 4.7209E-16 |
| ENSG00000210049\|MT-TF | 97.0406 | 1.6571 | ##### | 8.3392 | 7.4775E-17 | 4.8695E-16 |
| ENSG00000168564\|CDKN2AIP | 1121.6598 | -1.0783 | ##### | -8.3390 | 7.4913E-17 | 4.8775E-16 |
| ENSG00000070047\|PHRF1 | 7498.2851 | 0.9828 | ##### | 8.3385 | 7.5251E-17 | 4.8984E-16 |
| ENSG00000145388\|METTL14 | 1152.8429 | -1.1592 | ##### | -8.3379 | 7.5612E-17 | 4.9208E-16 |
| ENSG00000132128\|LRRC41 | 7756.1602 | -1.0523 | ##### | -8.3369 | 7.6256E-17 | 4.9616E-16 |
| ENSG00000107672\|NSMCE4A | 2247.6645 | -1.0675 | ##### | -8.3350 | 7.7500E-17 | 5.0414E-16 |
| ENSG00000187164\|SHTN1 | 6823.6048 | -0.9201 | ##### | -8.3338 | 7.8265E-17 | 5.0901E-16 |
| ENSG00000085433\|WDR47 | 2376.4729 | 1.0457 | ##### | 8.3328 | 7.8966E-17 | 5.1346E-16 |
| ENSG00000114251\|WNT5A | 377.8214 | -1.3265 | ##### | -8.3325 | 7.9166E-17 | 5.1465E-16 |
| ENSG00000185761\|ADAMTSL5 | 1361.0292 | 1.1120 | ##### | 8.3324 | 7.9200E-17 | 5.1476E-16 |
| ENSG00000155906\|RMND1 | 2377.0636 | 1.0605 | ##### | 8.3310 | 8.0191E-17 | 5.2108E-16 |
| ENSG00000204843\|DCTN1 | 23266.9505 | 0.9700 | ##### | 8.3294 | 8.1276E-17 | 5.2800E-16 |
| ENSG00000020256\|ZFP64 | 3477.1703 | 0.8101 | ##### | 8.3293 | 8.1291E-17 | 5.2800E-16 |

| | | | | | | |
|---|---|---|---|---|---|---|
| ENSG00000280273 | 159.7561 | -1.6541 | ##### | -8.3289 | 8.1566E-17 | 5.2968E-16 |
| ENSG00000138160\|KIF11 | 6805.0666 | 1.0300 | ##### | 8.3286 | 8.1776E-17 | 5.3092E-16 |
| ENSG00000065328\|MCM10 | 5569.4401 | 1.0180 | ##### | 8.3264 | 8.3307E-17 | 5.4074E-16 |
| ENSG00000233024\|NPIPA9 | 1137.6093 | 1.1920 | ##### | 8.3249 | 8.4388E-17 | 5.4764E-16 |
| ENSG00000041880\|PARP3 | 1864.4529 | -1.0572 | ##### | -8.3246 | 8.4586E-17 | 5.4880E-16 |
| ENSG00000153310\|CYRIB | 7865.5702 | -0.8612 | ##### | -8.3239 | 8.5089E-17 | 5.5195E-16 |
| ENSG00000182158\|CREB3L2 | 4188.0207 | -1.0166 | ##### | -8.3228 | 8.5915E-17 | 5.5719E-16 |
| ENSG00000205534\|SMG1P2 | 1572.9805 | 1.1787 | ##### | 8.3216 | 8.6803E-17 | 5.6282E-16 |
| ENSG00000122884\|P4HA1 | 4222.6760 | -1.0499 | ##### | -8.3207 | 8.7422E-17 | 5.6672E-16 |
| ENSG00000158122\|PRXL2C | 1218.0526 | -1.0836 | ##### | -8.3200 | 8.7977E-17 | 5.7019E-16 |
| ENSG00000120915\|EPHX2 | 763.6385 | -1.1669 | ##### | -8.3136 | 9.2874E-17 | 6.0179E-16 |
| ENSG00000204060\|FOXO6 | 215.8893 | 1.5619 | ##### | 8.3112 | 9.4743E-17 | 6.1377E-16 |
| ENSG00000226416\|MRPL23-AS1 | 61.1052 | -8.2659 | ##### | -8.3107 | 9.5117E-17 | 6.1606E-16 |
| ENSG00000103150\|MLYCD | 719.5957 | -1.2082 | ##### | -8.3071 | 9.8059E-17 | 6.3498E-16 |
| ENSG00000171790\|SLFNL1 | 272.4497 | -1.7323 | ##### | -8.3069 | 9.8219E-17 | 6.3588E-16 |
| ENSG00000138669\|PRKG2 | 430.8093 | -1.3991 | ##### | -8.3061 | 9.8907E-17 | 6.4019E-16 |
| ENSG00000132382\|MYBBP1A | 16812.1509 | 0.9537 | ##### | 8.3057 | 9.9236E-17 | 6.4218E-16 |
| ENSG00000048828\|FAM120A | 19969.7957 | -0.9641 | ##### | -8.3036 | 1.0098E-16 | 6.5332E-16 |
| ENSG00000269378\|ITGB1P1 | 2899.2469 | -1.0511 | ##### | -8.3030 | 1.0151E-16 | 6.5664E-16 |
| ENSG00000181773\|GPR3 | 470.0669 | 1.2785 | ##### | 8.3010 | 1.0328E-16 | 6.6790E-16 |
| ENSG00000171017\|LRRC8E | 779.9327 | -1.1589 | ##### | -8.3003 | 1.0388E-16 | 6.7166E-16 |
| ENSG00000150093\|ITGB1 | ########## | -0.9683 | ##### | -8.2982 | 1.0570E-16 | 6.8325E-16 |
| ENSG00000132122\|SPATA6 | 515.4873 | -1.2070 | ##### | -8.2954 | 1.0824E-16 | 6.9951E-16 |
| ENSG00000152503\|TRIM36 | 255.8661 | 2.0617 | ##### | 8.2947 | 1.0883E-16 | 7.0319E-16 |
| ENSG00000007047\|MARK4 | 2327.5877 | -0.8789 | ##### | -8.2902 | 1.1308E-16 | 7.3048E-16 |
| ENSG00000116209\|TMEM59 | 5682.0208 | -1.0112 | ##### | -8.2897 | 1.1353E-16 | 7.3329E-16 |
| ENSG00000165716\|DIPK1B | 1673.8142 | -1.1348 | ##### | -8.2881 | 1.1508E-16 | 7.4309E-16 |
| ENSG00000169896\|ITGAM | 334.9912 | 1.3636 | ##### | 8.2864 | 1.1673E-16 | 7.5362E-16 |
| ENSG00000110047\|EHD1 | 7730.8508 | -1.0283 | ##### | -8.2849 | 1.1823E-16 | 7.6297E-16 |
| ENSG00000129315\|CCNT1 | 7954.0964 | 1.0305 | ##### | 8.2849 | 1.1823E-16 | 7.6297E-16 |
| ENSG00000114739\|ACVR2B | 530.6056 | 1.2865 | ##### | 8.2848 | 1.1830E-16 | 7.6324E-16 |
| ENSG00000002016\|RAD52 | 1555.4142 | 1.0674 | ##### | 8.2843 | 1.1880E-16 | 7.6631E-16 |
| ENSG00000076321\|KLHL20 | 1467.3736 | -1.0597 | ##### | -8.2838 | 1.1930E-16 | 7.6933E-16 |
| ENSG00000151322\|NPAS3 | 176.6916 | 8.7996 | ##### | 8.2831 | 1.2004E-16 | 7.7395E-16 |
| ENSG00000171097\|KYAT1 | 407.4376 | -1.9645 | ##### | -8.2830 | 1.2014E-16 | 7.7444E-16 |
| ENSG00000125633\|CCDC93 | 5068.8874 | -1.0285 | ##### | -8.2815 | 1.2161E-16 | 7.8375E-16 |
| ENSG00000026025\|VIM | ########## | 0.9950 | ##### | 8.2805 | 1.2265E-16 | 7.9025E-16 |
| ENSG00000164944\|VIRMA | 7246.9060 | -0.9799 | ##### | -8.2787 | 1.2449E-16 | 8.0196E-16 |
| ENSG00000179115\|FARSA | 12781.6606 | 0.9137 | ##### | 8.2776 | 1.2568E-16 | 8.0944E-16 |
| ENSG00000126259\|KIRREL2 | 297.3366 | -1.5111 | ##### | -8.2774 | 1.2590E-16 | 8.1068E-16 |
| ENSG00000100836\|PABPN1 | 11843.9241 | 1.0119 | ##### | 8.2762 | 1.2715E-16 | 8.1856E-16 |
| ENSG00000143149\|ALDH9A1 | 4679.3053 | -1.0137 | ##### | -8.2758 | 1.2761E-16 | 8.2137E-16 |
| ENSG00000232093\|DCST1-AS1 | 164.7021 | 1.6120 | ##### | 8.2740 | 1.2953E-16 | 8.3352E-16 |
| ENSG00000234997\|LINC02966 | 125.0032 | -1.8434 | ##### | -8.2737 | 1.2987E-16 | 8.3553E-16 |
| ENSG00000267254\|ZNF790-AS1 | 305.4812 | -1.4308 | ##### | -8.2732 | 1.3046E-16 | 8.3917E-16 |
| ENSG00000111678\|C12orf57 | 10908.7579 | 1.0036 | ##### | 8.2729 | 1.3071E-16 | 8.4055E-16 |
| ENSG00000121064\|SCPEP1 | 2852.1068 | -1.0190 | ##### | -8.2699 | 1.3408E-16 | 8.6205E-16 |
| ENSG00000127838\|PNKD | 4327.3638 | -1.0552 | ##### | -8.2686 | 1.3549E-16 | 8.7096E-16 |
| ENSG00000115520\|COQ10B | 2756.9427 | 1.0405 | ##### | 8.2667 | 1.3768E-16 | 8.8482E-16 |
| ENSG00000241399\|CD302 | 152.2402 | -1.6710 | ##### | -8.2666 | 1.3788E-16 | 8.8572E-16 |
| ENSG00000196937\|FAM3C | 6896.0989 | 0.9855 | ##### | 8.2666 | 1.3787E-16 | 8.8572E-16 |
| ENSG00000126243\|LRFN3 | 1128.6239 | -1.1782 | ##### | -8.2656 | 1.3901E-16 | 8.9280E-16 |
| ENSG00000290917\|RAET1K | 117.4169 | 2.5589 | ##### | 8.2654 | 1.3917E-16 | 8.9367E-16 |
| ENSG00000000003\|TSPAN6 | 2398.4504 | 1.0327 | ##### | 8.2653 | 1.3936E-16 | 8.9465E-16 |
| ENSG00000143344\|RGL1 | 401.6401 | -1.2730 | ##### | -8.2649 | 1.3979E-16 | 8.9720E-16 |
| ENSG00000151422\|FER | 2071.5146 | 1.0690 | ##### | 8.2629 | 1.4214E-16 | 9.1215E-16 |
| ENSG00000169490\|TM2D2 | 4168.7486 | -1.0003 | ##### | -8.2588 | 1.4720E-16 | 9.4441E-16 |
| ENSG00000050393\|MCUR1 | 3607.2262 | 0.9876 | ##### | 8.2587 | 1.4730E-16 | 9.4482E-16 |
| ENSG00000184007\|PTP4A2 | 29029.4865 | -0.5911 | ##### | -8.2558 | 1.5087E-16 | 9.6750E-16 |
| ENSG00000131778\|CHD1L | 8001.9271 | 0.8789 | ##### | 8.2532 | 1.5423E-16 | 9.8887E-16 |
| ENSG00000198794\|SCAMP5 | 656.1323 | -1.1594 | ##### | -8.2528 | 1.5478E-16 | 9.9217E-16 |
| ENSG00000215041\|NEURL4 | 4930.9953 | 0.9944 | ##### | 8.2516 | 1.5629E-16 | 1.0017E-15 |
| ENSG00000188107\|EYS | 71.4286 | -2.6198 | ##### | -8.2487 | 1.6013E-16 | 1.0260E-15 |

| | | | | | | |
|---|---|---|---|---|---|---|
| ENSG00000090621\|PABPC4 | 25798.8403 | 0.6531 | ##### | 8.2472 | 1.6217E-16 | 1.0389E-15 |
| ENSG00000232040\|SCAND3 | 387.7595 | 1.4362 | ##### | 8.2471 | 1.6231E-16 | 1.0395E-15 |
| ENSG00000106066\|CPVL | 595.4571 | 6.4029 | ##### | 8.2441 | 1.6643E-16 | 1.0657E-15 |
| ENSG00000156273\|BACH1 | 2897.0476 | -1.0218 | ##### | -8.2422 | 1.6905E-16 | 1.0823E-15 |
| ENSG00000139899\|CBLN3 | 317.9716 | 1.4169 | ##### | 8.2369 | 1.7667E-16 | 1.1308E-15 |
| ENSG00000102445\|RUBCNL | 91.8731 | -2.4266 | ##### | -8.2369 | 1.7672E-16 | 1.1309E-15 |
| ENSG00000239445\|ST3GAL6-AS1 | 43.1469 | 5.0628 | ##### | 8.2367 | 1.7701E-16 | 1.1325E-15 |
| ENSG00000037474\|NSUN2 | 27382.4251 | 0.7480 | ##### | 8.2364 | 1.7744E-16 | 1.1350E-15 |
| ENSG00000289971 | 172.7219 | -1.7351 | ##### | -8.2361 | 1.7790E-16 | 1.1377E-15 |
| ENSG00000196151\|WDSUB1 | 745.1113 | -1.1875 | ##### | -8.2356 | 1.7867E-16 | 1.1424E-15 |
| ENSG00000068745\|IP6K2 | 3455.5869 | 1.0151 | ##### | 8.2350 | 1.7960E-16 | 1.1481E-15 |
| ENSG00000164920\|OSR2 | 274.2155 | -1.6501 | ##### | -8.2347 | 1.8008E-16 | 1.1509E-15 |
| ENSG00000144451\|SPAG16 | 620.2804 | -1.2486 | ##### | -8.2345 | 1.8027E-16 | 1.1519E-15 |
| ENSG00000261488\|TBILA | 107.4998 | 2.7927 | ##### | 8.2343 | 1.8064E-16 | 1.1540E-15 |
| ENSG00000246263\|UBR5-DT | 117.0007 | -1.6875 | ##### | -8.2342 | 1.8071E-16 | 1.1542E-15 |
| ENSG00000235545\|DOCK7-DT | 100.9034 | -8.9199 | ##### | -8.2281 | 1.9026E-16 | 1.2149E-15 |
| ENSG00000162944\|RFTN2 | 150.0987 | -1.7301 | ##### | -8.2280 | 1.9038E-16 | 1.2154E-15 |
| ENSG00000213983\|AP1G2 | 3754.8647 | 1.0199 | ##### | 8.2270 | 1.9195E-16 | 1.2252E-15 |
| ENSG00000224152\|BAZ2B-AS1 | 173.1058 | -1.4767 | ##### | -8.2268 | 1.9222E-16 | 1.2267E-15 |
| ENSG00000137845\|ADAM10 | 9188.6250 | -1.0034 | ##### | -8.2265 | 1.9272E-16 | 1.2295E-15 |
| ENSG00000239672\|NME1 | 19490.1325 | 1.0144 | ##### | 8.2255 | 1.9444E-16 | 1.2402E-15 |
| ENSG00000101928\|MOSPD1 | 1119.4012 | 1.1060 | ##### | 8.2250 | 1.9525E-16 | 1.2452E-15 |
| ENSG00000167258\|CDK12 | 9347.6468 | 0.9783 | ##### | 8.2227 | 1.9905E-16 | 1.2691E-15 |
| ENSG00000100479\|POLE2 | 1365.8092 | 1.1381 | ##### | 8.2225 | 1.9928E-16 | 1.2703E-15 |
| ENSG00000123159\|GIPC1 | 9864.7646 | -0.9360 | ##### | -8.2212 | 2.0142E-16 | 1.2837E-15 |
| ENSG00000213185\|FAM24B | 360.5952 | -1.3637 | ##### | -8.2206 | 2.0255E-16 | 1.2906E-15 |
| ENSG00000261308\|FIGNL2 | 110.5128 | 2.3942 | ##### | 8.2205 | 2.0269E-16 | 1.2912E-15 |
| ENSG00000101473\|ACOT8 | 1894.0122 | 1.0662 | ##### | 8.2198 | 2.0381E-16 | 1.2981E-15 |
| ENSG00000171812\|COL8A2 | 47.9446 | -2.9940 | ##### | -8.2136 | 2.1471E-16 | 1.3673E-15 |
| ENSG00000074590\|NUAK1 | 2837.6250 | -1.0494 | ##### | -8.2122 | 2.1712E-16 | 1.3823E-15 |
| ENSG00000113552\|GNPDA1 | 6870.5029 | -0.9957 | ##### | -8.2119 | 2.1769E-16 | 1.3856E-15 |
| ENSG00000134758\|RNF138 | 3875.3106 | 1.0055 | ##### | 8.2115 | 2.1851E-16 | 1.3905E-15 |
| ENSG00000122779\|TRIM24 | 3049.2146 | 1.0187 | ##### | 8.2114 | 2.1857E-16 | 1.3906E-15 |
| ENSG00000267244 | 133.3741 | 1.7213 | ##### | 8.2096 | 2.2184E-16 | 1.4111E-15 |
| ENSG00000137094\|DNAJB5 | 874.4357 | 1.0974 | ##### | 8.2083 | 2.2441E-16 | 1.4272E-15 |
| ENSG00000117450\|PRDX1 | 91260.7517 | -0.9792 | ##### | -8.2074 | 2.2600E-16 | 1.4370E-15 |
| ENSG00000115020\|PIKFYVE | 5612.3741 | 0.9812 | ##### | 8.2072 | 2.2645E-16 | 1.4395E-15 |
| ENSG00000103316\|CRYM | 86.1428 | -4.6119 | ##### | -8.2036 | 2.3333E-16 | 1.4830E-15 |
| ENSG00000154096\|THY1 | 2062.3642 | ####### | ##### | -8.2034 | 2.3361E-16 | 1.4844E-15 |
| ENSG00000203879\|GDI1 | 15976.9532 | 0.9854 | ##### | 8.2029 | 2.3471E-16 | 1.4911E-15 |
| ENSG00000119599\|DCAF4 | 1117.5351 | 1.1774 | ##### | 8.2024 | 2.3558E-16 | 1.4963E-15 |
| ENSG00000003509\|NDUFAF7 | 1416.7593 | 0.8004 | ##### | 8.1986 | 2.4326E-16 | 1.5447E-15 |
| ENSG00000228716\|DHFR | 5559.3451 | 1.2065 | ##### | 8.1968 | 2.4680E-16 | 1.5669E-15 |
| ENSG00000223478\|ZDHHC12-DT | 241.1806 | 1.4606 | ##### | 8.1959 | 2.4863E-16 | 1.5782E-15 |
| ENSG00000060140\|STYK1 | 1134.7442 | 1.0654 | ##### | 8.1951 | 2.5038E-16 | 1.5890E-15 |
| ENSG00000248092\|NNT-AS1 | 425.4922 | -8.6794 | ##### | -8.1938 | 2.5320E-16 | 1.6065E-15 |
| ENSG00000257167\|TMPO-AS1 | 1497.7519 | 1.1173 | ##### | 8.1933 | 2.5406E-16 | 1.6116E-15 |
| ENSG00000099860\|GADD45B | 3247.8888 | 0.9873 | ##### | 8.1910 | 2.5903E-16 | 1.6428E-15 |
| ENSG00000197863\|ZNF790 | 241.5929 | -1.4307 | ##### | -8.1902 | 2.6089E-16 | 1.6542E-15 |
| ENSG00000259146\|SIPA1L1-AS1 | 71.5846 | -9.2460 | ##### | -8.1880 | 2.6551E-16 | 1.6832E-15 |
| ENSG00000226479\|TMEM185B | 5562.5479 | -0.9632 | ##### | -8.1878 | 2.6606E-16 | 1.6863E-15 |
| ENSG00000010404\|IDS | 11541.5591 | -1.0156 | ##### | -8.1850 | 2.7224E-16 | 1.7251E-15 |
| ENSG00000172890\|NADSYN1 | 3407.9154 | 1.0418 | ##### | 8.1845 | 2.7338E-16 | 1.7320E-15 |
| ENSG00000129003\|VPS13C | 1473.4323 | -1.0856 | ##### | -8.1838 | 2.7504E-16 | 1.7421E-15 |
| ENSG00000154222\|CC2D1B | 4150.5837 | -0.9849 | ##### | -8.1838 | 2.7510E-16 | 1.7422E-15 |
| ENSG00000260934 | 91.2608 | -1.7493 | ##### | -8.1837 | 2.7526E-16 | 1.7428E-15 |
| ENSG00000236924 | 122.0709 | -2.8586 | ##### | -8.1829 | 2.7699E-16 | 1.7534E-15 |
| ENSG00000188157\|AGRN | 35502.2872 | -0.9589 | ##### | -8.1819 | 2.7936E-16 | 1.7680E-15 |
| ENSG00000119682\|AREL1 | 6177.7857 | 0.9764 | ##### | 8.1812 | 2.8094E-16 | 1.7776E-15 |
| ENSG00000151414\|NEK7 | 5041.8769 | -0.9899 | ##### | -8.1812 | 2.8104E-16 | 1.7779E-15 |
| ENSG00000196911\|KPNA5 | 723.2384 | 1.1299 | ##### | 8.1808 | 2.8205E-16 | 1.7839E-15 |
| ENSG00000110318\|CEP126 | 907.0510 | -1.0685 | ##### | -8.1777 | 2.8924E-16 | 1.8290E-15 |
| ENSG00000128284\|APOL3 | 150.7288 | -9.6337 | ##### | -8.1777 | 2.8930E-16 | 1.8290E-15 |

| | | | | | | |
|---|---|---|---|---|---|---|
| ENSG00000170836\|PPM1D | 1473.5272 | 1.0850 | ##### | 8.1777 | 2.8942E-16 | 1.8293E-15 |
| ENSG00000171365\|CLCN5 | 1126.3894 | -1.1939 | ##### | -8.1765 | 2.9232E-16 | 1.8473E-15 |
| ENSG00000117593\|DARS2 | 3644.1078 | -1.0215 | ##### | -8.1758 | 2.9386E-16 | 1.8566E-15 |
| ENSG00000109063\|MYH3 | 191.9222 | 2.0647 | ##### | 8.1756 | 2.9435E-16 | 1.8593E-15 |
| ENSG00000148288\|GBGT1 | 110.9804 | -2.6061 | ##### | -8.1756 | 2.9442E-16 | 1.8593E-15 |
| ENSG00000112299\|VNN1 | 56.0095 | -2.8573 | ##### | -8.1750 | 2.9579E-16 | 1.8676E-15 |
| ENSG00000061656\|SPAG4 | 657.8500 | 1.1515 | ##### | 8.1749 | 2.9608E-16 | 1.8690E-15 |
| ENSG00000185198\|PRSS57 | 25.8365 | -7.6205 | ##### | -8.1738 | 2.9892E-16 | 1.8865E-15 |
| ENSG00000162687\|KCNT2 | 131.6228 | -1.9671 | ##### | -8.1716 | 3.0437E-16 | 1.9205E-15 |
| ENSG00000164463\|CREBRF | 401.4208 | 1.2952 | ##### | 8.1704 | 3.0739E-16 | 1.9392E-15 |
| ENSG00000006740\|ARHGAP44 | 830.3511 | 1.0803 | ##### | 8.1691 | 3.1075E-16 | 1.9600E-15 |
| ENSG00000152944\|MED21 | 3454.5642 | 1.0332 | ##### | 8.1649 | 3.2182E-16 | 2.0294E-15 |
| ENSG00000159216\|RUNX1 | 3022.2707 | 1.0251 | ##### | 8.1648 | 3.2198E-16 | 2.0300E-15 |
| ENSG00000141314\|RHBDL3 | 214.5266 | -1.5281 | ##### | -8.1642 | 3.2346E-16 | 2.0389E-15 |
| ENSG00000055044\|NOP58 | 16535.1713 | 0.8663 | ##### | 8.1629 | 3.2712E-16 | 2.0615E-15 |
| ENSG00000143847\|PPFIA4 | 144.9232 | 1.6933 | ##### | 8.1600 | 3.3511E-16 | 2.1114E-15 |
| ENSG00000154978\|VOPP1 | 8948.8008 | -0.9697 | ##### | -8.1599 | 3.3532E-16 | 2.1123E-15 |
| ENSG00000089195\|TRMT6 | 3708.7132 | 0.8595 | ##### | 8.1584 | 3.3936E-16 | 2.1373E-15 |
| ENSG00000006652\|IFRD1 | 7340.4566 | -1.0429 | ##### | -8.1566 | 3.4445E-16 | 2.1689E-15 |
| ENSG00000143771\|CNIH4 | 3231.0692 | -0.8014 | ##### | -8.1565 | 3.4501E-16 | 2.1720E-15 |
| ENSG00000113532\|ST8SIA4 | 241.5783 | 1.4979 | ##### | 8.1554 | 3.4811E-16 | 2.1910E-15 |
| ENSG00000105520\|PLPPR2 | 3668.1750 | -0.9749 | ##### | -8.1553 | 3.4843E-16 | 2.1925E-15 |
| ENSG00000136883\|KIF12 | 190.2013 | -5.9321 | ##### | -8.1543 | 3.5110E-16 | 2.2089E-15 |
| ENSG00000290988\|ZNF781 | 65.2210 | -2.7977 | ##### | -8.1542 | 3.5159E-16 | 2.2115E-15 |
| ENSG00000139266\|MARCHF9 | 379.9950 | -1.2988 | ##### | -8.1541 | 3.5179E-16 | 2.2123E-15 |
| ENSG00000260000 | 156.5927 | -1.8040 | ##### | -8.1537 | 3.5291E-16 | 2.2189E-15 |
| ENSG00000138629\|UBL7 | 3634.5507 | 1.0432 | ##### | 8.1533 | 3.5425E-16 | 2.2268E-15 |
| ENSG00000163344\|PMVK | 3416.1387 | 1.0358 | ##### | 8.1527 | 3.5600E-16 | 2.2374E-15 |
| ENSG00000115194\|SLC30A3 | 98.2222 | 2.4205 | ##### | 8.1523 | 3.5714E-16 | 2.2441E-15 |
| ENSG00000138640\|FAM13A | 1229.6275 | -1.0765 | ##### | -8.1483 | 3.6894E-16 | 2.3177E-15 |
| ENSG00000181666\|ZNF875 | 1102.2524 | -1.0994 | ##### | -8.1473 | 3.7203E-16 | 2.3366E-15 |
| ENSG00000111328\|CDK2AP1 | 12112.4336 | 0.9816 | ##### | 8.1470 | 3.7312E-16 | 2.3430E-15 |
| ENSG00000162461\|SLC25A34 | 53.5651 | -2.0834 | ##### | -8.1448 | 3.7996E-16 | 2.3854E-15 |
| ENSG00000293478\|TP73-AS1 | 2563.8622 | 0.9838 | ##### | 8.1438 | 3.8315E-16 | 2.4050E-15 |
| ENSG00000111845\|PAK1IP1 | 3292.1195 | 0.9837 | ##### | 8.1429 | 3.8586E-16 | 2.4214E-15 |
| ENSG00000107872\|FBXL15 | 1226.9761 | -1.0361 | ##### | -8.1424 | 3.8740E-16 | 2.4306E-15 |
| ENSG00000164318\|EGFLAM | 117.9439 | 8.6957 | ##### | 8.1412 | 3.9146E-16 | 2.4556E-15 |
| ENSG00000070778\|PTPN21 | 2006.9280 | -1.0850 | ##### | -8.1394 | 3.9736E-16 | 2.4921E-15 |
| ENSG00000003400\|CASP10 | 215.5410 | -5.5635 | ##### | -8.1383 | 4.0093E-16 | 2.5139E-15 |
| ENSG00000267520 | 404.4758 | 3.4649 | ##### | 8.1377 | 4.0272E-16 | 2.5246E-15 |
| ENSG00000176884\|GRIN1 | 288.9162 | 1.5688 | ##### | 8.1373 | 4.0409E-16 | 2.5327E-15 |
| ENSG00000130054\|NALF2 | 1229.5318 | 1.2048 | ##### | 8.1368 | 4.0586E-16 | 2.5432E-15 |
| ENSG00000134709\|HOOK1 | 805.6426 | -1.1181 | ##### | -8.1364 | 4.0714E-16 | 2.5507E-15 |
| ENSG00000007255\|TRAPPC6A | 783.4135 | -1.1083 | ##### | -8.1359 | 4.0899E-16 | 2.5618E-15 |
| ENSG00000144283\|PKP4 | 10024.6358 | -0.9723 | ##### | -8.1356 | 4.1005E-16 | 2.5679E-15 |
| ENSG00000164327\|RICTOR | 6612.0477 | 0.9796 | ##### | 8.1348 | 4.1249E-16 | 2.5826E-15 |
| ENSG00000234614\|C2CD4D-AS1 | 98.0929 | -8.8766 | ##### | -8.1342 | 4.1468E-16 | 2.5958E-15 |
| ENSG00000184702\|SEPTIN5 | 1211.6505 | -1.5103 | ##### | -8.1317 | 4.2347E-16 | 2.6502E-15 |
| ENSG00000261087\|ZNNT1 | 65.1224 | -2.2055 | ##### | -8.1316 | 4.2365E-16 | 2.6508E-15 |
| ENSG00000152056\|AP1S3 | 4699.7872 | -0.9496 | ##### | -8.1264 | 4.4224E-16 | 2.7666E-15 |
| ENSG00000130150\|MOSPD2 | 1238.8937 | -1.1201 | ##### | -8.1248 | 4.4797E-16 | 2.8018E-15 |
| ENSG00000081177\|EXD2 | 2480.1221 | -1.0064 | ##### | -8.1220 | 4.5850E-16 | 2.8671E-15 |
| ENSG00000187801\|ZFP69B | 358.8128 | -1.2020 | ##### | -8.1209 | 4.6263E-16 | 2.8923E-15 |
| ENSG00000116138\|DNAJC16 | 2247.6988 | -1.0119 | ##### | -8.1202 | 4.6556E-16 | 2.9100E-15 |
| ENSG00000142002\|DPP9 | 23548.0870 | 0.5498 | ##### | 8.1162 | 4.8116E-16 | 3.0069E-15 |
| ENSG00000132475\|H3-3B | 38456.6131 | 0.9653 | ##### | 8.1155 | 4.8396E-16 | 3.0237E-15 |
| ENSG00000117020\|AKT3 | 3353.7318 | 0.9901 | ##### | 8.1143 | 4.8852E-16 | 3.0516E-15 |
| ENSG00000136059\|VILL | 375.9028 | 4.9130 | ##### | 8.1134 | 4.9221E-16 | 3.0740E-15 |
| ENSG00000263551 | 87.7397 | -2.0150 | ##### | -8.1098 | 5.0691E-16 | 3.1652E-15 |
| ENSG00000102385\|DRP2 | 172.7438 | 2.6216 | ##### | 8.1074 | 5.1735E-16 | 3.2296E-15 |
| ENSG00000198171\|DDRGK1 | 3054.6594 | -0.9799 | ##### | -8.1071 | 5.1823E-16 | 3.2345E-15 |
| ENSG00000116521\|SCAMP3 | 14508.1034 | 0.9682 | ##### | 8.1056 | 5.2484E-16 | 3.2750E-15 |
| ENSG00000171295\|ZNF440 | 308.9502 | -1.3021 | ##### | -8.1041 | 5.3150E-16 | 3.3159E-15 |

| | | | | | | |
|---|---|---|---|---|---|---|
| ENSG00000183723\|CMTM4 | 5367.3435 | -0.9884 | ##### | -8.1017 | 5.4194E-16 | 3.3803E-15 |
| ENSG00000169057\|MECP2 | 6679.7532 | 0.9806 | ##### | 8.0963 | 5.6643E-16 | 3.5323E-15 |
| ENSG00000090372\|STRN4 | 11815.2431 | -0.9552 | ##### | -8.0948 | 5.7373E-16 | 3.5771E-15 |
| ENSG00000281991\|TMEM265 | 130.5727 | -5.3331 | ##### | -8.0939 | 5.7801E-16 | 3.6030E-15 |
| ENSG00000249242\|TMEM150C | 131.4357 | 1.9584 | ##### | 8.0938 | 5.7825E-16 | 3.6038E-15 |
| ENSG00000173889\|PHC3 | 2189.8267 | -1.0034 | ##### | -8.0936 | 5.7938E-16 | 3.6101E-15 |
| ENSG00000123297\|TSFM | 3925.3686 | 1.1138 | ##### | 8.0925 | 5.8440E-16 | 3.6405E-15 |
| ENSG00000179859\|RNF227 | 496.7318 | 1.3422 | ##### | 8.0925 | 5.8450E-16 | 3.6405E-15 |
| ENSG00000137364\|TPMT | 2066.1052 | 1.0206 | ##### | 8.0917 | 5.8830E-16 | 3.6634E-15 |
| ENSG00000159753\|CARMIL2 | 308.4214 | 1.4931 | ##### | 8.0917 | 5.8858E-16 | 3.6644E-15 |
| ENSG00000284116 | 200.9330 | 1.8395 | ##### | 8.0910 | 5.9160E-16 | 3.6824E-15 |
| ENSG00000165819\|METTL3 | 5508.5816 | 0.9683 | ##### | 8.0901 | 5.9636E-16 | 3.7113E-15 |
| ENSG00000142892\|PIGK | 3028.5075 | -1.0099 | ##### | -8.0884 | 6.0436E-16 | 3.7602E-15 |
| ENSG00000151090\|THRB | 1139.8819 | -1.1132 | ##### | -8.0882 | 6.0546E-16 | 3.7663E-15 |
| ENSG00000108828\|VAT1 | 16324.0288 | -0.7323 | ##### | -8.0859 | 6.1705E-16 | 3.8376E-15 |
| ENSG00000144476\|ACKR3 | 148.4943 | -3.8159 | ##### | -8.0854 | 6.1952E-16 | 3.8521E-15 |
| ENSG00000178026\|LRRC75B | 304.8077 | -1.3757 | ##### | -8.0848 | 6.2269E-16 | 3.8711E-15 |
| ENSG00000127124\|HIVEP3 | 601.6383 | 1.2382 | ##### | 8.0846 | 6.2350E-16 | 3.8745E-15 |
| ENSG00000234127\|TRIM26 | 4526.6794 | -0.9505 | ##### | -8.0846 | 6.2346E-16 | 3.8745E-15 |
| ENSG00000267645 | 1445.1533 | 1.0546 | ##### | 8.0843 | 6.2538E-16 | 3.8854E-15 |
| ENSG00000130518\|IQCN | 86.8868 | -2.3320 | ##### | -8.0833 | 6.3045E-16 | 3.9160E-15 |
| ENSG00000177432\|NAP1L5 | 270.7508 | -1.3451 | ##### | -8.0832 | 6.3091E-16 | 3.9181E-15 |
| ENSG00000132109\|TRIM21 | 630.6176 | -1.2510 | ##### | -8.0831 | 6.3157E-16 | 3.9214E-15 |
| ENSG00000288885 | 153.9448 | 1.7838 | ##### | 8.0812 | 6.4115E-16 | 3.9801E-15 |
| ENSG00000162889\|MAPKAPK2 | 6566.1959 | -0.9442 | ##### | -8.0804 | 6.4528E-16 | 4.0049E-15 |
| ENSG00000253187\|HOXA10-AS | 136.9561 | -3.0694 | ##### | -8.0803 | 6.4585E-16 | 4.0076E-15 |
| ENSG00000149273\|RPS3 | 91040.4521 | 0.9496 | ##### | 8.0789 | 6.5342E-16 | 4.0537E-15 |
| ENSG00000099821\|POLRMT | 11489.1088 | 0.9514 | ##### | 8.0781 | 6.5810E-16 | 4.0818E-15 |
| ENSG00000170275\|CRTAP | 9955.5494 | -1.0153 | ##### | -8.0762 | 6.6817E-16 | 4.1435E-15 |
| ENSG00000131495\|NDUFA2 | 4288.6661 | 0.9796 | ##### | 8.0696 | 7.0527E-16 | 4.3726E-15 |
| ENSG00000152578\|GRIA4 | 59.3929 | 3.8399 | ##### | 8.0689 | 7.0911E-16 | 4.3955E-15 |
| ENSG00000269937 | 649.7942 | 1.3760 | ##### | 8.0688 | 7.1012E-16 | 4.4008E-15 |
| ENSG00000125755\|SYMPK | 12875.6598 | -0.7851 | ##### | -8.0682 | 7.1366E-16 | 4.4219E-15 |
| ENSG00000258405\|ZNF578 | 351.3163 | ####### | ##### | -8.0675 | 7.1781E-16 | 4.4467E-15 |
| ENSG00000123485\|HJURP | 10268.4093 | 0.9724 | ##### | 8.0636 | 7.4088E-16 | 4.5886E-15 |
| ENSG00000221821\|C6orf226 | 188.8941 | -1.5341 | ##### | -8.0628 | 7.4547E-16 | 4.6161E-15 |
| ENSG00000231170\|PDK4-AS1 | 35.8259 | -7.9987 | ##### | -8.0615 | 7.5366E-16 | 4.6659E-15 |
| ENSG00000228623\|ZNF883 | 1399.3396 | ####### | ##### | -8.0608 | 7.5792E-16 | 4.6912E-15 |
| ENSG00000065060\|BLTP3A | 2462.6932 | -1.0227 | ##### | -8.0592 | 7.6770E-16 | 4.7508E-15 |
| ENSG00000179277\|MEIS3P1 | 1703.1045 | ####### | ##### | -8.0588 | 7.7076E-16 | 4.7688E-15 |
| ENSG00000240429\|LRRFIP1P1 | 126.4421 | -1.6870 | ##### | -8.0586 | 7.7152E-16 | 4.7725E-15 |
| ENSG00000142409\|ZNF787 | 5324.7797 | -0.9738 | ##### | -8.0571 | 7.8137E-16 | 4.8324E-15 |
| ENSG00000142611\|PRDM16 | 72.5018 | 2.2521 | ##### | 8.0569 | 7.8273E-16 | 4.8398E-15 |
| ENSG00000290833 | 728.4854 | 1.1331 | ##### | 8.0542 | 7.9979E-16 | 4.9443E-15 |
| ENSG00000162231\|NXF1 | 4742.7520 | 0.9756 | ##### | 8.0540 | 8.0100E-16 | 4.9507E-15 |
| ENSG00000189046\|ALKBH2 | 1968.5199 | 0.6179 | ##### | 8.0532 | 8.0682E-16 | 4.9857E-15 |
| ENSG00000133794\|BMAL1 | 713.3465 | -1.1428 | ##### | -8.0505 | 8.2424E-16 | 5.0923E-15 |
| ENSG00000160007\|ARHGAP35 | 13528.8542 | -0.9825 | ##### | -8.0478 | 8.4301E-16 | 5.2072E-15 |
| ENSG00000026036\|RTEL1-TNFRSF6B | 1022.8756 | 1.3325 | ##### | 8.0458 | 8.5716E-16 | 5.2934E-15 |
| ENSG00000186187\|ZNRF1 | 2766.8229 | -0.9609 | ##### | -8.0454 | 8.5937E-16 | 5.3060E-15 |
| ENSG00000101199\|ARFGAP1 | 13180.6125 | 0.9547 | ##### | 8.0433 | 8.7424E-16 | 5.3967E-15 |
| ENSG00000244731\|C4A | 181.9216 | -1.4891 | ##### | -8.0409 | 8.9199E-16 | 5.5051E-15 |
| ENSG00000161800\|RACGAP1 | 11089.3875 | 0.9769 | ##### | 8.0395 | 9.0173E-16 | 5.5641E-15 |
| ENSG00000258072 | 78.2998 | 2.5678 | ##### | 8.0391 | 9.0496E-16 | 5.5828E-15 |
| ENSG00000144840\|RABL3 | 1652.8416 | -0.7833 | ##### | -8.0385 | 9.0977E-16 | 5.6114E-15 |
| ENSG00000136816\|TOR1B | 3979.0020 | -1.0187 | ##### | -8.0366 | 9.2373E-16 | 5.6963E-15 |
| ENSG00000115935\|WIPF1 | 1342.1566 | 1.0554 | ##### | 8.0336 | 9.4616E-16 | 5.8334E-15 |
| ENSG00000131584\|ACAP3 | 2569.1538 | -0.9875 | ##### | -8.0312 | 9.6539E-16 | 5.9507E-15 |
| ENSG00000106052\|TAX1BP1 | 21973.8833 | -0.8528 | ##### | -8.0300 | 9.7497E-16 | 6.0086E-15 |
| ENSG00000247081\|BAALC-AS1 | 121.8750 | -1.9387 | ##### | -8.0274 | 9.9546E-16 | 6.1336E-15 |
| ENSG00000278594 | 138.8946 | 1.6144 | ##### | 8.0265 | 1.0033E-15 | 6.1807E-15 |
| ENSG00000291085\|TMEM191A | 309.7708 | 1.3639 | ##### | 8.0255 | 1.0109E-15 | 6.2264E-15 |
| ENSG00000073756\|PTGS2 | 1366.6626 | -1.0520 | ##### | -8.0251 | 1.0143E-15 | 6.2456E-15 |

| | | | | | | |
|---|---|---|---|---|---|---|
| ENSG00000155903\|RASA2 | 2088.1962 | 1.0150 | ##### | 8.0240 | 1.0239E-15 | 6.3035E-15 |
| ENSG00000139974\|SLC38A6 | 1027.8188 | -0.9917 | ##### | -8.0195 | 1.0620E-15 | 6.5368E-15 |
| ENSG00000130724\|CHMP2A | 4861.3281 | -1.0062 | ##### | -8.0190 | 1.0659E-15 | 6.5598E-15 |
| ENSG00000066583\|ISOC1 | 2318.8326 | 1.0552 | ##### | 8.0190 | 1.0663E-15 | 6.5609E-15 |
| ENSG00000126878\|AIF1L | 1204.0967 | 1.0175 | ##### | 8.0189 | 1.0671E-15 | 6.5642E-15 |
| ENSG00000142687\|KIAA0319L | 5272.4599 | -0.9683 | ##### | -8.0184 | 1.0710E-15 | 6.5871E-15 |
| ENSG00000101474\|APMAP | 10355.7636 | -0.3928 | ##### | -8.0175 | 1.0790E-15 | 6.6344E-15 |
| ENSG00000159784\|FAM131B | 481.8811 | 1.2568 | ##### | 8.0172 | 1.0820E-15 | 6.6515E-15 |
| ENSG00000272047\|GTF2H5 | 1495.5663 | -1.1058 | ##### | -8.0171 | 1.0829E-15 | 6.6561E-15 |
| ENSG00000184828\|ZBTB7C | 29.2856 | -3.3309 | ##### | -8.0165 | 1.0879E-15 | 6.6853E-15 |
| ENSG00000186666\|BCDIN3D | 634.4881 | -1.2989 | ##### | -8.0152 | 1.0993E-15 | 6.7542E-15 |
| ENSG00000249378\|LINC01060 | 848.2372 | ####### | ##### | -8.0147 | 1.1043E-15 | 6.7830E-15 |
| ENSG00000100425\|BRD1 | 2453.7266 | 1.0196 | ##### | 8.0146 | 1.1046E-15 | 6.7839E-15 |
| ENSG00000165490\|DDIAS | 2336.4389 | 1.0479 | ##### | 8.0125 | 1.1236E-15 | 6.8988E-15 |
| ENSG00000221914\|PPP2R2A | 5834.8541 | 0.5844 | ##### | 8.0119 | 1.1291E-15 | 6.9314E-15 |
| ENSG00000249353\|NPM1P27 | 1396.8018 | 1.0461 | ##### | 8.0119 | 1.1295E-15 | 6.9325E-15 |
| ENSG00000183826\|BTBD9 | 1083.2648 | -1.0501 | ##### | -8.0116 | 1.1324E-15 | 6.9486E-15 |
| ENSG00000181929\|PRKAG1 | 3190.7992 | -1.0508 | ##### | -8.0114 | 1.1339E-15 | 6.9563E-15 |
| ENSG00000244462\|RBM12 | 8876.4042 | 0.6771 | ##### | 8.0083 | 1.1634E-15 | 7.1358E-15 |
| ENSG00000291233 | 50.4140 | 2.6357 | ##### | 8.0078 | 1.1673E-15 | 7.1571E-15 |
| ENSG00000241316\|SUCLG2-DT | 589.9397 | -1.2526 | ##### | -8.0079 | 1.1672E-15 | 7.1571E-15 |
| ENSG00000163956\|LRPAP1 | 7349.8907 | 0.8225 | ##### | 8.0078 | 1.1680E-15 | 7.1599E-15 |
| ENSG00000156787\|TBC1D31 | 1787.1787 | -0.9797 | ##### | -8.0055 | 1.1899E-15 | 7.2922E-15 |
| ENSG00000267598 | 1407.5743 | 1.9224 | ##### | 8.0039 | 1.2049E-15 | 7.3831E-15 |
| ENSG00000188883\|KLRG2 | 225.5644 | 2.0089 | ##### | 8.0038 | 1.2060E-15 | 7.3880E-15 |
| ENSG00000204392\|LSM2 | 3267.5765 | 1.0295 | ##### | 8.0035 | 1.2097E-15 | 7.4094E-15 |
| ENSG00000167476\|JSRP1 | 3185.6284 | 1.6505 | ##### | 8.0031 | 1.2134E-15 | 7.4302E-15 |
| ENSG00000235092\|ID2-AS1 | 399.7148 | 1.2900 | ##### | 8.0020 | 1.2237E-15 | 7.4918E-15 |
| ENSG00000168490\|PHYHIP | 109.4212 | -9.2161 | ##### | -8.0019 | 1.2248E-15 | 7.4973E-15 |
| ENSG00000265241\|RBM8A | 21932.6390 | 0.9428 | ##### | 8.0019 | 1.2255E-15 | 7.5001E-15 |
| ENSG00000161547\|SRSF2 | 26801.6487 | 0.9735 | ##### | 7.9986 | 1.2580E-15 | 7.6969E-15 |
| ENSG00000115252\|PDE1A | 149.5997 | -5.0311 | ##### | -7.9955 | 1.2903E-15 | 7.8932E-15 |
| ENSG00000142949\|PTPRF | 38895.5858 | -0.9455 | ##### | -7.9933 | 1.3140E-15 | 8.0363E-15 |
| ENSG00000140022\|STON2 | 264.1141 | -1.3751 | ##### | -7.9915 | 1.3329E-15 | 8.1506E-15 |
| ENSG00000176108\|CHMP6 | 1594.8645 | -1.0535 | ##### | -7.9913 | 1.3349E-15 | 8.1610E-15 |
| ENSG00000143554\|SLC27A3 | 1832.9601 | -0.9932 | ##### | -7.9873 | 1.3796E-15 | 8.4309E-15 |
| ENSG00000123870\|ZNF137P | 107.7966 | -2.6597 | ##### | -7.9873 | 1.3794E-15 | 8.4309E-15 |
| ENSG00000107789\|MINPP1 | 2534.2952 | -0.6068 | ##### | -7.9872 | 1.3804E-15 | 8.4343E-15 |
| ENSG00000290970 | 23.2914 | -4.5950 | ##### | -7.9857 | 1.3968E-15 | 8.5327E-15 |
| ENSG00000099901\|RANBP1 | 13653.3841 | 0.9540 | ##### | 7.9826 | 1.4326E-15 | 8.7493E-15 |
| ENSG00000291223\|FAM86DP | 909.4006 | -1.1214 | ##### | -7.9815 | 1.4453E-15 | 8.8253E-15 |
| ENSG00000178982\|EIF3K | 9902.1486 | -0.9691 | ##### | -7.9814 | 1.4468E-15 | 8.8326E-15 |
| ENSG00000174130\|TLR6 | 1148.1765 | -1.0439 | ##### | -7.9793 | 1.4722E-15 | 8.9859E-15 |
| ENSG00000213638\|ADAT3 | 548.1511 | 1.1962 | ##### | 7.9767 | 1.5024E-15 | 9.1683E-15 |
| ENSG00000117139\|KDM5B | 5364.1103 | -0.9418 | ##### | -7.9734 | 1.5441E-15 | 9.4209E-15 |
| ENSG00000171940\|ZNF217 | 9419.5575 | 0.9217 | ##### | 7.9707 | 1.5783E-15 | 9.6274E-15 |
| ENSG00000129473\|BCL2L2 | 3279.9715 | 0.8361 | ##### | 7.9706 | 1.5794E-15 | 9.6322E-15 |
| ENSG00000175536\|LIPT2 | 190.7499 | 1.4924 | ##### | 7.9700 | 1.5865E-15 | 9.6737E-15 |
| ENSG00000108691\|CCL2 | 3421.0146 | 1.0785 | ##### | 7.9686 | 1.6051E-15 | 9.7846E-15 |
| ENSG00000147044\|CASK | 2782.1709 | -1.0130 | ##### | -7.9684 | 1.6074E-15 | 9.7971E-15 |
| ENSG00000101638\|ST8SIA5 | 58.7918 | 2.3596 | ##### | 7.9683 | 1.6089E-15 | 9.8041E-15 |
| ENSG00000131149\|GSE1 | 9663.1373 | 0.9258 | ##### | 7.9658 | 1.6421E-15 | 1.0004E-14 |
| ENSG00000228843 | 64.5560 | 2.0801 | ##### | 7.9652 | 1.6499E-15 | 1.0050E-14 |
| ENSG00000117984\|CTSD | 20222.7066 | -0.9627 | ##### | -7.9647 | 1.6562E-15 | 1.0086E-14 |
| ENSG00000136379\|ABHD17C | 1201.7595 | -1.0643 | ##### | -7.9641 | 1.6639E-15 | 1.0131E-14 |
| ENSG00000104529\|EEF1D | 44752.3091 | 0.9644 | ##### | 7.9620 | 1.6923E-15 | 1.0302E-14 |
| ENSG00000175772\|LINC01106 | 865.5306 | -1.0989 | ##### | -7.9613 | 1.7022E-15 | 1.0360E-14 |
| ENSG00000150867\|PIP4K2A | 2560.3991 | -0.9923 | ##### | -7.9591 | 1.7328E-15 | 1.0544E-14 |
| ENSG00000144136\|SLC20A1 | 13198.1030 | -0.9573 | ##### | -7.9586 | 1.7394E-15 | 1.0582E-14 |
| ENSG00000170667\|RASA4B | 459.2648 | 1.1606 | ##### | 7.9564 | 1.7713E-15 | 1.0774E-14 |
| ENSG00000164961\|WASHC5 | 5678.5716 | -0.9345 | ##### | -7.9533 | 1.8159E-15 | 1.1043E-14 |
| ENSG00000107201\|RIGI | 474.2361 | -1.1814 | ##### | -7.9527 | 1.8252E-15 | 1.1097E-14 |
| ENSG00000276075 | 78.3806 | -2.2321 | ##### | -7.9525 | 1.8280E-15 | 1.1112E-14 |

| | | | | | | |
|---|---|---|---|---|---|---|
| ENSG00000150459\|SAP18 | 9744.4218 | 0.7281 | ##### | 7.9509 | 1.8514E-15 | 1.1252E-14 |
| ENSG00000290879 | 115.0722 | -1.7586 | ##### | -7.9506 | 1.8554E-15 | 1.1274E-14 |
| ENSG00000175567\|UCP2 | 1663.1872 | -1.0700 | ##### | -7.9488 | 1.8832E-15 | 1.1441E-14 |
| ENSG00000209082\|MT-TL1 | 2071.8622 | -0.9764 | ##### | -7.9458 | 1.9299E-15 | 1.1722E-14 |
| ENSG00000293516\|ZNF271P | 2254.1674 | -0.9919 | ##### | -7.9450 | 1.9425E-15 | 1.1796E-14 |
| ENSG00000123243\|ITIH5 | 126.7336 | 8.5952 | ##### | 7.9432 | 1.9709E-15 | 1.1966E-14 |
| ENSG00000118804\|STBD1 | 519.2067 | -1.3884 | ##### | -7.9420 | 1.9900E-15 | 1.2080E-14 |
| ENSG00000120265\|PCMT1 | 10198.2872 | 0.9611 | ##### | 7.9413 | 2.0007E-15 | 1.2142E-14 |
| ENSG00000064989\|CALCRL | 78.5281 | -3.4145 | ##### | -7.9411 | 2.0034E-15 | 1.2156E-14 |
| ENSG00000176406\|RIMS2 | 259.2708 | 1.5138 | ##### | 7.9398 | 2.0248E-15 | 1.2283E-14 |
| ENSG00000056097\|ZFR | 17596.4916 | 0.8975 | ##### | 7.9370 | 2.0714E-15 | 1.2563E-14 |
| ENSG00000141540\|TTYH2 | 521.9116 | 1.2170 | ##### | 7.9363 | 2.0838E-15 | 1.2636E-14 |
| ENSG00000253837\|LOXL2-AS1 | 50.4317 | 2.7675 | ##### | 7.9362 | 2.0846E-15 | 1.2639E-14 |
| ENSG00000082014\|SMARCD3 | 3683.5423 | -0.7804 | ##### | -7.9352 | 2.1024E-15 | 1.2744E-14 |
| ENSG00000181649\|PHLDA2 | 5203.0019 | 1.0218 | ##### | 7.9349 | 2.1059E-15 | 1.2762E-14 |
| ENSG00000149260\|CAPN5 | 2224.3597 | -0.9818 | ##### | -7.9344 | 2.1157E-15 | 1.2819E-14 |
| ENSG00000152661\|GJA1 | 3936.0959 | 0.9594 | ##### | 7.9317 | 2.1617E-15 | 1.3095E-14 |
| ENSG00000106100\|NOD1 | 925.4265 | -1.0552 | ##### | -7.9311 | 2.1723E-15 | 1.3157E-14 |
| ENSG00000157741\|UBN2 | 1950.0515 | 0.9982 | ##### | 7.9310 | 2.1731E-15 | 1.3159E-14 |
| ENSG00000122707\|RECK | 1569.4923 | 1.0675 | ##### | 7.9295 | 2.2011E-15 | 1.3326E-14 |
| ENSG00000115761\|NOL10 | 7785.2360 | 0.9218 | ##### | 7.9278 | 2.2300E-15 | 1.3498E-14 |
| ENSG00000269210 | 81.9227 | -2.1772 | ##### | -7.9230 | 2.3179E-15 | 1.4027E-14 |
| ENSG00000250616\|YPEL3-DT | 58.1158 | -2.9126 | ##### | -7.9202 | 2.3721E-15 | 1.4352E-14 |
| ENSG00000186318\|BACE1 | 1324.0831 | -1.1976 | ##### | -7.9162 | 2.4488E-15 | 1.4813E-14 |
| ENSG00000159596\|TMEM69 | 2877.1256 | -0.7747 | ##### | -7.9161 | 2.4511E-15 | 1.4824E-14 |
| ENSG00000165512\|ZNF22 | 565.1459 | -1.1518 | ##### | -7.9160 | 2.4537E-15 | 1.4837E-14 |
| ENSG00000181523\|SGSH | 1793.1521 | -1.0166 | ##### | -7.9152 | 2.4686E-15 | 1.4924E-14 |
| ENSG00000107175\|CREB3 | 6539.2191 | 0.8216 | ##### | 7.9136 | 2.4999E-15 | 1.5110E-14 |
| ENSG00000059691\|GATB | 1265.3109 | -1.0537 | ##### | -7.9132 | 2.5082E-15 | 1.5158E-14 |
| ENSG00000128944\|KNSTRN | 3809.5694 | 1.0202 | ##### | 7.9122 | 2.5289E-15 | 1.5280E-14 |
| ENSG00000165476\|REEP3 | 4265.3237 | -1.0035 | ##### | -7.9112 | 2.5494E-15 | 1.5400E-14 |
| ENSG00000214435\|AS3MT | 406.3501 | -5.2567 | ##### | -7.9070 | 2.6373E-15 | 1.5928E-14 |
| ENSG00000141837\|CACNA1A | 1258.0056 | 8.1553 | ##### | 7.9060 | 2.6589E-15 | 1.6055E-14 |
| ENSG00000213741\|RPS29 | 7497.9373 | 1.0951 | ##### | 7.8990 | 2.8109E-15 | 1.6969E-14 |
| ENSG00000172500\|FIBP | 4448.7995 | 0.9895 | ##### | 7.8980 | 2.8349E-15 | 1.7111E-14 |
| ENSG00000136877\|FPGS | 7172.5960 | -0.8868 | ##### | -7.8978 | 2.8383E-15 | 1.7128E-14 |
| ENSG00000260641\|TSPAN5-DT | 115.6855 | 8.5497 | ##### | 7.8977 | 2.8419E-15 | 1.7146E-14 |
| ENSG00000111669\|TPI1 | 91014.6417 | 0.8898 | ##### | 7.8976 | 2.8438E-15 | 1.7154E-14 |
| ENSG00000137877\|SPTBN5 | 403.9209 | -1.2518 | ##### | -7.8974 | 2.8471E-15 | 1.7170E-14 |
| ENSG00000163596\|ICA1L | 261.2173 | -1.4584 | ##### | -7.8973 | 2.8502E-15 | 1.7186E-14 |
| ENSG00000126749\|EMG1 | 18154.5659 | 0.9609 | ##### | 7.8956 | 2.8890E-15 | 1.7416E-14 |
| ENSG00000143093\|STRIP1 | 4714.9719 | 0.9314 | ##### | 7.8907 | 3.0050E-15 | 1.8112E-14 |
| ENSG00000270757\|HSPE1-MOB4 | 50.3619 | 22.7970 | ##### | 7.8885 | 3.0595E-15 | 1.8436E-14 |
| ENSG00000260192\|LINC02240 | 66.2113 | -7.2181 | ##### | -7.8881 | 3.0673E-15 | 1.8480E-14 |
| ENSG00000196422\|PPP1R26 | 2365.7908 | -0.9659 | ##### | -7.8881 | 3.0688E-15 | 1.8485E-14 |
| ENSG00000261359\|PYCARD-AS1 | 107.3543 | -1.9648 | ##### | -7.8870 | 3.0955E-15 | 1.8640E-14 |
| ENSG00000236753\|MKLN1-AS | 818.2634 | -1.0909 | ##### | -7.8870 | 3.0958E-15 | 1.8640E-14 |
| ENSG00000230530\|LIMD1-AS1 | 2059.2144 | -0.9907 | ##### | -7.8848 | 3.1497E-15 | 1.8961E-14 |
| ENSG00000170027\|YWHAG | 34235.8578 | -0.8550 | ##### | -7.8845 | 3.1569E-15 | 1.9000E-14 |
| ENSG00000285612 | 265.6240 | 1.6732 | ##### | 7.8845 | 3.1577E-15 | 1.9001E-14 |
| ENSG00000100883\|SRP54 | 6509.3505 | 0.8419 | ##### | 7.8839 | 3.1722E-15 | 1.9085E-14 |
| ENSG00000168237\|GLYCTK | 362.3342 | 1.3698 | ##### | 7.8806 | 3.2583E-15 | 1.9599E-14 |
| ENSG00000198258\|UBL5 | 9058.3579 | 0.9728 | ##### | 7.8803 | 3.2654E-15 | 1.9638E-14 |
| ENSG00000172590\|MRPL52 | 3476.5622 | 1.0050 | ##### | 7.8779 | 3.3300E-15 | 2.0022E-14 |
| ENSG00000140848\|CPNE2 | 4232.9911 | -0.9672 | ##### | -7.8773 | 3.3451E-15 | 2.0109E-14 |
| ENSG00000108448\|TRIM16L | 193.2217 | -3.8077 | ##### | -7.8761 | 3.3789E-15 | 2.0308E-14 |
| ENSG00000146242\|TPBG | 6187.0539 | 0.9725 | ##### | 7.8742 | 3.4282E-15 | 2.0600E-14 |
| ENSG00000127564\|PKMYT1 | 6365.8959 | 1.0001 | ##### | 7.8742 | 3.4295E-15 | 2.0604E-14 |
| ENSG00000261342 | 338.3558 | 1.4438 | ##### | 7.8740 | 3.4342E-15 | 2.0628E-14 |
| ENSG00000224831\|TMEM183BP | 306.7274 | -1.2950 | ##### | -7.8716 | 3.5010E-15 | 2.1025E-14 |
| ENSG00000204852\|TCTN1 | 1562.1925 | -1.0236 | ##### | -7.8712 | 3.5120E-15 | 2.1087E-14 |
| ENSG00000164548\|TRA2A | 7408.9163 | 0.9785 | ##### | 7.8712 | 3.5131E-15 | 2.1089E-14 |
| ENSG00000182568\|SATB1 | 71.3624 | -2.0531 | ##### | -7.8694 | 3.5644E-15 | 2.1393E-14 |

| | | | | | | |
|---|---|---|---|---|---|---|
| ENSG00000111445\|RFC5 | 4817.2252 | 0.9287 | ##### | 7.8682 | 3.5964E-15 | 2.1581E-14 |
| ENSG00000232453\|LINC02777 | 52.0560 | -2.6697 | ##### | -7.8675 | 3.6169E-15 | 2.1699E-14 |
| ENSG00000065618\|COL17A1 | 120.4151 | -2.2730 | ##### | -7.8645 | 3.7062E-15 | 2.2230E-14 |
| ENSG00000111837\|MAK | 65.0165 | 2.4836 | ##### | 7.8616 | 3.7915E-15 | 2.2738E-14 |
| ENSG00000168807\|SNTB2 | 3561.2687 | -0.9876 | ##### | -7.8612 | 3.8061E-15 | 2.2821E-14 |
| ENSG00000165084\|C8orf34 | 34.1247 | -4.5760 | ##### | -7.8608 | 3.8168E-15 | 2.2880E-14 |
| ENSG00000278817 | 175.0406 | -1.5022 | ##### | -7.8605 | 3.8250E-15 | 2.2924E-14 |
| ENSG00000269190\|FBXO17 | 4465.4936 | -0.8956 | ##### | -7.8591 | 3.8687E-15 | 2.3182E-14 |
| ENSG00000156030\|MIDEAS | 3992.6955 | 0.9514 | ##### | 7.8568 | 3.9423E-15 | 2.3618E-14 |
| ENSG00000198298\|ZNF485 | 174.1656 | -3.9127 | ##### | -7.8563 | 3.9550E-15 | 2.3690E-14 |
| ENSG00000144747\|TMF1 | 3036.1960 | -1.0016 | ##### | -7.8554 | 3.9835E-15 | 2.3855E-14 |
| ENSG00000197444\|OGDHL | 155.9057 | 1.6670 | ##### | 7.8554 | 3.9861E-15 | 2.3866E-14 |
| ENSG00000115896\|PLCL1 | 65.8460 | -2.2683 | ##### | -7.8551 | 3.9956E-15 | 2.3918E-14 |
| ENSG00000105376\|ICAM5 | 1052.4696 | 1.0285 | ##### | 7.8540 | 4.0303E-15 | 2.4121E-14 |
| ENSG00000282304 | 371.3087 | -7.9497 | ##### | -7.8535 | 4.0468E-15 | 2.4215E-14 |
| ENSG00000232638 | 34.3084 | 3.0862 | ##### | 7.8528 | 4.0673E-15 | 2.4333E-14 |
| ENSG00000143434\|SEMA6C | 449.0053 | 1.3063 | ##### | 7.8526 | 4.0758E-15 | 2.4379E-14 |
| ENSG00000111231\|GPN3 | 3896.6267 | 1.0070 | ##### | 7.8518 | 4.1013E-15 | 2.4522E-14 |
| ENSG00000196290\|NIF3L1 | 2326.7604 | 1.0562 | ##### | 7.8518 | 4.1007E-15 | 2.4522E-14 |
| ENSG00000253767\|PCDHGA8 | 118.2969 | -5.3218 | ##### | -7.8516 | 4.1093E-15 | 2.4564E-14 |
| ENSG00000130768\|SMPDL3B | 346.2292 | -1.3853 | ##### | -7.8484 | 4.2128E-15 | 2.5178E-14 |
| ENSG00000173706\|HEG1 | 3534.5287 | -1.0161 | ##### | -7.8482 | 4.2189E-15 | 2.5209E-14 |
| ENSG00000247853 | 161.2307 | 1.6499 | ##### | 7.8474 | 4.2488E-15 | 2.5383E-14 |
| ENSG00000145920\|CPLX2 | 5457.7287 | ####### | ##### | -7.8469 | 4.2638E-15 | 2.5468E-14 |
| ENSG00000038295\|TLL1 | 371.7565 | -1.3488 | ##### | -7.8449 | 4.3321E-15 | 2.5871E-14 |
| ENSG00000115525\|ST3GAL5 | 563.6366 | -1.1971 | ##### | -7.8434 | 4.3860E-15 | 2.6187E-14 |
| ENSG00000103942\|HOMER2 | 905.5557 | -1.1331 | ##### | -7.8383 | 4.5651E-15 | 2.7251E-14 |
| ENSG00000161558\|TMEM143 | 374.6522 | -1.2867 | ##### | -7.8382 | 4.5698E-15 | 2.7274E-14 |
| ENSG00000172007\|RAB33B | 593.7606 | -1.1655 | ##### | -7.8380 | 4.5760E-15 | 2.7305E-14 |
| ENSG00000142089\|IFITM3 | 7253.3164 | -0.9834 | ##### | -7.8372 | 4.6083E-15 | 2.7492E-14 |
| ENSG00000137133\|HINT2 | 2577.4077 | 1.0221 | ##### | 7.8362 | 4.6453E-15 | 2.7708E-14 |
| ENSG00000236204\|LINC01376 | 105.4017 | -1.7239 | ##### | -7.8351 | 4.6836E-15 | 2.7931E-14 |
| ENSG00000079435\|LIPE | 338.3659 | -1.3406 | ##### | -7.8340 | 4.7251E-15 | 2.8172E-14 |
| ENSG00000101294\|HM13 | 14473.5695 | -0.9128 | ##### | -7.8338 | 4.7331E-15 | 2.8214E-14 |
| ENSG00000167554\|ZNF610 | 106.3086 | -2.0382 | ##### | -7.8332 | 4.7542E-15 | 2.8335E-14 |
| ENSG00000163273\|NPPC | 33.1198 | 4.3621 | ##### | 7.8329 | 4.7687E-15 | 2.8415E-14 |
| ENSG00000198677\|SKIC3 | 5259.1378 | -0.9303 | ##### | -7.8327 | 4.7765E-15 | 2.8456E-14 |
| ENSG00000133393\|CEP20 | 6136.5546 | -0.6496 | ##### | -7.8318 | 4.8077E-15 | 2.8636E-14 |
| ENSG00000164978\|NUDT2 | 697.3734 | -1.1021 | ##### | -7.8317 | 4.8143E-15 | 2.8670E-14 |
| ENSG00000150636\|CCDC102B | 238.4473 | -1.4044 | ##### | -7.8315 | 4.8216E-15 | 2.8708E-14 |
| ENSG00000178773\|CPNE7 | 3023.8736 | 1.0037 | ##### | 7.8312 | 4.8315E-15 | 2.8761E-14 |
| ENSG00000151702\|FLI1 | 41.8638 | 3.2456 | ##### | 7.8309 | 4.8423E-15 | 2.8819E-14 |
| ENSG00000293161 | 1547.4708 | -0.9881 | ##### | -7.8294 | 4.9021E-15 | 2.9169E-14 |
| ENSG00000245614\|DDX11-AS1 | 151.9283 | 1.9043 | ##### | 7.8280 | 4.9588E-15 | 2.9501E-14 |
| ENSG00000101146\|RAE1 | 5801.9233 | 0.9477 | ##### | 7.8278 | 4.9660E-15 | 2.9538E-14 |
| ENSG00000248727\|LINC01948 | 170.3497 | -1.9922 | ##### | -7.8254 | 5.0615E-15 | 3.0100E-14 |
| ENSG00000179292\|TMEM151A | 151.9591 | -1.8210 | ##### | -7.8240 | 5.1155E-15 | 3.0415E-14 |
| ENSG00000100604\|CHGA | 1083.5234 | ####### | ##### | -7.8236 | 5.1344E-15 | 3.0521E-14 |
| ENSG00000168461\|RAB31 | 9393.8666 | 0.9365 | ##### | 7.8232 | 5.1492E-15 | 3.0603E-14 |
| ENSG00000125505\|MBOAT7 | 14335.6418 | -0.9304 | ##### | -7.8224 | 5.1842E-15 | 3.0805E-14 |
| ENSG00000100413\|POLR3H | 7589.0469 | 0.9783 | ##### | 7.8216 | 5.2151E-15 | 3.0982E-14 |
| ENSG00000141448\|GATA6 | 335.5548 | -1.4630 | ##### | -7.8199 | 5.2870E-15 | 3.1403E-14 |
| ENSG00000146232\|NFKBIE | 1875.6062 | -1.0027 | ##### | -7.8197 | 5.2933E-15 | 3.1434E-14 |
| ENSG00000145014\|TMEM44 | 1154.4534 | -1.0304 | ##### | -7.8196 | 5.2982E-15 | 3.1458E-14 |
| ENSG00000196436\|NPIPB15 | 27.4228 | -3.1851 | ##### | -7.8193 | 5.3126E-15 | 3.1537E-14 |
| ENSG00000122733\|PHF24 | 59.8558 | 3.2243 | ##### | 7.8185 | 5.3443E-15 | 3.1719E-14 |
| ENSG00000147140\|NONO | 43794.6902 | 0.7133 | ##### | 7.8181 | 5.3618E-15 | 3.1816E-14 |
| ENSG00000184863\|RBM33 | 4902.6324 | 0.9668 | ##### | 7.8163 | 5.4396E-15 | 3.2271E-14 |
| ENSG00000127540\|UQCR11 | 6841.9601 | 0.9434 | ##### | 7.8156 | 5.4722E-15 | 3.2458E-14 |
| ENSG00000179833\|SERTAD2 | 3157.6109 | -0.5861 | ##### | -7.8151 | 5.4911E-15 | 3.2564E-14 |
| ENSG00000069943\|PIGB | 595.7637 | -1.1358 | ##### | -7.8144 | 5.5232E-15 | 3.2748E-14 |
| ENSG00000293159\|POM121L15P | 35.0404 | -2.4190 | ##### | -7.8131 | 5.5785E-15 | 3.3069E-14 |
| ENSG00000189362\|NEMP2 | 1215.9140 | -1.0720 | ##### | -7.8131 | 5.5797E-15 | 3.3070E-14 |

| | | | | | | |
|---|---|---|---|---|---|---|
| ENSG00000273532 | 30.7862 | -2.4027 | ##### | -7.8116 | 5.6476E-15 | 3.3465E-14 |
| ENSG00000100823\|APEX1 | 12586.7545 | 0.8450 | ##### | 7.8112 | 5.6653E-15 | 3.3564E-14 |
| ENSG00000177182\|CLVS1 | 103.1587 | -3.7641 | ##### | -7.8100 | 5.7174E-15 | 3.3865E-14 |
| ENSG00000133983\|COX16 | 6238.9739 | 0.7972 | ##### | 7.8092 | 5.7544E-15 | 3.4078E-14 |
| ENSG00000072135\|PTPN18 | 2357.7890 | -0.9588 | ##### | -7.8089 | 5.7704E-15 | 3.4166E-14 |
| ENSG00000125304\|TM9SF2 | 9176.5074 | -0.9753 | ##### | -7.8083 | 5.7966E-15 | 3.4315E-14 |
| ENSG00000167380\|ZNF226 | 929.2162 | -1.1137 | ##### | -7.8058 | 5.9144E-15 | 3.5005E-14 |
| ENSG00000198839\|ZNF277 | 2075.3255 | -0.9544 | ##### | -7.8051 | 5.9457E-15 | 3.5183E-14 |
| ENSG00000074211\|PPP2R2C | 5472.4461 | -0.9301 | ##### | -7.8049 | 5.9547E-15 | 3.5230E-14 |
| ENSG00000146109\|ABT1 | 2218.9827 | 1.0236 | ##### | 7.8048 | 5.9605E-15 | 3.5257E-14 |
| ENSG00000073711\|PPP2R3A | 619.8899 | -1.0641 | ##### | -7.8041 | 5.9921E-15 | 3.5436E-14 |
| ENSG00000147454\|SLC25A37 | 8914.3699 | 0.9723 | ##### | 7.8017 | 6.1056E-15 | 3.6101E-14 |
| ENSG00000138709\|LARP1B | 1150.0767 | -1.0694 | ##### | -7.8008 | 6.1498E-15 | 3.6355E-14 |
| ENSG00000129810\|SGO1 | 1705.2714 | 1.0207 | ##### | 7.8005 | 6.1684E-15 | 3.6458E-14 |
| ENSG00000142102\|PGGHG | 2638.1693 | 0.9876 | ##### | 7.7999 | 6.1956E-15 | 3.6611E-14 |
| ENSG00000270959\|LPP-AS2 | 169.2806 | -1.8707 | ##### | -7.7996 | 6.2110E-15 | 3.6695E-14 |
| ENSG00000180535\|BHLHA15 | 93.2764 | -3.2145 | ##### | -7.7995 | 6.2149E-15 | 3.6710E-14 |
| ENSG00000163322\|ABRAXAS1 | 1035.3305 | -1.0724 | ##### | -7.7985 | 6.2626E-15 | 3.6985E-14 |
| ENSG00000025039\|RRAGD | 125.3108 | -1.6875 | ##### | -7.7980 | 6.2916E-15 | 3.7149E-14 |
| ENSG00000140650\|PMM2 | 7597.5610 | -0.9533 | ##### | -7.7974 | 6.3210E-15 | 3.7315E-14 |
| ENSG00000198954\|KIFBP | 3327.8179 | -0.9721 | ##### | -7.7968 | 6.3489E-15 | 3.7472E-14 |
| ENSG00000168481\|LGI3 | 25.0073 | -3.8647 | ##### | -7.7960 | 6.3891E-15 | 3.7702E-14 |
| ENSG00000154639\|CXADR | 6576.4290 | 0.9329 | ##### | 7.7956 | 6.4080E-15 | 3.7806E-14 |
| ENSG00000139343\|SNRPF | 7963.1021 | 0.9468 | ##### | 7.7931 | 6.5380E-15 | 3.8566E-14 |
| ENSG00000119685\|TTLL5 | 2000.7285 | 0.9965 | ##### | 7.7930 | 6.5417E-15 | 3.8572E-14 |
| ENSG00000109458\|GAB1 | 359.7239 | -1.2125 | ##### | -7.7930 | 6.5413E-15 | 3.8572E-14 |
| ENSG00000142459\|EVI5L | 963.2078 | -1.0456 | ##### | -7.7894 | 6.7311E-15 | 3.9681E-14 |
| ENSG00000246477 | 55.0977 | -2.4131 | ##### | -7.7889 | 6.7606E-15 | 3.9847E-14 |
| ENSG00000020426\|MNAT1 | 2506.8063 | 0.9839 | ##### | 7.7877 | 6.8219E-15 | 4.0200E-14 |
| ENSG00000138172\|CALHM2 | 840.4567 | 1.0966 | ##### | 7.7853 | 6.9531E-15 | 4.0966E-14 |
| ENSG00000266969\|CEP192-DT | 56.5439 | 2.3001 | ##### | 7.7851 | 6.9634E-15 | 4.1018E-14 |
| ENSG00000272841\|MAP3K4-AS1 | 163.0592 | 1.7491 | ##### | 7.7846 | 6.9939E-15 | 4.1189E-14 |
| ENSG00000168477\|TNXB | 318.4412 | 1.5936 | ##### | 7.7833 | 7.0654E-15 | 4.1603E-14 |
| ENSG00000160013\|PTGIR | 33.0568 | -3.6431 | ##### | -7.7829 | 7.0900E-15 | 4.1739E-14 |
| ENSG00000160360\|GPSM1 | 2244.1938 | -0.9937 | ##### | -7.7826 | 7.1032E-15 | 4.1809E-14 |
| ENSG00000147059\|SPIN2A | 66.8752 | -1.9643 | ##### | -7.7815 | 7.1662E-15 | 4.2171E-14 |
| ENSG00000103494\|RPGRIP1L | 1631.4498 | -0.9739 | ##### | -7.7809 | 7.1989E-15 | 4.2355E-14 |
| ENSG00000163558\|PRKCI | 4640.6178 | -0.9594 | ##### | -7.7779 | 7.3714E-15 | 4.3362E-14 |
| ENSG00000087903\|RFX2 | 973.9754 | 1.0752 | ##### | 7.7773 | 7.4084E-15 | 4.3570E-14 |
| ENSG00000087111\|PIGS | 4161.2982 | -0.9866 | ##### | -7.7763 | 7.4699E-15 | 4.3923E-14 |
| ENSG00000167085\|PHB1 | 35718.6482 | 0.6546 | ##### | 7.7757 | 7.5003E-15 | 4.4094E-14 |
| ENSG00000126803\|HSPA2 | 5675.8117 | -0.9283 | ##### | -7.7740 | 7.6069E-15 | 4.4712E-14 |
| ENSG00000104880\|ARHGEF18 | 6295.8746 | 0.9042 | ##### | 7.7723 | 7.7075E-15 | 4.5294E-14 |
| ENSG00000166669\|ATF7IP2 | 418.7393 | -1.1613 | ##### | -7.7722 | 7.7164E-15 | 4.5337E-14 |
| ENSG00000023318\|ERP44 | 3084.7763 | -0.9348 | ##### | -7.7691 | 7.9065E-15 | 4.6445E-14 |
| ENSG00000160179\|ABCG1 | 182.0181 | 1.6781 | ##### | 7.7678 | 7.9851E-15 | 4.6898E-14 |
| ENSG00000184226\|PCDH9 | 1864.2603 | -1.0002 | ##### | -7.7672 | 8.0236E-15 | 4.7115E-14 |
| ENSG00000114331\|ACAP2 | 4180.4445 | -0.9607 | ##### | -7.7669 | 8.0427E-15 | 4.7218E-14 |
| ENSG00000183513\|COA5 | 1299.7313 | 1.0290 | ##### | 7.7660 | 8.1007E-15 | 4.7548E-14 |
| ENSG00000186376\|ZNF75D | 834.1275 | 1.0949 | ##### | 7.7654 | 8.1380E-15 | 4.7758E-14 |
| ENSG00000185904\|LINC00839 | 243.2427 | -1.5412 | ##### | -7.7625 | 8.3265E-15 | 4.8855E-14 |
| ENSG00000183978\|COA3 | 2540.5187 | -0.9538 | ##### | -7.7608 | 8.4387E-15 | 4.9503E-14 |
| ENSG00000120784\|ZFP30 | 1282.4427 | -1.0861 | ##### | -7.7604 | 8.4673E-15 | 4.9661E-14 |
| ENSG00000153563\|CD8A | 108.6935 | 2.9452 | ##### | 7.7578 | 8.6396E-15 | 5.0662E-14 |
| ENSG00000226251\|LINC02608 | 84.9147 | -8.4942 | ##### | -7.7576 | 8.6535E-15 | 5.0734E-14 |
| ENSG00000180884\|ZNF792 | 114.0410 | -1.7366 | ##### | -7.7561 | 8.7609E-15 | 5.1346E-14 |
| ENSG00000068024\|HDAC4 | 3832.8807 | -0.9486 | ##### | -7.7561 | 8.7614E-15 | 5.1346E-14 |
| ENSG00000179889\|PDXDC1 | 8715.8326 | -0.9263 | ##### | -7.7540 | 8.9055E-15 | 5.2180E-14 |
| ENSG00000172534\|HCFC1 | 14702.6861 | 0.9175 | ##### | 7.7524 | 9.0172E-15 | 5.2824E-14 |
| ENSG00000235530\|TMEM11-DT | 50.8945 | 2.4527 | ##### | 7.7509 | 9.1230E-15 | 5.3434E-14 |
| ENSG00000055609\|KMT2C | 3068.7912 | 1.0331 | ##### | 7.7502 | 9.1715E-15 | 5.3707E-14 |
| ENSG00000247271\|ZBED5-AS1 | 325.6551 | -1.2242 | ##### | -7.7480 | 9.3355E-15 | 5.4657E-14 |
| ENSG00000117114\|ADGRL2 | 6352.6164 | -0.9508 | ##### | -7.7476 | 9.3669E-15 | 5.4830E-14 |

| | | | | | | |
|---|---|---|---|---|---|---|
| ENSG00000155380\|SLC16A1 | 14358.8699 | 0.8396 | ##### | 7.7468 | 9.4254E-15 | 5.5162E-14 |
| ENSG00000077458\|FAM76B | 1071.1451 | 1.0543 | ##### | 7.7460 | 9.4800E-15 | 5.5470E-14 |
| ENSG00000229320\|KRT8P12 | 582.1056 | 1.1528 | ##### | 7.7456 | 9.5129E-15 | 5.5652E-14 |
| ENSG00000133661\|SFTPD | 189.0981 | -4.5267 | ##### | -7.7435 | 9.6685E-15 | 5.6551E-14 |
| ENSG00000157890\|MEGF11 | 196.4165 | 3.5281 | ##### | 7.7432 | 9.6915E-15 | 5.6674E-14 |
| ENSG00000124429\|POF1B | 659.7508 | -7.9953 | ##### | -7.7431 | 9.7045E-15 | 5.6739E-14 |
| ENSG00000265399 | 609.3510 | 1.3287 | ##### | 7.7429 | 9.7143E-15 | 5.6785E-14 |
| ENSG00000141449\|GREB1L | 1324.9702 | -1.0693 | ##### | -7.7416 | 9.8207E-15 | 5.7396E-14 |
| ENSG00000178295\|GEN1 | 2602.4005 | 0.9977 | ##### | 7.7400 | 9.9441E-15 | 5.8106E-14 |
| ENSG00000103510\|KAT8 | 2228.5500 | -0.9990 | ##### | -7.7394 | 9.9895E-15 | 5.8360E-14 |
| ENSG00000204569\|PPP1R10 | 5526.5032 | 0.8765 | ##### | 7.7390 | 1.0017E-14 | 5.8508E-14 |
| ENSG00000188807\|TMEM201 | 3434.3074 | 0.9548 | ##### | 7.7376 | 1.0129E-14 | 5.9149E-14 |
| ENSG00000177455\|CD19 | 97.9024 | -9.0545 | ##### | -7.7365 | 1.0215E-14 | 5.9645E-14 |
| ENSG00000171984\|SHLD1 | 149.4865 | -1.4874 | ##### | -7.7344 | 1.0392E-14 | 6.0664E-14 |
| ENSG00000167460\|TPM4 | 51436.9382 | -0.6024 | ##### | -7.7333 | 1.0482E-14 | 6.1175E-14 |
| ENSG00000160219\|GAB3 | 156.8426 | -1.7644 | ##### | -7.7328 | 1.0520E-14 | 6.1388E-14 |
| ENSG00000141696\|P3H4 | 3927.4582 | -0.9013 | ##### | -7.7327 | 1.0532E-14 | 6.1444E-14 |
| ENSG00000157833\|GAREM2 | 947.7911 | -1.0422 | ##### | -7.7296 | 1.0793E-14 | 6.2954E-14 |
| ENSG00000162604\|TM2D1 | 1827.6703 | -1.0499 | ##### | -7.7282 | 1.0905E-14 | 6.3594E-14 |
| ENSG00000248538\|PPP1R3B-DT | 96.7797 | -2.5367 | ##### | -7.7270 | 1.1008E-14 | 6.4187E-14 |
| ENSG00000127920\|GNG11 | 8661.5229 | 0.9454 | ##### | 7.7265 | 1.1056E-14 | 6.4453E-14 |
| ENSG00000158825\|CDA | 806.6435 | -7.2858 | ##### | -7.7239 | 1.1283E-14 | 6.5762E-14 |
| ENSG00000095951\|HIVEP1 | 1661.6256 | 1.0108 | ##### | 7.7225 | 1.1409E-14 | 6.6486E-14 |
| ENSG00000254986\|DPP3 | 5800.1432 | 0.9227 | ##### | 7.7182 | 1.1797E-14 | 6.8731E-14 |
| ENSG00000167291\|TBC1D16 | 4049.4398 | -0.9368 | ##### | -7.7181 | 1.1806E-14 | 6.8772E-14 |
| ENSG00000155100\|OTUD6B | 2703.0696 | -0.9736 | ##### | -7.7175 | 1.1865E-14 | 6.9101E-14 |
| ENSG00000161958\|FGF11 | 263.3227 | 2.3672 | ##### | 7.7168 | 1.1924E-14 | 6.9433E-14 |
| ENSG00000180015 | 185.9728 | 1.3386 | ##### | 7.7167 | 1.1939E-14 | 6.9503E-14 |
| ENSG00000261799 | 448.9893 | 1.1663 | ##### | 7.7161 | 1.1997E-14 | 6.9827E-14 |
| ENSG00000076641\|PAG1 | 542.9871 | -1.1481 | ##### | -7.7160 | 1.2002E-14 | 6.9845E-14 |
| ENSG00000130997\|POLN | 82.8284 | 2.5216 | ##### | 7.7155 | 1.2048E-14 | 7.0096E-14 |
| ENSG00000130517\|PGPEP1 | 845.3452 | -1.0841 | ##### | -7.7116 | 1.2425E-14 | 7.2276E-14 |
| ENSG00000293105 | 74.5344 | -2.8038 | ##### | -7.7115 | 1.2430E-14 | 7.2292E-14 |
| ENSG00000111142\|METAP2 | 16919.9431 | 0.9071 | ##### | 7.7111 | 1.2478E-14 | 7.2560E-14 |
| ENSG00000087460\|GNAS | 82559.7024 | 0.8150 | ##### | 7.7095 | 1.2630E-14 | 7.3427E-14 |
| ENSG00000134769\|DTNA | 211.5818 | 2.1924 | ##### | 7.7082 | 1.2765E-14 | 7.4198E-14 |
| ENSG00000170608\|FOXA3 | 101.0808 | -2.3951 | ##### | -7.7061 | 1.2976E-14 | 7.5406E-14 |
| ENSG00000196954\|CASP4 | 640.7541 | -1.2985 | ##### | -7.7054 | 1.3046E-14 | 7.5799E-14 |
| ENSG00000265688\|MILIP | 591.4376 | -1.0677 | ##### | -7.7049 | 1.3093E-14 | 7.6057E-14 |
| ENSG00000275954\|TBC1D3F | 55.2726 | 2.4966 | ##### | 7.7045 | 1.3140E-14 | 7.6318E-14 |
| ENSG00000062598\|ELMO2 | 4674.4749 | 0.9293 | ##### | 7.7036 | 1.3233E-14 | 7.6846E-14 |
| ENSG00000089775\|ZBTB25 | 1958.7080 | 0.9340 | ##### | 7.7026 | 1.3333E-14 | 7.7406E-14 |
| ENSG00000291186 | 1790.3066 | 1.0017 | ##### | 7.7023 | 1.3365E-14 | 7.7580E-14 |
| ENSG00000130775\|THEMIS2 | 195.8136 | 1.5468 | ##### | 7.7014 | 1.3463E-14 | 7.8134E-14 |
| ENSG00000279520 | 60.4659 | 2.1531 | ##### | 7.7010 | 1.3503E-14 | 7.8350E-14 |
| ENSG00000083814\|ZNF671 | 86.7149 | 2.0183 | ##### | 7.7007 | 1.3533E-14 | 7.8510E-14 |
| ENSG00000164331\|ANKRA2 | 395.5289 | -1.2721 | ##### | -7.6997 | 1.3642E-14 | 7.9126E-14 |
| ENSG00000137878\|GCOM1 | 63.4910 | -3.9420 | ##### | -7.6988 | 1.3732E-14 | 7.9630E-14 |
| ENSG00000006634\|DBF4 | 3405.1981 | 0.9549 | ##### | 7.6939 | 1.4276E-14 | 8.2769E-14 |
| ENSG00000188917\|TRMT2B | 2369.5776 | 1.0187 | ##### | 7.6933 | 1.4339E-14 | 8.3119E-14 |
| ENSG00000158402\|CDC25C | 766.0855 | 1.1158 | ##### | 7.6928 | 1.4399E-14 | 8.3450E-14 |
| ENSG00000162639\|HENMT1 | 3107.0472 | 0.9628 | ##### | 7.6925 | 1.4431E-14 | 8.3619E-14 |
| ENSG00000075856\|SART3 | 12771.8633 | 0.9140 | ##### | 7.6911 | 1.4585E-14 | 8.4494E-14 |
| ENSG00000135677\|GNS | 14673.4521 | -0.9162 | ##### | -7.6904 | 1.4662E-14 | 8.4928E-14 |
| ENSG00000177337\|DLGAP1-AS1 | 1047.5205 | 1.1026 | ##### | 7.6886 | 1.4871E-14 | 8.6117E-14 |
| ENSG00000197632\|SERPINB2 | 5587.9463 | 13.7644 | ##### | 7.6876 | 1.4990E-14 | 8.6794E-14 |
| ENSG00000134901\|POGLUT2 | 659.1091 | -1.0920 | ##### | -7.6874 | 1.5017E-14 | 8.6930E-14 |
| ENSG00000128709\|HOXD9 | 594.2539 | 1.1085 | ##### | 7.6867 | 1.5092E-14 | 8.7351E-14 |
| ENSG00000145996\|CDKAL1 | 1283.0862 | 1.1119 | ##### | 7.6849 | 1.5313E-14 | 8.8614E-14 |
| ENSG00000126391\|FRMD8 | 3793.4759 | 0.9824 | ##### | 7.6843 | 1.5379E-14 | 8.8978E-14 |
| ENSG00000136378\|ADAMTS7 | 857.6865 | 1.0529 | ##### | 7.6842 | 1.5398E-14 | 8.9071E-14 |
| ENSG00000287528 | 34.7231 | 5.1740 | ##### | 7.6833 | 1.5502E-14 | 8.9653E-14 |
| ENSG00000077514\|POLD3 | 3471.2788 | 0.7862 | ##### | 7.6819 | 1.5677E-14 | 9.0650E-14 |

| | | | | | | |
|---|---|---|---|---|---|---|
| ENSG00000146038\|DCDC2 | 137.2795 | -3.8610 | ##### | -7.6816 | 1.5713E-14 | 9.0839E-14 |
| ENSG00000168826\|ZBTB49 | 340.1236 | 1.3281 | ##### | 7.6805 | 1.5847E-14 | 9.1594E-14 |
| ENSG00000180787\|ZFP3 | 917.9294 | ####### | ##### | -7.6801 | 1.5901E-14 | 9.1889E-14 |
| ENSG00000260025\|CRIM1-DT | 464.5447 | -1.1340 | ##### | -7.6799 | 1.5927E-14 | 9.2023E-14 |
| ENSG00000070061\|ELP1 | 6537.9866 | -0.9065 | ##### | -7.6779 | 1.6171E-14 | 9.3411E-14 |
| ENSG00000225465\|RFPL1S | 324.3628 | -1.2862 | ##### | -7.6778 | 1.6186E-14 | 9.3483E-14 |
| ENSG00000177663\|IL17RA | 1738.3362 | -1.0494 | ##### | -7.6761 | 1.6401E-14 | 9.4706E-14 |
| ENSG00000249715\|FER1L5 | 24.5397 | 4.4552 | ##### | 7.6745 | 1.6604E-14 | 9.5858E-14 |
| ENSG00000117569\|PTBP2 | 707.1871 | -1.0869 | ##### | -7.6726 | 1.6851E-14 | 9.7264E-14 |
| ENSG00000092295\|TGM1 | 177.8282 | 1.6389 | ##### | 7.6725 | 1.6866E-14 | 9.7335E-14 |
| ENSG00000254307 | 65.3984 | -1.7979 | ##### | -7.6716 | 1.6992E-14 | 9.8041E-14 |
| ENSG00000002549\|LAP3 | 5201.7760 | 0.6416 | ##### | 7.6708 | 1.7092E-14 | 9.8599E-14 |
| ENSG00000134259\|NGF | 280.6771 | 1.6575 | ##### | 7.6675 | 1.7542E-14 | 1.0118E-13 |
| ENSG00000110871\|COQ5 | 2325.7355 | -0.9762 | ##### | -7.6663 | 1.7698E-14 | 1.0206E-13 |
| ENSG00000152942\|RAD17 | 2489.0960 | 0.8912 | ##### | 7.6645 | 1.7947E-14 | 1.0347E-13 |
| ENSG00000236536 | 92.6249 | 8.1773 | ##### | 7.6629 | 1.8178E-14 | 1.0478E-13 |
| ENSG00000277196 | 60.9020 | -2.4810 | ##### | -7.6629 | 1.8183E-14 | 1.0479E-13 |
| ENSG00000148384\|INPP5E | 1310.4633 | -0.9600 | ##### | -7.6597 | 1.8643E-14 | 1.0742E-13 |
| ENSG00000172037\|LAMB2 | 15867.4426 | -0.9021 | ##### | -7.6596 | 1.8657E-14 | 1.0749E-13 |
| ENSG00000125611\|CHCHD5 | 1934.6809 | -0.9501 | ##### | -7.6555 | 1.9253E-14 | 1.1089E-13 |
| ENSG00000147874\|HAUS6 | 3140.6561 | 0.9463 | ##### | 7.6524 | 1.9729E-14 | 1.1362E-13 |
| ENSG00000172336\|POP7 | 3473.0484 | 0.9312 | ##### | 7.6521 | 1.9780E-14 | 1.1389E-13 |
| ENSG00000179941\|BBS10 | 1686.1808 | -0.9868 | ##### | -7.6518 | 1.9822E-14 | 1.1411E-13 |
| ENSG00000144227\|NXPH2 | 99.4118 | 6.3177 | ##### | 7.6514 | 1.9874E-14 | 1.1438E-13 |
| ENSG00000180304\|OAZ2 | 5112.5379 | 0.9425 | ##### | 7.6506 | 1.9999E-14 | 1.1508E-13 |
| ENSG00000132305\|IMMT | 14138.3341 | 0.5875 | ##### | 7.6506 | 2.0007E-14 | 1.1510E-13 |
| ENSG00000267774 | 93.7671 | 8.4648 | ##### | 7.6505 | 2.0026E-14 | 1.1517E-13 |
| ENSG00000170482\|SLC23A1 | 314.0221 | -6.8119 | ##### | -7.6505 | 2.0026E-14 | 1.1517E-13 |
| ENSG00000160877\|NACC1 | 17854.5299 | 0.7540 | ##### | 7.6487 | 2.0295E-14 | 1.1669E-13 |
| ENSG00000188219\|POTEE | 45.4031 | 2.2658 | ##### | 7.6485 | 2.0330E-14 | 1.1687E-13 |
| ENSG00000267815 | 34.0608 | -2.7528 | ##### | -7.6472 | 2.0533E-14 | 1.1802E-13 |
| ENSG00000254681\|PKD1P5 | 2563.0468 | 1.0013 | ##### | 7.6457 | 2.0786E-14 | 1.1945E-13 |
| ENSG00000285043 | 8411.6424 | -2.6543 | ##### | -7.6451 | 2.0872E-14 | 1.1992E-13 |
| ENSG00000131378\|RFTN1 | 397.4889 | 1.2351 | ##### | 7.6445 | 2.0981E-14 | 1.2052E-13 |
| ENSG00000271321\|CTAGE6 | 32.0163 | -2.5661 | ##### | -7.6444 | 2.0999E-14 | 1.2060E-13 |
| ENSG00000255857\|PXN-AS1 | 9574.1280 | 1.0379 | ##### | 7.6427 | 2.1271E-14 | 1.2214E-13 |
| ENSG00000103978\|TMEM87A | 5384.6112 | -0.7312 | ##### | -7.6423 | 2.1334E-14 | 1.2248E-13 |
| ENSG00000286950 | 299.8875 | 1.2265 | ##### | 7.6403 | 2.1678E-14 | 1.2443E-13 |
| ENSG00000182621\|PLCB1 | 1729.3015 | -1.0024 | ##### | -7.6402 | 2.1690E-14 | 1.2447E-13 |
| ENSG00000163808\|KIF15 | 1928.5318 | 0.9488 | ##### | 7.6382 | 2.2035E-14 | 1.2643E-13 |
| ENSG00000267978\|MAGEA9B | 324.7208 | -9.3264 | ##### | -7.6375 | 2.2146E-14 | 1.2704E-13 |
| ENSG00000136856\|SLC2A8 | 2596.1387 | 0.9898 | ##### | 7.6362 | 2.2380E-14 | 1.2836E-13 |
| ENSG00000197324\|LRP10 | 14206.9034 | -0.5465 | ##### | -7.6359 | 2.2426E-14 | 1.2860E-13 |
| ENSG00000169635\|HIC2 | 871.9115 | 1.0707 | ##### | 7.6357 | 2.2464E-14 | 1.2879E-13 |
| ENSG00000126003\|PLAGL2 | 3141.5467 | 0.9684 | ##### | 7.6346 | 2.2651E-14 | 1.2984E-13 |
| ENSG00000179051\|RCC2 | 18984.8852 | -0.7039 | ##### | -7.6345 | 2.2676E-14 | 1.2996E-13 |
| ENSG00000156469\|MTERF3 | 1772.6139 | -0.9607 | ##### | -7.6344 | 2.2695E-14 | 1.3005E-13 |
| ENSG00000139746\|RBM26 | 4368.9740 | 0.6596 | ##### | 7.6324 | 2.3041E-14 | 1.3200E-13 |
| ENSG00000171931\|FBXW10 | 87.9446 | -4.2724 | ##### | -7.6319 | 2.3129E-14 | 1.3248E-13 |
| ENSG00000131725\|WDR44 | 2354.5531 | 0.9763 | ##### | 7.6312 | 2.3249E-14 | 1.3314E-13 |
| ENSG00000114346\|ECT2 | 8343.6604 | -0.9274 | ##### | -7.6291 | 2.3646E-14 | 1.3539E-13 |
| ENSG00000141441\|GAREM1 | 1205.4707 | -1.0246 | ##### | -7.6268 | 2.4068E-14 | 1.3778E-13 |
| ENSG00000152413\|HOMER1 | 2013.0009 | 0.9350 | ##### | 7.6267 | 2.4075E-14 | 1.3779E-13 |
| ENSG00000112078\|KCTD20 | 8884.9319 | -0.8789 | ##### | -7.6259 | 2.4228E-14 | 1.3864E-13 |
| ENSG00000166272\|WBP1L | 3289.9143 | -0.9484 | ##### | -7.6251 | 2.4393E-14 | 1.3956E-13 |
| ENSG00000186603\|HPDL | 1003.4962 | -1.0261 | ##### | -7.6240 | 2.4585E-14 | 1.4063E-13 |
| ENSG00000159579\|RSPRY1 | 3863.7062 | -0.9475 | ##### | -7.6225 | 2.4878E-14 | 1.4228E-13 |
| ENSG00000115486\|GGCX | 6875.1933 | 0.9101 | ##### | 7.6207 | 2.5239E-14 | 1.4431E-13 |
| ENSG00000101343\|CRNKL1 | 2890.7188 | 0.8972 | ##### | 7.6201 | 2.5356E-14 | 1.4496E-13 |
| ENSG00000073670\|ADAM11 | 336.8853 | 1.3032 | ##### | 7.6193 | 2.5506E-14 | 1.4579E-13 |
| ENSG00000137274\|BPHL | 909.7269 | -1.0252 | ##### | -7.6178 | 2.5806E-14 | 1.4748E-13 |
| ENSG00000138035\|PNPT1 | 11354.5874 | 0.9210 | ##### | 7.6160 | 2.6169E-14 | 1.4952E-13 |
| ENSG00000123329\|ARHGAP9 | 51.8183 | 2.3604 | ##### | 7.6154 | 2.6293E-14 | 1.5020E-13 |

| | | | | | | |
|---|---|---|---|---|---|---|
| ENSG00000109846\|CRYAB | 267.3538 | -7.0816 | ##### | -7.6144 | 2.6483E-14 | 1.5126E-13 |
| ENSG00000169402\|RSPH10B2 | 88.0544 | -1.9873 | ##### | -7.6134 | 2.6704E-14 | 1.5249E-13 |
| ENSG00000258512\|LINC00239 | 82.6200 | -8.8101 | ##### | -7.6123 | 2.6922E-14 | 1.5367E-13 |
| ENSG00000138674\|SEC31A | 10303.8011 | -0.7451 | ##### | -7.6123 | 2.6920E-14 | 1.5367E-13 |
| ENSG00000214456\|PLIN5 | 81.0435 | -8.7367 | ##### | -7.6121 | 2.6978E-14 | 1.5397E-13 |
| ENSG00000204435\|CSNK2B | 11255.7099 | 0.9715 | ##### | 7.6118 | 2.7034E-14 | 1.5425E-13 |
| ENSG00000085511\|MAP3K4 | 2428.2767 | 0.9571 | ##### | 7.6102 | 2.7374E-14 | 1.5617E-13 |
| ENSG00000269318 | 68.7658 | 2.0857 | ##### | 7.6101 | 2.7380E-14 | 1.5617E-13 |
| ENSG00000259529 | 2546.9851 | 1.4456 | ##### | 7.6096 | 2.7503E-14 | 1.5684E-13 |
| ENSG00000143867\|OSR1 | 544.3409 | 1.3494 | ##### | 7.6087 | 2.7697E-14 | 1.5792E-13 |
| ENSG00000230870\|FBXW11P1 | 39.4537 | -3.6491 | ##### | -7.6074 | 2.7961E-14 | 1.5939E-13 |
| ENSG00000246889\|CTTN-DT | 179.7709 | 1.8157 | ##### | 7.6069 | 2.8076E-14 | 1.6002E-13 |
| ENSG00000120457\|KCNJ5 | 101.7538 | 8.1432 | ##### | 7.6067 | 2.8118E-14 | 1.6022E-13 |
| ENSG00000147065\|MSN | 41172.3734 | -0.7226 | ##### | -7.6051 | 2.8467E-14 | 1.6218E-13 |
| ENSG00000167654\|ATCAY | 870.7988 | 6.6868 | ##### | 7.6049 | 2.8511E-14 | 1.6240E-13 |
| ENSG00000289347 | 60.3379 | 2.9926 | ##### | 7.6044 | 2.8629E-14 | 1.6304E-13 |
| ENSG00000173040\|EVC2 | 742.3040 | 1.1503 | ##### | 7.6013 | 2.9311E-14 | 1.6690E-13 |
| ENSG00000099800\|TIMM13 | 6570.9032 | 0.9210 | ##### | 7.6010 | 2.9394E-14 | 1.6733E-13 |
| ENSG00000100284\|TOM1 | 1834.5875 | -0.9592 | ##### | -7.6006 | 2.9471E-14 | 1.6775E-13 |
| ENSG00000119396\|RAB14 | 7356.4361 | -0.9138 | ##### | -7.6001 | 2.9579E-14 | 1.6833E-13 |
| ENSG00000156206\|CFAP161 | 65.3924 | 8.3984 | ##### | 7.5986 | 2.9933E-14 | 1.7031E-13 |
| ENSG00000111144\|LTA4H | 11208.6636 | 0.9361 | ##### | 7.5985 | 2.9963E-14 | 1.7045E-13 |
| ENSG00000158528\|PPP1R9A | 1193.5624 | -0.9917 | ##### | -7.5982 | 3.0020E-14 | 1.7074E-13 |
| ENSG00000127948\|POR | 9677.6874 | -0.8996 | ##### | -7.5980 | 3.0077E-14 | 1.7103E-13 |
| ENSG00000137561\|TTPA | 226.1013 | -1.5168 | ##### | -7.5975 | 3.0180E-14 | 1.7158E-13 |
| ENSG00000177600\|RPLP2 | 28213.5960 | 0.9435 | ##### | 7.5955 | 3.0666E-14 | 1.7431E-13 |
| ENSG00000286623 | 142.2424 | 1.6833 | ##### | 7.5951 | 3.0744E-14 | 1.7472E-13 |
| ENSG00000198276\|UCKL1 | 3514.8280 | 0.9762 | ##### | 7.5909 | 3.1771E-14 | 1.8053E-13 |
| ENSG00000156958\|GALK2 | 1441.9103 | -0.9622 | ##### | -7.5873 | 3.2658E-14 | 1.8553E-13 |
| ENSG00000189120\|SP6 | 542.8909 | -1.1679 | ##### | -7.5845 | 3.3368E-14 | 1.8953E-13 |
| ENSG00000146376\|ARHGAP18 | 767.3584 | -1.0182 | ##### | -7.5842 | 3.3444E-14 | 1.8992E-13 |
| ENSG00000154122\|ANKH | 1790.3526 | -0.9389 | ##### | -7.5814 | 3.4196E-14 | 1.9416E-13 |
| ENSG00000198455\|ZXDB | 1103.3368 | -1.0649 | ##### | -7.5797 | 3.4644E-14 | 1.9666E-13 |
| ENSG00000104321\|TRPA1 | 476.4661 | -1.2177 | ##### | -7.5761 | 3.5615E-14 | 2.0214E-13 |
| ENSG00000135437\|RDH5 | 72.4082 | -2.0013 | ##### | -7.5742 | 3.6122E-14 | 2.0497E-13 |
| ENSG00000135048\|CEMIP2 | 12335.3848 | -0.9063 | ##### | -7.5740 | 3.6189E-14 | 2.0532E-13 |
| ENSG00000099958\|DERL3 | 39.6407 | -3.1512 | ##### | -7.5730 | 3.6458E-14 | 2.0680E-13 |
| ENSG00000160688\|FLAD1 | 6792.8325 | 0.9180 | ##### | 7.5729 | 3.6496E-14 | 2.0698E-13 |
| ENSG00000185437\|SH3BGR | 173.7248 | -1.8220 | ##### | -7.5690 | 3.7607E-14 | 2.1324E-13 |
| ENSG00000291120 | 182.0172 | 1.5187 | ##### | 7.5681 | 3.7878E-14 | 2.1474E-13 |
| ENSG00000136935\|GOLGA1 | 2439.2161 | -0.9374 | ##### | -7.5669 | 3.8215E-14 | 2.1660E-13 |
| ENSG00000103066\|PLA2G15 | 1750.3448 | -1.0228 | ##### | -7.5669 | 3.8232E-14 | 2.1666E-13 |
| ENSG00000143437\|ARNT | 5658.8033 | 0.9295 | ##### | 7.5665 | 3.8352E-14 | 2.1730E-13 |
| ENSG00000170464\|DNAJC18 | 538.1149 | 1.0716 | ##### | 7.5610 | 3.9984E-14 | 2.2650E-13 |
| ENSG00000163734\|CXCL3 | 245.1380 | -1.4774 | ##### | -7.5602 | 4.0231E-14 | 2.2786E-13 |
| ENSG00000290762 | 116.2498 | -1.7057 | ##### | -7.5560 | 4.1563E-14 | 2.3536E-13 |
| ENSG00000175274\|TP53I11 | 2571.7776 | -0.9860 | ##### | -7.5507 | 4.3296E-14 | 2.4512E-13 |
| ENSG00000176809\|LRRC37A3 | 689.5606 | -1.1333 | ##### | -7.5495 | 4.3683E-14 | 2.4727E-13 |
| ENSG00000265479\|DTX2P1-UPK3BP1-PMS2P11 | 545.6444 | -1.0969 | ##### | -7.5494 | 4.3742E-14 | 2.4755E-13 |
| ENSG00000134690\|CDCA8 | 6004.6900 | 0.9541 | ##### | 7.5493 | 4.3753E-14 | 2.4757E-13 |
| ENSG00000270276\|H4C15 | 1318.4436 | 0.9962 | ##### | 7.5436 | 4.5725E-14 | 2.5868E-13 |
| ENSG00000171453\|POLR1C | 5468.5414 | 0.8277 | ##### | 7.5434 | 4.5773E-14 | 2.5890E-13 |
| ENSG00000162735\|PEX19 | 8336.3686 | 0.8477 | ##### | 7.5429 | 4.5955E-14 | 2.5988E-13 |
| ENSG00000169129\|AFAP1L2 | 1060.9226 | -0.9403 | ##### | -7.5427 | 4.6033E-14 | 2.6027E-13 |
| ENSG00000274611\|TBC1D3 | 220.6891 | 1.3505 | ##### | 7.5426 | 4.6077E-14 | 2.6047E-13 |
| ENSG00000204642\|HLA-F | 519.8006 | -1.5012 | ##### | -7.5426 | 4.6084E-14 | 2.6047E-13 |
| ENSG00000166821\|PEX11A | 397.7841 | -1.2103 | ##### | -7.5418 | 4.6362E-14 | 2.6198E-13 |
| ENSG00000289981 | 48.5438 | 3.2957 | ##### | 7.5415 | 4.6443E-14 | 2.6240E-13 |
| ENSG00000145390\|USP53 | 1917.4187 | -0.9972 | ##### | -7.5397 | 4.7092E-14 | 2.6601E-13 |
| ENSG00000121454\|LHX4 | 218.4424 | 1.4632 | ##### | 7.5371 | 4.8061E-14 | 2.7143E-13 |
| ENSG00000243137\|PSG4 | 77.7546 | -3.2213 | ##### | -7.5359 | 4.8513E-14 | 2.7393E-13 |
| ENSG00000165118\|QNG1 | 2434.1911 | -0.9713 | ##### | -7.5354 | 4.8689E-14 | 2.7487E-13 |
| ENSG00000158669\|GPAT4 | 10824.8094 | 0.8847 | ##### | 7.5342 | 4.9135E-14 | 2.7734E-13 |

| | | | | | | |
|---|---|---|---|---|---|---|
| ENSG00000243811\|APOBEC3D | 86.0649 | 4.2059 | ##### | 7.5316 | 5.0104E-14 | 2.8276E-13 |
| ENSG00000102882\|MAPK3 | 2543.0051 | -0.9397 | ##### | -7.5309 | 5.0376E-14 | 2.8424E-13 |
| ENSG00000233476\|EEF1A1P6 | 3820.1126 | 0.9837 | ##### | 7.5273 | 5.1783E-14 | 2.9212E-13 |
| ENSG00000228315\|GUSBP11 | 824.1563 | 1.0996 | ##### | 7.5267 | 5.2032E-14 | 2.9347E-13 |
| ENSG00000135316\|SYNCRIP | 23982.6850 | 0.6214 | ##### | 7.5264 | 5.2166E-14 | 2.9417E-13 |
| ENSG00000225400\|RAB28P5 | 36.6715 | 2.7199 | ##### | 7.5259 | 5.2341E-14 | 2.9510E-13 |
| ENSG00000129028\|THAP10 | 614.7512 | 1.1769 | ##### | 7.5251 | 5.2661E-14 | 2.9685E-13 |
| ENSG00000247134 | 66.4619 | -2.8608 | ##### | -7.5244 | 5.2945E-14 | 2.9840E-13 |
| ENSG00000203896\|LIME1 | 958.6267 | 1.0778 | ##### | 7.5219 | 5.3973E-14 | 3.0413E-13 |
| ENSG00000058668\|ATP2B4 | 6275.4372 | -0.9154 | ##### | -7.5180 | 5.5629E-14 | 3.1340E-13 |
| ENSG00000169255\|B3GALNT1 | 921.4446 | -1.0199 | ##### | -7.5172 | 5.5959E-14 | 3.1521E-13 |
| ENSG00000071626\|DAZAP1 | 17627.9969 | 0.7222 | ##### | 7.5147 | 5.7025E-14 | 3.2115E-13 |
| ENSG00000223797\|ENTPD3-AS1 | 84.6604 | -2.3343 | ##### | -7.5133 | 5.7634E-14 | 3.2452E-13 |
| ENSG00000138613\|APH1B | 691.9026 | -1.1157 | ##### | -7.5124 | 5.8031E-14 | 3.2669E-13 |
| ENSG00000072571\|HMMR | 6080.6320 | 0.9214 | ##### | 7.5121 | 5.8200E-14 | 3.2758E-13 |
| ENSG00000287865 | 86.1400 | 7.9740 | ##### | 7.5115 | 5.8445E-14 | 3.2890E-13 |
| ENSG00000143126\|CELSR2 | 1199.3314 | -1.0567 | ##### | -7.5111 | 5.8636E-14 | 3.2991E-13 |
| ENSG00000155329\|ZCCHC10 | 1353.2727 | 1.0017 | ##### | 7.5073 | 6.0362E-14 | 3.3956E-13 |
| ENSG00000180592\|SKIDA1 | 396.5017 | 1.2521 | ##### | 7.5071 | 6.0449E-14 | 3.3998E-13 |
| ENSG00000117748\|RPA2 | 5437.4612 | 0.9314 | ##### | 7.5070 | 6.0512E-14 | 3.4027E-13 |
| ENSG00000164128\|NPY1R | 371.4820 | -6.8324 | ##### | -7.5062 | 6.0877E-14 | 3.4226E-13 |
| ENSG00000188647\|PTAR1 | 3701.8835 | -0.9367 | ##### | -7.5054 | 6.1244E-14 | 3.4426E-13 |
| ENSG00000287263 | 235.4774 | -1.3572 | ##### | -7.5047 | 6.1581E-14 | 3.4609E-13 |
| ENSG00000109436\|TBC1D9 | 2595.0190 | -0.9583 | ##### | -7.5044 | 6.1715E-14 | 3.4678E-13 |
| ENSG00000256928 | 238.4776 | 1.4269 | ##### | 7.5033 | 6.2251E-14 | 3.4972E-13 |
| ENSG00000168303\|MPLKIP | 2618.5585 | -0.9590 | ##### | -7.5011 | 6.3280E-14 | 3.5544E-13 |
| ENSG00000109220\|CHIC2 | 884.6170 | 1.1351 | ##### | 7.5008 | 6.3453E-14 | 3.5634E-13 |
| ENSG00000287496 | 66.9005 | -2.0681 | ##### | -7.5007 | 6.3482E-14 | 3.5644E-13 |
| ENSG00000168439\|STIP1 | 25987.8443 | 0.6686 | ##### | 7.4994 | 6.4087E-14 | 3.5976E-13 |
| ENSG00000121022\|COPS5 | 6237.0348 | -0.9312 | ##### | -7.4962 | 6.5681E-14 | 3.6864E-13 |
| ENSG00000198774\|RASSF9 | 489.6844 | -6.5271 | ##### | -7.4959 | 6.5843E-14 | 3.6948E-13 |
| ENSG00000136891\|TEX10 | 3338.2589 | 0.9444 | ##### | 7.4948 | 6.6416E-14 | 3.7263E-13 |
| ENSG00000049249\|TNFRSF9 | 71.1486 | -3.2086 | ##### | -7.4944 | 6.6602E-14 | 3.7360E-13 |
| ENSG00000276272 | 60.8654 | 2.4567 | ##### | 7.4943 | 6.6628E-14 | 3.7368E-13 |
| ENSG00000236993\|GAPDHP21 | 22.2965 | -3.9302 | ##### | -7.4937 | 6.6964E-14 | 3.7543E-13 |
| ENSG00000109189\|USP46 | 3135.1490 | 0.9253 | ##### | 7.4937 | 6.6965E-14 | 3.7543E-13 |
| ENSG00000196455\|PIK3R4 | 3999.8918 | 0.8912 | ##### | 7.4921 | 6.7799E-14 | 3.8003E-13 |
| ENSG00000110921\|MVK | 1489.7867 | -0.9742 | ##### | -7.4916 | 6.8047E-14 | 3.8135E-13 |
| ENSG00000179241\|LDLRAD3 | 2911.9904 | 0.7016 | ##### | 7.4910 | 6.8345E-14 | 3.8295E-13 |
| ENSG00000113924\|HGD | 975.2366 | -9.1072 | ##### | -7.4899 | 6.8925E-14 | 3.8613E-13 |
| ENSG00000261616\|TTC23-AS1 | 56.7374 | 2.6328 | ##### | 7.4888 | 6.9504E-14 | 3.8930E-13 |
| ENSG00000084444\|FAM234B | 1074.9632 | 0.9754 | ##### | 7.4880 | 6.9919E-14 | 3.9155E-13 |
| ENSG00000112290\|WASF1 | 2847.6670 | 0.9801 | ##### | 7.4878 | 7.0063E-14 | 3.9228E-13 |
| ENSG00000100519\|PSMC6 | 8643.9613 | 0.6869 | ##### | 7.4856 | 7.1204E-14 | 3.9860E-13 |
| ENSG00000105364\|MRPL4 | 8595.4794 | 0.9266 | ##### | 7.4854 | 7.1330E-14 | 3.9923E-13 |
| ENSG00000132522\|GPS2 | 5468.9228 | 0.9078 | ##### | 7.4847 | 7.1691E-14 | 4.0117E-13 |
| ENSG00000186529\|CYP4F3 | 11832.2936 | ####### | ##### | -7.4846 | 7.1754E-14 | 4.0145E-13 |
| ENSG00000282057 | 209.1476 | 1.7290 | ##### | 7.4835 | 7.2347E-14 | 4.0469E-13 |
| ENSG00000272325\|NUDT3 | 5056.2692 | -0.8841 | ##### | -7.4834 | 7.2420E-14 | 4.0502E-13 |
| ENSG00000179010\|MRFAP1 | 28905.2889 | 0.7715 | ##### | 7.4825 | 7.2925E-14 | 4.0777E-13 |
| ENSG00000176410\|DNAJC30 | 1065.1410 | -0.9721 | ##### | -7.4800 | 7.4318E-14 | 4.1548E-13 |
| ENSG00000113013\|HSPA9 | 46638.0860 | 0.7543 | ##### | 7.4800 | 7.4342E-14 | 4.1554E-13 |
| ENSG00000023445\|BIRC3 | 17275.5036 | -0.9082 | ##### | -7.4759 | 7.6690E-14 | 4.2858E-13 |
| ENSG00000180875\|GREM2 | 93.9642 | 1.5882 | ##### | 7.4749 | 7.7234E-14 | 4.3154E-13 |
| ENSG00000155368\|DBI | 7277.7703 | -0.8815 | ##### | -7.4744 | 7.7538E-14 | 4.3316E-13 |
| ENSG00000217128\|FNIP1 | 1844.7421 | 0.9321 | ##### | 7.4721 | 7.8904E-14 | 4.4071E-13 |
| ENSG00000285188\|PDE4C | 80.6205 | 2.2741 | ##### | 7.4716 | 7.9247E-14 | 4.4254E-13 |
| ENSG00000182712\|CMC4 | 847.2206 | 1.0477 | ##### | 7.4705 | 7.9883E-14 | 4.4601E-13 |
| ENSG00000100077\|GRK3 | 2472.4128 | 0.9027 | ##### | 7.4699 | 8.0238E-14 | 4.4791E-13 |
| ENSG00000285278\|TFAP2A-AS2 | 277.4600 | 2.0008 | ##### | 7.4686 | 8.1028E-14 | 4.5224E-13 |
| ENSG00000163635\|ATXN7 | 2693.1490 | 0.9574 | ##### | 7.4671 | 8.1995E-14 | 4.5755E-13 |
| ENSG00000227619 | 1320.5732 | ####### | ##### | -7.4669 | 8.2120E-14 | 4.5816E-13 |
| ENSG00000132639\|SNAP25 | 689.4003 | 1.1000 | ##### | 7.4665 | 8.2374E-14 | 4.5949E-13 |

| | | | | | | |
|---|---|---|---|---|---|---|
| ENSG00000172728\|FUT10 | 777.9232 | -1.1044 | ##### | -7.4650 | 8.3329E-14 | 4.6473E-13 |
| ENSG00000134987\|WDR36 | 6167.5664 | 0.9284 | ##### | 7.4645 | 8.3648E-14 | 4.6642E-13 |
| ENSG00000148248\|SURF4 | 28130.9075 | -0.5315 | ##### | -7.4643 | 8.3735E-14 | 4.6682E-13 |
| ENSG00000164187\|LMBRD2 | 2939.6309 | -0.9484 | ##### | -7.4633 | 8.4408E-14 | 4.7048E-13 |
| ENSG00000291168 | 35.8329 | 4.2279 | ##### | 7.4629 | 8.4639E-14 | 4.7168E-13 |
| ENSG00000205339\|IPO7 | 31814.2448 | 0.8759 | ##### | 7.4608 | 8.5991E-14 | 4.7913E-13 |
| ENSG00000228695\|CES1P1 | 57.5462 | ####### | ##### | -7.4607 | 8.6056E-14 | 4.7940E-13 |
| ENSG00000187024\|PTRH1 | 2119.6987 | -1.0000 | ##### | -7.4598 | 8.6648E-14 | 4.8261E-13 |
| ENSG00000152484\|USP12 | 8729.5931 | 0.8983 | ##### | 7.4598 | 8.6684E-14 | 4.8272E-13 |
| ENSG00000126067\|PSMB2 | 15240.8480 | -0.6790 | ##### | -7.4582 | 8.7696E-14 | 4.8826E-13 |
| ENSG00000166295\|ANAPC16 | 5324.1524 | -0.8785 | ##### | -7.4566 | 8.8787E-14 | 4.9425E-13 |
| ENSG00000126775\|ATG14 | 2837.5654 | 0.9525 | ##### | 7.4562 | 8.9058E-14 | 4.9566E-13 |
| ENSG00000145217\|SLC26A1 | 231.1424 | -1.5552 | ##### | -7.4544 | 9.0268E-14 | 5.0230E-13 |
| ENSG00000273513\|TBC1D3K | 119.6104 | 1.6077 | ##### | 7.4528 | 9.1408E-14 | 5.0855E-13 |
| ENSG00000134698\|AGO4 | 968.6321 | -1.0311 | ##### | -7.4493 | 9.3858E-14 | 5.2209E-13 |
| ENSG00000198887\|SMC5 | 4053.9058 | -0.9266 | ##### | -7.4489 | 9.4100E-14 | 5.2333E-13 |
| ENSG00000181135\|ZNF707 | 1350.2428 | 0.9866 | ##### | 7.4476 | 9.5057E-14 | 5.2856E-13 |
| ENSG00000186222\|BLOC1S4 | 1326.0415 | -0.9786 | ##### | -7.4465 | 9.5875E-14 | 5.3301E-13 |
| ENSG00000127588\|GNG13 | 74.8638 | 1.8138 | ##### | 7.4454 | 9.6619E-14 | 5.3704E-13 |
| ENSG00000147124\|ZNF41 | 843.4681 | -1.0456 | ##### | -7.4452 | 9.6826E-14 | 5.3809E-13 |
| ENSG00000172955\|ADH6 | 27.9461 | -3.4968 | ##### | -7.4449 | 9.7019E-14 | 5.3906E-13 |
| ENSG00000134852\|CLOCK | 5608.4552 | 0.9073 | ##### | 7.4441 | 9.7631E-14 | 5.4237E-13 |
| ENSG00000163126\|ANKRD23 | 197.9494 | 1.7716 | ##### | 7.4428 | 9.8609E-14 | 5.4769E-13 |
| ENSG00000121057\|AKAP1 | 6382.8828 | 0.9113 | ##### | 7.4427 | 9.8656E-14 | 5.4785E-13 |
| ENSG00000254901\|BORCS8 | 1069.4294 | 0.9945 | ##### | 7.4423 | 9.8942E-14 | 5.4934E-13 |
| ENSG00000155189\|AGPAT5 | 4387.1120 | 0.9285 | ##### | 7.4404 | 1.0037E-13 | 5.5717E-13 |
| ENSG00000182324\|KCNJ14 | 398.1513 | -1.1718 | ##### | -7.4399 | 1.0075E-13 | 5.5920E-13 |
| ENSG00000204397\|CARD16 | 111.5628 | -9.0890 | ##### | -7.4356 | 1.0413E-13 | 5.7780E-13 |
| ENSG00000197622\|CDC42SE1 | 15227.5363 | 0.7351 | ##### | 7.4326 | 1.0644E-13 | 5.9055E-13 |
| ENSG00000291078 | 504.8206 | -1.0952 | ##### | -7.4324 | 1.0665E-13 | 5.9158E-13 |
| ENSG00000173805\|HAP1 | 322.4365 | 1.2620 | ##### | 7.4312 | 1.0765E-13 | 5.9700E-13 |
| ENSG00000250033\|SLC7A11-AS1 | 45.9803 | -2.3373 | ##### | -7.4291 | 1.0934E-13 | 6.0627E-13 |
| ENSG00000139835\|GRTP1 | 529.9637 | -1.0674 | ##### | -7.4265 | 1.1148E-13 | 6.1802E-13 |
| ENSG00000182858\|ALG12 | 2216.8048 | -0.9136 | ##### | -7.4255 | 1.1232E-13 | 6.2260E-13 |
| ENSG00000101639\|CEP192 | 4227.1097 | 0.9157 | ##### | 7.4249 | 1.1289E-13 | 6.2564E-13 |
| ENSG00000164117\|FBXO8 | 551.1380 | -1.2469 | ##### | -7.4242 | 1.1344E-13 | 6.2857E-13 |
| ENSG00000154642\|C21orf91 | 943.4850 | 0.9642 | ##### | 7.4211 | 1.1618E-13 | 6.4359E-13 |
| ENSG00000261465\|COQ7-DT | 36.5220 | -2.6702 | ##### | -7.4192 | 1.1787E-13 | 6.5286E-13 |
| ENSG00000148187\|MRRF | 3035.4666 | -0.9363 | ##### | -7.4167 | 1.2011E-13 | 6.6514E-13 |
| ENSG00000148814\|LRRC27 | 474.3611 | -1.1252 | ##### | -7.4154 | 1.2123E-13 | 6.7123E-13 |
| ENSG00000037965\|HOXC8 | 448.5129 | -1.0882 | ##### | -7.4140 | 1.2250E-13 | 6.7814E-13 |
| ENSG00000176973\|FAM89B | 1988.3231 | 0.9746 | ##### | 7.4111 | 1.2525E-13 | 6.9320E-13 |
| ENSG00000271147\|ARMCX5-GPRASP2 | 372.2684 | 1.3284 | ##### | 7.4105 | 1.2583E-13 | 6.9631E-13 |
| ENSG00000109794\|FAM149A | 168.9352 | -1.3586 | ##### | -7.4084 | 1.2785E-13 | 7.0735E-13 |
| ENSG00000198892\|SHISA4 | 891.3693 | -1.0264 | ##### | -7.4081 | 1.2810E-13 | 7.0858E-13 |
| ENSG00000163346\|PBXIP1 | 2175.1509 | 0.9646 | ##### | 7.4080 | 1.2825E-13 | 7.0931E-13 |
| ENSG00000113209\|PCDHB5 | 55.4925 | -8.2367 | ##### | -7.4079 | 1.2828E-13 | 7.0931E-13 |
| ENSG00000135318\|NT5E | 11524.9333 | 0.9295 | ##### | 7.4077 | 1.2856E-13 | 7.1072E-13 |
| ENSG00000173465\|ZNRD2 | 2254.6406 | 0.9091 | ##### | 7.4075 | 1.2871E-13 | 7.1145E-13 |
| ENSG00000143320\|CRABP2 | 2294.0795 | 0.9564 | ##### | 7.4066 | 1.2957E-13 | 7.1607E-13 |
| ENSG00000101452\|DHX35 | 2160.1733 | 0.9484 | ##### | 7.4051 | 1.3102E-13 | 7.2395E-13 |
| ENSG00000133597\|ADCK2 | 2498.0392 | 0.9421 | ##### | 7.4048 | 1.3137E-13 | 7.2574E-13 |
| ENSG00000119227\|PIGZ | 221.6393 | -1.4372 | ##### | -7.4042 | 1.3196E-13 | 7.2887E-13 |
| ENSG00000183386\|FHL3 | 830.9356 | -0.9868 | ##### | -7.4040 | 1.3210E-13 | 7.2948E-13 |
| ENSG00000113739\|STC2 | 1958.1208 | 1.1034 | ##### | 7.4038 | 1.3238E-13 | 7.3092E-13 |
| ENSG00000196967\|ZNF585A | 718.3210 | -1.0126 | ##### | -7.4037 | 1.3244E-13 | 7.3113E-13 |
| ENSG00000188816\|HMX2 | 155.9260 | ####### | ##### | -7.4036 | 1.3251E-13 | 7.3135E-13 |
| ENSG00000100075\|SLC25A1 | 4209.4342 | -0.7834 | ##### | -7.4036 | 1.3256E-13 | 7.3149E-13 |
| ENSG00000118971\|CCND2 | 6442.9793 | 12.2053 | ##### | 7.4034 | 1.3274E-13 | 7.3239E-13 |
| ENSG00000139496\|NUP58 | 10842.1989 | 0.8807 | ##### | 7.4020 | 1.3416E-13 | 7.4004E-13 |
| ENSG00000135912\|TTLL4 | 4052.0420 | 0.9046 | ##### | 7.4010 | 1.3514E-13 | 7.4534E-13 |
| ENSG00000149403\|GRIK4 | 60.4186 | 3.8689 | ##### | 7.3992 | 1.3703E-13 | 7.5563E-13 |
| ENSG00000128731\|HERC2 | 5206.8371 | 0.9339 | ##### | 7.3991 | 1.3714E-13 | 7.5607E-13 |

| | | | | | | |
|---|---|---|---|---|---|---|
| ENSG00000128789\|PSMG2 | 4119.4684 | 0.9400 | ##### | 7.3990 | 1.3720E-13 | 7.5624E-13 |
| ENSG00000136295\|TTYH3 | 7190.6296 | 0.9128 | ##### | 7.3982 | 1.3800E-13 | 7.6056E-13 |
| ENSG00000182263\|FIGN | 1718.5509 | 0.9651 | ##### | 7.3977 | 1.3852E-13 | 7.6326E-13 |
| ENSG00000120800\|UTP20 | 11023.3380 | 0.8866 | ##### | 7.3976 | 1.3870E-13 | 7.6409E-13 |
| ENSG00000166226\|CCT2 | 39865.9486 | 0.5592 | ##### | 7.3972 | 1.3908E-13 | 7.6607E-13 |
| ENSG00000196586\|MYO6 | 3209.0076 | -0.9113 | ##### | -7.3950 | 1.4139E-13 | 7.7863E-13 |
| ENSG00000153157\|SYCP2L | 376.6197 | 1.2214 | ##### | 7.3932 | 1.4331E-13 | 7.8910E-13 |
| ENSG00000280177 | 105.6250 | 7.9343 | ##### | 7.3932 | 1.4338E-13 | 7.8929E-13 |
| ENSG00000108239\|TBC1D12 | 1075.2623 | -0.9912 | ##### | -7.3929 | 1.4369E-13 | 7.9090E-13 |
| ENSG00000121552\|CSTA | 664.5514 | ####### | ##### | -7.3908 | 1.4597E-13 | 8.0327E-13 |
| ENSG00000168255\|POLR2J3 | 1472.5424 | 1.0969 | ##### | 7.3900 | 1.4686E-13 | 8.0800E-13 |
| ENSG00000023572\|GLRX2 | 2377.1198 | -0.9077 | ##### | -7.3882 | 1.4882E-13 | 8.1865E-13 |
| ENSG00000157542\|KCNJ6 | 476.7652 | -6.7402 | ##### | -7.3880 | 1.4907E-13 | 8.1990E-13 |
| ENSG00000248927 | 68.1947 | 8.3479 | ##### | 7.3879 | 1.4915E-13 | 8.2016E-13 |
| ENSG00000183963\|SMTN | 3181.2591 | 0.9348 | ##### | 7.3876 | 1.4953E-13 | 8.2209E-13 |
| ENSG00000241343\|RPL36A | 14117.4365 | 0.9113 | ##### | 7.3870 | 1.5012E-13 | 8.2522E-13 |
| ENSG00000139433\|GLTP | 4203.7001 | -0.9015 | ##### | -7.3864 | 1.5084E-13 | 8.2901E-13 |
| ENSG00000071243\|ING3 | 1080.3858 | 0.9860 | ##### | 7.3852 | 1.5218E-13 | 8.3624E-13 |
| ENSG00000065457\|ADAT1 | 3144.5568 | -0.9145 | ##### | -7.3828 | 1.5498E-13 | 8.5142E-13 |
| ENSG00000198482\|ZNF808 | 477.8230 | -1.0943 | ##### | -7.3809 | 1.5725E-13 | 8.6376E-13 |
| ENSG00000183458\|PKD1P3 | 3781.9737 | 0.9784 | ##### | 7.3799 | 1.5837E-13 | 8.6974E-13 |
| ENSG00000273213\|H3-7 | 356.9701 | -7.9693 | ##### | -7.3776 | 1.6115E-13 | 8.8487E-13 |
| ENSG00000140391\|TSPAN3 | 7070.2755 | -0.9221 | ##### | -7.3754 | 1.6386E-13 | 8.9958E-13 |
| ENSG00000142197\|DOP1B | 1206.6339 | -0.9737 | ##### | -7.3742 | 1.6532E-13 | 9.0741E-13 |
| ENSG00000179673\|RPRML | 63.1851 | 2.4173 | ##### | 7.3738 | 1.6589E-13 | 9.1036E-13 |
| ENSG00000120278\|PLEKHG1 | 211.8705 | 1.3954 | ##### | 7.3737 | 1.6595E-13 | 9.1052E-13 |
| ENSG00000132423\|COQ3 | 618.2038 | -0.8987 | ##### | -7.3735 | 1.6617E-13 | 9.1160E-13 |
| ENSG00000205978\|NYNRIN | 99.9761 | 1.9536 | ##### | 7.3725 | 1.6743E-13 | 9.1833E-13 |
| ENSG00000267429 | 34.8073 | -7.5268 | ##### | -7.3724 | 1.6755E-13 | 9.1883E-13 |
| ENSG00000237945\|LINC00649 | 75.3829 | -3.1650 | ##### | -7.3708 | 1.6962E-13 | 9.3000E-13 |
| ENSG00000182378\|PLCXD1 | 2901.0363 | -0.8883 | ##### | -7.3695 | 1.7129E-13 | 9.3897E-13 |
| ENSG00000101624\|CEP76 | 2167.6283 | 0.9392 | ##### | 7.3678 | 1.7349E-13 | 9.5086E-13 |
| ENSG00000178882\|RFLNA | 66.7420 | 8.1886 | ##### | 7.3674 | 1.7400E-13 | 9.5347E-13 |
| ENSG00000271737 | 103.1372 | -1.9045 | ##### | -7.3660 | 1.7588E-13 | 9.6361E-13 |
| ENSG00000135272\|MDFIC | 4706.8007 | -0.8688 | ##### | -7.3622 | 1.8097E-13 | 9.9132E-13 |
| ENSG00000051128\|HOMER3 | 2780.3956 | -0.9225 | ##### | -7.3618 | 1.8150E-13 | 9.9402E-13 |
| ENSG00000100852\|ARHGAP5 | 9436.1383 | 0.8990 | ##### | 7.3609 | 1.8263E-13 | 1.0000E-12 |
| ENSG00000076248\|UNG | 10093.0930 | 0.8857 | ##### | 7.3595 | 1.8460E-13 | 1.0106E-12 |
| ENSG00000197063\|MAFG | 6303.8417 | -0.8830 | ##### | -7.3594 | 1.8481E-13 | 1.0116E-12 |
| ENSG00000279342 | 31.7255 | 4.0896 | ##### | 7.3593 | 1.8491E-13 | 1.0120E-12 |
| ENSG00000101298\|SNPH | 674.1316 | -1.0476 | ##### | -7.3586 | 1.8589E-13 | 1.0171E-12 |
| ENSG00000215032\|GNL3LP1 | 26.8397 | -3.2897 | ##### | -7.3557 | 1.8995E-13 | 1.0392E-12 |
| ENSG00000130589\|HELZ2 | 4297.8443 | 0.9517 | ##### | 7.3549 | 1.9102E-13 | 1.0448E-12 |
| ENSG00000135314\|KHDC1 | 533.9015 | 1.1495 | ##### | 7.3547 | 1.9138E-13 | 1.0466E-12 |
| ENSG00000125657\|TNFSF9 | 403.3150 | -1.2948 | ##### | -7.3524 | 1.9471E-13 | 1.0646E-12 |
| ENSG00000214783\|POLR2J4 | 564.6757 | -1.3487 | ##### | -7.3523 | 1.9489E-13 | 1.0654E-12 |
| ENSG00000270558 | 94.1477 | 1.9427 | ##### | 7.3515 | 1.9593E-13 | 1.0709E-12 |
| ENSG00000143507\|DUSP10 | 278.2758 | 1.5235 | ##### | 7.3513 | 1.9634E-13 | 1.0728E-12 |
| ENSG00000180801\|ARSJ | 1842.1986 | -0.9478 | ##### | -7.3513 | 1.9633E-13 | 1.0728E-12 |
| ENSG00000117533\|VAMP4 | 955.6449 | -0.9689 | ##### | -7.3497 | 1.9868E-13 | 1.0853E-12 |
| ENSG00000109381\|ELF2 | 1215.0742 | -0.8170 | ##### | -7.3495 | 1.9901E-13 | 1.0869E-12 |
| ENSG00000133858\|ZFC3H1 | 3724.9229 | 0.8803 | ##### | 7.3491 | 1.9947E-13 | 1.0893E-12 |
| ENSG00000162441\|LZIC | 3230.8069 | -0.9510 | ##### | -7.3488 | 2.0005E-13 | 1.0923E-12 |
| ENSG00000164683\|HEY1 | 142.7884 | -1.4972 | ##### | -7.3482 | 2.0088E-13 | 1.0966E-12 |
| ENSG00000257815\|PRANCR | 199.3579 | 1.4262 | ##### | 7.3475 | 2.0190E-13 | 1.1017E-12 |
| ENSG00000085224\|ATRX | 7548.0594 | 0.8751 | ##### | 7.3475 | 2.0189E-13 | 1.1017E-12 |
| ENSG00000156482\|RPL30 | 48691.6759 | -0.8806 | ##### | -7.3470 | 2.0276E-13 | 1.1062E-12 |
| ENSG00000033800\|PIAS1 | 3137.1285 | 0.6572 | ##### | 7.3458 | 2.0455E-13 | 1.1158E-12 |
| ENSG00000136100\|VPS36 | 3720.4137 | 0.4757 | ##### | 7.3449 | 2.0589E-13 | 1.1229E-12 |
| ENSG00000273983\|H3C8 | 184.4105 | 6.9185 | ##### | 7.3391 | 2.1509E-13 | 1.1728E-12 |
| ENSG00000185585\|OLFML2A | 970.3355 | -1.0317 | ##### | -7.3371 | 2.1819E-13 | 1.1895E-12 |
| ENSG00000168005\|SPINDOC | 2863.5244 | 0.9287 | ##### | 7.3363 | 2.1958E-13 | 1.1969E-12 |
| ENSG00000164338\|UTP15 | 3337.9711 | 0.9527 | ##### | 7.3325 | 2.2588E-13 | 1.2310E-12 |

| | | | | | | |
|---|---|---|---|---|---|---|
| ENSG00000136305\|CIDEB | 616.5911 | 1.0492 | ##### | 7.3323 | 2.2628E-13 | 1.2329E-12 |
| ENSG00000117152\|RGS4 | 48.1081 | 2.4412 | ##### | 7.3321 | 2.2654E-13 | 1.2342E-12 |
| ENSG00000278791 | 2289.4137 | 10.8719 | ##### | 7.3304 | 2.2952E-13 | 1.2502E-12 |
| ENSG00000171320\|ESCO2 | 1984.5490 | 0.9437 | ##### | 7.3298 | 2.3054E-13 | 1.2555E-12 |
| ENSG00000114383\|TUSC2 | 2402.4958 | 0.9411 | ##### | 7.3288 | 2.3221E-13 | 1.2643E-12 |
| ENSG00000255153\|TOLLIP-DT | 75.3828 | -2.6199 | ##### | -7.3286 | 2.3255E-13 | 1.2660E-12 |
| ENSG00000272269\|NUP153-AS1 | 330.5626 | 1.2122 | ##### | 7.3270 | 2.3538E-13 | 1.2812E-12 |
| ENSG00000140525\|FANCI | 9771.9007 | 0.8817 | ##### | 7.3269 | 2.3562E-13 | 1.2822E-12 |
| ENSG00000112425\|EPM2A | 266.5314 | -1.3526 | ##### | -7.3259 | 2.3733E-13 | 1.2913E-12 |
| ENSG00000210107\|MT-TQ | 306.0777 | 0.8761 | ##### | 7.3255 | 2.3793E-13 | 1.2943E-12 |
| ENSG00000218510\|LINC00339 | 439.5546 | -1.3169 | ##### | -7.3245 | 2.3988E-13 | 1.3047E-12 |
| ENSG00000215866\|LINC01356 | 196.8313 | 2.3847 | ##### | 7.3236 | 2.4134E-13 | 1.3124E-12 |
| ENSG00000257365\|FNTB | 3445.9810 | 0.9622 | ##### | 7.3231 | 2.4227E-13 | 1.3172E-12 |
| ENSG00000083290\|ULK2 | 1583.9809 | -0.9423 | ##### | -7.3222 | 2.4396E-13 | 1.3262E-12 |
| ENSG00000172273\|HINFP | 1148.0832 | 0.9521 | ##### | 7.3221 | 2.4406E-13 | 1.3264E-12 |
| ENSG00000254612\|DNAJB6P1 | 195.6732 | 1.3829 | ##### | 7.3210 | 2.4619E-13 | 1.3378E-12 |
| ENSG00000165733\|BMS1 | 9641.0879 | 0.8355 | ##### | 7.3206 | 2.4691E-13 | 1.3415E-12 |
| ENSG00000137802\|MAPKBP1 | 2422.5855 | 0.9013 | ##### | 7.3193 | 2.4934E-13 | 1.3544E-12 |
| ENSG00000275183\|LENG9 | 859.4413 | -0.9938 | ##### | -7.3175 | 2.5270E-13 | 1.3724E-12 |
| ENSG00000132801\|ZSWIM3 | 231.6200 | 1.3931 | ##### | 7.3173 | 2.5308E-13 | 1.3742E-12 |
| ENSG00000176428\|VPS37D | 134.5326 | -1.5519 | ##### | -7.3161 | 2.5536E-13 | 1.3863E-12 |
| ENSG00000122741\|DCAF10 | 5170.1922 | 0.8483 | ##### | 7.3133 | 2.6060E-13 | 1.4146E-12 |
| ENSG00000102316\|MAGED2 | 5849.6284 | -0.8650 | ##### | -7.3122 | 2.6281E-13 | 1.4263E-12 |
| ENSG00000088930\|XRN2 | 11151.4052 | 0.8307 | ##### | 7.3107 | 2.6581E-13 | 1.4423E-12 |
| ENSG00000129295\|DNAAF11 | 284.8799 | 1.2039 | ##### | 7.3079 | 2.7136E-13 | 1.4721E-12 |
| ENSG00000124523\|SIRT5 | 829.7129 | 1.0579 | ##### | 7.3069 | 2.7344E-13 | 1.4832E-12 |
| ENSG00000188878\|FBF1 | 1543.2135 | 1.0126 | ##### | 7.3067 | 2.7388E-13 | 1.4853E-12 |
| ENSG00000136574\|GATA4 | 508.6454 | 7.1935 | ##### | 7.3060 | 2.7515E-13 | 1.4919E-12 |
| ENSG00000151092\|NGLY1 | 2151.0541 | -0.9061 | ##### | -7.3051 | 2.7702E-13 | 1.5018E-12 |
| ENSG00000287769 | 173.3998 | -1.4288 | ##### | -7.3012 | 2.8530E-13 | 1.5464E-12 |
| ENSG00000188021\|UBQLN2 | 3741.9056 | -0.8703 | ##### | -7.3004 | 2.8690E-13 | 1.5548E-12 |
| ENSG00000204856\|FAM216A | 2369.8796 | 0.9294 | ##### | 7.3002 | 2.8742E-13 | 1.5573E-12 |
| ENSG00000167378\|IRGQ | 2353.1599 | -0.9128 | ##### | -7.2999 | 2.8792E-13 | 1.5597E-12 |
| ENSG00000146574\|CCZ1B | 6735.0537 | -0.8871 | ##### | -7.2999 | 2.8802E-13 | 1.5600E-12 |
| ENSG00000147274\|RBMX | 18840.9183 | 0.5308 | ##### | 7.2993 | 2.8937E-13 | 1.5670E-12 |
| ENSG00000196227\|FAM217B | 2810.4931 | 0.9194 | ##### | 7.2991 | 2.8980E-13 | 1.5691E-12 |
| ENSG00000267152 | 98.5843 | -1.7847 | ##### | -7.2962 | 2.9597E-13 | 1.6022E-12 |
| ENSG00000178752\|ERFE | 547.6643 | 1.0828 | ##### | 7.2962 | 2.9607E-13 | 1.6024E-12 |
| ENSG00000131043\|AAR2 | 4614.6331 | 0.9077 | ##### | 7.2961 | 2.9625E-13 | 1.6031E-12 |
| ENSG00000121988\|ZRANB3 | 778.8045 | -1.0332 | ##### | -7.2945 | 2.9968E-13 | 1.6214E-12 |
| ENSG00000269374 | 109.8571 | -1.8500 | ##### | -7.2932 | 3.0263E-13 | 1.6370E-12 |
| ENSG00000127191\|TRAF2 | 5213.9919 | -0.7502 | ##### | -7.2927 | 3.0385E-13 | 1.6433E-12 |
| ENSG00000177189\|RPS6KA3 | 6105.2401 | -0.8959 | ##### | -7.2923 | 3.0481E-13 | 1.6482E-12 |
| ENSG00000289604\|CTAGE8 | 114.5360 | -1.7931 | ##### | -7.2898 | 3.1045E-13 | 1.6782E-12 |
| ENSG00000106992\|AK1 | 2014.1815 | -0.9418 | ##### | -7.2898 | 3.1046E-13 | 1.6782E-12 |
| ENSG00000196353\|CPNE4 | 615.0959 | -8.6059 | ##### | -7.2888 | 3.1276E-13 | 1.6903E-12 |
| ENSG00000122674\|CCZ1 | 8074.2114 | 0.6943 | ##### | 7.2886 | 3.1316E-13 | 1.6922E-12 |
| ENSG00000108352\|RAPGEFL1 | 674.0085 | 1.1174 | ##### | 7.2882 | 3.1418E-13 | 1.6974E-12 |
| ENSG00000135452\|TSPAN31 | 3013.5657 | -1.0530 | ##### | -7.2878 | 3.1501E-13 | 1.7015E-12 |
| ENSG00000162409\|PRKAA2 | 2028.7676 | -0.8717 | ##### | -7.2856 | 3.2024E-13 | 1.7295E-12 |
| ENSG00000267470\|ZNF571-AS1 | 176.4784 | -7.3497 | ##### | -7.2852 | 3.2112E-13 | 1.7339E-12 |
| ENSG00000267042 | 49.1187 | 2.4198 | ##### | 7.2839 | 3.2430E-13 | 1.7507E-12 |
| ENSG00000005206\|SPPL2B | 4327.5376 | 0.8795 | ##### | 7.2835 | 3.2516E-13 | 1.7551E-12 |
| ENSG00000123728\|RAP2C | 4693.4771 | 0.8328 | ##### | 7.2808 | 3.3196E-13 | 1.7915E-12 |
| ENSG00000231925\|TAPBP | 9822.9311 | -0.8565 | ##### | -7.2789 | 3.3650E-13 | 1.8156E-12 |
| ENSG00000233178 | 62.2931 | -8.4019 | ##### | -7.2784 | 3.3769E-13 | 1.8217E-12 |
| ENSG00000181027\|FKRP | 1678.6054 | -1.3206 | ##### | -7.2784 | 3.3790E-13 | 1.8224E-12 |
| ENSG00000124198\|ARFGEF2 | 10170.4555 | 0.8865 | ##### | 7.2783 | 3.3794E-13 | 1.8224E-12 |
| ENSG00000186472\|PCLO | 611.2764 | 1.1122 | ##### | 7.2754 | 3.4550E-13 | 1.8629E-12 |
| ENSG00000117602\|RCAN3 | 2219.5799 | -0.9313 | ##### | -7.2750 | 3.4645E-13 | 1.8676E-12 |
| ENSG00000075131\|TIPIN | 1665.0420 | 0.9405 | ##### | 7.2745 | 3.4762E-13 | 1.8736E-12 |
| ENSG00000113140\|SPARC | 48.6940 | 4.0613 | ##### | 7.2739 | 3.4922E-13 | 1.8819E-12 |
| ENSG00000251365\|LINC02236 | 59.3350 | -1.6520 | ##### | -7.2730 | 3.5158E-13 | 1.8943E-12 |

| | | | | | | |
|---|---|---|---|---|---|---|
| ENSG00000198315\|ZKSCAN8 | 1526.7780 | 0.9400 | ##### | 7.2710 | 3.5676E-13 | 1.9219E-12 |
| ENSG00000167770\|OTUB1 | 7165.3259 | 0.7849 | ##### | 7.2710 | 3.5695E-13 | 1.9225E-12 |
| ENSG00000169252\|ADRB2 | 119.2999 | 1.9141 | ##### | 7.2697 | 3.6030E-13 | 1.9402E-12 |
| ENSG00000215114\|UBXN2B | 3002.9957 | -0.9099 | ##### | -7.2694 | 3.6098E-13 | 1.9435E-12 |
| ENSG00000103994\|ZNF106 | 6584.6235 | 0.8992 | ##### | 7.2694 | 3.6115E-13 | 1.9441E-12 |
| ENSG00000102053\|ZC3H12B | 135.9763 | -1.7656 | ##### | -7.2685 | 3.6361E-13 | 1.9570E-12 |
| ENSG00000039523\|RIPOR1 | 7267.5955 | -0.9049 | ##### | -7.2656 | 3.7149E-13 | 1.9990E-12 |
| ENSG00000230445\|LRRC37A6P | 164.7705 | 1.3572 | ##### | 7.2603 | 3.8613E-13 | 2.0774E-12 |
| ENSG00000166851\|PLK1 | 15420.6740 | 0.9134 | ##### | 7.2596 | 3.8836E-13 | 2.0891E-12 |
| ENSG00000143450\|OAZ3 | 315.6466 | -1.2983 | ##### | -7.2589 | 3.9021E-13 | 2.0986E-12 |
| ENSG00000130881\|LRP3 | 2267.8208 | -0.9414 | ##### | -7.2572 | 3.9513E-13 | 2.1247E-12 |
| ENSG00000081181\|ARG2 | 743.0316 | 1.0673 | ##### | 7.2565 | 3.9712E-13 | 2.1350E-12 |
| ENSG00000159685\|CHCHD6 | 1374.7856 | 0.9507 | ##### | 7.2565 | 3.9722E-13 | 2.1351E-12 |
| ENSG00000115446\|UNC50 | 2657.5385 | 0.9254 | ##### | 7.2555 | 4.0022E-13 | 2.1509E-12 |
| ENSG00000285641 | 1285.8094 | 1.1587 | ##### | 7.2545 | 4.0302E-13 | 2.1656E-12 |
| ENSG00000156170\|NDUFAF6 | 987.3391 | -0.9809 | ##### | -7.2534 | 4.0646E-13 | 2.1837E-12 |
| ENSG00000196159\|FAT4 | 996.0978 | -0.9523 | ##### | -7.2526 | 4.0884E-13 | 2.1960E-12 |
| ENSG00000176919\|C8G | 108.1131 | -1.5471 | ##### | -7.2519 | 4.1091E-13 | 2.2067E-12 |
| ENSG00000160325\|CACFD1 | 611.2399 | -1.0197 | ##### | -7.2517 | 4.1143E-13 | 2.2092E-12 |
| ENSG00000188542\|DUSP28 | 660.2362 | 0.8525 | ##### | 7.2517 | 4.1160E-13 | 2.2097E-12 |
| ENSG00000163818\|LZTFL1 | 1168.5189 | 0.9780 | ##### | 7.2502 | 4.1606E-13 | 2.2332E-12 |
| ENSG00000143951\|WDPCP | 486.4640 | 1.0332 | ##### | 7.2493 | 4.1894E-13 | 2.2483E-12 |
| ENSG00000162676\|GFI1 | 97.9550 | 2.5158 | ##### | 7.2486 | 4.2103E-13 | 2.2591E-12 |
| ENSG00000258659\|TRIM34 | 3543.8594 | ####### | ##### | -7.2474 | 4.2483E-13 | 2.2790E-12 |
| ENSG00000163947\|ARHGEF3 | 1193.4684 | -0.9640 | ##### | -7.2472 | 4.2558E-13 | 2.2827E-12 |
| ENSG00000170801\|HTRA3 | 294.5195 | -5.2952 | ##### | -7.2467 | 4.2705E-13 | 2.2901E-12 |
| ENSG00000197540\|GZMM | 42.8440 | -2.9560 | ##### | -7.2455 | 4.3073E-13 | 2.3095E-12 |
| ENSG00000233396\|LINC01719 | 154.7810 | -1.4063 | ##### | -7.2455 | 4.3082E-13 | 2.3095E-12 |
| ENSG00000288873 | 1028.6641 | -1.2033 | ##### | -7.2446 | 4.3385E-13 | 2.3254E-12 |
| ENSG00000145041\|DCAF1 | 5338.6174 | 0.8822 | ##### | 7.2429 | 4.3922E-13 | 2.3537E-12 |
| ENSG00000293498 | 85.6953 | 4.7064 | ##### | 7.2424 | 4.4084E-13 | 2.3620E-12 |
| ENSG00000146826\|TRAPPC14 | 2083.3838 | 0.9264 | ##### | 7.2401 | 4.4823E-13 | 2.4011E-12 |
| ENSG00000137177\|KIF13A | 3901.9304 | 0.8843 | ##### | 7.2399 | 4.4908E-13 | 2.4052E-12 |
| ENSG00000103042\|SLC38A7 | 2529.3729 | -0.8838 | ##### | -7.2383 | 4.5444E-13 | 2.4336E-12 |
| ENSG00000197362\|ZNF786 | 1116.0445 | 0.9842 | ##### | 7.2378 | 4.5593E-13 | 2.4411E-12 |
| ENSG00000188322\|SBK1 | 309.8665 | 1.3415 | ##### | 7.2375 | 4.5717E-13 | 2.4473E-12 |
| ENSG00000197153\|H3C12 | 20.1244 | 4.2514 | ##### | 7.2351 | 4.6509E-13 | 2.4892E-12 |
| ENSG00000153291\|SLC25A27 | 107.8176 | -2.0318 | ##### | -7.2341 | 4.6879E-13 | 2.5086E-12 |
| ENSG00000293476 | 532.5432 | 1.0402 | ##### | 7.2331 | 4.7219E-13 | 2.5263E-12 |
| ENSG00000205744\|DENND1C | 319.6231 | -1.1245 | ##### | -7.2329 | 4.7288E-13 | 2.5296E-12 |
| ENSG00000105953\|OGDH | 14963.0863 | -0.8330 | ##### | -7.2326 | 4.7396E-13 | 2.5349E-12 |
| ENSG00000285957 | 108.1427 | 1.6423 | ##### | 7.2325 | 4.7407E-13 | 2.5350E-12 |
| ENSG00000173588\|CEP83 | 2078.3531 | 0.9306 | ##### | 7.2313 | 4.7826E-13 | 2.5569E-12 |
| ENSG00000135127\|BICDL1 | 1743.3169 | -0.9499 | ##### | -7.2310 | 4.7928E-13 | 2.5620E-12 |
| ENSG00000136878\|USP20 | 1239.4447 | -0.9405 | ##### | -7.2306 | 4.8077E-13 | 2.5695E-12 |
| ENSG00000165678\|GHITM | 19958.8114 | -0.5254 | ##### | -7.2283 | 4.8892E-13 | 2.6126E-12 |
| ENSG00000236975\|LINC02814 | 46.1844 | 8.3499 | ##### | 7.2282 | 4.8936E-13 | 2.6144E-12 |
| ENSG00000125844\|RRBP1 | 11969.1317 | -0.8429 | ##### | -7.2267 | 4.9504E-13 | 2.6443E-12 |
| ENSG00000124459\|ZNF45 | 1493.6141 | -0.9554 | ##### | -7.2266 | 4.9522E-13 | 2.6448E-12 |
| ENSG00000137673\|MMP7 | 3168.4647 | ####### | ##### | -7.2257 | 4.9853E-13 | 2.6620E-12 |
| ENSG00000290712\|DSTNP2 | 1158.2984 | -0.9523 | ##### | -7.2250 | 5.0104E-13 | 2.6749E-12 |
| ENSG00000135423\|GLS2 | 36.8286 | 2.3738 | ##### | 7.2249 | 5.0155E-13 | 2.6772E-12 |
| ENSG00000075420\|FNDC3B | 9154.5263 | -0.8503 | ##### | -7.2240 | 5.0467E-13 | 2.6934E-12 |
| ENSG00000048342\|CC2D2A | 1745.0980 | 0.9251 | ##### | 7.2237 | 5.0601E-13 | 2.7000E-12 |
| ENSG00000136197\|C7orf25 | 839.4114 | -1.4695 | ##### | -7.2222 | 5.1163E-13 | 2.7295E-12 |
| ENSG00000153896\|ZNF599 | 276.3555 | -1.1902 | ##### | -7.2221 | 5.1173E-13 | 2.7296E-12 |
| ENSG00000106305\|AIMP2 | 4544.9784 | -0.9348 | ##### | -7.2215 | 5.1411E-13 | 2.7418E-12 |
| ENSG00000145860\|RNF145 | 13208.6282 | 0.7797 | ##### | 7.2214 | 5.1444E-13 | 2.7431E-12 |
| ENSG00000166448\|TMEM130 | 82.7667 | -4.9268 | ##### | -7.2211 | 5.1568E-13 | 2.7492E-12 |
| ENSG00000286782 | 100.7422 | 7.8018 | ##### | 7.2210 | 5.1623E-13 | 2.7516E-12 |
| ENSG00000146938\|NLGN4X | 5800.1115 | ####### | ##### | -7.2198 | 5.2072E-13 | 2.7750E-12 |
| ENSG00000111664\|GNB3 | 357.2093 | 1.5607 | ##### | 7.2186 | 5.2520E-13 | 2.7984E-12 |
| ENSG00000135845\|PIGC | 1512.7264 | -0.9345 | ##### | -7.2167 | 5.3250E-13 | 2.8368E-12 |

| | | | | | | |
|---|---|---|---|---|---|---|
| ENSG00000224969 | 39.1640 | 6.8960 | ##### | 7.2142 | 5.4243E-13 | 2.8892E-12 |
| ENSG00000214595\|EML6 | 325.1357 | -1.2994 | ##### | -7.2124 | 5.4961E-13 | 2.9269E-12 |
| ENSG00000005175\|RPAP3 | 5594.2768 | 0.8790 | ##### | 7.2096 | 5.6099E-13 | 2.9870E-12 |
| ENSG00000105705\|SUGP1 | 2116.4118 | 0.9234 | ##### | 7.2087 | 5.6492E-13 | 3.0074E-12 |
| ENSG00000108375\|RNF43 | 501.2182 | -1.0595 | ##### | -7.2048 | 5.8134E-13 | 3.0942E-12 |
| ENSG00000284862\|CCDC39 | 51.7069 | 2.3080 | ##### | 7.2044 | 5.8294E-13 | 3.1022E-12 |
| ENSG00000157554\|ERG | 2591.2886 | 12.8670 | ##### | 7.2001 | 6.0147E-13 | 3.2002E-12 |
| ENSG00000130643\|CALY | 55.3570 | 2.2018 | ##### | 7.1996 | 6.0378E-13 | 3.2120E-12 |
| ENSG00000185267\|CDNF | 44.1688 | -2.1223 | ##### | -7.1991 | 6.0596E-13 | 3.2230E-12 |
| ENSG00000188352\|FOCAD | 2777.6858 | 0.8782 | ##### | 7.1991 | 6.0624E-13 | 3.2239E-12 |
| ENSG00000254634\|SMG1P6 | 219.4766 | 1.3224 | ##### | 7.1979 | 6.1158E-13 | 3.2517E-12 |
| ENSG00000163399\|ATP1A1 | 79792.4944 | 0.4954 | ##### | 7.1968 | 6.1620E-13 | 3.2757E-12 |
| ENSG00000273015\|LINC00938 | 816.5096 | -1.0452 | ##### | -7.1966 | 6.1752E-13 | 3.2821E-12 |
| ENSG00000233901\|LINC01503 | 61.0844 | -2.3767 | ##### | -7.1952 | 6.2359E-13 | 3.3138E-12 |
| ENSG00000125753\|VASP | 13803.1480 | -0.8427 | ##### | -7.1950 | 6.2465E-13 | 3.3188E-12 |
| ENSG00000151835\|SACS | 6060.7156 | 0.8828 | ##### | 7.1888 | 6.5356E-13 | 3.4718E-12 |
| ENSG00000291090\|CCDC144BP | 98.3523 | -3.7241 | ##### | -7.1879 | 6.5806E-13 | 3.4951E-12 |
| ENSG00000101966\|XIAP | 6813.8105 | 0.8551 | ##### | 7.1878 | 6.5842E-13 | 3.4964E-12 |
| ENSG00000243678\|NME2 | 34783.3576 | 0.8747 | ##### | 7.1869 | 6.6292E-13 | 3.5197E-12 |
| ENSG00000040608\|RTN4R | 241.6411 | -1.3389 | ##### | -7.1867 | 6.6382E-13 | 3.5238E-12 |
| ENSG00000164902\|PHAX | 4464.3584 | 0.9256 | ##### | 7.1833 | 6.8054E-13 | 3.6120E-12 |
| ENSG00000121390\|PSPC1 | 3235.4118 | 0.8489 | ##### | 7.1826 | 6.8403E-13 | 3.6298E-12 |
| ENSG00000175899\|A2M | 52.0844 | 2.6675 | ##### | 7.1808 | 6.9330E-13 | 3.6783E-12 |
| ENSG00000175741\|RWDD4P2 | 239.5270 | -1.2988 | ##### | -7.1801 | 6.9662E-13 | 3.6953E-12 |
| ENSG00000259414 | 25.3886 | 6.3206 | ##### | 7.1789 | 7.0296E-13 | 3.7283E-12 |
| ENSG00000182580\|EPHB3 | 107.5366 | -1.6919 | ##### | -7.1787 | 7.0398E-13 | 3.7330E-12 |
| ENSG00000156931\|VPS8 | 1595.1830 | -0.9274 | ##### | -7.1778 | 7.0844E-13 | 3.7560E-12 |
| ENSG00000132780\|NASP | 10644.4827 | 0.6996 | ##### | 7.1771 | 7.1219E-13 | 3.7752E-12 |
| ENSG00000266412\|NCOA4 | 8285.4428 | -0.8556 | ##### | -7.1748 | 7.2430E-13 | 3.8387E-12 |
| ENSG00000174013\|FBXO45 | 2772.9830 | -0.8803 | ##### | -7.1745 | 7.2578E-13 | 3.8459E-12 |
| ENSG00000278048\|U2 | 23.5364 | 4.0233 | ##### | 7.1742 | 7.2741E-13 | 3.8538E-12 |
| ENSG00000186642\|PDE2A | 336.4718 | -1.1728 | ##### | -7.1739 | 7.2899E-13 | 3.8615E-12 |
| ENSG00000150394\|CDH8 | 67.0302 | -3.0381 | ##### | -7.1737 | 7.2997E-13 | 3.8660E-12 |
| ENSG00000196208\|GREB1 | 295.5040 | -1.1698 | ##### | -7.1706 | 7.4684E-13 | 3.9547E-12 |
| ENSG00000265401 | 22284.6158 | 0.9014 | ##### | 7.1704 | 7.4757E-13 | 3.9578E-12 |
| ENSG00000053371\|AKR7A2 | 2997.2865 | 0.7568 | ##### | 7.1697 | 7.5146E-13 | 3.9777E-12 |
| ENSG00000132952\|USPL1 | 1956.9878 | 0.9319 | ##### | 7.1696 | 7.5191E-13 | 3.9794E-12 |
| ENSG00000152463\|OLAH | 173.6888 | ####### | ##### | -7.1691 | 7.5480E-13 | 3.9940E-12 |
| ENSG00000106404\|CLDN15 | 582.3495 | -0.9937 | ##### | -7.1689 | 7.5589E-13 | 3.9991E-12 |
| ENSG00000260423\|LINC02367 | 104.8263 | 1.6211 | ##### | 7.1670 | 7.6684E-13 | 4.0562E-12 |
| ENSG00000206530\|CFAP44 | 600.4592 | 1.0361 | ##### | 7.1665 | 7.6952E-13 | 4.0697E-12 |
| ENSG00000101098\|RIMS4 | 3498.5218 | 0.8601 | ##### | 7.1662 | 7.7117E-13 | 4.0777E-12 |
| ENSG00000133895\|MEN1 | 4071.8635 | 0.8986 | ##### | 7.1656 | 7.7421E-13 | 4.0931E-12 |
| ENSG00000102870\|ZNF629 | 2730.2439 | -0.8824 | ##### | -7.1645 | 7.8079E-13 | 4.1271E-12 |
| ENSG00000162551\|ALPL | 148.6448 | 2.0212 | ##### | 7.1634 | 7.8698E-13 | 4.1591E-12 |
| ENSG00000182154\|MRPL41 | 4644.3690 | -0.8989 | ##### | -7.1633 | 7.8759E-13 | 4.1616E-12 |
| ENSG00000284966 | 62.0914 | -2.0449 | ##### | -7.1625 | 7.9206E-13 | 4.1845E-12 |
| ENSG00000174599\|TRAM1L1 | 182.6801 | -1.9563 | ##### | -7.1618 | 7.9625E-13 | 4.2059E-12 |
| ENSG00000177875\|CCDC184 | 896.3340 | 1.0053 | ##### | 7.1593 | 8.1099E-13 | 4.2830E-12 |
| ENSG00000259959 | 154.8159 | 1.3150 | ##### | 7.1574 | 8.2215E-13 | 4.3412E-12 |
| ENSG00000090316\|MAEA | 7511.4367 | 0.8164 | ##### | 7.1558 | 8.3203E-13 | 4.3925E-12 |
| ENSG00000215529\|EFCAB8 | 49.1779 | 2.9664 | ##### | 7.1548 | 8.3779E-13 | 4.4222E-12 |
| ENSG00000112159\|MDN1 | 4346.9123 | 0.9446 | ##### | 7.1545 | 8.3997E-13 | 4.4329E-12 |
| ENSG00000148672\|GLUD1 | 12591.5001 | -0.6211 | ##### | -7.1530 | 8.4884E-13 | 4.4789E-12 |
| ENSG00000175662\|TOM1L2 | 1893.7195 | -0.9023 | ##### | -7.1527 | 8.5116E-13 | 4.4904E-12 |
| ENSG00000140323\|DISP2 | 915.5511 | 1.0060 | ##### | 7.1517 | 8.5720E-13 | 4.5214E-12 |
| ENSG00000011422\|PLAUR | 4193.8334 | 0.8738 | ##### | 7.1510 | 8.6161E-13 | 4.5439E-12 |
| ENSG00000188089\|PLA2G4E | 51.9902 | 3.8123 | ##### | 7.1509 | 8.6193E-13 | 4.5440E-12 |
| ENSG00000182963\|GJC1 | 5659.2151 | 0.8376 | ##### | 7.1509 | 8.6183E-13 | 4.5440E-12 |
| ENSG00000227671\|ZNF731P | 581.1470 | -1.0063 | ##### | -7.1502 | 8.6658E-13 | 4.5677E-12 |
| ENSG00000197728\|RPS26 | 37418.1832 | 0.8779 | ##### | 7.1487 | 8.7578E-13 | 4.6154E-12 |
| ENSG00000107077\|KDM4C | 770.1254 | -0.9926 | ##### | -7.1471 | 8.8607E-13 | 4.6687E-12 |
| ENSG00000101605\|MYOM1 | 491.4018 | 1.2274 | ##### | 7.1456 | 8.9611E-13 | 4.7208E-12 |

| | | | | | | |
|---|---|---|---|---|---|---|
| ENSG00000161847\|RAVER1 | 8270.5048 | 0.8400 | ##### | 7.1447 | 9.0209E-13 | 4.7515E-12 |
| ENSG00000196652\|ZKSCAN5 | 2189.3450 | 0.8803 | ##### | 7.1442 | 9.0525E-13 | 4.7673E-12 |
| ENSG00000014138\|POLA2 | 3005.4020 | 0.9287 | ##### | 7.1437 | 9.0884E-13 | 4.7853E-12 |
| ENSG00000218739\|CEBPZOS | 4967.1831 | -0.8848 | ##### | -7.1422 | 9.1821E-13 | 4.8339E-12 |
| ENSG00000198342\|ZNF442 | 212.0853 | ####### | ##### | -7.1420 | 9.1993E-13 | 4.8420E-12 |
| ENSG00000103035\|PSMD7 | 13068.8959 | -0.7925 | ##### | -7.1416 | 9.2279E-13 | 4.8562E-12 |
| ENSG00000081721\|DUSP12 | 4591.4593 | 0.8813 | ##### | 7.1394 | 9.3757E-13 | 4.9332E-12 |
| ENSG00000106459\|NRF1 | 1736.3703 | 0.9397 | ##### | 7.1388 | 9.4154E-13 | 4.9531E-12 |
| ENSG00000293460\|LINC02210 | 979.1742 | 0.9737 | ##### | 7.1382 | 9.4529E-13 | 4.9720E-12 |
| ENSG00000230513\|THAP7-AS1 | 372.0928 | -1.2941 | ##### | -7.1381 | 9.4662E-13 | 4.9781E-12 |
| ENSG00000248846\|LINC02065 | 63.6568 | -8.4316 | ##### | -7.1373 | 9.5156E-13 | 5.0032E-12 |
| ENSG00000105887\|MTPN | 29059.1100 | 0.8597 | ##### | 7.1373 | 9.5193E-13 | 5.0043E-12 |
| ENSG00000261211 | 166.0824 | -1.6537 | ##### | -7.1372 | 9.5264E-13 | 5.0071E-12 |
| ENSG00000005189\|REXO5 | 403.5278 | -1.2067 | ##### | -7.1368 | 9.5550E-13 | 5.0213E-12 |
| ENSG00000164736\|SOX17 | 296.5856 | 6.5571 | ##### | 7.1365 | 9.5745E-13 | 5.0307E-12 |
| ENSG00000135245\|HILPDA | 1526.8316 | 0.9613 | ##### | 7.1360 | 9.6121E-13 | 5.0495E-12 |
| ENSG00000151725\|CENPU | 1729.2647 | -0.9284 | ##### | -7.1355 | 9.6434E-13 | 5.0651E-12 |
| ENSG00000137942\|FNBP1L | 4827.9017 | -0.8746 | ##### | -7.1347 | 9.6972E-13 | 5.0925E-12 |
| ENSG00000246985\|SOCS2-AS1 | 272.6049 | 1.2275 | ##### | 7.1347 | 9.7033E-13 | 5.0947E-12 |
| ENSG00000165879\|FRAT1 | 204.7316 | -1.3958 | ##### | -7.1344 | 9.7204E-13 | 5.1028E-12 |
| ENSG00000206527\|HACD2 | 8250.2231 | -0.8676 | ##### | -7.1339 | 9.7529E-13 | 5.1190E-12 |
| ENSG00000150764\|DIXDC1 | 1734.7399 | -0.7803 | ##### | -7.1314 | 9.9321E-13 | 5.2122E-12 |
| ENSG00000173852\|DPY19L1 | 13102.9068 | -0.8499 | ##### | -7.1306 | 9.9936E-13 | 5.2435E-12 |
| ENSG00000230896 | 77.2056 | -1.9247 | ##### | -7.1299 | 1.0045E-12 | 5.2696E-12 |
| ENSG00000293338 | 47.8009 | 2.5207 | ##### | 7.1289 | 1.0121E-12 | 5.3085E-12 |
| ENSG00000135931\|ARMC9 | 1802.6250 | 0.9368 | ##### | 7.1285 | 1.0145E-12 | 5.3202E-12 |
| ENSG00000095637\|SORBS1 | 213.0587 | 1.3208 | ##### | 7.1283 | 1.0159E-12 | 5.3267E-12 |
| ENSG00000204136\|GGTA1 | 45.7061 | -4.0408 | ##### | -7.1266 | 1.0290E-12 | 5.3940E-12 |
| ENSG00000135063\|ENTREP1 | 57.0239 | -3.5596 | ##### | -7.1262 | 1.0314E-12 | 5.4060E-12 |
| ENSG00000197702\|PARVA | 3695.8879 | -0.9022 | ##### | -7.1260 | 1.0334E-12 | 5.4157E-12 |
| ENSG00000174669\|SLC29A2 | 1980.0164 | -0.9415 | ##### | -7.1259 | 1.0338E-12 | 5.4167E-12 |
| ENSG00000151136\|ABTB3 | 270.6648 | -6.8814 | ##### | -7.1257 | 1.0353E-12 | 5.4237E-12 |
| ENSG00000274828 | 468.1770 | -1.2655 | ##### | -7.1256 | 1.0361E-12 | 5.4270E-12 |
| ENSG00000129173\|E2F8 | 1432.4451 | 0.9036 | ##### | 7.1246 | 1.0438E-12 | 5.4662E-12 |
| ENSG00000268521\|VN1R83P | 34.2406 | -7.7086 | ##### | -7.1240 | 1.0483E-12 | 5.4887E-12 |
| ENSG00000123472\|ATPAF1 | 3441.9523 | -0.8974 | ##### | -7.1203 | 1.0768E-12 | 5.6371E-12 |
| ENSG00000178821\|TMEM52 | 98.6597 | 1.7656 | ##### | 7.1188 | 1.0888E-12 | 5.6986E-12 |
| ENSG00000289901 | 317.8368 | 0.9338 | ##### | 7.1184 | 1.0917E-12 | 5.7130E-12 |
| ENSG00000143515\|ATP8B2 | 9015.8160 | 0.8514 | ##### | 7.1183 | 1.0924E-12 | 5.7158E-12 |
| ENSG00000171492\|LRRC8D | 3894.7655 | -0.8649 | ##### | -7.1181 | 1.0945E-12 | 5.7256E-12 |
| ENSG00000243479\|MNX1-AS1 | 150.0845 | 1.7598 | ##### | 7.1169 | 1.1041E-12 | 5.7747E-12 |
| ENSG00000155545\|MIER3 | 3370.4709 | -0.8412 | ##### | -7.1157 | 1.1133E-12 | 5.8216E-12 |
| ENSG00000176595\|KBTBD11 | 303.8431 | 1.1953 | ##### | 7.1157 | 1.1137E-12 | 5.8230E-12 |
| ENSG00000053770\|AP5M1 | 4121.2071 | 0.7311 | ##### | 7.1138 | 1.1291E-12 | 5.9023E-12 |
| ENSG00000165181\|SHOC1 | 125.4711 | -5.0699 | ##### | -7.1132 | 1.1340E-12 | 5.9272E-12 |
| ENSG00000247627\|MTND4P12 | 234.2932 | 1.3028 | ##### | 7.1121 | 1.1429E-12 | 5.9724E-12 |
| ENSG00000232456\|FAM131C2P | 111.6120 | -1.6734 | ##### | -7.1118 | 1.1454E-12 | 5.9843E-12 |
| ENSG00000237651\|C2orf74 | 689.7689 | -1.3973 | ##### | -7.1108 | 1.1539E-12 | 6.0279E-12 |
| ENSG00000048392\|RRM2B | 4473.1600 | -0.8708 | ##### | -7.1106 | 1.1554E-12 | 6.0346E-12 |
| ENSG00000260018\|MBTPS1-DT | 77.7482 | -1.6108 | ##### | -7.1083 | 1.1748E-12 | 6.1349E-12 |
| ENSG00000271894 | 48.5722 | 23.6823 | ##### | 7.1073 | 1.1831E-12 | 6.1773E-12 |
| ENSG00000170122\|FOXD4 | 121.8997 | 1.8724 | ##### | 7.1071 | 1.1853E-12 | 6.1876E-12 |
| ENSG00000291174 | 409.0075 | 1.1983 | ##### | 7.1070 | 1.1858E-12 | 6.1889E-12 |
| ENSG00000175874\|CREG2 | 108.2224 | -5.4721 | ##### | -7.1062 | 1.1926E-12 | 6.2234E-12 |
| ENSG00000108091\|CCDC6 | 6332.3677 | -0.8503 | ##### | -7.1062 | 1.1930E-12 | 6.2245E-12 |
| ENSG00000128510\|CPA4 | 407.7863 | -1.1560 | ##### | -7.1046 | 1.2067E-12 | 6.2951E-12 |
| ENSG00000173065\|FAM222B | 2616.1655 | -0.8668 | ##### | -7.1039 | 1.2127E-12 | 6.3251E-12 |
| ENSG00000269855\|RNF225 | 42.9936 | 7.5291 | ##### | 7.1017 | 1.2325E-12 | 6.4274E-12 |
| ENSG00000116062\|MSH6 | 22711.1205 | 0.8265 | ##### | 7.1014 | 1.2354E-12 | 6.4410E-12 |
| ENSG00000145425\|RPS3A | 56687.6958 | -0.8535 | ##### | -7.1010 | 1.2390E-12 | 6.4588E-12 |
| ENSG00000270882\|H4C14 | 1252.1575 | 0.9171 | ##### | 7.0978 | 1.2679E-12 | 6.6086E-12 |
| ENSG00000196456\|ZNF775 | 820.6070 | 0.9296 | ##### | 7.0968 | 1.2766E-12 | 6.6524E-12 |
| ENSG00000150527\|MIA2 | 3069.2062 | -0.8811 | ##### | -7.0958 | 1.2859E-12 | 6.6999E-12 |

| | | | | | | |
|---|---|---|---|---|---|---|
| ENSG00000162894\|FCMR | 65.8817 | -2.4840 | ##### | -7.0928 | 1.3141E-12 | 6.8453E-12 |
| ENSG00000123737\|EXOSC9 | 3757.0984 | 0.8733 | ##### | 7.0925 | 1.3174E-12 | 6.8617E-12 |
| ENSG00000142396\|ERVK3-1 | 5453.9159 | -0.9170 | ##### | -7.0911 | 1.3303E-12 | 6.9273E-12 |
| ENSG00000182180\|MRPS16 | 8980.6272 | -0.8575 | ##### | -7.0910 | 1.3317E-12 | 6.9336E-12 |
| ENSG00000205060\|SLC35B4 | 4945.2798 | -0.8605 | ##### | -7.0892 | 1.3491E-12 | 7.0230E-12 |
| ENSG00000095321\|CRAT | 2667.3679 | -0.8921 | ##### | -7.0888 | 1.3525E-12 | 7.0395E-12 |
| ENSG00000188706\|ZDHHC9 | 4129.8830 | -0.9027 | ##### | -7.0870 | 1.3701E-12 | 7.1299E-12 |
| ENSG00000189064\|GAGE2A | 25512.1346 | ####### | ##### | -7.0837 | 1.4033E-12 | 7.3012E-12 |
| ENSG00000108950\|FAM20A | 58.6743 | -3.6454 | ##### | -7.0833 | 1.4075E-12 | 7.3217E-12 |
| ENSG00000142279\|WTIP | 2631.5111 | -0.8716 | ##### | -7.0824 | 1.4170E-12 | 7.3699E-12 |
| ENSG00000173890\|GPR160 | 162.9222 | -1.4282 | ##### | -7.0821 | 1.4201E-12 | 7.3848E-12 |
| ENSG00000213977\|TAX1BP3 | 5118.8083 | -0.8847 | ##### | -7.0802 | 1.4399E-12 | 7.4863E-12 |
| ENSG00000129167\|TPH1 | 33.6972 | 2.7426 | ##### | 7.0779 | 1.4638E-12 | 7.6095E-12 |
| ENSG00000224080\|UBE2FP1 | 50.7384 | 2.3095 | ##### | 7.0777 | 1.4656E-12 | 7.6176E-12 |
| ENSG00000155324\|GRAMD2B | 180.0800 | 1.4659 | ##### | 7.0757 | 1.4868E-12 | 7.7262E-12 |
| ENSG00000162378\|ZYG11B | 3133.2337 | -0.6745 | ##### | -7.0753 | 1.4915E-12 | 7.7496E-12 |
| ENSG00000114646\|CSPG5 | 508.3593 | 1.1434 | ##### | 7.0750 | 1.4941E-12 | 7.7616E-12 |
| ENSG00000249992\|TMEM158 | 2427.6608 | 0.9686 | ##### | 7.0743 | 1.5022E-12 | 7.8023E-12 |
| ENSG00000290032 | 1563.9508 | 0.9418 | ##### | 7.0740 | 1.5049E-12 | 7.8147E-12 |
| ENSG00000227398\|KIF9-AS1 | 282.0901 | -1.2375 | ##### | -7.0733 | 1.5132E-12 | 7.8565E-12 |
| ENSG00000140463\|BBS4 | 1554.5858 | -0.9105 | ##### | -7.0731 | 1.5155E-12 | 7.8674E-12 |
| ENSG00000176208\|ATAD5 | 1849.4974 | 0.9077 | ##### | 7.0709 | 1.5398E-12 | 7.9920E-12 |
| ENSG00000170909\|OSCAR | 248.2313 | -1.3044 | ##### | -7.0698 | 1.5511E-12 | 8.0491E-12 |
| ENSG00000272667 | 204.5521 | -1.2498 | ##### | -7.0695 | 1.5546E-12 | 8.0661E-12 |
| ENSG00000141030\|COPS3 | 5842.2410 | 0.8641 | ##### | 7.0675 | 1.5778E-12 | 8.1842E-12 |
| ENSG00000163138\|PACRGL | 1553.5679 | 0.9310 | ##### | 7.0675 | 1.5779E-12 | 8.1842E-12 |
| ENSG00000114923\|SLC4A3 | 1073.9853 | 0.9130 | ##### | 7.0656 | 1.5988E-12 | 8.2910E-12 |
| ENSG00000172594\|SMPDL3A | 230.9338 | -1.6485 | ##### | -7.0646 | 1.6111E-12 | 8.3534E-12 |
| ENSG00000203761\|MSTO2P | 2215.4558 | 0.9508 | ##### | 7.0636 | 1.6225E-12 | 8.4109E-12 |
| ENSG00000067208\|EVI5 | 2241.7234 | -0.6961 | ##### | -7.0623 | 1.6375E-12 | 8.4875E-12 |
| ENSG00000177169\|ULK1 | 9568.3136 | -0.8548 | ##### | -7.0622 | 1.6392E-12 | 8.4946E-12 |
| ENSG00000008130\|NADK | 7986.1139 | -0.8274 | ##### | -7.0614 | 1.6486E-12 | 8.5420E-12 |
| ENSG00000082438\|COBLL1 | 821.2030 | -1.0605 | ##### | -7.0608 | 1.6558E-12 | 8.5777E-12 |
| ENSG00000213918\|DNASE1 | 1868.7923 | 1.7094 | ##### | 7.0607 | 1.6571E-12 | 8.5831E-12 |
| ENSG00000269834\|ZNF528-AS1 | 83.1161 | -2.3473 | ##### | -7.0596 | 1.6693E-12 | 8.6448E-12 |
| ENSG00000075240\|GRAMD4 | 1421.5868 | -0.8759 | ##### | -7.0594 | 1.6725E-12 | 8.6598E-12 |
| ENSG00000291118\|ZNF767P | 771.2240 | 0.9536 | ##### | 7.0593 | 1.6731E-12 | 8.6614E-12 |
| ENSG00000230724\|LINC01001 | 83.4933 | 1.6635 | ##### | 7.0591 | 1.6756E-12 | 8.6729E-12 |
| ENSG00000177873\|ZNF619 | 245.3943 | -1.3221 | ##### | -7.0585 | 1.6831E-12 | 8.7102E-12 |
| ENSG00000167110\|GOLGA2 | 6414.8195 | -0.8217 | ##### | -7.0577 | 1.6923E-12 | 8.7561E-12 |
| ENSG00000147533\|GOLGA7 | 5108.2985 | 0.8617 | ##### | 7.0566 | 1.7067E-12 | 8.8292E-12 |
| ENSG00000188523\|CFAP77 | 49.8941 | 7.9682 | ##### | 7.0562 | 1.7108E-12 | 8.8488E-12 |
| ENSG00000135407\|AVIL | 168.4449 | 1.2337 | ##### | 7.0549 | 1.7269E-12 | 8.9305E-12 |
| ENSG00000143970\|ASXL2 | 6372.8837 | 0.8478 | ##### | 7.0539 | 1.7399E-12 | 8.9964E-12 |
| ENSG00000142700\|DMRTA2 | 110.4354 | 5.2471 | ##### | 7.0526 | 1.7561E-12 | 9.0785E-12 |
| ENSG00000143353\|LYPLAL1 | 1184.6547 | -0.9346 | ##### | -7.0524 | 1.7588E-12 | 9.0906E-12 |
| ENSG00000143420\|ENSA | 12962.6462 | 0.7342 | ##### | 7.0520 | 1.7641E-12 | 9.1164E-12 |
| ENSG00000248479 | 106.4832 | 8.7140 | ##### | 7.0511 | 1.7748E-12 | 9.1702E-12 |
| ENSG00000158480\|SPATA2 | 2287.4995 | 0.8860 | ##### | 7.0504 | 1.7843E-12 | 9.2177E-12 |
| ENSG00000100997\|ABHD12 | 2732.4375 | -0.9031 | ##### | -7.0472 | 1.8252E-12 | 9.4274E-12 |
| ENSG00000143028\|SYPL2 | 81.9911 | 2.2942 | ##### | 7.0459 | 1.8427E-12 | 9.5164E-12 |
| ENSG00000187051\|RPS19BP1 | 4921.6080 | 0.8918 | ##### | 7.0458 | 1.8442E-12 | 9.5225E-12 |
| ENSG00000104343\|UBE2W | 2149.3529 | -0.9038 | ##### | -7.0451 | 1.8537E-12 | 9.5697E-12 |
| ENSG00000116679\|IVNS1ABP | 7409.7659 | 0.8120 | ##### | 7.0425 | 1.8885E-12 | 9.7471E-12 |
| ENSG00000128595\|CALU | 30686.4725 | -0.8252 | ##### | -7.0425 | 1.8887E-12 | 9.7471E-12 |
| ENSG00000145982\|FARS2 | 778.2075 | 0.9517 | ##### | 7.0422 | 1.8918E-12 | 9.7614E-12 |
| ENSG00000123584\|MAGEA9 | 145.6271 | -9.1752 | ##### | -7.0421 | 1.8942E-12 | 9.7719E-12 |
| ENSG00000274286\|ADRA2B | 1934.5955 | 12.4145 | ##### | 7.0414 | 1.9027E-12 | 9.8142E-12 |
| ENSG00000267023\|LRRC37A16P | 548.1236 | -1.0035 | ##### | -7.0405 | 1.9154E-12 | 9.8778E-12 |
| ENSG00000198947\|DMD | 3155.2609 | 0.9152 | ##### | 7.0402 | 1.9200E-12 | 9.9000E-12 |
| ENSG00000231528\|FAM225A | 1454.6912 | 12.0325 | ##### | 7.0401 | 1.9206E-12 | 9.9016E-12 |
| ENSG00000151025\|GPR158 | 70.9719 | -1.7915 | ##### | -7.0385 | 1.9429E-12 | 1.0015E-11 |
| ENSG00000104907\|TRMT1 | 5821.2201 | 0.6390 | ##### | 7.0383 | 1.9467E-12 | 1.0032E-11 |

| | | | | | | |
|---|---|---|---|---|---|---|
| ENSG00000291067 | 668.8279 | 1.0091 | ##### | 7.0374 | 1.9581E-12 | 1.0090E-11 |
| ENSG00000004534\|RBM6 | 4081.7632 | 0.8420 | ##### | 7.0351 | 1.9906E-12 | 1.0255E-11 |
| ENSG00000204628\|RACK1 | 66990.0297 | 0.8378 | ##### | 7.0346 | 1.9979E-12 | 1.0291E-11 |
| ENSG00000242110\|AMACR | 1360.9647 | -0.9400 | ##### | -7.0343 | 2.0030E-12 | 1.0315E-11 |
| ENSG00000251143 | 3529.5863 | 1.0151 | ##### | 7.0339 | 2.0089E-12 | 1.0344E-11 |
| ENSG00000214331\|PDPR2P | 280.3202 | 5.9220 | ##### | 7.0336 | 2.0122E-12 | 1.0359E-11 |
| ENSG00000119547\|ONECUT2 | 695.6524 | -1.0019 | ##### | -7.0330 | 2.0207E-12 | 1.0401E-11 |
| ENSG00000180626\|ZNF594 | 525.0451 | 1.0228 | ##### | 7.0330 | 2.0212E-12 | 1.0402E-11 |
| ENSG00000222012 | 82.6578 | 5.8117 | ##### | 7.0321 | 2.0350E-12 | 1.0471E-11 |
| ENSG00000177200\|CHD9 | 3708.2186 | -0.8956 | ##### | -7.0319 | 2.0373E-12 | 1.0481E-11 |
| ENSG00000291026 | 45.8457 | -2.0851 | ##### | -7.0317 | 2.0398E-12 | 1.0492E-11 |
| ENSG00000113300\|CNOT6 | 4028.0168 | 0.8510 | ##### | 7.0302 | 2.0617E-12 | 1.0603E-11 |
| ENSG00000136149\|RPL13AP25 | 288.5189 | -1.1164 | ##### | -7.0302 | 2.0622E-12 | 1.0604E-11 |
| ENSG00000170162\|VGLL2 | 38.7952 | 8.0073 | ##### | 7.0292 | 2.0767E-12 | 1.0677E-11 |
| ENSG00000162694\|EXTL2 | 1698.9718 | -0.9392 | ##### | -7.0290 | 2.0804E-12 | 1.0694E-11 |
| ENSG00000129197\|RPAIN | 2166.8597 | 1.0363 | ##### | 7.0282 | 2.0923E-12 | 1.0753E-11 |
| ENSG00000115365\|LANCL1 | 11153.9215 | -0.8609 | ##### | -7.0267 | 2.1146E-12 | 1.0866E-11 |
| ENSG00000184162\|NR2C2AP | 2144.6245 | 0.8538 | ##### | 7.0260 | 2.1255E-12 | 1.0920E-11 |
| ENSG00000108039\|XPNPEP1 | 4239.1811 | -0.8723 | ##### | -7.0247 | 2.1450E-12 | 1.1018E-11 |
| ENSG00000170471\|RALGAPB | 7419.3877 | 0.8497 | ##### | 7.0244 | 2.1492E-12 | 1.1038E-11 |
| ENSG00000290951 | 836.7370 | 0.9947 | ##### | 7.0231 | 2.1696E-12 | 1.1141E-11 |
| ENSG00000237515\|SHISA9 | 74.8032 | -2.5673 | ##### | -7.0224 | 2.1804E-12 | 1.1194E-11 |
| ENSG00000181722\|ZBTB20 | 160.8606 | -3.3609 | ##### | -7.0223 | 2.1830E-12 | 1.1206E-11 |
| ENSG00000156103\|MMP16 | 356.6201 | 1.1710 | ##### | 7.0209 | 2.2043E-12 | 1.1313E-11 |
| ENSG00000213062 | 231.7518 | -1.3166 | ##### | -7.0198 | 2.2218E-12 | 1.1401E-11 |
| ENSG00000267698 | 79.3045 | 1.7772 | ##### | 7.0169 | 2.2680E-12 | 1.1636E-11 |
| ENSG00000161048\|NAPEPLD | 1546.0061 | -0.9556 | ##### | -7.0151 | 2.2975E-12 | 1.1786E-11 |
| ENSG00000266265\|KLF14 | 189.0101 | 1.2987 | ##### | 7.0142 | 2.3123E-12 | 1.1857E-11 |
| ENSG00000180815\|MAP3K15 | 443.9734 | 1.1188 | ##### | 7.0142 | 2.3120E-12 | 1.1857E-11 |
| ENSG00000164306\|PRIMPOL | 694.7246 | -0.9755 | ##### | -7.0129 | 2.3337E-12 | 1.1965E-11 |
| ENSG00000065135\|GNAI3 | 13787.5514 | 0.5347 | ##### | 7.0127 | 2.3383E-12 | 1.1987E-11 |
| ENSG00000073111\|MCM2 | 21175.3121 | 0.8210 | ##### | 7.0106 | 2.3722E-12 | 1.2158E-11 |
| ENSG00000153774\|CFDP1 | 2129.7476 | 0.8922 | ##### | 7.0104 | 2.3763E-12 | 1.2177E-11 |
| ENSG00000237015 | 76.9642 | 1.7578 | ##### | 7.0100 | 2.3833E-12 | 1.2211E-11 |
| ENSG00000087494\|PTHLH | 109.0799 | 1.9587 | ##### | 7.0098 | 2.3867E-12 | 1.2226E-11 |
| ENSG00000123892\|RAB38 | 270.6425 | -7.0128 | ##### | -7.0096 | 2.3900E-12 | 1.2241E-11 |
| ENSG00000261188\|TFIP11-DT | 179.0638 | -1.9616 | ##### | -7.0085 | 2.4081E-12 | 1.2331E-11 |
| ENSG00000289207 | 82.7488 | -1.7287 | ##### | -7.0083 | 2.4127E-12 | 1.2353E-11 |
| ENSG00000177096\|PHETA2 | 47.8458 | 7.7152 | ##### | 7.0062 | 2.4483E-12 | 1.2533E-11 |
| ENSG00000288271 | 45.3212 | 3.0052 | ##### | 7.0019 | 2.5243E-12 | 1.2920E-11 |
| ENSG00000092931\|MFSD11 | 4282.9200 | -1.6004 | ##### | -7.0016 | 2.5304E-12 | 1.2949E-11 |
| ENSG00000290101 | 103.7624 | -4.7767 | ##### | -7.0015 | 2.5323E-12 | 1.2956E-11 |
| ENSG00000115839\|RAB3GAP1 | 7476.4882 | -0.8637 | ##### | -6.9999 | 2.5608E-12 | 1.3100E-11 |
| ENSG00000145247\|OCIAD2 | 1344.0378 | -0.9492 | ##### | -6.9992 | 2.5742E-12 | 1.3166E-11 |
| ENSG00000123119\|NECAB1 | 137.6018 | -2.4856 | ##### | -6.9982 | 2.5923E-12 | 1.3257E-11 |
| ENSG00000132646\|PCNA | 12182.5715 | 0.8214 | ##### | 6.9979 | 2.5988E-12 | 1.3288E-11 |
| ENSG00000136811\|ODF2 | 7642.7325 | 0.8138 | ##### | 6.9974 | 2.6082E-12 | 1.3333E-11 |
| ENSG00000166839\|ANKDD1A | 256.5249 | 1.1828 | ##### | 6.9973 | 2.6092E-12 | 1.3336E-11 |
| ENSG00000120458\|MSANTD2 | 797.4643 | 1.0063 | ##### | 6.9968 | 2.6181E-12 | 1.3379E-11 |
| ENSG00000188611\|ASAH2 | 283.0588 | -1.1626 | ##### | -6.9960 | 2.6328E-12 | 1.3452E-11 |
| ENSG00000116786\|PLEKHM2 | 7916.8551 | -0.8318 | ##### | -6.9940 | 2.6709E-12 | 1.3644E-11 |
| ENSG00000196696\|PDXDC2P-NPIPB14P | 491.0894 | -1.0650 | ##### | -6.9940 | 2.6723E-12 | 1.3650E-11 |
| ENSG00000143633\|C1orf131 | 1251.1826 | -0.9074 | ##### | -6.9935 | 2.6813E-12 | 1.3693E-11 |
| ENSG00000151332\|MBIP | 1507.6480 | 0.8667 | ##### | 6.9930 | 2.6909E-12 | 1.3740E-11 |
| ENSG00000177340\|FLJ13224 | 29.1438 | 3.4358 | ##### | 6.9913 | 2.7235E-12 | 1.3904E-11 |
| ENSG00000158169\|FANCC | 2809.6433 | -0.8798 | ##### | -6.9907 | 2.7354E-12 | 1.3962E-11 |
| ENSG00000100583\|SAMD15 | 229.8924 | 1.1589 | ##### | 6.9903 | 2.7435E-12 | 1.4001E-11 |
| ENSG00000118193\|KIF14 | 4190.5150 | 0.8702 | ##### | 6.9894 | 2.7599E-12 | 1.4083E-11 |
| ENSG00000162636\|EEIG2 | 5020.3011 | -0.8560 | ##### | -6.9890 | 2.7681E-12 | 1.4122E-11 |
| ENSG00000185100\|ADSS1 | 623.9238 | -1.0038 | ##### | -6.9883 | 2.7831E-12 | 1.4196E-11 |
| ENSG00000197586\|ENTPD6 | 4993.2975 | -0.8185 | ##### | -6.9880 | 2.7880E-12 | 1.4219E-11 |
| ENSG00000175582\|RAB6A | 13462.5932 | 0.7864 | ##### | 6.9866 | 2.8161E-12 | 1.4359E-11 |
| ENSG00000182162\|P2RY8 | 115.9646 | 8.7628 | ##### | 6.9864 | 2.8202E-12 | 1.4378E-11 |

| | | | | | | |
|---|---|---|---|---|---|---|
| ENSG00000169926\|KLF13 | 6955.7726 | -0.8487 | ##### | -6.9854 | 2.8396E-12 | 1.4474E-11 |
| ENSG00000143337\|TOR1AIP1 | 6368.6734 | -0.7599 | ##### | -6.9842 | 2.8648E-12 | 1.4600E-11 |
| ENSG00000153044\|CENPH | 1713.4224 | 0.9173 | ##### | 6.9833 | 2.8843E-12 | 1.4697E-11 |
| ENSG00000135953\|MFSD9 | 1811.2156 | -0.8828 | ##### | -6.9803 | 2.9464E-12 | 1.5011E-11 |
| ENSG00000229851\|ARSD-AS1 | 34.1871 | -2.3867 | ##### | -6.9800 | 2.9518E-12 | 1.5036E-11 |
| ENSG00000169372\|CRADD | 437.5318 | -1.1184 | ##### | -6.9777 | 3.0000E-12 | 1.5279E-11 |
| ENSG00000158006\|PAFAH2 | 1426.8800 | -0.9008 | ##### | -6.9771 | 3.0139E-12 | 1.5347E-11 |
| ENSG00000116254\|CHD5 | 626.3268 | 1.1008 | ##### | 6.9748 | 3.0620E-12 | 1.5589E-11 |
| ENSG00000135913\|USP37 | 3914.8933 | 0.8706 | ##### | 6.9747 | 3.0655E-12 | 1.5605E-11 |
| ENSG00000129521\|EGLN3 | 289.5276 | -1.3796 | ##### | -6.9741 | 3.0791E-12 | 1.5671E-11 |
| ENSG00000287921 | 23.9602 | -3.0994 | ##### | -6.9722 | 3.1193E-12 | 1.5873E-11 |
| ENSG00000111961\|SASH1 | 2025.6038 | 0.8805 | ##### | 6.9722 | 3.1212E-12 | 1.5880E-11 |
| ENSG00000142920\|AZIN2 | 229.2087 | -1.2465 | ##### | -6.9717 | 3.1304E-12 | 1.5924E-11 |
| ENSG00000291146\|RP9P | 908.6347 | -0.9891 | ##### | -6.9714 | 3.1384E-12 | 1.5962E-11 |
| ENSG00000160783\|PMF1 | 4078.2575 | 0.9095 | ##### | 6.9710 | 3.1461E-12 | 1.5998E-11 |
| ENSG00000120733\|KDM3B | 8088.6021 | 0.8522 | ##### | 6.9710 | 3.1478E-12 | 1.6004E-11 |
| ENSG00000163220\|S100A9 | 479.7895 | -7.2144 | ##### | -6.9705 | 3.1574E-12 | 1.6050E-11 |
| ENSG00000172339\|ALG14 | 680.3985 | -1.1289 | ##### | -6.9702 | 3.1642E-12 | 1.6082E-11 |
| ENSG00000088812\|ATRN | 5218.0038 | -0.8469 | ##### | -6.9702 | 3.1658E-12 | 1.6088E-11 |
| ENSG00000177409\|SAMD9L | 696.8827 | -9.6690 | ##### | -6.9686 | 3.2000E-12 | 1.6259E-11 |
| ENSG00000279673 | 72.0244 | -1.8687 | ##### | -6.9684 | 3.2061E-12 | 1.6287E-11 |
| ENSG00000143179\|UCK2 | 10420.4907 | -0.6880 | ##### | -6.9680 | 3.2158E-12 | 1.6333E-11 |
| ENSG00000290047 | 49.0710 | 7.6293 | ##### | 6.9668 | 3.2415E-12 | 1.6461E-11 |
| ENSG00000077585\|GPR137B | 178.2138 | 1.2656 | ##### | 6.9666 | 3.2471E-12 | 1.6487E-11 |
| ENSG00000184163\|C1QTNF12 | 101.7489 | -1.4144 | ##### | -6.9663 | 3.2544E-12 | 1.6521E-11 |
| ENSG00000225963 | 18.0766 | -4.7835 | ##### | -6.9662 | 3.2566E-12 | 1.6529E-11 |
| ENSG00000181450\|ZNF678 | 623.7159 | -0.9454 | ##### | -6.9640 | 3.3085E-12 | 1.6790E-11 |
| ENSG00000136492\|BRIP1 | 1651.1248 | 0.9238 | ##### | 6.9636 | 3.3165E-12 | 1.6828E-11 |
| ENSG00000100578\|KIAA0586 | 2379.3575 | 0.8765 | ##### | 6.9627 | 3.3383E-12 | 1.6935E-11 |
| ENSG00000179820\|MYADM | 20641.3164 | -0.8572 | ##### | -6.9608 | 3.3841E-12 | 1.7165E-11 |
| ENSG00000177045\|SIX5 | 4215.9725 | -0.8306 | ##### | -6.9592 | 3.4218E-12 | 1.7353E-11 |
| ENSG00000006327\|TNFRSF12A | 19328.5847 | -0.8492 | ##### | -6.9581 | 3.4494E-12 | 1.7490E-11 |
| ENSG00000124257\|NEURL2 | 55.7659 | 2.6592 | ##### | 6.9579 | 3.4550E-12 | 1.7515E-11 |
| ENSG00000159377\|PSMB4 | 27560.8307 | 0.8422 | ##### | 6.9576 | 3.4619E-12 | 1.7547E-11 |
| ENSG00000115159\|GPD2 | 9446.4633 | -0.8227 | ##### | -6.9574 | 3.4649E-12 | 1.7560E-11 |
| ENSG00000110713\|NUP98 | 13166.8607 | 0.8391 | ##### | 6.9562 | 3.4947E-12 | 1.7708E-11 |
| ENSG00000167136\|ENDOG | 2157.2694 | -0.9687 | ##### | -6.9542 | 3.5454E-12 | 1.7962E-11 |
| ENSG00000204072\|ARMCX7P | 100.0004 | 1.5041 | ##### | 6.9537 | 3.5576E-12 | 1.8020E-11 |
| ENSG00000264462\|MIR3648-2 | 209.8900 | 2.2746 | ##### | 6.9533 | 3.5689E-12 | 1.8074E-11 |
| ENSG00000182050\|MGAT4C | 128.8116 | -1.8836 | ##### | -6.9525 | 3.5885E-12 | 1.8171E-11 |
| ENSG00000107669\|ATE1 | 3285.9446 | -0.6991 | ##### | -6.9519 | 3.6052E-12 | 1.8252E-11 |
| ENSG00000104853\|CLPTM1 | 14151.5369 | -0.8238 | ##### | -6.9514 | 3.6163E-12 | 1.8305E-11 |
| ENSG00000172315\|TP53RK | 2517.2712 | 0.8702 | ##### | 6.9511 | 3.6247E-12 | 1.8344E-11 |
| ENSG00000247199\|FBXO38-DT | 90.8682 | -1.8172 | ##### | -6.9498 | 3.6580E-12 | 1.8510E-11 |
| ENSG00000143612\|C1orf43 | 23314.3031 | 0.6375 | ##### | 6.9497 | 3.6606E-12 | 1.8520E-11 |
| ENSG00000114446\|IFT57 | 3849.2355 | 0.8596 | ##### | 6.9489 | 3.6823E-12 | 1.8627E-11 |
| ENSG00000260005 | 790.2833 | 1.0246 | ##### | 6.9485 | 3.6919E-12 | 1.8672E-11 |
| ENSG00000060069\|CTDP1 | 5053.6934 | 0.8726 | ##### | 6.9477 | 3.7116E-12 | 1.8769E-11 |
| ENSG00000233728 | 453.4475 | -1.0616 | ##### | -6.9466 | 3.7423E-12 | 1.8921E-11 |
| ENSG00000170430\|MGMT | 847.0176 | -0.9476 | ##### | -6.9453 | 3.7778E-12 | 1.9097E-11 |
| ENSG00000268970 | 41.4029 | -2.5729 | ##### | -6.9436 | 3.8227E-12 | 1.9321E-11 |
| ENSG00000108883\|EFTUD2 | 22527.5124 | 0.7133 | ##### | 6.9434 | 3.8267E-12 | 1.9337E-11 |
| ENSG00000262831 | 4437.2796 | -0.8784 | ##### | -6.9429 | 3.8404E-12 | 1.9403E-11 |
| ENSG00000171148\|TADA3 | 6695.3089 | 0.8515 | ##### | 6.9405 | 3.9083E-12 | 1.9743E-11 |
| ENSG00000137038\|DMAC1 | 1319.9757 | -0.9778 | ##### | -6.9402 | 3.9153E-12 | 1.9775E-11 |
| ENSG00000204054\|LINC00963 | 3748.0124 | -0.8827 | ##### | -6.9383 | 3.9688E-12 | 2.0042E-11 |
| ENSG00000290839 | 39.6221 | -3.1131 | ##### | -6.9374 | 3.9937E-12 | 2.0165E-11 |
| ENSG00000167842\|MIS12 | 1724.8654 | 0.8944 | ##### | 6.9364 | 4.0227E-12 | 2.0307E-11 |
| ENSG00000170584\|NUDCD2 | 3216.6794 | 0.8788 | ##### | 6.9363 | 4.0239E-12 | 2.0310E-11 |
| ENSG00000100296\|THOC5 | 5122.0225 | 0.7434 | ##### | 6.9340 | 4.0906E-12 | 2.0643E-11 |
| ENSG00000145191\|EIF2B5 | 3877.6927 | -0.8130 | ##### | -6.9334 | 4.1091E-12 | 2.0733E-11 |
| ENSG00000135443\|KRT85 | 19781.7183 | ####### | ##### | -6.9314 | 4.1673E-12 | 2.1023E-11 |
| ENSG00000115902\|SLC1A4 | 1410.6511 | -0.9544 | ##### | -6.9278 | 4.2749E-12 | 2.1562E-11 |

| | | | | | | |
|---|---|---|---|---|---|---|
| ENSG00000110395\|CBL | 8134.3821 | 0.8512 | ##### | 6.9276 | 4.2816E-12 | 2.1592E-11 |
| ENSG00000273143\|DUSP5-DT | 289.8074 | ####### | ##### | -6.9274 | 4.2859E-12 | 2.1610E-11 |
| ENSG00000271533 | 128.8004 | 1.0987 | ##### | 6.9257 | 4.3395E-12 | 2.1877E-11 |
| ENSG00000153922\|CHD1 | 5412.0487 | 0.8418 | ##### | 6.9249 | 4.3632E-12 | 2.1993E-11 |
| ENSG00000131788\|PIAS3 | 6765.5330 | 0.8502 | ##### | 6.9243 | 4.3801E-12 | 2.2071E-11 |
| ENSG00000237357\|FAM88E | 94.4330 | -1.7868 | ##### | -6.9243 | 4.3802E-12 | 2.2071E-11 |
| ENSG00000161013\|MGAT4B | 11063.2847 | -0.7039 | ##### | -6.9234 | 4.4108E-12 | 2.2221E-11 |
| ENSG00000221994\|ZNF630 | 77.5277 | 1.8306 | ##### | 6.9221 | 4.4512E-12 | 2.2421E-11 |
| ENSG00000080802\|CNOT4 | 2917.4331 | 0.8375 | ##### | 6.9214 | 4.4734E-12 | 2.2529E-11 |
| ENSG00000198301\|SDAD1 | 4581.5466 | 0.8134 | ##### | 6.9213 | 4.4761E-12 | 2.2539E-11 |
| ENSG00000167103\|PIP5KL1 | 300.7965 | 1.1832 | ##### | 6.9202 | 4.5103E-12 | 2.2708E-11 |
| ENSG00000175792\|RUVBL1 | 14229.0569 | 0.7857 | ##### | 6.9196 | 4.5299E-12 | 2.2802E-11 |
| ENSG00000260808 | 40.3906 | 2.1350 | ##### | 6.9195 | 4.5339E-12 | 2.2818E-11 |
| ENSG00000264207 | 79.9920 | 1.5170 | ##### | 6.9189 | 4.5530E-12 | 2.2911E-11 |
| ENSG00000071189\|SNX13 | 4190.0240 | -0.8437 | ##### | -6.9171 | 4.6096E-12 | 2.3192E-11 |
| ENSG00000286522\|H3C2 | 49.4020 | 3.2516 | ##### | 6.9154 | 4.6662E-12 | 2.3472E-11 |
| ENSG00000170873\|MTSS1 | 1332.4254 | -0.9160 | ##### | -6.9145 | 4.6949E-12 | 2.3613E-11 |
| ENSG00000167705\|RILP | 522.5333 | 1.0737 | ##### | 6.9140 | 4.7129E-12 | 2.3700E-11 |
| ENSG00000152520\|PAN3 | 2250.9630 | 0.8979 | ##### | 6.9134 | 4.7303E-12 | 2.3783E-11 |
| ENSG00000066230\|SLC9A3 | 1014.6867 | -2.0496 | ##### | -6.9100 | 4.8464E-12 | 2.4362E-11 |
| ENSG00000154832\|CXXC1 | 6517.3332 | 0.8191 | ##### | 6.9096 | 4.8590E-12 | 2.4418E-11 |
| ENSG00000142544\|CTU1 | 1441.0544 | -0.8974 | ##### | -6.9096 | 4.8588E-12 | 2.4418E-11 |
| ENSG00000154743\|TSEN2 | 1558.0613 | 0.9106 | ##### | 6.9095 | 4.8623E-12 | 2.4430E-11 |
| ENSG00000203852\|H3C15 | 80.0199 | 1.7441 | ##### | 6.9086 | 4.8959E-12 | 2.4595E-11 |
| ENSG00000184903\|IMMP2L | 557.3653 | -1.0076 | ##### | -6.9072 | 4.9448E-12 | 2.4836E-11 |
| ENSG00000110074\|FOXRED1 | 1862.0582 | 0.8584 | ##### | 6.9066 | 4.9642E-12 | 2.4930E-11 |
| ENSG00000197608\|ZNF841 | 770.8011 | -0.9849 | ##### | -6.9062 | 4.9783E-12 | 2.4996E-11 |
| ENSG00000174469\|CNTNAP2 | 123.1157 | 7.2570 | ##### | 6.9040 | 5.0542E-12 | 2.5373E-11 |
| ENSG00000188315\|C3orf62 | 391.6401 | 1.1063 | ##### | 6.9038 | 5.0634E-12 | 2.5415E-11 |
| ENSG00000203811\|H3C14 | 81.2918 | 1.7338 | ##### | 6.9033 | 5.0802E-12 | 2.5495E-11 |
| ENSG00000256150\|ITFG2-AS1 | 30.6511 | 2.5424 | ##### | 6.9031 | 5.0880E-12 | 2.5530E-11 |
| ENSG00000292982\|WASH5P | 565.6235 | 1.1535 | ##### | 6.9026 | 5.1076E-12 | 2.5624E-11 |
| ENSG00000153904\|DDAH1 | 4227.1706 | -0.8455 | ##### | -6.9025 | 5.1094E-12 | 2.5628E-11 |
| ENSG00000166192\|SENP8 | 123.9441 | -1.5816 | ##### | -6.9020 | 5.1290E-12 | 2.5722E-11 |
| ENSG00000081479\|LRP2 | 1093.2218 | 11.9893 | ##### | 6.9002 | 5.1933E-12 | 2.6041E-11 |
| ENSG00000116791\|CRYZ | 9614.1306 | -0.8418 | ##### | -6.8985 | 5.2562E-12 | 2.6352E-11 |
| ENSG00000248223 | 23.1321 | -2.7422 | ##### | -6.8978 | 5.2798E-12 | 2.6465E-11 |
| ENSG00000183718\|TRIM52 | 736.7903 | 0.9341 | ##### | 6.8972 | 5.3021E-12 | 2.6573E-11 |
| ENSG00000140284\|SLC27A2 | 729.5363 | -1.0330 | ##### | -6.8971 | 5.3067E-12 | 2.6591E-11 |
| ENSG00000068489\|PRR11 | 9681.0932 | 0.8542 | ##### | 6.8971 | 5.3075E-12 | 2.6591E-11 |
| ENSG00000005075\|POLR2J | 4317.8003 | 0.7276 | ##### | 6.8963 | 5.3358E-12 | 2.6728E-11 |
| ENSG00000121741\|ZMYM2 | 6944.3616 | 0.8258 | ##### | 6.8963 | 5.3383E-12 | 2.6736E-11 |
| ENSG00000225663\|MCRIP1 | 2791.8740 | 0.8645 | ##### | 6.8959 | 5.3535E-12 | 2.6808E-11 |
| ENSG00000285943 | 237.6813 | 1.3670 | ##### | 6.8948 | 5.3931E-12 | 2.7002E-11 |
| ENSG00000102218\|RP2 | 1784.5375 | -0.9085 | ##### | -6.8947 | 5.3968E-12 | 2.7016E-11 |
| ENSG00000011198\|ABHD5 | 1530.8196 | -0.8547 | ##### | -6.8929 | 5.4650E-12 | 2.7352E-11 |
| ENSG00000152779\|SLC16A12 | 60.0806 | -5.4237 | ##### | -6.8923 | 5.4894E-12 | 2.7470E-11 |
| ENSG00000153233\|PTPRR | 78.3828 | 3.3905 | ##### | 6.8920 | 5.4995E-12 | 2.7516E-11 |
| ENSG00000242802\|AP5Z1 | 4617.4919 | 0.8577 | ##### | 6.8919 | 5.5036E-12 | 2.7532E-11 |
| ENSG00000117280\|RAB29 | 2055.6111 | -0.9021 | ##### | -6.8900 | 5.5790E-12 | 2.7904E-11 |
| ENSG00000163714\|U2SURP | 17701.2812 | 0.8209 | ##### | 6.8868 | 5.7073E-12 | 2.8541E-11 |
| ENSG00000101213\|PTK6 | 291.0396 | -1.1251 | ##### | -6.8860 | 5.7378E-12 | 2.8689E-11 |
| ENSG00000158427\|TMSB15B | 231.3199 | 1.5658 | ##### | 6.8853 | 5.7676E-12 | 2.8833E-11 |
| ENSG00000105926\|PALS2 | 4138.3928 | 0.8359 | ##### | 6.8817 | 5.9146E-12 | 2.9563E-11 |
| ENSG00000140400\|MAN2C1 | 3919.1682 | -0.7563 | ##### | -6.8807 | 5.9570E-12 | 2.9770E-11 |
| ENSG00000131435\|PDLIM4 | 1244.1635 | 11.9345 | ##### | 6.8805 | 5.9642E-12 | 2.9801E-11 |
| ENSG00000250186 | 5563.2075 | 0.9360 | ##### | 6.8777 | 6.0811E-12 | 3.0380E-11 |
| ENSG00000261408\|TEN1-CDK3 | 1302.3849 | 0.9752 | ##### | 6.8773 | 6.1004E-12 | 3.0471E-11 |
| ENSG00000103126\|AXIN1 | 2762.0553 | -0.8393 | ##### | -6.8764 | 6.1382E-12 | 3.0655E-11 |
| ENSG00000113272\|THG1L | 1215.9417 | 0.9612 | ##### | 6.8761 | 6.1508E-12 | 3.0713E-11 |
| ENSG00000290585 | 38.8360 | 2.8662 | ##### | 6.8753 | 6.1855E-12 | 3.0881E-11 |
| ENSG00000085563\|ABCB1 | 93.8750 | 1.8004 | ##### | 6.8737 | 6.2537E-12 | 3.1216E-11 |
| ENSG00000132424\|PNISR | 4279.8784 | 0.8474 | ##### | 6.8733 | 6.2738E-12 | 3.1311E-11 |

| | | | | | | |
|---|---|---|---|---|---|---|
| ENSG00000171004\|HS6ST2 | 1125.3153 | 5.9959 | ##### | 6.8720 | 6.3316E-12 | 3.1595E-11 |
| ENSG00000144134\|RABL2A | 440.8554 | -1.0379 | ##### | -6.8708 | 6.3844E-12 | 3.1853E-11 |
| ENSG00000290931 | 399.5386 | 0.9956 | ##### | 6.8704 | 6.4031E-12 | 3.1941E-11 |
| ENSG00000198857\|HSD3BP5 | 54.4287 | 7.3727 | ##### | 6.8703 | 6.4073E-12 | 3.1956E-11 |
| ENSG00000068079\|IFI35 | 828.7997 | -0.9345 | ##### | -6.8701 | 6.4168E-12 | 3.1998E-11 |
| ENSG00000173548\|SNX33 | 1572.8005 | -0.9036 | ##### | -6.8696 | 6.4371E-12 | 3.2094E-11 |
| ENSG00000106105\|GARS1 | 19686.7477 | -0.6617 | ##### | -6.8696 | 6.4391E-12 | 3.2099E-11 |
| ENSG00000234745\|HLA-B | 10498.3438 | 0.8188 | ##### | 6.8685 | 6.4873E-12 | 3.2334E-11 |
| ENSG00000143393\|PI4KB | 7414.7665 | 0.8364 | ##### | 6.8679 | 6.5160E-12 | 3.2471E-11 |
| ENSG00000159479\|MED8 | 4340.4636 | -0.8504 | ##### | -6.8663 | 6.5902E-12 | 3.2836E-11 |
| ENSG00000177425\|PAWR | 9753.0395 | -0.5948 | ##### | -6.8662 | 6.5922E-12 | 3.2840E-11 |
| ENSG00000174684\|B4GAT1 | 993.8737 | -0.8863 | ##### | -6.8637 | 6.7089E-12 | 3.3416E-11 |
| ENSG00000290871\|GTF2IP1 | 6434.5283 | -0.8261 | ##### | -6.8629 | 6.7459E-12 | 3.3595E-11 |
| ENSG00000187736\|NHEJ1 | 2882.3684 | 0.8793 | ##### | 6.8629 | 6.7486E-12 | 3.3602E-11 |
| ENSG00000144199\|FAHD2B | 1126.1999 | 0.9412 | ##### | 6.8624 | 6.7724E-12 | 3.3715E-11 |
| ENSG00000158055\|GRHL3 | 151.6625 | 2.7724 | ##### | 6.8622 | 6.7803E-12 | 3.3749E-11 |
| ENSG00000175764\|TTLL11 | 414.3007 | -1.0536 | ##### | -6.8613 | 6.8255E-12 | 3.3968E-11 |
| ENSG00000228463 | 27.2062 | -2.6300 | ##### | -6.8610 | 6.8397E-12 | 3.4033E-11 |
| ENSG00000104881\|PPP1R13L | 3420.3274 | -0.8344 | ##### | -6.8605 | 6.8633E-12 | 3.4145E-11 |
| ENSG00000124508\|BTN2A2 | 823.8053 | 0.9335 | ##### | 6.8593 | 6.9188E-12 | 3.4415E-11 |
| ENSG00000139620\|KANSL2 | 3189.6079 | 0.6582 | ##### | 6.8593 | 6.9219E-12 | 3.4425E-11 |
| ENSG00000162929\|SANBR | 1668.5181 | -0.8572 | ##### | -6.8592 | 6.9264E-12 | 3.4442E-11 |
| ENSG00000185896\|LAMP1 | 13785.3650 | -0.5439 | ##### | -6.8590 | 6.9368E-12 | 3.4488E-11 |
| ENSG00000188958\|UTS2B | 34.7553 | -2.1286 | ##### | -6.8587 | 6.9510E-12 | 3.4552E-11 |
| ENSG00000181619\|GPR135 | 287.7028 | 1.2385 | ##### | 6.8578 | 6.9912E-12 | 3.4747E-11 |
| ENSG00000156531\|PHF6 | 4907.4923 | 0.8359 | ##### | 6.8571 | 7.0266E-12 | 3.4917E-11 |
| ENSG00000233223\|MPDU1-AS1 | 332.3373 | 1.2615 | ##### | 6.8532 | 7.2228E-12 | 3.5886E-11 |
| ENSG00000176046\|NUPR1 | 1416.5475 | -8.7482 | ##### | -6.8524 | 7.2639E-12 | 3.6084E-11 |
| ENSG00000275055 | 67.0687 | -1.7388 | ##### | -6.8521 | 7.2756E-12 | 3.6136E-11 |
| ENSG00000164484\|TMEM200A | 412.8964 | 6.6355 | ##### | 6.8514 | 7.3143E-12 | 3.6322E-11 |
| ENSG00000151552\|QDPR | 1624.7511 | -0.8680 | ##### | -6.8500 | 7.3845E-12 | 3.6665E-11 |
| ENSG00000240498\|CDKN2B-AS1 | 229.3090 | 1.8837 | ##### | 6.8498 | 7.3978E-12 | 3.6724E-11 |
| ENSG00000006831\|ADIPOR2 | 17139.2588 | 0.7554 | ##### | 6.8493 | 7.4216E-12 | 3.6837E-11 |
| ENSG00000230844\|ZNF674-AS1 | 304.0005 | -1.1986 | ##### | -6.8451 | 7.6448E-12 | 3.7938E-11 |
| ENSG00000108561\|C1QBP | 21944.8853 | 0.7256 | ##### | 6.8440 | 7.6992E-12 | 3.8196E-11 |
| ENSG00000284308\|C2orf81 | 558.9644 | 1.2058 | ##### | 6.8440 | 7.6990E-12 | 3.8196E-11 |
| ENSG00000144659\|SLC25A38 | 2310.2089 | 0.7892 | ##### | 6.8437 | 7.7194E-12 | 3.8289E-11 |
| ENSG00000156140\|ADAMTS3 | 1179.7096 | 0.9442 | ##### | 6.8434 | 7.7324E-12 | 3.8348E-11 |
| ENSG00000187838\|PLSCR3 | 1988.2682 | -0.8969 | ##### | -6.8421 | 7.8066E-12 | 3.8709E-11 |
| ENSG00000072756\|TRNT1 | 3461.3279 | 0.8751 | ##### | 6.8399 | 7.9262E-12 | 3.9295E-11 |
| ENSG00000273514\|FOXD4L6 | 19.1284 | 5.2883 | ##### | 6.8362 | 8.1339E-12 | 4.0319E-11 |
| ENSG00000187514\|PTMA | 87304.3474 | 0.7967 | ##### | 6.8338 | 8.2683E-12 | 4.0978E-11 |
| ENSG00000126903\|SLC10A3 | 4020.7740 | 0.8558 | ##### | 6.8332 | 8.3014E-12 | 4.1135E-11 |
| ENSG00000124802\|EEF1E1 | 2188.6846 | 0.8920 | ##### | 6.8323 | 8.3549E-12 | 4.1393E-11 |
| ENSG00000272142\|LYRM4-AS1 | 444.2092 | 1.0584 | ##### | 6.8307 | 8.4491E-12 | 4.1853E-11 |
| ENSG00000154310\|TNIK | 429.3260 | -1.0329 | ##### | -6.8305 | 8.4607E-12 | 4.1904E-11 |
| ENSG00000109339\|MAPK10 | 138.0560 | -1.6512 | ##### | -6.8280 | 8.6107E-12 | 4.2639E-11 |
| ENSG00000165684\|SNAPC4 | 2661.4841 | 0.8426 | ##### | 6.8274 | 8.6474E-12 | 4.2814E-11 |
| ENSG00000122694\|GLIPR2 | 907.5444 | 0.9338 | ##### | 6.8270 | 8.6684E-12 | 4.2911E-11 |
| ENSG00000112852\|PCDHB2 | 301.9624 | -6.3610 | ##### | -6.8259 | 8.7355E-12 | 4.3236E-11 |
| ENSG00000133226\|SRRM1 | 9529.4538 | 0.8390 | ##### | 6.8259 | 8.7374E-12 | 4.3238E-11 |
| ENSG00000067704\|IARS2 | 14415.7786 | -0.4976 | ##### | -6.8257 | 8.7498E-12 | 4.3292E-11 |
| ENSG00000103642\|LACTB | 1700.8219 | -0.9090 | ##### | -6.8245 | 8.8214E-12 | 4.3640E-11 |
| ENSG00000164151\|ICE1 | 4892.3486 | -0.8275 | ##### | -6.8242 | 8.8413E-12 | 4.3731E-11 |
| ENSG00000162769\|FLVCR1 | 2862.4381 | -0.8655 | ##### | -6.8229 | 8.9231E-12 | 4.4128E-11 |
| ENSG00000101138\|CSTF1 | 4022.4335 | 0.8458 | ##### | 6.8226 | 8.9401E-12 | 4.4205E-11 |
| ENSG00000236279\|CLEC2L | 60.8829 | -2.2695 | ##### | -6.8226 | 8.9418E-12 | 4.4206E-11 |
| ENSG00000182952\|HMGN4 | 4469.0003 | 0.7019 | ##### | 6.8220 | 8.9796E-12 | 4.4385E-11 |
| ENSG00000286790 | 68.7769 | -2.3047 | ##### | -6.8219 | 8.9850E-12 | 4.4404E-11 |
| ENSG00000105088\|OLFM2 | 3063.5780 | 0.8727 | ##### | 6.8215 | 9.0077E-12 | 4.4509E-11 |
| ENSG00000258986\|TMEM179 | 180.8181 | 1.5306 | ##### | 6.8212 | 9.0306E-12 | 4.4615E-11 |
| ENSG00000144713\|RPL32 | 29123.2716 | 0.8318 | ##### | 6.8205 | 9.0751E-12 | 4.4827E-11 |
| ENSG00000187866\|PABIR1 | 1041.5015 | -0.9560 | ##### | -6.8204 | 9.0782E-12 | 4.4835E-11 |

| | | | | | | |
|---|---|---|---|---|---|---|
| ENSG00000156853\|ZNF689 | 1649.9010 | 0.8989 | ##### | 6.8191 | 9.1590E-12 | 4.5227E-11 |
| ENSG00000169641\|LUZP1 | 8081.7983 | 0.8134 | ##### | 6.8188 | 9.1812E-12 | 4.5329E-11 |
| ENSG00000128573\|FOXP2 | 73.2040 | -1.9963 | ##### | -6.8174 | 9.2681E-12 | 4.5751E-11 |
| ENSG00000256973 | 28.9183 | 6.5024 | ##### | 6.8152 | 9.4132E-12 | 4.6459E-11 |
| ENSG00000115504\|EHBP1 | 10178.4534 | -0.7842 | ##### | -6.8150 | 9.4252E-12 | 4.6511E-11 |
| ENSG00000111305\|GSG1 | 48.1852 | 7.6315 | ##### | 6.8147 | 9.4484E-12 | 4.6617E-11 |
| ENSG00000168575\|SLC20A2 | 2735.8656 | 0.9023 | ##### | 6.8144 | 9.4650E-12 | 4.6692E-11 |
| ENSG00000233922\|LINC01694 | 193.6710 | -1.2260 | ##### | -6.8143 | 9.4737E-12 | 4.6727E-11 |
| ENSG00000099956\|SMARCB1 | 7403.1395 | 0.8406 | ##### | 6.8139 | 9.4992E-12 | 4.6845E-11 |
| ENSG00000135185\|TMEM243 | 1077.2074 | 0.8346 | ##### | 6.8139 | 9.5012E-12 | 4.6847E-11 |
| ENSG00000008311\|AASS | 135.6315 | 1.7032 | ##### | 6.8134 | 9.5288E-12 | 4.6975E-11 |
| ENSG00000162670\|BRINP3 | 907.9028 | -0.9838 | ##### | -6.8122 | 9.6127E-12 | 4.7381E-11 |
| ENSG00000108349\|CASC3 | 10335.5480 | 0.8480 | ##### | 6.8116 | 9.6529E-12 | 4.7571E-11 |
| ENSG00000163393\|SLC22A15 | 166.7983 | -1.3006 | ##### | -6.8090 | 9.8278E-12 | 4.8425E-11 |
| ENSG00000183474\|GTF2H2C | 3051.3769 | 0.8269 | ##### | 6.8085 | 9.8626E-12 | 4.8589E-11 |
| ENSG00000154027\|AK5 | 413.6472 | -8.0368 | ##### | -6.8083 | 9.8751E-12 | 4.8642E-11 |
| ENSG00000011258\|MBTD1 | 1460.5558 | 0.9165 | ##### | 6.8077 | 9.9154E-12 | 4.8833E-11 |
| ENSG00000134283\|PPHLN1 | 7662.3847 | 0.6258 | ##### | 6.8068 | 9.9781E-12 | 4.9134E-11 |
| ENSG00000124120\|TTPAL | 6802.4486 | 0.8511 | ##### | 6.8055 | 1.0071E-11 | 4.9581E-11 |
| ENSG00000273274\|ZBTB8B | 79.1003 | 5.0381 | ##### | 6.8039 | 1.0181E-11 | 5.0115E-11 |
| ENSG00000225684\|FAM225B | 1152.7592 | 11.7017 | ##### | 6.8037 | 1.0198E-11 | 5.0191E-11 |
| ENSG00000196268\|ZNF493 | 44.2821 | -3.5069 | ##### | -6.8030 | 1.0247E-11 | 5.0424E-11 |
| ENSG00000107021\|TBC1D13 | 3323.0412 | -0.8094 | ##### | -6.8025 | 1.0284E-11 | 5.0598E-11 |
| ENSG00000069493\|CLEC2D | 176.1272 | 1.4335 | ##### | 6.8022 | 1.0305E-11 | 5.0695E-11 |
| ENSG00000233485\|FHAD1-AS1 | 50.2582 | 7.9707 | ##### | 6.7994 | 1.0502E-11 | 5.1656E-11 |
| ENSG00000141873\|SLC39A3 | 3280.7521 | 0.8118 | ##### | 6.7991 | 1.0531E-11 | 5.1789E-11 |
| ENSG00000175155\|YPEL2 | 193.2494 | -1.4067 | ##### | -6.7989 | 1.0539E-11 | 5.1820E-11 |
| ENSG00000163249\|CCNYL1 | 5346.9305 | -0.8193 | ##### | -6.7985 | 1.0575E-11 | 5.1986E-11 |
| ENSG00000156966\|B3GNT7 | 65.4438 | -1.9072 | ##### | -6.7984 | 1.0580E-11 | 5.2005E-11 |
| ENSG00000160111\|CPAMD8 | 53.7426 | 3.1019 | ##### | 6.7961 | 1.0749E-11 | 5.2823E-11 |
| ENSG00000183340\|JRKL | 846.6048 | -0.9417 | ##### | -6.7959 | 1.0761E-11 | 5.2874E-11 |
| ENSG00000103351\|CLUAP1 | 1091.4255 | -0.9013 | ##### | -6.7947 | 1.0854E-11 | 5.3322E-11 |
| ENSG00000188191\|PRKAR1B | 3428.0009 | -0.8647 | ##### | -6.7941 | 1.0897E-11 | 5.3527E-11 |
| ENSG00000244588\|RAD21L1 | 70.5298 | 3.2630 | ##### | 6.7941 | 1.0899E-11 | 5.3528E-11 |
| ENSG00000188060\|RAB42 | 636.4363 | -7.8306 | ##### | -6.7935 | 1.0945E-11 | 5.3746E-11 |
| ENSG00000136449\|MYCBPAP | 56.7661 | -2.3005 | ##### | -6.7919 | 1.1064E-11 | 5.4321E-11 |
| ENSG00000180855\|ZNF443 | 140.3882 | -1.6117 | ##### | -6.7907 | 1.1157E-11 | 5.4769E-11 |
| ENSG00000050405\|LIMA1 | 16594.4928 | -0.8294 | ##### | -6.7903 | 1.1189E-11 | 5.4914E-11 |
| ENSG00000206149\|HERC2P9 | 656.3570 | 0.9938 | ##### | 6.7895 | 1.1250E-11 | 5.5203E-11 |
| ENSG00000226564\|FTH1P20 | 29.4393 | -2.2376 | ##### | -6.7887 | 1.1312E-11 | 5.5501E-11 |
| ENSG00000101132\|PFDN4 | 3731.3189 | 0.8500 | ##### | 6.7887 | 1.1314E-11 | 5.5502E-11 |
| ENSG00000265994 | 43.4654 | -8.3757 | ##### | -6.7885 | 1.1329E-11 | 5.5564E-11 |
| ENSG00000271383\|NBPF19 | 2416.7899 | 0.9270 | ##### | 6.7884 | 1.1337E-11 | 5.5596E-11 |
| ENSG00000145287\|PLAC8 | 2222.2034 | 0.8924 | ##### | 6.7880 | 1.1371E-11 | 5.5755E-11 |
| ENSG00000286267 | 281.6570 | 9.6146 | ##### | 6.7876 | 1.1403E-11 | 5.5900E-11 |
| ENSG00000168243\|GNG4 | 1200.9878 | 0.9510 | ##### | 6.7860 | 1.1526E-11 | 5.6497E-11 |
| ENSG00000246876\|LINC02466 | 48.5552 | -8.0393 | ##### | -6.7846 | 1.1639E-11 | 5.7039E-11 |
| ENSG00000136986\|DERL1 | 10296.1881 | -0.8227 | ##### | -6.7825 | 1.1808E-11 | 5.7858E-11 |
| ENSG00000174236\|REP15 | 167.3646 | 1.4752 | ##### | 6.7825 | 1.1815E-11 | 5.7885E-11 |
| ENSG00000268307\|LINC02560 | 33.0313 | 7.5567 | ##### | 6.7804 | 1.1982E-11 | 5.8693E-11 |
| ENSG00000255949\|RPS6KB2-AS1 | 149.8376 | 1.5622 | ##### | 6.7794 | 1.2070E-11 | 5.9111E-11 |
| ENSG00000196550\|FAM72A | 996.1481 | 0.8805 | ##### | 6.7783 | 1.2163E-11 | 5.9557E-11 |
| ENSG00000173894\|CBX2 | 2405.1034 | -0.8503 | ##### | -6.7763 | 1.2328E-11 | 6.0354E-11 |
| ENSG00000156253\|RWDD2B | 1760.4149 | -0.8776 | ##### | -6.7761 | 1.2343E-11 | 6.0422E-11 |
| ENSG00000172974\|VDAC2P5 | 2598.4054 | -0.8106 | ##### | -6.7748 | 1.2459E-11 | 6.0979E-11 |
| ENSG00000205334\|LINC01460 | 71.8243 | 1.6982 | ##### | 6.7747 | 1.2463E-11 | 6.0989E-11 |
| ENSG00000100749\|VRK1 | 1863.9030 | 0.8620 | ##### | 6.7745 | 1.2484E-11 | 6.1083E-11 |
| ENSG00000237424\|FOXD2-AS1 | 555.1986 | 0.9774 | ##### | 6.7737 | 1.2549E-11 | 6.1390E-11 |
| ENSG00000149541\|B3GAT3 | 2388.8104 | -0.8603 | ##### | -6.7722 | 1.2680E-11 | 6.2017E-11 |
| ENSG00000177606\|JUN | 9422.8585 | 0.6414 | ##### | 6.7703 | 1.2849E-11 | 6.2835E-11 |
| ENSG00000226419\|SLC16A1-AS1 | 1050.5245 | -0.8770 | ##### | -6.7693 | 1.2940E-11 | 6.3269E-11 |
| ENSG00000116906\|GNPAT | 4657.0130 | -0.8324 | ##### | -6.7685 | 1.3012E-11 | 6.3612E-11 |
| ENSG00000258301\|VASH1-AS1 | 384.4282 | 1.1922 | ##### | 6.7664 | 1.3202E-11 | 6.4532E-11 |

| | | | | | | |
|---|---|---|---|---|---|---|
| ENSG00000125991\|ERGIC3 | 17568.9391 | 0.8130 | ##### | 6.7652 | 1.3314E-11 | 6.5066E-11 |
| ENSG00000250159\|PKD2L2-DT | 228.0763 | 7.5404 | ##### | 6.7638 | 1.3441E-11 | 6.5676E-11 |
| ENSG00000109654\|TRIM2 | 696.3213 | 0.9965 | ##### | 6.7637 | 1.3450E-11 | 6.5708E-11 |
| ENSG00000184445\|KNTC1 | 5538.6486 | 0.8400 | ##### | 6.7634 | 1.3482E-11 | 6.5857E-11 |
| ENSG00000163611\|SPICE1 | 1319.9707 | 0.8932 | ##### | 6.7632 | 1.3495E-11 | 6.5910E-11 |
| ENSG00000179542\|SLITRK4 | 133.3503 | 1.5051 | ##### | 6.7607 | 1.3733E-11 | 6.7059E-11 |
| ENSG00000092203\|TOX4 | 6311.5492 | 0.8310 | ##### | 6.7593 | 1.3867E-11 | 6.7701E-11 |
| ENSG00000197841\|ZNF181 | 666.2553 | -1.0693 | ##### | -6.7591 | 1.3889E-11 | 6.7801E-11 |
| ENSG00000158716\|DUSP23 | 4336.9794 | 0.8423 | ##### | 6.7589 | 1.3909E-11 | 6.7885E-11 |
| ENSG00000169760\|NLGN1 | 253.9365 | 1.2469 | ##### | 6.7587 | 1.3922E-11 | 6.7936E-11 |
| ENSG00000071859\|FAM50A | 8867.0916 | 0.8450 | ##### | 6.7582 | 1.3969E-11 | 6.8155E-11 |
| ENSG00000077454\|LRCH4 | 2525.3325 | 0.8906 | ##### | 6.7572 | 1.4069E-11 | 6.8635E-11 |
| ENSG00000225518\|LINC01703 | 60.0366 | -2.0993 | ##### | -6.7520 | 1.4578E-11 | 7.1106E-11 |
| ENSG00000275437 | 108.5363 | 2.2219 | ##### | 6.7517 | 1.4608E-11 | 7.1239E-11 |
| ENSG00000168661\|ZNF30 | 399.9782 | -1.1199 | ##### | -6.7512 | 1.4659E-11 | 7.1476E-11 |
| ENSG00000196177\|ACADSB | 1014.7140 | -0.8713 | ##### | -6.7510 | 1.4687E-11 | 7.1599E-11 |
| ENSG00000096968\|JAK2 | 494.8975 | -0.9855 | ##### | -6.7509 | 1.4694E-11 | 7.1623E-11 |
| ENSG00000126583\|PRKCG | 146.9064 | -1.3541 | ##### | -6.7495 | 1.4835E-11 | 7.2301E-11 |
| ENSG00000073050\|XRCC1 | 3320.4824 | -0.7729 | ##### | -6.7495 | 1.4838E-11 | 7.2304E-11 |
| ENSG00000118513\|MYB | 121.2471 | 1.9655 | ##### | 6.7484 | 1.4944E-11 | 7.2805E-11 |
| ENSG00000110200\|ANAPC15 | 2634.5892 | 0.9448 | ##### | 6.7467 | 1.5129E-11 | 7.3698E-11 |
| ENSG00000174547\|MRPL11 | 3955.5754 | 0.8522 | ##### | 6.7466 | 1.5138E-11 | 7.3728E-11 |
| ENSG00000197798\|FAM118B | 1266.3822 | 0.9328 | ##### | 6.7449 | 1.5313E-11 | 7.4566E-11 |
| ENSG00000130810\|PPAN | 5216.0440 | 0.8523 | ##### | 6.7422 | 1.5601E-11 | 7.5955E-11 |
| ENSG00000095261\|PSMD5 | 6245.1604 | -0.8346 | ##### | -6.7420 | 1.5621E-11 | 7.6042E-11 |
| ENSG00000184178\|SCFD2 | 2584.5954 | 0.8661 | ##### | 6.7399 | 1.5846E-11 | 7.7123E-11 |
| ENSG00000232630\|PRPS1P2 | 95.2460 | 1.9548 | ##### | 6.7396 | 1.5886E-11 | 7.7310E-11 |
| ENSG00000136709\|WDR33 | 10225.4042 | -0.7661 | ##### | -6.7390 | 1.5944E-11 | 7.7578E-11 |
| ENSG00000262919\|CCNQ | 2179.4022 | 0.8608 | ##### | 6.7390 | 1.5949E-11 | 7.7588E-11 |
| ENSG00000151687\|ANKAR | 188.0260 | -1.4987 | ##### | -6.7389 | 1.5959E-11 | 7.7626E-11 |
| ENSG00000289268 | 153.2746 | -1.4026 | ##### | -6.7383 | 1.6024E-11 | 7.7929E-11 |
| ENSG00000205562\|FLRT2-AS1 | 340.9392 | 6.4974 | ##### | 6.7380 | 1.6055E-11 | 7.8065E-11 |
| ENSG00000205659\|LIN52 | 1330.8465 | 0.9173 | ##### | 6.7375 | 1.6113E-11 | 7.8333E-11 |
| ENSG00000186073\|CDIN1 | 877.6506 | -0.9056 | ##### | -6.7374 | 1.6123E-11 | 7.8371E-11 |
| ENSG00000287271 | 575.2493 | -0.9904 | ##### | -6.7366 | 1.6211E-11 | 7.8786E-11 |
| ENSG00000264127\|SCML2P1 | 90.1560 | 3.2690 | ##### | 6.7350 | 1.6396E-11 | 7.9672E-11 |
| ENSG00000198270\|TMEM116 | 531.2758 | -1.1119 | ##### | -6.7333 | 1.6591E-11 | 8.0608E-11 |
| ENSG00000065413\|ANKRD44 | 314.6741 | -1.1433 | ##### | -6.7317 | 1.6771E-11 | 8.1466E-11 |
| ENSG00000244161\|FLNB-AS1 | 76.5930 | -2.0625 | ##### | -6.7313 | 1.6818E-11 | 8.1685E-11 |
| ENSG00000272834 | 40.4311 | 7.5530 | ##### | 6.7309 | 1.6857E-11 | 8.1860E-11 |
| ENSG00000144560\|VGLL4 | 2948.9310 | -0.8592 | ##### | -6.7306 | 1.6900E-11 | 8.2053E-11 |
| ENSG00000103227\|LMF1 | 687.1334 | -0.9089 | ##### | -6.7300 | 1.6971E-11 | 8.2384E-11 |
| ENSG00000293552 | 36.2855 | -5.5727 | ##### | -6.7287 | 1.7121E-11 | 8.3099E-11 |
| ENSG00000158092\|NCK1 | 3057.4584 | -0.8105 | ##### | -6.7278 | 1.7220E-11 | 8.3566E-11 |
| ENSG00000290461 | 318.8219 | ####### | ##### | -6.7277 | 1.7232E-11 | 8.3613E-11 |
| ENSG00000197837\|H4C16 | 122.6286 | 1.7253 | ##### | 6.7274 | 1.7272E-11 | 8.3794E-11 |
| ENSG00000288642\|CDR1 | 75.8077 | 5.4389 | ##### | 6.7266 | 1.7369E-11 | 8.4251E-11 |
| ENSG00000272808\|GCAWKR | 223.3642 | 2.2650 | ##### | 6.7265 | 1.7384E-11 | 8.4307E-11 |
| ENSG00000186575\|NF2 | 5035.8358 | 0.6337 | ##### | 6.7248 | 1.7587E-11 | 8.5280E-11 |
| ENSG00000240859\|LINC03014 | 71.1906 | 2.5057 | ##### | 6.7234 | 1.7759E-11 | 8.6100E-11 |
| ENSG00000279094\|LINC01670 | 50.5407 | -8.0960 | ##### | -6.7233 | 1.7771E-11 | 8.6143E-11 |
| ENSG00000234665\|LERFS | 373.4198 | -1.0979 | ##### | -6.7229 | 1.7816E-11 | 8.6348E-11 |
| ENSG00000266066\|POLRMTP1 | 247.2943 | 1.0698 | ##### | 6.7224 | 1.7880E-11 | 8.6645E-11 |
| ENSG00000187122\|SLIT1 | 77.8004 | 1.9635 | ##### | 6.7223 | 1.7890E-11 | 8.6680E-11 |
| ENSG00000274584 | 48.7250 | -7.8260 | ##### | -6.7222 | 1.7896E-11 | 8.6691E-11 |
| ENSG00000173818\|ENDOV | 1053.1063 | -0.9267 | ##### | -6.7212 | 1.8018E-11 | 8.7256E-11 |
| ENSG00000272347 | 69.4165 | -2.0002 | ##### | -6.7213 | 1.8016E-11 | 8.7256E-11 |
| ENSG00000261324 | 231.9495 | 1.2424 | ##### | 6.7209 | 1.8056E-11 | 8.7425E-11 |
| ENSG00000163737\|PF4 | 147.2142 | -7.0625 | ##### | -6.7204 | 1.8124E-11 | 8.7739E-11 |
| ENSG00000102904\|TSNAXIP1 | 75.2793 | 1.7971 | ##### | 6.7194 | 1.8250E-11 | 8.8337E-11 |
| ENSG00000123066\|MED13L | 8081.8046 | 0.8165 | ##### | 6.7192 | 1.8270E-11 | 8.8421E-11 |
| ENSG00000105699\|LSR | 3714.5023 | -0.8364 | ##### | -6.7184 | 1.8378E-11 | 8.8929E-11 |
| ENSG00000125257\|ABCC4 | 5390.8200 | -0.8199 | ##### | -6.7167 | 1.8585E-11 | 8.9914E-11 |

| | | | | | | |
|---|---|---|---|---|---|---|
| ENSG00000205763\|RP9P | 364.2454 | -1.3274 | ##### | -6.7158 | 1.8709E-11 | 9.0499E-11 |
| ENSG00000006015\|REX1BD | 1325.7442 | -0.8576 | ##### | -6.7156 | 1.8734E-11 | 9.0605E-11 |
| ENSG00000135775\|COG2 | 3229.7847 | -0.8012 | ##### | -6.7149 | 1.8825E-11 | 9.1032E-11 |
| ENSG00000132153\|DHX30 | 12452.3275 | 0.7762 | ##### | 6.7141 | 1.8923E-11 | 9.1492E-11 |
| ENSG00000159871\|LYPD5 | 94.2733 | -3.0617 | ##### | -6.7127 | 1.9110E-11 | 9.2381E-11 |
| ENSG00000173451\|THAP2 | 487.1130 | -1.0529 | ##### | -6.7109 | 1.9346E-11 | 9.3505E-11 |
| ENSG00000196139\|AKR1C3 | 13422.4626 | -9.8013 | ##### | -6.7108 | 1.9352E-11 | 9.3517E-11 |
| ENSG00000254558\|ANKRD33BP8 | 1071.0740 | 11.5391 | ##### | 6.7097 | 1.9509E-11 | 9.4260E-11 |
| ENSG00000110422\|HIPK3 | 5543.5360 | 0.8108 | ##### | 6.7096 | 1.9512E-11 | 9.4260E-11 |
| ENSG00000235454\|HAUS6P3 | 50.7578 | 2.9596 | ##### | 6.7093 | 1.9561E-11 | 9.4485E-11 |
| ENSG00000173868\|PHOSPHO1 | 172.8171 | 1.2953 | ##### | 6.7087 | 1.9639E-11 | 9.4846E-11 |
| ENSG00000144395\|CCDC150 | 616.9530 | 1.0065 | ##### | 6.7062 | 1.9978E-11 | 9.6465E-11 |
| ENSG00000269473\|ZNF132-DT | 81.1158 | 1.7947 | ##### | 6.7059 | 2.0023E-11 | 9.6666E-11 |
| ENSG00000122970\|IFT81 | 3306.9969 | -0.8176 | ##### | -6.7056 | 2.0057E-11 | 9.6817E-11 |
| ENSG00000267534\|S1PR2 | 342.4061 | -1.0253 | ##### | -6.7047 | 2.0175E-11 | 9.7372E-11 |
| ENSG00000169084\|DHRSX | 849.9826 | -0.9713 | ##### | -6.7041 | 2.0269E-11 | 9.7808E-11 |
| ENSG00000260287\|TBC1D3G | 92.4916 | 1.5600 | ##### | 6.7027 | 2.0461E-11 | 9.8716E-11 |
| ENSG00000104635\|SLC39A14 | 19839.0562 | 0.8128 | ##### | 6.7022 | 2.0533E-11 | 9.9050E-11 |
| ENSG00000130511\|SSBP4 | 4365.2672 | 0.8024 | ##### | 6.7020 | 2.0558E-11 | 9.9155E-11 |
| ENSG00000266053\|NDUFV2-AS1 | 74.1029 | 2.0415 | ##### | 6.7005 | 2.0767E-11 | 1.0014E-10 |
| ENSG00000103034\|NDRG4 | 158.8187 | -1.2766 | ##### | -6.6986 | 2.1047E-11 | 1.0148E-10 |
| ENSG00000167202\|TBC1D2B | 2947.1952 | 0.8167 | ##### | 6.6951 | 2.1555E-11 | 1.0391E-10 |
| ENSG00000259291\|ZNF710-AS1 | 125.7883 | -1.6145 | ##### | -6.6946 | 2.1621E-11 | 1.0422E-10 |
| ENSG00000090006\|LTBP4 | 9074.8759 | -0.8002 | ##### | -6.6941 | 2.1699E-11 | 1.0457E-10 |
| ENSG00000272282\|LINC02084 | 57.4941 | 2.2917 | ##### | 6.6938 | 2.1742E-11 | 1.0476E-10 |
| ENSG00000254245\|PCDHGA3 | 13263.3011 | ####### | ##### | -6.6920 | 2.2008E-11 | 1.0603E-10 |
| ENSG00000019485\|PRDM11 | 891.0761 | 0.9717 | ##### | 6.6916 | 2.2075E-11 | 1.0633E-10 |
| ENSG00000005882\|PDK2 | 1986.3129 | 0.8942 | ##### | 6.6914 | 2.2112E-11 | 1.0649E-10 |
| ENSG00000154040\|CABYR | 1705.5943 | -0.8176 | ##### | -6.6912 | 2.2129E-11 | 1.0656E-10 |
| ENSG00000272168\|CASC15 | 109.2500 | 2.0757 | ##### | 6.6911 | 2.2151E-11 | 1.0665E-10 |
| ENSG00000163435\|ELF3 | 4218.0132 | ####### | ##### | -6.6902 | 2.2285E-11 | 1.0727E-10 |
| ENSG00000101246\|ARFRP1 | 3493.0352 | 0.8093 | ##### | 6.6892 | 2.2446E-11 | 1.0804E-10 |
| ENSG00000100226\|GTPBP1 | 2963.3844 | 0.8026 | ##### | 6.6882 | 2.2587E-11 | 1.0870E-10 |
| ENSG00000065000\|AP3D1 | 24691.2756 | 0.7668 | ##### | 6.6864 | 2.2870E-11 | 1.1004E-10 |
| ENSG00000059122\|FLYWCH1 | 5571.3255 | -0.7959 | ##### | -6.6846 | 2.3161E-11 | 1.1142E-10 |
| ENSG00000088808\|PPP1R13B | 1305.2861 | -0.8700 | ##### | -6.6842 | 2.3220E-11 | 1.1169E-10 |
| ENSG00000279672 | 175.7964 | 1.3045 | ##### | 6.6836 | 2.3317E-11 | 1.1214E-10 |
| ENSG00000109618\|SEPSECS | 702.1873 | -0.9449 | ##### | -6.6834 | 2.3348E-11 | 1.1227E-10 |
| ENSG00000170681\|CAVIN4 | 219.6465 | 1.2637 | ##### | 6.6825 | 2.3495E-11 | 1.1295E-10 |
| ENSG00000229127\|KANSL1L-AS1 | 25.4197 | 3.6680 | ##### | 6.6816 | 2.3641E-11 | 1.1364E-10 |
| ENSG00000205572\|SERF1B | 613.9115 | -1.0777 | ##### | -6.6804 | 2.3824E-11 | 1.1450E-10 |
| ENSG00000162746\|FCRLB | 122.2330 | -1.4234 | ##### | -6.6784 | 2.4164E-11 | 1.1611E-10 |
| ENSG00000149136\|SSRP1 | 23174.5643 | 0.5859 | ##### | 6.6777 | 2.4273E-11 | 1.1662E-10 |
| ENSG00000125398\|SOX9 | 1060.6613 | -0.8953 | ##### | -6.6768 | 2.4428E-11 | 1.1735E-10 |
| ENSG00000290537 | 172.8665 | 1.5826 | ##### | 6.6766 | 2.4462E-11 | 1.1749E-10 |
| ENSG00000135926\|TMBIM1 | 11720.6517 | -0.7196 | ##### | -6.6762 | 2.4516E-11 | 1.1773E-10 |
| ENSG00000259838\|ELOCP2 | 2586.4413 | ####### | ##### | -6.6733 | 2.5019E-11 | 1.2013E-10 |
| ENSG00000268129 | 57.0889 | -1.9892 | ##### | -6.6730 | 2.5071E-11 | 1.2036E-10 |
| ENSG00000134202\|GSTM3 | 2613.6771 | -0.8593 | ##### | -6.6725 | 2.5153E-11 | 1.2073E-10 |
| ENSG00000130038\|CRACR2A | 29.7097 | 3.2330 | ##### | 6.6724 | 2.5167E-11 | 1.2078E-10 |
| ENSG00000151693\|ASAP2 | 5637.1102 | -0.8112 | ##### | -6.6720 | 2.5228E-11 | 1.2105E-10 |
| ENSG00000293434\|AHSA2P | 3496.6036 | 0.8732 | ##### | 6.6704 | 2.5508E-11 | 1.2238E-10 |
| ENSG00000127129\|EDN2 | 543.1541 | -5.7884 | ##### | -6.6703 | 2.5534E-11 | 1.2248E-10 |
| ENSG00000213901\|SLC23A3 | 141.9773 | 1.4677 | ##### | 6.6691 | 2.5735E-11 | 1.2343E-10 |
| ENSG00000171606\|ZNF274 | 2953.7577 | -0.8459 | ##### | -6.6688 | 2.5790E-11 | 1.2367E-10 |
| ENSG00000177733\|HNRNPA0 | 23369.1512 | 0.7852 | ##### | 6.6680 | 2.5923E-11 | 1.2429E-10 |
| ENSG00000240990\|HOXA11-AS | 187.0417 | -1.2527 | ##### | -6.6667 | 2.6162E-11 | 1.2541E-10 |
| ENSG00000040633\|PHF23 | 5228.6293 | 0.8317 | ##### | 6.6658 | 2.6330E-11 | 1.2620E-10 |
| ENSG00000196418\|ZNF124 | 366.5702 | -1.1312 | ##### | -6.6613 | 2.7136E-11 | 1.3004E-10 |
| ENSG00000274423\|SEC22B2P | 72.5985 | 1.8458 | ##### | 6.6601 | 2.7364E-11 | 1.3111E-10 |
| ENSG00000138002\|IFT172 | 1515.6067 | 0.8857 | ##### | 6.6601 | 2.7369E-11 | 1.3112E-10 |
| ENSG00000078967\|UBE2D4 | 1821.4009 | -0.8804 | ##### | -6.6597 | 2.7448E-11 | 1.3147E-10 |
| ENSG00000132471\|WBP2 | 6555.3135 | -0.8214 | ##### | -6.6592 | 2.7537E-11 | 1.3188E-10 |

| | | | | | | |
|---|---|---|---|---|---|---|
| ENSG00000196295\|GARS1-DT | 865.6139 | -0.9611 | ##### | -6.6589 | 2.7587E-11 | 1.3210E-10 |
| ENSG00000283375 | 96.3284 | -1.5962 | ##### | -6.6574 | 2.7880E-11 | 1.3348E-10 |
| ENSG00000162174\|ASRGL1 | 1981.8486 | 0.8472 | ##### | 6.6567 | 2.8005E-11 | 1.3405E-10 |
| ENSG00000289303 | 49.3297 | 7.2023 | ##### | 6.6560 | 2.8136E-11 | 1.3466E-10 |
| ENSG00000105085\|MED26 | 1095.1956 | 0.8841 | ##### | 6.6550 | 2.8327E-11 | 1.3555E-10 |
| ENSG00000259802\|MARCHF6-DT | 211.6125 | -1.1054 | ##### | -6.6526 | 2.8791E-11 | 1.3775E-10 |
| ENSG00000145725\|PPIP5K2 | 4768.7898 | 0.7265 | ##### | 6.6526 | 2.8795E-11 | 1.3775E-10 |
| ENSG00000172830\|SSH3 | 2245.2702 | 0.8547 | ##### | 6.6509 | 2.9128E-11 | 1.3932E-10 |
| ENSG00000169203\|NPIPB12 | 750.0454 | 0.9976 | ##### | 6.6501 | 2.9296E-11 | 1.4010E-10 |
| ENSG00000250479\|CHCHD10 | 4084.9951 | -0.8097 | ##### | -6.6495 | 2.9417E-11 | 1.4066E-10 |
| ENSG00000080546\|SESN1 | 734.7945 | 0.9778 | ##### | 6.6482 | 2.9678E-11 | 1.4188E-10 |
| ENSG00000078403\|MLLT10 | 3744.2021 | 0.7981 | ##### | 6.6478 | 2.9748E-11 | 1.4219E-10 |
| ENSG00000213963 | 811.2426 | -0.9234 | ##### | -6.6473 | 2.9842E-11 | 1.4262E-10 |
| ENSG00000184979\|USP18 | 449.6889 | -1.0322 | ##### | -6.6468 | 2.9959E-11 | 1.4316E-10 |
| ENSG00000155363\|MOV10 | 4615.3801 | -0.8067 | ##### | -6.6457 | 3.0169E-11 | 1.4414E-10 |
| ENSG00000145604\|SKP2 | 5057.2730 | 0.8071 | ##### | 6.6435 | 3.0627E-11 | 1.4630E-10 |
| ENSG00000177888\|ZBTB41 | 2653.2569 | -0.8365 | ##### | -6.6427 | 3.0809E-11 | 1.4715E-10 |
| ENSG00000160685\|ZBTB7B | 10720.5903 | 0.8034 | ##### | 6.6421 | 3.0928E-11 | 1.4769E-10 |
| ENSG00000241852\|C8orf58 | 1100.7121 | 0.9134 | ##### | 6.6420 | 3.0942E-11 | 1.4773E-10 |
| ENSG00000171700\|RGS19 | 2828.0784 | 0.8469 | ##### | 6.6410 | 3.1166E-11 | 1.4878E-10 |
| ENSG00000165264\|NDUFB6 | 1782.8335 | -0.8874 | ##### | -6.6408 | 3.1199E-11 | 1.4891E-10 |
| ENSG00000011105\|TSPAN9 | 3584.5229 | 0.6552 | ##### | 6.6408 | 3.1209E-11 | 1.4894E-10 |
| ENSG00000222041\|CYTOR | 3485.0176 | 0.8306 | ##### | 6.6389 | 3.1611E-11 | 1.5083E-10 |
| ENSG00000236761\|CTAGE9 | 30.2613 | -2.5291 | ##### | -6.6383 | 3.1727E-11 | 1.5136E-10 |
| ENSG00000100804\|PSMB5 | 15725.8737 | 0.8179 | ##### | 6.6367 | 3.2082E-11 | 1.5303E-10 |
| ENSG00000237489\|C10orf143 | 65.9119 | -2.6030 | ##### | -6.6343 | 3.2603E-11 | 1.5549E-10 |
| ENSG00000166415\|WDR72 | 1620.2209 | -8.7908 | ##### | -6.6341 | 3.2659E-11 | 1.5573E-10 |
| ENSG00000213859\|KCTD11 | 2052.5703 | 0.8516 | ##### | 6.6338 | 3.2707E-11 | 1.5592E-10 |
| ENSG00000179632\|MAF1 | 10616.6489 | 0.6855 | ##### | 6.6338 | 3.2708E-11 | 1.5592E-10 |
| ENSG00000184575\|XPOT | 26021.5386 | 0.7900 | ##### | 6.6338 | 3.2725E-11 | 1.5595E-10 |
| ENSG00000175061\|SNHG29 | 14250.7005 | -0.7853 | ##### | -6.6338 | 3.2724E-11 | 1.5595E-10 |
| ENSG00000244398\|RPL36AP37 | 3334.9469 | 0.8445 | ##### | 6.6337 | 3.2734E-11 | 1.5597E-10 |
| ENSG00000175387\|SMAD2 | 11575.8639 | -0.7415 | ##### | -6.6328 | 3.2935E-11 | 1.5690E-10 |
| ENSG00000177465\|ACOT4 | 144.5795 | -1.5927 | ##### | -6.6326 | 3.2988E-11 | 1.5713E-10 |
| ENSG00000178934\|LGALS7B | 49.7378 | 7.4266 | ##### | 6.6324 | 3.3034E-11 | 1.5732E-10 |
| ENSG00000135164\|DMTF1 | 3047.6915 | 0.8384 | ##### | 6.6320 | 3.3121E-11 | 1.5771E-10 |
| ENSG00000139178\|C1RL | 2291.2542 | -0.8554 | ##### | -6.6316 | 3.3209E-11 | 1.5810E-10 |
| ENSG00000154930\|ACSS1 | 408.8898 | -1.0576 | ##### | -6.6314 | 3.3256E-11 | 1.5830E-10 |
| ENSG00000139579\|NABP2 | 6009.8314 | 0.8148 | ##### | 6.6303 | 3.3496E-11 | 1.5942E-10 |
| ENSG00000203772\|SPRN | 309.9404 | 1.0968 | ##### | 6.6302 | 3.3532E-11 | 1.5956E-10 |
| ENSG00000125450\|NUP85 | 6051.6805 | 0.7959 | ##### | 6.6301 | 3.3543E-11 | 1.5959E-10 |
| ENSG00000251022\|THAP9-AS1 | 2440.4482 | 0.8543 | ##### | 6.6292 | 3.3757E-11 | 1.6059E-10 |
| ENSG00000150672\|DLG2 | 104.2643 | -1.4568 | ##### | -6.6287 | 3.3856E-11 | 1.6103E-10 |
| ENSG00000126091\|ST3GAL3 | 1258.5504 | -0.9044 | ##### | -6.6268 | 3.4309E-11 | 1.6316E-10 |
| ENSG00000186792\|HYAL3 | 1149.5128 | 0.8761 | ##### | 6.6237 | 3.5027E-11 | 1.6654E-10 |
| ENSG00000136237\|RAPGEF5 | 222.1808 | 5.4854 | ##### | 6.6209 | 3.5711E-11 | 1.6977E-10 |
| ENSG00000187325\|TAF9B | 2181.1708 | 0.8432 | ##### | 6.6188 | 3.6210E-11 | 1.7212E-10 |
| ENSG00000139163\|ETNK1 | 8306.0850 | 0.7596 | ##### | 6.6178 | 3.6457E-11 | 1.7326E-10 |
| ENSG00000111879\|FAM184A | 128.2798 | 2.1444 | ##### | 6.6170 | 3.6665E-11 | 1.7422E-10 |
| ENSG00000121579\|NAA50 | 20824.8712 | 0.5309 | ##### | 6.6168 | 3.6697E-11 | 1.7435E-10 |
| ENSG00000165487\|MICU2 | 3137.2881 | 0.6568 | ##### | 6.6148 | 3.7213E-11 | 1.7677E-10 |
| ENSG00000144791\|LIMD1 | 6400.0163 | -0.8107 | ##### | -6.6146 | 3.7251E-11 | 1.7692E-10 |
| ENSG00000142684\|ZNF593 | 5043.7262 | 0.8869 | ##### | 6.6141 | 3.7389E-11 | 1.7755E-10 |
| ENSG00000115568\|ZNF142 | 5048.2497 | 0.8016 | ##### | 6.6137 | 3.7485E-11 | 1.7798E-10 |
| ENSG00000174227\|PIGG | 3932.9169 | -0.8048 | ##### | -6.6129 | 3.7694E-11 | 1.7894E-10 |
| ENSG00000185033\|SEMA4B | 8704.7963 | 0.5994 | ##### | 6.6111 | 3.8151E-11 | 1.8109E-10 |
| ENSG00000293480\|TCAM1P | 50.6666 | 2.3481 | ##### | 6.6109 | 3.8198E-11 | 1.8128E-10 |
| ENSG00000100060\|MFNG | 66.4460 | -1.9367 | ##### | -6.6100 | 3.8430E-11 | 1.8235E-10 |
| ENSG00000240771\|ARHGEF25 | 1966.1009 | 0.8770 | ##### | 6.6094 | 3.8576E-11 | 1.8302E-10 |
| ENSG00000125462\|MIR9-1HG | 426.6326 | 1.1753 | ##### | 6.6085 | 3.8821E-11 | 1.8415E-10 |
| ENSG00000248774 | 56.3264 | -1.6032 | ##### | -6.6059 | 3.9512E-11 | 1.8740E-10 |
| ENSG00000062370\|ZNF112 | 342.1807 | -1.0209 | ##### | -6.6047 | 3.9825E-11 | 1.8885E-10 |
| ENSG00000180767\|CHST13 | 631.1661 | -0.9282 | ##### | -6.6014 | 4.0720E-11 | 1.9306E-10 |

| | | | | | | |
|---|---|---|---|---|---|---|
| ENSG00000089685\|BIRC5 | 11267.1495 | 0.8378 | ##### | 6.5988 | 4.1443E-11 | 1.9646E-10 |
| ENSG00000153827\|TRIP12 | 30186.4155 | 0.7761 | ##### | 6.5974 | 4.1844E-11 | 1.9833E-10 |
| ENSG00000180596\|H2BC4 | 917.2285 | 0.9252 | ##### | 6.5968 | 4.2019E-11 | 1.9913E-10 |
| ENSG00000288924 | 181.4072 | -1.7383 | ##### | -6.5950 | 4.2521E-11 | 2.0148E-10 |
| ENSG00000167565\|SERTAD3 | 2606.2971 | -0.8293 | ##### | -6.5925 | 4.3234E-11 | 2.0482E-10 |
| ENSG00000218617 | 22.5423 | -3.5349 | ##### | -6.5923 | 4.3297E-11 | 2.0508E-10 |
| ENSG00000247240\|UBL7-DT | 344.9054 | 1.0467 | ##### | 6.5894 | 4.4147E-11 | 2.0908E-10 |
| ENSG00000105401\|CDC37 | 21803.2129 | 0.7688 | ##### | 6.5886 | 4.4403E-11 | 2.1026E-10 |
| ENSG00000261135 | 51.2101 | -2.1912 | ##### | -6.5883 | 4.4485E-11 | 2.1061E-10 |
| ENSG00000178814\|OPLAH | 2009.0558 | 0.8234 | ##### | 6.5882 | 4.4509E-11 | 2.1069E-10 |
| ENSG00000227704 | 139.0244 | -1.3877 | ##### | -6.5882 | 4.4525E-11 | 2.1074E-10 |
| ENSG00000272610\|MAGI1-IT1 | 26.2218 | 7.4975 | ##### | 6.5874 | 4.4757E-11 | 2.1180E-10 |
| ENSG00000224165\|DNAJC27-AS1 | 52.7249 | -2.3760 | ##### | -6.5863 | 4.5100E-11 | 2.1336E-10 |
| ENSG00000196132\|MYT1 | 864.5388 | 11.1516 | ##### | 6.5863 | 4.5096E-11 | 2.1336E-10 |
| ENSG00000140688\|RUSF1 | 7555.1434 | -0.9012 | ##### | -6.5860 | 4.5174E-11 | 2.1367E-10 |
| ENSG00000144824\|PHLDB2 | 15093.7507 | 0.7954 | ##### | 6.5855 | 4.5337E-11 | 2.1441E-10 |
| ENSG00000197808\|ZNF461 | 460.8408 | -0.9909 | ##### | -6.5852 | 4.5435E-11 | 2.1484E-10 |
| ENSG00000259863\|SH3RF3-AS1 | 37.9578 | 3.1200 | ##### | 6.5850 | 4.5476E-11 | 2.1500E-10 |
| ENSG00000116198\|CEP104 | 3847.0627 | -0.8070 | ##### | -6.5823 | 4.6316E-11 | 2.1893E-10 |
| ENSG00000183605\|SFXN4 | 1905.9193 | -0.8156 | ##### | -6.5818 | 4.6492E-11 | 2.1973E-10 |
| ENSG00000109536\|FRG1 | 821.2719 | -0.9064 | ##### | -6.5813 | 4.6650E-11 | 2.2044E-10 |
| ENSG00000104611\|SH2D4A | 2329.2782 | -0.7927 | ##### | -6.5795 | 4.7194E-11 | 2.2298E-10 |
| ENSG00000118160\|SLC8A2 | 112.5238 | -1.6957 | ##### | -6.5787 | 4.7447E-11 | 2.2413E-10 |
| ENSG00000107036\|RIC1 | 1709.6576 | -0.8515 | ##### | -6.5787 | 4.7452E-11 | 2.2413E-10 |
| ENSG00000185246\|PRPF39 | 2210.5405 | 0.8344 | ##### | 6.5781 | 4.7660E-11 | 2.2507E-10 |
| ENSG00000133739\|LRRCC1 | 681.0444 | -0.9043 | ##### | -6.5769 | 4.8036E-11 | 2.2681E-10 |
| ENSG00000147439\|BIN3 | 1408.7893 | 0.8194 | ##### | 6.5767 | 4.8113E-11 | 2.2714E-10 |
| ENSG00000184205\|TSPYL2 | 2198.6741 | 0.8478 | ##### | 6.5760 | 4.8316E-11 | 2.2807E-10 |
| ENSG00000185009\|AP3M1 | 6047.1632 | -0.5174 | ##### | -6.5760 | 4.8338E-11 | 2.2813E-10 |
| ENSG00000113594\|LIFR | 6058.1662 | 0.7981 | ##### | 6.5750 | 4.8642E-11 | 2.2953E-10 |
| ENSG00000129128\|SPCS3 | 7326.4061 | -0.7921 | ##### | -6.5739 | 4.9023E-11 | 2.3129E-10 |
| ENSG00000115762\|PLEKHB2 | 16965.4792 | -0.6056 | ##### | -6.5738 | 4.9036E-11 | 2.3132E-10 |
| ENSG00000230778\|ANKRD63 | 136.5374 | 2.1182 | ##### | 6.5725 | 4.9487E-11 | 2.3340E-10 |
| ENSG00000099875\|MKNK2 | 10836.6343 | 0.7316 | ##### | 6.5716 | 4.9775E-11 | 2.3473E-10 |
| ENSG00000289236 | 35.3874 | 7.3715 | ##### | 6.5711 | 4.9946E-11 | 2.3550E-10 |
| ENSG00000174442\|ZWILCH | 5323.0541 | 0.8149 | ##### | 6.5708 | 5.0030E-11 | 2.3585E-10 |
| ENSG00000287934 | 168.4823 | 3.0431 | ##### | 6.5706 | 5.0114E-11 | 2.3621E-10 |
| ENSG00000214575\|CPEB1 | 941.9613 | 1.0217 | ##### | 6.5702 | 5.0263E-11 | 2.3688E-10 |
| ENSG00000280128 | 532.1331 | 0.9608 | ##### | 6.5687 | 5.0764E-11 | 2.3920E-10 |
| ENSG00000179344\|HLA-DQB1 | 73.6435 | 1.7737 | ##### | 6.5682 | 5.0933E-11 | 2.3996E-10 |
| ENSG00000155097\|ATP6V1C1 | 8220.3304 | -0.7738 | ##### | -6.5671 | 5.1289E-11 | 2.4160E-10 |
| ENSG00000279092 | 59.6596 | 5.1226 | ##### | 6.5669 | 5.1384E-11 | 2.4201E-10 |
| ENSG00000168944\|CEP120 | 2163.8069 | 0.8027 | ##### | 6.5667 | 5.1444E-11 | 2.4221E-10 |
| ENSG00000253948\|VPS13B-DT | 41.1483 | -2.9338 | ##### | -6.5667 | 5.1440E-11 | 2.4221E-10 |
| ENSG00000128923\|MINDY2 | 1934.7990 | -0.8492 | ##### | -6.5664 | 5.1558E-11 | 2.4271E-10 |
| ENSG00000159884\|CCDC107 | 2635.6930 | 0.8723 | ##### | 6.5661 | 5.1665E-11 | 2.4318E-10 |
| ENSG00000281856 | 34.3652 | -2.2496 | ##### | -6.5659 | 5.1725E-11 | 2.4342E-10 |
| ENSG00000123989\|CHPF | 5061.8965 | -0.7748 | ##### | -6.5657 | 5.1780E-11 | 2.4364E-10 |
| ENSG00000215021\|PHB2 | 30283.2601 | 0.7784 | ##### | 6.5647 | 5.2150E-11 | 2.4535E-10 |
| ENSG00000270629\|NBPF14 | 2242.1798 | 0.8565 | ##### | 6.5642 | 5.2310E-11 | 2.4606E-10 |
| ENSG00000115423\|DNAH6 | 44.8498 | -2.2856 | ##### | -6.5636 | 5.2530E-11 | 2.4706E-10 |
| ENSG00000152518\|ZFP36L2 | 7998.1741 | 0.7403 | ##### | 6.5635 | 5.2554E-11 | 2.4713E-10 |
| ENSG00000203880\|PCMTD2 | 3306.0913 | 0.8224 | ##### | 6.5629 | 5.2769E-11 | 2.4810E-10 |
| ENSG00000037757\|MRI1 | 2623.0170 | 0.8396 | ##### | 6.5613 | 5.3324E-11 | 2.5067E-10 |
| ENSG00000132256\|TRIM5 | 1637.1977 | -0.8777 | ##### | -6.5609 | 5.3494E-11 | 2.5143E-10 |
| ENSG00000100632\|ERH | 20146.0670 | 0.7660 | ##### | 6.5607 | 5.3565E-11 | 2.5173E-10 |
| ENSG00000174437\|ATP2A2 | 61256.2165 | -0.6602 | ##### | -6.5603 | 5.3694E-11 | 2.5229E-10 |
| ENSG00000141127\|PRPSAP2 | 3041.0565 | 0.8070 | ##### | 6.5602 | 5.3722E-11 | 2.5239E-10 |
| ENSG00000197822\|OCLN | 1855.9266 | 0.8210 | ##### | 6.5599 | 5.3844E-11 | 2.5292E-10 |
| ENSG00000157693\|TMEM268 | 1365.2669 | -0.8513 | ##### | -6.5593 | 5.4076E-11 | 2.5397E-10 |
| ENSG00000080823\|MOK | 1728.5656 | -0.8249 | ##### | -6.5581 | 5.4499E-11 | 2.5591E-10 |
| ENSG00000278615\|C11orf98 | 3162.7264 | 0.9344 | ##### | 6.5572 | 5.4833E-11 | 2.5744E-10 |
| ENSG00000101204\|CHRNA4 | 46.0163 | 2.3982 | ##### | 6.5561 | 5.5235E-11 | 2.5929E-10 |

| | | | | | | |
|---|---|---|---|---|---|---|
| ENSG00000186395\|KRT10 | 3104.4299 | -0.8251 | ##### | -6.5548 | 5.5726E-11 | 2.6155E-10 |
| ENSG00000144589\|STK11IP | 3152.8475 | 0.7862 | ##### | 6.5545 | 5.5827E-11 | 2.6199E-10 |
| ENSG00000113966\|ARL6 | 1094.8163 | 0.9218 | ##### | 6.5529 | 5.6449E-11 | 2.6486E-10 |
| ENSG00000055917\|PUM2 | 11657.1093 | 0.7796 | ##### | 6.5517 | 5.6891E-11 | 2.6689E-10 |
| ENSG00000137502\|RAB30 | 856.0024 | -0.8985 | ##### | -6.5512 | 5.7093E-11 | 2.6780E-10 |
| ENSG00000172428\|COPS9 | 3181.9021 | 0.8089 | ##### | 6.5509 | 5.7191E-11 | 2.6822E-10 |
| ENSG00000290069 | 28.4835 | 2.6676 | ##### | 6.5494 | 5.7756E-11 | 2.7082E-10 |
| ENSG00000046653\|GPM6B | 40.6288 | 2.0701 | ##### | 6.5492 | 5.7842E-11 | 2.7119E-10 |
| ENSG00000115419\|GLS | 22873.9945 | -0.7721 | ##### | -6.5491 | 5.7903E-11 | 2.7143E-10 |
| ENSG00000163075\|CFAP221 | 332.5273 | -8.6019 | ##### | -6.5486 | 5.8081E-11 | 2.7222E-10 |
| ENSG00000205710\|C17orf107 | 156.6269 | 1.1830 | ##### | 6.5478 | 5.8407E-11 | 2.7371E-10 |
| ENSG00000173678\|SPDYE2B | 81.7058 | 1.5161 | ##### | 6.5476 | 5.8452E-11 | 2.7387E-10 |
| ENSG00000137809\|ITGA11 | 41.7792 | -3.2294 | ##### | -6.5451 | 5.9457E-11 | 2.7854E-10 |
| ENSG00000179630\|LACC1 | 586.1419 | -0.9643 | ##### | -6.5450 | 5.9506E-11 | 2.7872E-10 |
| ENSG00000122335\|SERAC1 | 703.8764 | -0.9538 | ##### | -6.5434 | 6.0141E-11 | 2.8166E-10 |
| ENSG00000197665 | 37.5533 | -2.2256 | ##### | -6.5414 | 6.0964E-11 | 2.8546E-10 |
| ENSG00000270540 | 51.1257 | 8.3176 | ##### | 6.5413 | 6.0986E-11 | 2.8552E-10 |
| ENSG00000171724\|VAT1L | 150.1042 | 5.8312 | ##### | 6.5408 | 6.1170E-11 | 2.8634E-10 |
| ENSG00000100983\|GSS | 10048.8454 | 0.7998 | ##### | 6.5392 | 6.1839E-11 | 2.8943E-10 |
| ENSG00000196458\|ZNF605 | 1573.9673 | -0.8416 | ##### | -6.5381 | 6.2291E-11 | 2.9150E-10 |
| ENSG00000234432\|LINC02983 | 55.1333 | 2.5446 | ##### | 6.5378 | 6.2447E-11 | 2.9218E-10 |
| ENSG00000250902\|SMAD1-AS1 | 36.7981 | -2.3595 | ##### | -6.5372 | 6.2702E-11 | 2.9333E-10 |
| ENSG00000166452\|AKIP1 | 1478.0785 | -0.8641 | ##### | -6.5349 | 6.3668E-11 | 2.9780E-10 |
| ENSG00000168811\|IL12A | 103.2440 | 1.8238 | ##### | 6.5340 | 6.4043E-11 | 2.9951E-10 |
| ENSG00000128965\|CHAC1 | 361.6759 | -1.1726 | ##### | -6.5339 | 6.4090E-11 | 2.9968E-10 |
| ENSG00000142541\|RPL13A | 77693.0964 | -0.7777 | ##### | -6.5338 | 6.4130E-11 | 2.9982E-10 |
| ENSG00000131188\|PRR7 | 1570.3988 | 0.8378 | ##### | 6.5332 | 6.4399E-11 | 3.0103E-10 |
| ENSG00000148835\|TAF5 | 1121.7351 | 0.8492 | ##### | 6.5325 | 6.4677E-11 | 3.0228E-10 |
| ENSG00000160131\|VMA21 | 6943.4639 | 0.8040 | ##### | 6.5319 | 6.4921E-11 | 3.0337E-10 |
| ENSG00000164713\|BRI3 | 4579.1509 | -0.7956 | ##### | -6.5314 | 6.5159E-11 | 3.0444E-10 |
| ENSG00000181873\|IBA57 | 648.5791 | 0.8779 | ##### | 6.5299 | 6.5819E-11 | 3.0747E-10 |
| ENSG00000187550\|SBK2 | 52.7101 | -3.8277 | ##### | -6.5285 | 6.6442E-11 | 3.1033E-10 |
| ENSG00000204314\|PRRT1 | 302.7723 | 1.0938 | ##### | 6.5284 | 6.6486E-11 | 3.1049E-10 |
| ENSG00000120669\|SOHLH2 | 1492.2801 | -9.8283 | ##### | -6.5278 | 6.6756E-11 | 3.1168E-10 |
| ENSG00000105011\|ASF1B | 3236.0167 | 0.8081 | ##### | 6.5278 | 6.6762E-11 | 3.1168E-10 |
| ENSG00000267750\|RUNDC3A-AS1 | 217.8979 | -1.1314 | ##### | -6.5276 | 6.6842E-11 | 3.1201E-10 |
| ENSG00000109323\|MANBA | 883.2627 | -0.8605 | ##### | -6.5257 | 6.7694E-11 | 3.1594E-10 |
| ENSG00000100346\|CACNA1I | 26.7853 | -3.2569 | ##### | -6.5236 | 6.8651E-11 | 3.2035E-10 |
| ENSG00000130052\|STARD8 | 522.6005 | 0.9455 | ##### | 6.5233 | 6.8756E-11 | 3.2079E-10 |
| ENSG00000233527\|ZNF529-AS1 | 227.2164 | -1.4768 | ##### | -6.5227 | 6.9071E-11 | 3.2221E-10 |
| ENSG00000250874 | 57.9880 | -1.9977 | ##### | -6.5224 | 6.9171E-11 | 3.2263E-10 |
| ENSG00000135373\|EHF | 233.6936 | -4.2525 | ##### | -6.5215 | 6.9616E-11 | 3.2465E-10 |
| ENSG00000167840\|ZNF232 | 427.1074 | 1.0120 | ##### | 6.5215 | 6.9628E-11 | 3.2466E-10 |
| ENSG00000270147 | 43.5517 | 7.1037 | ##### | 6.5201 | 7.0238E-11 | 3.2745E-10 |
| ENSG00000275393 | 35.5142 | -7.5841 | ##### | -6.5185 | 7.1001E-11 | 3.3096E-10 |
| ENSG00000290921 | 173.1945 | 1.3925 | ##### | 6.5173 | 7.1606E-11 | 3.3373E-10 |
| ENSG00000185480\|PARPBP | 2975.3682 | 0.8498 | ##### | 6.5170 | 7.1748E-11 | 3.3433E-10 |
| ENSG00000256433 | 557.9999 | ####### | ##### | -6.5167 | 7.1866E-11 | 3.3483E-10 |
| ENSG00000175414\|ARL10 | 926.2372 | 0.9475 | ##### | 6.5163 | 7.2069E-11 | 3.3573E-10 |
| ENSG00000161677\|JOSD2 | 2254.4086 | -0.8185 | ##### | -6.5156 | 7.2384E-11 | 3.3714E-10 |
| ENSG00000272114 | 18.3742 | -2.8808 | ##### | -6.5152 | 7.2608E-11 | 3.3813E-10 |
| ENSG00000127511\|SIN3B | 5276.7821 | 0.7721 | ##### | 6.5139 | 7.3201E-11 | 3.4084E-10 |
| ENSG00000135424\|ITGA7 | 492.5210 | -0.9593 | ##### | -6.5128 | 7.3762E-11 | 3.4340E-10 |
| ENSG00000013306\|SLC25A39 | 16746.3237 | 0.7865 | ##### | 6.5124 | 7.3983E-11 | 3.4438E-10 |
| ENSG00000198816\|ZNF358 | 2531.2476 | -0.7240 | ##### | -6.5118 | 7.4260E-11 | 3.4561E-10 |
| ENSG00000276846 | 16.7264 | -5.0200 | ##### | -6.5113 | 7.4502E-11 | 3.4668E-10 |
| ENSG00000277287 | 43.2841 | -2.1663 | ##### | -6.5106 | 7.4865E-11 | 3.4832E-10 |
| ENSG00000205746\|PKD1P4 | 2535.3428 | 0.8824 | ##### | 6.5098 | 7.5251E-11 | 3.5006E-10 |
| ENSG00000171303\|KCNK3 | 172.2067 | -5.3532 | ##### | -6.5097 | 7.5302E-11 | 3.5024E-10 |
| ENSG00000289858 | 260.3161 | -1.3032 | ##### | -6.5087 | 7.5792E-11 | 3.5247E-10 |
| ENSG00000186020\|ZNF529 | 1152.9082 | -0.8465 | ##### | -6.5083 | 7.6014E-11 | 3.5343E-10 |
| ENSG00000054611\|TBC1D22A | 1750.3935 | -0.8333 | ##### | -6.5083 | 7.6023E-11 | 3.5343E-10 |
| ENSG00000286797 | 69.0165 | -4.5141 | ##### | -6.5081 | 7.6116E-11 | 3.5381E-10 |

| | | | | | | |
|---|---|---|---|---|---|---|
| ENSG00000164405\|UQCRQ | 8962.7693 | 0.8042 | ##### | 6.5074 | 7.6481E-11 | 3.5545E-10 |
| ENSG00000197265\|GTF2E2 | 4152.8170 | -0.8142 | ##### | -6.5069 | 7.6692E-11 | 3.5637E-10 |
| ENSG00000100592\|DAAM1 | 2011.1603 | 0.8630 | ##### | 6.5055 | 7.7421E-11 | 3.5971E-10 |
| ENSG00000107984\|DKK1 | 15687.5691 | -0.7854 | ##### | -6.5052 | 7.7577E-11 | 3.6037E-10 |
| ENSG00000185986\|SDHAP3 | 24.8228 | 2.7466 | ##### | 6.5023 | 7.9096E-11 | 3.6732E-10 |
| ENSG00000146063\|TRIM41 | 4395.7041 | 0.8159 | ##### | 6.5023 | 7.9088E-11 | 3.6732E-10 |
| ENSG00000101220\|ADISSP | 4621.3674 | 0.8191 | ##### | 6.5014 | 7.9584E-11 | 3.6952E-10 |
| ENSG00000130224\|LRCH2 | 534.3447 | 5.4995 | ##### | 6.5014 | 7.9597E-11 | 3.6953E-10 |
| ENSG00000224383\|PRR29 | 127.7652 | -1.8437 | ##### | -6.5012 | 7.9674E-11 | 3.6983E-10 |
| ENSG00000178074\|C2orf69 | 1800.2922 | 0.8971 | ##### | 6.5010 | 7.9789E-11 | 3.7030E-10 |
| ENSG00000166716\|ZNF592 | 4030.9835 | 0.7853 | ##### | 6.4982 | 8.1260E-11 | 3.7707E-10 |
| ENSG00000229967\|MAGEA4-AS1 | 281.8116 | 1.0990 | ##### | 6.4981 | 8.1358E-11 | 3.7747E-10 |
| ENSG00000176209\|SMIM19 | 853.7826 | -0.9463 | ##### | -6.4976 | 8.1604E-11 | 3.7855E-10 |
| ENSG00000133818\|RRAS2 | 3246.8513 | -0.8425 | ##### | -6.4969 | 8.1976E-11 | 3.8022E-10 |
| ENSG00000267984 | 101.1895 | -2.3906 | ##### | -6.4964 | 8.2268E-11 | 3.8151E-10 |
| ENSG00000115685\|PPP1R7 | 3395.3971 | -0.8165 | ##### | -6.4962 | 8.2388E-11 | 3.8200E-10 |
| ENSG00000147655\|RSPO2 | 580.7797 | 10.9537 | ##### | 6.4962 | 8.2398E-11 | 3.8200E-10 |
| ENSG00000100603\|SNW1 | 7994.9406 | 0.7617 | ##### | 6.4945 | 8.3318E-11 | 3.8620E-10 |
| ENSG00000249494\|DMXL1-DT | 49.6140 | -8.3040 | ##### | -6.4944 | 8.3373E-11 | 3.8640E-10 |
| ENSG00000137478\|FCHSD2 | 2708.8200 | 0.8126 | ##### | 6.4940 | 8.3608E-11 | 3.8742E-10 |
| ENSG00000101574\|METTL4 | 1610.2494 | 0.8477 | ##### | 6.4938 | 8.3691E-11 | 3.8775E-10 |
| ENSG00000183943\|PRKX | 1566.6535 | 0.8448 | ##### | 6.4927 | 8.4297E-11 | 3.9050E-10 |
| ENSG00000132357\|CARD6 | 288.9767 | -6.2708 | ##### | -6.4922 | 8.4582E-11 | 3.9176E-10 |
| ENSG00000139726\|DENR | 13559.4066 | 0.5225 | ##### | 6.4895 | 8.6113E-11 | 3.9878E-10 |
| ENSG00000100359\|SGSM3 | 2479.0549 | -0.8397 | ##### | -6.4887 | 8.6571E-11 | 4.0084E-10 |
| ENSG00000139370\|SLC15A4 | 2205.0800 | -0.7801 | ##### | -6.4874 | 8.7327E-11 | 4.0428E-10 |
| ENSG00000077420\|APBB1IP | 61.2331 | -6.6228 | ##### | -6.4863 | 8.7987E-11 | 4.0727E-10 |
| ENSG00000136352\|NKX2-1 | 100.8662 | 4.7565 | ##### | 6.4857 | 8.8346E-11 | 4.0887E-10 |
| ENSG00000175931\|UBE2O | 5879.2013 | 0.7829 | ##### | 6.4853 | 8.8570E-11 | 4.0985E-10 |
| ENSG00000268043\|NBPF12 | 1829.7876 | 0.8182 | ##### | 6.4847 | 8.8913E-11 | 4.1137E-10 |
| ENSG00000172809\|RPL38 | 18519.8867 | 0.8600 | ##### | 6.4837 | 8.9525E-11 | 4.1413E-10 |
| ENSG00000134762\|DSC3 | 1052.8753 | -0.8535 | ##### | -6.4834 | 8.9676E-11 | 4.1477E-10 |
| ENSG00000112294\|ALDH5A1 | 1058.1931 | 0.8772 | ##### | 6.4816 | 9.0734E-11 | 4.1960E-10 |
| ENSG00000239377\|ZNF775-AS1 | 40.3079 | 7.3109 | ##### | 6.4812 | 9.0973E-11 | 4.2064E-10 |
| ENSG00000181690\|PLAG1 | 118.1732 | 1.6198 | ##### | 6.4806 | 9.1388E-11 | 4.2249E-10 |
| ENSG00000168175\|MAPK1IP1L | 8818.1235 | 0.7894 | ##### | 6.4796 | 9.1943E-11 | 4.2499E-10 |
| ENSG00000166398\|GARRE1 | 1861.0322 | -0.8311 | ##### | -6.4770 | 9.3539E-11 | 4.3230E-10 |
| ENSG00000288773 | 142.9200 | -9.4581 | ##### | -6.4769 | 9.3633E-11 | 4.3267E-10 |
| ENSG00000130299\|GTPBP3 | 3064.3509 | 0.7919 | ##### | 6.4762 | 9.4058E-11 | 4.3457E-10 |
| ENSG00000136478\|TEX2 | 4594.8000 | 0.8146 | ##### | 6.4746 | 9.5059E-11 | 4.3912E-10 |
| ENSG00000001561\|ENPP4 | 727.8892 | -0.9020 | ##### | -6.4745 | 9.5127E-11 | 4.3937E-10 |
| ENSG00000204438\|GPANK1 | 1767.5778 | 0.8453 | ##### | 6.4731 | 9.6013E-11 | 4.4340E-10 |
| ENSG00000268049 | 93.6019 | -2.7058 | ##### | -6.4723 | 9.6532E-11 | 4.4572E-10 |
| ENSG00000118514\|ALDH8A1 | 47.9440 | 4.3354 | ##### | 6.4710 | 9.7337E-11 | 4.4937E-10 |
| ENSG00000164587\|RPS14 | 39050.1232 | 0.7700 | ##### | 6.4695 | 9.8297E-11 | 4.5373E-10 |
| ENSG00000196504\|PRPF40A | 17698.0713 | -0.6975 | ##### | -6.4687 | 9.8841E-11 | 4.5617E-10 |
| ENSG00000140379\|BCL2A1 | 47.7267 | -1.8771 | ##### | -6.4682 | 9.9170E-11 | 4.5762E-10 |
| ENSG00000143416\|SELENBP1 | 1756.2902 | ####### | ##### | -6.4665 | 1.0029E-10 | 4.6274E-10 |
| ENSG00000180185\|FAHD1 | 2638.9227 | -0.8111 | ##### | -6.4660 | 1.0065E-10 | 4.6429E-10 |
| ENSG00000220472 | 258.3957 | 1.1438 | ##### | 6.4657 | 1.0083E-10 | 4.6505E-10 |
| ENSG00000103187\|COTL1 | 38496.7023 | -0.7430 | ##### | -6.4656 | 1.0088E-10 | 4.6521E-10 |
| ENSG00000185164\|NOMO2 | 16193.4884 | -0.7610 | ##### | -6.4648 | 1.0143E-10 | 4.6770E-10 |
| ENSG00000164442\|CITED2 | 5452.0551 | 0.7107 | ##### | 6.4642 | 1.0181E-10 | 4.6938E-10 |
| ENSG00000178662\|CSRNP3 | 53.9682 | -3.3839 | ##### | -6.4627 | 1.0287E-10 | 4.7416E-10 |
| ENSG00000226124\|FTCDNL1 | 261.0958 | 1.1320 | ##### | 6.4612 | 1.0387E-10 | 4.7872E-10 |
| ENSG00000146054\|TRIM7 | 1662.5504 | -0.8244 | ##### | -6.4611 | 1.0393E-10 | 4.7891E-10 |
| ENSG00000103423\|DNAJA3 | 9572.9785 | -0.7165 | ##### | -6.4609 | 1.0409E-10 | 4.7957E-10 |
| ENSG00000140450\|ARRDC4 | 151.0104 | -1.5590 | ##### | -6.4598 | 1.0484E-10 | 4.8296E-10 |
| ENSG00000230091\|TMEM254-AS1 | 91.5410 | -1.6336 | ##### | -6.4596 | 1.0498E-10 | 4.8355E-10 |
| ENSG00000007341\|ST7L | 2218.1653 | 0.8283 | ##### | 6.4595 | 1.0508E-10 | 4.8394E-10 |
| ENSG00000179546\|HTR1D | 1333.7129 | 0.8698 | ##### | 6.4580 | 1.0611E-10 | 4.8853E-10 |
| ENSG00000166035\|LIPC | 22.8105 | -3.0576 | ##### | -6.4580 | 1.0611E-10 | 4.8853E-10 |
| ENSG00000010278\|CD9 | 16542.3790 | 0.8012 | ##### | 6.4577 | 1.0628E-10 | 4.8923E-10 |

| | | | | | | |
|---|---|---|---|---|---|---|
| ENSG00000182168\|UNC5C | 52.6767 | 2.3927 | ##### | 6.4575 | 1.0644E-10 | 4.8990E-10 |
| ENSG00000162620\|LRRIQ3 | 82.4378 | -1.6327 | ##### | -6.4567 | 1.0698E-10 | 4.9227E-10 |
| ENSG00000171827\|ZNF570 | 681.9314 | -0.9260 | ##### | -6.4548 | 1.0833E-10 | 4.9843E-10 |
| ENSG00000263001\|GTF2I | 15667.9795 | -0.7620 | ##### | -6.4540 | 1.0895E-10 | 5.0120E-10 |
| ENSG00000017797\|RALBP1 | 11623.5662 | 0.7434 | ##### | 6.4527 | 1.0989E-10 | 5.0544E-10 |
| ENSG00000111696\|NT5DC3 | 1965.2151 | 0.7777 | ##### | 6.4508 | 1.1127E-10 | 5.1172E-10 |
| ENSG00000165006\|UBAP1 | 5108.2146 | -0.6561 | ##### | -6.4506 | 1.1143E-10 | 5.1239E-10 |
| ENSG00000198909\|MAP3K3 | 1732.2098 | -0.8087 | ##### | -6.4500 | 1.1183E-10 | 5.1412E-10 |
| ENSG00000279561 | 96.4894 | -1.4284 | ##### | -6.4482 | 1.1318E-10 | 5.2025E-10 |
| ENSG00000089486\|CDIP1 | 1864.2336 | -0.8079 | ##### | -6.4479 | 1.1343E-10 | 5.2131E-10 |
| ENSG00000102189\|EEA1 | 1775.8297 | -0.8561 | ##### | -6.4463 | 1.1461E-10 | 5.2668E-10 |
| ENSG00000064309\|CDON | 1562.5132 | 0.8432 | ##### | 6.4460 | 1.1482E-10 | 5.2755E-10 |
| ENSG00000084093\|REST | 4550.0338 | 0.7810 | ##### | 6.4450 | 1.1557E-10 | 5.3092E-10 |
| ENSG00000230797\|YY2 | 349.3200 | 1.0709 | ##### | 6.4450 | 1.1562E-10 | 5.3109E-10 |
| ENSG00000291150\|LSP1P5 | 964.8074 | 0.8837 | ##### | 6.4448 | 1.1574E-10 | 5.3152E-10 |
| ENSG00000185055\|EFCAB10 | 333.0050 | 1.2820 | ##### | 6.4431 | 1.1708E-10 | 5.3761E-10 |
| ENSG00000110274\|CEP164 | 3520.8352 | 0.8064 | ##### | 6.4420 | 1.1789E-10 | 5.4125E-10 |
| ENSG00000118503\|TNFAIP3 | 4711.5262 | -0.7979 | ##### | -6.4420 | 1.1792E-10 | 5.4131E-10 |
| ENSG00000285402 | 27.5765 | -2.7318 | ##### | -6.4403 | 1.1923E-10 | 5.4724E-10 |
| ENSG00000287192 | 20.0199 | 4.8652 | ##### | 6.4392 | 1.2007E-10 | 5.5101E-10 |
| ENSG00000235257\|ITGA9-AS1 | 55.9178 | -1.8238 | ##### | -6.4383 | 1.2085E-10 | 5.5448E-10 |
| ENSG00000113658\|SMAD5 | 5147.0321 | 0.7698 | ##### | 6.4380 | 1.2102E-10 | 5.5518E-10 |
| ENSG00000159423\|ALDH4A1 | 1522.5739 | -0.8767 | ##### | -6.4376 | 1.2138E-10 | 5.5673E-10 |
| ENSG00000171970\|ZNF57 | 597.7039 | 2.0858 | ##### | 6.4374 | 1.2155E-10 | 5.5746E-10 |
| ENSG00000070444\|MNT | 2608.7366 | 0.8306 | ##### | 6.4364 | 1.2236E-10 | 5.6106E-10 |
| ENSG00000175866\|BAIAP2 | 1572.2273 | -0.8138 | ##### | -6.4363 | 1.2242E-10 | 5.6126E-10 |
| ENSG00000170802\|FOXN2 | 4469.9911 | 0.8032 | ##### | 6.4361 | 1.2260E-10 | 5.6201E-10 |
| ENSG00000172575\|RASGRP1 | 164.4072 | -1.5835 | ##### | -6.4352 | 1.2333E-10 | 5.6528E-10 |
| ENSG00000145779\|TNFAIP8 | 1526.0792 | 0.8781 | ##### | 6.4351 | 1.2338E-10 | 5.6539E-10 |
| ENSG00000290098 | 35.4998 | 7.4473 | ##### | 6.4344 | 1.2397E-10 | 5.6804E-10 |
| ENSG00000185163\|DDX51 | 3642.7442 | 0.8051 | ##### | 6.4341 | 1.2420E-10 | 5.6898E-10 |
| ENSG00000134597\|RBMX2 | 1891.8819 | 0.8177 | ##### | 6.4329 | 1.2519E-10 | 5.7343E-10 |
| ENSG00000180354\|MTURN | 930.6427 | 0.9269 | ##### | 6.4325 | 1.2553E-10 | 5.7493E-10 |
| ENSG00000105221\|AKT2 | 10319.5705 | -0.7400 | ##### | -6.4323 | 1.2567E-10 | 5.7545E-10 |
| ENSG00000230747 | 50.3308 | 5.4437 | ##### | 6.4265 | 1.3058E-10 | 5.9785E-10 |
| ENSG00000107736\|CDH23 | 153.5093 | 1.4749 | ##### | 6.4258 | 1.3117E-10 | 6.0045E-10 |
| ENSG00000234690\|EPCAM-DT | 44.0913 | -4.8193 | ##### | -6.4255 | 1.3147E-10 | 6.0173E-10 |
| ENSG00000198727\|MT-CYB | ########## | 0.7997 | ##### | 6.4250 | 1.3186E-10 | 6.0344E-10 |
| ENSG00000175216\|CKAP5 | 14496.9095 | 0.7681 | ##### | 6.4246 | 1.3225E-10 | 6.0511E-10 |
| ENSG00000103653\|CSK | 4583.6788 | 0.7932 | ##### | 6.4232 | 1.3346E-10 | 6.1058E-10 |
| ENSG00000185022\|MAFF | 3077.3304 | 0.8004 | ##### | 6.4231 | 1.3357E-10 | 6.1097E-10 |
| ENSG00000159314\|ARHGAP27 | 2780.8189 | 0.8134 | ##### | 6.4217 | 1.3472E-10 | 6.1615E-10 |
| ENSG00000138660\|AP1AR | 3573.2922 | 0.8046 | ##### | 6.4202 | 1.3611E-10 | 6.2240E-10 |
| ENSG00000138750\|NUP54 | 3342.9388 | 0.7660 | ##### | 6.4195 | 1.3672E-10 | 6.2511E-10 |
| ENSG00000065600\|PACC1 | 1303.8396 | -0.7894 | ##### | -6.4192 | 1.3697E-10 | 6.2614E-10 |
| ENSG00000086548\|CEACAM6 | 4892.2401 | ####### | ##### | -6.4180 | 1.3810E-10 | 6.3120E-10 |
| ENSG00000290710\|HLA-V | 58.1260 | -6.1930 | ##### | -6.4138 | 1.4195E-10 | 6.4870E-10 |
| ENSG00000277342 | 41.6968 | 2.3493 | ##### | 6.4134 | 1.4226E-10 | 6.4995E-10 |
| ENSG00000100814\|CCNB1IP1 | 1950.2736 | 0.8377 | ##### | 6.4135 | 1.4225E-10 | 6.4995E-10 |
| ENSG00000163932\|PRKCD | 1344.5987 | -0.8296 | ##### | -6.4115 | 1.4409E-10 | 6.5820E-10 |
| ENSG00000179855\|GIPC3 | 499.8000 | -1.0244 | ##### | -6.4098 | 1.4572E-10 | 6.6555E-10 |
| ENSG00000139865\|TTC6 | 49.7387 | -1.8904 | ##### | -6.4096 | 1.4593E-10 | 6.6640E-10 |
| ENSG00000236552\|RPL13AP5 | 3353.7192 | -0.8022 | ##### | -6.4081 | 1.4731E-10 | 6.7258E-10 |
| ENSG00000224729\|PCOLCE-AS1 | 48.4271 | 2.1144 | ##### | 6.4080 | 1.4744E-10 | 6.7309E-10 |
| ENSG00000175376\|EIF1AD | 2003.5904 | 0.6221 | ##### | 6.4077 | 1.4771E-10 | 6.7420E-10 |
| ENSG00000121297\|TSHZ3 | 323.2075 | 5.2527 | ##### | 6.4058 | 1.4957E-10 | 6.8258E-10 |
| ENSG00000100105\|PATZ1 | 2967.3065 | 0.8014 | ##### | 6.4053 | 1.5012E-10 | 6.8501E-10 |
| ENSG00000130813\|SHFL | 1621.2539 | -0.8092 | ##### | -6.4044 | 1.5094E-10 | 6.8863E-10 |
| ENSG00000180914\|OXTR | 376.8503 | -1.0429 | ##### | -6.4015 | 1.5390E-10 | 7.0201E-10 |
| ENSG00000141385\|AFG3L2 | 15081.9784 | 0.7256 | ##### | 6.4012 | 1.5419E-10 | 7.0324E-10 |
| ENSG00000138814\|PPP3CA | 3774.1745 | -0.6757 | ##### | -6.4012 | 1.5421E-10 | 7.0324E-10 |
| ENSG00000151773\|CCDC122 | 407.5411 | -1.0538 | ##### | -6.4003 | 1.5505E-10 | 7.0697E-10 |
| ENSG00000160194\|NDUFV3 | 2449.7207 | -0.7971 | ##### | -6.3991 | 1.5625E-10 | 7.1231E-10 |

| | | | | | | |
|---|---|---|---|---|---|---|
| ENSG00000178226\|PRSS36 | 77.1871 | -1.7162 | ##### | -6.3984 | 1.5698E-10 | 7.1552E-10 |
| ENSG00000213889\|PPM1N | 199.4322 | 1.3315 | ##### | 6.3983 | 1.5711E-10 | 7.1602E-10 |
| ENSG00000293192 | 365.6627 | 1.1037 | ##### | 6.3980 | 1.5740E-10 | 7.1723E-10 |
| ENSG00000132600\|PRMT7 | 2457.0895 | 0.7933 | ##### | 6.3966 | 1.5889E-10 | 7.2391E-10 |
| ENSG00000225953\|SATB2-AS1 | 300.9978 | 1.0490 | ##### | 6.3965 | 1.5894E-10 | 7.2405E-10 |
| ENSG00000184545\|DUSP8 | 673.6123 | 0.8498 | ##### | 6.3965 | 1.5900E-10 | 7.2420E-10 |
| ENSG00000047346\|ATOSA | 750.9176 | -0.9082 | ##### | -6.3963 | 1.5920E-10 | 7.2498E-10 |
| ENSG00000227857 | 37.5850 | -7.6688 | ##### | -6.3951 | 1.6043E-10 | 7.3046E-10 |
| ENSG00000293246\|ZNF321P | 153.5563 | -6.0144 | ##### | -6.3944 | 1.6121E-10 | 7.3393E-10 |
| ENSG00000119772\|DNMT3A | 1957.1066 | -0.8043 | ##### | -6.3924 | 1.6332E-10 | 7.4340E-10 |
| ENSG00000175279\|CENPS | 1808.3259 | -0.8655 | ##### | -6.3922 | 1.6352E-10 | 7.4420E-10 |
| ENSG00000127334\|DYRK2 | 3599.5277 | 0.7975 | ##### | 6.3916 | 1.6415E-10 | 7.4697E-10 |
| ENSG00000125970\|RALY | 12397.8032 | 0.7116 | ##### | 6.3915 | 1.6424E-10 | 7.4724E-10 |
| ENSG00000102977\|ACD | 2865.2935 | 0.8473 | ##### | 6.3872 | 1.6892E-10 | 7.6844E-10 |
| ENSG00000132323\|ILKAP | 3174.3669 | 0.7416 | ##### | 6.3864 | 1.6988E-10 | 7.7268E-10 |
| ENSG00000161992\|PRR35 | 504.3138 | 10.6793 | ##### | 6.3851 | 1.7131E-10 | 7.7906E-10 |
| ENSG00000237781\|ADAMTSL4-AS2 | 27.8985 | 2.6508 | ##### | 6.3845 | 1.7193E-10 | 7.8180E-10 |
| ENSG00000232149\|FERP1 | 309.1484 | 1.2675 | ##### | 6.3836 | 1.7294E-10 | 7.8624E-10 |
| ENSG00000230408\|BZW1-AS1 | 39.8830 | 1.8706 | ##### | 6.3798 | 1.7726E-10 | 8.0577E-10 |
| ENSG00000162542\|TMCO4 | 1133.0898 | -0.8960 | ##### | -6.3787 | 1.7865E-10 | 8.1195E-10 |
| ENSG00000074356\|NCBP3 | 5067.8265 | 0.7830 | ##### | 6.3782 | 1.7922E-10 | 8.1445E-10 |
| ENSG00000143374\|TARS2 | 4422.3713 | 0.7886 | ##### | 6.3780 | 1.7942E-10 | 8.1522E-10 |
| ENSG00000172831\|CES2 | 3238.6332 | -0.8243 | ##### | -6.3779 | 1.7952E-10 | 8.1543E-10 |
| ENSG00000164402\|SEPTIN8 | 1893.9447 | -0.7943 | ##### | -6.3779 | 1.7952E-10 | 8.1543E-10 |
| ENSG00000188603\|CLN3 | 2996.7867 | -0.8365 | ##### | -6.3776 | 1.7987E-10 | 8.1688E-10 |
| ENSG00000139826\|ABHD13 | 1594.9030 | -0.8577 | ##### | -6.3771 | 1.8049E-10 | 8.1957E-10 |
| ENSG00000262576\|PCDHGA4 | 607.7183 | -8.7491 | ##### | -6.3767 | 1.8095E-10 | 8.2156E-10 |
| ENSG00000170160\|CCDC144A | 50.7134 | -4.1099 | ##### | -6.3752 | 1.8278E-10 | 8.2972E-10 |
| ENSG00000174574\|AKIRIN1 | 14988.1048 | 0.5011 | ##### | 6.3739 | 1.8423E-10 | 8.3617E-10 |
| ENSG00000268104\|SLC6A14 | 46.0794 | -2.0755 | ##### | -6.3738 | 1.8434E-10 | 8.3656E-10 |
| ENSG00000134864\|GGACT | 236.4730 | -1.1105 | ##### | -6.3738 | 1.8440E-10 | 8.3671E-10 |
| ENSG00000126461\|SCAF1 | 13631.5046 | -0.7356 | ##### | -6.3730 | 1.8543E-10 | 8.4122E-10 |
| ENSG00000196072\|BLOC1S2 | 2587.2594 | -0.7809 | ##### | -6.3722 | 1.8632E-10 | 8.4514E-10 |
| ENSG00000165275\|TRMT10B | 906.9772 | 0.8881 | ##### | 6.3721 | 1.8650E-10 | 8.4582E-10 |
| ENSG00000254814 | 59.9566 | 1.6793 | ##### | 6.3720 | 1.8663E-10 | 8.4629E-10 |
| ENSG00000127980\|PEX1 | 1912.5807 | -0.6216 | ##### | -6.3716 | 1.8709E-10 | 8.4826E-10 |
| ENSG00000056291\|NPFFR2 | 533.2882 | -8.6105 | ##### | -6.3715 | 1.8724E-10 | 8.4882E-10 |
| ENSG00000131849\|ZNF132 | 362.8841 | -1.0570 | ##### | -6.3711 | 1.8762E-10 | 8.5040E-10 |
| ENSG00000137501\|SYTL2 | 815.1606 | -0.8525 | ##### | -6.3693 | 1.8987E-10 | 8.6045E-10 |
| ENSG00000214253\|FIS1 | 3292.5221 | 0.8013 | ##### | 6.3689 | 1.9033E-10 | 8.6231E-10 |
| ENSG00000287166 | 42.1926 | 2.4612 | ##### | 6.3689 | 1.9033E-10 | 8.6231E-10 |
| ENSG00000101391\|CDK5RAP1 | 2822.7319 | 0.7884 | ##### | 6.3688 | 1.9049E-10 | 8.6287E-10 |
| ENSG00000288398 | 480.6883 | -1.0235 | ##### | -6.3684 | 1.9103E-10 | 8.6519E-10 |
| ENSG00000267530\|LINC01836 | 225.2721 | -5.0637 | ##### | -6.3666 | 1.9323E-10 | 8.7504E-10 |
| ENSG00000198034\|RPS4X | 65384.7079 | 0.7049 | ##### | 6.3662 | 1.9379E-10 | 8.7742E-10 |
| ENSG00000291104\|BTN2A3P | 191.4929 | 1.8603 | ##### | 6.3648 | 1.9553E-10 | 8.8516E-10 |
| ENSG00000115363\|EVA1A | 90.0941 | 2.0759 | ##### | 6.3645 | 1.9591E-10 | 8.8677E-10 |
| ENSG00000010803\|SCMH1 | 2642.0411 | -0.7935 | ##### | -6.3644 | 1.9610E-10 | 8.8747E-10 |
| ENSG00000142556\|ZNF614 | 1295.5590 | -0.8328 | ##### | -6.3639 | 1.9668E-10 | 8.8997E-10 |
| ENSG00000257698\|GIHCG | 781.8529 | 0.8546 | ##### | 6.3634 | 1.9728E-10 | 8.9257E-10 |
| ENSG00000088543\|C3orf18 | 871.5754 | 0.8968 | ##### | 6.3628 | 1.9806E-10 | 8.9594E-10 |
| ENSG00000104998\|IL27RA | 3670.7394 | -0.7793 | ##### | -6.3614 | 1.9996E-10 | 9.0440E-10 |
| ENSG00000103995\|CEP152 | 1517.8505 | 0.8666 | ##### | 6.3609 | 2.0060E-10 | 9.0713E-10 |
| ENSG00000171135\|JAGN1 | 3200.1430 | -0.7712 | ##### | -6.3609 | 2.0062E-10 | 9.0713E-10 |
| ENSG00000203883\|SOX18 | 585.0158 | -0.9128 | ##### | -6.3594 | 2.0253E-10 | 9.1562E-10 |
| ENSG00000049245\|VAMP3 | 7868.6214 | -0.5382 | ##### | -6.3593 | 2.0268E-10 | 9.1614E-10 |
| ENSG00000281491\|DNAJB5-DT | 28.1365 | 7.3872 | ##### | 6.3579 | 2.0451E-10 | 9.2429E-10 |
| ENSG00000126215\|XRCC3 | 2945.5988 | 0.7951 | ##### | 6.3570 | 2.0568E-10 | 9.2942E-10 |
| ENSG00000273702 | 81.1246 | -1.7786 | ##### | -6.3560 | 2.0708E-10 | 9.3561E-10 |
| ENSG00000176393\|RNPEP | 9292.1427 | -0.7688 | ##### | -6.3558 | 2.0729E-10 | 9.3643E-10 |
| ENSG00000105402\|NAPA | 7121.2760 | -0.7662 | ##### | -6.3556 | 2.0760E-10 | 9.3766E-10 |
| ENSG00000268903 | 59.6210 | 1.6466 | ##### | 6.3550 | 2.0839E-10 | 9.4111E-10 |
| ENSG00000105976\|MET | 27179.0456 | -0.7539 | ##### | -6.3541 | 2.0972E-10 | 9.4696E-10 |

| | | | | | | |
|---|---|---|---|---|---|---|
| ENSG00000174501\|ANKRD36C | 187.9637 | 1.0907 | ##### | 6.3526 | 2.1165E-10 | 9.5556E-10 |
| ENSG00000136824\|SMC2 | 7923.8182 | 0.7654 | ##### | 6.3511 | 2.1372E-10 | 9.6476E-10 |
| ENSG00000106608\|URGCP | 2796.9793 | -0.8148 | ##### | -6.3499 | 2.1539E-10 | 9.7215E-10 |
| ENSG00000002746\|HECW1 | 867.2141 | ####### | ##### | -6.3498 | 2.1560E-10 | 9.7295E-10 |
| ENSG00000184185\|KCNJ12 | 289.3926 | 1.0939 | ##### | 6.3479 | 2.1831E-10 | 9.8501E-10 |
| ENSG00000119977\|TCTN3 | 4574.2399 | -0.8211 | ##### | -6.3469 | 2.1967E-10 | 9.9100E-10 |
| ENSG00000137198\|GMPR | 162.4322 | 5.5783 | ##### | 6.3454 | 2.2188E-10 | 1.0008E-09 |
| ENSG00000135587\|SMPD2 | 1265.6462 | 0.8289 | ##### | 6.3453 | 2.2202E-10 | 1.0013E-09 |
| ENSG00000168702\|LRP1B | 115.0065 | 1.5936 | ##### | 6.3440 | 2.2389E-10 | 1.0096E-09 |
| ENSG00000157870\|PRXL2B | 2330.5583 | 0.7982 | ##### | 6.3429 | 2.2551E-10 | 1.0167E-09 |
| ENSG00000099910\|KLHL22 | 693.6193 | -1.0045 | ##### | -6.3422 | 2.2649E-10 | 1.0210E-09 |
| ENSG00000250474\|WBP1LP2 | 109.8941 | -1.1893 | ##### | -6.3406 | 2.2887E-10 | 1.0316E-09 |
| ENSG00000162972\|MAIP1 | 1859.2706 | 0.8808 | ##### | 6.3400 | 2.2977E-10 | 1.0355E-09 |
| ENSG00000288851 | 152.3617 | 1.2253 | ##### | 6.3398 | 2.3002E-10 | 1.0364E-09 |
| ENSG00000137804\|NUSAP1 | 5811.2573 | 0.7638 | ##### | 6.3392 | 2.3103E-10 | 1.0408E-09 |
| ENSG00000288961 | 164.6504 | -1.2488 | ##### | -6.3380 | 2.3275E-10 | 1.0484E-09 |
| ENSG00000113758\|DBN1 | 14087.2064 | 0.4792 | ##### | 6.3378 | 2.3299E-10 | 1.0494E-09 |
| ENSG00000268739 | 14.0732 | -3.3338 | ##### | -6.3369 | 2.3446E-10 | 1.0558E-09 |
| ENSG00000115266\|APC2 | 1174.1143 | 0.8147 | ##### | 6.3366 | 2.3495E-10 | 1.0579E-09 |
| ENSG00000178177\|LCORL | 1371.5896 | -0.8337 | ##### | -6.3352 | 2.3699E-10 | 1.0669E-09 |
| ENSG00000138182\|KIF20B | 4384.1330 | 0.7920 | ##### | 6.3347 | 2.3786E-10 | 1.0706E-09 |
| ENSG00000089163\|SIRT4 | 99.5810 | 1.4123 | ##### | 6.3345 | 2.3812E-10 | 1.0716E-09 |
| ENSG00000105136\|ZNF419 | 504.8601 | -0.9715 | ##### | -6.3341 | 2.3869E-10 | 1.0740E-09 |
| ENSG00000251485\|NRBF2P6 | 49.8886 | 1.8030 | ##### | 6.3333 | 2.3995E-10 | 1.0796E-09 |
| ENSG00000105643\|ARRDC2 | 3577.6426 | -0.7466 | ##### | -6.3330 | 2.4045E-10 | 1.0817E-09 |
| ENSG00000229124\|VIM-AS1 | 17119.3318 | 0.8652 | ##### | 6.3327 | 2.4091E-10 | 1.0835E-09 |
| ENSG00000198707\|CEP290 | 2681.5771 | -0.7695 | ##### | -6.3321 | 2.4185E-10 | 1.0876E-09 |
| ENSG00000089558\|KCNH4 | 69.3110 | -1.8460 | ##### | -6.3320 | 2.4205E-10 | 1.0884E-09 |
| ENSG00000290474\|GUSBP2 | 60.7942 | 2.1984 | ##### | 6.3309 | 2.4372E-10 | 1.0957E-09 |
| ENSG00000292986\|DUXAP10 | 4869.0829 | 0.7451 | ##### | 6.3289 | 2.4698E-10 | 1.1102E-09 |
| ENSG00000230795\|HLA-K | 305.3967 | 0.9774 | ##### | 6.3282 | 2.4796E-10 | 1.1144E-09 |
| ENSG00000105778\|AVL9 | 6585.1585 | -0.7677 | ##### | -6.3273 | 2.4944E-10 | 1.1209E-09 |
| ENSG00000115596\|WNT6 | 63.1769 | 2.0667 | ##### | 6.3247 | 2.5369E-10 | 1.1398E-09 |
| ENSG00000127419\|TMEM175 | 697.4216 | -0.8357 | ##### | -6.3241 | 2.5470E-10 | 1.1442E-09 |
| ENSG00000149781\|FERMT3 | 135.5421 | 1.3244 | ##### | 6.3240 | 2.5493E-10 | 1.1450E-09 |
| ENSG00000119314\|PTBP3 | 10482.9412 | -0.7581 | ##### | -6.3229 | 2.5676E-10 | 1.1531E-09 |
| ENSG00000203722\|RAET1G | 104.9375 | -3.3123 | ##### | -6.3223 | 2.5765E-10 | 1.1569E-09 |
| ENSG00000251273\|LINC02228 | 68.2827 | -4.9244 | ##### | -6.3206 | 2.6061E-10 | 1.1700E-09 |
| ENSG00000169139\|UBE2V2 | 7182.9057 | -0.6019 | ##### | -6.3203 | 2.6101E-10 | 1.1716E-09 |
| ENSG00000134294\|SLC38A2 | 26788.8853 | 0.7594 | ##### | 6.3187 | 2.6383E-10 | 1.1841E-09 |
| ENSG00000229214\|LINC00242 | 42.0471 | 3.2557 | ##### | 6.3173 | 2.6621E-10 | 1.1947E-09 |
| ENSG00000242114\|MTFP1 | 2471.3416 | 0.7288 | ##### | 6.3158 | 2.6874E-10 | 1.2058E-09 |
| ENSG00000132300\|PTCD3 | 7759.4276 | 0.7896 | ##### | 6.3154 | 2.6940E-10 | 1.2086E-09 |
| ENSG00000134440\|NARS1 | 22245.5366 | -0.7183 | ##### | -6.3153 | 2.6966E-10 | 1.2096E-09 |
| ENSG00000256955 | 29.3780 | 2.9883 | ##### | 6.3141 | 2.7182E-10 | 1.2191E-09 |
| ENSG00000068394\|GPKOW | 2027.4534 | -0.8175 | ##### | -6.3134 | 2.7295E-10 | 1.2240E-09 |
| ENSG00000070731\|ST6GALNAC2 | 71.7596 | -2.2719 | ##### | -6.3127 | 2.7413E-10 | 1.2291E-09 |
| ENSG00000136274\|NACAD | 55.3208 | 1.9796 | ##### | 6.3122 | 2.7514E-10 | 1.2334E-09 |
| ENSG00000053900\|ANAPC4 | 2237.0188 | 0.5120 | ##### | 6.3108 | 2.7766E-10 | 1.2445E-09 |
| ENSG00000129083\|COPB1 | 13176.8120 | -0.7418 | ##### | -6.3100 | 2.7906E-10 | 1.2506E-09 |
| ENSG00000151914\|DST | 9527.9284 | -0.8019 | ##### | -6.3093 | 2.8023E-10 | 1.2557E-09 |
| ENSG00000230453\|ANKRD18B | 380.0347 | -1.0297 | ##### | -6.3090 | 2.8078E-10 | 1.2580E-09 |
| ENSG00000169609\|C15orf40 | 1108.8644 | -0.8253 | ##### | -6.3087 | 2.8144E-10 | 1.2607E-09 |
| ENSG00000164332\|UBLCP1 | 4336.6974 | 0.6317 | ##### | 6.3072 | 2.8414E-10 | 1.2726E-09 |
| ENSG00000188931\|CFAP126 | 431.1824 | 0.9448 | ##### | 6.3072 | 2.8419E-10 | 1.2727E-09 |
| ENSG00000274180\|NATD1 | 413.0283 | 1.1535 | ##### | 6.3061 | 2.8624E-10 | 1.2816E-09 |
| ENSG00000266903\|CEACAM16-AS1 | 52.7365 | -2.2525 | ##### | -6.3048 | 2.8856E-10 | 1.2919E-09 |
| ENSG00000141012\|GALNS | 4258.9340 | -0.6948 | ##### | -6.3047 | 2.8867E-10 | 1.2921E-09 |
| ENSG00000229047 | 24.2133 | -2.3357 | ##### | -6.3045 | 2.8918E-10 | 1.2942E-09 |
| ENSG00000167981\|ZNF597 | 246.1511 | -5.8433 | ##### | -6.3035 | 2.9092E-10 | 1.3018E-09 |
| ENSG00000234741\|GAS5 | 6217.8371 | -0.8108 | ##### | -6.3032 | 2.9158E-10 | 1.3046E-09 |
| ENSG00000140995\|DEF8 | 3753.0709 | -0.8052 | ##### | -6.3024 | 2.9309E-10 | 1.3111E-09 |
| ENSG00000116752\|BCAS2 | 5087.7757 | 0.6868 | ##### | 6.3018 | 2.9417E-10 | 1.3158E-09 |

| | | | | | | |
|---|---|---|---|---|---|---|
| ENSG00000188483\|IER5L | 7352.7566 | -0.7644 | ##### | -6.3011 | 2.9548E-10 | 1.3214E-09 |
| ENSG00000159556\|ISL2 | 484.2446 | 0.9974 | ##### | 6.3009 | 2.9590E-10 | 1.3231E-09 |
| ENSG00000144401\|METTL21A | 3884.8833 | 0.7566 | ##### | 6.3008 | 2.9613E-10 | 1.3239E-09 |
| ENSG00000101182\|PSMA7 | 25648.0540 | 0.7518 | ##### | 6.3005 | 2.9677E-10 | 1.3266E-09 |
| ENSG00000128951\|DUT | 7119.0128 | 0.7898 | ##### | 6.3004 | 2.9682E-10 | 1.3266E-09 |
| ENSG00000182481\|KPNA2 | 32888.3117 | 0.7542 | ##### | 6.2987 | 3.0018E-10 | 1.3415E-09 |
| ENSG00000261236\|BOP1 | 20013.2862 | 0.7509 | ##### | 6.2967 | 3.0403E-10 | 1.3584E-09 |
| ENSG00000131475\|VPS25 | 3613.8574 | -0.7829 | ##### | -6.2948 | 3.0789E-10 | 1.3755E-09 |
| ENSG00000124107\|SLPI | 2913.2551 | ####### | ##### | -6.2939 | 3.0954E-10 | 1.3826E-09 |
| ENSG00000105717\|PBX4 | 225.5603 | -1.2239 | ##### | -6.2915 | 3.1441E-10 | 1.4042E-09 |
| ENSG00000163026\|WDCP | 2436.4034 | 0.8169 | ##### | 6.2901 | 3.1731E-10 | 1.4169E-09 |
| ENSG00000197852\|INKA2 | 275.0656 | 1.2351 | ##### | 6.2899 | 3.1765E-10 | 1.4182E-09 |
| ENSG00000133069\|TMCC2 | 481.1328 | 0.9515 | ##### | 6.2896 | 3.1822E-10 | 1.4206E-09 |
| ENSG00000139354\|GAS2L3 | 1631.2411 | 1.0620 | ##### | 6.2895 | 3.1841E-10 | 1.4212E-09 |
| ENSG00000171729\|TMEM51 | 1715.7236 | -0.8191 | ##### | -6.2886 | 3.2026E-10 | 1.4293E-09 |
| ENSG00000206075\|SERPINB5 | 107.7097 | -6.5746 | ##### | -6.2884 | 3.2081E-10 | 1.4315E-09 |
| ENSG00000283696 | 34.5960 | 2.3074 | ##### | 6.2883 | 3.2098E-10 | 1.4320E-09 |
| ENSG00000196419\|XRCC6 | 37886.1875 | 0.4261 | ##### | 6.2882 | 3.2111E-10 | 1.4324E-09 |
| ENSG00000293324 | 52.4044 | 2.2800 | ##### | 6.2881 | 3.2128E-10 | 1.4330E-09 |
| ENSG00000169435\|RASSF6 | 3668.8735 | ####### | ##### | -6.2873 | 3.2313E-10 | 1.4410E-09 |
| ENSG00000226539\|MLXP1 | 1175.4409 | 9.7463 | ##### | 6.2847 | 3.2853E-10 | 1.4649E-09 |
| ENSG00000122547\|EEPD1 | 963.1724 | -0.8161 | ##### | -6.2844 | 3.2906E-10 | 1.4670E-09 |
| ENSG00000254422 | 460.4324 | 1.0254 | ##### | 6.2839 | 3.3024E-10 | 1.4720E-09 |
| ENSG00000185262\|UBALD2 | 2935.2402 | 0.7961 | ##### | 6.2837 | 3.3066E-10 | 1.4737E-09 |
| ENSG00000272447 | 254.2310 | -0.9841 | ##### | -6.2822 | 3.3391E-10 | 1.4880E-09 |
| ENSG00000152455\|SUV39H2 | 2576.4051 | 0.7891 | ##### | 6.2818 | 3.3467E-10 | 1.4911E-09 |
| ENSG00000124104\|SNX21 | 1787.3742 | 0.8517 | ##### | 6.2816 | 3.3515E-10 | 1.4930E-09 |
| ENSG00000125089\|SH3TC1 | 769.5448 | -9.5397 | ##### | -6.2798 | 3.3897E-10 | 1.5098E-09 |
| ENSG00000116701\|NCF2 | 62.1091 | -4.8318 | ##### | -6.2785 | 3.4183E-10 | 1.5223E-09 |
| ENSG00000182873\|PRKCZ-AS1 | 94.2917 | 1.7039 | ##### | 6.2767 | 3.4573E-10 | 1.5395E-09 |
| ENSG00000132005\|RFX1 | 1835.5040 | 0.7830 | ##### | 6.2767 | 3.4581E-10 | 1.5396E-09 |
| ENSG00000230918\|DPP4-DT | 17.6723 | -7.5588 | ##### | -6.2755 | 3.4856E-10 | 1.5516E-09 |
| ENSG00000069535\|MAOB | 3902.7210 | ####### | ##### | -6.2751 | 3.4949E-10 | 1.5555E-09 |
| ENSG00000169016\|E2F6 | 3163.2195 | 0.7806 | ##### | 6.2748 | 3.5000E-10 | 1.5576E-09 |
| ENSG00000214192\|UBE2V1P2 | 159.6665 | 1.2768 | ##### | 6.2745 | 3.5086E-10 | 1.5612E-09 |
| ENSG00000288947 | 72.0587 | 1.8829 | ##### | 6.2722 | 3.5606E-10 | 1.5840E-09 |
| ENSG00000239713\|APOBEC3G | 461.0780 | 10.4209 | ##### | 6.2717 | 3.5706E-10 | 1.5882E-09 |
| ENSG00000226210\|WASH8P | 2242.2927 | 0.8071 | ##### | 6.2711 | 3.5847E-10 | 1.5943E-09 |
| ENSG00000139625\|MAP3K12 | 3241.8176 | 0.7459 | ##### | 6.2702 | 3.6066E-10 | 1.6038E-09 |
| ENSG00000130475\|FCHO1 | 2755.4235 | 0.8033 | ##### | 6.2684 | 3.6483E-10 | 1.6221E-09 |
| ENSG00000168890\|TMEM150A | 981.2008 | -0.8406 | ##### | -6.2675 | 3.6679E-10 | 1.6306E-09 |
| ENSG00000159217\|IGF2BP1 | 10258.7332 | 0.7409 | ##### | 6.2668 | 3.6849E-10 | 1.6379E-09 |
| ENSG00000130762\|ARHGEF16 | 692.6424 | -0.8396 | ##### | -6.2650 | 3.7281E-10 | 1.6568E-09 |
| ENSG00000290033 | 62.5758 | -1.7164 | ##### | -6.2648 | 3.7319E-10 | 1.6583E-09 |
| ENSG00000113649\|TCERG1 | 8924.1599 | 0.7397 | ##### | 6.2639 | 3.7541E-10 | 1.6679E-09 |
| ENSG00000168936\|TMEM129 | 1906.7320 | -0.7852 | ##### | -6.2635 | 3.7647E-10 | 1.6724E-09 |
| ENSG00000085514\|PILRA | 99.4849 | 1.4965 | ##### | 6.2630 | 3.7767E-10 | 1.6774E-09 |
| ENSG00000144357\|UBR3 | 4231.3069 | 0.7759 | ##### | 6.2620 | 3.8016E-10 | 1.6883E-09 |
| ENSG00000289444 | 91.9198 | -1.7256 | ##### | -6.2617 | 3.8092E-10 | 1.6914E-09 |
| ENSG00000124613\|ZNF391 | 413.6774 | 1.0054 | ##### | 6.2614 | 3.8154E-10 | 1.6939E-09 |
| ENSG00000225973\|PIGBOS1 | 492.8824 | -1.0806 | ##### | -6.2610 | 3.8244E-10 | 1.6976E-09 |
| ENSG00000239382\|ALKBH6 | 1582.2312 | -0.8180 | ##### | -6.2598 | 3.8553E-10 | 1.7111E-09 |
| ENSG00000280347 | 138.3925 | 1.4634 | ##### | 6.2597 | 3.8584E-10 | 1.7122E-09 |
| ENSG00000169981\|ZNF35 | 261.9826 | -8.3010 | ##### | -6.2584 | 3.8899E-10 | 1.7259E-09 |
| ENSG00000267742\|SINHCAFP2 | 39.8548 | 2.3377 | ##### | 6.2568 | 3.9285E-10 | 1.7428E-09 |
| ENSG00000164532\|TBX20 | 64.3825 | 2.0516 | ##### | 6.2560 | 3.9500E-10 | 1.7521E-09 |
| ENSG00000144034\|TPRKB | 2395.1231 | 0.8006 | ##### | 6.2556 | 3.9595E-10 | 1.7560E-09 |
| ENSG00000101412\|E2F1 | 6013.2348 | 0.6868 | ##### | 6.2542 | 3.9949E-10 | 1.7715E-09 |
| ENSG00000164061\|BSN | 172.3961 | 1.3258 | ##### | 6.2542 | 3.9959E-10 | 1.7716E-09 |
| ENSG00000101230\|ISM1 | 85.5791 | -1.5626 | ##### | -6.2535 | 4.0135E-10 | 1.7792E-09 |
| ENSG00000187492\|CDHR4 | 154.3433 | -9.6942 | ##### | -6.2531 | 4.0239E-10 | 1.7835E-09 |
| ENSG00000140943\|MBTPS1 | 11464.7570 | -0.7285 | ##### | -6.2529 | 4.0282E-10 | 1.7852E-09 |
| ENSG00000178093\|TSSK6 | 176.5773 | 1.1937 | ##### | 6.2521 | 4.0507E-10 | 1.7949E-09 |

| | | | | | | |
|---|---|---|---|---|---|---|
| ENSG00000109519\|GRPEL1 | 5362.6496 | 0.7881 | ##### | 6.2517 | 4.0592E-10 | 1.7984E-09 |
| ENSG00000205981\|DNAJC19 | 2223.1473 | 0.8125 | ##### | 6.2516 | 4.0632E-10 | 1.7999E-09 |
| ENSG00000271425\|NBPF10 | 1154.8872 | 0.8673 | ##### | 6.2512 | 4.0727E-10 | 1.8038E-09 |
| ENSG00000187730\|GABRD | 149.3225 | 1.9187 | ##### | 6.2511 | 4.0754E-10 | 1.8047E-09 |
| ENSG00000263002\|ZNF234 | 624.7418 | -0.8963 | ##### | -6.2510 | 4.0785E-10 | 1.8058E-09 |
| ENSG00000183426\|NPIPA1 | 2122.5503 | 0.8598 | ##### | 6.2503 | 4.0960E-10 | 1.8133E-09 |
| ENSG00000177879\|AP3S1 | 4863.7298 | -0.7778 | ##### | -6.2500 | 4.1045E-10 | 1.8168E-09 |
| ENSG00000184220\|CMSS1 | 4669.2583 | 0.7776 | ##### | 6.2497 | 4.1131E-10 | 1.8204E-09 |
| ENSG00000089876\|DHX32 | 4479.8304 | -0.8001 | ##### | -6.2481 | 4.1551E-10 | 1.8387E-09 |
| ENSG00000144320\|LNPK | 2507.3219 | -0.7927 | ##### | -6.2455 | 4.2244E-10 | 1.8691E-09 |
| ENSG00000066827\|ZFAT | 917.4319 | 0.8514 | ##### | 6.2446 | 4.2493E-10 | 1.8798E-09 |
| ENSG00000084734\|GCKR | 1412.4018 | ####### | ##### | -6.2444 | 4.2542E-10 | 1.8817E-09 |
| ENSG00000114982\|KANSL3 | 9672.9773 | 0.7301 | ##### | 6.2430 | 4.2928E-10 | 1.8985E-09 |
| ENSG00000187210\|GCNT1 | 620.4223 | -0.9372 | ##### | -6.2417 | 4.3297E-10 | 1.9145E-09 |
| ENSG00000261934\|PCDHGA9 | 39.6108 | -7.0055 | ##### | -6.2404 | 4.3648E-10 | 1.9298E-09 |
| ENSG00000171219\|CDC42BPG | 126.1273 | -1.4981 | ##### | -6.2383 | 4.4244E-10 | 1.9558E-09 |
| ENSG00000243566\|UPK3B | 291.1164 | -8.2450 | ##### | -6.2379 | 4.4349E-10 | 1.9602E-09 |
| ENSG00000186205\|MTARC1 | 353.7828 | -1.0245 | ##### | -6.2363 | 4.4799E-10 | 1.9798E-09 |
| ENSG00000288796 | 73.0165 | -5.5142 | ##### | -6.2336 | 4.5583E-10 | 2.0141E-09 |
| ENSG00000165752\|STK32C | 1894.7048 | -0.8252 | ##### | -6.2324 | 4.5924E-10 | 2.0289E-09 |
| ENSG00000053524\|MCF2L2 | 283.8631 | -1.1384 | ##### | -6.2323 | 4.5952E-10 | 2.0298E-09 |
| ENSG00000122025\|FLT3 | 395.6831 | 10.3040 | ##### | 6.2323 | 4.5972E-10 | 2.0304E-09 |
| ENSG00000096696\|DSP | 3569.5661 | 0.7953 | ##### | 6.2320 | 4.6045E-10 | 2.0333E-09 |
| ENSG00000120137\|PANK3 | 8168.3648 | 0.7656 | ##### | 6.2309 | 4.6374E-10 | 2.0476E-09 |
| ENSG00000183155\|RABIF | 1439.8614 | -0.7803 | ##### | -6.2293 | 4.6861E-10 | 2.0685E-09 |
| ENSG00000168275\|COA6 | 2593.2293 | -0.7949 | ##### | -6.2292 | 4.6870E-10 | 2.0685E-09 |
| ENSG00000131508\|UBE2D2 | 7278.4386 | 0.7605 | ##### | 6.2293 | 4.6866E-10 | 2.0685E-09 |
| ENSG00000087470\|DNM1L | 13516.0455 | 0.5571 | ##### | 6.2285 | 4.7077E-10 | 2.0774E-09 |
| ENSG00000162408\|NOL9 | 3673.9530 | 0.7540 | ##### | 6.2278 | 4.7299E-10 | 2.0868E-09 |
| ENSG00000197943\|PLCG2 | 841.0160 | 0.9065 | ##### | 6.2273 | 4.7439E-10 | 2.0924E-09 |
| ENSG00000253200 | 636.0271 | 1.0654 | ##### | 6.2274 | 4.7436E-10 | 2.0924E-09 |
| ENSG00000197903\|H2BC12 | 5379.3992 | 0.7710 | ##### | 6.2271 | 4.7506E-10 | 2.0950E-09 |
| ENSG00000171811\|CFAP46 | 133.4758 | 1.5247 | ##### | 6.2260 | 4.7839E-10 | 2.1094E-09 |
| ENSG00000162086\|ZNF75A | 1407.1703 | -0.7894 | ##### | -6.2249 | 4.8189E-10 | 2.1245E-09 |
| ENSG00000165282\|PIGO | 4892.5376 | 0.7628 | ##### | 6.2232 | 4.8706E-10 | 2.1470E-09 |
| ENSG00000253978\|TENM2-AS1 | 33.7560 | 3.0946 | ##### | 6.2224 | 4.8959E-10 | 2.1578E-09 |
| ENSG00000106078\|COBL | 176.3245 | -1.7463 | ##### | -6.2223 | 4.8979E-10 | 2.1584E-09 |
| ENSG00000227359\|LINC02936 | 37.1106 | -1.9025 | ##### | -6.2216 | 4.9209E-10 | 2.1682E-09 |
| ENSG00000237264\|FTH1P11 | 26.4760 | -2.3896 | ##### | -6.2211 | 4.9360E-10 | 2.1746E-09 |
| ENSG00000166432\|ZMAT1 | 347.6669 | 1.0448 | ##### | 6.2191 | 5.0010E-10 | 2.2029E-09 |
| ENSG00000061273\|HDAC7 | 7102.7013 | -0.7661 | ##### | -6.2167 | 5.0763E-10 | 2.2357E-09 |
| ENSG00000130748\|TMEM160 | 1365.3692 | -0.8193 | ##### | -6.2119 | 5.2332E-10 | 2.3045E-09 |
| ENSG00000290718\|FAM95B1 | 31.5363 | -4.3434 | ##### | -6.2113 | 5.2536E-10 | 2.3131E-09 |
| ENSG00000172046\|USP19 | 6140.1594 | 0.7498 | ##### | 6.2112 | 5.2583E-10 | 2.3149E-09 |
| ENSG00000177483\|RBM44 | 81.8633 | 1.4760 | ##### | 6.2111 | 5.2626E-10 | 2.3164E-09 |
| ENSG00000187634\|SAMD11 | 2793.6431 | -0.9049 | ##### | -6.2105 | 5.2827E-10 | 2.3249E-09 |
| ENSG00000275234 | 59.1361 | -5.3513 | ##### | -6.2104 | 5.2854E-10 | 2.3258E-09 |
| ENSG00000147687\|TATDN1 | 3054.5293 | -0.8101 | ##### | -6.2103 | 5.2869E-10 | 2.3261E-09 |
| ENSG00000134882\|UBAC2 | 4856.8161 | -0.7782 | ##### | -6.2100 | 5.2980E-10 | 2.3306E-09 |
| ENSG00000264769\|MAFG-AS1 | 66.8785 | -1.7795 | ##### | -6.2094 | 5.3182E-10 | 2.3391E-09 |
| ENSG00000283849 | 28.4069 | 3.2375 | ##### | 6.2093 | 5.3214E-10 | 2.3402E-09 |
| ENSG00000276351 | 20.3421 | -2.2527 | ##### | -6.2088 | 5.3397E-10 | 2.3479E-09 |
| ENSG00000285108 | 59.1849 | 2.0252 | ##### | 6.2080 | 5.3646E-10 | 2.3585E-09 |
| ENSG00000133121\|STARD13 | 1364.1033 | 0.8290 | ##### | 6.2075 | 5.3842E-10 | 2.3668E-09 |
| ENSG00000136982\|DSCC1 | 2685.8036 | 0.7808 | ##### | 6.2061 | 5.4316E-10 | 2.3873E-09 |
| ENSG00000166949\|SMAD3 | 20572.8222 | 0.7365 | ##### | 6.2058 | 5.4402E-10 | 2.3907E-09 |
| ENSG00000233750\|CICP27 | 29.2334 | 2.4924 | ##### | 6.2058 | 5.4411E-10 | 2.3907E-09 |
| ENSG00000164022\|AIMP1 | 5894.8189 | -0.7447 | ##### | -6.2053 | 5.4606E-10 | 2.3989E-09 |
| ENSG00000134575\|ACP2 | 2279.2824 | -0.7977 | ##### | -6.2052 | 5.4628E-10 | 2.3996E-09 |
| ENSG00000198324\|PHETA1 | 988.6023 | -0.8138 | ##### | -6.2041 | 5.5010E-10 | 2.4160E-09 |
| ENSG00000184465\|WDR27 | 951.9655 | -0.7979 | ##### | -6.2026 | 5.5530E-10 | 2.4385E-09 |
| ENSG00000139197\|PEX5 | 6747.5802 | 0.6967 | ##### | 6.2021 | 5.5710E-10 | 2.4460E-09 |
| ENSG00000139684\|ESD | 4709.2083 | -0.7657 | ##### | -6.2020 | 5.5734E-10 | 2.4467E-09 |

| | | | | | | |
|---|---|---|---|---|---|---|
| ENSG00000162729\|IGSF8 | 3261.4496 | 0.7777 | ##### | 6.2020 | 5.5759E-10 | 2.4474E-09 |
| ENSG00000287966\|LARRPM | 81.6447 | 1.6260 | ##### | 6.2003 | 5.6361E-10 | 2.4735E-09 |
| ENSG00000104738\|MCM4 | 25914.4998 | 0.7009 | ##### | 6.1983 | 5.7089E-10 | 2.5051E-09 |
| ENSG00000102683\|SGCG | 43.7086 | 6.9300 | ##### | 6.1977 | 5.7282E-10 | 2.5132E-09 |
| ENSG00000175170\|FAM182B | 123.3363 | 1.5101 | ##### | 6.1972 | 5.7466E-10 | 2.5209E-09 |
| ENSG00000185716\|MOSMO | 1690.8782 | -0.6787 | ##### | -6.1972 | 5.7487E-10 | 2.5215E-09 |
| ENSG00000182912\|TSPEAR-AS2 | 74.4132 | 2.6422 | ##### | 6.1971 | 5.7524E-10 | 2.5227E-09 |
| ENSG00000177822\|TENM3-AS1 | 211.8095 | 6.4696 | ##### | 6.1968 | 5.7607E-10 | 2.5260E-09 |
| ENSG00000197170\|PSMD12 | 11351.1451 | 0.6079 | ##### | 6.1967 | 5.7657E-10 | 2.5278E-09 |
| ENSG00000282988 | 544.0009 | 1.0235 | ##### | 6.1961 | 5.7868E-10 | 2.5367E-09 |
| ENSG00000038382\|TRIO | 16554.2805 | 0.7530 | ##### | 6.1960 | 5.7902E-10 | 2.5378E-09 |
| ENSG00000141968\|VAV1 | 392.4421 | -0.9629 | ##### | -6.1955 | 5.8082E-10 | 2.5453E-09 |
| ENSG00000149636\|DSN1 | 3469.2816 | 0.7329 | ##### | 6.1952 | 5.8197E-10 | 2.5499E-09 |
| ENSG00000234337 | 89.6605 | 1.4814 | ##### | 6.1952 | 5.8211E-10 | 2.5502E-09 |
| ENSG00000104205\|SGK3 | 842.2639 | -0.8593 | ##### | -6.1949 | 5.8319E-10 | 2.5546E-09 |
| ENSG00000143776\|CDC42BPA | 5718.5206 | -0.7471 | ##### | -6.1940 | 5.8642E-10 | 2.5679E-09 |
| ENSG00000228170 | 116.2807 | -1.4111 | ##### | -6.1941 | 5.8636E-10 | 2.5679E-09 |
| ENSG00000174028\|FAM3C2P | 302.7449 | 1.0988 | ##### | 6.1938 | 5.8744E-10 | 2.5720E-09 |
| ENSG00000255398\|HCAR3 | 56.3768 | -7.9237 | ##### | -6.1918 | 5.9479E-10 | 2.6038E-09 |
| ENSG00000168291\|PDHB | 6520.9946 | 0.7602 | ##### | 6.1916 | 5.9573E-10 | 2.6076E-09 |
| ENSG00000006283\|CACNA1G | 650.1273 | 0.9488 | ##### | 6.1909 | 5.9808E-10 | 2.6175E-09 |
| ENSG00000290873\|LINC03009 | 934.7399 | 0.8700 | ##### | 6.1893 | 6.0449E-10 | 2.6452E-09 |
| ENSG00000065615\|CYB5R4 | 2936.4148 | 0.7459 | ##### | 6.1850 | 6.2105E-10 | 2.7172E-09 |
| ENSG00000272106 | 271.3559 | -0.9835 | ##### | -6.1848 | 6.2175E-10 | 2.7199E-09 |
| ENSG00000140307\|GTF2A2 | 4499.9647 | 0.7647 | ##### | 6.1845 | 6.2293E-10 | 2.7246E-09 |
| ENSG00000175785\|PRIMA1 | 68.8690 | 4.4879 | ##### | 6.1830 | 6.2903E-10 | 2.7509E-09 |
| ENSG00000186532\|SMYD4 | 1640.1685 | 0.7709 | ##### | 6.1827 | 6.3013E-10 | 2.7553E-09 |
| ENSG00000275395\|FCGBP | 210.8354 | -1.2063 | ##### | -6.1824 | 6.3145E-10 | 2.7607E-09 |
| ENSG00000128000\|ZNF780B | 693.9707 | -0.8585 | ##### | -6.1819 | 6.3317E-10 | 2.7678E-09 |
| ENSG00000049246\|PER3 | 696.9688 | 0.9646 | ##### | 6.1803 | 6.3999E-10 | 2.7968E-09 |
| ENSG00000272189\|MAP7-AS1 | 94.8302 | -1.7545 | ##### | -6.1803 | 6.3992E-10 | 2.7968E-09 |
| ENSG00000122203\|KIAA1191 | 9571.6225 | 0.6117 | ##### | 6.1796 | 6.4280E-10 | 2.8087E-09 |
| ENSG00000171720\|HDAC3 | 8025.9536 | -0.5175 | ##### | -6.1793 | 6.4371E-10 | 2.8122E-09 |
| ENSG00000242247\|ARFGAP3 | 2924.1638 | -0.7575 | ##### | -6.1783 | 6.4795E-10 | 2.8303E-09 |
| ENSG00000090273\|NUDC | 15715.5981 | 0.7324 | ##### | 6.1779 | 6.4958E-10 | 2.8371E-09 |
| ENSG00000129493\|HEATR5A | 5920.3657 | -0.7596 | ##### | -6.1766 | 6.5517E-10 | 2.8610E-09 |
| ENSG00000068878\|PSME4 | 8399.4293 | 0.7374 | ##### | 6.1764 | 6.5593E-10 | 2.8639E-09 |
| ENSG00000130255\|RPL36 | 16446.9355 | 0.7710 | ##### | 6.1748 | 6.6245E-10 | 2.8920E-09 |
| ENSG00000197587\|DMBX1 | 46.7731 | 2.5901 | ##### | 6.1741 | 6.6546E-10 | 2.9043E-09 |
| ENSG00000221916\|C19orf73 | 76.1233 | -1.5432 | ##### | -6.1741 | 6.6538E-10 | 2.9043E-09 |
| ENSG00000213839\|TMX2P1 | 511.0700 | 0.8951 | ##### | 6.1736 | 6.6767E-10 | 2.9135E-09 |
| ENSG00000131196\|NFATC1 | 1510.0165 | -0.7825 | ##### | -6.1729 | 6.7067E-10 | 2.9262E-09 |
| ENSG00000236641 | 24.6236 | -8.0372 | ##### | -6.1727 | 6.7147E-10 | 2.9292E-09 |
| ENSG00000262691 | 116.3183 | -1.5795 | ##### | -6.1720 | 6.7441E-10 | 2.9416E-09 |
| ENSG00000095380\|NANS | 5961.1003 | -0.7414 | ##### | -6.1712 | 6.7784E-10 | 2.9562E-09 |
| ENSG00000138018\|SELENOI | 10320.2096 | -0.7523 | ##### | -6.1693 | 6.8592E-10 | 2.9909E-09 |
| ENSG00000123570\|RAB9B | 117.5582 | 1.2367 | ##### | 6.1691 | 6.8662E-10 | 2.9936E-09 |
| ENSG00000290993 | 589.1193 | 0.8897 | ##### | 6.1689 | 6.8771E-10 | 2.9979E-09 |
| ENSG00000273284 | 33.4902 | 2.6976 | ##### | 6.1684 | 6.8981E-10 | 3.0066E-09 |
| ENSG00000137818\|RPLP1 | 49435.4726 | 0.7551 | ##### | 6.1674 | 6.9430E-10 | 3.0257E-09 |
| ENSG00000255760\|LINC02422 | 58.0660 | -1.6165 | ##### | -6.1666 | 6.9754E-10 | 3.0394E-09 |
| ENSG00000115866\|DARS1 | 10822.0418 | -0.5994 | ##### | -6.1653 | 7.0366E-10 | 3.0656E-09 |
| ENSG00000100027\|YPEL1 | 176.1043 | 1.1757 | ##### | 6.1649 | 7.0535E-10 | 3.0725E-09 |
| ENSG00000236675\|MTX1P1 | 1982.9510 | 0.7849 | ##### | 6.1648 | 7.0582E-10 | 3.0741E-09 |
| ENSG00000150779\|TIMM8B | 2421.5850 | -0.7607 | ##### | -6.1639 | 7.0957E-10 | 3.0900E-09 |
| ENSG00000144043\|TEX261 | 10666.4008 | 0.7605 | ##### | 6.1635 | 7.1148E-10 | 3.0979E-09 |
| ENSG00000027869\|SH2D2A | 386.1504 | 1.0682 | ##### | 6.1634 | 7.1204E-10 | 3.0992E-09 |
| ENSG00000026508\|CD44 | 19859.9705 | -0.7575 | ##### | -6.1634 | 7.1203E-10 | 3.0992E-09 |
| ENSG00000281832\|LINC00602 | 127.5391 | -1.3654 | ##### | -6.1634 | 7.1209E-10 | 3.0992E-09 |
| ENSG00000122420\|PTGFR | 123.3003 | -4.2633 | ##### | -6.1630 | 7.1372E-10 | 3.1058E-09 |
| ENSG00000255874\|PRECSIT | 62.1514 | 2.6424 | ##### | 6.1628 | 7.1470E-10 | 3.1096E-09 |
| ENSG00000280402 | 25.2785 | 2.7759 | ##### | 6.1624 | 7.1658E-10 | 3.1174E-09 |
| ENSG00000083635\|NUFIP1 | 1333.0651 | 0.8371 | ##### | 6.1621 | 7.1779E-10 | 3.1222E-09 |

| | | | | | | |
|---|---|---|---|---|---|---|
| ENSG00000013523\|ANGEL1 | 2841.0382 | 0.7604 | ##### | 6.1606 | 7.2486E-10 | 3.1525E-09 |
| ENSG00000168621\|GDNF | 82.7709 | 4.0547 | ##### | 6.1601 | 7.2693E-10 | 3.1610E-09 |
| ENSG00000197406\|DIO3 | 3248.4971 | ####### | ##### | -6.1598 | 7.2855E-10 | 3.1676E-09 |
| ENSG00000102738\|MRPS31 | 1226.2551 | 0.8595 | ##### | 6.1596 | 7.2950E-10 | 3.1713E-09 |
| ENSG00000198840\|MT-ND3 | 59318.5726 | 0.8120 | ##### | 6.1594 | 7.3043E-10 | 3.1749E-09 |
| ENSG00000163728\|TTC14 | 3315.8166 | 0.7399 | ##### | 6.1592 | 7.3127E-10 | 3.1780E-09 |
| ENSG00000100243\|CYB5R3 | 9347.8401 | -0.7431 | ##### | -6.1588 | 7.3314E-10 | 3.1857E-09 |
| ENSG00000172638\|EFEMP2 | 222.2233 | 1.3501 | ##### | 6.1582 | 7.3566E-10 | 3.1962E-09 |
| ENSG00000005812\|FBXL3 | 2507.8086 | 0.8802 | ##### | 6.1566 | 7.4338E-10 | 3.2292E-09 |
| ENSG00000144026\|ZNF514 | 1229.6859 | 0.9129 | ##### | 6.1564 | 7.4438E-10 | 3.2331E-09 |
| ENSG00000081665\|ZNF506 | 310.0578 | -1.0727 | ##### | -6.1552 | 7.4968E-10 | 3.2557E-09 |
| ENSG00000167701\|GPT | 186.0232 | -1.1605 | ##### | -6.1543 | 7.5431E-10 | 3.2753E-09 |
| ENSG00000263235 | 112.2426 | 1.8123 | ##### | 6.1538 | 7.5644E-10 | 3.2840E-09 |
| ENSG00000229728 | 40.8380 | 7.9967 | ##### | 6.1537 | 7.5689E-10 | 3.2856E-09 |
| ENSG00000165338\|HECTD2 | 1238.3069 | -0.8369 | ##### | -6.1536 | 7.5741E-10 | 3.2873E-09 |
| ENSG00000139117\|CPNE8 | 3813.3380 | 0.7863 | ##### | 6.1531 | 7.5972E-10 | 3.2969E-09 |
| ENSG00000101333\|PLCB4 | 3093.4671 | 0.7734 | ##### | 6.1522 | 7.6395E-10 | 3.3147E-09 |
| ENSG00000173208\|ABCD2 | 317.7848 | 10.0657 | ##### | 6.1506 | 7.7194E-10 | 3.3489E-09 |
| ENSG00000290439\|HSD17B7P2 | 51.5263 | -2.2688 | ##### | -6.1494 | 7.7764E-10 | 3.3731E-09 |
| ENSG00000277693 | 53.7098 | -2.0811 | ##### | -6.1476 | 7.8668E-10 | 3.4119E-09 |
| ENSG00000275494 | 52.9426 | -2.2174 | ##### | -6.1459 | 7.9520E-10 | 3.4483E-09 |
| ENSG00000224109\|CENPVL3 | 87.4864 | 4.0547 | ##### | 6.1448 | 8.0086E-10 | 3.4724E-09 |
| ENSG00000165795\|NDRG2 | 376.3034 | 0.8571 | ##### | 6.1447 | 8.0139E-10 | 3.4742E-09 |
| ENSG00000259129\|LINC00648 | 192.9150 | -1.0651 | ##### | -6.1439 | 8.0514E-10 | 3.4899E-09 |
| ENSG00000167874\|TMEM88 | 35.6181 | 2.4615 | ##### | 6.1437 | 8.0627E-10 | 3.4943E-09 |
| ENSG00000148660\|CAMK2G | 3705.2322 | -0.7720 | ##### | -6.1434 | 8.0788E-10 | 3.5008E-09 |
| ENSG00000109065\|NAT9 | 3326.2001 | 0.7504 | ##### | 6.1433 | 8.0836E-10 | 3.5023E-09 |
| ENSG00000141027\|NCOR1 | 11355.4842 | 0.7400 | ##### | 6.1431 | 8.0922E-10 | 3.5056E-09 |
| ENSG00000185615\|PDIA2 | 61.5215 | -6.2234 | ##### | -6.1410 | 8.1996E-10 | 3.5516E-09 |
| ENSG00000089116\|LHX5 | 97.3577 | 4.8191 | ##### | 6.1386 | 8.3268E-10 | 3.6062E-09 |
| ENSG00000141510\|TP53 | 4961.9248 | 0.7847 | ##### | 6.1374 | 8.3894E-10 | 3.6327E-09 |
| ENSG00000276430\|FAM25C | 35.3504 | -8.4604 | ##### | -6.1364 | 8.4390E-10 | 3.6537E-09 |
| ENSG00000015133\|CCDC88C | 1333.6506 | 0.8041 | ##### | 6.1361 | 8.4586E-10 | 3.6616E-09 |
| ENSG00000266338\|NBPF15 | 4543.7440 | 0.7304 | ##### | 6.1360 | 8.4642E-10 | 3.6635E-09 |
| ENSG00000123384\|LRP1 | 4845.6708 | -0.7888 | ##### | -6.1357 | 8.4774E-10 | 3.6687E-09 |
| ENSG00000171634\|BPTF | 7246.3662 | 0.7581 | ##### | 6.1352 | 8.5034E-10 | 3.6794E-09 |
| ENSG00000144161\|ZC3H8 | 1375.7713 | -0.8237 | ##### | -6.1345 | 8.5429E-10 | 3.6960E-09 |
| ENSG00000213222\|TOMM40P4 | 95.1146 | 1.5172 | ##### | 6.1330 | 8.6213E-10 | 3.7294E-09 |
| ENSG00000136051\|WASHC4 | 7636.4557 | -0.7209 | ##### | -6.1326 | 8.6470E-10 | 3.7394E-09 |
| ENSG00000188559\|RALGAPA2 | 1684.7748 | -0.8241 | ##### | -6.1326 | 8.6467E-10 | 3.7394E-09 |
| ENSG00000166888\|STAT6 | 7482.9607 | -0.7169 | ##### | -6.1307 | 8.7517E-10 | 3.7841E-09 |
| ENSG00000153179\|RASSF3 | 7326.4587 | -0.5619 | ##### | -6.1299 | 8.7956E-10 | 3.8026E-09 |
| ENSG00000139915\|MDGA2 | 410.2944 | -1.0535 | ##### | -6.1292 | 8.8326E-10 | 3.8180E-09 |
| ENSG00000004897\|CDC27 | 12736.8998 | 0.7254 | ##### | 6.1290 | 8.8406E-10 | 3.8209E-09 |
| ENSG00000181409\|AATK | 321.8333 | 1.0485 | ##### | 6.1290 | 8.8422E-10 | 3.8210E-09 |
| ENSG00000274565 | 93.3748 | 1.4479 | ##### | 6.1285 | 8.8720E-10 | 3.8334E-09 |
| ENSG00000254166\|CASC19 | 164.7875 | -7.8369 | ##### | -6.1269 | 8.9581E-10 | 3.8700E-09 |
| ENSG00000279227 | 130.4068 | -1.2657 | ##### | -6.1269 | 8.9605E-10 | 3.8705E-09 |
| ENSG00000290600 | 179.1554 | -1.2461 | ##### | -6.1248 | 9.0819E-10 | 3.9223E-09 |
| ENSG00000107816\|LZTS2 | 3545.7977 | 0.6109 | ##### | 6.1243 | 9.1105E-10 | 3.9341E-09 |
| ENSG00000128578\|STRIP2 | 846.4364 | -0.8308 | ##### | -6.1241 | 9.1204E-10 | 3.9379E-09 |
| ENSG00000204624\|DISP3 | 50.8437 | 2.0679 | ##### | 6.1234 | 9.1606E-10 | 3.9546E-09 |
| ENSG00000259956\|RBM15B | 8199.6273 | 0.4386 | ##### | 6.1228 | 9.1917E-10 | 3.9675E-09 |
| ENSG00000079462\|PAFAH1B3 | 4363.8684 | -0.7757 | ##### | -6.1223 | 9.2212E-10 | 3.9796E-09 |
| ENSG00000213064\|SFT2D2 | 4331.3506 | -0.7757 | ##### | -6.1215 | 9.2697E-10 | 4.0000E-09 |
| ENSG00000147164\|SNX12 | 6777.4557 | 0.7585 | ##### | 6.1198 | 9.3679E-10 | 4.0418E-09 |
| ENSG00000261884 | 970.1438 | 0.8734 | ##### | 6.1197 | 9.3729E-10 | 4.0434E-09 |
| ENSG00000153162\|BMP6 | 700.4200 | 1.0543 | ##### | 6.1197 | 9.3743E-10 | 4.0434E-09 |
| ENSG00000138675\|FGF5 | 909.4704 | 9.3468 | ##### | 6.1193 | 9.3979E-10 | 4.0530E-09 |
| ENSG00000161270\|NPHS1 | 60.2870 | -1.8102 | ##### | -6.1192 | 9.4057E-10 | 4.0557E-09 |
| ENSG00000108061\|SHOC2 | 3585.5526 | -0.7307 | ##### | -6.1178 | 9.4873E-10 | 4.0904E-09 |
| ENSG00000277504 | 22.6839 | 3.5711 | ##### | 6.1174 | 9.5127E-10 | 4.1007E-09 |
| ENSG00000116750\|UCHL5 | 4481.9968 | -0.7663 | ##### | -6.1170 | 9.5344E-10 | 4.1095E-09 |

| | | | | | | |
|---|---|---|---|---|---|---|
| ENSG00000225339 | 935.6774 | -1.2455 | ##### | -6.1167 | 9.5531E-10 | 4.1169E-09 |
| ENSG00000154917\|RAB6B | 1829.2115 | 0.7574 | ##### | 6.1163 | 9.5802E-10 | 4.1280E-09 |
| ENSG00000106348\|IMPDH1 | 8469.6209 | 0.7356 | ##### | 6.1152 | 9.6421E-10 | 4.1541E-09 |
| ENSG00000086598\|TMED2 | 27755.9723 | -0.5773 | ##### | -6.1143 | 9.7009E-10 | 4.1788E-09 |
| ENSG00000170779\|CDCA4 | 4432.8521 | 0.7436 | ##### | 6.1123 | 9.8189E-10 | 4.2290E-09 |
| ENSG00000196678\|ERI2 | 1701.2623 | -0.8277 | ##### | -6.1119 | 9.8476E-10 | 4.2408E-09 |
| ENSG00000261054\|SYNM-AS2 | 451.6252 | 1.4328 | ##### | 6.1114 | 9.8781E-10 | 4.2533E-09 |
| ENSG00000014914\|MTMR11 | 1515.4937 | -0.7695 | ##### | -6.1113 | 9.8800E-10 | 4.2535E-09 |
| ENSG00000013563\|DNASE1L1 | 2154.5914 | -0.7808 | ##### | -6.1109 | 9.9054E-10 | 4.2638E-09 |
| ENSG00000137760\|ALKBH8 | 1363.2452 | 0.7880 | ##### | 6.1108 | 9.9139E-10 | 4.2669E-09 |
| ENSG00000253395\|LINC03044 | 17.5884 | -3.7848 | ##### | -6.1081 | 1.0083E-09 | 4.3390E-09 |
| ENSG00000015676\|NUDCD3 | 7285.6613 | -0.6484 | ##### | -6.1071 | 1.0148E-09 | 4.3662E-09 |
| ENSG00000176998\|HCG4 | 222.5550 | -6.9644 | ##### | -6.1045 | 1.0315E-09 | 4.4378E-09 |
| ENSG00000104522\|GFUS | 6883.8899 | 0.7561 | ##### | 6.1041 | 1.0339E-09 | 4.4474E-09 |
| ENSG00000182771\|GRID1 | 182.0241 | 1.5296 | ##### | 6.1036 | 1.0368E-09 | 4.4591E-09 |
| ENSG00000011304\|PTBP1 | 45724.0057 | 0.3703 | ##### | 6.1023 | 1.0453E-09 | 4.4952E-09 |
| ENSG00000196591\|HDAC2 | 11913.4359 | 0.7544 | ##### | 6.1009 | 1.0545E-09 | 4.5338E-09 |
| ENSG00000029993\|HMGB3 | 4523.1226 | 0.7553 | ##### | 6.0999 | 1.0616E-09 | 4.5636E-09 |
| ENSG00000289014 | 53.5487 | 6.6451 | ##### | 6.0998 | 1.0623E-09 | 4.5650E-09 |
| ENSG00000166130\|IKBIP | 6247.5676 | 0.6675 | ##### | 6.0998 | 1.0621E-09 | 4.5650E-09 |
| ENSG00000229358\|DPY19L1P1 | 184.3641 | -1.3912 | ##### | -6.0998 | 1.0623E-09 | 4.5650E-09 |
| ENSG00000143977\|SNRPG | 7084.8400 | 0.7454 | ##### | 6.0988 | 1.0687E-09 | 4.5911E-09 |
| ENSG00000100300\|TSPO | 6558.2753 | -0.7462 | ##### | -6.0988 | 1.0687E-09 | 4.5911E-09 |
| ENSG00000065989\|PDE4A | 1886.6344 | 0.8017 | ##### | 6.0984 | 1.0715E-09 | 4.6022E-09 |
| ENSG00000226471 | 47.7205 | -2.4586 | ##### | -6.0981 | 1.0732E-09 | 4.6092E-09 |
| ENSG00000130340\|SNX9 | 5418.7447 | 0.7382 | ##### | 6.0980 | 1.0739E-09 | 4.6115E-09 |
| ENSG00000250846\|EPHA5-AS1 | 38.5168 | 2.0630 | ##### | 6.0960 | 1.0872E-09 | 4.6679E-09 |
| ENSG00000165637\|VDAC2 | 15121.3251 | -0.7368 | ##### | -6.0951 | 1.0938E-09 | 4.6954E-09 |
| ENSG00000122592\|HOXA7 | 79.6295 | -3.0849 | ##### | -6.0930 | 1.1079E-09 | 4.7555E-09 |
| ENSG00000179222\|MAGED1 | 7072.4476 | -0.7256 | ##### | -6.0923 | 1.1132E-09 | 4.7773E-09 |
| ENSG00000127831\|VIL1 | 698.7807 | 1.0132 | ##### | 6.0914 | 1.1192E-09 | 4.8024E-09 |
| ENSG00000188747\|NOXA1 | 144.6871 | -1.5662 | ##### | -6.0906 | 1.1247E-09 | 4.8255E-09 |
| ENSG00000099998\|GGT5 | 43.9446 | -3.3186 | ##### | -6.0902 | 1.1275E-09 | 4.8367E-09 |
| ENSG00000100342\|APOL1 | 658.6558 | -7.8825 | ##### | -6.0869 | 1.1509E-09 | 4.9363E-09 |
| ENSG00000262528 | 79.2819 | 1.9657 | ##### | 6.0861 | 1.1571E-09 | 4.9620E-09 |
| ENSG00000205268\|PDE7A | 1615.7969 | -0.8099 | ##### | -6.0859 | 1.1585E-09 | 4.9674E-09 |
| ENSG00000213160\|KLHL23 | 1466.9724 | 0.8516 | ##### | 6.0855 | 1.1615E-09 | 4.9797E-09 |
| ENSG00000224616\|RTCA-AS1 | 79.7772 | -1.8177 | ##### | -6.0849 | 1.1659E-09 | 4.9978E-09 |
| ENSG00000135720\|DYNC1LI2 | 13313.4934 | -0.7028 | ##### | -6.0834 | 1.1764E-09 | 5.0419E-09 |
| ENSG00000163191\|S100A11 | 51500.7079 | 0.7544 | ##### | 6.0832 | 1.1783E-09 | 5.0494E-09 |
| ENSG00000230314\|ELOVL2-AS1 | 33.7258 | -7.7729 | ##### | -6.0815 | 1.1903E-09 | 5.1000E-09 |
| ENSG00000226200\|SGMS1-AS1 | 169.2645 | -1.1386 | ##### | -6.0811 | 1.1933E-09 | 5.1122E-09 |
| ENSG00000156313\|RPGR | 701.6534 | 0.9446 | ##### | 6.0810 | 1.1945E-09 | 5.1167E-09 |
| ENSG00000182676\|PPP1R27 | 37.6779 | 6.8548 | ##### | 6.0809 | 1.1952E-09 | 5.1190E-09 |
| ENSG00000169621\|APLF | 434.9496 | 0.9492 | ##### | 6.0798 | 1.2031E-09 | 5.1520E-09 |
| ENSG00000108559\|NUP88 | 8958.1367 | 0.7475 | ##### | 6.0791 | 1.2085E-09 | 5.1745E-09 |
| ENSG00000214517\|PPME1 | 7406.9603 | 0.4409 | ##### | 6.0790 | 1.2096E-09 | 5.1784E-09 |
| ENSG00000170561\|IRX2 | 76.9971 | 4.0944 | ##### | 6.0779 | 1.2175E-09 | 5.2113E-09 |
| ENSG00000163389\|POGLUT1 | 1649.1894 | 0.8348 | ##### | 6.0775 | 1.2206E-09 | 5.2241E-09 |
| ENSG00000248098\|BCKDHA | 3290.0429 | -0.7726 | ##### | -6.0764 | 1.2291E-09 | 5.2595E-09 |
| ENSG00000153391\|INO80C | 1833.1303 | -0.7902 | ##### | -6.0759 | 1.2326E-09 | 5.2740E-09 |
| ENSG00000188013\|MEIS3P2 | 161.1325 | -9.7528 | ##### | -6.0758 | 1.2336E-09 | 5.2773E-09 |
| ENSG00000156500\|PABIR3 | 155.2265 | 1.2455 | ##### | 6.0752 | 1.2385E-09 | 5.2977E-09 |
| ENSG00000206557\|TRIM71 | 351.0312 | 9.9815 | ##### | 6.0743 | 1.2452E-09 | 5.3257E-09 |
| ENSG00000122557\|HERPUD2 | 2550.8156 | -0.7699 | ##### | -6.0742 | 1.2463E-09 | 5.3294E-09 |
| ENSG00000150893\|FREM2 | 229.2284 | 5.0562 | ##### | 6.0738 | 1.2488E-09 | 5.3395E-09 |
| ENSG00000177076\|ACER2 | 96.2435 | 1.5854 | ##### | 6.0732 | 1.2539E-09 | 5.3605E-09 |
| ENSG00000189127\|ANKRD34B | 306.9353 | 9.8657 | ##### | 6.0714 | 1.2677E-09 | 5.4187E-09 |
| ENSG00000171502\|COL24A1 | 55.4581 | 1.6041 | ##### | 6.0701 | 1.2784E-09 | 5.4636E-09 |
| ENSG00000133059\|DSTYK | 2413.1275 | -0.7622 | ##### | -6.0689 | 1.2882E-09 | 5.5045E-09 |
| ENSG00000114978\|MOB1A | 17644.9144 | 0.6505 | ##### | 6.0667 | 1.3057E-09 | 5.5786E-09 |
| ENSG00000181544\|FANCB | 784.6715 | 0.8645 | ##### | 6.0632 | 1.3347E-09 | 5.7017E-09 |
| ENSG00000131408\|NR1H2 | 7007.6686 | -0.7338 | ##### | -6.0627 | 1.3386E-09 | 5.7176E-09 |

| | | | | | | |
|---|---|---|---|---|---|---|
| ENSG00000162819\|BROX | 3427.8177 | -0.7566 | ##### | -6.0619 | 1.3451E-09 | 5.7447E-09 |
| ENSG00000086504\|MRPL28 | 3654.9440 | -0.7641 | ##### | -6.0616 | 1.3481E-09 | 5.7567E-09 |
| ENSG00000106258\|CYP3A5 | 184.3196 | -1.1899 | ##### | -6.0613 | 1.3507E-09 | 5.7666E-09 |
| ENSG00000092199\|HNRNPC | 42436.0470 | 0.4795 | ##### | 6.0610 | 1.3526E-09 | 5.7740E-09 |
| ENSG00000180953\|ST20 | 533.1605 | -0.9161 | ##### | -6.0610 | 1.3531E-09 | 5.7754E-09 |
| ENSG00000121621\|KIF18A | 2423.4547 | 0.8071 | ##### | 6.0603 | 1.3590E-09 | 5.7998E-09 |
| ENSG00000151276\|MAGI1 | 2153.0241 | 0.7653 | ##### | 6.0598 | 1.3633E-09 | 5.8173E-09 |
| ENSG00000104081\|BMF | 182.9838 | 1.1911 | ##### | 6.0585 | 1.3737E-09 | 5.8598E-09 |
| ENSG00000151729\|SLC25A4 | 1420.4782 | -0.8236 | ##### | -6.0586 | 1.3735E-09 | 5.8598E-09 |
| ENSG00000100393\|EP300 | 5537.8435 | 0.7383 | ##### | 6.0585 | 1.3742E-09 | 5.8613E-09 |
| ENSG00000288007 | 87.0081 | -8.7492 | ##### | -6.0568 | 1.3888E-09 | 5.9225E-09 |
| ENSG00000225746\|MEG8 | 33.1448 | 5.8436 | ##### | 6.0567 | 1.3895E-09 | 5.9247E-09 |
| ENSG00000258728 | 366.8051 | 1.2445 | ##### | 6.0558 | 1.3968E-09 | 5.9552E-09 |
| ENSG00000135916\|ITM2C | 6517.9553 | 0.7534 | ##### | 6.0555 | 1.4001E-09 | 5.9684E-09 |
| ENSG00000165494\|PCF11 | 4286.4835 | 0.7201 | ##### | 6.0539 | 1.4136E-09 | 6.0249E-09 |
| ENSG00000204256\|BRD2 | 21879.5572 | 0.6010 | ##### | 6.0527 | 1.4243E-09 | 6.0698E-09 |
| ENSG00000232480\|TGFB2-AS1 | 28.5240 | -2.2035 | ##### | -6.0524 | 1.4267E-09 | 6.0791E-09 |
| ENSG00000068650\|ATP11A | 13743.7567 | 0.7372 | ##### | 6.0522 | 1.4284E-09 | 6.0857E-09 |
| ENSG00000262185\|LINC02861 | 336.9640 | -8.7728 | ##### | -6.0516 | 1.4342E-09 | 6.1093E-09 |
| ENSG00000133935\|ERG28 | 3011.5433 | -0.7267 | ##### | -6.0496 | 1.4517E-09 | 6.1829E-09 |
| ENSG00000149970\|CNKSR2 | 422.8584 | -1.1489 | ##### | -6.0492 | 1.4560E-09 | 6.2004E-09 |
| ENSG00000198885\|ITPRIPL1 | 1492.7964 | 11.9269 | ##### | 6.0491 | 1.4565E-09 | 6.2017E-09 |
| ENSG00000286670 | 70.5731 | -7.5153 | ##### | -6.0482 | 1.4650E-09 | 6.2370E-09 |
| ENSG00000233930\|KRTAP5-AS1 | 51.4387 | 2.5689 | ##### | 6.0473 | 1.4730E-09 | 6.2702E-09 |
| ENSG00000281468 | 70.0599 | 1.7536 | ##### | 6.0472 | 1.4738E-09 | 6.2726E-09 |
| ENSG00000162517\|PEF1 | 5010.0743 | -0.7473 | ##### | -6.0468 | 1.4771E-09 | 6.2859E-09 |
| ENSG00000197442\|MAP3K5 | 1692.4241 | 0.7179 | ##### | 6.0466 | 1.4790E-09 | 6.2919E-09 |
| ENSG00000290803 | 191.7045 | -9.9986 | ##### | -6.0466 | 1.4789E-09 | 6.2919E-09 |
| ENSG00000125618\|PAX8 | 351.3457 | 0.9868 | ##### | 6.0459 | 1.4854E-09 | 6.3185E-09 |
| ENSG00000139645\|ANKRD52 | 18080.6738 | -0.7235 | ##### | -6.0452 | 1.4919E-09 | 6.3451E-09 |
| ENSG00000123636\|BAZ2B | 2138.2754 | -0.7898 | ##### | -6.0449 | 1.4949E-09 | 6.3571E-09 |
| ENSG00000147174\|GCNA | 135.7796 | 1.5935 | ##### | 6.0442 | 1.5018E-09 | 6.3853E-09 |
| ENSG00000114107\|CEP70 | 2387.5185 | 0.7442 | ##### | 6.0435 | 1.5083E-09 | 6.4122E-09 |
| ENSG00000244879\|GABPB1-AS1 | 1062.3638 | 0.8108 | ##### | 6.0429 | 1.5134E-09 | 6.4330E-09 |
| ENSG00000165410\|CFL2 | 4978.3314 | -0.7224 | ##### | -6.0429 | 1.5138E-09 | 6.4338E-09 |
| ENSG00000197385\|ZNF860 | 211.8456 | -1.3341 | ##### | -6.0420 | 1.5225E-09 | 6.4699E-09 |
| ENSG00000168994\|PXDC1 | 2210.3392 | -0.7553 | ##### | -6.0419 | 1.5233E-09 | 6.4722E-09 |
| ENSG00000175175\|PPM1E | 338.4312 | 1.0458 | ##### | 6.0412 | 1.5293E-09 | 6.4969E-09 |
| ENSG00000110195\|FOLR1 | 37.0056 | -5.9666 | ##### | -6.0412 | 1.5302E-09 | 6.4995E-09 |
| ENSG00000075826\|SEC31B | 497.6868 | -0.8537 | ##### | -6.0404 | 1.5369E-09 | 6.5272E-09 |
| ENSG00000260588 | 126.0379 | 1.2600 | ##### | 6.0402 | 1.5393E-09 | 6.5363E-09 |
| ENSG00000114127\|XRN1 | 1825.0741 | -0.7894 | ##### | -6.0399 | 1.5424E-09 | 6.5486E-09 |
| ENSG00000132376\|INPP5K | 1663.3577 | 0.7669 | ##### | 6.0396 | 1.5453E-09 | 6.5599E-09 |
| ENSG00000270299 | 43.9984 | -2.7137 | ##### | -6.0386 | 1.5541E-09 | 6.5967E-09 |
| ENSG00000060749\|QSER1 | 6963.7237 | 0.7439 | ##### | 6.0371 | 1.5690E-09 | 6.6587E-09 |
| ENSG00000140993\|TIGD7 | 753.0030 | -0.9772 | ##### | -6.0371 | 1.5692E-09 | 6.6589E-09 |
| ENSG00000143545\|RAB13 | 7055.4446 | 0.7220 | ##### | 6.0364 | 1.5760E-09 | 6.6866E-09 |
| ENSG00000162390\|ACOT11 | 717.9835 | 0.8370 | ##### | 6.0361 | 1.5788E-09 | 6.6975E-09 |
| ENSG00000178531\|CTXN1 | 2998.6815 | 0.7685 | ##### | 6.0358 | 1.5815E-09 | 6.7080E-09 |
| ENSG00000229376\|CICP3 | 32.9450 | 2.3911 | ##### | 6.0345 | 1.5947E-09 | 6.7632E-09 |
| ENSG00000085491\|SLC25A24 | 7657.9923 | -0.7238 | ##### | -6.0329 | 1.6104E-09 | 6.8279E-09 |
| ENSG00000004838\|ZMYND10 | 105.7867 | 1.4103 | ##### | 6.0329 | 1.6103E-09 | 6.8279E-09 |
| ENSG00000170153\|RNF150 | 96.7436 | 1.4173 | ##### | 6.0321 | 1.6185E-09 | 6.8612E-09 |
| ENSG00000197044\|ZNF441 | 114.3637 | -2.2898 | ##### | -6.0315 | 1.6248E-09 | 6.8860E-09 |
| ENSG00000151576\|QTRT2 | 5071.6338 | 0.7355 | ##### | 6.0315 | 1.6246E-09 | 6.8860E-09 |
| ENSG00000131368\|MRPS25 | 5991.6161 | 1.0307 | ##### | 6.0306 | 1.6333E-09 | 6.9209E-09 |
| ENSG00000042429\|MED17 | 3110.9821 | 0.7201 | ##### | 6.0277 | 1.6631E-09 | 7.0463E-09 |
| ENSG00000135643\|KCNMB4 | 77.2164 | 4.0499 | ##### | 6.0253 | 1.6875E-09 | 7.1484E-09 |
| ENSG00000169045\|HNRNPH1 | 29971.3132 | 0.5684 | ##### | 6.0251 | 1.6901E-09 | 7.1586E-09 |
| ENSG00000127586\|CHTF18 | 3480.9520 | 0.7457 | ##### | 6.0234 | 1.7081E-09 | 7.2337E-09 |
| ENSG00000141519\|CCDC40 | 377.2193 | -0.9259 | ##### | -6.0229 | 1.7137E-09 | 7.2563E-09 |
| ENSG00000160886\|LY6K | 459.6792 | 1.0296 | ##### | 6.0225 | 1.7169E-09 | 7.2691E-09 |
| ENSG00000159860\|TCAF2P1 | 32.6201 | 2.2432 | ##### | 6.0221 | 1.7212E-09 | 7.2860E-09 |

| | | | | | | |
|---|---|---|---|---|---|---|
| ENSG00000151208\|DLG5 | 9798.9907 | -0.7190 | ##### | -6.0220 | 1.7228E-09 | 7.2919E-09 |
| ENSG00000129351\|ILF3 | 36277.7155 | 0.6865 | ##### | 6.0214 | 1.7291E-09 | 7.3173E-09 |
| ENSG00000001631\|KRIT1 | 3866.6660 | 0.7355 | ##### | 6.0210 | 1.7340E-09 | 7.3370E-09 |
| ENSG00000164900\|GBX1 | 47.8231 | 7.6055 | ##### | 6.0201 | 1.7430E-09 | 7.3743E-09 |
| ENSG00000288891 | 90.6161 | -1.3175 | ##### | -6.0200 | 1.7442E-09 | 7.3783E-09 |
| ENSG00000288844 | 197.0861 | -0.9961 | ##### | -6.0193 | 1.7521E-09 | 7.4107E-09 |
| ENSG00000118733\|OLFM3 | 1616.6045 | ####### | ##### | -6.0187 | 1.7582E-09 | 7.4355E-09 |
| ENSG00000113360\|DROSHA | 8918.8775 | 0.7044 | ##### | 6.0178 | 1.7685E-09 | 7.4779E-09 |
| ENSG00000284130\|TUBA3FP | 162.1673 | 1.8002 | ##### | 6.0172 | 1.7746E-09 | 7.5025E-09 |
| ENSG00000289089 | 46.2980 | 2.2074 | ##### | 6.0156 | 1.7917E-09 | 7.5737E-09 |
| ENSG00000172167\|MTBP | 1253.1995 | 0.7709 | ##### | 6.0154 | 1.7948E-09 | 7.5859E-09 |
| ENSG00000227057\|WDR46 | 7599.5939 | 0.7010 | ##### | 6.0150 | 1.7983E-09 | 7.5996E-09 |
| ENSG00000068654\|POLR1A | 14211.7265 | 0.7131 | ##### | 6.0146 | 1.8037E-09 | 7.6211E-09 |
| ENSG00000163530\|DPPA2 | 2577.9161 | ####### | ##### | -6.0139 | 1.8106E-09 | 7.6495E-09 |
| ENSG00000227693\|GSTM3P1 | 48.4592 | 6.9949 | ##### | 6.0116 | 1.8376E-09 | 7.7623E-09 |
| ENSG00000101882\|NKAP | 2587.3315 | 0.7698 | ##### | 6.0096 | 1.8602E-09 | 7.8568E-09 |
| ENSG00000269289 | 45.2843 | -7.5024 | ##### | -6.0085 | 1.8719E-09 | 7.9051E-09 |
| ENSG00000146729\|NIPSNAP2 | 6404.2394 | -0.5871 | ##### | -6.0073 | 1.8866E-09 | 7.9658E-09 |
| ENSG00000273010 | 58.9685 | -1.6319 | ##### | -6.0071 | 1.8888E-09 | 7.9743E-09 |
| ENSG00000170919\|TPT1-AS1 | 1139.2430 | 0.8317 | ##### | 6.0067 | 1.8939E-09 | 7.9947E-09 |
| ENSG00000124222\|STX16 | 7270.6673 | 0.6980 | ##### | 6.0065 | 1.8952E-09 | 7.9988E-09 |
| ENSG00000137338\|PGBD1 | 475.6374 | 0.8884 | ##### | 6.0060 | 1.9013E-09 | 8.0235E-09 |
| ENSG00000048405\|ZNF800 | 2544.6956 | 0.7298 | ##### | 6.0040 | 1.9256E-09 | 8.1248E-09 |
| ENSG00000259319\|JDP2-AS1 | 80.8158 | -2.3747 | ##### | -6.0036 | 1.9304E-09 | 8.1438E-09 |
| ENSG00000137055\|PLAA | 3079.2613 | -0.7749 | ##### | -6.0031 | 1.9362E-09 | 8.1673E-09 |
| ENSG00000137312\|FLOT1 | 11250.0352 | 0.6695 | ##### | 6.0030 | 1.9370E-09 | 8.1696E-09 |
| ENSG00000134910\|STT3A | 11150.0985 | -0.5730 | ##### | -6.0020 | 1.9494E-09 | 8.2206E-09 |
| ENSG00000198833\|UBE2J1 | 3870.8043 | -0.7619 | ##### | -6.0019 | 1.9506E-09 | 8.2246E-09 |
| ENSG00000287185 | 209.6785 | 1.1308 | ##### | 6.0014 | 1.9559E-09 | 8.2460E-09 |
| ENSG00000166924\|NYAP1 | 353.3286 | -0.9376 | ##### | -6.0009 | 1.9626E-09 | 8.2727E-09 |
| ENSG00000224985 | 509.6931 | 0.9196 | ##### | 6.0000 | 1.9726E-09 | 8.3139E-09 |
| ENSG00000250264 | 801.0083 | 1.2077 | ##### | 5.9995 | 1.9789E-09 | 8.3394E-09 |
| ENSG00000127220\|ABHD8 | 1578.0493 | -0.7760 | ##### | -5.9980 | 1.9980E-09 | 8.4185E-09 |
| ENSG00000100442\|FKBP3 | 6419.7767 | 0.7123 | ##### | 5.9978 | 2.0000E-09 | 8.4258E-09 |
| ENSG00000278558\|TMEM191B | 54.9326 | -2.0483 | ##### | -5.9977 | 2.0009E-09 | 8.4284E-09 |
| ENSG00000204147\|ASAH2B | 579.7291 | -0.9082 | ##### | -5.9965 | 2.0162E-09 | 8.4916E-09 |
| ENSG00000171174\|RBKS | 678.8843 | -0.9255 | ##### | -5.9960 | 2.0221E-09 | 8.5150E-09 |
| ENSG00000223546\|LINC00630 | 342.4065 | 1.0505 | ##### | 5.9959 | 2.0233E-09 | 8.5191E-09 |
| ENSG00000115325\|DOK1 | 686.2405 | -0.8623 | ##### | -5.9955 | 2.0285E-09 | 8.5397E-09 |
| ENSG00000243279\|PRAF2 | 1407.6877 | -0.8169 | ##### | -5.9952 | 2.0325E-09 | 8.5556E-09 |
| ENSG00000105668\|UPK1A | 87.6019 | 1.4315 | ##### | 5.9950 | 2.0347E-09 | 8.5633E-09 |
| ENSG00000115561\|CHMP3 | 10503.1168 | -0.3772 | ##### | -5.9949 | 2.0360E-09 | 8.5678E-09 |
| ENSG00000293486 | 101.3300 | 4.7371 | ##### | 5.9938 | 2.0497E-09 | 8.6241E-09 |
| ENSG00000085872\|CHERP | 9887.1774 | 0.6056 | ##### | 5.9937 | 2.0518E-09 | 8.6318E-09 |
| ENSG00000290315 | 7295.6546 | 0.8130 | ##### | 5.9933 | 2.0564E-09 | 8.6499E-09 |
| ENSG00000083223\|TUT7 | 2487.7960 | -0.7802 | ##### | -5.9926 | 2.0651E-09 | 8.6853E-09 |
| ENSG00000204070\|SYS1 | 3034.6468 | 0.7503 | ##### | 5.9917 | 2.0762E-09 | 8.7308E-09 |
| ENSG00000123612\|ACVR1C | 68.0392 | -2.0464 | ##### | -5.9913 | 2.0816E-09 | 8.7524E-09 |
| ENSG00000049769\|PPP1R3F | 1697.6684 | -0.7199 | ##### | -5.9893 | 2.1075E-09 | 8.8597E-09 |
| ENSG00000263884\|THOC1-DT | 45.3384 | 2.5146 | ##### | 5.9883 | 2.1210E-09 | 8.9153E-09 |
| ENSG00000068615\|REEP1 | 145.4497 | -1.3461 | ##### | -5.9879 | 2.1254E-09 | 8.9328E-09 |
| ENSG00000224940\|PRRT4 | 101.3886 | 2.9877 | ##### | 5.9863 | 2.1463E-09 | 9.0190E-09 |
| ENSG00000290421 | 24.3595 | 3.1975 | ##### | 5.9857 | 2.1543E-09 | 9.0501E-09 |
| ENSG00000137434\|C6orf52 | 56.0344 | -2.0625 | ##### | -5.9857 | 2.1543E-09 | 9.0501E-09 |
| ENSG00000139324\|TMTC3 | 4127.9465 | -0.7610 | ##### | -5.9855 | 2.1577E-09 | 9.0634E-09 |
| ENSG00000253276\|CCDC71L | 2179.9889 | -0.7549 | ##### | -5.9848 | 2.1660E-09 | 9.0970E-09 |
| ENSG00000189403\|HMGB1 | 30581.0679 | 0.6564 | ##### | 5.9848 | 2.1665E-09 | 9.0977E-09 |
| ENSG00000257097\|CLIP1-AS1 | 106.0890 | 2.9841 | ##### | 5.9830 | 2.1912E-09 | 9.2002E-09 |
| ENSG00000267493\|CIRBP-AS1 | 177.9238 | 1.1241 | ##### | 5.9826 | 2.1955E-09 | 9.2169E-09 |
| ENSG00000272079 | 146.8839 | -1.3195 | ##### | -5.9825 | 2.1969E-09 | 9.2215E-09 |
| ENSG00000167081\|PBX3 | 1352.3728 | -0.7631 | ##### | -5.9820 | 2.2038E-09 | 9.2489E-09 |
| ENSG00000073578\|SDHA | 7947.5617 | -0.7188 | ##### | -5.9820 | 2.2041E-09 | 9.2492E-09 |
| ENSG00000079156\|OSBPL6 | 1239.6483 | 0.8115 | ##### | 5.9819 | 2.2050E-09 | 9.2516E-09 |

| | | | | | | |
|---|---|---|---|---|---|---|
| ENSG00000267680\|ZNF224 | 997.1997 | -0.8491 | ##### | -5.9811 | 2.2165E-09 | 9.2987E-09 |
| ENSG00000100354\|TNRC6B | 2109.1879 | 0.7388 | ##### | 5.9809 | 2.2187E-09 | 9.3066E-09 |
| ENSG00000184254\|ALDH1A3 | 1176.5148 | -0.8007 | ##### | -5.9807 | 2.2216E-09 | 9.3172E-09 |
| ENSG00000119899\|SLC17A5 | 3194.3563 | 0.7669 | ##### | 5.9799 | 2.2323E-09 | 9.3607E-09 |
| ENSG00000291214 | 716.4586 | -0.8193 | ##### | -5.9796 | 2.2362E-09 | 9.3758E-09 |
| ENSG00000198081\|ZBTB14 | 1828.2828 | 0.7587 | ##### | 5.9769 | 2.2742E-09 | 9.5338E-09 |
| ENSG00000124575\|H1-3 | 67.5414 | 3.5963 | ##### | 5.9766 | 2.2780E-09 | 9.5483E-09 |
| ENSG00000290746\|CA5BP1 | 760.5658 | -0.8259 | ##### | -5.9748 | 2.3043E-09 | 9.6573E-09 |
| ENSG00000047621\|C12orf4 | 2794.2713 | 0.4733 | ##### | 5.9746 | 2.3059E-09 | 9.6627E-09 |
| ENSG00000204186\|ZDBF2 | 2593.7061 | 0.7473 | ##### | 5.9745 | 2.3082E-09 | 9.6711E-09 |
| ENSG00000165898\|ISCA2 | 1602.9705 | 0.8087 | ##### | 5.9743 | 2.3105E-09 | 9.6791E-09 |
| ENSG00000118096\|IFT46 | 437.8743 | -0.9279 | ##### | -5.9733 | 2.3246E-09 | 9.7370E-09 |
| ENSG00000267484 | 472.5206 | -0.9268 | ##### | -5.9719 | 2.3448E-09 | 9.8203E-09 |
| ENSG00000183495\|EP400 | 8032.9982 | 0.7340 | ##### | 5.9692 | 2.3847E-09 | 9.9859E-09 |
| ENSG00000274512\|TBC1D3L | 308.3773 | -0.9934 | ##### | -5.9691 | 2.3864E-09 | 9.9914E-09 |
| ENSG00000226549\|SCDP1 | 22.8129 | -2.6105 | ##### | -5.9682 | 2.3993E-09 | 1.0044E-08 |
| ENSG00000204560\|DHX16 | 6797.1999 | 0.6396 | ##### | 5.9680 | 2.4025E-09 | 1.0056E-08 |
| ENSG00000253719\|ATXN7L3B | 7334.2048 | -0.7192 | ##### | -5.9674 | 2.4112E-09 | 1.0091E-08 |
| ENSG00000236404\|VLDLR-AS1 | 308.4389 | ####### | ##### | -5.9671 | 2.4151E-09 | 1.0106E-08 |
| ENSG00000075539\|FRYL | 4153.3178 | 0.7303 | ##### | 5.9662 | 2.4290E-09 | 1.0163E-08 |
| ENSG00000112414\|ADGRG6 | 18111.0977 | -0.7104 | ##### | -5.9655 | 2.4391E-09 | 1.0203E-08 |
| ENSG00000269792 | 54.9007 | -2.0450 | ##### | -5.9654 | 2.4405E-09 | 1.0208E-08 |
| ENSG00000006432\|MAP3K9 | 3447.1633 | -0.7499 | ##### | -5.9653 | 2.4413E-09 | 1.0210E-08 |
| ENSG00000137700\|SLC37A4 | 2030.9781 | 0.7526 | ##### | 5.9645 | 2.4545E-09 | 1.0264E-08 |
| ENSG00000198561\|CTNND1 | 11155.2279 | -0.7180 | ##### | -5.9641 | 2.4592E-09 | 1.0282E-08 |
| ENSG00000276185 | 37.0391 | -2.1305 | ##### | -5.9638 | 2.4646E-09 | 1.0303E-08 |
| ENSG00000204463\|BAG6 | 22822.8923 | 0.5547 | ##### | 5.9634 | 2.4709E-09 | 1.0328E-08 |
| ENSG00000136240\|KDELR2 | 26377.7169 | -0.7489 | ##### | -5.9631 | 2.4750E-09 | 1.0344E-08 |
| ENSG00000260643 | 488.2312 | -1.7238 | ##### | -5.9624 | 2.4856E-09 | 1.0387E-08 |
| ENSG00000108468\|CBX1 | 9617.8494 | 0.7289 | ##### | 5.9622 | 2.4893E-09 | 1.0400E-08 |
| ENSG00000197122\|SRC | 3882.4985 | -0.7038 | ##### | -5.9613 | 2.5027E-09 | 1.0455E-08 |
| ENSG00000158290\|CUL4B | 8240.0357 | 0.7091 | ##### | 5.9608 | 2.5105E-09 | 1.0486E-08 |
| ENSG00000111358\|GTF2H3 | 4668.3504 | -0.7354 | ##### | -5.9607 | 2.5114E-09 | 1.0487E-08 |
| ENSG00000011028\|MRC2 | 12965.4683 | 0.6998 | ##### | 5.9607 | 2.5111E-09 | 1.0487E-08 |
| ENSG00000063176\|SPHK2 | 1716.5566 | -0.7227 | ##### | -5.9602 | 2.5195E-09 | 1.0519E-08 |
| ENSG00000142534\|RPS11 | 49317.7607 | -0.7025 | ##### | -5.9595 | 2.5308E-09 | 1.0565E-08 |
| ENSG00000197951\|ZNF71 | 1201.4203 | -0.7800 | ##### | -5.9581 | 2.5521E-09 | 1.0652E-08 |
| ENSG00000173473\|SMARCC1 | 10771.8595 | 0.6636 | ##### | 5.9558 | 2.5879E-09 | 1.0800E-08 |
| ENSG00000169684\|CHRNA5 | 1524.9853 | 0.7941 | ##### | 5.9547 | 2.6057E-09 | 1.0873E-08 |
| ENSG00000185379\|RAD51D | 2116.2662 | -0.7157 | ##### | -5.9520 | 2.6488E-09 | 1.1052E-08 |
| ENSG00000166275\|BORCS7 | 1497.8698 | -0.8005 | ##### | -5.9508 | 2.6677E-09 | 1.1129E-08 |
| ENSG00000107551\|RASSF4 | 112.8996 | -1.4597 | ##### | -5.9497 | 2.6865E-09 | 1.1206E-08 |
| ENSG00000179562\|GCC1 | 3546.7148 | 0.7335 | ##### | 5.9496 | 2.6885E-09 | 1.1212E-08 |
| ENSG00000117592\|PRDX6 | 18845.2664 | -0.7278 | ##### | -5.9481 | 2.7128E-09 | 1.1312E-08 |
| ENSG00000253305\|PCDHGB6 | 38.6009 | -2.8361 | ##### | -5.9481 | 2.7135E-09 | 1.1314E-08 |
| ENSG00000215845\|TSTD1 | 324.3768 | 5.7805 | ##### | 5.9476 | 2.7207E-09 | 1.1342E-08 |
| ENSG00000131100\|ATP6V1E1 | 4679.9338 | -0.7314 | ##### | -5.9473 | 2.7257E-09 | 1.1361E-08 |
| ENSG00000067596\|DHX8 | 6908.0352 | 0.7026 | ##### | 5.9441 | 2.7798E-09 | 1.1585E-08 |
| ENSG00000173113\|TRMT112 | 10299.8555 | 0.7105 | ##### | 5.9438 | 2.7850E-09 | 1.1604E-08 |
| ENSG00000115816\|CEBPZ | 12164.9660 | 0.7140 | ##### | 5.9438 | 2.7849E-09 | 1.1604E-08 |
| ENSG00000168496\|FEN1 | 6532.0434 | 0.6569 | ##### | 5.9425 | 2.8068E-09 | 1.1693E-08 |
| ENSG00000143368\|SF3B4 | 16281.6372 | 0.7281 | ##### | 5.9420 | 2.8154E-09 | 1.1727E-08 |
| ENSG00000289874 | 32.0786 | 6.6346 | ##### | 5.9415 | 2.8250E-09 | 1.1765E-08 |
| ENSG00000142677\|IL22RA1 | 316.1635 | -0.9372 | ##### | -5.9413 | 2.8286E-09 | 1.1778E-08 |
| ENSG00000250305\|TRMT9B | 176.8599 | 5.0792 | ##### | 5.9409 | 2.8338E-09 | 1.1797E-08 |
| ENSG00000184709\|LRRC26 | 56.9188 | -2.3119 | ##### | -5.9410 | 2.8336E-09 | 1.1797E-08 |
| ENSG00000174231\|PRPF8 | 29655.2198 | 0.7016 | ##### | 5.9401 | 2.8490E-09 | 1.1859E-08 |
| ENSG00000116183\|PAPPA2 | 139.9198 | -9.3519 | ##### | -5.9387 | 2.8720E-09 | 1.1953E-08 |
| ENSG00000169180\|XPO6 | 18834.7063 | 0.4924 | ##### | 5.9387 | 2.8727E-09 | 1.1954E-08 |
| ENSG00000265590\|CFAP298-TCP10L | 565.0984 | -1.4012 | ##### | -5.9381 | 2.8832E-09 | 1.1996E-08 |
| ENSG00000167635\|ZNF146 | 8690.4798 | -0.7430 | ##### | -5.9368 | 2.9060E-09 | 1.2089E-08 |
| ENSG00000154370\|TRIM11 | 3770.3082 | 0.7187 | ##### | 5.9366 | 2.9094E-09 | 1.2101E-08 |
| ENSG00000283528\|TCAF2C | 28.8551 | 2.7241 | ##### | 5.9361 | 2.9187E-09 | 1.2139E-08 |

| | | | | | | |
|---|---|---|---|---|---|---|
| ENSG00000172062\|SMN1 | 4059.3594 | 0.7746 | ##### | 5.9353 | 2.9329E-09 | 1.2196E-08 |
| ENSG00000147121\|KRBOX4 | 627.0020 | -0.8883 | ##### | -5.9352 | 2.9353E-09 | 1.2204E-08 |
| ENSG00000135299\|ANKRD6 | 306.1447 | -1.1248 | ##### | -5.9343 | 2.9507E-09 | 1.2267E-08 |
| ENSG00000139292\|LGR5 | 338.0737 | 9.5082 | ##### | 5.9336 | 2.9638E-09 | 1.2319E-08 |
| ENSG00000176714\|CCDC121 | 434.2515 | 0.8865 | ##### | 5.9335 | 2.9659E-09 | 1.2325E-08 |
| ENSG00000115977\|AAK1 | 4746.4935 | 0.7081 | ##### | 5.9335 | 2.9657E-09 | 1.2325E-08 |
| ENSG00000226245\|ZNF32-AS1 | 19.5094 | -2.9123 | ##### | -5.9334 | 2.9671E-09 | 1.2328E-08 |
| ENSG00000109016\|DHRS7B | 1572.7852 | 0.8052 | ##### | 5.9322 | 2.9888E-09 | 1.2416E-08 |
| ENSG00000125826\|RBCK1 | 4874.7542 | -0.6975 | ##### | -5.9317 | 2.9984E-09 | 1.2455E-08 |
| ENSG00000035499\|DEPDC1B | 2157.5992 | 0.7842 | ##### | 5.9309 | 3.0119E-09 | 1.2509E-08 |
| ENSG00000158270\|COLEC12 | 86.5768 | 4.2673 | ##### | 5.9302 | 3.0261E-09 | 1.2565E-08 |
| ENSG00000170775\|GPR37 | 1022.4462 | 0.7714 | ##### | 5.9302 | 3.0264E-09 | 1.2565E-08 |
| ENSG00000116096\|SPR | 7670.3003 | 0.7116 | ##### | 5.9299 | 3.0306E-09 | 1.2581E-08 |
| ENSG00000233101\|HOXB-AS3 | 37.4668 | -2.1111 | ##### | -5.9286 | 3.0546E-09 | 1.2679E-08 |
| ENSG00000047634\|SCML1 | 442.5935 | -0.9403 | ##### | -5.9268 | 3.0881E-09 | 1.2816E-08 |
| ENSG00000266708 | 49.2436 | 7.4690 | ##### | 5.9255 | 3.1126E-09 | 1.2916E-08 |
| ENSG00000135838\|NPL | 240.8707 | -1.0460 | ##### | -5.9247 | 3.1292E-09 | 1.2983E-08 |
| ENSG00000177990\|DPY19L2 | 324.6168 | 1.0277 | ##### | 5.9230 | 3.1604E-09 | 1.3111E-08 |
| ENSG00000196850\|PPTC7 | 5943.5391 | 0.6315 | ##### | 5.9214 | 3.1922E-09 | 1.3241E-08 |
| ENSG00000176171\|BNIP3 | 4842.2962 | -0.7348 | ##### | -5.9202 | 3.2162E-09 | 1.3338E-08 |
| ENSG00000198440\|ZNF583 | 519.9106 | -0.8998 | ##### | -5.9199 | 3.2222E-09 | 1.3362E-08 |
| ENSG00000237714\|P4HA2-AS1 | 42.3120 | -1.6953 | ##### | -5.9190 | 3.2390E-09 | 1.3429E-08 |
| ENSG00000162520\|SYNC | 2180.9746 | -0.8182 | ##### | -5.9187 | 3.2457E-09 | 1.3455E-08 |
| ENSG00000283526\|PRRT1B | 1774.9338 | ####### | ##### | -5.9186 | 3.2462E-09 | 1.3456E-08 |
| ENSG00000175220\|ARHGAP1 | 4660.1633 | -0.7439 | ##### | -5.9182 | 3.2546E-09 | 1.3488E-08 |
| ENSG00000120802\|TMPO | 24754.5062 | 0.6120 | ##### | 5.9179 | 3.2613E-09 | 1.3514E-08 |
| ENSG00000169228\|RAB24 | 1287.9771 | 0.8036 | ##### | 5.9173 | 3.2718E-09 | 1.3556E-08 |
| ENSG00000142185\|TRPM2 | 2847.1507 | 0.7413 | ##### | 5.9170 | 3.2786E-09 | 1.3582E-08 |
| ENSG00000151849\|CENPJ | 1841.2756 | 0.7639 | ##### | 5.9157 | 3.3047E-09 | 1.3688E-08 |
| ENSG00000258285\|TESC-AS1 | 158.7185 | -9.7278 | ##### | -5.9156 | 3.3062E-09 | 1.3693E-08 |
| ENSG00000187257\|RSBN1L | 2941.5375 | 0.7451 | ##### | 5.9132 | 3.3557E-09 | 1.3896E-08 |
| ENSG00000050130\|JKAMP | 6712.5873 | 0.7345 | ##### | 5.9127 | 3.3657E-09 | 1.3936E-08 |
| ENSG00000280206 | 506.0225 | 0.9452 | ##### | 5.9117 | 3.3866E-09 | 1.4020E-08 |
| ENSG00000133110\|POSTN | 115.3393 | 1.6028 | ##### | 5.9116 | 3.3881E-09 | 1.4025E-08 |
| ENSG00000104059\|ENTREP2 | 203.1439 | -0.8783 | ##### | -5.9096 | 3.4304E-09 | 1.4197E-08 |
| ENSG00000237493 | 375.5199 | 0.9253 | ##### | 5.9089 | 3.4435E-09 | 1.4250E-08 |
| ENSG00000154162\|CDH12 | 198.6276 | 9.5654 | ##### | 5.9081 | 3.4612E-09 | 1.4321E-08 |
| ENSG00000151692\|RNF144A | 492.1481 | 0.9510 | ##### | 5.9078 | 3.4676E-09 | 1.4345E-08 |
| ENSG00000119820\|YIPF4 | 4691.8767 | 0.7308 | ##### | 5.9063 | 3.4995E-09 | 1.4475E-08 |
| ENSG00000198093\|ZNF649 | 298.7235 | 1.1947 | ##### | 5.9049 | 3.5296E-09 | 1.4598E-08 |
| ENSG00000259321 | 57.0857 | 1.8721 | ##### | 5.9046 | 3.5360E-09 | 1.4622E-08 |
| ENSG00000286078 | 466.6883 | -1.2902 | ##### | -5.9042 | 3.5429E-09 | 1.4649E-08 |
| ENSG00000214194\|SMIM30 | 2080.3597 | -0.7517 | ##### | -5.9038 | 3.5532E-09 | 1.4690E-08 |
| ENSG00000182628\|SKA2 | 3292.3498 | 0.7596 | ##### | 5.9033 | 3.5641E-09 | 1.4732E-08 |
| ENSG00000092330\|TINF2 | 4186.9969 | 0.7393 | ##### | 5.9021 | 3.5891E-09 | 1.4834E-08 |
| ENSG00000151748\|SAV1 | 3137.2946 | 0.7191 | ##### | 5.9013 | 3.6055E-09 | 1.4899E-08 |
| ENSG00000228794\|LINC01128 | 2268.7165 | 0.7673 | ##### | 5.9010 | 3.6129E-09 | 1.4928E-08 |
| ENSG00000260777 | 80.3122 | 8.2842 | ##### | 5.9006 | 3.6216E-09 | 1.4962E-08 |
| ENSG00000154175\|ABI3BP | 45.8430 | -2.0070 | ##### | -5.9001 | 3.6329E-09 | 1.5006E-08 |
| ENSG00000189079\|ARID2 | 4366.3807 | 0.7055 | ##### | 5.8999 | 3.6377E-09 | 1.5024E-08 |
| ENSG00000244005\|NFS1 | 4296.2748 | 0.7270 | ##### | 5.8993 | 3.6494E-09 | 1.5071E-08 |
| ENSG00000131480\|AOC2 | 589.6890 | 0.9209 | ##### | 5.8986 | 3.6650E-09 | 1.5133E-08 |
| ENSG00000136485\|DCAF7 | 11065.4175 | 0.7105 | ##### | 5.8977 | 3.6869E-09 | 1.5221E-08 |
| ENSG00000243244\|STON1 | 313.2736 | 1.0480 | ##### | 5.8975 | 3.6911E-09 | 1.5236E-08 |
| ENSG00000075702\|WDR62 | 5672.1533 | 0.7113 | ##### | 5.8972 | 3.6962E-09 | 1.5255E-08 |
| ENSG00000116489\|CAPZA1 | 25275.5579 | 0.4677 | ##### | 5.8972 | 3.6981E-09 | 1.5261E-08 |
| ENSG00000170860\|LSM3 | 3238.3326 | 0.7464 | ##### | 5.8969 | 3.7049E-09 | 1.5287E-08 |
| ENSG00000158792\|SPATA2L | 1707.6743 | 0.7591 | ##### | 5.8959 | 3.7261E-09 | 1.5372E-08 |
| ENSG00000138081\|FBXO11 | 10310.2157 | 0.7330 | ##### | 5.8957 | 3.7312E-09 | 1.5391E-08 |
| ENSG00000169885\|CALML6 | 49.4739 | 2.7989 | ##### | 5.8950 | 3.7458E-09 | 1.5449E-08 |
| ENSG00000162989\|KCNJ3 | 26.7433 | 6.6921 | ##### | 5.8939 | 3.7718E-09 | 1.5555E-08 |
| ENSG00000166734\|GOLM2 | 7818.5634 | -0.7017 | ##### | -5.8934 | 3.7829E-09 | 1.5598E-08 |
| ENSG00000102119\|EMD | 5836.9131 | 0.7315 | ##### | 5.8932 | 3.7877E-09 | 1.5616E-08 |

| | | | | | | |
|---|---|---|---|---|---|---|
| ENSG00000084110\|HAL | 68.4616 | -1.7150 | ##### | -5.8932 | 3.7889E-09 | 1.5619E-08 |
| ENSG00000070367\|EXOC5 | 10160.7413 | 0.7073 | ##### | 5.8931 | 3.7907E-09 | 1.5624E-08 |
| ENSG00000198818\|SFT2D1 | 3367.0748 | 0.7448 | ##### | 5.8921 | 3.8134E-09 | 1.5715E-08 |
| ENSG00000165271\|NOL6 | 12353.7502 | 0.4469 | ##### | 5.8919 | 3.8179E-09 | 1.5731E-08 |
| ENSG00000149577\|SIDT2 | 1887.2628 | -0.7616 | ##### | -5.8917 | 3.8233E-09 | 1.5752E-08 |
| ENSG00000162836\|ACP6 | 919.2488 | 0.8213 | ##### | 5.8915 | 3.8272E-09 | 1.5765E-08 |
| ENSG00000119689\|DLST | 8218.6541 | 0.5165 | ##### | 5.8908 | 3.8441E-09 | 1.5833E-08 |
| ENSG00000149292\|TTC12 | 507.3248 | -0.8234 | ##### | -5.8906 | 3.8469E-09 | 1.5842E-08 |
| ENSG00000104218\|CSPP1 | 2063.0995 | -0.7657 | ##### | -5.8905 | 3.8492E-09 | 1.5850E-08 |
| ENSG00000256263\|DDX11L8 | 212.2276 | 9.3866 | ##### | 5.8904 | 3.8526E-09 | 1.5861E-08 |
| ENSG00000197016\|ZNF470 | 807.5972 | -0.8214 | ##### | -5.8899 | 3.8651E-09 | 1.5911E-08 |
| ENSG00000105197\|TIMM50 | 9694.4529 | -0.7113 | ##### | -5.8888 | 3.8899E-09 | 1.6010E-08 |
| ENSG00000156463\|SH3RF2 | 1212.2925 | -0.8041 | ##### | -5.8854 | 3.9703E-09 | 1.6339E-08 |
| ENSG00000273619\|RPS21-DT | 176.1668 | 1.1878 | ##### | 5.8846 | 3.9911E-09 | 1.6423E-08 |
| ENSG00000136997\|MYC | 19273.7903 | 0.7052 | ##### | 5.8832 | 4.0241E-09 | 1.6556E-08 |
| ENSG00000204387\|SNHG32 | 3826.0983 | 0.7262 | ##### | 5.8831 | 4.0271E-09 | 1.6566E-08 |
| ENSG00000054118\|THRAP3 | 13602.4244 | 0.6966 | ##### | 5.8828 | 4.0331E-09 | 1.6589E-08 |
| ENSG00000129757\|CDKN1C | 90.7626 | -1.4869 | ##### | -5.8814 | 4.0676E-09 | 1.6728E-08 |
| ENSG00000167775\|CD320 | 5431.5163 | 0.5069 | ##### | 5.8810 | 4.0775E-09 | 1.6766E-08 |
| ENSG00000212907\|MT-ND4L | 53484.2340 | 0.7431 | ##### | 5.8789 | 4.1307E-09 | 1.6983E-08 |
| ENSG00000116288\|PARK7 | 18570.5020 | -0.4667 | ##### | -5.8771 | 4.1761E-09 | 1.7165E-08 |
| ENSG00000111676\|ATN1 | 19123.2593 | 0.6168 | ##### | 5.8771 | 4.1757E-09 | 1.7165E-08 |
| ENSG00000160633\|SAFB | 11499.6993 | 0.6691 | ##### | 5.8770 | 4.1789E-09 | 1.7174E-08 |
| ENSG00000065882\|TBC1D1 | 2976.4875 | 0.7290 | ##### | 5.8758 | 4.2085E-09 | 1.7293E-08 |
| ENSG00000106682\|EIF4H | 29224.3557 | -0.4940 | ##### | -5.8757 | 4.2106E-09 | 1.7299E-08 |
| ENSG00000123983\|ACSL3 | 8799.8282 | -0.6805 | ##### | -5.8749 | 4.2308E-09 | 1.7380E-08 |
| ENSG00000130165\|ELOF1 | 3532.6408 | 0.7371 | ##### | 5.8744 | 4.2450E-09 | 1.7436E-08 |
| ENSG00000112539\|C6orf118 | 167.7391 | 9.3613 | ##### | 5.8720 | 4.3045E-09 | 1.7678E-08 |
| ENSG00000141219\|C17orf80 | 3341.9181 | 0.6884 | ##### | 5.8709 | 4.3339E-09 | 1.7796E-08 |
| ENSG00000158457\|TSPAN33 | 805.1796 | 0.9990 | ##### | 5.8704 | 4.3478E-09 | 1.7851E-08 |
| ENSG00000160191\|PDE9A | 329.8899 | -0.9273 | ##### | -5.8700 | 4.3583E-09 | 1.7892E-08 |
| ENSG00000210144\|MT-TY | 7852.7551 | 0.7425 | ##### | 5.8689 | 4.3869E-09 | 1.8006E-08 |
| ENSG00000118707\|TGIF2 | 2118.3041 | 0.7764 | ##### | 5.8669 | 4.4401E-09 | 1.8222E-08 |
| ENSG00000113645\|WWC1 | 8731.4538 | -0.7181 | ##### | -5.8667 | 4.4455E-09 | 1.8242E-08 |
| ENSG00000135077\|HAVCR2 | 175.7390 | -9.5187 | ##### | -5.8667 | 4.4468E-09 | 1.8245E-08 |
| ENSG00000260097\|SPDYE6 | 65.0140 | 1.5260 | ##### | 5.8662 | 4.4599E-09 | 1.8296E-08 |
| ENSG00000257093\|DENND11 | 3420.3054 | -0.7149 | ##### | -5.8661 | 4.4611E-09 | 1.8298E-08 |
| ENSG00000278931 | 18.1949 | -4.1268 | ##### | -5.8651 | 4.4891E-09 | 1.8411E-08 |
| ENSG00000188610\|FAM72B | 1144.6310 | 0.8032 | ##### | 5.8649 | 4.4929E-09 | 1.8424E-08 |
| ENSG00000101266\|CSNK2A1 | 11745.4277 | 0.4882 | ##### | 5.8635 | 4.5330E-09 | 1.8586E-08 |
| ENSG00000289837 | 146.8091 | 1.1278 | ##### | 5.8629 | 4.5498E-09 | 1.8652E-08 |
| ENSG00000228393\|LINC01004 | 121.6824 | 1.2137 | ##### | 5.8611 | 4.5991E-09 | 1.8851E-08 |
| ENSG00000279631 | 78.0040 | -3.6776 | ##### | -5.8606 | 4.6125E-09 | 1.8904E-08 |
| ENSG00000286061\|RCAN3AS | 16.6387 | -2.7941 | ##### | -5.8587 | 4.6650E-09 | 1.9117E-08 |
| ENSG00000269893\|SNHG8 | 1228.2270 | -0.7571 | ##### | -5.8585 | 4.6720E-09 | 1.9143E-08 |
| ENSG00000273033\|LINC02035 | 344.3399 | -0.8688 | ##### | -5.8584 | 4.6734E-09 | 1.9146E-08 |
| ENSG00000180822\|PSMG4 | 1042.9413 | -0.8007 | ##### | -5.8582 | 4.6781E-09 | 1.9162E-08 |
| ENSG00000168778\|TCTN2 | 1196.1363 | -0.7437 | ##### | -5.8581 | 4.6812E-09 | 1.9172E-08 |
| ENSG00000117650\|NEK2 | 3944.2828 | 0.7225 | ##### | 5.8575 | 4.6992E-09 | 1.9243E-08 |
| ENSG00000244274\|DBNDD2 | 1426.1189 | 0.9055 | ##### | 5.8573 | 4.7047E-09 | 1.9263E-08 |
| ENSG00000116030\|SUMO1 | 11365.3874 | 0.6353 | ##### | 5.8573 | 4.7055E-09 | 1.9264E-08 |
| ENSG00000172732\|MUS81 | 3630.0420 | 0.7354 | ##### | 5.8565 | 4.7276E-09 | 1.9352E-08 |
| ENSG00000229932\|YWHAZP3 | 111.3634 | -1.2143 | ##### | -5.8544 | 4.7885E-09 | 1.9598E-08 |
| ENSG00000165506\|DNAAF2 | 1903.3610 | 0.7912 | ##### | 5.8540 | 4.7975E-09 | 1.9632E-08 |
| ENSG00000161904\|LEMD2 | 4575.7508 | -0.3876 | ##### | -5.8540 | 4.8002E-09 | 1.9641E-08 |
| ENSG00000150722\|PPP1R1C | 95.3944 | -1.4978 | ##### | -5.8535 | 4.8121E-09 | 1.9687E-08 |
| ENSG00000172893\|DHCR7 | 8293.3264 | -0.7336 | ##### | -5.8534 | 4.8175E-09 | 1.9706E-08 |
| ENSG00000271780 | 50.8654 | -1.7391 | ##### | -5.8520 | 4.8557E-09 | 1.9860E-08 |
| ENSG00000008128\|CDK11A | 3802.5977 | 0.5076 | ##### | 5.8506 | 4.8976E-09 | 2.0028E-08 |
| ENSG00000133119\|RFC3 | 4291.9497 | 0.4958 | ##### | 5.8501 | 4.9128E-09 | 2.0088E-08 |
| ENSG00000101306\|MYLK2 | 51.8253 | 2.6464 | ##### | 5.8487 | 4.9542E-09 | 2.0254E-08 |
| ENSG00000204574\|ABCF1 | 13200.3828 | 0.5280 | ##### | 5.8487 | 4.9550E-09 | 2.0255E-08 |
| ENSG00000286430 | 42.9260 | 7.4435 | ##### | 5.8477 | 4.9839E-09 | 2.0370E-08 |

| | | | | | | |
|---|---|---|---|---|---|---|
| ENSG00000135108\|FBXO21 | 5299.1556 | -0.7065 | ##### | -5.8476 | 4.9858E-09 | 2.0375E-08 |
| ENSG00000012048\|BRCA1 | 5883.5175 | 0.7216 | ##### | 5.8472 | 4.9978E-09 | 2.0421E-08 |
| ENSG00000050555\|LAMC3 | 3752.7407 | -0.7188 | ##### | -5.8468 | 5.0116E-09 | 2.0475E-08 |
| ENSG00000214389\|RPS3AP26 | 278.0393 | -1.0073 | ##### | -5.8465 | 5.0210E-09 | 2.0510E-08 |
| ENSG00000139187\|KLRG1 | 222.0086 | 1.2496 | ##### | 5.8464 | 5.0227E-09 | 2.0515E-08 |
| ENSG00000263956\|NBPF11 | 1407.8259 | 0.7949 | ##### | 5.8462 | 5.0284E-09 | 2.0535E-08 |
| ENSG00000065150\|IPO5 | 26765.7050 | 0.6813 | ##### | 5.8458 | 5.0407E-09 | 2.0582E-08 |
| ENSG00000180155\|LYNX1 | 1328.9645 | -0.7490 | ##### | -5.8445 | 5.0817E-09 | 2.0747E-08 |
| ENSG00000121851\|POLR3GL | 872.0369 | -0.8412 | ##### | -5.8444 | 5.0850E-09 | 2.0758E-08 |
| ENSG00000104412\|EMC2 | 2325.3963 | -0.7616 | ##### | -5.8443 | 5.0875E-09 | 2.0765E-08 |
| ENSG00000129991\|TNNI3 | 377.5714 | -0.9310 | ##### | -5.8437 | 5.1061E-09 | 2.0838E-08 |
| ENSG00000103037\|SETD6 | 2368.1812 | 0.7794 | ##### | 5.8431 | 5.1224E-09 | 2.0902E-08 |
| ENSG00000144810\|COL8A1 | 144.5113 | -4.6744 | ##### | -5.8424 | 5.1451E-09 | 2.0992E-08 |
| ENSG00000134278\|SPIRE1 | 9534.7453 | 0.7109 | ##### | 5.8423 | 5.1495E-09 | 2.1006E-08 |
| ENSG00000071539\|TRIP13 | 4317.5278 | -0.6653 | ##### | -5.8415 | 5.1748E-09 | 2.1107E-08 |
| ENSG00000245213\|GALNT7-DT | 134.1444 | -1.5574 | ##### | -5.8408 | 5.1936E-09 | 2.1181E-08 |
| ENSG00000148841\|ITPRIP | 6426.0361 | 0.6880 | ##### | 5.8405 | 5.2043E-09 | 2.1221E-08 |
| ENSG00000168264\|IRF2BP2 | 10668.4609 | -0.5308 | ##### | -5.8402 | 5.2149E-09 | 2.1262E-08 |
| ENSG00000136193\|SCRN1 | 15153.9152 | -0.6744 | ##### | -5.8396 | 5.2322E-09 | 2.1329E-08 |
| ENSG00000100033\|PRODH | 51.9751 | -2.4913 | ##### | -5.8392 | 5.2456E-09 | 2.1381E-08 |
| ENSG00000186648\|CARMIL3 | 93.5151 | 1.4089 | ##### | 5.8378 | 5.2903E-09 | 2.1560E-08 |
| ENSG00000274425 | 1246.3913 | 0.7848 | ##### | 5.8371 | 5.3123E-09 | 2.1647E-08 |
| ENSG00000147010\|SH3KBP1 | 3569.7968 | -0.7127 | ##### | -5.8367 | 5.3257E-09 | 2.1699E-08 |
| ENSG00000204386\|NEU1 | 3158.9315 | -0.7555 | ##### | -5.8362 | 5.3392E-09 | 2.1751E-08 |
| ENSG00000125249\|RAP2A | 2327.4138 | -0.7481 | ##### | -5.8359 | 5.3497E-09 | 2.1791E-08 |
| ENSG00000177291\|GJD4 | 23.9621 | 6.7854 | ##### | 5.8356 | 5.3586E-09 | 2.1824E-08 |
| ENSG00000105607\|GCDH | 1404.8153 | 0.7781 | ##### | 5.8348 | 5.3839E-09 | 2.1924E-08 |
| ENSG00000176014\|TUBB6 | 23526.7454 | 0.6788 | ##### | 5.8347 | 5.3872E-09 | 2.1934E-08 |
| ENSG00000260879 | 122.0971 | -1.3908 | ##### | -5.8343 | 5.4001E-09 | 2.1984E-08 |
| ENSG00000213398\|LCAT | 703.1389 | 0.9225 | ##### | 5.8340 | 5.4106E-09 | 2.2024E-08 |
| ENSG00000067992\|PDK3 | 753.7345 | -0.9050 | ##### | -5.8336 | 5.4258E-09 | 2.2083E-08 |
| ENSG00000107815\|TWNK | 4553.9431 | 0.7377 | ##### | 5.8335 | 5.4293E-09 | 2.2094E-08 |
| ENSG00000232434\|AJM1 | 923.9550 | -0.8506 | ##### | -5.8308 | 5.5168E-09 | 2.2447E-08 |
| ENSG00000176641\|RNF152 | 33.6860 | 3.9091 | ##### | 5.8294 | 5.5615E-09 | 2.2626E-08 |
| ENSG00000262880 | 27.0343 | 7.7957 | ##### | 5.8286 | 5.5901E-09 | 2.2739E-08 |
| ENSG00000196405\|EVL | 1429.1279 | -0.7817 | ##### | -5.8285 | 5.5924E-09 | 2.2745E-08 |
| ENSG00000145040\|UCN2 | 74.3278 | -1.6100 | ##### | -5.8285 | 5.5944E-09 | 2.2750E-08 |
| ENSG00000182389\|CACNB4 | 80.5770 | 2.4938 | ##### | 5.8280 | 5.6099E-09 | 2.2810E-08 |
| ENSG00000156110\|ADK | 3940.5008 | -0.7113 | ##### | -5.8279 | 5.6130E-09 | 2.2820E-08 |
| ENSG00000286155 | 104.5376 | 8.2639 | ##### | 5.8270 | 5.6441E-09 | 2.2943E-08 |
| ENSG00000184967\|NOC4L | 4125.6844 | 0.7050 | ##### | 5.8257 | 5.6889E-09 | 2.3122E-08 |
| ENSG00000102312\|PORCN | 785.4536 | -0.8267 | ##### | -5.8254 | 5.6962E-09 | 2.3149E-08 |
| ENSG00000293142 | 42.2130 | -3.7911 | ##### | -5.8241 | 5.7436E-09 | 2.3338E-08 |
| ENSG00000123965\|PMS2P5 | 193.2439 | -0.9823 | ##### | -5.8237 | 5.7542E-09 | 2.3378E-08 |
| ENSG00000115652\|UXS1 | 5016.9045 | -0.7013 | ##### | -5.8232 | 5.7726E-09 | 2.3449E-08 |
| ENSG00000066027\|PPP2R5A | 3465.0272 | -0.7040 | ##### | -5.8231 | 5.7749E-09 | 2.3455E-08 |
| ENSG00000170322\|NFRKB | 3253.9214 | 0.7023 | ##### | 5.8224 | 5.8007E-09 | 2.3557E-08 |
| ENSG00000235919\|ASH1L-AS1 | 226.6464 | 1.1672 | ##### | 5.8199 | 5.8887E-09 | 2.3911E-08 |
| ENSG00000133124\|IRS4 | 235.6569 | 9.3418 | ##### | 5.8194 | 5.9059E-09 | 2.3978E-08 |
| ENSG00000130312\|MRPL34 | 2386.0069 | 0.4149 | ##### | 5.8176 | 5.9703E-09 | 2.4236E-08 |
| ENSG00000254510 | 157.3422 | 9.3253 | ##### | 5.8165 | 6.0078E-09 | 2.4385E-08 |
| ENSG00000135945\|REV1 | 3495.7572 | 0.6305 | ##### | 5.8152 | 6.0559E-09 | 2.4577E-08 |
| ENSG00000169062\|UPF3A | 3008.6413 | -0.7181 | ##### | -5.8150 | 6.0628E-09 | 2.4601E-08 |
| ENSG00000197013\|ZNF429 | 221.3271 | -7.8845 | ##### | -5.8143 | 6.0886E-09 | 2.4702E-08 |
| ENSG00000044574\|HSPA5 | 67717.1085 | -0.5811 | ##### | -5.8137 | 6.1122E-09 | 2.4795E-08 |
| ENSG00000116266\|STXBP3 | 5390.4346 | 0.7116 | ##### | 5.8133 | 6.1264E-09 | 2.4849E-08 |
| ENSG00000289154 | 162.4330 | 1.7239 | ##### | 5.8125 | 6.1533E-09 | 2.4955E-08 |
| ENSG00000196693\|ZNF33B | 882.1276 | 0.8641 | ##### | 5.8107 | 6.2224E-09 | 2.5232E-08 |
| ENSG00000290976 | 1182.2499 | ####### | ##### | -5.8106 | 6.2237E-09 | 2.5233E-08 |
| ENSG00000140350\|ANP32A | 12680.2691 | 0.6336 | ##### | 5.8102 | 6.2395E-09 | 2.5294E-08 |
| ENSG00000070010\|UFD1 | 7812.4501 | 0.7156 | ##### | 5.8100 | 6.2466E-09 | 2.5319E-08 |
| ENSG00000141198\|TOM1L1 | 4206.0616 | -0.7483 | ##### | -5.8099 | 6.2525E-09 | 2.5340E-08 |
| ENSG00000109920\|FNBP4 | 3409.2834 | 0.7226 | ##### | 5.8096 | 6.2608E-09 | 2.5370E-08 |

| | | | | | | |
|---|---|---|---|---|---|---|
| ENSG00000006712\|PAF1 | 5136.5360 | -0.7138 | ##### | -5.8088 | 6.2910E-09 | 2.5489E-08 |
| ENSG00000071894\|CPSF1 | 12643.4113 | 0.6590 | ##### | 5.8077 | 6.3352E-09 | 2.5665E-08 |
| ENSG00000167525\|PROCA1 | 147.7437 | 1.2306 | ##### | 5.8075 | 6.3407E-09 | 2.5683E-08 |
| ENSG00000100416\|TRMU | 3309.4913 | 0.7136 | ##### | 5.8071 | 6.3575E-09 | 2.5748E-08 |
| ENSG00000183684\|ALYREF | 12430.4533 | 0.6915 | ##### | 5.8070 | 6.3585E-09 | 2.5748E-08 |
| ENSG00000239306\|RBM14 | 8199.7446 | 0.7080 | ##### | 5.8018 | 6.5618E-09 | 2.6568E-08 |
| ENSG00000011347\|SYT7 | 574.1959 | 0.8544 | ##### | 5.8013 | 6.5791E-09 | 2.6635E-08 |
| ENSG00000152763\|DNAI4 | 292.4282 | -1.0274 | ##### | -5.8004 | 6.6176E-09 | 2.6787E-08 |
| ENSG00000102309\|PIN4 | 1589.1156 | 0.7608 | ##### | 5.7989 | 6.6738E-09 | 2.7011E-08 |
| ENSG00000146007\|ZMAT2 | 5213.5919 | 0.7270 | ##### | 5.7987 | 6.6835E-09 | 2.7047E-08 |
| ENSG00000129951\|PLPPR3 | 1757.2808 | -0.7702 | ##### | -5.7985 | 6.6924E-09 | 2.7079E-08 |
| ENSG00000124155\|PIGT | 11689.9484 | 0.6990 | ##### | 5.7967 | 6.7619E-09 | 2.7356E-08 |
| ENSG00000185619\|PCGF3 | 5106.0864 | 0.6699 | ##### | 5.7966 | 6.7659E-09 | 2.7369E-08 |
| ENSG00000232112\|TMA7 | 9238.9294 | 0.7081 | ##### | 5.7965 | 6.7703E-09 | 2.7383E-08 |
| ENSG00000290082 | 363.9264 | -1.0063 | ##### | -5.7964 | 6.7754E-09 | 2.7400E-08 |
| ENSG00000125459\|MSTO1 | 7125.1161 | 0.7122 | ##### | 5.7963 | 6.7789E-09 | 2.7410E-08 |
| ENSG00000184378\|ACTRT3 | 155.3239 | -1.2393 | ##### | -5.7963 | 6.7810E-09 | 2.7415E-08 |
| ENSG00000125885\|MCM8 | 3605.6780 | 0.7199 | ##### | 5.7957 | 6.8049E-09 | 2.7508E-08 |
| ENSG00000272979 | 22.1509 | -3.2916 | ##### | -5.7947 | 6.8456E-09 | 2.7669E-08 |
| ENSG00000273181 | 22.8306 | -6.9457 | ##### | -5.7940 | 6.8732E-09 | 2.7776E-08 |
| ENSG00000180879\|SSR4 | 9758.0897 | 0.7077 | ##### | 5.7935 | 6.8924E-09 | 2.7850E-08 |
| ENSG00000153561\|RMND5A | 5487.3627 | 0.7293 | ##### | 5.7933 | 6.8998E-09 | 2.7877E-08 |
| ENSG00000235927\|NEXN-AS1 | 37.8840 | 2.1095 | ##### | 5.7932 | 6.9052E-09 | 2.7895E-08 |
| ENSG00000090971\|NAT14 | 1972.2726 | -0.7216 | ##### | -5.7931 | 6.9097E-09 | 2.7909E-08 |
| ENSG00000260992\|DOCK9-DT | 33.0868 | -3.1878 | ##### | -5.7926 | 6.9321E-09 | 2.7996E-08 |
| ENSG00000107521\|HPS1 | 2653.3272 | -0.7070 | ##### | -5.7919 | 6.9574E-09 | 2.8094E-08 |
| ENSG00000009830\|POMT2 | 3624.7441 | 0.6920 | ##### | 5.7911 | 6.9933E-09 | 2.8235E-08 |
| ENSG00000184381\|PLA2G6 | 564.2978 | 0.8787 | ##### | 5.7898 | 7.0463E-09 | 2.8445E-08 |
| ENSG00000231028\|AHI1-DT | 50.2853 | -2.0115 | ##### | -5.7885 | 7.1006E-09 | 2.8661E-08 |
| ENSG00000246174\|KCTD21-AS1 | 363.5197 | 1.0443 | ##### | 5.7876 | 7.1414E-09 | 2.8821E-08 |
| ENSG00000187522\|HSPA14 | 2584.0936 | 0.7349 | ##### | 5.7868 | 7.1729E-09 | 2.8945E-08 |
| ENSG00000285966 | 189.2842 | 9.2600 | ##### | 5.7860 | 7.2103E-09 | 2.9091E-08 |
| ENSG00000162419\|GMEB1 | 1357.3198 | 0.7867 | ##### | 5.7858 | 7.2164E-09 | 2.9112E-08 |
| ENSG00000171551\|ECEL1 | 614.8430 | 0.8633 | ##### | 5.7854 | 7.2341E-09 | 2.9180E-08 |
| ENSG00000166664\|CHRFAM7A | 388.3620 | -0.8951 | ##### | -5.7850 | 7.2521E-09 | 2.9248E-08 |
| ENSG00000100911\|PSME2 | 8290.5780 | 0.7060 | ##### | 5.7849 | 7.2567E-09 | 2.9263E-08 |
| ENSG00000137880\|GCHFR | 269.6514 | 1.5381 | ##### | 5.7845 | 7.2732E-09 | 2.9325E-08 |
| ENSG00000286608 | 35.2210 | 7.4343 | ##### | 5.7842 | 7.2873E-09 | 2.9378E-08 |
| ENSG00000204920\|ZNF155 | 593.2037 | -1.0528 | ##### | -5.7823 | 7.3690E-09 | 2.9704E-08 |
| ENSG00000116885\|OSCP1 | 231.3039 | -1.0142 | ##### | -5.7812 | 7.4161E-09 | 2.9890E-08 |
| ENSG00000123416\|TUBA1B | ########## | 0.6771 | ##### | 5.7784 | 7.5410E-09 | 3.0389E-08 |
| ENSG00000290548 | 194.1206 | 1.3939 | ##### | 5.7783 | 7.5465E-09 | 3.0407E-08 |
| ENSG00000228253\|MT-ATP8 | 30456.1047 | 0.7301 | ##### | 5.7771 | 7.5986E-09 | 3.0613E-08 |
| ENSG00000289966 | 55.3444 | -7.1023 | ##### | -5.7762 | 7.6405E-09 | 3.0777E-08 |
| ENSG00000085832\|EPS15 | 3285.7556 | -0.6411 | ##### | -5.7738 | 7.7525E-09 | 3.1224E-08 |
| ENSG00000155256\|ZFYVE27 | 2450.7011 | 0.7397 | ##### | 5.7732 | 7.7792E-09 | 3.1327E-08 |
| ENSG00000171346\|KRT15 | 207.1111 | -1.1670 | ##### | -5.7723 | 7.8192E-09 | 3.1484E-08 |
| ENSG00000171877\|FRMD5 | 105.0406 | 1.5220 | ##### | 5.7723 | 7.8204E-09 | 3.1485E-08 |
| ENSG00000145075 | 621.2526 | -9.4218 | ##### | -5.7716 | 7.8537E-09 | 3.1615E-08 |
| ENSG00000215837\|SDHAP2 | 210.9611 | 4.2369 | ##### | 5.7711 | 7.8744E-09 | 3.1694E-08 |
| ENSG00000169967\|MAP3K2 | 7298.9690 | -0.6983 | ##### | -5.7707 | 7.8950E-09 | 3.1773E-08 |
| ENSG00000198517\|MAFK | 18537.6573 | -0.6656 | ##### | -5.7698 | 7.9378E-09 | 3.1940E-08 |
| ENSG00000111269\|CREBL2 | 4211.1281 | -0.7251 | ##### | -5.7697 | 7.9404E-09 | 3.1947E-08 |
| ENSG00000289562 | 37.5813 | -2.0973 | ##### | -5.7686 | 7.9929E-09 | 3.2153E-08 |
| ENSG00000293057 | 331.0991 | 1.0009 | ##### | 5.7682 | 8.0110E-09 | 3.2218E-08 |
| ENSG00000284707 | 616.6564 | 1.1682 | ##### | 5.7682 | 8.0099E-09 | 3.2218E-08 |
| ENSG00000090263\|MRPS33 | 2527.2169 | -0.7009 | ##### | -5.7679 | 8.0275E-09 | 3.2280E-08 |
| ENSG00000188612\|SUMO2 | 16680.4360 | 0.6958 | ##### | 5.7661 | 8.1108E-09 | 3.2610E-08 |
| ENSG00000084628\|NKAIN1 | 271.0501 | -1.0095 | ##### | -5.7660 | 8.1167E-09 | 3.2630E-08 |
| ENSG00000135486\|HNRNPA1 | 67278.6321 | 0.5652 | ##### | 5.7645 | 8.1924E-09 | 3.2930E-08 |
| ENSG00000126106\|TMEM53 | 785.9112 | -0.8312 | ##### | -5.7639 | 8.2198E-09 | 3.3035E-08 |
| ENSG00000213809\|KLRK1 | 222.5252 | 9.1581 | ##### | 5.7631 | 8.2597E-09 | 3.3191E-08 |
| ENSG00000100532\|CGRRF1 | 935.1949 | 0.7976 | ##### | 5.7611 | 8.3553E-09 | 3.3571E-08 |

| | | | | | | |
|---|---|---|---|---|---|---|
| ENSG00000167608\|TMC4 | 99.2663 | -1.4975 | ##### | -5.7611 | 8.3565E-09 | 3.3571E-08 |
| ENSG00000289009 | 62.0844 | -1.7686 | ##### | -5.7605 | 8.3880E-09 | 3.3693E-08 |
| ENSG00000189014\|SHLD2P3 | 146.7945 | -1.0736 | ##### | -5.7602 | 8.4013E-09 | 3.3742E-08 |
| ENSG00000142166\|IFNAR1 | 5799.8430 | -0.6919 | ##### | -5.7582 | 8.5029E-09 | 3.4146E-08 |
| ENSG00000159069\|FBXW5 | 15778.6964 | -0.5422 | ##### | -5.7581 | 8.5057E-09 | 3.4152E-08 |
| ENSG00000237988\|OR2I1P | 1092.1889 | ####### | ##### | -5.7580 | 8.5098E-09 | 3.4164E-08 |
| ENSG00000260781\|ARHGAP23P1 | 283.9530 | ####### | ##### | -5.7571 | 8.5547E-09 | 3.4339E-08 |
| ENSG00000169629\|RGPD8 | 1189.2253 | -0.7810 | ##### | -5.7571 | 8.5584E-09 | 3.4350E-08 |
| ENSG00000067955\|CBFB | 5960.9138 | 0.4848 | ##### | 5.7569 | 8.5680E-09 | 3.4384E-08 |
| ENSG00000164825\|DEFB1 | 234.4486 | ####### | ##### | -5.7567 | 8.5764E-09 | 3.4413E-08 |
| ENSG00000262903\|CTNS-AS1 | 81.2853 | -1.4559 | ##### | -5.7566 | 8.5814E-09 | 3.4428E-08 |
| ENSG00000091409\|ITGA6 | 10806.6786 | 0.6830 | ##### | 5.7564 | 8.5913E-09 | 3.4463E-08 |
| ENSG00000169714\|CNBP | 29855.5943 | -0.5558 | ##### | -5.7561 | 8.6062E-09 | 3.4519E-08 |
| ENSG00000279399 | 152.6937 | -1.8397 | ##### | -5.7551 | 8.6586E-09 | 3.4724E-08 |
| ENSG00000166401\|SERPINB8 | 2268.9308 | 0.7261 | ##### | 5.7550 | 8.6629E-09 | 3.4736E-08 |
| ENSG00000075785\|RAB7A | 20851.3724 | 0.4594 | ##### | 5.7548 | 8.6753E-09 | 3.4781E-08 |
| ENSG00000183762\|KREMEN1 | 1119.8918 | -0.7500 | ##### | -5.7540 | 8.7135E-09 | 3.4928E-08 |
| ENSG00000160326\|SLC2A6 | 2071.6406 | -0.7147 | ##### | -5.7540 | 8.7142E-09 | 3.4928E-08 |
| ENSG00000187944\|C2orf66 | 98.7226 | 8.5829 | ##### | 5.7524 | 8.8011E-09 | 3.5272E-08 |
| ENSG00000168026\|TTC21A | 187.4119 | 1.1249 | ##### | 5.7508 | 8.8822E-09 | 3.5592E-08 |
| ENSG00000123992\|DNPEP | 6714.2706 | 0.7002 | ##### | 5.7503 | 8.9097E-09 | 3.5698E-08 |
| ENSG00000232618\|HIVEP2-DT | 49.7423 | 3.3476 | ##### | 5.7498 | 8.9325E-09 | 3.5784E-08 |
| ENSG00000288833 | 249.2367 | 1.0046 | ##### | 5.7497 | 8.9385E-09 | 3.5803E-08 |
| ENSG00000223486\|LINC03053 | 191.8919 | 9.0673 | ##### | 5.7486 | 8.9965E-09 | 3.6030E-08 |
| ENSG00000106554\|CHCHD3 | 8407.6649 | 0.6002 | ##### | 5.7467 | 9.0980E-09 | 3.6432E-08 |
| ENSG00000241015\|TPM3P9 | 25.2881 | -2.5880 | ##### | -5.7453 | 9.1756E-09 | 3.6738E-08 |
| ENSG00000288528 | 65.9370 | 1.8582 | ##### | 5.7452 | 9.1792E-09 | 3.6747E-08 |
| ENSG00000188554\|NBR1 | 9569.7560 | -0.6591 | ##### | -5.7443 | 9.2329E-09 | 3.6958E-08 |
| ENSG00000060566\|CREB3L3 | 97.1070 | -8.7582 | ##### | -5.7438 | 9.2576E-09 | 3.7052E-08 |
| ENSG00000089280\|FUS | 41377.5815 | 0.6690 | ##### | 5.7436 | 9.2678E-09 | 3.7087E-08 |
| ENSG00000173068\|BNC2 | 798.2216 | 0.7673 | ##### | 5.7429 | 9.3042E-09 | 3.7228E-08 |
| ENSG00000174567\|GOLT1A | 88.4070 | -1.8610 | ##### | -5.7424 | 9.3331E-09 | 3.7339E-08 |
| ENSG00000284874 | 1124.4024 | -3.0944 | ##### | -5.7422 | 9.3437E-09 | 3.7376E-08 |
| ENSG00000107771\|CCSER2 | 2270.0122 | -0.7319 | ##### | -5.7404 | 9.4470E-09 | 3.7784E-08 |
| ENSG00000260011 | 18.8982 | 6.9403 | ##### | 5.7400 | 9.4698E-09 | 3.7870E-08 |
| ENSG00000139537\|CCDC65 | 113.2359 | 1.2217 | ##### | 5.7385 | 9.5510E-09 | 3.8190E-08 |
| ENSG00000114993\|RTKN | 5199.1528 | 0.5986 | ##### | 5.7373 | 9.6188E-09 | 3.8456E-08 |
| ENSG00000112234\|FBXL4 | 1523.2846 | -0.7908 | ##### | -5.7368 | 9.6493E-09 | 3.8573E-08 |
| ENSG00000177479\|ARIH2 | 8975.8751 | 0.6942 | ##### | 5.7367 | 9.6514E-09 | 3.8576E-08 |
| ENSG00000184009\|ACTG1 | ########## | -0.7012 | ##### | -5.7366 | 9.6590E-09 | 3.8601E-08 |
| ENSG00000104147\|OIP5 | 2608.8625 | 0.7411 | ##### | 5.7346 | 9.7750E-09 | 3.9059E-08 |
| ENSG00000142149\|HUNK | 379.5279 | 0.9581 | ##### | 5.7326 | 9.8887E-09 | 3.9508E-08 |
| ENSG00000179832\|MROH1 | 2449.9722 | -0.7149 | ##### | -5.7326 | 9.8915E-09 | 3.9515E-08 |
| ENSG00000111667\|USP5 | 23755.7138 | 0.6253 | ##### | 5.7325 | 9.8935E-09 | 3.9517E-08 |
| ENSG00000224023\|EDRF1-DT | 67.1661 | -1.8055 | ##### | -5.7310 | 9.9833E-09 | 3.9870E-08 |
| ENSG00000067248\|DHX29 | 7113.4677 | -0.6987 | ##### | -5.7309 | 9.9924E-09 | 3.9902E-08 |
| ENSG00000075290\|WNT8B | 53.8845 | 7.3265 | ##### | 5.7307 | 9.9989E-09 | 3.9922E-08 |
| ENSG00000178386\|ZNF223 | 341.7267 | -5.4780 | ##### | -5.7303 | 1.0028E-08 | 4.0031E-08 |
| ENSG00000174243\|DDX23 | 18720.6811 | 0.6371 | ##### | 5.7302 | 1.0029E-08 | 4.0032E-08 |
| ENSG00000100897\|DCAF11 | 2551.6659 | 0.7255 | ##### | 5.7293 | 1.0083E-08 | 4.0240E-08 |
| ENSG00000258315\|C17orf49 | 2936.0904 | 0.9417 | ##### | 5.7258 | 1.0295E-08 | 4.1082E-08 |
| ENSG00000183840\|GPR39 | 2556.3338 | -0.7151 | ##### | -5.7258 | 1.0298E-08 | 4.1087E-08 |
| ENSG00000244313 | 1635.8291 | 0.7555 | ##### | 5.7249 | 1.0350E-08 | 4.1289E-08 |
| ENSG00000164794\|KCNV1 | 295.3739 | 1.0381 | ##### | 5.7248 | 1.0354E-08 | 4.1303E-08 |
| ENSG00000100139\|MICALL1 | 4177.0098 | 0.7160 | ##### | 5.7248 | 1.0356E-08 | 4.1303E-08 |
| ENSG00000105641\|SLC5A5 | 62.1602 | -2.1586 | ##### | -5.7247 | 1.0362E-08 | 4.1320E-08 |
| ENSG00000163539\|CLASP2 | 5518.6733 | 0.7059 | ##### | 5.7244 | 1.0382E-08 | 4.1397E-08 |
| ENSG00000110455\|ACCS | 411.5239 | -0.8571 | ##### | -5.7242 | 1.0393E-08 | 4.1435E-08 |
| ENSG00000289504 | 28.4019 | -2.5065 | ##### | -5.7226 | 1.0493E-08 | 4.1826E-08 |
| ENSG00000007866\|TEAD3 | 891.4203 | -0.7375 | ##### | -5.7214 | 1.0566E-08 | 4.2115E-08 |
| ENSG00000139613\|SMARCC2 | 13470.9702 | 0.5122 | ##### | 5.7210 | 1.0589E-08 | 4.2199E-08 |
| ENSG00000115694\|STK25 | 11016.4612 | 0.6910 | ##### | 5.7206 | 1.0615E-08 | 4.2298E-08 |
| ENSG00000271857 | 30.9601 | 6.8928 | ##### | 5.7202 | 1.0641E-08 | 4.2397E-08 |

| | | | | | | |
|---|---|---|---|---|---|---|
| ENSG00000272078 | 350.4889 | 1.0434 | ##### | 5.7164 | 1.0878E-08 | 4.3335E-08 |
| ENSG00000121060\|TRIM25 | 8143.0862 | -0.6846 | ##### | -5.7163 | 1.0884E-08 | 4.3346E-08 |
| ENSG00000136521\|NDUFB5 | 4092.4660 | -0.6937 | ##### | -5.7164 | 1.0883E-08 | 4.3346E-08 |
| ENSG00000236778\|INTS6-AS1 | 72.9364 | -1.6986 | ##### | -5.7162 | 1.0893E-08 | 4.3376E-08 |
| ENSG00000159593\|NAE1 | 6675.5140 | -0.5659 | ##### | -5.7162 | 1.0895E-08 | 4.3377E-08 |
| ENSG00000184441 | 521.8307 | 0.9201 | ##### | 5.7156 | 1.0930E-08 | 4.3512E-08 |
| ENSG00000160439\|RDH13 | 1815.3744 | -0.8158 | ##### | -5.7154 | 1.0944E-08 | 4.3563E-08 |
| ENSG00000237669 | 89.4679 | 1.2782 | ##### | 5.7151 | 1.0963E-08 | 4.3629E-08 |
| ENSG00000164841\|TMEM74 | 101.8877 | 1.7230 | ##### | 5.7145 | 1.1006E-08 | 4.3796E-08 |
| ENSG00000228078\|HLA-U | 134.0439 | 6.2254 | ##### | 5.7140 | 1.1036E-08 | 4.3910E-08 |
| ENSG00000228502\|EEF1A1P11 | 354.3215 | 0.8868 | ##### | 5.7119 | 1.1170E-08 | 4.4439E-08 |
| ENSG00000233436\|BTBD18 | 41.5129 | 7.3023 | ##### | 5.7115 | 1.1196E-08 | 4.4535E-08 |
| ENSG00000275236 | 18.8381 | -7.6493 | ##### | -5.7114 | 1.1207E-08 | 4.4571E-08 |
| ENSG00000137440\|FGFBP1 | 155.8289 | -4.5689 | ##### | -5.7113 | 1.1209E-08 | 4.4575E-08 |
| ENSG00000004660\|CAMKK1 | 582.1224 | 0.8186 | ##### | 5.7109 | 1.1236E-08 | 4.4678E-08 |
| ENSG00000246627\|CACNA1C-AS1 | 192.6671 | 9.1200 | ##### | 5.7080 | 1.1432E-08 | 4.5450E-08 |
| ENSG00000106263\|EIF3B | 49051.5968 | 0.3034 | ##### | 5.7076 | 1.1459E-08 | 4.5552E-08 |
| ENSG00000261071 | 38.9560 | -1.9888 | ##### | -5.7067 | 1.1522E-08 | 4.5794E-08 |
| ENSG00000182247\|UBE2E2 | 1671.7365 | -0.7593 | ##### | -5.7050 | 1.1631E-08 | 4.6222E-08 |
| ENSG00000077232\|DNAJC10 | 21490.0472 | -0.6835 | ##### | -5.7047 | 1.1653E-08 | 4.6302E-08 |
| ENSG00000152782\|PANK1 | 890.5442 | -0.7876 | ##### | -5.7036 | 1.1727E-08 | 4.6591E-08 |
| ENSG00000236751\|LINC01186 | 76.4784 | -1.5902 | ##### | -5.7036 | 1.1733E-08 | 4.6609E-08 |
| ENSG00000258525 | 93.4683 | -1.9262 | ##### | -5.7035 | 1.1741E-08 | 4.6633E-08 |
| ENSG00000127328\|RAB3IP | 1726.4673 | 0.7209 | ##### | 5.7027 | 1.1793E-08 | 4.6836E-08 |
| ENSG00000187688\|TRPV2 | 47.4757 | 2.7315 | ##### | 5.6997 | 1.1999E-08 | 4.7647E-08 |
| ENSG00000130177\|CDC16 | 4557.6727 | -0.6997 | ##### | -5.6996 | 1.2006E-08 | 4.7668E-08 |
| ENSG00000197006\|METTL9 | 6606.3620 | -0.4824 | ##### | -5.6989 | 1.2056E-08 | 4.7860E-08 |
| ENSG00000074370\|ATP2A3 | 684.3146 | 0.8316 | ##### | 5.6985 | 1.2088E-08 | 4.7981E-08 |
| ENSG00000198860\|TSEN15 | 2161.1042 | -0.7344 | ##### | -5.6984 | 1.2093E-08 | 4.7996E-08 |
| ENSG00000182934\|SRPRA | 11390.5173 | -0.5559 | ##### | -5.6977 | 1.2142E-08 | 4.8183E-08 |
| ENSG00000141391\|PRELID3A | 455.3807 | 0.8425 | ##### | 5.6974 | 1.2164E-08 | 4.8262E-08 |
| ENSG00000125898\|FAM110A | 634.7793 | 0.8407 | ##### | 5.6974 | 1.2168E-08 | 4.8272E-08 |
| ENSG00000263563\|UBBP4 | 246.5690 | 1.4665 | ##### | 5.6921 | 1.2546E-08 | 4.9766E-08 |
| ENSG00000118156\|ZNF541 | 204.9888 | ####### | ##### | -5.6917 | 1.2579E-08 | 4.9888E-08 |
| ENSG00000091732\|ZC3HC1 | 4616.8162 | 0.6959 | ##### | 5.6913 | 1.2610E-08 | 5.0008E-08 |
| ENSG00000217930\|PAM16 | 2660.7532 | 0.7690 | ##### | 5.6912 | 1.2615E-08 | 5.0019E-08 |
| ENSG00000086666\|ZFAND6 | 3641.9562 | -0.7052 | ##### | -5.6911 | 1.2620E-08 | 5.0034E-08 |
| ENSG00000261126\|RBFADN | 39.8108 | 2.2211 | ##### | 5.6904 | 1.2672E-08 | 5.0231E-08 |
| ENSG00000214432\|VPS33B-DT | 100.3598 | -8.8892 | ##### | -5.6891 | 1.2772E-08 | 5.0623E-08 |
| ENSG00000290846 | 1356.9502 | 0.7388 | ##### | 5.6877 | 1.2876E-08 | 5.1029E-08 |
| ENSG00000182379\|NXPH4 | 194.8508 | 1.1097 | ##### | 5.6874 | 1.2900E-08 | 5.1117E-08 |
| ENSG00000289006 | 117.6243 | -1.0666 | ##### | -5.6872 | 1.2914E-08 | 5.1164E-08 |
| ENSG00000005194\|CIAPIN1 | 6608.8307 | -0.6841 | ##### | -5.6866 | 1.2960E-08 | 5.1342E-08 |
| ENSG00000273472 | 60.8906 | -1.5039 | ##### | -5.6862 | 1.2991E-08 | 5.1455E-08 |
| ENSG00000133805\|AMPD3 | 202.7727 | -1.3192 | ##### | -5.6852 | 1.3068E-08 | 5.1753E-08 |
| ENSG00000117899\|MESD | 3777.3260 | -0.6337 | ##### | -5.6849 | 1.3092E-08 | 5.1843E-08 |
| ENSG00000286828 | 80.7090 | 8.4026 | ##### | 5.6847 | 1.3108E-08 | 5.1898E-08 |
| ENSG00000196547\|MAN2A2 | 4703.3857 | -0.6890 | ##### | -5.6838 | 1.3172E-08 | 5.2144E-08 |
| ENSG00000122386\|ZNF205 | 1542.7298 | 0.7348 | ##### | 5.6837 | 1.3180E-08 | 5.2164E-08 |
| ENSG00000165775\|FUNDC2 | 3133.4628 | 0.5123 | ##### | 5.6837 | 1.3179E-08 | 5.2164E-08 |
| ENSG00000146243\|IRAK1BP1 | 383.2475 | 0.9275 | ##### | 5.6836 | 1.3189E-08 | 5.2192E-08 |
| ENSG00000276966\|H4C5 | 33.9393 | 2.6918 | ##### | 5.6825 | 1.3272E-08 | 5.2512E-08 |
| ENSG00000236529 | 32.2348 | -2.2965 | ##### | -5.6823 | 1.3293E-08 | 5.2587E-08 |
| ENSG00000288920\|RPSA2 | 1278.2563 | 0.7234 | ##### | 5.6820 | 1.3315E-08 | 5.2668E-08 |
| ENSG00000172014\|ANKRD20A4P | 103.7291 | -8.9885 | ##### | -5.6810 | 1.3391E-08 | 5.2964E-08 |
| ENSG00000286846 | 33.1143 | 7.8127 | ##### | 5.6799 | 1.3476E-08 | 5.3291E-08 |
| ENSG00000178537\|SLC25A20 | 1433.0189 | -0.7304 | ##### | -5.6789 | 1.3558E-08 | 5.3610E-08 |
| ENSG00000119421\|NDUFA8 | 4064.0669 | -0.7095 | ##### | -5.6764 | 1.3758E-08 | 5.4391E-08 |
| ENSG00000279713 | 71.0811 | 1.4471 | ##### | 5.6755 | 1.3826E-08 | 5.4653E-08 |
| ENSG00000117411\|B4GALT2 | 6989.3786 | -0.6944 | ##### | -5.6752 | 1.3851E-08 | 5.4748E-08 |
| ENSG00000149187\|CELF1 | 8880.4258 | 0.6638 | ##### | 5.6749 | 1.3876E-08 | 5.4830E-08 |
| ENSG00000165675\|ENOX2 | 1926.1872 | 0.7701 | ##### | 5.6749 | 1.3875E-08 | 5.4830E-08 |
| ENSG00000178184\|PARD6G | 702.2680 | 0.8042 | ##### | 5.6731 | 1.4021E-08 | 5.5389E-08 |

| | | | | | | |
|---|---|---|---|---|---|---|
| ENSG00000124260\|MAGEA10 | 3494.4681 | ####### | ##### | -5.6731 | 1.4020E-08 | 5.5389E-08 |
| ENSG00000227141 | 67.7341 | -8.9595 | ##### | -5.6720 | 1.4113E-08 | 5.5743E-08 |
| ENSG00000197977\|ELOVL2 | 560.2364 | -0.8377 | ##### | -5.6718 | 1.4128E-08 | 5.5795E-08 |
| ENSG00000226944\|RNF207-AS1 | 17.8712 | -4.5851 | ##### | -5.6710 | 1.4198E-08 | 5.6066E-08 |
| ENSG00000149089\|APIP | 1065.0393 | -0.5314 | ##### | -5.6704 | 1.4246E-08 | 5.6246E-08 |
| ENSG00000114491\|UMPS | 5927.3286 | 0.5620 | ##### | 5.6700 | 1.4281E-08 | 5.6377E-08 |
| ENSG00000142512\|SIGLEC10 | 50.4242 | -2.0914 | ##### | -5.6695 | 1.4325E-08 | 5.6544E-08 |
| ENSG00000104237\|RP1 | 52.1589 | 4.6806 | ##### | 5.6694 | 1.4329E-08 | 5.6554E-08 |
| ENSG00000213147\|RPL23AP60 | 25.5168 | 2.1073 | ##### | 5.6682 | 1.4429E-08 | 5.6942E-08 |
| ENSG00000274825 | 34.0112 | 2.7528 | ##### | 5.6670 | 1.4534E-08 | 5.7346E-08 |
| ENSG00000081791\|DELE1 | 4860.7835 | -0.7196 | ##### | -5.6667 | 1.4556E-08 | 5.7425E-08 |
| ENSG00000161551\|ZNF577 | 176.7871 | 1.0959 | ##### | 5.6665 | 1.4572E-08 | 5.7481E-08 |
| ENSG00000140563\|MCTP2 | 55.4459 | -2.0510 | ##### | -5.6663 | 1.4589E-08 | 5.7539E-08 |
| ENSG00000235237 | 35.2120 | -2.1974 | ##### | -5.6661 | 1.4607E-08 | 5.7603E-08 |
| ENSG00000261089\|CDC37P2 | 32.0841 | -3.6108 | ##### | -5.6656 | 1.4652E-08 | 5.7773E-08 |
| ENSG00000139714\|MORN3 | 73.2322 | 1.4859 | ##### | 5.6654 | 1.4672E-08 | 5.7847E-08 |
| ENSG00000183765\|CHEK2 | 1877.3595 | 0.7562 | ##### | 5.6653 | 1.4679E-08 | 5.7866E-08 |
| ENSG00000178464\|RPL10P16 | 3220.8507 | 0.7258 | ##### | 5.6651 | 1.4696E-08 | 5.7924E-08 |
| ENSG00000169093\|ASMTL | 2567.7165 | 0.7175 | ##### | 5.6629 | 1.4886E-08 | 5.8667E-08 |
| ENSG00000286149 | 86.5130 | 7.9041 | ##### | 5.6621 | 1.4957E-08 | 5.8937E-08 |
| ENSG00000290767 | 57.6161 | 8.4984 | ##### | 5.6614 | 1.5010E-08 | 5.9141E-08 |
| ENSG00000106546\|AHR | 12282.0462 | -0.6698 | ##### | -5.6608 | 1.5070E-08 | 5.9366E-08 |
| ENSG00000072864\|NDE1 | 3209.5907 | 0.6631 | ##### | 5.6599 | 1.5148E-08 | 5.9667E-08 |
| ENSG00000129038\|LOXL1 | 266.7898 | 1.0612 | ##### | 5.6597 | 1.5163E-08 | 5.9718E-08 |
| ENSG00000221887\|HMSD | 146.2870 | 3.6660 | ##### | 5.6591 | 1.5220E-08 | 5.9933E-08 |
| ENSG00000198626\|RYR2 | 42.1424 | 2.8161 | ##### | 5.6584 | 1.5275E-08 | 6.0143E-08 |
| ENSG00000114638\|UPK1B | 401.0577 | -7.1405 | ##### | -5.6584 | 1.5280E-08 | 6.0153E-08 |
| ENSG00000123444\|KBTBD4 | 2312.8526 | 0.7285 | ##### | 5.6575 | 1.5356E-08 | 6.0448E-08 |
| ENSG00000163874\|ZC3H12A | 1111.3567 | -0.7661 | ##### | -5.6559 | 1.5501E-08 | 6.1008E-08 |
| ENSG00000286140\|DERPC | 3831.6255 | 0.8280 | ##### | 5.6555 | 1.5536E-08 | 6.1137E-08 |
| ENSG00000165406\|MARCHF8 | 1522.1341 | -0.7068 | ##### | -5.6543 | 1.5648E-08 | 6.1573E-08 |
| ENSG00000184719\|RNLS | 1018.3118 | ####### | ##### | -5.6531 | 1.5757E-08 | 6.1992E-08 |
| ENSG00000119523\|ALG2 | 3458.3468 | -0.5492 | ##### | -5.6528 | 1.5785E-08 | 6.2093E-08 |
| ENSG00000084754\|HADHA | 13871.5167 | -0.3547 | ##### | -5.6527 | 1.5792E-08 | 6.2114E-08 |
| ENSG00000112039\|FANCE | 1041.1729 | 0.7781 | ##### | 5.6522 | 1.5837E-08 | 6.2281E-08 |
| ENSG00000187010\|RHD | 44.9450 | 7.1970 | ##### | 5.6516 | 1.5895E-08 | 6.2493E-08 |
| ENSG00000291122\|CASTOR3P | 843.7225 | -0.8893 | ##### | -5.6516 | 1.5894E-08 | 6.2493E-08 |
| ENSG00000253326 | 50.6383 | 6.9434 | ##### | 5.6495 | 1.6091E-08 | 6.3256E-08 |
| ENSG00000163939\|PBRM1 | 5238.1151 | 0.6768 | ##### | 5.6483 | 1.6205E-08 | 6.3696E-08 |
| ENSG00000124370\|MCEE | 308.7967 | -0.9282 | ##### | -5.6480 | 1.6230E-08 | 6.3782E-08 |
| ENSG00000237260\|PPIAP67 | 46.5007 | 7.2048 | ##### | 5.6480 | 1.6231E-08 | 6.3782E-08 |
| ENSG00000142606\|MMEL1 | 107.1895 | 1.4521 | ##### | 5.6478 | 1.6247E-08 | 6.3835E-08 |
| ENSG00000272516 | 30.0190 | 6.2036 | ##### | 5.6475 | 1.6280E-08 | 6.3957E-08 |
| ENSG00000132205\|EMILIN2 | 484.6992 | -0.8744 | ##### | -5.6451 | 1.6507E-08 | 6.4840E-08 |
| ENSG00000151338\|MIPOL1 | 902.4624 | 0.7782 | ##### | 5.6451 | 1.6512E-08 | 6.4852E-08 |
| ENSG00000266473\|HELZ-AS1 | 18.0435 | -7.1018 | ##### | -5.6438 | 1.6633E-08 | 6.5318E-08 |
| ENSG00000198856\|OSTC | 6080.2041 | -0.6845 | ##### | -5.6437 | 1.6639E-08 | 6.5333E-08 |
| ENSG00000236439\|RPS27P8 | 85.0218 | 1.3507 | ##### | 5.6420 | 1.6806E-08 | 6.5980E-08 |
| ENSG00000256576\|LINC02361 | 108.0453 | 1.3234 | ##### | 5.6411 | 1.6893E-08 | 6.6312E-08 |
| ENSG00000174607\|UGT8 | 1571.3711 | 0.7396 | ##### | 5.6410 | 1.6904E-08 | 6.6347E-08 |
| ENSG00000272183 | 38.6988 | -1.7969 | ##### | -5.6408 | 1.6928E-08 | 6.6434E-08 |
| ENSG00000198156\|NPIPB6 | 89.1133 | -1.7264 | ##### | -5.6397 | 1.7036E-08 | 6.6849E-08 |
| ENSG00000244558\|KCNK15-AS1 | 42.6608 | 1.7579 | ##### | 5.6396 | 1.7044E-08 | 6.6872E-08 |
| ENSG00000139514\|SLC7A1 | 10620.9340 | 0.6874 | ##### | 5.6395 | 1.7050E-08 | 6.6886E-08 |
| ENSG00000230438\|SERPINB9P1 | 125.1638 | -3.7777 | ##### | -5.6386 | 1.7143E-08 | 6.7240E-08 |
| ENSG00000169750\|RAC3 | 853.5340 | -0.8190 | ##### | -5.6385 | 1.7155E-08 | 6.7280E-08 |
| ENSG00000228109\|MELTF-AS1 | 180.2218 | -1.5011 | ##### | -5.6384 | 1.7160E-08 | 6.7291E-08 |
| ENSG00000049247\|UTS2 | 61.0428 | 1.6546 | ##### | 5.6376 | 1.7245E-08 | 6.7613E-08 |
| ENSG00000203804\|ADAMTSL4-AS1 | 72.4038 | -1.3463 | ##### | -5.6371 | 1.7297E-08 | 6.7808E-08 |
| ENSG00000240849\|PEDS1 | 3972.0271 | 0.6471 | ##### | 5.6356 | 1.7443E-08 | 6.8373E-08 |
| ENSG00000224520\|KRT8P45 | 32.2362 | -2.1346 | ##### | -5.6356 | 1.7446E-08 | 6.8375E-08 |
| ENSG00000134686\|PHC2 | 6439.3784 | -0.6302 | ##### | -5.6354 | 1.7466E-08 | 6.8447E-08 |
| ENSG00000132773\|TOE1 | 2527.7328 | 0.7070 | ##### | 5.6337 | 1.7637E-08 | 6.9104E-08 |

| | | | | | | |
|---|---|---|---|---|---|---|
| ENSG00000172915\|NBEA | 1140.2242 | -0.7837 | ##### | -5.6337 | 1.7640E-08 | 6.9111E-08 |
| ENSG00000108021\|TASOR2 | 12389.2706 | 0.6939 | ##### | 5.6317 | 1.7848E-08 | 6.9916E-08 |
| ENSG00000159147\|DONSON | 2759.9948 | -0.7259 | ##### | -5.6313 | 1.7885E-08 | 7.0052E-08 |
| ENSG00000155744\|HYCC2 | 1520.7381 | 0.7430 | ##### | 5.6311 | 1.7902E-08 | 7.0108E-08 |
| ENSG00000070601\|FRMPD1 | 45.2160 | 2.7802 | ##### | 5.6301 | 1.8009E-08 | 7.0519E-08 |
| ENSG00000162873\|KLHDC8A | 59.4790 | 1.8255 | ##### | 5.6299 | 1.8034E-08 | 7.0606E-08 |
| ENSG00000082146\|STRADB | 1511.4163 | -0.7258 | ##### | -5.6294 | 1.8083E-08 | 7.0789E-08 |
| ENSG00000176020\|AMIGO3 | 305.6541 | -1.2929 | ##### | -5.6293 | 1.8095E-08 | 7.0827E-08 |
| ENSG00000147894\|C9orf72 | 455.7288 | -0.9621 | ##### | -5.6288 | 1.8149E-08 | 7.1029E-08 |
| ENSG00000135070\|ISCA1 | 4056.0387 | -0.7048 | ##### | -5.6265 | 1.8391E-08 | 7.1966E-08 |
| ENSG00000186704\|DTX2P1 | 97.0679 | -1.3762 | ##### | -5.6253 | 1.8515E-08 | 7.2441E-08 |
| ENSG00000204178\|MACO1 | 994.2163 | -0.7578 | ##### | -5.6245 | 1.8607E-08 | 7.2794E-08 |
| ENSG00000160539\|PLPP7 | 113.5958 | -6.5331 | ##### | -5.6233 | 1.8729E-08 | 7.3262E-08 |
| ENSG00000152049\|KCNE4 | 80.9953 | -3.9037 | ##### | -5.6218 | 1.8894E-08 | 7.3897E-08 |
| ENSG00000139372\|TDG | 4155.7929 | -0.5998 | ##### | -5.6217 | 1.8910E-08 | 7.3949E-08 |
| ENSG00000167264\|DUS2 | 1271.9340 | -0.7359 | ##### | -5.6216 | 1.8921E-08 | 7.3984E-08 |
| ENSG00000167419\|LPO | 45.9676 | 7.3829 | ##### | 5.6213 | 1.8951E-08 | 7.4089E-08 |
| ENSG00000145348\|TBCK | 947.2508 | -0.7732 | ##### | -5.6211 | 1.8973E-08 | 7.4165E-08 |
| ENSG00000182902\|SLC25A18 | 55.5605 | -1.7763 | ##### | -5.6210 | 1.8990E-08 | 7.4224E-08 |
| ENSG00000171943\|SRGAP2C | 2141.7676 | 0.7190 | ##### | 5.6207 | 1.9022E-08 | 7.4340E-08 |
| ENSG00000071205\|ARHGAP10 | 1029.1156 | -0.6624 | ##### | -5.6198 | 1.9113E-08 | 7.4682E-08 |
| ENSG00000132744\|ACY3 | 30.4224 | -2.4827 | ##### | -5.6192 | 1.9188E-08 | 7.4968E-08 |
| ENSG00000272217 | 32.5414 | 7.8078 | ##### | 5.6186 | 1.9255E-08 | 7.5221E-08 |
| ENSG00000162437\|RAVER2 | 2176.1106 | -0.6942 | ##### | -5.6180 | 1.9324E-08 | 7.5477E-08 |
| ENSG00000184515\|BEX5 | 72.6318 | 1.6314 | ##### | 5.6171 | 1.9423E-08 | 7.5855E-08 |
| ENSG00000135045\|C9orf40 | 1182.2271 | -0.7450 | ##### | -5.6161 | 1.9527E-08 | 7.6253E-08 |
| ENSG00000260912 | 66.5213 | 1.5521 | ##### | 5.6156 | 1.9589E-08 | 7.6485E-08 |
| ENSG00000115993\|TRAK2 | 5200.6942 | 0.6825 | ##### | 5.6152 | 1.9630E-08 | 7.6634E-08 |
| ENSG00000176105\|YES1 | 12718.9133 | 0.6670 | ##### | 5.6135 | 1.9829E-08 | 7.7401E-08 |
| ENSG00000138738\|PRDM5 | 565.6996 | -0.8075 | ##### | -5.6132 | 1.9861E-08 | 7.7517E-08 |
| ENSG00000286033 | 27.6856 | 1.7187 | ##### | 5.6122 | 1.9981E-08 | 7.7970E-08 |
| ENSG00000137959\|IFI44L | 876.7180 | ####### | ##### | -5.6122 | 1.9983E-08 | 7.7970E-08 |
| ENSG00000242396 | 67.8795 | -1.5561 | ##### | -5.6114 | 2.0072E-08 | 7.8287E-08 |
| ENSG00000165671\|NSD1 | 7324.3420 | 0.6792 | ##### | 5.6114 | 2.0069E-08 | 7.8287E-08 |
| ENSG00000197081\|IGF2R | 8945.1997 | -0.6883 | ##### | -5.6114 | 2.0070E-08 | 7.8287E-08 |
| ENSG00000288999 | 67.5536 | -1.5733 | ##### | -5.6102 | 2.0204E-08 | 7.8792E-08 |
| ENSG00000176928\|GCNT4 | 98.0069 | -1.9525 | ##### | -5.6099 | 2.0247E-08 | 7.8949E-08 |
| ENSG00000164118\|CEP44 | 1052.5747 | -0.7895 | ##### | -5.6089 | 2.0367E-08 | 7.9406E-08 |
| ENSG00000101365\|IDH3B | 7318.2358 | 0.6636 | ##### | 5.6064 | 2.0658E-08 | 8.0531E-08 |
| ENSG00000137815\|RTF1 | 5560.8913 | 0.6701 | ##### | 5.6056 | 2.0754E-08 | 8.0896E-08 |
| ENSG00000253616\|CHILL1 | 47.0964 | -4.8189 | ##### | -5.6038 | 2.0973E-08 | 8.1740E-08 |
| ENSG00000149782\|PLCB3 | 6075.4591 | 0.6692 | ##### | 5.6037 | 2.0985E-08 | 8.1774E-08 |
| ENSG00000042088\|TDP1 | 1372.2006 | 0.7457 | ##### | 5.6035 | 2.1013E-08 | 8.1871E-08 |
| ENSG00000232977\|LINC00327 | 57.7627 | 2.3147 | ##### | 5.6029 | 2.1083E-08 | 8.2136E-08 |
| ENSG00000270765\|GAS2L2 | 37.8086 | -5.4092 | ##### | -5.6028 | 2.1097E-08 | 8.2177E-08 |
| ENSG00000133773\|CCDC59 | 3082.4626 | 0.7187 | ##### | 5.6016 | 2.1239E-08 | 8.2722E-08 |
| ENSG00000100612\|DHRS7 | 3842.6364 | -0.6876 | ##### | -5.6014 | 2.1262E-08 | 8.2801E-08 |
| ENSG00000185532\|PRKG1 | 70.2560 | 1.8023 | ##### | 5.6006 | 2.1363E-08 | 8.3180E-08 |
| ENSG00000105186\|ANKRD27 | 3608.9124 | -0.6883 | ##### | -5.5999 | 2.1443E-08 | 8.3483E-08 |
| ENSG00000108785\|HSD17B1P1 | 62.8100 | -1.3851 | ##### | -5.5997 | 2.1474E-08 | 8.3594E-08 |
| ENSG00000104043\|ATP8B4 | 141.5562 | 1.0319 | ##### | 5.5991 | 2.1552E-08 | 8.3884E-08 |
| ENSG00000242516\|LINC00960 | 293.3291 | 5.9132 | ##### | 5.5990 | 2.1555E-08 | 8.3887E-08 |
| ENSG00000014216\|CAPN1 | 10239.9272 | -0.5081 | ##### | -5.5989 | 2.1570E-08 | 8.3935E-08 |
| ENSG00000068796\|KIF2A | 8065.1139 | 0.7585 | ##### | 5.5987 | 2.1593E-08 | 8.4011E-08 |
| ENSG00000163466\|ARPC2 | 28126.7765 | 0.6808 | ##### | 5.5979 | 2.1693E-08 | 8.4390E-08 |
| ENSG00000234263\|MAP3K5-AS1 | 111.3141 | 8.7841 | ##### | 5.5963 | 2.1899E-08 | 8.5180E-08 |
| ENSG00000248540 | 54.5418 | -3.3138 | ##### | -5.5957 | 2.1980E-08 | 8.5482E-08 |
| ENSG00000105193\|RPS16 | 46371.1833 | -0.6916 | ##### | -5.5955 | 2.2003E-08 | 8.5562E-08 |
| ENSG00000072364\|AFF4 | 8925.7175 | 0.6867 | ##### | 5.5952 | 2.2034E-08 | 8.5673E-08 |
| ENSG00000106330\|MOSPD3 | 1094.6580 | 0.7487 | ##### | 5.5946 | 2.2110E-08 | 8.5958E-08 |
| ENSG00000287917 | 91.4879 | -1.5663 | ##### | -5.5945 | 2.2125E-08 | 8.6002E-08 |
| ENSG00000273748 | 564.8787 | ####### | ##### | -5.5940 | 2.2185E-08 | 8.6224E-08 |
| ENSG00000109762\|SNX25 | 876.1885 | -0.8103 | ##### | -5.5940 | 2.2191E-08 | 8.6238E-08 |

| | | | | | | |
|---|---|---|---|---|---|---|
| ENSG00000109472\|CPE | 1659.1010 | 0.7380 | ##### | 5.5924 | 2.2391E-08 | 8.7002E-08 |
| ENSG00000256667\|KLRA1P | 90.5007 | 7.7164 | ##### | 5.5913 | 2.2534E-08 | 8.7546E-08 |
| ENSG00000215270\|TOMM40P2 | 284.0243 | 0.9148 | ##### | 5.5910 | 2.2574E-08 | 8.7691E-08 |
| ENSG00000115507\|OTX1 | 343.6644 | -0.8499 | ##### | -5.5910 | 2.2578E-08 | 8.7697E-08 |
| ENSG00000116106\|EPHA4 | 154.5513 | 1.5744 | ##### | 5.5892 | 2.2810E-08 | 8.8584E-08 |
| ENSG00000164125\|GASK1B | 442.2735 | -0.8739 | ##### | -5.5891 | 2.2827E-08 | 8.8640E-08 |
| ENSG00000280798\|LINC00294 | 1044.6714 | 0.7501 | ##### | 5.5884 | 2.2915E-08 | 8.8970E-08 |
| ENSG00000293076 | 262.7455 | -1.0167 | ##### | -5.5872 | 2.3076E-08 | 8.9582E-08 |
| ENSG00000140678\|ITGAX | 55.8082 | -8.7371 | ##### | -5.5871 | 2.3086E-08 | 8.9598E-08 |
| ENSG00000164953\|TMEM67 | 1151.1932 | -0.7479 | ##### | -5.5871 | 2.3085E-08 | 8.9598E-08 |
| ENSG00000250982\|GAPDHP35 | 17.4454 | 2.6440 | ##### | 5.5871 | 2.3090E-08 | 8.9602E-08 |
| ENSG00000159905\|ZNF221 | 135.4127 | -1.2868 | ##### | -5.5861 | 2.3223E-08 | 9.0108E-08 |
| ENSG00000099330\|OCEL1 | 436.5608 | -0.9375 | ##### | -5.5854 | 2.3323E-08 | 9.0482E-08 |
| ENSG00000289519 | 108.1208 | 1.5629 | ##### | 5.5832 | 2.3612E-08 | 9.1593E-08 |
| ENSG00000213082\|PPP1R14BP2 | 27.0438 | -2.4680 | ##### | -5.5831 | 2.3629E-08 | 9.1645E-08 |
| ENSG00000170004\|CHD3 | 15091.4505 | 0.6736 | ##### | 5.5830 | 2.3637E-08 | 9.1665E-08 |
| ENSG00000272518 | 34.0035 | -2.0617 | ##### | -5.5825 | 2.3714E-08 | 9.1952E-08 |
| ENSG00000256013 | 52.0293 | -1.7021 | ##### | -5.5803 | 2.4007E-08 | 9.3075E-08 |
| ENSG00000167676\|PLIN4 | 100.6579 | 1.2593 | ##### | 5.5801 | 2.4033E-08 | 9.3166E-08 |
| ENSG00000113356\|POLR3G | 2617.0486 | 0.7071 | ##### | 5.5800 | 2.4052E-08 | 9.3228E-08 |
| ENSG00000160963\|COL26A1 | 314.8741 | -1.0994 | ##### | -5.5799 | 2.4061E-08 | 9.3251E-08 |
| ENSG00000234284\|ZNF879 | 68.2636 | -1.6787 | ##### | -5.5796 | 2.4113E-08 | 9.3439E-08 |
| ENSG00000018869\|ZNF582 | 187.3739 | -1.2306 | ##### | -5.5793 | 2.4143E-08 | 9.3544E-08 |
| ENSG00000128973\|CLN6 | 3562.8539 | 0.6854 | ##### | 5.5787 | 2.4233E-08 | 9.3881E-08 |
| ENSG00000104228\|TRIM35 | 1506.7812 | 0.7583 | ##### | 5.5784 | 2.4277E-08 | 9.4038E-08 |
| ENSG00000287837 | 70.8397 | -1.8176 | ##### | -5.5782 | 2.4308E-08 | 9.4145E-08 |
| ENSG00000290596 | 234.9118 | 1.0272 | ##### | 5.5775 | 2.4405E-08 | 9.4508E-08 |
| ENSG00000160050\|CCDC28B | 439.9359 | -0.8153 | ##### | -5.5774 | 2.4414E-08 | 9.4534E-08 |
| ENSG00000290888 | 139.1215 | 8.7719 | ##### | 5.5768 | 2.4501E-08 | 9.4858E-08 |
| ENSG00000132964\|CDK8 | 2320.0137 | 0.6788 | ##### | 5.5756 | 2.4674E-08 | 9.5512E-08 |
| ENSG00000291317\|TMEM276 | 780.8760 | -0.8194 | ##### | -5.5751 | 2.4739E-08 | 9.5754E-08 |
| ENSG00000254604 | 39.1311 | 2.6122 | ##### | 5.5750 | 2.4760E-08 | 9.5820E-08 |
| ENSG00000270094 | 72.3129 | 1.2669 | ##### | 5.5742 | 2.4863E-08 | 9.6207E-08 |
| ENSG00000254154\|CRYZL2P-SEC16B | 76.8283 | -2.5330 | ##### | -5.5739 | 2.4907E-08 | 9.6367E-08 |
| ENSG00000075945\|KIFAP3 | 1785.8717 | -0.7459 | ##### | -5.5738 | 2.4931E-08 | 9.6446E-08 |
| ENSG00000293106 | 807.1161 | -0.8094 | ##### | -5.5734 | 2.4986E-08 | 9.6648E-08 |
| ENSG00000133789\|SWAP70 | 4921.7108 | 0.6673 | ##### | 5.5731 | 2.5022E-08 | 9.6774E-08 |
| ENSG00000197302\|KRBOX5 | 806.0284 | -0.7639 | ##### | -5.5721 | 2.5163E-08 | 9.7305E-08 |
| ENSG00000076003\|MCM6 | 12866.8162 | -0.6400 | ##### | -5.5717 | 2.5230E-08 | 9.7553E-08 |
| ENSG00000198722\|UNC13B | 4305.3758 | 0.6862 | ##### | 5.5714 | 2.5277E-08 | 9.7720E-08 |
| ENSG00000115514\|TXNDC9 | 5835.0982 | -0.6886 | ##### | -5.5708 | 2.5351E-08 | 9.7993E-08 |
| ENSG00000033867\|SLC4A7 | 9417.6563 | 0.6880 | ##### | 5.5706 | 2.5386E-08 | 9.8116E-08 |
| ENSG00000180035\|ZNF48 | 1296.0786 | -1.2409 | ##### | -5.5705 | 2.5404E-08 | 9.8173E-08 |
| ENSG00000104517\|UBR5 | 9631.5624 | -0.6674 | ##### | -5.5697 | 2.5516E-08 | 9.8593E-08 |
| ENSG00000136643\|RPS6KC1 | 3074.9310 | -0.7203 | ##### | -5.5695 | 2.5554E-08 | 9.8727E-08 |
| ENSG00000166435\|XRRA1 | 1232.6061 | 0.7306 | ##### | 5.5681 | 2.5753E-08 | 9.9486E-08 |
| ENSG00000169902\|TPST1 | 1152.3434 | -0.5059 | ##### | -5.5679 | 2.5779E-08 | 9.9570E-08 |
| ENSG00000129084\|PSMA1 | 10380.2969 | -0.6843 | ##### | -5.5668 | 2.5940E-08 | 1.0018E-07 |
| ENSG00000183199\|HSP90AB3P | 575.3258 | 0.7993 | ##### | 5.5667 | 2.5964E-08 | 1.0026E-07 |
| ENSG00000274210\|RNVU1-27 | 46.1743 | 7.0661 | ##### | 5.5666 | 2.5976E-08 | 1.0030E-07 |
| ENSG00000078053\|AMPH | 129.8139 | 4.8950 | ##### | 5.5651 | 2.6207E-08 | 1.0117E-07 |
| ENSG00000179761\|PIPOX | 35.6919 | 2.8742 | ##### | 5.5650 | 2.6219E-08 | 1.0121E-07 |
| ENSG00000144233\|AMMECR1L | 4626.5910 | -0.7001 | ##### | -5.5649 | 2.6233E-08 | 1.0125E-07 |
| ENSG00000120910\|PPP3CC | 726.3455 | -0.8078 | ##### | -5.5619 | 2.6687E-08 | 1.0298E-07 |
| ENSG00000220267\|ACTBP8 | 17.4537 | -3.9827 | ##### | -5.5617 | 2.6720E-08 | 1.0310E-07 |
| ENSG00000155959\|VBP1 | 7966.5847 | 0.6834 | ##### | 5.5614 | 2.6754E-08 | 1.0322E-07 |
| ENSG00000087076\|HSD17B14 | 721.3292 | -0.7792 | ##### | -5.5603 | 2.6924E-08 | 1.0386E-07 |
| ENSG00000180667\|YOD1 | 1801.8897 | -0.7189 | ##### | -5.5599 | 2.6988E-08 | 1.0409E-07 |
| ENSG00000230373\|GOLGA6L5P | 53.4288 | -1.4059 | ##### | -5.5595 | 2.7061E-08 | 1.0436E-07 |
| ENSG00000181513\|ACBD4 | 678.6170 | -0.8052 | ##### | -5.5592 | 2.7107E-08 | 1.0452E-07 |
| ENSG00000169026\|SLC49A3 | 40.4809 | -2.1901 | ##### | -5.5592 | 2.7108E-08 | 1.0452E-07 |
| ENSG00000174365\|SNHG11 | 915.7126 | 0.8100 | ##### | 5.5589 | 2.7154E-08 | 1.0468E-07 |
| ENSG00000261604 | 106.5003 | -1.7166 | ##### | -5.5581 | 2.7276E-08 | 1.0514E-07 |

| | | | | | | |
|---|---|---|---|---|---|---|
| ENSG00000135250\|SRPK2 | 4598.9950 | -0.6802 | ##### | -5.5580 | 2.7287E-08 | 1.0516E-07 |
| ENSG00000205238\|SPDYE2 | 99.5155 | 1.4131 | ##### | 5.5573 | 2.7398E-08 | 1.0558E-07 |
| ENSG00000258947\|TUBB3 | 19893.8934 | 0.6677 | ##### | 5.5566 | 2.7508E-08 | 1.0599E-07 |
| ENSG00000122417\|ODF2L | 2543.3386 | -0.7033 | ##### | -5.5564 | 2.7537E-08 | 1.0609E-07 |
| ENSG00000181038\|METTL23 | 2060.1112 | 0.7477 | ##### | 5.5561 | 2.7588E-08 | 1.0627E-07 |
| ENSG00000168028\|RPSA | 72115.5122 | 0.6676 | ##### | 5.5539 | 2.7929E-08 | 1.0757E-07 |
| ENSG00000257446\|ZNF878 | 50.7075 | -9.0812 | ##### | -5.5538 | 2.7947E-08 | 1.0763E-07 |
| ENSG00000243716\|NPIPB5 | 3311.2258 | 0.7282 | ##### | 5.5537 | 2.7963E-08 | 1.0767E-07 |
| ENSG00000149418\|ST14 | 186.3842 | -7.5482 | ##### | -5.5528 | 2.8112E-08 | 1.0823E-07 |
| ENSG00000141376\|BCAS3 | 981.4756 | 0.7366 | ##### | 5.5515 | 2.8321E-08 | 1.0902E-07 |
| ENSG00000171428\|NAT1 | 85.6827 | -1.6615 | ##### | -5.5515 | 2.8331E-08 | 1.0905E-07 |
| ENSG00000182134\|TDRKH | 2167.0136 | 0.6928 | ##### | 5.5504 | 2.8500E-08 | 1.0968E-07 |
| ENSG00000039600\|SOX30 | 37.7432 | 4.6458 | ##### | 5.5498 | 2.8597E-08 | 1.1004E-07 |
| ENSG00000224699\|LAMTOR5-AS1 | 246.8496 | 0.9764 | ##### | 5.5498 | 2.8604E-08 | 1.1005E-07 |
| ENSG00000144681\|STAC | 2458.1747 | 0.6847 | ##### | 5.5494 | 2.8668E-08 | 1.1029E-07 |
| ENSG00000100796\|PPP4R3A | 6383.6108 | 0.6490 | ##### | 5.5491 | 2.8713E-08 | 1.1045E-07 |
| ENSG00000249846\|LINC02021 | 38.8905 | 7.1362 | ##### | 5.5491 | 2.8720E-08 | 1.1046E-07 |
| ENSG00000160818\|GPATCH4 | 12997.4588 | 1.0555 | ##### | 5.5467 | 2.9115E-08 | 1.1196E-07 |
| ENSG00000171016\|PYGO1 | 759.0985 | -0.8039 | ##### | -5.5464 | 2.9156E-08 | 1.1211E-07 |
| ENSG00000101464\|PIGU | 1636.1879 | -0.7073 | ##### | -5.5446 | 2.9469E-08 | 1.1330E-07 |
| ENSG00000278903 | 30.8547 | -3.9602 | ##### | -5.5440 | 2.9564E-08 | 1.1365E-07 |
| ENSG00000144677\|CTDSPL | 1159.2318 | -0.7108 | ##### | -5.5416 | 2.9971E-08 | 1.1520E-07 |
| ENSG00000272462\|LINC02980 | 174.1503 | 1.3204 | ##### | 5.5404 | 3.0182E-08 | 1.1599E-07 |
| ENSG00000250072\|SH3TC2-DT | 36.1128 | 7.3960 | ##### | 5.5390 | 3.0413E-08 | 1.1687E-07 |
| ENSG00000198758\|EPS8L3 | 700.2632 | ####### | ##### | -5.5390 | 3.0424E-08 | 1.1689E-07 |
| ENSG00000115368\|WDR75 | 11078.7054 | 0.3667 | ##### | 5.5355 | 3.1027E-08 | 1.1919E-07 |
| ENSG00000229758\|DYNLT3P2 | 22.7081 | -6.9325 | ##### | -5.5354 | 3.1053E-08 | 1.1928E-07 |
| ENSG00000136261\|BZW2 | 11983.1471 | -0.6749 | ##### | -5.5350 | 3.1121E-08 | 1.1953E-07 |
| ENSG00000177082\|WDR73 | 2471.8352 | 0.7093 | ##### | 5.5349 | 3.1144E-08 | 1.1960E-07 |
| ENSG00000283525 | 136.4498 | 8.8164 | ##### | 5.5348 | 3.1163E-08 | 1.1966E-07 |
| ENSG00000152990\|ADGRA3 | 4887.2554 | 0.6948 | ##### | 5.5344 | 3.1236E-08 | 1.1992E-07 |
| ENSG00000089063\|TMEM230 | 6848.9361 | -0.6723 | ##### | -5.5341 | 3.1275E-08 | 1.2006E-07 |
| ENSG00000280013 | 13.7164 | 4.1073 | ##### | 5.5337 | 3.1347E-08 | 1.2032E-07 |
| ENSG00000131943\|C19orf12 | 2600.7331 | -0.6656 | ##### | -5.5337 | 3.1361E-08 | 1.2035E-07 |
| ENSG00000043143\|JADE2 | 5963.4215 | 0.6737 | ##### | 5.5336 | 3.1368E-08 | 1.2036E-07 |
| ENSG00000151364\|KCTD14 | 634.7145 | 8.2551 | ##### | 5.5324 | 3.1589E-08 | 1.2120E-07 |
| ENSG00000089234\|BRAP | 3200.6107 | 0.6766 | ##### | 5.5301 | 3.2004E-08 | 1.2277E-07 |
| ENSG00000168411\|RFWD3 | 10549.5221 | 0.6506 | ##### | 5.5293 | 3.2152E-08 | 1.2333E-07 |
| ENSG00000204366\|ZBTB12 | 705.5593 | 0.8101 | ##### | 5.5291 | 3.2191E-08 | 1.2346E-07 |
| ENSG00000164002\|EXO5 | 1160.3617 | -0.7168 | ##### | -5.5289 | 3.2232E-08 | 1.2360E-07 |
| ENSG00000048162\|NOP16 | 5465.4362 | 0.6886 | ##### | 5.5259 | 3.2774E-08 | 1.2566E-07 |
| ENSG00000070748\|CHAT | 126.9201 | 8.5863 | ##### | 5.5252 | 3.2914E-08 | 1.2619E-07 |
| ENSG00000170049\|KCNAB3 | 220.3426 | 1.1032 | ##### | 5.5250 | 3.2947E-08 | 1.2629E-07 |
| ENSG00000214248 | 770.1840 | -0.8163 | ##### | -5.5244 | 3.3069E-08 | 1.2674E-07 |
| ENSG00000005007\|UPF1 | 12138.4350 | 0.6527 | ##### | 5.5234 | 3.3251E-08 | 1.2743E-07 |
| ENSG00000170485\|NPAS2 | 3224.9609 | -0.7028 | ##### | -5.5230 | 3.3333E-08 | 1.2773E-07 |
| ENSG00000101888\|NXT2 | 1737.3216 | 0.7223 | ##### | 5.5205 | 3.3811E-08 | 1.2954E-07 |
| ENSG00000189144\|ZNF573 | 194.0241 | -1.2787 | ##### | -5.5199 | 3.3910E-08 | 1.2990E-07 |
| ENSG00000287031 | 127.3370 | 8.5108 | ##### | 5.5199 | 3.3917E-08 | 1.2991E-07 |
| ENSG00000293422\|PRORSD1P | 110.9511 | -1.0961 | ##### | -5.5189 | 3.4110E-08 | 1.3064E-07 |
| ENSG00000113721\|PDGFRB | 50.9424 | -2.2740 | ##### | -5.5185 | 3.4182E-08 | 1.3089E-07 |
| ENSG00000150457\|LATS2 | 5696.7153 | 0.6561 | ##### | 5.5176 | 3.4374E-08 | 1.3161E-07 |
| ENSG00000204179\|PTPN20 | 1999.7337 | ####### | ##### | -5.5175 | 3.4392E-08 | 1.3167E-07 |
| ENSG00000164080\|RAD54L2 | 2359.7348 | 0.7276 | ##### | 5.5154 | 3.4803E-08 | 1.3322E-07 |
| ENSG00000257390 | 734.4059 | 6.8429 | ##### | 5.5141 | 3.5062E-08 | 1.3420E-07 |
| ENSG00000226690 | 36.7640 | -2.4552 | ##### | -5.5139 | 3.5106E-08 | 1.3434E-07 |
| ENSG00000182093\|GET1 | 2257.8881 | -0.6458 | ##### | -5.5137 | 3.5130E-08 | 1.3442E-07 |
| ENSG00000233030 | 99.9253 | 1.3407 | ##### | 5.5131 | 3.5251E-08 | 1.3487E-07 |
| ENSG00000155893\|PXYLP1 | 503.1274 | -0.8853 | ##### | -5.5127 | 3.5339E-08 | 1.3519E-07 |
| ENSG00000111666\|CHPT1 | 6400.4798 | 0.6742 | ##### | 5.5113 | 3.5620E-08 | 1.3624E-07 |
| ENSG00000292285 | 140.5624 | 1.5358 | ##### | 5.5106 | 3.5768E-08 | 1.3679E-07 |
| ENSG00000197181\|PIWIL2 | 81.8080 | 3.3524 | ##### | 5.5101 | 3.5858E-08 | 1.3712E-07 |
| ENSG00000162510\|MATN1 | 83.4121 | 1.4679 | ##### | 5.5094 | 3.6012E-08 | 1.3769E-07 |

| | | | | | | |
|---|---|---|---|---|---|---|
| ENSG00000206573\|THUMPD3-AS1 | 704.1732 | 0.8052 | ##### | 5.5091 | 3.6061E-08 | 1.3786E-07 |
| ENSG00000228878\|SEPTIN7-DT | 79.3305 | -1.4565 | ##### | -5.5089 | 3.6110E-08 | 1.3803E-07 |
| ENSG00000148498\|PARD3 | 5780.1043 | -0.6591 | ##### | -5.5088 | 3.6135E-08 | 1.3811E-07 |
| ENSG00000188681\|TEKT4P2 | 34.6167 | 2.1279 | ##### | 5.5087 | 3.6153E-08 | 1.3816E-07 |
| ENSG00000140905\|GCSH | 3761.5007 | -0.6858 | ##### | -5.5082 | 3.6245E-08 | 1.3849E-07 |
| ENSG00000196557\|CACNA1H | 2605.1461 | -0.6940 | ##### | -5.5075 | 3.6395E-08 | 1.3905E-07 |
| ENSG00000142178\|SIK1 | 9538.7990 | -0.6519 | ##### | -5.5068 | 3.6534E-08 | 1.3956E-07 |
| ENSG00000270504 | 559.9571 | -0.7649 | ##### | -5.5065 | 3.6614E-08 | 1.3985E-07 |
| ENSG00000254941 | 17.2687 | 4.0078 | ##### | 5.5062 | 3.6673E-08 | 1.4005E-07 |
| ENSG00000227620\|ALG1L8P | 28.3869 | 2.1571 | ##### | 5.5060 | 3.6704E-08 | 1.4016E-07 |
| ENSG00000204381\|LAYN | 239.2308 | 4.9079 | ##### | 5.5057 | 3.6772E-08 | 1.4040E-07 |
| ENSG00000240137\|ERICH6-AS1 | 276.7778 | 1.0100 | ##### | 5.5038 | 3.7173E-08 | 1.4191E-07 |
| ENSG00000259169\|GNRHR2P1 | 331.9361 | 0.8646 | ##### | 5.5036 | 3.7220E-08 | 1.4207E-07 |
| ENSG00000173226\|IQCB1 | 1949.9894 | 0.7120 | ##### | 5.5031 | 3.7312E-08 | 1.4241E-07 |
| ENSG00000286975 | 88.0366 | 8.3801 | ##### | 5.5024 | 3.7470E-08 | 1.4299E-07 |
| ENSG00000244932\|IFT122P2 | 34.6277 | 1.6612 | ##### | 5.5007 | 3.7824E-08 | 1.4432E-07 |
| ENSG00000267755 | 121.4164 | 1.5181 | ##### | 5.5006 | 3.7840E-08 | 1.4436E-07 |
| ENSG00000188283\|ZNF383 | 613.6631 | -0.8650 | ##### | -5.4996 | 3.8071E-08 | 1.4523E-07 |
| ENSG00000256673 | 89.3560 | 2.3587 | ##### | 5.4995 | 3.8090E-08 | 1.4528E-07 |
| ENSG00000145979\|TBC1D7 | 783.7964 | 0.7697 | ##### | 5.4992 | 3.8148E-08 | 1.4548E-07 |
| ENSG00000198730\|CTR9 | 5945.1933 | 0.6165 | ##### | 5.4989 | 3.8215E-08 | 1.4572E-07 |
| ENSG00000177225\|GATD1 | 2341.4037 | 0.6717 | ##### | 5.4986 | 3.8292E-08 | 1.4600E-07 |
| ENSG00000164062\|APEH | 9471.7694 | 0.6454 | ##### | 5.4982 | 3.8376E-08 | 1.4630E-07 |
| ENSG00000235173\|HGH1 | 5146.2209 | 0.6605 | ##### | 5.4980 | 3.8417E-08 | 1.4644E-07 |
| ENSG00000198663\|C6orf89 | 5448.7089 | -0.6527 | ##### | -5.4968 | 3.8672E-08 | 1.4739E-07 |
| ENSG00000155761\|SPAG17 | 69.5938 | -1.5431 | ##### | -5.4967 | 3.8687E-08 | 1.4743E-07 |
| ENSG00000075711\|DLG1 | 3963.5010 | -0.6717 | ##### | -5.4966 | 3.8720E-08 | 1.4753E-07 |
| ENSG00000135148\|TRAFD1 | 2778.4778 | -0.7171 | ##### | -5.4966 | 3.8728E-08 | 1.4755E-07 |
| ENSG00000162006\|MSLNL | 736.4849 | ####### | ##### | -5.4964 | 3.8754E-08 | 1.4763E-07 |
| ENSG00000055950\|MRPL43 | 3372.0248 | -0.7002 | ##### | -5.4964 | 3.8766E-08 | 1.4765E-07 |
| ENSG00000289575 | 1360.6723 | ####### | ##### | -5.4949 | 3.9084E-08 | 1.4884E-07 |
| ENSG00000020922\|MRE11 | 2498.9506 | 0.6292 | ##### | 5.4948 | 3.9115E-08 | 1.4895E-07 |
| ENSG00000162585\|FAAP20 | 4007.8147 | 0.6812 | ##### | 5.4939 | 3.9314E-08 | 1.4968E-07 |
| ENSG00000289731 | 122.1346 | 8.5265 | ##### | 5.4930 | 3.9518E-08 | 1.5044E-07 |
| ENSG00000110844\|PRPF40B | 2587.4512 | 0.7075 | ##### | 5.4929 | 3.9538E-08 | 1.5048E-07 |
| ENSG00000064601\|CTSA | 20120.7303 | 0.6472 | ##### | 5.4929 | 3.9536E-08 | 1.5048E-07 |
| ENSG00000178764\|ZHX2 | 758.9517 | 0.7571 | ##### | 5.4918 | 3.9781E-08 | 1.5139E-07 |
| ENSG00000119402\|FBXW2 | 7541.8103 | -0.4707 | ##### | -5.4917 | 3.9799E-08 | 1.5143E-07 |
| ENSG00000170689\|HOXB9 | 225.4362 | 1.1147 | ##### | 5.4897 | 4.0253E-08 | 1.5314E-07 |
| ENSG00000198380\|GFPT1 | 9950.4574 | -0.6386 | ##### | -5.4883 | 4.0588E-08 | 1.5439E-07 |
| ENSG00000130811\|EIF3G | 13698.1203 | 0.6203 | ##### | 5.4878 | 4.0695E-08 | 1.5478E-07 |
| ENSG00000091483\|FH | 6556.3759 | -0.6433 | ##### | -5.4865 | 4.1007E-08 | 1.5595E-07 |
| ENSG00000180488\|MIGA1 | 4441.7684 | -0.6752 | ##### | -5.4837 | 4.1654E-08 | 1.5839E-07 |
| ENSG00000101695\|RNF125 | 479.1795 | 0.8078 | ##### | 5.4818 | 4.2109E-08 | 1.6010E-07 |
| ENSG00000169895\|SYAP1 | 6573.4897 | -0.5273 | ##### | -5.4804 | 4.2445E-08 | 1.6136E-07 |
| ENSG00000232956\|SNHG15 | 3368.2819 | 0.7180 | ##### | 5.4792 | 4.2725E-08 | 1.6240E-07 |
| ENSG00000117228\|GBP1 | 386.7858 | -7.7253 | ##### | -5.4789 | 4.2792E-08 | 1.6264E-07 |
| ENSG00000079101\|CLUL1 | 36.1903 | 3.9208 | ##### | 5.4786 | 4.2865E-08 | 1.6289E-07 |
| ENSG00000167302\|TEPSIN | 2559.7292 | 0.6917 | ##### | 5.4784 | 4.2917E-08 | 1.6307E-07 |
| ENSG00000166902\|MRPL16 | 2719.1091 | 0.6681 | ##### | 5.4781 | 4.2980E-08 | 1.6329E-07 |
| ENSG00000231840\|TMEM139-AS1 | 99.6451 | -1.9870 | ##### | -5.4776 | 4.3115E-08 | 1.6378E-07 |
| ENSG00000115602\|IL1RL1 | 40.9195 | -3.1984 | ##### | -5.4760 | 4.3502E-08 | 1.6523E-07 |
| ENSG00000136870\|ZNF189 | 858.9479 | -0.7575 | ##### | -5.4759 | 4.3536E-08 | 1.6534E-07 |
| ENSG00000198231\|DDX42 | 14162.7426 | 0.6444 | ##### | 5.4749 | 4.3787E-08 | 1.6627E-07 |
| ENSG00000262202 | 25.5966 | 5.0091 | ##### | 5.4745 | 4.3873E-08 | 1.6657E-07 |
| ENSG00000138641\|HERC3 | 975.1262 | -3.5906 | ##### | -5.4733 | 4.4162E-08 | 1.6765E-07 |
| ENSG00000244513\|EOGT-DT | 76.7937 | -1.6338 | ##### | -5.4731 | 4.4228E-08 | 1.6788E-07 |
| ENSG00000174939\|ASPHD1 | 2154.4441 | -0.6662 | ##### | -5.4726 | 4.4351E-08 | 1.6833E-07 |
| ENSG00000132016\|BRME1 | 255.6810 | 1.0128 | ##### | 5.4725 | 4.4378E-08 | 1.6841E-07 |
| ENSG00000163617\|CCDC191 | 371.2582 | -0.8082 | ##### | -5.4717 | 4.4567E-08 | 1.6910E-07 |
| ENSG00000163608\|NEPRO | 4246.6277 | 0.6283 | ##### | 5.4716 | 4.4608E-08 | 1.6923E-07 |
| ENSG00000101152\|DNAJC5 | 12182.5894 | 0.6444 | ##### | 5.4711 | 4.4724E-08 | 1.6965E-07 |
| ENSG00000244257\|PKD1P1 | 3833.8419 | 0.6812 | ##### | 5.4701 | 4.4986E-08 | 1.7062E-07 |

| | | | | | | |
|---|---|---|---|---|---|---|
| ENSG00000181315\|ZNF322 | 1004.3774 | 0.7481 | ##### | 5.4700 | 4.5006E-08 | 1.7068E-07 |
| ENSG00000171806\|METTL18 | 360.1113 | -0.8869 | ##### | -5.4698 | 4.5044E-08 | 1.7080E-07 |
| ENSG00000163870\|TPRA1 | 3451.6073 | -0.6879 | ##### | -5.4698 | 4.5064E-08 | 1.7086E-07 |
| ENSG00000171928\|TVP23B | 3685.7158 | -0.6999 | ##### | -5.4685 | 4.5379E-08 | 1.7203E-07 |
| ENSG00000022840\|RNF10 | 13449.7884 | 0.3677 | ##### | 5.4685 | 4.5395E-08 | 1.7207E-07 |
| ENSG00000029363\|BCLAF1 | 13095.0104 | 0.6693 | ##### | 5.4669 | 4.5792E-08 | 1.7355E-07 |
| ENSG00000101868\|POLA1 | 3073.2227 | 0.6661 | ##### | 5.4662 | 4.5976E-08 | 1.7423E-07 |
| ENSG00000108433\|GOSR2 | 5037.2773 | 0.6617 | ##### | 5.4651 | 4.6253E-08 | 1.7525E-07 |
| ENSG00000135090\|TAOK3 | 1534.4020 | -0.7147 | ##### | -5.4647 | 4.6365E-08 | 1.7566E-07 |
| ENSG00000119640\|ACYP1 | 468.1427 | 0.9035 | ##### | 5.4646 | 4.6386E-08 | 1.7571E-07 |
| ENSG00000234797\|RPS3AP6 | 277.1169 | -0.8614 | ##### | -5.4644 | 4.6442E-08 | 1.7590E-07 |
| ENSG00000010318\|PHF7 | 428.7074 | 0.8774 | ##### | 5.4639 | 4.6578E-08 | 1.7639E-07 |
| ENSG00000238227\|TMEM250 | 4452.2156 | -0.6526 | ##### | -5.4609 | 4.7363E-08 | 1.7934E-07 |
| ENSG00000261914\|RPL23AP84 | 26.8842 | 2.1074 | ##### | 5.4597 | 4.7687E-08 | 1.8055E-07 |
| ENSG00000100644\|HIF1A | 27586.4587 | -0.6563 | ##### | -5.4594 | 4.7784E-08 | 1.8089E-07 |
| ENSG00000248275\|TRIM52-AS1 | 233.2269 | -1.0209 | ##### | -5.4592 | 4.7832E-08 | 1.8105E-07 |
| ENSG00000145780\|FEM1C | 1670.4650 | 0.7127 | ##### | 5.4590 | 4.7874E-08 | 1.8119E-07 |
| ENSG00000164776\|PHKG1 | 547.8249 | -1.1830 | ##### | -5.4578 | 4.8206E-08 | 1.8242E-07 |
| ENSG00000183831\|ANKRD45 | 63.2926 | -1.9864 | ##### | -5.4552 | 4.8910E-08 | 1.8506E-07 |
| ENSG00000266640\|MIR4754 | 16.8283 | 6.2853 | ##### | 5.4551 | 4.8932E-08 | 1.8512E-07 |
| ENSG00000160404\|TOR2A | 699.3815 | -0.7962 | ##### | -5.4542 | 4.9190E-08 | 1.8608E-07 |
| ENSG00000271826\|PLS3-AS1 | 80.3206 | 1.3127 | ##### | 5.4535 | 4.9392E-08 | 1.8682E-07 |
| ENSG00000240053\|LY6G5B | 801.4675 | 0.9918 | ##### | 5.4532 | 4.9483E-08 | 1.8714E-07 |
| ENSG00000136026\|CKAP4 | 13391.7896 | -0.6754 | ##### | -5.4523 | 4.9734E-08 | 1.8806E-07 |
| ENSG00000090382\|LYZ | 37.6186 | 7.1535 | ##### | 5.4514 | 4.9981E-08 | 1.8897E-07 |
| ENSG00000261613 | 359.1217 | 0.9384 | ##### | 5.4511 | 5.0067E-08 | 1.8927E-07 |
| ENSG00000137573\|SULF1 | 141.6344 | -7.7078 | ##### | -5.4506 | 5.0206E-08 | 1.8977E-07 |
| ENSG00000134533\|RERG | 380.6170 | -6.6312 | ##### | -5.4499 | 5.0403E-08 | 1.9050E-07 |
| ENSG00000291041 | 29.2780 | 2.1307 | ##### | 5.4493 | 5.0581E-08 | 1.9114E-07 |
| ENSG00000175137\|SH3BP5L | 2711.3251 | -0.6881 | ##### | -5.4492 | 5.0611E-08 | 1.9123E-07 |
| ENSG00000290531\|TSPAN11-AS1 | 27.6579 | 3.1133 | ##### | 5.4483 | 5.0867E-08 | 1.9218E-07 |
| ENSG00000029153\|BMAL2 | 10750.9426 | -0.6717 | ##### | -5.4478 | 5.1005E-08 | 1.9267E-07 |
| ENSG00000164749\|HNF4G | 461.6525 | -6.3853 | ##### | -5.4476 | 5.1056E-08 | 1.9284E-07 |
| ENSG00000099617\|EFNA2 | 123.3187 | 1.1898 | ##### | 5.4474 | 5.1102E-08 | 1.9299E-07 |
| ENSG00000148053\|NTRK2 | 116.7780 | 1.8276 | ##### | 5.4466 | 5.1355E-08 | 1.9392E-07 |
| ENSG00000267323\|SLC25A1P5 | 221.8683 | -1.2740 | ##### | -5.4456 | 5.1643E-08 | 1.9498E-07 |
| ENSG00000255408\|PCDHA3 | 33.9607 | -4.5946 | ##### | -5.4453 | 5.1715E-08 | 1.9523E-07 |
| ENSG00000157168\|NRG1 | 5660.3945 | -0.6617 | ##### | -5.4449 | 5.1834E-08 | 1.9565E-07 |
| ENSG00000100412\|ACO2 | 11656.7766 | -0.6528 | ##### | -5.4447 | 5.1905E-08 | 1.9590E-07 |
| ENSG00000130545\|CRB3 | 190.7508 | -6.0587 | ##### | -5.4441 | 5.2071E-08 | 1.9650E-07 |
| ENSG00000278266 | 680.3806 | ####### | ##### | -5.4438 | 5.2147E-08 | 1.9676E-07 |
| ENSG00000266028\|SRGAP2 | 4342.6458 | 0.6359 | ##### | 5.4433 | 5.2314E-08 | 1.9737E-07 |
| ENSG00000131115\|ZNF227 | 1339.8767 | -0.6991 | ##### | -5.4423 | 5.2602E-08 | 1.9843E-07 |
| ENSG00000227473\|TSSK5P | 314.4819 | 0.8541 | ##### | 5.4417 | 5.2780E-08 | 1.9908E-07 |
| ENSG00000105821\|DNAJC2 | 6577.4902 | 0.6809 | ##### | 5.4412 | 5.2934E-08 | 1.9963E-07 |
| ENSG00000124444\|ZNF576 | 1066.5610 | -0.7553 | ##### | -5.4408 | 5.3033E-08 | 1.9998E-07 |
| ENSG00000100276\|RASL10A | 45.1096 | 1.8475 | ##### | 5.4400 | 5.3266E-08 | 2.0083E-07 |
| ENSG00000089356\|FXYD3 | 242.9989 | ####### | ##### | -5.4388 | 5.3634E-08 | 2.0219E-07 |
| ENSG00000141569\|TRIM65 | 4004.7389 | -0.6337 | ##### | -5.4387 | 5.3670E-08 | 2.0230E-07 |
| ENSG00000230311\|TOMM20P4 | 132.8780 | -1.0612 | ##### | -5.4387 | 5.3683E-08 | 2.0233E-07 |
| ENSG00000211445\|GPX3 | 64.6633 | -1.7913 | ##### | -5.4383 | 5.3778E-08 | 2.0266E-07 |
| ENSG00000248817\|PMPCAP1 | 47.1197 | -7.5296 | ##### | -5.4382 | 5.3828E-08 | 2.0282E-07 |
| ENSG00000113282\|CLINT1 | 11213.0719 | 0.6461 | ##### | 5.4381 | 5.3847E-08 | 2.0287E-07 |
| ENSG00000267260 | 144.4001 | -1.1186 | ##### | -5.4374 | 5.4069E-08 | 2.0368E-07 |
| ENSG00000285077\|ARHGAP11B | 178.2385 | 1.2401 | ##### | 5.4370 | 5.4180E-08 | 2.0407E-07 |
| ENSG00000234405 | 56.6091 | -1.5655 | ##### | -5.4364 | 5.4377E-08 | 2.0479E-07 |
| ENSG00000227766 | 326.5584 | 1.6138 | ##### | 5.4360 | 5.4485E-08 | 2.0517E-07 |
| ENSG00000145919\|BOD1 | 4333.5447 | 0.6602 | ##### | 5.4359 | 5.4521E-08 | 2.0528E-07 |
| ENSG00000121207\|LRAT | 60.3198 | -4.5532 | ##### | -5.4352 | 5.4745E-08 | 2.0610E-07 |
| ENSG00000225393 | 15.1533 | -3.0771 | ##### | -5.4348 | 5.4844E-08 | 2.0644E-07 |
| ENSG00000132763\|MMACHC | 5669.9169 | -0.7413 | ##### | -5.4329 | 5.5442E-08 | 2.0867E-07 |
| ENSG00000262209\|PCDHGB3 | 66.7007 | -4.1104 | ##### | -5.4326 | 5.5545E-08 | 2.0903E-07 |
| ENSG00000289418 | 109.5939 | 2.1741 | ##### | 5.4325 | 5.5570E-08 | 2.0910E-07 |

| | | | | | | |
|---|---|---|---|---|---|---|
| ENSG00000105549\|SPMAP2 | 33.7503 | -3.3105 | ##### | -5.4319 | 5.5760E-08 | 2.0979E-07 |
| ENSG00000251602\|MTA1-DT | 145.8605 | 1.1571 | ##### | 5.4312 | 5.5982E-08 | 2.1059E-07 |
| ENSG00000143436\|MRPL9 | 9423.0751 | 0.5922 | ##### | 5.4299 | 5.6381E-08 | 2.1207E-07 |
| ENSG00000177595\|PIDD1 | 1399.8596 | 0.7551 | ##### | 5.4296 | 5.6492E-08 | 2.1246E-07 |
| ENSG00000285600 | 2066.6325 | 1.0624 | ##### | 5.4284 | 5.6875E-08 | 2.1387E-07 |
| ENSG00000134545\|KLRC1 | 118.6581 | 8.4484 | ##### | 5.4264 | 5.7505E-08 | 2.1621E-07 |
| ENSG00000131067\|GGT7 | 1644.0145 | -0.7051 | ##### | -5.4260 | 5.7634E-08 | 2.1667E-07 |
| ENSG00000286689 | 46.3648 | 6.8849 | ##### | 5.4255 | 5.7796E-08 | 2.1725E-07 |
| ENSG00000198888\|MT-ND1 | ########## | 0.6804 | ##### | 5.4247 | 5.8054E-08 | 2.1820E-07 |
| ENSG00000186973\|CFAP144 | 83.3544 | 1.6157 | ##### | 5.4247 | 5.8066E-08 | 2.1821E-07 |
| ENSG00000128563\|PRKRIP1 | 2453.8886 | 0.6673 | ##### | 5.4233 | 5.8496E-08 | 2.1980E-07 |
| ENSG00000171204\|TMEM126B | 2060.1631 | -0.6761 | ##### | -5.4232 | 5.8552E-08 | 2.1998E-07 |
| ENSG00000266560 | 658.7324 | ####### | ##### | -5.4224 | 5.8818E-08 | 2.2096E-07 |
| ENSG00000290927 | 200.0182 | 1.2581 | ##### | 5.4221 | 5.8916E-08 | 2.2130E-07 |
| ENSG00000119326\|CTNNAL1 | 13309.6574 | 0.4839 | ##### | 5.4220 | 5.8939E-08 | 2.2136E-07 |
| ENSG00000086102\|NFX1 | 3537.5000 | -0.5891 | ##### | -5.4214 | 5.9145E-08 | 2.2210E-07 |
| ENSG00000168955\|TM4SF20 | 1140.3945 | ####### | ##### | -5.4207 | 5.9360E-08 | 2.2288E-07 |
| ENSG00000111011\|RSRC2 | 4548.2615 | 0.5774 | ##### | 5.4198 | 5.9649E-08 | 2.2394E-07 |
| ENSG00000169410\|PTPN9 | 3008.7043 | -0.6639 | ##### | -5.4190 | 5.9934E-08 | 2.2498E-07 |
| ENSG00000293183 | 129.1506 | -1.2163 | ##### | -5.4185 | 6.0114E-08 | 2.2563E-07 |
| ENSG00000133048\|CHI3L1 | 61.3882 | 8.4117 | ##### | 5.4183 | 6.0164E-08 | 2.2579E-07 |
| ENSG00000260231\|KDM7A-DT | 350.9077 | -0.8862 | ##### | -5.4177 | 6.0380E-08 | 2.2657E-07 |
| ENSG00000146963\|LUC7L2 | 10375.6810 | 0.6245 | ##### | 5.4174 | 6.0471E-08 | 2.2688E-07 |
| ENSG00000204540\|PSORS1C1 | 190.7945 | -7.4062 | ##### | -5.4159 | 6.0969E-08 | 2.2872E-07 |
| ENSG00000172817\|CYP7B1 | 56.4510 | -2.4837 | ##### | -5.4158 | 6.1011E-08 | 2.2885E-07 |
| ENSG00000036549\|ZZZ3 | 4259.8376 | -0.6616 | ##### | -5.4157 | 6.1044E-08 | 2.2894E-07 |
| ENSG00000290114 | 51.6173 | -2.2409 | ##### | -5.4156 | 6.1071E-08 | 2.2902E-07 |
| ENSG00000285722 | 22.3328 | -4.6913 | ##### | -5.4155 | 6.1103E-08 | 2.2911E-07 |
| ENSG00000158164\|TMSB15A | 128.5550 | 4.6776 | ##### | 5.4142 | 6.1556E-08 | 2.3078E-07 |
| ENSG00000160908\|ZNF394 | 1616.2072 | 0.7189 | ##### | 5.4132 | 6.1897E-08 | 2.3203E-07 |
| ENSG00000108759\|KRT32 | 40.0265 | 3.5684 | ##### | 5.4130 | 6.1977E-08 | 2.3230E-07 |
| ENSG00000215186\|GOLGA6B | 64.2309 | 7.9041 | ##### | 5.4124 | 6.2185E-08 | 2.3305E-07 |
| ENSG00000171566\|PLRG1 | 4401.3989 | -0.6777 | ##### | -5.4118 | 6.2411E-08 | 2.3386E-07 |
| ENSG00000232023\|LINC01807 | 557.6906 | ####### | ##### | -5.4114 | 6.2524E-08 | 2.3426E-07 |
| ENSG00000170604\|IRF2BP1 | 3084.6552 | -0.6640 | ##### | -5.4114 | 6.2533E-08 | 2.3426E-07 |
| ENSG00000111843\|TMEM14C | 8257.5553 | 0.6666 | ##### | 5.4109 | 6.2719E-08 | 2.3493E-07 |
| ENSG00000291080\|ZNF252P | 1749.2219 | -0.6953 | ##### | -5.4096 | 6.3159E-08 | 2.3655E-07 |
| ENSG00000174928\|C3orf33 | 565.2007 | -0.8139 | ##### | -5.4095 | 6.3214E-08 | 2.3673E-07 |
| ENSG00000163913\|IFT122 | 2903.0727 | 0.6736 | ##### | 5.4093 | 6.3264E-08 | 2.3688E-07 |
| ENSG00000243517\|RPL13AP22 | 65.0373 | -1.3201 | ##### | -5.4088 | 6.3461E-08 | 2.3759E-07 |
| ENSG00000096093\|EFHC1 | 1237.8016 | 0.7405 | ##### | 5.4086 | 6.3523E-08 | 2.3779E-07 |
| ENSG00000104886\|PLEKHJ1 | 2745.3003 | 0.6804 | ##### | 5.4056 | 6.4604E-08 | 2.4181E-07 |
| ENSG00000228175\|GEMIN8P4 | 66.4141 | -1.4820 | ##### | -5.4054 | 6.4669E-08 | 2.4202E-07 |
| ENSG00000003987\|MTMR7 | 301.1914 | -0.9477 | ##### | -5.4052 | 6.4754E-08 | 2.4231E-07 |
| ENSG00000197372\|ZNF675 | 226.8024 | -1.0316 | ##### | -5.4044 | 6.5041E-08 | 2.4336E-07 |
| ENSG00000104835\|SARS2 | 1951.0807 | -0.6205 | ##### | -5.4042 | 6.5099E-08 | 2.4354E-07 |
| ENSG00000268945 | 35.2483 | 7.2371 | ##### | 5.4041 | 6.5143E-08 | 2.4365E-07 |
| ENSG00000232949 | 140.5729 | 8.3134 | ##### | 5.4041 | 6.5136E-08 | 2.4365E-07 |
| ENSG00000244479\|OR2A1-AS1 | 83.0261 | -1.3119 | ##### | -5.4037 | 6.5278E-08 | 2.4412E-07 |
| ENSG00000118564\|FBXL5 | 2241.1650 | -0.6847 | ##### | -5.4037 | 6.5297E-08 | 2.4416E-07 |
| ENSG00000171385\|KCND3 | 36.6591 | 3.5120 | ##### | 5.4035 | 6.5362E-08 | 2.4437E-07 |
| ENSG00000234880\|LINC00163 | 64.1111 | 7.6903 | ##### | 5.4026 | 6.5682E-08 | 2.4554E-07 |
| ENSG00000144468\|RHBDD1 | 1872.7801 | -0.6964 | ##### | -5.4006 | 6.6402E-08 | 2.4820E-07 |
| ENSG00000170100\|ZNF778 | 2105.8652 | -0.7008 | ##### | -5.4006 | 6.6410E-08 | 2.4820E-07 |
| ENSG00000154133\|ROBO4 | 140.7434 | 8.1680 | ##### | 5.3998 | 6.6709E-08 | 2.4928E-07 |
| ENSG00000255036\|SUGT1P4-STRA6LP-CCDC18( | 173.0501 | -1.7307 | ##### | -5.3996 | 6.6792E-08 | 2.4956E-07 |
| ENSG00000157404\|KIT | 515.8967 | 6.5903 | ##### | 5.3991 | 6.6990E-08 | 2.5027E-07 |
| ENSG00000198932\|GPRASP1 | 150.5493 | 1.5002 | ##### | 5.3986 | 6.7155E-08 | 2.5086E-07 |
| ENSG00000291215 | 566.2899 | 0.7813 | ##### | 5.3978 | 6.7445E-08 | 2.5191E-07 |
| ENSG00000069329\|VPS35 | 11167.9214 | -0.6361 | ##### | -5.3971 | 6.7744E-08 | 2.5299E-07 |
| ENSG00000105722\|ERF | 9398.3887 | 0.6312 | ##### | 5.3970 | 6.7771E-08 | 2.5306E-07 |
| ENSG00000171224\|FAM241B | 813.0091 | -0.7505 | ##### | -5.3968 | 6.7855E-08 | 2.5334E-07 |
| ENSG00000167130\|DOLPP1 | 1773.9516 | -0.6623 | ##### | -5.3965 | 6.7966E-08 | 2.5372E-07 |

| | | | | | | |
|---|---|---|---|---|---|---|
| ENSG00000185298\|CCDC137 | 5438.1464 | 0.6388 | ##### | 5.3961 | 6.8091E-08 | 2.5416E-07 |
| ENSG00000187486\|KCNJ11 | 117.8237 | 1.2307 | ##### | 5.3951 | 6.8471E-08 | 2.5555E-07 |
| ENSG00000183671\|CMKLR2 | 115.3274 | -5.5229 | ##### | -5.3948 | 6.8587E-08 | 2.5595E-07 |
| ENSG00000115317\|HTRA2 | 4216.0820 | 0.5920 | ##### | 5.3947 | 6.8639E-08 | 2.5611E-07 |
| ENSG00000134318\|ROCK2 | 12868.0657 | -0.6565 | ##### | -5.3945 | 6.8731E-08 | 2.5642E-07 |
| ENSG00000289156 | 31.3071 | 7.3055 | ##### | 5.3927 | 6.9423E-08 | 2.5897E-07 |
| ENSG00000249855\|EEF1A1P19 | 71.3356 | 1.3004 | ##### | 5.3924 | 6.9509E-08 | 2.5924E-07 |
| ENSG00000197582\|GPX1P1 | 130.1484 | -9.4017 | ##### | -5.3924 | 6.9514E-08 | 2.5924E-07 |
| ENSG00000288958 | 43.1353 | 1.7003 | ##### | 5.3916 | 6.9846E-08 | 2.6045E-07 |
| ENSG00000183291\|SELENOF | 14127.1205 | -0.6337 | ##### | -5.3910 | 7.0067E-08 | 2.6124E-07 |
| ENSG00000130734\|ATG4D | 1664.6780 | 0.7295 | ##### | 5.3893 | 7.0722E-08 | 2.6365E-07 |
| ENSG00000147003\|CLTRN | 1027.1031 | ####### | ##### | -5.3892 | 7.0784E-08 | 2.6385E-07 |
| ENSG00000277142\|LINC00235 | 127.7304 | 1.2191 | ##### | 5.3890 | 7.0862E-08 | 2.6411E-07 |
| ENSG00000167182\|SP2 | 1597.2956 | 0.7169 | ##### | 5.3883 | 7.1117E-08 | 2.6502E-07 |
| ENSG00000174343\|CHRNA9 | 106.7661 | 5.5103 | ##### | 5.3882 | 7.1162E-08 | 2.6516E-07 |
| ENSG00000289164 | 28.0349 | -2.0416 | ##### | -5.3879 | 7.1280E-08 | 2.6557E-07 |
| ENSG00000256250 | 100.7712 | 8.2122 | ##### | 5.3855 | 7.2256E-08 | 2.6917E-07 |
| ENSG00000116005\|PCYOX1 | 7440.1250 | -0.6480 | ##### | -5.3850 | 7.2444E-08 | 2.6984E-07 |
| ENSG00000279766 | 24.8727 | 2.3824 | ##### | 5.3847 | 7.2559E-08 | 2.7023E-07 |
| ENSG00000254837\|LIPT2-AS1 | 447.6077 | 0.8574 | ##### | 5.3846 | 7.2587E-08 | 2.7030E-07 |
| ENSG00000113648\|MACROH2A1 | 15847.4323 | 0.4252 | ##### | 5.3841 | 7.2802E-08 | 2.7107E-07 |
| ENSG00000228084 | 194.1892 | -9.9960 | ##### | -5.3841 | 7.2813E-08 | 2.7107E-07 |
| ENSG00000113312\|TTC1 | 5003.3640 | 0.6612 | ##### | 5.3837 | 7.2980E-08 | 2.7166E-07 |
| ENSG00000175707\|KDF1 | 118.7717 | -9.7848 | ##### | -5.3810 | 7.4077E-08 | 2.7571E-07 |
| ENSG00000135999\|EPC2 | 1814.0132 | 0.6892 | ##### | 5.3791 | 7.4862E-08 | 2.7857E-07 |
| ENSG00000137460\|FHDC1 | 62.0827 | -1.5342 | ##### | -5.3791 | 7.4863E-08 | 2.7857E-07 |
| ENSG00000132437\|DDC | 133.7416 | -6.8733 | ##### | -5.3790 | 7.4883E-08 | 2.7861E-07 |
| ENSG00000290877\|ROCK1P1 | 221.8307 | 0.9833 | ##### | 5.3785 | 7.5111E-08 | 2.7942E-07 |
| ENSG00000135047\|CTSL | 14731.2490 | -0.5871 | ##### | -5.3773 | 7.5601E-08 | 2.8121E-07 |
| ENSG00000281005\|LINC00921 | 47.2822 | -1.5365 | ##### | -5.3771 | 7.5693E-08 | 2.8152E-07 |
| ENSG00000136161\|RCBTB2 | 428.9301 | 0.8318 | ##### | 5.3766 | 7.5889E-08 | 2.8221E-07 |
| ENSG00000214113\|LYRM4 | 1454.2275 | 0.7145 | ##### | 5.3759 | 7.6218E-08 | 2.8340E-07 |
| ENSG00000289096 | 107.7885 | 8.2520 | ##### | 5.3748 | 7.6673E-08 | 2.8505E-07 |
| ENSG00000290687\|ZNF286B | 263.1573 | 0.8545 | ##### | 5.3747 | 7.6694E-08 | 2.8510E-07 |
| ENSG00000249348\|UGDH-AS1 | 246.7145 | -1.1376 | ##### | -5.3742 | 7.6923E-08 | 2.8591E-07 |
| ENSG00000163785\|RYK | 6153.0173 | 0.6704 | ##### | 5.3737 | 7.7126E-08 | 2.8663E-07 |
| ENSG00000248810\|LINC02432 | 611.8017 | ####### | ##### | -5.3733 | 7.7315E-08 | 2.8730E-07 |
| ENSG00000010270\|STARD3NL | 5399.9873 | -0.6509 | ##### | -5.3721 | 7.7811E-08 | 2.8911E-07 |
| ENSG00000110031\|LPXN | 127.3719 | -1.3256 | ##### | -5.3720 | 7.7861E-08 | 2.8925E-07 |
| ENSG00000090924\|PLEKHG2 | 4621.1012 | -0.5948 | ##### | -5.3718 | 7.7939E-08 | 2.8951E-07 |
| ENSG00000284753\|EEF1AKMT4 | 1476.0878 | -0.6494 | ##### | -5.3712 | 7.8232E-08 | 2.9054E-07 |
| ENSG00000106665\|CLIP2 | 4327.8870 | -0.6604 | ##### | -5.3712 | 7.8236E-08 | 2.9054E-07 |
| ENSG00000155629\|PIK3AP1 | 168.5944 | 1.1397 | ##### | 5.3711 | 7.8247E-08 | 2.9055E-07 |
| ENSG00000007952\|NOX1 | 40.2894 | 1.7115 | ##### | 5.3708 | 7.8386E-08 | 2.9102E-07 |
| ENSG00000273084\|LINC03073 | 37.1321 | 3.3948 | ##### | 5.3702 | 7.8650E-08 | 2.9197E-07 |
| ENSG00000267296\|CEBPA-DT | 181.3094 | -1.0096 | ##### | -5.3683 | 7.9474E-08 | 2.9499E-07 |
| ENSG00000186310\|NAP1L3 | 112.5528 | -1.2711 | ##### | -5.3674 | 7.9878E-08 | 2.9645E-07 |
| ENSG00000215717\|TMEM167B | 4325.9909 | -0.6666 | ##### | -5.3672 | 7.9984E-08 | 2.9681E-07 |
| ENSG00000198886\|MT-ND4 | ########## | 0.6696 | ##### | 5.3670 | 8.0078E-08 | 2.9712E-07 |
| ENSG00000140839\|CLEC18B | 37.6192 | -1.9475 | ##### | -5.3658 | 8.0573E-08 | 2.9892E-07 |
| ENSG00000236892 | 14.1961 | -3.8299 | ##### | -5.3652 | 8.0871E-08 | 2.9999E-07 |
| ENSG00000167771\|RCOR2 | 214.8444 | 1.0052 | ##### | 5.3647 | 8.1072E-08 | 3.0070E-07 |
| ENSG00000168631\|MUCL3 | 32.0503 | -5.2496 | ##### | -5.3646 | 8.1143E-08 | 3.0092E-07 |
| ENSG00000285331\|TCF12-DT | 352.4322 | -0.9481 | ##### | -5.3640 | 8.1412E-08 | 3.0189E-07 |
| ENSG00000163923\|RPL39L | 1151.5758 | -0.7311 | ##### | -5.3622 | 8.2194E-08 | 3.0475E-07 |
| ENSG00000182979\|MTA1 | 9991.1752 | 0.6047 | ##### | 5.3622 | 8.2209E-08 | 3.0476E-07 |
| ENSG00000145088\|EAF2 | 296.3414 | 1.0586 | ##### | 5.3613 | 8.2645E-08 | 3.0634E-07 |
| ENSG00000258515 | 32.7584 | 2.5689 | ##### | 5.3596 | 8.3388E-08 | 3.0906E-07 |
| ENSG00000135525\|MAP7 | 6283.5228 | -0.6491 | ##### | -5.3594 | 8.3512E-08 | 3.0948E-07 |
| ENSG00000187957\|DNER | 283.6943 | -0.8733 | ##### | -5.3588 | 8.3780E-08 | 3.1043E-07 |
| ENSG00000182199\|SHMT2 | 15770.9130 | 0.6139 | ##### | 5.3585 | 8.3916E-08 | 3.1090E-07 |
| ENSG00000235191\|NUCB1-AS1 | 26.1439 | -7.0102 | ##### | -5.3578 | 8.4262E-08 | 3.1214E-07 |
| ENSG00000250575 | 110.0084 | 1.0678 | ##### | 5.3574 | 8.4408E-08 | 3.1264E-07 |

| | | | | | | |
|---|---|---|---|---|---|---|
| ENSG00000004766\|VPS50 | 1938.3185 | -0.6530 | ##### | -5.3550 | 8.5550E-08 | 3.1683E-07 |
| ENSG00000005302\|MSL3 | 1421.9030 | -0.6645 | ##### | -5.3549 | 8.5583E-08 | 3.1692E-07 |
| ENSG00000160310\|PRMT2 | 5342.0572 | -0.4597 | ##### | -5.3537 | 8.6178E-08 | 3.1908E-07 |
| ENSG00000162664\|ZNF326 | 3108.7066 | 0.6502 | ##### | 5.3533 | 8.6376E-08 | 3.1977E-07 |
| ENSG00000267469 | 3084.3483 | -0.7290 | ##### | -5.3524 | 8.6811E-08 | 3.2135E-07 |
| ENSG00000179943\|FIZ1 | 1951.5222 | -0.6818 | ##### | -5.3518 | 8.7063E-08 | 3.2224E-07 |
| ENSG00000250571\|GLI4 | 1378.4720 | -0.7314 | ##### | -5.3515 | 8.7233E-08 | 3.2283E-07 |
| ENSG00000166321\|NUDT13 | 518.6955 | 0.8102 | ##### | 5.3502 | 8.7836E-08 | 3.2502E-07 |
| ENSG00000179331\|RAB39A | 203.3708 | 1.2929 | ##### | 5.3499 | 8.7983E-08 | 3.2552E-07 |
| ENSG00000114738\|MAPKAPK3 | 2767.0157 | -0.6688 | ##### | -5.3498 | 8.8057E-08 | 3.2576E-07 |
| ENSG00000291194 | 116.4797 | -1.1611 | ##### | -5.3484 | 8.8723E-08 | 3.2818E-07 |
| ENSG00000123200\|ZC3H13 | 6200.8407 | 0.6443 | ##### | 5.3478 | 8.9029E-08 | 3.2927E-07 |
| ENSG00000210164\|MT-TG | 147.7037 | 1.0398 | ##### | 5.3477 | 8.9066E-08 | 3.2937E-07 |
| ENSG00000290083 | 29.9699 | 6.7078 | ##### | 5.3472 | 8.9311E-08 | 3.3023E-07 |
| ENSG00000207425\|Y_RNA | 13.2834 | -2.6727 | ##### | -5.3461 | 8.9857E-08 | 3.3217E-07 |
| ENSG00000106268\|NUDT1 | 3104.4075 | 0.6814 | ##### | 5.3461 | 8.9850E-08 | 3.3217E-07 |
| ENSG00000127125\|PPCS | 2455.2093 | -0.6723 | ##### | -5.3451 | 9.0362E-08 | 3.3399E-07 |
| ENSG00000272894\|MIOS-DT | 38.9935 | -2.0228 | ##### | -5.3448 | 9.0507E-08 | 3.3449E-07 |
| ENSG00000051523\|CYBA | 5619.6901 | -0.6656 | ##### | -5.3447 | 9.0551E-08 | 3.3461E-07 |
| ENSG00000203814\|H2BC18 | 154.1648 | 1.1412 | ##### | 5.3447 | 9.0588E-08 | 3.3471E-07 |
| ENSG00000139921\|TMX1 | 9164.9903 | 0.5790 | ##### | 5.3446 | 9.0617E-08 | 3.3477E-07 |
| ENSG00000185787\|MORF4L1 | 14559.5575 | -0.6623 | ##### | -5.3442 | 9.0839E-08 | 3.3555E-07 |
| ENSG00000129682\|FGF13 | 170.2076 | 8.1216 | ##### | 5.3441 | 9.0854E-08 | 3.3556E-07 |
| ENSG00000072954\|TMEM38A | 280.6796 | -0.9158 | ##### | -5.3418 | 9.2026E-08 | 3.3985E-07 |
| ENSG00000174106\|LEMD3 | 3635.5144 | 0.6376 | ##### | 5.3406 | 9.2619E-08 | 3.4200E-07 |
| ENSG00000205571\|SMN2 | 2326.5616 | -0.7373 | ##### | -5.3400 | 9.2929E-08 | 3.4310E-07 |
| ENSG00000233246\|PHC2-AS1 | 73.4093 | 8.1987 | ##### | 5.3396 | 9.3137E-08 | 3.4382E-07 |
| ENSG00000266709 | 41.7374 | 1.9603 | ##### | 5.3386 | 9.3658E-08 | 3.4571E-07 |
| ENSG00000223612 | 111.4848 | -1.0012 | ##### | -5.3383 | 9.3802E-08 | 3.4619E-07 |
| ENSG00000171033\|PKIA | 600.6572 | 0.7788 | ##### | 5.3381 | 9.3919E-08 | 3.4658E-07 |
| ENSG00000172775\|PSME3IP1 | 4023.5727 | -0.6463 | ##### | -5.3373 | 9.4350E-08 | 3.4813E-07 |
| ENSG00000267583\|ZNF24TR | 45.9795 | 5.6866 | ##### | 5.3369 | 9.4569E-08 | 3.4890E-07 |
| ENSG00000080824\|HSP90AA1 | ########## | 0.4917 | ##### | 5.3368 | 9.4587E-08 | 3.4892E-07 |
| ENSG00000111196\|MAGOHB | 2803.8976 | 0.6943 | ##### | 5.3359 | 9.5062E-08 | 3.5063E-07 |
| ENSG00000088035\|ALG6 | 1321.9063 | -0.6965 | ##### | -5.3355 | 9.5286E-08 | 3.5141E-07 |
| ENSG00000172667\|ZMAT3 | 2349.1188 | 0.6970 | ##### | 5.3342 | 9.5982E-08 | 3.5393E-07 |
| ENSG00000183072\|NKX2-5 | 599.8311 | 7.1591 | ##### | 5.3326 | 9.6841E-08 | 3.5706E-07 |
| ENSG00000210176\|MT-TH | 159.3795 | 1.0519 | ##### | 5.3322 | 9.7052E-08 | 3.5779E-07 |
| ENSG00000197912\|SPG7 | 9303.7966 | -0.5515 | ##### | -5.3318 | 9.7267E-08 | 3.5854E-07 |
| ENSG00000188886\|ASTL | 59.3049 | 3.8950 | ##### | 5.3310 | 9.7662E-08 | 3.5995E-07 |
| ENSG00000140557\|ST8SIA2 | 119.4985 | 8.1101 | ##### | 5.3309 | 9.7709E-08 | 3.6008E-07 |
| ENSG00000040531\|CTNS | 1968.1981 | 0.7059 | ##### | 5.3296 | 9.8427E-08 | 3.6268E-07 |
| ENSG00000068137\|PLEKHH3 | 2565.4035 | -0.6451 | ##### | -5.3295 | 9.8459E-08 | 3.6275E-07 |
| ENSG00000237399\|PITRM1-AS1 | 170.5592 | -1.0914 | ##### | -5.3282 | 9.9198E-08 | 3.6543E-07 |
| ENSG00000185245\|GP1BA | 116.5574 | 1.2824 | ##### | 5.3278 | 9.9413E-08 | 3.6618E-07 |
| ENSG00000177150\|FAM210A | 2641.3495 | 0.6783 | ##### | 5.3263 | 1.0025E-07 | 3.6921E-07 |
| ENSG00000106785\|TRIM14 | 3062.7033 | -0.6535 | ##### | -5.3258 | 1.0050E-07 | 3.7008E-07 |
| ENSG00000188493\|ACTMAP | 895.6411 | -0.7511 | ##### | -5.3257 | 1.0059E-07 | 3.7037E-07 |
| ENSG00000248740\|LINC02428 | 71.6298 | -4.5504 | ##### | -5.3243 | 1.0137E-07 | 3.7319E-07 |
| ENSG00000246100\|LINC00900 | 44.3332 | 7.1568 | ##### | 5.3238 | 1.0161E-07 | 3.7406E-07 |
| ENSG00000253315\|LINC01932 | 27.6975 | 4.6064 | ##### | 5.3237 | 1.0170E-07 | 3.7433E-07 |
| ENSG00000116128\|BCL9 | 4565.0443 | 0.6391 | ##### | 5.3229 | 1.0213E-07 | 3.7587E-07 |
| ENSG00000188243\|COMMD6 | 1336.5631 | 0.7465 | ##### | 5.3228 | 1.0220E-07 | 3.7607E-07 |
| ENSG00000289502 | 50.0740 | -3.6143 | ##### | -5.3220 | 1.0264E-07 | 3.7765E-07 |
| ENSG00000010322\|NISCH | 4170.5850 | 0.5811 | ##### | 5.3206 | 1.0342E-07 | 3.8047E-07 |
| ENSG00000187105\|HEATR4 | 74.9486 | 1.9344 | ##### | 5.3205 | 1.0346E-07 | 3.8058E-07 |
| ENSG00000143314\|MRPL24 | 8557.2097 | 0.6455 | ##### | 5.3198 | 1.0390E-07 | 3.8212E-07 |
| ENSG00000258498\|DIO3OS | 234.1529 | -8.4988 | ##### | -5.3195 | 1.0406E-07 | 3.8266E-07 |
| ENSG00000227906\|SNAP25-AS1 | 104.0873 | 1.7409 | ##### | 5.3191 | 1.0427E-07 | 3.8339E-07 |
| ENSG00000092847\|AGO1 | 4407.5793 | -0.6659 | ##### | -5.3188 | 1.0448E-07 | 3.8413E-07 |
| ENSG00000169105\|CHST14 | 1821.7672 | 0.6970 | ##### | 5.3165 | 1.0579E-07 | 3.8891E-07 |
| ENSG00000258038\|LINC02327 | 379.5448 | ####### | ##### | -5.3164 | 1.0583E-07 | 3.8901E-07 |
| ENSG00000069812\|HES2 | 125.5232 | -1.1884 | ##### | -5.3158 | 1.0620E-07 | 3.9031E-07 |

| | | | | | | |
|---|---|---|---|---|---|---|
| ENSG00000186866\|POFUT2 | 4647.5556 | -0.5333 | ##### | -5.3152 | 1.0654E-07 | 3.9153E-07 |
| ENSG00000277879 | 138.9106 | -1.0682 | ##### | -5.3145 | 1.0696E-07 | 3.9299E-07 |
| ENSG00000213096\|ZNF254 | 475.4330 | -0.8854 | ##### | -5.3143 | 1.0705E-07 | 3.9330E-07 |
| ENSG00000036257\|CUL3 | 14073.3636 | 0.4754 | ##### | 5.3132 | 1.0773E-07 | 3.9575E-07 |
| ENSG00000110400\|NECTIN1 | 2770.1803 | 0.6660 | ##### | 5.3131 | 1.0779E-07 | 3.9592E-07 |
| ENSG00000011007\|ELOA | 8042.6188 | 0.5488 | ##### | 5.3116 | 1.0869E-07 | 3.9916E-07 |
| ENSG00000176473\|WDR25 | 465.6927 | 0.7468 | ##### | 5.3111 | 1.0899E-07 | 4.0022E-07 |
| ENSG00000171747\|LGALS4 | 46.3283 | -3.8132 | ##### | -5.3108 | 1.0916E-07 | 4.0080E-07 |
| ENSG00000007923\|DNAJC11 | 9662.4798 | -0.4220 | ##### | -5.3099 | 1.0968E-07 | 4.0266E-07 |
| ENSG00000291129\|PPIEL | 104.0714 | 1.9465 | ##### | 5.3098 | 1.0973E-07 | 4.0278E-07 |
| ENSG00000078246\|TULP3 | 4787.8595 | 0.5857 | ##### | 5.3097 | 1.0981E-07 | 4.0304E-07 |
| ENSG00000206195\|DUXAP8 | 2204.4254 | -0.6705 | ##### | -5.3095 | 1.0995E-07 | 4.0351E-07 |
| ENSG00000142065\|ZFP14 | 362.7110 | -0.8804 | ##### | -5.3092 | 1.1011E-07 | 4.0403E-07 |
| ENSG00000167604\|NFKBID | 505.6319 | 0.8845 | ##### | 5.3090 | 1.1022E-07 | 4.0440E-07 |
| ENSG00000287248 | 83.6459 | 3.4788 | ##### | 5.3086 | 1.1049E-07 | 4.0534E-07 |
| ENSG00000196437\|ZNF569 | 1125.2283 | -0.7195 | ##### | -5.3071 | 1.1141E-07 | 4.0864E-07 |
| ENSG00000066044\|ELAVL1 | 10399.7037 | 0.5292 | ##### | 5.3068 | 1.1154E-07 | 4.0904E-07 |
| ENSG00000175182\|FAM131A | 1021.3446 | -0.6781 | ##### | -5.3068 | 1.1154E-07 | 4.0904E-07 |
| ENSG00000132622\|HSPA12B | 88.1788 | 1.3816 | ##### | 5.3068 | 1.1158E-07 | 4.0912E-07 |
| ENSG00000223802\|CERS1 | 674.1108 | 0.7320 | ##### | 5.3065 | 1.1174E-07 | 4.0968E-07 |
| ENSG00000167986\|DDB1 | 21347.0424 | 0.6286 | ##### | 5.3061 | 1.1201E-07 | 4.1061E-07 |
| ENSG00000155085\|AK9 | 129.4986 | -1.5641 | ##### | -5.3058 | 1.1217E-07 | 4.1112E-07 |
| ENSG00000151623\|NR3C2 | 64.8156 | -8.9204 | ##### | -5.3057 | 1.1224E-07 | 4.1128E-07 |
| ENSG00000281091 | 49.6561 | -1.8124 | ##### | -5.3057 | 1.1223E-07 | 4.1128E-07 |
| ENSG00000130957\|FBP2 | 17.1670 | -2.7582 | ##### | -5.3048 | 1.1283E-07 | 4.1339E-07 |
| ENSG00000204316\|MRPL38 | 7043.9713 | 0.6288 | ##### | 5.3047 | 1.1286E-07 | 4.1346E-07 |
| ENSG00000170396\|ZNF804A | 120.6948 | 5.6644 | ##### | 5.3043 | 1.1309E-07 | 4.1426E-07 |
| ENSG00000105948\|IFT56 | 2202.5289 | 0.6535 | ##### | 5.3039 | 1.1336E-07 | 4.1520E-07 |
| ENSG00000101577\|LPIN2 | 6066.5007 | -0.6374 | ##### | -5.3028 | 1.1402E-07 | 4.1755E-07 |
| ENSG00000172380\|GNG12 | 15142.2922 | -0.6613 | ##### | -5.3025 | 1.1425E-07 | 4.1835E-07 |
| ENSG00000187713\|TMEM203 | 2620.2624 | -0.7143 | ##### | -5.2999 | 1.1584E-07 | 4.2414E-07 |
| ENSG00000236809\|SNX25P1 | 60.1845 | -1.7138 | ##### | -5.2999 | 1.1586E-07 | 4.2415E-07 |
| ENSG00000183979\|NPB | 156.8972 | -1.0713 | ##### | -5.2991 | 1.1640E-07 | 4.2606E-07 |
| ENSG00000235726 | 45.4579 | 6.8633 | ##### | 5.2990 | 1.1645E-07 | 4.2620E-07 |
| ENSG00000166578\|IQCD | 353.0652 | 0.9340 | ##### | 5.2986 | 1.1667E-07 | 4.2694E-07 |
| ENSG00000276023\|DUSP14 | 2485.8436 | 0.6936 | ##### | 5.2986 | 1.1671E-07 | 4.2705E-07 |
| ENSG00000225855\|RUSC1-AS1 | 568.3340 | 1.0576 | ##### | 5.2983 | 1.1687E-07 | 4.2754E-07 |
| ENSG00000127252\|PLAAT1 | 383.7557 | 0.9086 | ##### | 5.2983 | 1.1687E-07 | 4.2754E-07 |
| ENSG00000214293\|APTR | 430.9905 | -0.8111 | ##### | -5.2976 | 1.1736E-07 | 4.2928E-07 |
| ENSG00000185129\|PURA | 2301.9195 | 0.6629 | ##### | 5.2975 | 1.1739E-07 | 4.2933E-07 |
| ENSG00000232811 | 158.4285 | -1.2739 | ##### | -5.2954 | 1.1878E-07 | 4.3437E-07 |
| ENSG00000107147\|KCNT1 | 46.7307 | 6.7400 | ##### | 5.2951 | 1.1897E-07 | 4.3499E-07 |
| ENSG00000224914\|LINC00863 | 380.4136 | -0.8458 | ##### | -5.2940 | 1.1966E-07 | 4.3748E-07 |
| ENSG00000126562\|WNK4 | 189.0361 | -0.9076 | ##### | -5.2936 | 1.1991E-07 | 4.3832E-07 |
| ENSG00000105341\|DMAC2 | 4381.2543 | -0.6559 | ##### | -5.2936 | 1.1994E-07 | 4.3838E-07 |
| ENSG00000259856\|RAB43P1 | 70.0167 | -1.3687 | ##### | -5.2927 | 1.2052E-07 | 4.4039E-07 |
| ENSG00000132846\|ZBED3 | 637.2402 | -0.7663 | ##### | -5.2927 | 1.2051E-07 | 4.4039E-07 |
| ENSG00000226562\|CYP4F26P | 118.7136 | -1.2869 | ##### | -5.2926 | 1.2060E-07 | 4.4063E-07 |
| ENSG00000213030\|CGB8 | 52.7803 | -1.7535 | ##### | -5.2913 | 1.2147E-07 | 4.4376E-07 |
| ENSG00000133606\|MKRN1 | 8130.9035 | 0.5351 | ##### | 5.2905 | 1.2197E-07 | 4.4551E-07 |
| ENSG00000171056\|SOX7 | 509.0391 | 0.8436 | ##### | 5.2889 | 1.2302E-07 | 4.4933E-07 |
| ENSG00000083807\|SLC27A5 | 567.6487 | -0.8459 | ##### | -5.2888 | 1.2315E-07 | 4.4972E-07 |
| ENSG00000258749 | 99.0827 | -1.2864 | ##### | -5.2885 | 1.2330E-07 | 4.5023E-07 |
| ENSG00000176273\|SLC35G1 | 1500.1037 | -0.6996 | ##### | -5.2881 | 1.2360E-07 | 4.5128E-07 |
| ENSG00000175550\|DRAP1 | 13714.1558 | 0.6433 | ##### | 5.2878 | 1.2378E-07 | 4.5185E-07 |
| ENSG00000197813 | 56.6806 | -3.7040 | ##### | -5.2878 | 1.2380E-07 | 4.5190E-07 |
| ENSG00000078795\|PKD2L2 | 58.4211 | 1.5499 | ##### | 5.2869 | 1.2440E-07 | 4.5402E-07 |
| ENSG00000266904\|LINC00663 | 59.5507 | -1.7502 | ##### | -5.2857 | 1.2526E-07 | 4.5709E-07 |
| ENSG00000107263\|RAPGEF1 | 7903.0842 | -0.6340 | ##### | -5.2852 | 1.2553E-07 | 4.5804E-07 |
| ENSG00000008018\|PSMB1 | 11257.8513 | -0.5044 | ##### | -5.2850 | 1.2568E-07 | 4.5854E-07 |
| ENSG00000226650\|KIF4B | 38.7845 | 1.6495 | ##### | 5.2849 | 1.2575E-07 | 4.5871E-07 |
| ENSG00000114019\|AMOTL2 | 25643.3980 | 0.4771 | ##### | 5.2845 | 1.2602E-07 | 4.5964E-07 |
| ENSG00000204351\|SKIC2 | 6705.0911 | 0.5654 | ##### | 5.2826 | 1.2733E-07 | 4.6439E-07 |

| | | | | | | |
|---|---|---|---|---|---|---|
| ENSG00000090020\|SLC9A1 | 3359.7517 | 0.5205 | ##### | 5.2825 | 1.2745E-07 | 4.6474E-07 |
| ENSG00000131966\|ACTR10 | 4891.4715 | 0.5789 | ##### | 5.2824 | 1.2749E-07 | 4.6482E-07 |
| ENSG00000172986\|GXYLT2 | 115.6824 | -1.2133 | ##### | -5.2819 | 1.2786E-07 | 4.6612E-07 |
| ENSG00000118482\|PHF3 | 5355.7213 | 0.6485 | ##### | 5.2799 | 1.2928E-07 | 4.7126E-07 |
| ENSG00000198858\|R3HDM4 | 4640.1281 | 0.6570 | ##### | 5.2796 | 1.2945E-07 | 4.7181E-07 |
| ENSG00000136908\|DPM2 | 3875.1201 | -0.6392 | ##### | -5.2784 | 1.3031E-07 | 4.7488E-07 |
| ENSG00000289840 | 77.5963 | 8.3931 | ##### | 5.2776 | 1.3090E-07 | 4.7699E-07 |
| ENSG00000162384\|CZIB | 2002.9309 | -0.6952 | ##### | -5.2775 | 1.3097E-07 | 4.7717E-07 |
| ENSG00000187772\|LIN28B | 96.9392 | 1.2081 | ##### | 5.2768 | 1.3146E-07 | 4.7890E-07 |
| ENSG00000009335\|UBE3C | 17936.2585 | 0.5023 | ##### | 5.2761 | 1.3195E-07 | 4.8061E-07 |
| ENSG00000197417\|SHPK | 1459.2498 | 0.7078 | ##### | 5.2758 | 1.3214E-07 | 4.8128E-07 |
| ENSG00000066422\|ZBTB11 | 5036.9305 | 0.6500 | ##### | 5.2758 | 1.3217E-07 | 4.8132E-07 |
| ENSG00000134262\|AP4B1 | 3114.3569 | 0.6666 | ##### | 5.2757 | 1.3223E-07 | 4.8146E-07 |
| ENSG00000111249\|CUX2 | 89.2094 | 7.9329 | ##### | 5.2753 | 1.3253E-07 | 4.8252E-07 |
| ENSG00000103023\|PRSS54 | 33.9969 | -2.2441 | ##### | -5.2751 | 1.3271E-07 | 4.8310E-07 |
| ENSG00000250536\|ABHD17AP3 | 17.9167 | 6.1128 | ##### | 5.2746 | 1.3304E-07 | 4.8425E-07 |
| ENSG00000188176\|SMTNL2 | 83.9502 | 7.9822 | ##### | 5.2741 | 1.3338E-07 | 4.8541E-07 |
| ENSG00000173114\|LRRN3 | 58.7045 | 8.4109 | ##### | 5.2730 | 1.3424E-07 | 4.8850E-07 |
| ENSG00000099783\|HNRNPM | 32713.3098 | 0.5412 | ##### | 5.2729 | 1.3429E-07 | 4.8863E-07 |
| ENSG00000101079\|NDRG3 | 5524.7938 | 0.6564 | ##### | 5.2712 | 1.3553E-07 | 4.9308E-07 |
| ENSG00000213782\|DDX47 | 13887.8796 | 0.5737 | ##### | 5.2686 | 1.3751E-07 | 5.0019E-07 |
| ENSG00000285791 | 201.8320 | 2.5579 | ##### | 5.2677 | 1.3814E-07 | 5.0246E-07 |
| ENSG00000079931\|MOXD1 | 212.5675 | -6.8050 | ##### | -5.2677 | 1.3818E-07 | 5.0251E-07 |
| ENSG00000101203\|COL20A1 | 87.8137 | 1.4571 | ##### | 5.2670 | 1.3867E-07 | 5.0423E-07 |
| ENSG00000225265\|TAF1A-AS1 | 88.9448 | -1.4720 | ##### | -5.2650 | 1.4018E-07 | 5.0967E-07 |
| ENSG00000175166\|PSMD2 | 29701.6867 | -0.5455 | ##### | -5.2628 | 1.4189E-07 | 5.1581E-07 |
| ENSG00000198934\|MAGEE1 | 305.6420 | -0.9577 | ##### | -5.2625 | 1.4211E-07 | 5.1658E-07 |
| ENSG00000163357\|DCST1 | 88.9938 | 1.4030 | ##### | 5.2615 | 1.4291E-07 | 5.1942E-07 |
| ENSG00000175575\|PAAF1 | 1539.1812 | 0.7148 | ##### | 5.2612 | 1.4314E-07 | 5.2019E-07 |
| ENSG00000197162\|ZNF785 | 585.0988 | 0.9852 | ##### | 5.2600 | 1.4402E-07 | 5.2331E-07 |
| ENSG00000172602\|RND1 | 136.9298 | -1.1158 | ##### | -5.2577 | 1.4591E-07 | 5.3009E-07 |
| ENSG00000293037 | 31.4606 | -4.1259 | ##### | -5.2576 | 1.4592E-07 | 5.3009E-07 |
| ENSG00000166317\|SYNPO2L | 54.4752 | 6.4026 | ##### | 5.2575 | 1.4603E-07 | 5.3044E-07 |
| ENSG00000290760\|UBE2Q2P13 | 56.3265 | 1.6161 | ##### | 5.2552 | 1.4787E-07 | 5.3705E-07 |
| ENSG00000171444\|MCC | 378.0505 | -0.7938 | ##### | -5.2542 | 1.4864E-07 | 5.3970E-07 |
| ENSG00000081842\|PCDHA6 | 348.0470 | ####### | ##### | -5.2543 | 1.4862E-07 | 5.3970E-07 |
| ENSG00000141556\|TBCD | 9341.7940 | 0.5427 | ##### | 5.2538 | 1.4902E-07 | 5.4103E-07 |
| ENSG00000100811\|YY1 | 12554.4832 | 0.6049 | ##### | 5.2537 | 1.4904E-07 | 5.4104E-07 |
| ENSG00000148848\|ADAM12 | 616.5383 | -6.5748 | ##### | -5.2529 | 1.4969E-07 | 5.4332E-07 |
| ENSG00000161021\|MAML1 | 4152.0238 | 0.6373 | ##### | 5.2529 | 1.4974E-07 | 5.4344E-07 |
| ENSG00000099814\|CEP170B | 4597.2575 | -0.6283 | ##### | -5.2527 | 1.4991E-07 | 5.4399E-07 |
| ENSG00000237437\|ASS1P12 | 40.7375 | 2.3757 | ##### | 5.2526 | 1.4998E-07 | 5.4417E-07 |
| ENSG00000044459\|CNTLN | 619.4439 | -0.7401 | ##### | -5.2514 | 1.5097E-07 | 5.4771E-07 |
| ENSG00000038219\|BOD1L1 | 4652.0643 | 0.6597 | ##### | 5.2503 | 1.5183E-07 | 5.5076E-07 |
| ENSG00000268996\|MAN1B1-DT | 382.8225 | -0.8697 | ##### | -5.2489 | 1.5298E-07 | 5.5487E-07 |
| ENSG00000161091\|MFSD12 | 14357.2378 | 0.6206 | ##### | 5.2485 | 1.5337E-07 | 5.5616E-07 |
| ENSG00000170185\|USP38 | 2043.6885 | -0.6707 | ##### | -5.2485 | 1.5338E-07 | 5.5616E-07 |
| ENSG00000136720\|HS6ST1 | 3034.7646 | -0.6641 | ##### | -5.2479 | 1.5388E-07 | 5.5792E-07 |
| ENSG00000183914\|DNAH2 | 81.8452 | 7.8568 | ##### | 5.2471 | 1.5454E-07 | 5.6024E-07 |
| ENSG00000058729\|RIOK2 | 2922.5435 | 0.4965 | ##### | 5.2461 | 1.5534E-07 | 5.6308E-07 |
| ENSG00000170178\|HOXD12 | 87.0728 | 7.9495 | ##### | 5.2461 | 1.5538E-07 | 5.6317E-07 |
| ENSG00000057608\|GDI2 | 17102.1045 | -0.2681 | ##### | -5.2459 | 1.5552E-07 | 5.6360E-07 |
| ENSG00000215105\|TTC3P1 | 159.5958 | -0.9688 | ##### | -5.2445 | 1.5670E-07 | 5.6782E-07 |
| ENSG00000134352\|IL6ST | 32788.5203 | 0.6343 | ##### | 5.2438 | 1.5730E-07 | 5.6990E-07 |
| ENSG00000156876\|SASS6 | 1200.1212 | -0.7072 | ##### | -5.2433 | 1.5777E-07 | 5.7155E-07 |
| ENSG00000163501\|IHH | 38.3385 | -2.1194 | ##### | -5.2432 | 1.5783E-07 | 5.7170E-07 |
| ENSG00000124557\|BTN1A1 | 32.0614 | 5.6865 | ##### | 5.2430 | 1.5801E-07 | 5.7227E-07 |
| ENSG00000129071\|MBD4 | 4020.9962 | 0.6347 | ##### | 5.2427 | 1.5820E-07 | 5.7290E-07 |
| ENSG00000169126\|ODAD2 | 151.9916 | 1.1360 | ##### | 5.2416 | 1.5916E-07 | 5.7629E-07 |
| ENSG00000080007\|DDX43 | 45.0055 | 6.8598 | ##### | 5.2412 | 1.5954E-07 | 5.7761E-07 |
| ENSG00000145022\|TCTA | 1889.0045 | -0.6851 | ##### | -5.2406 | 1.6007E-07 | 5.7945E-07 |
| ENSG00000171497\|PPID | 4232.7464 | -0.6765 | ##### | -5.2397 | 1.6085E-07 | 5.8222E-07 |
| ENSG00000293507\|FBXL9P | 115.5006 | -1.1723 | ##### | -5.2395 | 1.6101E-07 | 5.8270E-07 |

| | | | | | | |
|---|---|---|---|---|---|---|
| ENSG00000068323\|TFE3 | 7440.8856 | -0.5606 | ##### | -5.2394 | 1.6107E-07 | 5.8287E-07 |
| ENSG00000165572\|KBTBD6 | 2113.6269 | 0.6438 | ##### | 5.2393 | 1.6119E-07 | 5.8321E-07 |
| ENSG00000157152\|SYN2 | 75.3252 | 1.6910 | ##### | 5.2389 | 1.6156E-07 | 5.8451E-07 |
| ENSG00000100330\|MTMR3 | 3521.4524 | 0.8994 | ##### | 5.2381 | 1.6225E-07 | 5.8692E-07 |
| ENSG00000168393\|DTYMK | 7119.0934 | 0.6462 | ##### | 5.2378 | 1.6253E-07 | 5.8786E-07 |
| ENSG00000288649\|ACTL10 | 32.1637 | 4.3821 | ##### | 5.2376 | 1.6266E-07 | 5.8825E-07 |
| ENSG00000204315\|FKBPL | 559.6624 | 0.8035 | ##### | 5.2373 | 1.6297E-07 | 5.8932E-07 |
| ENSG00000105605\|CACNG7 | 589.3180 | 0.8096 | ##### | 5.2367 | 1.6351E-07 | 5.9118E-07 |
| ENSG00000239620\|PRR20G | 32.5382 | 7.0698 | ##### | 5.2347 | 1.6525E-07 | 5.9740E-07 |
| ENSG00000162188\|GNG3 | 33.9855 | 2.0485 | ##### | 5.2343 | 1.6562E-07 | 5.9867E-07 |
| ENSG00000188234\|AGAP4 | 496.4829 | 0.7956 | ##### | 5.2341 | 1.6582E-07 | 5.9932E-07 |
| ENSG00000268858 | 1175.6478 | 0.7032 | ##### | 5.2328 | 1.6695E-07 | 6.0335E-07 |
| ENSG00000258602\|LINC01629 | 88.2094 | 7.8112 | ##### | 5.2316 | 1.6804E-07 | 6.0720E-07 |
| ENSG00000099139\|PCSK5 | 96.2727 | -1.3486 | ##### | -5.2316 | 1.6807E-07 | 6.0724E-07 |
| ENSG00000180398\|MCFD2 | 19933.8294 | -0.6220 | ##### | -5.2307 | 1.6886E-07 | 6.1000E-07 |
| ENSG00000281332\|LINC00997 | 174.4382 | -0.9195 | ##### | -5.2294 | 1.7008E-07 | 6.1435E-07 |
| ENSG00000122861\|PLAU | 12345.8519 | 0.6404 | ##### | 5.2291 | 1.7029E-07 | 6.1505E-07 |
| ENSG00000214295\|FOXO1B | 43.9470 | 2.6567 | ##### | 5.2284 | 1.7100E-07 | 6.1752E-07 |
| ENSG00000180264\|ADGRD2 | 44.9674 | 1.7405 | ##### | 5.2278 | 1.7158E-07 | 6.1953E-07 |
| ENSG00000278864 | 92.5255 | -1.3619 | ##### | -5.2270 | 1.7230E-07 | 6.2207E-07 |
| ENSG00000066322\|ELOVL1 | 8920.1905 | -0.6544 | ##### | -5.2269 | 1.7242E-07 | 6.2243E-07 |
| ENSG00000127423\|AUNIP | 1123.0869 | 0.7134 | ##### | 5.2268 | 1.7247E-07 | 6.2255E-07 |
| ENSG00000162148\|SAXO4 | 27.3801 | -1.9053 | ##### | -5.2262 | 1.7300E-07 | 6.2436E-07 |
| ENSG00000048544\|MRPS10 | 4987.0632 | -0.6512 | ##### | -5.2251 | 1.7402E-07 | 6.2797E-07 |
| ENSG00000237187\|NR2F1-AS1 | 692.6443 | -0.7641 | ##### | -5.2251 | 1.7407E-07 | 6.2809E-07 |
| ENSG00000283498\|MIR1244-2 | 448.8718 | 0.7669 | ##### | 5.2247 | 1.7444E-07 | 6.2932E-07 |
| ENSG00000110799\|VWF | 157.7308 | 7.8579 | ##### | 5.2238 | 1.7525E-07 | 6.3218E-07 |
| ENSG00000107864\|CPEB3 | 281.5306 | 0.9533 | ##### | 5.2230 | 1.7608E-07 | 6.3509E-07 |
| ENSG00000115306\|SPTBN1 | 25521.2832 | -0.6170 | ##### | -5.2228 | 1.7623E-07 | 6.3556E-07 |
| ENSG00000226648\|PLCG1-AS1 | 42.9957 | 2.4511 | ##### | 5.2226 | 1.7643E-07 | 6.3620E-07 |
| ENSG00000160014\|CALM3 | 25086.7396 | -0.5854 | ##### | -5.2216 | 1.7735E-07 | 6.3945E-07 |
| ENSG00000140600\|SH3GL3 | 37.3295 | 4.5623 | ##### | 5.2213 | 1.7769E-07 | 6.4058E-07 |
| ENSG00000102878\|HSF4 | 674.2932 | -0.7412 | ##### | -5.2192 | 1.7973E-07 | 6.4789E-07 |
| ENSG00000100994\|PYGB | 16259.1485 | -0.6003 | ##### | -5.2181 | 1.8079E-07 | 6.5161E-07 |
| ENSG00000289353 | 650.8809 | -0.7680 | ##### | -5.2178 | 1.8110E-07 | 6.5265E-07 |
| ENSG00000267007 | 91.6811 | 1.2836 | ##### | 5.2176 | 1.8127E-07 | 6.5320E-07 |
| ENSG00000162702\|ZNF281 | 7952.7680 | -0.6054 | ##### | -5.2168 | 1.8203E-07 | 6.5586E-07 |
| ENSG00000197958\|RPL12 | 52633.6267 | 0.5188 | ##### | 5.2162 | 1.8263E-07 | 6.5792E-07 |
| ENSG00000280239 | 158.6066 | 1.1014 | ##### | 5.2160 | 1.8286E-07 | 6.5868E-07 |
| ENSG00000286835 | 21.3997 | 3.5967 | ##### | 5.2159 | 1.8297E-07 | 6.5898E-07 |
| ENSG00000159023\|EPB41 | 2071.2028 | -0.7190 | ##### | -5.2150 | 1.8385E-07 | 6.6207E-07 |
| ENSG00000242125\|SNHG3 | 2279.8324 | 0.6791 | ##### | 5.2142 | 1.8460E-07 | 6.6471E-07 |
| ENSG00000165061\|ZMAT4 | 140.4335 | -7.1486 | ##### | -5.2134 | 1.8542E-07 | 6.6756E-07 |
| ENSG00000129535\|NRL | 71.3753 | 1.4274 | ##### | 5.2119 | 1.8696E-07 | 6.7303E-07 |
| ENSG00000272071 | 55.1581 | -7.9434 | ##### | -5.2114 | 1.8746E-07 | 6.7476E-07 |
| ENSG00000244239\|ICA1-AS1 | 15.9424 | -3.8200 | ##### | -5.2110 | 1.8779E-07 | 6.7588E-07 |
| ENSG00000198563\|DDX39B | 22444.0274 | 0.6092 | ##### | 5.2104 | 1.8845E-07 | 6.7815E-07 |
| ENSG00000100982\|PCIF1 | 3773.0781 | 0.6481 | ##### | 5.2102 | 1.8863E-07 | 6.7874E-07 |
| ENSG00000276570 | 827.2498 | 0.7173 | ##### | 5.2101 | 1.8878E-07 | 6.7919E-07 |
| ENSG00000204531\|POU5F1 | 153.1866 | -4.7488 | ##### | -5.2094 | 1.8947E-07 | 6.8157E-07 |
| ENSG00000259868 | 70.6829 | 7.9358 | ##### | 5.2090 | 1.8984E-07 | 6.8282E-07 |
| ENSG00000229020\|AKR7A2P1 | 21.2352 | 6.5304 | ##### | 5.2088 | 1.9007E-07 | 6.8359E-07 |
| ENSG00000254087\|LYN | 2078.7078 | -0.6778 | ##### | -5.2080 | 1.9086E-07 | 6.8634E-07 |
| ENSG00000183696\|UPP1 | 6370.7824 | -0.6433 | ##### | -5.2073 | 1.9158E-07 | 6.8885E-07 |
| ENSG00000227268\|KLLN | 189.9411 | 1.0569 | ##### | 5.2072 | 1.9173E-07 | 6.8929E-07 |
| ENSG00000154764\|WNT7A | 217.7873 | ####### | ##### | -5.2049 | 1.9407E-07 | 6.9764E-07 |
| ENSG00000257704\|INAFM1 | 627.3031 | -0.8055 | ##### | -5.2043 | 1.9471E-07 | 6.9986E-07 |
| ENSG00000141086\|CTRL | 108.5291 | 1.5400 | ##### | 5.2032 | 1.9590E-07 | 7.0405E-07 |
| ENSG00000267265\|GP6-AS1 | 160.2844 | -1.0840 | ##### | -5.2032 | 1.9593E-07 | 7.0406E-07 |
| ENSG00000114942\|EEF1B2 | 24314.2003 | 0.6279 | ##### | 5.2026 | 1.9657E-07 | 7.0627E-07 |
| ENSG00000287707 | 107.6349 | 1.6287 | ##### | 5.2023 | 1.9689E-07 | 7.0733E-07 |
| ENSG00000223799\|IL10RB-DT | 903.3650 | -0.7869 | ##### | -5.2012 | 1.9804E-07 | 7.1137E-07 |
| ENSG00000168924\|LETM1 | 7720.0209 | 0.6037 | ##### | 5.2011 | 1.9809E-07 | 7.1149E-07 |

| | | | | | | |
|---|---|---|---|---|---|---|
| ENSG00000185088\|RPS27L | 3520.5763 | 0.6416 | ##### | 5.2006 | 1.9869E-07 | 7.1356E-07 |
| ENSG00000187961\|KLHL17 | 2734.3438 | 0.6538 | ##### | 5.1997 | 1.9964E-07 | 7.1686E-07 |
| ENSG00000204439\|C6orf47 | 1906.6203 | 0.6404 | ##### | 5.1996 | 1.9974E-07 | 7.1715E-07 |
| ENSG00000176978\|DPP7 | 6034.3938 | 0.6288 | ##### | 5.1970 | 2.0258E-07 | 7.2726E-07 |
| ENSG00000250251\|PKD1P6 | 1052.8365 | 0.8987 | ##### | 5.1946 | 2.0516E-07 | 7.3624E-07 |
| ENSG00000197496\|SLC2A10 | 336.2364 | -0.9536 | ##### | -5.1946 | 2.0512E-07 | 7.3624E-07 |
| ENSG00000248690\|HAS2-AS1 | 263.8682 | 0.9146 | ##### | 5.1946 | 2.0516E-07 | 7.3624E-07 |
| ENSG00000118762\|PKD2 | 3938.7966 | -0.6378 | ##### | -5.1941 | 2.0570E-07 | 7.3810E-07 |
| ENSG00000255154\|HTD2 | 220.5979 | 1.5780 | ##### | 5.1939 | 2.0589E-07 | 7.3868E-07 |
| ENSG00000229132\|EIF4A1P10 | 872.3146 | 0.7106 | ##### | 5.1934 | 2.0649E-07 | 7.4075E-07 |
| ENSG00000088205\|DDX18 | 15444.0949 | -0.5586 | ##### | -5.1929 | 2.0699E-07 | 7.4248E-07 |
| ENSG00000289924 | 65.6129 | 3.4116 | ##### | 5.1928 | 2.0720E-07 | 7.4313E-07 |
| ENSG00000227615 | 301.9907 | 0.9161 | ##### | 5.1926 | 2.0737E-07 | 7.4366E-07 |
| ENSG00000224950 | 60.4057 | 1.8685 | ##### | 5.1924 | 2.0764E-07 | 7.4451E-07 |
| ENSG00000082074\|FYB1 | 36.9077 | -3.2774 | ##### | -5.1923 | 2.0769E-07 | 7.4464E-07 |
| ENSG00000258289\|CHURC1 | 5677.4680 | 0.6350 | ##### | 5.1922 | 2.0780E-07 | 7.4494E-07 |
| ENSG00000160183\|TMPRSS3 | 452.4213 | ####### | ##### | -5.1920 | 2.0801E-07 | 7.4557E-07 |
| ENSG00000126934\|MAP2K2 | 18212.9943 | 0.5032 | ##### | 5.1917 | 2.0837E-07 | 7.4680E-07 |
| ENSG00000121104\|FAM117A | 731.0161 | 0.7483 | ##### | 5.1915 | 2.0864E-07 | 7.4765E-07 |
| ENSG00000128422\|KRT17 | 31.4181 | -2.2244 | ##### | -5.1909 | 2.0926E-07 | 7.4978E-07 |
| ENSG00000242520\|MAGEA5P | 34.3771 | -6.3387 | ##### | -5.1900 | 2.1029E-07 | 7.5339E-07 |
| ENSG00000172318\|B3GALT1 | 93.5663 | 7.8358 | ##### | 5.1898 | 2.1052E-07 | 7.5414E-07 |
| ENSG00000163950\|SLBP | 6273.7914 | 0.6384 | ##### | 5.1893 | 2.1106E-07 | 7.5598E-07 |
| ENSG00000136504\|KAT7 | 4514.6206 | 0.5953 | ##### | 5.1885 | 2.1202E-07 | 7.5931E-07 |
| ENSG00000125779\|PANK2 | 4210.1634 | 0.6067 | ##### | 5.1882 | 2.1234E-07 | 7.6038E-07 |
| ENSG00000179029\|TMEM107 | 503.4166 | -0.7586 | ##### | -5.1882 | 2.1238E-07 | 7.6044E-07 |
| ENSG00000259083\|PNN-AS1 | 22.7891 | 2.1762 | ##### | 5.1867 | 2.1410E-07 | 7.6650E-07 |
| ENSG00000068697\|LAPTM4A | 24997.2396 | 0.4029 | ##### | 5.1864 | 2.1436E-07 | 7.6735E-07 |
| ENSG00000184677\|ZBTB40 | 3370.3530 | 0.6455 | ##### | 5.1861 | 2.1479E-07 | 7.6878E-07 |
| ENSG00000197744\|PTMAP2 | 62.6217 | 1.4608 | ##### | 5.1855 | 2.1549E-07 | 7.7120E-07 |
| ENSG00000175104\|TRAF6 | 1035.4418 | 0.6946 | ##### | 5.1847 | 2.1632E-07 | 7.7406E-07 |
| ENSG00000285608 | 1959.5591 | 0.7347 | ##### | 5.1845 | 2.1659E-07 | 7.7495E-07 |
| ENSG00000126267\|COX6B1 | 8570.7143 | -0.6271 | ##### | -5.1824 | 2.1901E-07 | 7.8352E-07 |
| ENSG00000130227\|XPO7 | 8844.1859 | 0.6402 | ##### | 5.1822 | 2.1929E-07 | 7.8442E-07 |
| ENSG00000183087\|GAS6 | 979.5464 | -0.6978 | ##### | -5.1821 | 2.1936E-07 | 7.8457E-07 |
| ENSG00000255641 | 522.3093 | 7.9233 | ##### | 5.1819 | 2.1968E-07 | 7.8563E-07 |
| ENSG00000235560\|ZNF747-DT | 115.0163 | -1.1449 | ##### | -5.1817 | 2.1986E-07 | 7.8618E-07 |
| ENSG00000185203\|WASIR1 | 83.7550 | 7.7231 | ##### | 5.1803 | 2.2152E-07 | 7.9200E-07 |
| ENSG00000285889 | 118.6885 | 8.0930 | ##### | 5.1802 | 2.2160E-07 | 7.9221E-07 |
| ENSG00000115484\|CCT4 | 55160.8697 | 0.5979 | ##### | 5.1789 | 2.2322E-07 | 7.9790E-07 |
| ENSG00000075426\|FOSL2 | 16189.2947 | -0.6209 | ##### | -5.1787 | 2.2343E-07 | 7.9854E-07 |
| ENSG00000259642\|ST20-AS1 | 563.4108 | -0.8010 | ##### | -5.1780 | 2.2432E-07 | 8.0164E-07 |
| ENSG00000162631\|NTNG1 | 192.8690 | ####### | ##### | -5.1772 | 2.2519E-07 | 8.0464E-07 |
| ENSG00000167395\|ZNF646 | 3010.6517 | 0.6311 | ##### | 5.1769 | 2.2560E-07 | 8.0603E-07 |
| ENSG00000109790\|KLHL5 | 13117.9476 | 0.5258 | ##### | 5.1767 | 2.2586E-07 | 8.0685E-07 |
| ENSG00000258378 | 19.6266 | 3.8047 | ##### | 5.1747 | 2.2826E-07 | 8.1533E-07 |
| ENSG00000279159 | 4014.5086 | 0.7291 | ##### | 5.1743 | 2.2875E-07 | 8.1699E-07 |
| ENSG00000198225\|FKBP1C | 404.2855 | 0.8592 | ##### | 5.1737 | 2.2954E-07 | 8.1972E-07 |
| ENSG00000248206 | 175.0262 | -9.8598 | ##### | -5.1730 | 2.3040E-07 | 8.2270E-07 |
| ENSG00000157259\|GATAD1 | 3221.2544 | -0.6248 | ##### | -5.1725 | 2.3098E-07 | 8.2465E-07 |
| ENSG00000235296 | 78.7298 | 7.9096 | ##### | 5.1724 | 2.3111E-07 | 8.2502E-07 |
| ENSG00000134046\|MBD2 | 5346.7828 | -0.6440 | ##### | -5.1714 | 2.3241E-07 | 8.2955E-07 |
| ENSG00000203875\|SNHG5 | 10018.3696 | 0.6732 | ##### | 5.1713 | 2.3248E-07 | 8.2970E-07 |
| ENSG00000213285\|RPL23AP45 | 19.2465 | 2.1436 | ##### | 5.1712 | 2.3258E-07 | 8.2997E-07 |
| ENSG00000184611\|KCNH7 | 58.2639 | -3.8869 | ##### | -5.1710 | 2.3282E-07 | 8.3073E-07 |
| ENSG00000188517\|COL25A1 | 45.6981 | 6.4231 | ##### | 5.1702 | 2.3383E-07 | 8.3424E-07 |
| ENSG00000138785\|INTS12 | 2004.6559 | -0.6846 | ##### | -5.1685 | 2.3594E-07 | 8.4167E-07 |
| ENSG00000237877\|LINC01473 | 59.4602 | -1.4992 | ##### | -5.1678 | 2.3688E-07 | 8.4491E-07 |
| ENSG00000154928\|EPHB1 | 86.5348 | 7.7641 | ##### | 5.1667 | 2.3829E-07 | 8.4984E-07 |
| ENSG00000286319 | 19.1737 | 2.2470 | ##### | 5.1658 | 2.3941E-07 | 8.5374E-07 |
| ENSG00000226193 | 20.2656 | -2.4063 | ##### | -5.1655 | 2.3974E-07 | 8.5480E-07 |
| ENSG00000163655\|GMPS | 10902.2294 | 0.5624 | ##### | 5.1653 | 2.4009E-07 | 8.5596E-07 |
| ENSG00000177462\|OR2T8 | 37.4399 | 6.5038 | ##### | 5.1650 | 2.4041E-07 | 8.5700E-07 |

| | | | | | | |
|---|---|---|---|---|---|---|
| ENSG00000204434\|POTEKP | 12.8564 | -2.5737 | ##### | -5.1649 | 2.4061E-07 | 8.5759E-07 |
| ENSG00000147654\|EBAG9 | 1752.4517 | -0.6508 | ##### | -5.1643 | 2.4132E-07 | 8.6001E-07 |
| ENSG00000160948\|VPS28 | 4787.3163 | -0.6243 | ##### | -5.1642 | 2.4143E-07 | 8.6033E-07 |
| ENSG00000152409\|JMY | 897.3953 | 0.6982 | ##### | 5.1627 | 2.4338E-07 | 8.6715E-07 |
| ENSG00000205302\|SNX2 | 3611.6618 | 0.6480 | ##### | 5.1626 | 2.4360E-07 | 8.6783E-07 |
| ENSG00000244716\|RPL17P7 | 1081.9147 | -0.6785 | ##### | -5.1616 | 2.4483E-07 | 8.7212E-07 |
| ENSG00000171791\|BCL2 | 574.2879 | -0.8049 | ##### | -5.1611 | 2.4548E-07 | 8.7434E-07 |
| ENSG00000111554\|MDM1 | 1090.9915 | 0.6502 | ##### | 5.1597 | 2.4740E-07 | 8.8106E-07 |
| ENSG00000269386\|RAB11B-AS1 | 62.0060 | -1.4327 | ##### | -5.1589 | 2.4840E-07 | 8.8452E-07 |
| ENSG00000269405\|NXF2 | 525.0750 | -9.6861 | ##### | -5.1583 | 2.4923E-07 | 8.8738E-07 |
| ENSG00000198853\|RUSC2 | 8517.6526 | 0.4376 | ##### | 5.1580 | 2.4957E-07 | 8.8849E-07 |
| ENSG00000251138\|LINC02882 | 95.5550 | -9.0052 | ##### | -5.1574 | 2.5042E-07 | 8.9141E-07 |
| ENSG00000130695\|CEP85 | 3941.3063 | 0.5449 | ##### | 5.1553 | 2.5319E-07 | 9.0115E-07 |
| ENSG00000163006\|CCDC138 | 1281.5168 | -0.6993 | ##### | -5.1551 | 2.5347E-07 | 9.0205E-07 |
| ENSG00000174953\|DHX36 | 8873.0117 | 0.5693 | ##### | 5.1523 | 2.5725E-07 | 9.1538E-07 |
| ENSG00000124688\|MAD2L1BP | 1788.3464 | 0.6568 | ##### | 5.1521 | 2.5763E-07 | 9.1661E-07 |
| ENSG00000134001\|EIF2S1 | 16541.0081 | 0.4961 | ##### | 5.1520 | 2.5779E-07 | 9.1708E-07 |
| ENSG00000215475\|SIAH3 | 50.5327 | 8.1531 | ##### | 5.1513 | 2.5875E-07 | 9.2037E-07 |
| ENSG00000154678\|PDE1C | 139.9980 | -6.2815 | ##### | -5.1509 | 2.5924E-07 | 9.2203E-07 |
| ENSG00000188636\|RTL6 | 4249.5469 | -0.6041 | ##### | -5.1507 | 2.5956E-07 | 9.2306E-07 |
| ENSG00000177485\|ZBTB33 | 3428.0396 | 0.6591 | ##### | 5.1496 | 2.6100E-07 | 9.2807E-07 |
| ENSG00000204677\|FAM153CP | 32.3783 | 2.2909 | ##### | 5.1496 | 2.6108E-07 | 9.2824E-07 |
| ENSG00000164089\|ETNPPL | 89.2791 | 7.6441 | ##### | 5.1491 | 2.6178E-07 | 9.3061E-07 |
| ENSG00000137216\|TMEM63B | 4170.5011 | -0.6171 | ##### | -5.1484 | 2.6265E-07 | 9.3359E-07 |
| ENSG00000105676\|ARMC6 | 4249.1004 | 0.6279 | ##### | 5.1477 | 2.6368E-07 | 9.3714E-07 |
| ENSG00000174744\|BRMS1 | 4034.5506 | 0.6624 | ##### | 5.1466 | 2.6529E-07 | 9.4274E-07 |
| ENSG00000146205\|ANO7 | 514.7839 | 0.7951 | ##### | 5.1465 | 2.6544E-07 | 9.4319E-07 |
| ENSG00000250302\|LINC01618 | 33.8396 | 3.9943 | ##### | 5.1463 | 2.6573E-07 | 9.4410E-07 |
| ENSG00000130703\|OSBPL2 | 2955.0219 | 0.6211 | ##### | 5.1455 | 2.6685E-07 | 9.4796E-07 |
| ENSG00000207547\|MIR25 | 33.8906 | 1.6727 | ##### | 5.1448 | 2.6774E-07 | 9.5099E-07 |
| ENSG00000140691\|ARMC5 | 1582.1527 | 0.6470 | ##### | 5.1443 | 2.6855E-07 | 9.5376E-07 |
| ENSG00000052723\|SIKE1 | 5456.9761 | 0.6167 | ##### | 5.1424 | 2.7123E-07 | 9.6305E-07 |
| ENSG00000167850\|CD300C | 33.1325 | -6.8595 | ##### | -5.1424 | 2.7122E-07 | 9.6305E-07 |
| ENSG00000065978\|YBX1 | 98379.9303 | 0.4957 | ##### | 5.1424 | 2.7130E-07 | 9.6321E-07 |
| ENSG00000290285 | 136.8798 | 1.0263 | ##### | 5.1421 | 2.7175E-07 | 9.6468E-07 |
| ENSG00000133612\|AGAP3 | 5817.6122 | 0.4690 | ##### | 5.1411 | 2.7309E-07 | 9.6931E-07 |
| ENSG00000136840\|ST6GALNAC4 | 3159.0862 | -0.6492 | ##### | -5.1410 | 2.7335E-07 | 9.7013E-07 |
| ENSG00000234155\|LINC02535 | 30.3504 | -2.2272 | ##### | -5.1404 | 2.7421E-07 | 9.7308E-07 |
| ENSG00000081913\|PHLPP1 | 1179.8865 | -0.6946 | ##### | -5.1400 | 2.7467E-07 | 9.7458E-07 |
| ENSG00000186814\|ZSCAN30 | 1034.3049 | -0.6727 | ##### | -5.1397 | 2.7515E-07 | 9.7617E-07 |
| ENSG00000135624\|CCT7 | 44544.9306 | 0.5276 | ##### | 5.1395 | 2.7544E-07 | 9.7709E-07 |
| ENSG00000289198 | 20.7275 | -2.8176 | ##### | -5.1389 | 2.7628E-07 | 9.7994E-07 |
| ENSG00000241764\|CBLL1-AS1 | 234.2540 | 0.9247 | ##### | 5.1389 | 2.7641E-07 | 9.8030E-07 |
| ENSG00000172965\|MIR4435-2HG | 3633.0526 | -0.5935 | ##### | -5.1386 | 2.7686E-07 | 9.8177E-07 |
| ENSG00000272092 | 40.4844 | -1.6662 | ##### | -5.1375 | 2.7837E-07 | 9.8701E-07 |
| ENSG00000096401\|CDC5L | 5922.3623 | 0.6221 | ##### | 5.1367 | 2.7967E-07 | 9.9150E-07 |
| ENSG00000182472\|CAPN12 | 142.6833 | -1.5644 | ##### | -5.1362 | 2.8030E-07 | 9.9361E-07 |
| ENSG00000182173\|TSEN54 | 2743.7623 | 0.4380 | ##### | 5.1361 | 2.8050E-07 | 9.9419E-07 |
| ENSG00000112139\|MDGA1 | 204.8526 | -1.0651 | ##### | -5.1352 | 2.8183E-07 | 9.9881E-07 |
| ENSG00000269313\|MAGIX | 91.1907 | -1.3345 | ##### | -5.1347 | 2.8259E-07 | 1.0014E-06 |
| ENSG00000198933\|TBKBP1 | 2239.7368 | -0.6427 | ##### | -5.1346 | 2.8274E-07 | 1.0018E-06 |
| ENSG00000101216\|GMEB2 | 3015.8486 | 0.6140 | ##### | 5.1346 | 2.8281E-07 | 1.0019E-06 |
| ENSG00000149257\|SERPINH1 | 18243.6684 | 0.5474 | ##### | 5.1338 | 2.8389E-07 | 1.0056E-06 |
| ENSG00000290394 | 161.2521 | 0.9563 | ##### | 5.1325 | 2.8598E-07 | 1.0129E-06 |
| ENSG00000260001\|TGFBR3L | 375.9021 | -0.8670 | ##### | -5.1320 | 2.8668E-07 | 1.0153E-06 |
| ENSG00000006530\|AGK | 3829.2351 | 0.6516 | ##### | 5.1313 | 2.8770E-07 | 1.0188E-06 |
| ENSG00000134802\|SLC43A3 | 742.6319 | -9.7430 | ##### | -5.1312 | 2.8795E-07 | 1.0195E-06 |
| ENSG00000143952\|VPS54 | 3184.4094 | -0.6246 | ##### | -5.1301 | 2.8952E-07 | 1.0250E-06 |
| ENSG00000291040\|CD99P1 | 116.9169 | -1.2033 | ##### | -5.1291 | 2.9115E-07 | 1.0306E-06 |
| ENSG00000196642\|RABL6 | 11226.7500 | -0.5917 | ##### | -5.1290 | 2.9134E-07 | 1.0312E-06 |
| ENSG00000138758\|SEPTIN11 | 11576.7501 | 0.6216 | ##### | 5.1284 | 2.9226E-07 | 1.0343E-06 |
| ENSG00000167470\|MIDN | 9630.2864 | 0.5750 | ##### | 5.1283 | 2.9238E-07 | 1.0346E-06 |
| ENSG00000231292\|IGKV1OR2-108 | 20.4959 | -6.7826 | ##### | -5.1283 | 2.9244E-07 | 1.0347E-06 |

| | | | | | | |
|---|---|---|---|---|---|---|
| ENSG00000141295\|SCRN2 | 949.0997 | -0.6802 | ##### | -5.1261 | 2.9584E-07 | 1.0466E-06 |
| ENSG00000254842\|LINC02551 | 23.7579 | 2.7915 | ##### | 5.1256 | 2.9653E-07 | 1.0489E-06 |
| ENSG00000189184\|PCDH18 | 73.2328 | 3.7052 | ##### | 5.1252 | 2.9717E-07 | 1.0511E-06 |
| ENSG00000116874\|WARS2 | 1716.3119 | 0.6590 | ##### | 5.1251 | 2.9736E-07 | 1.0516E-06 |
| ENSG00000141384\|TAF4B | 1459.6096 | 0.6795 | ##### | 5.1210 | 3.0393E-07 | 1.0747E-06 |
| ENSG00000227212\|PFN1P6 | 76.9265 | 1.3314 | ##### | 5.1209 | 3.0414E-07 | 1.0753E-06 |
| ENSG00000189409\|MMP23B | 77.1664 | 1.2043 | ##### | 5.1208 | 3.0428E-07 | 1.0757E-06 |
| ENSG00000253541\|SEPTIN10P1 | 24.9913 | -1.8740 | ##### | -5.1203 | 3.0505E-07 | 1.0783E-06 |
| ENSG00000235001\|EIF4A1P2 | 96.9807 | 1.3847 | ##### | 5.1195 | 3.0638E-07 | 1.0828E-06 |
| ENSG00000206417\|H1-10-AS1 | 897.2804 | 0.7483 | ##### | 5.1192 | 3.0679E-07 | 1.0842E-06 |
| ENSG00000110955\|ATP5F1B | 93509.3209 | 0.5118 | ##### | 5.1187 | 3.0769E-07 | 1.0872E-06 |
| ENSG00000231908\|IDH1-AS1 | 35.2650 | -5.3854 | ##### | -5.1187 | 3.0772E-07 | 1.0872E-06 |
| ENSG00000127337\|YEATS4 | 1779.5089 | 0.6399 | ##### | 5.1184 | 3.0808E-07 | 1.0883E-06 |
| ENSG00000183317\|EPHA10 | 393.1373 | 0.7948 | ##### | 5.1183 | 3.0837E-07 | 1.0892E-06 |
| ENSG00000215374\|FAM66B | 33.9100 | 4.5669 | ##### | 5.1169 | 3.1058E-07 | 1.0969E-06 |
| ENSG00000167747\|C19orf48P | 121.5808 | -3.1692 | ##### | -5.1150 | 3.1381E-07 | 1.1082E-06 |
| ENSG00000213930\|GALT | 941.2902 | 0.7798 | ##### | 5.1149 | 3.1399E-07 | 1.1087E-06 |
| ENSG00000258643\|BCL2L2-PABPN1 | 1175.0074 | 1.1617 | ##### | 5.1147 | 3.1427E-07 | 1.1096E-06 |
| ENSG00000264578 | 591.2998 | 0.8336 | ##### | 5.1143 | 3.1498E-07 | 1.1119E-06 |
| ENSG00000159788\|RGS12 | 945.2156 | 0.6995 | ##### | 5.1142 | 3.1510E-07 | 1.1122E-06 |
| ENSG00000159199\|ATP5MC1 | 7157.7279 | -0.6126 | ##### | -5.1135 | 3.1625E-07 | 1.1162E-06 |
| ENSG00000123473\|STIL | 4878.3906 | 0.6150 | ##### | 5.1132 | 3.1678E-07 | 1.1179E-06 |
| ENSG00000112658\|SRF | 6554.1396 | 0.6058 | ##### | 5.1129 | 3.1732E-07 | 1.1196E-06 |
| ENSG00000164970\|FAM219A | 1211.4565 | -0.6794 | ##### | -5.1127 | 3.1764E-07 | 1.1206E-06 |
| ENSG00000163964\|PIGX | 2775.1110 | -0.6237 | ##### | -5.1126 | 3.1771E-07 | 1.1208E-06 |
| ENSG00000134853\|PDGFRA | 133.5737 | 5.2712 | ##### | 5.1114 | 3.1984E-07 | 1.1282E-06 |
| ENSG00000163814\|CDCP1 | 7233.2152 | 0.6568 | ##### | 5.1112 | 3.2008E-07 | 1.1288E-06 |
| ENSG00000199053\|MIR324 | 39.8350 | 1.4242 | ##### | 5.1106 | 3.2106E-07 | 1.1322E-06 |
| ENSG00000138780\|GSTCD | 2775.3701 | -0.5276 | ##### | -5.1106 | 3.2112E-07 | 1.1323E-06 |
| ENSG00000218027 | 87.7732 | -8.8826 | ##### | -5.1104 | 3.2149E-07 | 1.1334E-06 |
| ENSG00000250067\|YJEFN3 | 412.3609 | -1.2590 | ##### | -5.1103 | 3.2169E-07 | 1.1340E-06 |
| ENSG00000087008\|ACOX3 | 975.6684 | -0.6723 | ##### | -5.1090 | 3.2390E-07 | 1.1417E-06 |
| ENSG00000173456\|RNF26 | 7153.9917 | 0.6241 | ##### | 5.1089 | 3.2400E-07 | 1.1419E-06 |
| ENSG00000269821\|KCNQ1OT1 | 29.9868 | -5.2256 | ##### | -5.1088 | 3.2430E-07 | 1.1428E-06 |
| ENSG00000184489\|PTP4A3 | 786.4363 | -0.7001 | ##### | -5.1068 | 3.2760E-07 | 1.1543E-06 |
| ENSG00000112081\|SRSF3 | 23735.8671 | 0.6260 | ##### | 5.1065 | 3.2810E-07 | 1.1559E-06 |
| ENSG00000198586\|TLK1 | 5633.7205 | 0.6102 | ##### | 5.1061 | 3.2883E-07 | 1.1584E-06 |
| ENSG00000291075 | 464.5720 | -0.7839 | ##### | -5.1056 | 3.2975E-07 | 1.1615E-06 |
| ENSG00000275180 | 99.2780 | 1.3095 | ##### | 5.1051 | 3.3058E-07 | 1.1642E-06 |
| ENSG00000158615\|PPP1R15B | 9366.9855 | -0.6288 | ##### | -5.1050 | 3.3074E-07 | 1.1647E-06 |
| ENSG00000204271\|SPIN3 | 776.2819 | -0.6595 | ##### | -5.1040 | 3.3261E-07 | 1.1711E-06 |
| ENSG00000113141\|IK | 7717.6510 | 0.6149 | ##### | 5.1038 | 3.3288E-07 | 1.1719E-06 |
| ENSG00000031823\|RANBP3 | 4866.8732 | 0.6208 | ##### | 5.1030 | 3.3438E-07 | 1.1771E-06 |
| ENSG00000196365\|LONP1 | 11042.3222 | 0.5796 | ##### | 5.1025 | 3.3516E-07 | 1.1796E-06 |
| ENSG00000241484\|ARHGAP8 | 385.5436 | 1.0946 | ##### | 5.1025 | 3.3518E-07 | 1.1796E-06 |
| ENSG00000100441\|KHNYN | 8655.7446 | 0.6182 | ##### | 5.1021 | 3.3588E-07 | 1.1819E-06 |
| ENSG00000149743\|TRPT1 | 1282.4449 | 0.6977 | ##### | 5.1017 | 3.3670E-07 | 1.1847E-06 |
| ENSG00000059728\|MXD1 | 370.6237 | -0.7699 | ##### | -5.0998 | 3.3997E-07 | 1.1960E-06 |
| ENSG00000119673\|ACOT2 | 2144.7262 | 0.6400 | ##### | 5.0990 | 3.4137E-07 | 1.2008E-06 |
| ENSG00000227992 | 52.7663 | -1.2990 | ##### | -5.0990 | 3.4139E-07 | 1.2008E-06 |
| ENSG00000271614\|ATP2B1-AS1 | 472.5355 | -0.7970 | ##### | -5.0982 | 3.4287E-07 | 1.2058E-06 |
| ENSG00000187741\|FANCA | 4632.7961 | 0.6636 | ##### | 5.0978 | 3.4357E-07 | 1.2081E-06 |
| ENSG00000203667\|COX20 | 6345.1868 | -0.7475 | ##### | -5.0959 | 3.4711E-07 | 1.2204E-06 |
| ENSG00000230825\|LINC03016 | 27.2843 | -1.8550 | ##### | -5.0959 | 3.4716E-07 | 1.2205E-06 |
| ENSG00000259577\|CERNA1 | 22.9872 | -2.2047 | ##### | -5.0958 | 3.4731E-07 | 1.2208E-06 |
| ENSG00000070761\|CFAP20 | 3361.8990 | 0.6063 | ##### | 5.0957 | 3.4738E-07 | 1.2209E-06 |
| ENSG00000130770\|ATP5IF1 | 3646.5078 | -0.6414 | ##### | -5.0955 | 3.4786E-07 | 1.2225E-06 |
| ENSG00000146373\|RNF217 | 1061.3184 | -0.6954 | ##### | -5.0946 | 3.4949E-07 | 1.2281E-06 |
| ENSG00000278376 | 40.8009 | -1.9202 | ##### | -5.0945 | 3.4970E-07 | 1.2287E-06 |
| ENSG00000288062 | 32.7551 | 4.4948 | ##### | 5.0941 | 3.5034E-07 | 1.2308E-06 |
| ENSG00000228278\|ORM2 | 215.4697 | ####### | ##### | -5.0935 | 3.5153E-07 | 1.2348E-06 |
| ENSG00000290943\|ABCC6P2 | 30.6111 | -2.9937 | ##### | -5.0934 | 3.5175E-07 | 1.2355E-06 |
| ENSG00000166847\|DCTN5 | 6782.4640 | -0.6101 | ##### | -5.0933 | 3.5185E-07 | 1.2357E-06 |

| | | | | | | |
|---|---|---|---|---|---|---|
| ENSG00000175029\|CTBP2 | 10078.4888 | -0.4241 | ##### | -5.0922 | 3.5399E-07 | 1.2430E-06 |
| ENSG00000287787 | 50.7153 | -1.6342 | ##### | -5.0917 | 3.5480E-07 | 1.2457E-06 |
| ENSG00000129235\|TXNDC17 | 4631.8689 | 0.6324 | ##### | 5.0908 | 3.5654E-07 | 1.2517E-06 |
| ENSG00000119912\|IDE | 5372.1010 | -0.6317 | ##### | -5.0907 | 3.5683E-07 | 1.2525E-06 |
| ENSG00000120656\|TAF12 | 1875.2921 | 0.6425 | ##### | 5.0904 | 3.5728E-07 | 1.2540E-06 |
| ENSG00000134955\|SLC37A2 | 45.1008 | 1.8976 | ##### | 5.0900 | 3.5813E-07 | 1.2568E-06 |
| ENSG00000111481\|COPZ1 | 15680.1713 | -0.6299 | ##### | -5.0896 | 3.5880E-07 | 1.2590E-06 |
| ENSG00000290385 | 113.3420 | 1.0525 | ##### | 5.0883 | 3.6122E-07 | 1.2674E-06 |
| ENSG00000235241 | 73.4500 | 1.2805 | ##### | 5.0880 | 3.6193E-07 | 1.2697E-06 |
| ENSG00000184076\|UQCR10 | 4635.8541 | -0.6139 | ##### | -5.0877 | 3.6238E-07 | 1.2711E-06 |
| ENSG00000244128\|LINC01322 | 114.9950 | -6.2032 | ##### | -5.0874 | 3.6299E-07 | 1.2731E-06 |
| ENSG00000285589 | 5619.5049 | 0.6273 | ##### | 5.0854 | 3.6693E-07 | 1.2868E-06 |
| ENSG00000238113\|LINC01410 | 39.5530 | -1.6837 | ##### | -5.0848 | 3.6799E-07 | 1.2904E-06 |
| ENSG00000289100 | 3412.6370 | -0.6930 | ##### | -5.0843 | 3.6902E-07 | 1.2938E-06 |
| ENSG00000281333 | 27.6392 | -5.2179 | ##### | -5.0825 | 3.7241E-07 | 1.3056E-06 |
| ENSG00000259205\|PRKXP1 | 58.4841 | 7.6166 | ##### | 5.0817 | 3.7406E-07 | 1.3112E-06 |
| ENSG00000109255\|NMU | 79.7668 | 3.1747 | ##### | 5.0811 | 3.7528E-07 | 1.3153E-06 |
| ENSG00000109917\|ZPR1 | 5751.6517 | 0.5983 | ##### | 5.0805 | 3.7635E-07 | 1.3189E-06 |
| ENSG00000251196 | 46.0209 | -1.5770 | ##### | -5.0798 | 3.7779E-07 | 1.3238E-06 |
| ENSG00000259974\|LINC00261 | 35.1981 | -5.2300 | ##### | -5.0797 | 3.7811E-07 | 1.3248E-06 |
| ENSG00000173917\|HOXB2 | 55.2898 | 1.8558 | ##### | 5.0795 | 3.7851E-07 | 1.3260E-06 |
| ENSG00000141034\|GID4 | 1215.9302 | 0.6736 | ##### | 5.0779 | 3.8156E-07 | 1.3365E-06 |
| ENSG00000008710\|PKD1 | 6664.4459 | 0.6512 | ##### | 5.0778 | 3.8180E-07 | 1.3372E-06 |
| ENSG00000271776 | 31.5584 | 6.4267 | ##### | 5.0760 | 3.8555E-07 | 1.3502E-06 |
| ENSG00000274026\|FAM27E3 | 119.8253 | -1.0442 | ##### | -5.0756 | 3.8636E-07 | 1.3529E-06 |
| ENSG00000189052\|CGB5 | 19.1378 | -2.9120 | ##### | -5.0755 | 3.8653E-07 | 1.3533E-06 |
| ENSG00000078618\|NRDC | 25775.7784 | -0.3394 | ##### | -5.0743 | 3.8890E-07 | 1.3614E-06 |
| ENSG00000104714\|ERICH1 | 763.2346 | 0.6959 | ##### | 5.0741 | 3.8931E-07 | 1.3627E-06 |
| ENSG00000122958\|VPS26A | 5901.0131 | -0.6373 | ##### | -5.0740 | 3.8956E-07 | 1.3634E-06 |
| ENSG00000158161\|EYA3 | 2001.2714 | 0.6304 | ##### | 5.0738 | 3.8995E-07 | 1.3646E-06 |
| ENSG00000225422\|RBMS1P1 | 90.3797 | -1.1493 | ##### | -5.0736 | 3.9034E-07 | 1.3658E-06 |
| ENSG00000140374\|ETFA | 7825.2423 | -0.5048 | ##### | -5.0730 | 3.9162E-07 | 1.3702E-06 |
| ENSG00000266258\|LINC01909 | 60.3203 | 1.8696 | ##### | 5.0729 | 3.9187E-07 | 1.3709E-06 |
| ENSG00000140365\|COMMD4 | 4553.0783 | 0.6337 | ##### | 5.0723 | 3.9305E-07 | 1.3748E-06 |
| ENSG00000185479\|KRT6B | 37.9732 | -6.3630 | ##### | -5.0723 | 3.9309E-07 | 1.3748E-06 |
| ENSG00000165474\|GJB2 | 117.2779 | 3.1371 | ##### | 5.0719 | 3.9381E-07 | 1.3772E-06 |
| ENSG00000110104\|CCDC86 | 9596.3509 | 0.6220 | ##### | 5.0718 | 3.9407E-07 | 1.3779E-06 |
| ENSG00000234264\|DEPDC1-AS1 | 37.3818 | 2.0769 | ##### | 5.0700 | 3.9776E-07 | 1.3907E-06 |
| ENSG00000181085\|MAPK15 | 202.6453 | 1.0393 | ##### | 5.0699 | 3.9805E-07 | 1.3915E-06 |
| ENSG00000167721\|TSR1 | 16240.6795 | 0.4164 | ##### | 5.0698 | 3.9830E-07 | 1.3922E-06 |
| ENSG00000273828 | 46.5778 | 1.6532 | ##### | 5.0692 | 3.9958E-07 | 1.3965E-06 |
| ENSG00000258311 | 418.3259 | -0.8691 | ##### | -5.0682 | 4.0157E-07 | 1.4033E-06 |
| ENSG00000196700\|ZNF512B | 4031.6009 | 0.6011 | ##### | 5.0675 | 4.0302E-07 | 1.4082E-06 |
| ENSG00000276517 | 34.4748 | 2.0734 | ##### | 5.0675 | 4.0311E-07 | 1.4084E-06 |
| ENSG00000130803\|ZNF317 | 3331.4148 | 0.6305 | ##### | 5.0673 | 4.0357E-07 | 1.4098E-06 |
| ENSG00000218281\|H2AC9P | 13.3444 | 3.4608 | ##### | 5.0660 | 4.0634E-07 | 1.4193E-06 |
| ENSG00000225778\|PROSER2-AS1 | 35.0456 | -5.1534 | ##### | -5.0659 | 4.0651E-07 | 1.4198E-06 |
| ENSG00000170270\|GON7 | 801.5436 | -0.7021 | ##### | -5.0656 | 4.0716E-07 | 1.4219E-06 |
| ENSG00000203721\|LINC00862 | 61.9860 | -1.7539 | ##### | -5.0652 | 4.0797E-07 | 1.4245E-06 |
| ENSG00000172379\|ARNT2 | 217.5647 | 0.8673 | ##### | 5.0649 | 4.0867E-07 | 1.4268E-06 |
| ENSG00000267255 | 26.2679 | 6.5600 | ##### | 5.0646 | 4.0927E-07 | 1.4287E-06 |
| ENSG00000169689\|CENPX | 4883.1591 | 0.6187 | ##### | 5.0642 | 4.1016E-07 | 1.4317E-06 |
| ENSG00000114933\|INO80D | 1373.7491 | 0.6528 | ##### | 5.0634 | 4.1181E-07 | 1.4373E-06 |
| ENSG00000204394\|VARS1 | 23604.8398 | 0.5018 | ##### | 5.0626 | 4.1363E-07 | 1.4435E-06 |
| ENSG00000166845\|C18orf54 | 1217.1548 | -0.6673 | ##### | -5.0623 | 4.1428E-07 | 1.4456E-06 |
| ENSG00000165861\|ZFYVE1 | 1289.9538 | 0.6547 | ##### | 5.0622 | 4.1453E-07 | 1.4461E-06 |
| ENSG00000162040\|HS3ST6 | 35.8385 | -1.8239 | ##### | -5.0622 | 4.1450E-07 | 1.4461E-06 |
| ENSG00000236194 | 47.3571 | 6.3146 | ##### | 5.0619 | 4.1518E-07 | 1.4482E-06 |
| ENSG00000203778\|FAM229B | 298.1055 | -0.9334 | ##### | -5.0615 | 4.1593E-07 | 1.4506E-06 |
| ENSG00000196361\|ELAVL3 | 27.2057 | 6.2447 | ##### | 5.0609 | 4.1735E-07 | 1.4554E-06 |
| ENSG00000003989\|SLC7A2 | 925.1708 | 0.6983 | ##### | 5.0608 | 4.1744E-07 | 1.4555E-06 |
| ENSG00000213277\|MARCKSL1P1 | 19.7807 | -6.6693 | ##### | -5.0605 | 4.1826E-07 | 1.4582E-06 |
| ENSG00000183401\|CCDC159 | 348.5522 | -0.9108 | ##### | -5.0595 | 4.2029E-07 | 1.4651E-06 |

| | | | | | | |
|---|---|---|---|---|---|---|
| ENSG00000171456\|ASXL1 | 9512.5724 | 0.6064 | ##### | 5.0592 | 4.2110E-07 | 1.4678E-06 |
| ENSG00000267221\|C17orf113 | 49.2829 | -1.8312 | ##### | -5.0591 | 4.2131E-07 | 1.4684E-06 |
| ENSG00000291132 | 204.6298 | 1.1552 | ##### | 5.0585 | 4.2248E-07 | 1.4723E-06 |
| ENSG00000079277\|MKNK1 | 1705.1926 | -0.6639 | ##### | -5.0572 | 4.2540E-07 | 1.4823E-06 |
| ENSG00000249839 | 482.9301 | -0.7658 | ##### | -5.0571 | 4.2578E-07 | 1.4834E-06 |
| ENSG00000251003\|ZFPM2-AS1 | 201.1694 | -4.0177 | ##### | -5.0570 | 4.2592E-07 | 1.4837E-06 |
| ENSG00000265630 | 18.7457 | -6.6783 | ##### | -5.0564 | 4.2735E-07 | 1.4886E-06 |
| ENSG00000072818\|ACAP1 | 132.9294 | 0.9838 | ##### | 5.0549 | 4.3064E-07 | 1.4998E-06 |
| ENSG00000119718\|EIF2B2 | 4060.3774 | 0.6125 | ##### | 5.0538 | 4.3303E-07 | 1.5080E-06 |
| ENSG00000158406\|H4C8 | 991.4310 | 0.7130 | ##### | 5.0535 | 4.3386E-07 | 1.5107E-06 |
| ENSG00000189306\|RRP7A | 9211.9314 | 0.5973 | ##### | 5.0532 | 4.3458E-07 | 1.5130E-06 |
| ENSG00000288927 | 50.7941 | -5.1442 | ##### | -5.0509 | 4.3966E-07 | 1.5305E-06 |
| ENSG00000123395\|ATG101 | 8178.9930 | 0.6161 | ##### | 5.0509 | 4.3982E-07 | 1.5309E-06 |
| ENSG00000172348\|RCAN2 | 32.5570 | 2.6797 | ##### | 5.0507 | 4.4015E-07 | 1.5319E-06 |
| ENSG00000257086 | 195.6333 | 0.9117 | ##### | 5.0503 | 4.4113E-07 | 1.5351E-06 |
| ENSG00000159388\|BTG2 | 471.3713 | 0.7628 | ##### | 5.0499 | 4.4198E-07 | 1.5379E-06 |
| ENSG00000189129\|PLAC9 | 30.3605 | -7.3516 | ##### | -5.0490 | 4.4407E-07 | 1.5450E-06 |
| ENSG00000257327 | 17.9152 | -3.1011 | ##### | -5.0480 | 4.4657E-07 | 1.5535E-06 |
| ENSG00000006459\|KDM7A | 1323.8516 | 0.6769 | ##### | 5.0471 | 4.4867E-07 | 1.5606E-06 |
| ENSG00000006125\|AP2B1 | 32819.1559 | -0.2963 | ##### | -5.0467 | 4.4960E-07 | 1.5637E-06 |
| ENSG00000164122\|ASB5 | 36.6579 | 3.7970 | ##### | 5.0460 | 4.5111E-07 | 1.5687E-06 |
| ENSG00000261150\|EPPK1 | 355.7536 | 1.0198 | ##### | 5.0451 | 4.5340E-07 | 1.5765E-06 |
| ENSG00000147408\|CSGALNACT1 | 4796.8913 | -0.6138 | ##### | -5.0449 | 4.5374E-07 | 1.5775E-06 |
| ENSG00000221823\|PPP3R1 | 9231.5887 | 0.4660 | ##### | 5.0440 | 4.5586E-07 | 1.5847E-06 |
| ENSG00000006042\|TMEM98 | 512.9532 | -0.7589 | ##### | -5.0440 | 4.5593E-07 | 1.5848E-06 |
| ENSG00000115128\|SF3B6 | 7911.0514 | 0.6005 | ##### | 5.0438 | 4.5642E-07 | 1.5863E-06 |
| ENSG00000187537\|POTEG | 17.4420 | -5.7773 | ##### | -5.0437 | 4.5658E-07 | 1.5867E-06 |
| ENSG00000285996 | 41.6928 | -1.8452 | ##### | -5.0422 | 4.6029E-07 | 1.5994E-06 |
| ENSG00000204514\|ZNF814 | 1062.7301 | -0.6930 | ##### | -5.0417 | 4.6136E-07 | 1.6029E-06 |
| ENSG00000243094\|RPL32P2 | 16.9029 | 6.6426 | ##### | 5.0416 | 4.6166E-07 | 1.6038E-06 |
| ENSG00000118307\|DNAI7 | 33.3579 | 4.4375 | ##### | 5.0412 | 4.6265E-07 | 1.6070E-06 |
| ENSG00000099246\|RAB18 | 6209.7711 | -0.5965 | ##### | -5.0409 | 4.6332E-07 | 1.6092E-06 |
| ENSG00000262097\|LINC02185 | 25.0031 | 2.3375 | ##### | 5.0402 | 4.6496E-07 | 1.6147E-06 |
| ENSG00000186111\|PIP5K1C | 3798.9338 | 0.4618 | ##### | 5.0397 | 4.6621E-07 | 1.6188E-06 |
| ENSG00000178385\|PLEKHM3 | 1078.9219 | -0.6814 | ##### | -5.0393 | 4.6723E-07 | 1.6222E-06 |
| ENSG00000174837\|ADGRE1 | 644.7015 | 0.7271 | ##### | 5.0375 | 4.7162E-07 | 1.6372E-06 |
| ENSG00000147509\|RGS20 | 2888.7082 | 0.6511 | ##### | 5.0374 | 4.7202E-07 | 1.6384E-06 |
| ENSG00000135093\|USP30 | 780.5148 | -0.7038 | ##### | -5.0370 | 4.7280E-07 | 1.6409E-06 |
| ENSG00000281392\|LINC00506 | 242.1381 | ####### | ##### | -5.0366 | 4.7387E-07 | 1.6445E-06 |
| ENSG00000173825\|TIGD3 | 40.2559 | 1.7626 | ##### | 5.0365 | 4.7424E-07 | 1.6456E-06 |
| ENSG00000215908\|CROCCP2 | 166.5958 | 1.0816 | ##### | 5.0363 | 4.7473E-07 | 1.6469E-06 |
| ENSG00000231903 | 30.5212 | 2.0684 | ##### | 5.0362 | 4.7475E-07 | 1.6469E-06 |
| ENSG00000163104\|SMARCAD1 | 4937.3083 | 0.6247 | ##### | 5.0359 | 4.7569E-07 | 1.6500E-06 |
| ENSG00000130827\|PLXNA3 | 1796.2750 | -0.6357 | ##### | -5.0357 | 4.7618E-07 | 1.6515E-06 |
| ENSG00000165195\|PIGA | 955.7303 | -0.6054 | ##### | -5.0351 | 4.7748E-07 | 1.6558E-06 |
| ENSG00000125998\|FAM83C | 70.7934 | 8.4045 | ##### | 5.0349 | 4.7813E-07 | 1.6579E-06 |
| ENSG00000267124 | 103.6434 | 1.0538 | ##### | 5.0346 | 4.7884E-07 | 1.6602E-06 |
| ENSG00000250790 | 59.0123 | -1.6996 | ##### | -5.0341 | 4.8022E-07 | 1.6647E-06 |
| ENSG00000158296\|SLC13A3 | 18.5189 | 2.4792 | ##### | 5.0338 | 4.8083E-07 | 1.6667E-06 |
| ENSG00000103353\|UBFD1 | 9784.4667 | -0.5901 | ##### | -5.0335 | 4.8164E-07 | 1.6693E-06 |
| ENSG00000180616\|SSTR2 | 55.8828 | 2.2465 | ##### | 5.0325 | 4.8416E-07 | 1.6778E-06 |
| ENSG00000289915 | 34.7454 | -7.3313 | ##### | -5.0315 | 4.8674E-07 | 1.6866E-06 |
| ENSG00000229291\|LINC02768 | 13.0661 | 6.9738 | ##### | 5.0294 | 4.9197E-07 | 1.7045E-06 |
| ENSG00000270039 | 37.2012 | 2.0616 | ##### | 5.0290 | 4.9302E-07 | 1.7079E-06 |
| ENSG00000166341\|DCHS1 | 87.1870 | -1.3734 | ##### | -5.0288 | 4.9357E-07 | 1.7096E-06 |
| ENSG00000147403\|RPL10 | 66792.0124 | 0.6080 | ##### | 5.0287 | 4.9378E-07 | 1.7101E-06 |
| ENSG00000156831\|NSMCE2 | 1415.4769 | -0.6377 | ##### | -5.0281 | 4.9542E-07 | 1.7156E-06 |
| ENSG00000241635\|UGT1A1 | 251.5715 | ####### | ##### | -5.0280 | 4.9561E-07 | 1.7161E-06 |
| ENSG00000129474\|AJUBA | 6322.3228 | -0.6118 | ##### | -5.0278 | 4.9620E-07 | 1.7180E-06 |
| ENSG00000137707\|BTG4 | 56.2576 | 8.0280 | ##### | 5.0275 | 4.9697E-07 | 1.7204E-06 |
| ENSG00000204859\|ZBTB48 | 1353.2693 | 0.6624 | ##### | 5.0268 | 4.9886E-07 | 1.7268E-06 |
| ENSG00000108262\|GIT1 | 10193.0469 | 0.6018 | ##### | 5.0258 | 5.0123E-07 | 1.7348E-06 |
| ENSG00000131979\|GCH1 | 670.4210 | -0.7579 | ##### | -5.0247 | 5.0420E-07 | 1.7448E-06 |

| | | | | | | |
|---|---|---|---|---|---|---|
| ENSG00000267169\|ADGRL1-AS1 | 134.7132 | 1.1277 | ##### | 5.0244 | 5.0489E-07 | 1.7470E-06 |
| ENSG00000138030\|KHK | 766.5900 | 0.6973 | ##### | 5.0225 | 5.1001E-07 | 1.7645E-06 |
| ENSG00000159713\|TPPP3 | 64.3019 | -1.7584 | ##### | -5.0216 | 5.1243E-07 | 1.7727E-06 |
| ENSG00000226415\|TPI1P1 | 666.1234 | 0.6996 | ##### | 5.0200 | 5.1675E-07 | 1.7874E-06 |
| ENSG00000088298\|EDEM2 | 4450.1301 | -0.6142 | ##### | -5.0197 | 5.1750E-07 | 1.7898E-06 |
| ENSG00000259607 | 12.8824 | -2.4062 | ##### | -5.0190 | 5.1934E-07 | 1.7960E-06 |
| ENSG00000185641 | 940.2971 | 0.6899 | ##### | 5.0190 | 5.1952E-07 | 1.7964E-06 |
| ENSG00000125656\|CLPP | 4496.7208 | 0.6270 | ##### | 5.0186 | 5.2058E-07 | 1.7999E-06 |
| ENSG00000243943\|ZNF512 | 1580.5635 | 0.6657 | ##### | 5.0161 | 5.2736E-07 | 1.8231E-06 |
| ENSG00000077254\|USP33 | 5462.8718 | -0.5804 | ##### | -5.0160 | 5.2758E-07 | 1.8236E-06 |
| ENSG00000168701\|TMEM208 | 3352.6320 | -0.6296 | ##### | -5.0157 | 5.2851E-07 | 1.8266E-06 |
| ENSG00000169813\|HNRNPF | 18391.5134 | 0.4642 | ##### | 5.0153 | 5.2962E-07 | 1.8303E-06 |
| ENSG00000163703\|CRELD1 | 1317.6390 | -0.6687 | ##### | -5.0149 | 5.3057E-07 | 1.8333E-06 |
| ENSG00000172757\|CFL1 | 82629.3938 | 0.5958 | ##### | 5.0145 | 5.3163E-07 | 1.8368E-06 |
| ENSG00000250182\|EEF1A1P13 | 140.8847 | 0.9783 | ##### | 5.0126 | 5.3690E-07 | 1.8548E-06 |
| ENSG00000167333\|TRIM68 | 828.3224 | -0.6528 | ##### | -5.0121 | 5.3835E-07 | 1.8596E-06 |
| ENSG00000258806\|OR11H7 | 64.4653 | 7.8198 | ##### | 5.0099 | 5.4452E-07 | 1.8807E-06 |
| ENSG00000267041\|ZNF850 | 929.5168 | 0.6662 | ##### | 5.0090 | 5.4725E-07 | 1.8899E-06 |
| ENSG00000189043\|NDUFA4 | 11814.6581 | -0.6026 | ##### | -5.0051 | 5.5827E-07 | 1.9277E-06 |
| ENSG00000161638\|ITGA5 | 14641.7653 | -0.6022 | ##### | -5.0047 | 5.5958E-07 | 1.9320E-06 |
| ENSG00000044446\|PHKA2 | 1701.3599 | -0.6532 | ##### | -5.0043 | 5.6070E-07 | 1.9357E-06 |
| ENSG00000258471 | 90.6277 | 2.3347 | ##### | 5.0040 | 5.6151E-07 | 1.9382E-06 |
| ENSG00000237595\|LINC01275 | 23.0960 | 2.5489 | ##### | 5.0038 | 5.6205E-07 | 1.9399E-06 |
| ENSG00000169136\|ATF5 | 5617.4884 | -0.6164 | ##### | -5.0037 | 5.6241E-07 | 1.9409E-06 |
| ENSG00000187720\|THSD4 | 5757.3109 | 0.6120 | ##### | 5.0034 | 5.6330E-07 | 1.9437E-06 |
| ENSG00000196810\|CTBP1-DT | 1826.4857 | 0.6550 | ##### | 5.0030 | 5.6438E-07 | 1.9470E-06 |
| ENSG00000187391\|MAGI2 | 248.7614 | -0.8159 | ##### | -5.0030 | 5.6436E-07 | 1.9470E-06 |
| ENSG00000273142\|LINC02604 | 147.1945 | -0.9896 | ##### | -5.0027 | 5.6519E-07 | 1.9496E-06 |
| ENSG00000224184\|MIR3681HG | 21.6553 | 6.8173 | ##### | 5.0019 | 5.6764E-07 | 1.9578E-06 |
| ENSG00000290851 | 45.8029 | 1.4954 | ##### | 5.0011 | 5.7001E-07 | 1.9657E-06 |
| ENSG00000159915\|ZNF233 | 41.9855 | -1.9741 | ##### | -5.0010 | 5.7031E-07 | 1.9666E-06 |
| ENSG00000177917\|ARL6IP6 | 4079.4224 | -0.6351 | ##### | -4.9999 | 5.7365E-07 | 1.9778E-06 |
| ENSG00000184203\|PPP1R2 | 2498.2651 | -0.6354 | ##### | -4.9995 | 5.7476E-07 | 1.9814E-06 |
| ENSG00000170448\|NFXL1 | 2341.2029 | 0.6478 | ##### | 4.9994 | 5.7504E-07 | 1.9822E-06 |
| ENSG00000231527\|FAM27C | 141.5726 | -0.9525 | ##### | -4.9973 | 5.8139E-07 | 2.0038E-06 |
| ENSG00000183598\|H3C13 | 44.2870 | 1.9940 | ##### | 4.9967 | 5.8316E-07 | 2.0097E-06 |
| ENSG00000288983 | 123.4916 | -1.1863 | ##### | -4.9963 | 5.8444E-07 | 2.0139E-06 |
| ENSG00000204588\|LINC01123 | 1028.5260 | -0.6404 | ##### | -4.9955 | 5.8695E-07 | 2.0223E-06 |
| ENSG00000168282\|MGAT2 | 4376.9729 | 0.6175 | ##### | 4.9940 | 5.9154E-07 | 2.0376E-06 |
| ENSG00000197646\|PDCD1LG2 | 37.3681 | -5.1497 | ##### | -4.9940 | 5.9149E-07 | 2.0376E-06 |
| ENSG00000234456\|MAGI2-AS3 | 1452.1365 | -0.6394 | ##### | -4.9939 | 5.9163E-07 | 2.0377E-06 |
| ENSG00000119616\|FCF1 | 5381.4116 | 0.5951 | ##### | 4.9937 | 5.9247E-07 | 2.0404E-06 |
| ENSG00000250697 | 194.6280 | ####### | ##### | -4.9913 | 5.9966E-07 | 2.0649E-06 |
| ENSG00000131941\|RHPN2 | 3136.5393 | -0.5594 | ##### | -4.9913 | 5.9984E-07 | 2.0653E-06 |
| ENSG00000197275\|RAD54B | 1701.0899 | -0.6275 | ##### | -4.9901 | 6.0363E-07 | 2.0781E-06 |
| ENSG00000256453\|DND1 | 344.2454 | 0.9085 | ##### | 4.9899 | 6.0415E-07 | 2.0796E-06 |
| ENSG00000111906\|HDDC2 | 3733.5092 | -0.6014 | ##### | -4.9896 | 6.0508E-07 | 2.0826E-06 |
| ENSG00000138092\|CENPO | 7483.6902 | 0.6144 | ##### | 4.9890 | 6.0706E-07 | 2.0892E-06 |
| ENSG00000272523\|LINC01023 | 94.3565 | -1.5086 | ##### | -4.9884 | 6.0877E-07 | 2.0948E-06 |
| ENSG00000263590\|LINC02591 | 113.3980 | 1.0220 | ##### | 4.9881 | 6.0991E-07 | 2.0985E-06 |
| ENSG00000198466\|ZNF587 | 3117.2485 | -0.6204 | ##### | -4.9878 | 6.1069E-07 | 2.1007E-06 |
| ENSG00000138798\|EGF | 193.1311 | -6.5784 | ##### | -4.9878 | 6.1064E-07 | 2.1007E-06 |
| ENSG00000130540\|SULT4A1 | 369.5398 | 0.8774 | ##### | 4.9876 | 6.1142E-07 | 2.1030E-06 |
| ENSG00000197093\|GAL3ST4 | 45.1897 | 2.6389 | ##### | 4.9865 | 6.1486E-07 | 2.1146E-06 |
| ENSG00000198826\|ARHGAP11A | 8082.1391 | 1.4764 | ##### | 4.9860 | 6.1627E-07 | 2.1192E-06 |
| ENSG00000213397\|HAUS7 | 1414.5493 | 0.7882 | ##### | 4.9850 | 6.1952E-07 | 2.1301E-06 |
| ENSG00000279381 | 15.3841 | 3.2857 | ##### | 4.9845 | 6.2116E-07 | 2.1355E-06 |
| ENSG00000214274\|ANG | 320.0922 | -0.8424 | ##### | -4.9842 | 6.2227E-07 | 2.1388E-06 |
| ENSG00000115464\|USP34 | 12873.6706 | 0.6029 | ##### | 4.9842 | 6.2221E-07 | 2.1388E-06 |
| ENSG00000290460 | 45.2610 | -1.5738 | ##### | -4.9839 | 6.2300E-07 | 2.1411E-06 |
| ENSG00000130940\|CASZ1 | 1454.6458 | 0.6936 | ##### | 4.9838 | 6.2335E-07 | 2.1418E-06 |
| ENSG00000258568\|RHOQP1 | 86.6087 | 1.3321 | ##### | 4.9838 | 6.2335E-07 | 2.1418E-06 |
| ENSG00000218073 | 25.4245 | 5.5809 | ##### | 4.9833 | 6.2496E-07 | 2.1471E-06 |

| | | | | | | |
|---|---|---|---|---|---|---|
| ENSG00000245532\|NEAT1 | 12139.3297 | 0.6257 | ##### | 4.9826 | 6.2723E-07 | 2.1546E-06 |
| ENSG00000171450\|CDK5R2 | 186.9032 | -0.9095 | ##### | -4.9816 | 6.3053E-07 | 2.1657E-06 |
| ENSG00000135638\|EMX1 | 42.2654 | 5.5523 | ##### | 4.9810 | 6.3264E-07 | 2.1727E-06 |
| ENSG00000110660\|SLC35F2 | 5707.9116 | 0.6271 | ##### | 4.9798 | 6.3635E-07 | 2.1851E-06 |
| ENSG00000286153\|RUNX1-AS1 | 19.0556 | 6.7523 | ##### | 4.9798 | 6.3640E-07 | 2.1851E-06 |
| ENSG00000145107\|TM4SF19 | 24.1151 | -6.1111 | ##### | -4.9796 | 6.3706E-07 | 2.1871E-06 |
| ENSG00000070614\|NDST1 | 8079.7498 | -0.6004 | ##### | -4.9792 | 6.3839E-07 | 2.1914E-06 |
| ENSG00000106991\|ENG | 64.9880 | -1.3906 | ##### | -4.9788 | 6.3969E-07 | 2.1957E-06 |
| ENSG00000261586 | 29.5708 | 3.0107 | ##### | 4.9788 | 6.3981E-07 | 2.1958E-06 |
| ENSG00000109111\|SUPT6H | 17862.5266 | 0.5859 | ##### | 4.9778 | 6.4319E-07 | 2.2072E-06 |
| ENSG00000198478\|SH3BGRL2 | 489.6658 | -0.7160 | ##### | -4.9765 | 6.4749E-07 | 2.2217E-06 |
| ENSG00000164258\|NDUFS4 | 1956.0376 | -0.6740 | ##### | -4.9758 | 6.4982E-07 | 2.2294E-06 |
| ENSG00000171723\|GPHN | 2113.4557 | 0.6341 | ##### | 4.9750 | 6.5243E-07 | 2.2381E-06 |
| ENSG00000119203\|CPSF3 | 6203.7149 | 0.3914 | ##### | 4.9744 | 6.5446E-07 | 2.2448E-06 |
| ENSG00000197930\|ERO1A | 7685.3468 | -0.5391 | ##### | -4.9742 | 6.5529E-07 | 2.2474E-06 |
| ENSG00000184368\|MAP7D2 | 273.1238 | -7.5412 | ##### | -4.9740 | 6.5583E-07 | 2.2490E-06 |
| ENSG00000188026\|RILPL1 | 1526.2761 | -0.6465 | ##### | -4.9740 | 6.5591E-07 | 2.2490E-06 |
| ENSG00000109606\|DHX15 | 16934.3507 | 0.5484 | ##### | 4.9735 | 6.5741E-07 | 2.2539E-06 |
| ENSG00000235314\|LINC00957 | 131.4334 | -1.0499 | ##### | -4.9734 | 6.5777E-07 | 2.2548E-06 |
| ENSG00000167895\|TMC8 | 378.1249 | -7.8846 | ##### | -4.9721 | 6.6244E-07 | 2.2706E-06 |
| ENSG00000232388\|SMIM26 | 1067.3392 | -0.6312 | ##### | -4.9718 | 6.6350E-07 | 2.2740E-06 |
| ENSG00000258334\|TROAP-AS1 | 98.3801 | 1.5404 | ##### | 4.9712 | 6.6526E-07 | 2.2797E-06 |
| ENSG00000240303\|ACAD11 | 1514.6358 | -0.9245 | ##### | -4.9710 | 6.6603E-07 | 2.2821E-06 |
| ENSG00000176261\|ZBTB8OS | 2094.3821 | -0.6119 | ##### | -4.9702 | 6.6874E-07 | 2.2911E-06 |
| ENSG00000130684\|ZNF337 | 895.2302 | 0.7570 | ##### | 4.9702 | 6.6882E-07 | 2.2912E-06 |
| ENSG00000251258\|RFPL4B | 153.7053 | -7.1065 | ##### | -4.9700 | 6.6965E-07 | 2.2937E-06 |
| ENSG00000114021\|NIT2 | 4173.2000 | -0.5486 | ##### | -4.9697 | 6.7068E-07 | 2.2970E-06 |
| ENSG00000253485\|PCDHGA5 | 57.3791 | -4.8846 | ##### | -4.9696 | 6.7092E-07 | 2.2976E-06 |
| ENSG00000188368\|PRR19 | 556.2658 | -1.1202 | ##### | -4.9694 | 6.7154E-07 | 2.2994E-06 |
| ENSG00000105974\|CAV1 | 27917.4510 | -0.5823 | ##### | -4.9694 | 6.7164E-07 | 2.2995E-06 |
| ENSG00000140675\|SLC5A2 | 1077.5026 | -3.1360 | ##### | -4.9686 | 6.7438E-07 | 2.3086E-06 |
| ENSG00000168283\|BMI1 | 8403.6805 | -0.5938 | ##### | -4.9685 | 6.7461E-07 | 2.3091E-06 |
| ENSG00000091656\|ZFHX4 | 297.5252 | 0.8664 | ##### | 4.9677 | 6.7768E-07 | 2.3194E-06 |
| ENSG00000224292\|PORCN-DT | 36.2695 | -7.1898 | ##### | -4.9675 | 6.7811E-07 | 2.3206E-06 |
| ENSG00000214338\|SOGA3 | 265.9555 | 1.3649 | ##### | 4.9669 | 6.8041E-07 | 2.3282E-06 |
| ENSG00000114503\|NCBP2 | 6003.8466 | -0.5222 | ##### | -4.9664 | 6.8194E-07 | 2.3331E-06 |
| ENSG00000214413\|BBIP1 | 1348.3967 | -0.6656 | ##### | -4.9630 | 6.9417E-07 | 2.3747E-06 |
| ENSG00000290398\|ASNSP1 | 194.1133 | ####### | ##### | -4.9624 | 6.9632E-07 | 2.3818E-06 |
| ENSG00000186815\|TPCN1 | 6412.9701 | -0.5952 | ##### | -4.9623 | 6.9656E-07 | 2.3823E-06 |
| ENSG00000162300\|ZFPL1 | 3194.8985 | -0.7656 | ##### | -4.9611 | 7.0096E-07 | 2.3971E-06 |
| ENSG00000132507\|EIF5A | 93057.3090 | 0.4959 | ##### | 4.9600 | 7.0509E-07 | 2.4110E-06 |
| ENSG00000005893\|LAMP2 | 10168.5404 | 0.5845 | ##### | 4.9592 | 7.0791E-07 | 2.4204E-06 |
| ENSG00000116830\|TTF2 | 3531.0253 | -0.5938 | ##### | -4.9590 | 7.0874E-07 | 2.4229E-06 |
| ENSG00000187372\|PCDHB13 | 27.1190 | -2.8853 | ##### | -4.9589 | 7.0890E-07 | 2.4232E-06 |
| ENSG00000146904\|EPHA1 | 141.2933 | 1.2635 | ##### | 4.9588 | 7.0915E-07 | 2.4238E-06 |
| ENSG00000111615\|KRR1 | 11582.5261 | 0.6086 | ##### | 4.9587 | 7.0965E-07 | 2.4252E-06 |
| ENSG00000168906\|MAT2A | 26105.4150 | 0.5247 | ##### | 4.9573 | 7.1485E-07 | 2.4427E-06 |
| ENSG00000262678\|CAMTA2-AS1 | 13.6111 | 3.8783 | ##### | 4.9571 | 7.1553E-07 | 2.4447E-06 |
| ENSG00000143183\|TMCO1 | 6316.7843 | -0.6019 | ##### | -4.9550 | 7.2329E-07 | 2.4710E-06 |
| ENSG00000085998\|POMGNT1 | 10849.8567 | -0.4690 | ##### | -4.9547 | 7.2451E-07 | 2.4748E-06 |
| ENSG00000148459\|PDSS1 | 1338.7336 | 0.6892 | ##### | 4.9524 | 7.3300E-07 | 2.5036E-06 |
| ENSG00000149573\|MPZL2 | 113.7215 | -6.1097 | ##### | -4.9520 | 7.3463E-07 | 2.5085E-06 |
| ENSG00000278107 | 38.5850 | -2.2649 | ##### | -4.9520 | 7.3462E-07 | 2.5085E-06 |
| ENSG00000048740\|CELF2 | 125.4712 | -5.2814 | ##### | -4.9515 | 7.3654E-07 | 2.5148E-06 |
| ENSG00000115540\|MOB4 | 3971.2774 | 0.6134 | ##### | 4.9501 | 7.4178E-07 | 2.5321E-06 |
| ENSG00000154822\|PLCL2 | 201.2086 | -1.0705 | ##### | -4.9501 | 7.4173E-07 | 2.5321E-06 |
| ENSG00000122484\|RPAP2 | 2252.6420 | -0.5952 | ##### | -4.9498 | 7.4297E-07 | 2.5359E-06 |
| ENSG00000285679 | 28.7230 | -7.2703 | ##### | -4.9491 | 7.4538E-07 | 2.5438E-06 |
| ENSG00000121053\|EPX | 19.6311 | 6.6912 | ##### | 4.9482 | 7.4910E-07 | 2.5562E-06 |
| ENSG00000166068\|SPRED1 | 3283.4760 | 0.6268 | ##### | 4.9479 | 7.5023E-07 | 2.5598E-06 |
| ENSG00000122477\|LRRC39 | 61.1069 | -1.3366 | ##### | -4.9475 | 7.5179E-07 | 2.5648E-06 |
| ENSG00000099326\|MZF1 | 1028.0635 | -0.6771 | ##### | -4.9472 | 7.5285E-07 | 2.5681E-06 |
| ENSG00000171695\|LKAAEAR1 | 31.5811 | 2.1445 | ##### | 4.9461 | 7.5720E-07 | 2.5827E-06 |

| | | | | | | |
|---|---|---|---|---|---|---|
| ENSG00000198901\|PRC1 | 12611.1989 | 0.6206 | ##### | 4.9449 | 7.6165E-07 | 2.5975E-06 |
| ENSG00000138433\|CIR1 | 1172.7547 | 0.5980 | ##### | 4.9439 | 7.6587E-07 | 2.6116E-06 |
| ENSG00000184470\|TXNRD2 | 4113.3541 | 0.6159 | ##### | 4.9423 | 7.7210E-07 | 2.6326E-06 |
| ENSG00000142864\|SERBP1 | 39203.4120 | -0.4247 | ##### | -4.9420 | 7.7311E-07 | 2.6357E-06 |
| ENSG00000280049 | 21.8002 | -5.2823 | ##### | -4.9419 | 7.7384E-07 | 2.6379E-06 |
| ENSG00000256073\|URB1-AS1 | 272.1172 | -0.7951 | ##### | -4.9418 | 7.7424E-07 | 2.6390E-06 |
| ENSG00000119801\|YPEL5 | 3387.4854 | -0.6101 | ##### | -4.9412 | 7.7638E-07 | 2.6459E-06 |
| ENSG00000156150\|ALX3 | 149.8931 | 0.9654 | ##### | 4.9407 | 7.7834E-07 | 2.6523E-06 |
| ENSG00000188004\|SNHG28 | 187.8211 | 1.9023 | ##### | 4.9405 | 7.7911E-07 | 2.6547E-06 |
| ENSG00000249755 | 30.4975 | 7.4199 | ##### | 4.9404 | 7.7963E-07 | 2.6561E-06 |
| ENSG00000288935 | 43.0837 | -2.0693 | ##### | -4.9403 | 7.8011E-07 | 2.6575E-06 |
| ENSG00000025423\|HSD17B6 | 156.3305 | 0.9630 | ##### | 4.9395 | 7.8308E-07 | 2.6673E-06 |
| ENSG00000241388\|HNF1A-AS1 | 157.4448 | -9.7238 | ##### | -4.9393 | 7.8386E-07 | 2.6696E-06 |
| ENSG00000182459\|TEX19 | 46.4002 | 1.8318 | ##### | 4.9389 | 7.8556E-07 | 2.6751E-06 |
| ENSG00000244242\|IFITM10 | 40.6768 | -3.2200 | ##### | -4.9388 | 7.8595E-07 | 2.6761E-06 |
| ENSG00000198056\|PRIM1 | 3241.5138 | 0.6247 | ##### | 4.9386 | 7.8672E-07 | 2.6785E-06 |
| ENSG00000257663 | 1555.6858 | 0.6293 | ##### | 4.9386 | 7.8690E-07 | 2.6788E-06 |
| ENSG00000143224\|PPOX | 1223.3085 | 0.6259 | ##### | 4.9373 | 7.9218E-07 | 2.6964E-06 |
| ENSG00000228703 | 60.7810 | 1.3936 | ##### | 4.9370 | 7.9339E-07 | 2.6999E-06 |
| ENSG00000272572 | 36.3140 | -1.6538 | ##### | -4.9370 | 7.9331E-07 | 2.6999E-06 |
| ENSG00000162645\|GBP2 | 129.8985 | -4.2589 | ##### | -4.9368 | 7.9426E-07 | 2.7026E-06 |
| ENSG00000283294 | 52.9920 | -8.0876 | ##### | -4.9346 | 8.0321E-07 | 2.7327E-06 |
| ENSG00000174405\|LIG4 | 1622.0240 | -0.6247 | ##### | -4.9337 | 8.0686E-07 | 2.7448E-06 |
| ENSG00000236498 | 255.8906 | 0.8753 | ##### | 4.9326 | 8.1154E-07 | 2.7604E-06 |
| ENSG00000176986\|SEC24C | 15115.5766 | -0.5807 | ##### | -4.9318 | 8.1495E-07 | 2.7717E-06 |
| ENSG00000156427\|FGF18 | 67.1130 | 1.2558 | ##### | 4.9313 | 8.1673E-07 | 2.7775E-06 |
| ENSG00000151292\|CSNK1G3 | 3113.4757 | 0.6322 | ##### | 4.9295 | 8.2454E-07 | 2.8037E-06 |
| ENSG00000102935\|ZNF423 | 76.9000 | 7.9663 | ##### | 4.9294 | 8.2473E-07 | 2.8040E-06 |
| ENSG00000185338\|SOCS1 | 205.5327 | 0.9197 | ##### | 4.9294 | 8.2500E-07 | 2.8046E-06 |
| ENSG00000065923\|SLC9A7 | 2384.6764 | -0.6112 | ##### | -4.9278 | 8.3146E-07 | 2.8263E-06 |
| ENSG00000267940 | 17.9482 | 2.7565 | ##### | 4.9265 | 8.3733E-07 | 2.8459E-06 |
| ENSG00000213380\|COG8 | 3555.1274 | -0.6387 | ##### | -4.9264 | 8.3750E-07 | 2.8461E-06 |
| ENSG00000120438\|TCP1 | 28994.6896 | 0.5115 | ##### | 4.9258 | 8.4025E-07 | 2.8552E-06 |
| ENSG00000286905 | 141.3923 | 1.0951 | ##### | 4.9248 | 8.4432E-07 | 2.8687E-06 |
| ENSG00000117868\|ESYT2 | 15578.2661 | 0.4618 | ##### | 4.9247 | 8.4508E-07 | 2.8709E-06 |
| ENSG00000203523\|TAS2R2P | 22.7151 | -7.2447 | ##### | -4.9233 | 8.5080E-07 | 2.8900E-06 |
| ENSG00000091428\|RAPGEF4 | 93.3164 | 3.2540 | ##### | 4.9233 | 8.5114E-07 | 2.8909E-06 |
| ENSG00000168781\|PPIP5K1 | 1206.7217 | -0.6291 | ##### | -4.9232 | 8.5138E-07 | 2.8914E-06 |
| ENSG00000114770\|ABCC5 | 2807.9150 | -0.6282 | ##### | -4.9231 | 8.5198E-07 | 2.8931E-06 |
| ENSG00000183336\|BOLA2 | 1250.5559 | 0.6480 | ##### | 4.9222 | 8.5570E-07 | 2.9053E-06 |
| ENSG00000196502\|SULT1A1 | 204.4458 | -1.4318 | ##### | -4.9218 | 8.5772E-07 | 2.9119E-06 |
| ENSG00000125355\|TMEM255A | 65.6782 | 1.5610 | ##### | 4.9217 | 8.5782E-07 | 2.9119E-06 |
| ENSG00000184486\|POU3F2 | 44.4419 | 3.1504 | ##### | 4.9197 | 8.6688E-07 | 2.9423E-06 |
| ENSG00000178440\|TIMM23B-AGAP6 | 186.2624 | 1.4439 | ##### | 4.9196 | 8.6715E-07 | 2.9429E-06 |
| ENSG00000046647\|GEMIN8 | 534.2166 | -0.7470 | ##### | -4.9196 | 8.6741E-07 | 2.9434E-06 |
| ENSG00000174652\|ZNF266 | 1310.7798 | -0.6407 | ##### | -4.9189 | 8.7026E-07 | 2.9528E-06 |
| ENSG00000160752\|FDPS | 41809.8771 | -0.5704 | ##### | -4.9188 | 8.7066E-07 | 2.9538E-06 |
| ENSG00000290113 | 187.0609 | -9.9719 | ##### | -4.9182 | 8.7343E-07 | 2.9629E-06 |
| ENSG00000183010\|PYCR1 | 11649.5015 | -0.5903 | ##### | -4.9180 | 8.7425E-07 | 2.9653E-06 |
| ENSG00000259407 | 24.2960 | 2.3351 | ##### | 4.9170 | 8.7869E-07 | 2.9800E-06 |
| ENSG00000293107 | 75.5315 | -1.2712 | ##### | -4.9170 | 8.7879E-07 | 2.9800E-06 |
| ENSG00000112812\|PRSS16 | 263.6039 | 0.8884 | ##### | 4.9167 | 8.8009E-07 | 2.9841E-06 |
| ENSG00000225808\|DNAJC19P5 | 15.2738 | 3.9124 | ##### | 4.9160 | 8.8333E-07 | 2.9948E-06 |
| ENSG00000203288\|TDRKH-AS1 | 64.0660 | -1.4881 | ##### | -4.9158 | 8.8421E-07 | 2.9974E-06 |
| ENSG00000188785\|ZNF548 | 588.4510 | -0.7214 | ##### | -4.9157 | 8.8457E-07 | 2.9983E-06 |
| ENSG00000103319\|EEF2K | 4184.3221 | -0.5995 | ##### | -4.9154 | 8.8623E-07 | 3.0035E-06 |
| ENSG00000230204\|FTH1P5 | 13.0961 | -2.3080 | ##### | -4.9152 | 8.8706E-07 | 3.0060E-06 |
| ENSG00000168078\|PBK | 4241.7144 | 0.6071 | ##### | 4.9147 | 8.8902E-07 | 3.0123E-06 |
| ENSG00000149761\|NUDT22 | 2344.9387 | 0.6038 | ##### | 4.9134 | 8.9512E-07 | 3.0326E-06 |
| ENSG00000256001 | 175.5978 | -9.8808 | ##### | -4.9113 | 9.0489E-07 | 3.0651E-06 |
| ENSG00000278540\|ACACA | 8336.5789 | -0.5889 | ##### | -4.9113 | 9.0485E-07 | 3.0651E-06 |
| ENSG00000215251\|FASTKD5 | 2405.3886 | 0.5973 | ##### | 4.9109 | 9.0671E-07 | 3.0709E-06 |
| ENSG00000139546\|TARBP2 | 4092.6218 | 0.6161 | ##### | 4.9108 | 9.0698E-07 | 3.0714E-06 |

| | | | | | | |
|---|---|---|---|---|---|---|
| ENSG00000122035\|RASL11A | 615.6936 | 0.7497 | ##### | 4.9105 | 9.0853E-07 | 3.0763E-06 |
| ENSG00000263535\|AK4P1 | 255.2835 | 0.8567 | ##### | 4.9099 | 9.1115E-07 | 3.0849E-06 |
| ENSG00000242850\|RPL23AP68 | 28.3382 | -7.2493 | ##### | -4.9093 | 9.1407E-07 | 3.0944E-06 |
| ENSG00000145700\|ANKRD31 | 128.9857 | -1.4171 | ##### | -4.9080 | 9.2022E-07 | 3.1149E-06 |
| ENSG00000140406\|TLNRD1 | 3147.8688 | 0.5989 | ##### | 4.9077 | 9.2162E-07 | 3.1192E-06 |
| ENSG00000264443\|NCMAP-DT | 55.3213 | -1.4663 | ##### | -4.9076 | 9.2199E-07 | 3.1201E-06 |
| ENSG00000095713\|CRTAC1 | 34.9350 | 2.2465 | ##### | 4.9074 | 9.2297E-07 | 3.1227E-06 |
| ENSG00000258429\|PDF | 1090.0730 | -0.6795 | ##### | -4.9074 | 9.2292E-07 | 3.1227E-06 |
| ENSG00000170743\|SYT9 | 22.9454 | -7.4511 | ##### | -4.9073 | 9.2321E-07 | 3.1232E-06 |
| ENSG00000041515\|MYO16 | 129.1570 | -9.2452 | ##### | -4.9070 | 9.2462E-07 | 3.1276E-06 |
| ENSG00000185864\|NPIPB4 | 731.7020 | 0.7027 | ##### | 4.9067 | 9.2620E-07 | 3.1326E-06 |
| ENSG00000224126\|UBE2SP2 | 86.6382 | 1.0973 | ##### | 4.9060 | 9.2937E-07 | 3.1430E-06 |
| ENSG00000164287\|CDC20B | 38.1778 | -2.0605 | ##### | -4.9056 | 9.3144E-07 | 3.1496E-06 |
| ENSG00000196367\|TRRAP | 9314.1133 | 0.5889 | ##### | 4.9056 | 9.3157E-07 | 3.1497E-06 |
| ENSG00000205765\|RIMOC1 | 5209.7428 | 0.5903 | ##### | 4.9055 | 9.3199E-07 | 3.1508E-06 |
| ENSG00000289440 | 70.2649 | -1.2793 | ##### | -4.9054 | 9.3234E-07 | 3.1516E-06 |
| ENSG00000214736\|TOMM6 | 4345.5845 | 0.9215 | ##### | 4.9041 | 9.3837E-07 | 3.1716E-06 |
| ENSG00000237039\|RPS28P4 | 18.7375 | -2.0773 | ##### | -4.9039 | 9.3944E-07 | 3.1749E-06 |
| ENSG00000065613\|SLK | 8714.9817 | -0.5845 | ##### | -4.9023 | 9.4700E-07 | 3.2001E-06 |
| ENSG00000104047\|DTWD1 | 894.6519 | -0.6726 | ##### | -4.9023 | 9.4722E-07 | 3.2005E-06 |
| ENSG00000182578\|CSF1R | 39.0778 | -1.9461 | ##### | -4.9020 | 9.4888E-07 | 3.2057E-06 |
| ENSG00000205642\|VCX3B | 161.0283 | -9.7560 | ##### | -4.9017 | 9.5017E-07 | 3.2097E-06 |
| ENSG00000152620\|NADK2 | 5445.5363 | 0.5879 | ##### | 4.9015 | 9.5104E-07 | 3.2122E-06 |
| ENSG00000270091 | 12.0473 | 3.1203 | ##### | 4.9008 | 9.5461E-07 | 3.2240E-06 |
| ENSG00000249264\|EEF1A1P9 | 53.3047 | 1.2318 | ##### | 4.9006 | 9.5523E-07 | 3.2257E-06 |
| ENSG00000176399\|DMRTA1 | 193.5809 | -0.9550 | ##### | -4.9004 | 9.5651E-07 | 3.2296E-06 |
| ENSG00000230982\|DSTNP1 | 28.7409 | 2.1663 | ##### | 4.9002 | 9.5740E-07 | 3.2323E-06 |
| ENSG00000118473\|SGIP1 | 164.2618 | 8.5493 | ##### | 4.8990 | 9.6308E-07 | 3.2507E-06 |
| ENSG00000118017\|A4GNT | 159.7370 | -9.7444 | ##### | -4.8990 | 9.6307E-07 | 3.2507E-06 |
| ENSG00000254632 | 179.4674 | -9.9121 | ##### | -4.8983 | 9.6687E-07 | 3.2631E-06 |
| ENSG00000247157\|LINC01252 | 24.9174 | 2.1264 | ##### | 4.8980 | 9.6831E-07 | 3.2677E-06 |
| ENSG00000113712\|CSNK1A1 | 23917.4833 | -0.5379 | ##### | -4.8979 | 9.6866E-07 | 3.2684E-06 |
| ENSG00000286057 | 45.6577 | 4.9700 | ##### | 4.8976 | 9.7036E-07 | 3.2738E-06 |
| ENSG00000140093\|SERPINA10 | 28.2249 | 6.2281 | ##### | 4.8963 | 9.7639E-07 | 3.2938E-06 |
| ENSG00000283959 | 200.0564 | -0.8739 | ##### | -4.8951 | 9.8250E-07 | 3.3140E-06 |
| ENSG00000144843\|ADPRH | 558.9657 | -0.7141 | ##### | -4.8943 | 9.8659E-07 | 3.3274E-06 |
| ENSG00000293278 | 45.0011 | -1.8979 | ##### | -4.8937 | 9.8956E-07 | 3.3371E-06 |
| ENSG00000271698 | 57.1881 | -1.4215 | ##### | -4.8933 | 9.9152E-07 | 3.3433E-06 |
| ENSG00000030582\|GRN | 17972.1557 | 0.5776 | ##### | 4.8925 | 9.9587E-07 | 3.3576E-06 |
| ENSG00000110841\|PPFIBP1 | 9412.3768 | 0.5802 | ##### | 4.8920 | 9.9797E-07 | 3.3643E-06 |
| ENSG00000272851 | 201.1240 | -0.8439 | ##### | -4.8913 | 1.0015E-06 | 3.3757E-06 |
| ENSG00000093010\|COMT | 6594.5820 | -0.5943 | ##### | -4.8913 | 1.0019E-06 | 3.3767E-06 |
| ENSG00000088451\|TGDS | 1023.8238 | -0.6920 | ##### | -4.8910 | 1.0032E-06 | 3.3808E-06 |
| ENSG00000089157\|RPLP0 | ########## | 0.5908 | ##### | 4.8910 | 1.0035E-06 | 3.3813E-06 |
| ENSG00000166197\|NOLC1 | 30281.3805 | 0.4701 | ##### | 4.8901 | 1.0080E-06 | 3.3963E-06 |
| ENSG00000165029\|ABCA1 | 720.4442 | -0.6823 | ##### | -4.8896 | 1.0105E-06 | 3.4043E-06 |
| ENSG00000198814\|GK | 258.9803 | 0.8036 | ##### | 4.8886 | 1.0154E-06 | 3.4203E-06 |
| ENSG00000063177\|RPL18 | 43736.3115 | -0.5867 | ##### | -4.8875 | 1.0211E-06 | 3.4391E-06 |
| ENSG00000119777\|TMEM214 | 9050.7388 | -0.5946 | ##### | -4.8871 | 1.0233E-06 | 3.4463E-06 |
| ENSG00000139988\|RDH12 | 26.6305 | -3.7752 | ##### | -4.8863 | 1.0277E-06 | 3.4601E-06 |
| ENSG00000285744\|MITA1 | 159.6556 | -9.7436 | ##### | -4.8863 | 1.0276E-06 | 3.4601E-06 |
| ENSG00000233653\|CICP7 | 14.6926 | 2.2567 | ##### | 4.8861 | 1.0286E-06 | 3.4629E-06 |
| ENSG00000124151\|NCOA3 | 12018.8763 | -0.5855 | ##### | -4.8861 | 1.0287E-06 | 3.4629E-06 |
| ENSG00000133302\|SLF1 | 2099.5194 | 0.6306 | ##### | 4.8858 | 1.0300E-06 | 3.4668E-06 |
| ENSG00000171388\|APLN | 84.9508 | -1.4065 | ##### | -4.8857 | 1.0308E-06 | 3.4690E-06 |
| ENSG00000248996\|PDLIM7-AS1 | 15.8193 | 3.0332 | ##### | 4.8856 | 1.0309E-06 | 3.4692E-06 |
| ENSG00000010610\|CD4 | 20.2009 | 2.0868 | ##### | 4.8852 | 1.0332E-06 | 3.4763E-06 |
| ENSG00000225342\|LRRK2-DT | 16.6157 | -2.7504 | ##### | -4.8847 | 1.0361E-06 | 3.4858E-06 |
| ENSG00000257556\|LINC02298 | 372.9642 | -0.8342 | ##### | -4.8839 | 1.0402E-06 | 3.4992E-06 |
| ENSG00000289173 | 13.6238 | 4.6799 | ##### | 4.8814 | 1.0532E-06 | 3.5425E-06 |
| ENSG00000014164\|ZC3H3 | 3363.7295 | 0.6159 | ##### | 4.8812 | 1.0547E-06 | 3.5471E-06 |
| ENSG00000132182\|NUP210 | 4297.2792 | 0.5945 | ##### | 4.8810 | 1.0558E-06 | 3.5503E-06 |
| ENSG00000141574\|SECTM1 | 928.7691 | 0.6651 | ##### | 4.8801 | 1.0606E-06 | 3.5661E-06 |

| | | | | | | |
|---|---|---|---|---|---|---|
| ENSG00000168685\|IL7R | 585.5516 | 6.5927 | ##### | 4.8796 | 1.0630E-06 | 3.5738E-06 |
| ENSG00000258644\|SYNJ2BP-COX16 | 31.1067 | -2.8845 | ##### | -4.8792 | 1.0649E-06 | 3.5800E-06 |
| ENSG00000180773\|SLC36A4 | 3113.6646 | -0.6096 | ##### | -4.8784 | 1.0694E-06 | 3.5947E-06 |
| ENSG00000079432\|CIC | 6783.5992 | -0.5758 | ##### | -4.8782 | 1.0709E-06 | 3.5990E-06 |
| ENSG00000218175 | 246.5625 | 0.8772 | ##### | 4.8775 | 1.0746E-06 | 3.6114E-06 |
| ENSG00000264982 | 32.8379 | -1.6429 | ##### | -4.8773 | 1.0754E-06 | 3.6136E-06 |
| ENSG00000118217\|ATF6 | 8967.3874 | 0.5774 | ##### | 4.8760 | 1.0827E-06 | 3.6378E-06 |
| ENSG00000106636\|YKT6 | 20702.4719 | -0.3600 | ##### | -4.8757 | 1.0843E-06 | 3.6424E-06 |
| ENSG00000145414\|NAF1 | 1301.2460 | -0.6253 | ##### | -4.8749 | 1.0886E-06 | 3.6567E-06 |
| ENSG00000262050 | 71.0499 | -1.2770 | ##### | -4.8742 | 1.0924E-06 | 3.6689E-06 |
| ENSG00000185168\|LINC00482 | 52.6728 | -3.0394 | ##### | -4.8731 | 1.0987E-06 | 3.6899E-06 |
| ENSG00000185352\|HS6ST3 | 44.2129 | 3.7707 | ##### | 4.8730 | 1.0991E-06 | 3.6902E-06 |
| ENSG00000267309\|ZNF566-AS1 | 138.0972 | -1.2033 | ##### | -4.8730 | 1.0990E-06 | 3.6902E-06 |
| ENSG00000183604\|SMG1P5 | 467.0421 | 1.4074 | ##### | 4.8725 | 1.1018E-06 | 3.6989E-06 |
| ENSG00000088320\|REM1 | 61.3616 | -7.4977 | ##### | -4.8715 | 1.1075E-06 | 3.7177E-06 |
| ENSG00000271984 | 48.9855 | 2.3410 | ##### | 4.8712 | 1.1094E-06 | 3.7236E-06 |
| ENSG00000283674 | 90.6210 | 1.2804 | ##### | 4.8704 | 1.1138E-06 | 3.7379E-06 |
| ENSG00000267248 | 213.8300 | 0.8183 | ##### | 4.8701 | 1.1152E-06 | 3.7422E-06 |
| ENSG00000135637\|CCDC142 | 1634.9538 | 0.7317 | ##### | 4.8693 | 1.1201E-06 | 3.7582E-06 |
| ENSG00000085365\|SCAMP1 | 3139.7939 | -0.6058 | ##### | -4.8688 | 1.1230E-06 | 3.7676E-06 |
| ENSG00000005810\|MYCBP2 | 2940.2179 | 0.6417 | ##### | 4.8686 | 1.1241E-06 | 3.7707E-06 |
| ENSG00000277531\|PNMA8C | 29.3654 | -4.4470 | ##### | -4.8673 | 1.1312E-06 | 3.7942E-06 |
| ENSG00000211450\|SELENOH | 2552.1433 | -0.6170 | ##### | -4.8663 | 1.1369E-06 | 3.8124E-06 |
| ENSG00000104872\|PIH1D1 | 3890.6358 | 0.6037 | ##### | 4.8663 | 1.1369E-06 | 3.8124E-06 |
| ENSG00000258526 | 46.6674 | 2.1423 | ##### | 4.8662 | 1.1374E-06 | 3.8136E-06 |
| ENSG00000212724\|KRTAP2-3 | 192.6943 | 6.4242 | ##### | 4.8649 | 1.1451E-06 | 3.8390E-06 |
| ENSG00000211452\|DIO1 | 25.6561 | 7.6316 | ##### | 4.8646 | 1.1472E-06 | 3.8453E-06 |
| ENSG00000099942\|CRKL | 9710.6418 | 0.5698 | ##### | 4.8645 | 1.1472E-06 | 3.8453E-06 |
| ENSG00000269741 | 33.8357 | -6.0015 | ##### | -4.8645 | 1.1476E-06 | 3.8463E-06 |
| ENSG00000188811\|NHLRC3 | 664.9200 | -0.7564 | ##### | -4.8641 | 1.1500E-06 | 3.8538E-06 |
| ENSG00000181396\|OGFOD3 | 2935.4005 | -0.6019 | ##### | -4.8636 | 1.1529E-06 | 3.8630E-06 |
| ENSG00000271396 | 22.9211 | -6.5126 | ##### | -4.8629 | 1.1568E-06 | 3.8757E-06 |
| ENSG00000198681\|MAGEA1 | 488.7386 | ####### | ##### | -4.8619 | 1.1627E-06 | 3.8949E-06 |
| ENSG00000203865\|ATP1A1-AS1 | 746.5334 | 1.0296 | ##### | 4.8615 | 1.1649E-06 | 3.9019E-06 |
| ENSG00000254912\|MPHOSPH10P2 | 165.3889 | -9.7942 | ##### | -4.8612 | 1.1665E-06 | 3.9068E-06 |
| ENSG00000094804\|CDC6 | 9001.8358 | 0.5590 | ##### | 4.8612 | 1.1666E-06 | 3.9068E-06 |
| ENSG00000182919\|C11orf54 | 1833.8243 | -0.6162 | ##### | -4.8603 | 1.1722E-06 | 3.9252E-06 |
| ENSG00000233785\|SAT1-DT | 72.0303 | 3.4704 | ##### | 4.8577 | 1.1878E-06 | 3.9770E-06 |
| ENSG00000127314\|RAP1B | 17026.6860 | 0.5905 | ##### | 4.8575 | 1.1889E-06 | 3.9797E-06 |
| ENSG00000100605\|ITPK1 | 4783.4383 | -0.5787 | ##### | -4.8575 | 1.1889E-06 | 3.9797E-06 |
| ENSG00000168872\|DDX19A | 7185.5078 | -0.5047 | ##### | -4.8569 | 1.1923E-06 | 3.9906E-06 |
| ENSG00000117318\|ID3 | 32914.8559 | 0.5810 | ##### | 4.8562 | 1.1964E-06 | 4.0038E-06 |
| ENSG00000141526\|SLC16A3 | 10943.1718 | -0.5693 | ##### | -4.8531 | 1.2155E-06 | 4.0672E-06 |
| ENSG00000235631\|RNF148 | 78.2105 | -8.6216 | ##### | -4.8530 | 1.2162E-06 | 4.0694E-06 |
| ENSG00000162817\|C1orf115 | 407.8804 | -0.6888 | ##### | -4.8528 | 1.2175E-06 | 4.0729E-06 |
| ENSG00000293143 | 202.8702 | -0.8415 | ##### | -4.8519 | 1.2226E-06 | 4.0897E-06 |
| ENSG00000156521\|TYSND1 | 2334.9496 | -0.6119 | ##### | -4.8516 | 1.2248E-06 | 4.0961E-06 |
| ENSG00000054654\|SYNE2 | 2853.5456 | 0.6411 | ##### | 4.8516 | 1.2248E-06 | 4.0961E-06 |
| ENSG00000115526\|CHST10 | 2267.5051 | 0.5967 | ##### | 4.8514 | 1.2260E-06 | 4.0996E-06 |
| ENSG00000127947\|PTPN12 | 13768.8931 | 0.5198 | ##### | 4.8511 | 1.2277E-06 | 4.1050E-06 |
| ENSG00000126107\|HECTD3 | 3633.4293 | -0.6029 | ##### | -4.8510 | 1.2281E-06 | 4.1058E-06 |
| ENSG00000085231\|AK6 | 2055.0706 | 0.6259 | ##### | 4.8499 | 1.2354E-06 | 4.1296E-06 |
| ENSG00000196449\|YRDC | 6117.8484 | 0.5844 | ##### | 4.8498 | 1.2359E-06 | 4.1309E-06 |
| ENSG00000257181 | 96.4995 | 1.2905 | ##### | 4.8491 | 1.2403E-06 | 4.1451E-06 |
| ENSG00000288741 | 68.9817 | -1.5764 | ##### | -4.8491 | 1.2404E-06 | 4.1452E-06 |
| ENSG00000100554\|ATP6V1D | 5961.0806 | 0.5278 | ##### | 4.8489 | 1.2415E-06 | 4.1482E-06 |
| ENSG00000102158\|MAGT1 | 7206.2661 | -0.5840 | ##### | -4.8485 | 1.2438E-06 | 4.1555E-06 |
| ENSG00000053501\|USE1 | 855.6097 | -0.6479 | ##### | -4.8484 | 1.2445E-06 | 4.1573E-06 |
| ENSG00000186010\|NDUFA13 | 5523.3094 | 0.5901 | ##### | 4.8484 | 1.2447E-06 | 4.1576E-06 |
| ENSG00000160318\|CLDND2 | 63.3596 | -1.2282 | ##### | -4.8479 | 1.2474E-06 | 4.1663E-06 |
| ENSG00000078808\|SDF4 | 15169.0221 | -0.4469 | ##### | -4.8476 | 1.2497E-06 | 4.1735E-06 |
| ENSG00000267125 | 202.1643 | 0.9856 | ##### | 4.8468 | 1.2549E-06 | 4.1902E-06 |
| ENSG00000267565 | 9.9468 | -4.8834 | ##### | -4.8466 | 1.2561E-06 | 4.1937E-06 |

| | | | | | | |
|---|---|---|---|---|---|---|
| ENSG00000213420\|GPC2 | 211.0010 | 0.8993 | ##### | 4.8462 | 1.2587E-06 | 4.2018E-06 |
| ENSG00000279555 | 32.0546 | -6.7467 | ##### | -4.8442 | 1.2710E-06 | 4.2426E-06 |
| ENSG00000228436\|RRAGC-DT | 33.3930 | -2.1917 | ##### | -4.8437 | 1.2745E-06 | 4.2537E-06 |
| ENSG00000064489\|BORCS8-MEF2B | 431.0020 | -0.8768 | ##### | -4.8434 | 1.2763E-06 | 4.2593E-06 |
| ENSG00000124743\|KLHL31 | 103.9882 | -1.3769 | ##### | -4.8428 | 1.2799E-06 | 4.2710E-06 |
| ENSG00000288812 | 26.2293 | 3.8267 | ##### | 4.8422 | 1.2843E-06 | 4.2849E-06 |
| ENSG00000156232\|WHAMM | 1000.2570 | -0.6999 | ##### | -4.8420 | 1.2853E-06 | 4.2879E-06 |
| ENSG00000264278\|ZNF236-DT | 134.6311 | -1.0250 | ##### | -4.8419 | 1.2862E-06 | 4.2905E-06 |
| ENSG00000289281 | 34.1952 | -1.7083 | ##### | -4.8417 | 1.2872E-06 | 4.2932E-06 |
| ENSG00000139718\|SETD1B | 4169.5954 | 0.5942 | ##### | 4.8400 | 1.2986E-06 | 4.3307E-06 |
| ENSG00000167526\|RPL13 | 50139.7760 | -0.5754 | ##### | -4.8378 | 1.3129E-06 | 4.3780E-06 |
| ENSG00000289106 | 372.3617 | 0.7547 | ##### | 4.8373 | 1.3161E-06 | 4.3883E-06 |
| ENSG00000115415\|STAT1 | 20669.2148 | -0.5098 | ##### | -4.8335 | 1.3415E-06 | 4.4725E-06 |
| ENSG00000286614 | 216.3457 | -0.8699 | ##### | -4.8331 | 1.3442E-06 | 4.4809E-06 |
| ENSG00000250802\|ZBED3-AS1 | 106.1076 | 1.2169 | ##### | 4.8330 | 1.3450E-06 | 4.4832E-06 |
| ENSG00000188372\|ZP3 | 1432.4196 | -0.6956 | ##### | -4.8328 | 1.3464E-06 | 4.4874E-06 |
| ENSG00000132963\|POMP | 11368.5727 | 0.5850 | ##### | 4.8326 | 1.3474E-06 | 4.4900E-06 |
| ENSG00000183250\|LINC01547 | 428.8194 | -0.7313 | ##### | -4.8315 | 1.3549E-06 | 4.5145E-06 |
| ENSG00000131236\|CAP1 | 37747.6586 | -0.2691 | ##### | -4.8295 | 1.3688E-06 | 4.5603E-06 |
| ENSG00000166961\|MS4A15 | 23.8401 | -7.5066 | ##### | -4.8295 | 1.3690E-06 | 4.5606E-06 |
| ENSG00000236682\|MAP3K2-DT | 421.9248 | 0.7460 | ##### | 4.8291 | 1.3716E-06 | 4.5685E-06 |
| ENSG00000100246\|DNAL4 | 638.4956 | -0.7086 | ##### | -4.8282 | 1.3776E-06 | 4.5881E-06 |
| ENSG00000248839 | 16.8702 | 2.2944 | ##### | 4.8273 | 1.3838E-06 | 4.6082E-06 |
| ENSG00000133112\|TPT1 | 65365.6398 | 0.5183 | ##### | 4.8254 | 1.3974E-06 | 4.6532E-06 |
| ENSG00000048991\|R3HDM1 | 7133.7050 | -0.5853 | ##### | -4.8251 | 1.3994E-06 | 4.6593E-06 |
| ENSG00000149798\|CDC42EP2 | 552.1389 | -0.9328 | ##### | -4.8249 | 1.4008E-06 | 4.6632E-06 |
| ENSG00000120805\|ARL1 | 6047.3834 | -0.5897 | ##### | -4.8245 | 1.4038E-06 | 4.6728E-06 |
| ENSG00000109775\|UFSP2 | 1152.8310 | -0.6533 | ##### | -4.8241 | 1.4064E-06 | 4.6809E-06 |
| ENSG00000280594\|BTG3-AS1 | 113.7191 | 1.1424 | ##### | 4.8240 | 1.4074E-06 | 4.6836E-06 |
| ENSG00000287735 | 26.4684 | 3.9498 | ##### | 4.8237 | 1.4088E-06 | 4.6880E-06 |
| ENSG00000022556\|NLRP2 | 8238.5213 | -0.5688 | ##### | -4.8234 | 1.4110E-06 | 4.6946E-06 |
| ENSG00000182325\|FBXL6 | 5353.6816 | 0.6319 | ##### | 4.8220 | 1.4213E-06 | 4.7286E-06 |
| ENSG00000164162\|ANAPC10 | 556.4607 | -0.7959 | ##### | -4.8218 | 1.4228E-06 | 4.7329E-06 |
| ENSG00000139168\|ZCRB1 | 5393.6724 | 0.6047 | ##### | 4.8215 | 1.4247E-06 | 4.7385E-06 |
| ENSG00000157014\|TATDN2 | 5848.1010 | 0.5782 | ##### | 4.8205 | 1.4322E-06 | 4.7631E-06 |
| ENSG00000168938\|PPIC | 2854.4327 | -0.6282 | ##### | -4.8196 | 1.4384E-06 | 4.7830E-06 |
| ENSG00000149480\|MTA2 | 8516.0413 | 0.5735 | ##### | 4.8193 | 1.4408E-06 | 4.7905E-06 |
| ENSG00000169241\|SLC50A1 | 4083.4009 | 0.5840 | ##### | 4.8190 | 1.4430E-06 | 4.7975E-06 |
| ENSG00000205240\|OR7E36P | 15.6931 | 6.1185 | ##### | 4.8188 | 1.4442E-06 | 4.8009E-06 |
| ENSG00000136933\|RABEPK | 4249.3052 | 0.5915 | ##### | 4.8184 | 1.4469E-06 | 4.8093E-06 |
| ENSG00000273841\|TAF9 | 5056.8796 | 0.5997 | ##### | 4.8175 | 1.4539E-06 | 4.8319E-06 |
| ENSG00000168000\|BSCL2 | 1339.4553 | -1.1273 | ##### | -4.8165 | 1.4613E-06 | 4.8561E-06 |
| ENSG00000268041\|ERFL | 100.4474 | 1.2020 | ##### | 4.8161 | 1.4639E-06 | 4.8642E-06 |
| ENSG00000271966\|ARFGEF1-DT | 21.1132 | -2.4340 | ##### | -4.8159 | 1.4657E-06 | 4.8696E-06 |
| ENSG00000230565\|ZNF32-AS2 | 21.8858 | -3.0562 | ##### | -4.8154 | 1.4692E-06 | 4.8808E-06 |
| ENSG00000214021\|TTLL3 | 513.1646 | -0.7402 | ##### | -4.8142 | 1.4782E-06 | 4.9101E-06 |
| ENSG00000286431 | 224.5986 | -0.7760 | ##### | -4.8116 | 1.4975E-06 | 4.9736E-06 |
| ENSG00000184374\|COLEC10 | 40.4484 | 5.8754 | ##### | 4.8109 | 1.5026E-06 | 4.9900E-06 |
| ENSG00000052841\|TTC17 | 5327.3609 | -0.5810 | ##### | -4.8107 | 1.5040E-06 | 4.9940E-06 |
| ENSG00000117472\|TSPAN1 | 204.4326 | -0.9468 | ##### | -4.8099 | 1.5101E-06 | 5.0137E-06 |
| ENSG00000143155\|TIPRL | 4472.7403 | -0.5885 | ##### | -4.8095 | 1.5130E-06 | 5.0228E-06 |
| ENSG00000179242\|CDH4 | 614.4132 | 0.7752 | ##### | 4.8092 | 1.5152E-06 | 5.0295E-06 |
| ENSG00000078319\|PMS2P1 | 58.0629 | 2.9929 | ##### | 4.8089 | 1.5173E-06 | 5.0361E-06 |
| ENSG00000128408\|RIBC2 | 307.2571 | -0.7838 | ##### | -4.8089 | 1.5179E-06 | 5.0373E-06 |
| ENSG00000121879\|PIK3CA | 2492.1611 | -0.6209 | ##### | -4.8088 | 1.5187E-06 | 5.0396E-06 |
| ENSG00000238164\|TNFRSF14-AS1 | 55.7803 | 7.5298 | ##### | 4.8084 | 1.5214E-06 | 5.0479E-06 |
| ENSG00000186017\|ZNF566 | 1048.7854 | -0.6284 | ##### | -4.8075 | 1.5280E-06 | 5.0694E-06 |
| ENSG00000188529\|SRSF10 | 11446.6337 | 0.5568 | ##### | 4.8060 | 1.5397E-06 | 5.1076E-06 |
| ENSG00000111348\|ARHGDIB | 78.8028 | 2.4698 | ##### | 4.8053 | 1.5454E-06 | 5.1260E-06 |
| ENSG00000004059\|ARF5 | 10508.7613 | 0.5672 | ##### | 4.8050 | 1.5475E-06 | 5.1321E-06 |
| ENSG00000109101\|FOXN1 | 25.6329 | -7.1037 | ##### | -4.8049 | 1.5485E-06 | 5.1349E-06 |
| ENSG00000236137\|CD101-AS1 | 42.3195 | 1.4471 | ##### | 4.8029 | 1.5638E-06 | 5.1850E-06 |
| ENSG00000197933\|ZNF823 | 272.6930 | -0.7319 | ##### | -4.8029 | 1.5639E-06 | 5.1850E-06 |

| | | | | | | |
|---|---|---|---|---|---|---|
| ENSG00000163516\|ANKZF1 | 2652.2831 | 0.6044 | ##### | 4.8028 | 1.5648E-06 | 5.1875E-06 |
| ENSG00000210112\|MT-TM | 258.3389 | 0.8161 | ##### | 4.8023 | 1.5684E-06 | 5.1988E-06 |
| ENSG00000264575\|LINC00526 | 403.1887 | -0.7579 | ##### | -4.8006 | 1.5816E-06 | 5.2418E-06 |
| ENSG00000268309 | 78.4787 | -1.3651 | ##### | -4.8005 | 1.5824E-06 | 5.2439E-06 |
| ENSG00000141627\|DYM | 2557.2707 | -0.5453 | ##### | -4.8001 | 1.5856E-06 | 5.2540E-06 |
| ENSG00000119703\|ZC2HC1C | 347.4768 | 0.7252 | ##### | 4.7995 | 1.5910E-06 | 5.2712E-06 |
| ENSG00000214654\|B3GALT9 | 535.1579 | -0.7354 | ##### | -4.7993 | 1.5920E-06 | 5.2740E-06 |
| ENSG00000197223\|C1D | 2050.3856 | 0.6238 | ##### | 4.7993 | 1.5925E-06 | 5.2749E-06 |
| ENSG00000220161\|LINC02076 | 45.7687 | 1.4104 | ##### | 4.7990 | 1.5944E-06 | 5.2806E-06 |
| ENSG00000170502\|NUDT9 | 1713.0651 | -0.6096 | ##### | -4.7988 | 1.5961E-06 | 5.2858E-06 |
| ENSG00000165886\|UBTD1 | 1248.4506 | -0.6489 | ##### | -4.7981 | 1.6015E-06 | 5.3030E-06 |
| ENSG00000178623\|GPR35 | 159.1919 | -1.0939 | ##### | -4.7981 | 1.6020E-06 | 5.3042E-06 |
| ENSG00000124614\|RPS10 | 39522.2570 | 0.5925 | ##### | 4.7976 | 1.6055E-06 | 5.3151E-06 |
| ENSG00000108384\|RAD51C | 2044.5208 | 0.6201 | ##### | 4.7974 | 1.6077E-06 | 5.3220E-06 |
| ENSG00000206181\|ELOA2 | 23.0030 | -7.4531 | ##### | -4.7972 | 1.6087E-06 | 5.3246E-06 |
| ENSG00000066557\|LRRC40 | 2554.4685 | -0.6183 | ##### | -4.7963 | 1.6160E-06 | 5.3481E-06 |
| ENSG00000127870\|RNF6 | 5626.8211 | 0.5701 | ##### | 4.7960 | 1.6183E-06 | 5.3551E-06 |
| ENSG00000111644\|ACRBP | 43.3148 | 1.6278 | ##### | 4.7950 | 1.6264E-06 | 5.3815E-06 |
| ENSG00000240038\|AMY2B | 47.9142 | -3.7218 | ##### | -4.7949 | 1.6276E-06 | 5.3847E-06 |
| ENSG00000113643\|RARS1 | 11591.4644 | 0.5510 | ##### | 4.7944 | 1.6315E-06 | 5.3970E-06 |
| ENSG00000146352\|CLVS2 | 55.5807 | -1.8501 | ##### | -4.7936 | 1.6379E-06 | 5.4177E-06 |
| ENSG00000236263 | 63.7026 | 1.3189 | ##### | 4.7935 | 1.6392E-06 | 5.4214E-06 |
| ENSG00000110046\|ATG2A | 2725.8507 | 0.5668 | ##### | 4.7933 | 1.6405E-06 | 5.4251E-06 |
| ENSG00000248333\|CDK11B | 5753.1554 | 0.5632 | ##### | 4.7927 | 1.6458E-06 | 5.4418E-06 |
| ENSG00000175787\|ZNF169 | 184.0633 | 0.8231 | ##### | 4.7924 | 1.6478E-06 | 5.4480E-06 |
| ENSG00000100926\|TM9SF1 | 6434.2513 | 0.6031 | ##### | 4.7918 | 1.6529E-06 | 5.4644E-06 |
| ENSG00000214753\|HNRNPUL2 | 8421.9229 | 0.6033 | ##### | 4.7890 | 1.6765E-06 | 5.5412E-06 |
| ENSG00000267213\|DPY19L3-DT | 77.6319 | -1.0327 | ##### | -4.7890 | 1.6765E-06 | 5.5412E-06 |
| ENSG00000275084\|SNORD91B | 32.0459 | 7.0731 | ##### | 4.7878 | 1.6865E-06 | 5.5735E-06 |
| ENSG00000153902\|LGI4 | 67.9573 | 2.9124 | ##### | 4.7876 | 1.6877E-06 | 5.5768E-06 |
| ENSG00000291032 | 167.9902 | 3.5860 | ##### | 4.7875 | 1.6888E-06 | 5.5798E-06 |
| ENSG00000136247\|ZDHHC4 | 3691.5564 | -0.5948 | ##### | -4.7875 | 1.6890E-06 | 5.5800E-06 |
| ENSG00000135917\|SLC19A3 | 811.0237 | 0.6291 | ##### | 4.7874 | 1.6898E-06 | 5.5818E-06 |
| ENSG00000182185\|RAD51B | 352.5372 | 0.7277 | ##### | 4.7863 | 1.6987E-06 | 5.6107E-06 |
| ENSG00000095587\|TLL2 | 86.5253 | 1.6492 | ##### | 4.7862 | 1.6997E-06 | 5.6126E-06 |
| ENSG00000185633\|NDUFA4L2 | 48.9327 | -2.0229 | ##### | -4.7862 | 1.6996E-06 | 5.6126E-06 |
| ENSG00000233913\|RPL10P9 | 1044.1674 | 0.6671 | ##### | 4.7860 | 1.7010E-06 | 5.6165E-06 |
| ENSG00000092200\|RPGRIP1 | 26.9197 | -1.9354 | ##### | -4.7856 | 1.7047E-06 | 5.6280E-06 |
| ENSG00000101134\|DOK5 | 95.2825 | 4.8743 | ##### | 4.7841 | 1.7176E-06 | 5.6699E-06 |
| ENSG00000166261\|ZNF202 | 1434.4863 | 0.6247 | ##### | 4.7841 | 1.7180E-06 | 5.6705E-06 |
| ENSG00000139190\|VAMP1 | 878.2814 | 0.9677 | ##### | 4.7829 | 1.7279E-06 | 5.7025E-06 |
| ENSG00000174788\|PCP2 | 12.9869 | 4.2404 | ##### | 4.7827 | 1.7295E-06 | 5.7074E-06 |
| ENSG00000260947 | 114.9912 | -1.3741 | ##### | -4.7811 | 1.7430E-06 | 5.7514E-06 |
| ENSG00000291127\|ANKRD36BP2 | 128.0093 | 1.3430 | ##### | 4.7796 | 1.7562E-06 | 5.7941E-06 |
| ENSG00000132003\|ZSWIM4 | 1982.5733 | -0.6303 | ##### | -4.7791 | 1.7610E-06 | 5.8093E-06 |
| ENSG00000204160\|ZDHHC18 | 4599.1553 | -0.5960 | ##### | -4.7786 | 1.7648E-06 | 5.8212E-06 |
| ENSG00000145740\|SLC30A5 | 5716.7954 | -0.5148 | ##### | -4.7785 | 1.7662E-06 | 5.8251E-06 |
| ENSG00000077063\|CTTNBP2 | 120.4076 | 1.1621 | ##### | 4.7764 | 1.7850E-06 | 5.8865E-06 |
| ENSG00000117153\|KLHL12 | 3045.5880 | -0.5761 | ##### | -4.7759 | 1.7890E-06 | 5.8992E-06 |
| ENSG00000221838\|AP4M1 | 3158.4795 | 0.7190 | ##### | 4.7754 | 1.7931E-06 | 5.9119E-06 |
| ENSG00000269609\|C10orf95-AS1 | 352.2688 | -0.7795 | ##### | -4.7751 | 1.7961E-06 | 5.9204E-06 |
| ENSG00000134539\|KLRD1 | 30.5089 | 1.9588 | ##### | 4.7751 | 1.7962E-06 | 5.9204E-06 |
| ENSG00000198046\|ZNF667 | 383.4938 | -8.7843 | ##### | -4.7751 | 1.7959E-06 | 5.9204E-06 |
| ENSG00000118181\|RPS25 | 25316.8725 | 0.5738 | ##### | 4.7750 | 1.7967E-06 | 5.9213E-06 |
| ENSG00000210117\|MT-TW | 76.6203 | 1.1068 | ##### | 4.7748 | 1.7990E-06 | 5.9283E-06 |
| ENSG00000290911 | 34.9970 | 3.0446 | ##### | 4.7747 | 1.8001E-06 | 5.9313E-06 |
| ENSG00000236883 | 47.9767 | -1.3478 | ##### | -4.7740 | 1.8062E-06 | 5.9506E-06 |
| ENSG00000090863\|GLG1 | 12334.9226 | -0.5732 | ##### | -4.7738 | 1.8076E-06 | 5.9544E-06 |
| ENSG00000083844\|ZNF264 | 1868.5133 | -0.6023 | ##### | -4.7737 | 1.8084E-06 | 5.9567E-06 |
| ENSG00000172456\|FGGY | 377.5568 | -0.7679 | ##### | -4.7735 | 1.8107E-06 | 5.9635E-06 |
| ENSG00000185252\|ZNF74 | 1357.2892 | 0.6296 | ##### | 4.7702 | 1.8401E-06 | 6.0596E-06 |
| ENSG00000170242\|USP47 | 4504.2943 | -0.5721 | ##### | -4.7695 | 1.8471E-06 | 6.0822E-06 |
| ENSG00000127241\|MASP1 | 131.1467 | -6.4748 | ##### | -4.7691 | 1.8505E-06 | 6.0926E-06 |

| | | | | | | |
|---|---|---|---|---|---|---|
| ENSG00000156735\|BAG4 | 2270.5442 | 0.6139 | ##### | 4.7690 | 1.8512E-06 | 6.0941E-06 |
| ENSG00000288095 | 22.5416 | -4.5315 | ##### | -4.7685 | 1.8557E-06 | 6.1084E-06 |
| ENSG00000287387 | 246.8301 | 1.2767 | ##### | 4.7677 | 1.8638E-06 | 6.1341E-06 |
| ENSG00000280054 | 32.2114 | -2.2229 | ##### | -4.7675 | 1.8657E-06 | 6.1397E-06 |
| ENSG00000254870\|ATP6V1G2-DDX39B | 775.5186 | -2.1786 | ##### | -4.7662 | 1.8771E-06 | 6.1768E-06 |
| ENSG00000176788\|BASP1 | 37991.4301 | -0.5567 | ##### | -4.7655 | 1.8843E-06 | 6.1998E-06 |
| ENSG00000106038\|EVX1 | 127.5663 | -9.4197 | ##### | -4.7653 | 1.8856E-06 | 6.2033E-06 |
| ENSG00000268895\|A1BG-AS1 | 509.8740 | -1.0402 | ##### | -4.7652 | 1.8862E-06 | 6.2041E-06 |
| ENSG00000185049\|NELFA | 3372.8949 | 0.5896 | ##### | 4.7652 | 1.8862E-06 | 6.2041E-06 |
| ENSG00000283345\|MPHOSPH10P3 | 51.7822 | -8.6286 | ##### | -4.7647 | 1.8914E-06 | 6.2202E-06 |
| ENSG00000140263\|SORD | 5931.0050 | 0.5795 | ##### | 4.7646 | 1.8924E-06 | 6.2228E-06 |
| ENSG00000178105\|DDX10 | 3836.6927 | 0.4695 | ##### | 4.7645 | 1.8932E-06 | 6.2249E-06 |
| ENSG00000253898\|LINC01419 | 125.7168 | -9.3986 | ##### | -4.7639 | 1.8986E-06 | 6.2420E-06 |
| ENSG00000276672 | 195.5914 | 0.9334 | ##### | 4.7630 | 1.9070E-06 | 6.2690E-06 |
| ENSG00000154263\|ABCA10 | 85.5944 | 1.5395 | ##### | 4.7626 | 1.9108E-06 | 6.2806E-06 |
| ENSG00000061936\|SFSWAP | 4991.4025 | 0.5767 | ##### | 4.7626 | 1.9115E-06 | 6.2824E-06 |
| ENSG00000236859\|NIFK-AS1 | 508.3662 | -0.6977 | ##### | -4.7617 | 1.9197E-06 | 6.3086E-06 |
| ENSG00000259172\|SNRPA1-DT | 313.9521 | -1.4881 | ##### | -4.7611 | 1.9257E-06 | 6.3275E-06 |
| ENSG00000136940\|PDCL | 2166.9957 | -0.6083 | ##### | -4.7610 | 1.9263E-06 | 6.3287E-06 |
| ENSG00000233381\|AK4P3 | 206.6232 | 0.8833 | ##### | 4.7599 | 1.9372E-06 | 6.3638E-06 |
| ENSG00000196584\|XRCC2 | 2856.2578 | 0.5864 | ##### | 4.7586 | 1.9490E-06 | 6.4020E-06 |
| ENSG00000206028 | 17.9308 | 6.8375 | ##### | 4.7586 | 1.9497E-06 | 6.4034E-06 |
| ENSG00000272872 | 115.8779 | 1.1248 | ##### | 4.7573 | 1.9619E-06 | 6.4430E-06 |
| ENSG00000160932\|LY6E | 8497.0770 | 0.5990 | ##### | 4.7567 | 1.9676E-06 | 6.4610E-06 |
| ENSG00000268584 | 35.6925 | -2.4511 | ##### | -4.7561 | 1.9734E-06 | 6.4792E-06 |
| ENSG00000123064\|DDX54 | 16385.5757 | 0.5446 | ##### | 4.7556 | 1.9783E-06 | 6.4947E-06 |
| ENSG00000215544\|BCRP7 | 128.6562 | -9.4228 | ##### | -4.7555 | 1.9796E-06 | 6.4982E-06 |
| ENSG00000137841\|PLCB2 | 128.9738 | 1.1592 | ##### | 4.7553 | 1.9816E-06 | 6.5041E-06 |
| ENSG00000231087\|FDPSP7 | 9.2869 | -2.9506 | ##### | -4.7551 | 1.9834E-06 | 6.5094E-06 |
| ENSG00000087365\|SF3B2 | 18717.9492 | 0.4332 | ##### | 4.7547 | 1.9875E-06 | 6.5215E-06 |
| ENSG00000287996 | 133.1759 | -9.4815 | ##### | -4.7547 | 1.9876E-06 | 6.5215E-06 |
| ENSG00000185829\|ARL17A | 1528.8607 | 0.6287 | ##### | 4.7545 | 1.9890E-06 | 6.5254E-06 |
| ENSG00000163467\|TSACC | 111.9618 | 1.0544 | ##### | 4.7535 | 1.9995E-06 | 6.5591E-06 |
| ENSG00000253669\|GASAL1 | 434.2844 | 0.7744 | ##### | 4.7534 | 2.0004E-06 | 6.5615E-06 |
| ENSG00000155926\|SLA | 48.2612 | 5.6456 | ##### | 4.7531 | 2.0031E-06 | 6.5697E-06 |
| ENSG00000289305 | 115.0307 | -9.2706 | ##### | -4.7529 | 2.0054E-06 | 6.5763E-06 |
| ENSG00000130487\|KLHDC7B | 37.0551 | -2.8206 | ##### | -4.7528 | 2.0061E-06 | 6.5779E-06 |
| ENSG00000240731 | 110.7118 | 1.0461 | ##### | 4.7525 | 2.0096E-06 | 6.5889E-06 |
| ENSG00000153936\|HS2ST1 | 6394.1965 | -0.5821 | ##### | -4.7518 | 2.0158E-06 | 6.6083E-06 |
| ENSG00000204967\|PCDHA4 | 91.6325 | -1.6995 | ##### | -4.7504 | 2.0297E-06 | 6.6532E-06 |
| ENSG00000248858\|FLJ46284 | 120.5639 | -9.3382 | ##### | -4.7497 | 2.0367E-06 | 6.6755E-06 |
| ENSG00000160271\|RALGDS | 2110.2620 | 0.5902 | ##### | 4.7495 | 2.0390E-06 | 6.6823E-06 |
| ENSG00000003096\|KLHL13 | 203.4266 | 0.9006 | ##### | 4.7491 | 2.0428E-06 | 6.6938E-06 |
| ENSG00000260111 | 50.5687 | -1.1875 | ##### | -4.7477 | 2.0572E-06 | 6.7404E-06 |
| ENSG00000093144\|ECHDC1 | 3771.8106 | -0.5938 | ##### | -4.7467 | 2.0671E-06 | 6.7721E-06 |
| ENSG00000196167\|COLCA1 | 119.0139 | -9.3196 | ##### | -4.7465 | 2.0694E-06 | 6.7788E-06 |
| ENSG00000175294\|CATSPER1 | 19.7499 | -5.1710 | ##### | -4.7463 | 2.0720E-06 | 6.7867E-06 |
| ENSG00000169184\|MN1 | 101.6743 | -1.2543 | ##### | -4.7447 | 2.0883E-06 | 6.8393E-06 |
| ENSG00000163171\|CDC42EP3 | 1531.7981 | 0.6096 | ##### | 4.7440 | 2.0950E-06 | 6.8603E-06 |
| ENSG00000168883\|USP39 | 9024.1706 | 0.5133 | ##### | 4.7414 | 2.1220E-06 | 6.9482E-06 |
| ENSG00000166685\|COG1 | 3891.1678 | 0.6142 | ##### | 4.7414 | 2.1224E-06 | 6.9488E-06 |
| ENSG00000196337\|CGB7 | 119.8571 | 0.9450 | ##### | 4.7413 | 2.1233E-06 | 6.9508E-06 |
| ENSG00000133619\|KRBA1 | 860.9169 | 0.6411 | ##### | 4.7406 | 2.1313E-06 | 6.9764E-06 |
| ENSG00000124440\|HIF3A | 20.1059 | -3.6928 | ##### | -4.7405 | 2.1319E-06 | 6.9774E-06 |
| ENSG00000227486 | 39.7434 | -5.4649 | ##### | -4.7391 | 2.1466E-06 | 7.0248E-06 |
| ENSG00000279058\|AGAP14P | 29.5013 | -1.7487 | ##### | -4.7373 | 2.1659E-06 | 7.0873E-06 |
| ENSG00000077157\|PPP1R12B | 791.7290 | -0.6573 | ##### | -4.7372 | 2.1670E-06 | 7.0899E-06 |
| ENSG00000166762\|CATSPER2 | 77.7460 | -1.3427 | ##### | -4.7370 | 2.1693E-06 | 7.0967E-06 |
| ENSG00000152822\|GRM1 | 41.2761 | -7.9625 | ##### | -4.7367 | 2.1720E-06 | 7.1049E-06 |
| ENSG00000232202\|CHORDC1P1 | 25.7952 | 7.0053 | ##### | 4.7366 | 2.1731E-06 | 7.1077E-06 |
| ENSG00000287625 | 162.4990 | 0.9696 | ##### | 4.7366 | 2.1736E-06 | 7.1083E-06 |
| ENSG00000083799\|CYLD | 2481.4511 | -0.6137 | ##### | -4.7347 | 2.1934E-06 | 7.1724E-06 |
| ENSG00000244490\|RWDD4P1 | 84.0662 | -1.2692 | ##### | -4.7341 | 2.2000E-06 | 7.1933E-06 |

| | | | | | | |
|---|---|---|---|---|---|---|
| ENSG00000134461\|ANKRD16 | 577.5904 | 0.6737 | ##### | 4.7338 | 2.2035E-06 | 7.2040E-06 |
| ENSG00000147234\|FRMPD3 | 121.3329 | 4.1038 | ##### | 4.7338 | 2.2038E-06 | 7.2040E-06 |
| ENSG00000184343\|SRPK3 | 82.8482 | 1.3094 | ##### | 4.7334 | 2.2075E-06 | 7.2155E-06 |
| ENSG00000162975\|KCNF1 | 446.0701 | 0.7435 | ##### | 4.7304 | 2.2404E-06 | 7.3220E-06 |
| ENSG00000130449\|ZSWIM6 | 1903.3996 | 0.6053 | ##### | 4.7296 | 2.2501E-06 | 7.3531E-06 |
| ENSG00000290816 | 164.3029 | -1.4073 | ##### | -4.7286 | 2.2610E-06 | 7.3878E-06 |
| ENSG00000269929\|MIRLET7A1HG | 272.3119 | 0.8724 | ##### | 4.7282 | 2.2650E-06 | 7.4000E-06 |
| ENSG00000136381\|IREB2 | 5796.5434 | -0.5726 | ##### | -4.7271 | 2.2771E-06 | 7.4387E-06 |
| ENSG00000184584\|STING1 | 97.6925 | 1.1411 | ##### | 4.7269 | 2.2798E-06 | 7.4468E-06 |
| ENSG00000009950\|MLXIPL | 176.7469 | -0.9012 | ##### | -4.7261 | 2.2890E-06 | 7.4761E-06 |
| ENSG00000106617\|PRKAG2 | 3193.7754 | -0.3627 | ##### | -4.7259 | 2.2907E-06 | 7.4809E-06 |
| ENSG00000249790\|LINC02972 | 40.7846 | -5.7809 | ##### | -4.7255 | 2.2958E-06 | 7.4966E-06 |
| ENSG00000197746\|PSAP | 78477.7712 | -0.4707 | ##### | -4.7254 | 2.2966E-06 | 7.4984E-06 |
| ENSG00000127080\|IPPK | 2087.8934 | -0.6208 | ##### | -4.7245 | 2.3068E-06 | 7.5309E-06 |
| ENSG00000135333\|EPHA7 | 51.3141 | 4.7532 | ##### | 4.7242 | 2.3103E-06 | 7.5416E-06 |
| ENSG00000133997\|MED6 | 2875.6148 | 0.5896 | ##### | 4.7240 | 2.3122E-06 | 7.5470E-06 |
| ENSG00000292434\|KCNJ6-AS1 | 29.7925 | -5.5353 | ##### | -4.7229 | 2.3246E-06 | 7.5866E-06 |
| ENSG00000182885\|ADGRG3 | 124.1743 | -9.3806 | ##### | -4.7229 | 2.3249E-06 | 7.5868E-06 |
| ENSG00000163463\|KRTCAP2 | 9601.8155 | 0.5806 | ##### | 4.7220 | 2.3354E-06 | 7.6200E-06 |
| ENSG00000185340\|GAS2L1 | 4609.6112 | 0.5817 | ##### | 4.7187 | 2.3731E-06 | 7.7420E-06 |
| ENSG00000113946\|CLDN16 | 165.5658 | 2.9370 | ##### | 4.7187 | 2.3733E-06 | 7.7420E-06 |
| ENSG00000268001\|CARD8-AS1 | 72.2513 | -1.1819 | ##### | -4.7174 | 2.3888E-06 | 7.7917E-06 |
| ENSG00000261693 | 22.5263 | 2.3558 | ##### | 4.7171 | 2.3926E-06 | 7.8034E-06 |
| ENSG00000136717\|BIN1 | 5742.4975 | -0.5808 | ##### | -4.7167 | 2.3971E-06 | 7.8171E-06 |
| ENSG00000166801\|FAM111A | 1681.8386 | 0.6189 | ##### | 4.7157 | 2.4089E-06 | 7.8549E-06 |
| ENSG00000140015\|KCNH5 | 204.5108 | -7.8704 | ##### | -4.7156 | 2.4095E-06 | 7.8560E-06 |
| ENSG00000286178 | 92.2876 | 6.4610 | ##### | 4.7149 | 2.4182E-06 | 7.8833E-06 |
| ENSG00000290121 | 23.2969 | 3.3855 | ##### | 4.7139 | 2.4300E-06 | 7.9210E-06 |
| ENSG00000128340\|RAC2 | 45.8826 | 1.5831 | ##### | 4.7137 | 2.4331E-06 | 7.9303E-06 |
| ENSG00000272156 | 24.9695 | -2.7932 | ##### | -4.7130 | 2.4415E-06 | 7.9566E-06 |
| ENSG00000227252\|COPS8-DT | 92.0029 | -1.6587 | ##### | -4.7107 | 2.4691E-06 | 8.0449E-06 |
| ENSG00000271499 | 22.1288 | -7.3983 | ##### | -4.7107 | 2.4690E-06 | 8.0449E-06 |
| ENSG00000261587\|TMEM249 | 50.1889 | -1.6358 | ##### | -4.7085 | 2.4956E-06 | 8.1304E-06 |
| ENSG00000149823\|VPS51 | 5661.4771 | 0.5285 | ##### | 4.7082 | 2.4993E-06 | 8.1414E-06 |
| ENSG00000237978\|KCNMB2-AS1 | 33.3507 | -3.9090 | ##### | -4.7080 | 2.5019E-06 | 8.1493E-06 |
| ENSG00000109689\|STIM2 | 2400.3089 | 0.5741 | ##### | 4.7076 | 2.5070E-06 | 8.1649E-06 |
| ENSG00000183798\|EMILIN3 | 50.8501 | 1.3698 | ##### | 4.7075 | 2.5082E-06 | 8.1679E-06 |
| ENSG00000164330\|EBF1 | 103.5399 | -1.3646 | ##### | -4.7073 | 2.5106E-06 | 8.1749E-06 |
| ENSG00000254701 | 126.3849 | 1.0693 | ##### | 4.7069 | 2.5147E-06 | 8.1872E-06 |
| ENSG00000261455\|LINC01003 | 100.0851 | -1.1099 | ##### | -4.7062 | 2.5243E-06 | 8.2177E-06 |
| ENSG00000185559\|DLK1 | 22.9468 | 3.4666 | ##### | 4.7053 | 2.5353E-06 | 8.2524E-06 |
| ENSG00000253683 | 134.5614 | 0.9308 | ##### | 4.7052 | 2.5356E-06 | 8.2527E-06 |
| ENSG00000255435\|TTC36-AS1 | 13.2451 | -4.4576 | ##### | -4.7046 | 2.5432E-06 | 8.2765E-06 |
| ENSG00000105053\|VRK3 | 1950.7983 | -0.5979 | ##### | -4.7046 | 2.5439E-06 | 8.2770E-06 |
| ENSG00000125971\|DYNLRB1 | 4154.3292 | -0.6246 | ##### | -4.7046 | 2.5437E-06 | 8.2770E-06 |
| ENSG00000064655\|EYA2 | 150.6189 | -1.0293 | ##### | -4.7034 | 2.5590E-06 | 8.3250E-06 |
| ENSG00000057019\|DCBLD2 | 14660.1732 | -0.5630 | ##### | -4.7028 | 2.5666E-06 | 8.3490E-06 |
| ENSG00000229809\|ZNF688 | 442.8275 | -0.7279 | ##### | -4.7027 | 2.5671E-06 | 8.3497E-06 |
| ENSG00000159714\|ZDHHC1 | 1031.2844 | -0.6523 | ##### | -4.7017 | 2.5801E-06 | 8.3911E-06 |
| ENSG00000122643\|NT5C3A | 2786.8643 | -0.5909 | ##### | -4.7017 | 2.5805E-06 | 8.3915E-06 |
| ENSG00000259717\|LINC00677 | 24.6233 | -7.0444 | ##### | -4.7016 | 2.5811E-06 | 8.3923E-06 |
| ENSG00000080189\|SLC35C2 | 4785.5888 | 0.5260 | ##### | 4.7011 | 2.5871E-06 | 8.4110E-06 |
| ENSG00000005100\|DHX33 | 6289.1790 | 0.5790 | ##### | 4.7008 | 2.5913E-06 | 8.4236E-06 |
| ENSG00000253133\|GPAT4-AS1 | 1882.6024 | 0.8076 | ##### | 4.6999 | 2.6033E-06 | 8.4618E-06 |
| ENSG00000240254\|B4GALT4-AS1 | 8.9810 | -2.5317 | ##### | -4.6980 | 2.6271E-06 | 8.5384E-06 |
| ENSG00000124733\|MEA1 | 5677.2336 | 0.5832 | ##### | 4.6972 | 2.6371E-06 | 8.5699E-06 |
| ENSG00000121310\|ECHDC2 | 1666.8224 | -0.6225 | ##### | -4.6967 | 2.6436E-06 | 8.5899E-06 |
| ENSG00000259354\|SLC30A4-AS1 | 73.7439 | 1.3429 | ##### | 4.6965 | 2.6467E-06 | 8.5991E-06 |
| ENSG00000129007\|CALML4 | 215.1553 | 0.9684 | ##### | 4.6963 | 2.6498E-06 | 8.6082E-06 |
| ENSG00000064763\|FAR2 | 333.7159 | 0.8799 | ##### | 4.6961 | 2.6512E-06 | 8.6121E-06 |
| ENSG00000121058\|COIL | 2914.2362 | 0.5939 | ##### | 4.6961 | 2.6523E-06 | 8.6146E-06 |
| ENSG00000006744\|ELAC2 | 10357.5693 | 0.3593 | ##### | 4.6957 | 2.6576E-06 | 8.6307E-06 |
| ENSG00000289609 | 47.7230 | -1.7170 | ##### | -4.6942 | 2.6769E-06 | 8.6924E-06 |

| | | | | | | |
|---|---|---|---|---|---|---|
| ENSG00000216895\|RBM33-DT | 219.9883 | -0.8319 | ##### | -4.6933 | 2.6882E-06 | 8.7284E-06 |
| ENSG00000128590\|DNAJB9 | 1674.0589 | -0.6470 | ##### | -4.6932 | 2.6891E-06 | 8.7302E-06 |
| ENSG00000237773 | 86.0472 | -1.1707 | ##### | -4.6913 | 2.7149E-06 | 8.8131E-06 |
| ENSG00000177666\|PNPLA2 | 7365.1696 | 0.5718 | ##### | 4.6905 | 2.7256E-06 | 8.8469E-06 |
| ENSG00000106603\|COA1 | 4243.8286 | -0.5548 | ##### | -4.6898 | 2.7352E-06 | 8.8771E-06 |
| ENSG00000139116\|KIF21A | 6765.2916 | -0.5544 | ##### | -4.6880 | 2.7587E-06 | 8.9525E-06 |
| ENSG00000272933\|TRIM8-DT | 66.0552 | -1.1366 | ##### | -4.6876 | 2.7637E-06 | 8.9676E-06 |
| ENSG00000144485\|HES6 | 896.1811 | 0.6194 | ##### | 4.6870 | 2.7721E-06 | 8.9938E-06 |
| ENSG00000152767\|FARP1 | 3638.0217 | -0.7466 | ##### | -4.6867 | 2.7760E-06 | 9.0057E-06 |
| ENSG00000128791\|TWSG1 | 5555.2747 | 0.5771 | ##### | 4.6857 | 2.7900E-06 | 9.0501E-06 |
| ENSG00000132965\|ALOX5AP | 71.2906 | 1.2327 | ##### | 4.6855 | 2.7927E-06 | 9.0576E-06 |
| ENSG00000170370\|EMX2 | 28.2247 | 4.0158 | ##### | 4.6825 | 2.8338E-06 | 9.1899E-06 |
| ENSG00000164615\|CAMLG | 2113.1627 | 0.5769 | ##### | 4.6822 | 2.8376E-06 | 9.2015E-06 |
| ENSG00000204219\|TCEA3 | 202.4343 | -0.8205 | ##### | -4.6818 | 2.8432E-06 | 9.2187E-06 |
| ENSG00000237605\|DYRK3-AS1 | 23.3174 | 2.0947 | ##### | 4.6818 | 2.8435E-06 | 9.2187E-06 |
| ENSG00000122877\|EGR2 | 61.9234 | 3.1315 | ##### | 4.6815 | 2.8478E-06 | 9.2316E-06 |
| ENSG00000101337\|TM9SF4 | 8907.2593 | -0.5184 | ##### | -4.6811 | 2.8530E-06 | 9.2474E-06 |
| ENSG00000112787\|FBRSL1 | 8054.6773 | 0.5652 | ##### | 4.6808 | 2.8570E-06 | 9.2593E-06 |
| ENSG00000171055\|FEZ2 | 5038.7471 | -0.5720 | ##### | -4.6804 | 2.8635E-06 | 9.2792E-06 |
| ENSG00000143801\|PSEN2 | 2136.8847 | -0.6105 | ##### | -4.6803 | 2.8649E-06 | 9.2829E-06 |
| ENSG00000111266\|DUSP16 | 4675.2561 | -0.5749 | ##### | -4.6794 | 2.8766E-06 | 9.3196E-06 |
| ENSG00000279089 | 67.8920 | 1.2908 | ##### | 4.6792 | 2.8806E-06 | 9.3319E-06 |
| ENSG00000196663\|TECPR2 | 1722.5445 | -0.5960 | ##### | -4.6784 | 2.8906E-06 | 9.3632E-06 |
| ENSG00000167711\|SERPINF2 | 38.2070 | -1.5493 | ##### | -4.6783 | 2.8925E-06 | 9.3683E-06 |
| ENSG00000149564\|ESAM | 162.1359 | 0.9852 | ##### | 4.6769 | 2.9127E-06 | 9.4326E-06 |
| ENSG00000076201\|PTPN23 | 6723.2252 | 0.3752 | ##### | 4.6763 | 2.9209E-06 | 9.4581E-06 |
| ENSG00000160584\|SIK3 | 3833.0668 | 0.5589 | ##### | 4.6742 | 2.9504E-06 | 9.5528E-06 |
| ENSG00000034053\|APBA2 | 1103.5716 | -0.6342 | ##### | -4.6742 | 2.9516E-06 | 9.5556E-06 |
| ENSG00000151023\|ENKUR | 55.6873 | -1.3486 | ##### | -4.6734 | 2.9622E-06 | 9.5886E-06 |
| ENSG00000261226 | 22.9551 | 6.1889 | ##### | 4.6720 | 2.9827E-06 | 9.6540E-06 |
| ENSG00000110330\|BIRC2 | 10389.0026 | 0.5928 | ##### | 4.6715 | 2.9894E-06 | 9.6747E-06 |
| ENSG00000178028\|DMAP1 | 2661.0392 | -0.5847 | ##### | -4.6709 | 2.9986E-06 | 9.7035E-06 |
| ENSG00000213760\|ATP6V1G2 | 42.4086 | 1.6083 | ##### | 4.6707 | 3.0022E-06 | 9.7141E-06 |
| ENSG00000245146\|MALINC1 | 113.5367 | 1.2063 | ##### | 4.6704 | 3.0061E-06 | 9.7257E-06 |
| ENSG00000196873\|ZNG1C | 1245.1405 | -0.6163 | ##### | -4.6691 | 3.0257E-06 | 9.7881E-06 |
| ENSG00000260122 | 46.2813 | 3.2391 | ##### | 4.6688 | 3.0296E-06 | 9.7996E-06 |
| ENSG00000197858\|GPAA1 | 8998.7898 | -0.5587 | ##### | -4.6672 | 3.0535E-06 | 9.8756E-06 |
| ENSG00000225285\|LINC01770 | 23.3053 | -3.0291 | ##### | -4.6667 | 3.0604E-06 | 9.8969E-06 |
| ENSG00000291072 | 1710.7233 | 0.6076 | ##### | 4.6657 | 3.0754E-06 | 9.9443E-06 |
| ENSG00000169239\|CA5B | 390.0576 | -0.7299 | ##### | -4.6650 | 3.0865E-06 | 9.9791E-06 |
| ENSG00000143207\|COP1 | 2739.2054 | -0.5766 | ##### | -4.6642 | 3.0979E-06 | 1.0015E-05 |
| ENSG00000224848 | 46.1162 | -1.5125 | ##### | -4.6639 | 3.1020E-06 | 1.0027E-05 |
| ENSG00000180096\|SEPTIN1 | 104.5666 | -1.1565 | ##### | -4.6636 | 3.1076E-06 | 1.0044E-05 |
| ENSG00000260761 | 101.2233 | -9.0857 | ##### | -4.6631 | 3.1152E-06 | 1.0068E-05 |
| ENSG00000258708\|SLC25A21-AS1 | 71.1320 | -1.4546 | ##### | -4.6627 | 3.1201E-06 | 1.0083E-05 |
| ENSG00000185869\|ZNF829 | 496.8782 | -0.6815 | ##### | -4.6616 | 3.1376E-06 | 1.0138E-05 |
| ENSG00000165695\|AK8 | 43.8437 | -1.6796 | ##### | -4.6614 | 3.1407E-06 | 1.0147E-05 |
| ENSG00000179454\|KLHL28 | 1097.9399 | 0.6195 | ##### | 4.6613 | 3.1424E-06 | 1.0151E-05 |
| ENSG00000157107\|FCHO2 | 1745.5560 | -0.6036 | ##### | -4.6594 | 3.1710E-06 | 1.0243E-05 |
| ENSG00000136045\|PWP1 | 8611.4689 | 0.4343 | ##### | 4.6591 | 3.1762E-06 | 1.0258E-05 |
| ENSG00000228120 | 18.1764 | 2.5930 | ##### | 4.6588 | 3.1810E-06 | 1.0273E-05 |
| ENSG00000087191\|PSMC5 | 26058.4570 | 0.4224 | ##### | 4.6586 | 3.1842E-06 | 1.0282E-05 |
| ENSG00000196230\|TUBB | ########## | 0.4312 | ##### | 4.6584 | 3.1865E-06 | 1.0288E-05 |
| ENSG00000228784\|LINC00954 | 11.1277 | 6.2491 | ##### | 4.6577 | 3.1981E-06 | 1.0324E-05 |
| ENSG00000233966\|UBE2SP1 | 323.3222 | 0.8092 | ##### | 4.6574 | 3.2030E-06 | 1.0339E-05 |
| ENSG00000100038\|TOP3B | 2915.0746 | 0.8975 | ##### | 4.6560 | 3.2237E-06 | 1.0405E-05 |
| ENSG00000169299\|PGM2 | 2326.5030 | 0.6055 | ##### | 4.6555 | 3.2318E-06 | 1.0430E-05 |
| ENSG00000166979\|EVA1C | 214.9701 | -0.9087 | ##### | -4.6545 | 3.2472E-06 | 1.0478E-05 |
| ENSG00000274310 | 19.0840 | 6.1718 | ##### | 4.6540 | 3.2553E-06 | 1.0503E-05 |
| ENSG00000160813\|PPP1R35 | 4065.4995 | 0.5608 | ##### | 4.6537 | 3.2600E-06 | 1.0518E-05 |
| ENSG00000094880\|CDC23 | 4213.9444 | 0.5571 | ##### | 4.6535 | 3.2642E-06 | 1.0530E-05 |
| ENSG00000156009\|MAGEA8 | 185.1737 | -9.9280 | ##### | -4.6534 | 3.2645E-06 | 1.0530E-05 |
| ENSG00000272345 | 165.8902 | 0.9442 | ##### | 4.6526 | 3.2772E-06 | 1.0570E-05 |

| | | | | | | |
|---|---|---|---|---|---|---|
| ENSG00000135900\|MRPL44 | 4054.2613 | 0.6005 | ##### | 4.6526 | 3.2782E-06 | 1.0571E-05 |
| ENSG00000107611\|CUBN | 57.5762 | 3.7258 | ##### | 4.6525 | 3.2786E-06 | 1.0572E-05 |
| ENSG00000176771\|NCKAP5 | 83.0294 | 1.6354 | ##### | 4.6522 | 3.2838E-06 | 1.0587E-05 |
| ENSG00000166133\|RPUSD2 | 1183.5673 | 0.6023 | ##### | 4.6515 | 3.2959E-06 | 1.0625E-05 |
| ENSG00000231663\|COA6-AS1 | 70.1828 | -1.6980 | ##### | -4.6500 | 3.3192E-06 | 1.0699E-05 |
| ENSG00000293234 | 77.4955 | 6.8322 | ##### | 4.6498 | 3.3230E-06 | 1.0710E-05 |
| ENSG00000254852\|NPIPA2 | 121.3389 | 1.0910 | ##### | 4.6468 | 3.3708E-06 | 1.0863E-05 |
| ENSG00000277437 | 27.6328 | 1.7503 | ##### | 4.6459 | 3.3854E-06 | 1.0909E-05 |
| ENSG00000144837\|PLA1A | 113.8671 | -9.2552 | ##### | -4.6453 | 3.3954E-06 | 1.0940E-05 |
| ENSG00000280120 | 252.0221 | 0.8537 | ##### | 4.6440 | 3.4173E-06 | 1.1009E-05 |
| ENSG00000101489\|CELF4 | 31.4908 | -1.6381 | ##### | -4.6430 | 3.4333E-06 | 1.1060E-05 |
| ENSG00000101558\|VAPA | 15644.4164 | 0.4819 | ##### | 4.6429 | 3.4359E-06 | 1.1067E-05 |
| ENSG00000148690\|FRA10AC1 | 1913.9078 | -0.6113 | ##### | -4.6428 | 3.4374E-06 | 1.1070E-05 |
| ENSG00000214954\|LRRC69 | 85.7332 | -1.1051 | ##### | -4.6427 | 3.4381E-06 | 1.1072E-05 |
| ENSG00000187690\|EZHIP | 104.2497 | -9.1280 | ##### | -4.6426 | 3.4398E-06 | 1.1076E-05 |
| ENSG00000225899\|FRG2B | 22.5517 | 6.0063 | ##### | 4.6419 | 3.4523E-06 | 1.1115E-05 |
| ENSG00000107625\|DDX50 | 3735.3775 | -0.5743 | ##### | -4.6413 | 3.4631E-06 | 1.1149E-05 |
| ENSG00000258325\|ITFG2-AS1 | 21.4873 | 3.2412 | ##### | 4.6402 | 3.4812E-06 | 1.1206E-05 |
| ENSG00000167562\|ZNF701 | 502.6470 | -1.1918 | ##### | -4.6394 | 3.4935E-06 | 1.1244E-05 |
| ENSG00000196993\|NPIPB9 | 73.9773 | -1.0879 | ##### | -4.6375 | 3.5266E-06 | 1.1349E-05 |
| ENSG00000142192\|APP | 50424.0306 | -0.5088 | ##### | -4.6374 | 3.5275E-06 | 1.1351E-05 |
| ENSG00000255966\|GAPDH-DT | 22.9678 | 3.0095 | ##### | 4.6373 | 3.5291E-06 | 1.1355E-05 |
| ENSG00000174738\|NR1D2 | 2069.7088 | 0.5683 | ##### | 4.6371 | 3.5341E-06 | 1.1370E-05 |
| ENSG00000112394\|SLC16A10 | 51.9052 | 1.3400 | ##### | 4.6368 | 3.5382E-06 | 1.1382E-05 |
| ENSG00000234377\|OBI1-AS1 | 261.3645 | 0.8092 | ##### | 4.6358 | 3.5555E-06 | 1.1436E-05 |
| ENSG00000100290\|BIK | 93.1630 | -1.2257 | ##### | -4.6345 | 3.5772E-06 | 1.1505E-05 |
| ENSG00000232713\|RPS12P3 | 33.4702 | 1.8872 | ##### | 4.6341 | 3.5851E-06 | 1.1529E-05 |
| ENSG00000290948 | 924.1745 | -0.6025 | ##### | -4.6337 | 3.5923E-06 | 1.1551E-05 |
| ENSG00000043514\|TRIT1 | 2119.2267 | 0.5988 | ##### | 4.6331 | 3.6021E-06 | 1.1580E-05 |
| ENSG00000290535 | 286.0128 | -0.7476 | ##### | -4.6331 | 3.6019E-06 | 1.1580E-05 |
| ENSG00000123552\|USP45 | 1061.5509 | -0.6977 | ##### | -4.6320 | 3.6222E-06 | 1.1643E-05 |
| ENSG00000173376\|NDNF | 57.2690 | -2.9793 | ##### | -4.6314 | 3.6315E-06 | 1.1672E-05 |
| ENSG00000204052\|LRRC73 | 181.9217 | -0.8975 | ##### | -4.6305 | 3.6481E-06 | 1.1724E-05 |
| ENSG00000173715\|C11orf80 | 871.6584 | -0.6595 | ##### | -4.6303 | 3.6522E-06 | 1.1736E-05 |
| ENSG00000141179\|PCTP | 1179.9162 | -0.6018 | ##### | -4.6293 | 3.6685E-06 | 1.1787E-05 |
| ENSG00000266964\|FXYD1 | 37.3202 | 3.3698 | ##### | 4.6282 | 3.6888E-06 | 1.1851E-05 |
| ENSG00000146707\|POMZP3 | 1484.4402 | -0.5959 | ##### | -4.6281 | 3.6904E-06 | 1.1855E-05 |
| ENSG00000146410\|MTFR2 | 888.5520 | 0.6255 | ##### | 4.6265 | 3.7183E-06 | 1.1943E-05 |
| ENSG00000243364\|EFNA4 | 738.1566 | 0.6578 | ##### | 4.6265 | 3.7191E-06 | 1.1944E-05 |
| ENSG00000154144\|TBRG1 | 2697.7492 | 0.5764 | ##### | 4.6263 | 3.7220E-06 | 1.1952E-05 |
| ENSG00000293427 | 1050.8089 | 0.7258 | ##### | 4.6258 | 3.7317E-06 | 1.1982E-05 |
| ENSG00000166004\|CEP295 | 2046.0483 | 0.6046 | ##### | 4.6254 | 3.7394E-06 | 1.2006E-05 |
| ENSG00000257137\|LINC02874 | 22.7133 | -7.2983 | ##### | -4.6252 | 3.7423E-06 | 1.2014E-05 |
| ENSG00000254978\|ALG1L9P | 72.9885 | 1.1402 | ##### | 4.6251 | 3.7447E-06 | 1.2020E-05 |
| ENSG00000102038\|SMARCA1 | 7720.1046 | -0.5638 | ##### | -4.6235 | 3.7739E-06 | 1.2113E-05 |
| ENSG00000163257\|DCAF16 | 1742.0916 | 0.5788 | ##### | 4.6229 | 3.7832E-06 | 1.2141E-05 |
| ENSG00000249087\|ZNF436-AS1 | 110.3921 | 1.3767 | ##### | 4.6225 | 3.7912E-06 | 1.2164E-05 |
| ENSG00000291336 | 281.4959 | 0.9610 | ##### | 4.6225 | 3.7909E-06 | 1.2164E-05 |
| ENSG00000183148\|ANKRD20A2P | 109.3290 | -9.1858 | ##### | -4.6215 | 3.8094E-06 | 1.2221E-05 |
| ENSG00000141741\|MIEN1 | 3053.3312 | -0.4834 | ##### | -4.6209 | 3.8214E-06 | 1.2259E-05 |
| ENSG00000204272\|NBDY | 2001.7392 | -0.5919 | ##### | -4.6207 | 3.8241E-06 | 1.2266E-05 |
| ENSG00000111775\|COX6A1 | 15051.4480 | 0.5552 | ##### | 4.6201 | 3.8350E-06 | 1.2300E-05 |
| ENSG00000284395\|PERCC1 | 13.9118 | 3.5465 | ##### | 4.6193 | 3.8502E-06 | 1.2347E-05 |
| ENSG00000165646\|SLC18A2 | 199.6357 | -1.1625 | ##### | -4.6187 | 3.8608E-06 | 1.2379E-05 |
| ENSG00000105647\|PIK3R2 | 4828.4262 | -0.5468 | ##### | -4.6183 | 3.8689E-06 | 1.2404E-05 |
| ENSG00000125430\|HS3ST3B1 | 144.3418 | -0.9063 | ##### | -4.6171 | 3.8918E-06 | 1.2476E-05 |
| ENSG00000210195\|MT-TT | 236.1455 | 0.7548 | ##### | 4.6169 | 3.8947E-06 | 1.2484E-05 |
| ENSG00000135972\|MRPS9 | 5549.6196 | -0.5593 | ##### | -4.6163 | 3.9064E-06 | 1.2520E-05 |
| ENSG00000114026\|OGG1 | 1659.9744 | 0.6211 | ##### | 4.6138 | 3.9534E-06 | 1.2670E-05 |
| ENSG00000073614\|KDM5A | 5923.0381 | 0.5681 | ##### | 4.6134 | 3.9615E-06 | 1.2694E-05 |
| ENSG00000237281\|CATIP-AS2 | 32.5237 | -2.2614 | ##### | -4.6123 | 3.9824E-06 | 1.2760E-05 |
| ENSG00000186566\|GPATCH8 | 5815.1164 | 0.5616 | ##### | 4.6101 | 4.0245E-06 | 1.2893E-05 |
| ENSG00000174456\|C12orf76 | 663.3205 | -0.6649 | ##### | -4.6097 | 4.0329E-06 | 1.2919E-05 |

| | | | | | | |
|---|---|---|---|---|---|---|
| ENSG00000273314 | 574.4310 | 0.7044 | ##### | 4.6094 | 4.0374E-06 | 1.2932E-05 |
| ENSG00000204965\|PCDHA5 | 48.1141 | -5.0882 | ##### | -4.6092 | 4.0423E-06 | 1.2946E-05 |
| ENSG00000241525 | 35.9959 | 1.3501 | ##### | 4.6074 | 4.0768E-06 | 1.3055E-05 |
| ENSG00000267631\|CGB1 | 19.3195 | -2.5676 | ##### | -4.6058 | 4.1086E-06 | 1.3156E-05 |
| ENSG00000228824\|MIR4500HG | 47.8708 | -1.7316 | ##### | -4.6022 | 4.1798E-06 | 1.3382E-05 |
| ENSG00000167384\|ZNF180 | 990.1734 | -0.5265 | ##### | -4.6012 | 4.2016E-06 | 1.3451E-05 |
| ENSG00000114098\|ARMC8 | 3105.4051 | 0.5940 | ##### | 4.6011 | 4.2035E-06 | 1.3455E-05 |
| ENSG00000126254\|RBM42 | 10044.0385 | -0.5470 | ##### | -4.6008 | 4.2088E-06 | 1.3471E-05 |
| ENSG00000186193\|SAPCD2 | 4847.5932 | 0.5516 | ##### | 4.5991 | 4.2429E-06 | 1.3578E-05 |
| ENSG00000149295\|DRD2 | 100.1574 | 8.3421 | ##### | 4.5979 | 4.2676E-06 | 1.3656E-05 |
| ENSG00000232450 | 60.3487 | 1.1923 | ##### | 4.5974 | 4.2774E-06 | 1.3686E-05 |
| ENSG00000227258\|SMIM2-AS1 | 90.4552 | 1.1947 | ##### | 4.5973 | 4.2809E-06 | 1.3696E-05 |
| ENSG00000146859\|TMEM140 | 1733.5602 | -0.5950 | ##### | -4.5970 | 4.2867E-06 | 1.3712E-05 |
| ENSG00000236268\|LINC01361 | 87.8343 | -8.8809 | ##### | -4.5970 | 4.2871E-06 | 1.3712E-05 |
| ENSG00000189280\|GJB5 | 156.1729 | -9.6917 | ##### | -4.5968 | 4.2898E-06 | 1.3720E-05 |
| ENSG00000227725\|POLR2MP1 | 18.2855 | -2.0576 | ##### | -4.5968 | 4.2908E-06 | 1.3721E-05 |
| ENSG00000239335\|LLPH-DT | 60.6230 | 1.1941 | ##### | 4.5965 | 4.2968E-06 | 1.3739E-05 |
| ENSG00000256061\|DNAAF4 | 296.8305 | -0.8031 | ##### | -4.5964 | 4.2988E-06 | 1.3744E-05 |
| ENSG00000261572 | 34.9025 | -1.9824 | ##### | -4.5963 | 4.3011E-06 | 1.3750E-05 |
| ENSG00000134330\|IAH1 | 3632.4733 | -0.5783 | ##### | -4.5962 | 4.3028E-06 | 1.3754E-05 |
| ENSG00000095485\|CWF19L1 | 2898.9377 | -0.5719 | ##### | -4.5959 | 4.3080E-06 | 1.3769E-05 |
| ENSG00000246575 | 19.9572 | 2.3822 | ##### | 4.5959 | 4.3090E-06 | 1.3771E-05 |
| ENSG00000227080 | 11.8689 | 4.9266 | ##### | 4.5957 | 4.3132E-06 | 1.3783E-05 |
| ENSG00000125538\|IL1B | 103.1088 | -8.8456 | ##### | -4.5929 | 4.3715E-06 | 1.3967E-05 |
| ENSG00000136866\|ZFP37 | 429.7225 | -0.7787 | ##### | -4.5929 | 4.3718E-06 | 1.3967E-05 |
| ENSG00000287817 | 27.1854 | -1.6754 | ##### | -4.5928 | 4.3735E-06 | 1.3971E-05 |
| ENSG00000259442 | 24.1575 | 6.6510 | ##### | 4.5907 | 4.4182E-06 | 1.4112E-05 |
| ENSG00000197150\|ABCB8 | 4304.2180 | 0.3692 | ##### | 4.5901 | 4.4294E-06 | 1.4147E-05 |
| ENSG00000167491\|GATAD2A | 8325.7645 | 0.4380 | ##### | 4.5899 | 4.4341E-06 | 1.4160E-05 |
| ENSG00000240344\|PPIL3 | 785.6718 | 0.6404 | ##### | 4.5899 | 4.4347E-06 | 1.4160E-05 |
| ENSG00000214694\|ARHGEF33 | 168.6800 | -1.0150 | ##### | -4.5884 | 4.4672E-06 | 1.4263E-05 |
| ENSG00000267748 | 128.2008 | -9.0365 | ##### | -4.5883 | 4.4688E-06 | 1.4266E-05 |
| ENSG00000143889\|HNRNPLL | 5506.1105 | 0.5665 | ##### | 4.5869 | 4.4985E-06 | 1.4359E-05 |
| ENSG00000261801\|LOXL1-AS1 | 313.3544 | -0.7765 | ##### | -4.5863 | 4.5122E-06 | 1.4402E-05 |
| ENSG00000225205\|PDK1-AS1 | 36.0559 | 1.5928 | ##### | 4.5861 | 4.5153E-06 | 1.4410E-05 |
| ENSG00000100350\|FOXRED2 | 2366.6494 | -0.5253 | ##### | -4.5859 | 4.5199E-06 | 1.4423E-05 |
| ENSG00000162736\|NCSTN | 11678.6947 | 0.4174 | ##### | 4.5853 | 4.5326E-06 | 1.4461E-05 |
| ENSG00000140750\|ARHGAP17 | 6092.8528 | 0.5718 | ##### | 4.5853 | 4.5323E-06 | 1.4461E-05 |
| ENSG00000166482\|MFAP4 | 41.5571 | -1.4349 | ##### | -4.5846 | 4.5481E-06 | 1.4509E-05 |
| ENSG00000169884\|WNT10B | 403.3640 | -0.6728 | ##### | -4.5845 | 4.5513E-06 | 1.4517E-05 |
| ENSG00000205045\|SLFN12L | 26.1078 | 4.2975 | ##### | 4.5843 | 4.5547E-06 | 1.4527E-05 |
| ENSG00000214223\|HNRNPA1P10 | 485.8978 | 0.6485 | ##### | 4.5841 | 4.5593E-06 | 1.4540E-05 |
| ENSG00000073536\|NLE1 | 4313.4425 | 0.5533 | ##### | 4.5841 | 4.5603E-06 | 1.4541E-05 |
| ENSG00000198331\|HYLS1 | 1172.5980 | 0.5958 | ##### | 4.5839 | 4.5639E-06 | 1.4551E-05 |
| ENSG00000168779\|SHOX2 | 696.9063 | 0.6454 | ##### | 4.5837 | 4.5681E-06 | 1.4563E-05 |
| ENSG00000100028\|SNRPD3 | 8313.9702 | 0.5120 | ##### | 4.5834 | 4.5752E-06 | 1.4584E-05 |
| ENSG00000185436\|IFNLR1 | 145.7198 | 1.1439 | ##### | 4.5820 | 4.6061E-06 | 1.4679E-05 |
| ENSG00000203666\|EFCAB2 | 890.1874 | -0.6253 | ##### | -4.5820 | 4.6056E-06 | 1.4679E-05 |
| ENSG00000128534\|LSM8 | 2544.3874 | 0.5820 | ##### | 4.5818 | 4.6093E-06 | 1.4688E-05 |
| ENSG00000102786\|INTS6 | 4042.2233 | 0.5711 | ##### | 4.5813 | 4.6200E-06 | 1.4721E-05 |
| ENSG00000108592\|FTSJ3 | 12538.7779 | 0.5440 | ##### | 4.5806 | 4.6359E-06 | 1.4770E-05 |
| ENSG00000141522\|ARHGDIA | 25528.1343 | -0.5330 | ##### | -4.5804 | 4.6420E-06 | 1.4787E-05 |
| ENSG00000165526\|RPUSD4 | 1999.2880 | 0.5612 | ##### | 4.5803 | 4.6430E-06 | 1.4789E-05 |
| ENSG00000101224\|CDC25B | 11154.1828 | 0.5437 | ##### | 4.5803 | 4.6434E-06 | 1.4789E-05 |
| ENSG00000139304\|PTPRQ | 127.1260 | -4.5846 | ##### | -4.5802 | 4.6463E-06 | 1.4797E-05 |
| ENSG00000261286\|ATP2C2-AS1 | 133.7261 | -1.1869 | ##### | -4.5801 | 4.6472E-06 | 1.4798E-05 |
| ENSG00000255441\|SIGLEC10-AS1 | 28.1534 | -1.6516 | ##### | -4.5799 | 4.6524E-06 | 1.4813E-05 |
| ENSG00000185507\|IRF7 | 275.5104 | 0.8611 | ##### | 4.5790 | 4.6724E-06 | 1.4875E-05 |
| ENSG00000178921\|PFAS | 8742.5148 | 0.5453 | ##### | 4.5786 | 4.6816E-06 | 1.4902E-05 |
| ENSG00000177030\|DEAF1 | 1378.7044 | -0.6181 | ##### | -4.5783 | 4.6870E-06 | 1.4918E-05 |
| ENSG00000180448\|ARHGAP45 | 3476.4658 | 0.5539 | ##### | 4.5775 | 4.7057E-06 | 1.4976E-05 |
| ENSG00000185158\|LRRC37B | 518.6146 | -0.6766 | ##### | -4.5773 | 4.7093E-06 | 1.4986E-05 |
| ENSG00000162227\|TAF6L | 1492.9738 | 0.6036 | ##### | 4.5772 | 4.7126E-06 | 1.4993E-05 |

| | | | | | | |
|---|---|---|---|---|---|---|
| ENSG00000145734\|BDP1 | 4138.5138 | 0.5921 | ##### | 4.5772 | 4.7125E-06 | 1.4993E-05 |
| ENSG00000198791\|CNOT7 | 7651.6829 | 0.5572 | ##### | 4.5766 | 4.7263E-06 | 1.5035E-05 |
| ENSG00000106701\|FSD1L | 706.7099 | -0.6717 | ##### | -4.5758 | 4.7433E-06 | 1.5088E-05 |
| ENSG00000260871 | 11.1613 | 6.2530 | ##### | 4.5746 | 4.7712E-06 | 1.5175E-05 |
| ENSG00000219529 | 61.5745 | 1.9718 | ##### | 4.5745 | 4.7734E-06 | 1.5180E-05 |
| ENSG00000169679\|BUB1 | 11122.3331 | 0.5465 | ##### | 4.5736 | 4.7949E-06 | 1.5247E-05 |
| ENSG00000288765 | 23.5095 | -3.9419 | ##### | -4.5727 | 4.8137E-06 | 1.5305E-05 |
| ENSG00000169220\|RGS14 | 746.3963 | -0.6441 | ##### | -4.5722 | 4.8271E-06 | 1.5346E-05 |
| ENSG00000170191\|NANP | 1063.8937 | -0.6062 | ##### | -4.5716 | 4.8403E-06 | 1.5386E-05 |
| ENSG00000116285\|ERRFI1 | 27009.1050 | -0.3255 | ##### | -4.5706 | 4.8638E-06 | 1.5460E-05 |
| ENSG00000071994\|PDCD2 | 5367.1689 | -0.5245 | ##### | -4.5700 | 4.8767E-06 | 1.5498E-05 |
| ENSG00000274718 | 120.5644 | -9.2776 | ##### | -4.5700 | 4.8769E-06 | 1.5498E-05 |
| ENSG00000181061\|HIGD1A | 7076.1690 | 0.5369 | ##### | 4.5700 | 4.8775E-06 | 1.5498E-05 |
| ENSG00000108389\|MTMR4 | 3375.7490 | 0.5720 | ##### | 4.5699 | 4.8789E-06 | 1.5501E-05 |
| ENSG00000182504\|CEP97 | 2178.3703 | 0.5926 | ##### | 4.5687 | 4.9083E-06 | 1.5593E-05 |
| ENSG00000197872\|CYRIA | 41.5222 | -5.1311 | ##### | -4.5686 | 4.9095E-06 | 1.5595E-05 |
| ENSG00000257218\|GATC | 12225.0633 | 0.5570 | ##### | 4.5681 | 4.9211E-06 | 1.5630E-05 |
| ENSG00000235990\|RPL23AP20 | 21.4099 | 5.8767 | ##### | 4.5677 | 4.9302E-06 | 1.5657E-05 |
| ENSG00000154957\|ZNF18 | 419.8001 | 0.6850 | ##### | 4.5677 | 4.9316E-06 | 1.5660E-05 |
| ENSG00000280311 | 20.3714 | 3.8097 | ##### | 4.5675 | 4.9365E-06 | 1.5674E-05 |
| ENSG00000276603 | 20.0760 | 2.6455 | ##### | 4.5674 | 4.9373E-06 | 1.5675E-05 |
| ENSG00000115641\|FHL2 | 9021.4371 | 0.5711 | ##### | 4.5671 | 4.9463E-06 | 1.5702E-05 |
| ENSG00000127804\|METTL16 | 4693.6186 | 0.5586 | ##### | 4.5666 | 4.9563E-06 | 1.5732E-05 |
| ENSG00000282381 | 108.8845 | 8.7152 | ##### | 4.5663 | 4.9651E-06 | 1.5758E-05 |
| ENSG00000111912\|NCOA7 | 16558.7777 | 0.5406 | ##### | 4.5645 | 5.0067E-06 | 1.5888E-05 |
| ENSG00000267232 | 66.1217 | 1.2330 | ##### | 4.5645 | 5.0080E-06 | 1.5891E-05 |
| ENSG00000100034\|PPM1F | 6985.6475 | 0.5392 | ##### | 4.5642 | 5.0150E-06 | 1.5911E-05 |
| ENSG00000161980\|POLR3K | 1983.9496 | -0.6255 | ##### | -4.5633 | 5.0348E-06 | 1.5972E-05 |
| ENSG00000100711\|ZFYVE21 | 1184.6781 | -0.6141 | ##### | -4.5633 | 5.0364E-06 | 1.5976E-05 |
| ENSG00000123352\|SPATS2 | 5049.1884 | -0.4364 | ##### | -4.5618 | 5.0706E-06 | 1.6083E-05 |
| ENSG00000233461 | 621.1620 | -0.6844 | ##### | -4.5602 | 5.1105E-06 | 1.6207E-05 |
| ENSG00000003756\|RBM5 | 5960.3742 | 0.5347 | ##### | 4.5599 | 5.1172E-06 | 1.6227E-05 |
| ENSG00000135829\|DHX9 | 25723.8668 | 0.5389 | ##### | 4.5598 | 5.1207E-06 | 1.6236E-05 |
| ENSG00000293331 | 81.1301 | 1.0696 | ##### | 4.5594 | 5.1293E-06 | 1.6262E-05 |
| ENSG00000263412\|NFE2L1-DT | 68.9262 | 1.1707 | ##### | 4.5590 | 5.1401E-06 | 1.6294E-05 |
| ENSG00000179097\|HTR1F | 59.6272 | -2.4871 | ##### | -4.5586 | 5.1496E-06 | 1.6323E-05 |
| ENSG00000168140\|VASN | 579.3612 | -0.6639 | ##### | -4.5585 | 5.1514E-06 | 1.6327E-05 |
| ENSG00000121101\|TEX14 | 227.3300 | 4.3230 | ##### | 4.5573 | 5.1818E-06 | 1.6420E-05 |
| ENSG00000159079\|CFAP298 | 4517.6660 | 0.5585 | ##### | 4.5573 | 5.1815E-06 | 1.6420E-05 |
| ENSG00000076662\|ICAM3 | 967.5836 | 0.7229 | ##### | 4.5568 | 5.1948E-06 | 1.6459E-05 |
| ENSG00000078487\|ZCWPW1 | 509.4746 | 0.8365 | ##### | 4.5565 | 5.2010E-06 | 1.6477E-05 |
| ENSG00000217624\|YWHAZP10 | 31.7325 | -1.4654 | ##### | -4.5562 | 5.2086E-06 | 1.6499E-05 |
| ENSG00000217078 | 22.0125 | 5.3505 | ##### | 4.5554 | 5.2287E-06 | 1.6561E-05 |
| ENSG00000148655\|LRMDA | 392.1979 | ####### | ##### | -4.5553 | 5.2311E-06 | 1.6567E-05 |
| ENSG00000134905\|CARS2 | 2519.2325 | -0.5802 | ##### | -4.5552 | 5.2335E-06 | 1.6573E-05 |
| ENSG00000081087\|OSTM1 | 5031.9062 | -0.5668 | ##### | -4.5550 | 5.2394E-06 | 1.6590E-05 |
| ENSG00000102805\|CLN5 | 989.6205 | -0.6150 | ##### | -4.5548 | 5.2442E-06 | 1.6603E-05 |
| ENSG00000100445\|SDR39U1 | 1392.3591 | 0.9051 | ##### | 4.5547 | 5.2449E-06 | 1.6604E-05 |
| ENSG00000165434\|PGM2L1 | 1491.2943 | 0.6022 | ##### | 4.5544 | 5.2543E-06 | 1.6632E-05 |
| ENSG00000260273 | 37.4687 | 1.6044 | ##### | 4.5533 | 5.2816E-06 | 1.6716E-05 |
| ENSG00000236234 | 15.4084 | 5.8342 | ##### | 4.5530 | 5.2887E-06 | 1.6737E-05 |
| ENSG00000120333\|MRPS14 | 1395.0959 | -0.6265 | ##### | -4.5524 | 5.3045E-06 | 1.6785E-05 |
| ENSG00000134371\|CDC73 | 6155.4720 | -0.4575 | ##### | -4.5518 | 5.3177E-06 | 1.6825E-05 |
| ENSG00000148291\|SURF2 | 2069.5376 | 0.6135 | ##### | 4.5518 | 5.3182E-06 | 1.6825E-05 |
| ENSG00000235023 | 32.3675 | 6.7240 | ##### | 4.5517 | 5.3207E-06 | 1.6831E-05 |
| ENSG00000148719\|DNAJB12 | 3835.4112 | -0.5601 | ##### | -4.5511 | 5.3372E-06 | 1.6882E-05 |
| ENSG00000227827\|PKD1P2 | 935.7537 | 0.6662 | ##### | 4.5503 | 5.3561E-06 | 1.6939E-05 |
| ENSG00000196204\|RNF216P1 | 1998.5194 | -0.5942 | ##### | -4.5501 | 5.3618E-06 | 1.6956E-05 |
| ENSG00000172349\|IL16 | 42.0353 | 2.3707 | ##### | 4.5499 | 5.3680E-06 | 1.6973E-05 |
| ENSG00000290013 | 17.5066 | 5.9612 | ##### | 4.5495 | 5.3782E-06 | 1.7004E-05 |
| ENSG00000215126\|ZNG1F | 930.2157 | -0.6587 | ##### | -4.5492 | 5.3863E-06 | 1.7028E-05 |
| ENSG00000135605\|TEC | 309.0337 | -0.6883 | ##### | -4.5480 | 5.4167E-06 | 1.7122E-05 |
| ENSG00000265185\|SNORD3B-1 | 29.2826 | 4.8142 | ##### | 4.5475 | 5.4277E-06 | 1.7155E-05 |

| | | | | | | |
|---|---|---|---|---|---|---|
| ENSG00000175854\|SWI5 | 1296.8575 | -0.5522 | ##### | -4.5474 | 5.4320E-06 | 1.7167E-05 |
| ENSG00000187398\|LUZP2 | 43.2812 | 4.6880 | ##### | 4.5470 | 5.4405E-06 | 1.7192E-05 |
| ENSG00000145741\|BTF3 | 26304.0767 | 0.5531 | ##### | 4.5466 | 5.4519E-06 | 1.7226E-05 |
| ENSG00000257261\|SLC38A4-AS1 | 67.1216 | 1.1210 | ##### | 4.5465 | 5.4547E-06 | 1.7233E-05 |
| ENSG00000185015\|CA13 | 296.1417 | -0.7744 | ##### | -4.5453 | 5.4847E-06 | 1.7326E-05 |
| ENSG00000168569\|TMEM223 | 1299.3951 | 0.6060 | ##### | 4.5444 | 5.5098E-06 | 1.7404E-05 |
| ENSG00000176912\|TYMSOS | 230.5266 | 0.8205 | ##### | 4.5442 | 5.5148E-06 | 1.7417E-05 |
| ENSG00000214708 | 17.8896 | -1.9407 | ##### | -4.5435 | 5.5325E-06 | 1.7472E-05 |
| ENSG00000015568\|RGPD5 | 1602.1214 | -0.5791 | ##### | -4.5429 | 5.5495E-06 | 1.7523E-05 |
| ENSG00000257386 | 116.9148 | -0.8807 | ##### | -4.5425 | 5.5600E-06 | 1.7555E-05 |
| ENSG00000293047\|WTAPP1 | 37.0281 | -5.0538 | ##### | -4.5420 | 5.5712E-06 | 1.7588E-05 |
| ENSG00000215421\|ZNF407 | 1391.3027 | 0.5800 | ##### | 4.5418 | 5.5786E-06 | 1.7610E-05 |
| ENSG00000108187\|PBLD | 294.7794 | -0.7956 | ##### | -4.5417 | 5.5794E-06 | 1.7610E-05 |
| ENSG00000278768\|BACE1-AS | 871.7633 | -0.8491 | ##### | -4.5401 | 5.6239E-06 | 1.7749E-05 |
| ENSG00000133872\|SARAF | 10319.9099 | -0.4162 | ##### | -4.5397 | 5.6338E-06 | 1.7778E-05 |
| ENSG00000196470\|SIAH1 | 2157.1482 | -0.6143 | ##### | -4.5395 | 5.6389E-06 | 1.7793E-05 |
| ENSG00000084072\|PPIE | 4000.7208 | -0.5662 | ##### | -4.5393 | 5.6435E-06 | 1.7805E-05 |
| ENSG00000132793\|LPIN3 | 1493.5368 | 0.5938 | ##### | 4.5383 | 5.6713E-06 | 1.7891E-05 |
| ENSG00000108578\|BLMH | 3563.8391 | -0.5656 | ##### | -4.5380 | 5.6795E-06 | 1.7915E-05 |
| ENSG00000151715\|TMEM45B | 20.0115 | -4.8477 | ##### | -4.5373 | 5.6969E-06 | 1.7968E-05 |
| ENSG00000259353 | 29.2775 | 2.8902 | ##### | 4.5373 | 5.6981E-06 | 1.7970E-05 |
| ENSG00000163959\|SLC51A | 109.7552 | -1.1275 | ##### | -4.5365 | 5.7202E-06 | 1.8038E-05 |
| ENSG00000063169\|BICRA | 1940.2824 | -0.5768 | ##### | -4.5357 | 5.7409E-06 | 1.8101E-05 |
| ENSG00000259952 | 1211.1394 | -0.6307 | ##### | -4.5347 | 5.7684E-06 | 1.8186E-05 |
| ENSG00000264247\|ZNF407-AS1 | 562.5568 | -0.6805 | ##### | -4.5343 | 5.7803E-06 | 1.8221E-05 |
| ENSG00000104133\|SPG11 | 2968.2874 | -0.5749 | ##### | -4.5342 | 5.7817E-06 | 1.8224E-05 |
| ENSG00000133424\|LARGE1 | 283.7955 | ####### | ##### | -4.5340 | 5.7863E-06 | 1.8236E-05 |
| ENSG00000171735\|CAMTA1 | 2836.9119 | 0.5857 | ##### | 4.5339 | 5.7902E-06 | 1.8247E-05 |
| ENSG00000253741\|LNCOC1 | 347.8655 | -0.8053 | ##### | -4.5336 | 5.7999E-06 | 1.8275E-05 |
| ENSG00000102886\|GDPD3 | 127.8817 | -0.9900 | ##### | -4.5335 | 5.8014E-06 | 1.8278E-05 |
| ENSG00000215156 | 112.1088 | 6.0918 | ##### | 4.5327 | 5.8228E-06 | 1.8344E-05 |
| ENSG00000235852 | 498.7254 | -0.7396 | ##### | -4.5326 | 5.8270E-06 | 1.8355E-05 |
| ENSG00000283352\|LINC00680-GUSBP4 | 13.7009 | -4.6936 | ##### | -4.5322 | 5.8368E-06 | 1.8384E-05 |
| ENSG00000275708\|MIR3648-1 | 488.1489 | 1.5628 | ##### | 4.5318 | 5.8491E-06 | 1.8421E-05 |
| ENSG00000198521\|ZNF43 | 323.4311 | -0.8373 | ##### | -4.5310 | 5.8693E-06 | 1.8482E-05 |
| ENSG00000240541\|TM4SF1-AS1 | 65.1655 | -1.8089 | ##### | -4.5309 | 5.8730E-06 | 1.8492E-05 |
| ENSG00000127554\|GFER | 2686.4415 | -0.6261 | ##### | -4.5301 | 5.8964E-06 | 1.8564E-05 |
| ENSG00000159176\|CSRP1 | 16229.4786 | -0.5113 | ##### | -4.5300 | 5.8988E-06 | 1.8569E-05 |
| ENSG00000276855 | 51.3272 | -6.9090 | ##### | -4.5285 | 5.9412E-06 | 1.8701E-05 |
| ENSG00000198938\|MT-CO3 | ########## | 0.5642 | ##### | 4.5283 | 5.9456E-06 | 1.8713E-05 |
| ENSG00000064933\|PMS1 | 2754.2475 | -0.5706 | ##### | -4.5282 | 5.9475E-06 | 1.8717E-05 |
| ENSG00000287191 | 83.8024 | -8.8127 | ##### | -4.5282 | 5.9485E-06 | 1.8718E-05 |
| ENSG00000260804\|LINC01963 | 566.6512 | 0.7094 | ##### | 4.5278 | 5.9597E-06 | 1.8751E-05 |
| ENSG00000196498\|NCOR2 | 22537.7613 | 0.5383 | ##### | 4.5277 | 5.9628E-06 | 1.8759E-05 |
| ENSG00000104691\|UBXN8 | 862.5652 | -0.6150 | ##### | -4.5274 | 5.9713E-06 | 1.8784E-05 |
| ENSG00000279047\|PCDHB1-AS1 | 83.5975 | -7.0201 | ##### | -4.5269 | 5.9842E-06 | 1.8822E-05 |
| ENSG00000157514\|TSC22D3 | 631.5974 | -0.6475 | ##### | -4.5268 | 5.9884E-06 | 1.8834E-05 |
| ENSG00000102924\|CBLN1 | 35.4478 | 1.6934 | ##### | 4.5267 | 5.9919E-06 | 1.8842E-05 |
| ENSG00000223573\|TINCR | 505.7494 | -0.6311 | ##### | -4.5262 | 6.0057E-06 | 1.8884E-05 |
| ENSG00000145491\|ROPN1L | 96.5127 | 1.2313 | ##### | 4.5255 | 6.0260E-06 | 1.8946E-05 |
| ENSG00000283930\|PLD5P1 | 26.0713 | -3.9179 | ##### | -4.5252 | 6.0331E-06 | 1.8966E-05 |
| ENSG00000149646\|CNBD2 | 113.5164 | 1.2432 | ##### | 4.5248 | 6.0456E-06 | 1.9004E-05 |
| ENSG00000234028\|EIF2AK3-DT | 125.7017 | -1.0772 | ##### | -4.5247 | 6.0481E-06 | 1.9009E-05 |
| ENSG00000166575\|TMEM135 | 1065.1378 | -0.5662 | ##### | -4.5246 | 6.0515E-06 | 1.9018E-05 |
| ENSG00000069956\|MAPK6 | 6024.2688 | -0.5485 | ##### | -4.5245 | 6.0534E-06 | 1.9022E-05 |
| ENSG00000291298 | 52.0417 | -1.2629 | ##### | -4.5241 | 6.0661E-06 | 1.9060E-05 |
| ENSG00000263711\|LINC02864 | 1081.6757 | -9.5759 | ##### | -4.5236 | 6.0788E-06 | 1.9098E-05 |
| ENSG00000211459\|MT-RNR1 | 47593.4067 | 0.5589 | ##### | 4.5231 | 6.0947E-06 | 1.9146E-05 |
| ENSG00000183735\|TBK1 | 4936.6588 | 0.3966 | ##### | 4.5229 | 6.0992E-06 | 1.9158E-05 |
| ENSG00000293268 | 70.4604 | 1.1711 | ##### | 4.5228 | 6.1032E-06 | 1.9168E-05 |
| ENSG00000092148\|HECTD1 | 11977.9610 | 0.5447 | ##### | 4.5227 | 6.1063E-06 | 1.9174E-05 |
| ENSG00000148942\|SLC5A12 | 154.2124 | -7.9134 | ##### | -4.5227 | 6.1060E-06 | 1.9174E-05 |
| ENSG00000107951\|MTPAP | 3184.8799 | 0.4557 | ##### | 4.5213 | 6.1454E-06 | 1.9295E-05 |

| | | | | | | |
|---|---|---|---|---|---|---|
| ENSG00000038274\|MAT2B | 4747.8298 | 0.5719 | ##### | 4.5209 | 6.1567E-06 | 1.9328E-05 |
| ENSG00000167543\|TP53I13 | 2902.9528 | -0.5578 | ##### | -4.5198 | 6.1894E-06 | 1.9429E-05 |
| ENSG00000268287 | 19.9588 | 5.6601 | ##### | 4.5197 | 6.1919E-06 | 1.9435E-05 |
| ENSG00000165632\|TAF3 | 2156.9323 | 0.5665 | ##### | 4.5197 | 6.1941E-06 | 1.9440E-05 |
| ENSG00000287559 | 207.5024 | 4.6724 | ##### | 4.5187 | 6.2219E-06 | 1.9525E-05 |
| ENSG00000156860\|FBRS | 7958.2759 | 0.3325 | ##### | 4.5177 | 6.2517E-06 | 1.9616E-05 |
| ENSG00000106615\|RHEB | 10784.9970 | 0.5515 | ##### | 4.5168 | 6.2788E-06 | 1.9699E-05 |
| ENSG00000115289\|PCGF1 | 2185.3552 | 0.5729 | ##### | 4.5166 | 6.2828E-06 | 1.9710E-05 |
| ENSG00000227355 | 40.7230 | -2.6835 | ##### | -4.5152 | 6.3251E-06 | 1.9840E-05 |
| ENSG00000235205\|TATDN2P3 | 71.2715 | -8.5791 | ##### | -4.5150 | 6.3330E-06 | 1.9863E-05 |
| ENSG00000241634\|RPL13P8 | 67.6762 | 1.3615 | ##### | 4.5144 | 6.3500E-06 | 1.9914E-05 |
| ENSG00000155621\|C9orf85 | 738.4740 | -0.6278 | ##### | -4.5144 | 6.3510E-06 | 1.9915E-05 |
| ENSG00000105750\|ZNF85 | 55.0888 | -8.0087 | ##### | -4.5134 | 6.3800E-06 | 2.0004E-05 |
| ENSG00000289083\|LINC02256 | 18.6274 | -2.4777 | ##### | -4.5133 | 6.3813E-06 | 2.0006E-05 |
| ENSG00000205155\|PSENEN | 3172.8886 | -0.5706 | ##### | -4.5127 | 6.4019E-06 | 2.0068E-05 |
| ENSG00000124302\|CHST8 | 175.1105 | 5.6181 | ##### | 4.5123 | 6.4126E-06 | 2.0100E-05 |
| ENSG00000110063\|DCPS | 2999.9533 | 0.5405 | ##### | 4.5121 | 6.4203E-06 | 2.0122E-05 |
| ENSG00000180479\|ZNF571 | 159.8544 | -0.9326 | ##### | -4.5120 | 6.4231E-06 | 2.0129E-05 |
| ENSG00000077327\|SPAG6 | 176.2732 | -6.4674 | ##### | -4.5078 | 6.5504E-06 | 2.0525E-05 |
| ENSG00000168256\|NKIRAS2 | 3666.0537 | -0.5594 | ##### | -4.5072 | 6.5704E-06 | 2.0586E-05 |
| ENSG00000257576\|HSPD1P4 | 61.5293 | 1.1621 | ##### | 4.5068 | 6.5826E-06 | 2.0622E-05 |
| ENSG00000244414\|CFHR1 | 141.2857 | -6.1615 | ##### | -4.5056 | 6.6196E-06 | 2.0736E-05 |
| ENSG00000158486\|DNAH3 | 75.1821 | -1.3360 | ##### | -4.5048 | 6.6438E-06 | 2.0809E-05 |
| ENSG00000075624\|ACTB | ########## | -0.5571 | ##### | -4.5047 | 6.6465E-06 | 2.0816E-05 |
| ENSG00000293155 | 24.4175 | -7.0276 | ##### | -4.5047 | 6.6472E-06 | 2.0816E-05 |
| ENSG00000008294\|SPAG9 | 16839.4376 | 0.5365 | ##### | 4.5038 | 6.6751E-06 | 2.0901E-05 |
| ENSG00000286230 | 35.5516 | 1.4105 | ##### | 4.5033 | 6.6918E-06 | 2.0951E-05 |
| ENSG00000291159 | 2667.8264 | 0.5836 | ##### | 4.5031 | 6.6972E-06 | 2.0966E-05 |
| ENSG00000110169\|HPX | 20.3322 | -2.8194 | ##### | -4.5027 | 6.7101E-06 | 2.1004E-05 |
| ENSG00000171302\|CANT1 | 10101.6914 | 0.5627 | ##### | 4.5022 | 6.7248E-06 | 2.1048E-05 |
| ENSG00000198712\|MT-CO2 | ########## | 0.5593 | ##### | 4.5018 | 6.7382E-06 | 2.1087E-05 |
| ENSG00000254910 | 35.0245 | -1.5967 | ##### | -4.5012 | 6.7561E-06 | 2.1141E-05 |
| ENSG00000189091\|SF3B3 | 20070.5624 | -0.5216 | ##### | -4.5003 | 6.7858E-06 | 2.1232E-05 |
| ENSG00000135457\|TFCP2 | 5051.3069 | 0.5383 | ##### | 4.5000 | 6.7966E-06 | 2.1263E-05 |
| ENSG00000131462\|TUBG1 | 16014.0552 | 0.5548 | ##### | 4.4998 | 6.8002E-06 | 2.1273E-05 |
| ENSG00000141485\|SLC13A5 | 29.2899 | 3.5820 | ##### | 4.4997 | 6.8055E-06 | 2.1287E-05 |
| ENSG00000152465\|NMT2 | 2975.3186 | 0.5573 | ##### | 4.4988 | 6.8345E-06 | 2.1375E-05 |
| ENSG00000105723\|GSK3A | 7578.3373 | -0.5424 | ##### | -4.4986 | 6.8399E-06 | 2.1390E-05 |
| ENSG00000291124 | 357.4460 | 0.7255 | ##### | 4.4980 | 6.8604E-06 | 2.1452E-05 |
| ENSG00000205220\|PSMB10 | 1206.5483 | -0.6002 | ##### | -4.4976 | 6.8717E-06 | 2.1485E-05 |
| ENSG00000163995\|ABLIM2 | 219.7441 | -0.8736 | ##### | -4.4966 | 6.9049E-06 | 2.1586E-05 |
| ENSG00000286012 | 70.4246 | 7.4393 | ##### | 4.4965 | 6.9096E-06 | 2.1599E-05 |
| ENSG00000286148 | 101.9023 | 6.0290 | ##### | 4.4957 | 6.9330E-06 | 2.1668E-05 |
| ENSG00000170381\|SEMA3E | 601.1563 | -0.6666 | ##### | -4.4957 | 6.9327E-06 | 2.1668E-05 |
| ENSG00000225976\|CKS1BP2 | 55.2948 | 1.2485 | ##### | 4.4947 | 6.9654E-06 | 2.1766E-05 |
| ENSG00000160408\|ST6GALNAC6 | 1975.1505 | 0.7000 | ##### | 4.4944 | 6.9773E-06 | 2.1801E-05 |
| ENSG00000182670\|TTC3 | 11689.7612 | -0.5424 | ##### | -4.4940 | 6.9886E-06 | 2.1834E-05 |
| ENSG00000136603\|SKIL | 4794.0971 | -0.5331 | ##### | -4.4934 | 7.0110E-06 | 2.1902E-05 |
| ENSG00000258689\|LINC01269 | 67.2838 | -8.4960 | ##### | -4.4932 | 7.0160E-06 | 2.1915E-05 |
| ENSG00000154274\|C4orf19 | 125.3610 | 0.9473 | ##### | 4.4927 | 7.0342E-06 | 2.1970E-05 |
| ENSG00000271254 | 2925.1329 | -0.5471 | ##### | -4.4923 | 7.0458E-06 | 2.2004E-05 |
| ENSG00000120875\|DUSP4 | 7939.4719 | -0.5301 | ##### | -4.4922 | 7.0506E-06 | 2.2017E-05 |
| ENSG00000260261 | 12.8160 | -6.9393 | ##### | -4.4918 | 7.0625E-06 | 2.2051E-05 |
| ENSG00000236829 | 17.4703 | -2.9530 | ##### | -4.4910 | 7.0891E-06 | 2.2132E-05 |
| ENSG00000125148\|MT2A | 15675.3079 | 0.7032 | ##### | 4.4906 | 7.1033E-06 | 2.2174E-05 |
| ENSG00000213221\|DNLZ | 1194.5301 | 0.6312 | ##### | 4.4894 | 7.1422E-06 | 2.2293E-05 |
| ENSG00000268204 | 33.2299 | 6.2062 | ##### | 4.4894 | 7.1439E-06 | 2.2296E-05 |
| ENSG00000135679\|MDM2 | 9102.8343 | 0.5444 | ##### | 4.4888 | 7.1634E-06 | 2.2355E-05 |
| ENSG00000261386 | 9.2350 | 3.4342 | ##### | 4.4885 | 7.1726E-06 | 2.2381E-05 |
| ENSG00000217646\|H2BC16P | 13.4333 | 4.5438 | ##### | 4.4881 | 7.1872E-06 | 2.2424E-05 |
| ENSG00000186812\|ZNF397 | 1467.1037 | -0.5807 | ##### | -4.4876 | 7.2039E-06 | 2.2474E-05 |
| ENSG00000275557 | 48.3399 | -1.3441 | ##### | -4.4867 | 7.2352E-06 | 2.2567E-05 |
| ENSG00000242474\|LINC03015 | 334.0743 | -0.7109 | ##### | -4.4867 | 7.2350E-06 | 2.2567E-05 |

| | | | | | | |
|---|---|---|---|---|---|---|
| ENSG00000279901 | 12.5126 | 2.6278 | ##### | 4.4866 | 7.2364E-06 | 2.2568E-05 |
| ENSG00000177311\|ZBTB38 | 2894.4854 | -0.8139 | ##### | -4.4864 | 7.2436E-06 | 2.2588E-05 |
| ENSG00000124535\|WRNIP1 | 6703.7273 | 0.4951 | ##### | 4.4858 | 7.2636E-06 | 2.2649E-05 |
| ENSG00000139641\|ESYT1 | 17571.1880 | 0.5136 | ##### | 4.4857 | 7.2670E-06 | 2.2657E-05 |
| ENSG00000187535\|IFT140 | 1444.5066 | -0.5895 | ##### | -4.4855 | 7.2727E-06 | 2.2672E-05 |
| ENSG00000127743\|IL17B | 19.3813 | -7.2049 | ##### | -4.4848 | 7.2988E-06 | 2.2751E-05 |
| ENSG00000289511 | 86.9796 | -1.1191 | ##### | -4.4835 | 7.3430E-06 | 2.2887E-05 |
| ENSG00000175505\|CLCF1 | 2414.9752 | -0.5711 | ##### | -4.4834 | 7.3460E-06 | 2.2894E-05 |
| ENSG00000065675\|PRKCQ | 73.9443 | 1.5388 | ##### | 4.4828 | 7.3675E-06 | 2.2958E-05 |
| ENSG00000289075 | 155.5270 | 7.4419 | ##### | 4.4826 | 7.3731E-06 | 2.2973E-05 |
| ENSG00000286901 | 15.3848 | -6.8605 | ##### | -4.4824 | 7.3811E-06 | 2.2996E-05 |
| ENSG00000151929\|BAG3 | 11437.1280 | 0.4303 | ##### | 4.4808 | 7.4349E-06 | 2.3161E-05 |
| ENSG00000111291\|GPRC5D | 46.0757 | 1.5970 | ##### | 4.4802 | 7.4568E-06 | 2.3227E-05 |
| ENSG00000162073\|PAQR4 | 6671.0740 | -0.5579 | ##### | -4.4790 | 7.5006E-06 | 2.3361E-05 |
| ENSG00000165269\|AQP7 | 103.8044 | -9.0984 | ##### | -4.4789 | 7.5044E-06 | 2.3370E-05 |
| ENSG00000112667\|DNPH1 | 2707.6465 | -0.5472 | ##### | -4.4784 | 7.5216E-06 | 2.3421E-05 |
| ENSG00000272735 | 35.7617 | 1.5990 | ##### | 4.4782 | 7.5281E-06 | 2.3437E-05 |
| ENSG00000165288\|BRWD3 | 3303.8488 | 0.5644 | ##### | 4.4782 | 7.5282E-06 | 2.3437E-05 |
| ENSG00000084693\|AGBL5 | 5249.6950 | 0.5500 | ##### | 4.4770 | 7.5716E-06 | 2.3570E-05 |
| ENSG00000271601\|LIX1L | 5435.3266 | 0.4728 | ##### | 4.4769 | 7.5749E-06 | 2.3577E-05 |
| ENSG00000136436\|CALCOCO2 | 4372.4435 | -0.5552 | ##### | -4.4767 | 7.5801E-06 | 2.3591E-05 |
| ENSG00000287180 | 18.6334 | -7.4939 | ##### | -4.4764 | 7.5897E-06 | 2.3619E-05 |
| ENSG00000260267 | 506.6798 | -0.6382 | ##### | -4.4763 | 7.5951E-06 | 2.3633E-05 |
| ENSG00000081386\|ZNF510 | 1072.6543 | -0.6066 | ##### | -4.4762 | 7.5969E-06 | 2.3636E-05 |
| ENSG00000290971 | 16.9341 | -4.4296 | ##### | -4.4761 | 7.6021E-06 | 2.3650E-05 |
| ENSG00000144061\|NPHP1 | 473.9664 | -0.6325 | ##### | -4.4753 | 7.6315E-06 | 2.3739E-05 |
| ENSG00000269910 | 226.1644 | -0.7893 | ##### | -4.4749 | 7.6440E-06 | 2.3775E-05 |
| ENSG00000133835\|HSD17B4 | 5870.2221 | 0.4670 | ##### | 4.4744 | 7.6639E-06 | 2.3835E-05 |
| ENSG00000225595\|LINC01623 | 22.6870 | 7.1341 | ##### | 4.4743 | 7.6659E-06 | 2.3838E-05 |
| ENSG00000119650\|IFT43 | 1317.6334 | -0.5953 | ##### | -4.4739 | 7.6810E-06 | 2.3883E-05 |
| ENSG00000256417 | 43.3109 | -1.7335 | ##### | -4.4737 | 7.6870E-06 | 2.3899E-05 |
| ENSG00000205790\|DPP9-AS1 | 178.6211 | 1.1150 | ##### | 4.4737 | 7.6890E-06 | 2.3903E-05 |
| ENSG00000284693\|LINC02606 | 47.4864 | -3.4878 | ##### | -4.4735 | 7.6941E-06 | 2.3916E-05 |
| ENSG00000281880\|PAUPAR | 26.0910 | 6.6207 | ##### | 4.4724 | 7.7353E-06 | 2.4042E-05 |
| ENSG00000289042 | 60.0414 | -1.6449 | ##### | -4.4716 | 7.7647E-06 | 2.4130E-05 |
| ENSG00000279364 | 734.8368 | 0.6848 | ##### | 4.4713 | 7.7728E-06 | 2.4153E-05 |
| ENSG00000104973\|MED25 | 6136.7435 | -0.5406 | ##### | -4.4704 | 7.8087E-06 | 2.4262E-05 |
| ENSG00000196083\|IL1RAP | 1107.1139 | -0.5870 | ##### | -4.4702 | 7.8135E-06 | 2.4275E-05 |
| ENSG00000258128\|MKRN9P | 24.4063 | -5.8191 | ##### | -4.4699 | 7.8269E-06 | 2.4311E-05 |
| ENSG00000073712\|FERMT2 | 6658.7056 | 0.5299 | ##### | 4.4699 | 7.8268E-06 | 2.4311E-05 |
| ENSG00000106392\|C1GALT1 | 3928.1360 | -0.5573 | ##### | -4.4691 | 7.8531E-06 | 2.4390E-05 |
| ENSG00000269858\|EGLN2 | 5454.0472 | -0.6243 | ##### | -4.4690 | 7.8596E-06 | 2.4408E-05 |
| ENSG00000265681\|RPL17 | 38727.1731 | -0.5178 | ##### | -4.4688 | 7.8665E-06 | 2.4426E-05 |
| ENSG00000255642\|PABPC1P4 | 46.0081 | -1.4704 | ##### | -4.4684 | 7.8807E-06 | 2.4468E-05 |
| ENSG00000184058\|TBX1 | 305.7558 | 0.7531 | ##### | 4.4683 | 7.8833E-06 | 2.4473E-05 |
| ENSG00000054523\|KIF1B | 6446.3958 | -0.5492 | ##### | -4.4675 | 7.9138E-06 | 2.4566E-05 |
| ENSG00000196505\|GDAP2 | 2332.8635 | 0.5523 | ##### | 4.4673 | 7.9218E-06 | 2.4588E-05 |
| ENSG00000159363\|ATP13A2 | 7808.3754 | 0.4737 | ##### | 4.4672 | 7.9242E-06 | 2.4593E-05 |
| ENSG00000071967\|CYBRD1 | 1012.2108 | 0.5702 | ##### | 4.4665 | 7.9512E-06 | 2.4674E-05 |
| ENSG00000213648\|SULT1A4 | 158.4707 | -1.9953 | ##### | -4.4664 | 7.9549E-06 | 2.4683E-05 |
| ENSG00000183054\|RGPD6 | 1660.1825 | -0.5801 | ##### | -4.4662 | 7.9611E-06 | 2.4700E-05 |
| ENSG00000196284\|SUPT3H | 598.7009 | 0.6228 | ##### | 4.4654 | 7.9920E-06 | 2.4793E-05 |
| ENSG00000268199 | 40.6871 | 1.2497 | ##### | 4.4651 | 8.0018E-06 | 2.4821E-05 |
| ENSG00000255524\|NPIPB8 | 70.0627 | -1.2101 | ##### | -4.4651 | 8.0046E-06 | 2.4827E-05 |
| ENSG00000130856\|ZNF236 | 1303.9098 | 0.5736 | ##### | 4.4646 | 8.0230E-06 | 2.4881E-05 |
| ENSG00000124657\|OR2B6 | 20.8079 | 3.1293 | ##### | 4.4643 | 8.0325E-06 | 2.4908E-05 |
| ENSG00000188001\|TPRG1 | 76.8370 | -2.3813 | ##### | -4.4642 | 8.0380E-06 | 2.4923E-05 |
| ENSG00000261879\|ZNF594-DT | 103.5414 | 1.0134 | ##### | 4.4631 | 8.0791E-06 | 2.5048E-05 |
| ENSG00000070495\|JMJD6 | 3523.9268 | 0.5360 | ##### | 4.4618 | 8.1280E-06 | 2.5197E-05 |
| ENSG00000255529\|POLR2M | 2903.7453 | -0.5679 | ##### | -4.4613 | 8.1480E-06 | 2.5256E-05 |
| ENSG00000254851 | 44.8830 | 1.4929 | ##### | 4.4612 | 8.1509E-06 | 2.5262E-05 |
| ENSG00000230836\|LINC01293 | 21.7684 | 7.0552 | ##### | 4.4608 | 8.1663E-06 | 2.5307E-05 |
| ENSG00000123349\|PFDN5 | 8948.4974 | 0.5937 | ##### | 4.4602 | 8.1891E-06 | 2.5375E-05 |

| | | | | | | |
|---|---|---|---|---|---|---|
| ENSG00000293412\|GUCY1B2 | 69.9835 | 2.5297 | ##### | 4.4599 | 8.1983E-06 | 2.5401E-05 |
| ENSG00000161798\|AQP5 | 15.7713 | -7.1344 | ##### | -4.4591 | 8.2319E-06 | 2.5503E-05 |
| ENSG00000272931\|LRRC8D-DT | 30.9730 | -1.8483 | ##### | -4.4587 | 8.2467E-06 | 2.5546E-05 |
| ENSG00000272264 | 31.6429 | -1.5037 | ##### | -4.4585 | 8.2529E-06 | 2.5562E-05 |
| ENSG00000117385\|P3H1 | 6259.2443 | -0.5240 | ##### | -4.4584 | 8.2591E-06 | 2.5579E-05 |
| ENSG00000285991 | 2595.5024 | -4.6103 | ##### | -4.4575 | 8.2924E-06 | 2.5679E-05 |
| ENSG00000124126\|PREX1 | 1886.5048 | -0.5610 | ##### | -4.4568 | 8.3174E-06 | 2.5754E-05 |
| ENSG00000166780\|BMERB1 | 3542.9543 | 0.5288 | ##### | 4.4562 | 8.3408E-06 | 2.5824E-05 |
| ENSG00000237550 | 4651.4197 | -0.6753 | ##### | -4.4562 | 8.3422E-06 | 2.5826E-05 |
| ENSG00000105355\|PLIN3 | 11081.7569 | -0.5241 | ##### | -4.4560 | 8.3513E-06 | 2.5851E-05 |
| ENSG00000289614 | 235.1319 | -0.7582 | ##### | -4.4547 | 8.4008E-06 | 2.6002E-05 |
| ENSG00000114784\|EIF1B | 1992.3947 | 0.5591 | ##### | 4.4544 | 8.4121E-06 | 2.6034E-05 |
| ENSG00000259374\|NDUFB4P11 | 25.3425 | -7.0040 | ##### | -4.4541 | 8.4230E-06 | 2.6065E-05 |
| ENSG00000150967\|ABCB9 | 2726.8158 | 0.5739 | ##### | 4.4538 | 8.4361E-06 | 2.6103E-05 |
| ENSG00000105983\|LMBR1 | 6325.6127 | 0.5420 | ##### | 4.4533 | 8.4554E-06 | 2.6160E-05 |
| ENSG00000186716\|BCR | 3057.0716 | -0.5396 | ##### | -4.4514 | 8.5324E-06 | 2.6395E-05 |
| ENSG00000011451\|WIZ | 5153.0823 | 0.5166 | ##### | 4.4513 | 8.5372E-06 | 2.6407E-05 |
| ENSG00000119632\|IFI27L2 | 432.6895 | -0.6856 | ##### | -4.4510 | 8.5469E-06 | 2.6435E-05 |
| ENSG00000120306\|CYSTM1 | 2639.6730 | -0.5784 | ##### | -4.4506 | 8.5650E-06 | 2.6488E-05 |
| ENSG00000111530\|CAND1 | 21029.2269 | 0.4954 | ##### | 4.4500 | 8.5857E-06 | 2.6549E-05 |
| ENSG00000286732 | 19.2048 | -4.8277 | ##### | -4.4493 | 8.6166E-06 | 2.6642E-05 |
| ENSG00000166736\|HTR3A | 16.3395 | -7.4417 | ##### | -4.4488 | 8.6338E-06 | 2.6692E-05 |
| ENSG00000115806\|GORASP2 | 9215.7518 | 0.5467 | ##### | 4.4486 | 8.6440E-06 | 2.6720E-05 |
| ENSG00000231776\|LINC01611 | 33.8591 | -2.5524 | ##### | -4.4486 | 8.6447E-06 | 2.6720E-05 |
| ENSG00000054219\|LY75 | 37.9314 | -3.0590 | ##### | -4.4476 | 8.6834E-06 | 2.6837E-05 |
| ENSG00000183018\|SPNS2 | 284.2971 | -0.7677 | ##### | -4.4471 | 8.7039E-06 | 2.6898E-05 |
| ENSG00000222033\|LINC01124 | 108.8548 | -1.3120 | ##### | -4.4469 | 8.7113E-06 | 2.6918E-05 |
| ENSG00000129472\|RAB2B | 1705.4968 | 0.5752 | ##### | 4.4464 | 8.7310E-06 | 2.6976E-05 |
| ENSG00000081870\|IFT25 | 1618.5565 | 0.5753 | ##### | 4.4458 | 8.7547E-06 | 2.7046E-05 |
| ENSG00000196119\|OR8A1 | 68.4021 | -8.5192 | ##### | -4.4458 | 8.7564E-06 | 2.7049E-05 |
| ENSG00000117505\|DR1 | 7868.0709 | 0.5339 | ##### | 4.4457 | 8.7598E-06 | 2.7057E-05 |
| ENSG00000288984 | 20.9804 | 2.3504 | ##### | 4.4452 | 8.7791E-06 | 2.7114E-05 |
| ENSG00000253190 | 19.0986 | 4.0203 | ##### | 4.4448 | 8.7980E-06 | 2.7169E-05 |
| ENSG00000127184\|COX7C | 12142.3175 | 0.5407 | ##### | 4.4445 | 8.8081E-06 | 2.7197E-05 |
| ENSG00000161860\|SYCE2 | 86.1878 | 1.0359 | ##### | 4.4436 | 8.8457E-06 | 2.7311E-05 |
| ENSG00000226688\|ENTPD1-AS1 | 2508.0717 | -1.5213 | ##### | -4.4433 | 8.8592E-06 | 2.7349E-05 |
| ENSG00000249776 | 48.5522 | 7.7828 | ##### | 4.4432 | 8.8613E-06 | 2.7353E-05 |
| ENSG00000224905 | 29.4660 | 3.9732 | ##### | 4.4425 | 8.8936E-06 | 2.7450E-05 |
| ENSG00000229267\|SNHG31 | 43.9787 | 2.2027 | ##### | 4.4417 | 8.9263E-06 | 2.7548E-05 |
| ENSG00000161642\|ZNF385A | 1960.4170 | -0.5803 | ##### | -4.4415 | 8.9329E-06 | 2.7566E-05 |
| ENSG00000140832\|MARVELD3 | 55.7195 | -1.7818 | ##### | -4.4412 | 8.9448E-06 | 2.7599E-05 |
| ENSG00000023516\|AKAP11 | 4251.1945 | 0.5317 | ##### | 4.4406 | 8.9725E-06 | 2.7679E-05 |
| ENSG00000147206\|NXF3 | 106.9751 | 8.3675 | ##### | 4.4406 | 8.9722E-06 | 2.7679E-05 |
| ENSG00000132702\|HAPLN2 | 47.6936 | 1.2643 | ##### | 4.4404 | 8.9772E-06 | 2.7691E-05 |
| ENSG00000204516\|MICB | 2429.3857 | 0.5665 | ##### | 4.4396 | 9.0130E-06 | 2.7799E-05 |
| ENSG00000289407 | 58.2648 | -1.2872 | ##### | -4.4393 | 9.0245E-06 | 2.7831E-05 |
| ENSG00000267629 | 50.2628 | 2.0921 | ##### | 4.4386 | 9.0529E-06 | 2.7916E-05 |
| ENSG00000131233\|GJA9 | 27.2427 | -2.3684 | ##### | -4.4384 | 9.0625E-06 | 2.7942E-05 |
| ENSG00000142784\|WDTC1 | 2982.7980 | 0.5663 | ##### | 4.4382 | 9.0724E-06 | 2.7970E-05 |
| ENSG00000124593 | 4020.3215 | -0.9293 | ##### | -4.4378 | 9.0873E-06 | 2.8013E-05 |
| ENSG00000196670\|ZFP62 | 876.5666 | 0.6256 | ##### | 4.4377 | 9.0924E-06 | 2.8026E-05 |
| ENSG00000089154\|GCN1 | 29185.1391 | 0.5232 | ##### | 4.4374 | 9.1071E-06 | 2.8069E-05 |
| ENSG00000059377\|TBXAS1 | 99.3905 | -2.8582 | ##### | -4.4372 | 9.1145E-06 | 2.8088E-05 |
| ENSG00000255351 | 329.7867 | 0.7522 | ##### | 4.4363 | 9.1528E-06 | 2.8203E-05 |
| ENSG00000196338\|NLGN3 | 137.9797 | 1.0647 | ##### | 4.4361 | 9.1616E-06 | 2.8228E-05 |
| ENSG00000214353\|VAC14-AS1 | 24.2816 | 3.4924 | ##### | 4.4356 | 9.1834E-06 | 2.8292E-05 |
| ENSG00000283093\|CENPVL2 | 62.4036 | 7.3156 | ##### | 4.4352 | 9.1968E-06 | 2.8330E-05 |
| ENSG00000223559 | 98.5882 | 0.8873 | ##### | 4.4351 | 9.2044E-06 | 2.8351E-05 |
| ENSG00000167280\|ENGASE | 2224.3294 | 0.5609 | ##### | 4.4345 | 9.2267E-06 | 2.8417E-05 |
| ENSG00000129158\|SERGEF | 664.7230 | -0.6664 | ##### | -4.4344 | 9.2312E-06 | 2.8427E-05 |
| ENSG00000084073\|ZMPSTE24 | 10130.5773 | -0.5487 | ##### | -4.4341 | 9.2446E-06 | 2.8466E-05 |
| ENSG00000269194 | 40.8498 | 1.3095 | ##### | 4.4335 | 9.2712E-06 | 2.8545E-05 |
| ENSG00000101367\|MAPRE1 | 19056.9157 | 0.4135 | ##### | 4.4334 | 9.2739E-06 | 2.8550E-05 |

| | | | | | | |
|---|---|---|---|---|---|---|
| ENSG00000206878 | 7.7493 | 5.8517 | ##### | 4.4330 | 9.2927E-06 | 2.8605E-05 |
| ENSG00000214530\|STARD10 | 1104.9910 | 0.6170 | ##### | 4.4328 | 9.3012E-06 | 2.8628E-05 |
| ENSG00000246273\|SBF2-AS1 | 715.4325 | 0.6392 | ##### | 4.4318 | 9.3465E-06 | 2.8765E-05 |
| ENSG00000126945\|HNRNPH2 | 7800.7245 | 0.5328 | ##### | 4.4317 | 9.3510E-06 | 2.8776E-05 |
| ENSG00000197479\|PCDHB11 | 27.9327 | -3.4100 | ##### | -4.4307 | 9.3941E-06 | 2.8905E-05 |
| ENSG00000162009\|SSTR5 | 50.7591 | -8.4001 | ##### | -4.4305 | 9.4015E-06 | 2.8925E-05 |
| ENSG00000102271\|KLHL4 | 856.7109 | 0.5956 | ##### | 4.4298 | 9.4323E-06 | 2.9017E-05 |
| ENSG00000187097\|ENTPD5 | 1975.3894 | -0.5863 | ##### | -4.4298 | 9.4334E-06 | 2.9017E-05 |
| ENSG00000135966\|TGFBRAP1 | 4287.5574 | -0.5126 | ##### | -4.4289 | 9.4710E-06 | 2.9128E-05 |
| ENSG00000256463\|SALL3 | 69.0582 | -8.5330 | ##### | -4.4289 | 9.4713E-06 | 2.9128E-05 |
| ENSG00000248489\|CHD1-DT | 90.7418 | 1.0258 | ##### | 4.4285 | 9.4871E-06 | 2.9173E-05 |
| ENSG00000158019\|BABAM2 | 2793.0859 | -0.5570 | ##### | -4.4282 | 9.5008E-06 | 2.9213E-05 |
| ENSG00000108379\|WNT3 | 728.4685 | -0.5991 | ##### | -4.4281 | 9.5070E-06 | 2.9229E-05 |
| ENSG00000183283\|DAZAP2 | 16914.6426 | 0.5414 | ##### | 4.4279 | 9.5176E-06 | 2.9258E-05 |
| ENSG00000197579\|TOPORS | 1540.6844 | -0.5649 | ##### | -4.4278 | 9.5217E-06 | 2.9268E-05 |
| ENSG00000236384\|LINC00479 | 23.3929 | -6.9102 | ##### | -4.4272 | 9.5446E-06 | 2.9335E-05 |
| ENSG00000281912\|LINC01144 | 42.2536 | -1.2991 | ##### | -4.4265 | 9.5765E-06 | 2.9430E-05 |
| ENSG00000132825\|PPP1R3D | 649.4361 | -0.6399 | ##### | -4.4263 | 9.5862E-06 | 2.9457E-05 |
| ENSG00000072121\|ZFYVE26 | 3064.3672 | 0.5712 | ##### | 4.4261 | 9.5941E-06 | 2.9475E-05 |
| ENSG00000185115\|NSMCE3 | 1761.5752 | 0.5836 | ##### | 4.4261 | 9.5939E-06 | 2.9475E-05 |
| ENSG00000261971\|MMP25-AS1 | 392.4182 | -0.7631 | ##### | -4.4261 | 9.5970E-06 | 2.9481E-05 |
| ENSG00000196968\|FUT11 | 1471.4537 | -0.4157 | ##### | -4.4258 | 9.6106E-06 | 2.9520E-05 |
| ENSG00000100889\|PCK2 | 1866.4688 | -0.5540 | ##### | -4.4257 | 9.6135E-06 | 2.9526E-05 |
| ENSG00000279806 | 64.2346 | -8.4287 | ##### | -4.4255 | 9.6200E-06 | 2.9543E-05 |
| ENSG00000143811\|PYCR2 | 5742.9929 | -0.5314 | ##### | -4.4253 | 9.6287E-06 | 2.9566E-05 |
| ENSG00000135686\|KLHL36 | 4109.8260 | -0.5544 | ##### | -4.4248 | 9.6539E-06 | 2.9641E-05 |
| ENSG00000169682\|SPNS1 | 5293.6365 | -0.5319 | ##### | -4.4246 | 9.6623E-06 | 2.9663E-05 |
| ENSG00000204618\|RNF39 | 164.4796 | -0.8405 | ##### | -4.4239 | 9.6917E-06 | 2.9750E-05 |
| ENSG00000122873\|CISD1 | 2224.6202 | -0.5971 | ##### | -4.4235 | 9.7103E-06 | 2.9805E-05 |
| ENSG00000083896\|YTHDC1 | 6103.1267 | 0.5368 | ##### | 4.4231 | 9.7273E-06 | 2.9854E-05 |
| ENSG00000272476 | 141.2261 | -0.9075 | ##### | -4.4231 | 9.7289E-06 | 2.9855E-05 |
| ENSG00000241553\|ARPC4 | 10242.7949 | 0.5358 | ##### | 4.4225 | 9.7559E-06 | 2.9932E-05 |
| ENSG00000147041\|SYTL5 | 13.0573 | 3.9786 | ##### | 4.4225 | 9.7555E-06 | 2.9932E-05 |
| ENSG00000133315\|MACROD1 | 622.7893 | 0.6355 | ##### | 4.4221 | 9.7757E-06 | 2.9990E-05 |
| ENSG00000144711\|IQSEC1 | 2562.9821 | 0.5566 | ##### | 4.4220 | 9.7806E-06 | 3.0002E-05 |
| ENSG00000196126\|HLA-DRB1 | 37.2932 | -3.4622 | ##### | -4.4218 | 9.7903E-06 | 3.0025E-05 |
| ENSG00000270012 | 66.8863 | 1.1549 | ##### | 4.4218 | 9.7894E-06 | 3.0025E-05 |
| ENSG00000187860\|CCDC157 | 250.3359 | 0.8752 | ##### | 4.4211 | 9.8212E-06 | 3.0117E-05 |
| ENSG00000151413\|NUBPL | 1005.8557 | 0.5758 | ##### | 4.4204 | 9.8532E-06 | 3.0212E-05 |
| ENSG00000249115\|HAUS5 | 1901.1589 | -0.5410 | ##### | -4.4203 | 9.8563E-06 | 3.0218E-05 |
| ENSG00000137171\|KLC4 | 797.6505 | -0.6163 | ##### | -4.4201 | 9.8636E-06 | 3.0238E-05 |
| ENSG00000273151 | 325.1778 | 0.7779 | ##### | 4.4198 | 9.8788E-06 | 3.0278E-05 |
| ENSG00000254198 | 73.1280 | 1.6424 | ##### | 4.4198 | 9.8779E-06 | 3.0278E-05 |
| ENSG00000184635\|ZNF93 | 65.5490 | -8.4578 | ##### | -4.4197 | 9.8853E-06 | 3.0295E-05 |
| ENSG00000134940\|ACRV1 | 42.6783 | 1.8303 | ##### | 4.4195 | 9.8913E-06 | 3.0310E-05 |
| ENSG00000276087 | 16.1175 | -6.9378 | ##### | -4.4187 | 9.9282E-06 | 3.0420E-05 |
| ENSG00000168096\|ANKS3 | 1528.2245 | -0.5529 | ##### | -4.4183 | 9.9477E-06 | 3.0477E-05 |
| ENSG00000129988\|LBP | 75.4996 | -8.6617 | ##### | -4.4181 | 9.9559E-06 | 3.0499E-05 |
| ENSG00000284738 | 45.2598 | 1.3343 | ##### | 4.4178 | 9.9690E-06 | 3.0536E-05 |
| ENSG00000254858\|MPV17L2 | 1574.5548 | 0.5707 | ##### | 4.4177 | 9.9752E-06 | 3.0552E-05 |
| ENSG00000174851\|YIF1A | 5123.3168 | -0.5532 | ##### | -4.4176 | 9.9802E-06 | 3.0564E-05 |
| ENSG00000197860\|SGTB | 1632.5318 | -0.6122 | ##### | -4.4175 | 9.9831E-06 | 3.0570E-05 |
| ENSG00000083720\|OXCT1 | 2053.9566 | 0.5554 | ##### | 4.4173 | 9.9925E-06 | 3.0595E-05 |
| ENSG00000234719\|NPIPB2 | 44.4457 | -1.3841 | ##### | -4.4173 | 9.9936E-06 | 3.0596E-05 |
| ENSG00000233232\|NPIPB7 | 29.5936 | -1.6569 | ##### | -4.4169 | 1.0015E-05 | 3.0657E-05 |
| ENSG00000284773 | 128.6230 | -2.9584 | ##### | -4.4166 | 1.0026E-05 | 3.0688E-05 |
| ENSG00000137486\|ARRB1 | 4829.7757 | -0.5335 | ##### | -4.4165 | 1.0031E-05 | 3.0702E-05 |
| ENSG00000198865\|CCDC152 | 38.4735 | -2.3446 | ##### | -4.4161 | 1.0049E-05 | 3.0753E-05 |
| ENSG00000105447\|GRWD1 | 9407.4154 | -0.4154 | ##### | -4.4145 | 1.0122E-05 | 3.0974E-05 |
| ENSG00000273623 | 36.7506 | -1.6401 | ##### | -4.4140 | 1.0148E-05 | 3.1048E-05 |
| ENSG00000272694 | 18.1064 | -2.0193 | ##### | -4.4138 | 1.0156E-05 | 3.1071E-05 |
| ENSG00000287477 | 49.6734 | 1.7293 | ##### | 4.4137 | 1.0160E-05 | 3.1079E-05 |
| ENSG00000070366\|SMG6 | 2871.2168 | 0.5383 | ##### | 4.4137 | 1.0162E-05 | 3.1083E-05 |

| | | | | | | |
|---|---|---|---|---|---|---|
| ENSG00000060237\|WNK1 | 26701.9502 | 0.5300 | ##### | 4.4133 | 1.0182E-05 | 3.1140E-05 |
| ENSG00000110090\|CPT1A | 6228.7392 | -0.5304 | ##### | -4.4125 | 1.0216E-05 | 3.1242E-05 |
| ENSG00000132768\|DPH2 | 8513.6666 | 0.4964 | ##### | 4.4115 | 1.0263E-05 | 3.1383E-05 |
| ENSG00000169976\|SF3B5 | 4935.2317 | -0.5418 | ##### | -4.4114 | 1.0269E-05 | 3.1396E-05 |
| ENSG00000226065\|ZBTB45P2 | 750.1208 | -0.6128 | ##### | -4.4112 | 1.0280E-05 | 3.1428E-05 |
| ENSG00000114279\|FGF12 | 676.1572 | 0.6533 | ##### | 4.4109 | 1.0296E-05 | 3.1474E-05 |
| ENSG00000259006 | 485.5411 | 0.7889 | ##### | 4.4108 | 1.0301E-05 | 3.1486E-05 |
| ENSG00000289291 | 69.5163 | 1.2247 | ##### | 4.4105 | 1.0311E-05 | 3.1514E-05 |
| ENSG00000182141\|ZNF708 | 171.5325 | -0.8521 | ##### | -4.4104 | 1.0319E-05 | 3.1532E-05 |
| ENSG00000087884\|AAMDC | 384.5328 | 0.6826 | ##### | 4.4078 | 1.0440E-05 | 3.1901E-05 |
| ENSG00000169059\|VCX3A | 58.8210 | -8.3016 | ##### | -4.4071 | 1.0475E-05 | 3.2003E-05 |
| ENSG00000123405\|NFE2 | 34.7232 | -4.5555 | ##### | -4.4067 | 1.0494E-05 | 3.2060E-05 |
| ENSG00000162706\|CADM3 | 94.1522 | -8.9524 | ##### | -4.4067 | 1.0497E-05 | 3.2065E-05 |
| ENSG00000175894\|TSPEAR | 90.1817 | 5.8298 | ##### | 4.4060 | 1.0530E-05 | 3.2163E-05 |
| ENSG00000284543\|LINC01226 | 18.5628 | -7.5664 | ##### | -4.4056 | 1.0547E-05 | 3.2210E-05 |
| ENSG00000004948\|CALCR | 67.8648 | -4.0965 | ##### | -4.4055 | 1.0553E-05 | 3.2226E-05 |
| ENSG00000114388\|NPRL2 | 1135.2227 | 0.6062 | ##### | 4.4054 | 1.0557E-05 | 3.2235E-05 |
| ENSG00000281398\|SNHG4 | 1181.1508 | 0.6114 | ##### | 4.4047 | 1.0592E-05 | 3.2339E-05 |
| ENSG00000084774\|CAD | 17905.0159 | 0.5177 | ##### | 4.4041 | 1.0624E-05 | 3.2434E-05 |
| ENSG00000283667 | 39.5433 | 1.4222 | ##### | 4.4036 | 1.0647E-05 | 3.2501E-05 |
| ENSG00000142875\|PRKACB | 3780.9425 | -0.5518 | ##### | -4.4021 | 1.0722E-05 | 3.2725E-05 |
| ENSG00000236104\|ZBTB22 | 745.7706 | -0.6742 | ##### | -4.4018 | 1.0734E-05 | 3.2759E-05 |
| ENSG00000226055\|PAICSP1 | 31.8072 | 3.0467 | ##### | 4.4017 | 1.0741E-05 | 3.2775E-05 |
| ENSG00000196821\|ILRUN | 8911.4438 | -0.4785 | ##### | -4.4012 | 1.0764E-05 | 3.2842E-05 |
| ENSG00000169189\|NSMCE1 | 1847.8904 | -0.5804 | ##### | -4.3999 | 1.0829E-05 | 3.3038E-05 |
| ENSG00000143252\|SDHC | 10324.9578 | 0.5309 | ##### | 4.3996 | 1.0846E-05 | 3.3087E-05 |
| ENSG00000288880 | 51.7846 | 1.1931 | ##### | 4.3995 | 1.0852E-05 | 3.3098E-05 |
| ENSG00000106290\|TAF6 | 13560.3752 | 0.5519 | ##### | 4.3995 | 1.0851E-05 | 3.3098E-05 |
| ENSG00000232368\|FTLP2 | 10.3401 | -3.1694 | ##### | -4.3984 | 1.0907E-05 | 3.3261E-05 |
| ENSG00000230612 | 40.9824 | 7.4635 | ##### | 4.3973 | 1.0960E-05 | 3.3421E-05 |
| ENSG00000156239\|N6AMT1 | 804.8436 | -0.6632 | ##### | -4.3962 | 1.1016E-05 | 3.3587E-05 |
| ENSG00000012124\|CD22 | 124.2469 | -9.3340 | ##### | -4.3959 | 1.1033E-05 | 3.3634E-05 |
| ENSG00000283602 | 21.8492 | 6.5014 | ##### | 4.3959 | 1.1034E-05 | 3.3634E-05 |
| ENSG00000170166\|HOXD4 | 31.7959 | 1.7586 | ##### | 4.3956 | 1.1049E-05 | 3.3677E-05 |
| ENSG00000120509\|PDZD11 | 3371.1037 | 0.5384 | ##### | 4.3954 | 1.1058E-05 | 3.3701E-05 |
| ENSG00000104866\|PPP1R37 | 2325.6710 | -0.4652 | ##### | -4.3945 | 1.1100E-05 | 3.3828E-05 |
| ENSG00000101974\|ATP11C | 2536.0645 | 0.5701 | ##### | 4.3932 | 1.1169E-05 | 3.4033E-05 |
| ENSG00000106009\|BRAT1 | 6950.8235 | 0.3780 | ##### | 4.3929 | 1.1183E-05 | 3.4072E-05 |
| ENSG00000268635 | 21.8356 | 1.8813 | ##### | 4.3926 | 1.1201E-05 | 3.4125E-05 |
| ENSG00000112679\|DUSP22 | 1529.3591 | 0.5813 | ##### | 4.3926 | 1.1203E-05 | 3.4126E-05 |
| ENSG00000080709\|KCNN2 | 18.2396 | 6.0835 | ##### | 4.3919 | 1.1236E-05 | 3.4223E-05 |
| ENSG00000286585 | 29.1009 | -1.9679 | ##### | -4.3917 | 1.1248E-05 | 3.4255E-05 |
| ENSG00000229018\|PMS2P7 | 224.7847 | -0.7621 | ##### | -4.3916 | 1.1254E-05 | 3.4270E-05 |
| ENSG00000204130\|RUFY2 | 1129.6202 | -0.5710 | ##### | -4.3915 | 1.1258E-05 | 3.4279E-05 |
| ENSG00000059573\|ALDH18A1 | 8712.4078 | -0.4951 | ##### | -4.3913 | 1.1268E-05 | 3.4306E-05 |
| ENSG00000101997\|CCDC22 | 1320.6021 | -0.5840 | ##### | -4.3902 | 1.1323E-05 | 3.4470E-05 |
| ENSG00000108312\|UBTF | 18083.5553 | 0.5044 | ##### | 4.3898 | 1.1347E-05 | 3.4540E-05 |
| ENSG00000137414\|FAM8A1 | 1464.4456 | 0.5785 | ##### | 4.3884 | 1.1419E-05 | 3.4756E-05 |
| ENSG00000122034\|GTF3A | 7857.1811 | 0.5161 | ##### | 4.3875 | 1.1464E-05 | 3.4889E-05 |
| ENSG00000177842\|ZNF620 | 237.4232 | 0.8933 | ##### | 4.3873 | 1.1475E-05 | 3.4920E-05 |
| ENSG00000290784 | 22.0585 | -2.2501 | ##### | -4.3873 | 1.1478E-05 | 3.4925E-05 |
| ENSG00000198064\|NPIPB13 | 428.6762 | 0.7275 | ##### | 4.3868 | 1.1501E-05 | 3.4991E-05 |
| ENSG00000166582\|CENPV | 3407.8717 | 0.3898 | ##### | 4.3864 | 1.1525E-05 | 3.5061E-05 |
| ENSG00000008056\|SYN1 | 35.5859 | 1.7765 | ##### | 4.3849 | 1.1602E-05 | 3.5290E-05 |
| ENSG00000285410\|GABPB1-IT1 | 549.0391 | -0.6681 | ##### | -4.3848 | 1.1609E-05 | 3.5309E-05 |
| ENSG00000171401\|KRT13 | 97.1687 | -9.5385 | ##### | -4.3840 | 1.1652E-05 | 3.5438E-05 |
| ENSG00000288658 | 86.3460 | 1.1695 | ##### | 4.3832 | 1.1697E-05 | 3.5571E-05 |
| ENSG00000242021 | 112.6155 | -9.2104 | ##### | -4.3828 | 1.1717E-05 | 3.5627E-05 |
| ENSG00000259738\|ZNF444P1 | 18.7447 | 5.1447 | ##### | 4.3813 | 1.1796E-05 | 3.5864E-05 |
| ENSG00000165323\|FAT3 | 83.6306 | -4.8421 | ##### | -4.3811 | 1.1806E-05 | 3.5890E-05 |
| ENSG00000160299\|PCNT | 4607.2983 | 0.5413 | ##### | 4.3807 | 1.1830E-05 | 3.5959E-05 |
| ENSG00000107902\|LHPP | 896.2766 | -0.6011 | ##### | -4.3806 | 1.1837E-05 | 3.5976E-05 |
| ENSG00000123700\|KCNJ2 | 162.7710 | -1.0626 | ##### | -4.3805 | 1.1843E-05 | 3.5993E-05 |

| | | | | | | |
|---|---|---|---|---|---|---|
| ENSG00000145817\|YIPF5 | 5257.2722 | -0.3980 | ##### | -4.3801 | 1.1863E-05 | 3.6048E-05 |
| ENSG00000293199\|PTENP1 | 83.1049 | -1.0227 | ##### | -4.3792 | 1.1910E-05 | 3.6189E-05 |
| ENSG00000138688\|BLTP1 | 2750.6815 | 0.5480 | ##### | 4.3790 | 1.1922E-05 | 3.6221E-05 |
| ENSG00000256977\|LIMS3 | 86.0274 | -2.2146 | ##### | -4.3789 | 1.1926E-05 | 3.6230E-05 |
| ENSG00000117971\|CHRNB4 | 32.0049 | 1.5798 | ##### | 4.3788 | 1.1933E-05 | 3.6248E-05 |
| ENSG00000130287\|NCAN | 57.1250 | 3.8901 | ##### | 4.3781 | 1.1975E-05 | 3.6369E-05 |
| ENSG00000286104 | 400.0630 | -0.8202 | ##### | -4.3774 | 1.2010E-05 | 3.6472E-05 |
| ENSG00000147996\|ZNG1E | 1436.6157 | -0.5717 | ##### | -4.3756 | 1.2110E-05 | 3.6772E-05 |
| ENSG00000259755 | 14.3879 | -4.0510 | ##### | -4.3755 | 1.2115E-05 | 3.6784E-05 |
| ENSG00000048140\|TSPAN17 | 4216.9716 | 0.5293 | ##### | 4.3750 | 1.2145E-05 | 3.6871E-05 |
| ENSG00000251179\|TMEM92-AS1 | 60.9072 | -8.3515 | ##### | -4.3741 | 1.2194E-05 | 3.7018E-05 |
| ENSG00000168878\|SFTPB | 33.1063 | -6.5289 | ##### | -4.3740 | 1.2197E-05 | 3.7023E-05 |
| ENSG00000184047\|DIABLO | 4719.5760 | -0.6267 | ##### | -4.3730 | 1.2255E-05 | 3.7194E-05 |
| ENSG00000102057\|KCND1 | 255.5089 | -0.7560 | ##### | -4.3724 | 1.2289E-05 | 3.7294E-05 |
| ENSG00000119969\|HELLS | 4178.9694 | 0.5168 | ##### | 4.3719 | 1.2315E-05 | 3.7369E-05 |
| ENSG00000245025 | 309.6161 | 0.7319 | ##### | 4.3714 | 1.2344E-05 | 3.7454E-05 |
| ENSG00000176715\|ACSF3 | 1834.4021 | -0.5746 | ##### | -4.3708 | 1.2381E-05 | 3.7562E-05 |
| ENSG00000151304\|SRFBP1 | 1667.2164 | 0.5905 | ##### | 4.3705 | 1.2396E-05 | 3.7604E-05 |
| ENSG00000171223\|JUNB | 14324.2088 | 0.5063 | ##### | 4.3696 | 1.2446E-05 | 3.7751E-05 |
| ENSG00000138107\|ACTR1A | 10489.7466 | -0.4322 | ##### | -4.3694 | 1.2460E-05 | 3.7789E-05 |
| ENSG00000270095 | 27.0287 | 6.5499 | ##### | 4.3694 | 1.2461E-05 | 3.7789E-05 |
| ENSG00000153294\|ADGRF4 | 103.6799 | -1.1081 | ##### | -4.3692 | 1.2471E-05 | 3.7816E-05 |
| ENSG00000138867\|GUCD1 | 3814.3319 | -0.4264 | ##### | -4.3690 | 1.2483E-05 | 3.7847E-05 |
| ENSG00000258377 | 3024.5149 | 0.5575 | ##### | 4.3682 | 1.2529E-05 | 3.7985E-05 |
| ENSG00000281706\|LINC01012 | 98.8141 | 0.9881 | ##### | 4.3674 | 1.2576E-05 | 3.8122E-05 |
| ENSG00000054179\|ENTPD2 | 32.4062 | -1.6402 | ##### | -4.3668 | 1.2606E-05 | 3.8210E-05 |
| ENSG00000124641\|MED20 | 2040.7210 | -0.5585 | ##### | -4.3659 | 1.2658E-05 | 3.8365E-05 |
| ENSG00000204138\|PHACTR4 | 4690.7700 | 0.4689 | ##### | 4.3652 | 1.2701E-05 | 3.8489E-05 |
| ENSG00000224543\|SNRPGP15 | 111.1121 | 0.9869 | ##### | 4.3649 | 1.2720E-05 | 3.8544E-05 |
| ENSG00000213145\|CRIP1 | 5892.3815 | 0.5411 | ##### | 4.3646 | 1.2736E-05 | 3.8589E-05 |
| ENSG00000243710\|CFAP57 | 167.4543 | 0.8638 | ##### | 4.3639 | 1.2779E-05 | 3.8713E-05 |
| ENSG00000230805 | 47.7990 | 7.7730 | ##### | 4.3636 | 1.2796E-05 | 3.8762E-05 |
| ENSG00000187534\|PRR13P5 | 290.0799 | 0.6983 | ##### | 4.3635 | 1.2803E-05 | 3.8778E-05 |
| ENSG00000247735\|KCTD13-DT | 262.0293 | -0.7927 | ##### | -4.3627 | 1.2846E-05 | 3.8897E-05 |
| ENSG00000087152\|ATXN7L3 | 7239.2263 | 0.5188 | ##### | 4.3627 | 1.2845E-05 | 3.8897E-05 |
| ENSG00000183741\|CBX6 | 8181.5934 | -0.5238 | ##### | -4.3627 | 1.2845E-05 | 3.8897E-05 |
| ENSG00000153165\|RGPD3 | 94.3634 | -0.9679 | ##### | -4.3625 | 1.2859E-05 | 3.8932E-05 |
| ENSG00000147677\|EIF3H | 13455.3703 | -0.4998 | ##### | -4.3624 | 1.2867E-05 | 3.8952E-05 |
| ENSG00000104883\|PEX11G | 57.7321 | -1.4983 | ##### | -4.3623 | 1.2871E-05 | 3.8961E-05 |
| ENSG00000290832 | 1204.7437 | 0.5937 | ##### | 4.3619 | 1.2891E-05 | 3.9018E-05 |
| ENSG00000187554\|TLR5 | 65.8052 | -3.3837 | ##### | -4.3617 | 1.2906E-05 | 3.9060E-05 |
| ENSG00000289516 | 55.3495 | -8.2135 | ##### | -4.3613 | 1.2929E-05 | 3.9124E-05 |
| ENSG00000174945\|AMZ1 | 39.4753 | -1.9023 | ##### | -4.3609 | 1.2953E-05 | 3.9195E-05 |
| ENSG00000285280 | 57.5945 | -7.5601 | ##### | -4.3602 | 1.2997E-05 | 3.9323E-05 |
| ENSG00000183298\|RPSAP19 | 361.8678 | 0.7147 | ##### | 4.3600 | 1.3005E-05 | 3.9342E-05 |
| ENSG00000020129\|NCDN | 5075.7480 | 0.5326 | ##### | 4.3597 | 1.3023E-05 | 3.9394E-05 |
| ENSG00000125482\|TTF1 | 1950.2397 | 0.5473 | ##### | 4.3594 | 1.3042E-05 | 3.9447E-05 |
| ENSG00000164086\|DUSP7 | 3316.7834 | 0.5603 | ##### | 4.3590 | 1.3067E-05 | 3.9519E-05 |
| ENSG00000146701\|MDH2 | 19172.0415 | -0.5243 | ##### | -4.3577 | 1.3146E-05 | 3.9755E-05 |
| ENSG00000247809\|NR2F2-AS1 | 77.7670 | -1.2107 | ##### | -4.3569 | 1.3190E-05 | 3.9881E-05 |
| ENSG00000120693\|SMAD9 | 278.0751 | -0.7961 | ##### | -4.3569 | 1.3194E-05 | 3.9885E-05 |
| ENSG00000214510\|SPINK13 | 75.8157 | -4.7541 | ##### | -4.3569 | 1.3193E-05 | 3.9885E-05 |
| ENSG00000173064\|HECTD4 | 2979.9739 | 0.5666 | ##### | 4.3567 | 1.3202E-05 | 3.9907E-05 |
| ENSG00000006451\|RALA | 4169.9677 | -0.5326 | ##### | -4.3564 | 1.3220E-05 | 3.9956E-05 |
| ENSG00000163064\|EN1 | 68.1936 | 1.5744 | ##### | 4.3561 | 1.3237E-05 | 4.0005E-05 |
| ENSG00000268144\|NIFKP6 | 12.6027 | -7.0603 | ##### | -4.3557 | 1.3262E-05 | 4.0075E-05 |
| ENSG00000075151\|EIF4G3 | 9099.1358 | 0.5221 | ##### | 4.3555 | 1.3279E-05 | 4.0118E-05 |
| ENSG00000248905\|FMN1 | 415.1866 | 0.7479 | ##### | 4.3555 | 1.3279E-05 | 4.0118E-05 |
| ENSG00000119283\|TRIM67 | 112.0885 | -5.0718 | ##### | -4.3554 | 1.3280E-05 | 4.0118E-05 |
| ENSG00000229401\|MIR5689HG | 15.3713 | 6.3040 | ##### | 4.3551 | 1.3300E-05 | 4.0174E-05 |
| ENSG00000253626\|EIF5AL1 | 1202.3580 | 0.5667 | ##### | 4.3545 | 1.3338E-05 | 4.0284E-05 |
| ENSG00000122390\|NAA60 | 4320.5611 | -0.5407 | ##### | -4.3528 | 1.3441E-05 | 4.0592E-05 |
| ENSG00000166823\|MESP1 | 293.3826 | -0.6993 | ##### | -4.3522 | 1.3475E-05 | 4.0691E-05 |

| | | | | | | |
|---|---|---|---|---|---|---|
| ENSG00000159086\|PAXBP1 | 3972.7438 | 0.5411 | ##### | 4.3519 | 1.3498E-05 | 4.0756E-05 |
| ENSG00000111605\|CPSF6 | 8180.8497 | 0.5336 | ##### | 4.3502 | 1.3598E-05 | 4.1055E-05 |
| ENSG00000250132 | 267.4150 | -0.7508 | ##### | -4.3502 | 1.3603E-05 | 4.1066E-05 |
| ENSG00000293175 | 27.7914 | -2.0182 | ##### | -4.3501 | 1.3608E-05 | 4.1075E-05 |
| ENSG00000246548\|LINC02288 | 30.7862 | 3.4153 | ##### | 4.3501 | 1.3609E-05 | 4.1076E-05 |
| ENSG00000223865\|HLA-DPB1 | 52.6422 | -8.1410 | ##### | -4.3499 | 1.3621E-05 | 4.1106E-05 |
| ENSG00000227963\|RBM15-AS1 | 40.4540 | 1.7932 | ##### | 4.3496 | 1.3638E-05 | 4.1153E-05 |
| ENSG00000104888\|SLC17A7 | 141.2673 | 0.9919 | ##### | 4.3489 | 1.3680E-05 | 4.1278E-05 |
| ENSG00000272657 | 11.1322 | 5.6641 | ##### | 4.3481 | 1.3735E-05 | 4.1437E-05 |
| ENSG00000270580\|PKD1P6-NPIPP1 | 821.0922 | 0.6473 | ##### | 4.3478 | 1.3752E-05 | 4.1485E-05 |
| ENSG00000271401 | 16.5032 | 5.6038 | ##### | 4.3477 | 1.3760E-05 | 4.1505E-05 |
| ENSG00000242457\|RBBP4P2 | 23.1017 | -1.8472 | ##### | -4.3466 | 1.3825E-05 | 4.1696E-05 |
| ENSG00000204620 | 47.2239 | -1.3217 | ##### | -4.3464 | 1.3841E-05 | 4.1740E-05 |
| ENSG00000168348\|INSM2 | 19.3070 | 3.9860 | ##### | 4.3463 | 1.3844E-05 | 4.1746E-05 |
| ENSG00000108395\|TRIM37 | 6119.8075 | 0.5354 | ##### | 4.3459 | 1.3873E-05 | 4.1828E-05 |
| ENSG00000169429\|CXCL8 | 1402.4683 | -0.6667 | ##### | -4.3454 | 1.3903E-05 | 4.1914E-05 |
| ENSG00000214381\|LINC00488 | 51.2057 | -2.2532 | ##### | -4.3449 | 1.3932E-05 | 4.1998E-05 |
| ENSG00000177464\|GPR4 | 88.4012 | -0.9783 | ##### | -4.3433 | 1.4037E-05 | 4.2309E-05 |
| ENSG00000114857\|NKTR | 3437.3980 | 0.5223 | ##### | 4.3431 | 1.4050E-05 | 4.2346E-05 |
| ENSG00000132554\|RGS22 | 32.9172 | -7.8104 | ##### | -4.3426 | 1.4078E-05 | 4.2425E-05 |
| ENSG00000202538\|RNU4-2 | 14.9742 | 5.6739 | ##### | 4.3422 | 1.4105E-05 | 4.2503E-05 |
| ENSG00000163736\|PPBP | 35.2398 | -7.9560 | ##### | -4.3421 | 1.4116E-05 | 4.2531E-05 |
| ENSG00000105855\|ITGB8 | 1084.6543 | 0.5811 | ##### | 4.3416 | 1.4145E-05 | 4.2616E-05 |
| ENSG00000112697\|TMEM30A | 24019.2300 | 0.4447 | ##### | 4.3412 | 1.4170E-05 | 4.2687E-05 |
| ENSG00000258537\|FRMD6-AS2 | 125.0262 | -1.0750 | ##### | -4.3411 | 1.4180E-05 | 4.2710E-05 |
| ENSG00000120885\|CLU | 36250.7250 | -0.5094 | ##### | -4.3409 | 1.4189E-05 | 4.2736E-05 |
| ENSG00000215481 | 24.0693 | -3.3759 | ##### | -4.3408 | 1.4198E-05 | 4.2758E-05 |
| ENSG00000140854\|KATNB1 | 3582.0998 | 0.5473 | ##### | 4.3406 | 1.4209E-05 | 4.2785E-05 |
| ENSG00000147571\|CRH | 220.2399 | 5.5113 | ##### | 4.3395 | 1.4282E-05 | 4.3003E-05 |
| ENSG00000125686\|MED1 | 11729.2332 | 0.5123 | ##### | 4.3389 | 1.4319E-05 | 4.3110E-05 |
| ENSG00000164066\|INTU | 695.5997 | -0.6232 | ##### | -4.3385 | 1.4344E-05 | 4.3180E-05 |
| ENSG00000218283\|MORF4L1P1 | 2118.7032 | -0.5429 | ##### | -4.3378 | 1.4389E-05 | 4.3311E-05 |
| ENSG00000175879\|HOXD8 | 359.2016 | -0.7067 | ##### | -4.3377 | 1.4395E-05 | 4.3325E-05 |
| ENSG00000218418 | 138.0937 | 0.9003 | ##### | 4.3372 | 1.4432E-05 | 4.3433E-05 |
| ENSG00000142655\|PEX14 | 2349.3044 | -0.5497 | ##### | -4.3371 | 1.4434E-05 | 4.3434E-05 |
| ENSG00000271858 | 17.5778 | -4.1444 | ##### | -4.3370 | 1.4446E-05 | 4.3463E-05 |
| ENSG00000159374\|M1AP | 37.3049 | -7.4713 | ##### | -4.3359 | 1.4514E-05 | 4.3664E-05 |
| ENSG00000249279\|LINC02057 | 69.8654 | -6.8375 | ##### | -4.3358 | 1.4526E-05 | 4.3696E-05 |
| ENSG00000267855\|NDUFA7 | 1761.5339 | -0.6127 | ##### | -4.3338 | 1.4653E-05 | 4.4073E-05 |
| ENSG00000202415\|RN7SKP269 | 12.7969 | -2.0813 | ##### | -4.3336 | 1.4670E-05 | 4.4120E-05 |
| ENSG00000104979\|C19orf53 | 5542.2616 | 0.5433 | ##### | 4.3333 | 1.4692E-05 | 4.4181E-05 |
| ENSG00000148400\|NOTCH1 | 4062.7257 | -0.5352 | ##### | -4.3332 | 1.4693E-05 | 4.4181E-05 |
| ENSG00000137273\|FOXF2 | 943.1416 | -0.5970 | ##### | -4.3331 | 1.4700E-05 | 4.4196E-05 |
| ENSG00000232306 | 48.3528 | -8.0184 | ##### | -4.3329 | 1.4715E-05 | 4.4238E-05 |
| ENSG00000276744 | 20.1937 | 4.0314 | ##### | 4.3328 | 1.4726E-05 | 4.4266E-05 |
| ENSG00000289446 | 16.0012 | 7.2650 | ##### | 4.3317 | 1.4799E-05 | 4.4481E-05 |
| ENSG00000164111\|ANXA5 | 42632.9717 | 0.5168 | ##### | 4.3312 | 1.4828E-05 | 4.4564E-05 |
| ENSG00000259153\|MAP3K9-DT | 485.6596 | -0.6552 | ##### | -4.3312 | 1.4831E-05 | 4.4569E-05 |
| ENSG00000089220\|PEBP1 | 10234.6769 | 0.4561 | ##### | 4.3312 | 1.4833E-05 | 4.4570E-05 |
| ENSG00000179776\|CDH5 | 100.2817 | 8.0720 | ##### | 4.3311 | 1.4834E-05 | 4.4570E-05 |
| ENSG00000150991\|UBC | ########## | 0.2944 | ##### | 4.3310 | 1.4841E-05 | 4.4585E-05 |
| ENSG00000269399 | 78.3615 | 1.2014 | ##### | 4.3307 | 1.4866E-05 | 4.4656E-05 |
| ENSG00000188342\|GTF2F2 | 3028.3498 | 0.5526 | ##### | 4.3296 | 1.4937E-05 | 4.4857E-05 |
| ENSG00000178502\|KLHL11 | 1600.7794 | 0.5927 | ##### | 4.3296 | 1.4937E-05 | 4.4857E-05 |
| ENSG00000273192 | 435.7686 | -0.7713 | ##### | -4.3296 | 1.4935E-05 | 4.4857E-05 |
| ENSG00000163597\|SNHG16 | 6662.1849 | 0.6198 | ##### | 4.3296 | 1.4939E-05 | 4.4857E-05 |
| ENSG00000221818\|EBF2 | 34.8501 | 2.2635 | ##### | 4.3287 | 1.5002E-05 | 4.5042E-05 |
| ENSG00000167759\|KLK13 | 14.5082 | -6.7866 | ##### | -4.3277 | 1.5069E-05 | 4.5240E-05 |
| ENSG00000286545 | 62.7191 | -1.3194 | ##### | -4.3276 | 1.5075E-05 | 4.5253E-05 |
| ENSG00000105329\|TGFB1 | 6878.9506 | 0.5453 | ##### | 4.3269 | 1.5120E-05 | 4.5384E-05 |
| ENSG00000276545\|PCDHGB9P | 16.3100 | -4.5842 | ##### | -4.3256 | 1.5211E-05 | 4.5651E-05 |
| ENSG00000250471\|GMPSP1 | 105.8711 | 1.1028 | ##### | 4.3255 | 1.5216E-05 | 4.5662E-05 |
| ENSG00000285728 | 142.2085 | 1.0489 | ##### | 4.3253 | 1.5235E-05 | 4.5716E-05 |

| | | | | | | |
|---|---|---|---|---|---|---|
| ENSG00000018280\|SLC11A1 | 91.8511 | -1.7308 | ##### | -4.3250 | 1.5251E-05 | 4.5759E-05 |
| ENSG00000183889 | 1859.0652 | 0.5731 | ##### | 4.3247 | 1.5277E-05 | 4.5830E-05 |
| ENSG00000274211\|SOCS7 | 4699.8213 | 0.5283 | ##### | 4.3245 | 1.5290E-05 | 4.5865E-05 |
| ENSG00000278635 | 44.6335 | 1.5215 | ##### | 4.3244 | 1.5293E-05 | 4.5870E-05 |
| ENSG00000291273 | 71.3819 | 1.0274 | ##### | 4.3238 | 1.5339E-05 | 4.6002E-05 |
| ENSG00000246465 | 90.6538 | 1.1006 | ##### | 4.3237 | 1.5342E-05 | 4.6002E-05 |
| ENSG00000196236\|XPNPEP3 | 1932.1179 | -0.5447 | ##### | -4.3237 | 1.5341E-05 | 4.6002E-05 |
| ENSG00000258548\|LINC00645 | 51.6138 | -8.1125 | ##### | -4.3210 | 1.5533E-05 | 4.6572E-05 |
| ENSG00000284800 | 81.5516 | -7.8736 | ##### | -4.3208 | 1.5545E-05 | 4.6602E-05 |
| ENSG00000135436\|FAM186B | 109.4581 | 1.0520 | ##### | 4.3201 | 1.5594E-05 | 4.6744E-05 |
| ENSG00000119965\|C10orf88 | 1182.6856 | -0.5776 | ##### | -4.3199 | 1.5607E-05 | 4.6780E-05 |
| ENSG00000103184\|SEC14L5 | 117.1957 | 4.1706 | ##### | 4.3195 | 1.5640E-05 | 4.6874E-05 |
| ENSG00000165660\|ABRAXAS2 | 2336.9477 | -0.5508 | ##### | -4.3188 | 1.5685E-05 | 4.7002E-05 |
| ENSG00000213232\|PPP1R2P10 | 165.0507 | -9.7900 | ##### | -4.3184 | 1.5717E-05 | 4.7095E-05 |
| ENSG00000289056 | 89.8700 | -1.1753 | ##### | -4.3164 | 1.5858E-05 | 4.7512E-05 |
| ENSG00000287156 | 14.7860 | 6.9204 | ##### | 4.3153 | 1.5941E-05 | 4.7756E-05 |
| ENSG00000125454\|SLC25A19 | 1601.1670 | 0.4804 | ##### | 4.3152 | 1.5945E-05 | 4.7764E-05 |
| ENSG00000112874\|NUDT12 | 1291.7049 | -0.5844 | ##### | -4.3149 | 1.5966E-05 | 4.7821E-05 |
| ENSG00000090061\|CCNK | 4550.4753 | 0.6632 | ##### | 4.3145 | 1.5995E-05 | 4.7902E-05 |
| ENSG00000243150\|MLF1-DT | 45.5416 | 1.5958 | ##### | 4.3141 | 1.6022E-05 | 4.7979E-05 |
| ENSG00000174238\|PITPNA | 7100.0962 | 0.3866 | ##### | 4.3134 | 1.6076E-05 | 4.8136E-05 |
| ENSG00000134982\|APC | 2647.8313 | 0.5521 | ##### | 4.3120 | 1.6177E-05 | 4.8432E-05 |
| ENSG00000177951\|BET1L | 4326.4566 | -0.5419 | ##### | -4.3119 | 1.6186E-05 | 4.8456E-05 |
| ENSG00000266947 | 74.6190 | -1.1630 | ##### | -4.3117 | 1.6199E-05 | 4.8490E-05 |
| ENSG00000248187 | 49.1402 | -8.0415 | ##### | -4.3116 | 1.6207E-05 | 4.8510E-05 |
| ENSG00000143799\|PARP1 | 24900.0392 | 0.4540 | ##### | 4.3099 | 1.6332E-05 | 4.8872E-05 |
| ENSG00000291203 | 777.3328 | 0.6103 | ##### | 4.3099 | 1.6330E-05 | 4.8872E-05 |
| ENSG00000205476\|CCDC85C | 9730.9568 | -0.5149 | ##### | -4.3098 | 1.6344E-05 | 4.8903E-05 |
| ENSG00000105619\|TFPT | 1187.8379 | -0.5834 | ##### | -4.3089 | 1.6407E-05 | 4.9089E-05 |
| ENSG00000222448\|RN7SKP150 | 10.4105 | -6.1279 | ##### | -4.3071 | 1.6544E-05 | 4.9492E-05 |
| ENSG00000154889\|MPPE1 | 3197.6440 | -0.5213 | ##### | -4.3067 | 1.6568E-05 | 4.9560E-05 |
| ENSG00000229618 | 58.0943 | -6.0299 | ##### | -4.3067 | 1.6574E-05 | 4.9574E-05 |
| ENSG00000100228\|RAB36 | 781.1438 | 0.5813 | ##### | 4.3066 | 1.6579E-05 | 4.9582E-05 |
| ENSG00000113068\|PFDN1 | 4626.9517 | 0.5428 | ##### | 4.3060 | 1.6620E-05 | 4.9701E-05 |
| ENSG00000251819\|RNU6-322P | 14.2417 | 6.2822 | ##### | 4.3057 | 1.6648E-05 | 4.9778E-05 |
| ENSG00000229590\|MSX2P1 | 19.4781 | 3.5921 | ##### | 4.3056 | 1.6650E-05 | 4.9779E-05 |
| ENSG00000144406\|UNC80 | 29.1723 | 2.0322 | ##### | 4.3042 | 1.6756E-05 | 5.0093E-05 |
| ENSG00000213730\|POLD2P1 | 13.8595 | -4.2801 | ##### | -4.3039 | 1.6782E-05 | 5.0164E-05 |
| ENSG00000280739\|EIF1B-AS1 | 39.4364 | -2.0187 | ##### | -4.3038 | 1.6792E-05 | 5.0190E-05 |
| ENSG00000198185\|ZNF334 | 94.2324 | 5.2385 | ##### | 4.3035 | 1.6809E-05 | 5.0236E-05 |
| ENSG00000105467\|SYNGR4 | 32.0515 | -3.7253 | ##### | -4.3031 | 1.6839E-05 | 5.0320E-05 |
| ENSG00000094975\|SUCO | 3914.5099 | -0.5101 | ##### | -4.3028 | 1.6862E-05 | 5.0385E-05 |
| ENSG00000267128\|RNF157-AS1 | 356.6122 | -3.0669 | ##### | -4.3025 | 1.6884E-05 | 5.0445E-05 |
| ENSG00000243725\|TTC4 | 3760.9824 | -0.5428 | ##### | -4.3009 | 1.7007E-05 | 5.0808E-05 |
| ENSG00000245571\|FAM111A-DT | 87.0908 | -1.1115 | ##### | -4.3005 | 1.7043E-05 | 5.0910E-05 |
| ENSG00000234817\|ECI2-DT | 22.9057 | -2.3926 | ##### | -4.3004 | 1.7048E-05 | 5.0920E-05 |
| ENSG00000151532\|VTI1A | 2564.0355 | -0.5485 | ##### | -4.3004 | 1.7052E-05 | 5.0926E-05 |
| ENSG00000232973\|CYP1B1-AS1 | 358.1044 | 1.4353 | ##### | 4.2997 | 1.7103E-05 | 5.1074E-05 |
| ENSG00000161647\|MPP3 | 608.6504 | -0.6364 | ##### | -4.2990 | 1.7155E-05 | 5.1223E-05 |
| ENSG00000103489\|XYLT1 | 111.3453 | -7.5519 | ##### | -4.2985 | 1.7196E-05 | 5.1340E-05 |
| ENSG00000196812\|ZSCAN16 | 25.3223 | -3.7742 | ##### | -4.2981 | 1.7226E-05 | 5.1425E-05 |
| ENSG00000126858\|RHOT1 | 3668.8466 | -0.5396 | ##### | -4.2972 | 1.7301E-05 | 5.1643E-05 |
| ENSG00000280184 | 18.7688 | 3.1242 | ##### | 4.2968 | 1.7328E-05 | 5.1718E-05 |
| ENSG00000198356\|GET3 | 7808.4455 | 0.5329 | ##### | 4.2965 | 1.7351E-05 | 5.1783E-05 |
| ENSG00000172432\|GTPBP2 | 3086.6143 | 0.5249 | ##### | 4.2964 | 1.7361E-05 | 5.1806E-05 |
| ENSG00000204949\|FAM83A-AS1 | 114.8523 | -7.6295 | ##### | -4.2961 | 1.7383E-05 | 5.1869E-05 |
| ENSG00000290867\|LINC01145 | 585.0055 | 0.6537 | ##### | 4.2945 | 1.7510E-05 | 5.2242E-05 |
| ENSG00000224051\|CPTP | 2748.5657 | -0.5312 | ##### | -4.2944 | 1.7517E-05 | 5.2257E-05 |
| ENSG00000125386\|FAM193A | 2253.8562 | 0.4734 | ##### | 4.2936 | 1.7581E-05 | 5.2444E-05 |
| ENSG00000187848\|P2RX2 | 64.2301 | 8.3365 | ##### | 4.2931 | 1.7616E-05 | 5.2541E-05 |
| ENSG00000243766\|HOTTIP | 31.1328 | -2.8500 | ##### | -4.2922 | 1.7691E-05 | 5.2760E-05 |
| ENSG00000266980 | 14.3655 | 2.3895 | ##### | 4.2904 | 1.7832E-05 | 5.3176E-05 |
| ENSG00000113240\|CLK4 | 806.9739 | 0.6321 | ##### | 4.2901 | 1.7856E-05 | 5.3242E-05 |

| | | | | | | |
|---|---|---|---|---|---|---|
| ENSG00000111786\|SRSF9 | 19512.2826 | 0.5220 | ##### | 4.2900 | 1.7870E-05 | 5.3277E-05 |
| ENSG00000105176\|URI1 | 8129.1449 | -0.3778 | ##### | -4.2892 | 1.7929E-05 | 5.3448E-05 |
| ENSG00000290643 | 14.1960 | -2.4660 | ##### | -4.2889 | 1.7954E-05 | 5.3518E-05 |
| ENSG00000257270 | 148.0778 | 0.8990 | ##### | 4.2887 | 1.7975E-05 | 5.3576E-05 |
| ENSG00000165416\|SUGT1 | 4803.1372 | 0.5282 | ##### | 4.2881 | 1.8019E-05 | 5.3696E-05 |
| ENSG00000242259\|C22orf39 | 1028.4625 | 0.5879 | ##### | 4.2881 | 1.8018E-05 | 5.3696E-05 |
| ENSG00000167183\|PRR15L | 30.2680 | -4.8656 | ##### | -4.2869 | 1.8116E-05 | 5.3981E-05 |
| ENSG00000018189\|RUFY3 | 1563.7213 | -0.5942 | ##### | -4.2864 | 1.8156E-05 | 5.4092E-05 |
| ENSG00000270066 | 58.0134 | 2.5422 | ##### | 4.2855 | 1.8229E-05 | 5.4302E-05 |
| ENSG00000232842\|KAT7P1 | 9.9915 | 5.7721 | ##### | 4.2855 | 1.8230E-05 | 5.4302E-05 |
| ENSG00000057593\|F7 | 22.5834 | -5.6467 | ##### | -4.2837 | 1.8383E-05 | 5.4752E-05 |
| ENSG00000144891\|AGTR1 | 25.9344 | 2.2061 | ##### | 4.2829 | 1.8449E-05 | 5.4945E-05 |
| ENSG00000116815\|CD58 | 1393.2018 | 0.5559 | ##### | 4.2828 | 1.8458E-05 | 5.4966E-05 |
| ENSG00000291250 | 45.5753 | 4.6730 | ##### | 4.2822 | 1.8510E-05 | 5.5113E-05 |
| ENSG00000109390\|NDUFC1 | 1761.2375 | -0.5626 | ##### | -4.2821 | 1.8513E-05 | 5.5118E-05 |
| ENSG00000108924\|HLF | 71.6980 | -1.1986 | ##### | -4.2818 | 1.8535E-05 | 5.5178E-05 |
| ENSG00000112902\|SEMA5A | 162.0122 | 4.8369 | ##### | 4.2811 | 1.8599E-05 | 5.5363E-05 |
| ENSG00000255112\|CHMP1B | 16224.6760 | 0.3781 | ##### | 4.2804 | 1.8653E-05 | 5.5518E-05 |
| ENSG00000170881\|RNF139 | 2653.4540 | -0.5273 | ##### | -4.2798 | 1.8705E-05 | 5.5668E-05 |
| ENSG00000185989\|RASA3 | 2098.8673 | -0.5459 | ##### | -4.2795 | 1.8733E-05 | 5.5746E-05 |
| ENSG00000184986\|TMEM121 | 413.7761 | -0.6446 | ##### | -4.2784 | 1.8821E-05 | 5.5996E-05 |
| ENSG00000115840\|SLC25A12 | 1999.7696 | 0.5491 | ##### | 4.2784 | 1.8820E-05 | 5.5996E-05 |
| ENSG00000286054 | 13.5955 | 4.9127 | ##### | 4.2773 | 1.8917E-05 | 5.6275E-05 |
| ENSG00000166225\|FRS2 | 3407.6113 | 0.5365 | ##### | 4.2772 | 1.8926E-05 | 5.6297E-05 |
| ENSG00000170044\|ZPLD1 | 84.0925 | 9.3169 | ##### | 4.2769 | 1.8954E-05 | 5.6374E-05 |
| ENSG00000074696\|HACD3 | 9799.8395 | -0.5094 | ##### | -4.2766 | 1.8975E-05 | 5.6433E-05 |
| ENSG00000263301 | 21.0366 | 2.1127 | ##### | 4.2765 | 1.8983E-05 | 5.6450E-05 |
| ENSG00000083444\|PLOD1 | 11009.8179 | -0.4182 | ##### | -4.2753 | 1.9092E-05 | 5.6768E-05 |
| ENSG00000145868\|FBXO38 | 2121.1938 | 0.5442 | ##### | 4.2750 | 1.9117E-05 | 5.6836E-05 |
| ENSG00000124299\|PEPD | 8136.6720 | 0.5114 | ##### | 4.2736 | 1.9236E-05 | 5.7186E-05 |
| ENSG00000153714\|LURAP1L | 325.8186 | -0.6987 | ##### | -4.2733 | 1.9260E-05 | 5.7251E-05 |
| ENSG00000265808\|SEC22B | 8983.5997 | -0.3508 | ##### | -4.2730 | 1.9290E-05 | 5.7333E-05 |
| ENSG00000121957\|GPSM2 | 5771.7339 | 0.6764 | ##### | 4.2724 | 1.9339E-05 | 5.7472E-05 |
| ENSG00000124380\|SNRNP27 | 2517.7757 | 0.5461 | ##### | 4.2724 | 1.9340E-05 | 5.7472E-05 |
| ENSG00000128487\|SPECC1 | 3189.9940 | -0.5117 | ##### | -4.2721 | 1.9360E-05 | 5.7525E-05 |
| ENSG00000267107\|PCAT19 | 70.6413 | 5.0791 | ##### | 4.2718 | 1.9391E-05 | 5.7611E-05 |
| ENSG00000287306 | 37.5147 | -1.4186 | ##### | -4.2705 | 1.9501E-05 | 5.7932E-05 |
| ENSG00000250988\|SNHG21 | 177.6725 | -0.7744 | ##### | -4.2699 | 1.9558E-05 | 5.8096E-05 |
| ENSG00000215910\|C1orf167 | 40.3120 | -1.5575 | ##### | -4.2697 | 1.9570E-05 | 5.8127E-05 |
| ENSG00000290415 | 103.0139 | 0.9006 | ##### | 4.2696 | 1.9585E-05 | 5.8165E-05 |
| ENSG00000168488\|ATXN2L | 20776.1034 | 0.3431 | ##### | 4.2689 | 1.9640E-05 | 5.8322E-05 |
| ENSG00000163121\|NEURL3 | 35.4572 | -1.7342 | ##### | -4.2683 | 1.9696E-05 | 5.8484E-05 |
| ENSG00000188786\|MTF1 | 2781.1727 | 0.5200 | ##### | 4.2679 | 1.9732E-05 | 5.8584E-05 |
| ENSG00000168610\|STAT3 | 33332.5327 | -0.4957 | ##### | -4.2676 | 1.9759E-05 | 5.8659E-05 |
| ENSG00000182446\|NPLOC4 | 19307.7014 | -0.4596 | ##### | -4.2674 | 1.9775E-05 | 5.8694E-05 |
| ENSG00000145248\|SLC10A4 | 327.2648 | -3.4759 | ##### | -4.2674 | 1.9774E-05 | 5.8694E-05 |
| ENSG00000182985\|CADM1 | 2907.2580 | -0.5324 | ##### | -4.2671 | 1.9807E-05 | 5.8783E-05 |
| ENSG00000101290\|CDS2 | 2946.1618 | -0.5254 | ##### | -4.2666 | 1.9845E-05 | 5.8890E-05 |
| ENSG00000153487\|ING1 | 1133.9272 | 0.4880 | ##### | 4.2661 | 1.9892E-05 | 5.9024E-05 |
| ENSG00000127922\|SEM1 | 7331.8251 | 0.5300 | ##### | 4.2652 | 1.9972E-05 | 5.9256E-05 |
| ENSG00000227848\|SUCLA2-AS1 | 15.7885 | -4.4124 | ##### | -4.2650 | 1.9987E-05 | 5.9293E-05 |
| ENSG00000249568 | 11.6424 | -2.4475 | ##### | -4.2646 | 2.0026E-05 | 5.9403E-05 |
| ENSG00000110852\|CLEC2B | 86.6780 | 3.5258 | ##### | 4.2643 | 2.0052E-05 | 5.9474E-05 |
| ENSG00000275924\|MIR6807 | 23.2461 | 1.5662 | ##### | 4.2641 | 2.0070E-05 | 5.9524E-05 |
| ENSG00000058262\|SEC61A1 | 36565.3334 | -0.4071 | ##### | -4.2640 | 2.0077E-05 | 5.9536E-05 |
| ENSG00000254995\|STX16-NPEPL1 | 57.4365 | 1.8757 | ##### | 4.2634 | 2.0135E-05 | 5.9702E-05 |
| ENSG00000198018\|ENTPD7 | 2782.4545 | -0.5526 | ##### | -4.2632 | 2.0150E-05 | 5.9741E-05 |
| ENSG00000108424\|KPNB1 | 52068.4156 | 0.4954 | ##### | 4.2627 | 2.0197E-05 | 5.9876E-05 |
| ENSG00000161956\|SENP3 | 9182.0091 | 0.4548 | ##### | 4.2624 | 2.0222E-05 | 5.9943E-05 |
| ENSG00000234350\|ERICH2-DT | 24.0367 | -1.9605 | ##### | -4.2620 | 2.0257E-05 | 6.0041E-05 |
| ENSG00000122756\|CNTFR | 16.7697 | -3.3191 | ##### | -4.2619 | 2.0269E-05 | 6.0072E-05 |
| ENSG00000286125 | 51.5219 | -7.1398 | ##### | -4.2611 | 2.0345E-05 | 6.0290E-05 |
| ENSG00000165782\|PIP4P1 | 2266.4716 | 0.5373 | ##### | 4.2609 | 2.0357E-05 | 6.0321E-05 |

| | | | | | | |
|---|---|---|---|---|---|---|
| ENSG00000123975\|CKS2 | 5445.2997 | 0.5402 | ##### | 4.2602 | 2.0426E-05 | 6.0518E-05 |
| ENSG00000164542\|MATCAP2 | 2038.9923 | -0.5511 | ##### | -4.2601 | 2.0430E-05 | 6.0525E-05 |
| ENSG00000057935\|MTA3 | 4900.0395 | 0.5354 | ##### | 4.2600 | 2.0444E-05 | 6.0560E-05 |
| ENSG00000088356\|PDRG1 | 2874.0397 | 0.5225 | ##### | 4.2594 | 2.0494E-05 | 6.0703E-05 |
| ENSG00000233806\|LINC01237 | 40.3492 | 1.7493 | ##### | 4.2589 | 2.0542E-05 | 6.0837E-05 |
| ENSG00000173681\|BCLAF3 | 485.9000 | -0.6297 | ##### | -4.2589 | 2.0545E-05 | 6.0840E-05 |
| ENSG00000259342 | 83.7537 | 1.1951 | ##### | 4.2584 | 2.0591E-05 | 6.0969E-05 |
| ENSG00000163626\|COX18 | 1376.5670 | -0.5552 | ##### | -4.2581 | 2.0613E-05 | 6.1030E-05 |
| ENSG00000289586 | 364.2826 | 0.6995 | ##### | 4.2578 | 2.0643E-05 | 6.1113E-05 |
| ENSG00000230872\|MFSD13B | 26.5338 | -6.7300 | ##### | -4.2573 | 2.0691E-05 | 6.1249E-05 |
| ENSG00000291071\|FAM86JP | 546.9671 | -0.6335 | ##### | -4.2562 | 2.0798E-05 | 6.1558E-05 |
| ENSG00000152642\|GPD1L | 1366.4926 | -0.5627 | ##### | -4.2561 | 2.0801E-05 | 6.1562E-05 |
| ENSG00000132359\|RAP1GAP2 | 2684.6504 | -0.5242 | ##### | -4.2557 | 2.0841E-05 | 6.1674E-05 |
| ENSG00000103274\|NUBP1 | 2235.0199 | -0.5402 | ##### | -4.2553 | 2.0876E-05 | 6.1771E-05 |
| ENSG00000228536\|LYPLAL1-AS1 | 58.8673 | 7.6252 | ##### | 4.2552 | 2.0885E-05 | 6.1791E-05 |
| ENSG00000184697\|CLDN6 | 19.8131 | 1.6885 | ##### | 4.2552 | 2.0888E-05 | 6.1795E-05 |
| ENSG00000260740 | 13.5489 | -6.0513 | ##### | -4.2542 | 2.0982E-05 | 6.2067E-05 |
| ENSG00000141505\|ASGR1 | 58.9497 | 2.7859 | ##### | 4.2533 | 2.1068E-05 | 6.2314E-05 |
| ENSG00000291232 | 71.7644 | 1.3070 | ##### | 4.2524 | 2.1147E-05 | 6.2544E-05 |
| ENSG00000257433\|RPAP3-DT | 76.5898 | -1.1153 | ##### | -4.2522 | 2.1167E-05 | 6.2595E-05 |
| ENSG00000100348\|TXN2 | 3224.3655 | -0.5323 | ##### | -4.2490 | 2.1474E-05 | 6.3497E-05 |
| ENSG00000235663\|SAPCD1-AS1 | 44.4977 | -1.3642 | ##### | -4.2489 | 2.1485E-05 | 6.3525E-05 |
| ENSG00000178809\|TRIM73 | 103.1188 | -1.1223 | ##### | -4.2487 | 2.1505E-05 | 6.3577E-05 |
| ENSG00000165905\|LARGE2 | 139.5403 | 1.0013 | ##### | 4.2474 | 2.1628E-05 | 6.3932E-05 |
| ENSG00000158711\|ELK4 | 3317.2465 | -0.5409 | ##### | -4.2472 | 2.1645E-05 | 6.3978E-05 |
| ENSG00000269983 | 173.0587 | 0.7693 | ##### | 4.2471 | 2.1660E-05 | 6.4015E-05 |
| ENSG00000010219\|DYRK4 | 1140.7418 | -0.5536 | ##### | -4.2470 | 2.1667E-05 | 6.4029E-05 |
| ENSG00000150712\|MTMR12 | 5681.4770 | 0.5122 | ##### | 4.2466 | 2.1702E-05 | 6.4126E-05 |
| ENSG00000100596\|SPTLC2 | 4306.1739 | 0.5190 | ##### | 4.2466 | 2.1705E-05 | 6.4131E-05 |
| ENSG00000260698 | 18.7896 | 2.3214 | ##### | 4.2461 | 2.1749E-05 | 6.4253E-05 |
| ENSG00000266126 | 19.4486 | 6.2224 | ##### | 4.2457 | 2.1790E-05 | 6.4369E-05 |
| ENSG00000142230\|SAE1 | 14712.2470 | -0.5064 | ##### | -4.2457 | 2.1793E-05 | 6.4369E-05 |
| ENSG00000288907 | 7.3667 | -2.7526 | ##### | -4.2452 | 2.1838E-05 | 6.4498E-05 |
| ENSG00000132664\|POLR3F | 1281.7869 | 0.5551 | ##### | 4.2447 | 2.1889E-05 | 6.4641E-05 |
| ENSG00000138795\|LEF1 | 150.9868 | 5.8827 | ##### | 4.2443 | 2.1926E-05 | 6.4743E-05 |
| ENSG00000115297\|TLX2 | 57.7723 | 2.9099 | ##### | 4.2442 | 2.1933E-05 | 6.4760E-05 |
| ENSG00000160570\|DEDD2 | 1602.5210 | -0.5595 | ##### | -4.2437 | 2.1982E-05 | 6.4896E-05 |
| ENSG00000171100\|MTM1 | 1208.3211 | 0.5964 | ##### | 4.2434 | 2.2018E-05 | 6.4996E-05 |
| ENSG00000188986\|NELFB | 6720.1497 | -0.2571 | ##### | -4.2428 | 2.2079E-05 | 6.5171E-05 |
| ENSG00000270069\|MIR222HG | 110.3323 | 1.1243 | ##### | 4.2418 | 2.2175E-05 | 6.5446E-05 |
| ENSG00000213553\|RPLP0P6 | 4683.4739 | 0.5365 | ##### | 4.2416 | 2.2189E-05 | 6.5482E-05 |
| ENSG00000233885\|YEATS2-AS1 | 32.5805 | -1.5707 | ##### | -4.2414 | 2.2209E-05 | 6.5534E-05 |
| ENSG00000176402\|GJC3 | 36.4809 | 6.0879 | ##### | 4.2401 | 2.2343E-05 | 6.5924E-05 |
| ENSG00000149806\|FAU | 16074.4495 | 0.5228 | ##### | 4.2400 | 2.2352E-05 | 6.5944E-05 |
| ENSG00000256525\|POLG2 | 1531.2871 | 0.5717 | ##### | 4.2399 | 2.2367E-05 | 6.5979E-05 |
| ENSG00000124762\|CDKN1A | 10433.3212 | 0.5275 | ##### | 4.2387 | 2.2480E-05 | 6.6304E-05 |
| ENSG00000257523\|LINC02326 | 15.3026 | -6.3532 | ##### | -4.2387 | 2.2481E-05 | 6.6304E-05 |
| ENSG00000164983\|TMEM65 | 2197.8904 | -0.5135 | ##### | -4.2362 | 2.2737E-05 | 6.7052E-05 |
| ENSG00000133392\|MYH11 | 92.9476 | 1.1240 | ##### | 4.2361 | 2.2741E-05 | 6.7059E-05 |
| ENSG00000269940 | 285.5866 | -0.6433 | ##### | -4.2359 | 2.2760E-05 | 6.7106E-05 |
| ENSG00000254231\|WWP1-AS1 | 12.0714 | -3.6680 | ##### | -4.2348 | 2.2874E-05 | 6.7438E-05 |
| ENSG00000170903\|MSANTD4 | 1634.5980 | 0.5531 | ##### | 4.2341 | 2.2947E-05 | 6.7646E-05 |
| ENSG00000258940\|MIA2-AS1 | 49.8551 | -1.6281 | ##### | -4.2314 | 2.3224E-05 | 6.8455E-05 |
| ENSG00000123689\|G0S2 | 56.1819 | -3.2997 | ##### | -4.2313 | 2.3230E-05 | 6.8465E-05 |
| ENSG00000279155 | 37.0839 | 7.8528 | ##### | 4.2308 | 2.3283E-05 | 6.8615E-05 |
| ENSG00000184669\|OR7E14P | 46.4976 | -7.9612 | ##### | -4.2307 | 2.3298E-05 | 6.8652E-05 |
| ENSG00000275140\|SEC22B3P | 28.1442 | 2.0464 | ##### | 4.2305 | 2.3316E-05 | 6.8700E-05 |
| ENSG00000123353\|ORMDL2 | 5292.6388 | -0.5155 | ##### | -4.2303 | 2.3343E-05 | 6.8771E-05 |
| ENSG00000223547\|ZNF844 | 41.3190 | -7.0867 | ##### | -4.2297 | 2.3399E-05 | 6.8930E-05 |
| ENSG00000127481\|UBR4 | 12608.7724 | 0.5125 | ##### | 4.2296 | 2.3409E-05 | 6.8952E-05 |
| ENSG00000120088\|CRHR1 | 32.0322 | 3.8619 | ##### | 4.2288 | 2.3495E-05 | 6.9199E-05 |
| ENSG00000235586 | 15.6876 | 3.9347 | ##### | 4.2275 | 2.3633E-05 | 6.9599E-05 |
| ENSG00000291287 | 14.3647 | -4.5026 | ##### | -4.2272 | 2.3657E-05 | 6.9663E-05 |

| | | | | | | |
|---|---|---|---|---|---|---|
| ENSG00000268649 | 51.1274 | 3.2861 | ##### | 4.2267 | 2.3712E-05 | 6.9816E-05 |
| ENSG00000204790 | 75.1038 | -8.6015 | ##### | -4.2262 | 2.3766E-05 | 6.9968E-05 |
| ENSG00000120658\|ENOX1 | 417.2326 | -0.6129 | ##### | -4.2252 | 2.3870E-05 | 7.0270E-05 |
| ENSG00000107518\|ATRNL1 | 133.3173 | 1.1108 | ##### | 4.2243 | 2.3969E-05 | 7.0553E-05 |
| ENSG00000287620 | 64.3368 | 7.7665 | ##### | 4.2241 | 2.3989E-05 | 7.0605E-05 |
| ENSG00000165244\|ZNF367 | 2598.9234 | 0.5336 | ##### | 4.2235 | 2.4055E-05 | 7.0793E-05 |
| ENSG00000110076\|NRXN2 | 69.3526 | 1.0401 | ##### | 4.2219 | 2.4222E-05 | 7.1277E-05 |
| ENSG00000230894 | 74.8575 | -8.5966 | ##### | -4.2217 | 2.4251E-05 | 7.1355E-05 |
| ENSG00000290964 | 21.5041 | -6.6021 | ##### | -4.2213 | 2.4286E-05 | 7.1451E-05 |
| ENSG00000230499 | 43.6513 | -1.4303 | ##### | -4.2209 | 2.4335E-05 | 7.1588E-05 |
| ENSG00000176542\|USF3 | 1562.4403 | 0.5506 | ##### | 4.2198 | 2.4447E-05 | 7.1910E-05 |
| ENSG00000276101 | 207.9737 | 0.8210 | ##### | 4.2198 | 2.4452E-05 | 7.1919E-05 |
| ENSG00000109685\|NSD2 | 13474.6985 | 0.4971 | ##### | 4.2195 | 2.4483E-05 | 7.2002E-05 |
| ENSG00000225087 | 130.0300 | -9.4446 | ##### | -4.2189 | 2.4547E-05 | 7.2185E-05 |
| ENSG00000260198 | 34.1493 | 6.1591 | ##### | 4.2174 | 2.4712E-05 | 7.2663E-05 |
| ENSG00000234494\|SP2-AS1 | 263.3849 | -0.7437 | ##### | -4.2171 | 2.4751E-05 | 7.2763E-05 |
| ENSG00000286613 | 22.6150 | -1.9234 | ##### | -4.2171 | 2.4751E-05 | 7.2763E-05 |
| ENSG00000186510\|CLCNKA | 26.5406 | -6.5151 | ##### | -4.2166 | 2.4798E-05 | 7.2892E-05 |
| ENSG00000152894\|PTPRK | 4145.8560 | -0.5046 | ##### | -4.2155 | 2.4928E-05 | 7.3267E-05 |
| ENSG00000153253\|SCN3A | 82.0283 | 7.8544 | ##### | 4.2150 | 2.4977E-05 | 7.3403E-05 |
| ENSG00000085117\|CD82 | 1514.4978 | 0.5713 | ##### | 4.2150 | 2.4980E-05 | 7.3406E-05 |
| ENSG00000011590\|ZBTB32 | 24.2068 | -2.1045 | ##### | -4.2149 | 2.4985E-05 | 7.3415E-05 |
| ENSG00000284505\|LYNX1-SLURP2 | 22.9811 | 1.6553 | ##### | 4.2149 | 2.4988E-05 | 7.3416E-05 |
| ENSG00000174332\|GLIS1 | 126.3395 | 2.0906 | ##### | 4.2145 | 2.5036E-05 | 7.3548E-05 |
| ENSG00000148153\|INIP | 2485.6524 | -0.5254 | ##### | -4.2127 | 2.5236E-05 | 7.4130E-05 |
| ENSG00000159259\|CHAF1B | 2933.7758 | -0.3574 | ##### | -4.2124 | 2.5263E-05 | 7.4202E-05 |
| ENSG00000270179 | 15.3504 | -2.3542 | ##### | -4.2121 | 2.5305E-05 | 7.4319E-05 |
| ENSG00000269984 | 201.5422 | -0.9582 | ##### | -4.2120 | 2.5316E-05 | 7.4343E-05 |
| ENSG00000290366 | 15.3502 | 6.8201 | ##### | 4.2118 | 2.5334E-05 | 7.4387E-05 |
| ENSG00000107362\|ABHD17B | 1605.4918 | 0.5704 | ##### | 4.2118 | 2.5339E-05 | 7.4396E-05 |
| ENSG00000249492 | 124.6240 | 1.0118 | ##### | 4.2114 | 2.5384E-05 | 7.4521E-05 |
| ENSG00000065371\|ROPN1 | 8.3308 | 5.8288 | ##### | 4.2113 | 2.5392E-05 | 7.4537E-05 |
| ENSG00000151498\|ACAD8 | 1381.6898 | 0.5460 | ##### | 4.2108 | 2.5451E-05 | 7.4696E-05 |
| ENSG00000149634\|SPATA25 | 29.3281 | 1.6793 | ##### | 4.2108 | 2.5452E-05 | 7.4696E-05 |
| ENSG00000188460\|ACTBP11 | 220.2567 | -0.7666 | ##### | -4.2104 | 2.5494E-05 | 7.4814E-05 |
| ENSG00000254501 | 252.0574 | 0.9482 | ##### | 4.2102 | 2.5511E-05 | 7.4857E-05 |
| ENSG00000291055 | 286.7333 | -0.7809 | ##### | -4.2093 | 2.5622E-05 | 7.5173E-05 |
| ENSG00000288911 | 61.7014 | 1.2292 | ##### | 4.2076 | 2.5809E-05 | 7.5716E-05 |
| ENSG00000253629 | 13.2720 | -6.1452 | ##### | -4.2072 | 2.5858E-05 | 7.5851E-05 |
| ENSG00000236714\|LINC01844 | 32.3508 | 7.6701 | ##### | 4.2070 | 2.5876E-05 | 7.5896E-05 |
| ENSG00000178425\|NT5DC1 | 2440.1162 | -0.5188 | ##### | -4.2067 | 2.5913E-05 | 7.5999E-05 |
| ENSG00000210077\|MT-TV | 184.2270 | 0.7634 | ##### | 4.2062 | 2.5973E-05 | 7.6168E-05 |
| ENSG00000164070\|HSPA4L | 2351.2741 | -0.5337 | ##### | -4.2060 | 2.5995E-05 | 7.6223E-05 |
| ENSG00000216359 | 20.6829 | -6.5009 | ##### | -4.2058 | 2.6011E-05 | 7.6264E-05 |
| ENSG00000118007\|STAG1 | 5264.7103 | 0.4893 | ##### | 4.2036 | 2.6268E-05 | 7.7010E-05 |
| ENSG00000274253 | 70.0506 | 1.4820 | ##### | 4.2032 | 2.6313E-05 | 7.7133E-05 |
| ENSG00000179295\|PTPN11 | 26687.8149 | 0.4940 | ##### | 4.2032 | 2.6321E-05 | 7.7149E-05 |
| ENSG00000124251\|TP53TG5 | 75.1799 | 1.0123 | ##### | 4.2026 | 2.6388E-05 | 7.7338E-05 |
| ENSG00000111596\|CNOT2 | 4666.0827 | 0.5209 | ##### | 4.2025 | 2.6401E-05 | 7.7354E-05 |
| ENSG00000115263\|GCG | 163.5564 | 7.6522 | ##### | 4.2025 | 2.6399E-05 | 7.7354E-05 |
| ENSG00000188152\|NUTM2G | 49.3758 | 2.7569 | ##### | 4.2025 | 2.6397E-05 | 7.7354E-05 |
| ENSG00000247903 | 196.1057 | 0.7593 | ##### | 4.2023 | 2.6425E-05 | 7.7415E-05 |
| ENSG00000170289\|CNGB3 | 38.1284 | -7.6746 | ##### | -4.2020 | 2.6458E-05 | 7.7506E-05 |
| ENSG00000163116\|STPG2 | 29.5905 | -4.8096 | ##### | -4.2019 | 2.6464E-05 | 7.7516E-05 |
| ENSG00000291073 | 1326.9186 | 0.6123 | ##### | 4.2010 | 2.6575E-05 | 7.7833E-05 |
| ENSG00000279187 | 10.8732 | 2.5528 | ##### | 4.2001 | 2.6677E-05 | 7.8123E-05 |
| ENSG00000255836 | 490.1523 | -0.7287 | ##### | -4.1998 | 2.6721E-05 | 7.8244E-05 |
| ENSG00000279140 | 24.5632 | -1.5804 | ##### | -4.1990 | 2.6811E-05 | 7.8500E-05 |
| ENSG00000146197\|SCUBE3 | 35.6800 | 1.8123 | ##### | 4.1989 | 2.6826E-05 | 7.8536E-05 |
| ENSG00000289041 | 22.0753 | 1.8601 | ##### | 4.1985 | 2.6872E-05 | 7.8665E-05 |
| ENSG00000226084\|RPL17P6 | 911.9555 | -0.5508 | ##### | -4.1977 | 2.6959E-05 | 7.8911E-05 |
| ENSG00000235946\|H2BW3P | 26.2661 | 1.9129 | ##### | 4.1971 | 2.7033E-05 | 7.9120E-05 |
| ENSG00000164744\|SUN3 | 80.3017 | -6.1555 | ##### | -4.1967 | 2.7088E-05 | 7.9272E-05 |

| | | | | | | |
|---|---|---|---|---|---|---|
| ENSG00000142208\|AKT1 | 12342.5530 | 0.4788 | ##### | 4.1963 | 2.7132E-05 | 7.9394E-05 |
| ENSG00000184357\|H1-5 | 47.6103 | 1.4511 | ##### | 4.1962 | 2.7138E-05 | 7.9405E-05 |
| ENSG00000067836\|ROGDI | 891.1582 | -0.5949 | ##### | -4.1959 | 2.7184E-05 | 7.9531E-05 |
| ENSG00000203780\|FANK1 | 141.8322 | -0.8720 | ##### | -4.1949 | 2.7294E-05 | 7.9844E-05 |
| ENSG00000198830\|HMGN2 | 16461.3375 | 0.5100 | ##### | 4.1947 | 2.7324E-05 | 7.9923E-05 |
| ENSG00000110218\|PANX1 | 1753.4000 | -0.5305 | ##### | -4.1939 | 2.7414E-05 | 8.0180E-05 |
| ENSG00000132000\|PODNL1 | 83.9168 | -1.3480 | ##### | -4.1939 | 2.7424E-05 | 8.0202E-05 |
| ENSG00000289463 | 19.9865 | 2.7278 | ##### | 4.1938 | 2.7436E-05 | 8.0229E-05 |
| ENSG00000129467\|ADCY4 | 49.4474 | 6.7350 | ##### | 4.1923 | 2.7611E-05 | 8.0733E-05 |
| ENSG00000123977\|DAW1 | 293.7839 | 1.1506 | ##### | 4.1922 | 2.7622E-05 | 8.0758E-05 |
| ENSG00000288906 | 42.3346 | 1.6443 | ##### | 4.1921 | 2.7633E-05 | 8.0780E-05 |
| ENSG00000235904\|RBMS3-AS3 | 52.4411 | -3.7554 | ##### | -4.1914 | 2.7721E-05 | 8.1031E-05 |
| ENSG00000271869\|DCTN6-DT | 60.1713 | -1.3022 | ##### | -4.1911 | 2.7763E-05 | 8.1145E-05 |
| ENSG00000286091 | 19.5814 | 4.7842 | ##### | 4.1909 | 2.7791E-05 | 8.1218E-05 |
| ENSG00000165806\|CASP7 | 1538.4886 | -0.5419 | ##### | -4.1906 | 2.7827E-05 | 8.1316E-05 |
| ENSG00000197756\|RPL37A | 41967.1863 | 0.5527 | ##### | 4.1905 | 2.7839E-05 | 8.1344E-05 |
| ENSG00000152689\|RASGRP3 | 57.2290 | -1.2361 | ##### | -4.1901 | 2.7881E-05 | 8.1458E-05 |
| ENSG00000204758\|RPL26L1-AS1 | 131.8737 | -0.8295 | ##### | -4.1900 | 2.7894E-05 | 8.1479E-05 |
| ENSG00000102003\|SYP | 462.9906 | 0.7446 | ##### | 4.1900 | 2.7893E-05 | 8.1479E-05 |
| ENSG00000151239\|TWF1 | 23167.1977 | 0.3759 | ##### | 4.1895 | 2.7958E-05 | 8.1659E-05 |
| ENSG00000198001\|IRAK4 | 1297.3723 | -0.5597 | ##### | -4.1893 | 2.7979E-05 | 8.1713E-05 |
| ENSG00000166529\|ZSCAN21 | 737.8105 | 0.5924 | ##### | 4.1892 | 2.7991E-05 | 8.1739E-05 |
| ENSG00000198142\|SOWAHC | 4162.4910 | -0.4951 | ##### | -4.1890 | 2.8013E-05 | 8.1796E-05 |
| ENSG00000124356\|STAMBP | 4763.9881 | 0.4871 | ##### | 4.1887 | 2.8055E-05 | 8.1911E-05 |
| ENSG00000167113\|COQ4 | 1967.3215 | -0.5205 | ##### | -4.1872 | 2.8241E-05 | 8.2445E-05 |
| ENSG00000210184\|MT-TS2 | 132.9800 | 0.8828 | ##### | 4.1871 | 2.8254E-05 | 8.2476E-05 |
| ENSG00000118412\|CASP8AP2 | 2446.3755 | 0.5309 | ##### | 4.1868 | 2.8287E-05 | 8.2564E-05 |
| ENSG00000224817\|LINC03046 | 409.7407 | -0.6257 | ##### | -4.1866 | 2.8314E-05 | 8.2635E-05 |
| ENSG00000261829 | 22.4413 | -6.9041 | ##### | -4.1859 | 2.8408E-05 | 8.2900E-05 |
| ENSG00000113391\|FAM172A | 1770.0741 | 0.5625 | ##### | 4.1855 | 2.8456E-05 | 8.3032E-05 |
| ENSG00000239523\|MYLK-AS1 | 123.6470 | 0.9089 | ##### | 4.1851 | 2.8509E-05 | 8.3179E-05 |
| ENSG00000064270\|ATP2C2 | 66.8299 | -1.2191 | ##### | -4.1841 | 2.8627E-05 | 8.3516E-05 |
| ENSG00000267106\|ZNF561-AS1 | 334.5645 | -2.5427 | ##### | -4.1839 | 2.8653E-05 | 8.3583E-05 |
| ENSG00000121766\|ZCCHC17 | 2831.6118 | -0.5152 | ##### | -4.1836 | 2.8689E-05 | 8.3680E-05 |
| ENSG00000287104 | 48.4886 | -1.5663 | ##### | -4.1836 | 2.8698E-05 | 8.3699E-05 |
| ENSG00000148082\|SHC3 | 165.8742 | 0.8161 | ##### | 4.1829 | 2.8787E-05 | 8.3950E-05 |
| ENSG00000258701\|LINC00638 | 94.9469 | -1.0514 | ##### | -4.1828 | 2.8793E-05 | 8.3958E-05 |
| ENSG00000177108\|ZDHHC22 | 144.1344 | 5.8643 | ##### | 4.1827 | 2.8803E-05 | 8.3979E-05 |
| ENSG00000124383\|MPHOSPH10 | 5169.1462 | 0.3791 | ##### | 4.1824 | 2.8844E-05 | 8.4092E-05 |
| ENSG00000168300\|PCMTD1 | 1774.1065 | -0.5437 | ##### | -4.1818 | 2.8917E-05 | 8.4297E-05 |
| ENSG00000119737\|GPR75 | 150.1321 | 0.8278 | ##### | 4.1811 | 2.9008E-05 | 8.4553E-05 |
| ENSG00000160392\|C19orf47 | 1457.3712 | -0.5215 | ##### | -4.1810 | 2.9029E-05 | 8.4605E-05 |
| ENSG00000273784 | 483.0526 | 0.9396 | ##### | 4.1803 | 2.9109E-05 | 8.4830E-05 |
| ENSG00000137825\|ITPKA | 121.5722 | 0.8406 | ##### | 4.1802 | 2.9127E-05 | 8.4875E-05 |
| ENSG00000121898\|CPXM2 | 13.4641 | 7.0177 | ##### | 4.1794 | 2.9222E-05 | 8.5143E-05 |
| ENSG00000004975\|DVL2 | 3480.1201 | 0.5210 | ##### | 4.1786 | 2.9333E-05 | 8.5458E-05 |
| ENSG00000113319\|RASGRF2 | 124.5553 | 1.0858 | ##### | 4.1778 | 2.9437E-05 | 8.5751E-05 |
| ENSG00000223382\|LINC01778 | 57.2130 | 7.6107 | ##### | 4.1772 | 2.9510E-05 | 8.5957E-05 |
| ENSG00000132881\|CPLANE2 | 362.9161 | -0.6548 | ##### | -4.1768 | 2.9560E-05 | 8.6086E-05 |
| ENSG00000261487 | 27.6173 | -1.5726 | ##### | -4.1768 | 2.9558E-05 | 8.6086E-05 |
| ENSG00000128045\|RASL11B | 38.0728 | 4.1999 | ##### | 4.1752 | 2.9775E-05 | 8.6704E-05 |
| ENSG00000111540\|RAB5B | 6123.3584 | -0.4852 | ##### | -4.1745 | 2.9870E-05 | 8.6971E-05 |
| ENSG00000248405\|PRR5-ARHGAP8 | 41.7846 | 4.4605 | ##### | 4.1744 | 2.9873E-05 | 8.6971E-05 |
| ENSG00000290793 | 110.2720 | 0.9538 | ##### | 4.1743 | 2.9888E-05 | 8.7007E-05 |
| ENSG00000136153\|LMO7 | 1565.8411 | -0.5422 | ##### | -4.1741 | 2.9916E-05 | 8.7080E-05 |
| ENSG00000108423\|TUBD1 | 891.7114 | 0.5631 | ##### | 4.1734 | 3.0011E-05 | 8.7347E-05 |
| ENSG00000249572\|TARS1-DT | 183.8843 | -0.8689 | ##### | -4.1732 | 3.0032E-05 | 8.7400E-05 |
| ENSG00000273455 | 19.0763 | 6.4129 | ##### | 4.1722 | 3.0171E-05 | 8.7796E-05 |
| ENSG00000174384\|PMS2P6 | 170.3133 | -0.7735 | ##### | -4.1721 | 3.0180E-05 | 8.7815E-05 |
| ENSG00000262728\|ARHGAP11A-DT | 34.1624 | -1.5039 | ##### | -4.1720 | 3.0200E-05 | 8.7863E-05 |
| ENSG00000269729 | 12.8048 | -6.6038 | ##### | -4.1718 | 3.0221E-05 | 8.7916E-05 |
| ENSG00000118263\|KLF7 | 2447.1174 | 0.5307 | ##### | 4.1717 | 3.0232E-05 | 8.7941E-05 |
| ENSG00000148337\|CIZ1 | 10216.5804 | -0.4735 | ##### | -4.1715 | 3.0258E-05 | 8.8006E-05 |

| | | | | | | |
|---|---|---|---|---|---|---|
| ENSG00000258451\|EGILA | 36.7184 | -1.2928 | ##### | -4.1714 | 3.0276E-05 | 8.8052E-05 |
| ENSG00000099991\|CABIN1 | 3945.7719 | 0.5041 | ##### | 4.1701 | 3.0448E-05 | 8.8543E-05 |
| ENSG00000005238\|ATOSB | 1979.8195 | 0.5421 | ##### | 4.1697 | 3.0494E-05 | 8.8669E-05 |
| ENSG00000119725\|ZNF410 | 3589.4236 | 1.3073 | ##### | 4.1670 | 3.0866E-05 | 8.9740E-05 |
| ENSG00000137965\|IFI44 | 65.4011 | -5.3585 | ##### | -4.1667 | 3.0909E-05 | 8.9858E-05 |
| ENSG00000167785\|ZNF558 | 1783.1815 | 0.5336 | ##### | 4.1653 | 3.1095E-05 | 9.0388E-05 |
| ENSG00000110811\|P3H3 | 1761.1783 | 0.5175 | ##### | 4.1647 | 3.1175E-05 | 9.0612E-05 |
| ENSG00000232803\|SLCO4A1-AS1 | 56.4222 | 1.7257 | ##### | 4.1633 | 3.1364E-05 | 9.1154E-05 |
| ENSG00000177889\|UBE2N | 9775.7569 | 0.5057 | ##### | 4.1631 | 3.1400E-05 | 9.1249E-05 |
| ENSG00000236432\|MFF-DT | 171.9250 | -0.8819 | ##### | -4.1629 | 3.1430E-05 | 9.1326E-05 |
| ENSG00000254230 | 51.0155 | 1.4968 | ##### | 4.1628 | 3.1434E-05 | 9.1330E-05 |
| ENSG00000274523\|RCC1L | 6274.8872 | -0.4929 | ##### | -4.1628 | 3.1444E-05 | 9.1348E-05 |
| ENSG00000231742\|LINC01273 | 87.6738 | 1.2510 | ##### | 4.1625 | 3.1482E-05 | 9.1452E-05 |
| ENSG00000283689 | 98.5402 | 1.1790 | ##### | 4.1618 | 3.1574E-05 | 9.1710E-05 |
| ENSG00000131781\|FMO5 | 128.2918 | -0.8791 | ##### | -4.1612 | 3.1655E-05 | 9.1936E-05 |
| ENSG00000131269\|ABCB7 | 1644.7785 | 0.5405 | ##### | 4.1611 | 3.1669E-05 | 9.1967E-05 |
| ENSG00000008323\|PLEKHG6 | 169.7582 | 0.8490 | ##### | 4.1610 | 3.1686E-05 | 9.2008E-05 |
| ENSG00000203799\|CCDC162P | 34.1020 | -3.0941 | ##### | -4.1607 | 3.1722E-05 | 9.2103E-05 |
| ENSG00000253893\|FAM85B | 16.3845 | -3.5870 | ##### | -4.1607 | 3.1728E-05 | 9.2111E-05 |
| ENSG00000167114\|SLC27A4 | 5225.0493 | -0.4996 | ##### | -4.1594 | 3.1906E-05 | 9.2620E-05 |
| ENSG00000214077\|GNAQP1 | 29.5281 | 1.9030 | ##### | 4.1591 | 3.1944E-05 | 9.2723E-05 |
| ENSG00000163159\|VPS72 | 5047.2496 | 0.4988 | ##### | 4.1588 | 3.1987E-05 | 9.2837E-05 |
| ENSG00000008324\|SS18L2 | 799.9145 | 0.5657 | ##### | 4.1585 | 3.2036E-05 | 9.2971E-05 |
| ENSG00000213079\|SCAF8 | 3660.0007 | 0.5076 | ##### | 4.1572 | 3.2223E-05 | 9.3504E-05 |
| ENSG00000170634\|ACYP2 | 662.6646 | -0.5740 | ##### | -4.1566 | 3.2298E-05 | 9.3711E-05 |
| ENSG00000234776\|FREY1 | 36.3370 | 1.8828 | ##### | 4.1566 | 3.2306E-05 | 9.3725E-05 |
| ENSG00000254531\|FLJ20021 | 99.9219 | -1.1970 | ##### | -4.1558 | 3.2412E-05 | 9.4024E-05 |
| ENSG00000077549\|CAPZB | 17497.3143 | 0.5123 | ##### | 4.1556 | 3.2442E-05 | 9.4104E-05 |
| ENSG00000148832\|PAOX | 148.9093 | -0.8685 | ##### | -4.1549 | 3.2542E-05 | 9.4383E-05 |
| ENSG00000005073\|HOXA11 | 634.8829 | -0.5685 | ##### | -4.1547 | 3.2572E-05 | 9.4463E-05 |
| ENSG00000125744\|RTN2 | 925.2633 | -0.5685 | ##### | -4.1546 | 3.2591E-05 | 9.4506E-05 |
| ENSG00000204149\|AGAP6 | 884.9080 | 0.5898 | ##### | 4.1542 | 3.2643E-05 | 9.4650E-05 |
| ENSG00000272482 | 50.5338 | -5.6498 | ##### | -4.1536 | 3.2728E-05 | 9.4885E-05 |
| ENSG00000184481\|FOXO4 | 388.4032 | 0.6922 | ##### | 4.1531 | 3.2806E-05 | 9.5102E-05 |
| ENSG00000263823 | 59.0912 | 1.1714 | ##### | 4.1521 | 3.2949E-05 | 9.5509E-05 |
| ENSG00000132661\|NXT1 | 1791.2074 | 0.5471 | ##### | 4.1520 | 3.2959E-05 | 9.5528E-05 |
| ENSG00000111490\|TBC1D30 | 558.4317 | -0.5911 | ##### | -4.1506 | 3.3162E-05 | 9.6107E-05 |
| ENSG00000260903\|XKR7 | 32.9025 | 1.7422 | ##### | 4.1503 | 3.3199E-05 | 9.6204E-05 |
| ENSG00000196366\|C9orf163 | 71.9585 | 1.2589 | ##### | 4.1485 | 3.3472E-05 | 9.6988E-05 |
| ENSG00000180900\|SCRIB | 18084.6450 | 0.4909 | ##### | 4.1483 | 3.3489E-05 | 9.7028E-05 |
| ENSG00000123411\|IKZF4 | 462.2633 | 0.6007 | ##### | 4.1473 | 3.3637E-05 | 9.7446E-05 |
| ENSG00000158623\|COPG2 | 3568.3785 | 0.5123 | ##### | 4.1473 | 3.3645E-05 | 9.7461E-05 |
| ENSG00000205269\|TMEM170B | 597.7213 | 0.6188 | ##### | 4.1471 | 3.3672E-05 | 9.7527E-05 |
| ENSG00000130307\|USHBP1 | 28.2907 | 2.0426 | ##### | 4.1468 | 3.3712E-05 | 9.7634E-05 |
| ENSG00000184560\|SPEM2 | 60.1114 | 7.8034 | ##### | 4.1467 | 3.3732E-05 | 9.7683E-05 |
| ENSG00000138380\|CARF | 297.9605 | 0.6503 | ##### | 4.1462 | 3.3811E-05 | 9.7902E-05 |
| ENSG00000110583\|NAA40 | 2574.0777 | 0.5232 | ##### | 4.1458 | 3.3861E-05 | 9.8036E-05 |
| ENSG00000157578\|LCA5L | 140.2157 | -1.2104 | ##### | -4.1453 | 3.3931E-05 | 9.8231E-05 |
| ENSG00000258663 | 12.6423 | 5.5454 | ##### | 4.1452 | 3.3951E-05 | 9.8280E-05 |
| ENSG00000125356\|NDUFA1 | 4064.5166 | 0.5642 | ##### | 4.1444 | 3.4065E-05 | 9.8599E-05 |
| ENSG00000196136\|SERPINA3 | 41.5149 | -4.5107 | ##### | -4.1443 | 3.4092E-05 | 9.8669E-05 |
| ENSG00000243819\|RN7SL832P | 37.7097 | -1.3876 | ##### | -4.1437 | 3.4172E-05 | 9.8891E-05 |
| ENSG00000212296\|SNORD72 | 13.0189 | 5.0202 | ##### | 4.1431 | 3.4264E-05 | 9.9147E-05 |
| ENSG00000114115\|RBP1 | 24.4961 | -1.7968 | ##### | -4.1430 | 3.4274E-05 | 9.9166E-05 |
| ENSG00000154719\|MRPL39 | 2226.2293 | 0.5287 | ##### | 4.1427 | 3.4327E-05 | 9.9309E-05 |
| ENSG00000148180\|GSN | 5969.8511 | -0.4866 | ##### | -4.1425 | 3.4349E-05 | 9.9364E-05 |
| ENSG00000275464 | 3038.6865 | -0.5589 | ##### | -4.1422 | 3.4392E-05 | 9.9478E-05 |
| ENSG00000185710\|SMG1P4 | 508.7853 | 0.7410 | ##### | 4.1421 | 3.4419E-05 | 9.9547E-05 |
| ENSG00000124549\|BTN2A3P | 510.8726 | 6.0895 | ##### | 4.1419 | 3.4437E-05 | 9.9589E-05 |
| ENSG00000172061\|LRRC15 | 50.4368 | 2.2383 | ##### | 4.1417 | 3.4474E-05 | 9.9688E-05 |
| ENSG00000140986\|RPL3L | 22.7716 | -4.1347 | ##### | -4.1415 | 3.4511E-05 | 9.9784E-05 |
| ENSG00000242261 | 25.8243 | 1.5172 | ##### | 4.1405 | 3.4662E-05 | 1.0021E-04 |
| ENSG00000168556\|ING2 | 622.2698 | -0.6257 | ##### | -4.1404 | 3.4672E-05 | 1.0023E-04 |

| | | | | | | |
|---|---|---|---|---|---|---|
| ENSG00000056050\|HPF1 | 1413.2770 | -0.5464 | ##### | -4.1403 | 3.4678E-05 | 1.0024E-04 |
| ENSG00000227039\|ITGB2-AS1 | 22.0124 | -2.1890 | ##### | -4.1399 | 3.4749E-05 | 1.0043E-04 |
| ENSG00000188263\|IL17REL | 39.3871 | -7.7209 | ##### | -4.1392 | 3.4853E-05 | 1.0073E-04 |
| ENSG00000132478\|UNK | 2804.7473 | 0.5240 | ##### | 4.1390 | 3.4882E-05 | 1.0080E-04 |
| ENSG00000170498\|KISS1 | 50.1945 | 2.9035 | ##### | 4.1388 | 3.4918E-05 | 1.0089E-04 |
| ENSG00000124006\|OBSL1 | 4069.2214 | -0.4996 | ##### | -4.1385 | 3.4954E-05 | 1.0099E-04 |
| ENSG00000127329\|PTPRB | 157.7548 | 0.9621 | ##### | 4.1380 | 3.5040E-05 | 1.0123E-04 |
| ENSG00000168502\|MTCL1 | 4933.4530 | -0.5024 | ##### | -4.1378 | 3.5069E-05 | 1.0130E-04 |
| ENSG00000205937\|RNPS1 | 17239.7918 | 0.4821 | ##### | 4.1377 | 3.5085E-05 | 1.0133E-04 |
| ENSG00000140265\|ZSCAN29 | 2088.4131 | 0.5217 | ##### | 4.1375 | 3.5107E-05 | 1.0139E-04 |
| ENSG00000243708\|PLA2G4B | 47.4098 | 3.6162 | ##### | 4.1372 | 3.5153E-05 | 1.0151E-04 |
| ENSG00000198189\|HSD17B11 | 4232.8059 | 0.5087 | ##### | 4.1371 | 3.5165E-05 | 1.0154E-04 |
| ENSG00000239559\|RPL37P2 | 64.6910 | 1.0748 | ##### | 4.1371 | 3.5177E-05 | 1.0156E-04 |
| ENSG00000122068\|FYTTD1 | 5945.5642 | -0.5166 | ##### | -4.1370 | 3.5190E-05 | 1.0159E-04 |
| ENSG00000167186\|COQ7 | 911.8211 | -0.5391 | ##### | -4.1358 | 3.5366E-05 | 1.0209E-04 |
| ENSG00000179058\|C9orf50 | 12.4518 | -4.3607 | ##### | -4.1348 | 3.5524E-05 | 1.0253E-04 |
| ENSG00000273387 | 17.5801 | 5.2365 | ##### | 4.1344 | 3.5591E-05 | 1.0272E-04 |
| ENSG00000152818\|UTRN | 2940.4293 | -0.5409 | ##### | -4.1334 | 3.5746E-05 | 1.0315E-04 |
| ENSG00000230371\|KARS1P2 | 15.8191 | 6.7612 | ##### | 4.1333 | 3.5757E-05 | 1.0318E-04 |
| ENSG00000141428\|C18orf21 | 1178.1453 | -0.5458 | ##### | -4.1327 | 3.5854E-05 | 1.0345E-04 |
| ENSG00000135722\|FBXL8 | 514.5842 | -0.6006 | ##### | -4.1325 | 3.5889E-05 | 1.0354E-04 |
| ENSG00000175344\|CHRNA7 | 456.2159 | -0.6324 | ##### | -4.1324 | 3.5899E-05 | 1.0356E-04 |
| ENSG00000171530\|TBCA | 6677.2223 | 0.4976 | ##### | 4.1320 | 3.5965E-05 | 1.0374E-04 |
| ENSG00000248099\|INSL3 | 55.1011 | -1.1894 | ##### | -4.1318 | 3.5988E-05 | 1.0379E-04 |
| ENSG00000109680\|TBC1D19 | 271.7195 | -0.7069 | ##### | -4.1309 | 3.6128E-05 | 1.0419E-04 |
| ENSG00000146833\|TRIM4 | 2795.6599 | -0.3862 | ##### | -4.1303 | 3.6236E-05 | 1.0449E-04 |
| ENSG00000293316\|PLEKHA8P1 | 562.2339 | -0.6113 | ##### | -4.1301 | 3.6267E-05 | 1.0457E-04 |
| ENSG00000198868\|MTND4LP30 | 16.4739 | 2.0002 | ##### | 4.1296 | 3.6338E-05 | 1.0476E-04 |
| ENSG00000181904\|C5orf24 | 6512.8695 | -0.5007 | ##### | -4.1292 | 3.6406E-05 | 1.0495E-04 |
| ENSG00000142632\|ARHGEF19 | 2076.7108 | 0.5422 | ##### | 4.1289 | 3.6443E-05 | 1.0504E-04 |
| ENSG00000181638\|ZFP41 | 2072.6174 | 0.5284 | ##### | 4.1289 | 3.6447E-05 | 1.0505E-04 |
| ENSG00000138028\|CGREF1 | 1560.7165 | -0.5261 | ##### | -4.1286 | 3.6502E-05 | 1.0519E-04 |
| ENSG00000147180\|ZNF711 | 593.5011 | 0.6364 | ##### | 4.1285 | 3.6509E-05 | 1.0520E-04 |
| ENSG00000255568\|BRWD1-AS2 | 27.8294 | -2.0797 | ##### | -4.1285 | 3.6513E-05 | 1.0521E-04 |
| ENSG00000162367\|TAL1 | 10.4014 | -3.7393 | ##### | -4.1281 | 3.6584E-05 | 1.0540E-04 |
| ENSG00000293098 | 110.4332 | 0.9094 | ##### | 4.1280 | 3.6595E-05 | 1.0542E-04 |
| ENSG00000100014\|SPECC1L | 5152.8860 | 0.4905 | ##### | 4.1275 | 3.6666E-05 | 1.0562E-04 |
| ENSG00000290890 | 363.5304 | -0.9910 | ##### | -4.1274 | 3.6694E-05 | 1.0569E-04 |
| ENSG00000279386 | 17.4556 | 3.8247 | ##### | 4.1271 | 3.6737E-05 | 1.0580E-04 |
| ENSG00000246250 | 106.7761 | 5.0003 | ##### | 4.1270 | 3.6758E-05 | 1.0585E-04 |
| ENSG00000163291\|PAQR3 | 3669.9727 | 0.5149 | ##### | 4.1259 | 3.6929E-05 | 1.0633E-04 |
| ENSG00000117751\|PPP1R8 | 4301.5796 | 0.4266 | ##### | 4.1258 | 3.6949E-05 | 1.0638E-04 |
| ENSG00000079263\|SP140 | 101.0365 | -9.0916 | ##### | -4.1251 | 3.7054E-05 | 1.0667E-04 |
| ENSG00000144048\|DUSP11 | 2575.6273 | 0.4793 | ##### | 4.1245 | 3.7157E-05 | 1.0696E-04 |
| ENSG00000246334\|PRR7-AS1 | 113.5495 | 0.8250 | ##### | 4.1243 | 3.7189E-05 | 1.0704E-04 |
| ENSG00000136960\|ENPP2 | 79.6588 | -5.7493 | ##### | -4.1239 | 3.7248E-05 | 1.0720E-04 |
| ENSG00000145623\|OSMR | 29673.0301 | 0.4497 | ##### | 4.1236 | 3.7299E-05 | 1.0734E-04 |
| ENSG00000198492\|YTHDF2 | 7815.6704 | 0.5078 | ##### | 4.1235 | 3.7321E-05 | 1.0739E-04 |
| ENSG00000139651\|ZNF740 | 4147.0741 | 0.4872 | ##### | 4.1233 | 3.7347E-05 | 1.0745E-04 |
| ENSG00000148824\|MTG1 | 3250.0609 | 0.3739 | ##### | 4.1223 | 3.7509E-05 | 1.0790E-04 |
| ENSG00000137764\|MAP2K5 | 855.6197 | 0.5913 | ##### | 4.1223 | 3.7511E-05 | 1.0790E-04 |
| ENSG00000171988\|JMJD1C | 4685.1483 | 0.5010 | ##### | 4.1215 | 3.7647E-05 | 1.0828E-04 |
| ENSG00000145439\|CBR4 | 948.6521 | -0.5560 | ##### | -4.1212 | 3.7695E-05 | 1.0841E-04 |
| ENSG00000249592\|PCGF3-AS1 | 618.8678 | 0.6377 | ##### | 4.1209 | 3.7740E-05 | 1.0853E-04 |
| ENSG00000037897\|METTL1 | 2757.0512 | 0.5049 | ##### | 4.1206 | 3.7781E-05 | 1.0864E-04 |
| ENSG00000154217\|PITPNC1 | 1833.2258 | 0.5478 | ##### | 4.1206 | 3.7791E-05 | 1.0866E-04 |
| ENSG00000277791\|PSMB3 | 13332.5885 | 0.5010 | ##### | 4.1205 | 3.7813E-05 | 1.0871E-04 |
| ENSG00000198218\|QRICH1 | 3804.9030 | 0.5547 | ##### | 4.1204 | 3.7819E-05 | 1.0872E-04 |
| ENSG00000286626 | 16.1702 | 2.3236 | ##### | 4.1204 | 3.7825E-05 | 1.0872E-04 |
| ENSG00000130175\|PRKCSH | 24294.7621 | 0.4310 | ##### | 4.1196 | 3.7955E-05 | 1.0909E-04 |
| ENSG00000174915\|PTDSS2 | 3274.6827 | 0.5174 | ##### | 4.1192 | 3.8022E-05 | 1.0927E-04 |
| ENSG00000228305\|PRELID1P6 | 704.3228 | 0.5685 | ##### | 4.1191 | 3.8032E-05 | 1.0929E-04 |
| ENSG00000084676\|NCOA1 | 2108.5616 | -0.5291 | ##### | -4.1191 | 3.8042E-05 | 1.0931E-04 |

| | | | | | | |
|---|---|---|---|---|---|---|
| ENSG00000231160\|KLF3-AS1 | 38.5030 | -1.2203 | ##### | -4.1187 | 3.8103E-05 | 1.0947E-04 |
| ENSG00000072310\|SREBF1 | 6631.9124 | 0.4888 | ##### | 4.1175 | 3.8302E-05 | 1.1003E-04 |
| ENSG00000004961\|HCCS | 3586.5858 | -0.3601 | ##### | -4.1175 | 3.8307E-05 | 1.1004E-04 |
| ENSG00000077380\|DYNC1I2 | 8807.7666 | 0.4783 | ##### | 4.1174 | 3.8320E-05 | 1.1006E-04 |
| ENSG00000068028\|RASSF1 | 1872.6410 | 0.5338 | ##### | 4.1174 | 3.8325E-05 | 1.1007E-04 |
| ENSG00000241360\|PDXP | 2369.3030 | 0.5285 | ##### | 4.1168 | 3.8422E-05 | 1.1033E-04 |
| ENSG00000159904\|ZNF890P | 16.4734 | 3.4019 | ##### | 4.1166 | 3.8451E-05 | 1.1041E-04 |
| ENSG00000204055\|IER5L-AS1 | 266.7084 | -3.6223 | ##### | -4.1163 | 3.8500E-05 | 1.1054E-04 |
| ENSG00000009844\|VTA1 | 7612.0644 | -0.5091 | ##### | -4.1158 | 3.8586E-05 | 1.1077E-04 |
| ENSG00000154781\|CCDC174 | 1138.7288 | 0.5594 | ##### | 4.1156 | 3.8613E-05 | 1.1084E-04 |
| ENSG00000214121\|PRDX1P1 | 12.6968 | -3.0764 | ##### | -4.1155 | 3.8641E-05 | 1.1091E-04 |
| ENSG00000271662\|PPP4R2P6 | 16.5244 | -1.8811 | ##### | -4.1149 | 3.8735E-05 | 1.1117E-04 |
| ENSG00000271913\|TAGAP-AS1 | 26.9758 | 1.7562 | ##### | 4.1149 | 3.8738E-05 | 1.1117E-04 |
| ENSG00000104331\|BPNT2 | 13923.2744 | -0.4739 | ##### | -4.1144 | 3.8824E-05 | 1.1140E-04 |
| ENSG00000234233\|KCNH1-IT1 | 15.4330 | 6.3417 | ##### | 4.1143 | 3.8834E-05 | 1.1142E-04 |
| ENSG00000095015\|MAP3K1 | 2416.6554 | -0.5180 | ##### | -4.1143 | 3.8842E-05 | 1.1143E-04 |
| ENSG00000258890\|CEP95 | 1884.6328 | 0.4834 | ##### | 4.1139 | 3.8904E-05 | 1.1160E-04 |
| ENSG00000176438\|SYNE3 | 87.5620 | -0.9712 | ##### | -4.1137 | 3.8938E-05 | 1.1169E-04 |
| ENSG00000111679\|PTPN6 | 717.4840 | 0.5677 | ##### | 4.1126 | 3.9116E-05 | 1.1219E-04 |
| ENSG00000104290\|FZD3 | 737.7870 | 0.5780 | ##### | 4.1124 | 3.9161E-05 | 1.1231E-04 |
| ENSG00000235010\|JAKMIP3-AS1 | 6.9715 | -3.1038 | ##### | -4.1123 | 3.9177E-05 | 1.1234E-04 |
| ENSG00000159712\|ANKRD18CP | 28.9180 | -4.4968 | ##### | -4.1122 | 3.9192E-05 | 1.1237E-04 |
| ENSG00000290835 | 1478.5314 | 0.5064 | ##### | 4.1121 | 3.9208E-05 | 1.1241E-04 |
| ENSG00000201684\|RN7SKP239 | 12.5099 | -7.0565 | ##### | -4.1116 | 3.9286E-05 | 1.1262E-04 |
| ENSG00000131471\|AOC3 | 279.7302 | 0.6986 | ##### | 4.1108 | 3.9432E-05 | 1.1303E-04 |
| ENSG00000165138\|ANKS6 | 3827.4801 | -0.4864 | ##### | -4.1098 | 3.9603E-05 | 1.1351E-04 |
| ENSG00000170222\|ADPRM | 267.5938 | 0.6988 | ##### | 4.1094 | 3.9663E-05 | 1.1367E-04 |
| ENSG00000120068\|HOXB8 | 39.4176 | -4.2620 | ##### | -4.1079 | 3.9931E-05 | 1.1442E-04 |
| ENSG00000164188\|RANBP3L | 48.1685 | 7.3979 | ##### | 4.1078 | 3.9941E-05 | 1.1444E-04 |
| ENSG00000159110\|IFNAR2 | 4031.6234 | -0.4992 | ##### | -4.1076 | 3.9978E-05 | 1.1454E-04 |
| ENSG00000228836\|CT45A5 | 10.0261 | 5.9817 | ##### | 4.1073 | 4.0027E-05 | 1.1467E-04 |
| ENSG00000187796\|CARD9 | 145.5567 | 0.7842 | ##### | 4.1070 | 4.0076E-05 | 1.1480E-04 |
| ENSG00000066926\|FECH | 5207.8041 | -0.5002 | ##### | -4.1065 | 4.0163E-05 | 1.1504E-04 |
| ENSG00000021776\|AQR | 5644.1080 | 0.4965 | ##### | 4.1062 | 4.0224E-05 | 1.1518E-04 |
| ENSG00000087589\|CASS4 | 20.1140 | -3.1634 | ##### | -4.1062 | 4.0222E-05 | 1.1518E-04 |
| ENSG00000180233\|ZNRF2 | 1389.2997 | -0.5347 | ##### | -4.1062 | 4.0217E-05 | 1.1518E-04 |
| ENSG00000263155\|MYZAP | 89.1806 | -5.2232 | ##### | -4.1060 | 4.0253E-05 | 1.1525E-04 |
| ENSG00000249328 | 14.9514 | 6.6801 | ##### | 4.1059 | 4.0282E-05 | 1.1532E-04 |
| ENSG00000100626\|GALNT16 | 45.6876 | -1.1528 | ##### | -4.1058 | 4.0291E-05 | 1.1534E-04 |
| ENSG00000145936\|KCNMB1 | 26.1669 | -6.5844 | ##### | -4.1055 | 4.0339E-05 | 1.1546E-04 |
| ENSG00000204410\|MSH5 | 661.3413 | 0.6413 | ##### | 4.1055 | 4.0348E-05 | 1.1548E-04 |
| ENSG00000241973\|PI4KA | 5325.7744 | 0.4938 | ##### | 4.1054 | 4.0368E-05 | 1.1552E-04 |
| ENSG00000166189\|HPS6 | 2168.2627 | -0.5078 | ##### | -4.1053 | 4.0377E-05 | 1.1554E-04 |
| ENSG00000100722\|ZC3H14 | 5935.1090 | 0.4981 | ##### | 4.1051 | 4.0405E-05 | 1.1560E-04 |
| ENSG00000164291\|ARSK | 779.6633 | -0.5668 | ##### | -4.1052 | 4.0403E-05 | 1.1560E-04 |
| ENSG00000153048\|CARHSP1 | 6631.2863 | 0.4910 | ##### | 4.1049 | 4.0457E-05 | 1.1573E-04 |
| ENSG00000080644\|CHRNA3 | 95.4487 | 1.0531 | ##### | 4.1048 | 4.0472E-05 | 1.1577E-04 |
| ENSG00000226942\|IL9RP3 | 68.6474 | -8.4372 | ##### | -4.1045 | 4.0514E-05 | 1.1587E-04 |
| ENSG00000128253\|RFPL2 | 31.9953 | -3.2112 | ##### | -4.1042 | 4.0577E-05 | 1.1604E-04 |
| ENSG00000287061 | 24.5173 | 1.8253 | ##### | 4.1037 | 4.0667E-05 | 1.1629E-04 |
| ENSG00000187626\|ZKSCAN4 | 197.3067 | 0.6944 | ##### | 4.1031 | 4.0770E-05 | 1.1657E-04 |
| ENSG00000213920\|MDP1 | 319.5023 | 0.6958 | ##### | 4.1027 | 4.0832E-05 | 1.1674E-04 |
| ENSG00000276592\|PHB1P6 | 10.6022 | 2.3693 | ##### | 4.1025 | 4.0865E-05 | 1.1682E-04 |
| ENSG00000286875 | 15.5433 | 5.2298 | ##### | 4.1018 | 4.0990E-05 | 1.1717E-04 |
| ENSG00000129219\|PLD2 | 2206.3161 | 0.5241 | ##### | 4.1018 | 4.1003E-05 | 1.1719E-04 |
| ENSG00000160445\|ZER1 | 2573.2316 | -0.5179 | ##### | -4.1015 | 4.1041E-05 | 1.1729E-04 |
| ENSG00000290893 | 35.8826 | -7.5858 | ##### | -4.1014 | 4.1065E-05 | 1.1735E-04 |
| ENSG00000293167 | 9.9176 | 4.3350 | ##### | 4.1006 | 4.1214E-05 | 1.1776E-04 |
| ENSG00000288766 | 18.8709 | 1.9838 | ##### | 4.1003 | 4.1263E-05 | 1.1789E-04 |
| ENSG00000247373\|TMED2-DT | 625.5596 | 0.8809 | ##### | 4.1001 | 4.1296E-05 | 1.1798E-04 |
| ENSG00000228506\|PNISR-AS1 | 310.9130 | 0.6767 | ##### | 4.1000 | 4.1311E-05 | 1.1801E-04 |
| ENSG00000290230 | 39.4927 | -2.9300 | ##### | -4.0998 | 4.1356E-05 | 1.1812E-04 |
| ENSG00000288856 | 21.1138 | -1.8886 | ##### | -4.0994 | 4.1426E-05 | 1.1831E-04 |

| | | | | | | |
|---|---|---|---|---|---|---|
| ENSG00000243406\|MRPS31P5 | 57.3064 | 8.6941 | ##### | 4.0990 | 4.1486E-05 | 1.1847E-04 |
| ENSG00000238083\|LRRC37A2 | 1076.2359 | 0.5581 | ##### | 4.0989 | 4.1519E-05 | 1.1856E-04 |
| ENSG00000229337 | 37.9959 | 2.3417 | ##### | 4.0987 | 4.1550E-05 | 1.1863E-04 |
| ENSG00000148735\|PLEKHS1 | 12.4840 | -6.5668 | ##### | -4.0982 | 4.1630E-05 | 1.1885E-04 |
| ENSG00000213753\|CENPBD2P | 21.9456 | -2.1918 | ##### | -4.0971 | 4.1843E-05 | 1.1945E-04 |
| ENSG00000171786\|NHLH1 | 52.5458 | 1.3016 | ##### | 4.0970 | 4.1862E-05 | 1.1949E-04 |
| ENSG00000250091\|DNAH10OS | 313.6651 | 0.6969 | ##### | 4.0968 | 4.1887E-05 | 1.1955E-04 |
| ENSG00000280173 | 113.5770 | -0.9249 | ##### | -4.0968 | 4.1893E-05 | 1.1955E-04 |
| ENSG00000118961\|LDAH | 1561.2933 | -0.5478 | ##### | -4.0966 | 4.1924E-05 | 1.1963E-04 |
| ENSG00000172367\|NHERF4 | 68.4125 | -8.4301 | ##### | -4.0956 | 4.2109E-05 | 1.2015E-04 |
| ENSG00000293314\|SDAD1P1 | 123.0448 | 0.9505 | ##### | 4.0952 | 4.2180E-05 | 1.2034E-04 |
| ENSG00000171150\|SOCS5 | 4116.6853 | 0.4976 | ##### | 4.0951 | 4.2190E-05 | 1.2036E-04 |
| ENSG00000156925\|ZIC3 | 34.7822 | 4.3749 | ##### | 4.0947 | 4.2278E-05 | 1.2059E-04 |
| ENSG00000230186 | 13.9867 | -6.2210 | ##### | -4.0946 | 4.2296E-05 | 1.2063E-04 |
| ENSG00000214940\|NPIPA8 | 354.8345 | -1.0401 | ##### | -4.0943 | 4.2338E-05 | 1.2074E-04 |
| ENSG00000120896\|SORBS3 | 5369.6249 | -0.5167 | ##### | -4.0937 | 4.2461E-05 | 1.2108E-04 |
| ENSG00000286177 | 113.8313 | -0.9041 | ##### | -4.0935 | 4.2484E-05 | 1.2114E-04 |
| ENSG00000228135\|LINC01494 | 16.3322 | 5.7382 | ##### | 4.0934 | 4.2510E-05 | 1.2120E-04 |
| ENSG00000112306\|RPS12 | 38064.5302 | 0.4966 | ##### | 4.0930 | 4.2584E-05 | 1.2140E-04 |
| ENSG00000164008\|C1orf50 | 759.8014 | -0.5708 | ##### | -4.0925 | 4.2667E-05 | 1.2162E-04 |
| ENSG00000079950\|STX7 | 2618.6606 | -0.5062 | ##### | -4.0925 | 4.2669E-05 | 1.2162E-04 |
| ENSG00000184949\|FAM227A | 209.9769 | 0.7565 | ##### | 4.0920 | 4.2767E-05 | 1.2189E-04 |
| ENSG00000102974\|CTCF | 5805.5342 | 0.4285 | ##### | 4.0918 | 4.2803E-05 | 1.2198E-04 |
| ENSG00000228384 | 12.6630 | 1.9261 | ##### | 4.0914 | 4.2879E-05 | 1.2218E-04 |
| ENSG00000274386\|TMEM269 | 33.3840 | -1.2866 | ##### | -4.0908 | 4.2987E-05 | 1.2247E-04 |
| ENSG00000203739\|PRDX6-AS1 | 103.7869 | -0.9111 | ##### | -4.0908 | 4.2988E-05 | 1.2247E-04 |
| ENSG00000257923\|CUX1 | 8362.2743 | -0.5004 | ##### | -4.0901 | 4.3111E-05 | 1.2281E-04 |
| ENSG00000287851 | 23.4213 | 2.9388 | ##### | 4.0901 | 4.3119E-05 | 1.2282E-04 |
| ENSG00000176018\|LYSMD3 | 2290.4085 | 0.5208 | ##### | 4.0899 | 4.3148E-05 | 1.2289E-04 |
| ENSG00000101444\|AHCY | 17808.4642 | 0.5112 | ##### | 4.0873 | 4.3640E-05 | 1.2428E-04 |
| ENSG00000134333\|LDHA | ########## | 0.4190 | ##### | 4.0863 | 4.3829E-05 | 1.2480E-04 |
| ENSG00000293382\|GPR199P | 12.3169 | -3.7787 | ##### | -4.0862 | 4.3843E-05 | 1.2483E-04 |
| ENSG00000134874\|DZIP1 | 2688.9561 | 0.3850 | ##### | 4.0858 | 4.3932E-05 | 1.2508E-04 |
| ENSG00000154920\|EME1 | 1411.4992 | 0.5545 | ##### | 4.0844 | 4.4191E-05 | 1.2580E-04 |
| ENSG00000100304\|TTLL12 | 5813.4133 | 0.4902 | ##### | 4.0832 | 4.4412E-05 | 1.2642E-04 |
| ENSG00000066697\|MSANTD3 | 4312.7466 | 0.4992 | ##### | 4.0829 | 4.4482E-05 | 1.2660E-04 |
| ENSG00000002330\|BAD | 1900.6829 | -0.4224 | ##### | -4.0828 | 4.4500E-05 | 1.2664E-04 |
| ENSG00000125846\|ZNF133 | 621.8376 | 0.5482 | ##### | 4.0827 | 4.4525E-05 | 1.2670E-04 |
| ENSG00000231920\|NEBL-AS1 | 17.9943 | -2.7333 | ##### | -4.0823 | 4.4598E-05 | 1.2690E-04 |
| ENSG00000258702 | 24.8851 | 3.2607 | ##### | 4.0821 | 4.4627E-05 | 1.2697E-04 |
| ENSG00000106012\|IQCE | 5037.8629 | -0.4854 | ##### | -4.0816 | 4.4730E-05 | 1.2725E-04 |
| ENSG00000250995\|TMEM30BP1 | 25.7856 | 2.0617 | ##### | 4.0813 | 4.4791E-05 | 1.2741E-04 |
| ENSG00000166819\|PLIN1 | 27.0903 | -2.8188 | ##### | -4.0810 | 4.4848E-05 | 1.2756E-04 |
| ENSG00000282393 | 48.2555 | 1.6810 | ##### | 4.0806 | 4.4923E-05 | 1.2776E-04 |
| ENSG00000138231\|DBR1 | 2487.5887 | 0.5075 | ##### | 4.0802 | 4.5000E-05 | 1.2797E-04 |
| ENSG00000226237\|GAS1RR | 64.6604 | -8.3523 | ##### | -4.0801 | 4.5020E-05 | 1.2801E-04 |
| ENSG00000136147\|PHF11 | 534.1673 | -0.5763 | ##### | -4.0796 | 4.5107E-05 | 1.2825E-04 |
| ENSG00000274330 | 10.0427 | -3.5019 | ##### | -4.0796 | 4.5116E-05 | 1.2826E-04 |
| ENSG00000174206\|KICS2 | 1378.7583 | -0.5299 | ##### | -4.0795 | 4.5130E-05 | 1.2829E-04 |
| ENSG00000196199\|MPHOSPH8 | 3335.8941 | 0.5158 | ##### | 4.0793 | 4.5163E-05 | 1.2837E-04 |
| ENSG00000070785\|EIF2B3 | 2899.5487 | -0.5263 | ##### | -4.0787 | 4.5294E-05 | 1.2873E-04 |
| ENSG00000228570\|NUTM2E | 119.1341 | -0.8589 | ##### | -4.0769 | 4.5632E-05 | 1.2968E-04 |
| ENSG00000204632\|HLA-G | 50.2118 | 8.3392 | ##### | 4.0766 | 4.5691E-05 | 1.2984E-04 |
| ENSG00000171236\|LRG1 | 14.4776 | -4.5675 | ##### | -4.0760 | 4.5822E-05 | 1.3020E-04 |
| ENSG00000158156\|XKR8 | 750.4116 | -0.6064 | ##### | -4.0757 | 4.5866E-05 | 1.3031E-04 |
| ENSG00000172782\|FADS6 | 40.2774 | 7.1507 | ##### | 4.0754 | 4.5933E-05 | 1.3048E-04 |
| ENSG00000146676\|PURB | 9742.5097 | 0.4905 | ##### | 4.0748 | 4.6053E-05 | 1.3081E-04 |
| ENSG00000248109\|MARCOL | 34.1707 | -3.1075 | ##### | -4.0748 | 4.6063E-05 | 1.3083E-04 |
| ENSG00000237491\|LINC01409 | 77.3947 | 1.2163 | ##### | 4.0745 | 4.6119E-05 | 1.3098E-04 |
| ENSG00000278505\|C17orf78 | 11.9851 | -5.8449 | ##### | -4.0744 | 4.6132E-05 | 1.3100E-04 |
| ENSG00000197959\|DNM3 | 102.5032 | -0.9944 | ##### | -4.0737 | 4.6274E-05 | 1.3138E-04 |
| ENSG00000129480\|DTD2 | 2418.3603 | -0.5313 | ##### | -4.0737 | 4.6273E-05 | 1.3138E-04 |
| ENSG00000105373\|NOP53 | 8532.3578 | -0.4826 | ##### | -4.0734 | 4.6329E-05 | 1.3152E-04 |

| | | | | | | |
|---|---|---|---|---|---|---|
| ENSG00000212994\|RPS26P6 | 176.3618 | 0.7345 | ##### | 4.0729 | 4.6440E-05 | 1.3183E-04 |
| ENSG00000213995\|NAXD | 2211.6363 | -0.5078 | ##### | -4.0728 | 4.6458E-05 | 1.3186E-04 |
| ENSG00000197670 | 29.1584 | 1.4136 | ##### | 4.0717 | 4.6662E-05 | 1.3243E-04 |
| ENSG00000152439\|ZNF773 | 570.7096 | -0.5943 | ##### | -4.0717 | 4.6679E-05 | 1.3247E-04 |
| ENSG00000140104\|CLBA1 | 1022.9085 | 0.5760 | ##### | 4.0715 | 4.6714E-05 | 1.3255E-04 |
| ENSG00000082269\|FAM135A | 4054.1782 | 0.4894 | ##### | 4.0713 | 4.6757E-05 | 1.3266E-04 |
| ENSG00000176903\|PNMA1 | 7039.3805 | 0.4981 | ##### | 4.0710 | 4.6812E-05 | 1.3280E-04 |
| ENSG00000022976\|ZNF839 | 931.7924 | -0.5550 | ##### | -4.0703 | 4.6958E-05 | 1.3320E-04 |
| ENSG00000062716\|VMP1 | 15399.0779 | 0.4561 | ##### | 4.0703 | 4.6960E-05 | 1.3320E-04 |
| ENSG00000155957\|TMBIM4 | 3320.3025 | -0.5178 | ##### | -4.0702 | 4.6978E-05 | 1.3324E-04 |
| ENSG00000135763\|URB2 | 3643.7756 | 0.4278 | ##### | 4.0696 | 4.7087E-05 | 1.3354E-04 |
| ENSG00000036054\|TBC1D23 | 4257.4219 | 0.4830 | ##### | 4.0694 | 4.7126E-05 | 1.3363E-04 |
| ENSG00000164916\|FOXK1 | 9552.2033 | 0.4818 | ##### | 4.0694 | 4.7130E-05 | 1.3363E-04 |
| ENSG00000215492\|HNRNPA1P7 | 1582.8481 | 0.5291 | ##### | 4.0668 | 4.7660E-05 | 1.3512E-04 |
| ENSG00000177613\|CSTF2T | 3769.6508 | 0.5005 | ##### | 4.0667 | 4.7690E-05 | 1.3519E-04 |
| ENSG00000182575\|NXPH3 | 49.9440 | -1.2193 | ##### | -4.0666 | 4.7697E-05 | 1.3520E-04 |
| ENSG00000113555\|PCDH12 | 70.9455 | -1.0392 | ##### | -4.0661 | 4.7809E-05 | 1.3551E-04 |
| ENSG00000264668 | 446.8996 | ####### | ##### | -4.0657 | 4.7894E-05 | 1.3573E-04 |
| ENSG00000184995\|IFNE | 16.7805 | 3.5750 | ##### | 4.0654 | 4.7941E-05 | 1.3585E-04 |
| ENSG00000168970\|JMJD7-PLA2G4B | 719.8469 | 0.6092 | ##### | 4.0651 | 4.8002E-05 | 1.3601E-04 |
| ENSG00000246203 | 216.7631 | 0.7540 | ##### | 4.0651 | 4.8020E-05 | 1.3605E-04 |
| ENSG00000183579\|ZNRF3 | 2226.6143 | -0.5184 | ##### | -4.0649 | 4.8051E-05 | 1.3613E-04 |
| ENSG00000158186\|MRAS | 2563.2004 | -0.4980 | ##### | -4.0644 | 4.8147E-05 | 1.3638E-04 |
| ENSG00000264538\|SUZ12P1 | 172.2235 | 1.0094 | ##### | 4.0644 | 4.8165E-05 | 1.3642E-04 |
| ENSG00000237352\|LINC01358 | 13.3859 | -6.1577 | ##### | -4.0642 | 4.8201E-05 | 1.3651E-04 |
| ENSG00000132406\|TMEM128 | 843.8086 | -0.5994 | ##### | -4.0636 | 4.8325E-05 | 1.3685E-04 |
| ENSG00000173914\|RBM4B | 1114.2820 | 0.5542 | ##### | 4.0633 | 4.8379E-05 | 1.3699E-04 |
| ENSG00000161999\|JMJD8 | 6531.7429 | -0.5128 | ##### | -4.0614 | 4.8779E-05 | 1.3810E-04 |
| ENSG00000281903\|ASMER1 | 14.2094 | -3.6632 | ##### | -4.0614 | 4.8780E-05 | 1.3810E-04 |
| ENSG00000221843\|C2orf16 | 123.9599 | 1.0174 | ##### | 4.0613 | 4.8792E-05 | 1.3812E-04 |
| ENSG00000293198 | 8.5783 | 5.3556 | ##### | 4.0590 | 4.9281E-05 | 1.3949E-04 |
| ENSG00000011465\|DCN | 16.2720 | 6.5607 | ##### | 4.0588 | 4.9335E-05 | 1.3963E-04 |
| ENSG00000180525\|DIP2C-AS1 | 45.3147 | -8.4357 | ##### | -4.0582 | 4.9463E-05 | 1.3998E-04 |
| ENSG00000119041\|GTF3C3 | 6145.5277 | -0.4805 | ##### | -4.0574 | 4.9623E-05 | 1.4042E-04 |
| ENSG00000204839\|MROH6 | 407.1964 | -0.6441 | ##### | -4.0569 | 4.9736E-05 | 1.4073E-04 |
| ENSG00000233611\|GBX2-AS1 | 311.4669 | 0.6833 | ##### | 4.0566 | 4.9789E-05 | 1.4086E-04 |
| ENSG00000197614\|MFAP5 | 62.6553 | -1.2839 | ##### | -4.0563 | 4.9859E-05 | 1.4105E-04 |
| ENSG00000144857\|BOC | 71.8640 | 1.1057 | ##### | 4.0559 | 4.9947E-05 | 1.4128E-04 |
| ENSG00000123505\|AMD1 | 12420.3847 | 0.4283 | ##### | 4.0548 | 5.0170E-05 | 1.4190E-04 |
| ENSG00000289505 | 20.7222 | 6.0191 | ##### | 4.0545 | 5.0242E-05 | 1.4209E-04 |
| ENSG00000286237\|ARMCX5-GPRASP2 | 217.1112 | 1.7381 | ##### | 4.0541 | 5.0334E-05 | 1.4234E-04 |
| ENSG00000144579\|CTDSP1 | 7149.7659 | 0.5074 | ##### | 4.0536 | 5.0435E-05 | 1.4261E-04 |
| ENSG00000198783\|ZNF830 | 1350.6919 | 0.5357 | ##### | 4.0531 | 5.0545E-05 | 1.4291E-04 |
| ENSG00000187908\|DMBT1 | 41.7848 | 7.6506 | ##### | 4.0520 | 5.0787E-05 | 1.4358E-04 |
| ENSG00000189114\|BLOC1S3 | 693.4666 | -0.5659 | ##### | -4.0509 | 5.1022E-05 | 1.4423E-04 |
| ENSG00000232732 | 23.1774 | 3.0538 | ##### | 4.0504 | 5.1139E-05 | 1.4455E-04 |
| ENSG00000107679\|PLEKHA1 | 3398.3153 | -0.5170 | ##### | -4.0501 | 5.1185E-05 | 1.4466E-04 |
| ENSG00000289007 | 24.7371 | -2.4546 | ##### | -4.0495 | 5.1333E-05 | 1.4507E-04 |
| ENSG00000245112\|SMARCA5-AS1 | 212.9562 | -0.8246 | ##### | -4.0494 | 5.1341E-05 | 1.4508E-04 |
| ENSG00000143322\|ABL2 | 5637.4810 | 0.5016 | ##### | 4.0488 | 5.1481E-05 | 1.4546E-04 |
| ENSG00000132485\|ZRANB2 | 7873.1874 | 0.2688 | ##### | 4.0479 | 5.1689E-05 | 1.4603E-04 |
| ENSG00000133193\|FAM104A | 4465.6946 | 0.4968 | ##### | 4.0478 | 5.1711E-05 | 1.4606E-04 |
| ENSG00000279453 | 96.6975 | 0.8841 | ##### | 4.0478 | 5.1706E-05 | 1.4606E-04 |
| ENSG00000103932\|RPAP1 | 4744.4139 | 0.5021 | ##### | 4.0476 | 5.1738E-05 | 1.4613E-04 |
| ENSG00000269335\|IKBKG | 2521.9934 | 0.5137 | ##### | 4.0468 | 5.1926E-05 | 1.4665E-04 |
| ENSG00000101331\|CCM2L | 179.0537 | 0.8010 | ##### | 4.0465 | 5.1985E-05 | 1.4680E-04 |
| ENSG00000167566\|NCKAP5L | 2745.0426 | 0.4274 | ##### | 4.0461 | 5.2083E-05 | 1.4706E-04 |
| ENSG00000158865\|SLC5A11 | 28.8333 | -2.2705 | ##### | -4.0458 | 5.2155E-05 | 1.4725E-04 |
| ENSG00000173269\|MMRN2 | 70.3428 | -3.4985 | ##### | -4.0443 | 5.2488E-05 | 1.4818E-04 |
| ENSG00000219545\|UMAD1 | 1800.4466 | -0.5251 | ##### | -4.0427 | 5.2832E-05 | 1.4913E-04 |
| ENSG00000121653\|MAPK8IP1 | 1231.8415 | 0.5248 | ##### | 4.0425 | 5.2889E-05 | 1.4928E-04 |
| ENSG00000274993 | 76.0155 | -1.5482 | ##### | -4.0424 | 5.2902E-05 | 1.4930E-04 |
| ENSG00000223466\|LINC01825 | 16.2084 | -7.2176 | ##### | -4.0419 | 5.3021E-05 | 1.4962E-04 |

| | | | | | | |
|---|---|---|---|---|---|---|
| ENSG00000117877\|POLR1G | 6220.5212 | 0.5240 | ##### | 4.0405 | 5.3344E-05 | 1.5052E-04 |
| ENSG00000187764\|SEMA4D | 835.4015 | 0.5543 | ##### | 4.0377 | 5.3982E-05 | 1.5231E-04 |
| ENSG00000279009 | 15.5831 | -1.6834 | ##### | -4.0374 | 5.4041E-05 | 1.5246E-04 |
| ENSG00000188316\|ENO4 | 61.8022 | 1.1769 | ##### | 4.0370 | 5.4138E-05 | 1.5271E-04 |
| ENSG00000102547\|CAB39L | 986.2406 | 0.5448 | ##### | 4.0370 | 5.4139E-05 | 1.5271E-04 |
| ENSG00000289985 | 19.6925 | 5.6140 | ##### | 4.0369 | 5.4164E-05 | 1.5276E-04 |
| ENSG00000171989\|LDHAL6B | 14.2988 | -4.5224 | ##### | -4.0365 | 5.4261E-05 | 1.5302E-04 |
| ENSG00000167272\|POP5 | 2387.4084 | 0.5300 | ##### | 4.0360 | 5.4377E-05 | 1.5334E-04 |
| ENSG00000182117\|NOP10 | 4634.9447 | 0.4856 | ##### | 4.0359 | 5.4394E-05 | 1.5337E-04 |
| ENSG00000205670\|SMIM11 | 564.0070 | -0.6132 | ##### | -4.0355 | 5.4496E-05 | 1.5364E-04 |
| ENSG00000225108\|ZBTB45P1 | 583.0217 | -0.6001 | ##### | -4.0353 | 5.4541E-05 | 1.5375E-04 |
| ENSG00000275426 | 12.1646 | -6.4077 | ##### | -4.0342 | 5.4780E-05 | 1.5441E-04 |
| ENSG00000075340\|ADD2 | 1300.7412 | 0.5547 | ##### | 4.0336 | 5.4918E-05 | 1.5479E-04 |
| ENSG00000166803\|PCLAF | 2520.7826 | 0.5270 | ##### | 4.0333 | 5.5011E-05 | 1.5503E-04 |
| ENSG00000273305 | 11.6399 | 3.6385 | ##### | 4.0330 | 5.5078E-05 | 1.5521E-04 |
| ENSG00000221883\|ARIH2OS | 48.0750 | -1.1970 | ##### | -4.0322 | 5.5246E-05 | 1.5567E-04 |
| ENSG00000169246\|NPIPB3 | 1082.1923 | 0.5350 | ##### | 4.0318 | 5.5352E-05 | 1.5595E-04 |
| ENSG00000164134\|NAA15 | 9082.6626 | -0.4809 | ##### | -4.0310 | 5.5531E-05 | 1.5644E-04 |
| ENSG00000241370\|RPP21 | 2133.3646 | 0.5100 | ##### | 4.0307 | 5.5615E-05 | 1.5666E-04 |
| ENSG00000274615\|NPEPPSP1 | 409.6885 | 0.6720 | ##### | 4.0302 | 5.5738E-05 | 1.5700E-04 |
| ENSG00000134830\|C5AR2 | 117.8435 | -6.8831 | ##### | -4.0296 | 5.5863E-05 | 1.5733E-04 |
| ENSG00000268230 | 134.1301 | -0.9233 | ##### | -4.0289 | 5.6036E-05 | 1.5780E-04 |
| ENSG00000185800\|DMWD | 5086.0487 | -0.4940 | ##### | -4.0286 | 5.6107E-05 | 1.5799E-04 |
| ENSG00000258592\|LINC03059 | 50.7503 | 1.2388 | ##### | 4.0281 | 5.6229E-05 | 1.5832E-04 |
| ENSG00000113369\|ARRDC3 | 2939.3121 | 0.4949 | ##### | 4.0279 | 5.6286E-05 | 1.5846E-04 |
| ENSG00000260597 | 47.5700 | 1.3998 | ##### | 4.0276 | 5.6348E-05 | 1.5862E-04 |
| ENSG00000257267\|ZNF271P | 71.7330 | -1.8320 | ##### | -4.0265 | 5.6606E-05 | 1.5934E-04 |
| ENSG00000134058\|CDK7 | 2395.0736 | 0.4681 | ##### | 4.0257 | 5.6815E-05 | 1.5991E-04 |
| ENSG00000248636 | 88.2571 | -0.7507 | ##### | -4.0245 | 5.7091E-05 | 1.6067E-04 |
| ENSG00000284672 | 94.6280 | 8.7436 | ##### | 4.0241 | 5.7183E-05 | 1.6091E-04 |
| ENSG00000256995\|LINC02955 | 35.0639 | 7.4558 | ##### | 4.0237 | 5.7290E-05 | 1.6120E-04 |
| ENSG00000290445\|CCNYL2 | 21.5598 | -4.9914 | ##### | -4.0228 | 5.7519E-05 | 1.6183E-04 |
| ENSG00000253405\|EVX1-AS | 46.2271 | -8.4642 | ##### | -4.0226 | 5.7557E-05 | 1.6192E-04 |
| ENSG00000103067\|ESRP2 | 307.2811 | -0.6388 | ##### | -4.0225 | 5.7586E-05 | 1.6199E-04 |
| ENSG00000088367\|EPB41L1 | 4708.1145 | -0.4060 | ##### | -4.0216 | 5.7813E-05 | 1.6261E-04 |
| ENSG00000141867\|BRD4 | 8428.6871 | 0.4671 | ##### | 4.0210 | 5.7953E-05 | 1.6299E-04 |
| ENSG00000159352\|PSMD4 | 21342.9588 | 0.4845 | ##### | 4.0210 | 5.7959E-05 | 1.6299E-04 |
| ENSG00000280193 | 51.7520 | -1.3636 | ##### | -4.0207 | 5.8017E-05 | 1.6314E-04 |
| ENSG00000257337\|TNS2-AS1 | 535.8705 | -0.6766 | ##### | -4.0201 | 5.8165E-05 | 1.6354E-04 |
| ENSG00000155115\|GTF3C6 | 3669.9000 | -0.5003 | ##### | -4.0180 | 5.8707E-05 | 1.6505E-04 |
| ENSG00000263968\|RN7SL381P | 12.2492 | 2.8787 | ##### | 4.0176 | 5.8803E-05 | 1.6530E-04 |
| ENSG00000138495\|COX17 | 3304.7143 | 0.5153 | ##### | 4.0160 | 5.9193E-05 | 1.6638E-04 |
| ENSG00000257354\|MIRLET7IHG | 292.7888 | 0.6461 | ##### | 4.0158 | 5.9242E-05 | 1.6650E-04 |
| ENSG00000239407 | 140.8128 | 0.7391 | ##### | 4.0152 | 5.9387E-05 | 1.6689E-04 |
| ENSG00000212232\|SNORD17 | 43.7929 | 1.1714 | ##### | 4.0147 | 5.9519E-05 | 1.6725E-04 |
| ENSG00000233155\|HMGA1P8 | 47.9121 | 1.2274 | ##### | 4.0143 | 5.9618E-05 | 1.6751E-04 |
| ENSG00000160716\|CHRNB2 | 39.9603 | 1.8407 | ##### | 4.0139 | 5.9722E-05 | 1.6779E-04 |
| ENSG00000264456 | 55.0685 | -1.1359 | ##### | -4.0127 | 6.0017E-05 | 1.6860E-04 |
| ENSG00000118454\|ANKRD13C | 2898.5247 | -0.5141 | ##### | -4.0127 | 6.0040E-05 | 1.6865E-04 |
| ENSG00000241544\|LINC02029 | 25.1675 | 4.3558 | ##### | 4.0124 | 6.0099E-05 | 1.6880E-04 |
| ENSG00000171960\|PPIH | 3594.9752 | 0.5056 | ##### | 4.0115 | 6.0336E-05 | 1.6945E-04 |
| ENSG00000267757\|EML2-AS1 | 205.5210 | 1.3907 | ##### | 4.0114 | 6.0364E-05 | 1.6951E-04 |
| ENSG00000271287\|BCRP9 | 11.6756 | -5.0813 | ##### | -4.0110 | 6.0455E-05 | 1.6975E-04 |
| ENSG00000054267\|ARID4B | 2749.5852 | 0.5199 | ##### | 4.0103 | 6.0633E-05 | 1.7024E-04 |
| ENSG00000088726\|TMEM40 | 207.5799 | 2.9812 | ##### | 4.0099 | 6.0748E-05 | 1.7054E-04 |
| ENSG00000179772\|FOXS1 | 46.3476 | -7.8946 | ##### | -4.0098 | 6.0762E-05 | 1.7057E-04 |
| ENSG00000134363\|FST | 116.7993 | -5.1146 | ##### | -4.0097 | 6.0790E-05 | 1.7061E-04 |
| ENSG00000128714\|HOXD13 | 68.1199 | -6.7097 | ##### | -4.0097 | 6.0788E-05 | 1.7061E-04 |
| ENSG00000228203\|GRASLND | 53.3119 | 2.4947 | ##### | 4.0096 | 6.0809E-05 | 1.7065E-04 |
| ENSG00000241990\|PRR34-AS1 | 155.6575 | -0.8789 | ##### | -4.0092 | 6.0920E-05 | 1.7095E-04 |
| ENSG00000109083\|IFT20 | 1591.8969 | -0.5457 | ##### | -4.0078 | 6.1277E-05 | 1.7192E-04 |
| ENSG00000112339\|HBS1L | 5501.8202 | 0.3878 | ##### | 4.0078 | 6.1278E-05 | 1.7192E-04 |
| ENSG00000175745\|NR2F1 | 1818.7906 | 0.5236 | ##### | 4.0078 | 6.1297E-05 | 1.7196E-04 |

| | | | | | | |
|---|---|---|---|---|---|---|
| ENSG00000259426\|KIF23-AS1 | 214.9916 | 0.7498 | ##### | 4.0075 | 6.1358E-05 | 1.7211E-04 |
| ENSG00000214960\|CRPPA | 136.1278 | -0.8810 | ##### | -4.0074 | 6.1387E-05 | 1.7217E-04 |
| ENSG00000283041 | 11259.2246 | 0.4703 | ##### | 4.0071 | 6.1465E-05 | 1.7238E-04 |
| ENSG00000289486 | 106.3448 | -1.0351 | ##### | -4.0070 | 6.1489E-05 | 1.7243E-04 |
| ENSG00000290124 | 33.0646 | -1.3893 | ##### | -4.0065 | 6.1633E-05 | 1.7282E-04 |
| ENSG00000089327\|FXYD5 | 4393.7690 | -0.5170 | ##### | -4.0054 | 6.1917E-05 | 1.7360E-04 |
| ENSG00000165672\|PRDX3 | 17072.6091 | -0.4368 | ##### | -4.0052 | 6.1978E-05 | 1.7375E-04 |
| ENSG00000236123\|CEACAMP11 | 14.4822 | 5.8997 | ##### | 4.0048 | 6.2077E-05 | 1.7401E-04 |
| ENSG00000102984\|ZNF821 | 361.5734 | -0.6493 | ##### | -4.0045 | 6.2154E-05 | 1.7421E-04 |
| ENSG00000173960\|UBXN2A | 2987.3493 | -0.5127 | ##### | -4.0039 | 6.2312E-05 | 1.7464E-04 |
| ENSG00000067182\|TNFRSF1A | 18316.1811 | -0.4777 | ##### | -4.0035 | 6.2422E-05 | 1.7493E-04 |
| ENSG00000196743\|GM2A | 5350.0717 | 0.4850 | ##### | 4.0026 | 6.2650E-05 | 1.7555E-04 |
| ENSG00000182551\|ADI1 | 6817.8035 | 0.5064 | ##### | 4.0022 | 6.2760E-05 | 1.7585E-04 |
| ENSG00000231927 | 16.7016 | -6.9619 | ##### | -4.0017 | 6.2886E-05 | 1.7618E-04 |
| ENSG00000287884 | 11.1333 | 3.7039 | ##### | 4.0014 | 6.2967E-05 | 1.7639E-04 |
| ENSG00000228302 | 36.1879 | 1.4717 | ##### | 4.0012 | 6.3024E-05 | 1.7654E-04 |
| ENSG00000226342\|NMD3P1 | 9.0181 | -5.1187 | ##### | -4.0011 | 6.3048E-05 | 1.7658E-04 |
| ENSG00000258745 | 13.9998 | 6.1861 | ##### | 4.0011 | 6.3054E-05 | 1.7658E-04 |
| ENSG00000148334\|PTGES2 | 5684.6306 | 0.4519 | ##### | 4.0010 | 6.3085E-05 | 1.7666E-04 |
| ENSG00000176678\|FOXL1 | 20.2485 | -2.6546 | ##### | -4.0002 | 6.3280E-05 | 1.7718E-04 |
| ENSG00000267078 | 46.3678 | 7.7559 | ##### | 3.9999 | 6.3367E-05 | 1.7741E-04 |
| ENSG00000082515\|MRPL22 | 2912.9984 | -0.5082 | ##### | -3.9996 | 6.3450E-05 | 1.7763E-04 |
| ENSG00000108518\|PFN1 | ########## | 0.4783 | ##### | 3.9990 | 6.3606E-05 | 1.7805E-04 |
| ENSG00000172172\|MRPL13 | 3701.5389 | -0.5007 | ##### | -3.9989 | 6.3626E-05 | 1.7809E-04 |
| ENSG00000260329\|TMEM263-DT | 148.6696 | -0.8980 | ##### | -3.9985 | 6.3736E-05 | 1.7838E-04 |
| ENSG00000168350\|DEGS2 | 124.1818 | -1.4246 | ##### | -3.9980 | 6.3866E-05 | 1.7873E-04 |
| ENSG00000274220 | 61.7052 | -1.7768 | ##### | -3.9975 | 6.4028E-05 | 1.7916E-04 |
| ENSG00000257379 | 2475.7881 | 1.0350 | ##### | 3.9972 | 6.4101E-05 | 1.7935E-04 |
| ENSG00000260465 | 1046.9442 | -0.5483 | ##### | -3.9965 | 6.4298E-05 | 1.7988E-04 |
| ENSG00000205879\|FAM90A2P | 16.4510 | -1.8613 | ##### | -3.9964 | 6.4314E-05 | 1.7991E-04 |
| ENSG00000250722\|SELENOP | 116.4859 | -1.1192 | ##### | -3.9952 | 6.4651E-05 | 1.8084E-04 |
| ENSG00000058056\|USP13 | 2887.6172 | -0.5122 | ##### | -3.9951 | 6.4668E-05 | 1.8087E-04 |
| ENSG00000257252 | 20.4683 | -2.9078 | ##### | -3.9951 | 6.4676E-05 | 1.8087E-04 |
| ENSG00000103591\|AAGAB | 5319.2409 | 0.4739 | ##### | 3.9945 | 6.4837E-05 | 1.8131E-04 |
| ENSG00000197989\|SNHG12 | 1266.9757 | 0.5407 | ##### | 3.9942 | 6.4925E-05 | 1.8154E-04 |
| ENSG00000288187 | 10.5912 | -2.1371 | ##### | -3.9941 | 6.4931E-05 | 1.8154E-04 |
| ENSG00000224635 | 80.8142 | 0.9526 | ##### | 3.9935 | 6.5108E-05 | 1.8201E-04 |
| ENSG00000178127\|NDUFV2 | 8115.5762 | 0.4834 | ##### | 3.9934 | 6.5125E-05 | 1.8204E-04 |
| ENSG00000105404\|RABAC1 | 5561.9486 | -0.4917 | ##### | -3.9933 | 6.5151E-05 | 1.8210E-04 |
| ENSG00000273355 | 77.9852 | 1.0017 | ##### | 3.9927 | 6.5318E-05 | 1.8255E-04 |
| ENSG00000288066 | 440.4219 | -0.5930 | ##### | -3.9923 | 6.5438E-05 | 1.8287E-04 |
| ENSG00000167107\|ACSF2 | 900.2813 | -0.5795 | ##### | -3.9918 | 6.5563E-05 | 1.8320E-04 |
| ENSG00000123572\|NRK | 15.9003 | -4.3260 | ##### | -3.9912 | 6.5730E-05 | 1.8365E-04 |
| ENSG00000148541\|FAM13C | 18.5361 | 2.6326 | ##### | 3.9906 | 6.5910E-05 | 1.8414E-04 |
| ENSG00000197976\|AKAP17A | 3176.6819 | 0.4539 | ##### | 3.9905 | 6.5921E-05 | 1.8415E-04 |
| ENSG00000233589 | 53.6441 | 1.0965 | ##### | 3.9895 | 6.6200E-05 | 1.8491E-04 |
| ENSG00000279057 | 25.9562 | 2.2878 | ##### | 3.9885 | 6.6503E-05 | 1.8574E-04 |
| ENSG00000104369\|JPH1 | 72.4385 | -1.3231 | ##### | -3.9883 | 6.6559E-05 | 1.8588E-04 |
| ENSG00000227908\|IL6ST-DT | 54.8761 | 1.5065 | ##### | 3.9877 | 6.6710E-05 | 1.8626E-04 |
| ENSG00000134480\|CCNH | 3019.7365 | 0.4810 | ##### | 3.9877 | 6.6706E-05 | 1.8626E-04 |
| ENSG00000198734\|F5 | 90.2209 | -4.9080 | ##### | -3.9876 | 6.6758E-05 | 1.8636E-04 |
| ENSG00000262648\|PHB1P15 | 8.8087 | 2.8228 | ##### | 3.9876 | 6.6753E-05 | 1.8636E-04 |
| ENSG00000234928\|LINC01659 | 23.2402 | 1.9226 | ##### | 3.9870 | 6.6927E-05 | 1.8682E-04 |
| ENSG00000101004\|NINL | 1290.5717 | 0.5038 | ##### | 3.9866 | 6.7030E-05 | 1.8709E-04 |
| ENSG00000287808\|DHX9-AS1 | 56.6537 | 1.0406 | ##### | 3.9857 | 6.7281E-05 | 1.8777E-04 |
| ENSG00000105771\|SMG9 | 3010.3277 | -0.5083 | ##### | -3.9851 | 6.7451E-05 | 1.8823E-04 |
| ENSG00000248724\|NPHP3-AS1 | 24.5947 | 1.8179 | ##### | 3.9845 | 6.7625E-05 | 1.8870E-04 |
| ENSG00000169371\|SNUPN | 2108.0705 | 0.4860 | ##### | 3.9843 | 6.7680E-05 | 1.8883E-04 |
| ENSG00000186501\|TMEM222 | 2550.2232 | 0.4800 | ##### | 3.9840 | 6.7765E-05 | 1.8903E-04 |
| ENSG00000148798\|INA | 10239.9876 | -0.4322 | ##### | -3.9840 | 6.7764E-05 | 1.8903E-04 |
| ENSG00000176058\|TPRN | 900.7638 | 0.5437 | ##### | 3.9840 | 6.7774E-05 | 1.8904E-04 |
| ENSG00000211460\|TSN | 11646.5727 | -0.4904 | ##### | -3.9835 | 6.7914E-05 | 1.8942E-04 |
| ENSG00000198795\|ZNF521 | 11.1226 | 6.2376 | ##### | 3.9832 | 6.7996E-05 | 1.8962E-04 |

| | | | | | | |
|---|---|---|---|---|---|---|
| ENSG00000169627\|BOLA2B | 2492.3161 | 0.4962 | ##### | 3.9831 | 6.8016E-05 | 1.8966E-04 |
| ENSG00000278784 | 90.5175 | 0.9571 | ##### | 3.9820 | 6.8337E-05 | 1.9054E-04 |
| ENSG00000123999\|INHA | 50.9977 | -1.8016 | ##### | -3.9819 | 6.8371E-05 | 1.9062E-04 |
| ENSG00000285480\|H2BK1 | 11.9850 | 3.0320 | ##### | 3.9818 | 6.8398E-05 | 1.9068E-04 |
| ENSG00000279786 | 19.5597 | 6.8480 | ##### | 3.9816 | 6.8444E-05 | 1.9079E-04 |
| ENSG00000100263\|RHBDD3 | 2077.5356 | 0.5164 | ##### | 3.9808 | 6.8680E-05 | 1.9143E-04 |
| ENSG00000171861\|MRM3 | 2020.4467 | 0.5361 | ##### | 3.9808 | 6.8690E-05 | 1.9144E-04 |
| ENSG00000023697\|DERA | 3499.4915 | -0.4902 | ##### | -3.9807 | 6.8724E-05 | 1.9151E-04 |
| ENSG00000110107\|PRPF19 | 12258.5217 | 0.4149 | ##### | 3.9802 | 6.8848E-05 | 1.9184E-04 |
| ENSG00000250938\|MAD2L1-DT | 36.5694 | 1.5919 | ##### | 3.9802 | 6.8861E-05 | 1.9186E-04 |
| ENSG00000228050\|TOP3BP1 | 69.2611 | 1.0515 | ##### | 3.9799 | 6.8944E-05 | 1.9207E-04 |
| ENSG00000286722 | 46.3076 | -8.4230 | ##### | -3.9798 | 6.8982E-05 | 1.9216E-04 |
| ENSG00000187840\|EIF4EBP1 | 3733.3762 | 0.5011 | ##### | 3.9797 | 6.8999E-05 | 1.9219E-04 |
| ENSG00000288940 | 25.7581 | 2.5452 | ##### | 3.9787 | 6.9292E-05 | 1.9299E-04 |
| ENSG00000178081\|ULK4P3 | 12.0179 | -5.8448 | ##### | -3.9783 | 6.9397E-05 | 1.9326E-04 |
| ENSG00000119787\|ATL2 | 7090.1991 | 0.4769 | ##### | 3.9773 | 6.9690E-05 | 1.9406E-04 |
| ENSG00000123600\|METTL8 | 2608.8515 | 0.4978 | ##### | 3.9771 | 6.9747E-05 | 1.9420E-04 |
| ENSG00000099904\|ZDHHC8 | 3168.0960 | 0.5006 | ##### | 3.9770 | 6.9791E-05 | 1.9431E-04 |
| ENSG00000235244\|DANT2 | 19.6365 | 3.1914 | ##### | 3.9769 | 6.9832E-05 | 1.9440E-04 |
| ENSG00000283538 | 10.3359 | 3.3998 | ##### | 3.9761 | 7.0046E-05 | 1.9498E-04 |
| ENSG00000260942\|CAPN10-DT | 222.2868 | 0.7310 | ##### | 3.9760 | 7.0091E-05 | 1.9509E-04 |
| ENSG00000173214\|MFSD4B | 939.8528 | 0.5083 | ##### | 3.9757 | 7.0162E-05 | 1.9526E-04 |
| ENSG00000156049\|GNA14 | 45.3296 | -8.3935 | ##### | -3.9756 | 7.0209E-05 | 1.9538E-04 |
| ENSG00000214941\|ZSWIM7 | 822.8582 | -0.5427 | ##### | -3.9754 | 7.0252E-05 | 1.9548E-04 |
| ENSG00000140320\|BAHD1 | 3405.8116 | 0.4741 | ##### | 3.9752 | 7.0314E-05 | 1.9561E-04 |
| ENSG00000144057\|ST6GAL2 | 32.0590 | -4.6366 | ##### | -3.9752 | 7.0310E-05 | 1.9561E-04 |
| ENSG00000132434\|LANCL2 | 2670.8733 | -0.5039 | ##### | -3.9751 | 7.0349E-05 | 1.9570E-04 |
| ENSG00000286433 | 12.5973 | 2.0406 | ##### | 3.9750 | 7.0384E-05 | 1.9577E-04 |
| ENSG00000007545\|CRAMP1 | 2583.3363 | 0.5273 | ##### | 3.9743 | 7.0572E-05 | 1.9628E-04 |
| ENSG00000141298\|SSH2 | 5464.7786 | 0.4812 | ##### | 3.9742 | 7.0603E-05 | 1.9635E-04 |
| ENSG00000137193\|PIM1 | 3204.0645 | 0.4878 | ##### | 3.9742 | 7.0621E-05 | 1.9638E-04 |
| ENSG00000214429\|CYCSP6 | 72.0240 | -8.5912 | ##### | -3.9736 | 7.0808E-05 | 1.9688E-04 |
| ENSG00000291018 | 28.2852 | -1.4448 | ##### | -3.9735 | 7.0837E-05 | 1.9694E-04 |
| ENSG00000172794\|RAB37 | 58.9820 | 2.1198 | ##### | 3.9720 | 7.1257E-05 | 1.9809E-04 |
| ENSG00000163221\|S100A12 | 33.7934 | -8.0110 | ##### | -3.9711 | 7.1532E-05 | 1.9884E-04 |
| ENSG00000002745\|WNT16 | 21.8870 | 3.1022 | ##### | 3.9706 | 7.1700E-05 | 1.9929E-04 |
| ENSG00000282870\|FRG1DP | 44.8579 | 1.2127 | ##### | 3.9704 | 7.1766E-05 | 1.9945E-04 |
| ENSG00000231503\|PTMAP4 | 79.2653 | 0.8448 | ##### | 3.9702 | 7.1815E-05 | 1.9957E-04 |
| ENSG00000143226\|FCGR2A | 36.9632 | -7.9213 | ##### | -3.9701 | 7.1843E-05 | 1.9963E-04 |
| ENSG00000290073 | 42.1509 | -8.3316 | ##### | -3.9696 | 7.1979E-05 | 1.9998E-04 |
| ENSG00000288103 | 51.6471 | 7.4706 | ##### | 3.9696 | 7.1987E-05 | 1.9999E-04 |
| ENSG00000131591\|C1orf159 | 1639.5520 | 0.4552 | ##### | 3.9690 | 7.2185E-05 | 2.0052E-04 |
| ENSG00000123908\|AGO2 | 8491.3245 | 0.4807 | ##### | 3.9685 | 7.2315E-05 | 2.0086E-04 |
| ENSG00000279368 | 13.9892 | -2.7998 | ##### | -3.9683 | 7.2399E-05 | 2.0108E-04 |
| ENSG00000196408\|NOXO1 | 87.2903 | 1.0389 | ##### | 3.9680 | 7.2466E-05 | 2.0124E-04 |
| ENSG00000138031\|ADCY3 | 7724.0063 | 0.4480 | ##### | 3.9679 | 7.2515E-05 | 2.0136E-04 |
| ENSG00000172939\|OXSR1 | 4953.8461 | 0.4889 | ##### | 3.9678 | 7.2548E-05 | 2.0144E-04 |
| ENSG00000223714\|LINC02601 | 53.2925 | -8.0552 | ##### | -3.9676 | 7.2590E-05 | 2.0153E-04 |
| ENSG00000188613\|NANOS1 | 338.8008 | -0.6726 | ##### | -3.9668 | 7.2842E-05 | 2.0221E-04 |
| ENSG00000068097\|HEATR6 | 2596.8742 | 0.4869 | ##### | 3.9663 | 7.2983E-05 | 2.0259E-04 |
| ENSG00000271810 | 39.7407 | 7.5679 | ##### | 3.9663 | 7.2995E-05 | 2.0260E-04 |
| ENSG00000260955 | 41.1954 | 7.5678 | ##### | 3.9663 | 7.3003E-05 | 2.0261E-04 |
| ENSG00000143458\|GABPB2 | 1128.6947 | 0.5507 | ##### | 3.9657 | 7.3187E-05 | 2.0310E-04 |
| ENSG00000162105\|SHANK2 | 2591.2957 | 0.4949 | ##### | 3.9644 | 7.3584E-05 | 2.0418E-04 |
| ENSG00000125703\|ATG4C | 1034.3009 | 0.5279 | ##### | 3.9637 | 7.3790E-05 | 2.0473E-04 |
| ENSG00000141577\|CEP131 | 1838.2326 | 0.4969 | ##### | 3.9631 | 7.3978E-05 | 2.0523E-04 |
| ENSG00000171206\|TRIM8 | 3590.3310 | -0.4727 | ##### | -3.9624 | 7.4208E-05 | 2.0585E-04 |
| ENSG00000198604\|BAZ1A | 6079.5218 | 0.4801 | ##### | 3.9622 | 7.4258E-05 | 2.0597E-04 |
| ENSG00000221866\|PLXNA4 | 78.3549 | 4.7403 | ##### | 3.9617 | 7.4416E-05 | 2.0639E-04 |
| ENSG00000227581\|STAU2P1 | 19.9918 | -6.5514 | ##### | -3.9594 | 7.5128E-05 | 2.0835E-04 |
| ENSG00000167588\|GPD1 | 43.9156 | -1.6033 | ##### | -3.9587 | 7.5365E-05 | 2.0899E-04 |
| ENSG00000143318\|CASQ1 | 55.0682 | -7.9975 | ##### | -3.9582 | 7.5512E-05 | 2.0937E-04 |
| ENSG00000155636\|RBM45 | 1062.6307 | -0.5382 | ##### | -3.9575 | 7.5731E-05 | 2.0996E-04 |

| | | | | | | |
|---|---|---|---|---|---|---|
| ENSG00000083750\|RRAGB | 349.9389 | -0.6660 | ##### | -3.9569 | 7.5932E-05 | 2.1050E-04 |
| ENSG00000287126 | 310.4121 | 0.6740 | ##### | 3.9567 | 7.5981E-05 | 2.1062E-04 |
| ENSG00000159882\|ZNF230 | 323.1580 | -0.5950 | ##### | -3.9563 | 7.6111E-05 | 2.1096E-04 |
| ENSG00000235526 | 34.7551 | 1.3657 | ##### | 3.9562 | 7.6161E-05 | 2.1107E-04 |
| ENSG00000233830\|EIF4HP1 | 246.8804 | -0.6476 | ##### | -3.9560 | 7.6209E-05 | 2.1119E-04 |
| ENSG00000071655\|MBD3 | 9588.9379 | 0.4716 | ##### | 3.9551 | 7.6507E-05 | 2.1199E-04 |
| ENSG00000214087\|ARL16 | 2352.0048 | -0.4797 | ##### | -3.9530 | 7.7179E-05 | 2.1384E-04 |
| ENSG00000157600\|TMEM164 | 6684.5223 | -0.4887 | ##### | -3.9527 | 7.7267E-05 | 2.1406E-04 |
| ENSG00000204599\|TRIM39 | 1248.6318 | 0.5482 | ##### | 3.9527 | 7.7289E-05 | 2.1410E-04 |
| ENSG00000167755\|KLK6 | 18.2579 | -6.6016 | ##### | -3.9506 | 7.7964E-05 | 2.1595E-04 |
| ENSG00000267058 | 149.9454 | -2.2492 | ##### | -3.9504 | 7.8015E-05 | 2.1607E-04 |
| ENSG00000061337\|LZTS1 | 24.9335 | 3.7895 | ##### | 3.9498 | 7.8226E-05 | 2.1664E-04 |
| ENSG00000103460\|TOX3 | 36.3189 | -4.7620 | ##### | -3.9491 | 7.8451E-05 | 2.1724E-04 |
| ENSG00000235605 | 45.5606 | 1.1351 | ##### | 3.9486 | 7.8601E-05 | 2.1764E-04 |
| ENSG00000168298\|H1-4 | 48.9165 | 2.0588 | ##### | 3.9480 | 7.8811E-05 | 2.1820E-04 |
| ENSG00000287539 | 29.8816 | 1.7105 | ##### | 3.9470 | 7.9151E-05 | 2.1912E-04 |
| ENSG00000124587\|PEX6 | 1613.4089 | -0.5173 | ##### | -3.9465 | 7.9306E-05 | 2.1953E-04 |
| ENSG00000162980\|ARL5A | 6706.7476 | -0.4928 | ##### | -3.9454 | 7.9669E-05 | 2.2051E-04 |
| ENSG00000179886\|TIGD5 | 2745.4264 | 0.4438 | ##### | 3.9451 | 7.9763E-05 | 2.2075E-04 |
| ENSG00000181896\|ZNF101 | 765.7249 | 0.5622 | ##### | 3.9449 | 7.9824E-05 | 2.2090E-04 |
| ENSG00000156218\|ADAMTSL3 | 30.1077 | -8.2301 | ##### | -3.9448 | 7.9870E-05 | 2.2101E-04 |
| ENSG00000270800\|RPS10-NUDT3 | 4397.5044 | 0.6619 | ##### | 3.9443 | 8.0039E-05 | 2.2145E-04 |
| ENSG00000034693\|PEX3 | 1177.3771 | -0.5016 | ##### | -3.9442 | 8.0057E-05 | 2.2148E-04 |
| ENSG00000225507 | 83.3445 | -1.0920 | ##### | -3.9441 | 8.0094E-05 | 2.2154E-04 |
| ENSG00000213927\|CCL27 | 11.4684 | 3.0717 | ##### | 3.9441 | 8.0094E-05 | 2.2154E-04 |
| ENSG00000145495\|MARCHF6 | 12221.9904 | -0.4643 | ##### | -3.9438 | 8.0191E-05 | 2.2179E-04 |
| ENSG00000148484\|RSU1 | 8296.1205 | 0.4888 | ##### | 3.9435 | 8.0303E-05 | 2.2208E-04 |
| ENSG00000196747\|H2AC13 | 93.4403 | 1.0591 | ##### | 3.9429 | 8.0488E-05 | 2.2257E-04 |
| ENSG00000279382 | 139.7881 | 0.9710 | ##### | 3.9426 | 8.0614E-05 | 2.2290E-04 |
| ENSG00000100473\|COCH | 725.4456 | -0.6815 | ##### | -3.9425 | 8.0649E-05 | 2.2297E-04 |
| ENSG00000173988\|LRRC63 | 63.3777 | -0.9706 | ##### | -3.9421 | 8.0775E-05 | 2.2330E-04 |
| ENSG00000169548\|ZNF280A | 88.6928 | -0.9740 | ##### | -3.9398 | 8.1545E-05 | 2.2541E-04 |
| ENSG00000185347\|TEDC1 | 2217.5512 | 0.4880 | ##### | 3.9385 | 8.1998E-05 | 2.2664E-04 |
| ENSG00000287382 | 13.4119 | 6.3530 | ##### | 3.9383 | 8.2068E-05 | 2.2681E-04 |
| ENSG00000167617\|CDC42EP5 | 142.2489 | 0.9115 | ##### | 3.9369 | 8.2525E-05 | 2.2806E-04 |
| ENSG00000169919\|GUSB | 3112.4841 | 0.4673 | ##### | 3.9368 | 8.2579E-05 | 2.2818E-04 |
| ENSG00000165113\|GKAP1 | 337.3789 | -0.5971 | ##### | -3.9367 | 8.2598E-05 | 2.2822E-04 |
| ENSG00000249413 | 22.6761 | -2.6988 | ##### | -3.9363 | 8.2740E-05 | 2.2859E-04 |
| ENSG00000174950\|CD164L2 | 11.9869 | -6.9944 | ##### | -3.9360 | 8.2837E-05 | 2.2884E-04 |
| ENSG00000224680\|PLA2G12AP1 | 21.6054 | 1.7030 | ##### | 3.9350 | 8.3197E-05 | 2.2981E-04 |
| ENSG00000100138\|SNU13 | 8784.5997 | 0.3805 | ##### | 3.9349 | 8.3243E-05 | 2.2991E-04 |
| ENSG00000228696\|ARL17B | 1313.4634 | 0.4923 | ##### | 3.9347 | 8.3318E-05 | 2.3010E-04 |
| ENSG00000224376 | 15.8584 | -3.5619 | ##### | -3.9336 | 8.3687E-05 | 2.3110E-04 |
| ENSG00000159958\|TNFRSF13C | 83.4557 | 0.9395 | ##### | 3.9333 | 8.3787E-05 | 2.3135E-04 |
| ENSG00000103429\|BFAR | 5964.0487 | -0.4777 | ##### | -3.9321 | 8.4200E-05 | 2.3247E-04 |
| ENSG00000215187\|CIMIP2B | 57.2576 | 1.0627 | ##### | 3.9292 | 8.5218E-05 | 2.3526E-04 |
| ENSG00000125484\|GTF3C4 | 8236.7968 | 0.4714 | ##### | 3.9289 | 8.5338E-05 | 2.3557E-04 |
| ENSG00000248429\|GASK1B-AS1 | 37.6213 | -1.5418 | ##### | -3.9282 | 8.5595E-05 | 2.3626E-04 |
| ENSG00000015479\|MATR3 | 18854.5307 | 1.0660 | ##### | 3.9281 | 8.5634E-05 | 2.3634E-04 |
| ENSG00000108509\|CAMTA2 | 3504.5005 | -0.4799 | ##### | -3.9271 | 8.5986E-05 | 2.3729E-04 |
| ENSG00000249471\|ZNF324B | 467.4256 | -0.5685 | ##### | -3.9266 | 8.6152E-05 | 2.3773E-04 |
| ENSG00000284070 | 28.2315 | -2.5467 | ##### | -3.9264 | 8.6210E-05 | 2.3787E-04 |
| ENSG00000213406\|ANXA2P1 | 21.7426 | -1.4403 | ##### | -3.9255 | 8.6541E-05 | 2.3876E-04 |
| ENSG00000104671\|DCTN6 | 2286.4319 | -0.5004 | ##### | -3.9252 | 8.6666E-05 | 2.3908E-04 |
| ENSG00000198252\|STYX | 2551.5868 | 0.4896 | ##### | 3.9249 | 8.6756E-05 | 2.3931E-04 |
| ENSG00000178338\|ZNF354B | 466.6375 | 0.6094 | ##### | 3.9242 | 8.7020E-05 | 2.4001E-04 |
| ENSG00000141425\|RPRD1A | 10444.4325 | -0.4654 | ##### | -3.9218 | 8.7886E-05 | 2.4238E-04 |
| ENSG00000239389\|PCDHA13 | 183.4289 | -1.0239 | ##### | -3.9217 | 8.7926E-05 | 2.4247E-04 |
| ENSG00000116350\|SRSF4 | 6445.1806 | 0.4591 | ##### | 3.9214 | 8.8024E-05 | 2.4272E-04 |
| ENSG00000225792\|SNX10-AS1 | 22.0550 | -1.7792 | ##### | -3.9211 | 8.8158E-05 | 2.4306E-04 |
| ENSG00000205208\|C4orf46 | 1899.5158 | -0.4832 | ##### | -3.9207 | 8.8278E-05 | 2.4337E-04 |
| ENSG00000291342 | 12.6250 | 2.0456 | ##### | 3.9186 | 8.9059E-05 | 2.4550E-04 |
| ENSG00000139531\|SUOX | 1358.5843 | -0.5067 | ##### | -3.9179 | 8.9336E-05 | 2.4624E-04 |

| | | | | | | |
|---|---|---|---|---|---|---|
| ENSG00000042062\|RIPOR3 | 26.2330 | 2.2275 | ##### | 3.9177 | 8.9381E-05 | 2.4634E-04 |
| ENSG00000177694\|NAALADL2 | 65.1967 | -1.2665 | ##### | -3.9171 | 8.9631E-05 | 2.4701E-04 |
| ENSG00000227500\|SCAMP4 | 6186.5577 | 0.4853 | ##### | 3.9169 | 8.9689E-05 | 2.4714E-04 |
| ENSG00000140836\|ZFHX3 | 2193.3591 | -0.5029 | ##### | -3.9168 | 8.9737E-05 | 2.4725E-04 |
| ENSG00000228021 | 32.7526 | -4.4261 | ##### | -3.9164 | 8.9874E-05 | 2.4761E-04 |
| ENSG00000112406\|HECA | 676.8265 | -0.5522 | ##### | -3.9161 | 8.9986E-05 | 2.4789E-04 |
| ENSG00000075234\|TTC38 | 979.0341 | -0.4827 | ##### | -3.9160 | 9.0013E-05 | 2.4795E-04 |
| ENSG00000156261\|CCT8 | 34701.1140 | 0.3840 | ##### | 3.9160 | 9.0047E-05 | 2.4802E-04 |
| ENSG00000164989\|CCDC171 | 130.3160 | 0.8611 | ##### | 3.9156 | 9.0168E-05 | 2.4833E-04 |
| ENSG00000100216\|TOMM22 | 5506.9200 | 0.4195 | ##### | 3.9152 | 9.0324E-05 | 2.4873E-04 |
| ENSG00000108306\|FBXL20 | 1366.7631 | -0.4946 | ##### | -3.9147 | 9.0520E-05 | 2.4925E-04 |
| ENSG00000260828\|HMGB3P32 | 16.2456 | 5.5756 | ##### | 3.9146 | 9.0549E-05 | 2.4931E-04 |
| ENSG00000117054\|ACADM | 4558.1045 | -0.4766 | ##### | -3.9143 | 9.0671E-05 | 2.4962E-04 |
| ENSG00000130741\|EIF2S3 | 17996.5044 | -0.4623 | ##### | -3.9136 | 9.0926E-05 | 2.5030E-04 |
| ENSG00000196923\|PDLIM7 | 5854.8450 | 0.4789 | ##### | 3.9132 | 9.1090E-05 | 2.5073E-04 |
| ENSG00000159409\|CELF3 | 19.7364 | 3.4854 | ##### | 3.9131 | 9.1122E-05 | 2.5079E-04 |
| ENSG00000230426\|LINC01036 | 62.9408 | -4.9967 | ##### | -3.9128 | 9.1215E-05 | 2.5103E-04 |
| ENSG00000203668\|CHML | 5129.7598 | -0.4790 | ##### | -3.9127 | 9.1264E-05 | 2.5114E-04 |
| ENSG00000172183\|ISG20 | 1151.8210 | -0.5406 | ##### | -3.9121 | 9.1515E-05 | 2.5181E-04 |
| ENSG00000203877\|RIPPLY2 | 237.7739 | 0.6772 | ##### | 3.9117 | 9.1651E-05 | 2.5216E-04 |
| ENSG00000143110\|C1orf162 | 28.3380 | 1.7853 | ##### | 3.9112 | 9.1823E-05 | 2.5261E-04 |
| ENSG00000159592\|GPBP1L1 | 6583.3837 | 0.4886 | ##### | 3.9104 | 9.2158E-05 | 2.5350E-04 |
| ENSG00000112855\|HARS2 | 3194.1168 | 0.4965 | ##### | 3.9101 | 9.2246E-05 | 2.5372E-04 |
| ENSG00000198746\|GPATCH3 | 1224.2353 | 0.5407 | ##### | 3.9093 | 9.2580E-05 | 2.5462E-04 |
| ENSG00000178031\|ADAMTSL1 | 17.8524 | -6.0415 | ##### | -3.9088 | 9.2746E-05 | 2.5505E-04 |
| ENSG00000148356\|LRSAM1 | 1857.3611 | -0.4746 | ##### | -3.9086 | 9.2816E-05 | 2.5522E-04 |
| ENSG00000260536 | 15.6521 | 6.0316 | ##### | 3.9084 | 9.2915E-05 | 2.5547E-04 |
| ENSG00000283662 | 13.0368 | 3.6783 | ##### | 3.9083 | 9.2940E-05 | 2.5551E-04 |
| ENSG00000204121\|ECEL1P1 | 26.4731 | 5.0078 | ##### | 3.9082 | 9.3003E-05 | 2.5566E-04 |
| ENSG00000250539\|KRT8P33 | 50.3052 | 1.3353 | ##### | 3.9076 | 9.3201E-05 | 2.5619E-04 |
| ENSG00000028839\|TBPL1 | 1216.0118 | 0.5332 | ##### | 3.9076 | 9.3235E-05 | 2.5625E-04 |
| ENSG00000165915\|SLC39A13 | 1941.2657 | -0.4995 | ##### | -3.9071 | 9.3395E-05 | 2.5667E-04 |
| ENSG00000256394\|ASIC5 | 26.8317 | -7.6774 | ##### | -3.9068 | 9.3516E-05 | 2.5698E-04 |
| ENSG00000270617\|URGCP-MRPS24 | 201.9067 | -9.2189 | ##### | -3.9067 | 9.3584E-05 | 2.5714E-04 |
| ENSG00000197769\|MAP1LC3C | 23.1436 | -5.0368 | ##### | -3.9062 | 9.3747E-05 | 2.5757E-04 |
| ENSG00000167625\|ZNF526 | 2046.1517 | -0.4938 | ##### | -3.9042 | 9.4534E-05 | 2.5971E-04 |
| ENSG00000108064\|TFAM | 8009.8708 | 0.4773 | ##### | 3.9039 | 9.4665E-05 | 2.6004E-04 |
| ENSG00000142609\|CFAP74 | 24.0529 | 3.4166 | ##### | 3.9037 | 9.4735E-05 | 2.6021E-04 |
| ENSG00000291206\|GTF2IRD1P1 | 298.9587 | -0.6628 | ##### | -3.9035 | 9.4828E-05 | 2.6044E-04 |
| ENSG00000196712\|NF1 | 8406.4913 | 0.4623 | ##### | 3.9029 | 9.5032E-05 | 2.6098E-04 |
| ENSG00000289135 | 209.9309 | 0.7589 | ##### | 3.9012 | 9.5714E-05 | 2.6283E-04 |
| ENSG00000250101 | 29.6887 | 7.8216 | ##### | 3.9009 | 9.5833E-05 | 2.6313E-04 |
| ENSG00000132463\|GRSF1 | 9103.9357 | -0.4712 | ##### | -3.9006 | 9.5967E-05 | 2.6347E-04 |
| ENSG00000270562\|FOXP1-DT | 15.6576 | -6.1136 | ##### | -3.8997 | 9.6298E-05 | 2.6436E-04 |
| ENSG00000136450\|SRSF1 | 32096.9699 | 0.4510 | ##### | 3.8995 | 9.6398E-05 | 2.6461E-04 |
| ENSG00000174109\|UQCC4 | 1037.5299 | -0.5230 | ##### | -3.8990 | 9.6603E-05 | 2.6515E-04 |
| ENSG00000116688\|MFN2 | 13824.1655 | -0.4541 | ##### | -3.8983 | 9.6869E-05 | 2.6583E-04 |
| ENSG00000137507\|LRRC32 | 59.5050 | -1.1139 | ##### | -3.8983 | 9.6864E-05 | 2.6583E-04 |
| ENSG00000131374\|TBC1D5 | 5077.2900 | -0.5491 | ##### | -3.8974 | 9.7225E-05 | 2.6678E-04 |
| ENSG00000128218\|VPREB3 | 40.3457 | -3.6467 | ##### | -3.8971 | 9.7363E-05 | 2.6713E-04 |
| ENSG00000136897\|MRPL50 | 2451.6103 | -0.4760 | ##### | -3.8969 | 9.7442E-05 | 2.6733E-04 |
| ENSG00000153207\|AHCTF1 | 11191.6606 | 0.4639 | ##### | 3.8966 | 9.7552E-05 | 2.6760E-04 |
| ENSG00000261549 | 12.3779 | -4.3768 | ##### | -3.8966 | 9.7560E-05 | 2.6760E-04 |
| ENSG00000275484 | 40.1910 | 3.2940 | ##### | 3.8964 | 9.7630E-05 | 2.6777E-04 |
| ENSG00000273611\|ZNHIT3 | 2398.8111 | -0.4802 | ##### | -3.8962 | 9.7705E-05 | 2.6795E-04 |
| ENSG00000158859\|ADAMTS4 | 26.6725 | 2.9918 | ##### | 3.8943 | 9.8473E-05 | 2.7003E-04 |
| ENSG00000101986\|ABCD1 | 2079.9597 | -0.4935 | ##### | -3.8938 | 9.8670E-05 | 2.7055E-04 |
| ENSG00000204625\|HCG9 | 19.2853 | 2.8527 | ##### | 3.8927 | 9.9119E-05 | 2.7175E-04 |
| ENSG00000165072\|MAMDC2 | 27.0340 | 2.9585 | ##### | 3.8927 | 9.9142E-05 | 2.7179E-04 |
| ENSG00000135211\|TMEM60 | 707.5173 | -0.5630 | ##### | -3.8926 | 9.9160E-05 | 2.7181E-04 |
| ENSG00000236352 | 23.0128 | 6.1068 | ##### | 3.8920 | 9.9436E-05 | 2.7254E-04 |
| ENSG00000142686\|C1orf216 | 2208.6892 | 0.4876 | ##### | 3.8914 | 9.9660E-05 | 2.7313E-04 |
| ENSG00000113494\|PRLR | 243.2374 | 4.3914 | ##### | 3.8912 | 9.9757E-05 | 2.7337E-04 |

| | | | | | | |
|---|---|---|---|---|---|---|
| ENSG00000277496\|SLCO4A1-AS2 | 73.6283 | 2.5411 | ##### | 3.8895 | 1.0047E-04 | 2.7530E-04 |
| ENSG00000149926\|TLCD3B | 50.1270 | -1.0720 | ##### | -3.8880 | 1.0108E-04 | 2.7694E-04 |
| ENSG00000146416\|AIG1 | 1186.7485 | -0.4824 | ##### | -3.8873 | 1.0137E-04 | 2.7771E-04 |
| ENSG00000122406\|RPL5 | 46079.3114 | -0.3746 | ##### | -3.8872 | 1.0143E-04 | 2.7785E-04 |
| ENSG00000167522\|ANKRD11 | 13538.4896 | -0.4614 | ##### | -3.8871 | 1.0145E-04 | 2.7788E-04 |
| ENSG00000244040\|IL12A-AS1 | 11.6665 | 6.3081 | ##### | 3.8869 | 1.0153E-04 | 2.7807E-04 |
| ENSG00000234742\|RPL17P18 | 47.0927 | -1.0725 | ##### | -3.8863 | 1.0178E-04 | 2.7875E-04 |
| ENSG00000255910 | 32.9875 | -7.9761 | ##### | -3.8847 | 1.0244E-04 | 2.8053E-04 |
| ENSG00000182512\|GLRX5 | 4353.0037 | 0.4875 | ##### | 3.8842 | 1.0265E-04 | 2.8108E-04 |
| ENSG00000260459\|FTLP14 | 31.0532 | -7.2038 | ##### | -3.8842 | 1.0266E-04 | 2.8108E-04 |
| ENSG00000088986\|DYNLL1 | 17221.0911 | 0.4571 | ##### | 3.8835 | 1.0298E-04 | 2.8192E-04 |
| ENSG00000133243\|BTBD2 | 11144.8983 | 0.4595 | ##### | 3.8830 | 1.0320E-04 | 2.8249E-04 |
| ENSG00000245482 | 78.1279 | 1.1828 | ##### | 3.8829 | 1.0323E-04 | 2.8256E-04 |
| ENSG00000144580\|CNOT9 | 7899.6358 | 0.4017 | ##### | 3.8827 | 1.0332E-04 | 2.8277E-04 |
| ENSG00000188582\|PAQR9 | 461.8760 | -0.6443 | ##### | -3.8823 | 1.0347E-04 | 2.8316E-04 |
| ENSG00000255730 | 21.9336 | 4.4692 | ##### | 3.8822 | 1.0354E-04 | 2.8332E-04 |
| ENSG00000164104\|HMGB2 | 5134.0322 | 0.4792 | ##### | 3.8821 | 1.0357E-04 | 2.8340E-04 |
| ENSG00000178950\|GAK | 6964.8761 | 0.4631 | ##### | 3.8814 | 1.0386E-04 | 2.8414E-04 |
| ENSG00000135625\|EGR4 | 46.8718 | 4.0371 | ##### | 3.8812 | 1.0396E-04 | 2.8440E-04 |
| ENSG00000160193\|WDR4 | 3357.6902 | -0.4647 | ##### | -3.8810 | 1.0403E-04 | 2.8456E-04 |
| ENSG00000120217\|CD274 | 59.7949 | -1.3677 | ##### | -3.8808 | 1.0410E-04 | 2.8474E-04 |
| ENSG00000197696\|NMB | 435.9027 | 0.6058 | ##### | 3.8800 | 1.0446E-04 | 2.8568E-04 |
| ENSG00000290790 | 8.0839 | 2.4541 | ##### | 3.8794 | 1.0473E-04 | 2.8640E-04 |
| ENSG00000180764\|PIPSL | 153.8716 | 1.2002 | ##### | 3.8791 | 1.0483E-04 | 2.8664E-04 |
| ENSG00000186628\|FSD2 | 95.1274 | 0.8970 | ##### | 3.8788 | 1.0497E-04 | 2.8701E-04 |
| ENSG00000148300\|REXO4 | 3699.1362 | -0.4851 | ##### | -3.8782 | 1.0524E-04 | 2.8771E-04 |
| ENSG00000198832\|SELENOM | 4053.9867 | -0.4705 | ##### | -3.8775 | 1.0552E-04 | 2.8844E-04 |
| ENSG00000267261 | 10.5993 | 6.1722 | ##### | 3.8773 | 1.0563E-04 | 2.8874E-04 |
| ENSG00000169067\|ACTBL2 | 22.6240 | -3.2034 | ##### | -3.8768 | 1.0582E-04 | 2.8923E-04 |
| ENSG00000156398\|SFXN2 | 612.5765 | 0.5823 | ##### | 3.8762 | 1.0611E-04 | 2.8999E-04 |
| ENSG00000268034\|TPM3P5 | 89.1795 | 0.9768 | ##### | 3.8761 | 1.0613E-04 | 2.9002E-04 |
| ENSG00000135469\|COQ10A | 1088.8968 | 0.5273 | ##### | 3.8747 | 1.0677E-04 | 2.9174E-04 |
| ENSG00000214719 | 785.3368 | -0.5403 | ##### | -3.8744 | 1.0687E-04 | 2.9198E-04 |
| ENSG00000223773\|CD99P1 | 23.4775 | -4.8262 | ##### | -3.8744 | 1.0688E-04 | 2.9198E-04 |
| ENSG00000227006 | 36.0243 | -5.0824 | ##### | -3.8741 | 1.0701E-04 | 2.9230E-04 |
| ENSG00000148908\|RGS10 | 1178.0157 | 0.5275 | ##### | 3.8740 | 1.0708E-04 | 2.9247E-04 |
| ENSG00000106028\|SSBP1 | 10140.0219 | 0.4702 | ##### | 3.8734 | 1.0731E-04 | 2.9308E-04 |
| ENSG00000136936\|XPA | 758.7343 | -0.5603 | ##### | -3.8724 | 1.0777E-04 | 2.9432E-04 |
| ENSG00000255329\|H2AZP4 | 16.3475 | 5.8215 | ##### | 3.8723 | 1.0779E-04 | 2.9435E-04 |
| ENSG00000286591 | 37.9912 | 2.4385 | ##### | 3.8723 | 1.0783E-04 | 2.9441E-04 |
| ENSG00000182327\|GLTPD2 | 88.3819 | -1.1361 | ##### | -3.8722 | 1.0787E-04 | 2.9450E-04 |
| ENSG00000251050\|RBX1P2 | 13.6138 | 6.2177 | ##### | 3.8720 | 1.0794E-04 | 2.9467E-04 |
| ENSG00000198585\|NUDT16 | 1798.2837 | -0.5009 | ##### | -3.8718 | 1.0802E-04 | 2.9485E-04 |
| ENSG00000152464\|RPP38 | 1957.4792 | 0.5030 | ##### | 3.8711 | 1.0834E-04 | 2.9570E-04 |
| ENSG00000230325 | 18.3721 | 2.2219 | ##### | 3.8709 | 1.0845E-04 | 2.9598E-04 |
| ENSG00000171262\|FAM98B | 3732.6742 | 0.4750 | ##### | 3.8704 | 1.0867E-04 | 2.9654E-04 |
| ENSG00000254815\|LMNTD2-AS1 | 141.5893 | -0.9120 | ##### | -3.8703 | 1.0870E-04 | 2.9660E-04 |
| ENSG00000291066 | 41.4038 | -5.8204 | ##### | -3.8701 | 1.0878E-04 | 2.9679E-04 |
| ENSG00000120727\|PAIP2 | 6659.9219 | 0.4612 | ##### | 3.8701 | 1.0880E-04 | 2.9681E-04 |
| ENSG00000235072\|ARNILA | 27.3133 | 1.7992 | ##### | 3.8694 | 1.0909E-04 | 2.9758E-04 |
| ENSG00000172780\|RAB43 | 1710.6063 | -0.5130 | ##### | -3.8674 | 1.0999E-04 | 3.0002E-04 |
| ENSG00000081138\|CDH7 | 256.6109 | -6.6003 | ##### | -3.8673 | 1.1004E-04 | 3.0012E-04 |
| ENSG00000280407 | 396.3899 | 1.2113 | ##### | 3.8672 | 1.1008E-04 | 3.0018E-04 |
| ENSG00000144909\|OSBPL11 | 3261.3944 | 0.4791 | ##### | 3.8672 | 1.1008E-04 | 3.0018E-04 |
| ENSG00000286886 | 21.3700 | 1.5916 | ##### | 3.8667 | 1.1033E-04 | 3.0084E-04 |
| ENSG00000187559\|FOXD4L3 | 14.7485 | 1.8746 | ##### | 3.8664 | 1.1045E-04 | 3.0114E-04 |
| ENSG00000169403\|PTAFR | 63.4053 | -0.9000 | ##### | -3.8663 | 1.1049E-04 | 3.0122E-04 |
| ENSG00000229688\|CRPPA-AS1 | 36.4618 | -7.8451 | ##### | -3.8660 | 1.1062E-04 | 3.0153E-04 |
| ENSG00000147912\|FBXO10 | 1148.0062 | 0.5364 | ##### | 3.8659 | 1.1068E-04 | 3.0168E-04 |
| ENSG00000236502\|SIX3-AS1 | 33.4345 | -7.4143 | ##### | -3.8653 | 1.1095E-04 | 3.0238E-04 |
| ENSG00000179262\|RAD23A | 11546.1261 | -0.4572 | ##### | -3.8646 | 1.1127E-04 | 3.0323E-04 |
| ENSG00000268189 | 225.5535 | 1.2707 | ##### | 3.8638 | 1.1162E-04 | 3.0415E-04 |
| ENSG00000256879\|PDE3A-AS1 | 11.2445 | -2.9113 | ##### | -3.8635 | 1.1178E-04 | 3.0455E-04 |

| | | | | | | |
|---|---|---|---|---|---|---|
| ENSG00000119688\|ABCD4 | 1463.3630 | 0.4904 | ##### | 3.8632 | 1.1192E-04 | 3.0491E-04 |
| ENSG00000185730\|ZNF696 | 1706.5249 | 0.5079 | ##### | 3.8628 | 1.1209E-04 | 3.0535E-04 |
| ENSG00000287629 | 45.5530 | -1.2820 | ##### | -3.8622 | 1.1236E-04 | 3.0606E-04 |
| ENSG00000269371 | 49.0850 | -1.2273 | ##### | -3.8620 | 1.1245E-04 | 3.0628E-04 |
| ENSG00000272508 | 9.9632 | 6.0777 | ##### | 3.8618 | 1.1254E-04 | 3.0649E-04 |
| ENSG00000286456 | 46.0530 | -1.1935 | ##### | -3.8599 | 1.1342E-04 | 3.0885E-04 |
| ENSG00000289285 | 24.1039 | -1.3875 | ##### | -3.8593 | 1.1372E-04 | 3.0966E-04 |
| ENSG00000168710\|AHCYL1 | 19185.5978 | -0.4232 | ##### | -3.8586 | 1.1404E-04 | 3.1049E-04 |
| ENSG00000127415\|IDUA | 373.0471 | -0.5976 | ##### | -3.8580 | 1.1430E-04 | 3.1119E-04 |
| ENSG00000162391\|FAM151A | 83.0889 | 0.8501 | ##### | 3.8577 | 1.1444E-04 | 3.1152E-04 |
| ENSG00000278299\|TBC1D3C | 205.7222 | 0.6655 | ##### | 3.8576 | 1.1451E-04 | 3.1169E-04 |
| ENSG00000188976\|NOC2L | 31492.8247 | 0.3675 | ##### | 3.8573 | 1.1466E-04 | 3.1207E-04 |
| ENSG00000293503 | 40.7073 | -1.6819 | ##### | -3.8571 | 1.1473E-04 | 3.1224E-04 |
| ENSG00000128645\|HOXD1 | 47.4287 | 4.2479 | ##### | 3.8559 | 1.1528E-04 | 3.1371E-04 |
| ENSG00000238245\|MYO5BP2 | 35.1730 | -1.8963 | ##### | -3.8557 | 1.1542E-04 | 3.1404E-04 |
| ENSG00000112851\|ERBIN | 8857.9464 | -0.4509 | ##### | -3.8552 | 1.1564E-04 | 3.1462E-04 |
| ENSG00000161281\|COX7A1 | 15.9720 | 6.0813 | ##### | 3.8549 | 1.1576E-04 | 3.1492E-04 |
| ENSG00000135577\|NMBR | 37.0122 | -8.1429 | ##### | -3.8548 | 1.1583E-04 | 3.1507E-04 |
| ENSG00000276742 | 5.6982 | -3.0290 | ##### | -3.8547 | 1.1586E-04 | 3.1514E-04 |
| ENSG00000282386 | 117.9336 | 0.8797 | ##### | 3.8545 | 1.1596E-04 | 3.1537E-04 |
| ENSG00000263585 | 763.0325 | -0.5480 | ##### | -3.8544 | 1.1600E-04 | 3.1546E-04 |
| ENSG00000197467\|COL13A1 | 29.3890 | 2.0701 | ##### | 3.8540 | 1.1618E-04 | 3.1592E-04 |
| ENSG00000113269\|RNF130 | 3770.4803 | 0.4678 | ##### | 3.8536 | 1.1640E-04 | 3.1647E-04 |
| ENSG00000198689\|SLC9A6 | 2334.3827 | 0.5040 | ##### | 3.8526 | 1.1685E-04 | 3.1768E-04 |
| ENSG00000136273\|HUS1 | 2143.1443 | -0.4951 | ##### | -3.8516 | 1.1733E-04 | 3.1896E-04 |
| ENSG00000224875\|EML4-AS1 | 25.8014 | 1.7446 | ##### | 3.8512 | 1.1753E-04 | 3.1947E-04 |
| ENSG00000116560\|SFPQ | 28458.1249 | 0.3258 | ##### | 3.8506 | 1.1781E-04 | 3.2018E-04 |
| ENSG00000204590\|GNL1 | 5855.8718 | 0.4514 | ##### | 3.8506 | 1.1781E-04 | 3.2018E-04 |
| ENSG00000165300\|SLITRK5 | 692.8380 | 0.5410 | ##### | 3.8505 | 1.1785E-04 | 3.2027E-04 |
| ENSG00000122545\|SEPTIN7 | 18848.8127 | -0.2932 | ##### | -3.8486 | 1.1879E-04 | 3.2277E-04 |
| ENSG00000104964\|TLE5 | 29689.0420 | -0.4690 | ##### | -3.8485 | 1.1887E-04 | 3.2296E-04 |
| ENSG00000124608\|AARS2 | 1812.5091 | -0.4932 | ##### | -3.8478 | 1.1920E-04 | 3.2383E-04 |
| ENSG00000228532\|SUMO2P21 | 41.3611 | 1.2325 | ##### | 3.8474 | 1.1937E-04 | 3.2426E-04 |
| ENSG00000162923\|WDR26 | 10324.5482 | 0.4689 | ##### | 3.8471 | 1.1953E-04 | 3.2466E-04 |
| ENSG00000188313\|PLSCR1 | 1634.5262 | -0.5018 | ##### | -3.8465 | 1.1980E-04 | 3.2537E-04 |
| ENSG00000231006\|RPL7P32 | 24.8906 | -1.3875 | ##### | -3.8461 | 1.2003E-04 | 3.2598E-04 |
| ENSG00000173039\|RELA | 7211.4456 | 0.4748 | ##### | 3.8451 | 1.2052E-04 | 3.2727E-04 |
| ENSG00000233859\|ADH5P4 | 44.1697 | -1.1491 | ##### | -3.8450 | 1.2054E-04 | 3.2729E-04 |
| ENSG00000164818\|DNAAF5 | 10868.7581 | -0.4399 | ##### | -3.8450 | 1.2055E-04 | 3.2729E-04 |
| ENSG00000228925 | 52.0159 | 1.4177 | ##### | 3.8447 | 1.2068E-04 | 3.2762E-04 |
| ENSG00000275221\|H2AC15 | 34.4276 | 1.5551 | ##### | 3.8446 | 1.2076E-04 | 3.2780E-04 |
| ENSG00000145087\|STXBP5L | 18.8595 | -1.9477 | ##### | -3.8443 | 1.2089E-04 | 3.2813E-04 |
| ENSG00000248626\|GAPDHP40 | 15.7764 | 1.6911 | ##### | 3.8438 | 1.2112E-04 | 3.2872E-04 |
| ENSG00000071082\|RPL31 | 33644.5038 | -0.4747 | ##### | -3.8429 | 1.2161E-04 | 3.3001E-04 |
| ENSG00000235954\|TTC28-AS1 | 919.9383 | 0.4945 | ##### | 3.8424 | 1.2185E-04 | 3.3063E-04 |
| ENSG00000278112 | 30.7614 | 1.4392 | ##### | 3.8423 | 1.2190E-04 | 3.3075E-04 |
| ENSG00000290107 | 47.7035 | 3.4697 | ##### | 3.8422 | 1.2194E-04 | 3.3084E-04 |
| ENSG00000280143 | 439.6842 | -0.6189 | ##### | -3.8420 | 1.2205E-04 | 3.3108E-04 |
| ENSG00000259405\|ISCA1P4 | 12.7090 | -5.9255 | ##### | -3.8417 | 1.2218E-04 | 3.3141E-04 |
| ENSG00000153015\|CWC27 | 2925.1050 | 0.4853 | ##### | 3.8409 | 1.2259E-04 | 3.3250E-04 |
| ENSG00000151014\|NOCT | 1142.4702 | 0.4892 | ##### | 3.8407 | 1.2270E-04 | 3.3276E-04 |
| ENSG00000206052\|DOK6 | 117.5671 | 0.8807 | ##### | 3.8394 | 1.2331E-04 | 3.3439E-04 |
| ENSG00000259982\|CDC37P1 | 13.6993 | -5.4491 | ##### | -3.8394 | 1.2336E-04 | 3.3449E-04 |
| ENSG00000235655\|H3P6 | 646.4934 | -0.5389 | ##### | -3.8388 | 1.2361E-04 | 3.3516E-04 |
| ENSG00000284060 | 39.0576 | -1.5804 | ##### | -3.8377 | 1.2420E-04 | 3.3672E-04 |
| ENSG00000106144\|CASP2 | 4429.3731 | 0.4753 | ##### | 3.8375 | 1.2428E-04 | 3.3690E-04 |
| ENSG00000141570\|CBX8 | 1141.9717 | 0.5462 | ##### | 3.8374 | 1.2435E-04 | 3.3705E-04 |
| ENSG00000089289\|IGBP1 | 3432.7553 | 0.4767 | ##### | 3.8370 | 1.2457E-04 | 3.3763E-04 |
| ENSG00000228043\|SPOPL-DT | 45.1694 | -1.3688 | ##### | -3.8363 | 1.2492E-04 | 3.3854E-04 |
| ENSG00000261118 | 22.5278 | -2.2910 | ##### | -3.8351 | 1.2552E-04 | 3.4015E-04 |
| ENSG00000138303\|ASCC1 | 3147.3652 | -0.4934 | ##### | -3.8350 | 1.2555E-04 | 3.4020E-04 |
| ENSG00000268530 | 30.2231 | -1.3649 | ##### | -3.8349 | 1.2561E-04 | 3.4033E-04 |
| ENSG00000154645\|CHODL | 21.5094 | 4.5531 | ##### | 3.8338 | 1.2616E-04 | 3.4179E-04 |

| | | | | | | |
|---|---|---|---|---|---|---|
| ENSG00000101421\|CHMP4B | 6707.3580 | -0.3750 | ##### | -3.8337 | 1.2621E-04 | 3.4187E-04 |
| ENSG00000072042\|RDH11 | 9422.7998 | 0.4625 | ##### | 3.8335 | 1.2631E-04 | 3.4213E-04 |
| ENSG00000281571 | 187.4066 | -0.6930 | ##### | -3.8332 | 1.2650E-04 | 3.4261E-04 |
| ENSG00000225774\|SIRPAP1 | 9.8083 | -4.2395 | ##### | -3.8328 | 1.2668E-04 | 3.4305E-04 |
| ENSG00000070159\|PTPN3 | 4045.8855 | -0.4688 | ##### | -3.8326 | 1.2682E-04 | 3.4341E-04 |
| ENSG00000273079\|GRIN2B | 61.8368 | -3.5848 | ##### | -3.8319 | 1.2715E-04 | 3.4426E-04 |
| ENSG00000268565 | 23.8167 | 1.6016 | ##### | 3.8318 | 1.2719E-04 | 3.4435E-04 |
| ENSG00000004455\|AK2 | 16645.0102 | 0.2380 | ##### | 3.8314 | 1.2740E-04 | 3.4489E-04 |
| ENSG00000167046 | 20.8956 | 3.2972 | ##### | 3.8310 | 1.2764E-04 | 3.4550E-04 |
| ENSG00000223855\|PDGFA-DT | 24.9249 | -4.1837 | ##### | -3.8309 | 1.2769E-04 | 3.4561E-04 |
| ENSG00000146425\|DYNLT1 | 2528.7818 | -0.4659 | ##### | -3.8308 | 1.2772E-04 | 3.4565E-04 |
| ENSG00000285446 | 183.8341 | -9.9429 | ##### | -3.8306 | 1.2785E-04 | 3.4599E-04 |
| ENSG00000285677 | 51.2671 | 1.6049 | ##### | 3.8303 | 1.2799E-04 | 3.4634E-04 |
| ENSG00000135744\|AGT | 280.1075 | -1.2858 | ##### | -3.8300 | 1.2816E-04 | 3.4675E-04 |
| ENSG00000272940 | 21.6692 | -6.0328 | ##### | -3.8295 | 1.2842E-04 | 3.4743E-04 |
| ENSG00000267737 | 31.6685 | -7.8680 | ##### | -3.8279 | 1.2922E-04 | 3.4957E-04 |
| ENSG00000278259\|MYO19 | 11866.5124 | 0.4622 | ##### | 3.8278 | 1.2929E-04 | 3.4973E-04 |
| ENSG00000090054\|SPTLC1 | 8313.8210 | -0.4521 | ##### | -3.8277 | 1.2933E-04 | 3.4980E-04 |
| ENSG00000162981\|LRATD1 | 27.0282 | 2.4746 | ##### | 3.8266 | 1.2992E-04 | 3.5136E-04 |
| ENSG00000228492\|RAB11FIP1P1 | 106.7512 | 0.7188 | ##### | 3.8265 | 1.2999E-04 | 3.5151E-04 |
| ENSG00000169083\|AR | 190.3081 | -0.8194 | ##### | -3.8263 | 1.3010E-04 | 3.5178E-04 |
| ENSG00000204941\|PSG5 | 31.6689 | -1.5032 | ##### | -3.8262 | 1.3015E-04 | 3.5189E-04 |
| ENSG00000104177\|MYEF2 | 3758.0527 | 0.4784 | ##### | 3.8258 | 1.3036E-04 | 3.5242E-04 |
| ENSG00000197381\|ADARB1 | 1036.3588 | 0.4973 | ##### | 3.8251 | 1.3070E-04 | 3.5331E-04 |
| ENSG00000156928\|MALSU1 | 7210.8579 | 0.4601 | ##### | 3.8246 | 1.3099E-04 | 3.5406E-04 |
| ENSG00000139926\|FRMD6 | 5755.6690 | 0.4761 | ##### | 3.8243 | 1.3115E-04 | 3.5446E-04 |
| ENSG00000163406\|SLC15A2 | 69.8285 | 1.2680 | ##### | 3.8237 | 1.3146E-04 | 3.5527E-04 |
| ENSG00000196141\|SPATS2L | 7302.1724 | -0.3053 | ##### | -3.8235 | 1.3158E-04 | 3.5557E-04 |
| ENSG00000108001\|EBF3 | 95.9305 | 5.0372 | ##### | 3.8234 | 1.3162E-04 | 3.5563E-04 |
| ENSG00000128626\|MRPS12 | 6293.2017 | -0.4532 | ##### | -3.8222 | 1.3229E-04 | 3.5740E-04 |
| ENSG00000091010\|POU4F3 | 11.3629 | 2.7687 | ##### | 3.8215 | 1.3263E-04 | 3.5829E-04 |
| ENSG00000259562 | 23.1351 | 1.7610 | ##### | 3.8210 | 1.3293E-04 | 3.5908E-04 |
| ENSG00000280279\|LINC02887 | 15.5431 | 2.0364 | ##### | 3.8197 | 1.3364E-04 | 3.6096E-04 |
| ENSG00000129562\|DAD1 | 9979.5696 | 0.4702 | ##### | 3.8194 | 1.3379E-04 | 3.6134E-04 |
| ENSG00000165496\|RPL10L | 9.1103 | -3.8850 | ##### | -3.8192 | 1.3387E-04 | 3.6153E-04 |
| ENSG00000284661 | 15.5575 | -1.8175 | ##### | -3.8186 | 1.3419E-04 | 3.6236E-04 |
| ENSG00000169035\|KLK7 | 45.4938 | -7.9275 | ##### | -3.8185 | 1.3429E-04 | 3.6259E-04 |
| ENSG00000184384\|MAML2 | 548.7253 | -0.5565 | ##### | -3.8184 | 1.3434E-04 | 3.6268E-04 |
| ENSG00000198198\|SZT2 | 5318.4943 | -0.4832 | ##### | -3.8176 | 1.3478E-04 | 3.6385E-04 |
| ENSG00000161914\|ZNF653 | 452.4324 | 0.5670 | ##### | 3.8169 | 1.3515E-04 | 3.6481E-04 |
| ENSG00000136536\|MARCHF7 | 9458.3560 | -0.4475 | ##### | -3.8158 | 1.3575E-04 | 3.6638E-04 |
| ENSG00000289697 | 142.9467 | -2.3184 | ##### | -3.8155 | 1.3588E-04 | 3.6671E-04 |
| ENSG00000289603 | 12.3424 | -6.8370 | ##### | -3.8155 | 1.3589E-04 | 3.6672E-04 |
| ENSG00000204947\|ZNF425 | 306.2029 | 0.6684 | ##### | 3.8154 | 1.3597E-04 | 3.6690E-04 |
| ENSG00000090674\|MCOLN1 | 2052.6859 | 0.4894 | ##### | 3.8145 | 1.3643E-04 | 3.6810E-04 |
| ENSG00000248710\|TRIM59-IFT80 | 82.0651 | -1.3162 | ##### | -3.8139 | 1.3681E-04 | 3.6909E-04 |
| ENSG00000011260\|UTP18 | 6479.5113 | 0.4406 | ##### | 3.8138 | 1.3687E-04 | 3.6922E-04 |
| ENSG00000078061\|ARAF | 3636.3678 | -0.4820 | ##### | -3.8125 | 1.3757E-04 | 3.7106E-04 |
| ENSG00000276644\|DACH1 | 26.1117 | 4.6653 | ##### | 3.8115 | 1.3815E-04 | 3.7261E-04 |
| ENSG00000224281\|SLC25A5-AS1 | 269.6079 | 0.9070 | ##### | 3.8113 | 1.3821E-04 | 3.7274E-04 |
| ENSG00000214872\|SMTNL1 | 31.6547 | -7.3245 | ##### | -3.8113 | 1.3824E-04 | 3.7279E-04 |
| ENSG00000104660\|LEPROTL1 | 3286.3990 | -0.4799 | ##### | -3.8112 | 1.3827E-04 | 3.7282E-04 |
| ENSG00000165626\|BEND7 | 727.9078 | -0.5560 | ##### | -3.8109 | 1.3846E-04 | 3.7332E-04 |
| ENSG00000213516\|RBMXL1 | 947.7836 | 0.5093 | ##### | 3.8102 | 1.3885E-04 | 3.7432E-04 |
| ENSG00000283629 | 10.7277 | 4.9439 | ##### | 3.8097 | 1.3915E-04 | 3.7510E-04 |
| ENSG00000005156\|LIG3 | 5499.7302 | 0.4799 | ##### | 3.8096 | 1.3918E-04 | 3.7515E-04 |
| ENSG00000166159\|LRTM2 | 29.0173 | 7.5108 | ##### | 3.8094 | 1.3931E-04 | 3.7546E-04 |
| ENSG00000259623 | 160.8252 | -1.6439 | ##### | -3.8091 | 1.3948E-04 | 3.7589E-04 |
| ENSG00000143033\|MTF2 | 3261.2582 | 0.4651 | ##### | 3.8089 | 1.3960E-04 | 3.7617E-04 |
| ENSG00000139734\|DIAPH3 | 3259.5113 | 0.4665 | ##### | 3.8086 | 1.3977E-04 | 3.7661E-04 |
| ENSG00000134324\|LPIN1 | 2680.0740 | 0.4840 | ##### | 3.8085 | 1.3979E-04 | 3.7663E-04 |
| ENSG00000275074\|NUDT18 | 497.6099 | -0.5661 | ##### | -3.8083 | 1.3990E-04 | 3.7688E-04 |
| ENSG00000089006\|SNX5 | 5216.9084 | 0.3043 | ##### | 3.8051 | 1.4175E-04 | 3.8184E-04 |

| | | | | | | |
|---|---|---|---|---|---|---|
| ENSG00000290997\|ULK4P3 | 60.9716 | -1.1182 | ##### | -3.8048 | 1.4192E-04 | 3.8226E-04 |
| ENSG00000225721 | 46.1607 | 0.9860 | ##### | 3.8036 | 1.4260E-04 | 3.8405E-04 |
| ENSG00000237638\|LINC02245 | 84.3339 | 2.3346 | ##### | 3.8036 | 1.4261E-04 | 3.8405E-04 |
| ENSG00000102543\|CDADC1 | 523.1577 | -0.5730 | ##### | -3.8035 | 1.4268E-04 | 3.8419E-04 |
| ENSG00000274629 | 210.8172 | 0.7145 | ##### | 3.8034 | 1.4275E-04 | 3.8434E-04 |
| ENSG00000183891\|TTC32 | 366.0567 | -0.6090 | ##### | -3.8029 | 1.4302E-04 | 3.8504E-04 |
| ENSG00000157778\|PSMG3 | 5009.2369 | -0.4577 | ##### | -3.8028 | 1.4305E-04 | 3.8509E-04 |
| ENSG00000289554 | 109.6552 | 1.1190 | ##### | 3.8020 | 1.4351E-04 | 3.8630E-04 |
| ENSG00000133265\|HSPBP1 | 6665.5869 | -0.3688 | ##### | -3.8019 | 1.4362E-04 | 3.8654E-04 |
| ENSG00000157916\|RER1 | 11114.3953 | -0.4499 | ##### | -3.8016 | 1.4374E-04 | 3.8683E-04 |
| ENSG00000289716\|XNDC1N-ZNF705EP-ALG1L9I | 75.5902 | 1.7013 | ##### | 3.8013 | 1.4393E-04 | 3.8731E-04 |
| ENSG00000271741 | 159.6835 | -1.1346 | ##### | -3.8008 | 1.4420E-04 | 3.8801E-04 |
| ENSG00000285830 | 1141.7582 | 0.5435 | ##### | 3.8000 | 1.4468E-04 | 3.8925E-04 |
| ENSG00000273275 | 761.3039 | 0.5322 | ##### | 3.7999 | 1.4478E-04 | 3.8951E-04 |
| ENSG00000250742\|LINC02381 | 17.7890 | -1.9597 | ##### | -3.7997 | 1.4486E-04 | 3.8968E-04 |
| ENSG00000286594 | 15.5576 | 6.9277 | ##### | 3.7993 | 1.4508E-04 | 3.9023E-04 |
| ENSG00000170558\|CDH2 | 6408.6804 | -0.2941 | ##### | -3.7989 | 1.4532E-04 | 3.9086E-04 |
| ENSG00000173928\|SWSAP1 | 323.7290 | 0.6231 | ##### | 3.7984 | 1.4566E-04 | 3.9173E-04 |
| ENSG00000126353\|CCR7 | 169.7441 | -9.8281 | ##### | -3.7982 | 1.4576E-04 | 3.9195E-04 |
| ENSG00000196411\|EPHB4 | 6826.4214 | -0.4430 | ##### | -3.7974 | 1.4624E-04 | 3.9322E-04 |
| ENSG00000165802\|NSMF | 7009.1254 | 0.4523 | ##### | 3.7972 | 1.4636E-04 | 3.9350E-04 |
| ENSG00000177575\|CD163 | 15.1808 | 6.5127 | ##### | 3.7971 | 1.4640E-04 | 3.9357E-04 |
| ENSG00000286216 | 17.3026 | -6.6541 | ##### | -3.7955 | 1.4733E-04 | 3.9603E-04 |
| ENSG00000168286\|THAP11 | 5019.7675 | 0.4542 | ##### | 3.7955 | 1.4737E-04 | 3.9610E-04 |
| ENSG00000057757\|PITHD1 | 3048.0650 | 0.4836 | ##### | 3.7951 | 1.4757E-04 | 3.9662E-04 |
| ENSG00000136854\|STXBP1 | 6439.6806 | -0.4666 | ##### | -3.7949 | 1.4769E-04 | 3.9690E-04 |
| ENSG00000163743\|RCHY1 | 1986.2780 | -0.4218 | ##### | -3.7946 | 1.4788E-04 | 3.9737E-04 |
| ENSG00000151778\|SERP2 | 12.1624 | -7.3830 | ##### | -3.7942 | 1.4813E-04 | 3.9801E-04 |
| ENSG00000243629\|LINC00880 | 16.5521 | 3.5536 | ##### | 3.7938 | 1.4838E-04 | 3.9865E-04 |
| ENSG00000245904\|BTG1-DT | 193.2024 | -3.5561 | ##### | -3.7934 | 1.4862E-04 | 3.9925E-04 |
| ENSG00000204946\|ZNF783 | 1778.3016 | 0.4563 | ##### | 3.7931 | 1.4876E-04 | 3.9960E-04 |
| ENSG00000213763\|ACTBP2 | 133.3348 | -0.7034 | ##### | -3.7931 | 1.4878E-04 | 3.9961E-04 |
| ENSG00000155275\|TRMT44 | 903.7141 | 0.5386 | ##### | 3.7930 | 1.4881E-04 | 3.9967E-04 |
| ENSG00000185728\|YTHDF3 | 9353.6470 | -0.4675 | ##### | -3.7924 | 1.4923E-04 | 4.0075E-04 |
| ENSG00000011243\|AKAP8L | 4351.5637 | 0.4624 | ##### | 3.7918 | 1.4954E-04 | 4.0155E-04 |
| ENSG00000105784\|RUNDC3B | 14.6025 | 2.8634 | ##### | 3.7914 | 1.4979E-04 | 4.0217E-04 |
| ENSG00000110435\|PDHX | 2997.9304 | 0.4857 | ##### | 3.7914 | 1.4981E-04 | 4.0220E-04 |
| ENSG00000197457\|STMN3 | 548.2839 | 0.5608 | ##### | 3.7909 | 1.5010E-04 | 4.0294E-04 |
| ENSG00000166342\|NETO1 | 33.3461 | -6.3162 | ##### | -3.7908 | 1.5018E-04 | 4.0311E-04 |
| ENSG00000259305\|ZHX1-C8orf76 | 153.8462 | -0.7999 | ##### | -3.7903 | 1.5049E-04 | 4.0391E-04 |
| ENSG00000267280\|TBX2-AS1 | 42.7590 | -2.5502 | ##### | -3.7902 | 1.5055E-04 | 4.0403E-04 |
| ENSG00000013275\|PSMC4 | 16734.2254 | -0.4423 | ##### | -3.7901 | 1.5060E-04 | 4.0409E-04 |
| ENSG00000259341\|LINC03080 | 24.9465 | -7.5714 | ##### | -3.7901 | 1.5060E-04 | 4.0409E-04 |
| ENSG00000132953\|XPO4 | 5149.0348 | 0.4465 | ##### | 3.7894 | 1.5102E-04 | 4.0520E-04 |
| ENSG00000255100\|TSKU-AS1 | 17.7155 | -3.0503 | ##### | -3.7886 | 1.5150E-04 | 4.0645E-04 |
| ENSG00000120162\|MOB3B | 232.4171 | 0.7219 | ##### | 3.7883 | 1.5166E-04 | 4.0685E-04 |
| ENSG00000187288\|CIDEC | 59.5017 | -5.0905 | ##### | -3.7883 | 1.5169E-04 | 4.0689E-04 |
| ENSG00000080618\|CPB2 | 24.6148 | -4.8576 | ##### | -3.7879 | 1.5192E-04 | 4.0746E-04 |
| ENSG00000117586\|TNFSF4 | 73.5739 | -0.8953 | ##### | -3.7878 | 1.5201E-04 | 4.0767E-04 |
| ENSG00000277945 | 17.0339 | 6.3388 | ##### | 3.7865 | 1.5279E-04 | 4.0973E-04 |
| ENSG00000125434\|SLC25A35 | 1314.9620 | -0.5255 | ##### | -3.7864 | 1.5285E-04 | 4.0984E-04 |
| ENSG00000290044 | 21.1074 | 6.3663 | ##### | 3.7860 | 1.5309E-04 | 4.1046E-04 |
| ENSG00000125375\|DMAC2L | 925.9936 | 0.5029 | ##### | 3.7859 | 1.5314E-04 | 4.1056E-04 |
| ENSG00000188467\|SLC24A5 | 140.0432 | 0.9449 | ##### | 3.7857 | 1.5327E-04 | 4.1086E-04 |
| ENSG00000159842\|ABR | 6502.5645 | -0.4441 | ##### | -3.7851 | 1.5363E-04 | 4.1178E-04 |
| ENSG00000214029\|ZNF891 | 1896.4260 | 0.4748 | ##### | 3.7847 | 1.5387E-04 | 4.1239E-04 |
| ENSG00000158887\|MPZ | 16.4452 | -3.6520 | ##### | -3.7847 | 1.5392E-04 | 4.1249E-04 |
| ENSG00000162526\|TSSK3 | 140.0686 | 0.7269 | ##### | 3.7845 | 1.5401E-04 | 4.1269E-04 |
| ENSG00000163545\|NUAK2 | 3942.7289 | -0.4784 | ##### | -3.7844 | 1.5410E-04 | 4.1289E-04 |
| ENSG00000185475\|TMEM179B | 2238.2806 | -0.4970 | ##### | -3.7838 | 1.5445E-04 | 4.1379E-04 |
| ENSG00000289885 | 27.9657 | 2.4882 | ##### | 3.7832 | 1.5485E-04 | 4.1483E-04 |
| ENSG00000177683\|THAP5 | 2541.8966 | 0.4759 | ##### | 3.7827 | 1.5514E-04 | 4.1559E-04 |
| ENSG00000125743\|SNRPD2 | 12991.0796 | -0.4538 | ##### | -3.7824 | 1.5533E-04 | 4.1606E-04 |

| | | | | | | |
|---|---|---|---|---|---|---|
| ENSG00000275894 | 13.4868 | 3.5709 | ##### | 3.7822 | 1.5544E-04 | 4.1631E-04 |
| ENSG00000110375\|UPK2 | 82.3671 | 1.1390 | ##### | 3.7811 | 1.5614E-04 | 4.1815E-04 |
| ENSG00000237149\|ZNF503-AS2 | 318.0864 | -0.6155 | ##### | -3.7810 | 1.5619E-04 | 4.1825E-04 |
| ENSG00000100341\|PNPLA5 | 44.3558 | 6.4463 | ##### | 3.7808 | 1.5635E-04 | 4.1864E-04 |
| ENSG00000167641\|PPP1R14A | 792.9374 | -0.5505 | ##### | -3.7791 | 1.5736E-04 | 4.2131E-04 |
| ENSG00000293475 | 43.3943 | -1.1533 | ##### | -3.7790 | 1.5748E-04 | 4.2159E-04 |
| ENSG00000117399\|CDC20 | 20044.7862 | 0.4705 | ##### | 3.7783 | 1.5789E-04 | 4.2263E-04 |
| ENSG00000143001\|TMEM61 | 20.9877 | -5.0910 | ##### | -3.7778 | 1.5824E-04 | 4.2353E-04 |
| ENSG00000269973 | 303.4692 | 0.6564 | ##### | 3.7776 | 1.5833E-04 | 4.2374E-04 |
| ENSG00000165209\|STRBP | 3929.9409 | 0.4712 | ##### | 3.7771 | 1.5864E-04 | 4.2454E-04 |
| ENSG00000107242\|PIP5K1B | 17.6594 | -4.1816 | ##### | -3.7771 | 1.5867E-04 | 4.2458E-04 |
| ENSG00000143669\|LYST | 1636.4331 | 0.5148 | ##### | 3.7768 | 1.5887E-04 | 4.2506E-04 |
| ENSG00000166181\|API5 | 12325.4845 | 0.4379 | ##### | 3.7767 | 1.5893E-04 | 4.2521E-04 |
| ENSG00000185614\|INKA1 | 56.7343 | -1.2089 | ##### | -3.7767 | 1.5895E-04 | 4.2521E-04 |
| ENSG00000172901\|LVRN | 49.5803 | -3.5063 | ##### | -3.7762 | 1.5924E-04 | 4.2595E-04 |
| ENSG00000176454\|LPCAT4 | 2607.1878 | -0.4231 | ##### | -3.7759 | 1.5942E-04 | 4.2640E-04 |
| ENSG00000183527\|PSMG1 | 2854.8887 | -0.4839 | ##### | -3.7759 | 1.5946E-04 | 4.2646E-04 |
| ENSG00000213866\|YBX1P10 | 659.5015 | 0.5108 | ##### | 3.7757 | 1.5958E-04 | 4.2673E-04 |
| ENSG00000161664\|ASB16 | 204.4929 | 0.6692 | ##### | 3.7753 | 1.5980E-04 | 4.2728E-04 |
| ENSG00000185504\|FAAP100 | 2674.7380 | -0.4650 | ##### | -3.7746 | 1.6026E-04 | 4.2849E-04 |
| ENSG00000099995\|SF3A1 | 15888.5292 | 0.4600 | ##### | 3.7745 | 1.6031E-04 | 4.2857E-04 |
| ENSG00000237476\|LINC01637 | 27.1146 | -1.3066 | ##### | -3.7741 | 1.6056E-04 | 4.2922E-04 |
| ENSG00000115956\|PLEK | 58.8094 | 7.8257 | ##### | 3.7728 | 1.6145E-04 | 4.3156E-04 |
| ENSG00000231574\|LINC02015 | 42.5054 | -2.5372 | ##### | -3.7718 | 1.6207E-04 | 4.3318E-04 |
| ENSG00000013619\|MAMLD1 | 392.3447 | -0.5602 | ##### | -3.7717 | 1.6212E-04 | 4.3327E-04 |
| ENSG00000255298\|OR8G5 | 44.9012 | -7.9078 | ##### | -3.7714 | 1.6231E-04 | 4.3373E-04 |
| ENSG00000217648\|PARLP2 | 55.2958 | -1.3038 | ##### | -3.7694 | 1.6365E-04 | 4.3728E-04 |
| ENSG00000277249\|MIR6784 | 8.1170 | -2.3812 | ##### | -3.7687 | 1.6413E-04 | 4.3852E-04 |
| ENSG00000117425\|PTCH2 | 60.9181 | -1.0382 | ##### | -3.7683 | 1.6434E-04 | 4.3905E-04 |
| ENSG00000093000\|NUP50 | 7215.2107 | 0.4487 | ##### | 3.7679 | 1.6461E-04 | 4.3971E-04 |
| ENSG00000177463\|NR2C2 | 5051.8587 | 0.6747 | ##### | 3.7670 | 1.6525E-04 | 4.4138E-04 |
| ENSG00000162302\|RPS6KA4 | 6337.2536 | 0.3580 | ##### | 3.7662 | 1.6578E-04 | 4.4269E-04 |
| ENSG00000145241\|CENPC | 1031.9665 | 0.5376 | ##### | 3.7662 | 1.6578E-04 | 4.4269E-04 |
| ENSG00000122507\|BBS9 | 967.6315 | -0.4982 | ##### | -3.7662 | 1.6575E-04 | 4.4269E-04 |
| ENSG00000293440 | 96.6431 | -0.8731 | ##### | -3.7660 | 1.6588E-04 | 4.4292E-04 |
| ENSG00000164932\|CTHRC1 | 164.3756 | 0.5629 | ##### | 3.7659 | 1.6596E-04 | 4.4304E-04 |
| ENSG00000253227 | 21.6171 | -2.1605 | ##### | -3.7659 | 1.6595E-04 | 4.4304E-04 |
| ENSG00000291234 | 20.0333 | 2.0478 | ##### | 3.7658 | 1.6601E-04 | 4.4314E-04 |
| ENSG00000278434 | 23.8951 | 5.3672 | ##### | 3.7655 | 1.6623E-04 | 4.4369E-04 |
| ENSG00000133874\|RNF122 | 192.6889 | -0.6956 | ##### | -3.7654 | 1.6632E-04 | 4.4388E-04 |
| ENSG00000121774\|KHDRBS1 | 18354.2837 | 0.3093 | ##### | 3.7650 | 1.6658E-04 | 4.4455E-04 |
| ENSG00000215012\|RTL10 | 2461.6561 | 0.4763 | ##### | 3.7647 | 1.6674E-04 | 4.4493E-04 |
| ENSG00000251593\|MSNP1 | 46.2224 | -1.2104 | ##### | -3.7637 | 1.6745E-04 | 4.4678E-04 |
| ENSG00000259972 | 530.0929 | -0.5783 | ##### | -3.7630 | 1.6785E-04 | 4.4783E-04 |
| ENSG00000117481\|NSUN4 | 2462.6001 | -0.4664 | ##### | -3.7630 | 1.6789E-04 | 4.4788E-04 |
| ENSG00000168961\|LGALS9 | 30.0012 | -2.7760 | ##### | -3.7628 | 1.6801E-04 | 4.4816E-04 |
| ENSG00000242252\|BGLAP | 20.0879 | 1.5318 | ##### | 3.7625 | 1.6822E-04 | 4.4868E-04 |
| ENSG00000222043 | 41.1790 | 1.2604 | ##### | 3.7624 | 1.6832E-04 | 4.4891E-04 |
| ENSG00000205090\|TMEM240 | 95.3962 | -1.3100 | ##### | -3.7618 | 1.6867E-04 | 4.4981E-04 |
| ENSG00000207445\|SNORD15B | 14.3779 | 1.7823 | ##### | 3.7616 | 1.6884E-04 | 4.5022E-04 |
| ENSG00000267892 | 18.9925 | -3.2625 | ##### | -3.7615 | 1.6892E-04 | 4.5039E-04 |
| ENSG00000224578\|HNRNPA1L3 | 583.8114 | 0.5403 | ##### | 3.7614 | 1.6898E-04 | 4.5050E-04 |
| ENSG00000113407\|TARS1 | 23714.4675 | 0.4385 | ##### | 3.7613 | 1.6902E-04 | 4.5056E-04 |
| ENSG00000207864\|MIR27B | 11.1580 | -5.8860 | ##### | -3.7612 | 1.6912E-04 | 4.5080E-04 |
| ENSG00000159648\|SPMIP8 | 33.5952 | -1.7269 | ##### | -3.7608 | 1.6940E-04 | 4.5150E-04 |
| ENSG00000175611\|ERCC6L2-AS1 | 218.0858 | -0.7705 | ##### | -3.7604 | 1.6963E-04 | 4.5209E-04 |
| ENSG00000149269\|PAK1 | 2281.0856 | -0.4867 | ##### | -3.7601 | 1.6982E-04 | 4.5249E-04 |
| ENSG00000128185\|DGCR6L | 1571.0315 | 0.4910 | ##### | 3.7601 | 1.6981E-04 | 4.5249E-04 |
| ENSG00000164362\|TERT | 778.7339 | 0.5234 | ##### | 3.7601 | 1.6987E-04 | 4.5259E-04 |
| ENSG00000162377\|COA7 | 4853.8492 | -0.4693 | ##### | -3.7600 | 1.6990E-04 | 4.5265E-04 |
| ENSG00000260288 | 24.1103 | 2.3754 | ##### | 3.7583 | 1.7107E-04 | 4.5572E-04 |
| ENSG00000287279 | 21.9433 | 3.8297 | ##### | 3.7583 | 1.7109E-04 | 4.5573E-04 |
| ENSG00000231728\|TMSB15B-AS1 | 47.4265 | 1.6120 | ##### | 3.7578 | 1.7143E-04 | 4.5658E-04 |

| | | | | | | |
|---|---|---|---|---|---|---|
| ENSG00000149929\|HIRIP3 | 1802.9170 | 0.4688 | ##### | 3.7573 | 1.7178E-04 | 4.5749E-04 |
| ENSG00000262413 | 4186.1890 | -0.5599 | ##### | -3.7568 | 1.7212E-04 | 4.5835E-04 |
| ENSG00000123268\|ATF1 | 3714.9732 | 0.4455 | ##### | 3.7566 | 1.7227E-04 | 4.5870E-04 |
| ENSG00000258645\|HSPE1P2 | 52.1979 | 1.2090 | ##### | 3.7563 | 1.7243E-04 | 4.5908E-04 |
| ENSG00000137770\|CTDSPL2 | 3722.1325 | 0.4624 | ##### | 3.7553 | 1.7314E-04 | 4.6094E-04 |
| ENSG00000143315\|PIGM | 3585.8821 | 0.4531 | ##### | 3.7552 | 1.7318E-04 | 4.6102E-04 |
| ENSG00000205810\|KLRC3 | 152.9868 | 3.9990 | ##### | 3.7551 | 1.7327E-04 | 4.6120E-04 |
| ENSG00000197558\|SSPOP | 274.0614 | 0.7697 | ##### | 3.7548 | 1.7347E-04 | 4.6171E-04 |
| ENSG00000164743\|C8orf48 | 65.3016 | 1.0117 | ##### | 3.7544 | 1.7378E-04 | 4.6248E-04 |
| ENSG00000133706\|LARS1 | 13835.1890 | 0.4378 | ##### | 3.7539 | 1.7408E-04 | 4.6324E-04 |
| ENSG00000268655\|SAXO3 | 55.5018 | 1.2397 | ##### | 3.7522 | 1.7531E-04 | 4.6648E-04 |
| ENSG00000224271\|EPIC1 | 193.5225 | 0.6905 | ##### | 3.7516 | 1.7573E-04 | 4.6756E-04 |
| ENSG00000169567\|HINT1 | 10719.9646 | -0.4571 | ##### | -3.7514 | 1.7587E-04 | 4.6787E-04 |
| ENSG00000204706\|MAMDC2-AS1 | 9.8089 | 6.5591 | ##### | 3.7508 | 1.7626E-04 | 4.6886E-04 |
| ENSG00000254598\|CSNK2A3 | 1203.7204 | 0.4813 | ##### | 3.7505 | 1.7645E-04 | 4.6934E-04 |
| ENSG00000286599 | 53.2297 | 2.0888 | ##### | 3.7504 | 1.7657E-04 | 4.6960E-04 |
| ENSG00000134627\|PIWIL4 | 80.4990 | -2.9893 | ##### | -3.7504 | 1.7658E-04 | 4.6960E-04 |
| ENSG00000164172\|MOCS2 | 2501.7345 | -0.4701 | ##### | -3.7499 | 1.7690E-04 | 4.7041E-04 |
| ENSG00000113296\|THBS4 | 41.0481 | 2.5155 | ##### | 3.7497 | 1.7703E-04 | 4.7071E-04 |
| ENSG00000286015 | 42.6308 | -7.8328 | ##### | -3.7494 | 1.7728E-04 | 4.7133E-04 |
| ENSG00000130338\|TULP4 | 1491.1527 | 0.4923 | ##### | 3.7491 | 1.7744E-04 | 4.7172E-04 |
| ENSG00000267105 | 153.0269 | 1.3375 | ##### | 3.7487 | 1.7778E-04 | 4.7257E-04 |
| ENSG00000186481\|ANKRD20A5P | 56.2524 | -9.1637 | ##### | -3.7472 | 1.7882E-04 | 4.7530E-04 |
| ENSG00000140464\|PML | 2197.8286 | -0.4760 | ##### | -3.7446 | 1.8067E-04 | 4.8018E-04 |
| ENSG00000267508\|ZNF285 | 139.5310 | -0.7313 | ##### | -3.7443 | 1.8092E-04 | 4.8080E-04 |
| ENSG00000270872\|SRGAP2D | 684.8824 | 0.5841 | ##### | 3.7441 | 1.8106E-04 | 4.8111E-04 |
| ENSG00000241431\|RPL37P6 | 54.2091 | 1.0834 | ##### | 3.7440 | 1.8110E-04 | 4.8119E-04 |
| ENSG00000166887\|VPS39 | 3941.2415 | -0.4586 | ##### | -3.7436 | 1.8139E-04 | 4.8191E-04 |
| ENSG00000278621\|THBS1-AS1 | 19.6964 | 1.8097 | ##### | 3.7426 | 1.8216E-04 | 4.8392E-04 |
| ENSG00000293405\|CYP2T1P | 48.7511 | -1.2061 | ##### | -3.7418 | 1.8269E-04 | 4.8529E-04 |
| ENSG00000138286\|FAM149B1 | 3429.2441 | -0.5305 | ##### | -3.7418 | 1.8272E-04 | 4.8531E-04 |
| ENSG00000290681\|ECEL1P2 | 13.4584 | 6.3784 | ##### | 3.7417 | 1.8281E-04 | 4.8551E-04 |
| ENSG00000167578\|RAB4B | 721.6771 | -0.7410 | ##### | -3.7414 | 1.8303E-04 | 4.8606E-04 |
| ENSG00000199568\|RNU5A-1 | 9.9755 | 4.5695 | ##### | 3.7413 | 1.8310E-04 | 4.8619E-04 |
| ENSG00000018510\|AGPS | 12595.9980 | -0.4399 | ##### | -3.7411 | 1.8321E-04 | 4.8645E-04 |
| ENSG00000102539\|MLNR | 31.7763 | -1.3666 | ##### | -3.7402 | 1.8386E-04 | 4.8808E-04 |
| ENSG00000291102\|GUSBP1 | 745.8801 | 0.5449 | ##### | 3.7402 | 1.8385E-04 | 4.8808E-04 |
| ENSG00000180113\|TDRD6 | 15.5897 | -4.2579 | ##### | -3.7402 | 1.8389E-04 | 4.8813E-04 |
| ENSG00000270194\|GOLGA4-AS1 | 57.9324 | -1.1333 | ##### | -3.7397 | 1.8425E-04 | 4.8902E-04 |
| ENSG00000087074\|PPP1R15A | 2727.9585 | -0.4808 | ##### | -3.7397 | 1.8426E-04 | 4.8903E-04 |
| ENSG00000271141 | 10.7289 | 2.4965 | ##### | 3.7396 | 1.8429E-04 | 4.8906E-04 |
| ENSG00000213465\|ARL2 | 3408.9760 | 0.4712 | ##### | 3.7392 | 1.8462E-04 | 4.8990E-04 |
| ENSG00000214425\|LRRC37A4P | 395.4965 | -0.6803 | ##### | -3.7390 | 1.8477E-04 | 4.9023E-04 |
| ENSG00000169764\|UGP2 | 7510.1487 | -0.4477 | ##### | -3.7380 | 1.8552E-04 | 4.9218E-04 |
| ENSG00000293034\|NAP1L6P | 40.8982 | -8.7707 | ##### | -3.7375 | 1.8589E-04 | 4.9312E-04 |
| ENSG00000155229\|MMS19 | 6058.4211 | 0.4593 | ##### | 3.7371 | 1.8616E-04 | 4.9379E-04 |
| ENSG00000144040\|SFXN5 | 1790.1725 | -0.4861 | ##### | -3.7342 | 1.8831E-04 | 4.9945E-04 |
| ENSG00000286912 | 20.0602 | -2.4792 | ##### | -3.7321 | 1.8991E-04 | 5.0366E-04 |
| ENSG00000003393\|ALS2 | 5469.1831 | 0.4469 | ##### | 3.7312 | 1.9055E-04 | 5.0530E-04 |
| ENSG00000124279\|FASTKD3 | 1433.1495 | 0.5127 | ##### | 3.7311 | 1.9061E-04 | 5.0543E-04 |
| ENSG00000132581\|SDF2 | 2510.0933 | -0.4545 | ##### | -3.7310 | 1.9070E-04 | 5.0562E-04 |
| ENSG00000130305\|NSUN5 | 3668.6927 | -0.4790 | ##### | -3.7310 | 1.9075E-04 | 5.0570E-04 |
| ENSG00000118689\|FOXO3 | 1818.4616 | 0.4650 | ##### | 3.7307 | 1.9096E-04 | 5.0621E-04 |
| ENSG00000141560\|FN3KRP | 2383.2985 | -0.4782 | ##### | -3.7300 | 1.9151E-04 | 5.0763E-04 |
| ENSG00000287765 | 1718.6416 | 0.5718 | ##### | 3.7283 | 1.9280E-04 | 5.1101E-04 |
| ENSG00000142408\|CACNG8 | 1052.8832 | -0.5098 | ##### | -3.7269 | 1.9382E-04 | 5.1367E-04 |
| ENSG00000095319\|NUP188 | 14338.1906 | 0.4347 | ##### | 3.7267 | 1.9402E-04 | 5.1414E-04 |
| ENSG00000232682 | 10.5895 | -5.0843 | ##### | -3.7266 | 1.9406E-04 | 5.1420E-04 |
| ENSG00000286580 | 18.7332 | -4.0365 | ##### | -3.7263 | 1.9430E-04 | 5.1478E-04 |
| ENSG00000137571\|SLCO5A1 | 86.9945 | 5.2752 | ##### | 3.7262 | 1.9436E-04 | 5.1490E-04 |
| ENSG00000189376\|C8orf76 | 1850.1064 | -0.4687 | ##### | -3.7262 | 1.9441E-04 | 5.1501E-04 |
| ENSG00000166153\|DEPDC4 | 147.8257 | 0.7042 | ##### | 3.7250 | 1.9534E-04 | 5.1741E-04 |
| ENSG00000103544\|VPS35L | 3904.5644 | -0.4486 | ##### | -3.7247 | 1.9554E-04 | 5.1788E-04 |

| | | | | | | |
|---|---|---|---|---|---|---|
| ENSG00000227855\|DPY19L2P3 | 161.4699 | -0.7072 | ##### | -3.7238 | 1.9623E-04 | 5.1967E-04 |
| ENSG00000231084\|RPL22P24 | 20.8112 | -1.6017 | ##### | -3.7237 | 1.9633E-04 | 5.1989E-04 |
| ENSG00000183255\|PTTG1IP | 22413.8497 | -0.3608 | ##### | -3.7236 | 1.9644E-04 | 5.2013E-04 |
| ENSG00000006047\|YBX2 | 2227.9782 | 0.4684 | ##### | 3.7229 | 1.9696E-04 | 5.2147E-04 |
| ENSG00000149531\|FRG1BP | 392.0848 | -0.6159 | ##### | -3.7222 | 1.9751E-04 | 5.2288E-04 |
| ENSG00000174485\|DENND4A | 1660.4083 | -0.4976 | ##### | -3.7215 | 1.9805E-04 | 5.2427E-04 |
| ENSG00000285665\|TOP1P1 | 20.5921 | 1.7253 | ##### | 3.7213 | 1.9823E-04 | 5.2470E-04 |
| ENSG00000169047\|IRS1 | 6461.8067 | 0.4476 | ##### | 3.7212 | 1.9825E-04 | 5.2470E-04 |
| ENSG00000293469\|CHMP1B2P | 364.8716 | 0.7648 | ##### | 3.7210 | 1.9842E-04 | 5.2509E-04 |
| ENSG00000188277\|C15orf62 | 116.9656 | 0.9395 | ##### | 3.7205 | 1.9885E-04 | 5.2620E-04 |
| ENSG00000188051\|TMEM221 | 50.9740 | 1.6881 | ##### | 3.7188 | 2.0013E-04 | 5.2954E-04 |
| ENSG00000204237\|OXLD1 | 1864.2309 | -0.4923 | ##### | -3.7187 | 2.0027E-04 | 5.2987E-04 |
| ENSG00000092036\|HAUS4 | 2162.2122 | 0.4627 | ##### | 3.7185 | 2.0043E-04 | 5.3024E-04 |
| ENSG00000244153\|WWP1P1 | 13.0131 | -2.0183 | ##### | -3.7181 | 2.0074E-04 | 5.3101E-04 |
| ENSG00000168434\|COG7 | 2662.9853 | -0.4682 | ##### | -3.7179 | 2.0088E-04 | 5.3132E-04 |
| ENSG00000235106\|BRD3OS | 1954.9605 | -0.4534 | ##### | -3.7171 | 2.0150E-04 | 5.3291E-04 |
| ENSG00000273335 | 9.7726 | 6.0466 | ##### | 3.7171 | 2.0155E-04 | 5.3301E-04 |
| ENSG00000279800\|BCLAF1P2 | 353.8397 | 0.5883 | ##### | 3.7169 | 2.0169E-04 | 5.3333E-04 |
| ENSG00000004777\|ARHGAP33 | 748.7731 | -0.5392 | ##### | -3.7163 | 2.0219E-04 | 5.3462E-04 |
| ENSG00000091127\|PUS7 | 4455.4673 | 0.4455 | ##### | 3.7161 | 2.0235E-04 | 5.3498E-04 |
| ENSG00000214050\|FBXO16 | 284.4276 | -0.6654 | ##### | -3.7153 | 2.0296E-04 | 5.3654E-04 |
| ENSG00000125954\|CHURC1-FNTB | 135.3274 | 10.4433 | ##### | 3.7152 | 2.0302E-04 | 5.3666E-04 |
| ENSG00000185875\|THNSL1 | 1565.2569 | 0.4323 | ##### | 3.7148 | 2.0333E-04 | 5.3742E-04 |
| ENSG00000235587\|GAPDHP65 | 30.8637 | 1.2002 | ##### | 3.7140 | 2.0401E-04 | 5.3919E-04 |
| ENSG00000066117\|SMARCD1 | 8954.8611 | 0.3690 | ##### | 3.7139 | 2.0410E-04 | 5.3937E-04 |
| ENSG00000040199\|PHLPP2 | 3078.5328 | -0.4626 | ##### | -3.7135 | 2.0439E-04 | 5.4009E-04 |
| ENSG00000260755 | 40.1958 | -1.8799 | ##### | -3.7135 | 2.0441E-04 | 5.4011E-04 |
| ENSG00000180739\|S1PR5 | 774.5702 | -0.5693 | ##### | -3.7127 | 2.0510E-04 | 5.4185E-04 |
| ENSG00000253256 | 28.1450 | 2.8991 | ##### | 3.7126 | 2.0511E-04 | 5.4185E-04 |
| ENSG00000288899 | 31.0452 | -1.3920 | ##### | -3.7122 | 2.0547E-04 | 5.4276E-04 |
| ENSG00000226453\|LINC02542 | 33.9701 | -1.4556 | ##### | -3.7119 | 2.0570E-04 | 5.4332E-04 |
| ENSG00000146281\|PM20D2 | 3506.3560 | 0.4521 | ##### | 3.7116 | 2.0597E-04 | 5.4397E-04 |
| ENSG00000150873\|C2orf50 | 25.5271 | 3.3505 | ##### | 3.7115 | 2.0603E-04 | 5.4405E-04 |
| ENSG00000172354\|GNB2 | 20186.1964 | 0.4499 | ##### | 3.7115 | 2.0603E-04 | 5.4405E-04 |
| ENSG00000149483\|TMEM138 | 1987.3399 | -0.4714 | ##### | -3.7106 | 2.0680E-04 | 5.4602E-04 |
| ENSG00000176896\|TCEANC | 89.8356 | -0.9592 | ##### | -3.7102 | 2.0712E-04 | 5.4683E-04 |
| ENSG00000177640\|CASC2 | 183.1717 | -0.8245 | ##### | -3.7101 | 2.0714E-04 | 5.4683E-04 |
| ENSG00000253144 | 55.9753 | 1.2024 | ##### | 3.7097 | 2.0752E-04 | 5.4778E-04 |
| ENSG00000226421\|SLC25A5P5 | 16.9460 | -2.1105 | ##### | -3.7090 | 2.0810E-04 | 5.4926E-04 |
| ENSG00000269444 | 18.2483 | 4.5275 | ##### | 3.7083 | 2.0867E-04 | 5.5073E-04 |
| ENSG00000134684\|YARS1 | 14815.4814 | -0.3802 | ##### | -3.7082 | 2.0873E-04 | 5.5082E-04 |
| ENSG00000113387\|SUB1 | 14168.9902 | 0.4373 | ##### | 3.7080 | 2.0892E-04 | 5.5129E-04 |
| ENSG00000103769\|RAB11A | 7655.3876 | -0.4356 | ##### | -3.7078 | 2.0907E-04 | 5.5163E-04 |
| ENSG00000114268\|PFKFB4 | 369.2133 | -0.6541 | ##### | -3.7073 | 2.0949E-04 | 5.5268E-04 |
| ENSG00000124226\|RNF114 | 9282.7607 | 0.3498 | ##### | 3.7072 | 2.0956E-04 | 5.5283E-04 |
| ENSG00000066427\|ATXN3 | 891.0122 | 0.5031 | ##### | 3.7072 | 2.0959E-04 | 5.5287E-04 |
| ENSG00000263069\|RNF213-AS1 | 102.9478 | -0.9839 | ##### | -3.7068 | 2.0988E-04 | 5.5356E-04 |
| ENSG00000083937\|CHMP2B | 3686.1810 | -0.4742 | ##### | -3.7065 | 2.1016E-04 | 5.5425E-04 |
| ENSG00000172476\|RAB40A | 134.9198 | -0.7716 | ##### | -3.7062 | 2.1040E-04 | 5.5483E-04 |
| ENSG00000147883\|CDKN2B | 1122.0231 | -0.4862 | ##### | -3.7058 | 2.1073E-04 | 5.5566E-04 |
| ENSG00000105518\|TMEM205 | 8095.9457 | -0.4647 | ##### | -3.7051 | 2.1134E-04 | 5.5721E-04 |
| ENSG00000078098\|FAP | 173.9602 | -0.7965 | ##### | -3.7050 | 2.1136E-04 | 5.5722E-04 |
| ENSG00000124782\|RREB1 | 2880.5124 | 0.4578 | ##### | 3.7049 | 2.1144E-04 | 5.5739E-04 |
| ENSG00000173436\|MICOS10 | 6281.4735 | 0.4563 | ##### | 3.7049 | 2.1149E-04 | 5.5748E-04 |
| ENSG00000165030\|NFIL3 | 1638.1618 | -0.4731 | ##### | -3.7046 | 2.1175E-04 | 5.5809E-04 |
| ENSG00000226261\|PIMREGP1 | 15.8791 | 5.2021 | ##### | 3.7043 | 2.1198E-04 | 5.5868E-04 |
| ENSG00000173137\|ADCK5 | 882.1164 | 0.5234 | ##### | 3.7040 | 2.1223E-04 | 5.5927E-04 |
| ENSG00000160951\|PTGER1 | 37.3509 | -1.7799 | ##### | -3.7033 | 2.1281E-04 | 5.6074E-04 |
| ENSG00000154358\|OBSCN | 1928.8605 | -0.5106 | ##### | -3.7021 | 2.1384E-04 | 5.6342E-04 |
| ENSG00000284669 | 44.8942 | 1.1399 | ##### | 3.7020 | 2.1387E-04 | 5.6344E-04 |
| ENSG00000214182\|PTMAP5 | 26.7769 | 1.6596 | ##### | 3.7016 | 2.1425E-04 | 5.6439E-04 |
| ENSG00000183049\|CAMK1D | 1978.6362 | -0.4803 | ##### | -3.7006 | 2.1509E-04 | 5.6656E-04 |
| ENSG00000188486\|H2AX | 8494.1215 | 0.4453 | ##### | 3.6995 | 2.1601E-04 | 5.6894E-04 |

| | | | | | | |
|---|---|---|---|---|---|---|
| ENSG00000153956\|CACNA2D1 | 5149.7545 | -0.4548 | ##### | -3.6993 | 2.1621E-04 | 5.6940E-04 |
| ENSG00000176732\|PFN4 | 65.9961 | -0.9710 | ##### | -3.6985 | 2.1684E-04 | 5.7102E-04 |
| ENSG00000139910\|NOVA1 | 937.5092 | 0.5021 | ##### | 3.6981 | 2.1718E-04 | 5.7186E-04 |
| ENSG00000008118\|CAMK1G | 17.0896 | -6.4492 | ##### | -3.6973 | 2.1793E-04 | 5.7378E-04 |
| ENSG00000125945\|ZNF436 | 1351.1916 | 0.4897 | ##### | 3.6963 | 2.1873E-04 | 5.7580E-04 |
| ENSG00000288746 | 41.1052 | -1.2920 | ##### | -3.6964 | 2.1872E-04 | 5.7580E-04 |
| ENSG00000260046 | 73.0170 | -8.1488 | ##### | -3.6960 | 2.1905E-04 | 5.7659E-04 |
| ENSG00000234352 | 12.4541 | -5.3142 | ##### | -3.6952 | 2.1970E-04 | 5.7824E-04 |
| ENSG00000120051\|CFAP58 | 55.8555 | 1.8306 | ##### | 3.6946 | 2.2020E-04 | 5.7951E-04 |
| ENSG00000231652\|CD109-AS1 | 85.5812 | 0.9039 | ##### | 3.6944 | 2.2037E-04 | 5.7990E-04 |
| ENSG00000110900\|TSPAN11 | 26.2202 | 2.8532 | ##### | 3.6940 | 2.2074E-04 | 5.8083E-04 |
| ENSG00000279133 | 58.1605 | 1.0641 | ##### | 3.6935 | 2.2115E-04 | 5.8187E-04 |
| ENSG00000261499 | 175.6607 | 0.9904 | ##### | 3.6925 | 2.2205E-04 | 5.8418E-04 |
| ENSG00000240163\|RPL36AP44 | 3.1550 | 4.7995 | ##### | 3.6909 | 2.2349E-04 | 5.8790E-04 |
| ENSG00000197168\|NEK5 | 72.8015 | 0.8478 | ##### | 3.6904 | 2.2391E-04 | 5.8897E-04 |
| ENSG00000279853 | 33.9736 | -7.4734 | ##### | -3.6900 | 2.2427E-04 | 5.8985E-04 |
| ENSG00000225791\|TRAM2-AS1 | 422.9213 | -0.6042 | ##### | -3.6899 | 2.2437E-04 | 5.9006E-04 |
| ENSG00000105409\|ATP1A3 | 642.1345 | -0.5300 | ##### | -3.6897 | 2.2449E-04 | 5.9032E-04 |
| ENSG00000118292\|C1orf54 | 103.9169 | 0.9126 | ##### | 3.6895 | 2.2471E-04 | 5.9086E-04 |
| ENSG00000236814 | 164.5483 | 0.6804 | ##### | 3.6893 | 2.2491E-04 | 5.9133E-04 |
| ENSG00000235863\|B3GALT4 | 41.9195 | 2.4449 | ##### | 3.6887 | 2.2537E-04 | 5.9248E-04 |
| ENSG00000291093\|SVIL-AS1 | 1624.9058 | -0.4751 | ##### | -3.6887 | 2.2541E-04 | 5.9255E-04 |
| ENSG00000243989\|ACY1 | 2109.8752 | 0.4931 | ##### | 3.6885 | 2.2557E-04 | 5.9291E-04 |
| ENSG00000164796\|CSMD3 | 44.4020 | -1.0311 | ##### | -3.6879 | 2.2612E-04 | 5.9429E-04 |
| ENSG00000144231\|POLR2D | 6655.6910 | -0.4056 | ##### | -3.6875 | 2.2647E-04 | 5.9518E-04 |
| ENSG00000278275 | 12.3636 | 3.0048 | ##### | 3.6872 | 2.2677E-04 | 5.9591E-04 |
| ENSG00000265683\|SYPL1P2 | 80.9676 | 1.1811 | ##### | 3.6870 | 2.2690E-04 | 5.9619E-04 |
| ENSG00000253352\|TUG1 | 11833.6704 | -0.4349 | ##### | -3.6870 | 2.2693E-04 | 5.9622E-04 |
| ENSG00000173557\|CIMIP2C | 38.5432 | -8.6849 | ##### | -3.6864 | 2.2741E-04 | 5.9744E-04 |
| ENSG00000185640\|KRT79 | 52.0600 | -8.4499 | ##### | -3.6862 | 2.2759E-04 | 5.9785E-04 |
| ENSG00000149922\|TBX6 | 56.4077 | -1.0916 | ##### | -3.6857 | 2.2806E-04 | 5.9903E-04 |
| ENSG00000224897\|POT1-AS1 | 98.5337 | -0.9326 | ##### | -3.6851 | 2.2862E-04 | 6.0046E-04 |
| ENSG00000203414\|BTBD7P1 | 12.7610 | -5.9570 | ##### | -3.6844 | 2.2922E-04 | 6.0196E-04 |
| ENSG00000203724\|C1orf53 | 193.5021 | -0.6851 | ##### | -3.6842 | 2.2941E-04 | 6.0242E-04 |
| ENSG00000206706\|Y_RNA | 9.8908 | 1.9505 | ##### | 3.6837 | 2.2986E-04 | 6.0354E-04 |
| ENSG00000115307\|AUP1 | 21417.6429 | 0.4280 | ##### | 3.6837 | 2.2992E-04 | 6.0365E-04 |
| ENSG00000169814\|BTD | 1505.8260 | 0.4911 | ##### | 3.6828 | 2.3065E-04 | 6.0552E-04 |
| ENSG00000293160 | 59.5166 | -1.3410 | ##### | -3.6827 | 2.3082E-04 | 6.0591E-04 |
| ENSG00000213658\|LAT | 87.3730 | 1.1373 | ##### | 3.6822 | 2.3123E-04 | 6.0694E-04 |
| ENSG00000142751\|GPN2 | 2212.2983 | 0.4803 | ##### | 3.6805 | 2.3275E-04 | 6.1087E-04 |
| ENSG00000260989 | 56.6813 | -7.9095 | ##### | -3.6798 | 2.3337E-04 | 6.1245E-04 |
| ENSG00000100150\|DEPDC5 | 813.9142 | 0.5018 | ##### | 3.6795 | 2.3373E-04 | 6.1332E-04 |
| ENSG00000293050 | 12.1331 | -6.0886 | ##### | -3.6789 | 2.3426E-04 | 6.1466E-04 |
| ENSG00000104903\|LYL1 | 63.7048 | 1.1210 | ##### | 3.6785 | 2.3459E-04 | 6.1548E-04 |
| ENSG00000074054\|CLASP1 | 5798.9653 | -0.4278 | ##### | -3.6776 | 2.3548E-04 | 6.1777E-04 |
| ENSG00000048471\|SNX29 | 1554.0941 | -0.5033 | ##### | -3.6751 | 2.3776E-04 | 6.2370E-04 |
| ENSG00000110881\|ASIC1 | 712.5672 | 0.5454 | ##### | 3.6751 | 2.3779E-04 | 6.2372E-04 |
| ENSG00000274104 | 51.5888 | -1.2966 | ##### | -3.6743 | 2.3852E-04 | 6.2557E-04 |
| ENSG00000167220\|HDHD2 | 2161.6285 | -0.4730 | ##### | -3.6742 | 2.3858E-04 | 6.2568E-04 |
| ENSG00000134490\|TMEM241 | 1353.9557 | -0.4728 | ##### | -3.6739 | 2.3891E-04 | 6.2649E-04 |
| ENSG00000115461\|IGFBP5 | 106.6921 | -3.7060 | ##### | -3.6736 | 2.3918E-04 | 6.2715E-04 |
| ENSG00000156639\|ZFAND3 | 6403.1089 | -0.4429 | ##### | -3.6735 | 2.3927E-04 | 6.2732E-04 |
| ENSG00000077522\|ACTN2 | 36.4539 | 3.6954 | ##### | 3.6732 | 2.3952E-04 | 6.2791E-04 |
| ENSG00000261222 | 24.5980 | 7.4681 | ##### | 3.6731 | 2.3966E-04 | 6.2824E-04 |
| ENSG00000163605\|PPP4R2 | 3575.9385 | -0.4034 | ##### | -3.6729 | 2.3978E-04 | 6.2850E-04 |
| ENSG00000273311\|DGCR11 | 88.9981 | 1.1842 | ##### | 3.6728 | 2.3994E-04 | 6.2886E-04 |
| ENSG00000137824\|RMDN3 | 2427.8423 | -0.4650 | ##### | -3.6718 | 2.4087E-04 | 6.3123E-04 |
| ENSG00000167195\|GOLGA6C | 9.3419 | 5.8584 | ##### | 3.6716 | 2.4105E-04 | 6.3166E-04 |
| ENSG00000137558\|PI15 | 67.6348 | -1.1224 | ##### | -3.6714 | 2.4123E-04 | 6.3208E-04 |
| ENSG00000163380\|LMOD3 | 29.1438 | -7.1681 | ##### | -3.6710 | 2.4160E-04 | 6.3298E-04 |
| ENSG00000198721\|ECI2 | 4041.1093 | 0.4496 | ##### | 3.6707 | 2.4185E-04 | 6.3357E-04 |
| ENSG00000276771 | 12.4617 | -5.9445 | ##### | -3.6706 | 2.4202E-04 | 6.3397E-04 |
| ENSG00000293465 | 105.5164 | 0.7727 | ##### | 3.6703 | 2.4229E-04 | 6.3463E-04 |

| | | | | | | |
|---|---|---|---|---|---|---|
| ENSG00000291077 | 78.1451 | -1.1264 | ##### | -3.6701 | 2.4242E-04 | 6.3490E-04 |
| ENSG00000214754\|SRSF8CP | 16.7241 | -6.4783 | ##### | -3.6699 | 2.4266E-04 | 6.3548E-04 |
| ENSG00000156603\|MED19 | 1064.2302 | 0.4943 | ##### | 3.6698 | 2.4274E-04 | 6.3564E-04 |
| ENSG00000099797\|TECR | 7952.7209 | 0.4569 | ##### | 3.6696 | 2.4292E-04 | 6.3605E-04 |
| ENSG00000163946\|TASOR | 8337.0335 | -0.4357 | ##### | -3.6694 | 2.4310E-04 | 6.3648E-04 |
| ENSG00000220875\|H3C9P | 7.2511 | 2.5535 | ##### | 3.6687 | 2.4380E-04 | 6.3825E-04 |
| ENSG00000277738 | 26.2244 | -4.7749 | ##### | -3.6678 | 2.4466E-04 | 6.4044E-04 |
| ENSG00000172840\|PDP2 | 1649.6924 | -0.4556 | ##### | -3.6676 | 2.4484E-04 | 6.4073E-04 |
| ENSG00000126453\|BCL2L12 | 4488.6991 | 0.4606 | ##### | 3.6676 | 2.4480E-04 | 6.4073E-04 |
| ENSG00000151967\|SCHIP1 | 134.4255 | 1.5202 | ##### | 3.6676 | 2.4483E-04 | 6.4073E-04 |
| ENSG00000112304\|ACOT13 | 3264.4944 | 0.4317 | ##### | 3.6671 | 2.4533E-04 | 6.4196E-04 |
| ENSG00000197050\|ZNF420 | 689.8536 | -0.9383 | ##### | -3.6671 | 2.4535E-04 | 6.4196E-04 |
| ENSG00000004848\|ARX | 38.1457 | -4.3239 | ##### | -3.6670 | 2.4540E-04 | 6.4205E-04 |
| ENSG00000291208 | 306.9272 | 0.6034 | ##### | 3.6668 | 2.4564E-04 | 6.4261E-04 |
| ENSG00000170949\|ZNF160 | 1535.6180 | -0.4894 | ##### | -3.6665 | 2.4585E-04 | 6.4311E-04 |
| ENSG00000130640\|TUBGCP2 | 6387.7315 | 0.4012 | ##### | 3.6665 | 2.4589E-04 | 6.4315E-04 |
| ENSG00000279302 | 11.8855 | -6.4938 | ##### | -3.6662 | 2.4616E-04 | 6.4380E-04 |
| ENSG00000077147\|TM9SF3 | 19613.4361 | -0.3817 | ##### | -3.6655 | 2.4683E-04 | 6.4550E-04 |
| ENSG00000224424\|PRKAR2A-AS1 | 80.5171 | -0.9096 | ##### | -3.6654 | 2.4694E-04 | 6.4573E-04 |
| ENSG00000196440\|ARMCX4 | 135.3722 | 4.0111 | ##### | 3.6654 | 2.4697E-04 | 6.4575E-04 |
| ENSG00000135040\|NAA35 | 3798.1924 | -0.4515 | ##### | -3.6646 | 2.4776E-04 | 6.4777E-04 |
| ENSG00000269524 | 9.6817 | 6.0495 | ##### | 3.6644 | 2.4794E-04 | 6.4817E-04 |
| ENSG00000117479\|SLC19A2 | 1523.2558 | -0.4682 | ##### | -3.6639 | 2.4843E-04 | 6.4940E-04 |
| ENSG00000279432 | 35.8273 | 6.3446 | ##### | 3.6638 | 2.4850E-04 | 6.4952E-04 |
| ENSG00000101557\|USP14 | 16919.0948 | 0.4482 | ##### | 3.6637 | 2.4858E-04 | 6.4968E-04 |
| ENSG00000148377\|IDI2 | 12.8056 | 1.7549 | ##### | 3.6636 | 2.4870E-04 | 6.4994E-04 |
| ENSG00000140795\|MYLK3 | 36.7720 | -2.5240 | ##### | -3.6632 | 2.4910E-04 | 6.5092E-04 |
| ENSG00000165891\|E2F7 | 1123.1804 | 0.5061 | ##### | 3.6631 | 2.4916E-04 | 6.5102E-04 |
| ENSG00000100242\|SUN2 | 5709.4742 | -0.4423 | ##### | -3.6617 | 2.5057E-04 | 6.5465E-04 |
| ENSG00000289520 | 46.9250 | -1.1187 | ##### | -3.6612 | 2.5099E-04 | 6.5569E-04 |
| ENSG00000187627\|RGPD1 | 40.0953 | -1.3061 | ##### | -3.6610 | 2.5124E-04 | 6.5627E-04 |
| ENSG00000229780\|UBE2Q1-AS1 | 18.2123 | 3.0883 | ##### | 3.6598 | 2.5239E-04 | 6.5924E-04 |
| ENSG00000114654\|EFCC1 | 64.8848 | -1.0415 | ##### | -3.6597 | 2.5252E-04 | 6.5952E-04 |
| ENSG00000229833\|PET100 | 678.4587 | 0.5267 | ##### | 3.6592 | 2.5304E-04 | 6.6081E-04 |
| ENSG00000141959\|PFKL | 8156.9566 | 0.4345 | ##### | 3.6589 | 2.5332E-04 | 6.6149E-04 |
| ENSG00000166619\|BLCAP | 4753.5754 | 0.4273 | ##### | 3.6587 | 2.5349E-04 | 6.6187E-04 |
| ENSG00000259940 | 81.3258 | 0.9385 | ##### | 3.6580 | 2.5422E-04 | 6.6372E-04 |
| ENSG00000280132 | 36.2587 | -1.5980 | ##### | -3.6578 | 2.5439E-04 | 6.6411E-04 |
| ENSG00000290829 | 19.3104 | 3.3768 | ##### | 3.6578 | 2.5441E-04 | 6.6411E-04 |
| ENSG00000132640\|BTBD3 | 2805.5926 | 0.4666 | ##### | 3.6577 | 2.5445E-04 | 6.6414E-04 |
| ENSG00000147606\|SLC26A7 | 20.7260 | -1.9223 | ##### | -3.6576 | 2.5455E-04 | 6.6436E-04 |
| ENSG00000011426\|ANLN | 21905.8801 | 0.4362 | ##### | 3.6574 | 2.5480E-04 | 6.6494E-04 |
| ENSG00000281756\|C2-AS1 | 18.7451 | 5.4523 | ##### | 3.6572 | 2.5503E-04 | 6.6549E-04 |
| ENSG00000270933 | 17.0610 | 5.8154 | ##### | 3.6567 | 2.5544E-04 | 6.6650E-04 |
| ENSG00000291117 | 533.8605 | -0.5873 | ##### | -3.6558 | 2.5641E-04 | 6.6897E-04 |
| ENSG00000178343\|SHISA3 | 29.0180 | 1.9811 | ##### | 3.6557 | 2.5649E-04 | 6.6912E-04 |
| ENSG00000129696\|TTI2 | 1467.6893 | 0.4753 | ##### | 3.6556 | 2.5661E-04 | 6.6937E-04 |
| ENSG00000284842\|FAM91A2P | 34.7665 | 2.0438 | ##### | 3.6544 | 2.5779E-04 | 6.7241E-04 |
| ENSG00000132635\|PCED1A | 1240.4090 | 0.5013 | ##### | 3.6543 | 2.5790E-04 | 6.7261E-04 |
| ENSG00000272288\|ILRUN-AS1 | 56.6841 | -1.3163 | ##### | -3.6532 | 2.5903E-04 | 6.7551E-04 |
| ENSG00000075891\|PAX2 | 48.7441 | 2.2249 | ##### | 3.6529 | 2.5925E-04 | 6.7601E-04 |
| ENSG00000204923\|FBXO48 | 269.3420 | 0.6286 | ##### | 3.6524 | 2.5976E-04 | 6.7729E-04 |
| ENSG00000179912\|R3HDM2 | 4667.1001 | 0.4312 | ##### | 3.6522 | 2.6003E-04 | 6.7794E-04 |
| ENSG00000169379\|ARL13B | 2530.4507 | 0.4678 | ##### | 3.6520 | 2.6022E-04 | 6.7837E-04 |
| ENSG00000106635\|BCL7B | 3728.4563 | -0.4551 | ##### | -3.6519 | 2.6035E-04 | 6.7866E-04 |
| ENSG00000188573\|FBLL1 | 32.2359 | -2.6359 | ##### | -3.6512 | 2.6100E-04 | 6.8028E-04 |
| ENSG00000120071\|KANSL1 | 3356.0337 | 0.4429 | ##### | 3.6507 | 2.6150E-04 | 6.8154E-04 |
| ENSG00000161813\|LARP4 | 13185.5161 | 0.4465 | ##### | 3.6507 | 2.6154E-04 | 6.8157E-04 |
| ENSG00000277235 | 7.5788 | -3.5173 | ##### | -3.6494 | 2.6280E-04 | 6.8481E-04 |
| ENSG00000255248\|MIR100HG | 341.4503 | -0.7384 | ##### | -3.6493 | 2.6294E-04 | 6.8511E-04 |
| ENSG00000277152 | 46.5904 | -6.9162 | ##### | -3.6492 | 2.6307E-04 | 6.8539E-04 |
| ENSG00000260163 | 21.1608 | 4.5113 | ##### | 3.6489 | 2.6338E-04 | 6.8614E-04 |
| ENSG00000276968 | 14.1260 | 6.8588 | ##### | 3.6471 | 2.6523E-04 | 6.9089E-04 |

| | | | | | | |
|---|---|---|---|---|---|---|
| ENSG00000108666\|C17orf75 | 1576.0685 | -0.4662 | ##### | -3.6470 | 2.6534E-04 | 6.9107E-04 |
| ENSG00000163738\|MTHFD2L | 1252.5227 | -0.6188 | ##### | -3.6470 | 2.6533E-04 | 6.9107E-04 |
| ENSG00000272501 | 124.1208 | -0.8795 | ##### | -3.6464 | 2.6592E-04 | 6.9250E-04 |
| ENSG00000163817\|SLC6A20 | 18.9215 | -2.8151 | ##### | -3.6458 | 2.6656E-04 | 6.9412E-04 |
| ENSG00000266101 | 4.7084 | 3.3740 | ##### | 3.6454 | 2.6700E-04 | 6.9519E-04 |
| ENSG00000277462\|ZNF670 | 410.8568 | -0.6613 | ##### | -3.6452 | 2.6722E-04 | 6.9571E-04 |
| ENSG00000165948\|IFI27L1 | 311.6415 | -0.5647 | ##### | -3.6446 | 2.6777E-04 | 6.9707E-04 |
| ENSG00000124334\|IL9R | 31.2606 | -7.2034 | ##### | -3.6441 | 2.6834E-04 | 6.9851E-04 |
| ENSG00000241749\|RPSAP52 | 11.2947 | 3.1050 | ##### | 3.6439 | 2.6857E-04 | 6.9904E-04 |
| ENSG00000154438\|ASZ1 | 23.0957 | 3.8578 | ##### | 3.6436 | 2.6890E-04 | 6.9985E-04 |
| ENSG00000154309\|DISP1 | 655.2186 | -0.5222 | ##### | -3.6433 | 2.6912E-04 | 7.0036E-04 |
| ENSG00000070190\|DAPP1 | 21.8648 | -3.5505 | ##### | -3.6428 | 2.6973E-04 | 7.0189E-04 |
| ENSG00000248830\|ZNF807P | 8.8279 | -2.7334 | ##### | -3.6426 | 2.6993E-04 | 7.0234E-04 |
| ENSG00000174514\|MFSD4A | 40.8720 | -1.6621 | ##### | -3.6423 | 2.7023E-04 | 7.0307E-04 |
| ENSG00000107643\|MAPK8 | 3363.4821 | 0.4407 | ##### | 3.6420 | 2.7055E-04 | 7.0382E-04 |
| ENSG00000184678\|H2BC21 | 2740.4017 | 0.4572 | ##### | 3.6418 | 2.7069E-04 | 7.0413E-04 |
| ENSG00000269713\|NBPF9 | 6869.2037 | 0.4199 | ##### | 3.6416 | 2.7092E-04 | 7.0466E-04 |
| ENSG00000134744\|TUT4 | 4253.2545 | -0.4449 | ##### | -3.6413 | 2.7131E-04 | 7.0562E-04 |
| ENSG00000255099 | 34.4590 | -8.5229 | ##### | -3.6411 | 2.7143E-04 | 7.0587E-04 |
| ENSG00000076555\|ACACB | 1616.1794 | 0.4974 | ##### | 3.6407 | 2.7191E-04 | 7.0701E-04 |
| ENSG00000240477\|EIF3JP2 | 21.3967 | -2.9187 | ##### | -3.6407 | 2.7190E-04 | 7.0701E-04 |
| ENSG00000182742\|HOXB4 | 109.9761 | 5.5056 | ##### | 3.6401 | 2.7253E-04 | 7.0854E-04 |
| ENSG00000173575\|CHD2 | 3409.2500 | -0.4503 | ##### | -3.6400 | 2.7262E-04 | 7.0871E-04 |
| ENSG00000162913\|OBSCN-AS1 | 103.2123 | -0.8591 | ##### | -3.6399 | 2.7271E-04 | 7.0889E-04 |
| ENSG00000170043\|TRAPPC1 | 6734.7912 | 0.4390 | ##### | 3.6397 | 2.7295E-04 | 7.0946E-04 |
| ENSG00000256294\|ZNF225 | 381.4023 | -0.5717 | ##### | -3.6390 | 2.7367E-04 | 7.1126E-04 |
| ENSG00000254602\|MIR130AHG | 35.2910 | 2.0940 | ##### | 3.6387 | 2.7397E-04 | 7.1200E-04 |
| ENSG00000266835\|GAPLINC | 35.2160 | 1.8911 | ##### | 3.6377 | 2.7504E-04 | 7.1469E-04 |
| ENSG00000227953\|LINC01341 | 42.2898 | 1.2168 | ##### | 3.6376 | 2.7522E-04 | 7.1512E-04 |
| ENSG00000228474\|OST4 | 8430.3172 | 0.4441 | ##### | 3.6371 | 2.7576E-04 | 7.1645E-04 |
| ENSG00000153898\|MCOLN2 | 796.0963 | 0.5083 | ##### | 3.6366 | 2.7623E-04 | 7.1760E-04 |
| ENSG00000219747\|RPL32P16 | 18.3626 | 1.4943 | ##### | 3.6364 | 2.7643E-04 | 7.1807E-04 |
| ENSG00000213790\|OLA1P1 | 168.9682 | 0.7487 | ##### | 3.6364 | 2.7650E-04 | 7.1817E-04 |
| ENSG00000290002 | 32.7088 | -3.8925 | ##### | -3.6363 | 2.7658E-04 | 7.1833E-04 |
| ENSG00000237637\|FRY-AS1 | 8.6737 | 5.1320 | ##### | 3.6359 | 2.7697E-04 | 7.1928E-04 |
| ENSG00000157335\|CLEC18C | 13.4350 | -2.6311 | ##### | -3.6357 | 2.7719E-04 | 7.1978E-04 |
| ENSG00000166987\|MBD6 | 4110.7376 | 0.4489 | ##### | 3.6332 | 2.7990E-04 | 7.2677E-04 |
| ENSG00000214870\|LINC02981 | 43.1003 | 7.5743 | ##### | 3.6328 | 2.8035E-04 | 7.2788E-04 |
| ENSG00000278829 | 54.3083 | 7.8598 | ##### | 3.6324 | 2.8083E-04 | 7.2904E-04 |
| ENSG00000290644 | 29.4233 | 6.8768 | ##### | 3.6322 | 2.8107E-04 | 7.2960E-04 |
| ENSG00000089737\|DDX24 | 9464.7805 | 0.4420 | ##### | 3.6317 | 2.8153E-04 | 7.3075E-04 |
| ENSG00000184675\|AMER1 | 1169.9446 | 0.4940 | ##### | 3.6313 | 2.8196E-04 | 7.3179E-04 |
| ENSG00000250486\|FAM218A | 24.9295 | -1.7371 | ##### | -3.6313 | 2.8205E-04 | 7.3196E-04 |
| ENSG00000108506\|INTS2 | 2876.9281 | 0.4463 | ##### | 3.6303 | 2.8307E-04 | 7.3454E-04 |
| ENSG00000108469\|RECQL5 | 2212.6471 | 0.4531 | ##### | 3.6299 | 2.8356E-04 | 7.3575E-04 |
| ENSG00000291005\|NBPF25P | 765.9483 | 0.5350 | ##### | 3.6291 | 2.8441E-04 | 7.3789E-04 |
| ENSG00000170264\|FAM161A | 1484.0580 | 0.4666 | ##### | 3.6288 | 2.8474E-04 | 7.3868E-04 |
| ENSG00000278311\|GGNBP2 | 6011.3386 | -0.4241 | ##### | -3.6282 | 2.8539E-04 | 7.4031E-04 |
| ENSG00000102780\|DGKH | 1541.4226 | -0.4789 | ##### | -3.6270 | 2.8672E-04 | 7.4370E-04 |
| ENSG00000125863\|MKKS | 3177.6903 | -0.4641 | ##### | -3.6269 | 2.8683E-04 | 7.4391E-04 |
| ENSG00000289155 | 31.0099 | 1.5006 | ##### | 3.6266 | 2.8713E-04 | 7.4462E-04 |
| ENSG00000248498\|ASNSP1 | 117.3808 | -9.2953 | ##### | -3.6258 | 2.8810E-04 | 7.4708E-04 |
| ENSG00000170515\|PA2G4 | 37059.2096 | 0.3351 | ##### | 3.6254 | 2.8850E-04 | 7.4805E-04 |
| ENSG00000105737\|GRIK5 | 57.7465 | 0.9750 | ##### | 3.6239 | 2.9017E-04 | 7.5231E-04 |
| ENSG00000087338\|GMCL1 | 2812.9207 | 0.4588 | ##### | 3.6237 | 2.9042E-04 | 7.5290E-04 |
| ENSG00000278385 | 30.5919 | 1.5352 | ##### | 3.6233 | 2.9083E-04 | 7.5390E-04 |
| ENSG00000055732\|MCOLN3 | 276.2124 | 0.6492 | ##### | 3.6228 | 2.9149E-04 | 7.5553E-04 |
| ENSG00000270133 | 5.4617 | 3.0654 | ##### | 3.6225 | 2.9173E-04 | 7.5610E-04 |
| ENSG00000228639\|ROCR | 16.6255 | 6.6189 | ##### | 3.6219 | 2.9246E-04 | 7.5793E-04 |
| ENSG00000163848\|ZNF148 | 4789.7324 | 0.4241 | ##### | 3.6218 | 2.9259E-04 | 7.5819E-04 |
| ENSG00000274265 | 751.1098 | -0.5375 | ##### | -3.6212 | 2.9324E-04 | 7.5982E-04 |
| ENSG00000254703\|SENCR | 58.3716 | 5.9608 | ##### | 3.6211 | 2.9333E-04 | 7.5998E-04 |
| ENSG00000174886\|NDUFA11 | 6600.0221 | 0.4271 | ##### | 3.6210 | 2.9347E-04 | 7.6028E-04 |

| | | | | | | |
|---|---|---|---|---|---|---|
| ENSG00000129933\|MAU2 | 2118.1856 | -0.4476 | ##### | -3.6207 | 2.9386E-04 | 7.6123E-04 |
| ENSG00000273382\|TMEM167B-DT | 221.3191 | -0.6661 | ##### | -3.6204 | 2.9410E-04 | 7.6178E-04 |
| ENSG00000279479 | 10.5297 | -1.9641 | ##### | -3.6196 | 2.9509E-04 | 7.6420E-04 |
| ENSG00000273329 | 304.7351 | 0.5863 | ##### | 3.6196 | 2.9508E-04 | 7.6420E-04 |
| ENSG00000188677\|PARVB | 2423.6252 | -0.4443 | ##### | -3.6191 | 2.9560E-04 | 7.6548E-04 |
| ENSG00000173846\|PLK3 | 658.5086 | 0.5451 | ##### | 3.6186 | 2.9615E-04 | 7.6681E-04 |
| ENSG00000157796\|WDR19 | 1239.4912 | 0.4539 | ##### | 3.6185 | 2.9637E-04 | 7.6731E-04 |
| ENSG00000184319\|RPL23AP82 | 7.2458 | -2.6380 | ##### | -3.6184 | 2.9644E-04 | 7.6744E-04 |
| ENSG00000259845\|HERC2P10 | 30.7792 | 2.0334 | ##### | 3.6184 | 2.9649E-04 | 7.6749E-04 |
| ENSG00000284713\|SMIM38 | 39.2303 | 1.8256 | ##### | 3.6182 | 2.9666E-04 | 7.6788E-04 |
| ENSG00000131944\|FAAP24 | 613.3246 | 0.5062 | ##### | 3.6181 | 2.9676E-04 | 7.6803E-04 |
| ENSG00000231690\|LINC00574 | 24.6848 | 2.2841 | ##### | 3.6181 | 2.9677E-04 | 7.6803E-04 |
| ENSG00000060718\|COL11A1 | 13.2911 | -4.4391 | ##### | -3.6178 | 2.9710E-04 | 7.6881E-04 |
| ENSG00000287974 | 44.0092 | 7.5451 | ##### | 3.6172 | 2.9781E-04 | 7.7059E-04 |
| ENSG00000100225\|FBXO7 | 5268.9021 | -0.4575 | ##### | -3.6156 | 2.9964E-04 | 7.7524E-04 |
| ENSG00000225419\|RPL21P3 | 14.6810 | 6.0659 | ##### | 3.6152 | 3.0009E-04 | 7.7635E-04 |
| ENSG00000104408\|EIF3E | 23503.7264 | -0.4472 | ##### | -3.6151 | 3.0023E-04 | 7.7664E-04 |
| ENSG00000163156\|SCNM1 | 2729.1688 | 0.4552 | ##### | 3.6149 | 3.0050E-04 | 7.7727E-04 |
| ENSG00000162511\|LAPTM5 | 97.8099 | -0.7719 | ##### | -3.6143 | 3.0117E-04 | 7.7895E-04 |
| ENSG00000115211\|EIF2B4 | 4578.1200 | 0.4403 | ##### | 3.6134 | 3.0218E-04 | 7.8149E-04 |
| ENSG00000289043 | 22.5516 | 1.6919 | ##### | 3.6125 | 3.0328E-04 | 7.8427E-04 |
| ENSG00000138663\|COPS4 | 3142.1054 | -0.3990 | ##### | -3.6123 | 3.0352E-04 | 7.8482E-04 |
| ENSG00000182057\|OGFRP1 | 459.5326 | 0.5595 | ##### | 3.6118 | 3.0403E-04 | 7.8606E-04 |
| ENSG00000112584\|FAM120B | 2257.2304 | -0.4289 | ##### | -3.6118 | 3.0410E-04 | 7.8618E-04 |
| ENSG00000198739\|LRRTM3 | 35.4348 | 7.8273 | ##### | 3.6116 | 3.0427E-04 | 7.8655E-04 |
| ENSG00000272416 | 12.5785 | -5.5132 | ##### | -3.6113 | 3.0469E-04 | 7.8757E-04 |
| ENSG00000152402\|GUCY1A2 | 268.6435 | 0.6368 | ##### | 3.6108 | 3.0524E-04 | 7.8892E-04 |
| ENSG00000144597\|EAF1 | 4929.6326 | 0.4400 | ##### | 3.6099 | 3.0634E-04 | 7.9169E-04 |
| ENSG00000230562\|FAM133DP | 97.5236 | 0.8995 | ##### | 3.6098 | 3.0646E-04 | 7.9195E-04 |
| ENSG00000291142 | 396.9646 | -0.5659 | ##### | -3.6089 | 3.0749E-04 | 7.9452E-04 |
| ENSG00000239305\|RNF103 | 3755.0965 | -0.4452 | ##### | -3.6088 | 3.0757E-04 | 7.9468E-04 |
| ENSG00000146021\|KLHL3 | 73.6828 | 3.2042 | ##### | 3.6085 | 3.0792E-04 | 7.9552E-04 |
| ENSG00000259661 | 168.9803 | -0.7686 | ##### | -3.6083 | 3.0824E-04 | 7.9626E-04 |
| ENSG00000262188\|LINC01978 | 20.6470 | -2.9795 | ##### | -3.6081 | 3.0848E-04 | 7.9682E-04 |
| ENSG00000116014\|KISS1R | 101.4366 | 4.2690 | ##### | 3.6079 | 3.0868E-04 | 7.9726E-04 |
| ENSG00000197210 | 30.1654 | -1.4031 | ##### | -3.6063 | 3.1064E-04 | 8.0226E-04 |
| ENSG00000204253\|HNRNPCP2 | 701.3725 | 0.5258 | ##### | 3.6060 | 3.1096E-04 | 8.0301E-04 |
| ENSG00000226445\|THBS2-AS1 | 27.3778 | -1.7023 | ##### | -3.6060 | 3.1100E-04 | 8.0304E-04 |
| ENSG00000289935 | 24.5066 | 1.4766 | ##### | 3.6059 | 3.1110E-04 | 8.0323E-04 |
| ENSG00000223459\|TCAF1P1 | 1746.8096 | 0.4929 | ##### | 3.6051 | 3.1199E-04 | 8.0547E-04 |
| ENSG00000100056\|ESS2 | 1561.1030 | 0.4560 | ##### | 3.6025 | 3.1517E-04 | 8.1360E-04 |
| ENSG00000102804\|TSC22D1 | 7858.9712 | -0.3611 | ##### | -3.6020 | 3.1573E-04 | 8.1498E-04 |
| ENSG00000166923\|GREM1 | 30.8057 | -3.8089 | ##### | -3.6018 | 3.1605E-04 | 8.1572E-04 |
| ENSG00000138587\|MNS1 | 360.9638 | 0.6802 | ##### | 3.6014 | 3.1652E-04 | 8.1687E-04 |
| ENSG00000231764\|DLX6-AS1 | 160.0526 | 0.7638 | ##### | 3.6012 | 3.1678E-04 | 8.1748E-04 |
| ENSG00000188171\|ZNF626 | 11.2907 | -6.9001 | ##### | -3.6010 | 3.1702E-04 | 8.1801E-04 |
| ENSG00000231249\|ITPR1-DT | 48.1530 | -1.2981 | ##### | -3.6009 | 3.1716E-04 | 8.1832E-04 |
| ENSG00000168734\|PKIG | 2076.1907 | 0.4499 | ##### | 3.6006 | 3.1743E-04 | 8.1893E-04 |
| ENSG00000243199\|RPL6P10 | 762.6374 | -0.5180 | ##### | -3.6006 | 3.1746E-04 | 8.1894E-04 |
| ENSG00000130731\|METTL26 | 2810.5440 | 0.4720 | ##### | 3.6000 | 3.1821E-04 | 8.2081E-04 |
| ENSG00000276197 | 11.7125 | 4.6838 | ##### | 3.5996 | 3.1874E-04 | 8.2210E-04 |
| ENSG00000165115\|KIF27 | 469.5581 | -0.5463 | ##### | -3.5994 | 3.1892E-04 | 8.2250E-04 |
| ENSG00000078902\|TOLLIP | 3114.1288 | -0.4368 | ##### | -3.5987 | 3.1978E-04 | 8.2465E-04 |
| ENSG00000204231\|RXRB | 2504.1633 | -0.3356 | ##### | -3.5986 | 3.1990E-04 | 8.2489E-04 |
| ENSG00000246331 | 12.0813 | 4.9280 | ##### | 3.5983 | 3.2032E-04 | 8.2584E-04 |
| ENSG00000287249 | 16.7515 | -2.5923 | ##### | -3.5983 | 3.2031E-04 | 8.2584E-04 |
| ENSG00000279160 | 11.6295 | 3.2152 | ##### | 3.5967 | 3.2229E-04 | 8.3082E-04 |
| ENSG00000100478\|AP4S1 | 607.3595 | 0.5835 | ##### | 3.5960 | 3.2312E-04 | 8.3290E-04 |
| ENSG00000291027\|UBE2Q2P1 | 70.5900 | -1.1683 | ##### | -3.5955 | 3.2375E-04 | 8.3442E-04 |
| ENSG00000082196\|C1QTNF3 | 83.6736 | 0.8933 | ##### | 3.5955 | 3.2377E-04 | 8.3442E-04 |
| ENSG00000289926 | 19.5130 | -2.7479 | ##### | -3.5954 | 3.2388E-04 | 8.3464E-04 |
| ENSG00000144647\|POMGNT2 | 1648.5138 | 0.4657 | ##### | 3.5951 | 3.2430E-04 | 8.3565E-04 |
| ENSG00000160200\|CBS | 3201.3098 | -0.4572 | ##### | -3.5949 | 3.2456E-04 | 8.3625E-04 |

| | | | | | | |
|---|---|---|---|---|---|---|
| ENSG00000225434\|LINC01504 | 41.0863 | -1.0647 | ##### | -3.5947 | 3.2480E-04 | 8.3681E-04 |
| ENSG00000254224\|UQCRB-AS1 | 11.2411 | -3.5127 | ##### | -3.5946 | 3.2486E-04 | 8.3687E-04 |
| ENSG00000130733\|YIPF2 | 3058.3971 | -0.4519 | ##### | -3.5944 | 3.2515E-04 | 8.3756E-04 |
| ENSG00000053438\|NNAT | 54.5647 | 1.0598 | ##### | 3.5936 | 3.2608E-04 | 8.3987E-04 |
| ENSG00000063601\|MTMR1 | 3229.7958 | 0.4495 | ##### | 3.5933 | 3.2647E-04 | 8.4081E-04 |
| ENSG00000272906 | 40.1361 | -1.2417 | ##### | -3.5933 | 3.2657E-04 | 8.4099E-04 |
| ENSG00000272159 | 22.1582 | -3.4095 | ##### | -3.5931 | 3.2675E-04 | 8.4138E-04 |
| ENSG00000213588\|ZBTB9 | 1977.5442 | 0.4342 | ##### | 3.5928 | 3.2716E-04 | 8.4229E-04 |
| ENSG00000226774\|HIKESHIP1 | 14.8789 | 5.7046 | ##### | 3.5928 | 3.2713E-04 | 8.4229E-04 |
| ENSG00000289103 | 25.8416 | 2.9309 | ##### | 3.5925 | 3.2756E-04 | 8.4325E-04 |
| ENSG00000142252\|GEMIN7 | 2368.2332 | -0.6294 | ##### | -3.5923 | 3.2781E-04 | 8.4383E-04 |
| ENSG00000184731\|FAM110C | 86.5460 | -1.0460 | ##### | -3.5914 | 3.2891E-04 | 8.4658E-04 |
| ENSG00000268149 | 48.4977 | 0.8888 | ##### | 3.5911 | 3.2930E-04 | 8.4752E-04 |
| ENSG00000227954\|TARID | 67.7042 | -4.4012 | ##### | -3.5902 | 3.3039E-04 | 8.5025E-04 |
| ENSG00000289327 | 29.8746 | -7.3177 | ##### | -3.5900 | 3.3066E-04 | 8.5086E-04 |
| ENSG00000143379\|SETDB1 | 4269.6985 | 0.4305 | ##### | 3.5900 | 3.3071E-04 | 8.5091E-04 |
| ENSG00000115998\|C2orf42 | 718.6066 | 0.5288 | ##### | 3.5900 | 3.3074E-04 | 8.5093E-04 |
| ENSG00000149357\|LAMTOR1 | 5808.2460 | 0.4435 | ##### | 3.5897 | 3.3108E-04 | 8.5172E-04 |
| ENSG00000226856\|THORLNC | 27.0586 | -3.3548 | ##### | -3.5897 | 3.3111E-04 | 8.5173E-04 |
| ENSG00000102468\|HTR2A | 25.0472 | 4.0360 | ##### | 3.5891 | 3.3187E-04 | 8.5361E-04 |
| ENSG00000287175 | 22.9299 | 1.4679 | ##### | 3.5887 | 3.3237E-04 | 8.5483E-04 |
| ENSG00000278907 | 12.1722 | -6.0144 | ##### | -3.5884 | 3.3265E-04 | 8.5548E-04 |
| ENSG00000010626\|LRRC23 | 1185.4474 | 0.4785 | ##### | 3.5882 | 3.3301E-04 | 8.5632E-04 |
| ENSG00000227888\|FAM66A | 25.0711 | -1.9961 | ##### | -3.5881 | 3.3306E-04 | 8.5637E-04 |
| ENSG00000188766\|SPRED3 | 347.1922 | -0.5983 | ##### | -3.5880 | 3.3319E-04 | 8.5663E-04 |
| ENSG00000173809\|TDRD12 | 28.4317 | 1.6359 | ##### | 3.5879 | 3.3333E-04 | 8.5694E-04 |
| ENSG00000269420\|PLA2G4C-AS1 | 83.2003 | -8.7724 | ##### | -3.5878 | 3.3345E-04 | 8.5709E-04 |
| ENSG00000285967\|NIPBL-DT | 1753.4657 | 0.4531 | ##### | 3.5878 | 3.3345E-04 | 8.5709E-04 |
| ENSG00000266401 | 137.1270 | 0.7399 | ##### | 3.5875 | 3.3389E-04 | 8.5814E-04 |
| ENSG00000169188\|APEX2 | 3364.1157 | 0.4113 | ##### | 3.5871 | 3.3432E-04 | 8.5917E-04 |
| ENSG00000176681\|LRRC37A | 592.8706 | 0.5712 | ##### | 3.5862 | 3.3556E-04 | 8.6229E-04 |
| ENSG00000205867\|KRTAP5-2 | 21.9087 | 2.4121 | ##### | 3.5856 | 3.3633E-04 | 8.6418E-04 |
| ENSG00000254034\|INTS9-AS1 | 25.4812 | 7.0873 | ##### | 3.5847 | 3.3749E-04 | 8.6709E-04 |
| ENSG00000204568\|MRPS18B | 5318.6745 | -0.4361 | ##### | -3.5843 | 3.3798E-04 | 8.6827E-04 |
| ENSG00000085644\|ZNF213 | 1102.9389 | -0.4806 | ##### | -3.5843 | 3.3804E-04 | 8.6837E-04 |
| ENSG00000167740\|CYB5D2 | 418.4850 | -0.5566 | ##### | -3.5841 | 3.3820E-04 | 8.6870E-04 |
| ENSG00000232024\|LSM12P1 | 819.6606 | 0.7709 | ##### | 3.5840 | 3.3834E-04 | 8.6899E-04 |
| ENSG00000254109\|RBPMS-AS1 | 53.4139 | -3.3362 | ##### | -3.5838 | 3.3860E-04 | 8.6957E-04 |
| ENSG00000279329 | 55.5134 | -1.4272 | ##### | -3.5836 | 3.3887E-04 | 8.7018E-04 |
| ENSG00000166747\|AP1G1 | 10980.5779 | -0.4291 | ##### | -3.5826 | 3.4013E-04 | 8.7333E-04 |
| ENSG00000285602 | 22.1744 | -2.8670 | ##### | -3.5826 | 3.4015E-04 | 8.7333E-04 |
| ENSG00000141858\|SAMD1 | 5059.6185 | 0.4415 | ##### | 3.5826 | 3.4023E-04 | 8.7345E-04 |
| ENSG00000179134\|SAMD4B | 10175.9279 | -0.4164 | ##### | -3.5825 | 3.4026E-04 | 8.7346E-04 |
| ENSG00000267424 | 27.4861 | 2.7576 | ##### | 3.5805 | 3.4297E-04 | 8.8033E-04 |
| ENSG00000100314\|CABP7 | 188.2649 | 0.6811 | ##### | 3.5801 | 3.4345E-04 | 8.8150E-04 |
| ENSG00000029639\|TFB1M | 693.9100 | -0.5165 | ##### | -3.5794 | 3.4439E-04 | 8.8383E-04 |
| ENSG00000102879\|CORO1A | 325.8393 | 0.6893 | ##### | 3.5789 | 3.4506E-04 | 8.8548E-04 |
| ENSG00000280145 | 27.8009 | -2.1641 | ##### | -3.5781 | 3.4604E-04 | 8.8793E-04 |
| ENSG00000289367 | 18.4876 | 5.1587 | ##### | 3.5778 | 3.4647E-04 | 8.8893E-04 |
| ENSG00000167985\|SDHAF2 | 1933.6212 | -0.4502 | ##### | -3.5775 | 3.4689E-04 | 8.8994E-04 |
| ENSG00000121210\|TMEM131L | 1088.8188 | -0.4474 | ##### | -3.5760 | 3.4883E-04 | 8.9484E-04 |
| ENSG00000240375\|VPS26AP1 | 11.1614 | -5.1983 | ##### | -3.5755 | 3.4950E-04 | 8.9649E-04 |
| ENSG00000168802\|CHTF8 | 4774.0878 | -0.4863 | ##### | -3.5750 | 3.5021E-04 | 8.9822E-04 |
| ENSG00000119715\|ESRRB | 28.4418 | 1.7078 | ##### | 3.5739 | 3.5175E-04 | 9.0209E-04 |
| ENSG00000099974\|DDTL | 737.0707 | 0.5371 | ##### | 3.5738 | 3.5188E-04 | 9.0236E-04 |
| ENSG00000234171\|RNASEH1-DT | 1602.5102 | 0.4706 | ##### | 3.5723 | 3.5390E-04 | 9.0746E-04 |
| ENSG00000224287\|MSL3P1 | 15.8458 | -1.5035 | ##### | -3.5722 | 3.5404E-04 | 9.0776E-04 |
| ENSG00000114735\|HEMK1 | 925.6851 | -0.4930 | ##### | -3.5716 | 3.5476E-04 | 9.0952E-04 |
| ENSG00000275857 | 128.1673 | 0.6768 | ##### | 3.5705 | 3.5625E-04 | 9.1326E-04 |
| ENSG00000281189\|GHET1 | 39.9889 | -1.4136 | ##### | -3.5705 | 3.5634E-04 | 9.1341E-04 |
| ENSG00000241547\|ACTG1P20 | 34.0065 | 1.1368 | ##### | 3.5699 | 3.5707E-04 | 9.1519E-04 |
| ENSG00000101162\|TUBB1 | 26.7387 | 1.5413 | ##### | 3.5698 | 3.5727E-04 | 9.1564E-04 |
| ENSG00000146066\|HIGD2A | 4425.4489 | 0.4459 | ##### | 3.5698 | 3.5732E-04 | 9.1567E-04 |

| | | | | | | |
|---|---|---|---|---|---|---|
| ENSG00000157800\|SLC37A3 | 4974.0574 | 0.4296 | ##### | 3.5693 | 3.5794E-04 | 9.1719E-04 |
| ENSG00000246863\|GPR176-DT | 15.1101 | -2.5838 | ##### | -3.5692 | 3.5808E-04 | 9.1746E-04 |
| ENSG00000101843\|PSMD10 | 4632.0842 | 0.4391 | ##### | 3.5691 | 3.5826E-04 | 9.1785E-04 |
| ENSG00000123219\|CENPK | 1216.3800 | 0.4795 | ##### | 3.5689 | 3.5843E-04 | 9.1822E-04 |
| ENSG00000256377\|MRPS35-DT | 49.0440 | 1.1066 | ##### | 3.5683 | 3.5929E-04 | 9.2033E-04 |
| ENSG00000122786\|CALD1 | 16788.7131 | 0.4304 | ##### | 3.5679 | 3.5981E-04 | 9.2159E-04 |
| ENSG00000126457\|PRMT1 | 24802.3189 | -0.3267 | ##### | -3.5672 | 3.6084E-04 | 9.2414E-04 |
| ENSG00000138185\|ENTPD1 | 1671.8126 | 1.4174 | ##### | 3.5670 | 3.6114E-04 | 9.2468E-04 |
| ENSG00000254479\|SLC25A1P1 | 10.2779 | 6.2123 | ##### | 3.5670 | 3.6113E-04 | 9.2468E-04 |
| ENSG00000186470\|BTN3A2 | 999.0609 | -0.5074 | ##### | -3.5670 | 3.6112E-04 | 9.2468E-04 |
| ENSG00000138744\|NAAA | 733.4725 | -0.5056 | ##### | -3.5665 | 3.6182E-04 | 9.2634E-04 |
| ENSG00000171121\|KCNMB3 | 231.1083 | -0.6901 | ##### | -3.5664 | 3.6189E-04 | 9.2644E-04 |
| ENSG00000196233\|LCOR | 2988.2903 | 0.5410 | ##### | 3.5659 | 3.6258E-04 | 9.2814E-04 |
| ENSG00000151176\|PLBD2 | 5809.3749 | -0.4424 | ##### | -3.5657 | 3.6290E-04 | 9.2887E-04 |
| ENSG00000228742\|LINC02577 | 42.2884 | -1.4908 | ##### | -3.5655 | 3.6313E-04 | 9.2938E-04 |
| ENSG00000143376\|SNX27 | 5945.1185 | 0.3393 | ##### | 3.5654 | 3.6333E-04 | 9.2981E-04 |
| ENSG00000006282\|SPATA20 | 6469.6150 | -0.4326 | ##### | -3.5653 | 3.6345E-04 | 9.3004E-04 |
| ENSG00000268798 | 35.3822 | 1.4064 | ##### | 3.5653 | 3.6350E-04 | 9.3009E-04 |
| ENSG00000109927\|TECTA | 70.3332 | 2.0131 | ##### | 3.5651 | 3.6365E-04 | 9.3039E-04 |
| ENSG00000139180\|NDUFA9 | 10365.5380 | 0.4112 | ##### | 3.5647 | 3.6428E-04 | 9.3184E-04 |
| ENSG00000099381\|SETD1A | 6193.4580 | 0.4255 | ##### | 3.5647 | 3.6428E-04 | 9.3184E-04 |
| ENSG00000167637\|ZNF283 | 688.0255 | -0.5541 | ##### | -3.5646 | 3.6438E-04 | 9.3201E-04 |
| ENSG00000152433\|ZNF547 | 149.3918 | -0.7292 | ##### | -3.5644 | 3.6465E-04 | 9.3263E-04 |
| ENSG00000114744\|COMMD2 | 3264.4322 | -0.4352 | ##### | -3.5634 | 3.6604E-04 | 9.3609E-04 |
| ENSG00000272430\|LINC02637 | 146.7346 | -0.6721 | ##### | -3.5634 | 3.6607E-04 | 9.3610E-04 |
| ENSG00000186493\|IRX2-DT | 27.7085 | 4.0144 | ##### | 3.5627 | 3.6706E-04 | 9.3855E-04 |
| ENSG00000291262 | 79.7577 | -4.4410 | ##### | -3.5626 | 3.6727E-04 | 9.3900E-04 |
| ENSG00000149571\|KIRREL3 | 17.0157 | -1.5635 | ##### | -3.5618 | 3.6828E-04 | 9.4152E-04 |
| ENSG00000289260 | 53.9596 | -1.4266 | ##### | -3.5615 | 3.6877E-04 | 9.4267E-04 |
| ENSG00000152291\|TGOLN2 | 30468.8857 | 0.4131 | ##### | 3.5614 | 3.6887E-04 | 9.4287E-04 |
| ENSG00000183837\|PNMA3 | 108.0019 | -0.7389 | ##### | -3.5610 | 3.6941E-04 | 9.4415E-04 |
| ENSG00000261584 | 14.9360 | 2.2173 | ##### | 3.5603 | 3.7046E-04 | 9.4677E-04 |
| ENSG00000226580\|RPL39P40 | 12.5795 | 1.9439 | ##### | 3.5601 | 3.7078E-04 | 9.4750E-04 |
| ENSG00000244701 | 71.0044 | -0.9471 | ##### | -3.5600 | 3.7083E-04 | 9.4754E-04 |
| ENSG00000105063\|PPP6R1 | 17663.1517 | 0.4162 | ##### | 3.5591 | 3.7218E-04 | 9.5091E-04 |
| ENSG00000197780\|TAF13 | 8553.7309 | 0.4188 | ##### | 3.5584 | 3.7318E-04 | 9.5337E-04 |
| ENSG00000135801\|TAF5L | 2607.5470 | -0.4503 | ##### | -3.5578 | 3.7401E-04 | 9.5542E-04 |
| ENSG00000207294\|Y_RNA | 8.1294 | 2.3004 | ##### | 3.5574 | 3.7456E-04 | 9.5674E-04 |
| ENSG00000025770\|NCAPH2 | 5886.0167 | 0.4367 | ##### | 3.5571 | 3.7498E-04 | 9.5774E-04 |
| ENSG00000176635\|HORMAD2 | 66.1297 | -5.2945 | ##### | -3.5570 | 3.7515E-04 | 9.5810E-04 |
| ENSG00000275632 | 51.3361 | 1.0025 | ##### | 3.5567 | 3.7553E-04 | 9.5899E-04 |
| ENSG00000226026\|RWDD3-DT | 17.2887 | -4.9402 | ##### | -3.5561 | 3.7639E-04 | 9.6108E-04 |
| ENSG00000230578\|TMLHEP1 | 23.3256 | -7.9149 | ##### | -3.5559 | 3.7675E-04 | 9.6191E-04 |
| ENSG00000134375\|TIMM17A | 8153.5968 | -0.4306 | ##### | -3.5555 | 3.7723E-04 | 9.6307E-04 |
| ENSG00000143222\|UFC1 | 6879.0122 | 0.4314 | ##### | 3.5555 | 3.7729E-04 | 9.6315E-04 |
| ENSG00000136631\|VPS45 | 3208.2338 | 0.4564 | ##### | 3.5554 | 3.7735E-04 | 9.6322E-04 |
| ENSG00000185291\|IL3RA | 22.3181 | -2.9259 | ##### | -3.5552 | 3.7770E-04 | 9.6401E-04 |
| ENSG00000286892 | 24.6881 | -8.0412 | ##### | -3.5550 | 3.7795E-04 | 9.6458E-04 |
| ENSG00000130396\|AFDN | 5676.2121 | -0.4260 | ##### | -3.5546 | 3.7857E-04 | 9.6607E-04 |
| ENSG00000287456 | 9.7134 | 6.3573 | ##### | 3.5544 | 3.7888E-04 | 9.6680E-04 |
| ENSG00000106327\|TFR2 | 1088.3503 | 0.4920 | ##### | 3.5543 | 3.7900E-04 | 9.6702E-04 |
| ENSG00000144730\|IL17RD | 631.1156 | 0.5011 | ##### | 3.5542 | 3.7912E-04 | 9.6723E-04 |
| ENSG00000113396\|SLC27A6 | 6.3229 | -3.3357 | ##### | -3.5540 | 3.7938E-04 | 9.6780E-04 |
| ENSG00000289192 | 27.3763 | -7.1909 | ##### | -3.5539 | 3.7958E-04 | 9.6824E-04 |
| ENSG00000126947\|ARMCX1 | 259.7976 | 0.6476 | ##### | 3.5531 | 3.8077E-04 | 9.7121E-04 |
| ENSG00000242419\|PCDHGC4 | 34.2108 | -7.8267 | ##### | -3.5529 | 3.8107E-04 | 9.7187E-04 |
| ENSG00000176920\|FUT2 | 54.3495 | -0.9790 | ##### | -3.5526 | 3.8151E-04 | 9.7291E-04 |
| ENSG00000152147\|GEMIN6 | 1993.4256 | 0.4705 | ##### | 3.5510 | 3.8372E-04 | 9.7848E-04 |
| ENSG00000128918\|ALDH1A2 | 22.7100 | -1.7961 | ##### | -3.5507 | 3.8428E-04 | 9.7980E-04 |
| ENSG00000161243\|FBXO27 | 9326.1085 | 0.4302 | ##### | 3.5500 | 3.8518E-04 | 9.8202E-04 |
| ENSG00000172765\|TMCC1 | 1636.3063 | -0.4496 | ##### | -3.5495 | 3.8596E-04 | 9.8391E-04 |
| ENSG00000234948\|LINC01524 | 47.5763 | -5.4155 | ##### | -3.5495 | 3.8601E-04 | 9.8397E-04 |
| ENSG00000257242\|LINC01619 | 36.6323 | -1.2407 | ##### | -3.5494 | 3.8615E-04 | 9.8423E-04 |

| | | | | | | |
|---|---|---|---|---|---|---|
| ENSG00000278367 | 119.1074 | -1.2027 | ##### | -3.5492 | 3.8636E-04 | 9.8471E-04 |
| ENSG00000109062\|NHERF1 | 4123.8475 | 0.4294 | ##### | 3.5488 | 3.8698E-04 | 9.8620E-04 |
| ENSG00000007908\|SELE | 49.2794 | 6.7376 | ##### | 3.5483 | 3.8777E-04 | 9.8803E-04 |
| ENSG00000166224\|SGPL1 | 5852.7310 | -0.4354 | ##### | -3.5483 | 3.8775E-04 | 9.8803E-04 |
| ENSG00000164329\|TENT2 | 2390.1042 | 0.4523 | ##### | 3.5482 | 3.8793E-04 | 9.8836E-04 |
| ENSG00000287097 | 24.3895 | -1.9825 | ##### | -3.5479 | 3.8825E-04 | 9.8908E-04 |
| ENSG00000160229\|ZNF66 | 26.2602 | -4.3593 | ##### | -3.5471 | 3.8954E-04 | 9.9228E-04 |
| ENSG00000079332\|SAR1A | 9104.4138 | -0.3595 | ##### | -3.5469 | 3.8973E-04 | 9.9268E-04 |
| ENSG00000279930 | 33.0891 | 1.3812 | ##### | 3.5460 | 3.9107E-04 | 9.9601E-04 |
| ENSG00000155463\|OXA1L | 7436.7732 | 0.4334 | ##### | 3.5459 | 3.9134E-04 | 9.9662E-04 |
| ENSG00000236431 | 11.0051 | 3.1419 | ##### | 3.5448 | 3.9295E-04 | 1.0006E-03 |
| ENSG00000176407\|KCMF1 | 13741.1086 | 0.4248 | ##### | 3.5441 | 3.9402E-04 | 1.0033E-03 |
| ENSG00000178896\|EXOSC4 | 4772.3084 | 0.4345 | ##### | 3.5434 | 3.9506E-04 | 1.0058E-03 |
| ENSG00000106244\|PDAP1 | 14790.8211 | 0.4172 | ##### | 3.5432 | 3.9532E-04 | 1.0064E-03 |
| ENSG00000268621\|IGFL2-AS1 | 26.7148 | -7.1554 | ##### | -3.5429 | 3.9582E-04 | 1.0076E-03 |
| ENSG00000240682\|ISY1 | 3403.3151 | 0.3087 | ##### | 3.5420 | 3.9713E-04 | 1.0108E-03 |
| ENSG00000238000\|PSME2P1 | 54.2738 | 0.9377 | ##### | 3.5419 | 3.9728E-04 | 1.0111E-03 |
| ENSG00000204308\|RNF5 | 3063.1741 | -0.4336 | ##### | -3.5419 | 3.9728E-04 | 1.0111E-03 |
| ENSG00000269621 | 356.5420 | 0.6390 | ##### | 3.5410 | 3.9856E-04 | 1.0142E-03 |
| ENSG00000233452\|STXBP5-AS1 | 27.6881 | -1.7823 | ##### | -3.5410 | 3.9862E-04 | 1.0143E-03 |
| ENSG00000222047\|C10orf55 | 285.6643 | 0.8959 | ##### | 3.5409 | 3.9870E-04 | 1.0144E-03 |
| ENSG00000102390\|PBDC1 | 1795.3923 | 0.4757 | ##### | 3.5405 | 3.9943E-04 | 1.0162E-03 |
| ENSG00000166347\|CYB5A | 3252.2550 | -0.4412 | ##### | -3.5403 | 3.9961E-04 | 1.0166E-03 |
| ENSG00000285614 | 34.5813 | -7.8779 | ##### | -3.5392 | 4.0133E-04 | 1.0208E-03 |
| ENSG00000101608\|MYL12A | 38947.7330 | 0.3570 | ##### | 3.5390 | 4.0160E-04 | 1.0214E-03 |
| ENSG00000288787 | 23.5802 | 3.8876 | ##### | 3.5388 | 4.0193E-04 | 1.0222E-03 |
| ENSG00000147485\|PXDNL | 57.3103 | 6.4723 | ##### | 3.5387 | 4.0209E-04 | 1.0225E-03 |
| ENSG00000160712\|IL6R | 2104.6726 | -0.4441 | ##### | -3.5386 | 4.0233E-04 | 1.0230E-03 |
| ENSG00000149554\|CHEK1 | 4480.9763 | 0.3528 | ##### | 3.5382 | 4.0280E-04 | 1.0241E-03 |
| ENSG00000225399 | 54.1821 | -5.1206 | ##### | -3.5374 | 4.0402E-04 | 1.0271E-03 |
| ENSG00000253174 | 554.5721 | 2.1918 | ##### | 3.5367 | 4.0520E-04 | 1.0301E-03 |
| ENSG00000147955\|SIGMAR1 | 10177.5494 | 0.4293 | ##### | 3.5365 | 4.0542E-04 | 1.0305E-03 |
| ENSG00000260317\|LINC02986 | 42.4007 | 1.9263 | ##### | 3.5364 | 4.0561E-04 | 1.0309E-03 |
| ENSG00000228140\|EFHD2-AS1 | 38.7414 | 1.1999 | ##### | 3.5362 | 4.0597E-04 | 1.0317E-03 |
| ENSG00000244485\|RPL18P13 | 11.3387 | -4.1254 | ##### | -3.5362 | 4.0596E-04 | 1.0317E-03 |
| ENSG00000125835\|SNRPB | 23007.3533 | 0.4214 | ##### | 3.5358 | 4.0657E-04 | 1.0331E-03 |
| ENSG00000242198 | 23.4079 | 4.6655 | ##### | 3.5353 | 4.0735E-04 | 1.0350E-03 |
| ENSG00000176125\|UFSP1 | 299.4502 | -0.5445 | ##### | -3.5352 | 4.0741E-04 | 1.0351E-03 |
| ENSG00000275740 | 71.3833 | 1.5769 | ##### | 3.5350 | 4.0781E-04 | 1.0360E-03 |
| ENSG00000052344\|PRSS8 | 16.9738 | -2.7768 | ##### | -3.5348 | 4.0803E-04 | 1.0365E-03 |
| ENSG00000269038\|SF1-DT | 10.0043 | 3.3775 | ##### | 3.5348 | 4.0809E-04 | 1.0365E-03 |
| ENSG00000116001\|TIA1 | 4346.9996 | 0.3291 | ##### | 3.5342 | 4.0895E-04 | 1.0385E-03 |
| ENSG00000164253\|WDR41 | 3246.3907 | -0.4468 | ##### | -3.5342 | 4.0894E-04 | 1.0385E-03 |
| ENSG00000205097\|FRG2 | 16.0959 | 5.3698 | ##### | 3.5337 | 4.0983E-04 | 1.0407E-03 |
| ENSG00000198146\|ZNF770 | 6178.3563 | 0.4201 | ##### | 3.5335 | 4.1013E-04 | 1.0414E-03 |
| ENSG00000290592 | 37.5105 | 7.2615 | ##### | 3.5333 | 4.1040E-04 | 1.0420E-03 |
| ENSG00000254854\|NECTIN1-DT | 50.5142 | 1.7734 | ##### | 3.5331 | 4.1065E-04 | 1.0425E-03 |
| ENSG00000247137\|ANKRD42-DT | 123.9326 | 0.7769 | ##### | 3.5326 | 4.1142E-04 | 1.0444E-03 |
| ENSG00000226440\|LAMA4-AS1 | 8.0041 | -6.4046 | ##### | -3.5324 | 4.1184E-04 | 1.0454E-03 |
| ENSG00000179532\|DNHD1 | 732.6742 | -0.5127 | ##### | -3.5322 | 4.1204E-04 | 1.0458E-03 |
| ENSG00000129055\|ANAPC13 | 4728.3612 | 0.4242 | ##### | 3.5321 | 4.1220E-04 | 1.0461E-03 |
| ENSG00000263826 | 63.2136 | -1.1210 | ##### | -3.5316 | 4.1312E-04 | 1.0483E-03 |
| ENSG00000082996\|RNF13 | 3632.5220 | -0.4536 | ##### | -3.5313 | 4.1350E-04 | 1.0492E-03 |
| ENSG00000180574\|EIF2S3B | 282.5383 | -0.6231 | ##### | -3.5313 | 4.1356E-04 | 1.0492E-03 |
| ENSG00000153721\|CNKSR3 | 908.8757 | -0.4787 | ##### | -3.5313 | 4.1358E-04 | 1.0492E-03 |
| ENSG00000260233\|ZNRD2-DT | 98.0794 | 0.8132 | ##### | 3.5311 | 4.1384E-04 | 1.0498E-03 |
| ENSG00000257138\|TAS2R38 | 43.4900 | -8.0302 | ##### | -3.5306 | 4.1457E-04 | 1.0516E-03 |
| ENSG00000267325\|LINC01415 | 45.9729 | 8.1438 | ##### | 3.5289 | 4.1726E-04 | 1.0583E-03 |
| ENSG00000236548\|RNF217-AS1 | 37.9476 | -1.1977 | ##### | -3.5288 | 4.1749E-04 | 1.0588E-03 |
| ENSG00000289878 | 37.0730 | -7.3316 | ##### | -3.5283 | 4.1823E-04 | 1.0606E-03 |
| ENSG00000227413 | 6.6391 | 5.3323 | ##### | 3.5271 | 4.2011E-04 | 1.0653E-03 |
| ENSG00000086717\|PPEF1 | 27.3161 | -2.4762 | ##### | -3.5263 | 4.2140E-04 | 1.0684E-03 |
| ENSG00000196235\|SUPT5H | 17575.7141 | -0.4226 | ##### | -3.5253 | 4.2299E-04 | 1.0724E-03 |

| | | | | | | |
|---|---|---|---|---|---|---|
| ENSG00000267577\|DNAAF3-AS1 | 253.5693 | 0.6436 | ##### | 3.5248 | 4.2378E-04 | 1.0743E-03 |
| ENSG00000158526\|TSR2 | 3919.6295 | 0.4456 | ##### | 3.5245 | 4.2422E-04 | 1.0753E-03 |
| ENSG00000104442\|ARMC1 | 5289.6664 | -0.4239 | ##### | -3.5236 | 4.2567E-04 | 1.0789E-03 |
| ENSG00000048707\|VPS13D | 2799.2505 | -0.4621 | ##### | -3.5234 | 4.2603E-04 | 1.0797E-03 |
| ENSG00000170385\|SLC30A1 | 4955.4886 | -0.4176 | ##### | -3.5230 | 4.2672E-04 | 1.0813E-03 |
| ENSG00000196396\|PTPN1 | 12423.5626 | 0.4098 | ##### | 3.5230 | 4.2671E-04 | 1.0813E-03 |
| ENSG00000290958 | 8.1858 | 2.5890 | ##### | 3.5227 | 4.2713E-04 | 1.0822E-03 |
| ENSG00000120675\|DNAJC15 | 2970.8846 | -0.4482 | ##### | -3.5209 | 4.3012E-04 | 1.0897E-03 |
| ENSG00000169288\|MRPL1 | 2122.3479 | 0.4427 | ##### | 3.5195 | 4.3233E-04 | 1.0952E-03 |
| ENSG00000250775 | 20.0075 | -4.5940 | ##### | -3.5183 | 4.3432E-04 | 1.1002E-03 |
| ENSG00000277559 | 141.5652 | 0.6526 | ##### | 3.5173 | 4.3593E-04 | 1.1042E-03 |
| ENSG00000106004\|HOXA5 | 255.8178 | 0.5729 | ##### | 3.5169 | 4.3655E-04 | 1.1056E-03 |
| ENSG00000234743\|EIF5AP4 | 727.8218 | 0.5083 | ##### | 3.5168 | 4.3684E-04 | 1.1063E-03 |
| ENSG00000148343\|MIGA2 | 1480.6476 | -0.4578 | ##### | -3.5166 | 4.3718E-04 | 1.1070E-03 |
| ENSG00000165502\|RPL36AL | 6678.1925 | 0.4311 | ##### | 3.5164 | 4.3743E-04 | 1.1076E-03 |
| ENSG00000273343 | 18.9087 | 2.4989 | ##### | 3.5163 | 4.3763E-04 | 1.1079E-03 |
| ENSG00000280187 | 271.5279 | -0.5837 | ##### | -3.5163 | 4.3764E-04 | 1.1079E-03 |
| ENSG00000242221\|PSG2 | 24.9847 | -7.4642 | ##### | -3.5154 | 4.3903E-04 | 1.1113E-03 |
| ENSG00000139626\|ITGB7 | 29.2805 | -1.3924 | ##### | -3.5153 | 4.3931E-04 | 1.1120E-03 |
| ENSG00000156265\|MAP3K7CL | 197.4309 | -0.6537 | ##### | -3.5151 | 4.3953E-04 | 1.1124E-03 |
| ENSG00000145569\|OTULINL | 410.3400 | -0.5138 | ##### | -3.5150 | 4.3976E-04 | 1.1129E-03 |
| ENSG00000125814\|NAPB | 619.5935 | -0.5073 | ##### | -3.5148 | 4.4003E-04 | 1.1135E-03 |
| ENSG00000118094\|TREH | 110.8361 | 0.7777 | ##### | 3.5148 | 4.4013E-04 | 1.1137E-03 |
| ENSG00000196796\|NPIPB10P | 40.9148 | 1.2694 | ##### | 3.5144 | 4.4070E-04 | 1.1150E-03 |
| ENSG00000087448\|KLHL42 | 5798.1004 | 0.4257 | ##### | 3.5141 | 4.4120E-04 | 1.1161E-03 |
| ENSG00000242759\|LINC00882 | 57.3466 | 2.5694 | ##### | 3.5141 | 4.4119E-04 | 1.1161E-03 |
| ENSG00000224843\|LINC00240 | 60.7633 | 1.0854 | ##### | 3.5139 | 4.4152E-04 | 1.1168E-03 |
| ENSG00000163644\|PPM1K | 383.6840 | -0.5478 | ##### | -3.5136 | 4.4203E-04 | 1.1180E-03 |
| ENSG00000166025\|AMOTL1 | 3207.0203 | 0.4226 | ##### | 3.5133 | 4.4252E-04 | 1.1191E-03 |
| ENSG00000162512\|SDC3 | 3954.6539 | -0.4295 | ##### | -3.5128 | 4.4346E-04 | 1.1214E-03 |
| ENSG00000129244\|ATP1B2 | 17.2269 | 3.1972 | ##### | 3.5127 | 4.4363E-04 | 1.1216E-03 |
| ENSG00000108587\|GOSR1 | 6382.2379 | -0.4184 | ##### | -3.5127 | 4.4359E-04 | 1.1216E-03 |
| ENSG00000169087\|HSPBAP1 | 671.7399 | 0.5434 | ##### | 3.5127 | 4.4358E-04 | 1.1216E-03 |
| ENSG00000137168\|PPIL1 | 3409.6148 | -0.4336 | ##### | -3.5124 | 4.4412E-04 | 1.1227E-03 |
| ENSG00000289222 | 53.1090 | 0.9529 | ##### | 3.5113 | 4.4587E-04 | 1.1270E-03 |
| ENSG00000138442\|WDR12 | 7270.2582 | 0.2584 | ##### | 3.5094 | 4.4906E-04 | 1.1350E-03 |
| ENSG00000066651\|TRMT11 | 1453.8481 | 0.4439 | ##### | 3.5092 | 4.4952E-04 | 1.1361E-03 |
| ENSG00000235706\|DICER1-AS1 | 71.8902 | -0.9286 | ##### | -3.5089 | 4.4992E-04 | 1.1370E-03 |
| ENSG00000153140\|CETN3 | 1195.3697 | 0.4793 | ##### | 3.5089 | 4.5002E-04 | 1.1371E-03 |
| ENSG00000245857\|GS1-24F4.2 | 39.3160 | 3.8232 | ##### | 3.5076 | 4.5221E-04 | 1.1426E-03 |
| ENSG00000170633\|RNF34 | 3270.7727 | 0.4473 | ##### | 3.5067 | 4.5376E-04 | 1.1464E-03 |
| ENSG00000167193\|CRK | 11843.5122 | 0.4056 | ##### | 3.5066 | 4.5394E-04 | 1.1468E-03 |
| ENSG00000243433 | 23.6151 | 2.1035 | ##### | 3.5063 | 4.5439E-04 | 1.1478E-03 |
| ENSG00000237188 | 164.5802 | -5.0342 | ##### | -3.5055 | 4.5568E-04 | 1.1510E-03 |
| ENSG00000241404\|EGFL8 | 162.1998 | 0.7070 | ##### | 3.5036 | 4.5909E-04 | 1.1595E-03 |
| ENSG00000239779\|WBP1 | 3098.4350 | -0.4183 | ##### | -3.5028 | 4.6046E-04 | 1.1628E-03 |
| ENSG00000283293\|RN7SK | 145.4817 | 0.9856 | ##### | 3.5026 | 4.6079E-04 | 1.1636E-03 |
| ENSG00000255310 | 100.0198 | 0.8318 | ##### | 3.5025 | 4.6086E-04 | 1.1636E-03 |
| ENSG00000099937\|SERPIND1 | 27.9665 | 6.9010 | ##### | 3.5024 | 4.6103E-04 | 1.1640E-03 |
| ENSG00000220201\|ZGLP1 | 89.8790 | -0.8214 | ##### | -3.5017 | 4.6228E-04 | 1.1670E-03 |
| ENSG00000076984\|MAP2K7 | 4747.6451 | 0.4178 | ##### | 3.5005 | 4.6435E-04 | 1.1722E-03 |
| ENSG00000166165\|CKB | 10572.5594 | -0.4199 | ##### | -3.5003 | 4.6469E-04 | 1.1729E-03 |
| ENSG00000250379 | 11.5385 | 2.0468 | ##### | 3.5000 | 4.6528E-04 | 1.1743E-03 |
| ENSG00000227617\|CERS6-AS1 | 926.8161 | -3.2614 | ##### | -3.4998 | 4.6559E-04 | 1.1750E-03 |
| ENSG00000286548 | 43.7440 | 3.7664 | ##### | 3.4996 | 4.6601E-04 | 1.1759E-03 |
| ENSG00000254485 | 33.0766 | -1.7823 | ##### | -3.4995 | 4.6619E-04 | 1.1763E-03 |
| ENSG00000227110\|LMCD1-AS1 | 55.7016 | -1.4090 | ##### | -3.4985 | 4.6790E-04 | 1.1805E-03 |
| ENSG00000237356 | 63.9754 | 1.0743 | ##### | 3.4982 | 4.6833E-04 | 1.1815E-03 |
| ENSG00000287296 | 47.3920 | 1.4793 | ##### | 3.4980 | 4.6869E-04 | 1.1823E-03 |
| ENSG00000224356 | 24.0159 | 3.1128 | ##### | 3.4979 | 4.6902E-04 | 1.1830E-03 |
| ENSG00000241307\|RPL7AP24 | 14.5655 | -6.1989 | ##### | -3.4978 | 4.6906E-04 | 1.1831E-03 |
| ENSG00000261731 | 13.6456 | -3.1397 | ##### | -3.4975 | 4.6966E-04 | 1.1845E-03 |
| ENSG00000057657\|PRDM1 | 560.7015 | 0.5244 | ##### | 3.4973 | 4.6992E-04 | 1.1850E-03 |

| | | | | | | |
|---|---|---|---|---|---|---|
| ENSG00000258761 | 10.3873 | 2.9482 | ##### | 3.4972 | 4.7017E-04 | 1.1856E-03 |
| ENSG00000287445 | 18.8787 | 1.4996 | ##### | 3.4968 | 4.7085E-04 | 1.1872E-03 |
| ENSG00000255929\|PIWIL4-AS1 | 40.3947 | -1.4302 | ##### | -3.4967 | 4.7101E-04 | 1.1875E-03 |
| ENSG00000291283 | 19.0053 | 1.6696 | ##### | 3.4962 | 4.7194E-04 | 1.1897E-03 |
| ENSG00000078124\|ACER3 | 4700.4929 | 0.4689 | ##### | 3.4956 | 4.7300E-04 | 1.1922E-03 |
| ENSG00000138434\|ITPRID2 | 16119.4580 | 0.4029 | ##### | 3.4956 | 4.7300E-04 | 1.1922E-03 |
| ENSG00000267764 | 40.2367 | 5.8278 | ##### | 3.4950 | 4.7411E-04 | 1.1949E-03 |
| ENSG00000181625\|SLX1B | 2018.3897 | 0.4454 | ##### | 3.4943 | 4.7530E-04 | 1.1978E-03 |
| ENSG00000122299\|ZC3H7A | 3938.4548 | -0.4185 | ##### | -3.4940 | 4.7577E-04 | 1.1988E-03 |
| ENSG00000134899\|ERCC5 | 1657.2813 | -0.4497 | ##### | -3.4939 | 4.7602E-04 | 1.1994E-03 |
| ENSG00000148737\|TCF7L2 | 1902.7006 | -0.4484 | ##### | -3.4938 | 4.7618E-04 | 1.1997E-03 |
| ENSG00000135049\|AGTPBP1 | 1824.8231 | -0.4398 | ##### | -3.4933 | 4.7702E-04 | 1.2017E-03 |
| ENSG00000205363\|INSYN1 | 72.4846 | -0.9611 | ##### | -3.4933 | 4.7717E-04 | 1.2020E-03 |
| ENSG00000272717 | 14.0556 | -4.2076 | ##### | -3.4928 | 4.7798E-04 | 1.2039E-03 |
| ENSG00000181097\|LINC02878 | 93.1016 | 0.9976 | ##### | 3.4928 | 4.7801E-04 | 1.2039E-03 |
| ENSG00000128482\|RNF112 | 33.4445 | 1.2066 | ##### | 3.4920 | 4.7941E-04 | 1.2073E-03 |
| ENSG00000288859\|H2AC19 | 1590.8441 | -0.4461 | ##### | -3.4919 | 4.7955E-04 | 1.2076E-03 |
| ENSG00000101181\|MTG2 | 3527.5831 | 0.3602 | ##### | 3.4916 | 4.8013E-04 | 1.2089E-03 |
| ENSG00000117016\|RIMS3 | 761.6121 | -0.5087 | ##### | -3.4915 | 4.8038E-04 | 1.2094E-03 |
| ENSG00000035687\|ADSS2 | 4607.1235 | -0.4416 | ##### | -3.4913 | 4.8072E-04 | 1.2102E-03 |
| ENSG00000233818\|CLDN14-AS1 | 80.3782 | -0.8830 | ##### | -3.4905 | 4.8206E-04 | 1.2135E-03 |
| ENSG00000119608\|PROX2 | 15.5525 | 2.3519 | ##### | 3.4899 | 4.8323E-04 | 1.2163E-03 |
| ENSG00000100764\|PSMC1 | 19293.0623 | 0.4280 | ##### | 3.4890 | 4.8483E-04 | 1.2202E-03 |
| ENSG00000217094\|PPIAP31 | 90.0632 | -0.6509 | ##### | -3.4888 | 4.8523E-04 | 1.2212E-03 |
| ENSG00000235978 | 28.5617 | -1.2803 | ##### | -3.4887 | 4.8533E-04 | 1.2213E-03 |
| ENSG00000116691\|MIIP | 2216.9151 | 0.4505 | ##### | 3.4883 | 4.8608E-04 | 1.2231E-03 |
| ENSG00000289301 | 10.8754 | 6.0557 | ##### | 3.4871 | 4.8825E-04 | 1.2284E-03 |
| ENSG00000105427\|CNFN | 141.4805 | -0.7753 | ##### | -3.4863 | 4.8979E-04 | 1.2322E-03 |
| ENSG00000235865\|GSN-AS1 | 30.2616 | -2.2511 | ##### | -3.4861 | 4.9015E-04 | 1.2330E-03 |
| ENSG00000144619\|CNTN4 | 26.2384 | -1.4330 | ##### | -3.4860 | 4.9036E-04 | 1.2334E-03 |
| ENSG00000127527\|EPS15L1 | 2656.7259 | 0.4307 | ##### | 3.4859 | 4.9057E-04 | 1.2339E-03 |
| ENSG00000148700\|ADD3 | 2540.2267 | 0.4539 | ##### | 3.4855 | 4.9130E-04 | 1.2356E-03 |
| ENSG00000148200\|NR6A1 | 363.1469 | 0.5292 | ##### | 3.4847 | 4.9269E-04 | 1.2390E-03 |
| ENSG00000087502\|ERGIC2 | 8301.6361 | 0.4072 | ##### | 3.4843 | 4.9348E-04 | 1.2409E-03 |
| ENSG00000289102 | 87.6027 | -0.8859 | ##### | -3.4842 | 4.9369E-04 | 1.2412E-03 |
| ENSG00000289296 | 45.0004 | 1.1921 | ##### | 3.4842 | 4.9369E-04 | 1.2412E-03 |
| ENSG00000270806\|C17orf50 | 19.4534 | -2.6387 | ##### | -3.4838 | 4.9436E-04 | 1.2428E-03 |
| ENSG00000287916 | 83.3622 | -0.8366 | ##### | -3.4835 | 4.9492E-04 | 1.2441E-03 |
| ENSG00000265750 | 147.2336 | -0.7343 | ##### | -3.4832 | 4.9537E-04 | 1.2451E-03 |
| ENSG00000196209\|SIRPB2 | 44.4595 | -8.4061 | ##### | -3.4831 | 4.9562E-04 | 1.2456E-03 |
| ENSG00000258399\|MIR493HG | 20.4784 | 3.9819 | ##### | 3.4828 | 4.9626E-04 | 1.2471E-03 |
| ENSG00000197024\|ZNF398 | 1792.6625 | 0.4524 | ##### | 3.4826 | 4.9666E-04 | 1.2480E-03 |
| ENSG00000198420\|TCAF1 | 5770.6726 | 0.4246 | ##### | 3.4822 | 4.9726E-04 | 1.2494E-03 |
| ENSG00000197301\|HMGA2-AS1 | 16.9847 | 3.9556 | ##### | 3.4819 | 4.9783E-04 | 1.2508E-03 |
| ENSG00000112599\|GUCA1B | 59.5304 | -0.9466 | ##### | -3.4806 | 5.0025E-04 | 1.2567E-03 |
| ENSG00000286214\|COPG2IT1 | 26.5093 | 1.6350 | ##### | 3.4799 | 5.0160E-04 | 1.2600E-03 |
| ENSG00000100218\|RSPH14 | 38.2939 | -1.1535 | ##### | -3.4798 | 5.0177E-04 | 1.2603E-03 |
| ENSG00000139291\|TMEM19 | 4249.6732 | 0.4268 | ##### | 3.4797 | 5.0199E-04 | 1.2608E-03 |
| ENSG00000196387\|ZNF140 | 1161.8800 | -0.4972 | ##### | -3.4794 | 5.0253E-04 | 1.2620E-03 |
| ENSG00000156802\|ATAD2 | 12562.9988 | 0.4048 | ##### | 3.4786 | 5.0397E-04 | 1.2655E-03 |
| ENSG00000101546\|RBFA | 2206.4707 | 0.4374 | ##### | 3.4782 | 5.0482E-04 | 1.2676E-03 |
| ENSG00000285854 | 105.3824 | 1.6517 | ##### | 3.4763 | 5.0833E-04 | 1.2763E-03 |
| ENSG00000293330 | 231.0046 | -0.5832 | ##### | -3.4761 | 5.0875E-04 | 1.2772E-03 |
| ENSG00000206562\|METTL6 | 1052.1531 | 0.4569 | ##### | 3.4758 | 5.0940E-04 | 1.2787E-03 |
| ENSG00000174740\|PABPC5 | 35.3562 | -5.6245 | ##### | -3.4756 | 5.0972E-04 | 1.2794E-03 |
| ENSG00000131116\|ZNF428 | 3626.3668 | -0.4253 | ##### | -3.4755 | 5.0995E-04 | 1.2799E-03 |
| ENSG00000108588\|CCDC47 | 15201.7091 | 0.4088 | ##### | 3.4754 | 5.1005E-04 | 1.2801E-03 |
| ENSG00000184304\|PRKD1 | 477.1921 | -0.5550 | ##### | -3.4734 | 5.1389E-04 | 1.2896E-03 |
| ENSG00000198650\|TAT | 33.9440 | -2.8967 | ##### | -3.4726 | 5.1536E-04 | 1.2932E-03 |
| ENSG00000288762 | 32.6367 | 3.4984 | ##### | 3.4720 | 5.1666E-04 | 1.2963E-03 |
| ENSG00000213939 | 15.6596 | -1.8319 | ##### | -3.4717 | 5.1708E-04 | 1.2973E-03 |
| ENSG00000285520 | 10.5757 | 4.2064 | ##### | 3.4715 | 5.1752E-04 | 1.2981E-03 |
| ENSG00000183444\|OR7E38P | 448.6777 | 1.0518 | ##### | 3.4715 | 5.1752E-04 | 1.2981E-03 |

| | | | | | | |
|---|---|---|---|---|---|---|
| ENSG00000187672\|ERC2 | 49.6542 | 2.1998 | ##### | 3.4710 | 5.1843E-04 | 1.3003E-03 |
| ENSG00000197894\|ADH5 | 10186.2914 | -0.4338 | ##### | -3.4710 | 5.1857E-04 | 1.3006E-03 |
| ENSG00000160838\|LRRC71 | 38.5411 | 1.6503 | ##### | 3.4703 | 5.1986E-04 | 1.3037E-03 |
| ENSG00000104219\|ZDHHC2 | 3121.6574 | 0.4635 | ##### | 3.4698 | 5.2078E-04 | 1.3059E-03 |
| ENSG00000156414\|TDRD9 | 25.1903 | -2.3288 | ##### | -3.4696 | 5.2124E-04 | 1.3069E-03 |
| ENSG00000082805\|ERC1 | 10914.8114 | 0.4173 | ##### | 3.4692 | 5.2199E-04 | 1.3087E-03 |
| ENSG00000132326\|PER2 | 923.8617 | -0.4640 | ##### | -3.4687 | 5.2292E-04 | 1.3109E-03 |
| ENSG00000284830 | 34.2408 | 1.2089 | ##### | 3.4685 | 5.2337E-04 | 1.3119E-03 |
| ENSG00000269151 | 40.6777 | 1.1810 | ##### | 3.4682 | 5.2391E-04 | 1.3132E-03 |
| ENSG00000224020\|MIR181A2HG | 61.0376 | -1.2212 | ##### | -3.4682 | 5.2402E-04 | 1.3134E-03 |
| ENSG00000178033\|CALHM5 | 60.1407 | 1.0142 | ##### | 3.4679 | 5.2449E-04 | 1.3144E-03 |
| ENSG00000228919 | 6.6125 | -4.0757 | ##### | -3.4675 | 5.2538E-04 | 1.3165E-03 |
| ENSG00000272734\|ADIRF-AS1 | 508.3609 | 0.5109 | ##### | 3.4674 | 5.2557E-04 | 1.3169E-03 |
| ENSG00000175264\|CHST1 | 41.9403 | 3.8843 | ##### | 3.4673 | 5.2562E-04 | 1.3169E-03 |
| ENSG00000051825\|MPHOSPH9 | 3052.4617 | 0.4292 | ##### | 3.4673 | 5.2577E-04 | 1.3172E-03 |
| ENSG00000141639\|MAPK4 | 24.0591 | 3.5017 | ##### | 3.4672 | 5.2595E-04 | 1.3175E-03 |
| ENSG00000138594\|TMOD3 | 11894.5485 | -0.4123 | ##### | -3.4670 | 5.2628E-04 | 1.3182E-03 |
| ENSG00000272994\|C2orf49-DT | 260.3661 | -0.6046 | ##### | -3.4669 | 5.2650E-04 | 1.3187E-03 |
| ENSG00000274092 | 39.4890 | -1.1205 | ##### | -3.4667 | 5.2698E-04 | 1.3198E-03 |
| ENSG00000143797\|MBOAT2 | 2369.9774 | 0.4606 | ##### | 3.4665 | 5.2733E-04 | 1.3205E-03 |
| ENSG00000198855\|FICD | 639.0581 | -0.4928 | ##### | -3.4660 | 5.2823E-04 | 1.3227E-03 |
| ENSG00000179915\|NRXN1 | 29.7465 | 2.0872 | ##### | 3.4657 | 5.2880E-04 | 1.3240E-03 |
| ENSG00000267554 | 11.0701 | 5.6029 | ##### | 3.4652 | 5.2987E-04 | 1.3266E-03 |
| ENSG00000219410 | 649.4407 | 0.8456 | ##### | 3.4644 | 5.3136E-04 | 1.3302E-03 |
| ENSG00000135372\|NAT10 | 6573.1214 | 0.4008 | ##### | 3.4639 | 5.3238E-04 | 1.3326E-03 |
| ENSG00000108733\|PEX12 | 595.0289 | -0.5110 | ##### | -3.4638 | 5.3253E-04 | 1.3329E-03 |
| ENSG00000275468 | 17.9962 | 2.3310 | ##### | 3.4627 | 5.3487E-04 | 1.3386E-03 |
| ENSG00000164211\|STARD4 | 1610.4053 | 0.4747 | ##### | 3.4624 | 5.3547E-04 | 1.3400E-03 |
| ENSG00000168395\|ING5 | 4365.2136 | 0.4356 | ##### | 3.4622 | 5.3582E-04 | 1.3408E-03 |
| ENSG00000147166\|ITGB1BP2 | 41.7932 | 1.7249 | ##### | 3.4619 | 5.3635E-04 | 1.3420E-03 |
| ENSG00000117595\|IRF6 | 59.8780 | 2.5561 | ##### | 3.4618 | 5.3652E-04 | 1.3423E-03 |
| ENSG00000149927\|DOC2A | 412.6530 | 0.5329 | ##### | 3.4615 | 5.3726E-04 | 1.3441E-03 |
| ENSG00000224645 | 54.6736 | 1.0111 | ##### | 3.4614 | 5.3739E-04 | 1.3443E-03 |
| ENSG00000260302\|LINC01882 | 17.4885 | 6.0481 | ##### | 3.4612 | 5.3784E-04 | 1.3453E-03 |
| ENSG00000231327\|LINC01816 | 138.6223 | 0.6810 | ##### | 3.4605 | 5.3917E-04 | 1.3484E-03 |
| ENSG00000124767\|GLO1 | 17010.9315 | 0.4130 | ##### | 3.4605 | 5.3917E-04 | 1.3484E-03 |
| ENSG00000290455 | 11.1365 | 2.2686 | ##### | 3.4601 | 5.3989E-04 | 1.3501E-03 |
| ENSG00000188921\|HACD4 | 19.5397 | -4.4124 | ##### | -3.4595 | 5.4117E-04 | 1.3531E-03 |
| ENSG00000051009\|FHIP1B | 1646.5687 | 0.4373 | ##### | 3.4589 | 5.4234E-04 | 1.3560E-03 |
| ENSG00000223653\|BCL10-AS1 | 23.9336 | 1.5242 | ##### | 3.4589 | 5.4241E-04 | 1.3560E-03 |
| ENSG00000114423\|CBLB | 986.7241 | -0.4622 | ##### | -3.4588 | 5.4256E-04 | 1.3563E-03 |
| ENSG00000180964\|TCEAL8 | 2162.4169 | 0.4343 | ##### | 3.4587 | 5.4277E-04 | 1.3567E-03 |
| ENSG00000288905\|LSP1P5 | 338.2621 | 0.6301 | ##### | 3.4583 | 5.4353E-04 | 1.3585E-03 |
| ENSG00000244036\|UBE2H-DT | 9.1259 | 3.2679 | ##### | 3.4582 | 5.4376E-04 | 1.3590E-03 |
| ENSG00000160298\|C21orf58 | 717.7342 | 0.4621 | ##### | 3.4574 | 5.4547E-04 | 1.3631E-03 |
| ENSG00000254042\|SLIT3-AS2 | 35.0265 | 6.4564 | ##### | 3.4573 | 5.4564E-04 | 1.3634E-03 |
| ENSG00000077150\|NFKB2 | 23170.6310 | 0.3218 | ##### | 3.4572 | 5.4584E-04 | 1.3638E-03 |
| ENSG00000181924\|COA4 | 5201.2972 | 0.4195 | ##### | 3.4566 | 5.4711E-04 | 1.3668E-03 |
| ENSG00000224138 | 28.9852 | 1.5359 | ##### | 3.4562 | 5.4794E-04 | 1.3688E-03 |
| ENSG00000259439\|LINC01833 | 143.0105 | 0.7827 | ##### | 3.4556 | 5.4909E-04 | 1.3716E-03 |
| ENSG00000284194\|SCO2 | 1140.2622 | -0.5092 | ##### | -3.4555 | 5.4918E-04 | 1.3717E-03 |
| ENSG00000174353\|STAG3L3 | 13.2960 | 1.9691 | ##### | 3.4551 | 5.4999E-04 | 1.3736E-03 |
| ENSG00000179044\|EXOC3L1 | 25.6118 | -2.3907 | ##### | -3.4543 | 5.5165E-04 | 1.3776E-03 |
| ENSG00000197386\|HTT | 6392.8637 | 0.4147 | ##### | 3.4542 | 5.5198E-04 | 1.3783E-03 |
| ENSG00000290292\|HOMEZ | 1571.1234 | 0.4561 | ##### | 3.4541 | 5.5214E-04 | 1.3786E-03 |
| ENSG00000223358\|EHHADH-AS1 | 5.1485 | -2.8191 | ##### | -3.4540 | 5.5229E-04 | 1.3789E-03 |
| ENSG00000289375 | 13.5444 | 3.2522 | ##### | 3.4537 | 5.5302E-04 | 1.3806E-03 |
| ENSG00000166477\|LEO1 | 2932.9634 | 0.4151 | ##### | 3.4535 | 5.5333E-04 | 1.3813E-03 |
| ENSG00000277501 | 33.8105 | -6.9095 | ##### | -3.4526 | 5.5521E-04 | 1.3858E-03 |
| ENSG00000188580\|NKAIN2 | 84.7142 | 1.0772 | ##### | 3.4524 | 5.5569E-04 | 1.3869E-03 |
| ENSG00000198087\|CD2AP | 9537.5043 | 0.4186 | ##### | 3.4521 | 5.5624E-04 | 1.3882E-03 |
| ENSG00000151229\|SLC2A13 | 1109.1030 | -0.4814 | ##### | -3.4510 | 5.5849E-04 | 1.3935E-03 |
| ENSG00000279518 | 28.8244 | 2.5057 | ##### | 3.4510 | 5.5848E-04 | 1.3935E-03 |

| | | | | | | |
|---|---|---|---|---|---|---|
| ENSG00000122482\|ZNF644 | 4954.7364 | 0.4244 | ##### | 3.4495 | 5.6172E-04 | 1.4015E-03 |
| ENSG00000293172 | 33.8026 | 1.7726 | ##### | 3.4480 | 5.6467E-04 | 1.4087E-03 |
| ENSG00000293069 | 59.2972 | 9.2395 | ##### | 3.4476 | 5.6559E-04 | 1.4109E-03 |
| ENSG00000260257 | 153.7055 | 0.9887 | ##### | 3.4468 | 5.6721E-04 | 1.4148E-03 |
| ENSG00000259523 | 48.8935 | -1.1271 | ##### | -3.4467 | 5.6751E-04 | 1.4155E-03 |
| ENSG00000235652\|EPM2A-DT | 84.4417 | -1.6232 | ##### | -3.4457 | 5.6955E-04 | 1.4204E-03 |
| ENSG00000253372 | 8.2044 | 5.8055 | ##### | 3.4457 | 5.6962E-04 | 1.4205E-03 |
| ENSG00000082213\|C5orf22 | 4435.0445 | -0.4198 | ##### | -3.4455 | 5.7010E-04 | 1.4216E-03 |
| ENSG00000226964\|RHEBP2 | 1383.9614 | 0.4385 | ##### | 3.4453 | 5.7038E-04 | 1.4221E-03 |
| ENSG00000225125\|RANP4 | 12.6018 | 2.5397 | ##### | 3.4450 | 5.7098E-04 | 1.4235E-03 |
| ENSG00000066279\|ASPM | 6754.2568 | 0.4203 | ##### | 3.4449 | 5.7128E-04 | 1.4241E-03 |
| ENSG00000223873\|SAP18P2 | 9.0446 | 2.4340 | ##### | 3.4443 | 5.7258E-04 | 1.4273E-03 |
| ENSG00000197279\|ZNF165 | 121.8908 | 0.9500 | ##### | 3.4439 | 5.7337E-04 | 1.4291E-03 |
| ENSG00000272695\|GAS6-DT | 31.8211 | -1.3104 | ##### | -3.4438 | 5.7360E-04 | 1.4296E-03 |
| ENSG00000272716\|SLC30A6-DT | 43.8642 | -1.4058 | ##### | -3.4434 | 5.7446E-04 | 1.4316E-03 |
| ENSG00000270607\|LEISA1 | 20.0365 | -2.0910 | ##### | -3.4434 | 5.7454E-04 | 1.4317E-03 |
| ENSG00000115760\|BIRC6 | 6397.5595 | 0.4319 | ##### | 3.4429 | 5.7543E-04 | 1.4338E-03 |
| ENSG00000104892\|KLC3 | 351.4615 | -1.0061 | ##### | -3.4426 | 5.7606E-04 | 1.4352E-03 |
| ENSG00000164603\|BMT2 | 821.2347 | -0.4934 | ##### | -3.4426 | 5.7620E-04 | 1.4355E-03 |
| ENSG00000263072\|ZNF213-AS1 | 446.9278 | -0.5239 | ##### | -3.4425 | 5.7630E-04 | 1.4356E-03 |
| ENSG00000196646\|ZNF136 | 535.1858 | 0.5263 | ##### | 3.4425 | 5.7634E-04 | 1.4356E-03 |
| ENSG00000236756\|DNAJC9-AS1 | 259.6829 | -0.6176 | ##### | -3.4424 | 5.7660E-04 | 1.4361E-03 |
| ENSG00000185046\|ANKS1B | 176.7964 | 0.7873 | ##### | 3.4423 | 5.7681E-04 | 1.4365E-03 |
| ENSG00000214108\|TPT1P5 | 6.3845 | -3.3389 | ##### | -3.4423 | 5.7688E-04 | 1.4365E-03 |
| ENSG00000197771\|MCMBP | 9477.2798 | 0.4281 | ##### | 3.4421 | 5.7726E-04 | 1.4374E-03 |
| ENSG00000135506\|OS9 | 24295.9881 | -0.4095 | ##### | -3.4419 | 5.7756E-04 | 1.4380E-03 |
| ENSG00000234390\|USP27X-DT | 122.5213 | -0.8021 | ##### | -3.4419 | 5.7771E-04 | 1.4383E-03 |
| ENSG00000157823\|AP3S2 | 2571.7969 | -0.4277 | ##### | -3.4418 | 5.7788E-04 | 1.4386E-03 |
| ENSG00000215795 | 24.4711 | -7.4890 | ##### | -3.4410 | 5.7948E-04 | 1.4424E-03 |
| ENSG00000106400\|ZNHIT1 | 6142.3360 | 0.4197 | ##### | 3.4410 | 5.7966E-04 | 1.4427E-03 |
| ENSG00000141013\|GAS8 | 2414.8842 | -0.4472 | ##### | -3.4409 | 5.7978E-04 | 1.4429E-03 |
| ENSG00000166278\|C2 | 54.0442 | -0.9171 | ##### | -3.4404 | 5.8076E-04 | 1.4453E-03 |
| ENSG00000275022\|MIR6753 | 17.9023 | 1.5529 | ##### | 3.4402 | 5.8124E-04 | 1.4463E-03 |
| ENSG00000272182 | 29.6367 | 2.0933 | ##### | 3.4401 | 5.8150E-04 | 1.4469E-03 |
| ENSG00000165633\|VSTM4 | 38.5353 | 1.3520 | ##### | 3.4388 | 5.8432E-04 | 1.4537E-03 |
| ENSG00000149476\|TKFC | 3307.9993 | -0.4107 | ##### | -3.4386 | 5.8467E-04 | 1.4545E-03 |
| ENSG00000135097\|MSI1 | 98.7077 | 2.1005 | ##### | 3.4382 | 5.8553E-04 | 1.4565E-03 |
| ENSG00000139874\|SSTR1 | 52.1132 | -2.2190 | ##### | -3.4375 | 5.8721E-04 | 1.4606E-03 |
| ENSG00000141337\|ARSG | 233.2855 | 0.8703 | ##### | 3.4373 | 5.8749E-04 | 1.4611E-03 |
| ENSG00000226243\|RPL37AP1 | 113.5309 | -0.6941 | ##### | -3.4370 | 5.8818E-04 | 1.4627E-03 |
| ENSG00000102001\|CACNA1F | 16.9915 | 3.1458 | ##### | 3.4365 | 5.8929E-04 | 1.4654E-03 |
| ENSG00000187555\|USP7 | 11950.3895 | 0.4040 | ##### | 3.4363 | 5.8965E-04 | 1.4661E-03 |
| ENSG00000227544\|LINC03013 | 12.4988 | -6.5663 | ##### | -3.4362 | 5.9004E-04 | 1.4670E-03 |
| ENSG00000279070 | 16.7928 | 5.6230 | ##### | 3.4358 | 5.9087E-04 | 1.4689E-03 |
| ENSG00000170832\|USP32 | 6366.8890 | 0.4294 | ##### | 3.4349 | 5.9267E-04 | 1.4733E-03 |
| ENSG00000227014\|FKBP14-AS1 | 100.9507 | -0.8030 | ##### | -3.4349 | 5.9278E-04 | 1.4734E-03 |
| ENSG00000233429\|HOTAIRM1 | 168.3355 | 0.7612 | ##### | 3.4349 | 5.9284E-04 | 1.4735E-03 |
| ENSG00000187867\|PALM3 | 313.2900 | 0.5390 | ##### | 3.4346 | 5.9337E-04 | 1.4747E-03 |
| ENSG00000116273\|PHF13 | 2766.1949 | 0.3306 | ##### | 3.4343 | 5.9407E-04 | 1.4763E-03 |
| ENSG00000276234\|TADA2A | 1778.9956 | 0.4390 | ##### | 3.4342 | 5.9426E-04 | 1.4766E-03 |
| ENSG00000051180\|RAD51 | 1541.5854 | 0.4599 | ##### | 3.4326 | 5.9788E-04 | 1.4855E-03 |
| ENSG00000214145\|LINC00887 | 37.2506 | -1.1227 | ##### | -3.4325 | 5.9806E-04 | 1.4858E-03 |
| ENSG00000271643\|PDCD6IP-DT | 259.3784 | -0.5742 | ##### | -3.4325 | 5.9815E-04 | 1.4859E-03 |
| ENSG00000100567\|PSMA3 | 9357.2403 | 0.4353 | ##### | 3.4324 | 5.9823E-04 | 1.4860E-03 |
| ENSG00000230551 | 3433.5756 | -0.4847 | ##### | -3.4322 | 5.9871E-04 | 1.4871E-03 |
| ENSG00000290555\|LINC00680 | 538.7534 | -0.5020 | ##### | -3.4319 | 5.9927E-04 | 1.4883E-03 |
| ENSG00000148154\|UGCG | 6063.2579 | -0.3953 | ##### | -3.4313 | 6.0069E-04 | 1.4917E-03 |
| ENSG00000165495\|PKNOX2 | 6.9316 | 5.5470 | ##### | 3.4299 | 6.0389E-04 | 1.4996E-03 |
| ENSG00000235043\|TECRP1 | 87.7443 | 0.7008 | ##### | 3.4297 | 6.0434E-04 | 1.5006E-03 |
| ENSG00000182545\|RNASE10 | 68.6790 | -0.8519 | ##### | -3.4296 | 6.0440E-04 | 1.5006E-03 |
| ENSG00000262652 | 10.1183 | 5.9300 | ##### | 3.4295 | 6.0459E-04 | 1.5009E-03 |
| ENSG00000229419\|RALGAPA1P1 | 122.8135 | 0.8155 | ##### | 3.4287 | 6.0654E-04 | 1.5056E-03 |
| ENSG00000102572\|STK24 | 6950.6327 | -0.4428 | ##### | -3.4282 | 6.0760E-04 | 1.5081E-03 |

| | | | | | | |
|---|---|---|---|---|---|---|
| ENSG00000242852\|ZNF709 | 60.7952 | -1.1165 | ##### | -3.4281 | 6.0775E-04 | 1.5084E-03 |
| ENSG00000286675 | 19.8802 | -7.7266 | ##### | -3.4279 | 6.0832E-04 | 1.5097E-03 |
| ENSG00000264349\|MIR4258 | 13.8018 | 1.7119 | ##### | 3.4278 | 6.0859E-04 | 1.5102E-03 |
| ENSG00000178607\|ERN1 | 2358.5373 | -0.4347 | ##### | -3.4275 | 6.0913E-04 | 1.5114E-03 |
| ENSG00000282851\|BISPR | 37.2750 | -5.9243 | ##### | -3.4274 | 6.0934E-04 | 1.5118E-03 |
| ENSG00000271781 | 57.8220 | 0.8657 | ##### | 3.4256 | 6.1340E-04 | 1.5218E-03 |
| ENSG00000130382\|MLLT1 | 6793.2336 | 0.4234 | ##### | 3.4240 | 6.1713E-04 | 1.5309E-03 |
| ENSG00000205922\|ONECUT3 | 44.5678 | 7.1932 | ##### | 3.4236 | 6.1793E-04 | 1.5328E-03 |
| ENSG00000112659\|CUL9 | 2318.4412 | 0.4358 | ##### | 3.4233 | 6.1872E-04 | 1.5346E-03 |
| ENSG00000176723\|ZNF843 | 36.0582 | -1.1162 | ##### | -3.4232 | 6.1881E-04 | 1.5347E-03 |
| ENSG00000249937\|LINC02223 | 57.9435 | -5.7594 | ##### | -3.4231 | 6.1917E-04 | 1.5354E-03 |
| ENSG00000230216\|HSPB1P2 | 16.1261 | 1.8648 | ##### | 3.4229 | 6.1966E-04 | 1.5365E-03 |
| ENSG00000273204 | 25.4750 | -1.4396 | ##### | -3.4219 | 6.2178E-04 | 1.5417E-03 |
| ENSG00000198899\|MT-ATP6 | ########## | 0.4262 | ##### | 3.4217 | 6.2229E-04 | 1.5428E-03 |
| ENSG00000116213\|WRAP73 | 2314.3552 | 0.4444 | ##### | 3.4217 | 6.2236E-04 | 1.5429E-03 |
| ENSG00000167118\|URM1 | 4867.6556 | -0.4064 | ##### | -3.4207 | 6.2451E-04 | 1.5481E-03 |
| ENSG00000233937\|CTC-338M12.4 | 495.9035 | 0.5440 | ##### | 3.4207 | 6.2458E-04 | 1.5481E-03 |
| ENSG00000064115\|TM7SF3 | 9694.5595 | 0.3142 | ##### | 3.4206 | 6.2487E-04 | 1.5487E-03 |
| ENSG00000269981 | 17.7407 | 2.1945 | ##### | 3.4205 | 6.2495E-04 | 1.5488E-03 |
| ENSG00000100109\|TFIP11 | 3937.2781 | 0.4390 | ##### | 3.4205 | 6.2506E-04 | 1.5489E-03 |
| ENSG00000166436\|TRIM66 | 570.6910 | 0.4973 | ##### | 3.4203 | 6.2541E-04 | 1.5496E-03 |
| ENSG00000178035\|IMPDH2 | 9464.5300 | 0.4642 | ##### | 3.4203 | 6.2551E-04 | 1.5498E-03 |
| ENSG00000254002 | 26.2845 | 2.9860 | ##### | 3.4200 | 6.2619E-04 | 1.5513E-03 |
| ENSG00000169733\|RFNG | 2251.6128 | -0.4336 | ##### | -3.4195 | 6.2729E-04 | 1.5539E-03 |
| ENSG00000145723\|GIN1 | 407.8078 | -0.5326 | ##### | -3.4193 | 6.2789E-04 | 1.5553E-03 |
| ENSG00000117906\|RCN2 | 4563.0624 | 0.4272 | ##### | 3.4190 | 6.2855E-04 | 1.5568E-03 |
| ENSG00000186132\|C2orf76 | 592.6602 | -0.4946 | ##### | -3.4187 | 6.2931E-04 | 1.5585E-03 |
| ENSG00000286223 | 61.5634 | 1.4691 | ##### | 3.4186 | 6.2954E-04 | 1.5590E-03 |
| ENSG00000230423 | 7.8621 | 3.1823 | ##### | 3.4185 | 6.2975E-04 | 1.5594E-03 |
| ENSG00000239002\|SCARNA10 | 15.2861 | 2.9196 | ##### | 3.4184 | 6.2985E-04 | 1.5595E-03 |
| ENSG00000112062\|MAPK14 | 4076.5414 | -0.4138 | ##### | -3.4181 | 6.3052E-04 | 1.5610E-03 |
| ENSG00000180357\|ZNF609 | 5457.9952 | 0.4129 | ##### | 3.4168 | 6.3362E-04 | 1.5686E-03 |
| ENSG00000126464\|PRR12 | 4854.4722 | -0.3997 | ##### | -3.4167 | 6.3377E-04 | 1.5688E-03 |
| ENSG00000140153\|WDR20 | 2032.6541 | 0.4342 | ##### | 3.4162 | 6.3503E-04 | 1.5718E-03 |
| ENSG00000286812 | 21.0725 | -3.8403 | ##### | -3.4161 | 6.3523E-04 | 1.5722E-03 |
| ENSG00000178163\|ZNF518B | 1679.5641 | -0.4379 | ##### | -3.4155 | 6.3670E-04 | 1.5756E-03 |
| ENSG00000126709\|IFI6 | 938.0383 | -0.5040 | ##### | -3.4154 | 6.3680E-04 | 1.5758E-03 |
| ENSG00000113328\|CCNG1 | 2606.6296 | 0.4230 | ##### | 3.4152 | 6.3729E-04 | 1.5769E-03 |
| ENSG00000165525\|NEMF | 5932.0383 | 0.3981 | ##### | 3.4141 | 6.3988E-04 | 1.5831E-03 |
| ENSG00000224837\|GCSHP5 | 277.6854 | -0.5312 | ##### | -3.4136 | 6.4106E-04 | 1.5859E-03 |
| ENSG00000122085\|MTERF4 | 2254.5755 | 0.4222 | ##### | 3.4135 | 6.4145E-04 | 1.5868E-03 |
| ENSG00000206344\|HCG27 | 41.4491 | 1.2425 | ##### | 3.4130 | 6.4244E-04 | 1.5891E-03 |
| ENSG00000277075\|H2AC8 | 142.1352 | -0.6936 | ##### | -3.4129 | 6.4275E-04 | 1.5897E-03 |
| ENSG00000168813\|ZNF507 | 3502.5079 | -0.4233 | ##### | -3.4125 | 6.4360E-04 | 1.5917E-03 |
| ENSG00000117614\|SYF2 | 2611.5854 | 0.4248 | ##### | 3.4123 | 6.4409E-04 | 1.5927E-03 |
| ENSG00000289132 | 47.0709 | 1.1574 | ##### | 3.4115 | 6.4609E-04 | 1.5976E-03 |
| ENSG00000103160\|HSDL1 | 2633.8275 | 0.4118 | ##### | 3.4109 | 6.4747E-04 | 1.6007E-03 |
| ENSG00000175309\|PHYKPL | 1203.8893 | 0.4673 | ##### | 3.4109 | 6.4747E-04 | 1.6007E-03 |
| ENSG00000269743\|SLC25A53 | 186.0421 | 0.6985 | ##### | 3.4108 | 6.4773E-04 | 1.6012E-03 |
| ENSG00000226383\|LINC01876 | 84.0144 | -0.9077 | ##### | -3.4106 | 6.4824E-04 | 1.6024E-03 |
| ENSG00000261098 | 57.9385 | 1.0794 | ##### | 3.4101 | 6.4948E-04 | 1.6053E-03 |
| ENSG00000110321\|EIF4G2 | 77740.2935 | 0.2771 | ##### | 3.4100 | 6.4967E-04 | 1.6055E-03 |
| ENSG00000280294 | 28.1339 | 3.7380 | ##### | 3.4100 | 6.4965E-04 | 1.6055E-03 |
| ENSG00000173327\|MAP3K11 | 5290.6036 | 0.3979 | ##### | 3.4098 | 6.5017E-04 | 1.6065E-03 |
| ENSG00000130429\|ARPC1B | 36604.5157 | -0.3873 | ##### | -3.4098 | 6.5014E-04 | 1.6065E-03 |
| ENSG00000255561\|FDXACB1 | 266.4765 | 0.5756 | ##### | 3.4095 | 6.5072E-04 | 1.6076E-03 |
| ENSG00000173559\|NABP1 | 5014.9113 | 0.3246 | ##### | 3.4095 | 6.5073E-04 | 1.6076E-03 |
| ENSG00000273001\|PFKP-DT | 22.5094 | -2.2207 | ##### | -3.4088 | 6.5240E-04 | 1.6116E-03 |
| ENSG00000132286\|TIMM10B | 2118.7748 | 0.4338 | ##### | 3.4084 | 6.5346E-04 | 1.6141E-03 |
| ENSG00000091164\|TXNL1 | 6742.9458 | -0.4138 | ##### | -3.4073 | 6.5609E-04 | 1.6204E-03 |
| ENSG00000231307\|RPS3P2 | 14.0282 | 6.0293 | ##### | 3.4070 | 6.5683E-04 | 1.6221E-03 |
| ENSG00000125503\|PPP1R12C | 6992.3507 | -0.3571 | ##### | -3.4063 | 6.5860E-04 | 1.6264E-03 |
| ENSG00000237685 | 30.7096 | -5.8037 | ##### | -3.4057 | 6.6006E-04 | 1.6298E-03 |

| | | | | | | |
|---|---|---|---|---|---|---|
| ENSG00000269958 | 263.6468 | -0.6178 | ##### | -3.4049 | 6.6191E-04 | 1.6341E-03 |
| ENSG00000130363\|RSPH3 | 1230.5217 | -0.4318 | ##### | -3.4049 | 6.6189E-04 | 1.6341E-03 |
| ENSG00000168754\|FAM178B | 17.6896 | 3.7532 | ##### | 3.4042 | 6.6360E-04 | 1.6382E-03 |
| ENSG00000244055 | 113.7156 | -1.5063 | ##### | -3.4040 | 6.6409E-04 | 1.6392E-03 |
| ENSG00000108443\|RPS6KB1 | 4635.9550 | 0.4198 | ##### | 3.4031 | 6.6618E-04 | 1.6443E-03 |
| ENSG00000264647 | 11.8790 | 4.8847 | ##### | 3.4027 | 6.6713E-04 | 1.6465E-03 |
| ENSG00000118939\|UCHL3 | 1384.2389 | 0.5506 | ##### | 3.4027 | 6.6731E-04 | 1.6468E-03 |
| ENSG00000240616\|RPS6P25 | 10.2175 | 2.8127 | ##### | 3.4025 | 6.6777E-04 | 1.6478E-03 |
| ENSG00000250838 | 15.3557 | -1.4900 | ##### | -3.4024 | 6.6796E-04 | 1.6481E-03 |
| ENSG00000170852\|KBTBD2 | 5299.4529 | 0.4285 | ##### | 3.4024 | 6.6801E-04 | 1.6481E-03 |
| ENSG00000124243\|BCAS4 | 858.8701 | 0.4456 | ##### | 3.4012 | 6.7087E-04 | 1.6550E-03 |
| ENSG00000171603\|CLSTN1 | 16739.7887 | -0.3921 | ##### | -3.4003 | 6.7322E-04 | 1.6607E-03 |
| ENSG00000289939 | 10.4155 | 5.7052 | ##### | 3.4001 | 6.7357E-04 | 1.6614E-03 |
| ENSG00000204779\|FOXD4L5 | 5.0004 | 3.4667 | ##### | 3.3998 | 6.7445E-04 | 1.6634E-03 |
| ENSG00000260792\|LINC02280 | 11.1936 | 5.7872 | ##### | 3.3994 | 6.7527E-04 | 1.6653E-03 |
| ENSG00000053254\|FOXN3 | 1303.6641 | 0.4497 | ##### | 3.3989 | 6.7667E-04 | 1.6686E-03 |
| ENSG00000230623 | 29.2052 | -2.5038 | ##### | -3.3974 | 6.8023E-04 | 1.6773E-03 |
| ENSG00000131931\|THAP1 | 1121.3072 | 0.4729 | ##### | 3.3966 | 6.8225E-04 | 1.6821E-03 |
| ENSG00000266036\|SLC9A3R1-AS1 | 98.5075 | 0.7060 | ##### | 3.3962 | 6.8326E-04 | 1.6845E-03 |
| ENSG00000113811\|SELENOK | 2135.9422 | 0.4472 | ##### | 3.3951 | 6.8611E-04 | 1.6914E-03 |
| ENSG00000254480\|LINC02749 | 55.3696 | -8.2089 | ##### | -3.3948 | 6.8669E-04 | 1.6926E-03 |
| ENSG00000179218\|CALR | 45909.2647 | -0.2955 | ##### | -3.3946 | 6.8718E-04 | 1.6937E-03 |
| ENSG00000182179\|UBA7 | 63.7915 | -1.7718 | ##### | -3.3940 | 6.8889E-04 | 1.6978E-03 |
| ENSG00000006453\|BAIAP2L1 | 11271.5941 | -0.4106 | ##### | -3.3937 | 6.8945E-04 | 1.6990E-03 |
| ENSG00000273599 | 141.7971 | 0.6756 | ##### | 3.3935 | 6.9017E-04 | 1.7006E-03 |
| ENSG00000143217\|NECTIN4 | 31.7539 | -1.3155 | ##### | -3.3934 | 6.9032E-04 | 1.7009E-03 |
| ENSG00000103248\|MTHFSD | 1100.3636 | -0.4686 | ##### | -3.3934 | 6.9036E-04 | 1.7009E-03 |
| ENSG00000212932\|RPL23AP4 | 30.5079 | -1.2128 | ##### | -3.3932 | 6.9089E-04 | 1.7020E-03 |
| ENSG00000230337\|EXOSC10-AS1 | 72.2703 | 0.9216 | ##### | 3.3931 | 6.9107E-04 | 1.7023E-03 |
| ENSG00000185963\|BICD2 | 6417.0854 | 0.4080 | ##### | 3.3925 | 6.9250E-04 | 1.7057E-03 |
| ENSG00000135298\|ADGRB3 | 19.5832 | 2.8162 | ##### | 3.3921 | 6.9357E-04 | 1.7082E-03 |
| ENSG00000288704 | 9.6114 | 3.5522 | ##### | 3.3920 | 6.9384E-04 | 1.7087E-03 |
| ENSG00000143742\|SRP9 | 14302.6145 | -0.3572 | ##### | -3.3914 | 6.9535E-04 | 1.7123E-03 |
| ENSG00000210100\|MT-TI | 290.6357 | 0.5566 | ##### | 3.3912 | 6.9599E-04 | 1.7137E-03 |
| ENSG00000182749\|PAQR7 | 5746.1577 | -0.4221 | ##### | -3.3910 | 6.9629E-04 | 1.7143E-03 |
| ENSG00000254340 | 22.3724 | -1.6604 | ##### | -3.3908 | 6.9685E-04 | 1.7156E-03 |
| ENSG00000128524\|ATP6V1F | 11090.6553 | 0.4050 | ##### | 3.3907 | 6.9709E-04 | 1.7160E-03 |
| ENSG00000135766\|EGLN1 | 8436.9398 | -0.3899 | ##### | -3.3903 | 6.9827E-04 | 1.7188E-03 |
| ENSG00000135341\|MAP3K7 | 6323.5156 | 0.4055 | ##### | 3.3900 | 6.9898E-04 | 1.7204E-03 |
| ENSG00000279108 | 20.2695 | -2.6335 | ##### | -3.3898 | 6.9939E-04 | 1.7211E-03 |
| ENSG00000010030\|ETV7 | 39.0520 | 1.4083 | ##### | 3.3898 | 6.9946E-04 | 1.7211E-03 |
| ENSG00000232044\|SILC1 | 9.0943 | -6.1000 | ##### | -3.3898 | 6.9942E-04 | 1.7211E-03 |
| ENSG00000266278\|LINC01910 | 51.5006 | -1.4347 | ##### | -3.3895 | 7.0016E-04 | 1.7227E-03 |
| ENSG00000215869 | 8.7697 | -5.5191 | ##### | -3.3892 | 7.0099E-04 | 1.7246E-03 |
| ENSG00000271787 | 22.6142 | 1.2446 | ##### | 3.3890 | 7.0140E-04 | 1.7255E-03 |
| ENSG00000143947\|RPS27A | 45098.1734 | 0.4133 | ##### | 3.3890 | 7.0159E-04 | 1.7258E-03 |
| ENSG00000166173\|LARP6 | 1416.0896 | 0.4308 | ##### | 3.3885 | 7.0280E-04 | 1.7286E-03 |
| ENSG00000070950\|RAD18 | 2355.8052 | 0.4407 | ##### | 3.3883 | 7.0330E-04 | 1.7297E-03 |
| ENSG00000109534\|GAR1 | 3232.7168 | 0.4225 | ##### | 3.3882 | 7.0359E-04 | 1.7303E-03 |
| ENSG00000162783\|IER5 | 4072.8401 | -0.4222 | ##### | -3.3877 | 7.0489E-04 | 1.7333E-03 |
| ENSG00000281501\|SEPSECS-AS1 | 55.0153 | -1.4017 | ##### | -3.3876 | 7.0499E-04 | 1.7334E-03 |
| ENSG00000189056\|RELN | 47.4933 | 2.6016 | ##### | 3.3876 | 7.0510E-04 | 1.7336E-03 |
| ENSG00000267575\|LINC02987 | 2029.0952 | -0.4193 | ##### | -3.3874 | 7.0549E-04 | 1.7343E-03 |
| ENSG00000159200\|RCAN1 | 3920.6306 | 0.4241 | ##### | 3.3874 | 7.0546E-04 | 1.7343E-03 |
| ENSG00000213411\|RBM22P2 | 9.2985 | 6.4763 | ##### | 3.3864 | 7.0814E-04 | 1.7406E-03 |
| ENSG00000163166\|IWS1 | 6361.0189 | -0.3906 | ##### | -3.3860 | 7.0920E-04 | 1.7431E-03 |
| ENSG00000130254\|SAFB2 | 6470.9579 | 0.3978 | ##### | 3.3847 | 7.1257E-04 | 1.7512E-03 |
| ENSG00000141028\|CDRT15P1 | 16.5756 | 1.6856 | ##### | 3.3831 | 7.1669E-04 | 1.7612E-03 |
| ENSG00000244363\|RPL7P23 | 33.3403 | -1.1148 | ##### | -3.3823 | 7.1891E-04 | 1.7665E-03 |
| ENSG00000137070\|IL11RA | 266.5102 | 0.5505 | ##### | 3.3822 | 7.1911E-04 | 1.7669E-03 |
| ENSG00000267283 | 67.0515 | 1.0295 | ##### | 3.3820 | 7.1948E-04 | 1.7676E-03 |
| ENSG00000275367 | 15.5223 | 3.6240 | ##### | 3.3820 | 7.1966E-04 | 1.7679E-03 |
| ENSG00000265618 | 297.0078 | -0.5720 | ##### | -3.3812 | 7.2173E-04 | 1.7729E-03 |

| | | | | | | |
|---|---|---|---|---|---|---|
| ENSG00000188801\|ZNF322P1 | 110.2579 | 0.7314 | ##### | 3.3811 | 7.2205E-04 | 1.7735E-03 |
| ENSG00000247934 | 82.0646 | -0.8698 | ##### | -3.3810 | 7.2229E-04 | 1.7738E-03 |
| ENSG00000249936\|RAC1P2 | 351.2447 | 0.5941 | ##### | 3.3810 | 7.2229E-04 | 1.7738E-03 |
| ENSG00000106686\|SPATA6L | 27.0807 | -1.4718 | ##### | -3.3808 | 7.2268E-04 | 1.7746E-03 |
| ENSG00000183682\|BMP8A | 124.3138 | 0.7013 | ##### | 3.3797 | 7.2553E-04 | 1.7815E-03 |
| ENSG00000288890 | 20.3556 | 2.5823 | ##### | 3.3788 | 7.2813E-04 | 1.7877E-03 |
| ENSG00000288917 | 12.9881 | 2.2578 | ##### | 3.3787 | 7.2836E-04 | 1.7881E-03 |
| ENSG00000169710\|FASN | 32971.4957 | -0.3892 | ##### | -3.3777 | 7.3108E-04 | 1.7946E-03 |
| ENSG00000101292\|PROKR2 | 66.7140 | 5.4628 | ##### | 3.3770 | 7.3293E-04 | 1.7990E-03 |
| ENSG00000000457\|SCYL3 | 776.1839 | -0.5128 | ##### | -3.3767 | 7.3365E-04 | 1.8007E-03 |
| ENSG00000174775\|HRAS | 2878.4152 | 0.4227 | ##### | 3.3758 | 7.3600E-04 | 1.8063E-03 |
| ENSG00000077721\|UBE2A | 6541.7560 | 0.4177 | ##### | 3.3751 | 7.3786E-04 | 1.8107E-03 |
| ENSG00000185453\|ZSWIM9 | 1460.6797 | -0.4242 | ##### | -3.3747 | 7.3900E-04 | 1.8134E-03 |
| ENSG00000158828\|PINK1 | 4326.0054 | -0.4128 | ##### | -3.3744 | 7.3973E-04 | 1.8150E-03 |
| ENSG00000289883\|ENEMAL | 9.1174 | 5.4492 | ##### | 3.3741 | 7.4070E-04 | 1.8172E-03 |
| ENSG00000200674\|RN7SKP160 | 11.8577 | 4.8219 | ##### | 3.3738 | 7.4132E-04 | 1.8186E-03 |
| ENSG00000167595\|PROSER3 | 1884.4208 | -0.4175 | ##### | -3.3728 | 7.4400E-04 | 1.8250E-03 |
| ENSG00000142748\|FCN3 | 28.3062 | 7.4875 | ##### | 3.3719 | 7.4652E-04 | 1.8311E-03 |
| ENSG00000099341\|PSMD8 | 14371.6556 | -0.3673 | ##### | -3.3716 | 7.4739E-04 | 1.8330E-03 |
| ENSG00000182923\|CEP63 | 2423.2280 | 0.4201 | ##### | 3.3715 | 7.4760E-04 | 1.8334E-03 |
| ENSG00000175548\|ALG10B | 1491.3962 | 0.4477 | ##### | 3.3713 | 7.4804E-04 | 1.8343E-03 |
| ENSG00000177728\|TMEM94 | 4499.0437 | -0.4152 | ##### | -3.3706 | 7.5003E-04 | 1.8390E-03 |
| ENSG00000267769 | 18.4494 | 1.8432 | ##### | 3.3691 | 7.5404E-04 | 1.8487E-03 |
| ENSG00000291084\|ALOX12P2 | 98.4853 | -0.9812 | ##### | -3.3683 | 7.5646E-04 | 1.8545E-03 |
| ENSG00000142546\|NOSIP | 4396.9798 | -0.4139 | ##### | -3.3681 | 7.5691E-04 | 1.8555E-03 |
| ENSG00000231439\|WASIR2 | 25.7904 | -1.4107 | ##### | -3.3677 | 7.5803E-04 | 1.8581E-03 |
| ENSG00000260456\|C16orf95 | 173.7224 | -0.6179 | ##### | -3.3670 | 7.5997E-04 | 1.8627E-03 |
| ENSG00000275661 | 6.4396 | -2.1617 | ##### | -3.3669 | 7.6025E-04 | 1.8632E-03 |
| ENSG00000212978\|C2orf74-DT | 1270.7185 | -0.4514 | ##### | -3.3666 | 7.6109E-04 | 1.8651E-03 |
| ENSG00000267174 | 16.1990 | 5.7580 | ##### | 3.3665 | 7.6122E-04 | 1.8653E-03 |
| ENSG00000182450\|KCNK4 | 13.1005 | 6.4888 | ##### | 3.3661 | 7.6252E-04 | 1.8683E-03 |
| ENSG00000115310\|RTN4 | 36960.2459 | -0.2134 | ##### | -3.3660 | 7.6268E-04 | 1.8685E-03 |
| ENSG00000172006\|ZNF554 | 139.8447 | -0.6876 | ##### | -3.3658 | 7.6316E-04 | 1.8694E-03 |
| ENSG00000169020\|ATP5ME | 2309.2552 | -0.4772 | ##### | -3.3658 | 7.6311E-04 | 1.8694E-03 |
| ENSG00000204934\|ATP6V0E2-AS1 | 653.0518 | 0.4680 | ##### | 3.3658 | 7.6332E-04 | 1.8696E-03 |
| ENSG00000286134 | 46.9404 | 1.1137 | ##### | 3.3657 | 7.6359E-04 | 1.8701E-03 |
| ENSG00000228486\|C2orf92 | 81.5069 | 1.1028 | ##### | 3.3641 | 7.6783E-04 | 1.8804E-03 |
| ENSG00000137098\|SPAG8 | 24.0946 | 1.6506 | ##### | 3.3633 | 7.7021E-04 | 1.8860E-03 |
| ENSG00000166526\|ZNF3 | 2035.5960 | 0.4248 | ##### | 3.3626 | 7.7200E-04 | 1.8903E-03 |
| ENSG00000289378 | 75.2941 | -0.8718 | ##### | -3.3626 | 7.7224E-04 | 1.8907E-03 |
| ENSG00000246283\|TTBK2-AS1 | 14.2128 | -2.9657 | ##### | -3.3620 | 7.7385E-04 | 1.8945E-03 |
| ENSG00000175203\|DCTN2 | 11217.3713 | -0.4003 | ##### | -3.3619 | 7.7410E-04 | 1.8949E-03 |
| ENSG00000229282 | 54.5252 | -8.1348 | ##### | -3.3619 | 7.7416E-04 | 1.8949E-03 |
| ENSG00000137509\|PRCP | 3350.1709 | -0.4248 | ##### | -3.3617 | 7.7455E-04 | 1.8957E-03 |
| ENSG00000289409 | 4.9225 | 5.6642 | ##### | 3.3615 | 7.7507E-04 | 1.8969E-03 |
| ENSG00000103707\|MTFMT | 1058.2635 | 0.4662 | ##### | 3.3614 | 7.7543E-04 | 1.8976E-03 |
| ENSG00000213676\|ATF6B | 3335.6666 | -0.4235 | ##### | -3.3611 | 7.7621E-04 | 1.8994E-03 |
| ENSG00000096717\|SIRT1 | 3988.5683 | 0.4294 | ##### | 3.3607 | 7.7749E-04 | 1.9023E-03 |
| ENSG00000131969\|ABHD12B | 32.5731 | 1.3259 | ##### | 3.3606 | 7.7786E-04 | 1.9031E-03 |
| ENSG00000072401\|UBE2D1 | 1708.5293 | -0.4300 | ##### | -3.3598 | 7.7993E-04 | 1.9079E-03 |
| ENSG00000254449\|SF3A3P2 | 14.4683 | 1.7444 | ##### | 3.3598 | 7.7996E-04 | 1.9079E-03 |
| ENSG00000134780\|DAGLA | 1154.9318 | -0.4475 | ##### | -3.3597 | 7.8020E-04 | 1.9083E-03 |
| ENSG00000273004 | 23.5015 | 1.7866 | ##### | 3.3595 | 7.8070E-04 | 1.9094E-03 |
| ENSG00000226465 | 24.9674 | -1.4301 | ##### | -3.3592 | 7.8173E-04 | 1.9117E-03 |
| ENSG00000269949 | 14.0738 | 2.1309 | ##### | 3.3577 | 7.8600E-04 | 1.9220E-03 |
| ENSG00000208028\|MIR616 | 11.4135 | -4.9154 | ##### | -3.3574 | 7.8684E-04 | 1.9239E-03 |
| ENSG00000167447\|SMG8 | 3796.0422 | 0.3899 | ##### | 3.3564 | 7.8976E-04 | 1.9309E-03 |
| ENSG00000215241\|LINC02449 | 30.2335 | 1.4107 | ##### | 3.3558 | 7.9133E-04 | 1.9346E-03 |
| ENSG00000166860\|ZBTB39 | 1412.0539 | 0.4197 | ##### | 3.3555 | 7.9223E-04 | 1.9366E-03 |
| ENSG00000187601\|MAGEH1 | 449.8631 | 0.4971 | ##### | 3.3554 | 7.9258E-04 | 1.9373E-03 |
| ENSG00000159140\|SON | 23659.2647 | 0.3944 | ##### | 3.3553 | 7.9292E-04 | 1.9380E-03 |
| ENSG00000242829\|RPS26P21 | 49.7311 | 8.7164 | ##### | 3.3552 | 7.9302E-04 | 1.9381E-03 |
| ENSG00000134590\|RTL8C | 5257.5728 | -0.4041 | ##### | -3.3544 | 7.9527E-04 | 1.9434E-03 |

| | | | | | | |
|---|---|---|---|---|---|---|
| ENSG00000151006\|PRSS53 | 368.0467 | -0.5271 | ##### | -3.3537 | 7.9745E-04 | 1.9486E-03 |
| ENSG00000157322\|CLEC18A | 12.2939 | -1.8059 | ##### | -3.3536 | 7.9759E-04 | 1.9488E-03 |
| ENSG00000159210\|SNF8 | 12270.3908 | 0.4035 | ##### | 3.3534 | 7.9820E-04 | 1.9501E-03 |
| ENSG00000125813\|PAX1 | 20.1920 | 1.4201 | ##### | 3.3531 | 7.9922E-04 | 1.9524E-03 |
| ENSG00000286898 | 13.6102 | 5.0159 | ##### | 3.3526 | 8.0054E-04 | 1.9555E-03 |
| ENSG00000283312 | 31.1951 | 1.1721 | ##### | 3.3521 | 8.0205E-04 | 1.9590E-03 |
| ENSG00000188229\|TUBB4B | 48463.6917 | 0.3056 | ##### | 3.3517 | 8.0327E-04 | 1.9619E-03 |
| ENSG00000285257 | 36.7116 | -1.1635 | ##### | -3.3515 | 8.0366E-04 | 1.9627E-03 |
| ENSG00000114023\|FAM162A | 2145.4313 | 0.4039 | ##### | 3.3514 | 8.0404E-04 | 1.9632E-03 |
| ENSG00000246366\|LACTB2-AS1 | 54.1016 | -8.1295 | ##### | -3.3514 | 8.0408E-04 | 1.9632E-03 |
| ENSG00000227071\|FOCAD-AS1 | 12.4135 | -6.0418 | ##### | -3.3514 | 8.0398E-04 | 1.9632E-03 |
| ENSG00000100991\|TRPC4AP | 8867.7435 | 0.3955 | ##### | 3.3505 | 8.0655E-04 | 1.9691E-03 |
| ENSG00000141562\|NARF | 2099.7848 | 0.4263 | ##### | 3.3503 | 8.0718E-04 | 1.9705E-03 |
| ENSG00000255136\|TPBGL-AS1 | 6.8424 | -3.1983 | ##### | -3.3497 | 8.0910E-04 | 1.9750E-03 |
| ENSG00000289074 | 67.6673 | 1.1840 | ##### | 3.3494 | 8.0975E-04 | 1.9764E-03 |
| ENSG00000197191\|CYSRT1 | 47.5784 | -1.2259 | ##### | -3.3491 | 8.1089E-04 | 1.9790E-03 |
| ENSG00000188010\|MORN2 | 880.6398 | 0.4558 | ##### | 3.3487 | 8.1190E-04 | 1.9813E-03 |
| ENSG00000160563\|MED27 | 2160.7939 | -0.4146 | ##### | -3.3484 | 8.1281E-04 | 1.9834E-03 |
| ENSG00000286788 | 56.3327 | 4.3845 | ##### | 3.3481 | 8.1372E-04 | 1.9854E-03 |
| ENSG00000111880\|RNGTT | 3045.1507 | 0.4132 | ##### | 3.3474 | 8.1580E-04 | 1.9903E-03 |
| ENSG00000279246 | 257.1644 | -1.2479 | ##### | -3.3474 | 8.1588E-04 | 1.9904E-03 |
| ENSG00000260526\|AP1AR-DT | 37.9407 | 1.1933 | ##### | 3.3473 | 8.1616E-04 | 1.9909E-03 |
| ENSG00000266921\|ZNF230-DT | 258.1154 | -1.2993 | ##### | -3.3466 | 8.1809E-04 | 1.9954E-03 |
| ENSG00000139133\|ALG10 | 940.2486 | 0.4901 | ##### | 3.3461 | 8.1957E-04 | 1.9989E-03 |
| ENSG00000127663\|KDM4B | 1527.5583 | 0.4108 | ##### | 3.3457 | 8.2062E-04 | 2.0013E-03 |
| ENSG00000072840\|EVC | 1926.5544 | -0.4286 | ##### | -3.3453 | 8.2189E-04 | 2.0041E-03 |
| ENSG00000289402 | 12.0467 | -6.5120 | ##### | -3.3453 | 8.2184E-04 | 2.0041E-03 |
| ENSG00000289064 | 10.2105 | 6.6022 | ##### | 3.3451 | 8.2248E-04 | 2.0053E-03 |
| ENSG00000251600\|GUSBP5 | 38.5767 | -2.3960 | ##### | -3.3450 | 8.2279E-04 | 2.0059E-03 |
| ENSG00000169064\|ZBBX | 48.8139 | -8.0266 | ##### | -3.3449 | 8.2325E-04 | 2.0069E-03 |
| ENSG00000183784\|DOCK8-AS1 | 31.2667 | -4.0342 | ##### | -3.3447 | 8.2366E-04 | 2.0077E-03 |
| ENSG00000233337\|UBE2FP3 | 41.7002 | 1.1207 | ##### | 3.3446 | 8.2392E-04 | 2.0082E-03 |
| ENSG00000168952\|STXBP6 | 462.5945 | -0.5914 | ##### | -3.3431 | 8.2852E-04 | 2.0192E-03 |
| ENSG00000290876 | 13.1621 | -3.1092 | ##### | -3.3426 | 8.2986E-04 | 2.0223E-03 |
| ENSG00000261371\|PECAM1 | 50.8276 | -8.0848 | ##### | -3.3425 | 8.3036E-04 | 2.0234E-03 |
| ENSG00000248896 | 22.2989 | -1.7550 | ##### | -3.3424 | 8.3045E-04 | 2.0235E-03 |
| ENSG00000224411\|HSP90AA2P | 828.8885 | 0.4517 | ##### | 3.3422 | 8.3119E-04 | 2.0249E-03 |
| ENSG00000143919\|CAMKMT | 552.9558 | -0.4816 | ##### | -3.3422 | 8.3118E-04 | 2.0249E-03 |
| ENSG00000187013\|LINC02875 | 20.2087 | -1.4677 | ##### | -3.3421 | 8.3138E-04 | 2.0252E-03 |
| ENSG00000118369\|USP35 | 779.3383 | 0.4897 | ##### | 3.3419 | 8.3203E-04 | 2.0266E-03 |
| ENSG00000272473 | 71.0685 | -4.0461 | ##### | -3.3413 | 8.3380E-04 | 2.0308E-03 |
| ENSG00000159202\|UBE2Z | 13112.3779 | 0.2444 | ##### | 3.3412 | 8.3404E-04 | 2.0312E-03 |
| ENSG00000288825\|H2AC18 | 1581.0962 | -0.4277 | ##### | -3.3398 | 8.3846E-04 | 2.0418E-03 |
| ENSG00000179152\|TCAIM | 1353.0086 | -0.4444 | ##### | -3.3393 | 8.3997E-04 | 2.0453E-03 |
| ENSG00000089682\|RBM41 | 1899.5170 | 0.4142 | ##### | 3.3390 | 8.4086E-04 | 2.0473E-03 |
| ENSG00000136104\|RNASEH2B | 1937.4994 | 0.4284 | ##### | 3.3384 | 8.4260E-04 | 2.0514E-03 |
| ENSG00000272666\|KLHDC7B-DT | 16.7881 | -4.4541 | ##### | -3.3374 | 8.4554E-04 | 2.0584E-03 |
| ENSG00000277304 | 62.9485 | -8.9381 | ##### | -3.3374 | 8.4580E-04 | 2.0588E-03 |
| ENSG00000291288 | 17.7081 | 3.0151 | ##### | 3.3373 | 8.4598E-04 | 2.0591E-03 |
| ENSG00000115282\|TTC31 | 2226.6819 | 0.4326 | ##### | 3.3369 | 8.4732E-04 | 2.0622E-03 |
| ENSG00000286808 | 99.6769 | 1.2983 | ##### | 3.3367 | 8.4778E-04 | 2.0632E-03 |
| ENSG00000176076\|KCNE5 | 23.0714 | 1.9029 | ##### | 3.3366 | 8.4817E-04 | 2.0639E-03 |
| ENSG00000292979 | 179.2062 | 0.6356 | ##### | 3.3364 | 8.4883E-04 | 2.0654E-03 |
| ENSG00000120832\|MTERF2 | 1863.8423 | -0.4277 | ##### | -3.3363 | 8.4896E-04 | 2.0655E-03 |
| ENSG00000105171\|POP4 | 3341.8462 | -0.4185 | ##### | -3.3360 | 8.4994E-04 | 2.0677E-03 |
| ENSG00000205352\|PRR13 | 13219.5049 | 0.4272 | ##### | 3.3358 | 8.5055E-04 | 2.0690E-03 |
| ENSG00000204183\|GDF5-AS1 | 25.8456 | 7.3318 | ##### | 3.3355 | 8.5134E-04 | 2.0708E-03 |
| ENSG00000286001 | 773.7283 | -1.1312 | ##### | -3.3351 | 8.5284E-04 | 2.0743E-03 |
| ENSG00000285637 | 44.1203 | -5.9062 | ##### | -3.3350 | 8.5309E-04 | 2.0747E-03 |
| ENSG00000273032\|DGCR5 | 777.5053 | -0.4613 | ##### | -3.3344 | 8.5493E-04 | 2.0790E-03 |
| ENSG00000267512 | 112.9161 | 0.8003 | ##### | 3.3343 | 8.5526E-04 | 2.0797E-03 |
| ENSG00000111897\|SERINC1 | 10645.1132 | -0.3968 | ##### | -3.3342 | 8.5543E-04 | 2.0799E-03 |
| ENSG00000130726\|TRIM28 | 57294.6678 | 0.3883 | ##### | 3.3341 | 8.5570E-04 | 2.0804E-03 |

| | | | | | | |
|---|---|---|---|---|---|---|
| ENSG00000284747 | 142.0455 | 0.5769 | ##### | 3.3329 | 8.5952E-04 | 2.0895E-03 |
| ENSG00000228158\|TLE1P1 | 11.8441 | -1.9246 | ##### | -3.3329 | 8.5960E-04 | 2.0895E-03 |
| ENSG00000260806\|PAPOLA-DT | 65.2558 | -1.1017 | ##### | -3.3325 | 8.6073E-04 | 2.0921E-03 |
| ENSG00000196834\|POTEI | 15.3550 | 1.5448 | ##### | 3.3323 | 8.6132E-04 | 2.0934E-03 |
| ENSG00000161544\|CYGB | 65.8384 | -0.8716 | ##### | -3.3321 | 8.6196E-04 | 2.0948E-03 |
| ENSG00000105669\|COPE | 7823.3012 | 0.4014 | ##### | 3.3320 | 8.6237E-04 | 2.0956E-03 |
| ENSG00000132024\|CC2D1A | 2337.2590 | -0.4017 | ##### | -3.3318 | 8.6300E-04 | 2.0969E-03 |
| ENSG00000261770 | 22.4691 | -1.4380 | ##### | -3.3315 | 8.6369E-04 | 2.0984E-03 |
| ENSG00000117461\|PIK3R3 | 1866.9329 | 0.4206 | ##### | 3.3309 | 8.6551E-04 | 2.1027E-03 |
| ENSG00000149313\|AASDHPPT | 6158.3916 | 0.4087 | ##### | 3.3301 | 8.6807E-04 | 2.1087E-03 |
| ENSG00000066185\|ZMYND12 | 54.0802 | -0.9738 | ##### | -3.3300 | 8.6836E-04 | 2.1093E-03 |
| ENSG00000100418\|DESI1 | 5184.6259 | -0.3927 | ##### | -3.3298 | 8.6911E-04 | 2.1109E-03 |
| ENSG00000063015\|SEZ6 | 18.9397 | 7.4369 | ##### | 3.3296 | 8.6958E-04 | 2.1119E-03 |
| ENSG00000147130\|ZMYM3 | 4679.4503 | 0.4149 | ##### | 3.3292 | 8.7098E-04 | 2.1151E-03 |
| ENSG00000275764 | 304.0535 | 0.6000 | ##### | 3.3290 | 8.7162E-04 | 2.1165E-03 |
| ENSG00000283317 | 15.4291 | 2.4586 | ##### | 3.3288 | 8.7229E-04 | 2.1180E-03 |
| ENSG00000059588\|TARBP1 | 2559.4587 | -0.3940 | ##### | -3.3287 | 8.7266E-04 | 2.1187E-03 |
| ENSG00000273398 | 287.8597 | 0.6604 | ##### | 3.3285 | 8.7328E-04 | 2.1200E-03 |
| ENSG00000262904\|TMPOP2 | 17.5540 | 1.5978 | ##### | 3.3280 | 8.7480E-04 | 2.1235E-03 |
| ENSG00000251247\|ZNF345 | 146.8782 | -0.6605 | ##### | -3.3267 | 8.7870E-04 | 2.1328E-03 |
| ENSG00000075886\|TUBA3D | 19.4029 | -1.6246 | ##### | -3.3266 | 8.7899E-04 | 2.1334E-03 |
| ENSG00000181227\|DLSTP1 | 19.8505 | 2.2004 | ##### | 3.3259 | 8.8122E-04 | 2.1386E-03 |
| ENSG00000174371\|EXO1 | 4112.7052 | -0.4081 | ##### | -3.3252 | 8.8343E-04 | 2.1438E-03 |
| ENSG00000066136\|NFYC | 6009.5634 | 0.3652 | ##### | 3.3250 | 8.8420E-04 | 2.1455E-03 |
| ENSG00000144445\|KANSL1L | 882.7996 | -0.4482 | ##### | -3.3246 | 8.8534E-04 | 2.1479E-03 |
| ENSG00000144567\|RETREG2 | 9033.3144 | 0.3919 | ##### | 3.3246 | 8.8531E-04 | 2.1479E-03 |
| ENSG00000141294\|LRRC46 | 130.5738 | 0.7781 | ##### | 3.3244 | 8.8622E-04 | 2.1499E-03 |
| ENSG00000117362\|APH1A | 13843.4911 | 0.4088 | ##### | 3.3243 | 8.8646E-04 | 2.1503E-03 |
| ENSG00000261136 | 41.1785 | 1.2240 | ##### | 3.3239 | 8.8759E-04 | 2.1528E-03 |
| ENSG00000159337\|PLA2G4D | 7.2074 | -6.0684 | ##### | -3.3238 | 8.8799E-04 | 2.1536E-03 |
| ENSG00000065559\|MAP2K4 | 2744.6167 | -0.4280 | ##### | -3.3230 | 8.9042E-04 | 2.1593E-03 |
| ENSG00000276851 | 33.9260 | -1.5404 | ##### | -3.3228 | 8.9133E-04 | 2.1614E-03 |
| ENSG00000175106\|TVP23C | 420.6474 | -0.5481 | ##### | -3.3218 | 8.9443E-04 | 2.1687E-03 |
| ENSG00000056736\|IL17RB | 284.8512 | 0.6800 | ##### | 3.3217 | 8.9469E-04 | 2.1692E-03 |
| ENSG00000157502\|PWWP3B | 11.1546 | -3.5967 | ##### | -3.3216 | 8.9502E-04 | 2.1698E-03 |
| ENSG00000186496\|ZNF396 | 114.8995 | -0.6824 | ##### | -3.3204 | 8.9885E-04 | 2.1789E-03 |
| ENSG00000168772\|CXXC4 | 287.1281 | 0.6539 | ##### | 3.3202 | 8.9938E-04 | 2.1800E-03 |
| ENSG00000075142\|SRI | 6791.0154 | -0.4007 | ##### | -3.3199 | 9.0042E-04 | 2.1824E-03 |
| ENSG00000078369\|GNB1 | 40686.6187 | -0.2486 | ##### | -3.3192 | 9.0281E-04 | 2.1879E-03 |
| ENSG00000112249\|ASCC3 | 6822.0104 | -0.4039 | ##### | -3.3192 | 9.0284E-04 | 2.1879E-03 |
| ENSG00000162144\|CYB561A3 | 4359.7107 | -0.4426 | ##### | -3.3191 | 9.0307E-04 | 2.1883E-03 |
| ENSG00000167094\|TTC16 | 29.7545 | -1.2373 | ##### | -3.3185 | 9.0509E-04 | 2.1930E-03 |
| ENSG00000258586\|LINC02274 | 7.9928 | 4.2926 | ##### | 3.3184 | 9.0533E-04 | 2.1934E-03 |
| ENSG00000267405 | 16.3567 | 1.9372 | ##### | 3.3177 | 9.0768E-04 | 2.1989E-03 |
| ENSG00000172586\|CHCHD1 | 2101.2539 | -0.4010 | ##### | -3.3176 | 9.0780E-04 | 2.1990E-03 |
| ENSG00000284606 | 21.7775 | 2.3174 | ##### | 3.3160 | 9.1324E-04 | 2.2120E-03 |
| ENSG00000159708\|LRRC36 | 25.4328 | 4.2788 | ##### | 3.3158 | 9.1393E-04 | 2.2135E-03 |
| ENSG00000255108 | 87.6970 | 0.7368 | ##### | 3.3156 | 9.1460E-04 | 2.2149E-03 |
| ENSG00000272799 | 8.9655 | 6.1643 | ##### | 3.3147 | 9.1726E-04 | 2.2212E-03 |
| ENSG00000277895 | 16.5209 | 2.0006 | ##### | 3.3145 | 9.1819E-04 | 2.2233E-03 |
| ENSG00000214546 | 11.0889 | -6.3916 | ##### | -3.3144 | 9.1828E-04 | 2.2233E-03 |
| ENSG00000238151\|MLLT10P1 | 9.6016 | 2.8825 | ##### | 3.3143 | 9.1883E-04 | 2.2244E-03 |
| ENSG00000177192\|PUS1 | 6752.1863 | 0.3918 | ##### | 3.3141 | 9.1947E-04 | 2.2258E-03 |
| ENSG00000110925\|CSRNP2 | 3806.8838 | 0.4135 | ##### | 3.3139 | 9.2011E-04 | 2.2272E-03 |
| ENSG00000228793 | 22.6380 | -1.2440 | ##### | -3.3136 | 9.2089E-04 | 2.2289E-03 |
| ENSG00000204576\|PRR3 | 1246.2151 | 0.4196 | ##### | 3.3136 | 9.2117E-04 | 2.2294E-03 |
| ENSG00000005339\|CREBBP | 7366.5557 | 0.3910 | ##### | 3.3130 | 9.2290E-04 | 2.2334E-03 |
| ENSG00000224861\|YBX1P1 | 739.2845 | 0.4630 | ##### | 3.3128 | 9.2369E-04 | 2.2351E-03 |
| ENSG00000240891\|PLCXD2 | 1051.0219 | 0.4280 | ##### | 3.3124 | 9.2496E-04 | 2.2380E-03 |
| ENSG00000164896\|FASTK | 7363.6069 | 0.3449 | ##### | 3.3123 | 9.2522E-04 | 2.2385E-03 |
| ENSG00000105514\|RAB3D | 2553.2874 | -0.3619 | ##### | -3.3121 | 9.2604E-04 | 2.2403E-03 |
| ENSG00000113558\|SKP1 | 12536.2355 | 0.4077 | ##### | 3.3115 | 9.2793E-04 | 2.2447E-03 |
| ENSG00000137413\|TAF8 | 1989.3240 | -0.4250 | ##### | -3.3114 | 9.2838E-04 | 2.2456E-03 |

| | | | | | | |
|---|---|---|---|---|---|---|
| ENSG00000148362\|PAXX | 2527.1411 | -0.2802 | ##### | -3.3111 | 9.2935E-04 | 2.2477E-03 |
| ENSG00000235105 | 69.2511 | -0.9761 | ##### | -3.3104 | 9.3146E-04 | 2.2527E-03 |
| ENSG00000200120\|Y_RNA | 4.9018 | 5.4993 | ##### | 3.3104 | 9.3158E-04 | 2.2527E-03 |
| ENSG00000144031\|ANKRD53 | 16.4495 | 1.6002 | ##### | 3.3104 | 9.3165E-04 | 2.2527E-03 |
| ENSG00000273203 | 38.4942 | -1.0326 | ##### | -3.3099 | 9.3329E-04 | 2.2565E-03 |
| ENSG00000167037\|SGSM1 | 174.8112 | -0.5769 | ##### | -3.3092 | 9.3561E-04 | 2.2620E-03 |
| ENSG00000203644 | 218.6137 | 0.6884 | ##### | 3.3083 | 9.3875E-04 | 2.2694E-03 |
| ENSG00000226053\|LINC01776 | 22.7342 | -1.7417 | ##### | -3.3081 | 9.3948E-04 | 2.2709E-03 |
| ENSG00000269486\|ERVK9-11 | 107.2995 | -0.7713 | ##### | -3.3080 | 9.3962E-04 | 2.2711E-03 |
| ENSG00000234327\|ZNF232-AS1 | 282.0005 | -0.5498 | ##### | -3.3073 | 9.4214E-04 | 2.2770E-03 |
| ENSG00000260196 | 298.4040 | 0.5893 | ##### | 3.3068 | 9.4355E-04 | 2.2802E-03 |
| ENSG00000259520\|SLC28A2-AS1 | 14.0104 | -2.6172 | ##### | -3.3061 | 9.4620E-04 | 2.2865E-03 |
| ENSG00000291136 | 1086.2585 | 0.4625 | ##### | 3.3060 | 9.4652E-04 | 2.2870E-03 |
| ENSG00000232815\|DUX4L50 | 14.5884 | -2.3862 | ##### | -3.3057 | 9.4733E-04 | 2.2888E-03 |
| ENSG00000289592 | 50.8317 | -1.6474 | ##### | -3.3053 | 9.4876E-04 | 2.2921E-03 |
| ENSG00000121905\|HPCA | 96.7783 | -1.0340 | ##### | -3.3048 | 9.5043E-04 | 2.2959E-03 |
| ENSG00000183807\|FAM162B | 12.4359 | -6.0408 | ##### | -3.3046 | 9.5108E-04 | 2.2973E-03 |
| ENSG00000257527 | 28.2465 | -1.2137 | ##### | -3.3041 | 9.5285E-04 | 2.3014E-03 |
| ENSG00000152380\|FAM151B | 83.3240 | -0.9718 | ##### | -3.3035 | 9.5494E-04 | 2.3063E-03 |
| ENSG00000159322\|ADPGK | 4503.2263 | -0.4102 | ##### | -3.3031 | 9.5608E-04 | 2.3088E-03 |
| ENSG00000286112\|KYAT1 | 422.7239 | -1.1163 | ##### | -3.3027 | 9.5751E-04 | 2.3121E-03 |
| ENSG00000188818\|ZDHHC11 | 582.4058 | -0.5116 | ##### | -3.3026 | 9.5803E-04 | 2.3132E-03 |
| ENSG00000174899\|SLC66A1L | 11.8096 | -6.3747 | ##### | -3.3025 | 9.5842E-04 | 2.3139E-03 |
| ENSG00000112182\|BACH2 | 76.7305 | 0.7982 | ##### | 3.3019 | 9.6044E-04 | 2.3186E-03 |
| ENSG00000163930\|BAP1 | 7336.5993 | 0.3880 | ##### | 3.3014 | 9.6212E-04 | 2.3223E-03 |
| ENSG00000145365\|TIFA | 766.5968 | -0.4614 | ##### | -3.3014 | 9.6212E-04 | 2.3223E-03 |
| ENSG00000259865 | 335.0558 | -0.5718 | ##### | -3.3013 | 9.6238E-04 | 2.3227E-03 |
| ENSG00000155592\|ZKSCAN2 | 1347.1703 | 0.4293 | ##### | 3.3004 | 9.6559E-04 | 2.3303E-03 |
| ENSG00000101310\|SEC23B | 7815.1774 | -0.3812 | ##### | -3.2998 | 9.6748E-04 | 2.3347E-03 |
| ENSG00000259945 | 62.7524 | -0.7749 | ##### | -3.2994 | 9.6892E-04 | 2.3380E-03 |
| ENSG00000170871\|KIAA0232 | 3448.0127 | -0.3993 | ##### | -3.2991 | 9.6982E-04 | 2.3399E-03 |
| ENSG00000212807\|OR2A42 | 9.2348 | -3.5162 | ##### | -3.2990 | 9.7023E-04 | 2.3407E-03 |
| ENSG00000114547\|ROPN1B | 25.4274 | 2.6324 | ##### | 3.2987 | 9.7138E-04 | 2.3433E-03 |
| ENSG00000173875\|ZNF791 | 1474.7892 | 0.4321 | ##### | 3.2982 | 9.7302E-04 | 2.3471E-03 |
| ENSG00000291006 | 460.6704 | -0.5376 | ##### | -3.2977 | 9.7477E-04 | 2.3511E-03 |
| ENSG00000255741 | 15.0603 | 1.8873 | ##### | 3.2969 | 9.7753E-04 | 2.3576E-03 |
| ENSG00000243449\|C4orf48 | 1616.3462 | 0.4284 | ##### | 3.2969 | 9.7767E-04 | 2.3577E-03 |
| ENSG00000225981\|MICALL2-DT | 27.5663 | 1.1470 | ##### | 3.2954 | 9.8267E-04 | 2.3696E-03 |
| ENSG00000199751\|Y_RNA | 6.2875 | -6.0595 | ##### | -3.2953 | 9.8309E-04 | 2.3704E-03 |
| ENSG00000141446\|ESCO1 | 2396.3227 | 0.4222 | ##### | 3.2948 | 9.8507E-04 | 2.3750E-03 |
| ENSG00000254300\|LINC01111 | 12.2407 | -5.0208 | ##### | -3.2947 | 9.8543E-04 | 2.3757E-03 |
| ENSG00000130711\|PRDM12 | 49.4980 | 1.3530 | ##### | 3.2946 | 9.8574E-04 | 2.3762E-03 |
| ENSG00000281357\|ARRDC3-AS1 | 69.0330 | -1.1675 | ##### | -3.2941 | 9.8724E-04 | 2.3796E-03 |
| ENSG00000137500\|CCDC90B | 2875.8760 | -0.4228 | ##### | -3.2938 | 9.8836E-04 | 2.3822E-03 |
| ENSG00000106483\|SFRP4 | 42.5650 | -4.6853 | ##### | -3.2930 | 9.9134E-04 | 2.3892E-03 |
| ENSG00000197136\|PCNX3 | 7302.4292 | 0.3909 | ##### | 3.2922 | 9.9389E-04 | 2.3951E-03 |
| ENSG00000071889\|FAM3A | 2559.4570 | 0.4096 | ##### | 3.2913 | 9.9721E-04 | 2.4029E-03 |
| ENSG00000198648\|STK39 | 2775.7225 | -0.3982 | ##### | -3.2911 | 9.9788E-04 | 2.4043E-03 |
| ENSG00000263873\|THY1-AS1 | 11.9915 | -4.4331 | ##### | -3.2911 | 9.9800E-04 | 2.4044E-03 |
| ENSG00000100324\|TAB1 | 1736.7478 | -0.3858 | ##### | -3.2909 | 9.9868E-04 | 2.4059E-03 |
| ENSG00000233577 | 11.7181 | 6.3283 | ##### | 3.2907 | 9.9934E-04 | 2.4073E-03 |
| ENSG00000291121 | 30.6569 | 1.1562 | ##### | 3.2905 | 1.0002E-03 | 2.4090E-03 |
| ENSG00000241095\|CYP51A1P1 | 16.7133 | -1.6516 | ##### | -3.2891 | 1.0052E-03 | 2.4210E-03 |
| ENSG00000255330 | 285.4273 | 0.9736 | ##### | 3.2887 | 1.0064E-03 | 2.4236E-03 |
| ENSG00000185122\|HSF1 | 14241.4945 | 0.2999 | ##### | 3.2883 | 1.0081E-03 | 2.4276E-03 |
| ENSG00000126767\|ELK1 | 3651.2165 | 0.4048 | ##### | 3.2879 | 1.0095E-03 | 2.4308E-03 |
| ENSG00000293223 | 10.7935 | -6.3528 | ##### | -3.2874 | 1.0111E-03 | 2.4345E-03 |
| ENSG00000221930\|DENND10P1 | 133.9064 | -0.9674 | ##### | -3.2871 | 1.0124E-03 | 2.4374E-03 |
| ENSG00000223510\|CDRT15 | 9.7964 | 3.9076 | ##### | 3.2864 | 1.0146E-03 | 2.4425E-03 |
| ENSG00000273485 | 76.1739 | 0.8324 | ##### | 3.2860 | 1.0162E-03 | 2.4461E-03 |
| ENSG00000275703 | 10.4799 | -3.3323 | ##### | -3.2860 | 1.0164E-03 | 2.4463E-03 |
| ENSG00000186940\|CHCHD2P9 | 40.2281 | 0.9773 | ##### | 3.2858 | 1.0168E-03 | 2.4472E-03 |
| ENSG00000110693\|SOX6 | 17.6540 | -1.4834 | ##### | -3.2857 | 1.0175E-03 | 2.4486E-03 |

| | | | | | | |
|---|---|---|---|---|---|---|
| ENSG00000132467\|UTP3 | 3662.4168 | 0.4097 | ##### | 3.2856 | 1.0178E-03 | 2.4492E-03 |
| ENSG00000117758\|STX12 | 1936.5370 | -0.4285 | ##### | -3.2853 | 1.0188E-03 | 2.4513E-03 |
| ENSG00000269925 | 24.4603 | -1.3879 | ##### | -3.2839 | 1.0238E-03 | 2.4631E-03 |
| ENSG00000200152\|RNU6-216P | 7.6948 | 2.7711 | ##### | 3.2834 | 1.0255E-03 | 2.4670E-03 |
| ENSG00000166598\|HSP90B1 | 72509.5718 | 0.1842 | ##### | 3.2814 | 1.0329E-03 | 2.4847E-03 |
| ENSG00000285130 | 646.9954 | -3.0983 | ##### | -3.2812 | 1.0336E-03 | 2.4861E-03 |
| ENSG00000286964 | 67.7711 | 1.0597 | ##### | 3.2812 | 1.0337E-03 | 2.4861E-03 |
| ENSG00000142765\|SYTL1 | 62.9028 | -0.8127 | ##### | -3.2811 | 1.0340E-03 | 2.4867E-03 |
| ENSG00000273363 | 38.1678 | 7.5487 | ##### | 3.2811 | 1.0340E-03 | 2.4867E-03 |
| ENSG00000256628\|ZBTB11-AS1 | 687.3399 | 0.4593 | ##### | 3.2799 | 1.0383E-03 | 2.4966E-03 |
| ENSG00000286379 | 41.1139 | -2.2311 | ##### | -3.2797 | 1.0391E-03 | 2.4984E-03 |
| ENSG00000111726\|CMAS | 6736.1965 | 0.3938 | ##### | 3.2796 | 1.0394E-03 | 2.4988E-03 |
| ENSG00000156968\|MPV17L | 1814.2248 | -0.4285 | ##### | -3.2796 | 1.0395E-03 | 2.4989E-03 |
| ENSG00000223749\|MIR503HG | 351.9016 | 0.5845 | ##### | 3.2796 | 1.0397E-03 | 2.4991E-03 |
| ENSG00000286873 | 47.0085 | -1.6112 | ##### | -3.2795 | 1.0399E-03 | 2.4994E-03 |
| ENSG00000158258\|CLSTN2 | 22.4969 | -2.5715 | ##### | -3.2795 | 1.0399E-03 | 2.4994E-03 |
| ENSG00000234818 | 28.1929 | 1.0516 | ##### | 3.2791 | 1.0414E-03 | 2.5027E-03 |
| ENSG00000180817\|PPA1 | 9283.8135 | -0.3955 | ##### | -3.2785 | 1.0437E-03 | 2.5080E-03 |
| ENSG00000290695 | 26.0114 | 4.1309 | ##### | 3.2781 | 1.0450E-03 | 2.5109E-03 |
| ENSG00000221886\|FAM200C | 444.4184 | 0.5298 | ##### | 3.2778 | 1.0462E-03 | 2.5136E-03 |
| ENSG00000260121 | 777.4003 | 0.5280 | ##### | 3.2774 | 1.0477E-03 | 2.5171E-03 |
| ENSG00000151135\|TMEM263 | 6669.3487 | -0.4002 | ##### | -3.2764 | 1.0512E-03 | 2.5253E-03 |
| ENSG00000198780\|FAM169A | 1045.9597 | -0.4276 | ##### | -3.2755 | 1.0546E-03 | 2.5333E-03 |
| ENSG00000143178\|TBX19 | 117.9121 | -0.7586 | ##### | -3.2750 | 1.0567E-03 | 2.5382E-03 |
| ENSG00000108179\|PPIF | 7608.9941 | -0.4039 | ##### | -3.2743 | 1.0591E-03 | 2.5437E-03 |
| ENSG00000198003\|ODAD3 | 142.6171 | 0.6571 | ##### | 3.2740 | 1.0603E-03 | 2.5463E-03 |
| ENSG00000279130 | 12.0776 | -2.8609 | ##### | -3.2738 | 1.0609E-03 | 2.5476E-03 |
| ENSG00000262902\|MTCO1P40 | 23.6006 | 1.1957 | ##### | 3.2737 | 1.0615E-03 | 2.5489E-03 |
| ENSG00000128271\|ADORA2A | 123.5111 | -2.3098 | ##### | -3.2733 | 1.0629E-03 | 2.5519E-03 |
| ENSG00000143793\|C1orf35 | 1771.2304 | -0.4370 | ##### | -3.2732 | 1.0634E-03 | 2.5529E-03 |
| ENSG00000092439\|TRPM7 | 4519.1454 | -0.4009 | ##### | -3.2728 | 1.0649E-03 | 2.5564E-03 |
| ENSG00000275895 | 4133.1640 | 0.3617 | ##### | 3.2723 | 1.0666E-03 | 2.5602E-03 |
| ENSG00000231443\|PPP4R2P2 | 16.3868 | -2.1342 | ##### | -3.2721 | 1.0673E-03 | 2.5618E-03 |
| ENSG00000272944 | 18.6476 | 2.0471 | ##### | 3.2715 | 1.0698E-03 | 2.5675E-03 |
| ENSG00000077616\|NAALAD2 | 70.5385 | 1.4283 | ##### | 3.2706 | 1.0732E-03 | 2.5754E-03 |
| ENSG00000253986 | 4.9261 | 3.8465 | ##### | 3.2701 | 1.0749E-03 | 2.5793E-03 |
| ENSG00000273891 | 58.1044 | -1.2862 | ##### | -3.2697 | 1.0768E-03 | 2.5836E-03 |
| ENSG00000245937\|SLC12A2-DT | 1663.6080 | -0.4305 | ##### | -3.2696 | 1.0770E-03 | 2.5839E-03 |
| ENSG00000204963\|PCDHA7 | 57.9724 | -4.9614 | ##### | -3.2695 | 1.0775E-03 | 2.5849E-03 |
| ENSG00000130876\|SLC7A10 | 14.4527 | 2.6535 | ##### | 3.2691 | 1.0789E-03 | 2.5881E-03 |
| ENSG00000167283\|ATP5MG | 11481.9954 | 0.3834 | ##### | 3.2687 | 1.0804E-03 | 2.5914E-03 |
| ENSG00000086159\|AQP6 | 30.1022 | 1.3369 | ##### | 3.2685 | 1.0813E-03 | 2.5933E-03 |
| ENSG00000255201 | 56.3770 | -1.2111 | ##### | -3.2683 | 1.0819E-03 | 2.5947E-03 |
| ENSG00000185024\|BRF1 | 2831.3294 | 0.3979 | ##### | 3.2682 | 1.0824E-03 | 2.5957E-03 |
| ENSG00000206633\|SNORA80B | 7.3761 | 2.7247 | ##### | 3.2680 | 1.0832E-03 | 2.5973E-03 |
| ENSG00000230532 | 21.4855 | 1.2004 | ##### | 3.2678 | 1.0839E-03 | 2.5988E-03 |
| ENSG00000147457\|CHMP7 | 4290.8620 | 0.4179 | ##### | 3.2671 | 1.0865E-03 | 2.6048E-03 |
| ENSG00000064692\|SNCAIP | 20.5406 | -1.7104 | ##### | -3.2667 | 1.0881E-03 | 2.6086E-03 |
| ENSG00000272008 | 34.8730 | -1.8443 | ##### | -3.2658 | 1.0916E-03 | 2.6166E-03 |
| ENSG00000067798\|NAV3 | 1594.2403 | 0.4671 | ##### | 3.2658 | 1.0917E-03 | 2.6166E-03 |
| ENSG00000019144\|PHLDB1 | 3523.7453 | 0.4002 | ##### | 3.2656 | 1.0922E-03 | 2.6177E-03 |
| ENSG00000234912\|SNHG20 | 664.6640 | 0.4782 | ##### | 3.2651 | 1.0942E-03 | 2.6221E-03 |
| ENSG00000153201\|RANBP2 | 14368.9716 | -0.3953 | ##### | -3.2651 | 1.0942E-03 | 2.6221E-03 |
| ENSG00000256442 | 20.6073 | 5.6533 | ##### | 3.2650 | 1.0945E-03 | 2.6225E-03 |
| ENSG00000253570\|RNF5P1 | 12.1831 | -3.0906 | ##### | -3.2645 | 1.0966E-03 | 2.6274E-03 |
| ENSG00000198824\|CHAMP1 | 2546.4083 | -0.4125 | ##### | -3.2643 | 1.0973E-03 | 2.6289E-03 |
| ENSG00000072518\|MARK2 | 4178.5026 | 0.3883 | ##### | 3.2638 | 1.0994E-03 | 2.6337E-03 |
| ENSG00000249158\|PCDHA11 | 490.6031 | -0.4794 | ##### | -3.2637 | 1.0996E-03 | 2.6340E-03 |
| ENSG00000226530 | 16.0850 | 2.6715 | ##### | 3.2636 | 1.1002E-03 | 2.6352E-03 |
| ENSG00000230251\|PHB1P4 | 21.3328 | -1.7612 | ##### | -3.2633 | 1.1011E-03 | 2.6370E-03 |
| ENSG00000090581\|GNPTG | 4589.3295 | -0.3934 | ##### | -3.2622 | 1.1055E-03 | 2.6475E-03 |
| ENSG00000147381\|MAGEA4 | 23884.6309 | 0.3415 | ##### | 3.2621 | 1.1059E-03 | 2.6481E-03 |
| ENSG00000169223\|LMAN2 | 11688.5097 | 0.3928 | ##### | 3.2614 | 1.1085E-03 | 2.6542E-03 |

| | | | | | | |
|---|---|---|---|---|---|---|
| ENSG00000287260 | 32.5982 | -1.9753 | ##### | -3.2596 | 1.1159E-03 | 2.6716E-03 |
| ENSG00000279069 | 797.4898 | 0.5096 | ##### | 3.2576 | 1.1238E-03 | 2.6903E-03 |
| ENSG00000275734 | 95.0734 | -0.8012 | ##### | -3.2575 | 1.1239E-03 | 2.6903E-03 |
| ENSG00000291039 | 28.2609 | 2.3599 | ##### | 3.2574 | 1.1242E-03 | 2.6907E-03 |
| ENSG00000225032 | 73.3909 | -0.7586 | ##### | -3.2575 | 1.1241E-03 | 2.6907E-03 |
| ENSG00000224621 | 27.7098 | 1.4972 | ##### | 3.2563 | 1.1289E-03 | 2.7017E-03 |
| ENSG00000233221\|ARHGEF4-AS1 | 32.5109 | 3.3799 | ##### | 3.2557 | 1.1311E-03 | 2.7067E-03 |
| ENSG00000251634\|NAIPP4 | 84.1237 | 0.7430 | ##### | 3.2557 | 1.1313E-03 | 2.7071E-03 |
| ENSG00000174473\|GALNTL6 | 57.6170 | -1.2057 | ##### | -3.2540 | 1.1380E-03 | 2.7228E-03 |
| ENSG00000205362\|MT1A | 211.0904 | -0.6993 | ##### | -3.2536 | 1.1397E-03 | 2.7267E-03 |
| ENSG00000106328\|FSCN3 | 12.5096 | 3.2556 | ##### | 3.2533 | 1.1407E-03 | 2.7288E-03 |
| ENSG00000232530\|LIF-AS1 | 24.5558 | 3.2958 | ##### | 3.2531 | 1.1416E-03 | 2.7307E-03 |
| ENSG00000157045\|NTAN1 | 1940.2570 | 0.4418 | ##### | 3.2524 | 1.1445E-03 | 2.7375E-03 |
| ENSG00000293311\|SMPD4BP | 1659.4571 | -0.4342 | ##### | -3.2520 | 1.1459E-03 | 2.7407E-03 |
| ENSG00000079482\|OPHN1 | 660.0336 | 0.4820 | ##### | 3.2517 | 1.1472E-03 | 2.7435E-03 |
| ENSG00000256982 | 28.9721 | -1.4063 | ##### | -3.2516 | 1.1477E-03 | 2.7445E-03 |
| ENSG00000114354\|TFG | 14929.1990 | -0.4112 | ##### | -3.2512 | 1.1493E-03 | 2.7481E-03 |
| ENSG00000145358\|DDIT4L | 86.3927 | -0.9001 | ##### | -3.2506 | 1.1514E-03 | 2.7529E-03 |
| ENSG00000286942 | 23.4439 | 7.0133 | ##### | 3.2502 | 1.1531E-03 | 2.7568E-03 |
| ENSG00000270469 | 10.9294 | -6.3701 | ##### | -3.2501 | 1.1538E-03 | 2.7581E-03 |
| ENSG00000166484\|MAPK7 | 1967.3835 | 0.4276 | ##### | 3.2498 | 1.1549E-03 | 2.7606E-03 |
| ENSG00000253816\|GUSBP14 | 168.0757 | 0.7175 | ##### | 3.2496 | 1.1558E-03 | 2.7626E-03 |
| ENSG00000288982 | 11.7230 | -6.3403 | ##### | -3.2492 | 1.1572E-03 | 2.7656E-03 |
| ENSG00000286786 | 12.4956 | 1.6476 | ##### | 3.2487 | 1.1595E-03 | 2.7709E-03 |
| ENSG00000280417 | 146.6605 | 0.6318 | ##### | 3.2478 | 1.1630E-03 | 2.7790E-03 |
| ENSG00000174680\|GRIK1-AS1 | 19.8666 | 7.5804 | ##### | 3.2474 | 1.1648E-03 | 2.7833E-03 |
| ENSG00000249493\|ANKRD20A18P | 20.2993 | -7.2714 | ##### | -3.2472 | 1.1654E-03 | 2.7843E-03 |
| ENSG00000267191\|ZNF45-AS1 | 34.2302 | -1.8861 | ##### | -3.2457 | 1.1716E-03 | 2.7990E-03 |
| ENSG00000145675\|PIK3R1 | 1212.8836 | -0.4442 | ##### | -3.2444 | 1.1769E-03 | 2.8113E-03 |
| ENSG00000167461\|RAB8A | 6810.8954 | 0.3890 | ##### | 3.2444 | 1.1771E-03 | 2.8118E-03 |
| ENSG00000269425 | 11.2294 | 3.7654 | ##### | 3.2439 | 1.1791E-03 | 2.8162E-03 |
| ENSG00000161835\|TAMALIN | 101.9638 | 1.0282 | ##### | 3.2437 | 1.1799E-03 | 2.8178E-03 |
| ENSG00000187955\|COL14A1 | 78.1072 | -3.4838 | ##### | -3.2430 | 1.1830E-03 | 2.8250E-03 |
| ENSG00000174123\|TLR10 | 20.3130 | -3.5942 | ##### | -3.2429 | 1.1831E-03 | 2.8251E-03 |
| ENSG00000226695\|ANKRD20A10P | 12.8010 | 2.7922 | ##### | 3.2427 | 1.1840E-03 | 2.8265E-03 |
| ENSG00000124574\|ABCC10 | 2514.5170 | 0.4082 | ##### | 3.2427 | 1.1840E-03 | 2.8265E-03 |
| ENSG00000170632\|ARMC10 | 4653.2313 | 0.3987 | ##### | 3.2427 | 1.1839E-03 | 2.8265E-03 |
| ENSG00000198373\|WWP2 | 3181.9066 | -0.4113 | ##### | -3.2425 | 1.1848E-03 | 2.8283E-03 |
| ENSG00000157764\|BRAF | 2434.1126 | 0.4080 | ##### | 3.2415 | 1.1888E-03 | 2.8376E-03 |
| ENSG00000204196\|RPL12P16 | 481.2210 | 0.5247 | ##### | 3.2413 | 1.1899E-03 | 2.8399E-03 |
| ENSG00000004866\|ST7 | 3016.6373 | 0.4083 | ##### | 3.2409 | 1.1914E-03 | 2.8432E-03 |
| ENSG00000147614\|ATP6V0D2 | 14.9159 | 5.5244 | ##### | 3.2405 | 1.1930E-03 | 2.8470E-03 |
| ENSG00000092208\|GEMIN2 | 1057.1822 | 0.4317 | ##### | 3.2400 | 1.1953E-03 | 2.8521E-03 |
| ENSG00000182768\|NGRN | 10793.1633 | -0.2351 | ##### | -3.2388 | 1.2003E-03 | 2.8638E-03 |
| ENSG00000210140\|MT-TC | 5241.7301 | 0.4206 | ##### | 3.2385 | 1.2014E-03 | 2.8663E-03 |
| ENSG00000262265 | 11.1249 | 5.4292 | ##### | 3.2384 | 1.2019E-03 | 2.8673E-03 |
| ENSG00000261505 | 154.4644 | 0.8834 | ##### | 3.2379 | 1.2040E-03 | 2.8720E-03 |
| ENSG00000205038\|PKHD1L1 | 38.4191 | -7.6797 | ##### | -3.2377 | 1.2050E-03 | 2.8741E-03 |
| ENSG00000171570\|RAB4B-EGLN2 | 755.7708 | -3.2472 | ##### | -3.2377 | 1.2051E-03 | 2.8743E-03 |
| ENSG00000254578 | 45.4352 | -0.9120 | ##### | -3.2375 | 1.2060E-03 | 2.8761E-03 |
| ENSG00000178297\|TMPRSS9 | 78.2101 | -0.7165 | ##### | -3.2374 | 1.2062E-03 | 2.8763E-03 |
| ENSG00000272948 | 16.6262 | 1.5369 | ##### | 3.2365 | 1.2100E-03 | 2.8852E-03 |
| ENSG00000291070\|AOC4P | 7.3608 | -2.8225 | ##### | -3.2363 | 1.2109E-03 | 2.8872E-03 |
| ENSG00000182463\|TSHZ2 | 28.0885 | 1.9404 | ##### | 3.2360 | 1.2120E-03 | 2.8894E-03 |
| ENSG00000096070\|BRPF3 | 3960.3515 | 0.3895 | ##### | 3.2360 | 1.2121E-03 | 2.8895E-03 |
| ENSG00000145536\|ADAMTS16 | 13.9482 | -3.5553 | ##### | -3.2360 | 1.2122E-03 | 2.8895E-03 |
| ENSG00000111860\|CEP85L | 320.6430 | -0.5136 | ##### | -3.2354 | 1.2149E-03 | 2.8957E-03 |
| ENSG00000232472\|EEF1B2P3 | 1077.7258 | 0.4399 | ##### | 3.2350 | 1.2162E-03 | 2.8986E-03 |
| ENSG00000196656 | 1672.5059 | 0.4607 | ##### | 3.2350 | 1.2164E-03 | 2.8988E-03 |
| ENSG00000226330\|PPP1R15B-AS1 | 16.7016 | -3.7934 | ##### | -3.2346 | 1.2179E-03 | 2.9022E-03 |
| ENSG00000089250\|NOS1 | 20.9529 | -4.4063 | ##### | -3.2338 | 1.2216E-03 | 2.9107E-03 |
| ENSG00000103168\|TAF1C | 4944.7985 | 0.4125 | ##### | 3.2338 | 1.2217E-03 | 2.9107E-03 |
| ENSG00000256806\|C17orf100 | 176.4716 | -0.6010 | ##### | -3.2336 | 1.2225E-03 | 2.9123E-03 |

| | | | | | | |
|---|---|---|---|---|---|---|
| ENSG00000100207\|TCF20 | 5736.1253 | 0.3900 | ##### | 3.2329 | 1.2253E-03 | 2.9189E-03 |
| ENSG00000166411\|IDH3A | 5438.7007 | 0.3778 | ##### | 3.2326 | 1.2265E-03 | 2.9214E-03 |
| ENSG00000156304\|SCAF4 | 3246.5138 | 0.4074 | ##### | 3.2325 | 1.2271E-03 | 2.9228E-03 |
| ENSG00000212694\|LINC01089 | 456.1831 | 0.5139 | ##### | 3.2324 | 1.2276E-03 | 2.9235E-03 |
| ENSG00000167523\|SPATA33 | 778.9949 | 0.4522 | ##### | 3.2324 | 1.2276E-03 | 2.9235E-03 |
| ENSG00000236130\|PTCSC2 | 18.0185 | 3.6212 | ##### | 3.2320 | 1.2293E-03 | 2.9273E-03 |
| ENSG00000218018\|RBM38-AS1 | 37.8410 | 2.1044 | ##### | 3.2308 | 1.2346E-03 | 2.9396E-03 |
| ENSG00000280269 | 17.9382 | 5.9392 | ##### | 3.2301 | 1.2373E-03 | 2.9457E-03 |
| ENSG00000141232\|TOB1 | 5303.4740 | -0.4022 | ##### | -3.2298 | 1.2388E-03 | 2.9491E-03 |
| ENSG00000280152 | 71.3616 | 1.0320 | ##### | 3.2297 | 1.2390E-03 | 2.9495E-03 |
| ENSG00000139193\|CD27 | 22.5230 | 2.2068 | ##### | 3.2293 | 1.2407E-03 | 2.9533E-03 |
| ENSG00000136925\|TSTD2 | 1888.9458 | -0.4189 | ##### | -3.2278 | 1.2476E-03 | 2.9693E-03 |
| ENSG00000149308\|NPAT | 1445.3699 | 0.4112 | ##### | 3.2272 | 1.2499E-03 | 2.9745E-03 |
| ENSG00000267365\|KCNJ2-AS1 | 52.0664 | -2.4807 | ##### | -3.2270 | 1.2512E-03 | 2.9774E-03 |
| ENSG00000180043\|GARIN5B | 23.3780 | -7.4731 | ##### | -3.2267 | 1.2523E-03 | 2.9798E-03 |
| ENSG00000228649\|SNHG26 | 483.0954 | 0.5487 | ##### | 3.2263 | 1.2539E-03 | 2.9833E-03 |
| ENSG00000111653\|ING4 | 978.1437 | 0.4321 | ##### | 3.2262 | 1.2545E-03 | 2.9845E-03 |
| ENSG00000155016\|CYP2U1 | 304.1353 | -0.5183 | ##### | -3.2254 | 1.2578E-03 | 2.9921E-03 |
| ENSG00000007541\|PIGQ | 3244.1315 | -0.3916 | ##### | -3.2254 | 1.2579E-03 | 2.9923E-03 |
| ENSG00000154845\|PPP4R1 | 12350.3344 | 0.3033 | ##### | 3.2252 | 1.2589E-03 | 2.9945E-03 |
| ENSG00000250365\|HEATR5A-DT | 715.5147 | -0.4626 | ##### | -3.2244 | 1.2622E-03 | 3.0020E-03 |
| ENSG00000233328\|PFN1P1 | 266.3211 | 0.5164 | ##### | 3.2241 | 1.2638E-03 | 3.0055E-03 |
| ENSG00000144045\|DQX1 | 41.5908 | 1.9208 | ##### | 3.2231 | 1.2680E-03 | 3.0154E-03 |
| ENSG00000229388\|TAF12-DT | 33.4091 | 1.2767 | ##### | 3.2223 | 1.2715E-03 | 3.0234E-03 |
| ENSG00000225096 | 37.6298 | 1.4213 | ##### | 3.2219 | 1.2735E-03 | 3.0278E-03 |
| ENSG00000269967 | 9.9651 | 5.3836 | ##### | 3.2215 | 1.2753E-03 | 3.0318E-03 |
| ENSG00000204323\|SMIM5 | 27.7170 | -1.8883 | ##### | -3.2214 | 1.2758E-03 | 3.0328E-03 |
| ENSG00000170703\|TTLL6 | 26.8616 | -2.4691 | ##### | -3.2212 | 1.2766E-03 | 3.0344E-03 |
| ENSG00000258780\|BMS1P15 | 9.1957 | -6.3567 | ##### | -3.2210 | 1.2773E-03 | 3.0359E-03 |
| ENSG00000114554\|PLXNA1 | 13805.3327 | 0.3856 | ##### | 3.2207 | 1.2790E-03 | 3.0397E-03 |
| ENSG00000142937\|RPS8 | 45401.9165 | -0.3925 | ##### | -3.2196 | 1.2837E-03 | 3.0506E-03 |
| ENSG00000108786\|HSD17B1 | 876.3955 | 0.4723 | ##### | 3.2196 | 1.2838E-03 | 3.0507E-03 |
| ENSG00000261997 | 7.8743 | 2.9990 | ##### | 3.2193 | 1.2851E-03 | 3.0535E-03 |
| ENSG00000267353 | 44.3665 | -1.0548 | ##### | -3.2184 | 1.2892E-03 | 3.0629E-03 |
| ENSG00000270804\|UBE2CP5 | 21.4863 | -1.7404 | ##### | -3.2181 | 1.2903E-03 | 3.0654E-03 |
| ENSG00000132386\|SERPINF1 | 38.5665 | 1.1222 | ##### | 3.2179 | 1.2912E-03 | 3.0672E-03 |
| ENSG00000120913\|PDLIM2 | 1827.8405 | -0.4213 | ##### | -3.2176 | 1.2925E-03 | 3.0702E-03 |
| ENSG00000163354\|DCST2 | 68.9421 | 0.8000 | ##### | 3.2174 | 1.2934E-03 | 3.0720E-03 |
| ENSG00000214022\|REPIN1 | 15244.5688 | 0.2923 | ##### | 3.2172 | 1.2945E-03 | 3.0745E-03 |
| ENSG00000183628\|DGCR6 | 158.6335 | -0.8028 | ##### | -3.2169 | 1.2957E-03 | 3.0769E-03 |
| ENSG00000138376\|BARD1 | 2121.5630 | 0.3967 | ##### | 3.2162 | 1.2991E-03 | 3.0848E-03 |
| ENSG00000242553 | 23.3323 | -1.7715 | ##### | -3.2162 | 1.2992E-03 | 3.0848E-03 |
| ENSG00000231711\|LINC00899 | 13.5843 | -3.8317 | ##### | -3.2157 | 1.3013E-03 | 3.0896E-03 |
| ENSG00000215417\|MIR17HG | 211.4570 | 1.2261 | ##### | 3.2145 | 1.3068E-03 | 3.1023E-03 |
| ENSG00000119318\|RAD23B | 19457.2783 | -0.2936 | ##### | -3.2145 | 1.3069E-03 | 3.1023E-03 |
| ENSG00000176381\|PRR18 | 16.4491 | 3.5383 | ##### | 3.2134 | 1.3116E-03 | 3.1133E-03 |
| ENSG00000172366\|MCRIP2 | 1563.8721 | -0.4124 | ##### | -3.2130 | 1.3134E-03 | 3.1172E-03 |
| ENSG00000134317\|GRHL1 | 169.8200 | 0.7404 | ##### | 3.2120 | 1.3180E-03 | 3.1279E-03 |
| ENSG00000238103\|RPL9P7 | 1764.3955 | -0.4338 | ##### | -3.2118 | 1.3192E-03 | 3.1306E-03 |
| ENSG00000133250\|ZNF414 | 706.0992 | -0.4511 | ##### | -3.2115 | 1.3205E-03 | 3.1333E-03 |
| ENSG00000213080 | 170.6467 | 0.6286 | ##### | 3.2114 | 1.3208E-03 | 3.1337E-03 |
| ENSG00000204217\|BMPR2 | 5840.2471 | 0.4005 | ##### | 3.2104 | 1.3253E-03 | 3.1443E-03 |
| ENSG00000103264\|FBXO31 | 2913.0334 | 0.4108 | ##### | 3.2096 | 1.3293E-03 | 3.1536E-03 |
| ENSG00000244349 | 21.5180 | 1.2664 | ##### | 3.2091 | 1.3315E-03 | 3.1584E-03 |
| ENSG00000289201 | 27.3173 | -1.6492 | ##### | -3.2085 | 1.3345E-03 | 3.1653E-03 |
| ENSG00000163319\|MRPS18C | 3988.2434 | 0.4155 | ##### | 3.2082 | 1.3356E-03 | 3.1676E-03 |
| ENSG00000178852\|EFCAB13 | 245.3448 | 0.5715 | ##### | 3.2076 | 1.3383E-03 | 3.1738E-03 |
| ENSG00000274340 | 11.5313 | -5.6269 | ##### | -3.2076 | 1.3386E-03 | 3.1743E-03 |
| ENSG00000204977\|TRIM13 | 1990.7578 | -0.4216 | ##### | -3.2071 | 1.3406E-03 | 3.1788E-03 |
| ENSG00000224505\|HEXIM2-AS1 | 76.5230 | -1.0195 | ##### | -3.2066 | 1.3433E-03 | 3.1849E-03 |
| ENSG00000260853\|NFATC2IP-AS1 | 343.1875 | 0.5582 | ##### | 3.2063 | 1.3447E-03 | 3.1881E-03 |
| ENSG00000228192\|ZNF691-DT | 153.2213 | -0.6695 | ##### | -3.2061 | 1.3453E-03 | 3.1892E-03 |
| ENSG00000280388 | 13.3494 | 1.6846 | ##### | 3.2058 | 1.3469E-03 | 3.1926E-03 |

| | | | | | | |
|---|---|---|---|---|---|---|
| ENSG00000232019 | 8.1045 | -5.6598 | ##### | -3.2056 | 1.3479E-03 | 3.1949E-03 |
| ENSG00000039650\|PNKP | 3129.2526 | -0.4080 | ##### | -3.2052 | 1.3498E-03 | 3.1990E-03 |
| ENSG00000240376 | 510.0897 | -0.4596 | ##### | -3.2050 | 1.3507E-03 | 3.2011E-03 |
| ENSG00000226510\|UPK1A-AS1 | 39.6749 | 1.0727 | ##### | 3.2048 | 1.3514E-03 | 3.2023E-03 |
| ENSG00000226355 | 13.2077 | -5.7610 | ##### | -3.2046 | 1.3524E-03 | 3.2046E-03 |
| ENSG00000229689\|ANKRD33BP1 | 165.9822 | 1.6398 | ##### | 3.2043 | 1.3538E-03 | 3.2075E-03 |
| ENSG00000135823\|STX6 | 3395.5337 | -0.3870 | ##### | -3.2040 | 1.3553E-03 | 3.2109E-03 |
| ENSG00000278775 | 8.2718 | 4.1616 | ##### | 3.2037 | 1.3567E-03 | 3.2138E-03 |
| ENSG00000233974 | 11.7513 | 3.7458 | ##### | 3.2035 | 1.3577E-03 | 3.2161E-03 |
| ENSG00000141052\|MYOCD | 757.5075 | 0.4640 | ##### | 3.2028 | 1.3612E-03 | 3.2241E-03 |
| ENSG00000272335 | 323.8810 | 0.5619 | ##### | 3.2024 | 1.3630E-03 | 3.2278E-03 |
| ENSG00000171044\|XKR6 | 132.8791 | 0.6762 | ##### | 3.2024 | 1.3630E-03 | 3.2278E-03 |
| ENSG00000234373\|SNX18P7 | 30.7472 | -1.1602 | ##### | -3.2024 | 1.3631E-03 | 3.2278E-03 |
| ENSG00000140830\|TXNL4B | 1680.8607 | -0.4077 | ##### | -3.2021 | 1.3642E-03 | 3.2302E-03 |
| ENSG00000155287\|SLC25A28 | 1713.2085 | -0.4111 | ##### | -3.2018 | 1.3659E-03 | 3.2340E-03 |
| ENSG00000286669 | 16.5764 | -2.3378 | ##### | -3.2015 | 1.3674E-03 | 3.2371E-03 |
| ENSG00000162959\|MEMO1 | 4654.8918 | 0.4107 | ##### | 3.2009 | 1.3702E-03 | 3.2435E-03 |
| ENSG00000138439\|FAM117B | 1357.3599 | 0.4062 | ##### | 3.2006 | 1.3716E-03 | 3.2467E-03 |
| ENSG00000267480\|LINC02979 | 68.1045 | -0.8231 | ##### | -3.2003 | 1.3727E-03 | 3.2489E-03 |
| ENSG00000131153\|GINS2 | 2843.6750 | 0.4012 | ##### | 3.1997 | 1.3755E-03 | 3.2554E-03 |
| ENSG00000139793\|MBNL2 | 3432.2798 | 0.3997 | ##### | 3.1996 | 1.3761E-03 | 3.2564E-03 |
| ENSG00000144369\|FAM171B | 952.4723 | 0.4270 | ##### | 3.1985 | 1.3813E-03 | 3.2685E-03 |
| ENSG00000232448 | 17.0982 | -3.2603 | ##### | -3.1984 | 1.3818E-03 | 3.2694E-03 |
| ENSG00000168899\|VAMP5 | 17.6351 | -2.0454 | ##### | -3.1983 | 1.3822E-03 | 3.2702E-03 |
| ENSG00000278576 | 15.3687 | 1.7016 | ##### | 3.1983 | 1.3824E-03 | 3.2704E-03 |
| ENSG00000074803\|SLC12A1 | 19.3435 | 7.3920 | ##### | 3.1982 | 1.3828E-03 | 3.2711E-03 |
| ENSG00000100664\|EIF5 | 24042.3606 | 0.3105 | ##### | 3.1980 | 1.3838E-03 | 3.2732E-03 |
| ENSG00000264522\|OTUD7B | 3537.7766 | 0.3192 | ##### | 3.1979 | 1.3843E-03 | 3.2742E-03 |
| ENSG00000184206\|GOLGA6L4 | 49.9910 | -0.8187 | ##### | -3.1973 | 1.3874E-03 | 3.2813E-03 |
| ENSG00000204310\|AGPAT1 | 11739.3836 | 0.3908 | ##### | 3.1962 | 1.3927E-03 | 3.2934E-03 |
| ENSG00000232389 | 262.0916 | 0.5446 | ##### | 3.1960 | 1.3937E-03 | 3.2955E-03 |
| ENSG00000224790\|HLCS-AS1 | 63.5753 | -1.1405 | ##### | -3.1950 | 1.3983E-03 | 3.3061E-03 |
| ENSG00000234286 | 29.6546 | -7.3256 | ##### | -3.1950 | 1.3984E-03 | 3.3061E-03 |
| ENSG00000178307\|TMEM11 | 2394.5169 | 0.4027 | ##### | 3.1948 | 1.3995E-03 | 3.3084E-03 |
| ENSG00000226976\|COX6A1P2 | 180.3792 | 0.5977 | ##### | 3.1947 | 1.3998E-03 | 3.3089E-03 |
| ENSG00000254996\|ANKHD1-EIF4EBP3 | 4421.8228 | 1.5502 | ##### | 3.1944 | 1.4012E-03 | 3.3120E-03 |
| ENSG00000101443\|WFDC2 | 57.9756 | 3.0960 | ##### | 3.1944 | 1.4013E-03 | 3.3120E-03 |
| ENSG00000234602\|MCIDAS | 380.9584 | -0.4883 | ##### | -3.1938 | 1.4039E-03 | 3.3178E-03 |
| ENSG00000167004\|PDIA3 | 28517.3087 | -0.3001 | ##### | -3.1926 | 1.4102E-03 | 3.3324E-03 |
| ENSG00000283265 | 10.7709 | -5.8302 | ##### | -3.1922 | 1.4120E-03 | 3.3365E-03 |
| ENSG00000267439 | 133.6044 | 0.9227 | ##### | 3.1906 | 1.4199E-03 | 3.3548E-03 |
| ENSG00000270482 | 15.0411 | -5.9705 | ##### | -3.1904 | 1.4209E-03 | 3.3570E-03 |
| ENSG00000290916 | 26.0595 | 2.7507 | ##### | 3.1901 | 1.4223E-03 | 3.3600E-03 |
| ENSG00000177854\|TMEM187 | 426.8698 | -0.5181 | ##### | -3.1900 | 1.4229E-03 | 3.3611E-03 |
| ENSG00000246640\|PICART1 | 72.5256 | -0.9221 | ##### | -3.1898 | 1.4239E-03 | 3.3633E-03 |
| ENSG00000272702 | 34.6819 | 1.2347 | ##### | 3.1896 | 1.4249E-03 | 3.3652E-03 |
| ENSG00000288913 | 94.5852 | -0.7000 | ##### | -3.1883 | 1.4311E-03 | 3.3794E-03 |
| ENSG00000168913\|ENHO | 113.0028 | 0.8559 | ##### | 3.1883 | 1.4311E-03 | 3.3794E-03 |
| ENSG00000144228\|SPOPL | 3052.3745 | -0.3924 | ##### | -3.1882 | 1.4315E-03 | 3.3802E-03 |
| ENSG00000278917 | 59.8989 | -0.8581 | ##### | -3.1877 | 1.4340E-03 | 3.3856E-03 |
| ENSG00000110060\|PUS3 | 674.3354 | -0.4713 | ##### | -3.1873 | 1.4362E-03 | 3.3907E-03 |
| ENSG00000223718\|RPL15P11 | 5.7189 | 3.6027 | ##### | 3.1871 | 1.4369E-03 | 3.3921E-03 |
| ENSG00000188725\|SMIM15 | 6135.6505 | 0.2973 | ##### | 3.1867 | 1.4389E-03 | 3.3966E-03 |
| ENSG00000207313\|SNORA2B | 5.3222 | 3.5804 | ##### | 3.1866 | 1.4396E-03 | 3.3979E-03 |
| ENSG00000167011\|NAT16 | 27.1888 | -1.0911 | ##### | -3.1863 | 1.4409E-03 | 3.4008E-03 |
| ENSG00000173338\|KCNK7 | 32.9664 | 1.0311 | ##### | 3.1861 | 1.4420E-03 | 3.4031E-03 |
| ENSG00000016391\|CHDH | 595.2258 | 0.4248 | ##### | 3.1861 | 1.4422E-03 | 3.4032E-03 |
| ENSG00000260498 | 9.9274 | -3.1949 | ##### | -3.1861 | 1.4423E-03 | 3.4032E-03 |
| ENSG00000261270 | 18.7957 | -1.3958 | ##### | -3.1857 | 1.4439E-03 | 3.4068E-03 |
| ENSG00000260918 | 13.2962 | -5.1175 | ##### | -3.1853 | 1.4460E-03 | 3.4113E-03 |
| ENSG00000063241\|ISOC2 | 3362.9229 | -0.3877 | ##### | -3.1853 | 1.4462E-03 | 3.4117E-03 |
| ENSG00000226049\|TLK2P1 | 106.5256 | -0.6843 | ##### | -3.1851 | 1.4469E-03 | 3.4130E-03 |
| ENSG00000105879\|CBLL1 | 3796.8876 | 0.4064 | ##### | 3.1851 | 1.4470E-03 | 3.4130E-03 |

| | | | | | | |
|---|---|---|---|---|---|---|
| ENSG00000229393 | 21.1314 | 5.4135 | ##### | 3.1850 | 1.4475E-03 | 3.4139E-03 |
| ENSG00000107779\|BMPR1A | 3502.9259 | -0.3999 | ##### | -3.1850 | 1.4477E-03 | 3.4140E-03 |
| ENSG00000285230\|RALY-AS1 | 162.5532 | -0.5935 | ##### | -3.1845 | 1.4501E-03 | 3.4194E-03 |
| ENSG00000280157 | 12.5850 | 5.2348 | ##### | 3.1831 | 1.4572E-03 | 3.4359E-03 |
| ENSG00000120889\|TNFRSF10B | 10902.2851 | -0.3960 | ##### | -3.1829 | 1.4582E-03 | 3.4380E-03 |
| ENSG00000255197\|SLC39A13-AS1 | 60.5169 | 7.8729 | ##### | 3.1820 | 1.4627E-03 | 3.4484E-03 |
| ENSG00000229204\|PTGES3P3 | 28.8831 | 1.0498 | ##### | 3.1816 | 1.4647E-03 | 3.4527E-03 |
| ENSG00000121542\|SEC22A | 1199.8866 | -0.4099 | ##### | -3.1810 | 1.4675E-03 | 3.4591E-03 |
| ENSG00000214561\|RBBP4P4 | 6.9751 | 2.2327 | ##### | 3.1808 | 1.4685E-03 | 3.4611E-03 |
| ENSG00000176087\|SLC35A4 | 8593.8353 | 0.3840 | ##### | 3.1800 | 1.4726E-03 | 3.4705E-03 |
| ENSG00000172969\|FRG2C | 7.9659 | 5.5931 | ##### | 3.1799 | 1.4734E-03 | 3.4723E-03 |
| ENSG00000084623\|EIF3I | 28508.0741 | 0.3010 | ##### | 3.1794 | 1.4757E-03 | 3.4774E-03 |
| ENSG00000224429\|LINC00539 | 21.1017 | 2.0940 | ##### | 3.1794 | 1.4760E-03 | 3.4778E-03 |
| ENSG00000291263\|SMG1P7 | 65.3560 | -0.7972 | ##### | -3.1793 | 1.4764E-03 | 3.4780E-03 |
| ENSG00000225173 | 30.6265 | 1.8629 | ##### | 3.1793 | 1.4762E-03 | 3.4780E-03 |
| ENSG00000273739 | 7.5729 | 4.1855 | ##### | 3.1793 | 1.4764E-03 | 3.4780E-03 |
| ENSG00000186615\|KTN1-AS1 | 754.5639 | 0.4399 | ##### | 3.1792 | 1.4770E-03 | 3.4791E-03 |
| ENSG00000205913\|SRRM2-AS1 | 151.5959 | -0.5952 | ##### | -3.1791 | 1.4773E-03 | 3.4796E-03 |
| ENSG00000230698 | 41.9693 | -8.3426 | ##### | -3.1791 | 1.4775E-03 | 3.4796E-03 |
| ENSG00000112245\|PTP4A1 | 16725.5053 | -0.5348 | ##### | -3.1787 | 1.4794E-03 | 3.4838E-03 |
| ENSG00000290146 | 115.9615 | 0.8184 | ##### | 3.1784 | 1.4807E-03 | 3.4866E-03 |
| ENSG00000131828\|PDHA1 | 9003.9216 | -0.3879 | ##### | -3.1782 | 1.4817E-03 | 3.4887E-03 |
| ENSG00000166340\|TPP1 | 5163.5953 | -0.3701 | ##### | -3.1776 | 1.4852E-03 | 3.4968E-03 |
| ENSG00000006071\|ABCC8 | 137.2364 | 3.4200 | ##### | 3.1770 | 1.4881E-03 | 3.5031E-03 |
| ENSG00000245498\|MSANTD2-AS1 | 130.5211 | 0.6963 | ##### | 3.1766 | 1.4899E-03 | 3.5073E-03 |
| ENSG00000272414\|FAM47E-STBD1 | 307.5103 | -1.0880 | ##### | -3.1764 | 1.4912E-03 | 3.5099E-03 |
| ENSG00000286186 | 11.7962 | -5.7931 | ##### | -3.1760 | 1.4932E-03 | 3.5143E-03 |
| ENSG00000248377\|HMGN1P9 | 11.4988 | 4.8392 | ##### | 3.1756 | 1.4954E-03 | 3.5193E-03 |
| ENSG00000100485\|SOS2 | 2454.3303 | 0.4115 | ##### | 3.1752 | 1.4973E-03 | 3.5234E-03 |
| ENSG00000117408\|IPO13 | 3447.6716 | -0.3898 | ##### | -3.1745 | 1.5007E-03 | 3.5312E-03 |
| ENSG00000188848\|BEND4 | 47.1744 | 2.8085 | ##### | 3.1745 | 1.5008E-03 | 3.5312E-03 |
| ENSG00000163125\|RPRD2 | 5715.8493 | 0.3803 | ##### | 3.1742 | 1.5023E-03 | 3.5343E-03 |
| ENSG00000120868\|APAF1 | 2498.0902 | 0.4047 | ##### | 3.1733 | 1.5070E-03 | 3.5451E-03 |
| ENSG00000158079\|PTPDC1 | 973.5667 | -0.4402 | ##### | -3.1730 | 1.5089E-03 | 3.5494E-03 |
| ENSG00000229539\|PANK2-AS1 | 76.6262 | 0.8396 | ##### | 3.1726 | 1.5108E-03 | 3.5536E-03 |
| ENSG00000239332\|LINC01119 | 31.9666 | 1.6577 | ##### | 3.1725 | 1.5114E-03 | 3.5547E-03 |
| ENSG00000225137\|DYNC1I2P1 | 436.1938 | 0.4985 | ##### | 3.1720 | 1.5137E-03 | 3.5599E-03 |
| ENSG00000146677\|RPL32P18 | 69.3463 | 0.9393 | ##### | 3.1719 | 1.5146E-03 | 3.5617E-03 |
| ENSG00000264235\|MYL12-AS1 | 91.3580 | 1.2728 | ##### | 3.1716 | 1.5160E-03 | 3.5648E-03 |
| ENSG00000128908\|INO80 | 4648.5447 | 0.3977 | ##### | 3.1713 | 1.5177E-03 | 3.5685E-03 |
| ENSG00000282849 | 269.2318 | -0.5143 | ##### | -3.1706 | 1.5214E-03 | 3.5768E-03 |
| ENSG00000196860\|TOMM20L | 12.6016 | 1.5560 | ##### | 3.1705 | 1.5215E-03 | 3.5768E-03 |
| ENSG00000125871\|MGME1 | 2040.4188 | 0.4046 | ##### | 3.1703 | 1.5226E-03 | 3.5791E-03 |
| ENSG00000140254\|DUOXA1 | 20.0636 | -4.1627 | ##### | -3.1701 | 1.5240E-03 | 3.5822E-03 |
| ENSG00000164284\|GRPEL2 | 3824.3525 | -0.3907 | ##### | -3.1692 | 1.5283E-03 | 3.5920E-03 |
| ENSG00000185627\|PSMD13 | 10176.1060 | 0.3836 | ##### | 3.1688 | 1.5308E-03 | 3.5975E-03 |
| ENSG00000260401 | 13.9974 | -2.8098 | ##### | -3.1686 | 1.5319E-03 | 3.5998E-03 |
| ENSG00000289365 | 27.4721 | 2.5489 | ##### | 3.1683 | 1.5333E-03 | 3.6029E-03 |
| ENSG00000173535\|TNFRSF10C | 31.8311 | -3.6669 | ##### | -3.1679 | 1.5354E-03 | 3.6075E-03 |
| ENSG00000138443\|ABI2 | 11013.1567 | 0.3790 | ##### | 3.1677 | 1.5365E-03 | 3.6098E-03 |
| ENSG00000176055\|MBLAC2 | 1096.7635 | -0.3045 | ##### | -3.1674 | 1.5378E-03 | 3.6124E-03 |
| ENSG00000186130\|ZBTB6 | 1460.9717 | 0.3987 | ##### | 3.1675 | 1.5377E-03 | 3.6124E-03 |
| ENSG00000149489\|ROM1 | 125.0815 | -0.6122 | ##### | -3.1660 | 1.5453E-03 | 3.6296E-03 |
| ENSG00000271824\|SMIM32 | 12.0114 | 5.7947 | ##### | 3.1650 | 1.5510E-03 | 3.6428E-03 |
| ENSG00000293546\|ALG1L9P | 44.9139 | 1.8583 | ##### | 3.1649 | 1.5516E-03 | 3.6439E-03 |
| ENSG00000237298\|TTN-AS1 | 591.1376 | 0.4513 | ##### | 3.1644 | 1.5540E-03 | 3.6492E-03 |
| ENSG00000225506\|CYP4A22-AS1 | 16.6266 | -1.6787 | ##### | -3.1644 | 1.5541E-03 | 3.6492E-03 |
| ENSG00000109084\|TMEM97 | 5357.7823 | -0.3244 | ##### | -3.1643 | 1.5545E-03 | 3.6499E-03 |
| ENSG00000259370 | 37.3789 | 1.5335 | ##### | 3.1640 | 1.5562E-03 | 3.6535E-03 |
| ENSG00000282933\|RHOXF1P3 | 267.7018 | 0.5511 | ##### | 3.1635 | 1.5591E-03 | 3.6600E-03 |
| ENSG00000185634\|SHC4 | 20.8171 | 2.4220 | ##### | 3.1634 | 1.5596E-03 | 3.6610E-03 |
| ENSG00000081154\|PCNP | 13047.4237 | 0.2824 | ##### | 3.1630 | 1.5617E-03 | 3.6654E-03 |
| ENSG00000122824\|NUDT10 | 24.1592 | 2.5094 | ##### | 3.1622 | 1.5660E-03 | 3.6754E-03 |

| | | | | | | |
|---|---|---|---|---|---|---|
| ENSG00000149231\|CCDC82 | 2037.0106 | 0.4003 | ##### | 3.1618 | 1.5681E-03 | 3.6799E-03 |
| ENSG00000285649 | 110.3286 | 1.0500 | ##### | 3.1617 | 1.5684E-03 | 3.6805E-03 |
| ENSG00000245573\|BDNF-AS | 48.3680 | -1.1520 | ##### | -3.1616 | 1.5692E-03 | 3.6821E-03 |
| ENSG00000132141\|CCT6B | 53.9129 | -0.8679 | ##### | -3.1609 | 1.5731E-03 | 3.6908E-03 |
| ENSG00000272861 | 8.4406 | -3.5005 | ##### | -3.1606 | 1.5744E-03 | 3.6935E-03 |
| ENSG00000290574 | 98.4639 | -0.7442 | ##### | -3.1606 | 1.5745E-03 | 3.6935E-03 |
| ENSG00000172239\|PAIP1 | 9329.4968 | 0.3532 | ##### | 3.1605 | 1.5749E-03 | 3.6943E-03 |
| ENSG00000154914\|USP43 | 695.3040 | 0.4777 | ##### | 3.1574 | 1.5919E-03 | 3.7332E-03 |
| ENSG00000116117\|PARD3B | 405.6654 | 0.5826 | ##### | 3.1574 | 1.5918E-03 | 3.7332E-03 |
| ENSG00000279501 | 13.1712 | 3.2414 | ##### | 3.1574 | 1.5918E-03 | 3.7332E-03 |
| ENSG00000176244\|ACBD7 | 173.2278 | 0.6206 | ##### | 3.1571 | 1.5934E-03 | 3.7364E-03 |
| ENSG00000272360 | 9.8552 | -3.5122 | ##### | -3.1567 | 1.5956E-03 | 3.7413E-03 |
| ENSG00000174695\|TMEM167A | 7747.5936 | -0.3105 | ##### | -3.1562 | 1.5983E-03 | 3.7473E-03 |
| ENSG00000212464\|SNORA12 | 6.5603 | 4.5301 | ##### | 3.1557 | 1.6013E-03 | 3.7539E-03 |
| ENSG00000241860 | 42.0846 | 0.8125 | ##### | 3.1555 | 1.6020E-03 | 3.7554E-03 |
| ENSG00000108958 | 67.4688 | 0.7345 | ##### | 3.1550 | 1.6049E-03 | 3.7620E-03 |
| ENSG00000242797\|GLYCTK-AS1 | 32.9253 | 1.3335 | ##### | 3.1549 | 1.6053E-03 | 3.7625E-03 |
| ENSG00000289695 | 6.3601 | 5.4305 | ##### | 3.1541 | 1.6098E-03 | 3.7729E-03 |
| ENSG00000100099\|HPS4 | 2045.1118 | -0.3836 | ##### | -3.1539 | 1.6112E-03 | 3.7754E-03 |
| ENSG00000230590\|FTX | 3080.4346 | 0.4625 | ##### | 3.1539 | 1.6111E-03 | 3.7754E-03 |
| ENSG00000129646\|QRICH2 | 270.3641 | -0.5098 | ##### | -3.1528 | 1.6173E-03 | 3.7896E-03 |
| ENSG00000070770\|CSNK2A2 | 9287.8549 | -0.4210 | ##### | -3.1527 | 1.6176E-03 | 3.7899E-03 |
| ENSG00000289151 | 28.1060 | -1.2552 | ##### | -3.1526 | 1.6185E-03 | 3.7917E-03 |
| ENSG00000203286\|Metazoa_SRP | 6.0365 | 5.5370 | ##### | 3.1524 | 1.6194E-03 | 3.7934E-03 |
| ENSG00000181856\|SLC2A4 | 127.7801 | 1.0517 | ##### | 3.1518 | 1.6227E-03 | 3.8009E-03 |
| ENSG00000287961 | 8.0259 | 5.4927 | ##### | 3.1497 | 1.6344E-03 | 3.8281E-03 |
| ENSG00000172059\|KLF11 | 2069.1478 | 0.3956 | ##### | 3.1496 | 1.6351E-03 | 3.8295E-03 |
| ENSG00000151838\|CCDC175 | 13.0275 | 1.5299 | ##### | 3.1495 | 1.6355E-03 | 3.8301E-03 |
| ENSG00000251402\|FAM90A25P | 26.0360 | -1.2169 | ##### | -3.1487 | 1.6398E-03 | 3.8399E-03 |
| ENSG00000234782\|TPT1P9 | 39.8568 | 0.9025 | ##### | 3.1484 | 1.6418E-03 | 3.8441E-03 |
| ENSG00000204304\|PBX2 | 4832.3054 | 0.3910 | ##### | 3.1473 | 1.6480E-03 | 3.8585E-03 |
| ENSG00000120008\|WDR11 | 5284.9066 | -0.3731 | ##### | -3.1468 | 1.6506E-03 | 3.8642E-03 |
| ENSG00000256139 | 15.8123 | 3.5843 | ##### | 3.1468 | 1.6508E-03 | 3.8644E-03 |
| ENSG00000259677 | 9.0247 | 3.0151 | ##### | 3.1464 | 1.6527E-03 | 3.8685E-03 |
| ENSG00000230701\|FBXW4P1 | 19.9130 | 1.9323 | ##### | 3.1463 | 1.6533E-03 | 3.8693E-03 |
| ENSG00000172795\|DCP2 | 2303.6751 | 0.4040 | ##### | 3.1463 | 1.6532E-03 | 3.8693E-03 |
| ENSG00000247033 | 46.2086 | -1.0431 | ##### | -3.1463 | 1.6536E-03 | 3.8698E-03 |
| ENSG00000272054 | 73.2354 | 1.1135 | ##### | 3.1459 | 1.6559E-03 | 3.8747E-03 |
| ENSG00000251364\|SYT9-AS1 | 30.0725 | 1.4227 | ##### | 3.1457 | 1.6572E-03 | 3.8774E-03 |
| ENSG00000113583\|C5orf15 | 6136.0138 | -0.3840 | ##### | -3.1444 | 1.6644E-03 | 3.8940E-03 |
| ENSG00000281530 | 6.5408 | 3.4081 | ##### | 3.1441 | 1.6658E-03 | 3.8971E-03 |
| ENSG00000270248 | 4.3237 | -3.7161 | ##### | -3.1439 | 1.6673E-03 | 3.9003E-03 |
| ENSG00000146263\|MMS22L | 2462.6806 | 0.3860 | ##### | 3.1438 | 1.6678E-03 | 3.9010E-03 |
| ENSG00000180228\|PRKRA | 5627.6265 | -0.4127 | ##### | -3.1431 | 1.6715E-03 | 3.9095E-03 |
| ENSG00000081692\|JMJD4 | 2957.6453 | -0.4110 | ##### | -3.1430 | 1.6725E-03 | 3.9115E-03 |
| ENSG00000256399 | 11.1907 | -5.8870 | ##### | -3.1428 | 1.6734E-03 | 3.9133E-03 |
| ENSG00000261242 | 17.8098 | 1.3506 | ##### | 3.1426 | 1.6748E-03 | 3.9163E-03 |
| ENSG00000130844\|ZNF331 | 1725.9972 | -0.4065 | ##### | -3.1425 | 1.6751E-03 | 3.9165E-03 |
| ENSG00000156650\|KAT6B | 2068.3542 | -0.3959 | ##### | -3.1422 | 1.6768E-03 | 3.9203E-03 |
| ENSG00000269807 | 29.0256 | 4.6667 | ##### | 3.1419 | 1.6783E-03 | 3.9235E-03 |
| ENSG00000091039\|OSBPL8 | 5472.8050 | -0.3829 | ##### | -3.1407 | 1.6853E-03 | 3.9394E-03 |
| ENSG00000187049\|TMEM216 | 717.8336 | -0.4518 | ##### | -3.1394 | 1.6927E-03 | 3.9565E-03 |
| ENSG00000289757 | 27.0613 | -1.1614 | ##### | -3.1387 | 1.6973E-03 | 3.9669E-03 |
| ENSG00000273312 | 21.4152 | 7.4172 | ##### | 3.1384 | 1.6989E-03 | 3.9703E-03 |
| ENSG00000139351\|SYCP3 | 20.4582 | 1.6524 | ##### | 3.1382 | 1.7000E-03 | 3.9726E-03 |
| ENSG00000224079 | 9.4046 | 2.3017 | ##### | 3.1376 | 1.7035E-03 | 3.9805E-03 |
| ENSG00000092529\|CAPN3 | 34.3428 | -1.6941 | ##### | -3.1373 | 1.7050E-03 | 3.9837E-03 |
| ENSG00000280213\|UCKL1-AS1 | 111.2875 | 0.7903 | ##### | 3.1369 | 1.7073E-03 | 3.9888E-03 |
| ENSG00000186594\|MIR22HG | 1466.8784 | 0.4131 | ##### | 3.1359 | 1.7131E-03 | 4.0021E-03 |
| ENSG00000119711\|ALDH6A1 | 503.9785 | -0.5083 | ##### | -3.1350 | 1.7183E-03 | 4.0138E-03 |
| ENSG00000069509\|FUNDC1 | 947.3396 | -0.4486 | ##### | -3.1349 | 1.7190E-03 | 4.0151E-03 |
| ENSG00000173163\|COMMD1 | 1979.5609 | 0.4092 | ##### | 3.1346 | 1.7211E-03 | 4.0197E-03 |
| ENSG00000280277 | 9.1652 | -5.4586 | ##### | -3.1345 | 1.7212E-03 | 4.0198E-03 |

| | | | | | | |
|---|---|---|---|---|---|---|
| ENSG00000274569 | 19.5379 | 2.2150 | ##### | 3.1343 | 1.7224E-03 | 4.0221E-03 |
| ENSG00000254990 | 24.7721 | -7.1652 | ##### | -3.1341 | 1.7237E-03 | 4.0248E-03 |
| ENSG00000232936\|PANK1-AS1 | 22.2770 | -1.8776 | ##### | -3.1338 | 1.7258E-03 | 4.0294E-03 |
| ENSG00000273007 | 20.4789 | 1.3909 | ##### | 3.1337 | 1.7259E-03 | 4.0294E-03 |
| ENSG00000125901\|MRPS26 | 3343.3028 | 0.3856 | ##### | 3.1333 | 1.7287E-03 | 4.0357E-03 |
| ENSG00000063438\|AHRR | 419.6339 | -2.4779 | ##### | -3.1326 | 1.7327E-03 | 4.0445E-03 |
| ENSG00000125931\|CITED1 | 35.1962 | 3.6229 | ##### | 3.1324 | 1.7337E-03 | 4.0466E-03 |
| ENSG00000115687\|PASK | 1715.3758 | 0.3914 | ##### | 3.1319 | 1.7369E-03 | 4.0539E-03 |
| ENSG00000249116\|IRX4-AS1 | 8.9337 | 5.9905 | ##### | 3.1316 | 1.7388E-03 | 4.0579E-03 |
| ENSG00000125352\|RNF113A | 989.8406 | 0.4513 | ##### | 3.1307 | 1.7441E-03 | 4.0700E-03 |
| ENSG00000169715\|MT1E | 3911.7661 | 0.4581 | ##### | 3.1306 | 1.7443E-03 | 4.0702E-03 |
| ENSG00000137642\|SORL1 | 1012.9842 | 0.4260 | ##### | 3.1297 | 1.7496E-03 | 4.0822E-03 |
| ENSG00000182774\|RPS17 | 30648.6808 | 0.3758 | ##### | 3.1295 | 1.7512E-03 | 4.0857E-03 |
| ENSG00000174446\|SNAPC5 | 1412.7671 | -0.4207 | ##### | -3.1293 | 1.7524E-03 | 4.0880E-03 |
| ENSG00000104856\|RELB | 4433.6100 | -0.3106 | ##### | -3.1289 | 1.7545E-03 | 4.0927E-03 |
| ENSG00000157184\|CPT2 | 2236.9733 | -0.4042 | ##### | -3.1284 | 1.7577E-03 | 4.0998E-03 |
| ENSG00000054148\|PHPT1 | 8605.3336 | -0.3798 | ##### | -3.1280 | 1.7600E-03 | 4.1048E-03 |
| ENSG00000079387\|SENP1 | 4824.2104 | 0.3830 | ##### | 3.1277 | 1.7621E-03 | 4.1094E-03 |
| ENSG00000151962\|RBM46 | 82.4454 | ####### | ##### | -3.1274 | 1.7638E-03 | 4.1132E-03 |
| ENSG00000287576 | 126.1548 | -0.6757 | ##### | -3.1270 | 1.7659E-03 | 4.1176E-03 |
| ENSG00000180481\|GLIPR1L2 | 40.4656 | -1.0087 | ##### | -3.1267 | 1.7677E-03 | 4.1215E-03 |
| ENSG00000100201\|DDX17 | 19776.9181 | 0.3666 | ##### | 3.1266 | 1.7683E-03 | 4.1225E-03 |
| ENSG00000288670 | 587.3711 | 0.4271 | ##### | 3.1261 | 1.7713E-03 | 4.1292E-03 |
| ENSG00000279637 | 10.0914 | 6.4979 | ##### | 3.1257 | 1.7740E-03 | 4.1353E-03 |
| ENSG00000206897\|SNORA9B | 8.4791 | 2.0173 | ##### | 3.1256 | 1.7747E-03 | 4.1364E-03 |
| ENSG00000171703\|TCEA2 | 2545.1511 | -0.3982 | ##### | -3.1251 | 1.7774E-03 | 4.1425E-03 |
| ENSG00000142733\|MAP3K6 | 2278.0661 | 0.3846 | ##### | 3.1251 | 1.7777E-03 | 4.1429E-03 |
| ENSG00000287193 | 30.0711 | -6.9165 | ##### | -3.1248 | 1.7794E-03 | 4.1465E-03 |
| ENSG00000287190 | 13.6014 | 3.1800 | ##### | 3.1243 | 1.7824E-03 | 4.1531E-03 |
| ENSG00000279726 | 22.8654 | -3.4017 | ##### | -3.1238 | 1.7854E-03 | 4.1599E-03 |
| ENSG00000163888\|CAMK2N2 | 124.9195 | 0.6016 | ##### | 3.1237 | 1.7858E-03 | 4.1605E-03 |
| ENSG00000231616 | 12.3967 | 5.4503 | ##### | 3.1236 | 1.7868E-03 | 4.1624E-03 |
| ENSG00000198520\|ARMH1 | 32.0798 | -1.9229 | ##### | -3.1235 | 1.7872E-03 | 4.1632E-03 |
| ENSG00000243696 | 95.1999 | -1.9943 | ##### | -3.1227 | 1.7919E-03 | 4.1734E-03 |
| ENSG00000146700\|SSC4D | 153.7603 | -0.6146 | ##### | -3.1227 | 1.7919E-03 | 4.1734E-03 |
| ENSG00000185745\|IFIT1 | 175.0550 | -0.5766 | ##### | -3.1226 | 1.7926E-03 | 4.1747E-03 |
| ENSG00000258944 | 37.7405 | 0.9437 | ##### | 3.1222 | 1.7947E-03 | 4.1793E-03 |
| ENSG00000087095\|NLK | 1638.0388 | 0.4149 | ##### | 3.1209 | 1.8029E-03 | 4.1980E-03 |
| ENSG00000246323\|FAM13B-AS1 | 10.8735 | 2.7357 | ##### | 3.1191 | 1.8140E-03 | 4.2234E-03 |
| ENSG00000165983\|PTER | 3176.7054 | 0.2676 | ##### | 3.1189 | 1.8151E-03 | 4.2257E-03 |
| ENSG00000134245\|WNT2B | 379.7497 | 0.6595 | ##### | 3.1177 | 1.8230E-03 | 4.2437E-03 |
| ENSG00000116459\|ATP5PB | 18356.7702 | 0.3733 | ##### | 3.1173 | 1.8252E-03 | 4.2486E-03 |
| ENSG00000268163 | 4.2821 | 4.8245 | ##### | 3.1169 | 1.8276E-03 | 4.2538E-03 |
| ENSG00000167513\|CDT1 | 4750.0537 | 0.3731 | ##### | 3.1161 | 1.8327E-03 | 4.2653E-03 |
| ENSG00000289987 | 82.3762 | -1.4044 | ##### | -3.1159 | 1.8338E-03 | 4.2676E-03 |
| ENSG00000290386 | 72.4935 | 0.6907 | ##### | 3.1155 | 1.8365E-03 | 4.2737E-03 |
| ENSG00000244144\|INAVAP1 | 11.4121 | -3.7385 | ##### | -3.1142 | 1.8446E-03 | 4.2921E-03 |
| ENSG00000275329 | 15.2708 | -2.4863 | ##### | -3.1141 | 1.8452E-03 | 4.2932E-03 |
| ENSG00000272009 | 13.0945 | 1.7516 | ##### | 3.1140 | 1.8458E-03 | 4.2942E-03 |
| ENSG00000169551\|CT55 | 25.0553 | -3.0949 | ##### | -3.1131 | 1.8511E-03 | 4.3063E-03 |
| ENSG00000116918\|TSNAX | 4547.2474 | -0.3822 | ##### | -3.1131 | 1.8515E-03 | 4.3067E-03 |
| ENSG00000170791\|CHCHD7 | 824.7062 | -0.4127 | ##### | -3.1130 | 1.8517E-03 | 4.3070E-03 |
| ENSG00000137944\|KYAT3 | 967.2949 | 0.4272 | ##### | 3.1127 | 1.8536E-03 | 4.3111E-03 |
| ENSG00000128191\|DGCR8 | 3581.4725 | -0.2698 | ##### | -3.1127 | 1.8538E-03 | 4.3111E-03 |
| ENSG00000258704\|SRP54-AS1 | 21.2282 | 7.1910 | ##### | 3.1125 | 1.8548E-03 | 4.3131E-03 |
| ENSG00000285730 | 7.6598 | 3.2576 | ##### | 3.1124 | 1.8555E-03 | 4.3144E-03 |
| ENSG00000145016\|RUBCN | 2834.1483 | -0.3807 | ##### | -3.1120 | 1.8584E-03 | 4.3208E-03 |
| ENSG00000226007\|FAM88F | 56.0565 | -0.8501 | ##### | -3.1114 | 1.8620E-03 | 4.3288E-03 |
| ENSG00000258359\|PCNPP1 | 26.2551 | 1.1966 | ##### | 3.1113 | 1.8628E-03 | 4.3304E-03 |
| ENSG00000277728 | 14.2875 | 2.3088 | ##### | 3.1110 | 1.8642E-03 | 4.3334E-03 |
| ENSG00000131015\|ULBP2 | 197.2103 | 0.6084 | ##### | 3.1110 | 1.8645E-03 | 4.3337E-03 |
| ENSG00000120837\|NFYB | 3799.1943 | 0.3827 | ##### | 3.1105 | 1.8676E-03 | 4.3404E-03 |
| ENSG00000166454\|ATMIN | 6322.0207 | -0.3912 | ##### | -3.1105 | 1.8680E-03 | 4.3410E-03 |

| | | | | | | |
|---|---|---|---|---|---|---|
| ENSG00000110048\|OSBP | 5235.4304 | -0.3709 | ##### | -3.1103 | 1.8691E-03 | 4.3433E-03 |
| ENSG00000103502\|CDIPT | 7910.5712 | -0.3582 | ##### | -3.1103 | 1.8692E-03 | 4.3433E-03 |
| ENSG00000131470\|PSMC3IP | 2169.7763 | 0.4136 | ##### | 3.1095 | 1.8743E-03 | 4.3547E-03 |
| ENSG00000243810\|CFAP61-AS1 | 9.8998 | -5.7068 | ##### | -3.1090 | 1.8769E-03 | 4.3605E-03 |
| ENSG00000089060\|SLC8B1 | 1441.6616 | -0.4266 | ##### | -3.1090 | 1.8775E-03 | 4.3615E-03 |
| ENSG00000136279\|DBNL | 8956.5481 | -0.3050 | ##### | -3.1083 | 1.8819E-03 | 4.3714E-03 |
| ENSG00000253356 | 554.0942 | 0.5218 | ##### | 3.1081 | 1.8828E-03 | 4.3732E-03 |
| ENSG00000240821\|RPL9P25 | 8.3763 | -2.0293 | ##### | -3.1076 | 1.8862E-03 | 4.3807E-03 |
| ENSG00000261915 | 28.4045 | 3.8337 | ##### | 3.1076 | 1.8864E-03 | 4.3807E-03 |
| ENSG00000217555\|CKLF | 3272.8511 | -0.3685 | ##### | -3.1068 | 1.8915E-03 | 4.3924E-03 |
| ENSG00000166912\|MTMR10 | 1683.9407 | -0.4177 | ##### | -3.1060 | 1.8964E-03 | 4.4031E-03 |
| ENSG00000101197\|BIRC7 | 51.7580 | 0.9731 | ##### | 3.1060 | 1.8964E-03 | 4.4031E-03 |
| ENSG00000217716\|RPS10P3 | 52.5294 | 0.9205 | ##### | 3.1060 | 1.8966E-03 | 4.4032E-03 |
| ENSG00000188306\|LRRIQ4 | 13.6599 | 5.5349 | ##### | 3.1053 | 1.9007E-03 | 4.4124E-03 |
| ENSG00000248778 | 5.9507 | -5.4655 | ##### | -3.1046 | 1.9056E-03 | 4.4234E-03 |
| ENSG00000254859 | 22.8570 | -1.4658 | ##### | -3.1043 | 1.9075E-03 | 4.4274E-03 |
| ENSG00000185345\|PRKN | 25.5286 | -2.8289 | ##### | -3.1038 | 1.9105E-03 | 4.4339E-03 |
| ENSG00000271360\|USP6NL-AS1 | 8.8551 | 1.9959 | ##### | 3.1036 | 1.9118E-03 | 4.4367E-03 |
| ENSG00000261349 | 10.0582 | 2.2143 | ##### | 3.1023 | 1.9200E-03 | 4.4554E-03 |
| ENSG00000159224\|GIP | 19.6420 | -3.8526 | ##### | -3.1022 | 1.9210E-03 | 4.4573E-03 |
| ENSG00000283071\|LBHD2 | 5.7556 | -5.4304 | ##### | -3.1010 | 1.9290E-03 | 4.4756E-03 |
| ENSG00000278356 | 17.6896 | 2.0239 | ##### | 3.1007 | 1.9306E-03 | 4.4789E-03 |
| ENSG00000264635 | 22.8714 | 2.4464 | ##### | 3.1006 | 1.9312E-03 | 4.4801E-03 |
| ENSG00000224885\|EIPR1-IT1 | 11.6281 | 6.1262 | ##### | 3.1005 | 1.9317E-03 | 4.4809E-03 |
| ENSG00000029725\|RABEP1 | 6356.6599 | -0.3683 | ##### | -3.1003 | 1.9334E-03 | 4.4840E-03 |
| ENSG00000063244\|U2AF2 | 23397.5388 | 0.2560 | ##### | 3.1003 | 1.9333E-03 | 4.4840E-03 |
| ENSG00000137822\|TUBGCP4 | 3910.1378 | 0.3858 | ##### | 3.0998 | 1.9363E-03 | 4.4904E-03 |
| ENSG00000071051\|NCK2 | 3049.4891 | -0.3763 | ##### | -3.0997 | 1.9374E-03 | 4.4925E-03 |
| ENSG00000123009\|NME2P1 | 46.0529 | 0.8570 | ##### | 3.0994 | 1.9388E-03 | 4.4956E-03 |
| ENSG00000185924\|RTN4RL1 | 105.9979 | 0.8035 | ##### | 3.0993 | 1.9400E-03 | 4.4981E-03 |
| ENSG00000214013\|GANC | 626.6793 | -0.4242 | ##### | -3.0989 | 1.9427E-03 | 4.5038E-03 |
| ENSG00000205108\|SPATA31F1 | 16.9669 | -7.2969 | ##### | -3.0977 | 1.9503E-03 | 4.5211E-03 |
| ENSG00000229081\|LINC01165 | 12.8453 | 3.0430 | ##### | 3.0976 | 1.9510E-03 | 4.5220E-03 |
| ENSG00000163510\|CWC22 | 4465.4747 | 0.3743 | ##### | 3.0976 | 1.9509E-03 | 4.5220E-03 |
| ENSG00000069275\|NUCKS1 | 24545.7542 | -0.3560 | ##### | -3.0975 | 1.9517E-03 | 4.5231E-03 |
| ENSG00000160284\|SPATC1L | 1173.1914 | 0.4173 | ##### | 3.0975 | 1.9517E-03 | 4.5231E-03 |
| ENSG00000197885\|NKIRAS1 | 1928.2106 | -0.6017 | ##### | -3.0974 | 1.9520E-03 | 4.5232E-03 |
| ENSG00000287431\|RENO1 | 139.0745 | -0.6507 | ##### | -3.0972 | 1.9538E-03 | 4.5272E-03 |
| ENSG00000163312\|HELQ | 737.5983 | 0.4231 | ##### | 3.0970 | 1.9547E-03 | 4.5290E-03 |
| ENSG00000184743\|ATL3 | 9052.6973 | -0.3118 | ##### | -3.0966 | 1.9576E-03 | 4.5353E-03 |
| ENSG00000089048\|ESF1 | 3394.9528 | 0.3742 | ##### | 3.0963 | 1.9594E-03 | 4.5390E-03 |
| ENSG00000178199\|ZC3H12D | 34.3400 | 1.3728 | ##### | 3.0963 | 1.9598E-03 | 4.5397E-03 |
| ENSG00000213117\|MEMO1P2 | 4.1634 | 3.7048 | ##### | 3.0962 | 1.9605E-03 | 4.5408E-03 |
| ENSG00000101346\|POFUT1 | 9272.4121 | -0.3710 | ##### | -3.0961 | 1.9607E-03 | 4.5411E-03 |
| ENSG00000177981\|ASB8 | 1583.7029 | -0.3263 | ##### | -3.0958 | 1.9627E-03 | 4.5450E-03 |
| ENSG00000218305\|CDC14C | 16.7041 | -2.0235 | ##### | -3.0958 | 1.9626E-03 | 4.5450E-03 |
| ENSG00000259151\|CAP2P1 | 22.8962 | 6.4116 | ##### | 3.0957 | 1.9635E-03 | 4.5464E-03 |
| ENSG00000166800\|LDHAL6A | 23.7981 | -5.0647 | ##### | -3.0953 | 1.9662E-03 | 4.5524E-03 |
| ENSG00000227695\|DNMBP-AS1 | 32.4092 | 1.2753 | ##### | 3.0932 | 1.9803E-03 | 4.5845E-03 |
| ENSG00000017260\|ATP2C1 | 14104.1931 | 0.2853 | ##### | 3.0929 | 1.9818E-03 | 4.5878E-03 |
| ENSG00000241782\|RPL21P95 | 9.9692 | 6.3631 | ##### | 3.0925 | 1.9846E-03 | 4.5938E-03 |
| ENSG00000163283\|ALPP | 90.6628 | -1.8487 | ##### | -3.0925 | 1.9849E-03 | 4.5941E-03 |
| ENSG00000125734\|GPR108 | 2701.1371 | -0.3004 | ##### | -3.0925 | 1.9851E-03 | 4.5942E-03 |
| ENSG00000228412\|LNC-LBCS | 131.1457 | 0.6162 | ##### | 3.0909 | 1.9954E-03 | 4.6178E-03 |
| ENSG00000086730\|LAT2 | 199.3027 | -0.5310 | ##### | -3.0909 | 1.9955E-03 | 4.6178E-03 |
| ENSG00000106609\|TMEM248 | 8068.4151 | -0.3659 | ##### | -3.0905 | 1.9980E-03 | 4.6231E-03 |
| ENSG00000260882 | 10.3849 | 5.8371 | ##### | 3.0902 | 2.0000E-03 | 4.6274E-03 |
| ENSG00000117013\|KCNQ4 | 74.0212 | -0.8427 | ##### | -3.0902 | 2.0004E-03 | 4.6280E-03 |
| ENSG00000287475 | 10.1956 | 5.8642 | ##### | 3.0896 | 2.0044E-03 | 4.6369E-03 |
| ENSG00000184281\|TSSC4 | 2971.5927 | 0.3869 | ##### | 3.0893 | 2.0060E-03 | 4.6397E-03 |
| ENSG00000171462\|DLK2 | 649.8427 | 0.4786 | ##### | 3.0894 | 2.0059E-03 | 4.6397E-03 |
| ENSG00000230005\|SNAP47-AS1 | 7.8502 | 6.0671 | ##### | 3.0885 | 2.0119E-03 | 4.6530E-03 |
| ENSG00000166262\|FAM227B | 256.0618 | -0.6940 | ##### | -3.0883 | 2.0133E-03 | 4.6559E-03 |

| | | | | | | |
|---|---|---|---|---|---|---|
| ENSG00000277586\|NEFL | 6023.1518 | 0.3091 | ##### | 3.0881 | 2.0143E-03 | 4.6580E-03 |
| ENSG00000103194\|USP10 | 11777.9934 | 0.3700 | ##### | 3.0871 | 2.0214E-03 | 4.6739E-03 |
| ENSG00000279204 | 7.8531 | -2.4463 | ##### | -3.0868 | 2.0234E-03 | 4.6782E-03 |
| ENSG00000111276\|CDKN1B | 4547.4132 | 0.3548 | ##### | 3.0864 | 2.0261E-03 | 4.6842E-03 |
| ENSG00000104497\|SNX16 | 482.5057 | -0.4735 | ##### | -3.0857 | 2.0310E-03 | 4.6951E-03 |
| ENSG00000196876\|SCN8A | 1033.3951 | 0.5007 | ##### | 3.0850 | 2.0352E-03 | 4.7044E-03 |
| ENSG00000293529 | 19.7144 | 4.6807 | ##### | 3.0842 | 2.0408E-03 | 4.7171E-03 |
| ENSG00000266967\|AARSD1 | 2566.7541 | 0.3979 | ##### | 3.0838 | 2.0438E-03 | 4.7237E-03 |
| ENSG00000170035\|UBE2E3 | 5626.7352 | -0.3783 | ##### | -3.0836 | 2.0451E-03 | 4.7262E-03 |
| ENSG00000215014 | 56.1272 | -1.3610 | ##### | -3.0835 | 2.0461E-03 | 4.7281E-03 |
| ENSG00000198740\|ZNF652 | 2631.8115 | -0.3763 | ##### | -3.0834 | 2.0462E-03 | 4.7281E-03 |
| ENSG00000290848 | 14.7721 | -1.4595 | ##### | -3.0834 | 2.0468E-03 | 4.7291E-03 |
| ENSG00000166822\|TMEM170A | 1909.1757 | 0.3898 | ##### | 3.0830 | 2.0496E-03 | 4.7350E-03 |
| ENSG00000075415\|SLC25A3 | 42107.7233 | 0.3677 | ##### | 3.0824 | 2.0536E-03 | 4.7441E-03 |
| ENSG00000271553 | 12.4154 | 2.6527 | ##### | 3.0818 | 2.0573E-03 | 4.7521E-03 |
| ENSG00000230366\|DSCR9 | 13.0745 | -1.6071 | ##### | -3.0818 | 2.0579E-03 | 4.7531E-03 |
| ENSG00000256060\|TRAPPC2B | 328.6059 | -0.5418 | ##### | -3.0815 | 2.0594E-03 | 4.7564E-03 |
| ENSG00000165669\|FAM204A | 2376.7895 | -0.3881 | ##### | -3.0811 | 2.0624E-03 | 4.7629E-03 |
| ENSG00000274944 | 18.9017 | -3.6450 | ##### | -3.0808 | 2.0642E-03 | 4.7666E-03 |
| ENSG00000163214\|DHX57 | 4026.0648 | 0.3683 | ##### | 3.0806 | 2.0656E-03 | 4.7694E-03 |
| ENSG00000100345\|MYH9 | 74595.1972 | 0.3619 | ##### | 3.0801 | 2.0693E-03 | 4.7776E-03 |
| ENSG00000111404\|RERGL | 15.1653 | 3.4928 | ##### | 3.0797 | 2.0723E-03 | 4.7842E-03 |
| ENSG00000290594 | 56.2183 | 9.7048 | ##### | 3.0795 | 2.0736E-03 | 4.7869E-03 |
| ENSG00000173209\|AHSA2P | 70.8901 | ####### | ##### | -3.0795 | 2.0738E-03 | 4.7870E-03 |
| ENSG00000239653\|PSMD6-AS2 | 37.8349 | 2.0003 | ##### | 3.0793 | 2.0747E-03 | 4.7887E-03 |
| ENSG00000139436\|GIT2 | 6330.1818 | 0.3584 | ##### | 3.0785 | 2.0802E-03 | 4.8009E-03 |
| ENSG00000137869\|CYP19A1 | 28.0301 | -7.5894 | ##### | -3.0785 | 2.0806E-03 | 4.8016E-03 |
| ENSG00000179743\|SPEN-AS1 | 322.8347 | -0.4599 | ##### | -3.0784 | 2.0812E-03 | 4.8026E-03 |
| ENSG00000260563 | 179.8928 | -0.7355 | ##### | -3.0784 | 2.0814E-03 | 4.8027E-03 |
| ENSG00000115137\|DNAJC27 | 335.3270 | 0.5544 | ##### | 3.0782 | 2.0825E-03 | 4.8049E-03 |
| ENSG00000272369 | 12.0456 | 2.5394 | ##### | 3.0781 | 2.0836E-03 | 4.8070E-03 |
| ENSG00000119185\|ITGB1BP1 | 5717.7593 | -0.3871 | ##### | -3.0778 | 2.0855E-03 | 4.8111E-03 |
| ENSG00000070729\|CNGB1 | 28.5595 | -2.5704 | ##### | -3.0777 | 2.0858E-03 | 4.8113E-03 |
| ENSG00000259024\|TVP23C-CDRT4 | 445.2122 | -0.9781 | ##### | -3.0774 | 2.0879E-03 | 4.8158E-03 |
| ENSG00000179598\|PLD6 | 638.7313 | 0.4523 | ##### | 3.0759 | 2.0990E-03 | 4.8406E-03 |
| ENSG00000270257 | 10.8532 | 4.6021 | ##### | 3.0759 | 2.0990E-03 | 4.8406E-03 |
| ENSG00000143514\|TP53BP2 | 5453.3977 | 0.3848 | ##### | 3.0757 | 2.0999E-03 | 4.8424E-03 |
| ENSG00000157020\|SEC13 | 8219.0697 | -0.3649 | ##### | -3.0752 | 2.1036E-03 | 4.8506E-03 |
| ENSG00000290677 | 23.3145 | -7.5497 | ##### | -3.0750 | 2.1054E-03 | 4.8542E-03 |
| ENSG00000205143\|ARID3C | 34.3225 | -1.0090 | ##### | -3.0745 | 2.1083E-03 | 4.8606E-03 |
| ENSG00000186300\|ZNF555 | 469.9240 | 0.4759 | ##### | 3.0745 | 2.1088E-03 | 4.8613E-03 |
| ENSG00000122257\|RBBP6 | 6187.7758 | 0.3654 | ##### | 3.0744 | 2.1092E-03 | 4.8619E-03 |
| ENSG00000240184\|PCDHGC3 | 889.6435 | -0.4393 | ##### | -3.0740 | 2.1124E-03 | 4.8689E-03 |
| ENSG00000205643\|CDPF1 | 607.1703 | -0.4445 | ##### | -3.0738 | 2.1136E-03 | 4.8714E-03 |
| ENSG00000184857\|TMEM186 | 953.2998 | -0.4019 | ##### | -3.0736 | 2.1149E-03 | 4.8740E-03 |
| ENSG00000178078\|STAP2 | 28.4384 | 2.5117 | ##### | 3.0736 | 2.1152E-03 | 4.8742E-03 |
| ENSG00000127952\|STYXL1 | 1692.8816 | -0.3945 | ##### | -3.0733 | 2.1170E-03 | 4.8780E-03 |
| ENSG00000120696\|KBTBD7 | 964.7497 | 0.4137 | ##### | 3.0733 | 2.1173E-03 | 4.8782E-03 |
| ENSG00000287670 | 57.7892 | -1.0289 | ##### | -3.0731 | 2.1185E-03 | 4.8808E-03 |
| ENSG00000277157\|H4C4 | 12.0939 | 2.4527 | ##### | 3.0730 | 2.1189E-03 | 4.8812E-03 |
| ENSG00000254165 | 13.0409 | 2.1856 | ##### | 3.0714 | 2.1303E-03 | 4.9071E-03 |
| ENSG00000287229 | 33.6481 | -7.3684 | ##### | -3.0711 | 2.1325E-03 | 4.9118E-03 |
| ENSG00000254244\|PAICSP4 | 11.6032 | 1.6378 | ##### | 3.0711 | 2.1327E-03 | 4.9120E-03 |
| ENSG00000034510\|TMSB10 | 99441.8797 | 0.3575 | ##### | 3.0705 | 2.1370E-03 | 4.9215E-03 |
| ENSG00000271021 | 9.8663 | 2.7255 | ##### | 3.0700 | 2.1402E-03 | 4.9285E-03 |
| ENSG00000261514\|LINC01976 | 13.2184 | 2.6326 | ##### | 3.0697 | 2.1430E-03 | 4.9345E-03 |
| ENSG00000244754\|N4BP2L2 | 2755.4924 | 0.3816 | ##### | 3.0694 | 2.1449E-03 | 4.9384E-03 |
| ENSG00000050344\|NFE2L3 | 4678.1485 | 0.4333 | ##### | 3.0692 | 2.1461E-03 | 4.9410E-03 |
| ENSG00000245261 | 19.2024 | 1.2968 | ##### | 3.0689 | 2.1484E-03 | 4.9458E-03 |
| ENSG00000135521\|LTV1 | 4385.1885 | 0.3740 | ##### | 3.0689 | 2.1486E-03 | 4.9460E-03 |
| ENSG00000092010\|PSME1 | 4256.1856 | 0.3738 | ##### | 3.0682 | 2.1532E-03 | 4.9557E-03 |
| ENSG00000243961\|PARAL1 | 21.8611 | -4.2615 | ##### | -3.0683 | 2.1532E-03 | 4.9557E-03 |
| ENSG00000086015\|MAST2 | 6267.5407 | -0.3627 | ##### | -3.0681 | 2.1540E-03 | 4.9572E-03 |

| | | | | | | |
|---|---|---|---|---|---|---|
| ENSG00000130590\|SAMD10 | 496.7037 | -0.4496 | ##### | -3.0676 | 2.1578E-03 | 4.9655E-03 |
| ENSG00000215018\|COL28A1 | 10.5805 | -3.5538 | ##### | -3.0665 | 2.1656E-03 | 4.9830E-03 |
| ENSG00000170745\|KCNS3 | 333.3306 | -0.4791 | ##### | -3.0663 | 2.1676E-03 | 4.9872E-03 |
| ENSG00000274859 | 24.6391 | -1.1826 | ##### | -3.0662 | 2.1678E-03 | 4.9874E-03 |
| ENSG00000197566\|ZNF624 | 357.8660 | 0.5438 | ##### | 3.0658 | 2.1711E-03 | 4.9946E-03 |
| ENSG00000259933\|PKD1-AS1 | 28.3890 | 1.1581 | ##### | 3.0654 | 2.1736E-03 | 5.0000E-03 |
| ENSG00000173409\|ARV1 | 1195.0824 | -0.4009 | ##### | -3.0653 | 2.1744E-03 | 5.0013E-03 |
| ENSG00000273272 | 16.6882 | -3.1786 | ##### | -3.0653 | 2.1745E-03 | 5.0013E-03 |
| ENSG00000139971\|ARMH4 | 73.8468 | 0.9132 | ##### | 3.0640 | 2.1840E-03 | 5.0228E-03 |
| ENSG00000146457\|WTAP | 6131.9859 | 0.2809 | ##### | 3.0635 | 2.1875E-03 | 5.0304E-03 |
| ENSG00000234608\|MAPKAPK5-AS1 | 840.9843 | -0.4250 | ##### | -3.0631 | 2.1907E-03 | 5.0373E-03 |
| ENSG00000279539 | 66.9832 | -0.9923 | ##### | -3.0628 | 2.1928E-03 | 5.0418E-03 |
| ENSG00000108256\|NUFIP2 | 13713.0516 | 0.3673 | ##### | 3.0625 | 2.1947E-03 | 5.0459E-03 |
| ENSG00000140259\|MFAP1 | 3564.8412 | 0.3544 | ##### | 3.0624 | 2.1956E-03 | 5.0474E-03 |
| ENSG00000159346\|ADIPOR1 | 6490.1627 | -0.3845 | ##### | -3.0622 | 2.1970E-03 | 5.0503E-03 |
| ENSG00000231811 | 28.4079 | 7.0768 | ##### | 3.0616 | 2.2015E-03 | 5.0602E-03 |
| ENSG00000197594\|ENPP1 | 768.2087 | -0.4285 | ##### | -3.0614 | 2.2031E-03 | 5.0635E-03 |
| ENSG00000134884\|ARGLU1 | 4423.1609 | 0.3848 | ##### | 3.0612 | 2.2043E-03 | 5.0660E-03 |
| ENSG00000169019\|COMMD8 | 2365.0392 | 0.3976 | ##### | 3.0608 | 2.2075E-03 | 5.0728E-03 |
| ENSG00000049449\|RCN1 | 11520.1730 | -0.3699 | ##### | -3.0608 | 2.2076E-03 | 5.0728E-03 |
| ENSG00000130313\|PGLS | 3409.3757 | -0.3827 | ##### | -3.0606 | 2.2086E-03 | 5.0747E-03 |
| ENSG00000142676\|RPL11 | 33257.2811 | 0.3599 | ##### | 3.0602 | 2.2122E-03 | 5.0821E-03 |
| ENSG00000288548 | 11.3448 | 5.4460 | ##### | 3.0602 | 2.2122E-03 | 5.0821E-03 |
| ENSG00000290683 | 174.3839 | -0.5578 | ##### | -3.0593 | 2.2182E-03 | 5.0955E-03 |
| ENSG00000283463\|HSFX4 | 38.3905 | 1.3438 | ##### | 3.0590 | 2.2205E-03 | 5.1003E-03 |
| ENSG00000212916\|MAP10 | 399.7067 | -0.5767 | ##### | -3.0574 | 2.2328E-03 | 5.1282E-03 |
| ENSG00000275835\|TUBGCP5 | 1685.9517 | -0.4088 | ##### | -3.0571 | 2.2352E-03 | 5.1334E-03 |
| ENSG00000232613\|LINC02576 | 10.0407 | 6.4205 | ##### | 3.0570 | 2.2357E-03 | 5.1342E-03 |
| ENSG00000259692\|LINC01418 | 17.0180 | -1.4779 | ##### | -3.0567 | 2.2380E-03 | 5.1385E-03 |
| ENSG00000049541\|RFC2 | 4199.6833 | 0.3825 | ##### | 3.0567 | 2.2380E-03 | 5.1385E-03 |
| ENSG00000258682 | 17.8306 | 1.3757 | ##### | 3.0566 | 2.2389E-03 | 5.1404E-03 |
| ENSG00000112640\|PPP2R5D | 5234.4298 | 0.3584 | ##### | 3.0563 | 2.2411E-03 | 5.1450E-03 |
| ENSG00000163154\|TNFAIP8L2 | 21.6463 | -7.2969 | ##### | -3.0549 | 2.2514E-03 | 5.1683E-03 |
| ENSG00000287750\|LINC03026 | 54.7640 | -1.1373 | ##### | -3.0546 | 2.2536E-03 | 5.1728E-03 |
| ENSG00000257557\|PPP1R12A-AS1 | 159.3244 | -0.6235 | ##### | -3.0543 | 2.2556E-03 | 5.1771E-03 |
| ENSG00000278223\|MIR6783 | 9.4930 | 1.6631 | ##### | 3.0543 | 2.2560E-03 | 5.1777E-03 |
| ENSG00000279691 | 8.1138 | 2.6527 | ##### | 3.0542 | 2.2565E-03 | 5.1782E-03 |
| ENSG00000115539\|PDCL3 | 3190.8884 | -0.3702 | ##### | -3.0541 | 2.2574E-03 | 5.1801E-03 |
| ENSG00000113196\|HAND1 | 32.8328 | 1.3859 | ##### | 3.0536 | 2.2609E-03 | 5.1877E-03 |
| ENSG00000164951\|PDP1 | 1221.2546 | -0.4182 | ##### | -3.0527 | 2.2677E-03 | 5.2028E-03 |
| ENSG00000221953\|LINC02897 | 10.4833 | -4.3537 | ##### | -3.0526 | 2.2688E-03 | 5.2049E-03 |
| ENSG00000216331\|H1-12P | 42.1501 | 0.9197 | ##### | 3.0518 | 2.2749E-03 | 5.2184E-03 |
| ENSG00000101439\|CST3 | 5902.0577 | -0.3811 | ##### | -3.0516 | 2.2760E-03 | 5.2199E-03 |
| ENSG00000241472\|PTPRG-AS1 | 236.9648 | 0.5187 | ##### | 3.0517 | 2.2757E-03 | 5.2199E-03 |
| ENSG00000237863\|IRS4-AS1 | 14.1165 | 4.9925 | ##### | 3.0516 | 2.2760E-03 | 5.2199E-03 |
| ENSG00000278238\|CKAP2-DT | 42.8605 | -0.8299 | ##### | -3.0515 | 2.2767E-03 | 5.2210E-03 |
| ENSG00000038210\|PI4K2B | 2171.2581 | 0.3857 | ##### | 3.0513 | 2.2784E-03 | 5.2245E-03 |
| ENSG00000232648\|ING2-DT | 43.9975 | -0.9480 | ##### | -3.0509 | 2.2813E-03 | 5.2307E-03 |
| ENSG00000268583 | 32.9998 | -1.0257 | ##### | -3.0508 | 2.2821E-03 | 5.2323E-03 |
| ENSG00000113163\|CERT1 | 2621.3722 | -0.3957 | ##### | -3.0506 | 2.2839E-03 | 5.2359E-03 |
| ENSG00000137161\|CNPY3 | 3190.4299 | 0.3777 | ##### | 3.0506 | 2.2841E-03 | 5.2360E-03 |
| ENSG00000185038\|MROH2A | 26.3489 | 7.4537 | ##### | 3.0502 | 2.2870E-03 | 5.2422E-03 |
| ENSG00000205129\|CFAP96 | 38.4770 | 2.6649 | ##### | 3.0499 | 2.2889E-03 | 5.2463E-03 |
| ENSG00000272791 | 18.0538 | -2.3641 | ##### | -3.0499 | 2.2894E-03 | 5.2469E-03 |
| ENSG00000184270\|H2AC21 | 18.1658 | 1.8398 | ##### | 3.0497 | 2.2904E-03 | 5.2489E-03 |
| ENSG00000215875\|ST13P20 | 15.2597 | 5.0418 | ##### | 3.0487 | 2.2985E-03 | 5.2671E-03 |
| ENSG00000115355\|CCDC88A | 10158.8845 | -0.3718 | ##### | -3.0483 | 2.3011E-03 | 5.2725E-03 |
| ENSG00000290966\|KRT87P | 873.3768 | -8.9267 | ##### | -3.0478 | 2.3051E-03 | 5.2812E-03 |
| ENSG00000110700\|RPS13 | 17976.7186 | 0.3828 | ##### | 3.0478 | 2.3053E-03 | 5.2814E-03 |
| ENSG00000267666 | 45.4485 | -1.9799 | ##### | -3.0475 | 2.3074E-03 | 5.2858E-03 |
| ENSG00000133316\|WDR74 | 4560.5816 | 0.3494 | ##### | 3.0469 | 2.3125E-03 | 5.2970E-03 |
| ENSG00000234693\|RIPOR3-AS1 | 11.2806 | 6.0460 | ##### | 3.0465 | 2.3155E-03 | 5.3035E-03 |
| ENSG00000130751\|NPAS1 | 435.8238 | -0.4830 | ##### | -3.0463 | 2.3165E-03 | 5.3054E-03 |

| | | | | | | |
|---|---|---|---|---|---|---|
| ENSG00000240204\|SMKR1 | 496.5240 | 0.4484 | ##### | 3.0461 | 2.3185E-03 | 5.3096E-03 |
| ENSG00000254331\|CKS1BP7 | 7.0077 | 2.5685 | ##### | 3.0448 | 2.3280E-03 | 5.3309E-03 |
| ENSG00000249456 | 51.8821 | -0.7929 | ##### | -3.0445 | 2.3303E-03 | 5.3359E-03 |
| ENSG00000285906 | 39.1557 | -3.3084 | ##### | -3.0435 | 2.3385E-03 | 5.3542E-03 |
| ENSG00000136040\|PLXNC1 | 98.9246 | 0.6350 | ##### | 3.0431 | 2.3412E-03 | 5.3599E-03 |
| ENSG00000236581\|STARD13-AS | 23.9755 | 1.2773 | ##### | 3.0431 | 2.3418E-03 | 5.3609E-03 |
| ENSG00000127445\|PIN1 | 3705.5648 | 0.3792 | ##### | 3.0429 | 2.3430E-03 | 5.3631E-03 |
| ENSG00000278224\|PRICKLE4 | 31.0552 | -1.1227 | ##### | -3.0429 | 2.3434E-03 | 5.3638E-03 |
| ENSG00000291217\|LINC01347 | 32.1171 | 0.9089 | ##### | 3.0427 | 2.3447E-03 | 5.3663E-03 |
| ENSG00000261575 | 8.2684 | -6.0008 | ##### | -3.0426 | 2.3455E-03 | 5.3677E-03 |
| ENSG00000243554\|CCZ1P1 | 184.0116 | 1.0348 | ##### | 3.0418 | 2.3518E-03 | 5.3816E-03 |
| ENSG00000175826\|CTDNEP1 | 10572.7436 | 0.3681 | ##### | 3.0413 | 2.3552E-03 | 5.3892E-03 |
| ENSG00000290040 | 25.2527 | 7.5079 | ##### | 3.0411 | 2.3574E-03 | 5.3936E-03 |
| ENSG00000135336\|ORC3 | 2418.0754 | 0.3796 | ##### | 3.0403 | 2.3634E-03 | 5.4070E-03 |
| ENSG00000281100 | 23.8674 | 6.8479 | ##### | 3.0399 | 2.3664E-03 | 5.4135E-03 |
| ENSG00000281358\|RASSF1-AS1 | 66.9000 | -0.9455 | ##### | -3.0391 | 2.3730E-03 | 5.4281E-03 |
| ENSG00000289478 | 51.6200 | -8.6719 | ##### | -3.0386 | 2.3772E-03 | 5.4372E-03 |
| ENSG00000174885\|NLRP6 | 18.8413 | 2.9830 | ##### | 3.0374 | 2.3859E-03 | 5.4568E-03 |
| ENSG00000185837\|HDHD5-AS1 | 43.4321 | -8.3741 | ##### | -3.0371 | 2.3883E-03 | 5.4619E-03 |
| ENSG00000174600\|CMKLR1 | 9.4205 | -3.9779 | ##### | -3.0370 | 2.3895E-03 | 5.4643E-03 |
| ENSG00000237596\|PDE7B-AS1 | 15.3396 | -1.5000 | ##### | -3.0369 | 2.3899E-03 | 5.4644E-03 |
| ENSG00000261334 | 5.8690 | 5.0274 | ##### | 3.0369 | 2.3899E-03 | 5.4644E-03 |
| ENSG00000149256\|TENM4 | 40.9075 | 2.9995 | ##### | 3.0366 | 2.3926E-03 | 5.4701E-03 |
| ENSG00000136522\|MRPL47 | 3013.3639 | -0.3775 | ##### | -3.0358 | 2.3989E-03 | 5.4839E-03 |
| ENSG00000166126\|AMN | 303.1075 | -0.6059 | ##### | -3.0345 | 2.4091E-03 | 5.5068E-03 |
| ENSG00000224728\|IMPDH1P8 | 20.4236 | 1.3788 | ##### | 3.0343 | 2.4111E-03 | 5.5110E-03 |
| ENSG00000033178\|UBA6 | 6347.3315 | 0.3766 | ##### | 3.0340 | 2.4131E-03 | 5.5152E-03 |
| ENSG00000283355 | 23.9875 | -7.4590 | ##### | -3.0339 | 2.4143E-03 | 5.5176E-03 |
| ENSG00000224660\|SH3BP5-AS1 | 479.8224 | -0.8643 | ##### | -3.0338 | 2.4152E-03 | 5.5191E-03 |
| ENSG00000168159\|RNF187 | 8503.3205 | -0.3727 | ##### | -3.0331 | 2.4203E-03 | 5.5305E-03 |
| ENSG00000149451\|ADAM33 | 28.3822 | -2.8464 | ##### | -3.0325 | 2.4252E-03 | 5.5411E-03 |
| ENSG00000119938\|PPP1R3C | 377.8311 | -0.4777 | ##### | -3.0320 | 2.4294E-03 | 5.5503E-03 |
| ENSG00000268670 | 15.6295 | 2.1837 | ##### | 3.0313 | 2.4348E-03 | 5.5619E-03 |
| ENSG00000135324\|MRAP2 | 136.8943 | -0.6585 | ##### | -3.0313 | 2.4350E-03 | 5.5619E-03 |
| ENSG00000253508 | 20.4189 | -2.2160 | ##### | -3.0313 | 2.4350E-03 | 5.5619E-03 |
| ENSG00000240898\|RPL21P125 | 15.2726 | 2.4441 | ##### | 3.0312 | 2.4361E-03 | 5.5639E-03 |
| ENSG00000124422\|USP22 | 30555.3114 | 0.3309 | ##### | 3.0307 | 2.4398E-03 | 5.5719E-03 |
| ENSG00000186654\|PRR5 | 844.6285 | -0.4163 | ##### | -3.0301 | 2.4445E-03 | 5.5823E-03 |
| ENSG00000176472\|ZNF575 | 158.7980 | -0.6919 | ##### | -3.0292 | 2.4523E-03 | 5.5996E-03 |
| ENSG00000154646\|TMPRSS15 | 32.0140 | -5.2245 | ##### | -3.0286 | 2.4567E-03 | 5.6094E-03 |
| ENSG00000265806\|MIR4292 | 16.5402 | -1.1660 | ##### | -3.0286 | 2.4571E-03 | 5.6099E-03 |
| ENSG00000177352\|CCDC71 | 1013.3280 | -0.4079 | ##### | -3.0284 | 2.4583E-03 | 5.6120E-03 |
| ENSG00000068438\|FTSJ1 | 3579.7303 | -0.3779 | ##### | -3.0275 | 2.4657E-03 | 5.6284E-03 |
| ENSG00000197124\|ZNF682 | 8.5164 | -5.4882 | ##### | -3.0273 | 2.4676E-03 | 5.6323E-03 |
| ENSG00000146267\|FAXC | 282.5128 | 0.5451 | ##### | 3.0263 | 2.4758E-03 | 5.6507E-03 |
| ENSG00000279348 | 255.1271 | 0.6473 | ##### | 3.0261 | 2.4773E-03 | 5.6537E-03 |
| ENSG00000160972\|PPP1R16A | 6953.9148 | 0.3682 | ##### | 3.0253 | 2.4835E-03 | 5.6675E-03 |
| ENSG00000131558\|EXOC4 | 6077.0346 | -0.3593 | ##### | -3.0244 | 2.4911E-03 | 5.6843E-03 |
| ENSG00000258757 | 67.3547 | 0.7047 | ##### | 3.0238 | 2.4961E-03 | 5.6953E-03 |
| ENSG00000203496 | 21.4366 | -6.8337 | ##### | -3.0238 | 2.4965E-03 | 5.6956E-03 |
| ENSG00000169862\|CTNND2 | 16.2853 | 2.3370 | ##### | 3.0238 | 2.4966E-03 | 5.6956E-03 |
| ENSG00000289428 | 25.8944 | -2.1165 | ##### | -3.0237 | 2.4974E-03 | 5.6970E-03 |
| ENSG00000104879\|CKM | 25.0017 | -7.3609 | ##### | -3.0230 | 2.5028E-03 | 5.7089E-03 |
| ENSG00000259005 | 12.9254 | -6.1884 | ##### | -3.0226 | 2.5059E-03 | 5.7155E-03 |
| ENSG00000166783\|MARF1 | 4247.7798 | 0.3697 | ##### | 3.0226 | 2.5061E-03 | 5.7155E-03 |
| ENSG00000184363\|PKP3 | 1947.8522 | -0.3743 | ##### | -3.0222 | 2.5092E-03 | 5.7220E-03 |
| ENSG00000168090\|COPS6 | 13594.8734 | 0.3684 | ##### | 3.0216 | 2.5146E-03 | 5.7339E-03 |
| ENSG00000289123 | 25.5811 | 6.2320 | ##### | 3.0212 | 2.5179E-03 | 5.7412E-03 |
| ENSG00000267147\|LINC01842 | 151.2315 | 0.6464 | ##### | 3.0211 | 2.5184E-03 | 5.7417E-03 |
| ENSG00000290842\|CROCCP3 | 152.3981 | 0.5635 | ##### | 3.0206 | 2.5231E-03 | 5.7522E-03 |
| ENSG00000263874\|LASP1NB | 40.8631 | -2.3592 | ##### | -3.0204 | 2.5243E-03 | 5.7543E-03 |
| ENSG00000162402\|USP24 | 7872.8308 | -0.3626 | ##### | -3.0204 | 2.5246E-03 | 5.7546E-03 |
| ENSG00000224712\|NPIPA3 | 128.8429 | 0.6307 | ##### | 3.0195 | 2.5319E-03 | 5.7707E-03 |

| | | | | | | |
|---|---|---|---|---|---|---|
| ENSG00000122218\|COPA | 32840.9486 | 0.3557 | ##### | 3.0194 | 2.5326E-03 | 5.7719E-03 |
| ENSG00000139624\|CERS5 | 2975.1709 | 0.3810 | ##### | 3.0193 | 2.5334E-03 | 5.7733E-03 |
| ENSG00000293093 | 4.7872 | -5.3415 | ##### | -3.0193 | 2.5337E-03 | 5.7737E-03 |
| ENSG00000273137\|SELENOO-AS1 | 52.5324 | 0.7720 | ##### | 3.0193 | 2.5340E-03 | 5.7737E-03 |
| ENSG00000145331\|TRMT10A | 713.3006 | -0.4124 | ##### | -3.0189 | 2.5365E-03 | 5.7792E-03 |
| ENSG00000286205 | 13.1656 | 2.8580 | ##### | 3.0187 | 2.5384E-03 | 5.7831E-03 |
| ENSG00000149131\|SERPING1 | 215.9980 | -0.5382 | ##### | -3.0183 | 2.5423E-03 | 5.7914E-03 |
| ENSG00000290397 | 52.1707 | -7.8549 | ##### | -3.0179 | 2.5456E-03 | 5.7985E-03 |
| ENSG00000272372 | 24.3298 | 6.6275 | ##### | 3.0170 | 2.5528E-03 | 5.8144E-03 |
| ENSG00000258092 | 133.3884 | 1.1958 | ##### | 3.0168 | 2.5545E-03 | 5.8179E-03 |
| ENSG00000163697\|APBB2 | 3573.6542 | 0.3802 | ##### | 3.0162 | 2.5598E-03 | 5.8295E-03 |
| ENSG00000134438\|RAX | 11.1126 | 4.7905 | ##### | 3.0157 | 2.5638E-03 | 5.8382E-03 |
| ENSG00000137776\|SLTM | 9719.3655 | 0.3547 | ##### | 3.0154 | 2.5660E-03 | 5.8428E-03 |
| ENSG00000171962\|DRC3 | 172.3176 | 0.5612 | ##### | 3.0148 | 2.5711E-03 | 5.8538E-03 |
| ENSG00000271943\|TPM4P2 | 31.2646 | -1.1740 | ##### | -3.0144 | 2.5745E-03 | 5.8612E-03 |
| ENSG00000176102\|CSTF3 | 4702.8970 | -0.3613 | ##### | -3.0143 | 2.5757E-03 | 5.8636E-03 |
| ENSG00000115705\|TPO | 3.5065 | -4.9791 | ##### | -3.0134 | 2.5836E-03 | 5.8812E-03 |
| ENSG00000154518\|ATP5MC3 | 12025.4441 | 0.2931 | ##### | 3.0129 | 2.5873E-03 | 5.8890E-03 |
| ENSG00000105254\|TBCB | 9656.5189 | -0.3461 | ##### | -3.0126 | 2.5900E-03 | 5.8947E-03 |
| ENSG00000230869\|AGAP10P | 76.5577 | 0.7702 | ##### | 3.0124 | 2.5920E-03 | 5.8988E-03 |
| ENSG00000237772\|LINC02631 | 21.0193 | 7.2867 | ##### | 3.0121 | 2.5947E-03 | 5.9045E-03 |
| ENSG00000132912\|DCTN4 | 4305.7966 | 0.3620 | ##### | 3.0103 | 2.6099E-03 | 5.9386E-03 |
| ENSG00000079999\|KEAP1 | 8255.4164 | -0.3497 | ##### | -3.0094 | 2.6172E-03 | 5.9549E-03 |
| ENSG00000168394\|TAP1 | 12943.0500 | 0.3079 | ##### | 3.0091 | 2.6199E-03 | 5.9606E-03 |
| ENSG00000290717 | 54.4521 | -0.8740 | ##### | -3.0090 | 2.6208E-03 | 5.9622E-03 |
| ENSG00000285763 | 19.1830 | 1.7671 | ##### | 3.0089 | 2.6223E-03 | 5.9651E-03 |
| ENSG00000162592\|CCDC27 | 15.4028 | 1.3212 | ##### | 3.0082 | 2.6278E-03 | 5.9772E-03 |
| ENSG00000267651 | 15.9909 | 4.6848 | ##### | 3.0082 | 2.6283E-03 | 5.9779E-03 |
| ENSG00000100197\|CYP2D6 | 95.7322 | 0.7688 | ##### | 3.0079 | 2.6308E-03 | 5.9827E-03 |
| ENSG00000164051\|CCDC51 | 1932.3559 | 0.4006 | ##### | 3.0079 | 2.6309E-03 | 5.9827E-03 |
| ENSG00000124209\|RAB22A | 4196.4715 | 0.3694 | ##### | 3.0075 | 2.6338E-03 | 5.9888E-03 |
| ENSG00000166825\|ANPEP | 262.1455 | 2.1229 | ##### | 3.0069 | 2.6391E-03 | 6.0006E-03 |
| ENSG00000111335\|OAS2 | 20.9958 | -4.5058 | ##### | -3.0066 | 2.6418E-03 | 6.0062E-03 |
| ENSG00000286106\|NOTCH2NLR | 23.0138 | -2.6118 | ##### | -3.0064 | 2.6435E-03 | 6.0096E-03 |
| ENSG00000010165\|METTL13 | 5501.8402 | 0.3407 | ##### | 3.0058 | 2.6492E-03 | 6.0222E-03 |
| ENSG00000105701\|FKBP8 | 11384.3908 | 0.3170 | ##### | 3.0054 | 2.6520E-03 | 6.0277E-03 |
| ENSG00000008853\|RHOBTB2 | 5281.5701 | -0.3567 | ##### | -3.0054 | 2.6521E-03 | 6.0277E-03 |
| ENSG00000272153 | 56.4580 | 0.8397 | ##### | 3.0054 | 2.6527E-03 | 6.0286E-03 |
| ENSG00000117600\|PLPPR4 | 68.1034 | 1.0018 | ##### | 3.0053 | 2.6532E-03 | 6.0293E-03 |
| ENSG00000147099\|HDAC8 | 2063.7639 | 0.3874 | ##### | 3.0051 | 2.6546E-03 | 6.0321E-03 |
| ENSG00000113621\|TXNDC15 | 2925.3275 | -0.3815 | ##### | -3.0051 | 2.6550E-03 | 6.0326E-03 |
| ENSG00000138101\|DTNB | 1180.7172 | 0.3968 | ##### | 3.0050 | 2.6556E-03 | 6.0327E-03 |
| ENSG00000283235 | 27.3184 | 7.0115 | ##### | 3.0050 | 2.6557E-03 | 6.0327E-03 |
| ENSG00000203993\|ARRDC1-AS1 | 973.4481 | -0.4024 | ##### | -3.0050 | 2.6556E-03 | 6.0327E-03 |
| ENSG00000263986 | 321.4754 | 0.4552 | ##### | 3.0049 | 2.6564E-03 | 6.0339E-03 |
| ENSG00000123144\|TRIR | 7848.6934 | 0.3553 | ##### | 3.0049 | 2.6566E-03 | 6.0339E-03 |
| ENSG00000224236\|MRRFP1 | 4.8444 | -4.5795 | ##### | -3.0046 | 2.6590E-03 | 6.0389E-03 |
| ENSG00000229781\|RRN3P4 | 14.8412 | -4.3525 | ##### | -3.0044 | 2.6612E-03 | 6.0434E-03 |
| ENSG00000126698\|DNAJC8 | 8632.5377 | 0.3723 | ##### | 3.0040 | 2.6649E-03 | 6.0513E-03 |
| ENSG00000151490\|PTPRO | 11.2261 | -1.5381 | ##### | -3.0034 | 2.6694E-03 | 6.0611E-03 |
| ENSG00000186583\|SPATC1 | 14.2196 | 3.2213 | ##### | 3.0030 | 2.6729E-03 | 6.0687E-03 |
| ENSG00000147437\|GNRH1 | 46.6486 | 0.9081 | ##### | 3.0029 | 2.6744E-03 | 6.0715E-03 |
| ENSG00000267938\|EIF1P6 | 13.7657 | 5.5663 | ##### | 3.0028 | 2.6752E-03 | 6.0730E-03 |
| ENSG00000177427\|MIEF2 | 518.1728 | -0.4338 | ##### | -3.0024 | 2.6782E-03 | 6.0793E-03 |
| ENSG00000285805 | 17.7583 | -1.6729 | ##### | -3.0020 | 2.6819E-03 | 6.0873E-03 |
| ENSG00000247950\|SEC24B-AS1 | 22.8229 | -2.0433 | ##### | -3.0019 | 2.6832E-03 | 6.0896E-03 |
| ENSG00000203760\|CENPW | 1124.5340 | 0.3914 | ##### | 3.0015 | 2.6868E-03 | 6.0973E-03 |
| ENSG00000266642 | 72.3085 | -1.1510 | ##### | -3.0010 | 2.6913E-03 | 6.1072E-03 |
| ENSG00000280431 | 6.3900 | 5.9310 | ##### | 3.0006 | 2.6946E-03 | 6.1140E-03 |
| ENSG00000213204 | 14.8743 | -2.2734 | ##### | -3.0005 | 2.6952E-03 | 6.1151E-03 |
| ENSG00000277806 | 17.1361 | -1.8392 | ##### | -2.9982 | 2.7158E-03 | 6.1607E-03 |
| ENSG00000114902\|SPCS1 | 5514.0828 | 0.3724 | ##### | 2.9982 | 2.7157E-03 | 6.1607E-03 |
| ENSG00000102098\|SCML2 | 1392.0855 | 0.4076 | ##### | 2.9981 | 2.7168E-03 | 6.1627E-03 |

| | | | | | | |
|---|---|---|---|---|---|---|
| ENSG00000245156 | 71.2508 | 2.0123 | ##### | 2.9971 | 2.7254E-03 | 6.1817E-03 |
| ENSG00000134086\|VHL | 6603.3238 | -0.3611 | ##### | -2.9967 | 2.7290E-03 | 6.1893E-03 |
| ENSG00000119778\|ATAD2B | 1305.0830 | -0.4087 | ##### | -2.9963 | 2.7324E-03 | 6.1965E-03 |
| ENSG00000260924\|LINC01311 | 78.0859 | 0.7511 | ##### | 2.9959 | 2.7362E-03 | 6.2047E-03 |
| ENSG00000114988\|LMAN2L | 3623.5320 | 0.3357 | ##### | 2.9954 | 2.7406E-03 | 6.2142E-03 |
| ENSG00000142599\|RERE | 5382.3589 | 0.3509 | ##### | 2.9948 | 2.7459E-03 | 6.2258E-03 |
| ENSG00000292994 | 66.4356 | -0.7575 | ##### | -2.9948 | 2.7461E-03 | 6.2259E-03 |
| ENSG00000253704\|VIRMA-DT | 18.3506 | -2.1640 | ##### | -2.9946 | 2.7480E-03 | 6.2297E-03 |
| ENSG00000188153\|COL4A5 | 3092.1692 | 0.3755 | ##### | 2.9940 | 2.7531E-03 | 6.2407E-03 |
| ENSG00000196655\|TRAPPC4 | 2859.3217 | 0.3888 | ##### | 2.9934 | 2.7590E-03 | 6.2535E-03 |
| ENSG00000135387\|CAPRIN1 | 22304.0356 | 0.3476 | ##### | 2.9933 | 2.7599E-03 | 6.2552E-03 |
| ENSG00000251013\|GAPDHP62 | 30.1114 | -1.9077 | ##### | -2.9919 | 2.7721E-03 | 6.2823E-03 |
| ENSG00000166710\|B2M | 20057.6359 | -0.3177 | ##### | -2.9916 | 2.7753E-03 | 6.2890E-03 |
| ENSG00000022277\|RTF2 | 7613.9905 | 0.3632 | ##### | 2.9913 | 2.7780E-03 | 6.2948E-03 |
| ENSG00000285804 | 21.0155 | -1.8515 | ##### | -2.9912 | 2.7788E-03 | 6.2960E-03 |
| ENSG00000138378\|STAT4 | 93.5839 | -3.7510 | ##### | -2.9909 | 2.7812E-03 | 6.3011E-03 |
| ENSG00000244405\|ETV5 | 3390.9474 | -0.3094 | ##### | -2.9907 | 2.7838E-03 | 6.3064E-03 |
| ENSG00000145375\|AFG2A | 2242.3258 | 0.3862 | ##### | 2.9901 | 2.7886E-03 | 6.3170E-03 |
| ENSG00000119950\|MXI1 | 1121.3266 | 0.3820 | ##### | 2.9898 | 2.7912E-03 | 6.3222E-03 |
| ENSG00000242578 | 3.5679 | 5.5882 | ##### | 2.9896 | 2.7932E-03 | 6.3264E-03 |
| ENSG00000263859 | 65.7155 | 0.8235 | ##### | 2.9895 | 2.7948E-03 | 6.3295E-03 |
| ENSG00000249740\|OSMR-DT | 145.1576 | 0.5710 | ##### | 2.9894 | 2.7951E-03 | 6.3298E-03 |
| ENSG00000215023 | 13.9628 | -5.9292 | ##### | -2.9893 | 2.7961E-03 | 6.3309E-03 |
| ENSG00000135205\|CCDC146 | 110.7867 | -0.6050 | ##### | -2.9893 | 2.7960E-03 | 6.3309E-03 |
| ENSG00000213742\|ZNF337-AS1 | 653.7647 | 0.6593 | ##### | 2.9892 | 2.7975E-03 | 6.3337E-03 |
| ENSG00000167515\|TRAPPC2L | 2691.8092 | 0.3587 | ##### | 2.9890 | 2.7989E-03 | 6.3363E-03 |
| ENSG00000150051\|MKX | 111.1024 | 0.8358 | ##### | 2.9889 | 2.7999E-03 | 6.3380E-03 |
| ENSG00000239789\|MRPS17 | 3240.3393 | -0.2810 | ##### | -2.9888 | 2.8005E-03 | 6.3390E-03 |
| ENSG00000254894\|NAV2-AS1 | 28.4249 | -1.9055 | ##### | -2.9885 | 2.8036E-03 | 6.3455E-03 |
| ENSG00000196459\|TRAPPC2 | 615.1193 | -0.4262 | ##### | -2.9881 | 2.8069E-03 | 6.3526E-03 |
| ENSG00000204682\|MIR1915HG | 2344.8525 | 0.3583 | ##### | 2.9880 | 2.8082E-03 | 6.3551E-03 |
| ENSG00000184068\|SREBF2-AS1 | 118.4392 | -0.9034 | ##### | -2.9879 | 2.8088E-03 | 6.3560E-03 |
| ENSG00000120937\|NPPB | 6.2587 | 4.3567 | ##### | 2.9867 | 2.8202E-03 | 6.3811E-03 |
| ENSG00000187546\|AGMO | 15.6477 | -3.5269 | ##### | -2.9865 | 2.8218E-03 | 6.3843E-03 |
| ENSG00000249889\|ALG1L11P | 18.7015 | -1.2710 | ##### | -2.9864 | 2.8228E-03 | 6.3861E-03 |
| ENSG00000188522\|FAM83G | 3706.9703 | -0.3729 | ##### | -2.9863 | 2.8233E-03 | 6.3868E-03 |
| ENSG00000227540\|DNAJC9-AS1 | 94.0535 | -0.9065 | ##### | -2.9861 | 2.8251E-03 | 6.3904E-03 |
| ENSG00000290824\|DDX11L16 | 9.5029 | 1.5327 | ##### | 2.9860 | 2.8263E-03 | 6.3923E-03 |
| ENSG00000182224\|CYB5D1 | 1874.8781 | 0.5858 | ##### | 2.9860 | 2.8264E-03 | 6.3923E-03 |
| ENSG00000158748\|HTR6 | 47.7068 | 2.4091 | ##### | 2.9858 | 2.8288E-03 | 6.3967E-03 |
| ENSG00000167862\|MRPL58 | 2049.8639 | -0.3917 | ##### | -2.9858 | 2.8288E-03 | 6.3967E-03 |
| ENSG00000253390\|ENTPD4-DT | 23.4827 | 2.3991 | ##### | 2.9856 | 2.8306E-03 | 6.4005E-03 |
| ENSG00000112531\|QKI | 7439.2124 | 0.3632 | ##### | 2.9855 | 2.8311E-03 | 6.4010E-03 |
| ENSG00000160201\|U2AF1 | 4083.3642 | 0.3581 | ##### | 2.9851 | 2.8353E-03 | 6.4100E-03 |
| ENSG00000269961\|ERBIN-DT | 26.8577 | -1.3582 | ##### | -2.9847 | 2.8385E-03 | 6.4168E-03 |
| ENSG00000164610\|RP9 | 1267.0346 | -0.3783 | ##### | -2.9840 | 2.8448E-03 | 6.4305E-03 |
| ENSG00000251692\|PTX4 | 24.5566 | -6.9015 | ##### | -2.9836 | 2.8492E-03 | 6.4400E-03 |
| ENSG00000291105 | 273.4006 | 3.9813 | ##### | 2.9820 | 2.8636E-03 | 6.4719E-03 |
| ENSG00000178691\|SUZ12 | 8270.1872 | 0.3171 | ##### | 2.9817 | 2.8662E-03 | 6.4775E-03 |
| ENSG00000226281 | 7.4110 | -4.6407 | ##### | -2.9813 | 2.8698E-03 | 6.4852E-03 |
| ENSG00000134077\|THUMPD3 | 4968.5259 | 0.3711 | ##### | 2.9812 | 2.8714E-03 | 6.4881E-03 |
| ENSG00000293196 | 37.0458 | -1.4785 | ##### | -2.9809 | 2.8737E-03 | 6.4930E-03 |
| ENSG00000231588\|SHQ1P1 | 15.9381 | -1.7689 | ##### | -2.9807 | 2.8762E-03 | 6.4979E-03 |
| ENSG00000136238\|RAC1 | 42020.4036 | 0.3026 | ##### | 2.9804 | 2.8786E-03 | 6.5031E-03 |
| ENSG00000185104\|FAF1 | 5368.3817 | -0.3665 | ##### | -2.9803 | 2.8800E-03 | 6.5055E-03 |
| ENSG00000138400\|MDH1B | 180.5049 | 0.6163 | ##### | 2.9802 | 2.8807E-03 | 6.5067E-03 |
| ENSG00000229409 | 6.7892 | 5.8601 | ##### | 2.9801 | 2.8817E-03 | 6.5085E-03 |
| ENSG00000169271\|HSPB3 | 9.9031 | 2.5477 | ##### | 2.9799 | 2.8832E-03 | 6.5114E-03 |
| ENSG00000215217\|CFAP90 | 12.1315 | -2.8926 | ##### | -2.9797 | 2.8852E-03 | 6.5155E-03 |
| ENSG00000274985\|PTCHD3P1 | 222.1438 | -0.8317 | ##### | -2.9795 | 2.8867E-03 | 6.5184E-03 |
| ENSG00000093167\|LRRFIP2 | 2998.6444 | -0.3687 | ##### | -2.9794 | 2.8877E-03 | 6.5200E-03 |
| ENSG00000105058\|FAM32A | 4565.6660 | 0.3716 | ##### | 2.9788 | 2.8942E-03 | 6.5338E-03 |
| ENSG00000186847\|KRT14 | 37.4328 | -8.1596 | ##### | -2.9788 | 2.8941E-03 | 6.5338E-03 |

| | | | | | | |
|---|---|---|---|---|---|---|
| ENSG00000188822\|CNR2 | 23.6338 | 7.3001 | ##### | 2.9787 | 2.8951E-03 | 6.5354E-03 |
| ENSG00000254872\|LINC02688 | 19.4708 | -2.4442 | ##### | -2.9783 | 2.8981E-03 | 6.5417E-03 |
| ENSG00000103005\|USB1 | 8901.8742 | -0.3525 | ##### | -2.9778 | 2.9028E-03 | 6.5518E-03 |
| ENSG00000276302 | 10.7885 | 5.2584 | ##### | 2.9777 | 2.9046E-03 | 6.5554E-03 |
| ENSG00000186710\|CFAP73 | 336.1434 | 0.6455 | ##### | 2.9776 | 2.9049E-03 | 6.5554E-03 |
| ENSG00000285948 | 878.0616 | 1.2322 | ##### | 2.9764 | 2.9168E-03 | 6.5818E-03 |
| ENSG00000135119\|RNFT2 | 392.2570 | 0.5499 | ##### | 2.9763 | 2.9173E-03 | 6.5825E-03 |
| ENSG00000226457\|RPL22P3 | 4.2990 | -2.9201 | ##### | -2.9761 | 2.9197E-03 | 6.5869E-03 |
| ENSG00000127993\|RBM48 | 1155.1731 | 0.4241 | ##### | 2.9761 | 2.9197E-03 | 6.5869E-03 |
| ENSG00000159256\|MORC3 | 2480.1906 | 0.3948 | ##### | 2.9760 | 2.9206E-03 | 6.5883E-03 |
| ENSG00000214132 | 7.6112 | -2.9060 | ##### | -2.9759 | 2.9211E-03 | 6.5889E-03 |
| ENSG00000293471 | 47.4394 | -1.0309 | ##### | -2.9759 | 2.9215E-03 | 6.5894E-03 |
| ENSG00000260815\|PPIAP47 | 4.6787 | -3.0990 | ##### | -2.9757 | 2.9228E-03 | 6.5919E-03 |
| ENSG00000100353\|EIF3D | 15819.4320 | -0.3752 | ##### | -2.9755 | 2.9247E-03 | 6.5957E-03 |
| ENSG00000288973 | 1115.2743 | -0.4027 | ##### | -2.9752 | 2.9283E-03 | 6.6032E-03 |
| ENSG00000219392\|ZNF602P | 18.2437 | 3.1761 | ##### | 2.9745 | 2.9350E-03 | 6.6178E-03 |
| ENSG00000234869\|LINC02925 | 44.5162 | -2.4537 | ##### | -2.9739 | 2.9409E-03 | 6.6306E-03 |
| ENSG00000275215\|RNA5-8SN3 | 195.7679 | 1.1398 | ##### | 2.9738 | 2.9411E-03 | 6.6306E-03 |
| ENSG00000176907\|TCIM | 4679.7484 | 0.3724 | ##### | 2.9738 | 2.9418E-03 | 6.6317E-03 |
| ENSG00000236609\|ZNF853 | 31.0639 | -2.0309 | ##### | -2.9733 | 2.9460E-03 | 6.6406E-03 |
| ENSG00000214706\|IFRD2 | 10309.1860 | 0.2370 | ##### | 2.9727 | 2.9518E-03 | 6.6531E-03 |
| ENSG00000187621\|TCL6 | 9.8344 | -6.2160 | ##### | -2.9727 | 2.9524E-03 | 6.6535E-03 |
| ENSG00000128383\|APOBEC3A | 20.8941 | 3.2222 | ##### | 2.9727 | 2.9523E-03 | 6.6535E-03 |
| ENSG00000228653\|HNRNPCP7 | 8.8085 | -2.7368 | ##### | -2.9726 | 2.9527E-03 | 6.6537E-03 |
| ENSG00000234268\|RPS27P19 | 13.5402 | 1.5093 | ##### | 2.9725 | 2.9536E-03 | 6.6554E-03 |
| ENSG00000275580 | 22.6784 | -6.5736 | ##### | -2.9723 | 2.9561E-03 | 6.6604E-03 |
| ENSG00000157191\|NECAP2 | 4168.1434 | -0.3531 | ##### | -2.9721 | 2.9580E-03 | 6.6643E-03 |
| ENSG00000198010\|DLGAP2 | 5.7512 | 4.8652 | ##### | 2.9720 | 2.9588E-03 | 6.6655E-03 |
| ENSG00000100299\|ARSA | 1928.1762 | -0.3713 | ##### | -2.9712 | 2.9667E-03 | 6.6828E-03 |
| ENSG00000105185\|PDCD5 | 7053.9831 | -0.3604 | ##### | -2.9706 | 2.9720E-03 | 6.6942E-03 |
| ENSG00000272787 | 24.5311 | 1.3232 | ##### | 2.9691 | 2.9870E-03 | 6.7274E-03 |
| ENSG00000213463\|SYNJ2BP | 4716.7831 | 0.3688 | ##### | 2.9689 | 2.9886E-03 | 6.7307E-03 |
| ENSG00000289235 | 23.6913 | 1.4092 | ##### | 2.9687 | 2.9910E-03 | 6.7354E-03 |
| ENSG00000231887\|PRH1 | 189.0158 | -0.5739 | ##### | -2.9686 | 2.9918E-03 | 6.7368E-03 |
| ENSG00000280195 | 322.8609 | 0.4974 | ##### | 2.9684 | 2.9932E-03 | 6.7394E-03 |
| ENSG00000289979 | 26.5597 | 1.0715 | ##### | 2.9678 | 2.9992E-03 | 6.7524E-03 |
| ENSG00000148090\|AUH | 970.7607 | -0.4007 | ##### | -2.9678 | 2.9996E-03 | 6.7528E-03 |
| ENSG00000183617\|MRPL54 | 1500.7132 | 0.3881 | ##### | 2.9677 | 3.0003E-03 | 6.7539E-03 |
| ENSG00000166908\|PIP4K2C | 5056.4759 | -0.3727 | ##### | -2.9673 | 3.0041E-03 | 6.7619E-03 |
| ENSG00000286302 | 20.7903 | -6.4728 | ##### | -2.9669 | 3.0084E-03 | 6.7711E-03 |
| ENSG00000198336\|MYL4 | 17.3964 | 3.6223 | ##### | 2.9667 | 3.0102E-03 | 6.7747E-03 |
| ENSG00000125388\|GRK4 | 194.5992 | -0.5250 | ##### | -2.9658 | 3.0188E-03 | 6.7934E-03 |
| ENSG00000163472\|TMEM79 | 1841.2044 | 0.3789 | ##### | 2.9655 | 3.0215E-03 | 6.7986E-03 |
| ENSG00000127955\|GNAI1 | 4573.5930 | -0.3214 | ##### | -2.9656 | 3.0214E-03 | 6.7986E-03 |
| ENSG00000263934\|SNORD3A | 10.4648 | 1.8074 | ##### | 2.9655 | 3.0218E-03 | 6.7988E-03 |
| ENSG00000228779 | 20.8485 | -6.8623 | ##### | -2.9652 | 3.0245E-03 | 6.8043E-03 |
| ENSG00000104818\|CGB2 | 43.7401 | -0.8952 | ##### | -2.9650 | 3.0272E-03 | 6.8099E-03 |
| ENSG00000225783\|MIAT | 26.0467 | 1.3041 | ##### | 2.9648 | 3.0287E-03 | 6.8127E-03 |
| ENSG00000213468\|FIRRE | 68.8498 | -2.1277 | ##### | -2.9643 | 3.0335E-03 | 6.8231E-03 |
| ENSG00000198841\|KTI12 | 1886.4883 | -0.3715 | ##### | -2.9638 | 3.0384E-03 | 6.8334E-03 |
| ENSG00000271537 | 8.9434 | 3.0705 | ##### | 2.9637 | 3.0396E-03 | 6.8356E-03 |
| ENSG00000234546\|LNCTAM34A | 44.6449 | 1.0092 | ##### | 2.9636 | 3.0405E-03 | 6.8373E-03 |
| ENSG00000124541\|RRP36 | 4086.6530 | 0.3561 | ##### | 2.9635 | 3.0417E-03 | 6.8394E-03 |
| ENSG00000166451\|CENPN | 6944.2423 | 0.3704 | ##### | 2.9632 | 3.0446E-03 | 6.8453E-03 |
| ENSG00000289152 | 8.6457 | -2.4556 | ##### | -2.9629 | 3.0473E-03 | 6.8509E-03 |
| ENSG00000226002\|GTF2IP14 | 67.2694 | 9.8713 | ##### | 2.9625 | 3.0513E-03 | 6.8593E-03 |
| ENSG00000107560\|RAB11FIP2 | 2837.4018 | -0.3575 | ##### | -2.9623 | 3.0539E-03 | 6.8648E-03 |
| ENSG00000168143\|FAM83B | 96.1715 | 3.6873 | ##### | 2.9609 | 3.0672E-03 | 6.8942E-03 |
| ENSG00000272359\|RNU4-89P | 11.2441 | -6.0074 | ##### | -2.9609 | 3.0678E-03 | 6.8950E-03 |
| ENSG00000103051\|COG4 | 4957.7464 | -0.3158 | ##### | -2.9607 | 3.0698E-03 | 6.8988E-03 |
| ENSG00000272333\|KMT2B | 7751.5437 | 0.3501 | ##### | 2.9594 | 3.0819E-03 | 6.9256E-03 |
| ENSG00000231131\|LNCAROD | 14.8013 | 3.0175 | ##### | 2.9578 | 3.0984E-03 | 6.9621E-03 |
| ENSG00000271547 | 6.5709 | -5.1053 | ##### | -2.9575 | 3.1016E-03 | 6.9688E-03 |

| | | | | | | |
|---|---|---|---|---|---|---|
| ENSG00000267185\|PTP4A2P1 | 23.1896 | -1.4264 | ##### | -2.9574 | 3.1027E-03 | 6.9707E-03 |
| ENSG00000290920 | 81.6859 | 0.7945 | ##### | 2.9573 | 3.1032E-03 | 6.9714E-03 |
| ENSG00000180370\|PAK2 | 15214.6201 | 0.3177 | ##### | 2.9569 | 3.1078E-03 | 6.9811E-03 |
| ENSG00000248881 | 150.2061 | -1.3682 | ##### | -2.9566 | 3.1108E-03 | 6.9875E-03 |
| ENSG00000277867 | 5.8602 | -4.6283 | ##### | -2.9565 | 3.1115E-03 | 6.9879E-03 |
| ENSG00000256951 | 5.8602 | -4.6283 | ##### | -2.9565 | 3.1115E-03 | 6.9879E-03 |
| ENSG00000266826\|IGBP1C | 7.3898 | 5.2090 | ##### | 2.9560 | 3.1161E-03 | 6.9977E-03 |
| ENSG00000153107\|ANAPC1 | 9901.1570 | -0.3529 | ##### | -2.9552 | 3.1243E-03 | 7.0156E-03 |
| ENSG00000196735\|HLA-DQA1 | 24.5402 | -6.9041 | ##### | -2.9546 | 3.1309E-03 | 7.0298E-03 |
| ENSG00000137221\|TJAP1 | 2990.6958 | 0.3504 | ##### | 2.9545 | 3.1321E-03 | 7.0320E-03 |
| ENSG00000274893 | 7.6943 | -5.8582 | ##### | -2.9541 | 3.1355E-03 | 7.0390E-03 |
| ENSG00000076043\|REXO2 | 4899.6265 | -0.3517 | ##### | -2.9538 | 3.1385E-03 | 7.0454E-03 |
| ENSG00000249641\|HOXC13-AS | 236.5945 | 0.5519 | ##### | 2.9536 | 3.1414E-03 | 7.0513E-03 |
| ENSG00000285685 | 8.7907 | 6.2596 | ##### | 2.9528 | 3.1488E-03 | 7.0675E-03 |
| ENSG00000157303\|SUSD3 | 26.8165 | -3.7553 | ##### | -2.9519 | 3.1579E-03 | 7.0873E-03 |
| ENSG00000143845\|ETNK2 | 566.8183 | 0.4348 | ##### | 2.9505 | 3.1729E-03 | 7.1205E-03 |
| ENSG00000236065 | 10.0544 | 6.3761 | ##### | 2.9504 | 3.1735E-03 | 7.1213E-03 |
| ENSG00000182013\|PNMA8A | 2443.6740 | 0.3753 | ##### | 2.9504 | 3.1739E-03 | 7.1216E-03 |
| ENSG00000272100 | 11.5325 | -1.4863 | ##### | -2.9502 | 3.1754E-03 | 7.1245E-03 |
| ENSG00000174132\|FAM174A | 847.0319 | -0.4397 | ##### | -2.9494 | 3.1838E-03 | 7.1426E-03 |
| ENSG00000131791\|PRKAB2 | 2991.7248 | 0.3637 | ##### | 2.9492 | 3.1858E-03 | 7.1461E-03 |
| ENSG00000166900\|STX3 | 1708.9500 | 0.3933 | ##### | 2.9492 | 3.1857E-03 | 7.1461E-03 |
| ENSG00000183808\|RBM12B | 3931.9381 | -0.3396 | ##### | -2.9491 | 3.1868E-03 | 7.1478E-03 |
| ENSG00000127528\|KLF2 | 875.4529 | -0.4126 | ##### | -2.9491 | 3.1872E-03 | 7.1483E-03 |
| ENSG00000241954 | 35.3516 | 7.3831 | ##### | 2.9490 | 3.1884E-03 | 7.1503E-03 |
| ENSG00000167553\|TUBA1C | ########## | 0.3574 | ##### | 2.9488 | 3.1904E-03 | 7.1544E-03 |
| ENSG00000226803\|ZNF451-AS1 | 67.1165 | -0.8064 | ##### | -2.9486 | 3.1925E-03 | 7.1585E-03 |
| ENSG00000261606 | 23.7560 | 2.0440 | ##### | 2.9482 | 3.1966E-03 | 7.1671E-03 |
| ENSG00000197548\|ATG7 | 4874.8095 | 0.3551 | ##### | 2.9471 | 3.2076E-03 | 7.1914E-03 |
| ENSG00000279430 | 10.2898 | 5.8898 | ##### | 2.9464 | 3.2148E-03 | 7.2069E-03 |
| ENSG00000131652\|THOC6 | 3402.1657 | 0.3647 | ##### | 2.9458 | 3.2217E-03 | 7.2217E-03 |
| ENSG00000100029\|PES1 | 12065.7914 | 0.3447 | ##### | 2.9451 | 3.2289E-03 | 7.2374E-03 |
| ENSG00000214199\|EEF1A1P12 | 25.2741 | 0.9248 | ##### | 2.9447 | 3.2329E-03 | 7.2459E-03 |
| ENSG00000125648\|SLC25A23 | 3679.2003 | 0.3575 | ##### | 2.9446 | 3.2342E-03 | 7.2483E-03 |
| ENSG00000236700\|LINC01010 | 19.0034 | 2.9450 | ##### | 2.9443 | 3.2365E-03 | 7.2527E-03 |
| ENSG00000136143\|SUCLA2 | 3861.4570 | -0.3629 | ##### | -2.9438 | 3.2419E-03 | 7.2645E-03 |
| ENSG00000279148 | 115.9964 | 0.6707 | ##### | 2.9436 | 3.2444E-03 | 7.2694E-03 |
| ENSG00000229970 | 113.1908 | -0.9048 | ##### | -2.9435 | 3.2456E-03 | 7.2716E-03 |
| ENSG00000171307\|ZDHHC16 | 3405.1052 | -0.3803 | ##### | -2.9431 | 3.2499E-03 | 7.2805E-03 |
| ENSG00000161618\|ALDH16A1 | 1560.5970 | -0.3848 | ##### | -2.9427 | 3.2538E-03 | 7.2888E-03 |
| ENSG00000233138 | 8.6887 | 1.6028 | ##### | 2.9425 | 3.2559E-03 | 7.2929E-03 |
| ENSG00000279259 | 53.7727 | -0.9230 | ##### | -2.9421 | 3.2595E-03 | 7.2999E-03 |
| ENSG00000290763\|SDHAP1 | 361.3304 | 0.6314 | ##### | 2.9422 | 3.2594E-03 | 7.2999E-03 |
| ENSG00000219665\|ZNF433-AS1 | 58.6774 | 0.8602 | ##### | 2.9411 | 3.2706E-03 | 7.3237E-03 |
| ENSG00000151005\|TKTL2 | 7.4960 | -6.3159 | ##### | -2.9411 | 3.2704E-03 | 7.3237E-03 |
| ENSG00000164241\|C5orf63 | 146.3552 | -0.5699 | ##### | -2.9409 | 3.2728E-03 | 7.3282E-03 |
| ENSG00000205293\|LINC01602 | 19.0752 | -6.9457 | ##### | -2.9397 | 3.2848E-03 | 7.3545E-03 |
| ENSG00000261320 | 9.4562 | -2.8548 | ##### | -2.9394 | 3.2881E-03 | 7.3607E-03 |
| ENSG00000265168 | 4.7842 | -2.1989 | ##### | -2.9395 | 3.2880E-03 | 7.3607E-03 |
| ENSG00000234509\|SOD1-DT | 11.0607 | -3.7201 | ##### | -2.9393 | 3.2892E-03 | 7.3625E-03 |
| ENSG00000108599\|AKAP10 | 2813.3353 | 0.3629 | ##### | 2.9389 | 3.2941E-03 | 7.3730E-03 |
| ENSG00000260115\|CDH8-AS1 | 6.9384 | -5.4092 | ##### | -2.9386 | 3.2970E-03 | 7.3789E-03 |
| ENSG00000293109\|ANKRD36BP1 | 76.2014 | -0.7617 | ##### | -2.9383 | 3.3005E-03 | 7.3858E-03 |
| ENSG00000204745 | 918.8290 | -0.4331 | ##### | -2.9383 | 3.3005E-03 | 7.3858E-03 |
| ENSG00000130714\|POMT1 | 2689.3777 | -0.2768 | ##### | -2.9382 | 3.3012E-03 | 7.3868E-03 |
| ENSG00000077097\|TOP2B | 10613.4995 | 0.3399 | ##### | 2.9373 | 3.3103E-03 | 7.4063E-03 |
| ENSG00000044090\|CUL7 | 3886.2552 | 0.3450 | ##### | 2.9373 | 3.3104E-03 | 7.4063E-03 |
| ENSG00000075914\|EXOSC7 | 2270.5654 | 0.3814 | ##### | 2.9372 | 3.3122E-03 | 7.4096E-03 |
| ENSG00000283415 | 18.0152 | -1.1519 | ##### | -2.9370 | 3.3141E-03 | 7.4134E-03 |
| ENSG00000242058\|RPS4XP19 | 11.9292 | 5.2040 | ##### | 2.9366 | 3.3185E-03 | 7.4221E-03 |
| ENSG00000035115\|SH3YL1 | 695.8860 | -0.4204 | ##### | -2.9366 | 3.3184E-03 | 7.4221E-03 |
| ENSG00000142233\|NTN5 | 29.2427 | -1.7220 | ##### | -2.9365 | 3.3190E-03 | 7.4226E-03 |
| ENSG00000100815\|TRIP11 | 3201.3777 | -0.3637 | ##### | -2.9363 | 3.3214E-03 | 7.4274E-03 |

| | | | | | | |
|---|---|---|---|---|---|---|
| ENSG00000060642\|PIGV | 1046.0435 | 0.3892 | ##### | 2.9363 | 3.3219E-03 | 7.4281E-03 |
| ENSG00000261594\|TPBGL | 110.9381 | 0.6505 | ##### | 2.9362 | 3.3224E-03 | 7.4286E-03 |
| ENSG00000261687 | 9.6972 | -2.5616 | ##### | -2.9360 | 3.3248E-03 | 7.4333E-03 |
| ENSG00000159625\|DRC7 | 32.0536 | -3.1939 | ##### | -2.9359 | 3.3259E-03 | 7.4353E-03 |
| ENSG00000267475\|NUDT19-DT | 71.6147 | -0.8545 | ##### | -2.9354 | 3.3313E-03 | 7.4468E-03 |
| ENSG00000279632 | 11.3333 | 4.7061 | ##### | 2.9346 | 3.3397E-03 | 7.4649E-03 |
| ENSG00000205702\|CYP2D7 | 148.4349 | 0.6515 | ##### | 2.9336 | 3.3510E-03 | 7.4896E-03 |
| ENSG00000248019\|FAM13A-AS1 | 87.6718 | -1.1254 | ##### | -2.9335 | 3.3513E-03 | 7.4897E-03 |
| ENSG00000160781\|PAQR6 | 811.7758 | 0.4159 | ##### | 2.9334 | 3.3525E-03 | 7.4919E-03 |
| ENSG00000172071\|EIF2AK3 | 2384.4247 | 0.3717 | ##### | 2.9332 | 3.3544E-03 | 7.4956E-03 |
| ENSG00000233912 | 15.3397 | -4.3386 | ##### | -2.9332 | 3.3546E-03 | 7.4956E-03 |
| ENSG00000095303\|PTGS1 | 23.0696 | -3.3131 | ##### | -2.9331 | 3.3565E-03 | 7.4991E-03 |
| ENSG00000126581\|BECN1 | 7235.2131 | 0.3418 | ##### | 2.9329 | 3.3582E-03 | 7.5024E-03 |
| ENSG00000139722\|VPS37B | 4188.3416 | 0.3697 | ##### | 2.9319 | 3.3692E-03 | 7.5264E-03 |
| ENSG00000278206 | 24.6619 | -1.5873 | ##### | -2.9317 | 3.3712E-03 | 7.5304E-03 |
| ENSG00000232762 | 14.5563 | 5.0830 | ##### | 2.9314 | 3.3746E-03 | 7.5375E-03 |
| ENSG00000271092\|TLCD4-RWDD3 | 42.9903 | -1.8125 | ##### | -2.9313 | 3.3759E-03 | 7.5392E-03 |
| ENSG00000105146\|AURKC | 19.4776 | -3.2937 | ##### | -2.9313 | 3.3759E-03 | 7.5392E-03 |
| ENSG00000233033\|CASK-AS1 | 20.8291 | -1.0774 | ##### | -2.9311 | 3.3781E-03 | 7.5436E-03 |
| ENSG00000260160 | 17.7736 | -1.4692 | ##### | -2.9309 | 3.3797E-03 | 7.5466E-03 |
| ENSG00000105662\|CRTC1 | 1317.4473 | 0.3336 | ##### | 2.9306 | 3.3829E-03 | 7.5530E-03 |
| ENSG00000211456\|SACM1L | 4153.2080 | 0.3186 | ##### | 2.9304 | 3.3856E-03 | 7.5586E-03 |
| ENSG00000232300\|FAM215B | 56.2973 | 0.9202 | ##### | 2.9302 | 3.3877E-03 | 7.5626E-03 |
| ENSG00000280282 | 6.2648 | 4.1278 | ##### | 2.9300 | 3.3896E-03 | 7.5663E-03 |
| ENSG00000163945\|UVSSA | 1885.3979 | 0.5252 | ##### | 2.9299 | 3.3910E-03 | 7.5689E-03 |
| ENSG00000276418 | 136.4466 | -5.4221 | ##### | -2.9298 | 3.3921E-03 | 7.5707E-03 |
| ENSG00000278233\|RNA5-8SN2 | 105.4170 | 1.1360 | ##### | 2.9297 | 3.3934E-03 | 7.5731E-03 |
| ENSG00000110172\|CHORDC1 | 6646.1194 | 0.3532 | ##### | 2.9295 | 3.3955E-03 | 7.5772E-03 |
| ENSG00000178922\|HYI | 1724.6047 | -0.5289 | ##### | -2.9292 | 3.3985E-03 | 7.5834E-03 |
| ENSG00000205795\|CYS1 | 185.3613 | -0.5308 | ##### | -2.9283 | 3.4079E-03 | 7.6038E-03 |
| ENSG00000206172\|HBA1 | 12.3429 | -3.0805 | ##### | -2.9282 | 3.4091E-03 | 7.6059E-03 |
| ENSG00000196453\|ZNF777 | 2030.2096 | -0.3691 | ##### | -2.9281 | 3.4099E-03 | 7.6071E-03 |
| ENSG00000184924\|PTRHD1 | 1992.9557 | -0.3927 | ##### | -2.9280 | 3.4111E-03 | 7.6093E-03 |
| ENSG00000102900\|NUP93 | 9908.4703 | 0.2966 | ##### | 2.9276 | 3.4156E-03 | 7.6187E-03 |
| ENSG00000182732\|RGS6 | 27.0381 | 3.8129 | ##### | 2.9271 | 3.4212E-03 | 7.6307E-03 |
| ENSG00000254893\|RAP1BL | 81.1317 | 0.6458 | ##### | 2.9262 | 3.4312E-03 | 7.6524E-03 |
| ENSG00000241839\|PLEKHO2 | 795.3803 | 0.3842 | ##### | 2.9261 | 3.4326E-03 | 7.6549E-03 |
| ENSG00000221533\|MIR1184-1 | 12.1813 | -1.2868 | ##### | -2.9248 | 3.4468E-03 | 7.6860E-03 |
| ENSG00000286656 | 17.9997 | -1.6933 | ##### | -2.9242 | 3.4530E-03 | 7.6993E-03 |
| ENSG00000291081 | 302.2783 | 0.5099 | ##### | 2.9238 | 3.4580E-03 | 7.7097E-03 |
| ENSG00000289453 | 25.4843 | 1.1831 | ##### | 2.9233 | 3.4634E-03 | 7.7212E-03 |
| ENSG00000158428\|CATIP | 121.1834 | -0.6256 | ##### | -2.9231 | 3.4657E-03 | 7.7258E-03 |
| ENSG00000146433\|TMEM181 | 4116.9263 | -0.3551 | ##### | -2.9230 | 3.4668E-03 | 7.7277E-03 |
| ENSG00000272638 | 55.0493 | -0.8552 | ##### | -2.9219 | 3.4787E-03 | 7.7531E-03 |
| ENSG00000210196\|MT-TP | 417.0621 | 0.4063 | ##### | 2.9219 | 3.4787E-03 | 7.7531E-03 |
| ENSG00000102890\|ELMO3 | 376.6802 | 0.4746 | ##### | 2.9219 | 3.4791E-03 | 7.7534E-03 |
| ENSG00000100083\|GGA1 | 3299.7829 | 0.3609 | ##### | 2.9216 | 3.4823E-03 | 7.7600E-03 |
| ENSG00000132383\|RPA1 | 10252.0611 | 0.2576 | ##### | 2.9209 | 3.4901E-03 | 7.7768E-03 |
| ENSG00000149968\|MMP3 | 30.9590 | 2.1733 | ##### | 2.9208 | 3.4915E-03 | 7.7792E-03 |
| ENSG00000105204\|DYRK1B | 639.4059 | -0.4307 | ##### | -2.9207 | 3.4922E-03 | 7.7803E-03 |
| ENSG00000172269\|DPAGT1 | 5275.9663 | -0.3626 | ##### | -2.9206 | 3.4931E-03 | 7.7816E-03 |
| ENSG00000105290\|APLP1 | 6412.9247 | -0.3502 | ##### | -2.9199 | 3.5009E-03 | 7.7985E-03 |
| ENSG00000127249\|ATP13A4 | 9.8648 | -4.3181 | ##### | -2.9199 | 3.5013E-03 | 7.7988E-03 |
| ENSG00000240401 | 18.0999 | -1.4721 | ##### | -2.9194 | 3.5065E-03 | 7.8099E-03 |
| ENSG00000115947\|ORC4 | 2331.6819 | 0.3687 | ##### | 2.9194 | 3.5068E-03 | 7.8099E-03 |
| ENSG00000180776\|ZDHHC20 | 7138.6537 | 0.3433 | ##### | 2.9189 | 3.5122E-03 | 7.8214E-03 |
| ENSG00000086848\|ALG9 | 1423.1842 | -0.3731 | ##### | -2.9177 | 3.5259E-03 | 7.8513E-03 |
| ENSG00000107186\|MPDZ | 1296.9446 | 0.3864 | ##### | 2.9174 | 3.5300E-03 | 7.8599E-03 |
| ENSG00000146070\|PLA2G7 | 30.2619 | -7.8512 | ##### | -2.9164 | 3.5406E-03 | 7.8829E-03 |
| ENSG00000119541\|VPS4B | 5746.1375 | 0.3672 | ##### | 2.9161 | 3.5445E-03 | 7.8908E-03 |
| ENSG00000152583\|SPARCL1 | 31.7391 | -2.6703 | ##### | -2.9160 | 3.5458E-03 | 7.8933E-03 |
| ENSG00000015285\|WAS | 26.5211 | 1.3699 | ##### | 2.9151 | 3.5559E-03 | 7.9151E-03 |
| ENSG00000100410\|PHF5A | 2970.3643 | 0.3692 | ##### | 2.9151 | 3.5562E-03 | 7.9152E-03 |

| | | | | | | |
|---|---|---|---|---|---|---|
| ENSG00000230289 | 8.9042 | 2.6772 | ##### | 2.9148 | 3.5592E-03 | 7.9212E-03 |
| ENSG00000253633\|LINC03047 | 23.6234 | -1.8942 | ##### | -2.9147 | 3.5605E-03 | 7.9235E-03 |
| ENSG00000275238\|MIR4734 | 4.6277 | -4.5581 | ##### | -2.9146 | 3.5609E-03 | 7.9239E-03 |
| ENSG00000198168\|SVIP | 1334.2639 | 0.3808 | ##### | 2.9145 | 3.5629E-03 | 7.9275E-03 |
| ENSG00000090266\|NDUFB2 | 12052.3028 | 0.3250 | ##### | 2.9145 | 3.5631E-03 | 7.9275E-03 |
| ENSG00000106526\|ACTR3C | 142.6742 | -0.6078 | ##### | -2.9143 | 3.5648E-03 | 7.9309E-03 |
| ENSG00000280543\|ASAP1-IT2 | 8.0535 | 5.8223 | ##### | 2.9141 | 3.5673E-03 | 7.9357E-03 |
| ENSG00000281490\|CICP14 | 66.6256 | 1.4544 | ##### | 2.9137 | 3.5714E-03 | 7.9442E-03 |
| ENSG00000104946\|TBC1D17 | 2617.5982 | -0.3769 | ##### | -2.9137 | 3.5721E-03 | 7.9453E-03 |
| ENSG00000289396 | 11.0224 | -2.3508 | ##### | -2.9133 | 3.5767E-03 | 7.9549E-03 |
| ENSG00000227646\|STEAP2-AS1 | 7.6750 | -6.3439 | ##### | -2.9131 | 3.5785E-03 | 7.9583E-03 |
| ENSG00000258725\|PRC1-AS1 | 335.2256 | 0.5931 | ##### | 2.9130 | 3.5799E-03 | 7.9607E-03 |
| ENSG00000270587\|FAM136DP | 22.4574 | -1.1778 | ##### | -2.9127 | 3.5834E-03 | 7.9681E-03 |
| ENSG00000236090\|LDHAP3 | 92.9214 | 0.6391 | ##### | 2.9125 | 3.5855E-03 | 7.9720E-03 |
| ENSG00000113522\|RAD50 | 10463.4018 | 0.3590 | ##### | 2.9119 | 3.5925E-03 | 7.9870E-03 |
| ENSG00000188825\|LINC00910 | 150.5921 | -0.5480 | ##### | -2.9115 | 3.5965E-03 | 7.9953E-03 |
| ENSG00000162433\|AK4 | 3933.1546 | -0.3640 | ##### | -2.9100 | 3.6140E-03 | 8.0337E-03 |
| ENSG00000258056\|CD63-AS1 | 540.2279 | -0.4385 | ##### | -2.9099 | 3.6156E-03 | 8.0366E-03 |
| ENSG00000265690 | 189.4629 | -0.7252 | ##### | -2.9098 | 3.6165E-03 | 8.0380E-03 |
| ENSG00000204021\|LIPK | 5.1915 | -6.1081 | ##### | -2.9095 | 3.6196E-03 | 8.0444E-03 |
| ENSG00000267586\|LINC00907 | 27.4245 | 4.4047 | ##### | 2.9093 | 3.6226E-03 | 8.0504E-03 |
| ENSG00000255893 | 17.5184 | -1.7075 | ##### | -2.9088 | 3.6281E-03 | 8.0621E-03 |
| ENSG00000120341\|SEC16B | 13.6803 | -4.7767 | ##### | -2.9087 | 3.6294E-03 | 8.0644E-03 |
| ENSG00000213024\|NUP62 | 13792.9396 | -0.3392 | ##### | -2.9082 | 3.6347E-03 | 8.0754E-03 |
| ENSG00000215068\|ANXA2R-AS1 | 221.9525 | -0.5053 | ##### | -2.9077 | 3.6406E-03 | 8.0879E-03 |
| ENSG00000277511 | 35.4826 | -1.0821 | ##### | -2.9072 | 3.6463E-03 | 8.1001E-03 |
| ENSG00000267383 | 17.8155 | -1.2669 | ##### | -2.9064 | 3.6556E-03 | 8.1201E-03 |
| ENSG00000183475\|ASB7 | 1788.4353 | 0.3689 | ##### | 2.9062 | 3.6583E-03 | 8.1256E-03 |
| ENSG00000210174\|MT-TR | 79.7044 | 0.7148 | ##### | 2.9061 | 3.6595E-03 | 8.1276E-03 |
| ENSG00000125877\|ITPA | 3967.8181 | -0.3603 | ##### | -2.9041 | 3.6832E-03 | 8.1795E-03 |
| ENSG00000290074 | 11.0960 | 3.2734 | ##### | 2.9039 | 3.6857E-03 | 8.1845E-03 |
| ENSG00000085840\|ORC1 | 4017.1002 | 0.2850 | ##### | 2.9035 | 3.6900E-03 | 8.1935E-03 |
| ENSG00000165322\|ARHGAP12 | 2918.9219 | -0.3663 | ##### | -2.9034 | 3.6911E-03 | 8.1954E-03 |
| ENSG00000055332\|EIF2AK2 | 6174.6611 | -0.3523 | ##### | -2.9032 | 3.6934E-03 | 8.1999E-03 |
| ENSG00000122644\|ARL4A | 2600.9515 | -0.3520 | ##### | -2.9029 | 3.6968E-03 | 8.2066E-03 |
| ENSG00000113194\|FAF2 | 9489.6065 | 0.2082 | ##### | 2.9027 | 3.6992E-03 | 8.2115E-03 |
| ENSG00000273136\|NBPF26 | 1828.2285 | -0.3843 | ##### | -2.9027 | 3.6995E-03 | 8.2115E-03 |
| ENSG00000092531\|SNAP23 | 5831.7111 | -0.3617 | ##### | -2.9023 | 3.7043E-03 | 8.2216E-03 |
| ENSG00000125449\|ARMC7 | 1319.7761 | -0.3707 | ##### | -2.9012 | 3.7178E-03 | 8.2509E-03 |
| ENSG00000167716\|WDR81 | 3398.9364 | 0.3545 | ##### | 2.9006 | 3.7250E-03 | 8.2662E-03 |
| ENSG00000289537 | 131.4715 | 0.6854 | ##### | 2.9000 | 3.7314E-03 | 8.2798E-03 |
| ENSG00000116473\|RAP1A | 5267.2087 | 0.2822 | ##### | 2.8999 | 3.7329E-03 | 8.2825E-03 |
| ENSG00000274929 | 28.1411 | -7.6280 | ##### | -2.8998 | 3.7340E-03 | 8.2844E-03 |
| ENSG00000235670\|RPL21P40 | 9.3977 | -6.4453 | ##### | -2.8997 | 3.7357E-03 | 8.2876E-03 |
| ENSG00000272911 | 34.2900 | -7.8474 | ##### | -2.8990 | 3.7441E-03 | 8.3056E-03 |
| ENSG00000215835 | 105.7624 | -0.7442 | ##### | -2.8987 | 3.7467E-03 | 8.3103E-03 |
| ENSG00000104129\|DNAJC17 | 1251.2225 | 0.3926 | ##### | 2.8987 | 3.7468E-03 | 8.3103E-03 |
| ENSG00000137103\|TMEM8B | 507.3309 | 0.3626 | ##### | 2.8982 | 3.7529E-03 | 8.3232E-03 |
| ENSG00000091542\|ALKBH5 | 7508.3355 | 0.2793 | ##### | 2.8978 | 3.7584E-03 | 8.3348E-03 |
| ENSG00000291213\|DPY19L2P3 | 132.5592 | -0.7713 | ##### | -2.8973 | 3.7638E-03 | 8.3463E-03 |
| ENSG00000186919\|ZACN | 2986.1126 | 0.5243 | ##### | 2.8972 | 3.7650E-03 | 8.3483E-03 |
| ENSG00000167984\|NLRC3 | 47.5073 | -0.7717 | ##### | -2.8967 | 3.7709E-03 | 8.3605E-03 |
| ENSG00000114573\|ATP6V1A | 4736.2681 | -0.5628 | ##### | -2.8967 | 3.7713E-03 | 8.3610E-03 |
| ENSG00000108590\|MED31 | 805.2319 | 0.3975 | ##### | 2.8959 | 3.7810E-03 | 8.3818E-03 |
| ENSG00000278924 | 32.0848 | -7.0664 | ##### | -2.8943 | 3.8005E-03 | 8.4243E-03 |
| ENSG00000100104\|SRRD | 2828.7623 | 0.3683 | ##### | 2.8932 | 3.8138E-03 | 8.4533E-03 |
| ENSG00000277453 | 11.6375 | 2.1897 | ##### | 2.8930 | 3.8159E-03 | 8.4572E-03 |
| ENSG00000247228 | 86.8841 | -3.6988 | ##### | -2.8927 | 3.8192E-03 | 8.4640E-03 |
| ENSG00000095564\|BTAF1 | 4513.8746 | 0.3534 | ##### | 2.8923 | 3.8246E-03 | 8.4753E-03 |
| ENSG00000140025\|EFCAB11 | 634.7392 | 0.4151 | ##### | 2.8914 | 3.8356E-03 | 8.4991E-03 |
| ENSG00000215572\|ESRRAP1 | 6.4342 | 4.9844 | ##### | 2.8892 | 3.8620E-03 | 8.5569E-03 |
| ENSG00000206077\|ZDHHC11B | 85.5254 | 0.7191 | ##### | 2.8891 | 3.8629E-03 | 8.5583E-03 |
| ENSG00000273014 | 117.1173 | -0.7903 | ##### | -2.8889 | 3.8655E-03 | 8.5634E-03 |

| | | | | | | |
|---|---|---|---|---|---|---|
| ENSG00000230943\|LINC02541 | 59.6340 | -1.1232 | ##### | -2.8888 | 3.8670E-03 | 8.5661E-03 |
| ENSG00000284954 | 68.6450 | 0.7410 | ##### | 2.8882 | 3.8746E-03 | 8.5822E-03 |
| ENSG00000204361\|NXPE2 | 13.0213 | 3.8124 | ##### | 2.8879 | 3.8785E-03 | 8.5902E-03 |
| ENSG00000134873\|CLDN10 | 169.0544 | -0.5937 | ##### | -2.8871 | 3.8881E-03 | 8.6110E-03 |
| ENSG00000120860\|WASHC3 | 916.5670 | -0.3920 | ##### | -2.8867 | 3.8927E-03 | 8.6204E-03 |
| ENSG00000261787\|TCF24 | 34.3714 | -1.0680 | ##### | -2.8866 | 3.8942E-03 | 8.6232E-03 |
| ENSG00000067225\|PKM | ########## | 0.1899 | ##### | 2.8863 | 3.8975E-03 | 8.6299E-03 |
| ENSG00000136950\|ARPC5L | 5147.5835 | -0.3551 | ##### | -2.8862 | 3.8988E-03 | 8.6320E-03 |
| ENSG00000145198\|VWA5B2 | 66.1437 | -0.7371 | ##### | -2.8857 | 3.9060E-03 | 8.6473E-03 |
| ENSG00000135378\|PRRG4 | 258.8273 | -0.5670 | ##### | -2.8850 | 3.9146E-03 | 8.6651E-03 |
| ENSG00000245849\|RAD51-AS1 | 158.6658 | 0.6756 | ##### | 2.8850 | 3.9145E-03 | 8.6651E-03 |
| ENSG00000119655\|NPC2 | 2395.5713 | 0.3676 | ##### | 2.8848 | 3.9165E-03 | 8.6686E-03 |
| ENSG00000167552\|TUBA1A | 24240.9349 | 0.3566 | ##### | 2.8847 | 3.9175E-03 | 8.6702E-03 |
| ENSG00000140443\|IGF1R | 13073.1635 | 0.3374 | ##### | 2.8837 | 3.9298E-03 | 8.6969E-03 |
| ENSG00000007376\|RPUSD1 | 3766.3084 | 0.3606 | ##### | 2.8836 | 3.9311E-03 | 8.6991E-03 |
| ENSG00000279168 | 24.1791 | 1.2439 | ##### | 2.8835 | 3.9329E-03 | 8.7024E-03 |
| ENSG00000254333\|NDST1-AS1 | 7.9751 | -2.0247 | ##### | -2.8833 | 3.9354E-03 | 8.7073E-03 |
| ENSG00000104957\|YJU2B | 1115.7239 | 0.3989 | ##### | 2.8832 | 3.9370E-03 | 8.7102E-03 |
| ENSG00000288804 | 42.2503 | 8.0743 | ##### | 2.8828 | 3.9415E-03 | 8.7188E-03 |
| ENSG00000124831\|LRRFIP1 | 15429.8663 | 0.3400 | ##### | 2.8828 | 3.9413E-03 | 8.7188E-03 |
| ENSG00000230202\|RPL29P4 | 796.0167 | 0.4267 | ##### | 2.8826 | 3.9436E-03 | 8.7228E-03 |
| ENSG00000174460\|ZCCHC12 | 23.0135 | 3.3289 | ##### | 2.8826 | 3.9439E-03 | 8.7228E-03 |
| ENSG00000104221\|BRF2 | 1683.0055 | 0.3694 | ##### | 2.8823 | 3.9474E-03 | 8.7298E-03 |
| ENSG00000124875\|CXCL6 | 68.0324 | 0.8346 | ##### | 2.8816 | 3.9560E-03 | 8.7483E-03 |
| ENSG00000240270\|RPL12P37 | 8.8300 | 3.0672 | ##### | 2.8816 | 3.9564E-03 | 8.7484E-03 |
| ENSG00000100902\|PSMA6 | 13171.3356 | 0.4106 | ##### | 2.8808 | 3.9660E-03 | 8.7692E-03 |
| ENSG00000238061 | 7.9304 | 6.2504 | ##### | 2.8804 | 3.9721E-03 | 8.7820E-03 |
| ENSG00000237748\|UQCRBP1 | 9.6690 | -2.0173 | ##### | -2.8802 | 3.9748E-03 | 8.7873E-03 |
| ENSG00000082516\|GEMIN5 | 8232.2511 | 0.3428 | ##### | 2.8799 | 3.9777E-03 | 8.7930E-03 |
| ENSG00000249899\|ITGA2-AS1 | 5.2034 | 5.2981 | ##### | 2.8796 | 3.9823E-03 | 8.8026E-03 |
| ENSG00000049768\|FOXP3 | 32.1234 | 1.1676 | ##### | 2.8793 | 3.9854E-03 | 8.8087E-03 |
| ENSG00000114648\|KLHL18 | 2820.4401 | 0.3540 | ##### | 2.8792 | 3.9873E-03 | 8.8121E-03 |
| ENSG00000198917\|SPOUT1 | 5160.1898 | 0.3551 | ##### | 2.8791 | 3.9875E-03 | 8.8121E-03 |
| ENSG00000181035\|SLC25A42 | 242.9204 | -0.4821 | ##### | -2.8789 | 3.9907E-03 | 8.8184E-03 |
| ENSG00000257727\|CNPY2 | 12030.7413 | 0.3558 | ##### | 2.8786 | 3.9941E-03 | 8.8254E-03 |
| ENSG00000170312\|CDK1 | 5373.4359 | 0.3436 | ##### | 2.8776 | 4.0066E-03 | 8.8523E-03 |
| ENSG00000256087\|ZNF432 | 650.1877 | -0.4107 | ##### | -2.8775 | 4.0084E-03 | 8.8556E-03 |
| ENSG00000148840\|PPRC1 | 9703.3592 | 0.2875 | ##### | 2.8773 | 4.0106E-03 | 8.8600E-03 |
| ENSG00000264058 | 14.9950 | 1.7482 | ##### | 2.8770 | 4.0143E-03 | 8.8674E-03 |
| ENSG00000286435 | 46.3418 | 1.0218 | ##### | 2.8767 | 4.0184E-03 | 8.8759E-03 |
| ENSG00000058335\|RASGRF1 | 9.4096 | 3.8256 | ##### | 2.8762 | 4.0245E-03 | 8.8887E-03 |
| ENSG00000290425\|PI4KAP1 | 615.0904 | -0.3918 | ##### | -2.8762 | 4.0254E-03 | 8.8898E-03 |
| ENSG00000090534\|THPO | 9.6084 | 2.9159 | ##### | 2.8760 | 4.0273E-03 | 8.8935E-03 |
| ENSG00000168314\|MOBP | 9.5301 | 6.1762 | ##### | 2.8757 | 4.0311E-03 | 8.9005E-03 |
| ENSG00000109736\|MFSD10 | 5332.4574 | 0.3458 | ##### | 2.8757 | 4.0308E-03 | 8.9005E-03 |
| ENSG00000257545 | 10.7685 | 2.4322 | ##### | 2.8750 | 4.0409E-03 | 8.9214E-03 |
| ENSG00000104827\|CGB3 | 34.0026 | -0.9829 | ##### | -2.8740 | 4.0532E-03 | 8.9481E-03 |
| ENSG00000253710\|ALG11 | 3822.7656 | 0.3638 | ##### | 2.8736 | 4.0577E-03 | 8.9573E-03 |
| ENSG00000137726\|FXYD6 | 44.0021 | 1.7909 | ##### | 2.8730 | 4.0653E-03 | 8.9734E-03 |
| ENSG00000253477 | 19.0589 | -1.9841 | ##### | -2.8729 | 4.0667E-03 | 8.9758E-03 |
| ENSG00000151704\|KCNJ1 | 10.3762 | 5.3079 | ##### | 2.8726 | 4.0711E-03 | 8.9844E-03 |
| ENSG00000025156\|HSF2 | 1081.1756 | 0.3972 | ##### | 2.8726 | 4.0712E-03 | 8.9844E-03 |
| ENSG00000251513\|LIX1-AS1 | 12.5326 | 5.4699 | ##### | 2.8724 | 4.0740E-03 | 8.9901E-03 |
| ENSG00000111237\|VPS29 | 6025.2595 | -0.3574 | ##### | -2.8721 | 4.0775E-03 | 8.9969E-03 |
| ENSG00000127578\|WFIKKN1 | 46.8265 | -1.3572 | ##### | -2.8706 | 4.0963E-03 | 9.0378E-03 |
| ENSG00000289067 | 14.1845 | 1.5250 | ##### | 2.8705 | 4.0979E-03 | 9.0406E-03 |
| ENSG00000275401 | 20.1067 | 1.4749 | ##### | 2.8703 | 4.1009E-03 | 9.0466E-03 |
| ENSG00000117640\|MTFR1L | 3830.4934 | -0.3578 | ##### | -2.8697 | 4.1082E-03 | 9.0615E-03 |
| ENSG00000224629\|RPL36AP30 | 6.4138 | 4.8522 | ##### | 2.8697 | 4.1083E-03 | 9.0615E-03 |
| ENSG00000168124\|OR1F1 | 94.1235 | -0.6984 | ##### | -2.8697 | 4.1091E-03 | 9.0621E-03 |
| ENSG00000275888 | 82.8081 | 0.7414 | ##### | 2.8697 | 4.1091E-03 | 9.0621E-03 |
| ENSG00000249306\|LINC01411 | 5.4217 | 6.1917 | ##### | 2.8694 | 4.1119E-03 | 9.0675E-03 |
| ENSG00000234520\|HRAT17 | 11.0472 | 5.0786 | ##### | 2.8693 | 4.1135E-03 | 9.0705E-03 |

| | | | | | | |
|---|---|---|---|---|---|---|
| ENSG00000293022 | 86.5596 | -0.7133 | ##### | -2.8689 | 4.1187E-03 | 9.0809E-03 |
| ENSG00000114904\|NEK4 | 2339.7851 | 0.3630 | ##### | 2.8689 | 4.1188E-03 | 9.0809E-03 |
| ENSG00000182836\|PLCXD3 | 51.0199 | 0.9240 | ##### | 2.8683 | 4.1269E-03 | 9.0980E-03 |
| ENSG00000219435\|CATSPERZ | 24.0391 | -1.9404 | ##### | -2.8681 | 4.1294E-03 | 9.1028E-03 |
| ENSG00000163161\|ERCC3 | 7054.5839 | -0.3432 | ##### | -2.8672 | 4.1410E-03 | 9.1278E-03 |
| ENSG00000149532\|CPSF7 | 7348.5573 | 0.2977 | ##### | 2.8671 | 4.1428E-03 | 9.1311E-03 |
| ENSG00000260350 | 177.8482 | -0.6192 | ##### | -2.8670 | 4.1437E-03 | 9.1322E-03 |
| ENSG00000148357\|HMCN2 | 10.2449 | 2.8151 | ##### | 2.8665 | 4.1510E-03 | 9.1477E-03 |
| ENSG00000253831\|ETV3L | 63.7099 | 3.9713 | ##### | 2.8661 | 4.1553E-03 | 9.1565E-03 |
| ENSG00000171094\|ALK | 55.7446 | 3.5964 | ##### | 2.8648 | 4.1733E-03 | 9.1954E-03 |
| ENSG00000238084 | 19.8044 | 1.2941 | ##### | 2.8647 | 4.1739E-03 | 9.1961E-03 |
| ENSG00000167034\|NKX3-1 | 736.7284 | 0.3800 | ##### | 2.8645 | 4.1768E-03 | 9.2018E-03 |
| ENSG00000111341\|MGP | 23.5665 | -2.5968 | ##### | -2.8642 | 4.1803E-03 | 9.2086E-03 |
| ENSG00000134107\|BHLHE40 | 9077.5188 | 0.3347 | ##### | 2.8642 | 4.1805E-03 | 9.2086E-03 |
| ENSG00000213453\|FTH1P3 | 13.9236 | -1.3824 | ##### | -2.8641 | 4.1821E-03 | 9.2115E-03 |
| ENSG00000109881\|CCDC34 | 1789.5563 | -0.3723 | ##### | -2.8638 | 4.1857E-03 | 9.2188E-03 |
| ENSG00000248746\|ACTN3 | 5.5559 | -5.0733 | ##### | -2.8638 | 4.1864E-03 | 9.2195E-03 |
| ENSG00000202461\|Y_RNA | 6.5656 | -5.3583 | ##### | -2.8632 | 4.1932E-03 | 9.2340E-03 |
| ENSG00000101003\|GINS1 | 2650.2890 | 0.3597 | ##### | 2.8631 | 4.1958E-03 | 9.2389E-03 |
| ENSG00000255355\|PATL1-DT | 6.3054 | -5.6309 | ##### | -2.8629 | 4.1980E-03 | 9.2431E-03 |
| ENSG00000222764\|RN7SKP106 | 7.0652 | -6.2282 | ##### | -2.8627 | 4.1998E-03 | 9.2465E-03 |
| ENSG00000137185\|ZSCAN9 | 411.7750 | 0.4929 | ##### | 2.8624 | 4.2042E-03 | 9.2554E-03 |
| ENSG00000198937\|CCDC167 | 1158.6438 | -0.3852 | ##### | -2.8624 | 4.2046E-03 | 9.2557E-03 |
| ENSG00000246922\|UBAP1L | 90.0890 | 0.6808 | ##### | 2.8622 | 4.2065E-03 | 9.2592E-03 |
| ENSG00000229117\|RPL41 | 64822.1016 | 0.3516 | ##### | 2.8621 | 4.2087E-03 | 9.2633E-03 |
| ENSG00000089818\|NECAP1 | 3700.4514 | 0.3664 | ##### | 2.8612 | 4.2198E-03 | 9.2869E-03 |
| ENSG00000160602\|NEK8 | 497.0005 | 0.4046 | ##### | 2.8608 | 4.2257E-03 | 9.2993E-03 |
| ENSG00000127526\|SLC35E1 | 6978.4130 | -0.3464 | ##### | -2.8606 | 4.2284E-03 | 9.3046E-03 |
| ENSG00000100426\|ZBED4 | 3610.7669 | 0.3462 | ##### | 2.8595 | 4.2430E-03 | 9.3352E-03 |
| ENSG00000250312\|ZNF718 | 594.3312 | -0.4105 | ##### | -2.8595 | 4.2430E-03 | 9.3352E-03 |
| ENSG00000122692\|SMU1 | 6052.0200 | 0.3471 | ##### | 2.8585 | 4.2568E-03 | 9.3649E-03 |
| ENSG00000259673\|IQCH-AS1 | 209.8408 | -0.6232 | ##### | -2.8584 | 4.2573E-03 | 9.3654E-03 |
| ENSG00000164591\|MYOZ3 | 21.1716 | -1.3344 | ##### | -2.8583 | 4.2590E-03 | 9.3684E-03 |
| ENSG00000102796\|DHRS12 | 180.3045 | -0.5078 | ##### | -2.8573 | 4.2726E-03 | 9.3976E-03 |
| ENSG00000233532\|LINC00460 | 27.9652 | -3.0152 | ##### | -2.8567 | 4.2801E-03 | 9.4135E-03 |
| ENSG00000134697\|GNL2 | 8425.4532 | 0.3463 | ##### | 2.8566 | 4.2821E-03 | 9.4171E-03 |
| ENSG00000274598 | 11.0738 | 4.3336 | ##### | 2.8558 | 4.2933E-03 | 9.4411E-03 |
| ENSG00000118298\|CA14 | 25.3175 | 1.1560 | ##### | 2.8556 | 4.2958E-03 | 9.4458E-03 |
| ENSG00000069399\|BCL3 | 4028.7857 | -0.3576 | ##### | -2.8552 | 4.3008E-03 | 9.4563E-03 |
| ENSG00000120685\|PROSER1 | 2743.6994 | 0.3604 | ##### | 2.8547 | 4.3079E-03 | 9.4710E-03 |
| ENSG00000100558\|PLEK2 | 1045.9479 | 0.4128 | ##### | 2.8542 | 4.3140E-03 | 9.4839E-03 |
| ENSG00000264063 | 13.1955 | 1.3607 | ##### | 2.8541 | 4.3166E-03 | 9.4888E-03 |
| ENSG00000232860\|SMG7-AS1 | 107.6465 | -0.7130 | ##### | -2.8537 | 4.3217E-03 | 9.4994E-03 |
| ENSG00000133101\|CCNA1 | 405.2468 | 0.4716 | ##### | 2.8533 | 4.3270E-03 | 9.5102E-03 |
| ENSG00000142794\|NBPF3 | 1352.0703 | 0.3809 | ##### | 2.8531 | 4.3290E-03 | 9.5138E-03 |
| ENSG00000235989\|MORC2-AS1 | 27.2845 | 1.3707 | ##### | 2.8531 | 4.3292E-03 | 9.5138E-03 |
| ENSG00000230409\|TCEA1P2 | 287.1561 | -0.5467 | ##### | -2.8529 | 4.3316E-03 | 9.5182E-03 |
| ENSG00000151116\|UEVLD | 2343.0206 | -0.3640 | ##### | -2.8523 | 4.3404E-03 | 9.5369E-03 |
| ENSG00000163935\|SFMBT1 | 1722.7298 | 0.3474 | ##### | 2.8519 | 4.3464E-03 | 9.5494E-03 |
| ENSG00000280604 | 36.0005 | 1.0118 | ##### | 2.8514 | 4.3529E-03 | 9.5629E-03 |
| ENSG00000111371\|SLC38A1 | 21577.1369 | -0.3445 | ##### | -2.8513 | 4.3542E-03 | 9.5651E-03 |
| ENSG00000136003\|ISCU | 4357.3824 | 0.3570 | ##### | 2.8513 | 4.3545E-03 | 9.5651E-03 |
| ENSG00000149182\|ARFGAP2 | 3296.8860 | 0.2372 | ##### | 2.8512 | 4.3560E-03 | 9.5676E-03 |
| ENSG00000109501\|WFS1 | 3722.1283 | 0.3670 | ##### | 2.8506 | 4.3639E-03 | 9.5844E-03 |
| ENSG00000083642\|PDS5B | 3231.3859 | 0.3510 | ##### | 2.8505 | 4.3650E-03 | 9.5860E-03 |
| ENSG00000286082 | 22.8001 | -2.1526 | ##### | -2.8502 | 4.3686E-03 | 9.5933E-03 |
| ENSG00000155252\|PI4K2A | 2821.4561 | -0.3515 | ##### | -2.8496 | 4.3774E-03 | 9.6120E-03 |
| ENSG00000134265\|NAPG | 4615.5998 | 0.3624 | ##### | 2.8496 | 4.3781E-03 | 9.6126E-03 |
| ENSG00000153975\|ZUP1 | 994.6579 | 0.3666 | ##### | 2.8495 | 4.3793E-03 | 9.6147E-03 |
| ENSG00000185304\|RGPD2 | 55.8970 | -0.6974 | ##### | -2.8490 | 4.3862E-03 | 9.6291E-03 |
| ENSG00000179165\|PXT1 | 9.9183 | 2.8246 | ##### | 2.8480 | 4.3991E-03 | 9.6567E-03 |
| ENSG00000291057\|ABCC6P1 | 29.3321 | -2.6406 | ##### | -2.8479 | 4.4002E-03 | 9.6585E-03 |
| ENSG00000100395\|L3MBTL2 | 4536.9069 | 0.3398 | ##### | 2.8468 | 4.4158E-03 | 9.6919E-03 |

| | | | | | | |
|---|---|---|---|---|---|---|
| ENSG00000153237\|CCDC148 | 77.4590 | -0.9731 | ##### | -2.8468 | 4.4165E-03 | 9.6926E-03 |
| ENSG00000151148\|UBE3B | 5386.4946 | -0.3415 | ##### | -2.8467 | 4.4178E-03 | 9.6950E-03 |
| ENSG00000182359\|KBTBD3 | 168.6442 | 0.6931 | ##### | 2.8466 | 4.4186E-03 | 9.6959E-03 |
| ENSG00000215196\|BASP1-AS1 | 344.7576 | 0.5369 | ##### | 2.8463 | 4.4233E-03 | 9.7056E-03 |
| ENSG00000232838\|PET117 | 256.2459 | -0.5651 | ##### | -2.8461 | 4.4254E-03 | 9.7094E-03 |
| ENSG00000100241\|SBF1 | 12086.2916 | 0.3434 | ##### | 2.8459 | 4.4290E-03 | 9.7167E-03 |
| ENSG00000163584\|RPL22L1 | 1274.8456 | -0.4194 | ##### | -2.8452 | 4.4387E-03 | 9.7371E-03 |
| ENSG00000100918\|REC8 | 196.7753 | 0.5092 | ##### | 2.8450 | 4.4417E-03 | 9.7430E-03 |
| ENSG00000257231\|DYNLL1P4 | 6.0131 | 5.3489 | ##### | 2.8449 | 4.4425E-03 | 9.7441E-03 |
| ENSG00000234241 | 18.9728 | 1.2660 | ##### | 2.8442 | 4.4526E-03 | 9.7655E-03 |
| ENSG00000267270\|PARD6G-AS1 | 26.4748 | -1.1842 | ##### | -2.8438 | 4.4581E-03 | 9.7768E-03 |
| ENSG00000250397 | 81.7636 | 0.6593 | ##### | 2.8436 | 4.4613E-03 | 9.7833E-03 |
| ENSG00000263786 | 40.1628 | 1.1898 | ##### | 2.8434 | 4.4635E-03 | 9.7873E-03 |
| ENSG00000136244\|IL6 | 43.6876 | 1.9072 | ##### | 2.8432 | 4.4667E-03 | 9.7935E-03 |
| ENSG00000126012\|KDM5C | 10339.7983 | 0.3290 | ##### | 2.8431 | 4.4671E-03 | 9.7936E-03 |
| ENSG00000278177 | 9.2886 | -2.4449 | ##### | -2.8429 | 4.4711E-03 | 9.8017E-03 |
| ENSG00000260361 | 17.8395 | 1.5848 | ##### | 2.8427 | 4.4735E-03 | 9.8063E-03 |
| ENSG00000276350\|MIR6858 | 7.2865 | 1.8316 | ##### | 2.8421 | 4.4816E-03 | 9.8233E-03 |
| ENSG00000257264 | 21.0861 | -1.6029 | ##### | -2.8417 | 4.4868E-03 | 9.8341E-03 |
| ENSG00000197008\|ZNF138 | 211.8018 | -0.5039 | ##### | -2.8417 | 4.4874E-03 | 9.8345E-03 |
| ENSG00000198682\|PAPSS2 | 12242.0021 | 0.3403 | ##### | 2.8415 | 4.4902E-03 | 9.8401E-03 |
| ENSG00000141293\|SKAP1 | 34.1490 | -1.0116 | ##### | -2.8414 | 4.4918E-03 | 9.8428E-03 |
| ENSG00000228238 | 5.0547 | 5.5966 | ##### | 2.8408 | 4.4994E-03 | 9.8587E-03 |
| ENSG00000179841\|AKAP5 | 252.0439 | -0.6351 | ##### | -2.8408 | 4.5004E-03 | 9.8603E-03 |
| ENSG00000271715 | 24.6601 | -7.5550 | ##### | -2.8407 | 4.5017E-03 | 9.8623E-03 |
| ENSG00000272279 | 12.3310 | -1.4529 | ##### | -2.8406 | 4.5024E-03 | 9.8631E-03 |
| ENSG00000234380\|LINC01426 | 20.3461 | 1.2823 | ##### | 2.8404 | 4.5062E-03 | 9.8708E-03 |
| ENSG00000260798 | 4.5796 | -3.2407 | ##### | -2.8403 | 4.5078E-03 | 9.8736E-03 |
| ENSG00000227107 | 10.6190 | 2.7280 | ##### | 2.8396 | 4.5175E-03 | 9.8940E-03 |
| ENSG00000260468\|LINC01290 | 8.0075 | -6.4093 | ##### | -2.8394 | 4.5199E-03 | 9.8981E-03 |
| ENSG00000253919\|THAP12P7 | 31.6632 | 1.0712 | ##### | 2.8394 | 4.5200E-03 | 9.8981E-03 |
| ENSG00000128604\|IRF5 | 378.4701 | -0.4175 | ##### | -2.8391 | 4.5245E-03 | 9.9072E-03 |
| ENSG00000251521\|IMPA1P1 | 8.0034 | -4.1383 | ##### | -2.8386 | 4.5319E-03 | 9.9226E-03 |
| ENSG00000160993\|ALKBH4 | 1119.0475 | 0.3860 | ##### | 2.8383 | 4.5357E-03 | 9.9303E-03 |
| ENSG00000155962\|CLIC2 | 27.0986 | -2.0148 | ##### | -2.8378 | 4.5423E-03 | 9.9440E-03 |
| ENSG00000183726\|TMEM50A | 6590.2676 | 0.3500 | ##### | 2.8371 | 4.5526E-03 | 9.9658E-03 |
| ENSG00000105679\|GAPDHS | 121.1893 | -0.6915 | ##### | -2.8366 | 4.5599E-03 | 9.9809E-03 |
| ENSG00000031081\|ARHGAP31 | 1422.7572 | 0.3828 | ##### | 2.8351 | 4.5811E-03 | 1.0027E-02 |
| ENSG00000277873 | 9.4368 | -1.5708 | ##### | -2.8350 | 4.5827E-03 | 1.0030E-02 |
| ENSG00000070669\|ASNS | 3935.8339 | -0.3685 | ##### | -2.8341 | 4.5956E-03 | 1.0057E-02 |
| ENSG00000260213\|CENPN-AS1 | 10.7937 | 2.4289 | ##### | 2.8338 | 4.5995E-03 | 1.0065E-02 |
| ENSG00000229152\|ANKRD10-IT1 | 75.7155 | 0.8322 | ##### | 2.8334 | 4.6056E-03 | 1.0077E-02 |
| ENSG00000248994 | 25.4934 | 7.1225 | ##### | 2.8332 | 4.6083E-03 | 1.0082E-02 |
| ENSG00000293320 | 6676.7661 | 0.3638 | ##### | 2.8331 | 4.6102E-03 | 1.0086E-02 |
| ENSG00000166428\|PLD4 | 52.1915 | -0.9489 | ##### | -2.8324 | 4.6203E-03 | 1.0107E-02 |
| ENSG00000007520\|TSR3 | 5087.1457 | -0.3482 | ##### | -2.8321 | 4.6242E-03 | 1.0115E-02 |
| ENSG00000092607\|TBX15 | 20.5861 | -2.6483 | ##### | -2.8302 | 4.6520E-03 | 1.0175E-02 |
| ENSG00000165985\|C1QL3 | 117.0866 | 0.6296 | ##### | 2.8300 | 4.6554E-03 | 1.0182E-02 |
| ENSG00000232527\|LINC02802 | 198.6732 | 0.6226 | ##### | 2.8297 | 4.6595E-03 | 1.0190E-02 |
| ENSG00000197142\|ACSL5 | 17.0136 | -2.9153 | ##### | -2.8293 | 4.6652E-03 | 1.0202E-02 |
| ENSG00000289457 | 6.0783 | 3.2011 | ##### | 2.8290 | 4.6689E-03 | 1.0209E-02 |
| ENSG00000149179\|CSTPP1 | 1346.3667 | -0.3817 | ##### | -2.8290 | 4.6695E-03 | 1.0210E-02 |
| ENSG00000196588\|MRTFA | 4734.7574 | -0.3323 | ##### | -2.8286 | 4.6751E-03 | 1.0221E-02 |
| ENSG00000197363\|ZNF517 | 712.5215 | 0.3920 | ##### | 2.8282 | 4.6816E-03 | 1.0235E-02 |
| ENSG00000283148 | 23.2653 | -7.4698 | ##### | -2.8279 | 4.6850E-03 | 1.0241E-02 |
| ENSG00000287944 | 24.8170 | -7.5642 | ##### | -2.8276 | 4.6895E-03 | 1.0251E-02 |
| ENSG00000287586 | 7.1450 | 4.8549 | ##### | 2.8269 | 4.6996E-03 | 1.0272E-02 |
| ENSG00000233980\|FDPSP2 | 7.6871 | -1.9558 | ##### | -2.8268 | 4.7014E-03 | 1.0275E-02 |
| ENSG00000172057\|ORMDL3 | 2938.5325 | 0.3407 | ##### | 2.8267 | 4.7028E-03 | 1.0277E-02 |
| ENSG00000254791\|FAR1-IT1 | 7.2333 | 5.4607 | ##### | 2.8266 | 4.7051E-03 | 1.0281E-02 |
| ENSG00000118816\|CCNI | 12603.1463 | 0.3488 | ##### | 2.8264 | 4.7069E-03 | 1.0285E-02 |
| ENSG00000129455\|KLK8 | 18.6042 | -1.8371 | ##### | -2.8252 | 4.7246E-03 | 1.0322E-02 |
| ENSG00000289109 | 54.8009 | 0.9683 | ##### | 2.8252 | 4.7248E-03 | 1.0322E-02 |

| | | | | | | |
|---|---|---|---|---|---|---|
| ENSG00000131848\|ZSCAN5A | 813.3281 | 0.4079 | ##### | 2.8251 | 4.7261E-03 | 1.0324E-02 |
| ENSG00000228857\|ACTR3-AS1 | 22.8902 | -2.7388 | ##### | -2.8251 | 4.7260E-03 | 1.0324E-02 |
| ENSG00000242866\|STRC | 16.5043 | -1.3048 | ##### | -2.8248 | 4.7312E-03 | 1.0334E-02 |
| ENSG00000105726\|ATP13A1 | 8112.9067 | 0.3210 | ##### | 2.8247 | 4.7321E-03 | 1.0335E-02 |
| ENSG00000235776\|RPL7AP70 | 111.1240 | -0.5833 | ##### | -2.8245 | 4.7348E-03 | 1.0340E-02 |
| ENSG00000258661 | 38.0266 | 2.7875 | ##### | 2.8240 | 4.7422E-03 | 1.0355E-02 |
| ENSG00000101665\|SMAD7 | 2163.5459 | -0.3642 | ##### | -2.8240 | 4.7423E-03 | 1.0355E-02 |
| ENSG00000181449\|SOX2 | 31.6291 | 2.4856 | ##### | 2.8240 | 4.7429E-03 | 1.0356E-02 |
| ENSG00000226981\|ABHD17AP6 | 20.4680 | 1.0212 | ##### | 2.8232 | 4.7542E-03 | 1.0380E-02 |
| ENSG00000165699\|TSC1 | 3213.9293 | -0.2979 | ##### | -2.8230 | 4.7577E-03 | 1.0387E-02 |
| ENSG00000236762\|RPL19P16 | 42.5236 | 7.9176 | ##### | 2.8227 | 4.7619E-03 | 1.0395E-02 |
| ENSG00000279149 | 4.4082 | 3.4367 | ##### | 2.8223 | 4.7674E-03 | 1.0406E-02 |
| ENSG00000291046 | 302.4996 | 0.4484 | ##### | 2.8219 | 4.7736E-03 | 1.0419E-02 |
| ENSG00000258813\|RBM25-AS1 | 22.3495 | 1.3591 | ##### | 2.8212 | 4.7852E-03 | 1.0444E-02 |
| ENSG00000165516\|KLHDC2 | 2273.6405 | -0.4097 | ##### | -2.8208 | 4.7901E-03 | 1.0454E-02 |
| ENSG00000258260 | 15.8824 | 1.4030 | ##### | 2.8198 | 4.8048E-03 | 1.0485E-02 |
| ENSG00000169085\|VXN | 234.7832 | 6.6316 | ##### | 2.8192 | 4.8144E-03 | 1.0505E-02 |
| ENSG00000237624\|OXCT2P1 | 90.5261 | 0.6079 | ##### | 2.8191 | 4.8164E-03 | 1.0509E-02 |
| ENSG00000134900\|TPP2 | 2919.9245 | -0.3418 | ##### | -2.8188 | 4.8208E-03 | 1.0518E-02 |
| ENSG00000184164\|CRELD2 | 2215.2098 | -0.3421 | ##### | -2.8187 | 4.8213E-03 | 1.0518E-02 |
| ENSG00000117148\|ACTL8 | 31.9157 | -1.0935 | ##### | -2.8185 | 4.8250E-03 | 1.0525E-02 |
| ENSG00000183423\|LRIT3 | 13.5936 | 3.8133 | ##### | 2.8178 | 4.8354E-03 | 1.0547E-02 |
| ENSG00000258303 | 9.6392 | -3.9380 | ##### | -2.8178 | 4.8360E-03 | 1.0548E-02 |
| ENSG00000162244\|RPL29 | 31693.4857 | 0.3438 | ##### | 2.8176 | 4.8387E-03 | 1.0553E-02 |
| ENSG00000164674\|SYTL3 | 116.8507 | -0.5999 | ##### | -2.8175 | 4.8398E-03 | 1.0554E-02 |
| ENSG00000198846\|TOX | 29.1004 | 2.6278 | ##### | 2.8173 | 4.8429E-03 | 1.0560E-02 |
| ENSG00000214975\|PPIAP29 | 262.2593 | -0.4805 | ##### | -2.8171 | 4.8452E-03 | 1.0565E-02 |
| ENSG00000289321 | 7.8191 | 4.3484 | ##### | 2.8168 | 4.8498E-03 | 1.0574E-02 |
| ENSG00000225131\|PSME2P2 | 15.5519 | 1.4411 | ##### | 2.8167 | 4.8523E-03 | 1.0578E-02 |
| ENSG00000289483 | 20.8372 | -1.2260 | ##### | -2.8164 | 4.8564E-03 | 1.0587E-02 |
| ENSG00000288602\|C8orf44-SGK3 | 17.7633 | -3.6685 | ##### | -2.8162 | 4.8597E-03 | 1.0593E-02 |
| ENSG00000271079\|CTAGE15 | 39.4558 | -0.9737 | ##### | -2.8157 | 4.8667E-03 | 1.0607E-02 |
| ENSG00000183801\|OLFML1 | 19.9914 | 7.2946 | ##### | 2.8157 | 4.8675E-03 | 1.0608E-02 |
| ENSG00000276390 | 37.8116 | 1.2925 | ##### | 2.8152 | 4.8741E-03 | 1.0622E-02 |
| ENSG00000226121\|AHCTF1P1 | 1011.1649 | 0.3851 | ##### | 2.8149 | 4.8786E-03 | 1.0631E-02 |
| ENSG00000235034\|C19orf81 | 252.9752 | -0.4349 | ##### | -2.8148 | 4.8807E-03 | 1.0635E-02 |
| ENSG00000201823\|SNORD48 | 13.7548 | 1.2858 | ##### | 2.8140 | 4.8923E-03 | 1.0660E-02 |
| ENSG00000137409\|MTCH1 | 16997.8394 | 0.3436 | ##### | 2.8137 | 4.8981E-03 | 1.0671E-02 |
| ENSG00000100490\|CDKL1 | 1272.2463 | -0.4012 | ##### | -2.8127 | 4.9123E-03 | 1.0702E-02 |
| ENSG00000225533\|PAWRP1 | 6.9123 | -5.7004 | ##### | -2.8125 | 4.9157E-03 | 1.0708E-02 |
| ENSG00000240975 | 6.5291 | -3.4837 | ##### | -2.8116 | 4.9292E-03 | 1.0737E-02 |
| ENSG00000101911\|PRPS2 | 3343.0308 | -0.3313 | ##### | -2.8116 | 4.9295E-03 | 1.0737E-02 |
| ENSG00000198176\|TFDP1 | 11771.7206 | 0.3454 | ##### | 2.8113 | 4.9347E-03 | 1.0747E-02 |
| ENSG00000254964 | 6.0338 | 4.5524 | ##### | 2.8112 | 4.9352E-03 | 1.0747E-02 |
| ENSG00000257279 | 9.8029 | -2.1428 | ##### | -2.8109 | 4.9406E-03 | 1.0758E-02 |
| ENSG00000289488 | 9.9278 | -5.2144 | ##### | -2.8107 | 4.9426E-03 | 1.0762E-02 |
| ENSG00000186994\|KANK3 | 80.8199 | 0.7084 | ##### | 2.8103 | 4.9491E-03 | 1.0775E-02 |
| ENSG00000224063\|CALCRL-AS1 | 142.6051 | 0.5279 | ##### | 2.8102 | 4.9516E-03 | 1.0780E-02 |
| ENSG00000169599\|NFU1 | 2780.4029 | -0.3508 | ##### | -2.8100 | 4.9546E-03 | 1.0786E-02 |
| ENSG00000232110 | 7.5123 | 6.1658 | ##### | 2.8095 | 4.9625E-03 | 1.0802E-02 |
| ENSG00000108055\|SMC3 | 10209.6857 | 0.3348 | ##### | 2.8091 | 4.9676E-03 | 1.0812E-02 |
| ENSG00000111727\|HCFC2 | 703.3473 | 0.3809 | ##### | 2.8089 | 4.9717E-03 | 1.0821E-02 |
| ENSG00000228495\|LINC01013 | 115.7180 | 0.6159 | ##### | 2.8087 | 4.9748E-03 | 1.0827E-02 |
| ENSG00000234396 | 8.3093 | -5.0053 | ##### | -2.8081 | 4.9831E-03 | 1.0844E-02 |
| ENSG00000252690 | 178.0176 | 0.6232 | ##### | 2.8079 | 4.9863E-03 | 1.0850E-02 |
| ENSG00000171109\|MFN1 | 5512.8744 | -0.3407 | ##### | -2.8076 | 4.9920E-03 | 1.0861E-02 |
| ENSG00000235944\|ZNF815P | 46.9023 | -2.7759 | ##### | -2.8069 | 5.0023E-03 | 1.0883E-02 |
| ENSG00000236397\|DDX11L2 | 87.6481 | -0.6787 | ##### | -2.8065 | 5.0081E-03 | 1.0895E-02 |
| ENSG00000257142 | 4.8868 | 2.9527 | ##### | 2.8063 | 5.0110E-03 | 1.0901E-02 |
| ENSG00000175509\|ACRP1 | 7.8950 | -6.3888 | ##### | -2.8059 | 5.0174E-03 | 1.0914E-02 |
| ENSG00000271751 | 7.0595 | -3.1169 | ##### | -2.8047 | 5.0368E-03 | 1.0955E-02 |
| ENSG00000237765\|FAM200B | 1342.9517 | 0.3693 | ##### | 2.8046 | 5.0386E-03 | 1.0958E-02 |
| ENSG00000177697\|CD151 | 22723.4774 | -0.3402 | ##### | -2.8035 | 5.0550E-03 | 1.0993E-02 |

| | | | | | | |
|---|---|---|---|---|---|---|
| ENSG00000004864\|SLC25A13 | 5751.5872 | 0.3519 | ##### | 2.8032 | 5.0593E-03 | 1.1002E-02 |
| ENSG00000187904 | 28.9170 | -7.0682 | ##### | -2.8030 | 5.0635E-03 | 1.1010E-02 |
| ENSG00000206140\|TMEM191C | 54.9527 | -0.9364 | ##### | -2.8029 | 5.0642E-03 | 1.1011E-02 |
| ENSG00000122952\|ZWINT | 6114.2471 | 0.3421 | ##### | 2.8024 | 5.0731E-03 | 1.1029E-02 |
| ENSG00000129993\|CBFA2T3 | 55.3698 | 0.7369 | ##### | 2.8023 | 5.0740E-03 | 1.1029E-02 |
| ENSG00000169100\|SLC25A6 | 12408.1449 | -0.3353 | ##### | -2.8023 | 5.0738E-03 | 1.1029E-02 |
| ENSG00000260439\|LMF1-AS1 | 20.2604 | 4.6971 | ##### | 2.8019 | 5.0798E-03 | 1.1041E-02 |
| ENSG00000189419\|SPATA41 | 8.3369 | 2.8186 | ##### | 2.8019 | 5.0805E-03 | 1.1042E-02 |
| ENSG00000273374 | 16.6730 | 2.5289 | ##### | 2.8019 | 5.0807E-03 | 1.1042E-02 |
| ENSG00000197045\|GMFB | 10449.6187 | 0.2177 | ##### | 2.8015 | 5.0860E-03 | 1.1052E-02 |
| ENSG00000262136\|NQO1-DT | 884.6428 | -0.3960 | ##### | -2.8003 | 5.1062E-03 | 1.1096E-02 |
| ENSG00000215788\|TNFRSF25 | 376.4947 | 0.4674 | ##### | 2.8002 | 5.1072E-03 | 1.1097E-02 |
| ENSG00000287642 | 39.1176 | -9.3310 | ##### | -2.8002 | 5.1074E-03 | 1.1097E-02 |
| ENSG00000231305\|IFT122P3 | 61.3288 | 0.9892 | ##### | 2.7999 | 5.1116E-03 | 1.1105E-02 |
| ENSG00000137941\|TTLL7 | 1650.1641 | -0.3771 | ##### | -2.7999 | 5.1123E-03 | 1.1106E-02 |
| ENSG00000288988 | 46.1285 | 0.7828 | ##### | 2.7991 | 5.1240E-03 | 1.1130E-02 |
| ENSG00000127603\|MACF1 | 10523.8680 | 0.3502 | ##### | 2.7988 | 5.1289E-03 | 1.1140E-02 |
| ENSG00000176953\|NFATC2IP | 6494.5793 | 0.3288 | ##### | 2.7984 | 5.1352E-03 | 1.1153E-02 |
| ENSG00000253463\|HMGB1P19 | 6.6485 | -5.6455 | ##### | -2.7984 | 5.1364E-03 | 1.1155E-02 |
| ENSG00000168928\|CTRB2 | 15.9433 | 1.7619 | ##### | 2.7974 | 5.1509E-03 | 1.1185E-02 |
| ENSG00000293528 | 7.0590 | 3.0657 | ##### | 2.7974 | 5.1514E-03 | 1.1186E-02 |
| ENSG00000132155\|RAF1 | 12115.3605 | 0.3539 | ##### | 2.7970 | 5.1574E-03 | 1.1198E-02 |
| ENSG00000235333\|PVRIG2P | 28.7830 | 2.2258 | ##### | 2.7964 | 5.1679E-03 | 1.1220E-02 |
| ENSG00000162618\|ADGRL4 | 9.3758 | -3.6594 | ##### | -2.7963 | 5.1696E-03 | 1.1223E-02 |
| ENSG00000259764\|PCSK6-AS1 | 22.0385 | -7.2234 | ##### | -2.7962 | 5.1711E-03 | 1.1225E-02 |
| ENSG00000248956\|HMGB1P44 | 5.7507 | 5.7628 | ##### | 2.7952 | 5.1863E-03 | 1.1257E-02 |
| ENSG00000172789\|HOXC5 | 40.4505 | 1.8678 | ##### | 2.7952 | 5.1872E-03 | 1.1258E-02 |
| ENSG00000078177\|N4BP2 | 2316.3280 | 0.3671 | ##### | 2.7933 | 5.2176E-03 | 1.1324E-02 |
| ENSG00000157450\|RNF111 | 3054.4271 | 0.3613 | ##### | 2.7932 | 5.2189E-03 | 1.1326E-02 |
| ENSG00000173705\|SUSD5 | 912.7639 | -0.3970 | ##### | -2.7918 | 5.2410E-03 | 1.1373E-02 |
| ENSG00000131730\|CKMT2 | 12.2748 | 1.8676 | ##### | 2.7918 | 5.2418E-03 | 1.1374E-02 |
| ENSG00000256282\|MTCH1P2 | 10.8146 | 3.7560 | ##### | 2.7910 | 5.2548E-03 | 1.1400E-02 |
| ENSG00000157637\|SLC38A10 | 8768.3878 | -0.3290 | ##### | -2.7910 | 5.2545E-03 | 1.1400E-02 |
| ENSG00000240666\|MME-AS1 | 9.9605 | 1.4953 | ##### | 2.7905 | 5.2627E-03 | 1.1417E-02 |
| ENSG00000276115 | 19.0636 | 2.0550 | ##### | 2.7905 | 5.2632E-03 | 1.1417E-02 |
| ENSG00000243477\|NAA80 | 427.0912 | 0.4184 | ##### | 2.7904 | 5.2645E-03 | 1.1419E-02 |
| ENSG00000198416\|ZNF658B | 40.2928 | -2.3839 | ##### | -2.7897 | 5.2750E-03 | 1.1441E-02 |
| ENSG00000110042\|DTX4 | 533.1428 | -0.4027 | ##### | -2.7895 | 5.2795E-03 | 1.1450E-02 |
| ENSG00000137463\|MGARP | 15.6678 | -1.6137 | ##### | -2.7892 | 5.2842E-03 | 1.1459E-02 |
| ENSG00000275700\|AATF | 8116.9483 | 0.3419 | ##### | 2.7886 | 5.2933E-03 | 1.1478E-02 |
| ENSG00000282961\|PRNCR1 | 37.9953 | 0.7622 | ##### | 2.7859 | 5.3387E-03 | 1.1575E-02 |
| ENSG00000144741\|SLC25A26 | 1465.1859 | 0.3772 | ##### | 2.7856 | 5.3437E-03 | 1.1585E-02 |
| ENSG00000135482\|ZC3H10 | 795.6699 | 0.6124 | ##### | 2.7854 | 5.3455E-03 | 1.1589E-02 |
| ENSG00000150540\|HNMT | 48.3020 | -4.0610 | ##### | -2.7852 | 5.3495E-03 | 1.1596E-02 |
| ENSG00000120942\|UBIAD1 | 3169.1102 | 0.3472 | ##### | 2.7849 | 5.3544E-03 | 1.1606E-02 |
| ENSG00000120690\|ELF1 | 2824.0697 | -0.2438 | ##### | -2.7848 | 5.3563E-03 | 1.1609E-02 |
| ENSG00000148671\|ADIRF | 8.7911 | -2.9556 | ##### | -2.7846 | 5.3596E-03 | 1.1616E-02 |
| ENSG00000260868\|LINC01960 | 10.5482 | -4.8715 | ##### | -2.7840 | 5.3695E-03 | 1.1636E-02 |
| ENSG00000271714\|FAM169A-AS1 | 17.5430 | -1.8186 | ##### | -2.7839 | 5.3714E-03 | 1.1640E-02 |
| ENSG00000247473\|CARS1-AS1 | 8.2874 | -5.0346 | ##### | -2.7835 | 5.3768E-03 | 1.1650E-02 |
| ENSG00000115207\|GTF3C2 | 13332.2763 | 0.2418 | ##### | 2.7834 | 5.3787E-03 | 1.1654E-02 |
| ENSG00000230304\|CICP6 | 6.0902 | 1.8888 | ##### | 2.7833 | 5.3803E-03 | 1.1656E-02 |
| ENSG00000120327\|PCDHB14 | 48.0634 | -1.0023 | ##### | -2.7832 | 5.3834E-03 | 1.1662E-02 |
| ENSG00000248873\|SERBP1P6 | 17.6972 | -1.1605 | ##### | -2.7825 | 5.3944E-03 | 1.1685E-02 |
| ENSG00000118260\|CREB1 | 3325.4909 | 0.3524 | ##### | 2.7822 | 5.3994E-03 | 1.1695E-02 |
| ENSG00000111725\|PRKAB1 | 2628.7948 | -0.3471 | ##### | -2.7821 | 5.4014E-03 | 1.1699E-02 |
| ENSG00000235363\|SNRPGP10 | 40.9058 | 0.9018 | ##### | 2.7820 | 5.4023E-03 | 1.1700E-02 |
| ENSG00000290602 | 26.8110 | 6.8685 | ##### | 2.7819 | 5.4040E-03 | 1.1703E-02 |
| ENSG00000225177\|NHSL1-AS1 | 113.9123 | 0.5784 | ##### | 2.7817 | 5.4076E-03 | 1.1709E-02 |
| ENSG00000224785 | 4.0356 | -3.5924 | ##### | -2.7815 | 5.4116E-03 | 1.1717E-02 |
| ENSG00000260577\|CDH3-AS1 | 20.6983 | -1.3265 | ##### | -2.7814 | 5.4129E-03 | 1.1719E-02 |
| ENSG00000234193\|ACSL3-AS1 | 7.0765 | -4.1366 | ##### | -2.7813 | 5.4135E-03 | 1.1720E-02 |
| ENSG00000125457\|MIF4GD | 1457.7583 | -0.3521 | ##### | -2.7811 | 5.4170E-03 | 1.1726E-02 |

| | | | | | | |
|---|---|---|---|---|---|---|
| ENSG00000121075\|TBX4 | 24.4841 | -2.6817 | ##### | -2.7811 | 5.4176E-03 | 1.1727E-02 |
| ENSG00000176049\|JAKMIP2 | 75.7527 | 4.0961 | ##### | 2.7810 | 5.4196E-03 | 1.1730E-02 |
| ENSG00000172548\|NIPAL4 | 34.5846 | -0.9893 | ##### | -2.7799 | 5.4372E-03 | 1.1768E-02 |
| ENSG00000198821\|CD247 | 26.4454 | -1.0436 | ##### | -2.7798 | 5.4392E-03 | 1.1771E-02 |
| ENSG00000121892\|PDS5A | 15208.4191 | 0.3345 | ##### | 2.7794 | 5.4460E-03 | 1.1785E-02 |
| ENSG00000230629\|RPS23P8 | 57.8290 | 0.7029 | ##### | 2.7792 | 5.4494E-03 | 1.1791E-02 |
| ENSG00000047249\|ATP6V1H | 3045.7924 | -0.3512 | ##### | -2.7789 | 5.4549E-03 | 1.1803E-02 |
| ENSG00000283122\|HYMAI | 5.0520 | 2.7007 | ##### | 2.7788 | 5.4568E-03 | 1.1806E-02 |
| ENSG00000288952 | 6.7824 | -6.1704 | ##### | -2.7782 | 5.4660E-03 | 1.1825E-02 |
| ENSG00000245848\|CEBPA | 762.3044 | 0.3940 | ##### | 2.7779 | 5.4716E-03 | 1.1836E-02 |
| ENSG00000286837 | 25.3113 | 1.2128 | ##### | 2.7774 | 5.4803E-03 | 1.1854E-02 |
| ENSG00000262823 | 19.2743 | -4.3309 | ##### | -2.7773 | 5.4815E-03 | 1.1856E-02 |
| ENSG00000075391\|RASAL2 | 3781.3281 | -0.3390 | ##### | -2.7767 | 5.4907E-03 | 1.1875E-02 |
| ENSG00000106803\|SEC61B | 5540.3370 | -0.3494 | ##### | -2.7759 | 5.5048E-03 | 1.1904E-02 |
| ENSG00000266527 | 17.1583 | -2.2983 | ##### | -2.7755 | 5.5114E-03 | 1.1918E-02 |
| ENSG00000058272\|PPP1R12A | 7734.2774 | 0.3343 | ##### | 2.7755 | 5.5123E-03 | 1.1919E-02 |
| ENSG00000131203\|IDO1 | 6.5308 | -5.6197 | ##### | -2.7749 | 5.5215E-03 | 1.1938E-02 |
| ENSG00000189367\|KIAA0408 | 235.2230 | 2.1129 | ##### | 2.7746 | 5.5266E-03 | 1.1948E-02 |
| ENSG00000270813\|NANOGNBP3 | 10.0829 | -3.9878 | ##### | -2.7742 | 5.5344E-03 | 1.1964E-02 |
| ENSG00000246982\|BRPF3-AS1 | 13.9631 | 2.0330 | ##### | 2.7739 | 5.5385E-03 | 1.1972E-02 |
| ENSG00000264070\|DND1P1 | 102.7034 | 0.5788 | ##### | 2.7734 | 5.5468E-03 | 1.1989E-02 |
| ENSG00000267231 | 7.2239 | 5.6206 | ##### | 2.7731 | 5.5517E-03 | 1.1999E-02 |
| ENSG00000278963 | 5.7489 | 3.5883 | ##### | 2.7729 | 5.5553E-03 | 1.2006E-02 |
| ENSG00000205307\|SAP25 | 18.9477 | 1.3909 | ##### | 2.7722 | 5.5676E-03 | 1.2032E-02 |
| ENSG00000139405\|RITA1 | 3504.4495 | 0.3409 | ##### | 2.7720 | 5.5719E-03 | 1.2039E-02 |
| ENSG00000155428\|TRIM74 | 38.9044 | -0.7878 | ##### | -2.7720 | 5.5717E-03 | 1.2039E-02 |
| ENSG00000265511 | 22.7785 | 1.1625 | ##### | 2.7718 | 5.5746E-03 | 1.2044E-02 |
| ENSG00000060971\|ACAA1 | 2480.1398 | 0.3442 | ##### | 2.7718 | 5.5751E-03 | 1.2044E-02 |
| ENSG00000174807\|CD248 | 21.1965 | 7.0510 | ##### | 2.7713 | 5.5827E-03 | 1.2060E-02 |
| ENSG00000260442\|ATP2A1-AS1 | 170.8101 | -0.5391 | ##### | -2.7712 | 5.5846E-03 | 1.2062E-02 |
| ENSG00000289045 | 34.8782 | 1.2519 | ##### | 2.7712 | 5.5845E-03 | 1.2062E-02 |
| ENSG00000181350\|LRRC75A | 4212.7592 | -0.3912 | ##### | -2.7707 | 5.5931E-03 | 1.2080E-02 |
| ENSG00000230847\|OCLNP1 | 18.3817 | 3.2049 | ##### | 2.7697 | 5.6101E-03 | 1.2116E-02 |
| ENSG00000075239\|ACAT1 | 4876.1501 | -0.3411 | ##### | -2.7694 | 5.6163E-03 | 1.2128E-02 |
| ENSG00000104368\|PLAT | 46.3287 | 1.2227 | ##### | 2.7693 | 5.6180E-03 | 1.2131E-02 |
| ENSG00000103832\|GOLGA8UP | 14.3242 | 2.8975 | ##### | 2.7692 | 5.6192E-03 | 1.2132E-02 |
| ENSG00000075407\|ZNF37A | 2646.1697 | 0.3333 | ##### | 2.7692 | 5.6197E-03 | 1.2133E-02 |
| ENSG00000272897 | 85.2473 | 9.2262 | ##### | 2.7690 | 5.6226E-03 | 1.2138E-02 |
| ENSG00000232310\|LINC03002 | 25.0194 | 7.4296 | ##### | 2.7683 | 5.6356E-03 | 1.2165E-02 |
| ENSG00000166441\|RPL27A | 31667.7382 | 0.3350 | ##### | 2.7676 | 5.6477E-03 | 1.2191E-02 |
| ENSG00000128266\|GNAZ | 756.9067 | 0.3747 | ##### | 2.7672 | 5.6547E-03 | 1.2204E-02 |
| ENSG00000161896\|IP6K3 | 28.4923 | -7.4478 | ##### | -2.7672 | 5.6546E-03 | 1.2204E-02 |
| ENSG00000279452 | 45.3742 | 1.0037 | ##### | 2.7667 | 5.6633E-03 | 1.2221E-02 |
| ENSG00000109171\|SLAIN2 | 5015.7649 | 0.3460 | ##### | 2.7667 | 5.6634E-03 | 1.2221E-02 |
| ENSG00000274553\|CICP15 | 8.2918 | 1.6118 | ##### | 2.7666 | 5.6653E-03 | 1.2224E-02 |
| ENSG00000233838\|DPH3P1 | 11.6017 | 1.4392 | ##### | 2.7664 | 5.6684E-03 | 1.2230E-02 |
| ENSG00000053372\|MRTO4 | 7639.7052 | 0.3370 | ##### | 2.7663 | 5.6697E-03 | 1.2232E-02 |
| ENSG00000235688\|SNTG2-AS1 | 3.6948 | 4.6489 | ##### | 2.7657 | 5.6793E-03 | 1.2252E-02 |
| ENSG00000182400\|TRAPPC6B | 2247.9052 | 0.3461 | ##### | 2.7655 | 5.6835E-03 | 1.2260E-02 |
| ENSG00000010072\|SPRTN | 1775.2226 | 0.3353 | ##### | 2.7650 | 5.6929E-03 | 1.2278E-02 |
| ENSG00000143702\|CEP170 | 8766.1811 | 0.3344 | ##### | 2.7650 | 5.6926E-03 | 1.2278E-02 |
| ENSG00000143858\|SYT2 | 19.6850 | -1.1742 | ##### | -2.7647 | 5.6981E-03 | 1.2289E-02 |
| ENSG00000176289\|IDSP1 | 19.5774 | 1.8769 | ##### | 2.7641 | 5.7074E-03 | 1.2308E-02 |
| ENSG00000273230 | 1298.2455 | 0.3623 | ##### | 2.7640 | 5.7089E-03 | 1.2310E-02 |
| ENSG00000189339\|SLC35E2B | 4726.4013 | -0.3330 | ##### | -2.7640 | 5.7101E-03 | 1.2312E-02 |
| ENSG00000225345\|SNX18P3 | 8.9482 | -1.8446 | ##### | -2.7636 | 5.7165E-03 | 1.2325E-02 |
| ENSG00000121406\|ZNF549 | 889.0667 | -0.3900 | ##### | -2.7625 | 5.7358E-03 | 1.2365E-02 |
| ENSG00000253438\|PCAT1 | 74.8808 | 1.3427 | ##### | 2.7614 | 5.7549E-03 | 1.2406E-02 |
| ENSG00000180902\|D2HGDH | 1471.7943 | -0.3458 | ##### | -2.7611 | 5.7600E-03 | 1.2416E-02 |
| ENSG00000162078\|ZG16B | 86.2758 | -0.8235 | ##### | -2.7608 | 5.7656E-03 | 1.2427E-02 |
| ENSG00000167311\|ART5 | 17.5420 | -4.0854 | ##### | -2.7608 | 5.7665E-03 | 1.2428E-02 |
| ENSG00000112200\|ZNF451 | 3882.0540 | 0.3439 | ##### | 2.7607 | 5.7684E-03 | 1.2431E-02 |
| ENSG00000188389\|PDCD1 | 9.7963 | 6.1449 | ##### | 2.7605 | 5.7712E-03 | 1.2437E-02 |

| | | | | | | |
|---|---|---|---|---|---|---|
| ENSG00000289114 | 5.5467 | 5.2307 | ##### | 2.7605 | 5.7721E-03 | 1.2438E-02 |
| ENSG00000138483\|CCDC54 | 7.6998 | 5.4822 | ##### | 2.7604 | 5.7736E-03 | 1.2440E-02 |
| ENSG00000123146\|ADGRE5 | 9053.1403 | 0.3365 | ##### | 2.7600 | 5.7798E-03 | 1.2452E-02 |
| ENSG00000186468\|RPS23 | 42037.3765 | 0.3345 | ##### | 2.7596 | 5.7872E-03 | 1.2467E-02 |
| ENSG00000232721 | 11.7223 | 3.1855 | ##### | 2.7594 | 5.7905E-03 | 1.2474E-02 |
| ENSG00000184988\|TMEM106A | 52.9496 | 2.1798 | ##### | 2.7592 | 5.7935E-03 | 1.2479E-02 |
| ENSG00000175390\|EIF3F | 8861.9844 | 0.3284 | ##### | 2.7589 | 5.7994E-03 | 1.2491E-02 |
| ENSG00000110367\|DDX6 | 8433.9122 | 0.3361 | ##### | 2.7588 | 5.8013E-03 | 1.2494E-02 |
| ENSG00000280789\|PAGR1 | 4685.4913 | 0.3543 | ##### | 2.7586 | 5.8046E-03 | 1.2500E-02 |
| ENSG00000246792 | 18.9432 | -2.3585 | ##### | -2.7574 | 5.8261E-03 | 1.2546E-02 |
| ENSG00000261595\|PPM1J-DT | 21.3644 | 7.1149 | ##### | 2.7566 | 5.8412E-03 | 1.2577E-02 |
| ENSG00000078081\|LAMP3 | 105.6432 | 0.6572 | ##### | 2.7564 | 5.8439E-03 | 1.2582E-02 |
| ENSG00000236489\|RPL34P10 | 5.3229 | -5.2754 | ##### | -2.7564 | 5.8446E-03 | 1.2583E-02 |
| ENSG00000289863 | 19.1500 | 2.4431 | ##### | 2.7557 | 5.8561E-03 | 1.2607E-02 |
| ENSG00000275638 | 11.6518 | 2.3583 | ##### | 2.7557 | 5.8573E-03 | 1.2608E-02 |
| ENSG00000147573\|TRIM55 | 36.3129 | 1.7307 | ##### | 2.7552 | 5.8650E-03 | 1.2624E-02 |
| ENSG00000284428 | 50.6039 | -1.1617 | ##### | -2.7548 | 5.8728E-03 | 1.2640E-02 |
| ENSG00000226783\|TLK1P1 | 5.2808 | -4.6587 | ##### | -2.7545 | 5.8773E-03 | 1.2649E-02 |
| ENSG00000267264 | 25.7246 | 1.0112 | ##### | 2.7531 | 5.9040E-03 | 1.2705E-02 |
| ENSG00000102908\|NFAT5 | 6492.5432 | 0.3391 | ##### | 2.7527 | 5.9111E-03 | 1.2720E-02 |
| ENSG00000272459\|FAM193B-DT | 14.1792 | 2.2301 | ##### | 2.7525 | 5.9147E-03 | 1.2727E-02 |
| ENSG00000169306\|IL1RAPL1 | 3.7698 | -4.6385 | ##### | -2.7521 | 5.9216E-03 | 1.2740E-02 |
| ENSG00000116981\|NT5C1A | 11.6448 | -1.8962 | ##### | -2.7515 | 5.9316E-03 | 1.2761E-02 |
| ENSG00000036448\|MYOM2 | 32.6678 | 3.2580 | ##### | 2.7512 | 5.9386E-03 | 1.2775E-02 |
| ENSG00000273384 | 9.7207 | -3.0370 | ##### | -2.7498 | 5.9640E-03 | 1.2829E-02 |
| ENSG00000138246\|DNAJC13 | 8375.9539 | 0.3347 | ##### | 2.7496 | 5.9671E-03 | 1.2835E-02 |
| ENSG00000244502\|HDAC11-AS1 | 37.8856 | -8.4977 | ##### | -2.7495 | 5.9686E-03 | 1.2837E-02 |
| ENSG00000160075\|SSU72 | 10538.9901 | -0.3421 | ##### | -2.7493 | 5.9717E-03 | 1.2842E-02 |
| ENSG00000139890\|REM2 | 41.8987 | 1.0147 | ##### | 2.7493 | 5.9716E-03 | 1.2842E-02 |
| ENSG00000138769\|CDKL2 | 40.1082 | 2.6281 | ##### | 2.7488 | 5.9807E-03 | 1.2860E-02 |
| ENSG00000261008\|LINC01572 | 130.9372 | 0.5296 | ##### | 2.7486 | 5.9846E-03 | 1.2867E-02 |
| ENSG00000273759 | 18.8079 | 1.5813 | ##### | 2.7485 | 5.9869E-03 | 1.2871E-02 |
| ENSG00000224789 | 10.2404 | -1.4215 | ##### | -2.7478 | 5.9992E-03 | 1.2897E-02 |
| ENSG00000251201\|TMED7-TICAM2 | 1519.8470 | -0.4224 | ##### | -2.7477 | 6.0018E-03 | 1.2902E-02 |
| ENSG00000147471\|PLPBP | 3269.1855 | -0.3383 | ##### | -2.7475 | 6.0044E-03 | 1.2906E-02 |
| ENSG00000269296 | 6.6103 | -2.5202 | ##### | -2.7472 | 6.0113E-03 | 1.2920E-02 |
| ENSG00000172232\|AZU1 | 26.3393 | -2.7240 | ##### | -2.7470 | 6.0148E-03 | 1.2927E-02 |
| ENSG00000035928\|RFC1 | 5817.8838 | 0.3248 | ##### | 2.7466 | 6.0218E-03 | 1.2941E-02 |
| ENSG00000234143\|UGT1A13P | 7.2828 | 6.6170 | ##### | 2.7463 | 6.0281E-03 | 1.2954E-02 |
| ENSG00000184708\|EIF4ENIF1 | 2724.2555 | -0.3312 | ##### | -2.7462 | 6.0289E-03 | 1.2954E-02 |
| ENSG00000122026\|RPL21 | 22509.6069 | 0.3367 | ##### | 2.7461 | 6.0306E-03 | 1.2957E-02 |
| ENSG00000260852\|FBXL19-AS1 | 463.3849 | -0.4084 | ##### | -2.7460 | 6.0323E-03 | 1.2960E-02 |
| ENSG00000176124\|DLEU1 | 1104.9283 | -0.3831 | ##### | -2.7454 | 6.0445E-03 | 1.2985E-02 |
| ENSG00000137996\|RTCA | 4761.0524 | -0.3296 | ##### | -2.7449 | 6.0528E-03 | 1.3002E-02 |
| ENSG00000067560\|RHOA | 39309.2357 | -0.3303 | ##### | -2.7446 | 6.0591E-03 | 1.3015E-02 |
| ENSG00000183690\|EFHC2 | 33.9168 | -2.3300 | ##### | -2.7441 | 6.0677E-03 | 1.3032E-02 |
| ENSG00000137337\|MDC1 | 6521.9183 | 0.3309 | ##### | 2.7438 | 6.0731E-03 | 1.3043E-02 |
| ENSG00000224067 | 18.5374 | -7.1405 | ##### | -2.7438 | 6.0736E-03 | 1.3043E-02 |
| ENSG00000104518\|GSDMD | 1152.1738 | -0.3723 | ##### | -2.7435 | 6.0794E-03 | 1.3054E-02 |
| ENSG00000281731\|METTL14-DT | 20.5975 | 7.0293 | ##### | 2.7432 | 6.0845E-03 | 1.3064E-02 |
| ENSG00000111850\|SMIM8 | 508.8327 | -0.4000 | ##### | -2.7427 | 6.0933E-03 | 1.3082E-02 |
| ENSG00000213711\|PHB1P7 | 7.5183 | 2.2768 | ##### | 2.7424 | 6.0983E-03 | 1.3092E-02 |
| ENSG00000148488\|ST8SIA6 | 40.2192 | -2.5685 | ##### | -2.7422 | 6.1024E-03 | 1.3100E-02 |
| ENSG00000103245\|CIAO3 | 2790.6598 | -0.3337 | ##### | -2.7422 | 6.1033E-03 | 1.3101E-02 |
| ENSG00000286492 | 3.6940 | 5.2732 | ##### | 2.7417 | 6.1130E-03 | 1.3121E-02 |
| ENSG00000287624 | 20.1211 | 1.1809 | ##### | 2.7415 | 6.1164E-03 | 1.3127E-02 |
| ENSG00000258388\|PPT2-EGFL8 | 2066.4172 | 0.3477 | ##### | 2.7406 | 6.1321E-03 | 1.3160E-02 |
| ENSG00000172936\|MYD88 | 1371.8300 | 0.3059 | ##### | 2.7404 | 6.1358E-03 | 1.3167E-02 |
| ENSG00000272514\|CFAP206 | 96.3702 | -1.1171 | ##### | -2.7401 | 6.1423E-03 | 1.3180E-02 |
| ENSG00000102606\|ARHGEF7 | 3275.0169 | -0.3399 | ##### | -2.7400 | 6.1433E-03 | 1.3181E-02 |
| ENSG00000286620 | 7.6838 | -3.0030 | ##### | -2.7396 | 6.1516E-03 | 1.3198E-02 |
| ENSG00000100124\|ANKRD54 | 1551.2713 | -0.3590 | ##### | -2.7389 | 6.1654E-03 | 1.3227E-02 |
| ENSG00000205236 | 206.5172 | 1.8603 | ##### | 2.7387 | 6.1678E-03 | 1.3231E-02 |

| | | | | | | |
|---|---|---|---|---|---|---|
| ENSG00000259322 | 3.3999 | 3.1183 | ##### | 2.7382 | 6.1783E-03 | 1.3253E-02 |
| ENSG00000125531\|FNDC11 | 67.0554 | -0.8210 | ##### | -2.7381 | 6.1798E-03 | 1.3255E-02 |
| ENSG00000178732\|GP5 | 8.7564 | 5.9729 | ##### | 2.7380 | 6.1821E-03 | 1.3259E-02 |
| ENSG00000185900\|POMK | 2409.8063 | 0.3648 | ##### | 2.7363 | 6.2131E-03 | 1.3324E-02 |
| ENSG00000224546\|EIF4BP3 | 41.6718 | -0.7603 | ##### | -2.7357 | 6.2246E-03 | 1.3348E-02 |
| ENSG00000250317\|SMIM20 | 1142.7482 | 0.3987 | ##### | 2.7355 | 6.2295E-03 | 1.3358E-02 |
| ENSG00000266111 | 21.6299 | -7.2317 | ##### | -2.7352 | 6.2337E-03 | 1.3366E-02 |
| ENSG00000231789\|PIK3CD-AS2 | 196.7783 | 0.5831 | ##### | 2.7347 | 6.2433E-03 | 1.3385E-02 |
| ENSG00000143363\|PRUNE1 | 2320.9244 | 0.3532 | ##### | 2.7342 | 6.2537E-03 | 1.3406E-02 |
| ENSG00000182993\|C12orf60 | 273.3695 | -0.4571 | ##### | -2.7342 | 6.2541E-03 | 1.3406E-02 |
| ENSG00000170075\|GPR37L1 | 11.7336 | -1.4809 | ##### | -2.7338 | 6.2602E-03 | 1.3419E-02 |
| ENSG00000289543 | 6.7553 | 3.6369 | ##### | 2.7335 | 6.2663E-03 | 1.3431E-02 |
| ENSG00000227470\|RPL39P16 | 11.0148 | -1.5003 | ##### | -2.7331 | 6.2745E-03 | 1.3447E-02 |
| ENSG00000128463\|EMC4 | 6572.1733 | 0.3321 | ##### | 2.7327 | 6.2818E-03 | 1.3460E-02 |
| ENSG00000276131 | 75.9229 | -0.6558 | ##### | -2.7327 | 6.2815E-03 | 1.3460E-02 |
| ENSG00000258839\|MC1R | 1679.8762 | 0.3515 | ##### | 2.7327 | 6.2817E-03 | 1.3460E-02 |
| ENSG00000254192 | 13.8509 | 1.4432 | ##### | 2.7326 | 6.2832E-03 | 1.3462E-02 |
| ENSG00000125878\|TCF15 | 83.1391 | 0.7576 | ##### | 2.7324 | 6.2866E-03 | 1.3468E-02 |
| ENSG00000255753 | 28.3683 | 1.7661 | ##### | 2.7322 | 6.2913E-03 | 1.3478E-02 |
| ENSG00000285292\|ABCF2-H2BK1 | 28.0772 | 3.6505 | ##### | 2.7321 | 6.2932E-03 | 1.3481E-02 |
| ENSG00000263756 | 290.6551 | 0.4627 | ##### | 2.7320 | 6.2960E-03 | 1.3485E-02 |
| ENSG00000164879\|CA3 | 80.1605 | -9.0423 | ##### | -2.7320 | 6.2957E-03 | 1.3485E-02 |
| ENSG00000171540\|OTP | 22.3973 | 2.7829 | ##### | 2.7317 | 6.3012E-03 | 1.3495E-02 |
| ENSG00000166896\|ATP23 | 892.4350 | 0.3686 | ##### | 2.7307 | 6.3203E-03 | 1.3535E-02 |
| ENSG00000287929 | 10.6032 | -1.8836 | ##### | -2.7306 | 6.3223E-03 | 1.3538E-02 |
| ENSG00000159658\|EFCAB14 | 8499.8168 | -0.3215 | ##### | -2.7304 | 6.3259E-03 | 1.3545E-02 |
| ENSG00000130725\|UBE2M | 12456.5535 | -0.3339 | ##### | -2.7300 | 6.3339E-03 | 1.3560E-02 |
| ENSG00000185972\|CCIN | 13.2146 | 1.7007 | ##### | 2.7300 | 6.3338E-03 | 1.3560E-02 |
| ENSG00000185522\|LMNTD2 | 250.0003 | -0.4971 | ##### | -2.7295 | 6.3428E-03 | 1.3578E-02 |
| ENSG00000293293 | 221.2140 | 0.5215 | ##### | 2.7281 | 6.3709E-03 | 1.3637E-02 |
| ENSG00000119684\|MLH3 | 1639.6442 | -0.3629 | ##### | -2.7279 | 6.3733E-03 | 1.3642E-02 |
| ENSG00000269959\|SPACA6-AS1 | 72.2747 | 0.6673 | ##### | 2.7278 | 6.3756E-03 | 1.3645E-02 |
| ENSG00000261351 | 118.0795 | 0.6864 | ##### | 2.7276 | 6.3806E-03 | 1.3655E-02 |
| ENSG00000289494 | 67.4228 | -0.7927 | ##### | -2.7269 | 6.3942E-03 | 1.3683E-02 |
| ENSG00000163938\|GNL3 | 10248.3902 | 0.2454 | ##### | 2.7264 | 6.4035E-03 | 1.3702E-02 |
| ENSG00000287032 | 34.3948 | -8.6608 | ##### | -2.7261 | 6.4093E-03 | 1.3714E-02 |
| ENSG00000232671\|ZNF687-AS1 | 91.4837 | 0.8159 | ##### | 2.7260 | 6.4115E-03 | 1.3717E-02 |
| ENSG00000117308\|GALE | 5252.5997 | -0.3230 | ##### | -2.7257 | 6.4158E-03 | 1.3725E-02 |
| ENSG00000253730 | 10.4309 | 2.3109 | ##### | 2.7257 | 6.4171E-03 | 1.3727E-02 |
| ENSG00000100253\|MIOX | 26.5326 | -6.6282 | ##### | -2.7251 | 6.4291E-03 | 1.3752E-02 |
| ENSG00000104825\|NFKBIB | 3946.8091 | -0.3411 | ##### | -2.7249 | 6.4327E-03 | 1.3759E-02 |
| ENSG00000198551\|ZNF627 | 728.0208 | 0.4052 | ##### | 2.7246 | 6.4383E-03 | 1.3770E-02 |
| ENSG00000270659 | 56.7741 | -2.4003 | ##### | -2.7243 | 6.4446E-03 | 1.3782E-02 |
| ENSG00000223482\|NUTM2A-AS1 | 2295.6522 | -0.3335 | ##### | -2.7235 | 6.4597E-03 | 1.3813E-02 |
| ENSG00000187479\|C11orf96 | 46.2414 | 1.4867 | ##### | 2.7233 | 6.4624E-03 | 1.3818E-02 |
| ENSG00000228897 | 8.2277 | 2.9553 | ##### | 2.7231 | 6.4672E-03 | 1.3828E-02 |
| ENSG00000143493\|INTS7 | 3465.8505 | -0.3364 | ##### | -2.7221 | 6.4875E-03 | 1.3869E-02 |
| ENSG00000179958\|DCTPP1 | 10776.2842 | 0.3044 | ##### | 2.7221 | 6.4877E-03 | 1.3869E-02 |
| ENSG00000215009\|ACSM4 | 8.1184 | 2.5445 | ##### | 2.7220 | 6.4898E-03 | 1.3873E-02 |
| ENSG00000142856\|ITGB3BP | 1365.3968 | -0.3538 | ##### | -2.7217 | 6.4951E-03 | 1.3881E-02 |
| ENSG00000270127 | 13.9083 | 2.6986 | ##### | 2.7217 | 6.4942E-03 | 1.3881E-02 |
| ENSG00000293180 | 22.0781 | 5.8052 | ##### | 2.7217 | 6.4951E-03 | 1.3881E-02 |
| ENSG00000176970\|RPL7L1P11 | 5.7143 | -5.4160 | ##### | -2.7216 | 6.4965E-03 | 1.3883E-02 |
| ENSG00000141002\|TCF25 | 7231.8912 | -0.3208 | ##### | -2.7216 | 6.4971E-03 | 1.3884E-02 |
| ENSG00000232237\|ASCL5 | 18.6607 | -1.7782 | ##### | -2.7211 | 6.5067E-03 | 1.3903E-02 |
| ENSG00000196417\|ZNF765 | 1334.4146 | -0.3591 | ##### | -2.7209 | 6.5098E-03 | 1.3909E-02 |
| ENSG00000286532\|PARTICL | 302.1132 | 0.4876 | ##### | 2.7199 | 6.5303E-03 | 1.3952E-02 |
| ENSG00000286281 | 36.7905 | 6.6338 | ##### | 2.7195 | 6.5377E-03 | 1.3966E-02 |
| ENSG00000108576\|SLC6A4 | 18.7439 | 2.3674 | ##### | 2.7194 | 6.5400E-03 | 1.3970E-02 |
| ENSG00000280063 | 67.0796 | 0.7300 | ##### | 2.7192 | 6.5440E-03 | 1.3978E-02 |
| ENSG00000197329\|PELI1 | 438.3358 | 0.4353 | ##### | 2.7188 | 6.5525E-03 | 1.3995E-02 |
| ENSG00000066455\|GOLGA5 | 3679.8043 | -0.3364 | ##### | -2.7182 | 6.5633E-03 | 1.4017E-02 |
| ENSG00000274767 | 21.1048 | 1.4962 | ##### | 2.7177 | 6.5731E-03 | 1.4037E-02 |

| | | | | | | |
|---|---|---|---|---|---|---|
| ENSG00000233175\|FMNL1-AS1 | 225.5765 | 0.5458 | ##### | 2.7174 | 6.5792E-03 | 1.4049E-02 |
| ENSG00000224738 | 167.6828 | 0.6079 | ##### | 2.7171 | 6.5859E-03 | 1.4062E-02 |
| ENSG00000207359\|RNU6-925P | 6.3704 | -2.7850 | ##### | -2.7168 | 6.5926E-03 | 1.4075E-02 |
| ENSG00000215347\|SLC25A5P1 | 5.4029 | -2.2928 | ##### | -2.7165 | 6.5973E-03 | 1.4084E-02 |
| ENSG00000006625\|GGCT | 3867.9544 | -0.3372 | ##### | -2.7165 | 6.5984E-03 | 1.4086E-02 |
| ENSG00000285508 | 25.0443 | -1.9137 | ##### | -2.7162 | 6.6033E-03 | 1.4095E-02 |
| ENSG00000272043 | 9.4409 | 2.1366 | ##### | 2.7155 | 6.6174E-03 | 1.4124E-02 |
| ENSG00000165169\|DYNLT3 | 1562.2888 | -0.3491 | ##### | -2.7154 | 6.6190E-03 | 1.4127E-02 |
| ENSG00000260368 | 16.2432 | 2.5138 | ##### | 2.7153 | 6.6215E-03 | 1.4131E-02 |
| ENSG00000273248 | 34.9374 | -0.9097 | ##### | -2.7147 | 6.6333E-03 | 1.4155E-02 |
| ENSG00000119771\|KLHL29 | 1329.7755 | -0.3596 | ##### | -2.7142 | 6.6443E-03 | 1.4178E-02 |
| ENSG00000264577 | 599.8418 | 0.4179 | ##### | 2.7140 | 6.6480E-03 | 1.4185E-02 |
| ENSG00000137713\|PPP2R1B | 3254.7508 | -0.3363 | ##### | -2.7129 | 6.6703E-03 | 1.4230E-02 |
| ENSG00000237499\|WAKMAR2 | 130.0756 | 0.5968 | ##### | 2.7129 | 6.6704E-03 | 1.4230E-02 |
| ENSG00000178935\|ZNF552 | 380.6503 | -0.4076 | ##### | -2.7128 | 6.6712E-03 | 1.4231E-02 |
| ENSG00000151466\|SCLT1 | 1040.7656 | -0.3654 | ##### | -2.7122 | 6.6842E-03 | 1.4258E-02 |
| ENSG00000255843 | 5.5899 | 5.0673 | ##### | 2.7119 | 6.6900E-03 | 1.4269E-02 |
| ENSG00000257624 | 4.1905 | 5.2780 | ##### | 2.7118 | 6.6926E-03 | 1.4274E-02 |
| ENSG00000137309\|HMGA1 | 37708.7160 | -0.3335 | ##### | -2.7113 | 6.7020E-03 | 1.4293E-02 |
| ENSG00000290588 | 73.1951 | 1.4912 | ##### | 2.7113 | 6.7029E-03 | 1.4294E-02 |
| ENSG00000291211 | 192.6191 | -0.4842 | ##### | -2.7111 | 6.7052E-03 | 1.4298E-02 |
| ENSG00000214203\|RPS4XP1 | 7.3367 | 1.9559 | ##### | 2.7106 | 6.7163E-03 | 1.4320E-02 |
| ENSG00000166889\|PATL1 | 5153.1151 | 0.3358 | ##### | 2.7104 | 6.7207E-03 | 1.4329E-02 |
| ENSG00000047056\|WDR37 | 1476.4422 | 0.3654 | ##### | 2.7103 | 6.7230E-03 | 1.4332E-02 |
| ENSG00000204120\|GIGYF2 | 7072.8330 | 0.3211 | ##### | 2.7101 | 6.7261E-03 | 1.4338E-02 |
| ENSG00000131871\|SELENOS | 4887.5376 | -0.3297 | ##### | -2.7092 | 6.7437E-03 | 1.4375E-02 |
| ENSG00000117983\|MUC5B | 38.3519 | -5.3346 | ##### | -2.7092 | 6.7450E-03 | 1.4376E-02 |
| ENSG00000288684 | 77.2551 | -2.1438 | ##### | -2.7086 | 6.7561E-03 | 1.4399E-02 |
| ENSG00000286703 | 9.6742 | 6.0719 | ##### | 2.7080 | 6.7690E-03 | 1.4425E-02 |
| ENSG00000278619\|MRM1 | 1210.7460 | 0.3596 | ##### | 2.7073 | 6.7827E-03 | 1.4454E-02 |
| ENSG00000237737\|DCTN1-AS1 | 11.0558 | 1.8484 | ##### | 2.7064 | 6.8021E-03 | 1.4494E-02 |
| ENSG00000197256\|KANK2 | 8993.9190 | -0.3126 | ##### | -2.7052 | 6.8257E-03 | 1.4543E-02 |
| ENSG00000284671 | 66.9629 | -9.0113 | ##### | -2.7045 | 6.8416E-03 | 1.4575E-02 |
| ENSG00000184307\|ZDHHC23 | 1575.0367 | 0.3778 | ##### | 2.7045 | 6.8416E-03 | 1.4575E-02 |
| ENSG00000135930\|EIF4E2 | 8489.4006 | -0.2806 | ##### | -2.7044 | 6.8425E-03 | 1.4576E-02 |
| ENSG00000284770\|TBCE | 3034.9072 | -0.5034 | ##### | -2.7042 | 6.8465E-03 | 1.4583E-02 |
| ENSG00000143815\|LBR | 5646.1010 | -0.3293 | ##### | -2.7041 | 6.8480E-03 | 1.4585E-02 |
| ENSG00000271270\|TMCC1-DT | 372.8565 | -0.5163 | ##### | -2.7039 | 6.8538E-03 | 1.4597E-02 |
| ENSG00000033122\|LRRC7 | 12.1769 | -1.4829 | ##### | -2.7035 | 6.8606E-03 | 1.4610E-02 |
| ENSG00000254593\|OR7E126P | 20.0976 | 1.0220 | ##### | 2.7034 | 6.8638E-03 | 1.4616E-02 |
| ENSG00000249971 | 67.4048 | -1.3981 | ##### | -2.7014 | 6.9053E-03 | 1.4703E-02 |
| ENSG00000067191\|CACNB1 | 889.9831 | 0.3497 | ##### | 2.7013 | 6.9064E-03 | 1.4704E-02 |
| ENSG00000136930\|PSMB7 | 15612.9726 | -0.3269 | ##### | -2.7011 | 6.9106E-03 | 1.4712E-02 |
| ENSG00000289047 | 1455.4087 | -0.3687 | ##### | -2.7010 | 6.9124E-03 | 1.4715E-02 |
| ENSG00000114455\|HHLA2 | 40.8801 | 7.4868 | ##### | 2.6979 | 6.9776E-03 | 1.4853E-02 |
| ENSG00000250734 | 11.2781 | 1.2932 | ##### | 2.6977 | 6.9821E-03 | 1.4861E-02 |
| ENSG00000225697\|SLC26A6 | 1360.3437 | 0.3638 | ##### | 2.6973 | 6.9904E-03 | 1.4878E-02 |
| ENSG00000232133\|IMPDH1P10 | 224.2111 | -0.5400 | ##### | -2.6971 | 6.9940E-03 | 1.4885E-02 |
| ENSG00000169905\|TOR1AIP2 | 5421.4496 | -0.3210 | ##### | -2.6971 | 6.9947E-03 | 1.4885E-02 |
| ENSG00000289021 | 53.8430 | -0.7720 | ##### | -2.6968 | 6.9999E-03 | 1.4895E-02 |
| ENSG00000184900\|SUMO3 | 8537.2607 | -0.2712 | ##### | -2.6961 | 7.0150E-03 | 1.4926E-02 |
| ENSG00000278816 | 8.5480 | -5.1647 | ##### | -2.6954 | 7.0308E-03 | 1.4959E-02 |
| ENSG00000289340 | 23.2488 | 5.6733 | ##### | 2.6953 | 7.0328E-03 | 1.4962E-02 |
| ENSG00000236048\|SSBL5P | 16.0020 | 1.0731 | ##### | 2.6949 | 7.0409E-03 | 1.4978E-02 |
| ENSG00000250934 | 50.3548 | 0.9079 | ##### | 2.6949 | 7.0414E-03 | 1.4978E-02 |
| ENSG00000167992\|VWCE | 29.9745 | 1.1213 | ##### | 2.6948 | 7.0431E-03 | 1.4981E-02 |
| ENSG00000277150\|F8A3 | 1632.7049 | 0.3494 | ##### | 2.6944 | 7.0511E-03 | 1.4997E-02 |
| ENSG00000099840\|IZUMO4 | 39.1736 | 1.2900 | ##### | 2.6943 | 7.0532E-03 | 1.5000E-02 |
| ENSG00000250629 | 15.2504 | -6.8581 | ##### | -2.6936 | 7.0680E-03 | 1.5030E-02 |
| ENSG00000242999\|RN7SL239P | 12.0045 | 5.3438 | ##### | 2.6935 | 7.0709E-03 | 1.5035E-02 |
| ENSG00000260335 | 4.8010 | 4.4613 | ##### | 2.6935 | 7.0715E-03 | 1.5036E-02 |
| ENSG00000261609\|GAN | 1454.0970 | -0.3506 | ##### | -2.6928 | 7.0849E-03 | 1.5062E-02 |
| ENSG00000273554 | 8.0341 | 4.4207 | ##### | 2.6928 | 7.0845E-03 | 1.5062E-02 |

| | | | | | | |
|---|---|---|---|---|---|---|
| ENSG00000278060 | 5.4118 | 4.2377 | ##### | 2.6928 | 7.0864E-03 | 1.5064E-02 |
| ENSG00000259032\|ENSAP2 | 144.9815 | 0.5959 | ##### | 2.6925 | 7.0927E-03 | 1.5076E-02 |
| ENSG00000075618\|FSCN1 | 26208.8577 | -0.3214 | ##### | -2.6923 | 7.0960E-03 | 1.5082E-02 |
| ENSG00000231125 | 14.2082 | 1.5763 | ##### | 2.6921 | 7.1014E-03 | 1.5093E-02 |
| ENSG00000116251\|RPL22 | 34245.4275 | 0.2892 | ##### | 2.6920 | 7.1028E-03 | 1.5095E-02 |
| ENSG00000178631\|ACTG1P1 | 15.8861 | -1.5093 | ##### | -2.6918 | 7.1062E-03 | 1.5101E-02 |
| ENSG00000134184\|GSTM1 | 64.8113 | 0.9835 | ##### | 2.6916 | 7.1108E-03 | 1.5109E-02 |
| ENSG00000232729\|GTF2I-AS1 | 39.2395 | -4.0249 | ##### | -2.6903 | 7.1392E-03 | 1.5169E-02 |
| ENSG00000170525\|PFKFB3 | 14085.7187 | -0.2733 | ##### | -2.6892 | 7.1632E-03 | 1.5219E-02 |
| ENSG00000257596\|SCAT2 | 17.9695 | 1.1268 | ##### | 2.6891 | 7.1646E-03 | 1.5221E-02 |
| ENSG00000130675\|MNX1 | 292.5962 | 0.4715 | ##### | 2.6890 | 7.1669E-03 | 1.5224E-02 |
| ENSG00000260898\|ADPGK-AS1 | 33.8450 | 1.0276 | ##### | 2.6888 | 7.1705E-03 | 1.5231E-02 |
| ENSG00000091972\|CD200 | 40.8900 | 2.6765 | ##### | 2.6883 | 7.1809E-03 | 1.5252E-02 |
| ENSG00000219201 | 1707.8399 | 0.3538 | ##### | 2.6883 | 7.1824E-03 | 1.5254E-02 |
| ENSG00000214401\|KANSL1-AS1 | 39.7509 | 0.8972 | ##### | 2.6882 | 7.1842E-03 | 1.5257E-02 |
| ENSG00000167962\|ZNF598 | 10586.6984 | 0.3286 | ##### | 2.6877 | 7.1948E-03 | 1.5278E-02 |
| ENSG00000148943\|LIN7C | 2217.3910 | -0.3537 | ##### | -2.6876 | 7.1961E-03 | 1.5279E-02 |
| ENSG00000287751\|FAM106B | 4.0723 | 5.2781 | ##### | 2.6876 | 7.1958E-03 | 1.5279E-02 |
| ENSG00000158683\|PKD1L1 | 41.8892 | 2.2503 | ##### | 2.6874 | 7.2007E-03 | 1.5287E-02 |
| ENSG00000148481\|MINDY3 | 1590.5853 | 0.3604 | ##### | 2.6871 | 7.2065E-03 | 1.5299E-02 |
| ENSG00000105640\|RPL18A | 36807.0715 | 0.3290 | ##### | 2.6868 | 7.2147E-03 | 1.5315E-02 |
| ENSG00000088386\|SLC15A1 | 17.0576 | -2.4536 | ##### | -2.6867 | 7.2158E-03 | 1.5316E-02 |
| ENSG00000184838\|PRR16 | 98.0656 | 0.7674 | ##### | 2.6867 | 7.2169E-03 | 1.5317E-02 |
| ENSG00000286367 | 9.4707 | -2.2052 | ##### | -2.6865 | 7.2201E-03 | 1.5323E-02 |
| ENSG00000134809\|TIMM10 | 1883.4393 | 0.3539 | ##### | 2.6861 | 7.2291E-03 | 1.5341E-02 |
| ENSG00000206192\|ANKRD20A9P | 6.7557 | 5.0440 | ##### | 2.6858 | 7.2352E-03 | 1.5353E-02 |
| ENSG00000124232\|RBPJL | 13.4991 | 4.0557 | ##### | 2.6853 | 7.2455E-03 | 1.5374E-02 |
| ENSG00000270006\|C16orf95-DT | 15.9725 | -1.6599 | ##### | -2.6849 | 7.2541E-03 | 1.5391E-02 |
| ENSG00000103740\|ACSBG1 | 1614.9011 | 0.3930 | ##### | 2.6837 | 7.2804E-03 | 1.5446E-02 |
| ENSG00000102387\|TAF7L | 25.2379 | 2.1749 | ##### | 2.6832 | 7.2925E-03 | 1.5470E-02 |
| ENSG00000197978\|GOLGA6L9 | 327.2333 | -0.4199 | ##### | -2.6831 | 7.2943E-03 | 1.5473E-02 |
| ENSG00000106771\|TMEM245 | 17086.5713 | -0.3213 | ##### | -2.6822 | 7.3134E-03 | 1.5512E-02 |
| ENSG00000251257 | 35.1351 | 8.3078 | ##### | 2.6822 | 7.3143E-03 | 1.5513E-02 |
| ENSG00000151151\|IPMK | 1204.9518 | -0.3748 | ##### | -2.6821 | 7.3160E-03 | 1.5516E-02 |
| ENSG00000108298\|RPL19 | 59658.5691 | 0.3264 | ##### | 2.6818 | 7.3232E-03 | 1.5530E-02 |
| ENSG00000100129\|EIF3L | 13676.7975 | 0.3208 | ##### | 2.6817 | 7.3249E-03 | 1.5532E-02 |
| ENSG00000279945 | 378.0988 | 0.4546 | ##### | 2.6814 | 7.3320E-03 | 1.5546E-02 |
| ENSG00000267249 | 103.5197 | 0.6506 | ##### | 2.6811 | 7.3385E-03 | 1.5559E-02 |
| ENSG00000129355\|CDKN2D | 327.1825 | -0.5128 | ##### | -2.6809 | 7.3420E-03 | 1.5565E-02 |
| ENSG00000140006\|WDR89 | 3148.4804 | 0.3262 | ##### | 2.6807 | 7.3479E-03 | 1.5577E-02 |
| ENSG00000116685\|KIAA2013 | 6670.3616 | -0.3303 | ##### | -2.6806 | 7.3496E-03 | 1.5579E-02 |
| ENSG00000168748\|CA7 | 27.3572 | 3.0203 | ##### | 2.6805 | 7.3509E-03 | 1.5581E-02 |
| ENSG00000285770 | 3.3552 | -5.2041 | ##### | -2.6802 | 7.3568E-03 | 1.5592E-02 |
| ENSG00000204482\|LST1 | 19.6554 | 2.0436 | ##### | 2.6798 | 7.3658E-03 | 1.5610E-02 |
| ENSG00000119729\|RHOQ | 5118.8355 | -0.3370 | ##### | -2.6798 | 7.3668E-03 | 1.5611E-02 |
| ENSG00000255725\|TDGP1 | 42.3999 | -1.0718 | ##### | -2.6795 | 7.3730E-03 | 1.5623E-02 |
| ENSG00000279064 | 6.3890 | -1.9429 | ##### | -2.6794 | 7.3764E-03 | 1.5629E-02 |
| ENSG00000099364\|FBXL19 | 5042.5321 | 0.3203 | ##### | 2.6789 | 7.3854E-03 | 1.5647E-02 |
| ENSG00000274943 | 32.6805 | 1.3121 | ##### | 2.6784 | 7.3979E-03 | 1.5673E-02 |
| ENSG00000228013\|IL6R-AS1 | 16.0443 | -2.7554 | ##### | -2.6783 | 7.4005E-03 | 1.5677E-02 |
| ENSG00000237870 | 11.5666 | -1.8978 | ##### | -2.6781 | 7.4030E-03 | 1.5681E-02 |
| ENSG00000115286\|NDUFS7 | 6256.8210 | 0.3337 | ##### | 2.6780 | 7.4056E-03 | 1.5686E-02 |
| ENSG00000031698\|SARS1 | 14006.7973 | 0.2902 | ##### | 2.6777 | 7.4139E-03 | 1.5702E-02 |
| ENSG00000177374\|HIC1 | 568.6748 | 0.9260 | ##### | 2.6775 | 7.4165E-03 | 1.5707E-02 |
| ENSG00000268951\|CDRT15P7 | 3.9205 | -2.8408 | ##### | -2.6772 | 7.4249E-03 | 1.5723E-02 |
| ENSG00000240914\|RPL15P2 | 9.7485 | -1.5500 | ##### | -2.6769 | 7.4308E-03 | 1.5734E-02 |
| ENSG00000271991\|TTC32-DT | 23.2739 | 2.1075 | ##### | 2.6768 | 7.4318E-03 | 1.5736E-02 |
| ENSG00000226352\|PSPC1-AS2 | 18.6142 | 1.2870 | ##### | 2.6766 | 7.4369E-03 | 1.5745E-02 |
| ENSG00000182473\|EXOC7 | 16448.9381 | 0.3141 | ##### | 2.6766 | 7.4378E-03 | 1.5746E-02 |
| ENSG00000235720\|GABPAP | 90.4635 | -0.5879 | ##### | -2.6760 | 7.4499E-03 | 1.5771E-02 |
| ENSG00000112511\|PHF1 | 2249.1035 | -0.3395 | ##### | -2.6758 | 7.4553E-03 | 1.5781E-02 |
| ENSG00000225526\|MKRN2OS | 11.9119 | 2.4349 | ##### | 2.6756 | 7.4600E-03 | 1.5790E-02 |
| ENSG00000165810\|BTNL9 | 652.5646 | -0.3878 | ##### | -2.6746 | 7.4822E-03 | 1.5836E-02 |

| | | | | | | |
|---|---|---|---|---|---|---|
| ENSG00000160613\|PCSK7 | 2948.2573 | -0.3366 | ##### | -2.6743 | 7.4878E-03 | 1.5846E-02 |
| ENSG00000055070\|SZRD1 | 19844.5500 | -0.3076 | ##### | -2.6733 | 7.5108E-03 | 1.5894E-02 |
| ENSG00000225486 | 6.6575 | -5.2830 | ##### | -2.6731 | 7.5146E-03 | 1.5901E-02 |
| ENSG00000278133 | 166.6360 | -0.4898 | ##### | -2.6729 | 7.5201E-03 | 1.5911E-02 |
| ENSG00000183208\|GDPGP1 | 363.1063 | -0.4242 | ##### | -2.6728 | 7.5216E-03 | 1.5913E-02 |
| ENSG00000138698\|RAP1GDS1 | 4435.8442 | -0.3239 | ##### | -2.6721 | 7.5389E-03 | 1.5949E-02 |
| ENSG00000122566\|HNRNPA2B1 | 86211.0242 | 0.3288 | ##### | 2.6720 | 7.5394E-03 | 1.5949E-02 |
| ENSG00000236308 | 15.8184 | 1.5335 | ##### | 2.6715 | 7.5514E-03 | 1.5972E-02 |
| ENSG00000180071\|ANKRD18A | 424.3700 | -0.4439 | ##### | -2.6715 | 7.5509E-03 | 1.5972E-02 |
| ENSG00000100580\|TMED8 | 2958.1454 | -0.3308 | ##### | -2.6714 | 7.5543E-03 | 1.5977E-02 |
| ENSG00000282978\|BTF3P16 | 23.1647 | 1.2603 | ##### | 2.6713 | 7.5562E-03 | 1.5980E-02 |
| ENSG00000185222\|TCEAL9 | 4943.7451 | 0.2872 | ##### | 2.6710 | 7.5621E-03 | 1.5991E-02 |
| ENSG00000103047\|TANGO6 | 2497.4598 | -0.3439 | ##### | -2.6704 | 7.5772E-03 | 1.6022E-02 |
| ENSG00000127418\|FGFRL1 | 1578.5414 | 0.3325 | ##### | 2.6703 | 7.5781E-03 | 1.6023E-02 |
| ENSG00000200966\|RN7SKP87 | 7.1925 | 5.8439 | ##### | 2.6701 | 7.5829E-03 | 1.6032E-02 |
| ENSG00000290956\|FAM21EP | 57.5887 | -0.7183 | ##### | -2.6700 | 7.5861E-03 | 1.6036E-02 |
| ENSG00000239614\|HMGN1P7 | 6.5182 | 5.4695 | ##### | 2.6700 | 7.5858E-03 | 1.6036E-02 |
| ENSG00000254162 | 24.5440 | 1.5961 | ##### | 2.6692 | 7.6027E-03 | 1.6070E-02 |
| ENSG00000227203\|SUB1P1 | 11.7623 | -1.5957 | ##### | -2.6691 | 7.6045E-03 | 1.6073E-02 |
| ENSG00000088876\|ZNF343 | 1065.3313 | 0.3690 | ##### | 2.6682 | 7.6257E-03 | 1.6116E-02 |
| ENSG00000102109\|PCSK1N | 2146.6814 | -0.3234 | ##### | -2.6682 | 7.6269E-03 | 1.6118E-02 |
| ENSG00000293482 | 52.3603 | 0.7214 | ##### | 2.6681 | 7.6282E-03 | 1.6119E-02 |
| ENSG00000152443\|ZNF776 | 996.9956 | -0.3846 | ##### | -2.6676 | 7.6401E-03 | 1.6143E-02 |
| ENSG00000262692 | 13.2201 | 2.1290 | ##### | 2.6673 | 7.6469E-03 | 1.6157E-02 |
| ENSG00000273720 | 8.6211 | 4.1911 | ##### | 2.6671 | 7.6518E-03 | 1.6166E-02 |
| ENSG00000213885\|RPL13AP7 | 16.0150 | -1.1835 | ##### | -2.6668 | 7.6577E-03 | 1.6177E-02 |
| ENSG00000271151 | 13.2318 | 2.4455 | ##### | 2.6659 | 7.6776E-03 | 1.6217E-02 |
| ENSG00000147394\|ZNF185 | 826.4643 | -0.3781 | ##### | -2.6660 | 7.6771E-03 | 1.6217E-02 |
| ENSG00000258284\|POLR2KP1 | 4.0252 | 5.0274 | ##### | 2.6654 | 7.6898E-03 | 1.6242E-02 |
| ENSG00000185247\|MAGEA11 | 9.4837 | -5.6436 | ##### | -2.6650 | 7.6982E-03 | 1.6258E-02 |
| ENSG00000169891\|REPS2 | 47.7575 | 0.9079 | ##### | 2.6648 | 7.7029E-03 | 1.6267E-02 |
| ENSG00000223646\|LINC03076 | 6.5833 | -2.2018 | ##### | -2.6648 | 7.7045E-03 | 1.6269E-02 |
| ENSG00000283167 | 5.1284 | -3.6953 | ##### | -2.6646 | 7.7084E-03 | 1.6276E-02 |
| ENSG00000258469\|CHMP4BP1 | 7.1683 | 3.0956 | ##### | 2.6642 | 7.7162E-03 | 1.6292E-02 |
| ENSG00000266079\|SNORA59B | 4.8466 | -3.1194 | ##### | -2.6642 | 7.7178E-03 | 1.6294E-02 |
| ENSG00000169744\|LDB2 | 9.6168 | -2.7750 | ##### | -2.6635 | 7.7340E-03 | 1.6327E-02 |
| ENSG00000187608\|ISG15 | 1083.6258 | 0.3499 | ##### | 2.6629 | 7.7479E-03 | 1.6355E-02 |
| ENSG00000289701 | 250.2773 | -0.5928 | ##### | -2.6628 | 7.7485E-03 | 1.6355E-02 |
| ENSG00000274925\|ZKSCAN2-DT | 95.5148 | 0.6525 | ##### | 2.6628 | 7.7500E-03 | 1.6357E-02 |
| ENSG00000083845\|RPS5 | 41194.6669 | -0.3122 | ##### | -2.6627 | 7.7514E-03 | 1.6359E-02 |
| ENSG00000168438\|CDC40 | 2455.2311 | 0.3299 | ##### | 2.6626 | 7.7536E-03 | 1.6362E-02 |
| ENSG00000213621\|RPSAP54 | 24.7490 | 0.9385 | ##### | 2.6623 | 7.7609E-03 | 1.6377E-02 |
| ENSG00000216867\|RPL22P12 | 11.4472 | 2.4444 | ##### | 2.6620 | 7.7677E-03 | 1.6390E-02 |
| ENSG00000129515\|SNX6 | 8941.5000 | -0.3090 | ##### | -2.6617 | 7.7750E-03 | 1.6404E-02 |
| ENSG00000126088\|UROD | 4101.0683 | -0.3274 | ##### | -2.6614 | 7.7825E-03 | 1.6419E-02 |
| ENSG00000233122\|CTAGE7P | 31.2300 | 1.2579 | ##### | 2.6611 | 7.7875E-03 | 1.6428E-02 |
| ENSG00000185818\|NAT8L | 2303.3060 | 0.3377 | ##### | 2.6607 | 7.7985E-03 | 1.6450E-02 |
| ENSG00000236830\|CBR3-AS1 | 118.9050 | -0.5656 | ##### | -2.6604 | 7.8050E-03 | 1.6463E-02 |
| ENSG00000131503\|ANKHD1 | 4570.8630 | 0.6270 | ##### | 2.6601 | 7.8120E-03 | 1.6476E-02 |
| ENSG00000177984\|LCN15 | 12.2036 | -2.7174 | ##### | -2.6600 | 7.8136E-03 | 1.6479E-02 |
| ENSG00000275552 | 19.0011 | -2.6148 | ##### | -2.6599 | 7.8162E-03 | 1.6483E-02 |
| ENSG00000182307\|C8orf33 | 8275.1002 | -0.2884 | ##### | -2.6597 | 7.8211E-03 | 1.6492E-02 |
| ENSG00000161542\|PRPSAP1 | 2727.5726 | 0.3457 | ##### | 2.6594 | 7.8274E-03 | 1.6504E-02 |
| ENSG00000013288\|MAN2B2 | 5254.4656 | 0.3129 | ##### | 2.6592 | 7.8335E-03 | 1.6516E-02 |
| ENSG00000262304 | 4.9896 | -6.1967 | ##### | -2.6590 | 7.8370E-03 | 1.6522E-02 |
| ENSG00000226396 | 29.2300 | 1.0700 | ##### | 2.6585 | 7.8497E-03 | 1.6548E-02 |
| ENSG00000230479\|LNCTSI | 67.4005 | -0.6930 | ##### | -2.6584 | 7.8521E-03 | 1.6552E-02 |
| ENSG00000135736\|CCDC102A | 1523.9203 | 0.3399 | ##### | 2.6583 | 7.8527E-03 | 1.6552E-02 |
| ENSG00000286587 | 22.5086 | -0.8211 | ##### | -2.6582 | 7.8560E-03 | 1.6558E-02 |
| ENSG00000177197\|PCNPP5 | 9.6996 | 1.8064 | ##### | 2.6579 | 7.8622E-03 | 1.6569E-02 |
| ENSG00000211563\|MIR3065 | 4.7860 | 4.3342 | ##### | 2.6578 | 7.8643E-03 | 1.6571E-02 |
| ENSG00000138459\|SLC35A5 | 1962.9124 | -0.3384 | ##### | -2.6579 | 7.8641E-03 | 1.6571E-02 |
| ENSG00000177971\|IMP3 | 4309.8401 | 0.3262 | ##### | 2.6575 | 7.8731E-03 | 1.6589E-02 |

| | | | | | | |
|---|---|---|---|---|---|---|
| ENSG00000258465 | 12.5685 | 2.6679 | ##### | 2.6574 | 7.8742E-03 | 1.6590E-02 |
| ENSG00000244097\|RPS4XP17 | 12.9493 | -2.3057 | ##### | -2.6572 | 7.8793E-03 | 1.6599E-02 |
| ENSG00000275709 | 9.2134 | 4.8920 | ##### | 2.6570 | 7.8842E-03 | 1.6609E-02 |
| ENSG00000212664\|RPL17P39 | 59.5990 | -0.7836 | ##### | -2.6553 | 7.9231E-03 | 1.6689E-02 |
| ENSG00000269915 | 4.9954 | -3.0217 | ##### | -2.6547 | 7.9385E-03 | 1.6721E-02 |
| ENSG00000285982 | 20.4458 | -7.7025 | ##### | -2.6545 | 7.9435E-03 | 1.6730E-02 |
| ENSG00000138463\|SLC49A4 | 1401.3837 | -0.3481 | ##### | -2.6539 | 7.9559E-03 | 1.6755E-02 |
| ENSG00000103415\|HMOX2 | 3451.6398 | -0.3269 | ##### | -2.6535 | 7.9657E-03 | 1.6774E-02 |
| ENSG00000122512\|PMS2 | 5100.9130 | 0.2518 | ##### | 2.6532 | 7.9726E-03 | 1.6788E-02 |
| ENSG00000279041 | 23.2566 | -2.3916 | ##### | -2.6529 | 7.9797E-03 | 1.6802E-02 |
| ENSG00000167377\|ZNF23 | 710.3400 | -0.5272 | ##### | -2.6513 | 8.0193E-03 | 1.6884E-02 |
| ENSG00000117242\|PINK1-AS | 3031.5090 | -0.3338 | ##### | -2.6511 | 8.0224E-03 | 1.6889E-02 |
| ENSG00000254706 | 6.7847 | -3.5041 | ##### | -2.6510 | 8.0247E-03 | 1.6893E-02 |
| ENSG00000121964\|GTDC1 | 828.9204 | 0.3723 | ##### | 2.6503 | 8.0412E-03 | 1.6926E-02 |
| ENSG00000185974\|GRK1 | 8.9486 | 5.4012 | ##### | 2.6494 | 8.0628E-03 | 1.6971E-02 |
| ENSG00000109670\|FBXW7 | 940.3238 | -0.3859 | ##### | -2.6494 | 8.0644E-03 | 1.6973E-02 |
| ENSG00000271555 | 4.6242 | 5.1978 | ##### | 2.6489 | 8.0753E-03 | 1.6994E-02 |
| ENSG00000251205 | 22.1379 | 1.5156 | ##### | 2.6488 | 8.0777E-03 | 1.6998E-02 |
| ENSG00000263432\|RN7SL689P | 20.5730 | -1.3247 | ##### | -2.6487 | 8.0813E-03 | 1.7005E-02 |
| ENSG00000259185 | 102.2628 | -0.6611 | ##### | -2.6484 | 8.0874E-03 | 1.7016E-02 |
| ENSG00000204564\|C6orf136 | 881.6881 | -0.3746 | ##### | -2.6483 | 8.0902E-03 | 1.7021E-02 |
| ENSG00000166793\|YPEL4 | 29.4835 | 1.6319 | ##### | 2.6475 | 8.1080E-03 | 1.7057E-02 |
| ENSG00000170425\|ADORA2B | 1136.1470 | 0.3309 | ##### | 2.6469 | 8.1222E-03 | 1.7086E-02 |
| ENSG00000259605 | 971.3181 | -0.3740 | ##### | -2.6467 | 8.1273E-03 | 1.7096E-02 |
| ENSG00000280927\|CTBP1-AS | 766.3124 | 0.3944 | ##### | 2.6466 | 8.1309E-03 | 1.7102E-02 |
| ENSG00000271978 | 57.8536 | 0.8209 | ##### | 2.6459 | 8.1464E-03 | 1.7133E-02 |
| ENSG00000270002 | 20.2139 | 2.3158 | ##### | 2.6456 | 8.1545E-03 | 1.7149E-02 |
| ENSG00000206992\|RNU6-574P | 8.5932 | 4.0169 | ##### | 2.6453 | 8.1620E-03 | 1.7164E-02 |
| ENSG00000258982 | 8.2902 | -1.9497 | ##### | -2.6446 | 8.1777E-03 | 1.7196E-02 |
| ENSG00000171873\|ADRA1D | 61.7278 | -1.1099 | ##### | -2.6444 | 8.1833E-03 | 1.7206E-02 |
| ENSG00000284691 | 353.0238 | -0.4239 | ##### | -2.6443 | 8.1866E-03 | 1.7212E-02 |
| ENSG00000290024 | 16.9567 | 2.7133 | ##### | 2.6434 | 8.2075E-03 | 1.7254E-02 |
| ENSG00000182220\|ATP6AP2 | 7153.0734 | -0.3126 | ##### | -2.6433 | 8.2101E-03 | 1.7259E-02 |
| ENSG00000124659\|TBCC | 1587.0616 | 0.3387 | ##### | 2.6432 | 8.2127E-03 | 1.7263E-02 |
| ENSG00000186047\|DLEU7 | 9.5614 | -3.3596 | ##### | -2.6425 | 8.2296E-03 | 1.7297E-02 |
| ENSG00000166796\|LDHC | 39.2166 | 0.9057 | ##### | 2.6424 | 8.2311E-03 | 1.7299E-02 |
| ENSG00000273893 | 65.8157 | 1.0091 | ##### | 2.6415 | 8.2539E-03 | 1.7343E-02 |
| ENSG00000237222\|LINC01968 | 11.1353 | 3.4499 | ##### | 2.6415 | 8.2535E-03 | 1.7343E-02 |
| ENSG00000105982\|RNF32 | 340.8631 | 0.5081 | ##### | 2.6415 | 8.2537E-03 | 1.7343E-02 |
| ENSG00000112514\|CUTA | 5417.9915 | -0.3286 | ##### | -2.6413 | 8.2589E-03 | 1.7353E-02 |
| ENSG00000110958\|PTGES3 | 43427.0395 | 0.3238 | ##### | 2.6410 | 8.2659E-03 | 1.7366E-02 |
| ENSG00000130584\|ZBTB46 | 449.0779 | -0.3885 | ##### | -2.6408 | 8.2720E-03 | 1.7378E-02 |
| ENSG00000163517\|HDAC11 | 534.9599 | 0.4016 | ##### | 2.6402 | 8.2845E-03 | 1.7403E-02 |
| ENSG00000289161 | 271.5814 | -0.4638 | ##### | -2.6401 | 8.2875E-03 | 1.7408E-02 |
| ENSG00000178567\|EPM2AIP1 | 1503.5173 | -0.3535 | ##### | -2.6400 | 8.2914E-03 | 1.7415E-02 |
| ENSG00000138071\|ACTR2 | 38480.3167 | 0.2457 | ##### | 2.6399 | 8.2933E-03 | 1.7418E-02 |
| ENSG00000012061\|ERCC1 | 9129.5340 | -0.3224 | ##### | -2.6397 | 8.2972E-03 | 1.7425E-02 |
| ENSG00000217889\|KRT18P48 | 4.2946 | -2.5566 | ##### | -2.6396 | 8.2992E-03 | 1.7428E-02 |
| ENSG00000070886\|EPHA8 | 11.8405 | -5.2198 | ##### | -2.6396 | 8.3014E-03 | 1.7431E-02 |
| ENSG00000168397\|ATG4B | 5194.4984 | 0.3182 | ##### | 2.6391 | 8.3115E-03 | 1.7450E-02 |
| ENSG00000165233\|CARD19 | 2213.1194 | -0.3452 | ##### | -2.6391 | 8.3116E-03 | 1.7450E-02 |
| ENSG00000269688 | 32.9898 | 0.9369 | ##### | 2.6385 | 8.3262E-03 | 1.7479E-02 |
| ENSG00000176095\|IP6K1 | 4170.9967 | 0.3245 | ##### | 2.6384 | 8.3288E-03 | 1.7483E-02 |
| ENSG00000107290\|SETX | 6936.3739 | 0.3312 | ##### | 2.6384 | 8.3289E-03 | 1.7483E-02 |
| ENSG00000272205\|POU2F1-DT | 14.5629 | 1.7185 | ##### | 2.6383 | 8.3311E-03 | 1.7486E-02 |
| ENSG00000172943\|PHF8 | 2579.0655 | -0.3350 | ##### | -2.6383 | 8.3335E-03 | 1.7490E-02 |
| ENSG00000127054\|INTS11 | 7536.4687 | 0.3044 | ##### | 2.6378 | 8.3450E-03 | 1.7512E-02 |
| ENSG00000235016\|SEMA3F-AS1 | 86.2295 | 0.7004 | ##### | 2.6378 | 8.3446E-03 | 1.7512E-02 |
| ENSG00000103852\|TTC23 | 944.4980 | -0.3519 | ##### | -2.6375 | 8.3530E-03 | 1.7527E-02 |
| ENSG00000117298\|ECE1 | 14091.4066 | -0.3093 | ##### | -2.6372 | 8.3595E-03 | 1.7539E-02 |
| ENSG00000272274\|LINC00551 | 6.7828 | 3.5871 | ##### | 2.6345 | 8.4260E-03 | 1.7678E-02 |
| ENSG00000153086\|ACMSD | 8.9932 | -2.9247 | ##### | -2.6344 | 8.4296E-03 | 1.7683E-02 |
| ENSG00000237693\|IRGM | 7.5505 | 4.5908 | ##### | 2.6344 | 8.4297E-03 | 1.7683E-02 |

| | | | | | | |
|---|---|---|---|---|---|---|
| ENSG00000261221\|ZNF865 | 3691.2716 | -0.3172 | ##### | -2.6343 | 8.4306E-03 | 1.7684E-02 |
| ENSG00000278743 | 22.7418 | 1.7942 | ##### | 2.6342 | 8.4326E-03 | 1.7684E-02 |
| ENSG00000113593\|PPWD1 | 2214.6509 | 0.3203 | ##### | 2.6342 | 8.4324E-03 | 1.7684E-02 |
| ENSG00000188163\|CIMIP2A | 29.8151 | 1.0287 | ##### | 2.6343 | 8.4322E-03 | 1.7684E-02 |
| ENSG00000014641\|MDH1 | 12815.4132 | 0.3230 | ##### | 2.6341 | 8.4364E-03 | 1.7689E-02 |
| ENSG00000136895\|GARNL3 | 289.3656 | 0.4688 | ##### | 2.6341 | 8.4365E-03 | 1.7689E-02 |
| ENSG00000122584\|NXPH1 | 5.7531 | -5.9238 | ##### | -2.6341 | 8.4361E-03 | 1.7689E-02 |
| ENSG00000266283 | 9.8022 | -1.3928 | ##### | -2.6335 | 8.4506E-03 | 1.7717E-02 |
| ENSG00000279568 | 20.6489 | 1.1533 | ##### | 2.6323 | 8.4804E-03 | 1.7778E-02 |
| ENSG00000087301\|TXNDC16 | 354.8553 | -0.4276 | ##### | -2.6318 | 8.4939E-03 | 1.7805E-02 |
| ENSG00000169118\|CSNK1G1 | 2712.3624 | 0.3351 | ##### | 2.6317 | 8.4947E-03 | 1.7806E-02 |
| ENSG00000261187 | 20.9188 | -6.5140 | ##### | -2.6307 | 8.5209E-03 | 1.7859E-02 |
| ENSG00000146856\|AGBL3 | 147.2940 | -0.5185 | ##### | -2.6304 | 8.5288E-03 | 1.7875E-02 |
| ENSG00000276997 | 58.1173 | -0.8393 | ##### | -2.6301 | 8.5349E-03 | 1.7886E-02 |
| ENSG00000152092\|ASTN1 | 8.4634 | -3.3643 | ##### | -2.6301 | 8.5363E-03 | 1.7888E-02 |
| ENSG00000232479\|KRT8P52 | 13.1750 | -1.4861 | ##### | -2.6298 | 8.5443E-03 | 1.7902E-02 |
| ENSG00000204227\|RING1 | 2716.0336 | -0.3089 | ##### | -2.6298 | 8.5444E-03 | 1.7902E-02 |
| ENSG00000011021\|CLCN6 | 1283.7909 | -0.3569 | ##### | -2.6295 | 8.5521E-03 | 1.7915E-02 |
| ENSG00000259546 | 5.6968 | -4.3576 | ##### | -2.6295 | 8.5520E-03 | 1.7915E-02 |
| ENSG00000230729 | 10.1813 | 1.5968 | ##### | 2.6295 | 8.5518E-03 | 1.7915E-02 |
| ENSG00000106245\|BUD31 | 6723.0283 | 0.3268 | ##### | 2.6287 | 8.5720E-03 | 1.7955E-02 |
| ENSG00000291300\|PRSS30P | 44.2489 | 2.3432 | ##### | 2.6285 | 8.5761E-03 | 1.7962E-02 |
| ENSG00000105996\|HOXA2 | 22.0082 | 1.8700 | ##### | 2.6284 | 8.5778E-03 | 1.7963E-02 |
| ENSG00000196150\|ZNF250 | 545.9810 | -0.3792 | ##### | -2.6285 | 8.5772E-03 | 1.7963E-02 |
| ENSG00000279592 | 126.7380 | -0.6358 | ##### | -2.6281 | 8.5856E-03 | 1.7978E-02 |
| ENSG00000141543\|EIF4A3 | 18786.5083 | -0.2288 | ##### | -2.6280 | 8.5884E-03 | 1.7983E-02 |
| ENSG00000169249\|ZRSR2 | 533.4921 | -0.4438 | ##### | -2.6280 | 8.5891E-03 | 1.7983E-02 |
| ENSG00000230162\|CT45A11P | 47.9165 | 8.4021 | ##### | 2.6280 | 8.5895E-03 | 1.7983E-02 |
| ENSG00000272721\|EIF2B5-DT | 8.8166 | -2.0848 | ##### | -2.6264 | 8.6301E-03 | 1.8067E-02 |
| ENSG00000293548 | 30.4451 | 1.4669 | ##### | 2.6260 | 8.6394E-03 | 1.8085E-02 |
| ENSG00000273302 | 19.4311 | 7.4964 | ##### | 2.6260 | 8.6399E-03 | 1.8085E-02 |
| ENSG00000116747\|RO60 | 3042.2605 | -0.3241 | ##### | -2.6259 | 8.6430E-03 | 1.8090E-02 |
| ENSG00000197561\|ELANE | 25.2484 | -2.5065 | ##### | -2.6256 | 8.6489E-03 | 1.8101E-02 |
| ENSG00000108684\|ASIC2 | 130.6191 | -0.6521 | ##### | -2.6251 | 8.6619E-03 | 1.8127E-02 |
| ENSG00000293256\|CBX3P2 | 31.5453 | 1.5923 | ##### | 2.6249 | 8.6666E-03 | 1.8135E-02 |
| ENSG00000184831\|APOO | 1402.2242 | 0.3558 | ##### | 2.6246 | 8.6759E-03 | 1.8153E-02 |
| ENSG00000136444\|RSAD1 | 3287.2585 | 0.3238 | ##### | 2.6244 | 8.6805E-03 | 1.8162E-02 |
| ENSG00000284882 | 11.4076 | -1.5978 | ##### | -2.6238 | 8.6949E-03 | 1.8191E-02 |
| ENSG00000230148\|HOXB-AS1 | 34.1785 | 1.0415 | ##### | 2.6237 | 8.6968E-03 | 1.8194E-02 |
| ENSG00000088038\|CNOT3 | 8322.5594 | 0.2896 | ##### | 2.6237 | 8.6978E-03 | 1.8194E-02 |
| ENSG00000257103\|LSM14A | 10245.6267 | -0.3207 | ##### | -2.6234 | 8.7052E-03 | 1.8208E-02 |
| ENSG00000149499\|EML3 | 3412.6006 | 0.3230 | ##### | 2.6232 | 8.7099E-03 | 1.8217E-02 |
| ENSG00000125967\|NECAB3 | 2140.0346 | -0.3462 | ##### | -2.6219 | 8.7432E-03 | 1.8285E-02 |
| ENSG00000232938\|RPL23AP87 | 10.0341 | -2.5308 | ##### | -2.6217 | 8.7491E-03 | 1.8296E-02 |
| ENSG00000233117\|LINC00702 | 26.4430 | 5.7573 | ##### | 2.6206 | 8.7765E-03 | 1.8352E-02 |
| ENSG00000181744\|DIPK2A | 1422.4785 | -0.3390 | ##### | -2.6200 | 8.7923E-03 | 1.8384E-02 |
| ENSG00000146411\|SLC2A12 | 47.7686 | -0.7714 | ##### | -2.6191 | 8.8161E-03 | 1.8433E-02 |
| ENSG00000291171 | 1391.2830 | 0.3946 | ##### | 2.6189 | 8.8202E-03 | 1.8440E-02 |
| ENSG00000251667\|BRCC3P1 | 26.8773 | 0.9798 | ##### | 2.6187 | 8.8255E-03 | 1.8450E-02 |
| ENSG00000267282 | 8.4097 | -5.0886 | ##### | -2.6184 | 8.8353E-03 | 1.8469E-02 |
| ENSG00000259705\|FBN1-DT | 14.6169 | -2.9842 | ##### | -2.6183 | 8.8364E-03 | 1.8470E-02 |
| ENSG00000173213\|TUBB8B | 10.0406 | 1.9291 | ##### | 2.6182 | 8.8397E-03 | 1.8476E-02 |
| ENSG00000274270 | 41.7822 | 0.9135 | ##### | 2.6181 | 8.8434E-03 | 1.8482E-02 |
| ENSG00000149548\|CCDC15 | 215.1336 | 0.5359 | ##### | 2.6166 | 8.8807E-03 | 1.8559E-02 |
| ENSG00000273777\|CEACAM20 | 24.3812 | -8.7187 | ##### | -2.6162 | 8.8918E-03 | 1.8581E-02 |
| ENSG00000260772 | 18.0486 | 1.1982 | ##### | 2.6155 | 8.9100E-03 | 1.8617E-02 |
| ENSG00000212789\|ST13P5 | 16.6203 | -1.1605 | ##### | -2.6155 | 8.9108E-03 | 1.8618E-02 |
| ENSG00000112218\|GPR63 | 522.1016 | 0.3891 | ##### | 2.6154 | 8.9120E-03 | 1.8619E-02 |
| ENSG00000288770 | 11.5177 | -2.5456 | ##### | -2.6148 | 8.9276E-03 | 1.8650E-02 |
| ENSG00000185105\|MYADML2 | 16.7601 | 1.8976 | ##### | 2.6148 | 8.9284E-03 | 1.8650E-02 |
| ENSG00000198042\|MAK16 | 4285.6916 | 0.3235 | ##### | 2.6148 | 8.9290E-03 | 1.8650E-02 |
| ENSG00000172081\|MOB3A | 2318.4140 | 0.3298 | ##### | 2.6145 | 8.9349E-03 | 1.8661E-02 |
| ENSG00000243323\|PTPRVP | 33.2515 | -8.1951 | ##### | -2.6144 | 8.9395E-03 | 1.8670E-02 |

| | | | | | | |
|---|---|---|---|---|---|---|
| ENSG00000210191\|MT-TL2 | 109.9644 | 0.6098 | ##### | 2.6141 | 8.9468E-03 | 1.8684E-02 |
| ENSG00000179523\|EIF3J-DT | 341.0723 | -0.4096 | ##### | -2.6137 | 8.9566E-03 | 1.8703E-02 |
| ENSG00000179774\|ATOH7 | 17.3050 | -2.7529 | ##### | -2.6135 | 8.9626E-03 | 1.8714E-02 |
| ENSG00000231560\|CLEC12A-AS1 | 6.2770 | 4.4045 | ##### | 2.6133 | 8.9662E-03 | 1.8720E-02 |
| ENSG00000270574 | 59.4203 | -0.7684 | ##### | -2.6133 | 8.9666E-03 | 1.8720E-02 |
| ENSG00000101440\|ASIP | 9.3977 | 2.1978 | ##### | 2.6128 | 8.9805E-03 | 1.8748E-02 |
| ENSG00000186056\|MATN1-AS1 | 216.4232 | -0.4951 | ##### | -2.6127 | 8.9840E-03 | 1.8754E-02 |
| ENSG00000243896\|OR2A7 | 30.8901 | -0.9838 | ##### | -2.6120 | 9.0027E-03 | 1.8791E-02 |
| ENSG00000284469\|MIR6721 | 4.9247 | 1.9519 | ##### | 2.6119 | 9.0045E-03 | 1.8794E-02 |
| ENSG00000207185\|RNU6-1157P | 3.6532 | 4.3363 | ##### | 2.6114 | 9.0174E-03 | 1.8819E-02 |
| ENSG00000233837\|EIF3LP2 | 10.0053 | -1.6393 | ##### | -2.6104 | 9.0436E-03 | 1.8873E-02 |
| ENSG00000267205 | 9.4422 | 2.2727 | ##### | 2.6101 | 9.0513E-03 | 1.8887E-02 |
| ENSG00000120594\|PLXDC2 | 91.4606 | -0.5759 | ##### | -2.6095 | 9.0669E-03 | 1.8918E-02 |
| ENSG00000278876 | 16.3999 | 1.7089 | ##### | 2.6095 | 9.0674E-03 | 1.8918E-02 |
| ENSG00000076604\|TRAF4 | 9363.3726 | -0.3193 | ##### | -2.6091 | 9.0773E-03 | 1.8938E-02 |
| ENSG00000232422\|KNOP1P4 | 11.6135 | 4.9520 | ##### | 2.6087 | 9.0884E-03 | 1.8960E-02 |
| ENSG00000064995\|TAF11 | 1755.8560 | -0.3443 | ##### | -2.6086 | 9.0902E-03 | 1.8962E-02 |
| ENSG00000183814\|LIN9 | 1631.9734 | 0.3499 | ##### | 2.6085 | 9.0949E-03 | 1.8971E-02 |
| ENSG00000085978\|ATG16L1 | 6415.0345 | -0.2397 | ##### | -2.6068 | 9.1388E-03 | 1.9059E-02 |
| ENSG00000236591\|UST-AS2 | 6.8352 | 3.1720 | ##### | 2.6068 | 9.1388E-03 | 1.9059E-02 |
| ENSG00000273145 | 69.5699 | -0.8693 | ##### | -2.6063 | 9.1520E-03 | 1.9086E-02 |
| ENSG00000142082\|SIRT3 | 967.5897 | -0.3548 | ##### | -2.6058 | 9.1648E-03 | 1.9111E-02 |
| ENSG00000233330 | 17.8771 | 2.0565 | ##### | 2.6057 | 9.1683E-03 | 1.9117E-02 |
| ENSG00000183648\|NDUFB1 | 1037.3549 | -0.3599 | ##### | -2.6052 | 9.1823E-03 | 1.9143E-02 |
| ENSG00000108591\|DRG2 | 3124.2075 | 0.3305 | ##### | 2.6052 | 9.1821E-03 | 1.9143E-02 |
| ENSG00000272779\|BMS1P20 | 383.5053 | 0.4248 | ##### | 2.6051 | 9.1841E-03 | 1.9146E-02 |
| ENSG00000260466 | 8.5146 | -3.1526 | ##### | -2.6047 | 9.1948E-03 | 1.9166E-02 |
| ENSG00000236039\|LINC02889 | 19.2989 | -1.8789 | ##### | -2.6047 | 9.1948E-03 | 1.9166E-02 |
| ENSG00000272970 | 29.9229 | -1.4933 | ##### | -2.6047 | 9.1956E-03 | 1.9166E-02 |
| ENSG00000284634 | 86.6819 | 1.0489 | ##### | 2.6044 | 9.2035E-03 | 1.9181E-02 |
| ENSG00000286018 | 49.9399 | 0.6904 | ##### | 2.6042 | 9.2082E-03 | 1.9189E-02 |
| ENSG00000183508\|TENT5C | 65.1086 | -0.9847 | ##### | -2.6033 | 9.2335E-03 | 1.9238E-02 |
| ENSG00000267515 | 4.7174 | 5.1372 | ##### | 2.6033 | 9.2326E-03 | 1.9238E-02 |
| ENSG00000134313\|KIDINS220 | 4835.9849 | -0.3107 | ##### | -2.6033 | 9.2331E-03 | 1.9238E-02 |
| ENSG00000238286\|SLC35E1P1 | 16.4533 | 1.8581 | ##### | 2.6032 | 9.2354E-03 | 1.9241E-02 |
| ENSG00000198130\|HIBCH | 2742.1201 | 0.3135 | ##### | 2.6031 | 9.2390E-03 | 1.9244E-02 |
| ENSG00000253333 | 10.5130 | 3.1323 | ##### | 2.6031 | 9.2383E-03 | 1.9244E-02 |
| ENSG00000113248\|PCDHB15 | 12.1661 | 1.7364 | ##### | 2.6031 | 9.2379E-03 | 1.9244E-02 |
| ENSG00000123106\|CCDC91 | 2724.3957 | 0.3187 | ##### | 2.6031 | 9.2397E-03 | 1.9244E-02 |
| ENSG00000279762 | 23.1917 | 1.0332 | ##### | 2.6029 | 9.2439E-03 | 1.9252E-02 |
| ENSG00000181260\|MTHFD2P7 | 15.7065 | 1.1000 | ##### | 2.6028 | 9.2461E-03 | 1.9255E-02 |
| ENSG00000291056\|SEPTIN7P9 | 7.3302 | -2.3623 | ##### | -2.6027 | 9.2504E-03 | 1.9263E-02 |
| ENSG00000240970\|RPL23AP64 | 11.0266 | 1.4854 | ##### | 2.6019 | 9.2718E-03 | 1.9304E-02 |
| ENSG00000260017 | 4.4774 | -5.3003 | ##### | -2.6019 | 9.2718E-03 | 1.9304E-02 |
| ENSG00000182518\|FAM104B | 564.4923 | -0.3855 | ##### | -2.6015 | 9.2822E-03 | 1.9325E-02 |
| ENSG00000073921\|PICALM | 10157.1215 | -0.2984 | ##### | -2.6014 | 9.2850E-03 | 1.9329E-02 |
| ENSG00000187902\|SHISA7 | 25.0824 | 2.0858 | ##### | 2.6013 | 9.2880E-03 | 1.9334E-02 |
| ENSG00000215630\|GUSBP9 | 47.1627 | 1.4726 | ##### | 2.6011 | 9.2931E-03 | 1.9343E-02 |
| ENSG00000004799\|PDK4 | 2748.9268 | -0.3589 | ##### | -2.6006 | 9.3052E-03 | 1.9367E-02 |
| ENSG00000273679 | 125.3671 | 0.5446 | ##### | 2.5997 | 9.3316E-03 | 1.9421E-02 |
| ENSG00000272848 | 13.6021 | 1.9581 | ##### | 2.5990 | 9.3499E-03 | 1.9458E-02 |
| ENSG00000230424\|EMC1-AS1 | 21.1075 | -1.0712 | ##### | -2.5986 | 9.3618E-03 | 1.9481E-02 |
| ENSG00000274460 | 63.6399 | 0.8395 | ##### | 2.5980 | 9.3760E-03 | 1.9509E-02 |
| ENSG00000281128\|PTENP1-AS | 40.0306 | -1.0989 | ##### | -2.5971 | 9.4014E-03 | 1.9561E-02 |
| ENSG00000160172\|FAM86C2P | 34.4905 | 1.1491 | ##### | 2.5967 | 9.4121E-03 | 1.9581E-02 |
| ENSG00000138138\|ATAD1 | 5932.6543 | 0.3197 | ##### | 2.5955 | 9.4442E-03 | 1.9647E-02 |
| ENSG00000178395\|CCDC185 | 33.7997 | 8.1252 | ##### | 2.5951 | 9.4570E-03 | 1.9671E-02 |
| ENSG00000201496\|RN7SKP275 | 5.6292 | 2.2512 | ##### | 2.5951 | 9.4567E-03 | 1.9671E-02 |
| ENSG00000249673\|NOP14-AS1 | 1297.7368 | 0.3446 | ##### | 2.5936 | 9.4979E-03 | 1.9752E-02 |
| ENSG00000261366\|MANEA-DT | 15.8091 | -1.5735 | ##### | -2.5936 | 9.4967E-03 | 1.9752E-02 |
| ENSG00000249156\|TAF11L12 | 6.1861 | -6.0364 | ##### | -2.5936 | 9.4976E-03 | 1.9752E-02 |
| ENSG00000219814\|RPL23AP47 | 7.5736 | 1.4931 | ##### | 2.5935 | 9.4991E-03 | 1.9753E-02 |
| ENSG00000234917 | 14.5055 | 1.7055 | ##### | 2.5930 | 9.5150E-03 | 1.9783E-02 |

| | | | | | | |
|---|---|---|---|---|---|---|
| ENSG00000273017 | 7.4150 | -2.5149 | ##### | -2.5930 | 9.5145E-03 | 1.9783E-02 |
| ENSG00000265840 | 74.5142 | -0.7050 | ##### | -2.5927 | 9.5226E-03 | 1.9798E-02 |
| ENSG00000116205\|TCEANC2 | 1228.0823 | -0.3422 | ##### | -2.5926 | 9.5247E-03 | 1.9801E-02 |
| ENSG00000136319\|TTC5 | 1485.0940 | 0.3390 | ##### | 2.5923 | 9.5332E-03 | 1.9817E-02 |
| ENSG00000280047 | 65.7715 | -0.8313 | ##### | -2.5920 | 9.5425E-03 | 1.9835E-02 |
| ENSG00000187607\|ZNF286A | 1446.3272 | 0.3246 | ##### | 2.5917 | 9.5509E-03 | 1.9851E-02 |
| ENSG00000287727 | 21.1204 | -7.6678 | ##### | -2.5915 | 9.5557E-03 | 1.9860E-02 |
| ENSG00000051382\|PIK3CB | 4003.1726 | -0.3034 | ##### | -2.5913 | 9.5627E-03 | 1.9873E-02 |
| ENSG00000254662 | 11.7158 | 1.8008 | ##### | 2.5903 | 9.5894E-03 | 1.9927E-02 |
| ENSG00000225071\|RPS26P58 | 143.2681 | 0.5112 | ##### | 2.5898 | 9.6034E-03 | 1.9954E-02 |
| ENSG00000274428\|RNVU1-25 | 5.5856 | 4.9234 | ##### | 2.5894 | 9.6131E-03 | 1.9973E-02 |
| ENSG00000291200\|RHOXF1P1 | 14.9923 | 2.8251 | ##### | 2.5893 | 9.6164E-03 | 1.9979E-02 |
| ENSG00000260704\|LINC00543 | 35.6399 | -1.4219 | ##### | -2.5893 | 9.6173E-03 | 1.9979E-02 |
| ENSG00000118690\|ARMC2 | 173.8939 | 0.4915 | ##### | 2.5891 | 9.6240E-03 | 1.9992E-02 |
| ENSG00000082258\|CCNT2 | 3264.1792 | -0.3270 | ##### | -2.5884 | 9.6427E-03 | 2.0029E-02 |
| ENSG00000112837\|TBX18 | 1896.0351 | -0.3368 | ##### | -2.5883 | 9.6462E-03 | 2.0035E-02 |
| ENSG00000285865 | 11.7027 | 5.5105 | ##### | 2.5881 | 9.6498E-03 | 2.0041E-02 |
| ENSG00000222020\|HDAC4-AS1 | 24.0490 | -1.1446 | ##### | -2.5880 | 9.6527E-03 | 2.0046E-02 |
| ENSG00000290800\|TBC1D3P2 | 5.9137 | 5.6697 | ##### | 2.5879 | 9.6574E-03 | 2.0053E-02 |
| ENSG00000293328\|NKAPP1 | 66.6198 | 0.7681 | ##### | 2.5879 | 9.6572E-03 | 2.0053E-02 |
| ENSG00000255126 | 68.6912 | 0.7394 | ##### | 2.5876 | 9.6635E-03 | 2.0064E-02 |
| ENSG00000163788\|SNRK | 1491.3486 | 0.3375 | ##### | 2.5875 | 9.6668E-03 | 2.0069E-02 |
| ENSG00000274751 | 5.6289 | 2.4928 | ##### | 2.5868 | 9.6875E-03 | 2.0111E-02 |
| ENSG00000182195\|LDOC1 | 4555.7305 | 0.3226 | ##### | 2.5865 | 9.6969E-03 | 2.0129E-02 |
| ENSG00000249209 | 8.7972 | -2.9288 | ##### | -2.5861 | 9.7070E-03 | 2.0149E-02 |
| ENSG00000275620\|FLJ16779 | 93.4196 | -1.6328 | ##### | -2.5860 | 9.7093E-03 | 2.0152E-02 |
| ENSG00000106723\|SPIN1 | 7669.5398 | -0.3119 | ##### | -2.5856 | 9.7214E-03 | 2.0176E-02 |
| ENSG00000213578\|CPLX3 | 22.4674 | 8.6161 | ##### | 2.5854 | 9.7259E-03 | 2.0184E-02 |
| ENSG00000079974\|RABL2B | 1556.7542 | -0.3282 | ##### | -2.5854 | 9.7275E-03 | 2.0186E-02 |
| ENSG00000268947 | 10.4838 | 1.8180 | ##### | 2.5852 | 9.7334E-03 | 2.0197E-02 |
| ENSG00000178403\|NEUROG2 | 9.8963 | 2.7015 | ##### | 2.5851 | 9.7361E-03 | 2.0201E-02 |
| ENSG00000141748\|ARL5C | 10.6422 | 1.8015 | ##### | 2.5850 | 9.7381E-03 | 2.0203E-02 |
| ENSG00000263394 | 4.6244 | 4.2532 | ##### | 2.5846 | 9.7504E-03 | 2.0228E-02 |
| ENSG00000212396\|RNA5SP323 | 2.6725 | -4.1776 | ##### | -2.5842 | 9.7614E-03 | 2.0249E-02 |
| ENSG00000112685\|EXOC2 | 4453.1290 | 0.3085 | ##### | 2.5836 | 9.7768E-03 | 2.0279E-02 |
| ENSG00000062582\|MRPS24 | 7868.8528 | 0.3080 | ##### | 2.5836 | 9.7774E-03 | 2.0279E-02 |
| ENSG00000175463\|TBC1D10C | 33.2380 | 1.9622 | ##### | 2.5830 | 9.7947E-03 | 2.0314E-02 |
| ENSG00000257341 | 13.1726 | 2.7333 | ##### | 2.5827 | 9.8039E-03 | 2.0331E-02 |
| ENSG00000213860\|RPL21P75 | 2925.8294 | 0.3324 | ##### | 2.5819 | 9.8249E-03 | 2.0374E-02 |
| ENSG00000073969\|NSF | 7541.9974 | 0.2176 | ##### | 2.5819 | 9.8271E-03 | 2.0377E-02 |
| ENSG00000102760\|RGCC | 37.5347 | 0.8772 | ##### | 2.5817 | 9.8306E-03 | 2.0383E-02 |
| ENSG00000273254 | 5.5417 | -2.7490 | ##### | -2.5816 | 9.8346E-03 | 2.0389E-02 |
| ENSG00000226919 | 23.8918 | -2.5830 | ##### | -2.5815 | 9.8379E-03 | 2.0395E-02 |
| ENSG00000011405\|PIK3C2A | 4187.7610 | 0.3204 | ##### | 2.5813 | 9.8443E-03 | 2.0407E-02 |
| ENSG00000114166\|KAT2B | 1005.3352 | -0.3586 | ##### | -2.5809 | 9.8531E-03 | 2.0424E-02 |
| ENSG00000185917\|SETD4 | 1171.5657 | 0.3571 | ##### | 2.5806 | 9.8621E-03 | 2.0441E-02 |
| ENSG00000198000\|NOL8 | 4359.4242 | 0.3011 | ##### | 2.5804 | 9.8677E-03 | 2.0451E-02 |
| ENSG00000243389 | 18.5839 | -1.0045 | ##### | -2.5804 | 9.8698E-03 | 2.0454E-02 |
| ENSG00000236723 | 98.5089 | -0.5829 | ##### | -2.5803 | 9.8707E-03 | 2.0454E-02 |
| ENSG00000234224\|TMEM229A | 2.9399 | -5.0709 | ##### | -2.5800 | 9.8807E-03 | 2.0474E-02 |
| ENSG00000267426 | 667.6486 | 0.9126 | ##### | 2.5796 | 9.8901E-03 | 2.0492E-02 |
| ENSG00000283541 | 7.9532 | -2.6114 | ##### | -2.5791 | 9.9052E-03 | 2.0521E-02 |
| ENSG00000001626\|CFTR | 40.1698 | 1.7692 | ##### | 2.5786 | 9.9198E-03 | 2.0550E-02 |
| ENSG00000280367 | 18.7873 | 1.3073 | ##### | 2.5782 | 9.9311E-03 | 2.0572E-02 |
| ENSG00000064490\|RFXANK | 3341.9969 | 0.3279 | ##### | 2.5780 | 9.9365E-03 | 2.0582E-02 |
| ENSG00000272742 | 440.4500 | 0.4052 | ##### | 2.5775 | 9.9506E-03 | 2.0610E-02 |
| ENSG00000151726\|ACSL1 | 1631.7038 | 0.3405 | ##### | 2.5774 | 9.9552E-03 | 2.0618E-02 |
| ENSG00000104044\|OCA2 | 71.3080 | -0.6961 | ##### | -2.5772 | 9.9601E-03 | 2.0627E-02 |
| ENSG00000117155\|SSX2IP | 5514.7636 | 0.3009 | ##### | 2.5769 | 9.9699E-03 | 2.0644E-02 |
| ENSG00000179564\|LSMEM2 | 652.3270 | 0.3816 | ##### | 2.5769 | 9.9700E-03 | 2.0644E-02 |
| ENSG00000198488\|B3GNT6 | 25.2138 | -8.5840 | ##### | -2.5757 | 1.0004E-02 | 2.0711E-02 |
| ENSG00000160396\|HIPK4 | 10.2891 | -1.5971 | ##### | -2.5757 | 1.0003E-02 | 2.0711E-02 |
| ENSG00000102221\|JADE3 | 1556.1717 | -0.3303 | ##### | -2.5755 | 1.0009E-02 | 2.0720E-02 |

| | | | | | | |
|---|---|---|---|---|---|---|
| ENSG00000255328 | 6.0023 | 5.2228 | ##### | 2.5754 | 1.0013E-02 | 2.0728E-02 |
| ENSG00000243709\|LEFTY1 | 11.9543 | -2.6320 | ##### | -2.5750 | 1.0024E-02 | 2.0749E-02 |
| ENSG00000258674 | 370.4810 | -0.7898 | ##### | -2.5750 | 1.0025E-02 | 2.0749E-02 |
| ENSG00000130348\|QRSL1 | 2486.0986 | -0.3201 | ##### | -2.5744 | 1.0043E-02 | 2.0785E-02 |
| ENSG00000260604 | 100.8260 | -0.5742 | ##### | -2.5739 | 1.0055E-02 | 2.0809E-02 |
| ENSG00000181938\|GINS3 | 1973.5430 | 0.3238 | ##### | 2.5733 | 1.0073E-02 | 2.0845E-02 |
| ENSG00000146223\|RPL7L1 | 21976.2160 | -0.3019 | ##### | -2.5733 | 1.0074E-02 | 2.0846E-02 |
| ENSG00000236307\|EEF1E1P1 | 16.8010 | -1.4835 | ##### | -2.5731 | 1.0079E-02 | 2.0855E-02 |
| ENSG00000182310\|SPACA6 | 472.3066 | -0.3817 | ##### | -2.5726 | 1.0094E-02 | 2.0885E-02 |
| ENSG00000049618\|ARID1B | 4856.5961 | 0.3113 | ##### | 2.5724 | 1.0098E-02 | 2.0892E-02 |
| ENSG00000249319 | 36.0055 | -4.1410 | ##### | -2.5717 | 1.0121E-02 | 2.0937E-02 |
| ENSG00000260325\|HSPB9 | 9.0541 | 2.7573 | ##### | 2.5715 | 1.0127E-02 | 2.0947E-02 |
| ENSG00000143147\|GPR161 | 1045.1197 | -0.3313 | ##### | -2.5714 | 1.0129E-02 | 2.0951E-02 |
| ENSG00000212283\|SNORD89 | 6.9795 | -1.4819 | ##### | -2.5711 | 1.0136E-02 | 2.0964E-02 |
| ENSG00000198515\|CNGA1 | 57.5000 | -0.8340 | ##### | -2.5711 | 1.0139E-02 | 2.0967E-02 |
| ENSG00000100271\|TTLL1 | 301.5182 | -0.4177 | ##### | -2.5709 | 1.0142E-02 | 2.0973E-02 |
| ENSG00000165731\|RET | 49.7802 | 0.7216 | ##### | 2.5707 | 1.0149E-02 | 2.0986E-02 |
| ENSG00000154478\|GPR26 | 9.4811 | -5.4771 | ##### | -2.5706 | 1.0151E-02 | 2.0989E-02 |
| ENSG00000257918 | 24.0479 | 1.8268 | ##### | 2.5705 | 1.0156E-02 | 2.0998E-02 |
| ENSG00000204950\|LRRC10B | 28.6138 | 1.4186 | ##### | 2.5702 | 1.0163E-02 | 2.1010E-02 |
| ENSG00000251870\|RNU2-69P | 3.4513 | -4.6376 | ##### | -2.5699 | 1.0172E-02 | 2.1028E-02 |
| ENSG00000290875\|GLUD1P2 | 11.3934 | -1.7782 | ##### | -2.5692 | 1.0192E-02 | 2.1067E-02 |
| ENSG00000198162\|MAN1A2 | 6330.3073 | -0.3148 | ##### | -2.5691 | 1.0196E-02 | 2.1075E-02 |
| ENSG00000249684 | 69.2329 | 0.9229 | ##### | 2.5689 | 1.0201E-02 | 2.1083E-02 |
| ENSG00000151617\|EDNRA | 25.2580 | -2.9840 | ##### | -2.5687 | 1.0208E-02 | 2.1097E-02 |
| ENSG00000227933 | 9.4997 | 4.2888 | ##### | 2.5684 | 1.0216E-02 | 2.1110E-02 |
| ENSG00000079246\|XRCC5 | 41875.9086 | 0.2473 | ##### | 2.5678 | 1.0234E-02 | 2.1147E-02 |
| ENSG00000163636\|PSMD6 | 4933.7039 | -0.3157 | ##### | -2.5678 | 1.0234E-02 | 2.1147E-02 |
| ENSG00000285367 | 41.8755 | -4.7116 | ##### | -2.5677 | 1.0237E-02 | 2.1151E-02 |
| ENSG00000226754\|MAGOH-DT | 108.4468 | -0.8008 | ##### | -2.5673 | 1.0248E-02 | 2.1172E-02 |
| ENSG00000286075\|SAGSIN1 | 2083.1590 | -0.3198 | ##### | -2.5671 | 1.0255E-02 | 2.1184E-02 |
| ENSG00000141756\|FKBP10 | 16979.0803 | -0.2977 | ##### | -2.5669 | 1.0260E-02 | 2.1194E-02 |
| ENSG00000227091\|SLC6A17-AS1 | 16.6352 | 1.4405 | ##### | 2.5669 | 1.0262E-02 | 2.1195E-02 |
| ENSG00000177238\|TRIM72 | 31.2954 | 5.2100 | ##### | 2.5669 | 1.0262E-02 | 2.1195E-02 |
| ENSG00000170373\|CST1 | 3.5216 | -5.5680 | ##### | -2.5666 | 1.0269E-02 | 2.1208E-02 |
| ENSG00000260941\|LINC00622 | 15.9719 | 2.0915 | ##### | 2.5666 | 1.0270E-02 | 2.1208E-02 |
| ENSG00000235453\|SMIM27 | 111.3253 | -0.7040 | ##### | -2.5664 | 1.0276E-02 | 2.1219E-02 |
| ENSG00000109320\|NFKB1 | 6667.7118 | 0.3044 | ##### | 2.5656 | 1.0298E-02 | 2.1264E-02 |
| ENSG00000131127\|ZNF141 | 734.6905 | 0.3585 | ##### | 2.5655 | 1.0302E-02 | 2.1270E-02 |
| ENSG00000265121 | 6.4808 | -5.8531 | ##### | -2.5655 | 1.0303E-02 | 2.1271E-02 |
| ENSG00000279088 | 132.0669 | -0.5119 | ##### | -2.5646 | 1.0329E-02 | 2.1322E-02 |
| ENSG00000100601\|ALKBH1 | 1196.4944 | 0.3539 | ##### | 2.5645 | 1.0332E-02 | 2.1328E-02 |
| ENSG00000143748\|NVL | 2948.5724 | 0.3074 | ##### | 2.5640 | 1.0346E-02 | 2.1355E-02 |
| ENSG00000174672\|BRSK2 | 417.2707 | -0.4124 | ##### | -2.5637 | 1.0356E-02 | 2.1373E-02 |
| ENSG00000277971 | 14.9645 | -1.0718 | ##### | -2.5633 | 1.0369E-02 | 2.1399E-02 |
| ENSG00000162885\|B3GALNT2 | 4150.3657 | -0.3299 | ##### | -2.5632 | 1.0371E-02 | 2.1400E-02 |
| ENSG00000180881\|CAPS2 | 282.7623 | -0.4510 | ##### | -2.5632 | 1.0372E-02 | 2.1400E-02 |
| ENSG00000119446\|RBM18 | 3132.3976 | -0.2645 | ##### | -2.5632 | 1.0371E-02 | 2.1400E-02 |
| ENSG00000164967\|RPP25L | 1141.2439 | 0.3249 | ##### | 2.5631 | 1.0375E-02 | 2.1406E-02 |
| ENSG00000267391\|MIR122HG | 7.2035 | -5.2193 | ##### | -2.5630 | 1.0376E-02 | 2.1407E-02 |
| ENSG00000015171\|ZMYND11 | 7461.4165 | -0.2329 | ##### | -2.5629 | 1.0381E-02 | 2.1413E-02 |
| ENSG00000088832\|FKBP1A | 18975.8953 | 0.3194 | ##### | 2.5629 | 1.0380E-02 | 2.1413E-02 |
| ENSG00000065485\|PDIA5 | 2771.5725 | 0.3129 | ##### | 2.5621 | 1.0403E-02 | 2.1458E-02 |
| ENSG00000261051 | 12.3080 | -2.3634 | ##### | -2.5619 | 1.0412E-02 | 2.1474E-02 |
| ENSG00000274835 | 4.8365 | -5.6557 | ##### | -2.5614 | 1.0425E-02 | 2.1499E-02 |
| ENSG00000267705 | 5.5080 | 2.0289 | ##### | 2.5611 | 1.0434E-02 | 2.1516E-02 |
| ENSG00000147050\|KDM6A | 1739.5060 | -0.3211 | ##### | -2.5611 | 1.0435E-02 | 2.1518E-02 |
| ENSG00000227630\|LINC01132 | 14.7586 | 2.6977 | ##### | 2.5608 | 1.0442E-02 | 2.1531E-02 |
| ENSG00000175189\|INHBC | 16.3785 | 2.6412 | ##### | 2.5602 | 1.0460E-02 | 2.1566E-02 |
| ENSG00000157064\|NMNAT2 | 260.9593 | -0.5274 | ##### | -2.5602 | 1.0462E-02 | 2.1568E-02 |
| ENSG00000213194\|CDK7P1 | 4.3000 | 4.1303 | ##### | 2.5602 | 1.0462E-02 | 2.1568E-02 |
| ENSG00000230916\|MTCO1P53 | 6.1728 | 1.6175 | ##### | 2.5598 | 1.0474E-02 | 2.1591E-02 |
| ENSG00000100429\|HDAC10 | 709.4132 | -0.3747 | ##### | -2.5597 | 1.0476E-02 | 2.1593E-02 |

| | | | | | | |
|---|---|---|---|---|---|---|
| ENSG00000160957\|RECQL4 | 4853.3188 | -0.3035 | ##### | -2.5591 | 1.0495E-02 | 2.1631E-02 |
| ENSG00000007168\|PAFAH1B1 | 14143.3483 | 0.2938 | ##### | 2.5588 | 1.0502E-02 | 2.1644E-02 |
| ENSG00000196605\|ZNF846 | 130.8291 | -0.5443 | ##### | -2.5588 | 1.0505E-02 | 2.1648E-02 |
| ENSG00000202408\|RNVU1-21 | 5.0136 | 5.0466 | ##### | 2.5569 | 1.0562E-02 | 2.1766E-02 |
| ENSG00000138668\|HNRNPD | 19222.4097 | 0.3053 | ##### | 2.5562 | 1.0582E-02 | 2.1804E-02 |
| ENSG00000293151 | 4.6963 | 3.4662 | ##### | 2.5560 | 1.0588E-02 | 2.1815E-02 |
| ENSG00000100522\|GNPNAT1 | 12545.0401 | 0.3183 | ##### | 2.5559 | 1.0592E-02 | 2.1822E-02 |
| ENSG00000267416\|HEATR6-DT | 26.3803 | -1.1665 | ##### | -2.5558 | 1.0593E-02 | 2.1823E-02 |
| ENSG00000198951\|NAGA | 2131.5294 | 0.3288 | ##### | 2.5558 | 1.0595E-02 | 2.1824E-02 |
| ENSG00000261611 | 222.1055 | -2.5529 | ##### | -2.5557 | 1.0597E-02 | 2.1829E-02 |
| ENSG00000198835\|GJC2 | 95.9565 | 0.5858 | ##### | 2.5555 | 1.0605E-02 | 2.1842E-02 |
| ENSG00000114439\|BBX | 7379.0992 | 0.3096 | ##### | 2.5549 | 1.0621E-02 | 2.1875E-02 |
| ENSG00000113318\|MSH3 | 1405.0949 | -0.3369 | ##### | -2.5541 | 1.0645E-02 | 2.1922E-02 |
| ENSG00000128609\|NDUFA5 | 4351.7707 | 0.2696 | ##### | 2.5541 | 1.0646E-02 | 2.1922E-02 |
| ENSG00000273333 | 138.0723 | 2.3952 | ##### | 2.5538 | 1.0654E-02 | 2.1938E-02 |
| ENSG00000186340\|THBS2 | 10.8164 | -3.0799 | ##### | -2.5527 | 1.0689E-02 | 2.2009E-02 |
| ENSG00000169962\|TAS1R3 | 92.1595 | -0.6000 | ##### | -2.5524 | 1.0697E-02 | 2.2024E-02 |
| ENSG00000274928\|KRT89P | 6.1877 | -5.5383 | ##### | -2.5523 | 1.0700E-02 | 2.2028E-02 |
| ENSG00000253399 | 5.1702 | 4.5864 | ##### | 2.5520 | 1.0711E-02 | 2.2048E-02 |
| ENSG00000155660\|PDIA4 | 33446.3374 | 0.2359 | ##### | 2.5520 | 1.0711E-02 | 2.2048E-02 |
| ENSG00000152117\|SMPD4BP | 24.2838 | -1.5077 | ##### | -2.5516 | 1.0722E-02 | 2.2069E-02 |
| ENSG00000225439\|BOLA3-DT | 291.7924 | -0.4188 | ##### | -2.5513 | 1.0734E-02 | 2.2091E-02 |
| ENSG00000213714\|FAM209B | 19.9006 | 1.2447 | ##### | 2.5511 | 1.0738E-02 | 2.2096E-02 |
| ENSG00000226085\|UQCRFS1P1 | 253.0397 | -0.4165 | ##### | -2.5511 | 1.0737E-02 | 2.2096E-02 |
| ENSG00000069764\|PLA2G10 | 17.6373 | -1.7681 | ##### | -2.5505 | 1.0756E-02 | 2.2132E-02 |
| ENSG00000213442\|RPL18AP3 | 4350.0779 | 0.3126 | ##### | 2.5505 | 1.0757E-02 | 2.2133E-02 |
| ENSG00000260710 | 10.3236 | 4.2041 | ##### | 2.5495 | 1.0788E-02 | 2.2195E-02 |
| ENSG00000236213\|CDC42EP3-AS1 | 9.2401 | 1.9000 | ##### | 2.5495 | 1.0789E-02 | 2.2195E-02 |
| ENSG00000124772\|CPNE5 | 16.4718 | 2.4816 | ##### | 2.5492 | 1.0797E-02 | 2.2212E-02 |
| ENSG00000277182 | 108.1346 | -0.5788 | ##### | -2.5490 | 1.0803E-02 | 2.2221E-02 |
| ENSG00000176624\|MEX3C | 3883.3776 | 0.3135 | ##### | 2.5490 | 1.0804E-02 | 2.2223E-02 |
| ENSG00000285020 | 5.7763 | 2.6903 | ##### | 2.5488 | 1.0808E-02 | 2.2229E-02 |
| ENSG00000196981\|WDR5B | 748.9661 | -0.3600 | ##### | -2.5485 | 1.0819E-02 | 2.2251E-02 |
| ENSG00000285269 | 87.4062 | -4.3897 | ##### | -2.5479 | 1.0837E-02 | 2.2285E-02 |
| ENSG00000286447 | 30.5973 | 0.8591 | ##### | 2.5477 | 1.0842E-02 | 2.2293E-02 |
| ENSG00000175003\|SLC22A1 | 21.2151 | -3.6925 | ##### | -2.5477 | 1.0842E-02 | 2.2293E-02 |
| ENSG00000143632\|ACTA1 | 24.1721 | 1.0766 | ##### | 2.5476 | 1.0846E-02 | 2.2298E-02 |
| ENSG00000166855\|CLPX | 4168.3722 | 0.3144 | ##### | 2.5476 | 1.0846E-02 | 2.2298E-02 |
| ENSG00000240048\|DDX50P2 | 11.6144 | -3.6238 | ##### | -2.5459 | 1.0899E-02 | 2.2405E-02 |
| ENSG00000260500 | 17.8332 | -1.0748 | ##### | -2.5456 | 1.0909E-02 | 2.2424E-02 |
| ENSG00000239264\|TXNDC5 | 14278.5006 | 0.2388 | ##### | 2.5450 | 1.0927E-02 | 2.2460E-02 |
| ENSG00000112210\|RAB23 | 2331.7212 | -0.3437 | ##### | -2.5447 | 1.0938E-02 | 2.2480E-02 |
| ENSG00000227896\|WAPL-DT | 68.5247 | -0.8200 | ##### | -2.5443 | 1.0950E-02 | 2.2503E-02 |
| ENSG00000178878\|APOLD1 | 672.1946 | 0.3699 | ##### | 2.5442 | 1.0952E-02 | 2.2506E-02 |
| ENSG00000101194\|SLC17A9 | 18.6199 | 4.5554 | ##### | 2.5441 | 1.0957E-02 | 2.2515E-02 |
| ENSG00000258599 | 5.1660 | -5.2744 | ##### | -2.5438 | 1.0964E-02 | 2.2528E-02 |
| ENSG00000109771\|LRP2BP | 52.0145 | -0.8161 | ##### | -2.5437 | 1.0967E-02 | 2.2533E-02 |
| ENSG00000063854\|HAGH | 1797.0463 | -0.3208 | ##### | -2.5429 | 1.0993E-02 | 2.2585E-02 |
| ENSG00000168234\|TTC39C | 1953.3808 | 0.3178 | ##### | 2.5426 | 1.1003E-02 | 2.2603E-02 |
| ENSG00000267395\|DM1-AS | 98.9162 | -0.5708 | ##### | -2.5422 | 1.1015E-02 | 2.2625E-02 |
| ENSG00000164466\|SFXN1 | 8592.0760 | 0.3029 | ##### | 2.5420 | 1.1021E-02 | 2.2637E-02 |
| ENSG00000231203\|KRT8P10 | 11.0298 | -1.7419 | ##### | -2.5419 | 1.1026E-02 | 2.2646E-02 |
| ENSG00000223356 | 11.1625 | 2.0395 | ##### | 2.5410 | 1.1054E-02 | 2.2701E-02 |
| ENSG00000267317 | 473.3285 | -0.3994 | ##### | -2.5409 | 1.1057E-02 | 2.2707E-02 |
| ENSG00000163428\|LRRC58 | 7512.6797 | 0.3087 | ##### | 2.5403 | 1.1076E-02 | 2.2744E-02 |
| ENSG00000285852 | 27.4586 | 7.8252 | ##### | 2.5401 | 1.1083E-02 | 2.2756E-02 |
| ENSG00000187837\|H1-2 | 2957.1084 | -0.2881 | ##### | -2.5401 | 1.1083E-02 | 2.2756E-02 |
| ENSG00000262456 | 125.7504 | 0.6028 | ##### | 2.5398 | 1.1092E-02 | 2.2773E-02 |
| ENSG00000105048\|TNNT1 | 8110.4427 | 0.3010 | ##### | 2.5395 | 1.1103E-02 | 2.2792E-02 |
| ENSG00000102225\|CDK16 | 9581.7004 | -0.3084 | ##### | -2.5391 | 1.1115E-02 | 2.2817E-02 |
| ENSG00000268889\|LIM2-AS1 | 19.9502 | -1.0399 | ##### | -2.5389 | 1.1120E-02 | 2.2825E-02 |
| ENSG00000083535\|PIBF1 | 871.2571 | 0.3417 | ##### | 2.5388 | 1.1125E-02 | 2.2833E-02 |
| ENSG00000196104\|SPOCK3 | 14.5394 | 1.7607 | ##### | 2.5386 | 1.1129E-02 | 2.2840E-02 |

| | | | | | | |
|---|---|---|---|---|---|---|
| ENSG00000293091 | 12.9639 | -3.0430 | ##### | -2.5376 | 1.1160E-02 | 2.2903E-02 |
| ENSG00000182648\|RNF32-DT | 1096.3590 | -0.3379 | ##### | -2.5376 | 1.1162E-02 | 2.2905E-02 |
| ENSG00000238280 | 16.1160 | -1.1100 | ##### | -2.5367 | 1.1190E-02 | 2.2960E-02 |
| ENSG00000234420\|ZNF37BP | 4.3543 | -1.8915 | ##### | -2.5366 | 1.1192E-02 | 2.2963E-02 |
| ENSG00000143061\|IGSF3 | 4164.3125 | -0.3209 | ##### | -2.5351 | 1.1242E-02 | 2.3064E-02 |
| ENSG00000286606 | 23.1511 | 0.9570 | ##### | 2.5346 | 1.1259E-02 | 2.3098E-02 |
| ENSG00000050628\|PTGER3 | 52.2861 | 0.8919 | ##### | 2.5343 | 1.1268E-02 | 2.3114E-02 |
| ENSG00000172878\|METAP1D | 1071.9259 | -0.3247 | ##### | -2.5339 | 1.1279E-02 | 2.3135E-02 |
| ENSG00000289028 | 20.8516 | 1.3237 | ##### | 2.5338 | 1.1284E-02 | 2.3144E-02 |
| ENSG00000162194\|LBHD1 | 1065.4488 | -1.2810 | ##### | -2.5336 | 1.1289E-02 | 2.3151E-02 |
| ENSG00000235885\|LINC01828 | 10.6089 | 2.4540 | ##### | 2.5336 | 1.1289E-02 | 2.3151E-02 |
| ENSG00000177757\|FAM87B | 16.2833 | -2.6326 | ##### | -2.5334 | 1.1295E-02 | 2.3161E-02 |
| ENSG00000260830 | 15.2213 | 2.0324 | ##### | 2.5334 | 1.1296E-02 | 2.3162E-02 |
| ENSG00000205838\|TTC23L | 39.9848 | 0.8406 | ##### | 2.5332 | 1.1304E-02 | 2.3177E-02 |
| ENSG00000198873\|GRK5 | 577.3318 | -0.4149 | ##### | -2.5331 | 1.1307E-02 | 2.3182E-02 |
| ENSG00000137265\|IRF4 | 9.7664 | 2.3897 | ##### | 2.5328 | 1.1315E-02 | 2.3196E-02 |
| ENSG00000185085\|INTS5 | 2365.9491 | 0.3187 | ##### | 2.5324 | 1.1328E-02 | 2.3220E-02 |
| ENSG00000139597\|N4BP2L1 | 41.2107 | 0.7402 | ##### | 2.5322 | 1.1336E-02 | 2.3236E-02 |
| ENSG00000166016\|ABTB2 | 3131.8081 | 0.2977 | ##### | 2.5319 | 1.1345E-02 | 2.3253E-02 |
| ENSG00000166169\|POLL | 1551.9020 | -0.3428 | ##### | -2.5316 | 1.1353E-02 | 2.3268E-02 |
| ENSG00000174197\|MGA | 3586.6100 | 0.3153 | ##### | 2.5316 | 1.1354E-02 | 2.3268E-02 |
| ENSG00000101856\|PGRMC1 | 14716.9789 | 0.3028 | ##### | 2.5308 | 1.1380E-02 | 2.3321E-02 |
| ENSG00000137574\|TGS1 | 3620.4424 | 0.2499 | ##### | 2.5302 | 1.1401E-02 | 2.3361E-02 |
| ENSG00000116127\|ALMS1 | 2385.7617 | 0.3257 | ##### | 2.5297 | 1.1417E-02 | 2.3392E-02 |
| ENSG00000166326\|TRIM44 | 7946.1338 | -0.2945 | ##### | -2.5296 | 1.1418E-02 | 2.3392E-02 |
| ENSG00000090432\|MUL1 | 3054.0606 | -0.3050 | ##### | -2.5296 | 1.1420E-02 | 2.3395E-02 |
| ENSG00000261474 | 20.9934 | -1.2728 | ##### | -2.5295 | 1.1421E-02 | 2.3397E-02 |
| ENSG00000149311\|ATM | 3228.5249 | 0.3236 | ##### | 2.5294 | 1.1426E-02 | 2.3405E-02 |
| ENSG00000265182\|SRP72P1 | 5.7925 | 3.0436 | ##### | 2.5289 | 1.1442E-02 | 2.3435E-02 |
| ENSG00000286207 | 72.4730 | 0.7720 | ##### | 2.5283 | 1.1460E-02 | 2.3471E-02 |
| ENSG00000235897\|TM4SF19-AS1 | 27.2614 | -0.9966 | ##### | -2.5280 | 1.1470E-02 | 2.3490E-02 |
| ENSG00000283740\|TAF11L11 | 12.6033 | -2.6697 | ##### | -2.5277 | 1.1483E-02 | 2.3514E-02 |
| ENSG00000185942\|NKAIN3 | 204.2753 | 0.5354 | ##### | 2.5276 | 1.1484E-02 | 2.3515E-02 |
| ENSG00000197576\|HOXA4 | 103.4281 | 0.5285 | ##### | 2.5275 | 1.1487E-02 | 2.3521E-02 |
| ENSG00000167720\|SRR | 1462.5196 | -0.3534 | ##### | -2.5275 | 1.1488E-02 | 2.3521E-02 |
| ENSG00000255050 | 183.6093 | -0.5287 | ##### | -2.5274 | 1.1493E-02 | 2.3528E-02 |
| ENSG00000261758\|STAG1-DT | 17.2990 | -7.5193 | ##### | -2.5264 | 1.1525E-02 | 2.3593E-02 |
| ENSG00000132432\|SEC61G | 7515.9803 | -0.3124 | ##### | -2.5262 | 1.1529E-02 | 2.3600E-02 |
| ENSG00000231864 | 247.8552 | 0.5035 | ##### | 2.5258 | 1.1543E-02 | 2.3627E-02 |
| ENSG00000254682\|DHCR7-DT | 112.4433 | -0.5503 | ##### | -2.5252 | 1.1563E-02 | 2.3664E-02 |
| ENSG00000138593\|SECISBP2L | 3509.9182 | 0.3125 | ##### | 2.5252 | 1.1563E-02 | 2.3664E-02 |
| ENSG00000165935\|SMCO2 | 29.8971 | -1.1954 | ##### | -2.5247 | 1.1580E-02 | 2.3696E-02 |
| ENSG00000234956\|LINC02539 | 5.4803 | -6.3534 | ##### | -2.5233 | 1.1627E-02 | 2.3791E-02 |
| ENSG00000255867\|DENND5B-AS1 | 27.0184 | 0.9734 | ##### | 2.5232 | 1.1630E-02 | 2.3794E-02 |
| ENSG00000184208\|C22orf46P | 177.8357 | 2.2072 | ##### | 2.5232 | 1.1629E-02 | 2.3794E-02 |
| ENSG00000197472\|ZNF695 | 240.4029 | -0.4403 | ##### | -2.5231 | 1.1631E-02 | 2.3795E-02 |
| ENSG00000115677\|HDLBP | 53109.3538 | -0.2852 | ##### | -2.5225 | 1.1653E-02 | 2.3839E-02 |
| ENSG00000167528\|ZNF641 | 1488.4108 | -0.3220 | ##### | -2.5222 | 1.1661E-02 | 2.3850E-02 |
| ENSG00000290382\|ZNF876P | 36.8663 | -4.9029 | ##### | -2.5223 | 1.1660E-02 | 2.3850E-02 |
| ENSG00000272428 | 9.7993 | 5.3719 | ##### | 2.5222 | 1.1661E-02 | 2.3850E-02 |
| ENSG00000263120 | 5.5486 | 5.7277 | ##### | 2.5216 | 1.1681E-02 | 2.3888E-02 |
| ENSG00000155269\|GPR78 | 101.8229 | 0.5123 | ##### | 2.5216 | 1.1681E-02 | 2.3888E-02 |
| ENSG00000267293 | 11.6595 | 1.5636 | ##### | 2.5216 | 1.1683E-02 | 2.3889E-02 |
| ENSG00000275720 | 11.5897 | 1.3453 | ##### | 2.5215 | 1.1686E-02 | 2.3894E-02 |
| ENSG00000162191\|UBXN1 | 4573.6615 | 0.3159 | ##### | 2.5215 | 1.1687E-02 | 2.3894E-02 |
| ENSG00000114859\|CLCN2 | 735.5479 | -0.3520 | ##### | -2.5213 | 1.1694E-02 | 2.3907E-02 |
| ENSG00000103111\|MON1B | 5630.7436 | -0.2932 | ##### | -2.5208 | 1.1710E-02 | 2.3939E-02 |
| ENSG00000280828 | 10.6434 | 1.4785 | ##### | 2.5207 | 1.1712E-02 | 2.3940E-02 |
| ENSG00000119321\|FKBP15 | 4295.4639 | -0.2969 | ##### | -2.5206 | 1.1715E-02 | 2.3946E-02 |
| ENSG00000184983\|NDUFA6 | 4349.4970 | -0.2935 | ##### | -2.5203 | 1.1726E-02 | 2.3966E-02 |
| ENSG00000247077\|PGAM5 | 14162.0165 | 0.2831 | ##### | 2.5201 | 1.1731E-02 | 2.3975E-02 |
| ENSG00000241838 | 6.9194 | -1.4487 | ##### | -2.5200 | 1.1737E-02 | 2.3985E-02 |
| ENSG00000224094\|RPS24P8 | 14.3163 | -1.2337 | ##### | -2.5192 | 1.1762E-02 | 2.4034E-02 |

| | | | | | | |
|---|---|---|---|---|---|---|
| ENSG00000219607\|PPP1R3G | 15.7414 | -1.3510 | ##### | -2.5192 | 1.1763E-02 | 2.4036E-02 |
| ENSG00000288989 | 17.8984 | -0.9827 | ##### | -2.5191 | 1.1765E-02 | 2.4037E-02 |
| ENSG00000188536\|HBA2 | 7.6636 | -2.6807 | ##### | -2.5187 | 1.1780E-02 | 2.4065E-02 |
| ENSG00000198908\|GPRASP3 | 243.2665 | -1.1705 | ##### | -2.5187 | 1.1779E-02 | 2.4065E-02 |
| ENSG00000090932\|DLL3 | 554.0952 | -0.3914 | ##### | -2.5184 | 1.1790E-02 | 2.4085E-02 |
| ENSG00000142405\|NLRP12 | 30.5690 | -8.3493 | ##### | -2.5177 | 1.1813E-02 | 2.4130E-02 |
| ENSG00000176531\|PHLDB3 | 999.8352 | -0.3207 | ##### | -2.5172 | 1.1831E-02 | 2.4163E-02 |
| ENSG00000104951\|IL4I1 | 33.8229 | 1.0445 | ##### | 2.5171 | 1.1831E-02 | 2.4163E-02 |
| ENSG00000274641\|H2BC17 | 36.4063 | 1.5524 | ##### | 2.5169 | 1.1840E-02 | 2.4179E-02 |
| ENSG00000130203\|APOE | 1831.6275 | 0.3121 | ##### | 2.5167 | 1.1847E-02 | 2.4192E-02 |
| ENSG00000232194 | 32.1478 | -2.1940 | ##### | -2.5165 | 1.1851E-02 | 2.4197E-02 |
| ENSG00000156345\|CDK20 | 634.8699 | -0.3648 | ##### | -2.5166 | 1.1851E-02 | 2.4197E-02 |
| ENSG00000173175\|ADCY5 | 141.1554 | 0.5451 | ##### | 2.5161 | 1.1868E-02 | 2.4229E-02 |
| ENSG00000226833\|PPP1CB-DT | 36.9079 | -0.9247 | ##### | -2.5157 | 1.1880E-02 | 2.4252E-02 |
| ENSG00000124615\|MOCS1 | 414.9599 | 0.4159 | ##### | 2.5153 | 1.1895E-02 | 2.4281E-02 |
| ENSG00000239920 | 9.8683 | -3.9457 | ##### | -2.5148 | 1.1909E-02 | 2.4309E-02 |
| ENSG00000272686\|WASL-DT | 333.9535 | -0.4269 | ##### | -2.5147 | 1.1913E-02 | 2.4316E-02 |
| ENSG00000166562\|SEC11C | 3242.9110 | -0.3036 | ##### | -2.5147 | 1.1915E-02 | 2.4317E-02 |
| ENSG00000280164 | 10.4316 | -1.4962 | ##### | -2.5144 | 1.1924E-02 | 2.4334E-02 |
| ENSG00000291299 | 104.8188 | 0.8378 | ##### | 2.5136 | 1.1951E-02 | 2.4389E-02 |
| ENSG00000128346\|C22orf23 | 387.0921 | 0.4107 | ##### | 2.5128 | 1.1979E-02 | 2.4443E-02 |
| ENSG00000189431\|RASSF10 | 16.6597 | -2.4173 | ##### | -2.5125 | 1.1988E-02 | 2.4459E-02 |
| ENSG00000278588\|H2BC10 | 12.9117 | 2.4304 | ##### | 2.5125 | 1.1989E-02 | 2.4460E-02 |
| ENSG00000178053\|MLF1 | 2571.0407 | -0.3159 | ##### | -2.5124 | 1.1991E-02 | 2.4463E-02 |
| ENSG00000227210\|WDR35-DT | 48.1560 | 0.8291 | ##### | 2.5124 | 1.1992E-02 | 2.4463E-02 |
| ENSG00000268536 | 27.9836 | 5.3189 | ##### | 2.5119 | 1.2009E-02 | 2.4497E-02 |
| ENSG00000189334\|S100A14 | 18.5423 | -3.6068 | ##### | -2.5112 | 1.2032E-02 | 2.4541E-02 |
| ENSG00000197763\|TXNRD3 | 765.2541 | -0.3505 | ##### | -2.5107 | 1.2050E-02 | 2.4577E-02 |
| ENSG00000249738 | 14.5615 | 1.8021 | ##### | 2.5099 | 1.2077E-02 | 2.4629E-02 |
| ENSG00000124216\|SNAI1 | 985.8938 | -0.3506 | ##### | -2.5098 | 1.2081E-02 | 2.4636E-02 |
| ENSG00000265100 | 15.7958 | 1.5360 | ##### | 2.5093 | 1.2097E-02 | 2.4668E-02 |
| ENSG00000124596\|OARD1 | 1261.4231 | -0.3240 | ##### | -2.5090 | 1.2108E-02 | 2.4689E-02 |
| ENSG00000183570\|PCBP3 | 695.2738 | 0.3669 | ##### | 2.5088 | 1.2113E-02 | 2.4697E-02 |
| ENSG00000235872 | 25.9643 | 0.8748 | ##### | 2.5086 | 1.2121E-02 | 2.4711E-02 |
| ENSG00000010539\|ZNF200 | 1332.3261 | 0.3363 | ##### | 2.5085 | 1.2125E-02 | 2.4718E-02 |
| ENSG00000149428\|HYOU1 | 27515.6453 | 0.2580 | ##### | 2.5079 | 1.2144E-02 | 2.4755E-02 |
| ENSG00000290907 | 36.4013 | -0.9736 | ##### | -2.5074 | 1.2163E-02 | 2.4792E-02 |
| ENSG00000279641 | 195.2684 | 0.5562 | ##### | 2.5072 | 1.2168E-02 | 2.4800E-02 |
| ENSG00000231345\|BEND3P1 | 29.9965 | 2.5104 | ##### | 2.5072 | 1.2169E-02 | 2.4801E-02 |
| ENSG00000064687\|ABCA7 | 3173.5302 | -0.3109 | ##### | -2.5066 | 1.2189E-02 | 2.4839E-02 |
| ENSG00000259469 | 9.1711 | 2.3941 | ##### | 2.5063 | 1.2199E-02 | 2.4859E-02 |
| ENSG00000108094\|CUL2 | 6765.2899 | 0.2983 | ##### | 2.5062 | 1.2203E-02 | 2.4864E-02 |
| ENSG00000237080\|EHMT2-AS1 | 56.5891 | 1.8671 | ##### | 2.5058 | 1.2218E-02 | 2.4894E-02 |
| ENSG00000259462\|CPEB1-AS1 | 31.0287 | 1.3460 | ##### | 2.5057 | 1.2221E-02 | 2.4898E-02 |
| ENSG00000173826\|KCNH6 | 11.7883 | 2.8366 | ##### | 2.5052 | 1.2237E-02 | 2.4929E-02 |
| ENSG00000259828 | 12.9963 | -2.3579 | ##### | -2.5050 | 1.2246E-02 | 2.4945E-02 |
| ENSG00000291029\|FKBP9P1 | 37.6176 | -1.2650 | ##### | -2.5046 | 1.2259E-02 | 2.4970E-02 |
| ENSG00000187824\|TMEM220 | 538.1452 | 0.3774 | ##### | 2.5044 | 1.2265E-02 | 2.4981E-02 |
| ENSG00000223313\|RNU6-516P | 4.4432 | 2.1107 | ##### | 2.5044 | 1.2266E-02 | 2.4981E-02 |
| ENSG00000169564\|PCBP1 | 59371.8145 | 0.3084 | ##### | 2.5042 | 1.2273E-02 | 2.4993E-02 |
| ENSG00000257681 | 50.1284 | 0.9388 | ##### | 2.5039 | 1.2282E-02 | 2.5011E-02 |
| ENSG00000134256\|CD101 | 20.7875 | -1.1119 | ##### | -2.5039 | 1.2285E-02 | 2.5015E-02 |
| ENSG00000144668\|ITGA9 | 21.2391 | 1.7615 | ##### | 2.5036 | 1.2292E-02 | 2.5027E-02 |
| ENSG00000125843\|AP5S1 | 952.0048 | -0.2852 | ##### | -2.5029 | 1.2320E-02 | 2.5082E-02 |
| ENSG00000183337\|BCOR | 5270.1911 | -0.3067 | ##### | -2.5025 | 1.2332E-02 | 2.5105E-02 |
| ENSG00000148468\|FAM171A1 | 3397.0120 | 0.2928 | ##### | 2.5025 | 1.2334E-02 | 2.5105E-02 |
| ENSG00000261652\|PIERCE2 | 145.9309 | 0.5632 | ##### | 2.5025 | 1.2334E-02 | 2.5105E-02 |
| ENSG00000224596\|ZMIZ1-AS1 | 8.8148 | -3.4823 | ##### | -2.5022 | 1.2342E-02 | 2.5121E-02 |
| ENSG00000185515\|BRCC3 | 3196.6915 | 0.3100 | ##### | 2.5021 | 1.2345E-02 | 2.5125E-02 |
| ENSG00000232485\|RPL37A-DT | 51.0646 | -0.8824 | ##### | -2.5019 | 1.2352E-02 | 2.5137E-02 |
| ENSG00000267321\|SNHG30 | 662.7984 | -0.3325 | ##### | -2.5018 | 1.2358E-02 | 2.5147E-02 |
| ENSG00000058091\|CDK14 | 219.1573 | -0.4682 | ##### | -2.5012 | 1.2377E-02 | 2.5185E-02 |
| ENSG00000164221\|CCDC112 | 639.4913 | 0.3816 | ##### | 2.5004 | 1.2404E-02 | 2.5238E-02 |

| | | | | | | |
|---|---|---|---|---|---|---|
| ENSG00000140090\|SLC24A4 | 13.0299 | 2.0279 | ##### | 2.5004 | 1.2406E-02 | 2.5240E-02 |
| ENSG00000182931\|WFDC10B | 20.3210 | 7.6817 | ##### | 2.5000 | 1.2421E-02 | 2.5267E-02 |
| ENSG00000081051\|AFP | 18.6519 | -8.3320 | ##### | -2.5000 | 1.2421E-02 | 2.5267E-02 |
| ENSG00000236008\|LINC01814 | 19.0802 | -5.7714 | ##### | -2.4996 | 1.2434E-02 | 2.5293E-02 |
| ENSG00000113763\|UNC5A | 43.6067 | 1.2446 | ##### | 2.4995 | 1.2437E-02 | 2.5296E-02 |
| ENSG00000172175\|MALT1 | 5964.6339 | 0.3103 | ##### | 2.4993 | 1.2444E-02 | 2.5310E-02 |
| ENSG00000256849\|TCP1P3 | 12.4541 | -1.1846 | ##### | -2.4990 | 1.2454E-02 | 2.5328E-02 |
| ENSG00000290578 | 125.5779 | -0.5016 | ##### | -2.4990 | 1.2456E-02 | 2.5330E-02 |
| ENSG00000135315\|CEP162 | 495.1200 | 0.3630 | ##### | 2.4987 | 1.2466E-02 | 2.5348E-02 |
| ENSG00000159917\|ZNF235 | 439.0628 | -0.3862 | ##### | -2.4985 | 1.2471E-02 | 2.5356E-02 |
| ENSG00000243870\|RN7SL236P | 4.5422 | 4.4864 | ##### | 2.4983 | 1.2478E-02 | 2.5369E-02 |
| ENSG00000184014\|DENND5A | 6358.8834 | -0.2770 | ##### | -2.4982 | 1.2482E-02 | 2.5376E-02 |
| ENSG00000261357 | 5.1983 | -3.7177 | ##### | -2.4980 | 1.2490E-02 | 2.5390E-02 |
| ENSG00000173153\|ESRRA | 4041.9904 | 0.3084 | ##### | 2.4978 | 1.2498E-02 | 2.5405E-02 |
| ENSG00000221164\|SNORA11F | 20.8687 | -0.9512 | ##### | -2.4977 | 1.2501E-02 | 2.5410E-02 |
| ENSG00000213780\|GTF2H4 | 1527.4383 | 0.4996 | ##### | 2.4975 | 1.2508E-02 | 2.5421E-02 |
| ENSG00000121417\|ZNF211 | 610.5199 | -0.3510 | ##### | -2.4970 | 1.2524E-02 | 2.5453E-02 |
| ENSG00000049089\|COL9A2 | 54.3311 | 1.5745 | ##### | 2.4967 | 1.2535E-02 | 2.5473E-02 |
| ENSG00000120057\|SFRP5 | 25.4098 | 5.2894 | ##### | 2.4965 | 1.2541E-02 | 2.5484E-02 |
| ENSG00000171435\|KSR2 | 48.9184 | -0.6728 | ##### | -2.4963 | 1.2550E-02 | 2.5500E-02 |
| ENSG00000077238\|IL4R | 4111.0112 | -0.2990 | ##### | -2.4962 | 1.2555E-02 | 2.5508E-02 |
| ENSG00000128805\|ARHGAP22 | 545.6288 | 0.3597 | ##### | 2.4954 | 1.2581E-02 | 2.5561E-02 |
| ENSG00000196844\|PATE2 | 18.4805 | 1.6616 | ##### | 2.4951 | 1.2591E-02 | 2.5579E-02 |
| ENSG00000197037\|ZSCAN25 | 1574.3979 | 0.3272 | ##### | 2.4949 | 1.2599E-02 | 2.5592E-02 |
| ENSG00000086232\|EIF2AK1 | 23597.8977 | -0.2205 | ##### | -2.4947 | 1.2608E-02 | 2.5610E-02 |
| ENSG00000260681 | 3.6463 | -3.7106 | ##### | -2.4946 | 1.2609E-02 | 2.5610E-02 |
| ENSG00000272599 | 396.2693 | -0.3700 | ##### | -2.4945 | 1.2613E-02 | 2.5617E-02 |
| ENSG00000124602\|UNC5CL | 70.0124 | -0.7259 | ##### | -2.4942 | 1.2623E-02 | 2.5636E-02 |
| ENSG00000289209 | 28.7795 | 0.8359 | ##### | 2.4942 | 1.2625E-02 | 2.5638E-02 |
| ENSG00000181264\|TLCD5 | 694.8317 | -0.3670 | ##### | -2.4940 | 1.2631E-02 | 2.5647E-02 |
| ENSG00000174373\|RALGAPA1 | 2448.0269 | 0.3121 | ##### | 2.4933 | 1.2658E-02 | 2.5700E-02 |
| ENSG00000187079\|TEAD1 | 14824.4419 | -0.2968 | ##### | -2.4932 | 1.2661E-02 | 2.5706E-02 |
| ENSG00000277491 | 18.4199 | 1.2687 | ##### | 2.4931 | 1.2665E-02 | 2.5711E-02 |
| ENSG00000169607\|CKAP2L | 3689.6480 | -0.3299 | ##### | -2.4925 | 1.2684E-02 | 2.5749E-02 |
| ENSG00000254979 | 7.5645 | -3.5624 | ##### | -2.4924 | 1.2687E-02 | 2.5752E-02 |
| ENSG00000131504\|DIAPH1 | 25406.2270 | -0.2893 | ##### | -2.4923 | 1.2691E-02 | 2.5759E-02 |
| ENSG00000272631 | 110.4036 | -0.5208 | ##### | -2.4919 | 1.2706E-02 | 2.5788E-02 |
| ENSG00000260659 | 7.3571 | -1.5474 | ##### | -2.4916 | 1.2719E-02 | 2.5811E-02 |
| ENSG00000243680\|RPL37P23 | 15.8366 | 1.1456 | ##### | 2.4914 | 1.2724E-02 | 2.5822E-02 |
| ENSG00000235082\|SUMO1P3 | 21.2529 | 1.0779 | ##### | 2.4914 | 1.2726E-02 | 2.5823E-02 |
| ENSG00000103546\|SLC6A2 | 5.3034 | -6.3062 | ##### | -2.4912 | 1.2732E-02 | 2.5833E-02 |
| ENSG00000154814\|OXNAD1 | 845.1428 | 0.3524 | ##### | 2.4911 | 1.2735E-02 | 2.5838E-02 |
| ENSG00000258988\|SF3B4P1 | 7.4716 | 1.9151 | ##### | 2.4910 | 1.2737E-02 | 2.5839E-02 |
| ENSG00000239948\|RN7SL368P | 5.3721 | 5.1791 | ##### | 2.4907 | 1.2748E-02 | 2.5861E-02 |
| ENSG00000126005\|MMP24OS | 2347.2772 | -0.3146 | ##### | -2.4906 | 1.2752E-02 | 2.5866E-02 |
| ENSG00000070501\|POLB | 829.3422 | 0.3531 | ##### | 2.4898 | 1.2781E-02 | 2.5924E-02 |
| ENSG00000137074\|APTX | 2542.5016 | 0.3174 | ##### | 2.4896 | 1.2790E-02 | 2.5941E-02 |
| ENSG00000270919 | 4.7610 | -5.1309 | ##### | -2.4894 | 1.2795E-02 | 2.5949E-02 |
| ENSG00000277349\|CICP25 | 9.9376 | 4.0280 | ##### | 2.4886 | 1.2825E-02 | 2.6008E-02 |
| ENSG00000258466 | 3.6132 | 2.4794 | ##### | 2.4881 | 1.2841E-02 | 2.6039E-02 |
| ENSG00000271253 | 7.7284 | 5.5205 | ##### | 2.4880 | 1.2845E-02 | 2.6045E-02 |
| ENSG00000291048 | 37.7026 | -1.1517 | ##### | -2.4877 | 1.2856E-02 | 2.6066E-02 |
| ENSG00000170959\|DCDC1 | 23.8929 | 2.1162 | ##### | 2.4869 | 1.2888E-02 | 2.6126E-02 |
| ENSG00000283511 | 15.3415 | -3.0401 | ##### | -2.4869 | 1.2888E-02 | 2.6126E-02 |
| ENSG00000115233\|PSMD14 | 9343.6676 | 0.2980 | ##### | 2.4866 | 1.2898E-02 | 2.6145E-02 |
| ENSG00000250166\|C2CD5-AS1 | 12.5712 | -4.6057 | ##### | -2.4861 | 1.2917E-02 | 2.6180E-02 |
| ENSG00000170776\|AKAP13 | 5610.9148 | -0.3063 | ##### | -2.4861 | 1.2917E-02 | 2.6180E-02 |
| ENSG00000143228\|NUF2 | 2131.1345 | 0.3188 | ##### | 2.4859 | 1.2923E-02 | 2.6191E-02 |
| ENSG00000164898\|FMC1 | 873.4851 | -0.3734 | ##### | -2.4858 | 1.2925E-02 | 2.6193E-02 |
| ENSG00000257493 | 5.5433 | 5.7198 | ##### | 2.4855 | 1.2938E-02 | 2.6217E-02 |
| ENSG00000186952\|TMEM232 | 8.1479 | 2.0970 | ##### | 2.4854 | 1.2942E-02 | 2.6223E-02 |
| ENSG00000150756\|ATPSCKMT | 1111.0128 | -0.3327 | ##### | -2.4849 | 1.2958E-02 | 2.6254E-02 |
| ENSG00000007516\|BAIAP3 | 259.6835 | 0.4612 | ##### | 2.4847 | 1.2964E-02 | 2.6265E-02 |

| | | | | | | |
|---|---|---|---|---|---|---|
| ENSG00000161654\|LSM12 | 4626.1215 | 0.3123 | ##### | 2.4847 | 1.2965E-02 | 2.6265E-02 |
| ENSG00000286181 | 69.9004 | -0.6845 | ##### | -2.4847 | 1.2966E-02 | 2.6266E-02 |
| ENSG00000279265 | 19.9981 | 2.3408 | ##### | 2.4846 | 1.2971E-02 | 2.6275E-02 |
| ENSG00000188938\|FAM120AOS | 3610.8601 | -0.3126 | ##### | -2.4842 | 1.2986E-02 | 2.6302E-02 |
| ENSG00000253327\|RAD21-AS1 | 19.8149 | -1.3003 | ##### | -2.4834 | 1.3013E-02 | 2.6355E-02 |
| ENSG00000169598\|DFFB | 470.3368 | -0.3643 | ##### | -2.4834 | 1.3014E-02 | 2.6355E-02 |
| ENSG00000162642\|C1orf52 | 1776.9104 | 0.3187 | ##### | 2.4821 | 1.3061E-02 | 2.6448E-02 |
| ENSG00000246308 | 27.7544 | 1.7577 | ##### | 2.4820 | 1.3066E-02 | 2.6457E-02 |
| ENSG00000137494\|ANKRD42 | 1241.4490 | -0.3087 | ##### | -2.4819 | 1.3069E-02 | 2.6462E-02 |
| ENSG00000006194\|ZNF263 | 3391.3509 | 0.3001 | ##### | 2.4812 | 1.3093E-02 | 2.6509E-02 |
| ENSG00000137218\|FRS3 | 281.5297 | 0.4194 | ##### | 2.4806 | 1.3117E-02 | 2.6554E-02 |
| ENSG00000232536 | 18.0129 | -1.3568 | ##### | -2.4802 | 1.3130E-02 | 2.6580E-02 |
| ENSG00000273447 | 9.7311 | 3.6671 | ##### | 2.4800 | 1.3137E-02 | 2.6592E-02 |
| ENSG00000237916\|RPL37P12 | 6.0258 | -2.8260 | ##### | -2.4799 | 1.3141E-02 | 2.6594E-02 |
| ENSG00000155657\|TTN | 1114.8267 | 0.3231 | ##### | 2.4799 | 1.3141E-02 | 2.6594E-02 |
| ENSG00000275882\|IKBKGP1 | 958.8479 | 0.3170 | ##### | 2.4799 | 1.3140E-02 | 2.6594E-02 |
| ENSG00000150275\|PCDH15 | 118.2807 | 0.5088 | ##### | 2.4792 | 1.3167E-02 | 2.6644E-02 |
| ENSG00000227288\|ARID3BP1 | 24.9041 | 1.4735 | ##### | 2.4792 | 1.3169E-02 | 2.6647E-02 |
| ENSG00000229558\|SACS-AS1 | 5.3380 | 5.6712 | ##### | 2.4782 | 1.3206E-02 | 2.6721E-02 |
| ENSG00000272701\|MESTIT1 | 8.6696 | 2.7812 | ##### | 2.4776 | 1.3228E-02 | 2.6763E-02 |
| ENSG00000198836\|OPA1 | 9703.2496 | 0.2997 | ##### | 2.4771 | 1.3245E-02 | 2.6796E-02 |
| ENSG00000134049\|IER3IP1 | 4595.9674 | -0.3033 | ##### | -2.4769 | 1.3254E-02 | 2.6812E-02 |
| ENSG00000276753\|MIR6821 | 26.0438 | 0.9042 | ##### | 2.4768 | 1.3257E-02 | 2.6816E-02 |
| ENSG00000104808\|DHDH | 33.5344 | -1.6102 | ##### | -2.4767 | 1.3260E-02 | 2.6821E-02 |
| ENSG00000113742\|CPEB4 | 1546.3306 | -0.3353 | ##### | -2.4765 | 1.3266E-02 | 2.6830E-02 |
| ENSG00000270761\|CD2AP-DT | 37.8924 | -0.8014 | ##### | -2.4766 | 1.3265E-02 | 2.6830E-02 |
| ENSG00000077713\|SLC25A43 | 3692.2415 | -0.2984 | ##### | -2.4765 | 1.3267E-02 | 2.6830E-02 |
| ENSG00000293190 | 18.0449 | 1.2768 | ##### | 2.4756 | 1.3303E-02 | 2.6900E-02 |
| ENSG00000177054\|ZDHHC13 | 1358.9261 | 0.3372 | ##### | 2.4753 | 1.3311E-02 | 2.6915E-02 |
| ENSG00000165055\|METTL2B | 5560.4728 | 0.3034 | ##### | 2.4751 | 1.3320E-02 | 2.6932E-02 |
| ENSG00000215458\|AATBC | 30.7514 | 1.0848 | ##### | 2.4748 | 1.3332E-02 | 2.6954E-02 |
| ENSG00000228929\|RPS13P2 | 91.2212 | 0.5183 | ##### | 2.4736 | 1.3377E-02 | 2.7041E-02 |
| ENSG00000273328 | 11.2238 | 4.8939 | ##### | 2.4736 | 1.3377E-02 | 2.7041E-02 |
| ENSG00000085377\|PREP | 4861.8928 | -0.3011 | ##### | -2.4734 | 1.3384E-02 | 2.7053E-02 |
| ENSG00000271032 | 24.0236 | -2.0403 | ##### | -2.4733 | 1.3386E-02 | 2.7056E-02 |
| ENSG00000251666\|ZNF346-IT1 | 8.8273 | 2.8895 | ##### | 2.4731 | 1.3393E-02 | 2.7068E-02 |
| ENSG00000114405\|CEP15 | 773.6143 | -0.3661 | ##### | -2.4729 | 1.3401E-02 | 2.7082E-02 |
| ENSG00000289031 | 6.3875 | -2.7824 | ##### | -2.4729 | 1.3404E-02 | 2.7086E-02 |
| ENSG00000130560\|UBAC1 | 2946.3485 | -0.3041 | ##### | -2.4727 | 1.3409E-02 | 2.7095E-02 |
| ENSG00000182108\|DEXI | 2800.8444 | -0.3155 | ##### | -2.4724 | 1.3419E-02 | 2.7113E-02 |
| ENSG00000157423\|HYDIN | 220.8775 | -0.4435 | ##### | -2.4722 | 1.3428E-02 | 2.7130E-02 |
| ENSG00000169856\|ONECUT1 | 20.8569 | 2.1845 | ##### | 2.4722 | 1.3430E-02 | 2.7131E-02 |
| ENSG00000255958\|GABARAPL1-AS1 | 7.6180 | 2.0761 | ##### | 2.4721 | 1.3434E-02 | 2.7138E-02 |
| ENSG00000185186\|LINC00313 | 39.2996 | -0.8155 | ##### | -2.4718 | 1.3443E-02 | 2.7154E-02 |
| ENSG00000134070\|IRAK2 | 2081.3241 | -0.3135 | ##### | -2.4714 | 1.3457E-02 | 2.7180E-02 |
| ENSG00000237840\|FAM21FP | 14.1533 | -2.9735 | ##### | -2.4714 | 1.3460E-02 | 2.7185E-02 |
| ENSG00000159885\|ZNF222 | 268.9700 | -0.5145 | ##### | -2.4710 | 1.3475E-02 | 2.7214E-02 |
| ENSG00000198680\|TUSC1 | 818.9563 | -0.3538 | ##### | -2.4709 | 1.3477E-02 | 2.7216E-02 |
| ENSG00000197061\|H4C3 | 15.9948 | 2.2872 | ##### | 2.4702 | 1.3503E-02 | 2.7265E-02 |
| ENSG00000272849 | 63.0481 | 0.6562 | ##### | 2.4699 | 1.3515E-02 | 2.7290E-02 |
| ENSG00000273026 | 46.1850 | 0.7843 | ##### | 2.4698 | 1.3517E-02 | 2.7291E-02 |
| ENSG00000264558 | 193.1603 | 0.4079 | ##### | 2.4698 | 1.3518E-02 | 2.7291E-02 |
| ENSG00000288710 | 75.1036 | 4.2774 | ##### | 2.4694 | 1.3534E-02 | 2.7321E-02 |
| ENSG00000260086 | 12.8587 | -1.3916 | ##### | -2.4691 | 1.3547E-02 | 2.7346E-02 |
| ENSG00000150456\|EEF1AKMT1 | 325.6385 | -0.4294 | ##### | -2.4689 | 1.3553E-02 | 2.7353E-02 |
| ENSG00000293499 | 47.0767 | 0.7088 | ##### | 2.4689 | 1.3553E-02 | 2.7353E-02 |
| ENSG00000115274\|INO80B | 2112.7639 | 0.3124 | ##### | 2.4689 | 1.3552E-02 | 2.7353E-02 |
| ENSG00000277702 | 20.0836 | -8.2277 | ##### | -2.4687 | 1.3560E-02 | 2.7364E-02 |
| ENSG00000293332 | 333.8905 | -0.8153 | ##### | -2.4683 | 1.3577E-02 | 2.7396E-02 |
| ENSG00000178409\|BEND3 | 1310.2548 | 0.3432 | ##### | 2.4676 | 1.3604E-02 | 2.7449E-02 |
| ENSG00000231113\|LINC02976 | 71.1552 | 0.5841 | ##### | 2.4672 | 1.3616E-02 | 2.7473E-02 |
| ENSG00000180834\|MAP6D1 | 171.6707 | -0.5832 | ##### | -2.4671 | 1.3620E-02 | 2.7479E-02 |
| ENSG00000214391\|TUBAP2 | 150.5813 | 0.4472 | ##### | 2.4668 | 1.3634E-02 | 2.7505E-02 |

| | | | | | | |
|---|---|---|---|---|---|---|
| ENSG00000277383\|BICRA-AS2 | 57.5097 | -0.7515 | ##### | -2.4664 | 1.3649E-02 | 2.7533E-02 |
| ENSG00000166263\|STXBP4 | 1207.6387 | -0.3173 | ##### | -2.4663 | 1.3653E-02 | 2.7540E-02 |
| ENSG00000196302\|GUSBP15 | 37.3202 | 1.1548 | ##### | 2.4660 | 1.3663E-02 | 2.7558E-02 |
| ENSG00000263272 | 305.9562 | 0.4183 | ##### | 2.4651 | 1.3697E-02 | 2.7625E-02 |
| ENSG00000176371\|ZSCAN2 | 451.0572 | 0.3811 | ##### | 2.4650 | 1.3701E-02 | 2.7630E-02 |
| ENSG00000241506\|PSMC1P1 | 166.3567 | 0.4617 | ##### | 2.4649 | 1.3705E-02 | 2.7637E-02 |
| ENSG00000273108 | 6.2437 | -5.5159 | ##### | -2.4648 | 1.3708E-02 | 2.7640E-02 |
| ENSG00000229292\|RFPL4AL1 | 8.5991 | 5.6700 | ##### | 2.4648 | 1.3708E-02 | 2.7640E-02 |
| ENSG00000277498 | 2.8286 | 4.9299 | ##### | 2.4644 | 1.3723E-02 | 2.7668E-02 |
| ENSG00000278654 | 12.8246 | 1.9505 | ##### | 2.4643 | 1.3728E-02 | 2.7676E-02 |
| ENSG00000241685\|ARPC1A | 12781.9973 | -0.2967 | ##### | -2.4641 | 1.3735E-02 | 2.7688E-02 |
| ENSG00000237118\|CYP2F2P | 18.8770 | -2.4588 | ##### | -2.4638 | 1.3747E-02 | 2.7710E-02 |
| ENSG00000237017\|PRPF31-AS1 | 371.1835 | 0.6925 | ##### | 2.4638 | 1.3749E-02 | 2.7712E-02 |
| ENSG00000235698\|PA2G4P2 | 124.0802 | 0.5503 | ##### | 2.4633 | 1.3767E-02 | 2.7746E-02 |
| ENSG00000246130 | 15.5198 | 2.4204 | ##### | 2.4631 | 1.3775E-02 | 2.7762E-02 |
| ENSG00000183853\|KIRREL1 | 22667.8408 | 0.2931 | ##### | 2.4626 | 1.3794E-02 | 2.7797E-02 |
| ENSG00000085721\|RRN3 | 7745.5128 | 0.2298 | ##### | 2.4621 | 1.3813E-02 | 2.7834E-02 |
| ENSG00000103343\|ZNF174 | 934.9770 | -0.3289 | ##### | -2.4620 | 1.3817E-02 | 2.7840E-02 |
| ENSG00000180712\|LINC02363 | 5.6582 | -4.2699 | ##### | -2.4619 | 1.3821E-02 | 2.7847E-02 |
| ENSG00000235138 | 12.6618 | 2.2335 | ##### | 2.4617 | 1.3829E-02 | 2.7860E-02 |
| ENSG00000272030 | 195.4506 | 1.8044 | ##### | 2.4613 | 1.3842E-02 | 2.7886E-02 |
| ENSG00000137947\|GTF2B | 1740.4056 | -0.3107 | ##### | -2.4608 | 1.3863E-02 | 2.7925E-02 |
| ENSG00000233231\|HNRNPA1P49 | 7.3883 | 2.3404 | ##### | 2.4606 | 1.3869E-02 | 2.7936E-02 |
| ENSG00000182489\|XKRX | 31.2425 | 1.9829 | ##### | 2.4605 | 1.3876E-02 | 2.7947E-02 |
| ENSG00000183688\|RFLNB | 3245.0487 | -0.3107 | ##### | -2.4596 | 1.3908E-02 | 2.8010E-02 |
| ENSG00000234921 | 6.6909 | 1.9414 | ##### | 2.4590 | 1.3933E-02 | 2.8060E-02 |
| ENSG00000251026\|NIHCOLE | 95.8262 | -0.7098 | ##### | -2.4587 | 1.3942E-02 | 2.8076E-02 |
| ENSG00000197245\|FAM110D | 10.3689 | -2.3440 | ##### | -2.4580 | 1.3969E-02 | 2.8128E-02 |
| ENSG00000134504\|KCTD1 | 3184.9475 | -0.3095 | ##### | -2.4579 | 1.3976E-02 | 2.8139E-02 |
| ENSG00000170340\|B3GNT2 | 2537.8878 | 0.3062 | ##### | 2.4577 | 1.3982E-02 | 2.8150E-02 |
| ENSG00000225695\|HNRNPA1P35 | 9.9496 | 1.3650 | ##### | 2.4576 | 1.3986E-02 | 2.8156E-02 |
| ENSG00000135406\|PRPH | 57.5646 | 0.6861 | ##### | 2.4574 | 1.3994E-02 | 2.8170E-02 |
| ENSG00000255959 | 103.3125 | 0.6493 | ##### | 2.4573 | 1.3999E-02 | 2.8178E-02 |
| ENSG00000280383 | 85.2082 | 0.6727 | ##### | 2.4572 | 1.4004E-02 | 2.8187E-02 |
| ENSG00000155111\|CDK19 | 1674.6268 | -0.3193 | ##### | -2.4571 | 1.4008E-02 | 2.8192E-02 |
| ENSG00000172661\|WASHC2C | 3274.3492 | -0.3054 | ##### | -2.4566 | 1.4027E-02 | 2.8228E-02 |
| ENSG00000144120\|TMEM177 | 1694.5040 | -0.3159 | ##### | -2.4565 | 1.4031E-02 | 2.8235E-02 |
| ENSG00000228452\|C1orf50-AS1 | 5.7588 | -1.7582 | ##### | -2.4562 | 1.4041E-02 | 2.8253E-02 |
| ENSG00000072778\|ACADVL | 24320.5251 | 0.1890 | ##### | 2.4556 | 1.4065E-02 | 2.8300E-02 |
| ENSG00000030419\|IKZF2 | 233.9521 | -0.4809 | ##### | -2.4551 | 1.4084E-02 | 2.8335E-02 |
| ENSG00000246859\|STARD4-AS1 | 532.1039 | 1.8790 | ##### | 2.4548 | 1.4098E-02 | 2.8362E-02 |
| ENSG00000222046\|DCDC2B | 18.4540 | -0.9597 | ##### | -2.4547 | 1.4101E-02 | 2.8366E-02 |
| ENSG00000272461 | 17.7642 | -2.3456 | ##### | -2.4531 | 1.4163E-02 | 2.8489E-02 |
| ENSG00000181908\|LINC02724 | 10.0755 | 2.4200 | ##### | 2.4521 | 1.4201E-02 | 2.8563E-02 |
| ENSG00000164346\|NSA2 | 6002.3053 | 0.3003 | ##### | 2.4521 | 1.4203E-02 | 2.8566E-02 |
| ENSG00000258673\|LINC01397 | 7.6906 | 3.5139 | ##### | 2.4516 | 1.4222E-02 | 2.8603E-02 |
| ENSG00000156509\|FBXO43 | 276.7875 | -0.4218 | ##### | -2.4516 | 1.4224E-02 | 2.8603E-02 |
| ENSG00000163866\|SMIM12 | 3315.9631 | -0.2970 | ##### | -2.4509 | 1.4249E-02 | 2.8653E-02 |
| ENSG00000167315\|ACAA2 | 5661.3910 | -0.3091 | ##### | -2.4508 | 1.4254E-02 | 2.8660E-02 |
| ENSG00000230280\|HNRNPA1P59 | 18.1103 | -1.6661 | ##### | -2.4507 | 1.4259E-02 | 2.8668E-02 |
| ENSG00000269680 | 26.1609 | -0.8853 | ##### | -2.4497 | 1.4298E-02 | 2.8744E-02 |
| ENSG00000229873\|OGFR-AS1 | 37.9093 | 0.9560 | ##### | 2.4492 | 1.4317E-02 | 2.8781E-02 |
| ENSG00000280214 | 3.5038 | -5.2045 | ##### | -2.4492 | 1.4319E-02 | 2.8784E-02 |
| ENSG00000251303\|CAB39P1 | 6.1632 | -2.7472 | ##### | -2.4486 | 1.4340E-02 | 2.8823E-02 |
| ENSG00000081041\|CXCL2 | 1650.1249 | 0.3076 | ##### | 2.4485 | 1.4347E-02 | 2.8836E-02 |
| ENSG00000247092\|SNHG10 | 581.3908 | -0.3862 | ##### | -2.4480 | 1.4366E-02 | 2.8872E-02 |
| ENSG00000269826 | 25.7411 | -8.1013 | ##### | -2.4479 | 1.4369E-02 | 2.8875E-02 |
| ENSG00000283703\|VSIG10L2 | 79.0199 | 0.6564 | ##### | 2.4476 | 1.4381E-02 | 2.8899E-02 |
| ENSG00000135597\|REPS1 | 2258.9117 | -0.3169 | ##### | -2.4469 | 1.4410E-02 | 2.8954E-02 |
| ENSG00000097046\|CDC7 | 2286.2718 | 0.3043 | ##### | 2.4466 | 1.4419E-02 | 2.8972E-02 |
| ENSG00000250240 | 11.4722 | -1.6238 | ##### | -2.4463 | 1.4431E-02 | 2.8994E-02 |
| ENSG00000186862\|PDZD7 | 84.8741 | -0.6636 | ##### | -2.4460 | 1.4446E-02 | 2.9020E-02 |
| ENSG00000152217\|SETBP1 | 34.3638 | 1.9392 | ##### | 2.4459 | 1.4448E-02 | 2.9021E-02 |

| | | | | | | |
|---|---|---|---|---|---|---|
| ENSG00000093183\|SEC22C | 3641.7733 | -0.3091 | ##### | -2.4459 | 1.4447E-02 | 2.9021E-02 |
| ENSG00000206913 | 4.3899 | 4.2082 | ##### | 2.4459 | 1.4450E-02 | 2.9023E-02 |
| ENSG00000258758 | 6.3493 | 2.6568 | ##### | 2.4444 | 1.4510E-02 | 2.9141E-02 |
| ENSG00000275645 | 20.2245 | 1.5753 | ##### | 2.4434 | 1.4550E-02 | 2.9221E-02 |
| ENSG00000256843 | 3.3449 | 2.9625 | ##### | 2.4430 | 1.4565E-02 | 2.9248E-02 |
| ENSG00000138617\|PARP16 | 444.8064 | -0.3785 | ##### | -2.4424 | 1.4591E-02 | 2.9299E-02 |
| ENSG00000283743 | 9.9396 | 4.0413 | ##### | 2.4422 | 1.4596E-02 | 2.9308E-02 |
| ENSG00000198876\|DCAF12 | 4373.2798 | -0.2724 | ##### | -2.4421 | 1.4602E-02 | 2.9316E-02 |
| ENSG00000234036\|TXNP6 | 22.5632 | -1.0495 | ##### | -2.4420 | 1.4606E-02 | 2.9324E-02 |
| ENSG00000289891 | 42.9937 | -0.8767 | ##### | -2.4415 | 1.4625E-02 | 2.9360E-02 |
| ENSG00000141644\|MBD1 | 6001.7039 | 0.2739 | ##### | 2.4414 | 1.4630E-02 | 2.9368E-02 |
| ENSG00000110427\|KIAA1549L | 791.0705 | -0.3495 | ##### | -2.4406 | 1.4662E-02 | 2.9430E-02 |
| ENSG00000266405\|CBX3P2 | 6.8642 | -1.9169 | ##### | -2.4405 | 1.4666E-02 | 2.9436E-02 |
| ENSG00000248240 | 6.8582 | 5.0104 | ##### | 2.4405 | 1.4667E-02 | 2.9436E-02 |
| ENSG00000100058\|CRYBB2P1 | 31.5451 | 2.1562 | ##### | 2.4404 | 1.4670E-02 | 2.9440E-02 |
| ENSG00000288093 | 30.7787 | -0.8296 | ##### | -2.4403 | 1.4676E-02 | 2.9450E-02 |
| ENSG00000185946\|RNPC3 | 977.0296 | 0.3463 | ##### | 2.4400 | 1.4688E-02 | 2.9471E-02 |
| ENSG00000264864\|MIR3613 | 7.8117 | 4.5000 | ##### | 2.4398 | 1.4694E-02 | 2.9482E-02 |
| ENSG00000268333 | 21.2795 | 1.1422 | ##### | 2.4378 | 1.4775E-02 | 2.9643E-02 |
| ENSG00000260907 | 12.1654 | 2.4441 | ##### | 2.4374 | 1.4793E-02 | 2.9676E-02 |
| ENSG00000271664 | 25.2725 | 1.1615 | ##### | 2.4372 | 1.4800E-02 | 2.9688E-02 |
| ENSG00000176390\|CRLF3 | 3420.4912 | 0.3050 | ##### | 2.4369 | 1.4814E-02 | 2.9715E-02 |
| ENSG00000183260\|ABHD16B | 321.8304 | 0.4580 | ##### | 2.4365 | 1.4831E-02 | 2.9747E-02 |
| ENSG00000211454\|AKR7L | 22.0886 | 1.8830 | ##### | 2.4364 | 1.4834E-02 | 2.9751E-02 |
| ENSG00000288551 | 2.3105 | 4.9634 | ##### | 2.4359 | 1.4855E-02 | 2.9792E-02 |
| ENSG00000248643\|RBM14-RBM4 | 526.6152 | -0.6053 | ##### | -2.4354 | 1.4877E-02 | 2.9832E-02 |
| ENSG00000276216 | 9.2257 | 2.8233 | ##### | 2.4353 | 1.4878E-02 | 2.9833E-02 |
| ENSG00000121989\|ACVR2A | 561.7779 | 0.3592 | ##### | 2.4351 | 1.4889E-02 | 2.9854E-02 |
| ENSG00000104299\|INTS9 | 1450.1996 | 0.3231 | ##### | 2.4350 | 1.4894E-02 | 2.9860E-02 |
| ENSG00000237343 | 5.3345 | 5.4231 | ##### | 2.4344 | 1.4915E-02 | 2.9902E-02 |
| ENSG00000196689\|TRPV1 | 377.1556 | -0.3639 | ##### | -2.4340 | 1.4934E-02 | 2.9937E-02 |
| ENSG00000021574\|SPAST | 4140.9088 | 0.3070 | ##### | 2.4336 | 1.4950E-02 | 2.9967E-02 |
| ENSG00000286943 | 7.4074 | 2.6102 | ##### | 2.4335 | 1.4953E-02 | 2.9971E-02 |
| ENSG00000082175\|PGR | 16.3013 | 7.1481 | ##### | 2.4335 | 1.4956E-02 | 2.9974E-02 |
| ENSG00000115107\|STEAP3 | 10509.6501 | -0.2945 | ##### | -2.4334 | 1.4957E-02 | 2.9975E-02 |
| ENSG00000254680 | 56.0699 | 0.8653 | ##### | 2.4333 | 1.4964E-02 | 2.9983E-02 |
| ENSG00000279428 | 253.7352 | 0.4517 | ##### | 2.4333 | 1.4962E-02 | 2.9983E-02 |
| ENSG00000169245\|CXCL10 | 28.6259 | 8.1085 | ##### | 2.4333 | 1.4964E-02 | 2.9983E-02 |
| ENSG00000291010\|MRPS31P5 | 258.7483 | 0.8104 | ##### | 2.4327 | 1.4987E-02 | 3.0026E-02 |
| ENSG00000223813\|PRR15-DT | 23.4290 | -2.0812 | ##### | -2.4326 | 1.4991E-02 | 3.0034E-02 |
| ENSG00000250994 | 6.5497 | -3.2259 | ##### | -2.4325 | 1.4994E-02 | 3.0037E-02 |
| ENSG00000224631\|RPS27AP16 | 739.2221 | 0.3484 | ##### | 2.4322 | 1.5008E-02 | 3.0062E-02 |
| ENSG00000091128\|LAMB4 | 31.9970 | -7.4112 | ##### | -2.4319 | 1.5022E-02 | 3.0089E-02 |
| ENSG00000232335 | 7.8347 | 5.2253 | ##### | 2.4318 | 1.5025E-02 | 3.0090E-02 |
| ENSG00000258177 | 11.7404 | 1.5801 | ##### | 2.4318 | 1.5024E-02 | 3.0090E-02 |
| ENSG00000275185 | 174.8886 | 0.4679 | ##### | 2.4318 | 1.5025E-02 | 3.0090E-02 |
| ENSG00000188038\|NRN1L | 2.1668 | -2.9070 | ##### | -2.4316 | 1.5031E-02 | 3.0099E-02 |
| ENSG00000269696\|ZNF470-DT | 40.7116 | -1.5315 | ##### | -2.4313 | 1.5045E-02 | 3.0126E-02 |
| ENSG00000274105 | 7.2913 | 2.1227 | ##### | 2.4310 | 1.5056E-02 | 3.0145E-02 |
| ENSG00000262769 | 13.8764 | -2.4898 | ##### | -2.4306 | 1.5072E-02 | 3.0176E-02 |
| ENSG00000244563\|RPS26P19 | 20.4243 | -1.4398 | ##### | -2.4291 | 1.5136E-02 | 3.0301E-02 |
| ENSG00000226045 | 3.7968 | 5.1709 | ##### | 2.4291 | 1.5137E-02 | 3.0301E-02 |
| ENSG00000133961\|NUMB | 4898.4392 | -0.2371 | ##### | -2.4289 | 1.5146E-02 | 3.0317E-02 |
| ENSG00000217027\|TPT1P4 | 4.7151 | -1.9740 | ##### | -2.4288 | 1.5151E-02 | 3.0324E-02 |
| ENSG00000279122 | 6.7075 | 4.8894 | ##### | 2.4285 | 1.5161E-02 | 3.0342E-02 |
| ENSG00000261140 | 58.0632 | 0.6250 | ##### | 2.4280 | 1.5184E-02 | 3.0387E-02 |
| ENSG00000141506\|PIK3R5 | 24.5418 | 3.0106 | ##### | 2.4277 | 1.5195E-02 | 3.0407E-02 |
| ENSG00000119139\|TJP2 | 441.4236 | -2.2301 | ##### | -2.4276 | 1.5201E-02 | 3.0417E-02 |
| ENSG00000197019\|SERTAD1 | 1591.7857 | -0.3042 | ##### | -2.4271 | 1.5220E-02 | 3.0453E-02 |
| ENSG00000289523 | 33.2331 | 0.8786 | ##### | 2.4267 | 1.5238E-02 | 3.0483E-02 |
| ENSG00000263218 | 35.2129 | -0.5938 | ##### | -2.4267 | 1.5239E-02 | 3.0483E-02 |
| ENSG00000228395 | 54.5063 | -1.0037 | ##### | -2.4267 | 1.5236E-02 | 3.0483E-02 |
| ENSG00000288956 | 12.2536 | 1.6740 | ##### | 2.4266 | 1.5239E-02 | 3.0483E-02 |

| | | | | | | |
|---|---|---|---|---|---|---|
| ENSG00000258819\|LINC02289 | 14.3580 | 2.5236 | ##### | 2.4265 | 1.5247E-02 | 3.0497E-02 |
| ENSG00000255549\|ENPP7P6 | 4.3077 | 5.3594 | ##### | 2.4260 | 1.5267E-02 | 3.0536E-02 |
| ENSG00000267030 | 8.4402 | 1.5335 | ##### | 2.4249 | 1.5312E-02 | 3.0622E-02 |
| ENSG00000273687 | 7.1758 | 4.6164 | ##### | 2.4245 | 1.5328E-02 | 3.0653E-02 |
| ENSG00000117834\|SLC5A9 | 23.6959 | -2.1614 | ##### | -2.4242 | 1.5342E-02 | 3.0680E-02 |
| ENSG00000257635 | 4.8557 | -3.2804 | ##### | -2.4241 | 1.5345E-02 | 3.0682E-02 |
| ENSG00000232901\|CYCSP10 | 5.1485 | 2.8090 | ##### | 2.4241 | 1.5348E-02 | 3.0687E-02 |
| ENSG00000238278\|ALG1L6P | 30.5115 | -0.8136 | ##### | -2.4240 | 1.5351E-02 | 3.0690E-02 |
| ENSG00000260774 | 33.9437 | -0.8147 | ##### | -2.4223 | 1.5422E-02 | 3.0830E-02 |
| ENSG00000125347\|IRF1 | 3979.9428 | 0.3130 | ##### | 2.4217 | 1.5446E-02 | 3.0877E-02 |
| ENSG00000282950 | 31.6292 | -7.8657 | ##### | -2.4215 | 1.5455E-02 | 3.0892E-02 |
| ENSG00000253476 | 15.7465 | 2.0242 | ##### | 2.4213 | 1.5465E-02 | 3.0910E-02 |
| ENSG00000260352\|SMG1-DT | 17.3937 | -2.3385 | ##### | -2.4211 | 1.5472E-02 | 3.0919E-02 |
| ENSG00000212452\|SNORD69 | 9.2358 | 1.3617 | ##### | 2.4211 | 1.5472E-02 | 3.0919E-02 |
| ENSG00000119401\|TRIM32 | 2527.0521 | -0.2997 | ##### | -2.4211 | 1.5472E-02 | 3.0919E-02 |
| ENSG00000113812\|ACTR8 | 1738.5129 | -0.3045 | ##### | -2.4210 | 1.5476E-02 | 3.0924E-02 |
| ENSG00000102753\|KPNA3 | 8131.2346 | 0.2659 | ##### | 2.4210 | 1.5478E-02 | 3.0926E-02 |
| ENSG00000291259 | 38.9010 | 1.0990 | ##### | 2.4208 | 1.5487E-02 | 3.0941E-02 |
| ENSG00000226266 | 73.3553 | 0.6341 | ##### | 2.4204 | 1.5503E-02 | 3.0971E-02 |
| ENSG00000163157\|TMOD4 | 38.7890 | 1.0172 | ##### | 2.4203 | 1.5506E-02 | 3.0975E-02 |
| ENSG00000229222\|KRT18P4 | 11.7993 | 1.2568 | ##### | 2.4201 | 1.5517E-02 | 3.0995E-02 |
| ENSG00000179335\|CLK3 | 2975.9149 | 0.3078 | ##### | 2.4199 | 1.5524E-02 | 3.1007E-02 |
| ENSG00000103253\|HAGHL | 1224.7680 | -0.3334 | ##### | -2.4198 | 1.5529E-02 | 3.1016E-02 |
| ENSG00000173275\|ZNF449 | 646.7870 | 0.3454 | ##### | 2.4191 | 1.5557E-02 | 3.1070E-02 |
| ENSG00000258443 | 2.3279 | 4.9738 | ##### | 2.4190 | 1.5565E-02 | 3.1083E-02 |
| ENSG00000146463\|ZMYM4 | 8070.8536 | -0.2891 | ##### | -2.4187 | 1.5574E-02 | 3.1099E-02 |
| ENSG00000273724 | 7.7875 | 2.9632 | ##### | 2.4183 | 1.5595E-02 | 3.1139E-02 |
| ENSG00000260430 | 17.5373 | -3.0470 | ##### | -2.4178 | 1.5615E-02 | 3.1176E-02 |
| ENSG00000203711\|LINC02901 | 80.4215 | -0.5896 | ##### | -2.4175 | 1.5628E-02 | 3.1200E-02 |
| ENSG00000260953 | 15.8662 | 2.5572 | ##### | 2.4174 | 1.5631E-02 | 3.1203E-02 |
| ENSG00000286729 | 27.0256 | 0.9730 | ##### | 2.4174 | 1.5630E-02 | 3.1203E-02 |
| ENSG00000158195\|WASF2 | 13856.6169 | 0.2896 | ##### | 2.4172 | 1.5640E-02 | 3.1219E-02 |
| ENSG00000221500\|SNORD100 | 8.9876 | 1.3321 | ##### | 2.4166 | 1.5665E-02 | 3.1267E-02 |
| ENSG00000153339\|TRAPPC8 | 6448.2496 | 0.2923 | ##### | 2.4165 | 1.5671E-02 | 3.1275E-02 |
| ENSG00000196705\|ZNF431 | 496.3420 | 0.3635 | ##### | 2.4165 | 1.5671E-02 | 3.1275E-02 |
| ENSG00000152332\|UHMK1 | 15920.3374 | -0.2927 | ##### | -2.4163 | 1.5680E-02 | 3.1289E-02 |
| ENSG00000280106 | 10.7616 | -1.4404 | ##### | -2.4161 | 1.5687E-02 | 3.1302E-02 |
| ENSG00000285851 | 17.6773 | -3.1236 | ##### | -2.4156 | 1.5709E-02 | 3.1344E-02 |
| ENSG00000182175\|RGMA | 17.2877 | 2.0439 | ##### | 2.4154 | 1.5718E-02 | 3.1359E-02 |
| ENSG00000272986 | 12.1821 | -3.1418 | ##### | -2.4154 | 1.5719E-02 | 3.1360E-02 |
| ENSG00000160883\|HK3 | 28.3534 | -6.4087 | ##### | -2.4153 | 1.5724E-02 | 3.1367E-02 |
| ENSG00000231473\|RB1-DT | 15.0309 | 6.9686 | ##### | 2.4149 | 1.5740E-02 | 3.1397E-02 |
| ENSG00000256116 | 175.1891 | 0.5551 | ##### | 2.4147 | 1.5747E-02 | 3.1409E-02 |
| ENSG00000236760 | 21.2474 | 5.7080 | ##### | 2.4146 | 1.5751E-02 | 3.1415E-02 |
| ENSG00000164715\|LMTK2 | 5892.2906 | 0.3011 | ##### | 2.4144 | 1.5761E-02 | 3.1433E-02 |
| ENSG00000066654\|THUMPD1 | 4225.2544 | -0.2955 | ##### | -2.4144 | 1.5762E-02 | 3.1433E-02 |
| ENSG00000125124\|BBS2 | 3154.7420 | 0.3178 | ##### | 2.4138 | 1.5786E-02 | 3.1478E-02 |
| ENSG00000228507\|DAP3P2 | 11.7509 | 1.9662 | ##### | 2.4136 | 1.5796E-02 | 3.1495E-02 |
| ENSG00000223725 | 33.9353 | 1.3217 | ##### | 2.4136 | 1.5797E-02 | 3.1495E-02 |
| ENSG00000261634\|PAQR5-DT | 20.3024 | -2.8500 | ##### | -2.4129 | 1.5825E-02 | 3.1549E-02 |
| ENSG00000099999\|RNF215 | 1644.9756 | 0.3088 | ##### | 2.4125 | 1.5846E-02 | 3.1589E-02 |
| ENSG00000140612\|SEC11A | 6374.8740 | -0.2953 | ##### | -2.4124 | 1.5850E-02 | 3.1594E-02 |
| ENSG00000279491 | 25.9349 | 1.0334 | ##### | 2.4122 | 1.5859E-02 | 3.1610E-02 |
| ENSG00000135409\|AMHR2 | 19.0903 | -1.1830 | ##### | -2.4121 | 1.5861E-02 | 3.1612E-02 |
| ENSG00000136280\|CCM2 | 4811.7943 | -0.2837 | ##### | -2.4121 | 1.5862E-02 | 3.1613E-02 |
| ENSG00000171863\|RPS7 | 38472.5949 | 0.2565 | ##### | 2.4120 | 1.5864E-02 | 3.1614E-02 |
| ENSG00000261456\|TUBB8 | 10.2295 | 1.4135 | ##### | 2.4116 | 1.5881E-02 | 3.1647E-02 |
| ENSG00000243777 | 17.1793 | 1.7360 | ##### | 2.4114 | 1.5893E-02 | 3.1667E-02 |
| ENSG00000213722\|DDAH2 | 2594.0149 | -0.3013 | ##### | -2.4113 | 1.5894E-02 | 3.1667E-02 |
| ENSG00000141744\|PNMT | 15.1751 | -1.4133 | ##### | -2.4113 | 1.5895E-02 | 3.1668E-02 |
| ENSG00000204099\|NEU4 | 28.6800 | 0.9521 | ##### | 2.4109 | 1.5914E-02 | 3.1704E-02 |
| ENSG00000119906\|SLF2 | 3217.8531 | 0.3045 | ##### | 2.4107 | 1.5921E-02 | 3.1715E-02 |
| ENSG00000181458\|TMEM45A | 1404.5439 | -0.3218 | ##### | -2.4107 | 1.5922E-02 | 3.1716E-02 |

| | | | | | | |
|---|---|---|---|---|---|---|
| ENSG00000234636\|MED14OS | 171.7023 | -0.4018 | ##### | -2.4106 | 1.5926E-02 | 3.1722E-02 |
| ENSG00000188305\|PEAK3 | 12.3407 | -6.8151 | ##### | -2.4100 | 1.5953E-02 | 3.1773E-02 |
| ENSG00000287510 | 11.4249 | -1.7752 | ##### | -2.4099 | 1.5955E-02 | 3.1775E-02 |
| ENSG00000234851\|RPL23AP42 | 1139.3913 | -0.3124 | ##### | -2.4098 | 1.5961E-02 | 3.1782E-02 |
| ENSG00000174007\|CEP19 | 539.3635 | 0.3605 | ##### | 2.4098 | 1.5960E-02 | 3.1782E-02 |
| ENSG00000175470\|PPP2R2D | 2498.3040 | -0.2870 | ##### | -2.4097 | 1.5968E-02 | 3.1794E-02 |
| ENSG00000116157\|GPX7 | 33.2733 | 4.4380 | ##### | 2.4095 | 1.5972E-02 | 3.1801E-02 |
| ENSG00000169515\|CCDC8 | 23.0852 | 1.0420 | ##### | 2.4089 | 1.6001E-02 | 3.1857E-02 |
| ENSG00000120337\|TNFSF18 | 8.2741 | 4.9438 | ##### | 2.4085 | 1.6016E-02 | 3.1884E-02 |
| ENSG00000187017\|ESPN | 61.4169 | 0.7780 | ##### | 2.4082 | 1.6033E-02 | 3.1915E-02 |
| ENSG00000058600\|POLR3E | 5991.5928 | -0.2898 | ##### | -2.4080 | 1.6039E-02 | 3.1925E-02 |
| ENSG00000213965\|NUDT19 | 4553.9341 | -0.3017 | ##### | -2.4079 | 1.6045E-02 | 3.1934E-02 |
| ENSG00000254847 | 7.5926 | 6.1874 | ##### | 2.4075 | 1.6062E-02 | 3.1967E-02 |
| ENSG00000287908 | 517.7549 | -3.2559 | ##### | -2.4070 | 1.6085E-02 | 3.2010E-02 |
| ENSG00000240240 | 6.4904 | -3.6090 | ##### | -2.4067 | 1.6097E-02 | 3.2033E-02 |
| ENSG00000139192\|TAPBPL | 548.2228 | 0.3441 | ##### | 2.4064 | 1.6109E-02 | 3.2054E-02 |
| ENSG00000290727\|ALMS1P1 | 46.4994 | 0.9309 | ##### | 2.4063 | 1.6113E-02 | 3.2058E-02 |
| ENSG00000164164\|OTUD4 | 4496.8946 | -0.2984 | ##### | -2.4063 | 1.6113E-02 | 3.2058E-02 |
| ENSG00000260447 | 7.2031 | -4.9720 | ##### | -2.4062 | 1.6119E-02 | 3.2067E-02 |
| ENSG00000280320 | 67.9607 | -0.5161 | ##### | -2.4056 | 1.6144E-02 | 3.2115E-02 |
| ENSG00000291091\|SDHAP4 | 840.8704 | -0.3396 | ##### | -2.4055 | 1.6149E-02 | 3.2122E-02 |
| ENSG00000260464 | 102.7895 | 0.5472 | ##### | 2.4053 | 1.6157E-02 | 3.2137E-02 |
| ENSG00000218980\|FTH1P15 | 4.2039 | -2.2845 | ##### | -2.4049 | 1.6178E-02 | 3.2175E-02 |
| ENSG00000288012 | 55.9594 | -0.6662 | ##### | -2.4043 | 1.6201E-02 | 3.2220E-02 |
| ENSG00000257950\|P2RX5-TAX1BP3 | 4632.2519 | 0.3449 | ##### | 2.4042 | 1.6210E-02 | 3.2235E-02 |
| ENSG00000206596\|RNU1-27P | 3.3683 | 2.8710 | ##### | 2.4033 | 1.6248E-02 | 3.2307E-02 |
| ENSG00000053328\|METTL24 | 7.6130 | 6.1849 | ##### | 2.4033 | 1.6248E-02 | 3.2307E-02 |
| ENSG00000165807\|PPP1R36 | 81.6598 | -0.5293 | ##### | -2.4031 | 1.6259E-02 | 3.2324E-02 |
| ENSG00000113615\|SEC24A | 3493.8887 | -0.2962 | ##### | -2.4031 | 1.6259E-02 | 3.2324E-02 |
| ENSG00000283239\|KBTBD11-OT1 | 5.1216 | -5.7620 | ##### | -2.4027 | 1.6276E-02 | 3.2356E-02 |
| ENSG00000213713\|PIGCP1 | 193.6875 | 0.5611 | ##### | 2.4020 | 1.6304E-02 | 3.2409E-02 |
| ENSG00000267140 | 16.8188 | 2.1745 | ##### | 2.4019 | 1.6308E-02 | 3.2415E-02 |
| ENSG00000259086\|KIAA1143P1 | 37.1455 | 0.8691 | ##### | 2.4016 | 1.6322E-02 | 3.2438E-02 |
| ENSG00000260565\|ERVK13-1 | 480.8033 | 0.3873 | ##### | 2.4016 | 1.6322E-02 | 3.2438E-02 |
| ENSG00000162572\|SCNN1D | 450.5830 | 0.3538 | ##### | 2.4010 | 1.6352E-02 | 3.2497E-02 |
| ENSG00000188710\|QRFP | 39.6885 | 0.8597 | ##### | 2.4007 | 1.6365E-02 | 3.2519E-02 |
| ENSG00000171817\|ZNF540 | 30.8512 | -1.0748 | ##### | -2.4006 | 1.6366E-02 | 3.2520E-02 |
| ENSG00000226167\|AP4B1-AS1 | 202.6407 | 0.4798 | ##### | 2.4005 | 1.6371E-02 | 3.2528E-02 |
| ENSG00000161265\|U2AF1L4 | 888.4128 | -0.3315 | ##### | -2.4004 | 1.6376E-02 | 3.2535E-02 |
| ENSG00000130522\|JUND | 4581.4605 | 0.2851 | ##### | 2.4003 | 1.6383E-02 | 3.2547E-02 |
| ENSG00000178971\|CTC1 | 3531.3934 | 0.2946 | ##### | 2.3995 | 1.6417E-02 | 3.2612E-02 |
| ENSG00000205871\|RPS3AP47 | 17.9019 | -1.2729 | ##### | -2.3994 | 1.6422E-02 | 3.2621E-02 |
| ENSG00000275318 | 28.8830 | 1.0021 | ##### | 2.3994 | 1.6423E-02 | 3.2621E-02 |
| ENSG00000090539\|CHRD | 130.2605 | 0.4960 | ##### | 2.3989 | 1.6446E-02 | 3.2663E-02 |
| ENSG00000112893\|MAN2A1 | 3944.4239 | -0.2980 | ##### | -2.3988 | 1.6447E-02 | 3.2663E-02 |
| ENSG00000213090\|SCYL2P1 | 6.6337 | 2.8485 | ##### | 2.3988 | 1.6449E-02 | 3.2665E-02 |
| ENSG00000237278\|RLIMP2 | 27.9567 | 0.8088 | ##### | 2.3987 | 1.6452E-02 | 3.2669E-02 |
| ENSG00000230303 | 17.3645 | -2.1877 | ##### | -2.3985 | 1.6464E-02 | 3.2689E-02 |
| ENSG00000255856 | 3.7441 | -2.6812 | ##### | -2.3983 | 1.6473E-02 | 3.2706E-02 |
| ENSG00000259448\|LINC02352 | 17.4439 | -1.2086 | ##### | -2.3982 | 1.6477E-02 | 3.2712E-02 |
| ENSG00000149547\|EI24 | 10331.6409 | 0.2917 | ##### | 2.3979 | 1.6491E-02 | 3.2737E-02 |
| ENSG00000273174 | 8.1057 | 2.2851 | ##### | 2.3977 | 1.6499E-02 | 3.2752E-02 |
| ENSG00000260136 | 186.8995 | -0.4616 | ##### | -2.3975 | 1.6507E-02 | 3.2765E-02 |
| ENSG00000224295\|OLFM5P | 5.8183 | -5.4424 | ##### | -2.3973 | 1.6515E-02 | 3.2779E-02 |
| ENSG00000268628 | 8.1518 | 2.0141 | ##### | 2.3968 | 1.6538E-02 | 3.2823E-02 |
| ENSG00000267767\|LINC01801 | 8.4143 | 5.5280 | ##### | 2.3967 | 1.6542E-02 | 3.2827E-02 |
| ENSG00000244968\|LIFR-AS1 | 175.2623 | -0.4732 | ##### | -2.3965 | 1.6554E-02 | 3.2850E-02 |
| ENSG00000178568\|ERBB4 | 25.7987 | 1.3915 | ##### | 2.3961 | 1.6572E-02 | 3.2883E-02 |
| ENSG00000197647\|ZNF433 | 27.6539 | -2.9462 | ##### | -2.3960 | 1.6574E-02 | 3.2884E-02 |
| ENSG00000234996\|ACTG1P25 | 4.1029 | 2.8178 | ##### | 2.3956 | 1.6591E-02 | 3.2916E-02 |
| ENSG00000138615\|CILP | 11.7862 | -2.4394 | ##### | -2.3956 | 1.6592E-02 | 3.2916E-02 |
| ENSG00000142330\|CAPN10 | 2498.9438 | 0.3076 | ##### | 2.3954 | 1.6601E-02 | 3.2933E-02 |
| ENSG00000241061\|RPL5P1 | 11.1450 | -1.2238 | ##### | -2.3952 | 1.6610E-02 | 3.2947E-02 |

| | | | | | | |
|---|---|---|---|---|---|---|
| ENSG00000129159\|KCNC1 | 25.1809 | 2.4220 | ##### | 2.3951 | 1.6618E-02 | 3.2961E-02 |
| ENSG00000176894\|PXMP2 | 868.6976 | -0.3310 | ##### | -2.3950 | 1.6622E-02 | 3.2967E-02 |
| ENSG00000261777\|DDX19A-DT | 52.3485 | -0.7197 | ##### | -2.3949 | 1.6626E-02 | 3.2973E-02 |
| ENSG00000270020 | 9.8097 | 2.4623 | ##### | 2.3939 | 1.6673E-02 | 3.3063E-02 |
| ENSG00000288945 | 25.3284 | -0.9548 | ##### | -2.3937 | 1.6678E-02 | 3.3071E-02 |
| ENSG00000213260\|YWHAZP5 | 13.6785 | -1.1080 | ##### | -2.3935 | 1.6690E-02 | 3.3093E-02 |
| ENSG00000157111\|TMEM171 | 136.2334 | -0.5581 | ##### | -2.3932 | 1.6701E-02 | 3.3113E-02 |
| ENSG00000164458\|TBXT | 7.2057 | 4.1707 | ##### | 2.3931 | 1.6708E-02 | 3.3124E-02 |
| ENSG00000204713\|TRIM27 | 5389.5322 | 0.2471 | ##### | 2.3928 | 1.6722E-02 | 3.3150E-02 |
| ENSG00000239883\|PARGP1 | 72.5357 | 0.7792 | ##### | 2.3927 | 1.6725E-02 | 3.3153E-02 |
| ENSG00000112110\|MRPL18 | 4286.6395 | 0.2866 | ##### | 2.3925 | 1.6732E-02 | 3.3165E-02 |
| ENSG00000284968 | 62.0163 | -2.1440 | ##### | -2.3925 | 1.6734E-02 | 3.3167E-02 |
| ENSG00000108797\|CNTNAP1 | 2773.1718 | -0.3026 | ##### | -2.3918 | 1.6764E-02 | 3.3225E-02 |
| ENSG00000261240 | 3.6888 | 4.3632 | ##### | 2.3918 | 1.6767E-02 | 3.3227E-02 |
| ENSG00000267404 | 3.7404 | 3.7081 | ##### | 2.3918 | 1.6768E-02 | 3.3227E-02 |
| ENSG00000233791\|BTG2-DT | 14.5690 | 2.4334 | ##### | 2.3917 | 1.6769E-02 | 3.3228E-02 |
| ENSG00000260751 | 37.8554 | 1.0126 | ##### | 2.3911 | 1.6800E-02 | 3.3287E-02 |
| ENSG00000119862\|LGALSL | 1073.4927 | -0.3414 | ##### | -2.3909 | 1.6808E-02 | 3.3298E-02 |
| ENSG00000042832\|TG | 17.4616 | -0.9881 | ##### | -2.3909 | 1.6807E-02 | 3.3298E-02 |
| ENSG00000165917\|RAPSN | 13.9646 | 1.7844 | ##### | 2.3906 | 1.6823E-02 | 3.3325E-02 |
| ENSG00000278514 | 17.9331 | -7.9668 | ##### | -2.3903 | 1.6835E-02 | 3.3346E-02 |
| ENSG00000136636\|KCTD3 | 4591.7353 | -0.2924 | ##### | -2.3902 | 1.6837E-02 | 3.3349E-02 |
| ENSG00000267827 | 2.9996 | -4.3751 | ##### | -2.3902 | 1.6839E-02 | 3.3351E-02 |
| ENSG00000226891\|LINC01359 | 5.5398 | 3.2569 | ##### | 2.3896 | 1.6867E-02 | 3.3403E-02 |
| ENSG00000212952 | 38.0849 | -0.9555 | ##### | -2.3896 | 1.6867E-02 | 3.3403E-02 |
| ENSG00000275549\|STPG3-AS1 | 21.5074 | -0.9248 | ##### | -2.3890 | 1.6895E-02 | 3.3455E-02 |
| ENSG00000127083\|OMD | 22.0159 | -7.8751 | ##### | -2.3887 | 1.6906E-02 | 3.3475E-02 |
| ENSG00000260394\|STUB1-DT | 8.3252 | 2.7523 | ##### | 2.3879 | 1.6944E-02 | 3.3547E-02 |
| ENSG00000168246\|UBTD2 | 1817.6721 | 0.3032 | ##### | 2.3879 | 1.6945E-02 | 3.3547E-02 |
| ENSG00000227227 | 10.6614 | -1.2312 | ##### | -2.3877 | 1.6954E-02 | 3.3563E-02 |
| ENSG00000139668\|WDFY2 | 2081.7247 | -0.3056 | ##### | -2.3872 | 1.6976E-02 | 3.3601E-02 |
| ENSG00000213888\|LINC01521 | 264.0538 | -0.3845 | ##### | -2.3872 | 1.6975E-02 | 3.3601E-02 |
| ENSG00000265402\|RSL24D1P9 | 3.1152 | -4.9906 | ##### | -2.3868 | 1.6998E-02 | 3.3643E-02 |
| ENSG00000289003 | 11.4335 | -1.6334 | ##### | -2.3866 | 1.7005E-02 | 3.3654E-02 |
| ENSG00000289318 | 458.3286 | 0.3857 | ##### | 2.3863 | 1.7019E-02 | 3.3680E-02 |
| ENSG00000171862\|PTEN | 6278.6619 | 0.2286 | ##### | 2.3862 | 1.7022E-02 | 3.3684E-02 |
| ENSG00000290770\|MFSD14CP | 1929.6710 | -0.3037 | ##### | -2.3859 | 1.7037E-02 | 3.3712E-02 |
| ENSG00000077800\|FKBP6 | 9.1017 | 1.3941 | ##### | 2.3852 | 1.7071E-02 | 3.3778E-02 |
| ENSG00000117697\|NSL1 | 1934.0192 | -0.3047 | ##### | -2.3850 | 1.7079E-02 | 3.3791E-02 |
| ENSG00000156050\|FAM161B | 999.9494 | 0.7056 | ##### | 2.3846 | 1.7099E-02 | 3.3827E-02 |
| ENSG00000229848 | 37.5001 | 0.7992 | ##### | 2.3845 | 1.7101E-02 | 3.3827E-02 |
| ENSG00000171865\|RNASEH1 | 8460.9958 | 0.2888 | ##### | 2.3845 | 1.7100E-02 | 3.3827E-02 |
| ENSG00000198597\|ZNF536 | 17.6578 | 2.8612 | ##### | 2.3844 | 1.7105E-02 | 3.3833E-02 |
| ENSG00000273723\|SUGT1-DT | 48.8977 | 0.6992 | ##### | 2.3839 | 1.7132E-02 | 3.3884E-02 |
| ENSG00000228612\|HK2P1 | 6.0039 | 3.8013 | ##### | 2.3838 | 1.7135E-02 | 3.3887E-02 |
| ENSG00000156535\|CD109 | 8708.9108 | 0.2932 | ##### | 2.3835 | 1.7150E-02 | 3.3916E-02 |
| ENSG00000142530\|GARIN5A | 69.0881 | 0.8222 | ##### | 2.3834 | 1.7155E-02 | 3.3923E-02 |
| ENSG00000110492\|MDK | 1720.4879 | -0.3146 | ##### | -2.3826 | 1.7193E-02 | 3.3996E-02 |
| ENSG00000179399\|GPC5 | 8.9344 | -3.2658 | ##### | -2.3825 | 1.7196E-02 | 3.4000E-02 |
| ENSG00000259098 | 2.4204 | -4.4458 | ##### | -2.3823 | 1.7204E-02 | 3.4012E-02 |
| ENSG00000162851\|TFB2M | 3280.1058 | 0.2877 | ##### | 2.3820 | 1.7217E-02 | 3.4033E-02 |
| ENSG00000269054\|ZNF497-AS1 | 18.3310 | -1.4340 | ##### | -2.3821 | 1.7216E-02 | 3.4033E-02 |
| ENSG00000225764\|P3H2-AS1 | 10.3723 | -1.2338 | ##### | -2.3820 | 1.7218E-02 | 3.4033E-02 |
| ENSG00000081818\|PCDHB4 | 4.1726 | -3.6583 | ##### | -2.3813 | 1.7253E-02 | 3.4100E-02 |
| ENSG00000267729 | 5.8407 | 4.9230 | ##### | 2.3811 | 1.7259E-02 | 3.4111E-02 |
| ENSG00000176236\|RPP38-DT | 19.1945 | 0.9647 | ##### | 2.3810 | 1.7265E-02 | 3.4119E-02 |
| ENSG00000142182\|DNMT3L | 7.2029 | 4.6687 | ##### | 2.3806 | 1.7284E-02 | 3.4156E-02 |
| ENSG00000290707\|MED15P9 | 5.0035 | 1.9798 | ##### | 2.3806 | 1.7286E-02 | 3.4158E-02 |
| ENSG00000086570\|FAT2 | 14.6874 | 3.5190 | ##### | 2.3801 | 1.7309E-02 | 3.4200E-02 |
| ENSG00000279096 | 9.0125 | 2.7766 | ##### | 2.3799 | 1.7318E-02 | 3.4216E-02 |
| ENSG00000113578\|FGF1 | 64.9171 | -0.7366 | ##### | -2.3796 | 1.7330E-02 | 3.4237E-02 |
| ENSG00000215022 | 27.8507 | 1.1333 | ##### | 2.3793 | 1.7346E-02 | 3.4267E-02 |
| ENSG00000287778 | 5.4973 | -4.8339 | ##### | -2.3792 | 1.7348E-02 | 3.4268E-02 |

| | | | | | | |
|---|---|---|---|---|---|---|
| ENSG00000196917\|HCAR1 | 4.6336 | -3.9823 | ##### | -2.3790 | 1.7359E-02 | 3.4287E-02 |
| ENSG00000225930\|LINC02249 | 12.1324 | 2.2408 | ##### | 2.3789 | 1.7366E-02 | 3.4298E-02 |
| ENSG00000278592 | 10.2788 | -1.6875 | ##### | -2.3788 | 1.7367E-02 | 3.4298E-02 |
| ENSG00000228323\|MYADM-AS1 | 13.5860 | 1.6203 | ##### | 2.3788 | 1.7371E-02 | 3.4305E-02 |
| ENSG00000104983\|CCDC61 | 447.0769 | 0.3394 | ##### | 2.3782 | 1.7396E-02 | 3.4352E-02 |
| ENSG00000290384 | 5.1387 | 2.0471 | ##### | 2.3779 | 1.7410E-02 | 3.4377E-02 |
| ENSG00000113851\|CRBN | 974.9649 | 0.3285 | ##### | 2.3777 | 1.7419E-02 | 3.4392E-02 |
| ENSG00000170145\|SIK2 | 3379.4253 | 0.2833 | ##### | 2.3776 | 1.7424E-02 | 3.4400E-02 |
| ENSG00000257607 | 85.1076 | 0.6302 | ##### | 2.3772 | 1.7445E-02 | 3.4439E-02 |
| ENSG00000105278\|ZFR2 | 33.0185 | 1.4245 | ##### | 2.3770 | 1.7453E-02 | 3.4453E-02 |
| ENSG00000279177\|CTBP2P9 | 4.8996 | -2.5613 | ##### | -2.3769 | 1.7458E-02 | 3.4460E-02 |
| ENSG00000280078 | 8.3632 | 5.4867 | ##### | 2.3767 | 1.7468E-02 | 3.4478E-02 |
| ENSG00000099822\|HCN2 | 992.5173 | 0.3116 | ##### | 2.3762 | 1.7493E-02 | 3.4525E-02 |
| ENSG00000213881\|NPM1P6 | 20.2461 | 0.9024 | ##### | 2.3755 | 1.7524E-02 | 3.4583E-02 |
| ENSG00000225062\|CATIP-AS1 | 22.3664 | -1.2096 | ##### | -2.3755 | 1.7526E-02 | 3.4585E-02 |
| ENSG00000130164\|LDLR | 7200.2972 | -0.2870 | ##### | -2.3747 | 1.7563E-02 | 3.4654E-02 |
| ENSG00000120156\|TEK | 7.7998 | -3.1449 | ##### | -2.3747 | 1.7563E-02 | 3.4654E-02 |
| ENSG00000224127 | 16.1724 | -7.9148 | ##### | -2.3747 | 1.7565E-02 | 3.4656E-02 |
| ENSG00000158062\|UBXN11 | 1505.2096 | 0.3362 | ##### | 2.3746 | 1.7567E-02 | 3.4658E-02 |
| ENSG00000198961\|PJA2 | 5782.0696 | 0.2964 | ##### | 2.3743 | 1.7583E-02 | 3.4687E-02 |
| ENSG00000245748 | 2890.9237 | 0.3018 | ##### | 2.3737 | 1.7612E-02 | 3.4740E-02 |
| ENSG00000137073\|UBAP2 | 4582.7639 | 0.2948 | ##### | 2.3737 | 1.7612E-02 | 3.4740E-02 |
| ENSG00000137955\|RABGGTB | 5254.3724 | 0.2793 | ##### | 2.3735 | 1.7619E-02 | 3.4748E-02 |
| ENSG00000267397 | 7.9504 | -3.0252 | ##### | -2.3735 | 1.7618E-02 | 3.4748E-02 |
| ENSG00000119636\|BBOF1 | 410.7960 | -0.3684 | ##### | -2.3734 | 1.7627E-02 | 3.4761E-02 |
| ENSG00000224769\|MUC20P1 | 13.7494 | -2.0524 | ##### | -2.3732 | 1.7633E-02 | 3.4771E-02 |
| ENSG00000067082\|KLF6 | 13207.9909 | 0.2819 | ##### | 2.3730 | 1.7644E-02 | 3.4791E-02 |
| ENSG00000271851 | 100.1327 | 0.7030 | ##### | 2.3727 | 1.7659E-02 | 3.4819E-02 |
| ENSG00000238171 | 39.9579 | -1.1013 | ##### | -2.3726 | 1.7664E-02 | 3.4825E-02 |
| ENSG00000209482\|SNORD83A | 19.2555 | 0.8711 | ##### | 2.3725 | 1.7669E-02 | 3.4834E-02 |
| ENSG00000168137\|SETD5 | 7169.7619 | 0.2790 | ##### | 2.3724 | 1.7671E-02 | 3.4834E-02 |
| ENSG00000068784\|SRBD1 | 1847.9271 | -0.3090 | ##### | -2.3724 | 1.7673E-02 | 3.4836E-02 |
| ENSG00000228809 | 7.2481 | 5.6213 | ##### | 2.3724 | 1.7674E-02 | 3.4836E-02 |
| ENSG00000047662\|FAM184B | 36.6202 | 0.9529 | ##### | 2.3723 | 1.7677E-02 | 3.4839E-02 |
| ENSG00000289727 | 101.1194 | -0.6108 | ##### | -2.3722 | 1.7682E-02 | 3.4848E-02 |
| ENSG00000272902\|TBC1D8-AS1 | 107.3128 | -0.5643 | ##### | -2.3721 | 1.7690E-02 | 3.4860E-02 |
| ENSG00000236792 | 26.1215 | -1.0709 | ##### | -2.3717 | 1.7705E-02 | 3.4887E-02 |
| ENSG00000173991\|TCAP | 52.8132 | 0.7353 | ##### | 2.3710 | 1.7738E-02 | 3.4951E-02 |
| ENSG00000137656\|BUD13 | 1407.1695 | 0.3002 | ##### | 2.3707 | 1.7755E-02 | 3.4982E-02 |
| ENSG00000288564 | 42.3516 | 7.9003 | ##### | 2.3703 | 1.7776E-02 | 3.5020E-02 |
| ENSG00000228218\|ATF4P3 | 25.8771 | -0.8790 | ##### | -2.3701 | 1.7784E-02 | 3.5034E-02 |
| ENSG00000005421\|PON1 | 14.2460 | -3.3430 | ##### | -2.3699 | 1.7795E-02 | 3.5054E-02 |
| ENSG00000010244\|ZNF207 | 17275.7905 | 0.2522 | ##### | 2.3697 | 1.7804E-02 | 3.5069E-02 |
| ENSG00000285599 | 57.0915 | -0.6814 | ##### | -2.3694 | 1.7815E-02 | 3.5088E-02 |
| ENSG00000137513\|NARS2 | 1923.7540 | 0.3054 | ##### | 2.3694 | 1.7818E-02 | 3.5093E-02 |
| ENSG00000279464 | 13.6957 | 2.1419 | ##### | 2.3692 | 1.7828E-02 | 3.5109E-02 |
| ENSG00000125651\|GTF2F1 | 8218.2818 | 0.2060 | ##### | 2.3676 | 1.7906E-02 | 3.5260E-02 |
| ENSG00000256006 | 10.1011 | 1.6044 | ##### | 2.3675 | 1.7909E-02 | 3.5264E-02 |
| ENSG00000287867 | 3.1177 | -5.5401 | ##### | -2.3674 | 1.7913E-02 | 3.5271E-02 |
| ENSG00000271849\|MAN2A1-DT | 3.8959 | 4.6984 | ##### | 2.3674 | 1.7915E-02 | 3.5272E-02 |
| ENSG00000091436\|MAP3K20 | 11821.3119 | 0.3149 | ##### | 2.3672 | 1.7925E-02 | 3.5289E-02 |
| ENSG00000135338\|LCA5 | 207.3875 | -0.4820 | ##### | -2.3665 | 1.7957E-02 | 3.5349E-02 |
| ENSG00000287619 | 13.9468 | 2.2297 | ##### | 2.3664 | 1.7960E-02 | 3.5353E-02 |
| ENSG00000287506 | 27.8432 | -7.8857 | ##### | -2.3663 | 1.7966E-02 | 3.5362E-02 |
| ENSG00000168959\|GRM5 | 30.5153 | 1.0459 | ##### | 2.3662 | 1.7974E-02 | 3.5373E-02 |
| ENSG00000060762\|MPC1 | 415.4825 | -0.3904 | ##### | -2.3662 | 1.7974E-02 | 3.5373E-02 |
| ENSG00000151657\|KIN | 1289.4667 | 0.3330 | ##### | 2.3656 | 1.8003E-02 | 3.5428E-02 |
| ENSG00000213599\|SLX1A-SULT1A3 | 1768.9833 | 0.5636 | ##### | 2.3639 | 1.8084E-02 | 3.5586E-02 |
| ENSG00000158467\|AHCYL2 | 2395.6632 | 0.2866 | ##### | 2.3636 | 1.8098E-02 | 3.5611E-02 |
| ENSG00000181817\|LSM10 | 2617.0063 | -0.2979 | ##### | -2.3635 | 1.8102E-02 | 3.5616E-02 |
| ENSG00000267077 | 7.7191 | -3.4576 | ##### | -2.3631 | 1.8122E-02 | 3.5652E-02 |
| ENSG00000233560\|KRT8P39 | 21.5434 | 1.0470 | ##### | 2.3622 | 1.8165E-02 | 3.5735E-02 |
| ENSG00000214922\|HLA-F-AS1 | 59.6400 | -0.8353 | ##### | -2.3622 | 1.8168E-02 | 3.5739E-02 |

| | | | | | | |
|---|---|---|---|---|---|---|
| ENSG00000224614\|TNK2-AS1 | 21.0003 | 1.1983 | ##### | 2.3620 | 1.8175E-02 | 3.5750E-02 |
| ENSG00000251615 | 68.0717 | -0.6386 | ##### | -2.3620 | 1.8179E-02 | 3.5756E-02 |
| ENSG00000275029\|HMGB1P24 | 4.1099 | 2.0871 | ##### | 2.3618 | 1.8185E-02 | 3.5765E-02 |
| ENSG00000196420\|S100A5 | 32.4915 | 1.8287 | ##### | 2.3603 | 1.8260E-02 | 3.5910E-02 |
| ENSG00000228519\|RANBP1P1 | 19.8573 | 0.9158 | ##### | 2.3598 | 1.8286E-02 | 3.5959E-02 |
| ENSG00000115944\|COX7A2L | 8040.2106 | 0.2875 | ##### | 2.3597 | 1.8289E-02 | 3.5962E-02 |
| ENSG00000277744 | 10.8666 | -2.0327 | ##### | -2.3597 | 1.8290E-02 | 3.5962E-02 |
| ENSG00000163141\|BNIPL | 44.1126 | 0.9099 | ##### | 2.3595 | 1.8301E-02 | 3.5981E-02 |
| ENSG00000203469 | 20.5431 | 1.3763 | ##### | 2.3591 | 1.8317E-02 | 3.6010E-02 |
| ENSG00000273680 | 38.9957 | 1.0139 | ##### | 2.3585 | 1.8347E-02 | 3.6067E-02 |
| ENSG00000261513 | 11.0924 | -2.1034 | ##### | -2.3582 | 1.8365E-02 | 3.6099E-02 |
| ENSG00000289702 | 2.1940 | 4.2015 | ##### | 2.3573 | 1.8407E-02 | 3.6180E-02 |
| ENSG00000247151\|CSTF3-DT | 26.2949 | 0.8947 | ##### | 2.3569 | 1.8430E-02 | 3.6224E-02 |
| ENSG00000267120 | 114.8016 | -1.1077 | ##### | -2.3567 | 1.8439E-02 | 3.6239E-02 |
| ENSG00000114030\|KPNA1 | 10743.4469 | 0.1459 | ##### | 2.3556 | 1.8492E-02 | 3.6339E-02 |
| ENSG00000260651 | 19.1626 | 1.0959 | ##### | 2.3556 | 1.8492E-02 | 3.6339E-02 |
| ENSG00000269918 | 43.2374 | 1.1135 | ##### | 2.3554 | 1.8503E-02 | 3.6358E-02 |
| ENSG00000106367\|AP1S1 | 8007.1753 | -0.2777 | ##### | -2.3547 | 1.8538E-02 | 3.6424E-02 |
| ENSG00000231969\|MMADHC-DT | 29.0172 | 1.0440 | ##### | 2.3545 | 1.8547E-02 | 3.6440E-02 |
| ENSG00000236297\|EEF1A1P23 | 6.3021 | -2.3217 | ##### | -2.3544 | 1.8550E-02 | 3.6443E-02 |
| ENSG00000115827\|DCAF17 | 2051.1863 | 0.3036 | ##### | 2.3539 | 1.8575E-02 | 3.6489E-02 |
| ENSG00000166682\|TMPRSS5 | 12.8002 | 1.6038 | ##### | 2.3534 | 1.8601E-02 | 3.6536E-02 |
| ENSG00000173083\|HPSE | 768.3870 | 0.3301 | ##### | 2.3534 | 1.8602E-02 | 3.6536E-02 |
| ENSG00000271924\|RNA5SP108 | 5.6631 | 4.8946 | ##### | 2.3534 | 1.8605E-02 | 3.6539E-02 |
| ENSG00000243176 | 186.4518 | 0.4261 | ##### | 2.3533 | 1.8605E-02 | 3.6539E-02 |
| ENSG00000164219\|PGGT1B | 1988.7153 | -0.3008 | ##### | -2.3531 | 1.8618E-02 | 3.6561E-02 |
| ENSG00000255138\|GLTPP1 | 6.7484 | 4.8673 | ##### | 2.3528 | 1.8634E-02 | 3.6591E-02 |
| ENSG00000185414\|MRPL30 | 6563.8133 | -0.2891 | ##### | -2.3522 | 1.8662E-02 | 3.6643E-02 |
| ENSG00000166971\|AKTIP | 1634.5833 | -0.3247 | ##### | -2.3521 | 1.8665E-02 | 3.6647E-02 |
| ENSG00000255152\|MSH5-SAPCD1 | 515.5049 | 0.3653 | ##### | 2.3518 | 1.8681E-02 | 3.6675E-02 |
| ENSG00000233426\|EIF3FP3 | 551.2453 | 0.3640 | ##### | 2.3515 | 1.8698E-02 | 3.6706E-02 |
| ENSG00000261669 | 6.5535 | 4.9170 | ##### | 2.3513 | 1.8708E-02 | 3.6724E-02 |
| ENSG00000129204\|USP6 | 44.9547 | 0.7595 | ##### | 2.3495 | 1.8801E-02 | 3.6903E-02 |
| ENSG00000152315\|KCNK13 | 29.2132 | -3.0393 | ##### | -2.3492 | 1.8815E-02 | 3.6928E-02 |
| ENSG00000266441\|ARHGAP28-AS1 | 10.6799 | -1.2400 | ##### | -2.3492 | 1.8816E-02 | 3.6928E-02 |
| ENSG00000133246\|PRAM1 | 22.7927 | -6.0661 | ##### | -2.3491 | 1.8818E-02 | 3.6929E-02 |
| ENSG00000173674\|EIF1AX | 9971.6763 | 0.2695 | ##### | 2.3490 | 1.8826E-02 | 3.6942E-02 |
| ENSG00000234424 | 13.3074 | -1.9822 | ##### | -2.3478 | 1.8882E-02 | 3.7051E-02 |
| ENSG00000280202 | 11.0598 | -3.7634 | ##### | -2.3478 | 1.8885E-02 | 3.7053E-02 |
| ENSG00000254812 | 13.1433 | 1.2103 | ##### | 2.3474 | 1.8905E-02 | 3.7091E-02 |
| ENSG00000140740\|UQCRC2 | 13893.6173 | -0.2642 | ##### | -2.3473 | 1.8910E-02 | 3.7098E-02 |
| ENSG00000117419\|ERI3 | 5597.1894 | 0.2888 | ##### | 2.3468 | 1.8938E-02 | 3.7150E-02 |
| ENSG00000283267\|FAM237B | 18.9134 | -7.6553 | ##### | -2.3467 | 1.8939E-02 | 3.7150E-02 |
| ENSG00000100320\|RBFOX2 | 10420.1003 | 0.2793 | ##### | 2.3462 | 1.8967E-02 | 3.7202E-02 |
| ENSG00000172115\|CYCS | 30993.9887 | -0.2803 | ##### | -2.3460 | 1.8977E-02 | 3.7220E-02 |
| ENSG00000262294\|RPH3AL-AS2 | 23.0056 | -6.9328 | ##### | -2.3459 | 1.8981E-02 | 3.7226E-02 |
| ENSG00000241170\|RPL31P46 | 6.0416 | -1.7233 | ##### | -2.3459 | 1.8984E-02 | 3.7228E-02 |
| ENSG00000113971\|NPHP3 | 1791.4651 | 0.3636 | ##### | 2.3457 | 1.8993E-02 | 3.7244E-02 |
| ENSG00000278993 | 97.1349 | 0.5781 | ##### | 2.3445 | 1.9052E-02 | 3.7357E-02 |
| ENSG00000256196 | 30.7979 | 0.8742 | ##### | 2.3444 | 1.9055E-02 | 3.7361E-02 |
| ENSG00000257027 | 77.9535 | 0.6500 | ##### | 2.3443 | 1.9065E-02 | 3.7378E-02 |
| ENSG00000289540 | 23.2952 | 1.0519 | ##### | 2.3437 | 1.9094E-02 | 3.7433E-02 |
| ENSG00000178082\|TWF1P1 | 616.1581 | 0.3344 | ##### | 2.3432 | 1.9119E-02 | 3.7478E-02 |
| ENSG00000139083\|ETV6 | 2655.2355 | 0.3017 | ##### | 2.3425 | 1.9155E-02 | 3.7547E-02 |
| ENSG00000259244 | 9.3682 | -2.3421 | ##### | -2.3425 | 1.9156E-02 | 3.7547E-02 |
| ENSG00000288877 | 11.9122 | 1.3162 | ##### | 2.3424 | 1.9158E-02 | 3.7548E-02 |
| ENSG00000086712\|TXLNG | 2965.3445 | 0.2638 | ##### | 2.3424 | 1.9160E-02 | 3.7549E-02 |
| ENSG00000226756 | 8.3834 | -3.6302 | ##### | -2.3421 | 1.9176E-02 | 3.7579E-02 |
| ENSG00000289474 | 4.4885 | 2.6481 | ##### | 2.3416 | 1.9202E-02 | 3.7625E-02 |
| ENSG00000197448\|GSTK1 | 5334.6912 | -0.2897 | ##### | -2.3416 | 1.9203E-02 | 3.7625E-02 |
| ENSG00000233588\|CYP51A1P2 | 34.4912 | -0.8619 | ##### | -2.3410 | 1.9230E-02 | 3.7677E-02 |
| ENSG00000244607\|CCDC13 | 30.4467 | 2.9618 | ##### | 2.3409 | 1.9236E-02 | 3.7685E-02 |
| ENSG00000095464\|PDE6C | 4.8491 | 5.2601 | ##### | 2.3406 | 1.9253E-02 | 3.7716E-02 |

| | | | | | | |
|---|---|---|---|---|---|---|
| ENSG00000165916\|PSMC3 | 11393.6029 | 0.2117 | ##### | 2.3399 | 1.9291E-02 | 3.7788E-02 |
| ENSG00000290514 | 6.9861 | -3.6403 | ##### | -2.3390 | 1.9337E-02 | 3.7876E-02 |
| ENSG00000289082 | 5.3459 | 5.1779 | ##### | 2.3387 | 1.9349E-02 | 3.7898E-02 |
| ENSG00000260144 | 17.0312 | 7.7943 | ##### | 2.3385 | 1.9361E-02 | 3.7919E-02 |
| ENSG00000185933\|CALHM1 | 10.9110 | 2.5795 | ##### | 2.3384 | 1.9368E-02 | 3.7930E-02 |
| ENSG00000107614\|TRDMT1 | 575.8856 | 0.3422 | ##### | 2.3379 | 1.9392E-02 | 3.7974E-02 |
| ENSG00000163634\|THOC7 | 2934.7313 | -0.2980 | ##### | -2.3378 | 1.9396E-02 | 3.7980E-02 |
| ENSG00000167281\|RBFOX3 | 54.5866 | 0.8881 | ##### | 2.3377 | 1.9404E-02 | 3.7991E-02 |
| ENSG00000277581 | 12.2792 | 2.6365 | ##### | 2.3377 | 1.9403E-02 | 3.7991E-02 |
| ENSG00000198754\|OXCT2 | 110.6801 | 0.5038 | ##### | 2.3376 | 1.9407E-02 | 3.7993E-02 |
| ENSG00000233136\|USP17L11 | 6.8681 | 4.8798 | ##### | 2.3376 | 1.9409E-02 | 3.7995E-02 |
| ENSG00000141349\|G6PC3 | 6986.8447 | 0.2927 | ##### | 2.3375 | 1.9414E-02 | 3.8003E-02 |
| ENSG00000177508\|IRX3 | 2218.9195 | 0.2936 | ##### | 2.3371 | 1.9432E-02 | 3.8035E-02 |
| ENSG00000265474 | 117.4942 | 0.5626 | ##### | 2.3370 | 1.9440E-02 | 3.8048E-02 |
| ENSG00000205221\|VIT | 40.6531 | 4.4836 | ##### | 2.3370 | 1.9441E-02 | 3.8048E-02 |
| ENSG00000279544 | 6.5072 | 3.8826 | ##### | 2.3365 | 1.9465E-02 | 3.8092E-02 |
| ENSG00000283154\|IQCJ-SCHIP1 | 1079.8037 | -0.3296 | ##### | -2.3357 | 1.9506E-02 | 3.8169E-02 |
| ENSG00000293220\|LINC01399 | 5.0632 | -3.8165 | ##### | -2.3356 | 1.9513E-02 | 3.8181E-02 |
| ENSG00000112782\|CLIC5 | 6.9729 | 6.0616 | ##### | 2.3345 | 1.9570E-02 | 3.8291E-02 |
| ENSG00000267092 | 27.1429 | 0.9069 | ##### | 2.3343 | 1.9580E-02 | 3.8307E-02 |
| ENSG00000289211 | 8.8844 | -1.8848 | ##### | -2.3341 | 1.9590E-02 | 3.8324E-02 |
| ENSG00000092020\|PPP2R3C | 2271.2342 | 0.3021 | ##### | 2.3339 | 1.9599E-02 | 3.8339E-02 |
| ENSG00000162669\|HFM1 | 14.5923 | -2.7485 | ##### | -2.3333 | 1.9634E-02 | 3.8404E-02 |
| ENSG00000233547 | 13.0329 | -1.1985 | ##### | -2.3326 | 1.9667E-02 | 3.8467E-02 |
| ENSG00000151502\|VPS26B | 3453.9499 | -0.2335 | ##### | -2.3318 | 1.9713E-02 | 3.8555E-02 |
| ENSG00000230457\|PA2G4P4 | 361.6666 | 0.3476 | ##### | 2.3316 | 1.9719E-02 | 3.8565E-02 |
| ENSG00000164815\|ORC5 | 2682.6294 | -0.2953 | ##### | -2.3313 | 1.9735E-02 | 3.8593E-02 |
| ENSG00000188428\|BLOC1S5 | 881.8407 | 0.3177 | ##### | 2.3304 | 1.9784E-02 | 3.8686E-02 |
| ENSG00000239697\|TNFSF12 | 578.5206 | -0.3467 | ##### | -2.3299 | 1.9812E-02 | 3.8735E-02 |
| ENSG00000114302\|PRKAR2A | 8872.9370 | -0.2784 | ##### | -2.3299 | 1.9812E-02 | 3.8735E-02 |
| ENSG00000229212 | 4.7601 | 2.7962 | ##### | 2.3295 | 1.9830E-02 | 3.8769E-02 |
| ENSG00000267080\|ASB16-AS1 | 695.7766 | 0.3732 | ##### | 2.3295 | 1.9835E-02 | 3.8775E-02 |
| ENSG00000105371\|ICAM4 | 5.4477 | 1.8110 | ##### | 2.3290 | 1.9857E-02 | 3.8817E-02 |
| ENSG00000057294\|PKP2 | 1138.4543 | -0.3054 | ##### | -2.3290 | 1.9860E-02 | 3.8820E-02 |
| ENSG00000223947 | 76.8434 | -0.5267 | ##### | -2.3286 | 1.9882E-02 | 3.8859E-02 |
| ENSG00000117625\|RCOR3 | 1580.0680 | -0.2831 | ##### | -2.3285 | 1.9884E-02 | 3.8861E-02 |
| ENSG00000162613\|FUBP1 | 10532.9823 | -0.2565 | ##### | -2.3283 | 1.9895E-02 | 3.8881E-02 |
| ENSG00000100376\|FAM118A | 1112.5859 | -0.3189 | ##### | -2.3278 | 1.9922E-02 | 3.8929E-02 |
| ENSG00000258763 | 1.9699 | 4.6629 | ##### | 2.3276 | 1.9931E-02 | 3.8946E-02 |
| ENSG00000278970\|HEIH | 1632.0482 | -0.3181 | ##### | -2.3267 | 1.9979E-02 | 3.9036E-02 |
| ENSG00000185420\|SMYD3 | 779.3164 | 0.3307 | ##### | 2.3266 | 1.9989E-02 | 3.9053E-02 |
| ENSG00000261556\|SMG1P7 | 94.4215 | -0.5075 | ##### | -2.3263 | 2.0003E-02 | 3.9077E-02 |
| ENSG00000136152\|COG3 | 2146.8729 | -0.3022 | ##### | -2.3261 | 2.0015E-02 | 3.9099E-02 |
| ENSG00000242156 | 8.0524 | 3.6889 | ##### | 2.3257 | 2.0032E-02 | 3.9130E-02 |
| ENSG00000289877 | 13.5560 | 2.6857 | ##### | 2.3254 | 2.0053E-02 | 3.9168E-02 |
| ENSG00000021762\|OSBPL5 | 1005.7462 | 0.3106 | ##### | 2.3251 | 2.0064E-02 | 3.9188E-02 |
| ENSG00000169857\|AVEN | 1075.3840 | -0.1970 | ##### | -2.3250 | 2.0071E-02 | 3.9198E-02 |
| ENSG00000280087 | 16.0681 | -1.0681 | ##### | -2.3248 | 2.0085E-02 | 3.9223E-02 |
| ENSG00000279789 | 130.5380 | 0.5082 | ##### | 2.3241 | 2.0121E-02 | 3.9291E-02 |
| ENSG00000104365\|IKBKB | 2720.5667 | 0.2869 | ##### | 2.3238 | 2.0139E-02 | 3.9323E-02 |
| ENSG00000189167\|ZAR1L | 3.0618 | 4.8490 | ##### | 2.3232 | 2.0170E-02 | 3.9378E-02 |
| ENSG00000173077\|DELEC1 | 7.5606 | 1.8695 | ##### | 2.3232 | 2.0169E-02 | 3.9378E-02 |
| ENSG00000244041\|LINC01011 | 165.4709 | 0.4370 | ##### | 2.3227 | 2.0193E-02 | 3.9421E-02 |
| ENSG00000227113 | 8.6908 | 4.9174 | ##### | 2.3226 | 2.0200E-02 | 3.9429E-02 |
| ENSG00000258116\|PPIAP45 | 7.2732 | -4.9598 | ##### | -2.3226 | 2.0200E-02 | 3.9429E-02 |
| ENSG00000176029\|C11orf16 | 6.1135 | -3.0878 | ##### | -2.3226 | 2.0203E-02 | 3.9433E-02 |
| ENSG00000074855\|ANO8 | 1875.7769 | 0.2973 | ##### | 2.3224 | 2.0214E-02 | 3.9451E-02 |
| ENSG00000255390\|DCHS1-AS1 | 3.3729 | -2.3076 | ##### | -2.3219 | 2.0237E-02 | 3.9495E-02 |
| ENSG00000167920\|KRT10-AS1 | 1138.4797 | -0.3163 | ##### | -2.3219 | 2.0241E-02 | 3.9499E-02 |
| ENSG00000285458\|C4orf36 | 56.4966 | -2.1141 | ##### | -2.3211 | 2.0283E-02 | 3.9578E-02 |
| ENSG00000064652\|SNX24 | 1512.3590 | -0.3176 | ##### | -2.3210 | 2.0289E-02 | 3.9588E-02 |
| ENSG00000107831\|FGF8 | 6.1564 | -5.5348 | ##### | -2.3206 | 2.0309E-02 | 3.9619E-02 |
| ENSG00000092470\|WDR76 | 2353.5780 | 0.2838 | ##### | 2.3206 | 2.0307E-02 | 3.9619E-02 |

| | | | | | | |
|---|---|---|---|---|---|---|
| ENSG00000287262 | 16.2072 | -2.3526 | ##### | -2.3206 | 2.0307E-02 | 3.9619E-02 |
| ENSG00000289385 | 11.8352 | 2.4692 | ##### | 2.3201 | 2.0334E-02 | 3.9663E-02 |
| ENSG00000213413\|PVRIG | 35.0037 | 7.7315 | ##### | 2.3201 | 2.0333E-02 | 3.9663E-02 |
| ENSG00000259209 | 18.3398 | 1.8063 | ##### | 2.3191 | 2.0392E-02 | 3.9772E-02 |
| ENSG00000261618\|LINC02605 | 7.8725 | -3.0356 | ##### | -2.3189 | 2.0398E-02 | 3.9783E-02 |
| ENSG00000278771\|RN7SL3 | 188.3301 | 1.1634 | ##### | 2.3185 | 2.0423E-02 | 3.9825E-02 |
| ENSG00000158516\|CPA2 | 3.5587 | 5.1851 | ##### | 2.3185 | 2.0422E-02 | 3.9825E-02 |
| ENSG00000262075\|DKFZP434A062 | 11.2328 | -1.9449 | ##### | -2.3185 | 2.0424E-02 | 3.9825E-02 |
| ENSG00000253167\|WASHC5-AS1 | 9.1556 | 2.4355 | ##### | 2.3175 | 2.0478E-02 | 3.9928E-02 |
| ENSG00000232160\|RAP2C-AS1 | 128.1324 | 0.4974 | ##### | 2.3174 | 2.0479E-02 | 3.9928E-02 |
| ENSG00000218350\|LYPLA1P3 | 38.4348 | -0.8755 | ##### | -2.3172 | 2.0492E-02 | 3.9950E-02 |
| ENSG00000172478\|MAB21L4 | 7.6165 | -3.1113 | ##### | -2.3172 | 2.0495E-02 | 3.9954E-02 |
| ENSG00000278899 | 39.9095 | 0.7077 | ##### | 2.3167 | 2.0522E-02 | 4.0003E-02 |
| ENSG00000260634 | 15.2704 | 2.5063 | ##### | 2.3166 | 2.0528E-02 | 4.0011E-02 |
| ENSG00000186230\|ZNF749 | 749.9935 | -0.3125 | ##### | -2.3165 | 2.0533E-02 | 4.0019E-02 |
| ENSG00000207650\|MIR570 | 14.0469 | 1.0048 | ##### | 2.3164 | 2.0535E-02 | 4.0022E-02 |
| ENSG00000213205\|STRADBP1 | 14.5238 | -7.1009 | ##### | -2.3151 | 2.0607E-02 | 4.0159E-02 |
| ENSG00000265145\|SNORD53 | 4.0129 | 4.4295 | ##### | 2.3148 | 2.0623E-02 | 4.0184E-02 |
| ENSG00000245067\|IGFBP7-AS1 | 12.9532 | -1.9348 | ##### | -2.3148 | 2.0622E-02 | 4.0184E-02 |
| ENSG00000139574\|NPFF | 48.6686 | 0.6643 | ##### | 2.3148 | 2.0626E-02 | 4.0188E-02 |
| ENSG00000141452\|RMC1 | 2963.8698 | -0.2206 | ##### | -2.3146 | 2.0635E-02 | 4.0202E-02 |
| ENSG00000234806\|RPL26P29 | 4.2689 | -4.9371 | ##### | -2.3145 | 2.0642E-02 | 4.0210E-02 |
| ENSG00000149346\|SLX4IP | 599.7598 | 0.3307 | ##### | 2.3145 | 2.0641E-02 | 4.0210E-02 |
| ENSG00000280408 | 5.5823 | 4.9963 | ##### | 2.3142 | 2.0659E-02 | 4.0241E-02 |
| ENSG00000288701\|PRRC2B | 24143.2058 | -0.2799 | ##### | -2.3139 | 2.0671E-02 | 4.0262E-02 |
| ENSG00000157399\|ARSL | 7.3775 | 5.7875 | ##### | 2.3136 | 2.0689E-02 | 4.0294E-02 |
| ENSG00000213996\|TM6SF2 | 34.9749 | 1.7832 | ##### | 2.3134 | 2.0699E-02 | 4.0306E-02 |
| ENSG00000250221\|KRT8P32 | 4.3934 | -1.7460 | ##### | -2.3134 | 2.0699E-02 | 4.0306E-02 |
| ENSG00000186407\|CD300E | 4.5988 | 5.4544 | ##### | 2.3134 | 2.0698E-02 | 4.0306E-02 |
| ENSG00000250910\|MAP9-AS1 | 8.6930 | -1.2113 | ##### | -2.3129 | 2.0731E-02 | 4.0366E-02 |
| ENSG00000289685 | 24.0649 | 6.9191 | ##### | 2.3120 | 2.0779E-02 | 4.0456E-02 |
| ENSG00000286501 | 7.4082 | 5.7877 | ##### | 2.3118 | 2.0790E-02 | 4.0476E-02 |
| ENSG00000105428\|ZNRF4 | 6.8562 | 6.0321 | ##### | 2.3114 | 2.0809E-02 | 4.0510E-02 |
| ENSG00000275454 | 770.3353 | 0.3294 | ##### | 2.3114 | 2.0812E-02 | 4.0512E-02 |
| ENSG00000153989\|NUS1 | 8610.4667 | 0.2808 | ##### | 2.3112 | 2.0824E-02 | 4.0534E-02 |
| ENSG00000162607\|USP1 | 8494.8031 | 0.2811 | ##### | 2.3103 | 2.0872E-02 | 4.0625E-02 |
| ENSG00000134243\|SORT1 | 2059.0910 | 0.2937 | ##### | 2.3092 | 2.0934E-02 | 4.0742E-02 |
| ENSG00000203943\|SAMD13 | 102.2831 | 0.5776 | ##### | 2.3089 | 2.0950E-02 | 4.0771E-02 |
| ENSG00000289729 | 7.7248 | 4.3438 | ##### | 2.3085 | 2.0970E-02 | 4.0806E-02 |
| ENSG00000248569 | 71.3582 | -0.5426 | ##### | -2.3082 | 2.0988E-02 | 4.0840E-02 |
| ENSG00000154710\|RABGEF1 | 3537.7628 | -0.2904 | ##### | -2.3078 | 2.1010E-02 | 4.0880E-02 |
| ENSG00000184984\|CHRM5 | 4.0614 | 4.3503 | ##### | 2.3076 | 2.1021E-02 | 4.0899E-02 |
| ENSG00000185721\|DRG1 | 5388.0317 | 0.2396 | ##### | 2.3056 | 2.1134E-02 | 4.1116E-02 |
| ENSG00000197054\|ZNF763 | 8.0462 | 2.9205 | ##### | 2.3055 | 2.1141E-02 | 4.1126E-02 |
| ENSG00000164626\|KCNK5 | 1231.4266 | 0.3111 | ##### | 2.3054 | 2.1143E-02 | 4.1128E-02 |
| ENSG00000276434 | 2.7184 | -3.8772 | ##### | -2.3050 | 2.1169E-02 | 4.1176E-02 |
| ENSG00000267952 | 46.8302 | -2.2362 | ##### | -2.3048 | 2.1180E-02 | 4.1192E-02 |
| ENSG00000231686\|ANKRD11P2 | 172.5644 | -0.4169 | ##### | -2.3048 | 2.1180E-02 | 4.1192E-02 |
| ENSG00000235079\|ZRANB2-AS1 | 11.9232 | -1.6365 | ##### | -2.3042 | 2.1209E-02 | 4.1246E-02 |
| ENSG00000293205\|TPI1P2 | 140.8600 | 0.4689 | ##### | 2.3042 | 2.1213E-02 | 4.1250E-02 |
| ENSG00000160255\|ITGB2 | 58.9476 | -0.6841 | ##### | -2.3041 | 2.1217E-02 | 4.1254E-02 |
| ENSG00000287394\|MAGEA3-DT | 318.8401 | -0.4173 | ##### | -2.3041 | 2.1217E-02 | 4.1254E-02 |
| ENSG00000110436\|SLC1A2 | 16.1947 | 1.9432 | ##### | 2.3040 | 2.1222E-02 | 4.1261E-02 |
| ENSG00000200795\|RNU4-1 | 6.6234 | 4.6463 | ##### | 2.3036 | 2.1243E-02 | 4.1299E-02 |
| ENSG00000284740 | 3.5431 | 3.0202 | ##### | 2.3034 | 2.1258E-02 | 4.1325E-02 |
| ENSG00000116977\|LGALS8 | 2612.6571 | -0.3079 | ##### | -2.3032 | 2.1266E-02 | 4.1338E-02 |
| ENSG00000293494\|ZPLD2P | 4.6262 | 4.9604 | ##### | 2.3031 | 2.1272E-02 | 4.1347E-02 |
| ENSG00000112367\|FIG4 | 1037.2541 | -0.3144 | ##### | -2.3030 | 2.1279E-02 | 4.1358E-02 |
| ENSG00000272916 | 590.2503 | 0.4499 | ##### | 2.3029 | 2.1286E-02 | 4.1369E-02 |
| ENSG00000234129\|HCCS-DT | 47.9778 | 0.7178 | ##### | 2.3026 | 2.1301E-02 | 4.1395E-02 |
| ENSG00000274620\|MIR378J | 4.0414 | 3.8517 | ##### | 2.3026 | 2.1303E-02 | 4.1396E-02 |
| ENSG00000267165\|CHMP1B-AS1 | 360.2975 | 0.2518 | ##### | 2.3025 | 2.1309E-02 | 4.1405E-02 |
| ENSG00000088766\|CRLS1 | 3062.9047 | 0.2819 | ##### | 2.3020 | 2.1334E-02 | 4.1451E-02 |

| | | | | | | |
|---|---|---|---|---|---|---|
| ENSG00000275481 | 17.0701 | 1.2390 | ##### | 2.3018 | 2.1344E-02 | 4.1467E-02 |
| ENSG00000277128 | 28.0780 | 0.9543 | ##### | 2.3018 | 2.1348E-02 | 4.1472E-02 |
| ENSG00000147383\|NSDHL | 3289.0446 | 0.2797 | ##### | 2.3017 | 2.1354E-02 | 4.1482E-02 |
| ENSG00000119919\|NKX2-3 | 10.7036 | 1.5638 | ##### | 2.3009 | 2.1396E-02 | 4.1561E-02 |
| ENSG00000114841\|DNAH1 | 278.7902 | -0.3728 | ##### | -2.3005 | 2.1419E-02 | 4.1603E-02 |
| ENSG00000279873\|LINC01126 | 26.9570 | 0.8303 | ##### | 2.2998 | 2.1459E-02 | 4.1678E-02 |
| ENSG00000293144 | 6.0143 | -2.4314 | ##### | -2.2998 | 2.1461E-02 | 4.1679E-02 |
| ENSG00000112146\|FBXO9 | 3863.4193 | -0.2713 | ##### | -2.2996 | 2.1474E-02 | 4.1700E-02 |
| ENSG00000275131\|PDE4DIPP2 | 313.8619 | -0.3742 | ##### | -2.2995 | 2.1475E-02 | 4.1701E-02 |
| ENSG00000270210 | 17.1143 | 1.3096 | ##### | 2.2994 | 2.1482E-02 | 4.1710E-02 |
| ENSG00000236810\|ELOA-AS1 | 128.8069 | -0.5277 | ##### | -2.2989 | 2.1508E-02 | 4.1759E-02 |
| ENSG00000197021\|EOLA2 | 1723.7339 | 0.3002 | ##### | 2.2989 | 2.1511E-02 | 4.1762E-02 |
| ENSG00000162438\|CTRC | 24.1039 | -6.5792 | ##### | -2.2988 | 2.1516E-02 | 4.1767E-02 |
| ENSG00000259595\|CTDSPL2-DT | 23.2080 | -0.9939 | ##### | -2.2988 | 2.1516E-02 | 4.1767E-02 |
| ENSG00000185000\|DGAT1 | 3045.3517 | 0.2812 | ##### | 2.2984 | 2.1542E-02 | 4.1813E-02 |
| ENSG00000252254\|Y_RNA | 3.4507 | 4.5297 | ##### | 2.2980 | 2.1561E-02 | 4.1849E-02 |
| ENSG00000018408\|WWTR1 | 10352.2028 | -0.2756 | ##### | -2.2978 | 2.1571E-02 | 4.1865E-02 |
| ENSG00000257139\|LINC02821 | 26.2056 | -0.7925 | ##### | -2.2977 | 2.1580E-02 | 4.1880E-02 |
| ENSG00000229422 | 44.2405 | -0.7350 | ##### | -2.2971 | 2.1615E-02 | 4.1944E-02 |
| ENSG00000232852\|CICP4 | 8.0651 | 1.4834 | ##### | 2.2969 | 2.1623E-02 | 4.1958E-02 |
| ENSG00000093217\|XYLB | 631.0474 | 0.3117 | ##### | 2.2968 | 2.1629E-02 | 4.1963E-02 |
| ENSG00000221740\|SNORD93 | 5.8445 | 1.6841 | ##### | 2.2968 | 2.1628E-02 | 4.1963E-02 |
| ENSG00000231233\|CFAP58-DT | 28.7910 | -1.1429 | ##### | -2.2965 | 2.1649E-02 | 4.2000E-02 |
| ENSG00000150667\|FSIP1 | 124.7536 | 0.6108 | ##### | 2.2963 | 2.1658E-02 | 4.2016E-02 |
| ENSG00000123453\|SARDH | 8.8331 | -2.0651 | ##### | -2.2962 | 2.1667E-02 | 4.2030E-02 |
| ENSG00000272969 | 17.6719 | 1.3333 | ##### | 2.2958 | 2.1687E-02 | 4.2065E-02 |
| ENSG00000273338 | 4.4779 | 4.9071 | ##### | 2.2948 | 2.1744E-02 | 4.2170E-02 |
| ENSG00000095139\|ARCN1 | 14963.9517 | -0.2595 | ##### | -2.2948 | 2.1742E-02 | 4.2170E-02 |
| ENSG00000174844\|DNAH12 | 6.9153 | -2.2669 | ##### | -2.2945 | 2.1763E-02 | 4.2199E-02 |
| ENSG00000002079\|MYH16 | 10.0650 | -2.4513 | ##### | -2.2945 | 2.1761E-02 | 4.2199E-02 |
| ENSG00000128585\|MKLN1 | 9415.9675 | -0.2700 | ##### | -2.2945 | 2.1763E-02 | 4.2199E-02 |
| ENSG00000157933\|SKI | 5442.3979 | -0.2704 | ##### | -2.2940 | 2.1791E-02 | 4.2252E-02 |
| ENSG00000138796\|HADH | 2016.6408 | -0.3018 | ##### | -2.2928 | 2.1859E-02 | 4.2380E-02 |
| ENSG00000050426\|LETMD1 | 2574.4017 | 0.3004 | ##### | 2.2926 | 2.1870E-02 | 4.2399E-02 |
| ENSG00000107890\|ANKRD26 | 1211.2678 | 0.3029 | ##### | 2.2922 | 2.1894E-02 | 4.2442E-02 |
| ENSG00000171759\|PAH | 16.2829 | -3.7436 | ##### | -2.2920 | 2.1906E-02 | 4.2460E-02 |
| ENSG00000259783\|LINC02259 | 16.5315 | -7.4594 | ##### | -2.2920 | 2.1906E-02 | 4.2460E-02 |
| ENSG00000176956\|LY6H | 8.7256 | 3.9707 | ##### | 2.2918 | 2.1918E-02 | 4.2481E-02 |
| ENSG00000250903\|GMDS-DT | 128.5241 | -0.5075 | ##### | -2.2917 | 2.1925E-02 | 4.2492E-02 |
| ENSG00000233956\|BTF3P6 | 5.5006 | 4.4432 | ##### | 2.2914 | 2.1938E-02 | 4.2513E-02 |
| ENSG00000230832\|SSBL4P | 15.9543 | 1.8750 | ##### | 2.2907 | 2.1980E-02 | 4.2592E-02 |
| ENSG00000100483\|VCPKMT | 646.5385 | 0.3263 | ##### | 2.2901 | 2.2018E-02 | 4.2660E-02 |
| ENSG00000175426\|PCSK1 | 91.8937 | 0.5274 | ##### | 2.2901 | 2.2017E-02 | 4.2660E-02 |
| ENSG00000169193\|CCDC126 | 610.2954 | -0.3493 | ##### | -2.2898 | 2.2035E-02 | 4.2691E-02 |
| ENSG00000256234\|ITPR2-AS1 | 3.3660 | 2.2470 | ##### | 2.2896 | 2.2047E-02 | 4.2711E-02 |
| ENSG00000170677\|SOCS6 | 5785.4384 | -0.2735 | ##### | -2.2892 | 2.2068E-02 | 4.2746E-02 |
| ENSG00000289443 | 24.6771 | 0.8796 | ##### | 2.2892 | 2.2066E-02 | 4.2746E-02 |
| ENSG00000171311\|EXOSC1 | 2062.9138 | -0.2964 | ##### | -2.2889 | 2.2087E-02 | 4.2780E-02 |
| ENSG00000258744 | 202.2677 | -0.4620 | ##### | -2.2881 | 2.2131E-02 | 4.2863E-02 |
| ENSG00000232581 | 126.4153 | -0.5007 | ##### | -2.2880 | 2.2138E-02 | 4.2873E-02 |
| ENSG00000130921\|MTRFR | 2426.2926 | 0.2958 | ##### | 2.2877 | 2.2154E-02 | 4.2902E-02 |
| ENSG00000283390\|PET100P1 | 52.5013 | 0.6661 | ##### | 2.2871 | 2.2191E-02 | 4.2970E-02 |
| ENSG00000185652\|NTF3 | 6.1696 | -6.0305 | ##### | -2.2870 | 2.2198E-02 | 4.2981E-02 |
| ENSG00000259814 | 7.0866 | 4.9961 | ##### | 2.2866 | 2.2220E-02 | 4.3022E-02 |
| ENSG00000178718\|RPP25 | 3103.7239 | 0.2894 | ##### | 2.2865 | 2.2225E-02 | 4.3026E-02 |
| ENSG00000198417\|MT1F | 140.7271 | -0.5231 | ##### | -2.2865 | 2.2226E-02 | 4.3026E-02 |
| ENSG00000286952 | 16.8819 | -6.7296 | ##### | -2.2862 | 2.2244E-02 | 4.3058E-02 |
| ENSG00000280119 | 7.9166 | 2.6707 | ##### | 2.2858 | 2.2265E-02 | 4.3098E-02 |
| ENSG00000099785\|MARCHF2 | 733.5320 | -0.3107 | ##### | -2.2855 | 2.2284E-02 | 4.3131E-02 |
| ENSG00000151093\|OXSM | 861.8475 | 0.3096 | ##### | 2.2841 | 2.2363E-02 | 4.3282E-02 |
| ENSG00000126456\|IRF3 | 6160.2024 | -0.2803 | ##### | -2.2840 | 2.2370E-02 | 4.3291E-02 |
| ENSG00000156475\|PPP2R2B | 16.6960 | 5.6336 | ##### | 2.2837 | 2.2391E-02 | 4.3330E-02 |
| ENSG00000116793\|PHTF1 | 2708.8314 | 0.2859 | ##### | 2.2836 | 2.2397E-02 | 4.3338E-02 |

| | | | | | | |
|---|---|---|---|---|---|---|
| ENSG00000256341 | 17.8581 | -1.2846 | ##### | -2.2835 | 2.2401E-02 | 4.3343E-02 |
| ENSG00000076706\|MCAM | 3261.2865 | -0.2890 | ##### | -2.2835 | 2.2403E-02 | 4.3345E-02 |
| ENSG00000261192\|RNF126P1 | 5.5016 | 2.8305 | ##### | 2.2832 | 2.2416E-02 | 4.3367E-02 |
| ENSG00000267149 | 16.6034 | 1.0281 | ##### | 2.2831 | 2.2424E-02 | 4.3380E-02 |
| ENSG00000137876\|RSL24D1 | 5766.0346 | 0.2832 | ##### | 2.2825 | 2.2458E-02 | 4.3443E-02 |
| ENSG00000182872\|RBM10 | 5656.9657 | 0.2705 | ##### | 2.2823 | 2.2471E-02 | 4.3465E-02 |
| ENSG00000179627\|ZBTB42 | 567.7968 | 0.3253 | ##### | 2.2819 | 2.2497E-02 | 4.3512E-02 |
| ENSG00000140968\|IRF8 | 26.1708 | 1.9373 | ##### | 2.2818 | 2.2500E-02 | 4.3515E-02 |
| ENSG00000272057\|ANKH-DT | 8.7430 | 4.6848 | ##### | 2.2812 | 2.2539E-02 | 4.3589E-02 |
| ENSG00000213218\|CSH2 | 8.9830 | 4.5559 | ##### | 2.2811 | 2.2544E-02 | 4.3595E-02 |
| ENSG00000235121 | 16.4952 | 6.7894 | ##### | 2.2809 | 2.2557E-02 | 4.3616E-02 |
| ENSG00000118508\|RAB32 | 2030.3138 | -0.2971 | ##### | -2.2803 | 2.2593E-02 | 4.3683E-02 |
| ENSG00000187037\|GPR141 | 24.7676 | -6.5304 | ##### | -2.2801 | 2.2599E-02 | 4.3693E-02 |
| ENSG00000179750\|APOBEC3B | 1481.4058 | 0.2957 | ##### | 2.2797 | 2.2625E-02 | 4.3736E-02 |
| ENSG00000254370 | 2.5275 | -5.1080 | ##### | -2.2797 | 2.2625E-02 | 4.3736E-02 |
| ENSG00000144182\|LIPT1 | 473.4584 | 0.3478 | ##### | 2.2797 | 2.2628E-02 | 4.3741E-02 |
| ENSG00000278518 | 8.6534 | 3.1550 | ##### | 2.2794 | 2.2644E-02 | 4.3768E-02 |
| ENSG00000289990 | 4.0019 | -5.4007 | ##### | -2.2792 | 2.2653E-02 | 4.3782E-02 |
| ENSG00000245888\|NSMCE1-DT | 9.3036 | 2.0257 | ##### | 2.2790 | 2.2665E-02 | 4.3803E-02 |
| ENSG00000223305\|RN7SKP30 | 9.7313 | 2.3189 | ##### | 2.2788 | 2.2677E-02 | 4.3824E-02 |
| ENSG00000100865\|CINP | 2425.0005 | -0.2724 | ##### | -2.2786 | 2.2689E-02 | 4.3843E-02 |
| ENSG00000158014\|SLC30A2 | 19.5950 | -6.6989 | ##### | -2.2786 | 2.2690E-02 | 4.3844E-02 |
| ENSG00000166105\|GLB1L3 | 34.3086 | 1.6500 | ##### | 2.2785 | 2.2696E-02 | 4.3852E-02 |
| ENSG00000174963\|ZIC4 | 236.9643 | -0.3749 | ##### | -2.2780 | 2.2726E-02 | 4.3907E-02 |
| ENSG00000287979 | 5.8260 | -2.8669 | ##### | -2.2776 | 2.2748E-02 | 4.3943E-02 |
| ENSG00000125733\|TRIP10 | 7070.7385 | -0.2122 | ##### | -2.2776 | 2.2748E-02 | 4.3943E-02 |
| ENSG00000196636\|SDHAF3 | 1590.2716 | 0.2879 | ##### | 2.2775 | 2.2759E-02 | 4.3962E-02 |
| ENSG00000154124\|OTULIN | 3436.5127 | -0.2808 | ##### | -2.2768 | 2.2800E-02 | 4.4039E-02 |
| ENSG00000268069 | 65.6943 | -0.9159 | ##### | -2.2765 | 2.2815E-02 | 4.4063E-02 |
| ENSG00000288797 | 7.1995 | 6.6012 | ##### | 2.2765 | 2.2816E-02 | 4.4063E-02 |
| ENSG00000169618\|PROKR1 | 7.5370 | 2.9957 | ##### | 2.2760 | 2.2844E-02 | 4.4115E-02 |
| ENSG00000146386\|ABRACL | 771.4895 | -0.3339 | ##### | -2.2757 | 2.2863E-02 | 4.4149E-02 |
| ENSG00000293147\|SLC9A7P1 | 447.1985 | -0.3818 | ##### | -2.2749 | 2.2914E-02 | 4.4241E-02 |
| ENSG00000171316\|CHD7 | 2867.8730 | -0.2894 | ##### | -2.2749 | 2.2914E-02 | 4.4241E-02 |
| ENSG00000033627\|ATP6V0A1 | 6084.6609 | 0.2124 | ##### | 2.2747 | 2.2922E-02 | 4.4254E-02 |
| ENSG00000287434 | 3.4398 | -4.8991 | ##### | -2.2746 | 2.2930E-02 | 4.4266E-02 |
| ENSG00000265354\|TIMM23 | 5875.9434 | -0.2785 | ##### | -2.2745 | 2.2936E-02 | 4.4276E-02 |
| ENSG00000259657\|PIGHP1 | 5.2492 | 2.1949 | ##### | 2.2745 | 2.2938E-02 | 4.4276E-02 |
| ENSG00000141504\|SAT2 | 1088.9586 | -0.3115 | ##### | -2.2739 | 2.2972E-02 | 4.4340E-02 |
| ENSG00000116819\|TFAP2E | 54.7979 | -1.4455 | ##### | -2.2738 | 2.2977E-02 | 4.4344E-02 |
| ENSG00000188868\|ZNF563 | 13.4468 | -1.7970 | ##### | -2.2738 | 2.2977E-02 | 4.4344E-02 |
| ENSG00000268713 | 3.9036 | 2.2173 | ##### | 2.2735 | 2.2993E-02 | 4.4370E-02 |
| ENSG00000080224\|EPHA6 | 117.0916 | 0.5063 | ##### | 2.2735 | 2.2994E-02 | 4.4370E-02 |
| ENSG00000140694\|PARN | 4700.1136 | 0.2759 | ##### | 2.2733 | 2.3006E-02 | 4.4388E-02 |
| ENSG00000087157\|PGS1 | 2397.4978 | 0.2718 | ##### | 2.2733 | 2.3005E-02 | 4.4388E-02 |
| ENSG00000288683 | 42.4232 | -2.1755 | ##### | -2.2733 | 2.3010E-02 | 4.4392E-02 |
| ENSG00000248871\|TNFSF12-TNFSF13 | 148.1979 | 3.6939 | ##### | 2.2731 | 2.3018E-02 | 4.4406E-02 |
| ENSG00000169989\|TIGD4 | 32.1556 | -0.8352 | ##### | -2.2727 | 2.3045E-02 | 4.4455E-02 |
| ENSG00000175087\|PDIK1L | 462.0980 | -0.3310 | ##### | -2.2727 | 2.3047E-02 | 4.4455E-02 |
| ENSG00000267355\|RPL9P29 | 10.2996 | 1.6869 | ##### | 2.2722 | 2.3072E-02 | 4.4501E-02 |
| ENSG00000131242\|RAB11FIP4 | 359.7917 | 0.3622 | ##### | 2.2720 | 2.3084E-02 | 4.4520E-02 |
| ENSG00000237178\|H3P7 | 13.8742 | -7.2058 | ##### | -2.2720 | 2.3085E-02 | 4.4521E-02 |
| ENSG00000019549\|SNAI2 | 223.4875 | -0.4201 | ##### | -2.2719 | 2.3094E-02 | 4.4535E-02 |
| ENSG00000150337\|FCGR1A | 20.9678 | 1.1605 | ##### | 2.2717 | 2.3107E-02 | 4.4556E-02 |
| ENSG00000180089\|TMEM86B | 254.8542 | 0.4214 | ##### | 2.2713 | 2.3127E-02 | 4.4592E-02 |
| ENSG00000204267\|TAP2 | 10320.2033 | 0.2745 | ##### | 2.2711 | 2.3138E-02 | 4.4611E-02 |
| ENSG00000273175 | 3033.0129 | 0.3509 | ##### | 2.2710 | 2.3150E-02 | 4.4630E-02 |
| ENSG00000169251\|NMD3 | 8061.2430 | -0.2310 | ##### | -2.2705 | 2.3175E-02 | 4.4677E-02 |
| ENSG00000151376\|ME3 | 1080.8109 | -0.3052 | ##### | -2.2697 | 2.3227E-02 | 4.4770E-02 |
| ENSG00000011677\|GABRA3 | 543.4031 | -0.3376 | ##### | -2.2697 | 2.3226E-02 | 4.4770E-02 |
| ENSG00000280809\|LINC00836 | 14.0393 | 7.5643 | ##### | 2.2695 | 2.3240E-02 | 4.4792E-02 |
| ENSG00000254373 | 3.4021 | -2.6413 | ##### | -2.2694 | 2.3243E-02 | 4.4795E-02 |
| ENSG00000293446 | 12.5284 | 6.0622 | ##### | 2.2693 | 2.3252E-02 | 4.4810E-02 |

| | | | | | | |
|---|---|---|---|---|---|---|
| ENSG00000258430 | 861.9244 | 0.3066 | ##### | 2.2692 | 2.3256E-02 | 4.4814E-02 |
| ENSG00000134825\|TMEM258 | 3917.2618 | 0.2811 | ##### | 2.2685 | 2.3299E-02 | 4.4896E-02 |
| ENSG00000290615\|RAB6C-AS1 | 5.7594 | 2.2393 | ##### | 2.2679 | 2.3333E-02 | 4.4957E-02 |
| ENSG00000139549\|DHH | 11.9863 | 2.5906 | ##### | 2.2679 | 2.3336E-02 | 4.4961E-02 |
| ENSG00000224687\|RASAL2-AS1 | 93.8561 | -0.7203 | ##### | -2.2677 | 2.3349E-02 | 4.4982E-02 |
| ENSG00000269794 | 4.9302 | -2.4771 | ##### | -2.2673 | 2.3372E-02 | 4.5024E-02 |
| ENSG00000267422 | 73.0697 | -0.6735 | ##### | -2.2671 | 2.3381E-02 | 4.5039E-02 |
| ENSG00000223969 | 6.4760 | 5.9501 | ##### | 2.2671 | 2.3384E-02 | 4.5042E-02 |
| ENSG00000233448\|PMS2P9 | 15.5448 | -0.9697 | ##### | -2.2670 | 2.3390E-02 | 4.5050E-02 |
| ENSG00000249565\|SERBP1P5 | 545.9619 | -0.3274 | ##### | -2.2670 | 2.3393E-02 | 4.5052E-02 |
| ENSG00000289437 | 12.1974 | -2.1935 | ##### | -2.2668 | 2.3402E-02 | 4.5068E-02 |
| ENSG00000160799\|CCDC12 | 1895.4337 | 0.2836 | ##### | 2.2665 | 2.3419E-02 | 4.5098E-02 |
| ENSG00000287743 | 18.4933 | -7.5554 | ##### | -2.2664 | 2.3428E-02 | 4.5111E-02 |
| ENSG00000131732\|ZCCHC9 | 2202.8021 | 0.2846 | ##### | 2.2661 | 2.3445E-02 | 4.5142E-02 |
| ENSG00000081985\|IL12RB2 | 29.9417 | 1.6762 | ##### | 2.2660 | 2.3451E-02 | 4.5149E-02 |
| ENSG00000162998\|FRZB | 22.8376 | 3.0899 | ##### | 2.2655 | 2.3481E-02 | 4.5205E-02 |
| ENSG00000131238\|PPT1 | 11961.4173 | -0.2277 | ##### | -2.2654 | 2.3490E-02 | 4.5218E-02 |
| ENSG00000255082\|GRM5-AS1 | 69.2191 | 0.6394 | ##### | 2.2653 | 2.3495E-02 | 4.5225E-02 |
| ENSG00000289130 | 2.9428 | 3.4463 | ##### | 2.2652 | 2.3498E-02 | 4.5228E-02 |
| ENSG00000278668 | 11.3874 | 4.1129 | ##### | 2.2651 | 2.3505E-02 | 4.5240E-02 |
| ENSG00000103043\|VAC14 | 5084.0751 | -0.2790 | ##### | -2.2651 | 2.3508E-02 | 4.5241E-02 |
| ENSG00000171612\|SLC25A33 | 2116.4774 | 0.2877 | ##### | 2.2648 | 2.3527E-02 | 4.5276E-02 |
| ENSG00000251189 | 2.9713 | 4.8094 | ##### | 2.2644 | 2.3551E-02 | 4.5318E-02 |
| ENSG00000172216\|CEBPB | 26373.7988 | 0.2689 | ##### | 2.2638 | 2.3587E-02 | 4.5386E-02 |
| ENSG00000168925\|CTRB1 | 8.5521 | 1.8667 | ##### | 2.2633 | 2.3620E-02 | 4.5445E-02 |
| ENSG00000199172\|MIR331 | 4.5489 | 5.1657 | ##### | 2.2631 | 2.3630E-02 | 4.5462E-02 |
| ENSG00000272886\|DCP1A | 2714.6782 | 0.2886 | ##### | 2.2630 | 2.3636E-02 | 4.5470E-02 |
| ENSG00000182541\|LIMK2 | 927.5575 | 0.3144 | ##### | 2.2628 | 2.3649E-02 | 4.5492E-02 |
| ENSG00000032389\|EIPR1 | 3264.0028 | -0.2219 | ##### | -2.2626 | 2.3663E-02 | 4.5517E-02 |
| ENSG00000118518\|RNF146 | 1399.3327 | -0.2924 | ##### | -2.2621 | 2.3690E-02 | 4.5565E-02 |
| ENSG00000139567\|ACVRL1 | 24.6393 | -1.5248 | ##### | -2.2621 | 2.3693E-02 | 4.5569E-02 |
| ENSG00000110756\|HPS5 | 5413.7678 | 0.2782 | ##### | 2.2607 | 2.3776E-02 | 4.5723E-02 |
| ENSG00000177335\|LINC02904 | 12.9936 | -7.1126 | ##### | -2.2607 | 2.3775E-02 | 4.5723E-02 |
| ENSG00000235448\|LURAP1L-AS1 | 4.8866 | 5.0361 | ##### | 2.2606 | 2.3786E-02 | 4.5739E-02 |
| ENSG00000289791 | 22.8374 | -1.5993 | ##### | -2.2603 | 2.3804E-02 | 4.5770E-02 |
| ENSG00000126705\|AHDC1 | 2625.7504 | 0.2817 | ##### | 2.2601 | 2.3815E-02 | 4.5786E-02 |
| ENSG00000145934\|TENM2 | 76.5723 | -0.8332 | ##### | -2.2601 | 2.3815E-02 | 4.5786E-02 |
| ENSG00000289317 | 7.1213 | 2.0772 | ##### | 2.2598 | 2.3833E-02 | 4.5817E-02 |
| ENSG00000161016\|RPL8 | ########## | 0.2694 | ##### | 2.2598 | 2.3835E-02 | 4.5817E-02 |
| ENSG00000232640 | 13.1159 | -1.0331 | ##### | -2.2596 | 2.3848E-02 | 4.5841E-02 |
| ENSG00000175318\|GRAMD2A | 27.0738 | 1.9451 | ##### | 2.2591 | 2.3876E-02 | 4.5890E-02 |
| ENSG00000266261 | 59.4420 | -1.8228 | ##### | -2.2590 | 2.3882E-02 | 4.5900E-02 |
| ENSG00000197099\|HRG-AS1 | 4.5547 | -2.6044 | ##### | -2.2587 | 2.3905E-02 | 4.5941E-02 |
| ENSG00000151062\|CACNA2D4 | 12.4091 | 1.7243 | ##### | 2.2576 | 2.3969E-02 | 4.6061E-02 |
| ENSG00000090060\|PAPOLA | 16194.1543 | 0.2680 | ##### | 2.2571 | 2.3999E-02 | 4.6116E-02 |
| ENSG00000225039\|LINC01058 | 22.2745 | 1.7767 | ##### | 2.2571 | 2.4001E-02 | 4.6117E-02 |
| ENSG00000216624\|GAPDHP72 | 10.7966 | 1.1727 | ##### | 2.2566 | 2.4034E-02 | 4.6177E-02 |
| ENSG00000272374\|SCUBE3-AS1 | 26.2687 | 0.8215 | ##### | 2.2564 | 2.4045E-02 | 4.6195E-02 |
| ENSG00000267360 | 5.9277 | -1.8658 | ##### | -2.2561 | 2.4062E-02 | 4.6225E-02 |
| ENSG00000128268\|MGAT3 | 111.4586 | 0.5983 | ##### | 2.2560 | 2.4070E-02 | 4.6237E-02 |
| ENSG00000132207\|SLX1A | 1751.5025 | 0.7778 | ##### | 2.2559 | 2.4077E-02 | 4.6247E-02 |
| ENSG00000181392\|SYNE4 | 40.2961 | -0.8803 | ##### | -2.2558 | 2.4081E-02 | 4.6253E-02 |
| ENSG00000006062\|MAP3K14 | 20366.9384 | 0.2673 | ##### | 2.2552 | 2.4118E-02 | 4.6321E-02 |
| ENSG00000289415 | 43.5143 | -0.6688 | ##### | -2.2549 | 2.4141E-02 | 4.6362E-02 |
| ENSG00000188419\|CHM | 3918.2727 | 0.2499 | ##### | 2.2544 | 2.4169E-02 | 4.6413E-02 |
| ENSG00000037241\|RPL26L1 | 3023.4799 | 0.2877 | ##### | 2.2540 | 2.4199E-02 | 4.6467E-02 |
| ENSG00000177646\|ACAD9 | 4562.1166 | 0.2650 | ##### | 2.2539 | 2.4203E-02 | 4.6472E-02 |
| ENSG00000234292 | 15.4793 | 1.5182 | ##### | 2.2539 | 2.4206E-02 | 4.6474E-02 |
| ENSG00000117400\|MPL | 31.7946 | -1.5453 | ##### | -2.2538 | 2.4212E-02 | 4.6482E-02 |
| ENSG00000165533\|TTC8 | 1394.0856 | -0.2873 | ##### | -2.2534 | 2.4233E-02 | 4.6520E-02 |
| ENSG00000160683\|CXCR5 | 1626.3930 | -0.4403 | ##### | -2.2532 | 2.4247E-02 | 4.6544E-02 |
| ENSG00000213246\|SUPT4H1 | 6433.5845 | 0.2839 | ##### | 2.2528 | 2.4269E-02 | 4.6584E-02 |
| ENSG00000286070 | 223.6141 | -2.5216 | ##### | -2.2526 | 2.4286E-02 | 4.6614E-02 |

| | | | | | | |
|---|---|---|---|---|---|---|
| ENSG00000142538\|PTH2 | 4.1541 | -5.9550 | ##### | -2.2522 | 2.4310E-02 | 4.6655E-02 |
| ENSG00000212607\|SNORA3B | 9.0660 | 1.2251 | ##### | 2.2517 | 2.4344E-02 | 4.6719E-02 |
| ENSG00000231665\|OGFOD1P1 | 13.9024 | -7.2095 | ##### | -2.2516 | 2.4351E-02 | 4.6728E-02 |
| ENSG00000293410\|CASP16P | 15.3063 | 2.1062 | ##### | 2.2512 | 2.4370E-02 | 4.6763E-02 |
| ENSG00000136699\|SMPD4 | 13917.5895 | -0.2692 | ##### | -2.2512 | 2.4373E-02 | 4.6764E-02 |
| ENSG00000227195\|MIR663AHG | 4.0033 | 3.0664 | ##### | 2.2504 | 2.4421E-02 | 4.6855E-02 |
| ENSG00000233009\|NALCN-AS1 | 2.8314 | -3.3891 | ##### | -2.2504 | 2.4427E-02 | 4.6859E-02 |
| ENSG00000254332\|TPM4P3 | 16.3181 | -0.9640 | ##### | -2.2504 | 2.4426E-02 | 4.6859E-02 |
| ENSG00000183742\|MACC1 | 24.3127 | -6.8054 | ##### | -2.2502 | 2.4438E-02 | 4.6877E-02 |
| ENSG00000223734\|LAPTM4A-DT | 7.2972 | 2.8177 | ##### | 2.2501 | 2.4442E-02 | 4.6882E-02 |
| ENSG00000232117\|LINC00384 | 6.3355 | 5.9222 | ##### | 2.2495 | 2.4478E-02 | 4.6948E-02 |
| ENSG00000233098\|CCDC144NL-AS1 | 167.4751 | -0.4257 | ##### | -2.2494 | 2.4487E-02 | 4.6962E-02 |
| ENSG00000225975\|ZNF567-DT | 81.7707 | -0.6984 | ##### | -2.2485 | 2.4546E-02 | 4.7073E-02 |
| ENSG00000165828\|PRAP1 | 5.2151 | -4.0201 | ##### | -2.2471 | 2.4636E-02 | 4.7242E-02 |
| ENSG00000048540\|LMO3 | 16.0750 | -2.6966 | ##### | -2.2470 | 2.4639E-02 | 4.7245E-02 |
| ENSG00000161981\|SNRNP25 | 1937.5373 | -0.3029 | ##### | -2.2470 | 2.4641E-02 | 4.7246E-02 |
| ENSG00000158411\|MITD1 | 2215.2638 | -0.2829 | ##### | -2.2467 | 2.4662E-02 | 4.7283E-02 |
| ENSG00000249216 | 5.7101 | -5.4232 | ##### | -2.2466 | 2.4667E-02 | 4.7289E-02 |
| ENSG00000236423\|LINC01134 | 87.6286 | 0.6349 | ##### | 2.2463 | 2.4683E-02 | 4.7318E-02 |
| ENSG00000253159\|PCDHGA12 | 4.6716 | -3.1048 | ##### | -2.2460 | 2.4704E-02 | 4.7354E-02 |
| ENSG00000272763\|LINC03057 | 8.4638 | -2.7338 | ##### | -2.2460 | 2.4707E-02 | 4.7357E-02 |
| ENSG00000165478\|HEPACAM | 6.3537 | 5.9258 | ##### | 2.2455 | 2.4736E-02 | 4.7409E-02 |
| ENSG00000214967\|NPIPA7 | 534.0035 | 0.3908 | ##### | 2.2454 | 2.4742E-02 | 4.7418E-02 |
| ENSG00000256537\|SMIM10L1 | 4809.8538 | 0.2743 | ##### | 2.2453 | 2.4747E-02 | 4.7424E-02 |
| ENSG00000179271\|GADD45GIP1 | 6763.4333 | 0.2728 | ##### | 2.2452 | 2.4753E-02 | 4.7434E-02 |
| ENSG00000146085\|MMUT | 1592.0618 | 0.2851 | ##### | 2.2451 | 2.4765E-02 | 4.7453E-02 |
| ENSG00000267633 | 67.6918 | 0.5251 | ##### | 2.2450 | 2.4771E-02 | 4.7461E-02 |
| ENSG00000290603\|CEACAM22P | 9.0584 | -2.8514 | ##### | -2.2443 | 2.4810E-02 | 4.7534E-02 |
| ENSG00000267127 | 20.4622 | 0.9716 | ##### | 2.2443 | 2.4816E-02 | 4.7542E-02 |
| ENSG00000275139 | 14.1759 | 5.7030 | ##### | 2.2436 | 2.4859E-02 | 4.7620E-02 |
| ENSG00000141551\|CSNK1D | 19592.7207 | 0.1884 | ##### | 2.2431 | 2.4888E-02 | 4.7673E-02 |
| ENSG00000143443\|C1orf56 | 1057.9227 | 0.3066 | ##### | 2.2430 | 2.4899E-02 | 4.7691E-02 |
| ENSG00000251409 | 12.1970 | -7.0200 | ##### | -2.2429 | 2.4904E-02 | 4.7697E-02 |
| ENSG00000236679\|RPL23AP24 | 11.9082 | 1.1161 | ##### | 2.2426 | 2.4923E-02 | 4.7732E-02 |
| ENSG00000212163\|SNORD91A | 5.0267 | -4.3106 | ##### | -2.2421 | 2.4958E-02 | 4.7792E-02 |
| ENSG00000187987\|ZSCAN23 | 55.2041 | 0.8243 | ##### | 2.2421 | 2.4957E-02 | 4.7792E-02 |
| ENSG00000169021\|UQCRFS1 | 10175.5783 | -0.2589 | ##### | -2.2413 | 2.5004E-02 | 4.7878E-02 |
| ENSG00000124562\|SNRPC | 6950.2297 | 0.2789 | ##### | 2.2412 | 2.5015E-02 | 4.7896E-02 |
| ENSG00000184659\|FOXD4L4 | 4.6527 | 2.8723 | ##### | 2.2409 | 2.5035E-02 | 4.7931E-02 |
| ENSG00000261335 | 17.7285 | -0.9404 | ##### | -2.2405 | 2.5060E-02 | 4.7975E-02 |
| ENSG00000147364\|FBXO25 | 719.1527 | 0.3153 | ##### | 2.2396 | 2.5119E-02 | 4.8085E-02 |
| ENSG00000172543\|CTSW | 8.2098 | 2.5021 | ##### | 2.2393 | 2.5135E-02 | 4.8112E-02 |
| ENSG00000235529\|AGAP1-IT1 | 30.0141 | -0.8021 | ##### | -2.2389 | 2.5159E-02 | 4.8156E-02 |
| ENSG00000157765\|SLC34A2 | 19.4379 | -3.3103 | ##### | -2.2388 | 2.5166E-02 | 4.8166E-02 |
| ENSG00000215861 | 175.1388 | 0.4036 | ##### | 2.2387 | 2.5174E-02 | 4.8178E-02 |
| ENSG00000266086 | 519.0302 | 1.9067 | ##### | 2.2386 | 2.5180E-02 | 4.8186E-02 |
| ENSG00000270866 | 5.7502 | -5.4307 | ##### | -2.2384 | 2.5193E-02 | 4.8209E-02 |
| ENSG00000107821\|KAZALD1 | 1841.9298 | -0.2945 | ##### | -2.2384 | 2.5197E-02 | 4.8213E-02 |
| ENSG00000279748 | 11.4160 | 2.5592 | ##### | 2.2382 | 2.5207E-02 | 4.8229E-02 |
| ENSG00000251579 | 7.1921 | 5.6367 | ##### | 2.2381 | 2.5216E-02 | 4.8242E-02 |
| ENSG00000107443\|CCNJ | 1421.6230 | -0.2999 | ##### | -2.2381 | 2.5218E-02 | 4.8243E-02 |
| ENSG00000112473\|SLC39A7 | 10069.6366 | -0.2689 | ##### | -2.2379 | 2.5231E-02 | 4.8265E-02 |
| ENSG00000288886 | 12.2031 | 2.1009 | ##### | 2.2374 | 2.5262E-02 | 4.8323E-02 |
| ENSG00000110880\|CORO1C | 19807.9542 | -0.2541 | ##### | -2.2370 | 2.5289E-02 | 4.8371E-02 |
| ENSG00000276840\|PMS2P13 | 82.3372 | -0.5706 | ##### | -2.2369 | 2.5294E-02 | 4.8376E-02 |
| ENSG00000286445 | 11.0421 | -2.7520 | ##### | -2.2368 | 2.5301E-02 | 4.8387E-02 |
| ENSG00000170296\|GABARAP | 6537.2222 | -0.2799 | ##### | -2.2365 | 2.5316E-02 | 4.8413E-02 |
| ENSG00000262619\|LINC00621 | 13.8477 | 2.1094 | ##### | 2.2359 | 2.5361E-02 | 4.8493E-02 |
| ENSG00000105219\|CCNP | 17.8591 | -1.3706 | ##### | -2.2359 | 2.5361E-02 | 4.8493E-02 |
| ENSG00000278009 | 8.9257 | 1.6914 | ##### | 2.2346 | 2.5445E-02 | 4.8649E-02 |
| ENSG00000291061 | 18.3738 | -1.2020 | ##### | -2.2346 | 2.5446E-02 | 4.8649E-02 |
| ENSG00000235272\|RAMACL | 394.0533 | 0.3431 | ##### | 2.2343 | 2.5464E-02 | 4.8680E-02 |
| ENSG00000131899\|LLGL1 | 3311.7106 | 0.2723 | ##### | 2.2342 | 2.5469E-02 | 4.8686E-02 |

| | | | | | | |
|---|---|---|---|---|---|---|
| ENSG00000278887 | 25.9102 | -2.8825 | ##### | -2.2341 | 2.5476E-02 | 4.8696E-02 |
| ENSG00000277159 | 35.9055 | 0.9366 | ##### | 2.2338 | 2.5495E-02 | 4.8730E-02 |
| ENSG00000255621 | 16.9580 | 1.5005 | ##### | 2.2337 | 2.5506E-02 | 4.8747E-02 |
| ENSG00000183621\|ZNF438 | 395.9602 | 0.3551 | ##### | 2.2336 | 2.5508E-02 | 4.8748E-02 |
| ENSG00000254772\|EEF1G | 49175.7674 | 0.3550 | ##### | 2.2335 | 2.5516E-02 | 4.8762E-02 |
| ENSG00000290010 | 4.4332 | -4.4919 | ##### | -2.2331 | 2.5543E-02 | 4.8810E-02 |
| ENSG00000272732 | 12.9338 | -1.7455 | ##### | -2.2327 | 2.5566E-02 | 4.8851E-02 |
| ENSG00000288560 | 16.6399 | -6.4612 | ##### | -2.2326 | 2.5574E-02 | 4.8862E-02 |
| ENSG00000275757\|5_8S_rRNA | 90.1922 | 0.9100 | ##### | 2.2325 | 2.5579E-02 | 4.8869E-02 |
| ENSG00000108671\|PSMD11 | 19952.8907 | -0.2402 | ##### | -2.2320 | 2.5616E-02 | 4.8936E-02 |
| ENSG00000204619\|PPP1R11 | 4907.8923 | -0.2782 | ##### | -2.2314 | 2.5653E-02 | 4.9003E-02 |
| ENSG00000006576\|PHTF2 | 4548.8698 | 0.2680 | ##### | 2.2311 | 2.5675E-02 | 4.9043E-02 |
| ENSG00000198711\|SSBP3-AS1 | 11.5122 | 1.5815 | ##### | 2.2310 | 2.5681E-02 | 4.9052E-02 |
| ENSG00000105202\|FBL | 16908.9343 | 0.2729 | ##### | 2.2310 | 2.5684E-02 | 4.9053E-02 |
| ENSG00000100100\|PIK3IP1 | 129.9660 | 0.4921 | ##### | 2.2309 | 2.5685E-02 | 4.9053E-02 |
| ENSG00000284664 | 12.4674 | 1.4071 | ##### | 2.2309 | 2.5691E-02 | 4.9061E-02 |
| ENSG00000284337 | 91.1514 | -3.0921 | ##### | -2.2306 | 2.5705E-02 | 4.9084E-02 |
| ENSG00000112130\|RNF8 | 1564.5224 | 0.2923 | ##### | 2.2305 | 2.5712E-02 | 4.9094E-02 |
| ENSG00000285923 | 28.2070 | -1.5149 | ##### | -2.2303 | 2.5724E-02 | 4.9112E-02 |
| ENSG00000259155\|STK16P1 | 4.6420 | 5.4610 | ##### | 2.2304 | 2.5723E-02 | 4.9112E-02 |
| ENSG00000279594 | 4.5567 | 2.4120 | ##### | 2.2302 | 2.5731E-02 | 4.9122E-02 |
| ENSG00000279118 | 9.3123 | 2.9158 | ##### | 2.2301 | 2.5739E-02 | 4.9134E-02 |
| ENSG00000255384 | 7.3489 | 4.7838 | ##### | 2.2301 | 2.5741E-02 | 4.9134E-02 |
| ENSG00000135018\|UBQLN1 | 20957.7447 | -0.2624 | ##### | -2.2301 | 2.5743E-02 | 4.9134E-02 |
| ENSG00000240877\|RN7SL521P | 12.1390 | 1.8183 | ##### | 2.2291 | 2.5805E-02 | 4.9250E-02 |
| ENSG00000289196 | 15.3825 | -1.4549 | ##### | -2.2291 | 2.5807E-02 | 4.9251E-02 |
| ENSG00000289280 | 13.7294 | -7.4299 | ##### | -2.2289 | 2.5823E-02 | 4.9279E-02 |
| ENSG00000166135\|HIF1AN | 5667.5625 | -0.2696 | ##### | -2.2287 | 2.5834E-02 | 4.9295E-02 |
| ENSG00000266171 | 20.8709 | 1.5641 | ##### | 2.2287 | 2.5835E-02 | 4.9295E-02 |
| ENSG00000150625\|GPM6A | 30.3581 | -1.6239 | ##### | -2.2285 | 2.5850E-02 | 4.9320E-02 |
| ENSG00000274297 | 6.2774 | 5.9035 | ##### | 2.2283 | 2.5860E-02 | 4.9337E-02 |
| ENSG00000269292 | 499.3480 | -0.4049 | ##### | -2.2281 | 2.5872E-02 | 4.9357E-02 |
| ENSG00000143761\|ARF1 | 28308.0545 | -0.2631 | ##### | -2.2277 | 2.5902E-02 | 4.9410E-02 |
| ENSG00000131373\|HACL1 | 1973.1566 | 0.2913 | ##### | 2.2274 | 2.5919E-02 | 4.9439E-02 |
| ENSG00000164024\|METAP1 | 8048.8502 | 0.2762 | ##### | 2.2271 | 2.5940E-02 | 4.9475E-02 |
| ENSG00000242493\|RN7SL37P | 2.9723 | 3.6708 | ##### | 2.2270 | 2.5946E-02 | 4.9484E-02 |
| ENSG00000105677\|TMEM147 | 10106.4201 | -0.2771 | ##### | -2.2269 | 2.5952E-02 | 4.9493E-02 |
| ENSG00000278763\|FAM27B | 86.8234 | -0.5022 | ##### | -2.2264 | 2.5987E-02 | 4.9557E-02 |
| ENSG00000044524\|EPHA3 | 23.4003 | 2.2961 | ##### | 2.2260 | 2.6013E-02 | 4.9602E-02 |
| ENSG00000241837\|ATP5PO | 10232.8681 | -0.2660 | ##### | -2.2260 | 2.6015E-02 | 4.9604E-02 |
| ENSG00000132517\|SLC52A1 | 21.8903 | -1.2983 | ##### | -2.2256 | 2.6042E-02 | 4.9652E-02 |
| ENSG00000230432 | 6.8181 | -2.7722 | ##### | -2.2250 | 2.6078E-02 | 4.9717E-02 |
| ENSG00000280387 | 9.8054 | 2.0489 | ##### | 2.2250 | 2.6080E-02 | 4.9718E-02 |
| ENSG00000109079\|TNFAIP1 | 7207.3047 | 0.2664 | ##### | 2.2249 | 2.6091E-02 | 4.9736E-02 |
| ENSG00000138079\|SLC3A1 | 1159.1779 | -2.0282 | ##### | -2.2245 | 2.6118E-02 | 4.9784E-02 |
| ENSG00000285171 | 12.4518 | -7.0489 | ##### | -2.2241 | 2.6143E-02 | 4.9829E-02 |
| ENSG00000181029\|TRAPPC5 | 3496.4768 | -0.2678 | ##### | -2.2238 | 2.6161E-02 | 4.9860E-02 |
| ENSG00000213799\|ZNF845 | 1827.7478 | 0.2721 | ##### | 2.2237 | 2.6169E-02 | 4.9871E-02 |
| ENSG00000273024\|INTS4P2 | 12.7461 | 1.6100 | ##### | 2.2235 | 2.6182E-02 | 4.9894E-02 |
| ENSG00000272449 | 82.5616 | 0.5084 | ##### | 2.2230 | 2.6213E-02 | 4.9948E-02 |
| ENSG00000122778\|KIAA1549 | 2712.5385 | 0.2778 | ##### | 2.2229 | 2.6223E-02 | 4.9964E-02 |
| ENSG00000233680\|HNRNPA1P27 | 3.7638 | -2.3121 | ##### | -2.2229 | 2.6226E-02 | 4.9967E-02 |
